# Biomechanical properties of a buzz-pollinated flower

**DOI:** 10.1101/2020.03.17.995746

**Authors:** Vinicius Lourenço Garcia Brito, Carlos Eduardo Pereira Nunes, Caique Rocha Resende, Fernando Montealegre-Zapata, Mario Vallejo-Marín

**Affiliations:** Instituto de Biologia, Universidade Federal de Uberlândia. Uberlândia, Brazil, 38405-315; Department of Biological and Environmental Sciences, University of Stirling. Stirling, Scotland, United Kingdom, FK9 4LA; School of Life Sciences, University of Lincoln, Lincoln, United Kingdom, LN67DL

**Keywords:** Apidae, bees, biomechanics, buzz pollination, laser vibrometry, pollen release, pollination, *Solanum*

## Abstract

Approximately half of all bee species use vibrations to remove pollen from plants with diverse floral morphologies. In many buzz-pollinated flowers, these mechanical vibrations generated by bees are transmitted through floral tissues, principally pollen-containing anthers, causing pollen to be ejected from small openings (pores or slits) at the tip of the stamen. Despite the importance of substrate-borne vibrations for both bees and plants, few studies to date have characterised the transmission properties of floral vibrations. In this study, we use contactless laser vibrometry to evaluate the transmission of vibrations in the corolla and anthers of buzz-pollinated flowers of *Solanum rostratum*, and measured vibrations in three spatial axes. We found that floral vibrations conserve their dominant frequency (300Hz) as they are transmitted through the flower, but that vibrations in anthers and petals can gain additional harmonics relative to the pure tone of input vibrations. We also found that vibrations are generally amplified (up to >400%) as they travel from the receptacle at the base of the flower to other floral structures, and that anthers vibrate with a higher amplitude velocity than petals. Together, these results suggest that vibrations travel differently through floral structures and across different spatial axes. As pollen release is a function of vibration amplitude, we conjecture that bees might benefit from applying vibrations in the axes associated with higher vibration amplification.

## Introduction

Vibrations play an important role in diverse biological interactions from animal communication to predation and pollination [1–3]. Communication in invertebrates often occurs through vibrations that are transmitted through the substrate, particularly plant structures [4,5]. For example, male and female wandering spiders use plant leaves to detect each other during pre-copulation and some hemipteran predators can detect vibrations produced by leaf-feeding caterpillars during prey search [6,7]. In these cases, substrate properties can affect the vibrations and mediate information transmitted from sender to receiver. To date, our understanding of how plant structures modify the transmission of vibrations is relatively limited [3,5].

Beyond communication, vibrations are also involved in another extraordinary interaction between invertebrates and plants. Some insects, specifically bees, use vibrations to extract pollen grains from certain types of flowers, in a phenomenon called floral buzzing or sonication, which gives rise to the buzz pollination syndrome [8–11]. Buzz pollination has evolved independently multiple times across flowering plants, being found in more than 65 plant families [8,10]. Although the morphology of buzz-pollinated flowers ranges widely [10,12], many buzz-pollinated flowers have repeatedly converged to similar morphologies (e.g. the *Solanum*-like flower type), with anthers that dehisce through small apical pores (poricidal anthers) arranged in a cone-like central structure, as exemplified by some species in genus *Dodecatheon* (= *Primula*, Primulaceae), *Miconia* (Melastomataceae), *Solanum* (Solanaceae) and many others [13–15]. In buzz-pollinated flowers, mechanical vibrations produced by the thoracic muscles of visiting bees are transmitted to floral tissues resulting in pollen ejection via the apical pores [8,10]. Understanding how bee vibrations are transmitted through different floral structures has the potential to illuminate the mechanistic function of buzz-pollinated flowers while providing the background for further ecological and evolutionary studies of buzz pollination from both bee and flower perspectives.

The behaviour of bees while producing floral vibrations is relatively stereotypical [10,12]. Usually, bees land on the flower and use their mandibles and legs to grasp one or multiple stamens, after which they begin rapidly contract their thoracic flight muscles transmitting vibrations to the whole flower [2,16]. Although bees may apply vibrations only to one stamen, pollen can be released from all stamens in the flower simultaneously. This effect is perhaps more dramatically demonstrated in flowers that possess two or more sets of morphologically distinct types of stamens, i.e. they are heterantherous [17–19]. In these heterantherous flowers, bees usually focus their vibration efforts into only one set of stamens, the feeding anthers, while a second set of stamens, the pollinating anthers, deposits pollen on a different part of the bee’s body disproportionally contributing to plant fertilisation [20–23]. Therefore, when bees collect pollen directly from centrally located feeding anthers, the vibrations applied there need to be simultaneously transmitted to the pollinating anthers to enable pollen release. In heterantherous species, the transmission of vibrations from one part of the flower to another has immediate and important consequences for their reproductive success.

In buzz-pollinated flowers of the *Solanum*-type, individual or fused petals form sheet-like, flattened and elastic structures that are projected perpendicular or reflexed from the central axis of the flower (Figure 1). In contrast, the stamens have a cylindrical or conical shape, are made of relatively rigid tissue, and are projected forward in parallel to the central axis of the flower (Figure 1). The different configuration of petals and anthers raises the hypothesis that petals and anthers might differ in the magnitude of vibrations experienced across different axes of the flower. Petals and stamens can be thought of as bending cantilever structures [3,24]. Both petals and stamens have a fixed end attached to the base of the flower and are subject to possible transversal and longitudinal bending vibrations applied by bees. Cantilevers can vary in length and shape being cylindrical, flat or even more complex, but they are defined as rigid or semi-rigid elongated structures often perpendicularly fixed by one end to a support while the other end is free [25]. In cantilever bending, the displacement of the structure’s free end is directly proportional to the cube of its length, but not to its mass, and inversely proportional to the material stiffness and its second moment of area [25]. Feeding and pollinating stamens have different sizes and shapes, suggesting that they might also have different vibrational properties [19]. Therefore, we predict that in heterantherous flowers, longer (pollinating) stamens should show higher vibration amplitudes at their tip during buzz-pollination compared to shorter (feeding) stamens, despite differences in mass.

**Figure 1.**
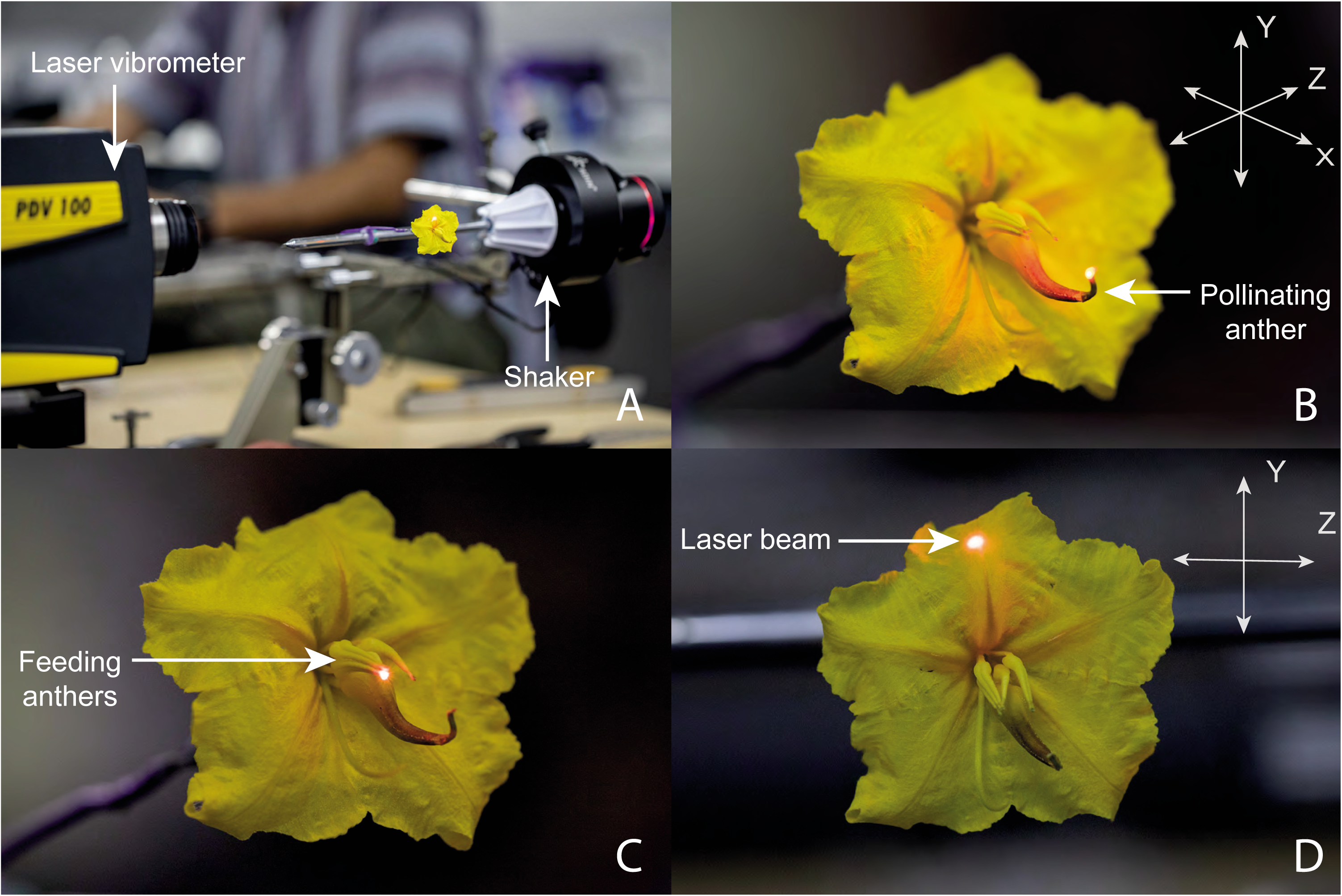
Experimental setup for contactless measurement of the transmission of vibrations through the flowers of a buzz-pollinated plant, *Solanum rostratum* (Solanaceae). (**A**) Experimental set up. The first laser vibrometer (PDV 100) is shown in the foreground, with the laser beam hitting the upper petal of the flower along the z-axis. The vibration transducer system is parallel to the laser beam and is shown in the background. The second laser is aimed at the flower receptacle behind the petals also along the z-axis; the second laser vibrometer is outside of the frame. The three measured axes (x, y and z) are shown as arrows at the top-right of panels B and D. The perspective of the x and z axes in panel B is exaggerated for illustration purposed. Panels B, C and D show the laser beam hitting the pore of the pollinating anther (**B**), the pore of the feeding anther (**C**) or the upper petal (**D**) with the laser beam parallel to the z-axis.

In this study, we aim to characterise the transmission of vibrations in different floral organs in buzz-pollinated plants. Specifically, we address two questions: (1) Does the differences in size and morphology of feeding and pollinating anthers translate into different capacities for transmitting vibrations? (2) Do petals and anthers differ in their transmission of vibrations across different axes of vibration (x, y, and z in Figure 1B)? We address these questions by applying artificial vibrations of similar characteristics of those produced by bees to the base of the flower (receptacle) and recording vibrations in petals or anther tips. Our study is the first to investigate vibration transmission across multiple flower organs and multiple axes of vibration.

## Material and Methods

### Study system

We used *Solanum rostratum* Dunal (Solanaceae), a buzz-pollinated annual herb with pollen-flowers, i.e. flowers that offer only pollen as resource for pollinators (Figure 1) [17,26]. The pollen grains are concealed in anthers that dehisce only through a small pore in their apex (poricidal anthers). The flowers of *S. rostratum* are heterantherous, with two morphologically and functionally distinct sets of stamens in the same flower [22]. One set consists of four anthers (feeding anthers) that are small, bright yellow and centrally located within the flower. The second set consists of a single anther (pollinating anther) that is large, yellow to brown, and positioned opposite to the deflected style [27]. Some bees visiting *S. rostratum* use vibrations to extract and collect pollen grains (floral vibrations; [26,28]). During floral visitation, bees curve their body around the anthers and vibrate their thoracic muscles [11]. In general, pollen release from flowers with poricidal anthers is related to the vibrational properties of floral vibrations [13,29,30].

### Plant growth and flower material

Plants of *S. rostratum* were grown from seeds collected in 2010 from a population near San Miguel De Allende, Queretaro, Mexico (20.902°N, 100.706°W, 2033m a.s.l.; accessions 10s77, 10s79, 10s81, and 10s86). Seeds were germinated and plants grown as described by Vallejo-Marín et al. [31]. Plants were transplanted either to 1.5 L pots or onto flowerbeds (1 m x 5 m, with plants spaced about 50 cm from each other) in the research glasshouses at the University of Stirling. Plants were grown with 16 h of supplemental lighting a day, and supplemental heating (25°/16° C, day/night). Experimental flowers were collected in the morning by cutting an entire inflorescence and placing it in flower foam (Oasis Floral Products, Washington, UK) and transported to the lab in a plastic container. In the lab, the plastic container was kept open in a room with controlled temperature (21°C) and humidity (50% RH). For each trial, we used a single flower with the pedicel cut immediately underneath the receptacle.

### Generation and playback of vibration signals

To study the biomechanical transmission of vibrations through flower structures, we synthesised vibration signals using *Audacity* (www.audacityteam.org). We produced sine waves with 300 Hz frequency, 5 minutes duration, maximum relative amplitude of one, sampling rate of 44.1 kHz, and phase ϕ = 0°. Using a computer, the vibrational signals produced were played in a custom designed playback system (Figure 1A). Briefly, this system consists on a vibration transducer speaker (Adin S8BT 26W, Shenzen, China) with a vibrating metal plate on which we attached (Loctite UltraGel Control, Hemel Hempstead, United Kingdom) a plastic base with a metal rod (15 cm tall and 0.5 cm diameter) was also glued. The vibration transducer was held with a three-pronged clamp attached to a metal base. We glued (Loctite UltraGel Control) a metal wire (6 cm length and 0.1 cm diameter) to the end of the metal rod, which could then be moved to the desired position. To transmit the vibrations to the flower, a single flower was pinned and glued (Loctite UltraGel Control) on the base of their receptacle perpendicularly to the plane formed by the corolla (Figure 1B). Finally, the pinned flower was attached to the metal wire of the vibration transducer system using a reusable adhesive (BluTack, Figure 1A). The system (vibration transducer and attached flower) could then be positioned as required. The definition of x, y and z axes in our study follows Vallejo-Marín [11] and is shown in Figure 1B.

### Measurement of vibrations

Once the flower was correctly positioned, we then deployed two laser Doppler vibrometers (PDV-100, Polytec, Waldbronn, Germany) facing each other (parallel laser beams) and parallel to the axis of the main displacement of the vibration transducer system (Figure 1A). One of the lasers (reference) was always aimed to the flower receptacle, behind the flower, while the other one was aimed either to (1) the pore of the pollinating anther (Figure 1B), (2) the pore of the feeding anther (Figure 1C), or (3) the third upper quarter of the upper corolla petal (Figure 1D). Since the receptacle of *S. rostratum* is covered by trichomes, we used a small amount (2 to 3 mm^2^) of reflective tape to improve the quality of the signal reflected to the vibrometer. Both laser vibrometers were connected to a second computer where the mechanical vibration of the receptacle and the other flower structures were simultaneously recorded using the VibSoft-20 software (Polytec, Waldbronn, Germany).

We set the laser vibrometers to a sensitivity of 500 mm/s and used a low-pass filter of 5 kHz and no high-pass filter. Since vibrations signals, specially amplitude, can damp or amplify during their transmission in plant tissues [5], we adjusted the input stimulus to the desired amplitudes in the flower receptacle using the volume control of the computer and visually checking the amplitude of the reference signal. The amplitude of the input vibration (reference signal in the receptacle) was set as close as possible to one of three amplitude values: 14 mm s^-1^, 28 mm s^-1^ or 57 mm s^-1^ root mean squared amplitude velocity (V_RMS_). The V_RMS_ of the vibrations measured at the receptacle are shown in the Supplementary Table S1. The three target amplitude values correspond to 20 mm s^-1^, 40 mm s^-1^ and 80 mm s^-1^ peak amplitude velocity (V_PEAK_), respectively, and are within the range of floral vibrations measured on flowers during buzz pollination by bumblebees. At a frequency of 300 Hz, these peak velocities correspond to peak accelerations of 37.7 m s^-2^, 75.4 m s^-2^, and 150.8 m s^-2^, and peak displacements of 10.61 µm, 21.22 µm, and 42.44 µm, respectively (Vallejo-Marín 2019).

Vibrations were simultaneously recorded at both receptacle and other flower structures at a sampling frequency of 12,000 samples per second, 781.25 mHz frequency resolution, using a 0-5 kHz bandpass filter, and recorded for 1.28 s. The input vibrations (at the receptacle) could be calibrated very close (± 3 mm s^-1^) to the target amplitude velocities of V_RMS_ = 14 mm s^-1^, 28 mm s^-1^ and 57 mm s^-1^ (Supplementary Table S1). We played the vibrations at each of the three spatial axes and in each of three amplitude levels in a set of 10 fresh, unvisited *S. rostratum* flowers (n = 2 recordings x 3 floral structures x 3 axes x 3 amplitudes levels x 10 flowers = 540 vibration recordings). One of the measurements (feeding anther, y-axis, input velocity 57 mm s^-1^, flower accession 10-s-77-19) was missed during the recording phase and thus not included in the analysis.

### Data analysis

The last 300 ms of each recording was used for analysis. The full recording is shown in Supplementary Figure S1. Frequency spectra were estimated using a Hamming window with length of 512 samples and an overlap of 70%. We estimated the dominant frequency and the number of harmonics from the frequency spectra. The number of harmonics was estimated as peaks in the frequency spectrum at multiples of 300 Hz with a relative amplitude of more than 10%. We also estimated the V_RMS_ as a measure of vibration amplitude. For this analysis we used the packages *seewave* [32] and *tuneR* [33].

We used Poisson linear mixed-effects models implemented in the *R* package *lme4* [34] to compare the number of harmonics on different floral structures. The number of harmonics measured in floral structures were the response variable while floral structure (receptacle, corolla, feeding or pollinating anther) and the axis of measurement (x, y or z) were considered fixed effects. In all models, plant accession (maternal family) was considered a random effect. We compared models with and without the interaction among fixed effects and a null model considering only the random effect using the Akaike information criterion (AIC) [35]. We compared the estimated ΔAIC (the difference between the AIC for the i^th^ model and the minimum AIC among all the models) of each model to choose the best fit. Values of ΔAIC within 0–2 are considered to have substantial support, within 4–7 considerably less support, and greater than 10 essentially no support [35]. The AIC as well as other model parameters for comparison were estimated using *AICcmodavg* [36].The best model was used for parameter estimation of the fixed effects and their interaction assessed using *lmerTest* [37]. Differences in number of peaks in floral structures were analysed a posteriori using pairwise t-tests with Holm correction [38]. The data were plotted using *sciplot* [39].

To determine how vibration amplitudes are transmitted to different floral structures, we used linear mixed effects models with Gaussian distribution also implemented in the *R* package *lme4* [34]. In these models, the vibration amplitude velocity (V_RMS_) measured in the corolla, feeding or pollinating anthers were the response variable. The input amplitude velocity (V_RMS_) measured at the receptacle, the type of floral structure (corolla, feeding anther or pollinating anther), and the axis of measurement (x, y or z) were considered fixed effects, and plant accession (maternal family) were considered a random effect. We also compared multiple models with and without interactions among fixed effects and the null model only with the random effect using the same AIC methodology described above [35]. Model predictions of the fixed effects in this analyses were plotted using the *pred* option in *sjPlot* [40]. All analyses were done in *R* ver. 3.6.2 (R Core Team, 2020). The code as well as the raw data will be made available upon publication.

## Results

### Frequency of floral vibrations

Floral organs (receptacle, petals, feeding and pollinating anthers) mostly presented the same dominant frequency of the input signal (300 Hz), no matter the vibration axes considered (Figure 2). Only 2.5% of the estimated dominant frequencies differed from the input frequency (Supplementary Figure S1). The number of harmonics tended to increase on corolla and anthers, but such increase depended on the axis of measurement. The statistical model considering the interaction between floral structure and axis of vibration was the best one in predicting the number of harmonics in floral structure (Supplementary Table S2 and S3). Although we found more harmonic peaks in anthers than in receptacles in all spatial axes, we found no difference in the number of harmonics between anther types (Figure 3). The number of harmonics was higher in the corolla than in receptacle only in the x-axis (Figure 3).

**Figure 2.**
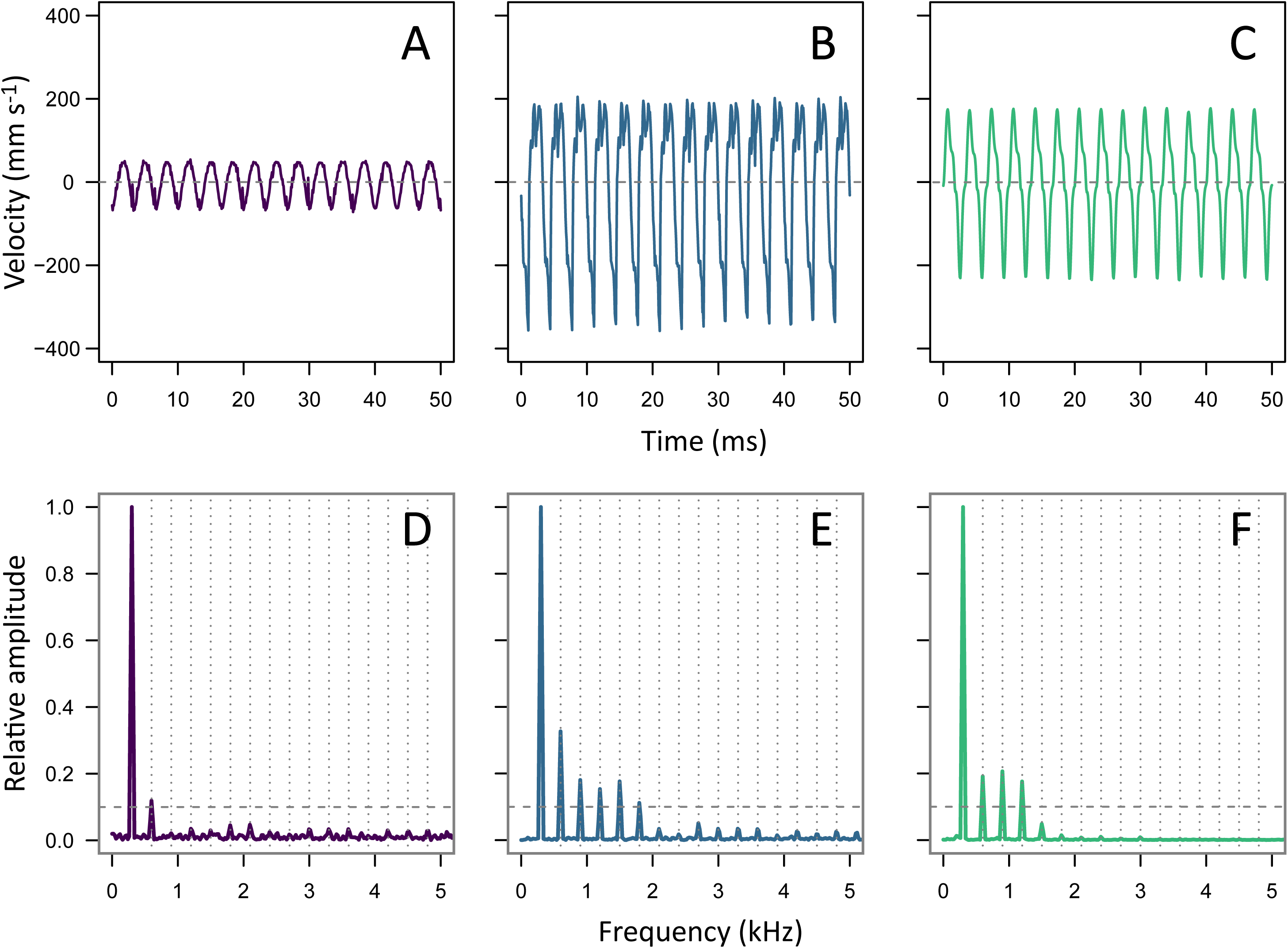
Floral vibrations transmitted from the receptacle at the base of the flowers to three different floral organs of the buzz-pollinated plant *Solanum rostratum*: Petal (A, D), feeding anther (B, E), and pollinating anther (C, F). The top panel shows a 50 ms section of the floral vibration in the time domain, and the bottom panels show the same floral vibration in the frequency domain. Vertical dotted grey lines indicate the considered harmonics of the dominant input frequency (300 Hz) while horizontal dashed grey lines indicate the 0.1 threshold to a given peak be considered a harmonic peak. All vibrations shown here were measured in flower accession 10-s-77-12, in the y-axis and with an input velocity of V_RMS_ = 28 mm s^-1^.

**Figure 3.**
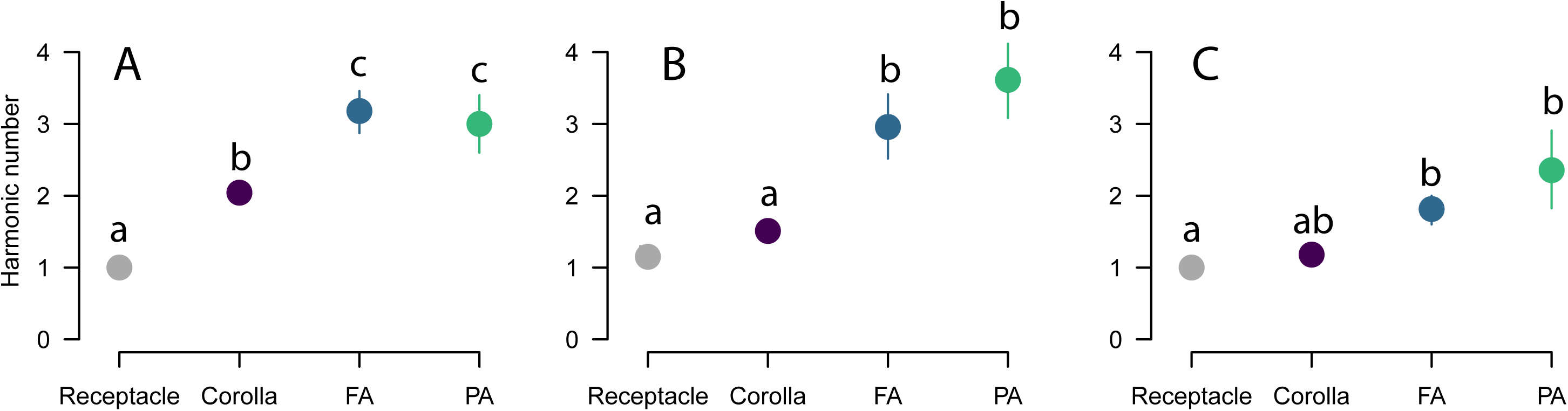
Number of harmonics in each spatial axis and floral structures of a buzz-pollinated plant, *Solanum rostratum* (Solanaceae). The number of harmonics was estimated as multiples of 300 Hz with a relative amplitude of more than 0.1. Symbols and bars represent the mean values and the 95% confidence intervals. Different letters indicate statistically significant differences as assessed in a post-hoc test with Holm correction. FA = Feeding anthers; PA = Pollinating anther.

### Amplitude modification in floral structures

The selected statistical model for amplitude velocity included two-way interactions between input velocity and both floral structure and axis of measurement (Supplementary Table S4 and S5). These statistical interactions suggest that differences in amplitude velocity depend on both floral structure and the axis of measurement. The amplitude velocity (V_RMS_) of vibrations transmitted from the receptacle to the petals closely resembled the input vibration amplitude, showing little evidence of either damping or resonance (Table 1, Figure 4). In contrast, the vibrations transmitted to the tip of both feeding and pollinating anthers were consistently of higher amplitude velocity than the input velocity, revealing amplitude increases that ranged from 69% to 443% (Table 1, Figure 4). The axis with the smallest velocity amplification was the x-axis (69%—110%), while the y- and z-axes showed a higher signal amplification (176%—397% in the y-axis and 229%—443% in the z-axis; Table 1, Figure 4). Within anthers, both pollinating and feeding anthers showed a very similar relationship with input velocity across the three axes of measurement (Figure 4).

**Table 1.**
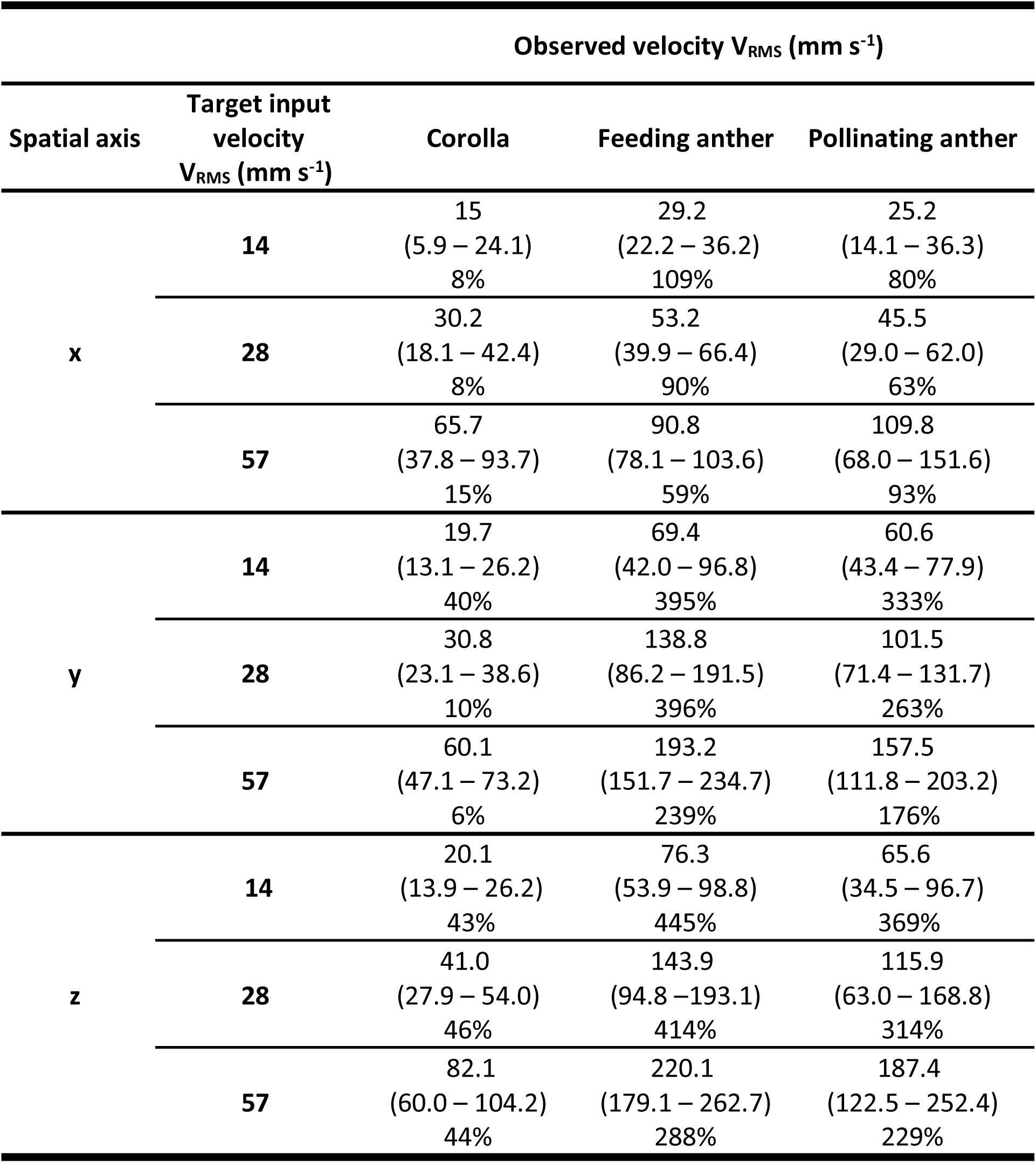
Transmission of vibrations in buzz-pollinated flowers of *Solanum rostratum* (Solanaceae). Input vibrations were applied with a mechanical shaker at the base of the flower (receptacle), and measured at one of three floral structures: petals, feeding anthers or pollinating anthers. The vibrations were applied and measured along the same axis (x, y or z; see Figure 1). Input vibrations had a frequency of 300 Hz, and a RMS amplitude velocity (V_RMS_) of either 14, 18 or 57 mm s^-1^. The table shows the mean V_RMS_ and the 95% confidence intervals (CI; in parenthesis) of measured floral vibrations in each of the three floral structures and axes. The percentage shows the relative change of the measured vibration compared to the targeted input vibration. Positive values indicate amplitude increases.

**Figure 4.**
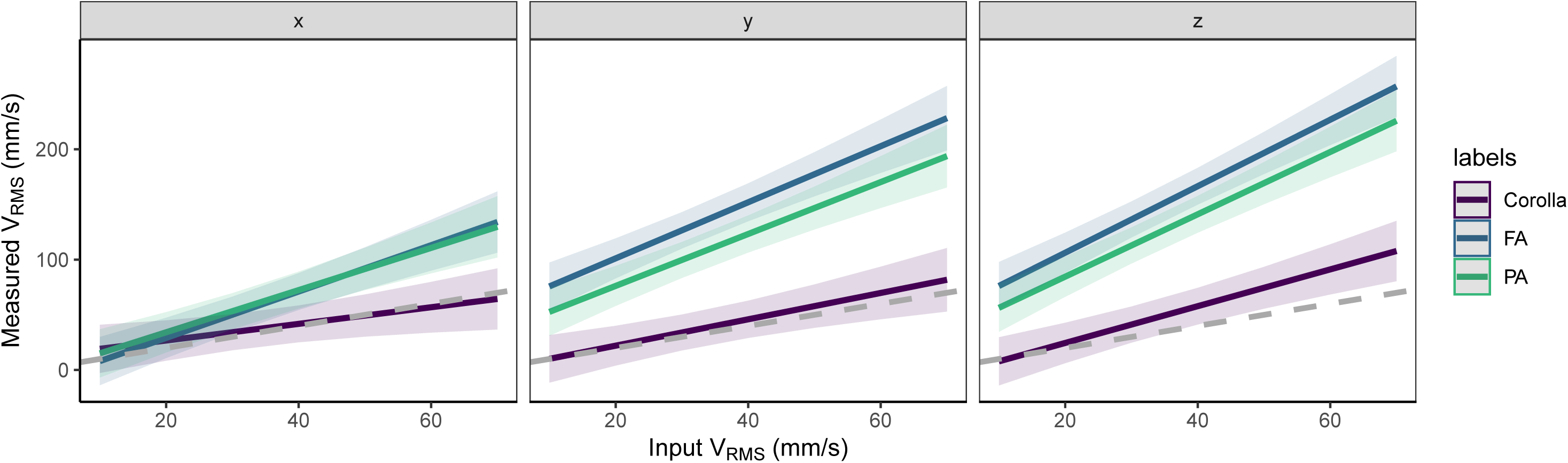
Model estimates of input amplitude velocity (V_RMS_) measured at the receptacle, type of floral structure (corolla, feeding anther or pollinating anther), and axis of measurement (x, y or z) on the root mean squared velocity (V_RMS_) of floral vibrations in *Solanum rostratum*. Lines show the model fit and shadowed areas indicate standard errors.

## Discussion

Our study contributes to understanding how vibrations are transmitted through floral tissues across multiple spatial axes and on multiple floral organs. We found that the spectral properties of floral vibrations are little affected during their transmission through flowers, with most vibrations retaining their fundamental frequency, and some vibrations, particularly on anthers, being enriched with additional harmonics. Our experiments indicate that the amplitude of floral vibrations (V_RMS_) are magnified up to >400% in anthers and to a much lesser extent in petals. However, this amplification depends on the axis on which the vibrations are applied. Together, our results suggest that floral structures differ in their capacity to transmit vibrations and raise the possibility that the way in which vibrations are applied to the flower (for example, where the vibrations are applied and along which axis) may influence their magnitude and thus their effect on pollen release.

### Frequency of floral vibrations

We found that the dominant frequency of floral vibrations remains mostly unchanged as it travels through the flower. Similarly, studies of vibration transmission through plant tissues during insect communication have also shown preservation of spectral characteristics, which is essential to intra- or inter-specific localisation and recognition [41–43]. Interestingly, we found that other floral structures, particularly anthers, show additional harmonics. Multiple harmonics of steeply decreasing amplitude have been recorded in buzz-pollinated flowers [11,29,44,45]. These harmonics also characterise bee vibrations when measured directly on the bee’s thorax [46]. Here we have shown that such harmonics arise in floral structures, particularly stamens, even when the input vibration is a sinusoidal wave with a pure tone (300 Hz).

Vibrations including harmonics are expected in some structures including strings and cantilever beams. Interestingly, the two contrasting types of anthers measured presented a similar number of harmonics. Since it is expected that longer and more flexible cantilever beam structures vibrate with a higher number of harmonics [25], we would predict that pollinating anthers have more harmonics than feeding anthers, since the latter are 37% shorter [26,31]. Differences in stiffness between anther types may compensate for the differences in the length of anthers and generate the observed pattern. In other words, our results suggest that the shorter feeding anthers might be more flexible than the longer pollinating anthers causing them to vibrate with the same number of harmonics under the same vibration source. Despite the occurrence of harmonics in floral vibrations [10], their functional significance is currently unknown. From a biomechanical perspective, harmonics may increase the energy of vibrations. Whether the rapidly decaying higher harmonics observed in floral vibrations contribute to pollen release or not remains to be tested empirically.

### Amplitude modification in anthers

Our finding that floral structures, particularly anthers, amplify the amplitude velocity of input vibrations, while frequency remains the same or is enriched by harmonics, suggest that anther pores are experiencing amplified accelerations [11] and thus higher forces at the anther tips (force = mass x acceleration; [47]). In the cantilever beams, it is expected that amplitude of oscillation at their tip is positively related to their length [25]. However, we found no difference in the recorded amplitude at different anther types. As these anthers also present different morphologies [48], it is possible that others stamen properties such as stiffness and the second moment of area compensates for differences in anther length, reducing the expected differences in measured amplitude.

Despite the amplification of the V_RMS_ measured in the corolla, vibrations transmitted to petals are lower than the vibrations transmitted to the anthers in the same axes. In *S. rostratum*, petals do not vibrate in higher amplitudes along the axis parallel to the central axis of the flower as it would be expected if the petal conformed to a simple cantilever [25]. When predicting displacement of the free end of cantilevers, it is important to consider possible counter forces restricting the bending experienced by the beam as a whole [25]. In *S. rostratum*, the fusion among petals together with the relatively large air-contact and attachment areas may explain the lack of amplitude amplification as well as of amplitude differences among the three spatial axes in corolla. Differences in biomechanical properties among petals and anthers suggest that vibrations measured in the corolla may not necessarily reflect the vibrations experienced at anther tips.

### Bee vibrations

Our study applied vibrations at the base of the flower, which allowed us to study the biomechanical properties of freely moving floral structures. However, during buzz-pollination, floral structures are in direct contact with the body of the vibrating bee. For instance, bees tend to grasp the anthers with their mandibles while curling their body around the anthers [10]. Such direct contact between bees and floral structures are likely to influence the vibration properties of the coupled bee-flower system. Further work is required to understand how the interaction between bee and flower during buzz pollination (e.g., where and how the bee holds the flower) affect the vibrations experienced by the anthers. Nevertheless, our study indicates that the axis in which floral vibrations are applied will influence the magnitude of the amplitude velocity experienced at the anther tips. As vibrations of higher amplitude are associated with increased pollen release [29], bees might benefit from exploiting the biomechanical properties of flowers described here and applying vibrations in the axes associated with higher vibration amplitudes. Previous work has shown that buzz-pollinating bees improve their pollen collection as they gain experience in manipulating flowers [49]. It is possible that such pollen-collection improvement involves adjustments on the handling of the flower, including the axis in which the bee applies floral vibrations. If this is the case, we would predict that vibrating bees adjust their position during flower visiting to match the axis of higher vibration amplitude of the anthers (i.e. y and z axes). Future work on how bees manipulate flowers during buzz pollination, particularly how they apply vibrations to anthers, will help elucidate if bees are able to exploit the biomechanical properties of flowers to maximise pollen collection.

## Acknowledgements

We thank Alistair Gordon for building the vibration playback system (B.Sc. Dissertation, University of Stirling, 2016) and the Vallejo-Marín Lab for help with plant growth and discussions on buzz pollination. We thank FAPEMIG (APQ02497-16) and UFU-CAPES-PrInt (88887.374220/2019-00) grants to VLGB supporting two visits to the University of Stirling, and a Research Grant from The Leverhulme Trust (RPG-2018-235) to MVM and FMZ.

## Supplementary material

**Figure S1**. Oscillograms and frequency spectra of all 539 analysed vibrations. Spectral analysis was done on the last 300 ms of the recorded vibration. Red dashed lines on the frequency spectrum indicates the 0.1 relative amplitude threshold to compute the number of harmonic peaks.

**Supplementary Table S1.** Transmission of vibrations in buzz-pollinated flowers of *Solanum rostratum* (Solanaceae). Input vibrations were applied with a mechanical shaker at the base of the flower (receptacle), and measured at one of three floral structures: petals, feeding anthers or pollinating anthers. The vibrations were applied and measured along the same axis (x, y or z; see Figure 1). Input vibrations had a frequency of 300 Hz, and a RMS amplitude velocity (V_RMS_) of either 14, 18 or 57 mm s^-1^. The table shows the mean V_RMS_ and the 95% confidence intervals (CI; in parenthesis) of measured floral vibrations in the flower’s receptacle. Negative or positive values indicate damping or amplifying effects during vibration transmission, respectively.

| Spatial axis | Target input $V_{\text{RMS}}$ (mm s<sup>-1</sup>) | Observed $V_{\text{RMS}}$ (mm s<sup>-1</sup>) at flower's receptacle |
| --- | --- | --- |
| x | 14 | 15.2 (14.8 – 15.5)<br>8% |
|  | 28 | 29 (28.5 – 29.4)<br>4% |
|  | 57 | 56.8 (56.0 – 57.5)<br>-0.4% |
| y | 14 | 14.2 (13.8 – 14.6)<br>1% |
|  | 28 | 29.2 (28.5 – 29.8)<br>4% |
|  | 57 | 53.4 (53.7 – 56.5)<br>-4% |
| z | 14 | 14.9 (14.4 – 15.4)<br>6% |
|  | 28 | 29 (28.4 – 29.6)<br>4% |
|  | 57 | 57.1 (56.4 – 57.8)<br>0.3% |

**Table S2.** Comparison of generalised mixed-effects models with Poisson distribution analysing the number of peaks in the frequency spectrum of floral structures of *Solanum rostratum*. In all models, number of peaks with a relative amplitude higher than 10% were considered the response variable, and floral structure (*str* = receptacle, corolla, feeding or pollinating anther) as well as axis of measurement (*axes* = x, y or z) were considered the fixed effects. Plant accession was considered a random effect. Models were built in a decreasing order of complexity from a full model including interactions. * = interaction; K = number of parameters; AIC = Akaike information criteria; ΔAIC = difference between the AIC for the considered model and the minimum AIC among all the models; Wt = model probabilities; LL = Log Likelihood.

| model | K | AIC | ΔAIC | Wt | LL |
| --- | --- | --- | --- | --- | --- |
| ***str\*axes*** | 13 | 1547.98 | 0.00 | 1 | 1 |
| ***axes*** | 5 | 1561.05 | 13.07 | 0 | 1 |
| ***str*** | 4 | 1725.21 | 177.23 | 0 | 1 |
| **null** | 2 | 1739.88 | 191.90 | 0 | 1 |

**Supplementary Table S3.** Statistical analysis of the effect of floral structure (corolla, feeding or pollinating anther) and axis of measurement (x, y or z) on number of frequency peaks of transmitted vibrations measured in different parts of a buzz-pollinated flower of *Solanum rostratum*. Floral vibrations were applied at the base of the flower using a mechanical shaker attached to the flower’s receptacle and measured either at the anther tips of feeding and pollinating anthers or at the distal end (1/4) of the upper petal. The model shown here was selected among competing linear mixed-effects models with Poisson distribution using AIC. The model includes the second-order interactions among floral structure and axis of measurement. In this model, both axis of measurement and floral structure were considered as fixed effects and plant accession as a random effect. The reference values for the model (intercept) are corolla and x-axis. Only fixed-effects are shown in the Table.

| | Estimate ± SD | F |
| --- | --- | --- |
| **Intercept** (Corolla, x-axis) | 0.72 ± 0.13 | |
| **Floral structure** |  | 69.56 |
| Feeding anther | 0.45 ± 0.16 | |
| Pollinating anther | 0.38 ± 0.16 | |
| Receptacle | -0.73 ± 0.16 | |
| **Spatial axis** |  | 18.88 |
| y | -0.28 ± 0.19 | |
| z | -0.54 ± 0.21 | |
| **Floral structure × Spatial axis** |  | 2.59 |
| Feeding anther × y | 0.22 ± 0.24 | |
| Pollinating anther × y | 0.64 ± 0.24 | |
| Receptacle × y | 0.47 ± 0.24 | |
| Feeding anther × z | -0.04 ± 0.27 | |
| Pollinating anther × z | 0.30 ± 0.26 | |
| Receptacle × z | 0.54 ± 0.26 | |

**Supplementary Table S4.** Comparison of mixed-effects models explaining the vibration amplitude recorded in floral structures of *Solanum rostratum*. In all models, V_RMS_ measured in floral structures was considered the response variable and floral structure (*str* = corolla, feeding or pollinating anther), axis of measurement (*axes* = x, y or z) and/or recorded V_RMS_ in the receptacle (*rec*) were considered fixed effects. Plant accession was considered a random effect. Models were built in a decreasing order of complexity from a full model including interactions. * = interaction; K = number of parameters; AIC = Akaike information criteria; ΔAIC = difference between the AIC for the considered model and the minimum AIC among all the models; Wt = model probabilities; LL = Log Likelihood.

| model | K | AIC | $\Delta AIC$ | Wt | LL |
| --- | --- | --- | --- | --- | --- |
| *str + axes + rec + str\*axes + str\*rec + str\*axes* | 16 | 2797.62 | 0.00 | 0.70 | 0.70 |
| *str + axes + rec + str\*axes + str\*rec* | 14 | 2800.11 | 2.49 | 0.20 | 0.91 |
| *str\*axes\*rec* | 20 | 2801.67 | 4.05 | 0.09 | 1.00 |
| *str + axes + rec + str\*axes + rec\*axes* | 14 | 2810.18 | 12.57 | 0.00 | 1.00 |
| *str + axes + rec + str\*axes* | 12 | 2812.21 | 14.59 | 0.00 | 1.00 |
| *str + axes + rec + str\*rec + rec\*axes* | 12 | 2824.71 | 27.09 | 0.00 | 1.00 |
| *str + axes + rec + str\*rec* | 10 | 2826.34 | 28.72 | 0.00 | 1.00 |
| *str + axes + rec + rec\*axes* | 10 | 2834.66 | 37.04 | 0.00 | 1.00 |
| *str + axes + rec* | 8 | 2835.96 | 38.34 | 0.00 | 1.00 |
| null | 3 | 3051.70 | 254.09 | 0.00 | 1.00 |

**Supplementary Table S5.**
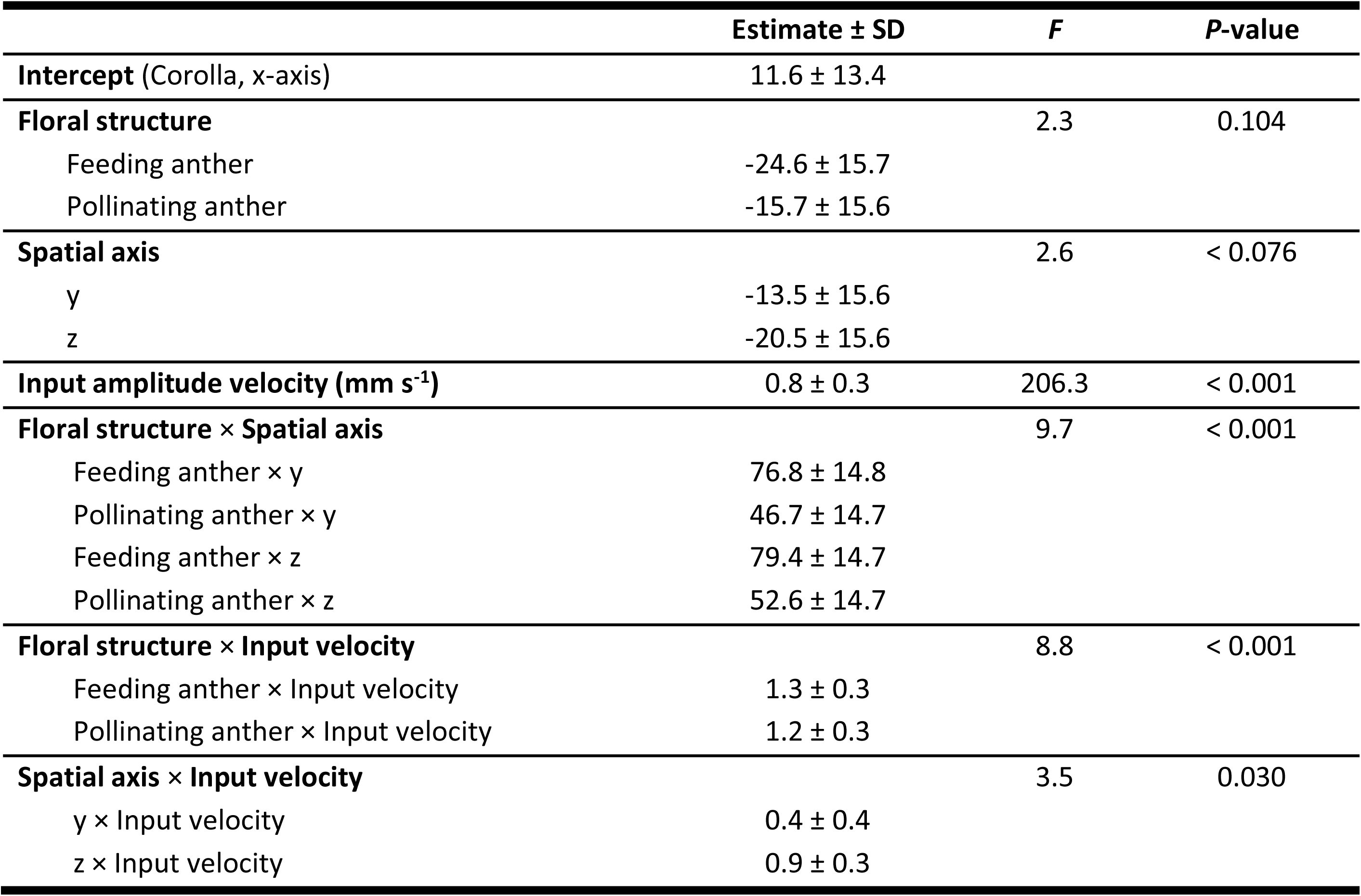
Statistical analysis of the effect of floral structure (corolla, feeding or pollinating anther), axis of measurement (x, y or z) and amplitude of input vibrations on the Root Mean Squared amplitude velocity (V_RMS_; mm s^-1^) of transmitted vibrations measured in different parts of a buzz-pollinated flower of *Solanum rostratum*. Floral vibrations were applied at the base of the flower using a mechanical shaker attached to the flower’s receptacle and measured either at the anther tips of feeding and pollinating anthers or at the distal end (1/4) of the upper petal. The model shown here was selected among competing linear mixed-effects models using AIC. The model includes all second-order interactions among input amplitude velocity (V_RMS_ measured in the receptacle), floral structure and axis of measurement. In this model, input amplitude velocity, axis of measurement, and floral structure measured were considered as fixed effects and plant accession as a random effect. The reference values for the model (intercept) are petal and z-axis. Only fixed-effects are shown in the table.

