## Supplementary Figure S1 for "Biomechanical properties of a buzz-pollinated flower"

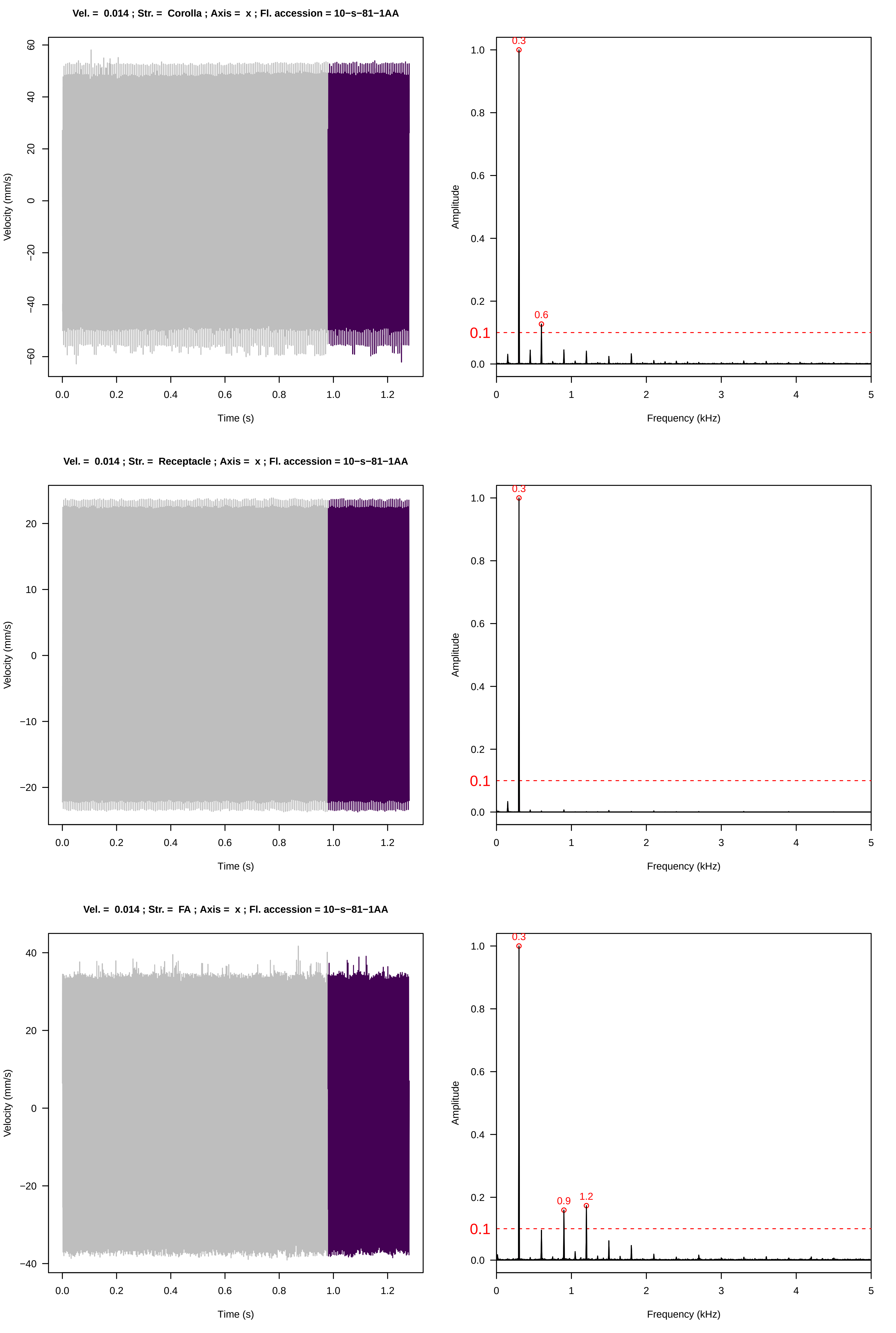

Vel. = 0.014 ; Str. = Corolla ; Axis = x ; Fl. accession = 10−s−81−1AA
Velocity (mm/s)
Time (s)
Amplitude
Frequency (kHz)
0.3
0.6
0.1
Vel. = 0.014 ; Str. = Receptacle ; Axis = x ; Fl. accession = 10−s−81−1AA
Velocity (mm/s)
Time (s)
Amplitude
Frequency (kHz)
0.3
0.1
Vel. = 0.014 ; Str. = FA ; Axis = x ; Fl. accession = 10−s−81−1AA
Velocity (mm/s)
Time (s)
Amplitude
Frequency (kHz)
0.3
0.9
1.2
0.1

**Vel. = 0.014 ; Str. = Receptacle ; Axis = x ; Fl. accession = 10−s−81−1AA**

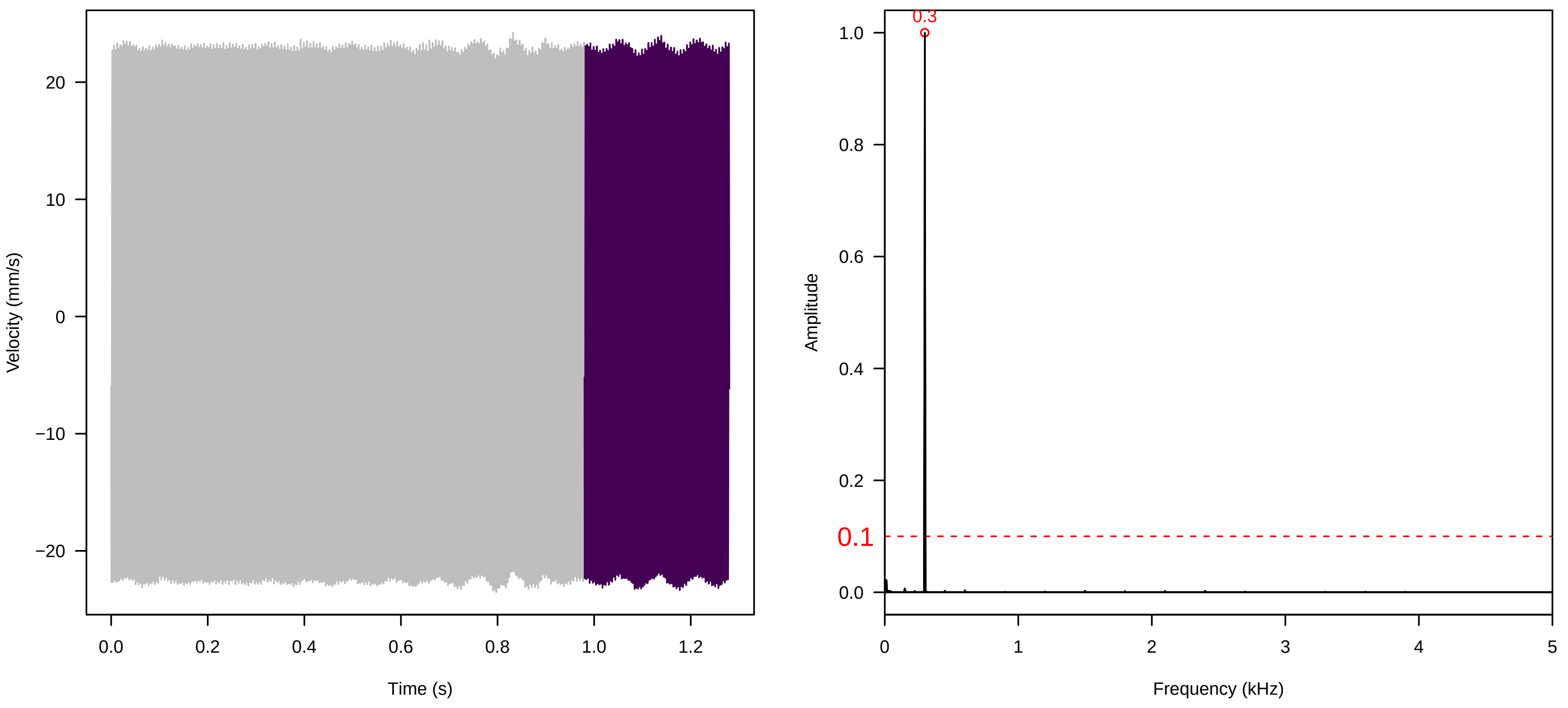


**Vel. = 0.014 ; Str. = PA ; Axis = x ; Fl. accession = 10−s−81−1AA**

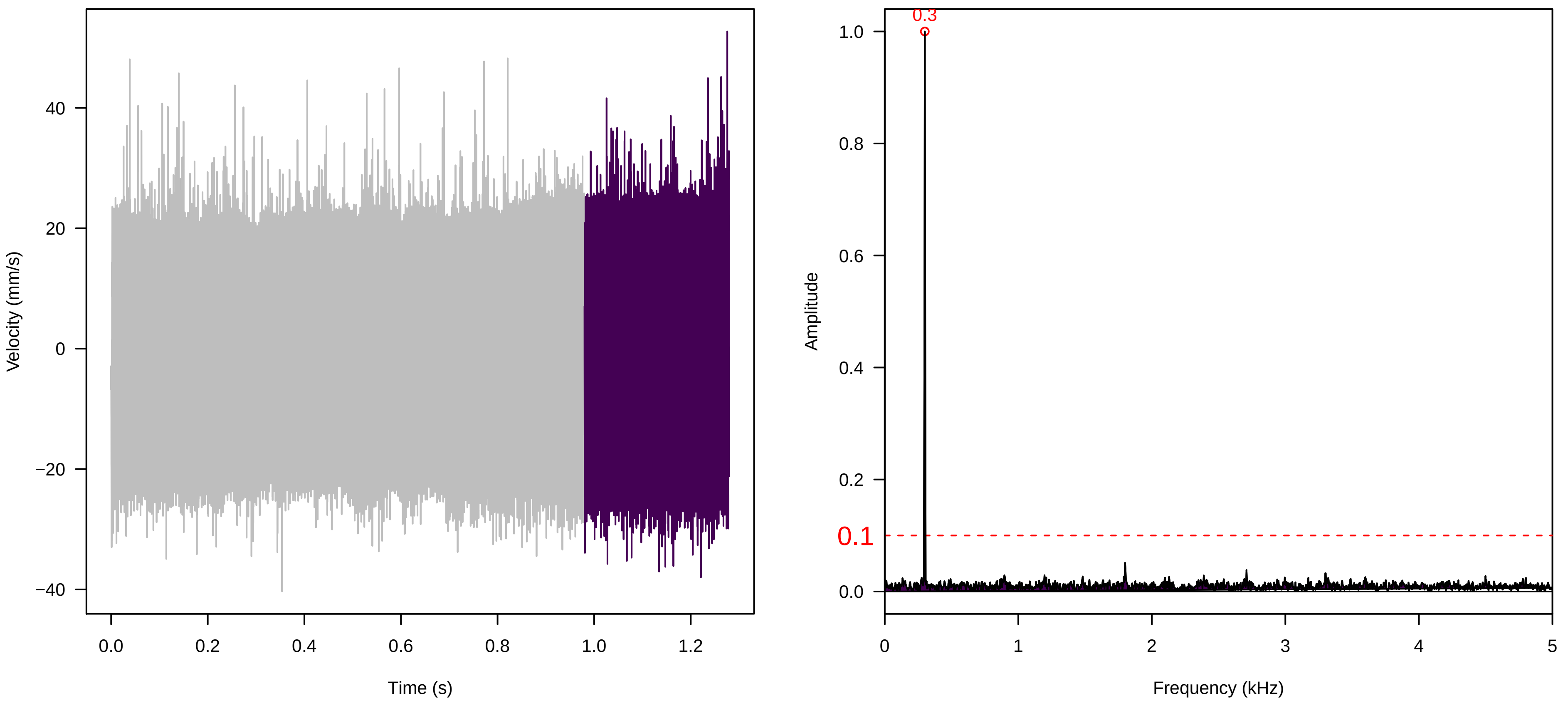


**Vel. = 0.014 ; Str. = Receptacle ; Axis = x ; Fl. accession = 10−s−81−1AA**

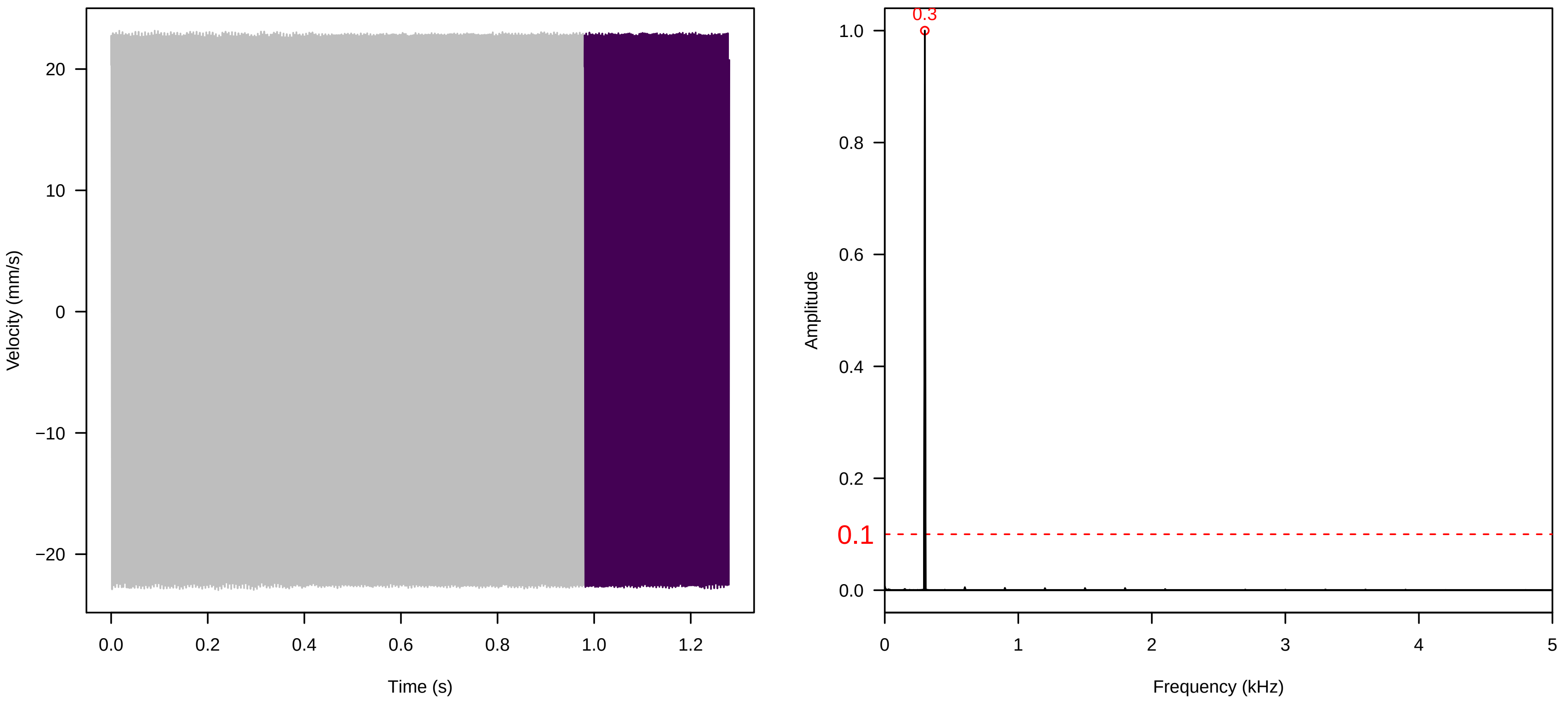

**Vel. =  0.028 ; Str. =  PA ; Axis =  x ; Fl. accession = 10−s−81−1AA**

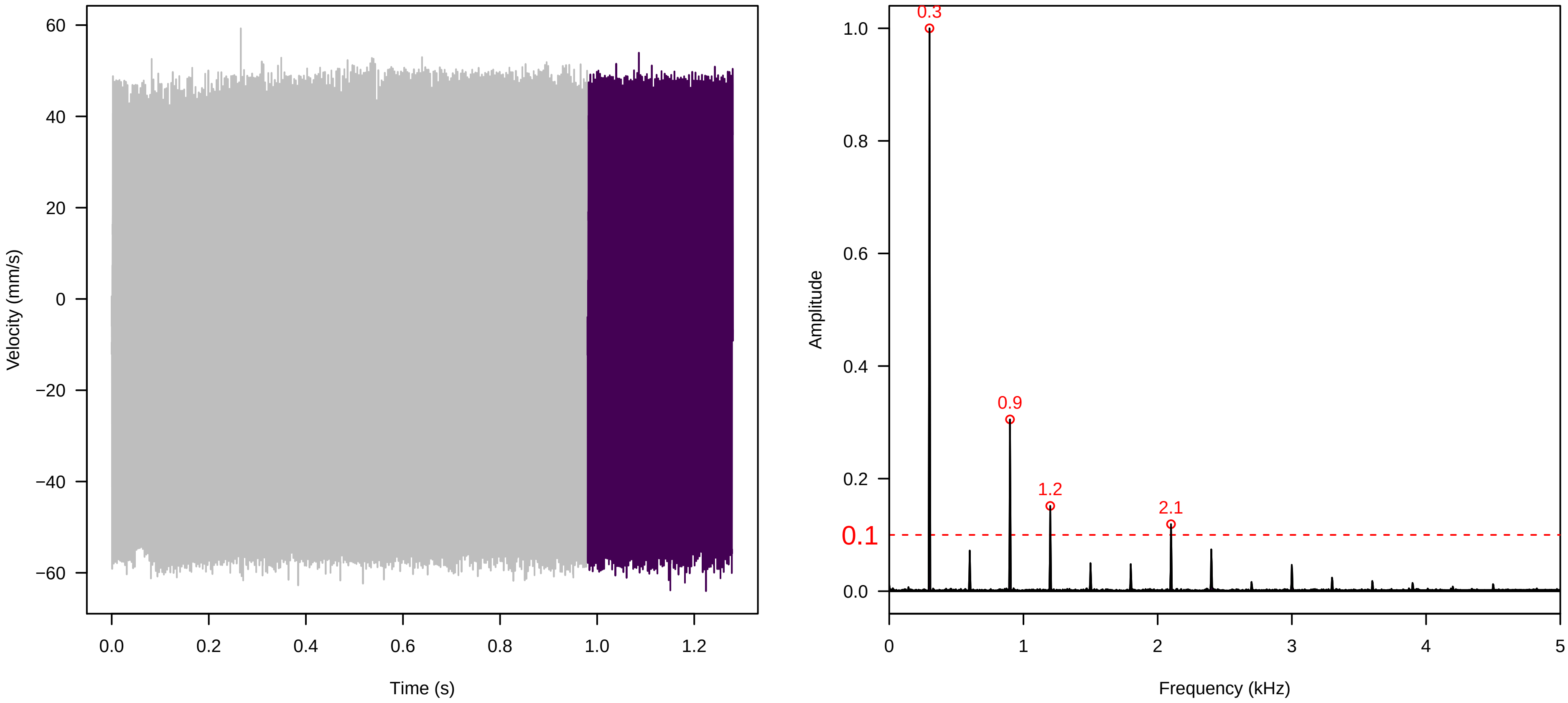


**Vel. =  0.028 ; Str. =  Receptacle ; Axis =  x ; Fl. accession = 10−s−81−1AA**

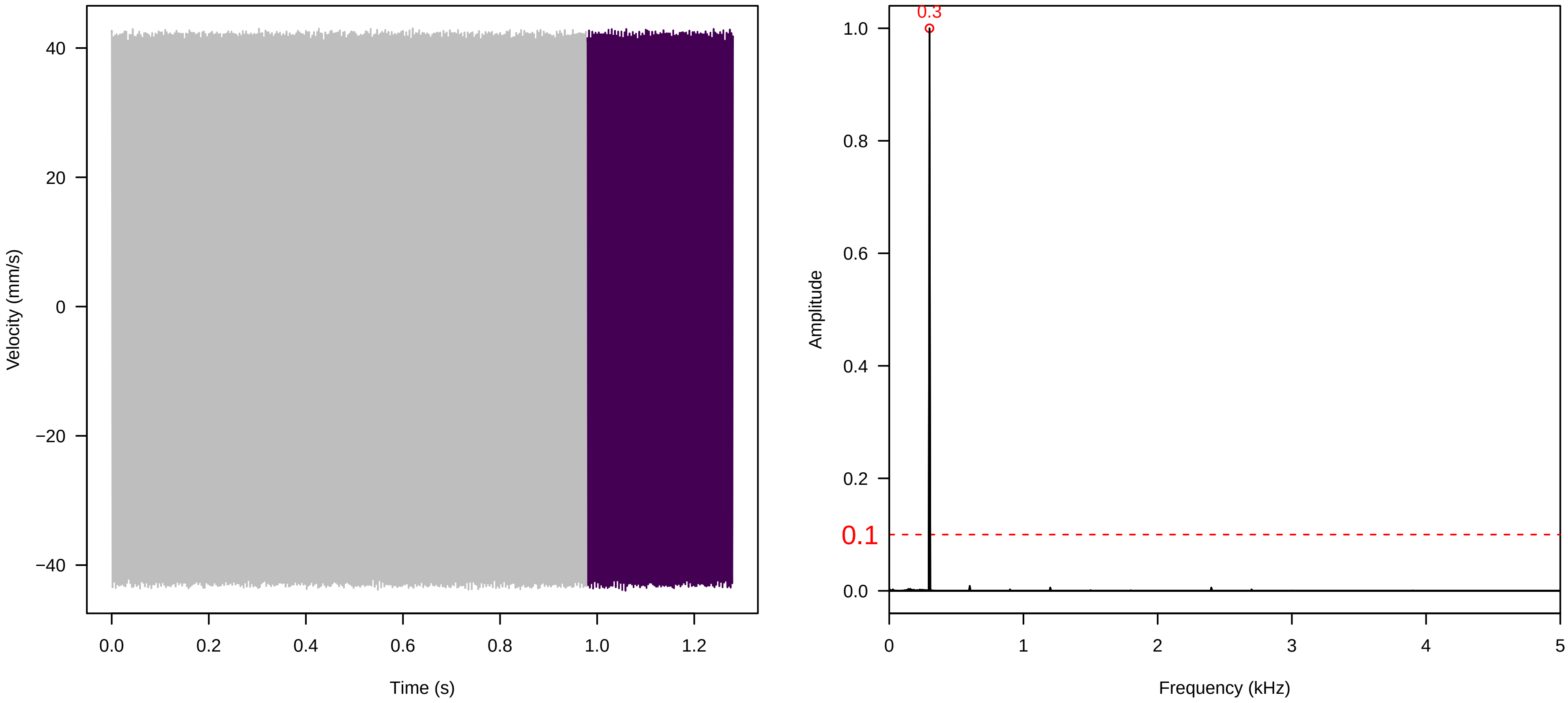


**Vel. =  0.028 ; Str. =  FA ; Axis =  x ; Fl. accession = 10−s−81−1AA**

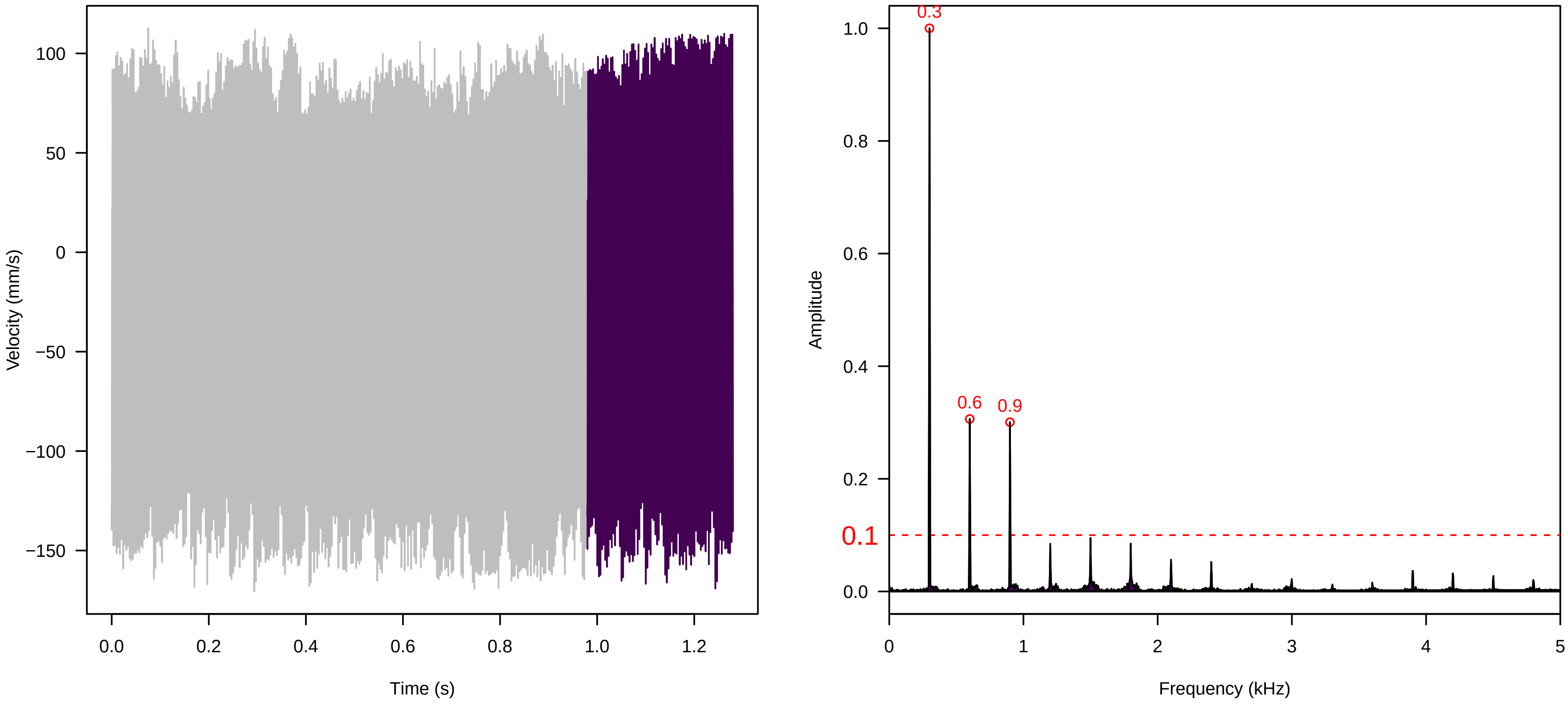

**Vel. = 0.028 ; Str. = Receptacle ; Axis = x ; Fl. accession = 10−s−81−1AA**

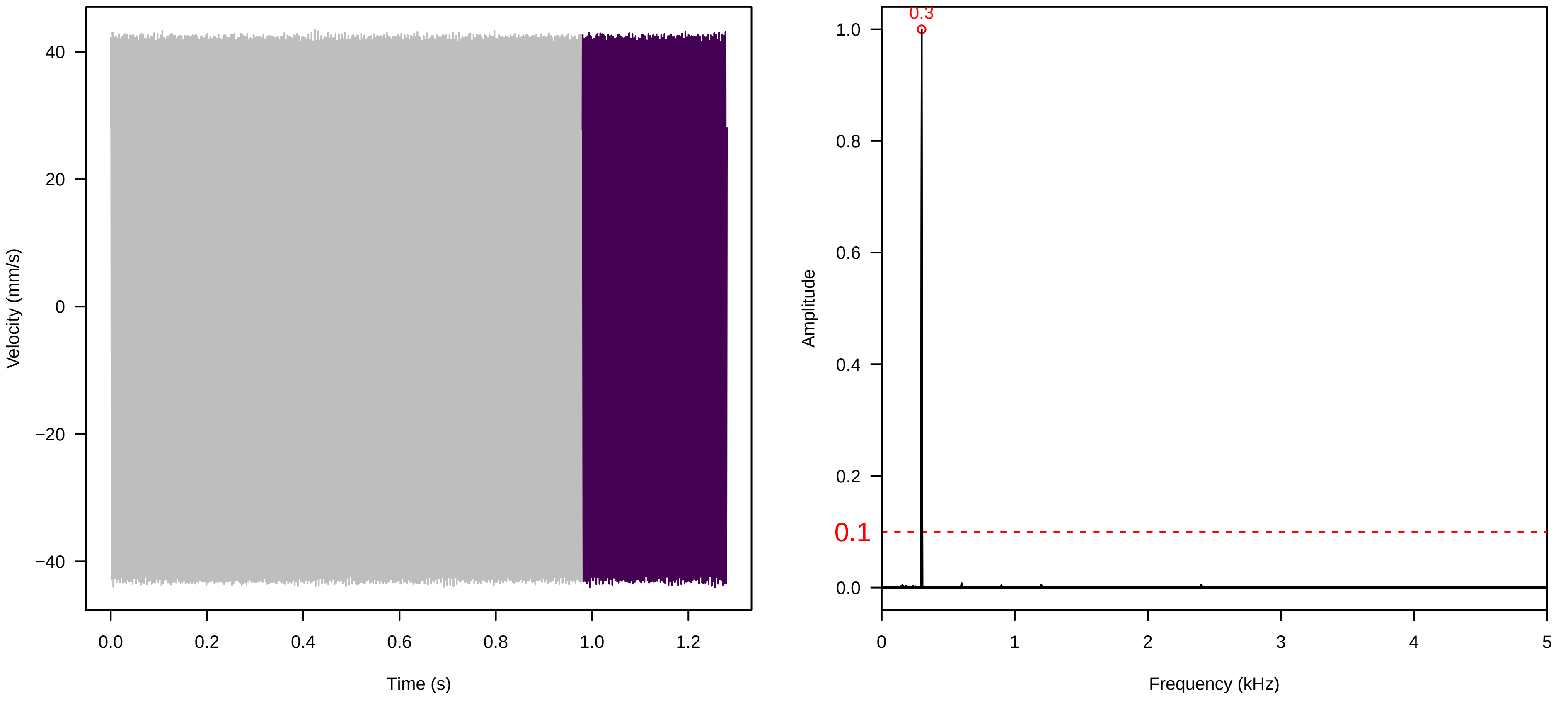


**Vel. = 0.028 ; Str. = Corolla ; Axis = x ; Fl. accession = 10−s−81−1AA**

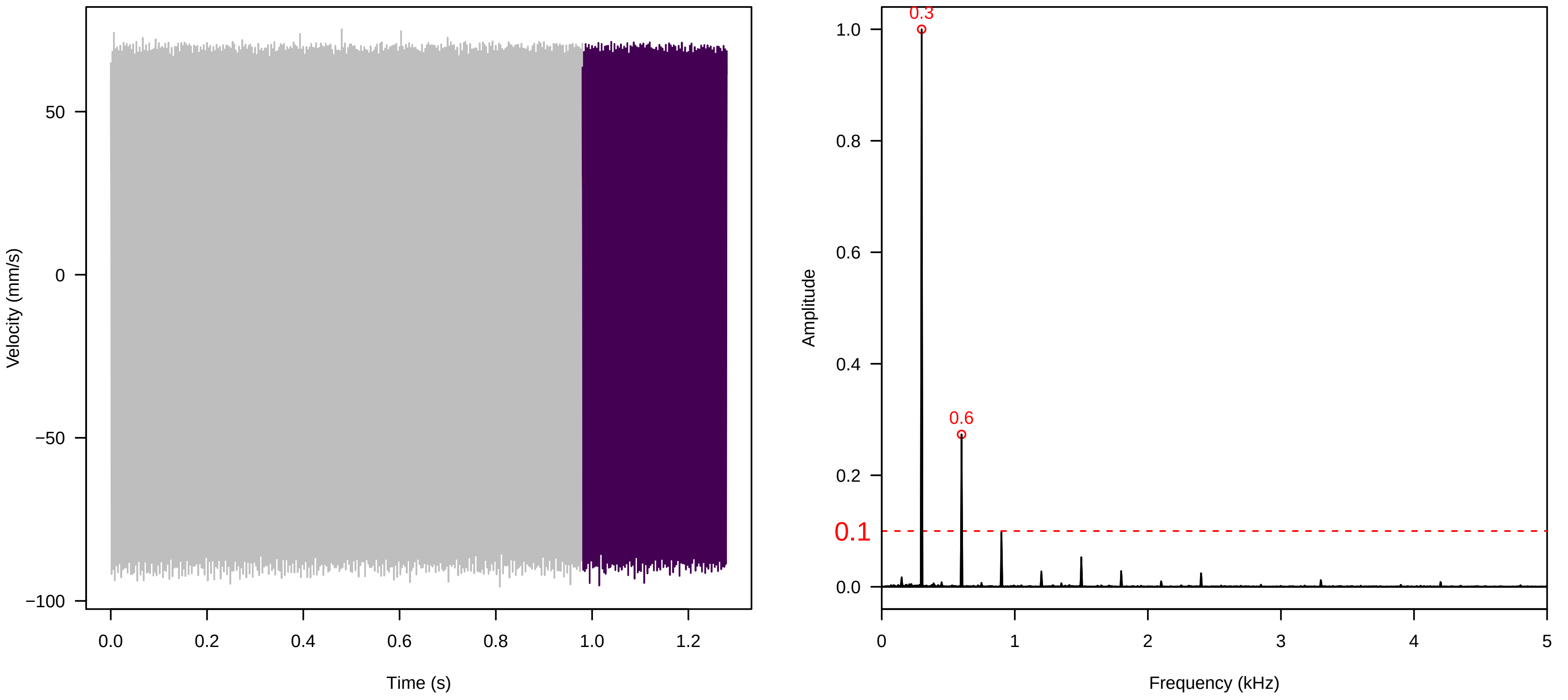


**Vel. = 0.028 ; Str. = Receptacle ; Axis = x ; Fl. accession = 10−s−81−1AA**

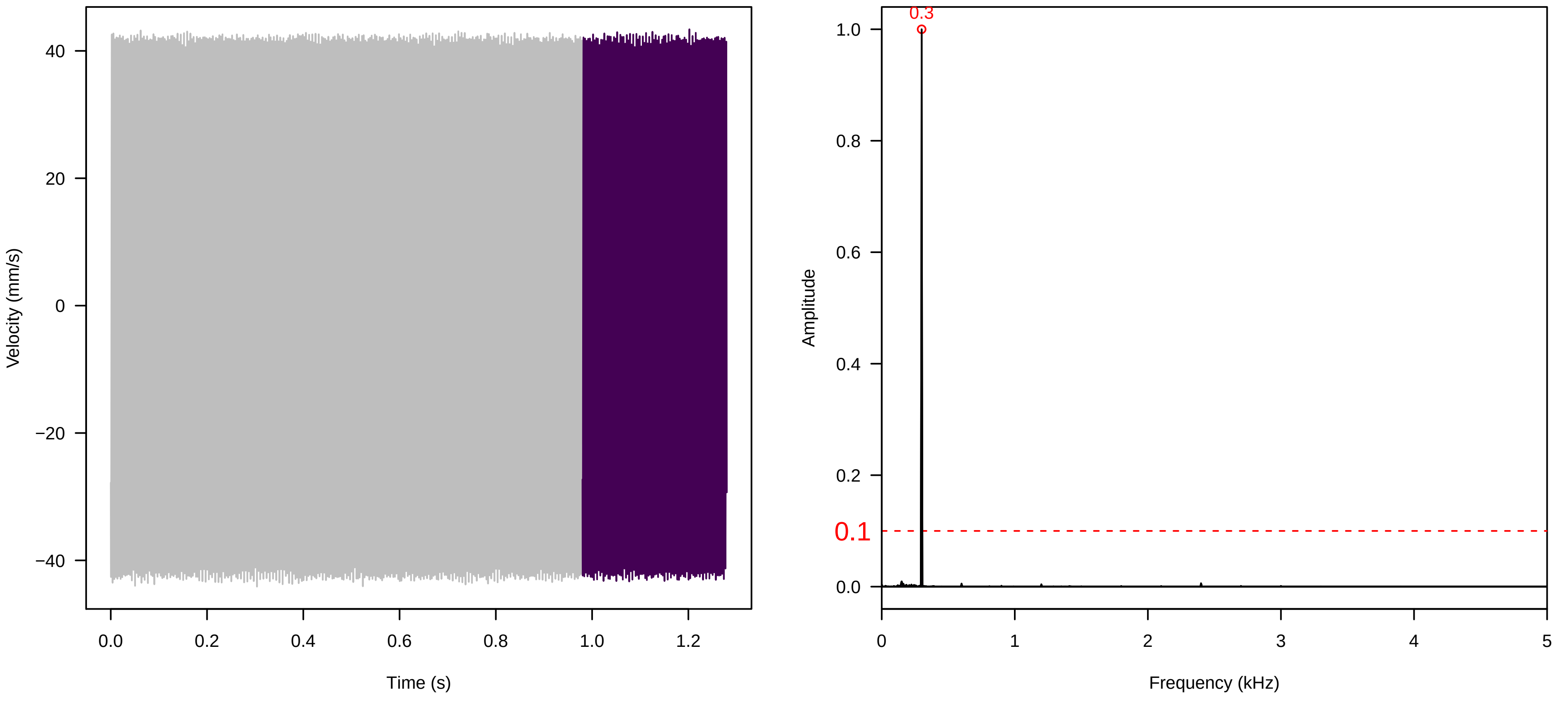

**Vel. = 0.057 ; Str. = Corolla ; Axis = x ; Fl. accession = 10−s−81−1AA**

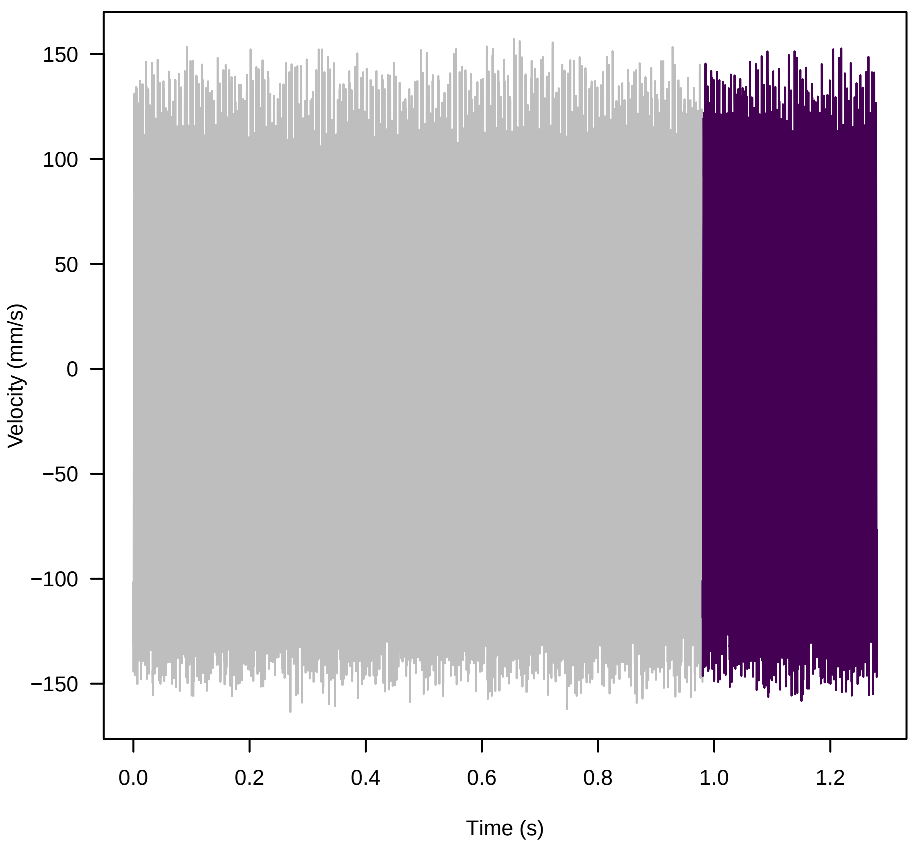


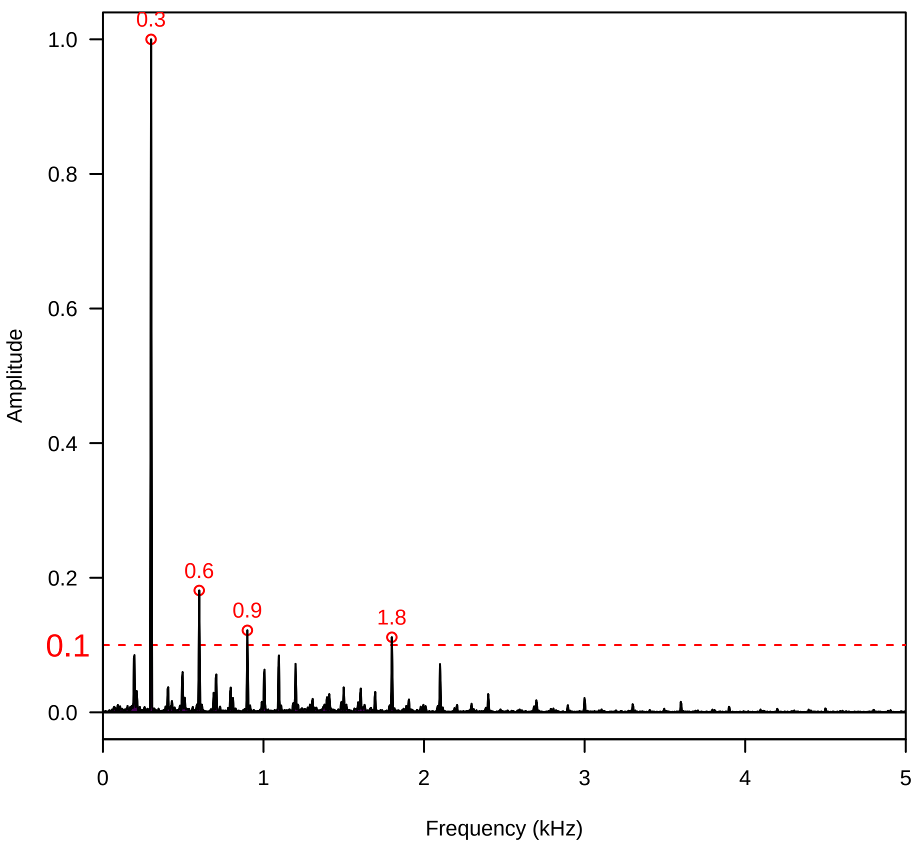


**Vel. = 0.057 ; Str. = Receptacle ; Axis = x ; Fl. accession = 10−s−81−1AA**

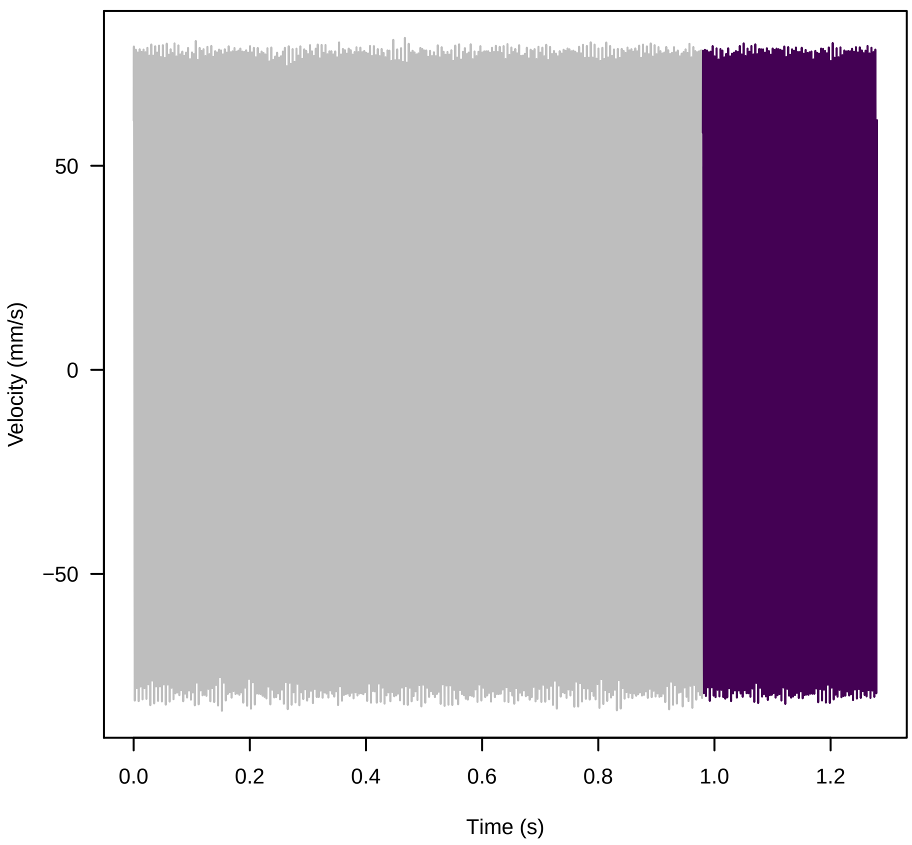


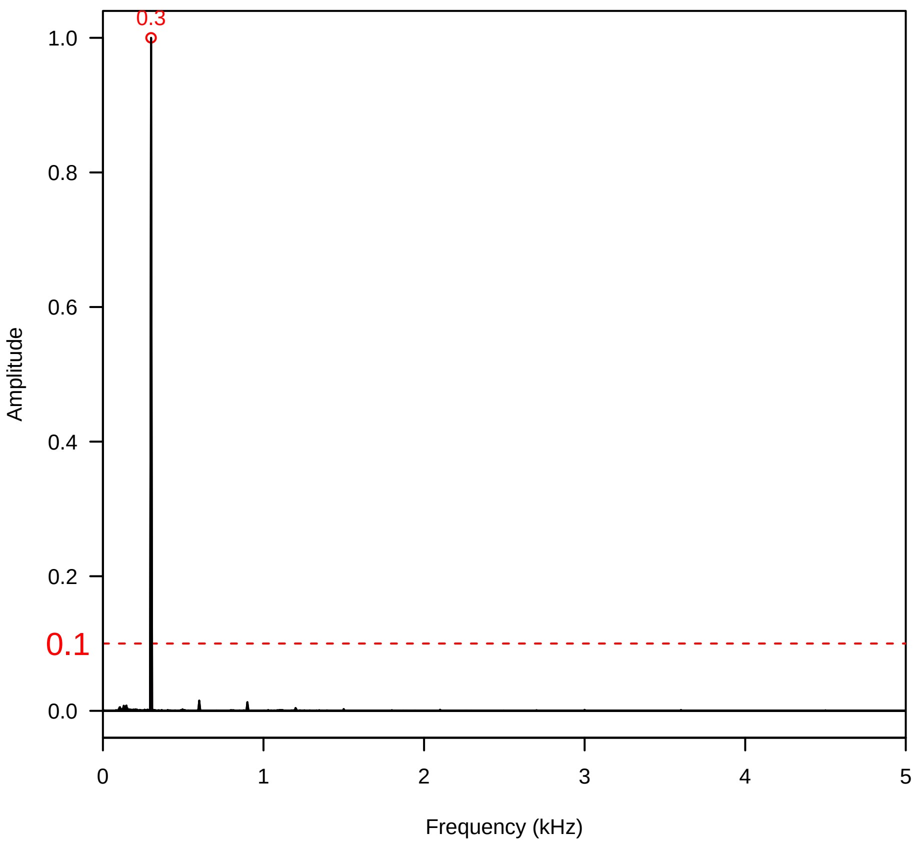


**Vel. = 0.057 ; Str. = FA ; Axis = x ; Fl. accession = 10−s−81−1AA**

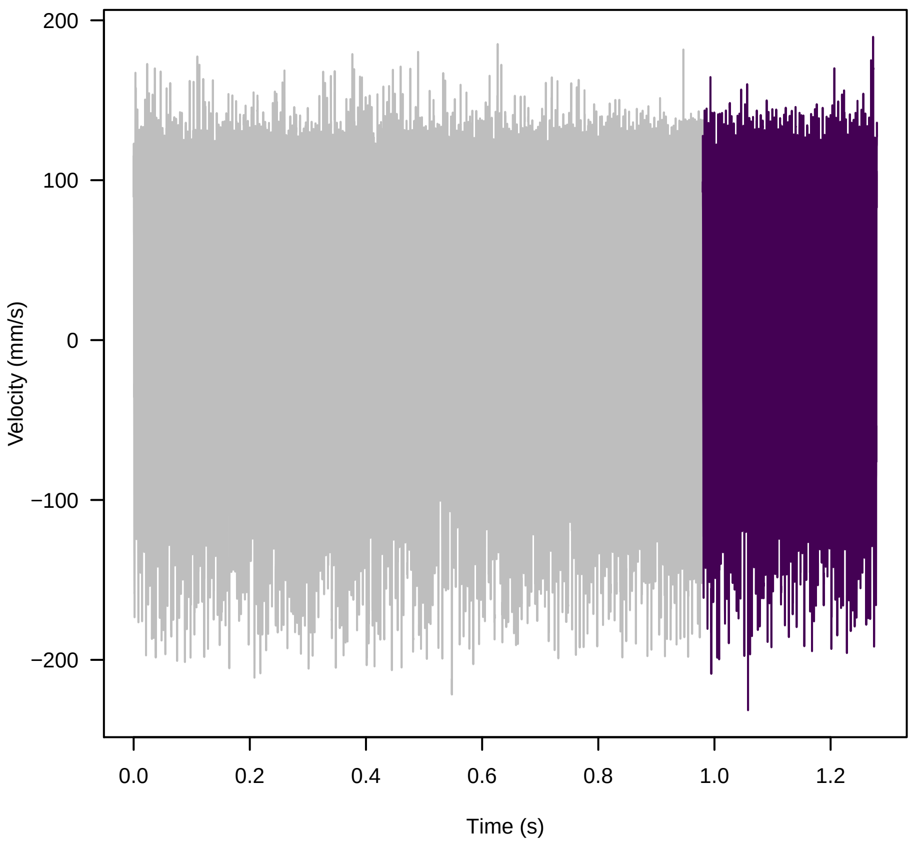


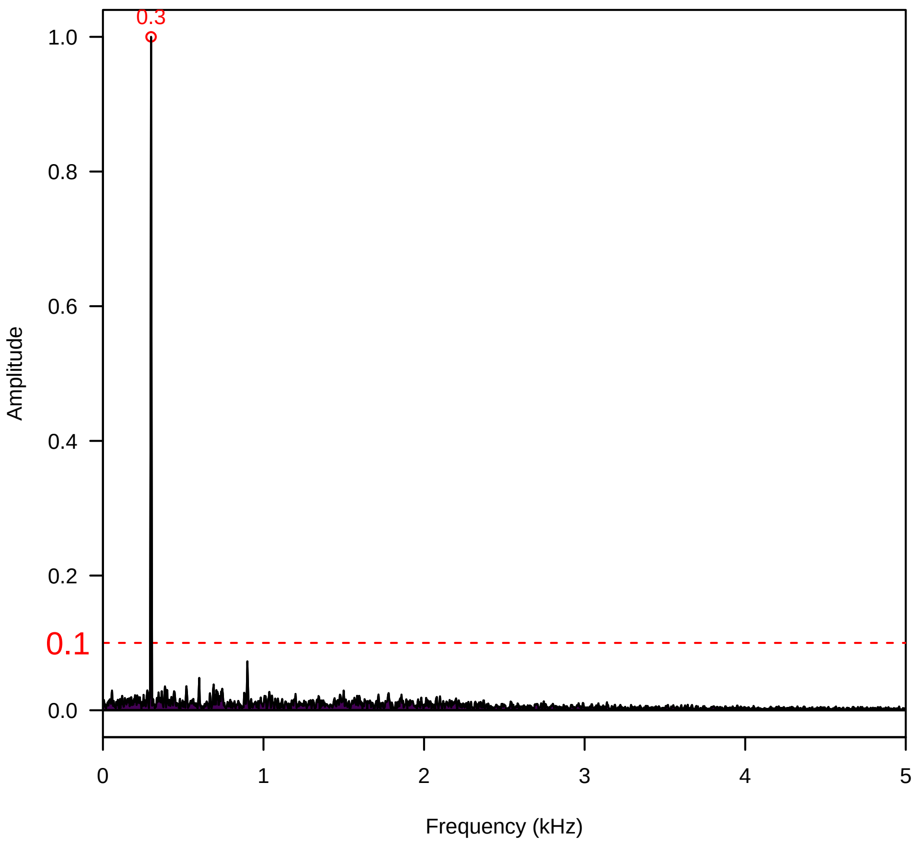

**Vel. = 0.057 ; Str. = Receptacle ; Axis = x ; Fl. accession = 10−s−81−1AA**

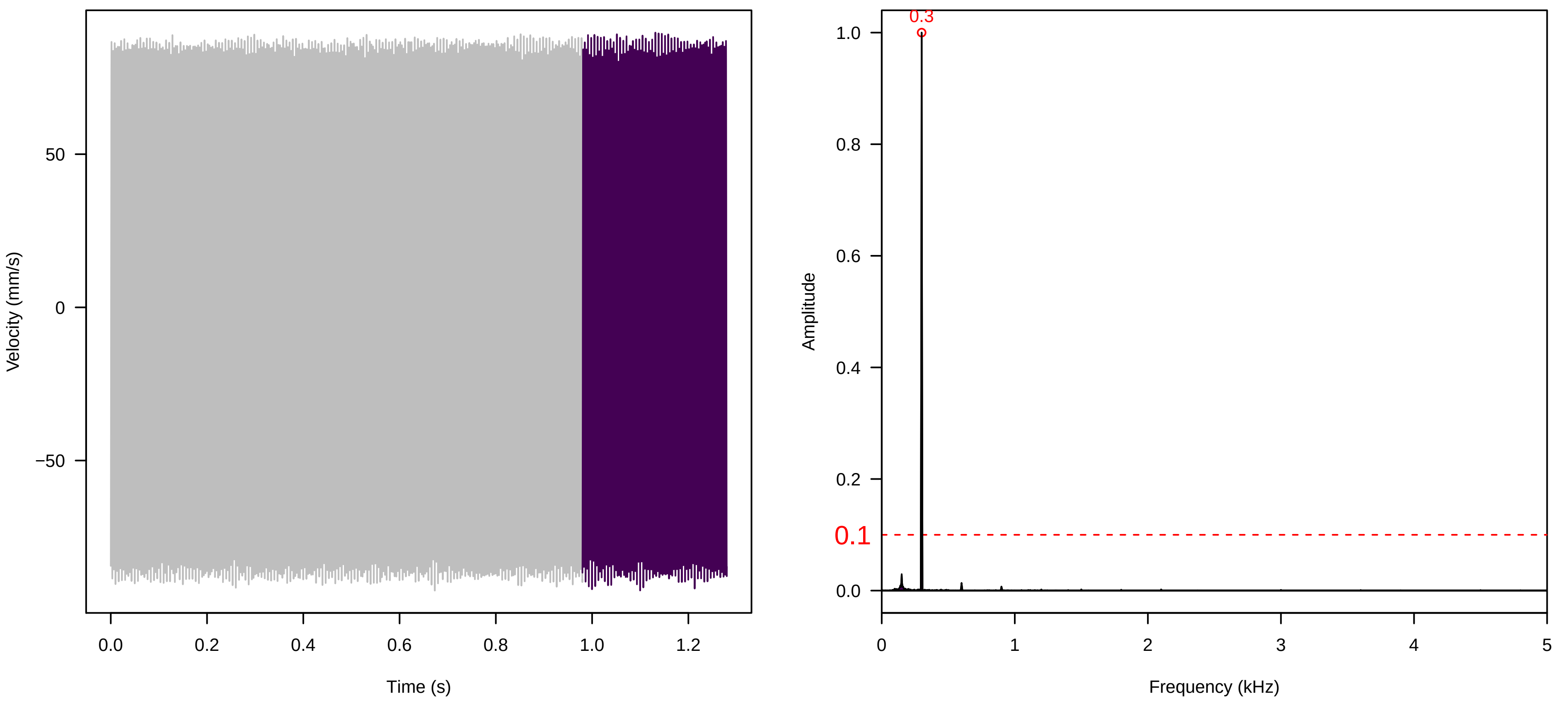


**Vel. = 0.057 ; Str. = PA ; Axis = x ; Fl. accession = 10−s−81−1AA**

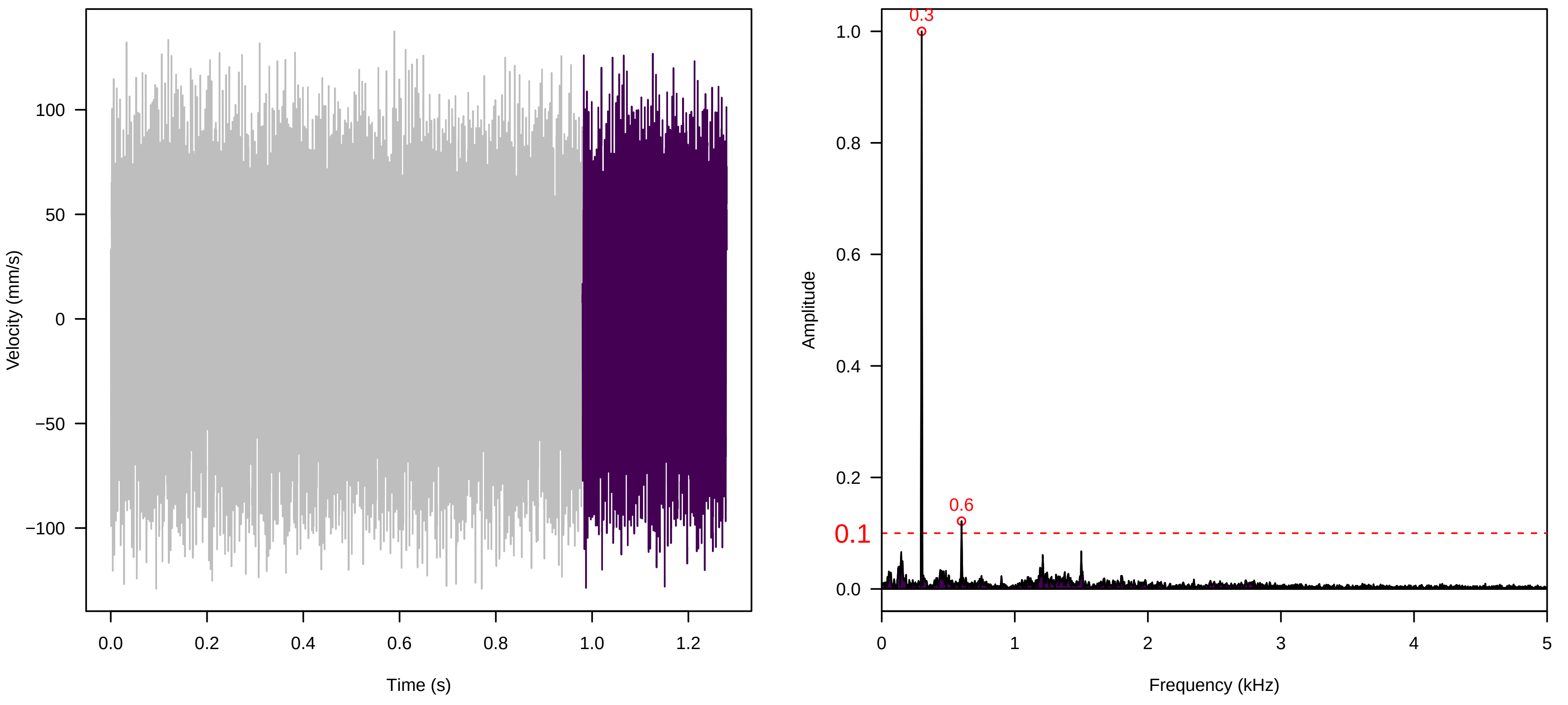


**Vel. = 0.057 ; Str. = Receptacle ; Axis = x ; Fl. accession = 10−s−81−1AA**

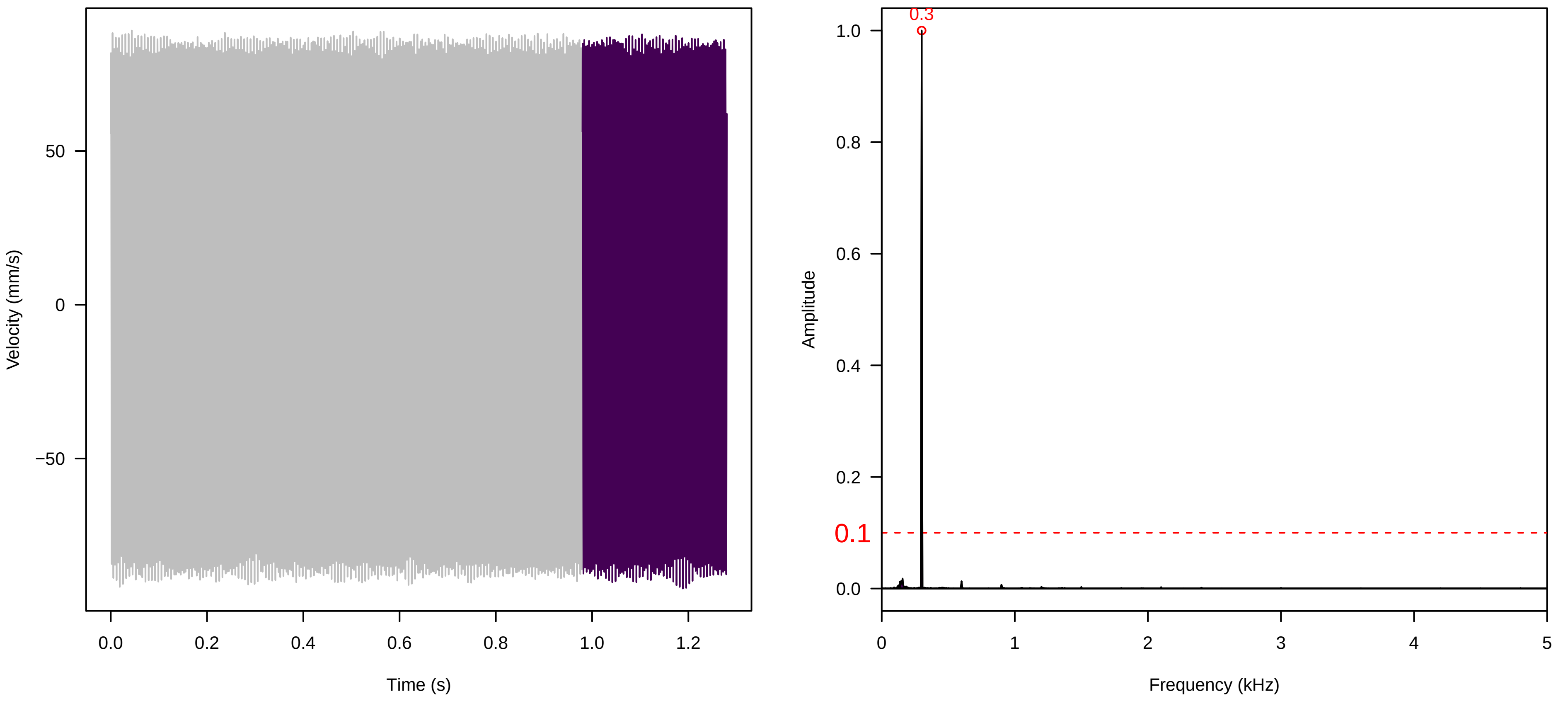

**Vel. = 0.057 ; Str. = Corolla ; Axis = z ; Fl. accession = 10−s−81−1AA**

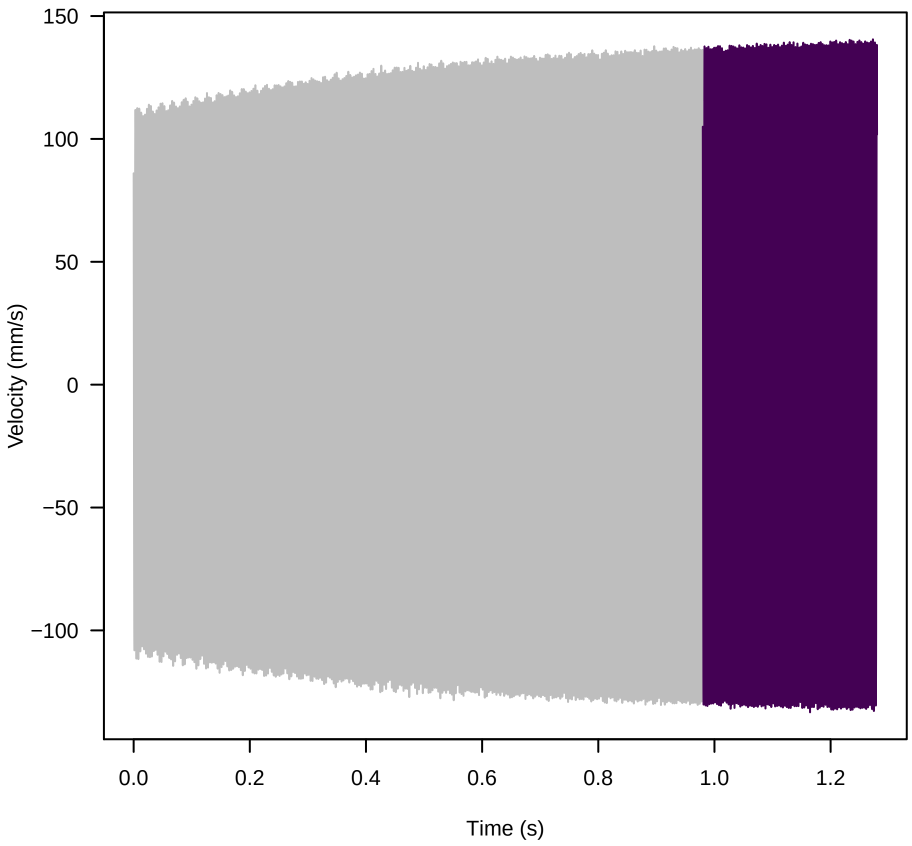


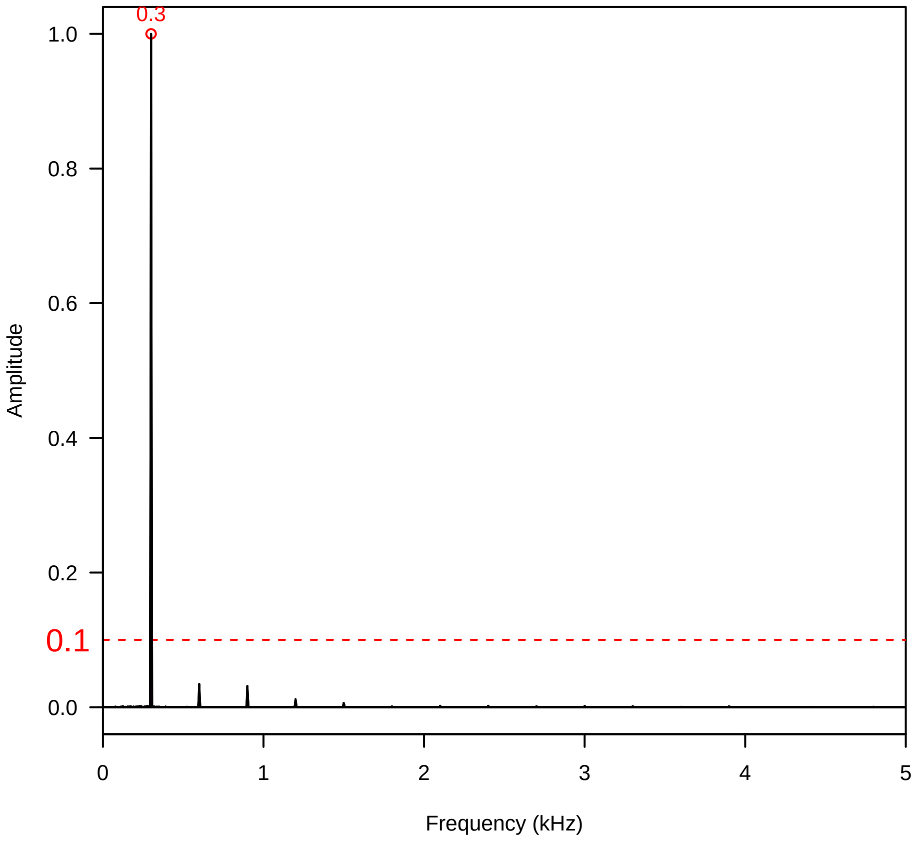


**Vel. = 0.057 ; Str. = Receptacle ; Axis = z ; Fl. accession = 10−s−81−1AA**

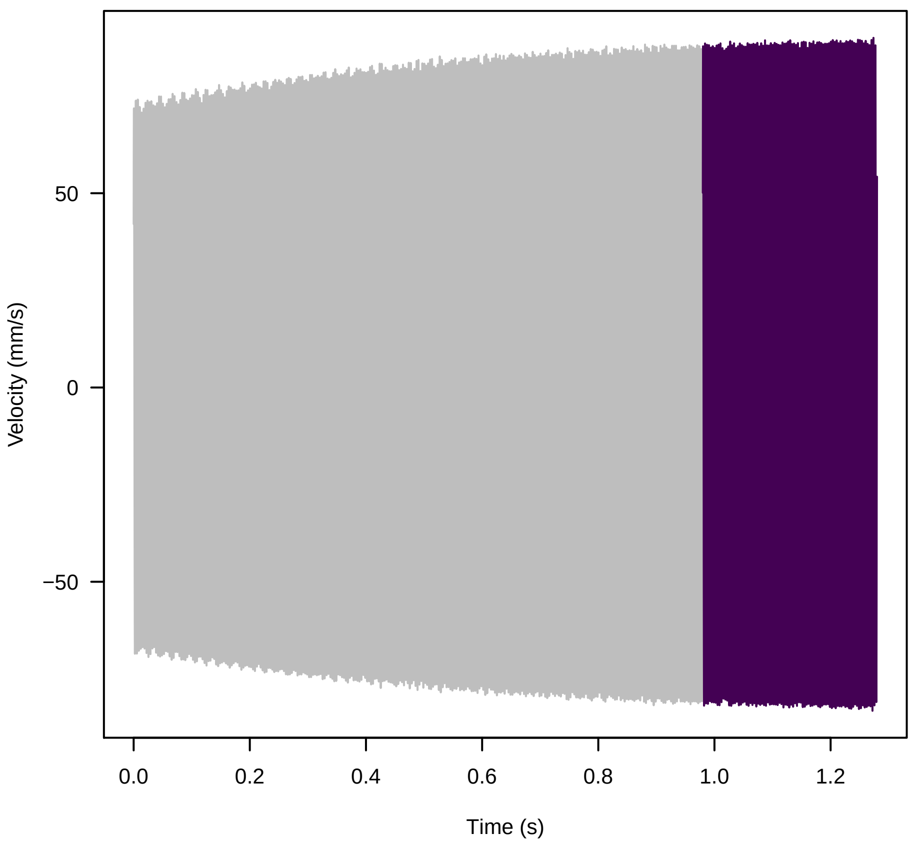


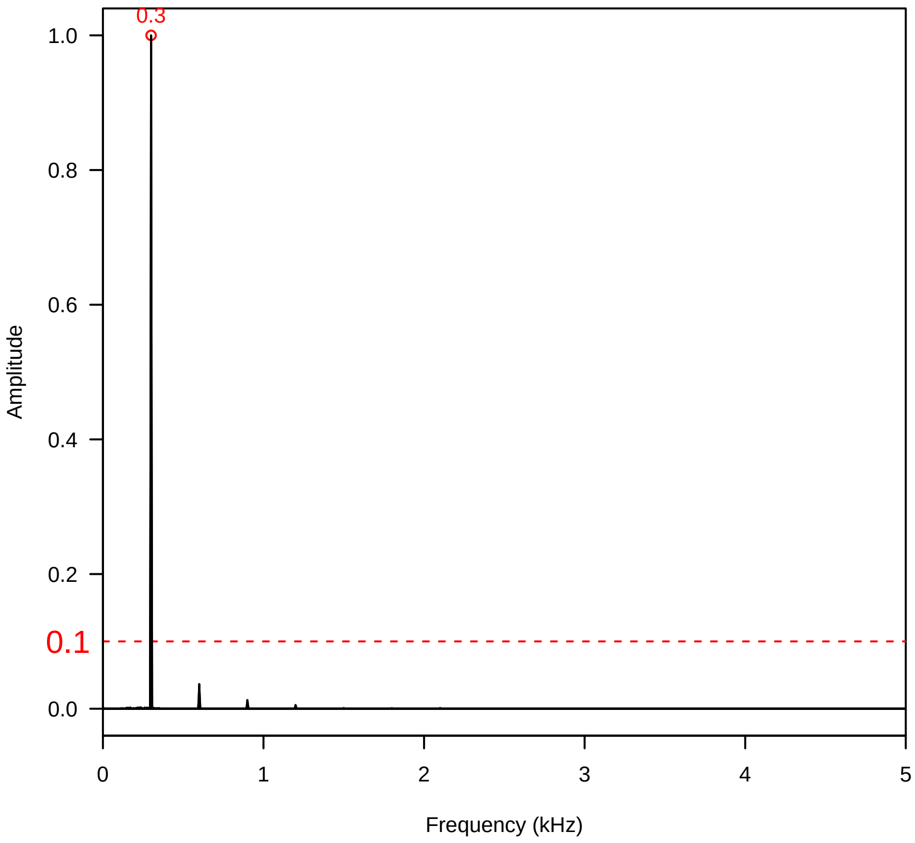


**Vel. = 0.057 ; Str. = FA ; Axis = z ; Fl. accession = 10−s−81−1AA**

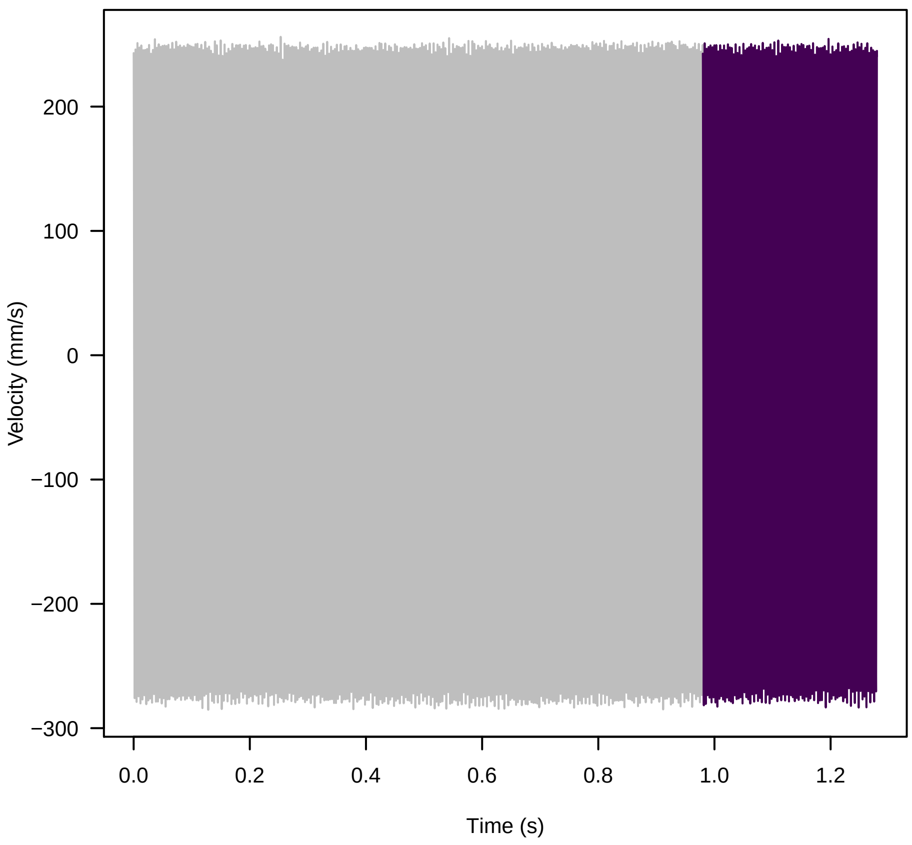


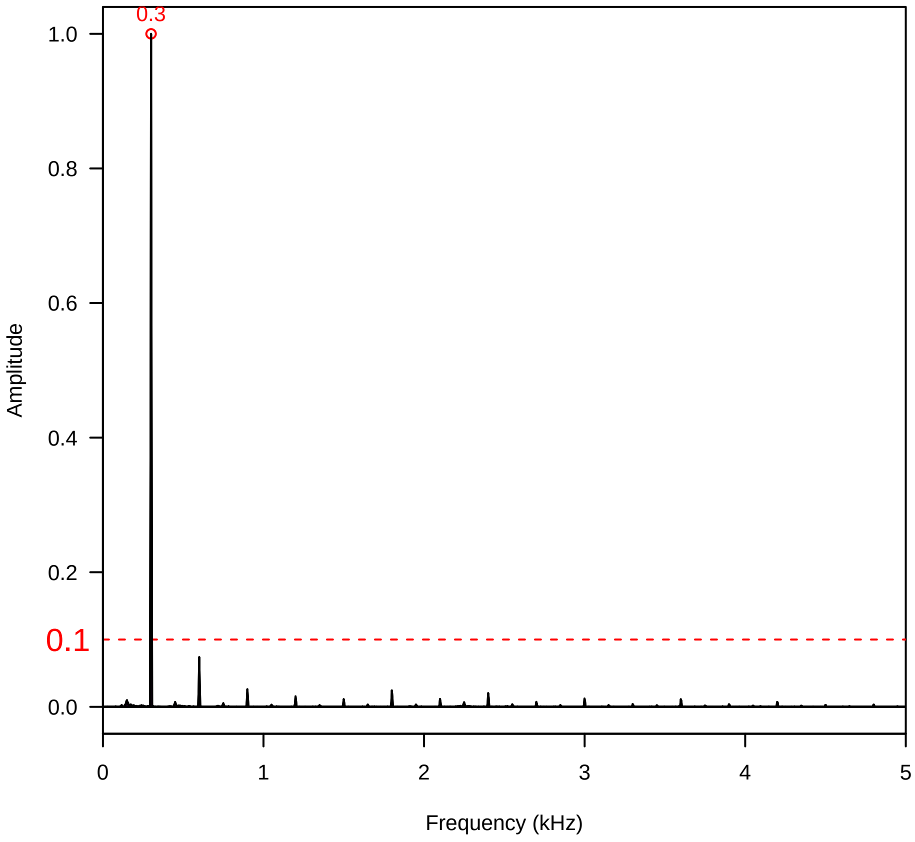

**Vel. = 0.057 ; Str. = Receptacle ; Axis = z ; Fl. accession = 10−s−81−1AA**

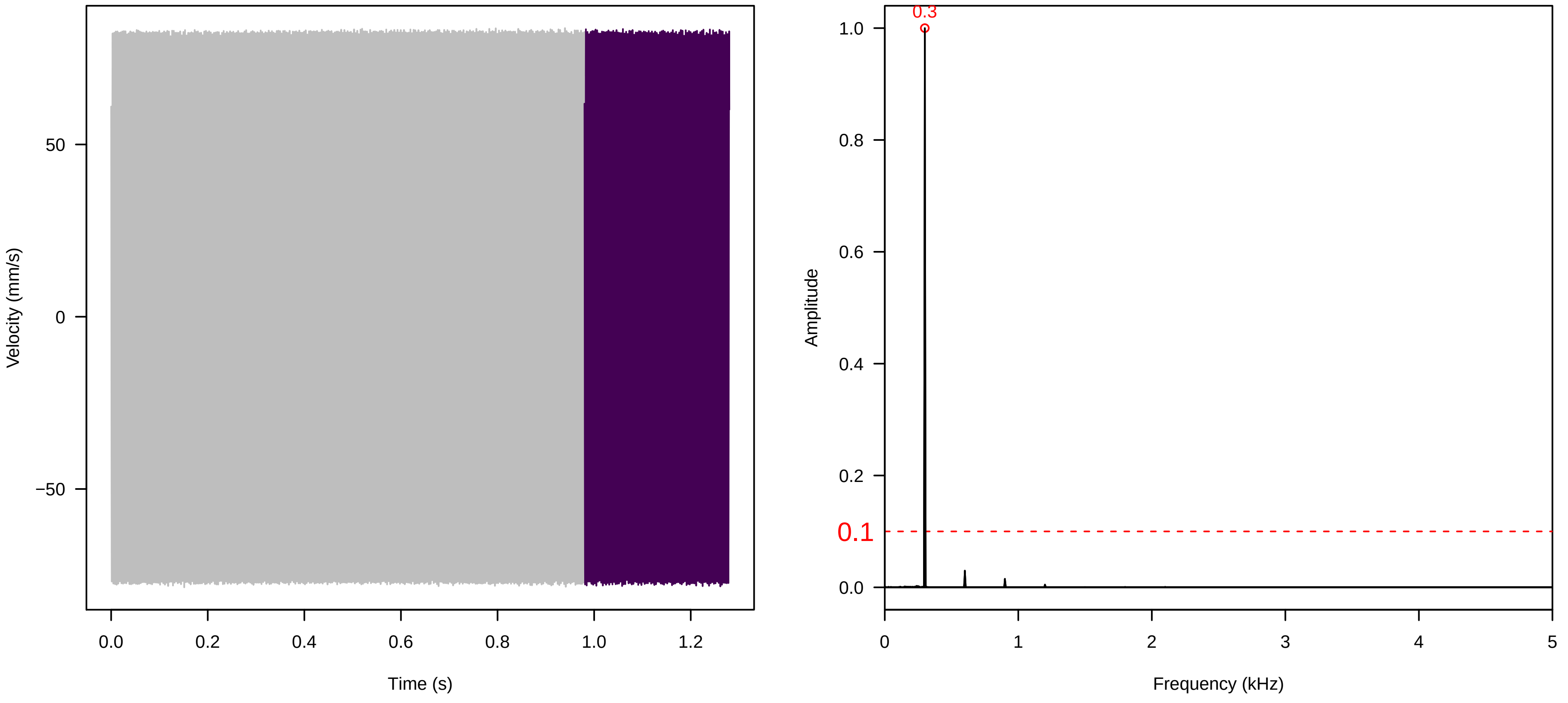


**Vel. = 0.057 ; Str. = PA ; Axis = z ; Fl. accession = 10−s−81−1AA**

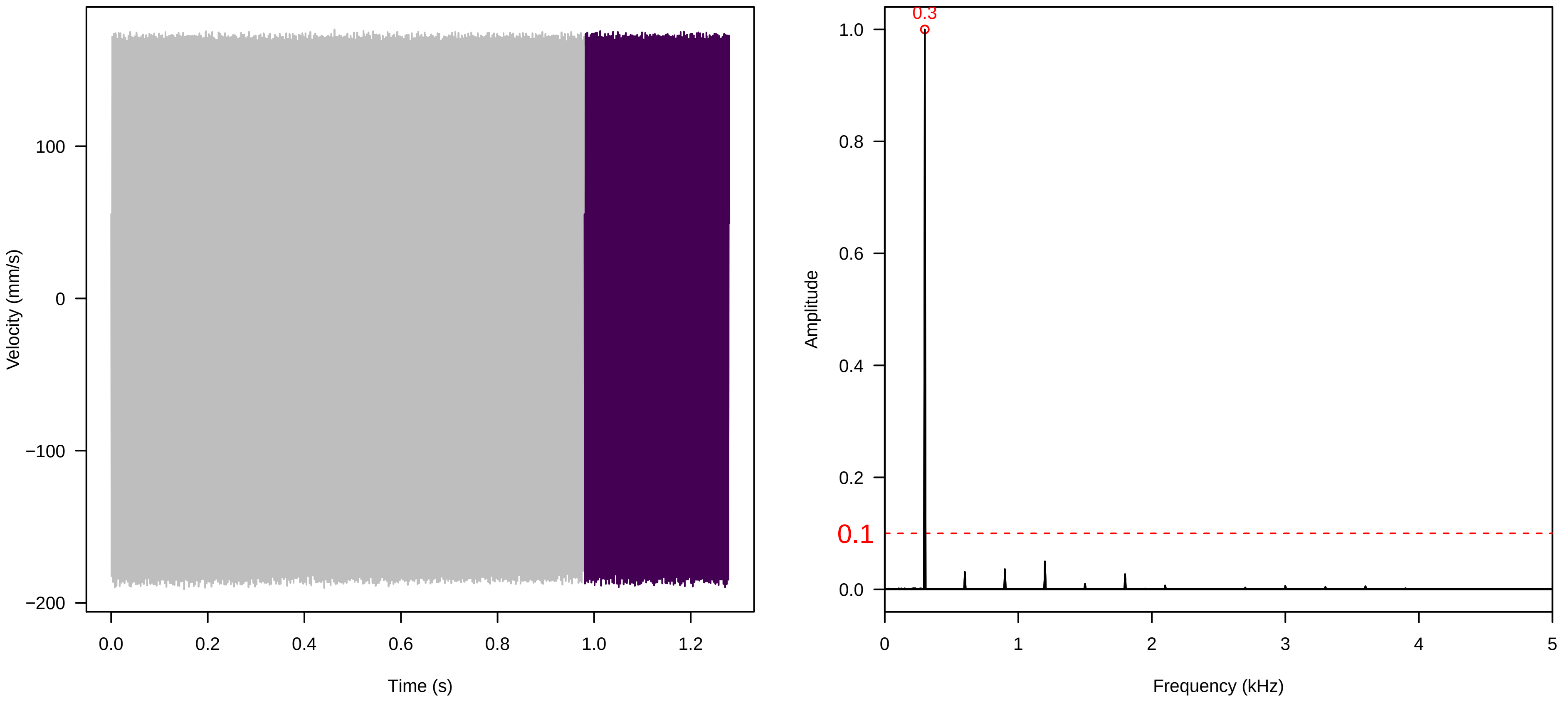


**Vel. = 0.057 ; Str. = Receptacle ; Axis = z ; Fl. accession = 10−s−81−1AA**

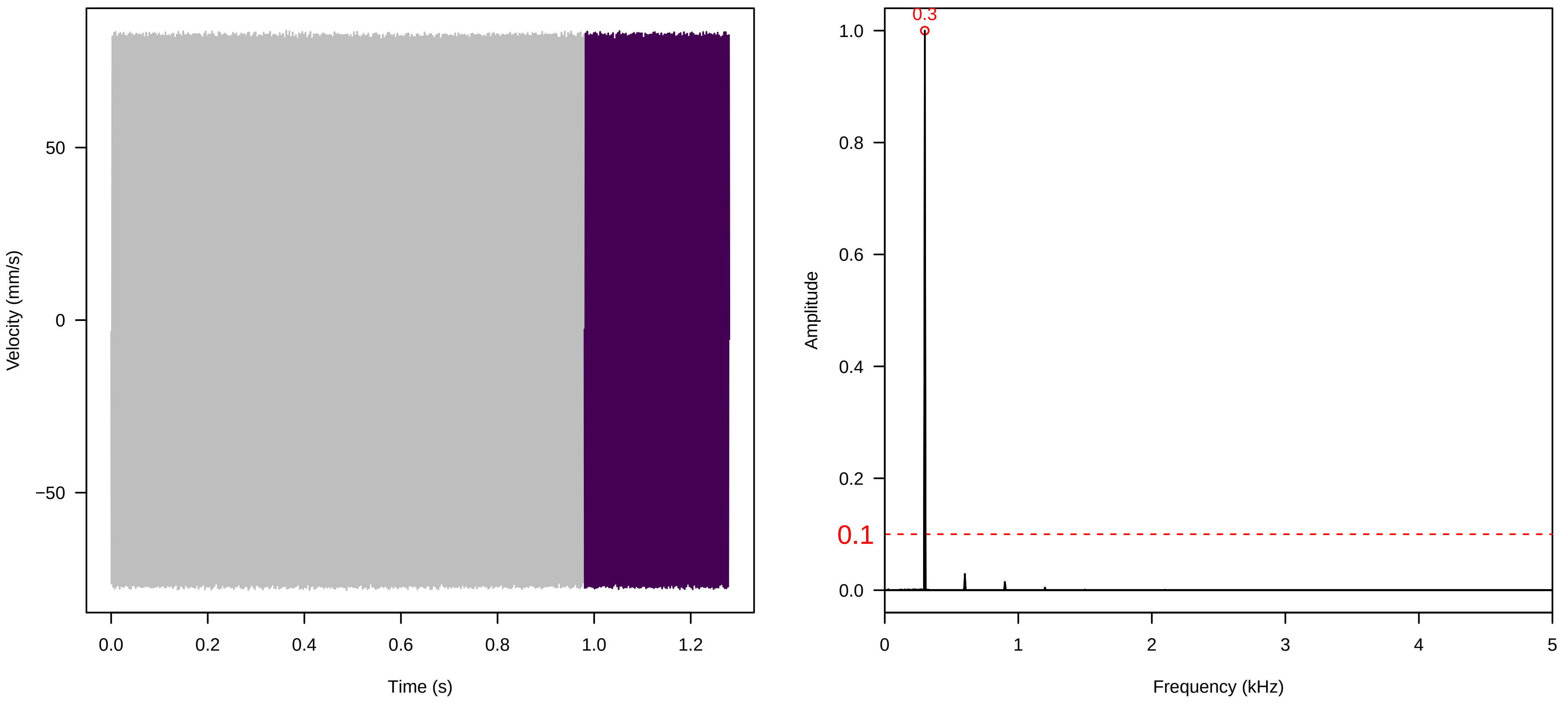

**Vel. = 0.028 ; Str. = PA ; Axis = z ; Fl. accession = 10−s−81−1AA**

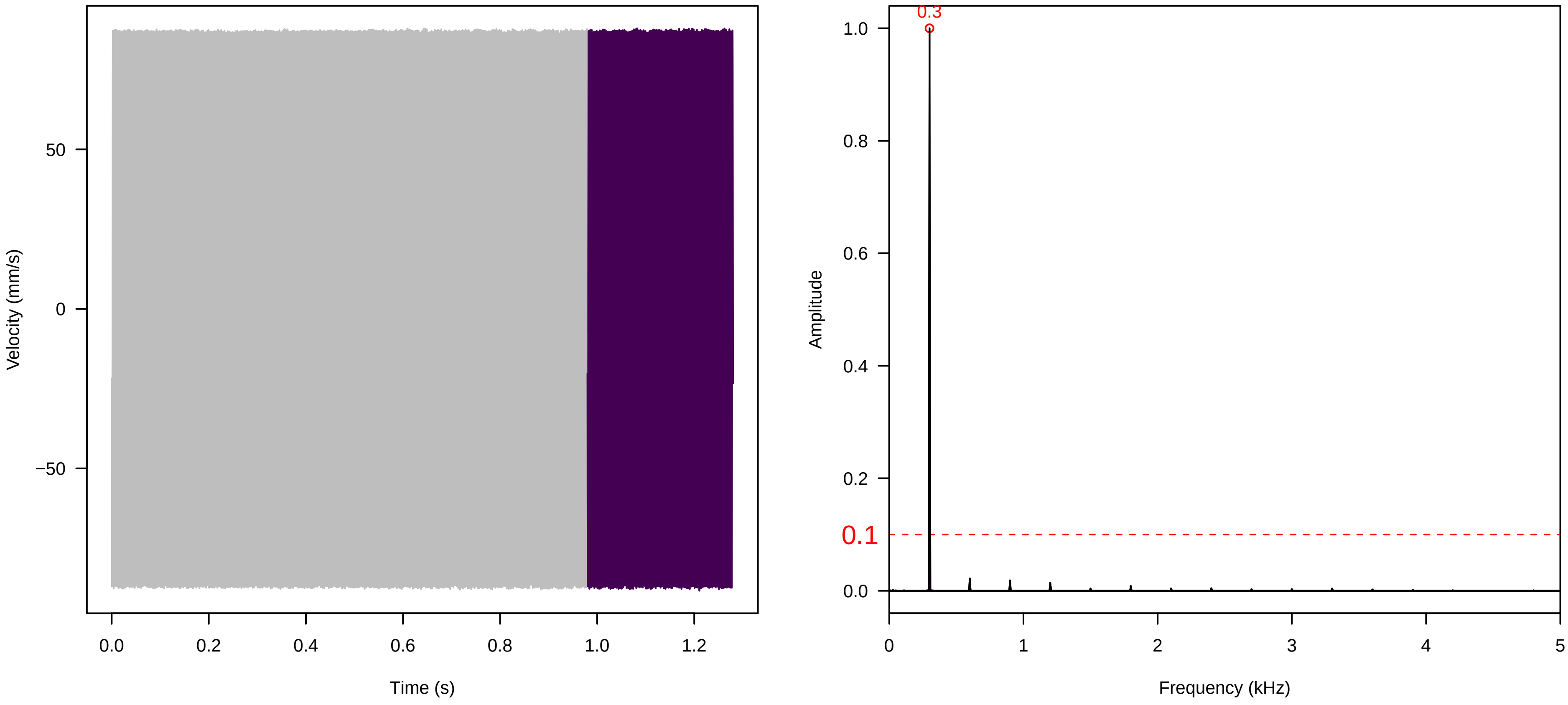


**Vel. = 0.028 ; Str. = Receptacle ; Axis = z ; Fl. accession = 10−s−81−1AA**

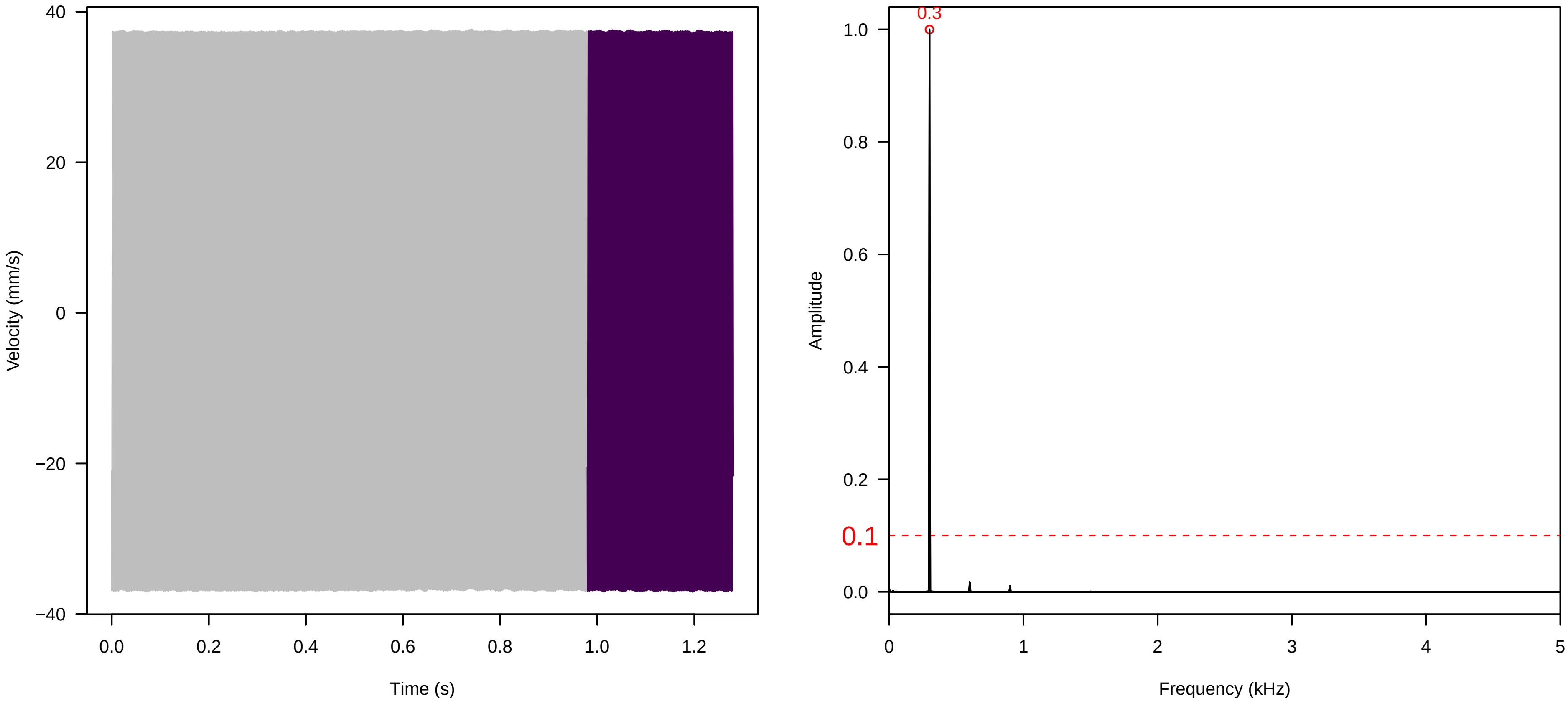


**Vel. = 0.028 ; Str. = FA ; Axis = z ; Fl. accession = 10−s−81−1AA**

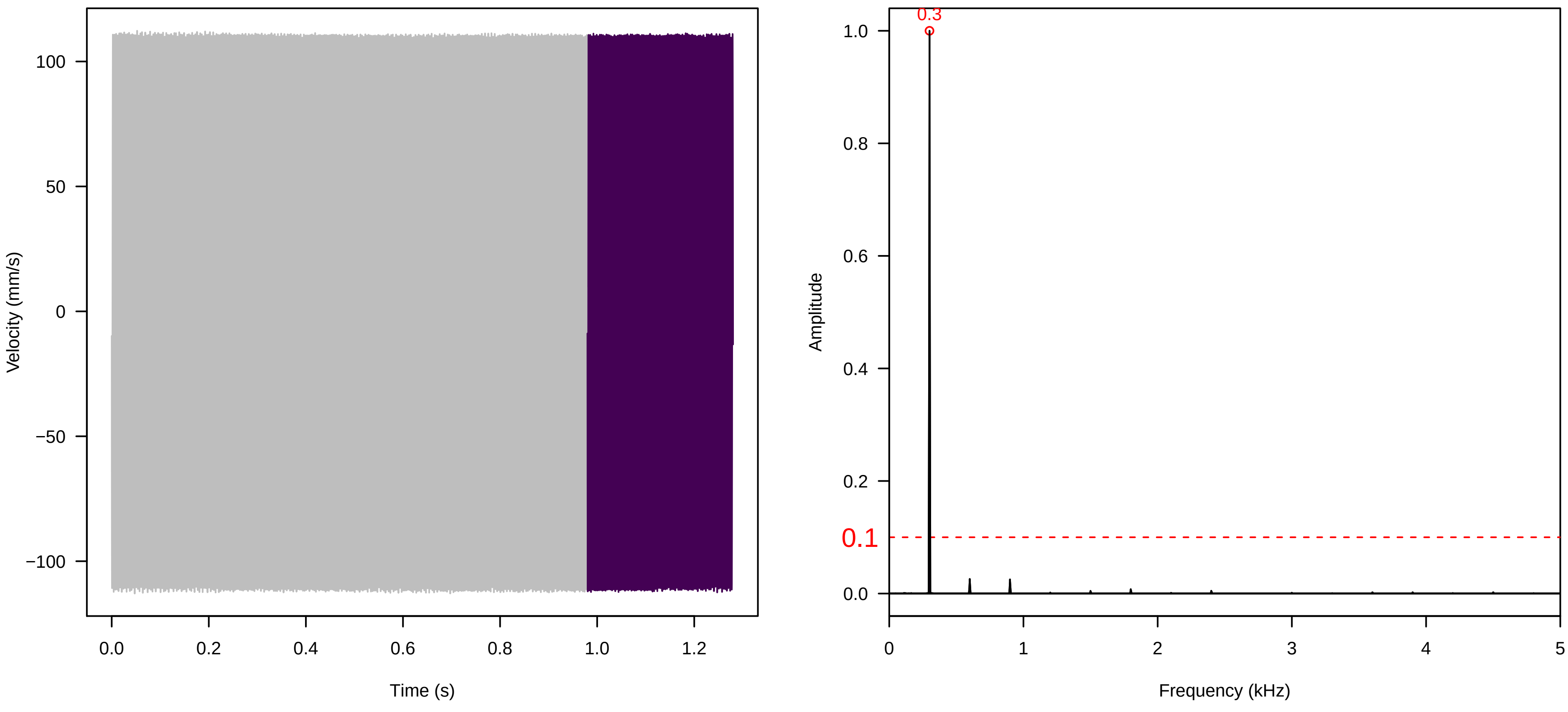

**Vel. = 0.028 ; Str. = Receptacle ; Axis = z ; Fl. accession = 10−s−81−1AA**

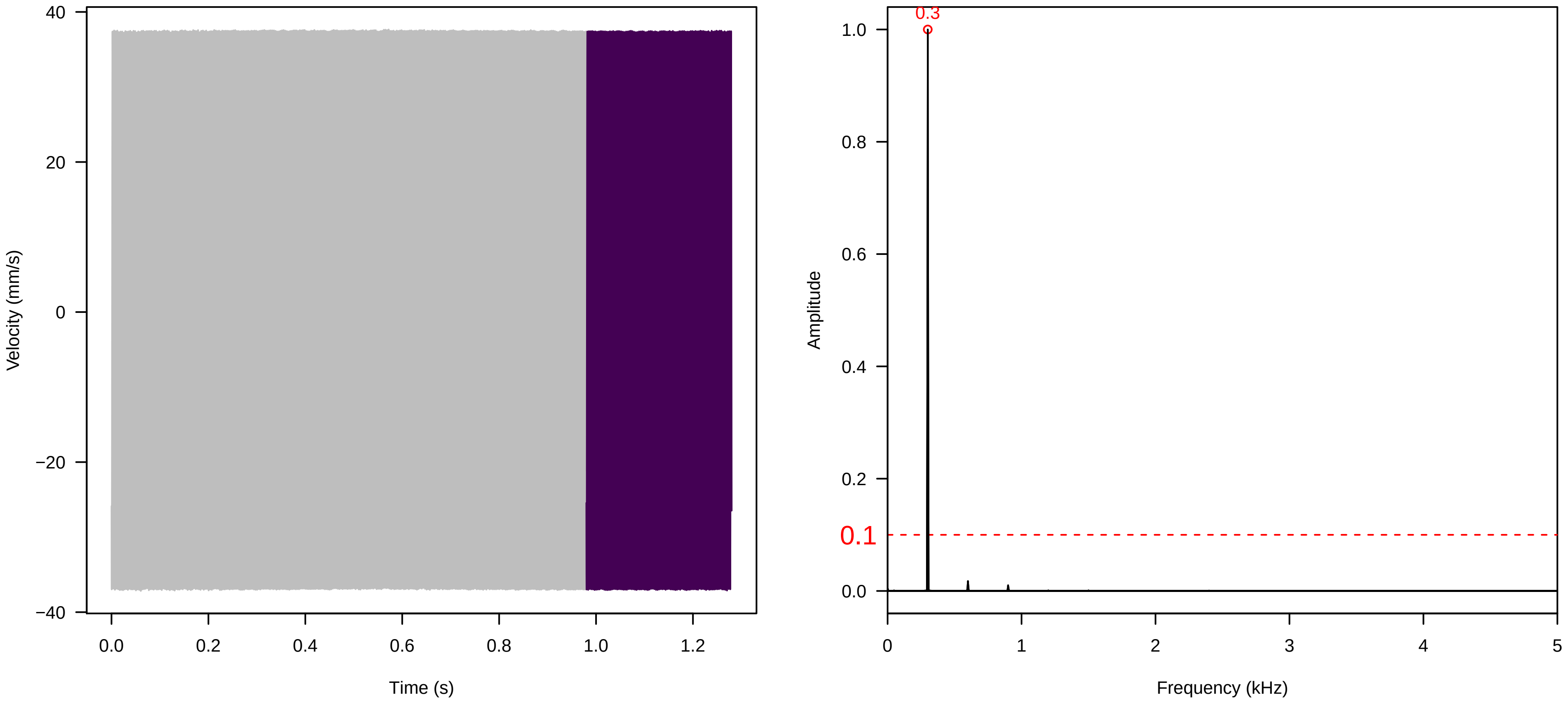


**Vel. = 0.028 ; Str. = Corolla ; Axis = z ; Fl. accession = 10−s−81−1AA**

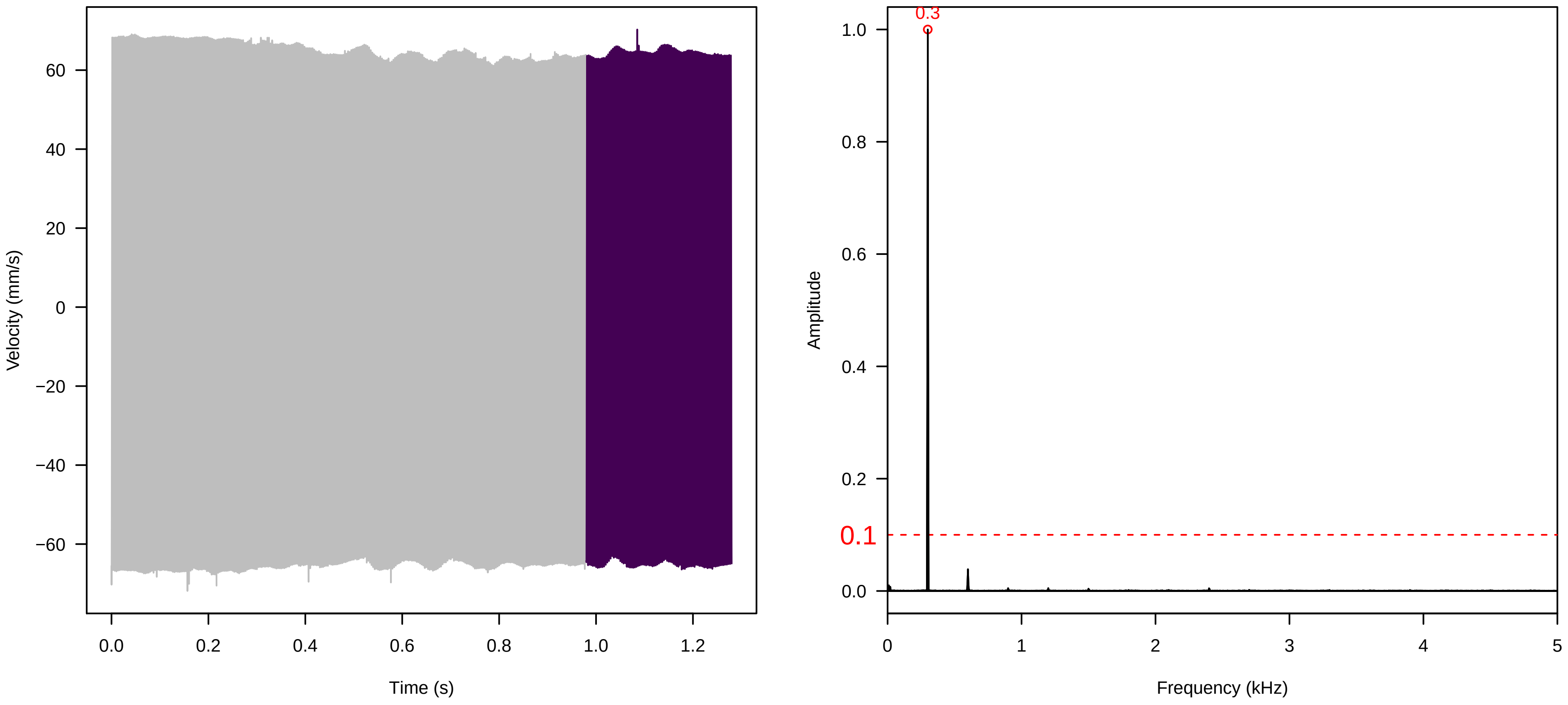


**Vel. = 0.028 ; Str. = Receptacle ; Axis = z ; Fl. accession = 10−s−81−1AA**

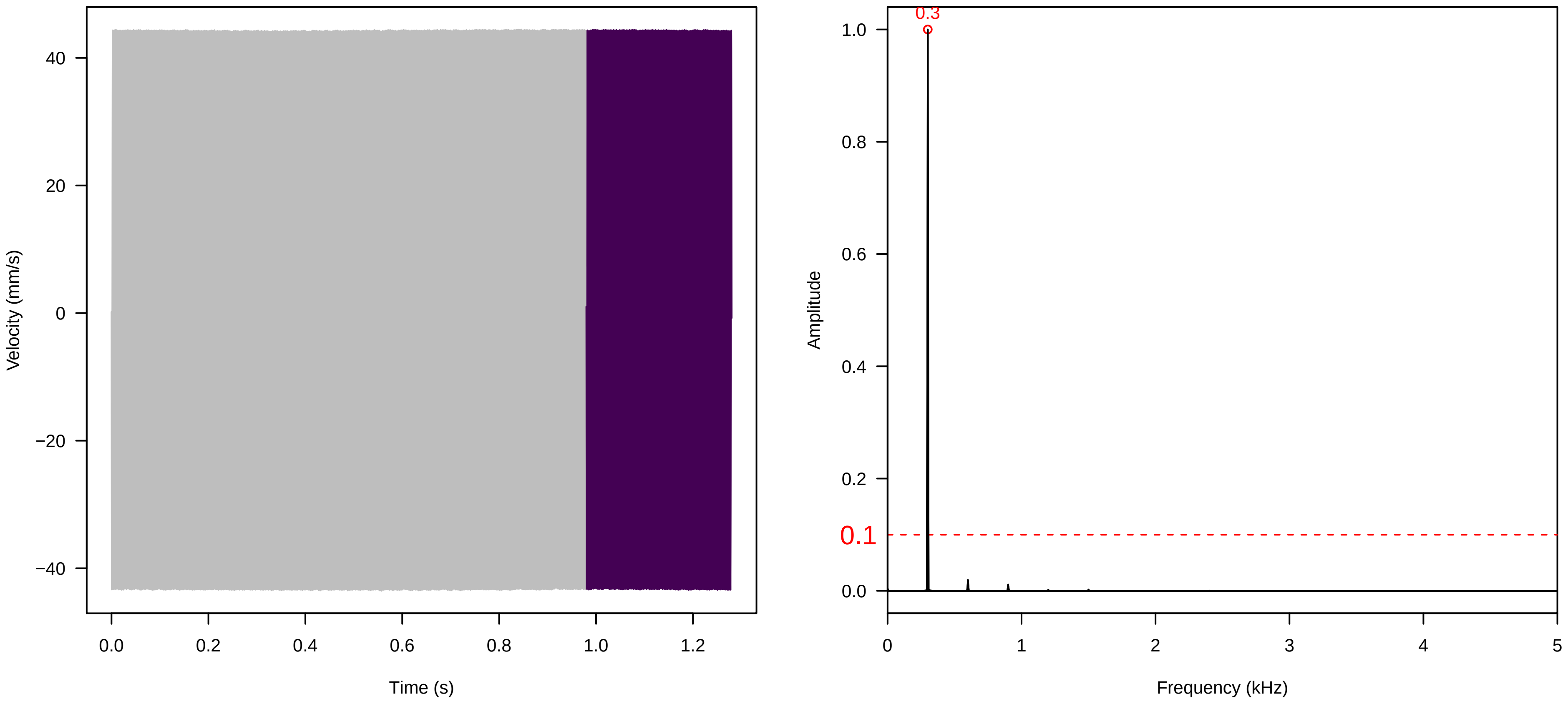

Vel. =  0.014 ; Str. =  Corolla ; Axis =  z ; Fl. accession = 10−s−81−1AA

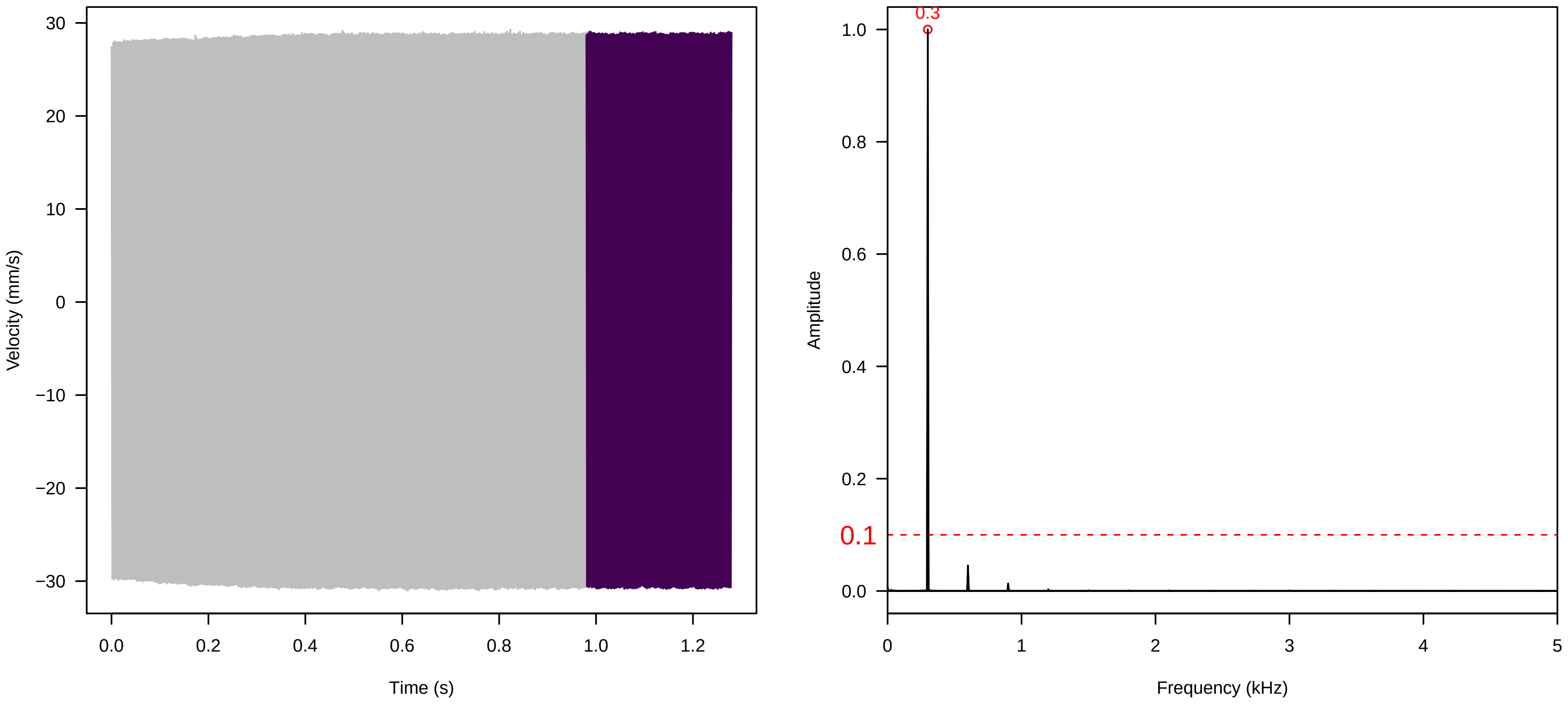


Vel. =  0.014 ; Str. =  Receptacle ; Axis =  z ; Fl. accession = 10−s−81−1AA

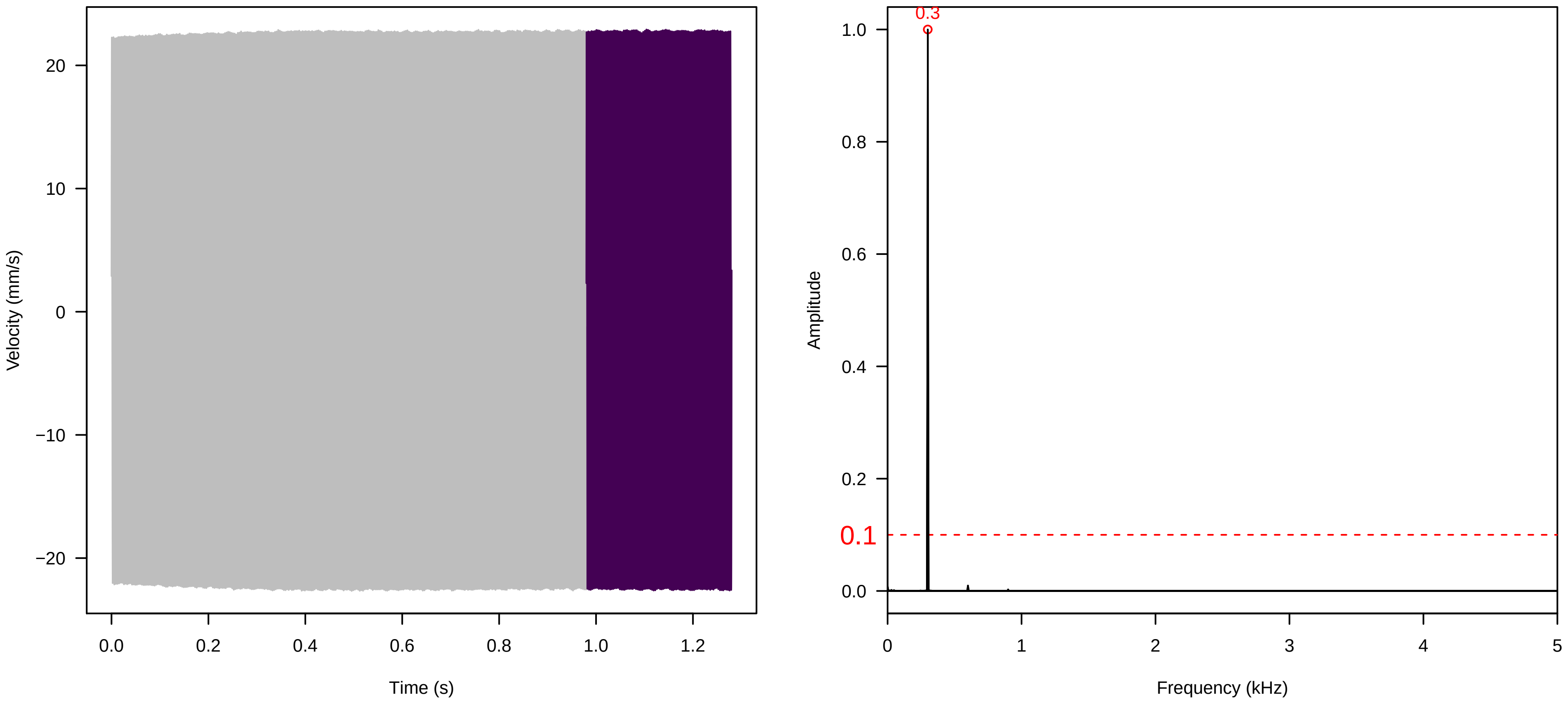


Vel. =  0.014 ; Str. =  FA ; Axis =  z ; Fl. accession = 10−s−81−1AA

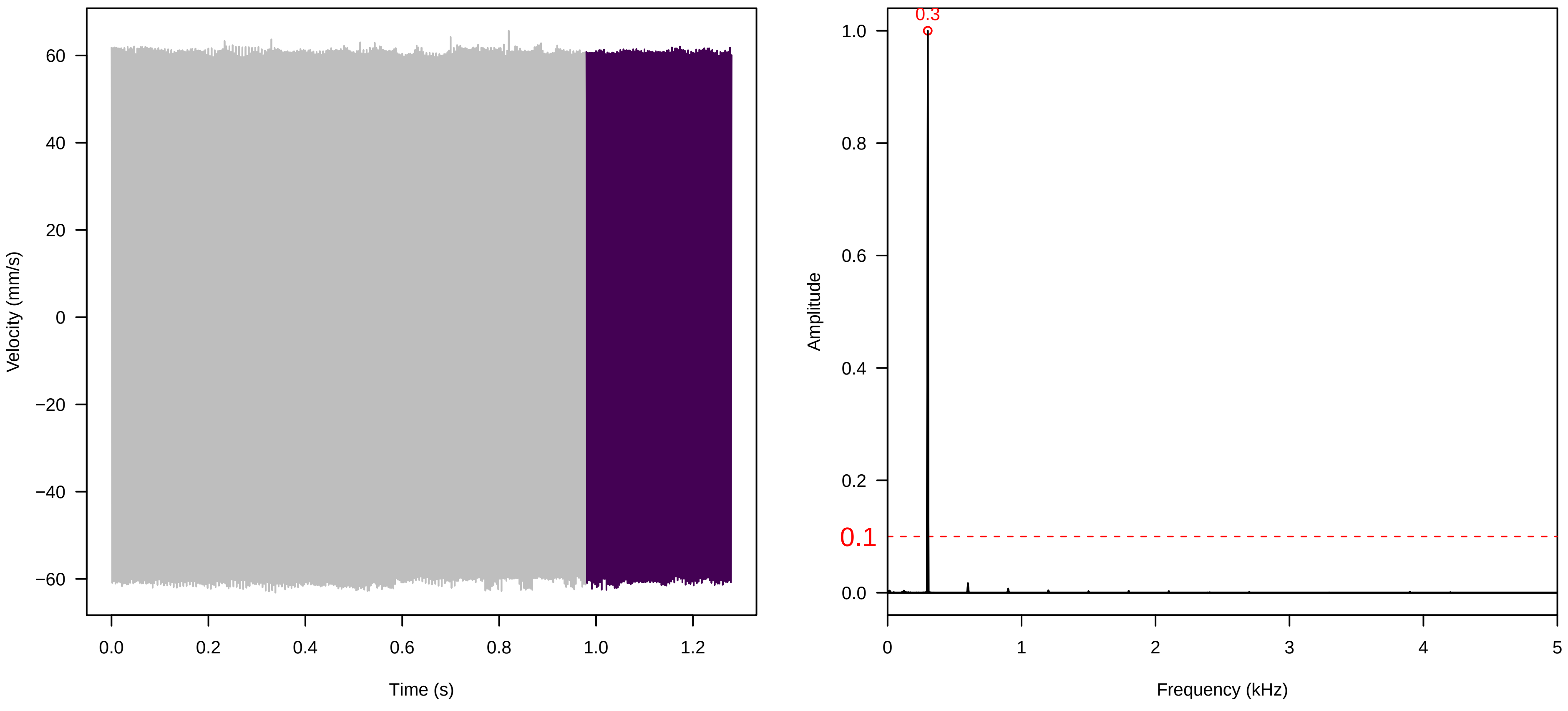

**Vel. =  0.014 ; Str. =  Receptacle ; Axis =  z ; Fl. accession = 10−s−81−1AA**

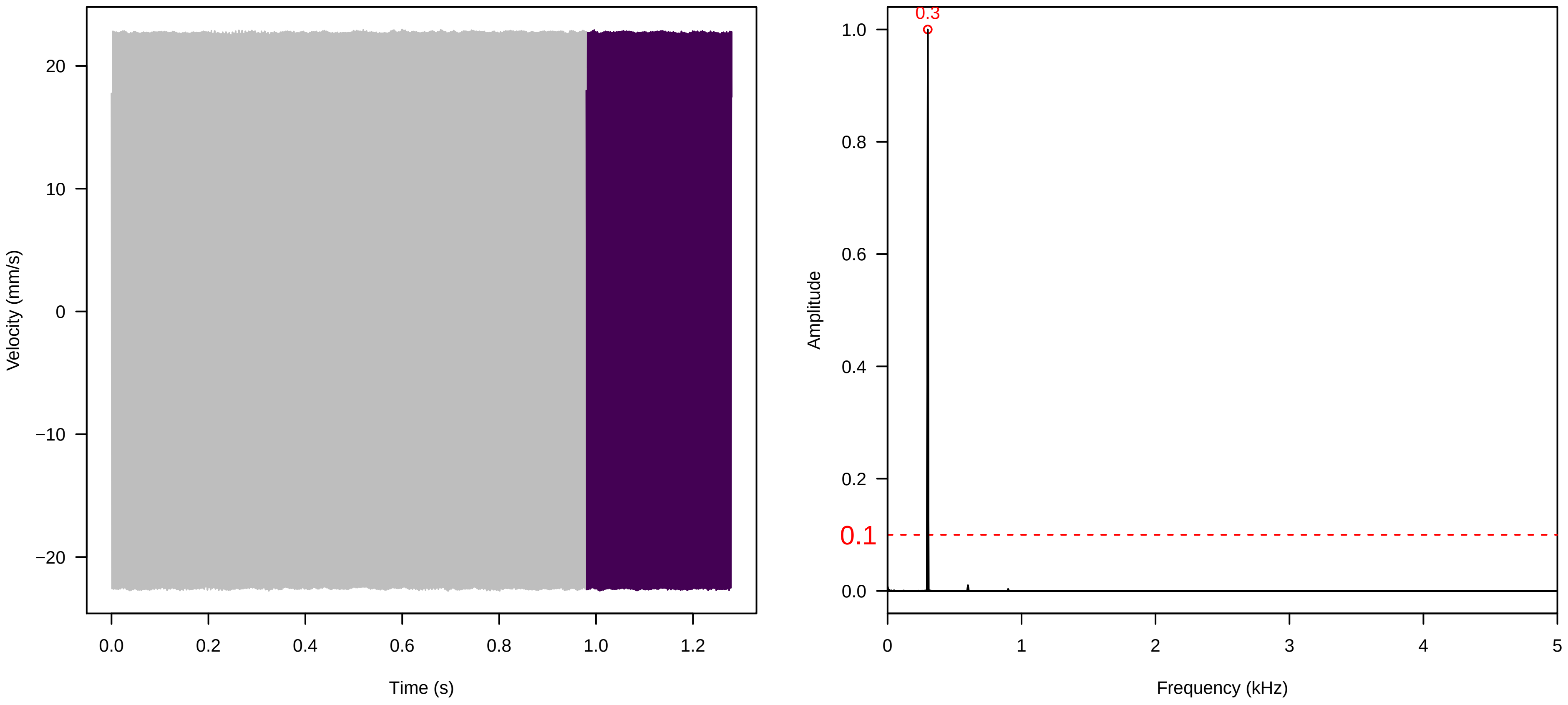


**Vel. =  0.014 ; Str. =  PA ; Axis =  z ; Fl. accession = 10−s−81−1AA**

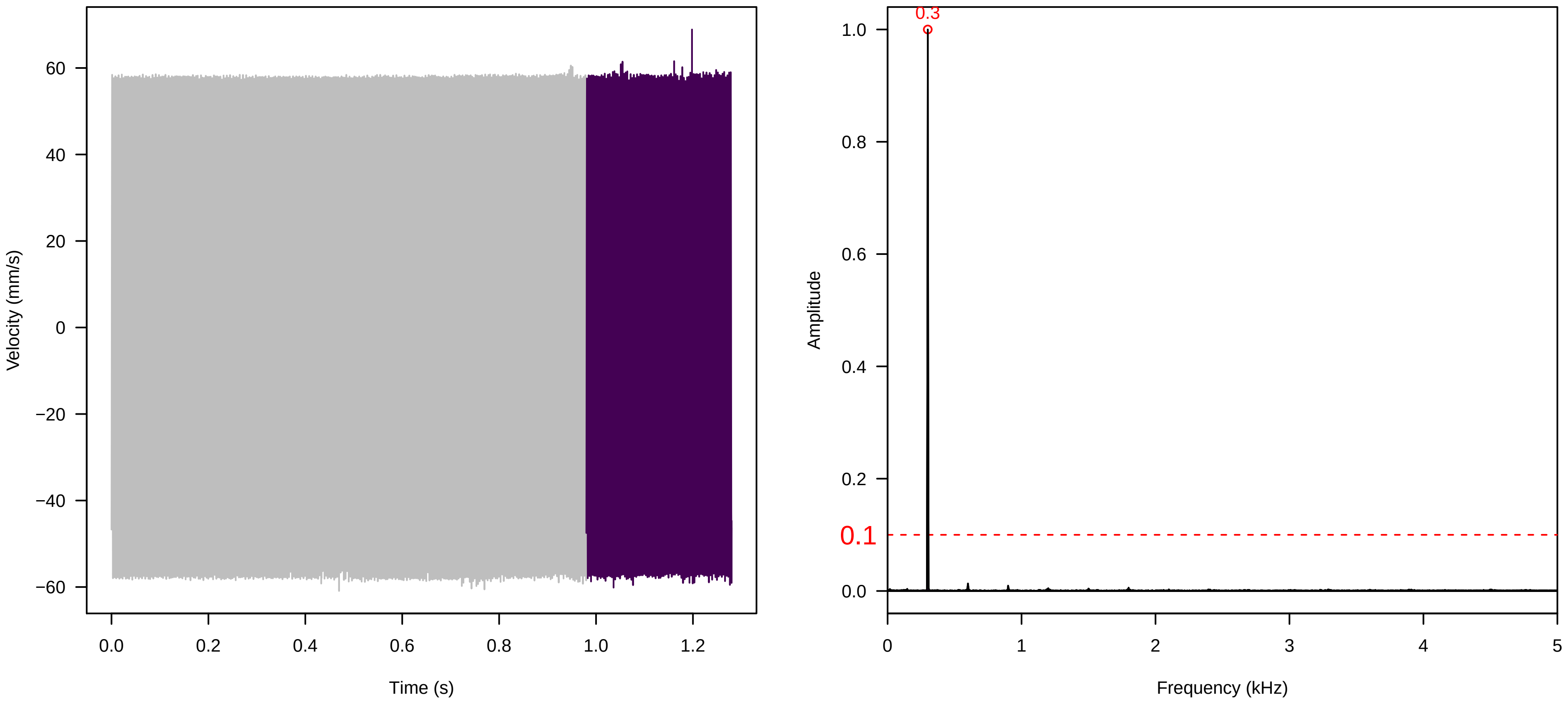


**Vel. =  0.014 ; Str. =  Receptacle ; Axis =  z ; Fl. accession = 10−s−81−1AA**

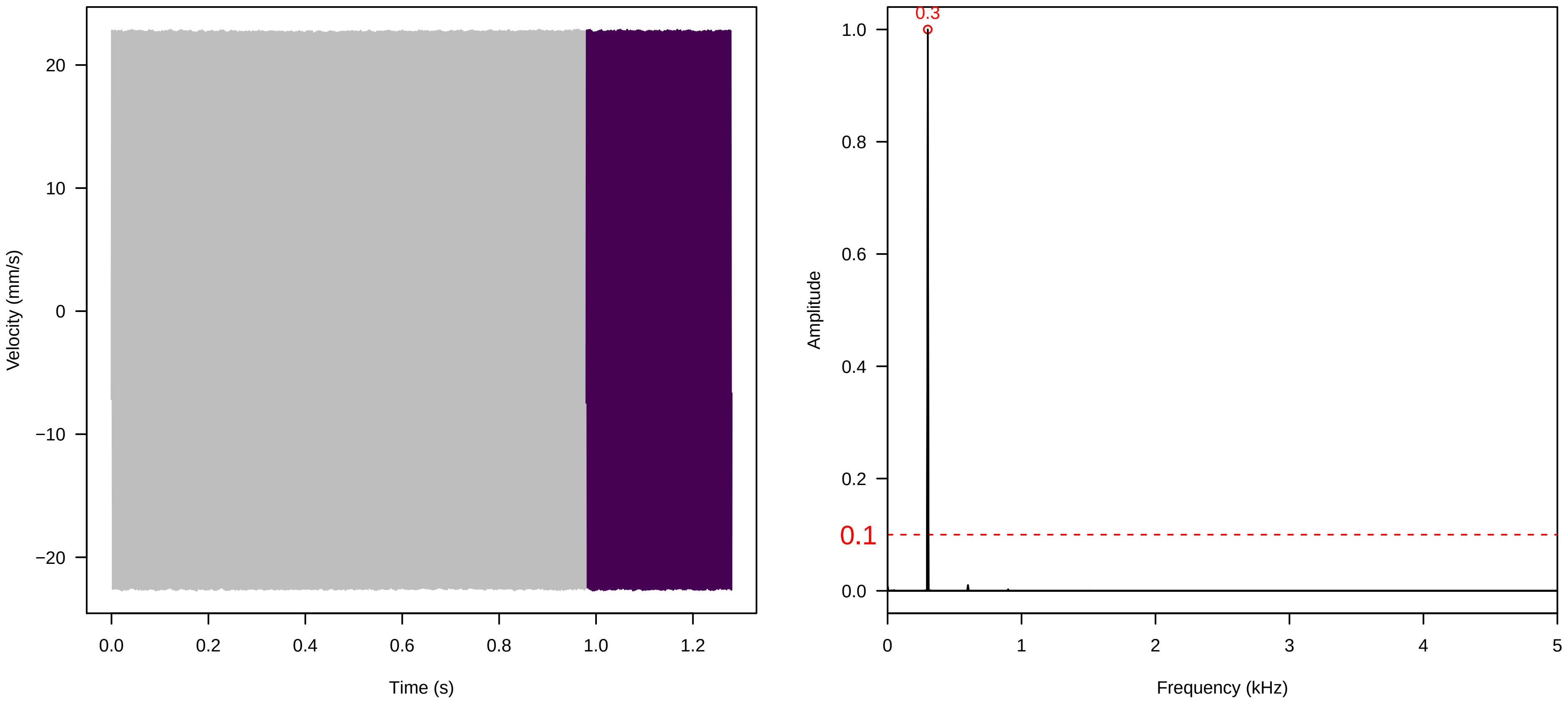

Vel. =  0.014 ; Str. =  Corolla ; Axis =  y ; Fl. accession = 10−s−81−1AA

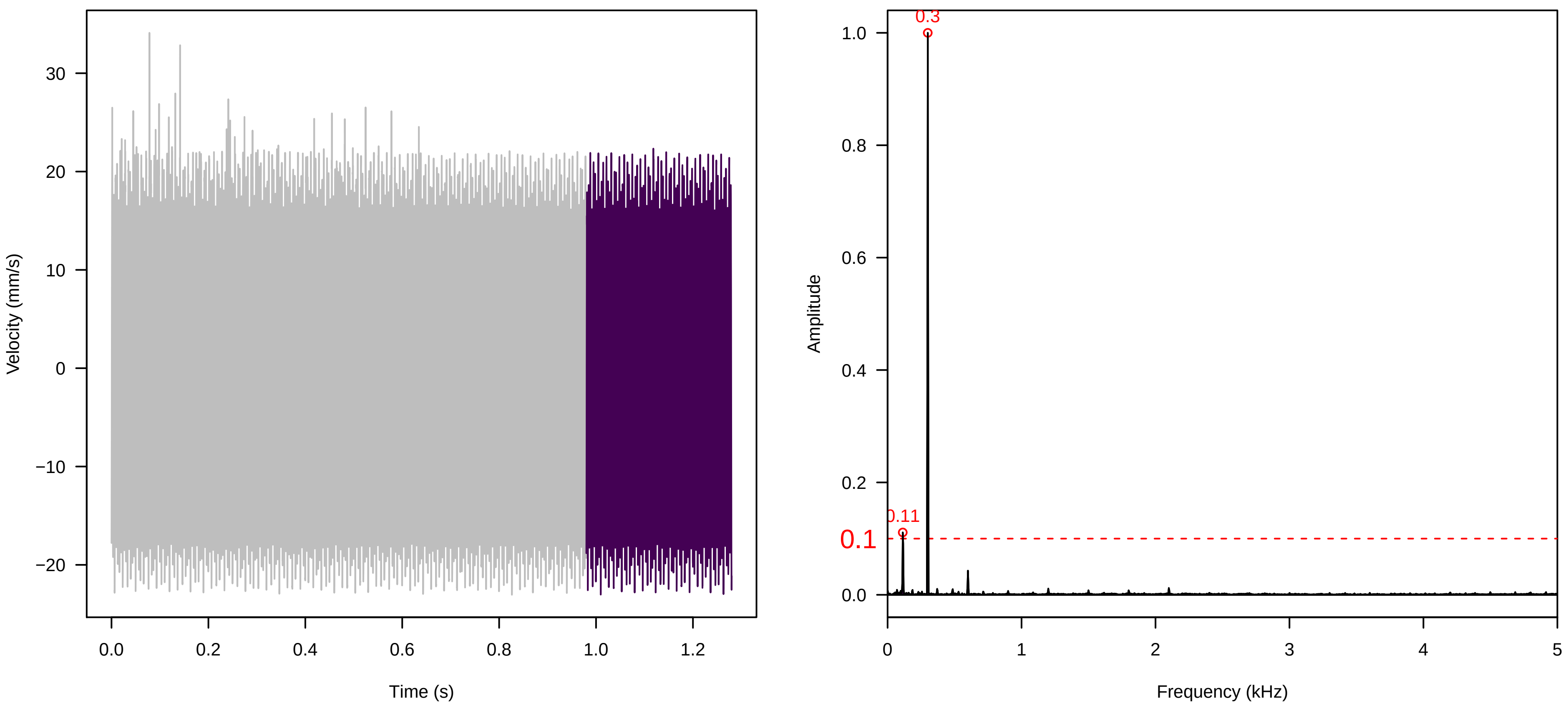


Vel. =  0.014 ; Str. =  Receptacle ; Axis =  y ; Fl. accession = 10−s−81−1AA

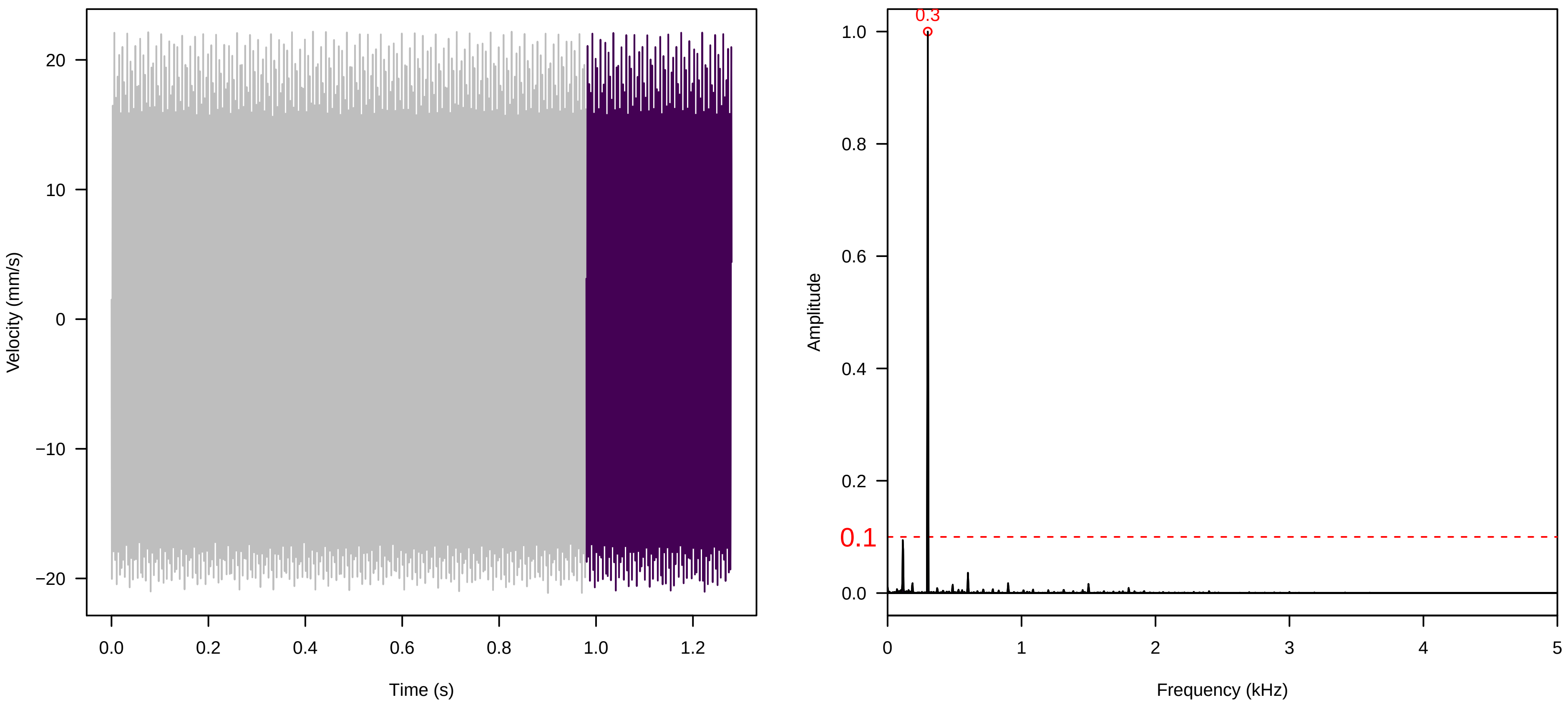


Vel. =  0.014 ; Str. =  FA ; Axis =  y ; Fl. accession = 10−s−81−1AA

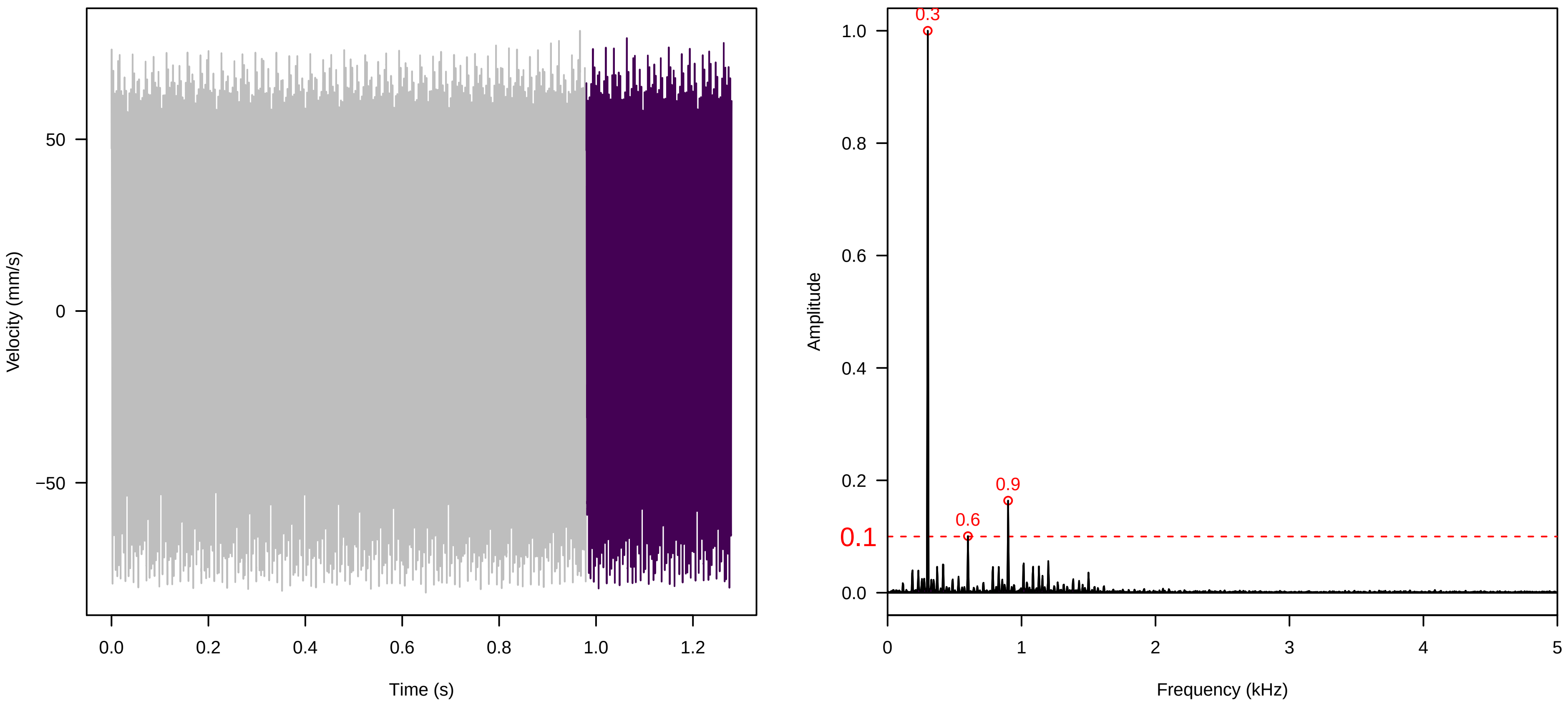

**Vel. =  0.014 ; Str. =  Receptacle ; Axis =  y ; Fl. accession = 10−s−81−1AA**

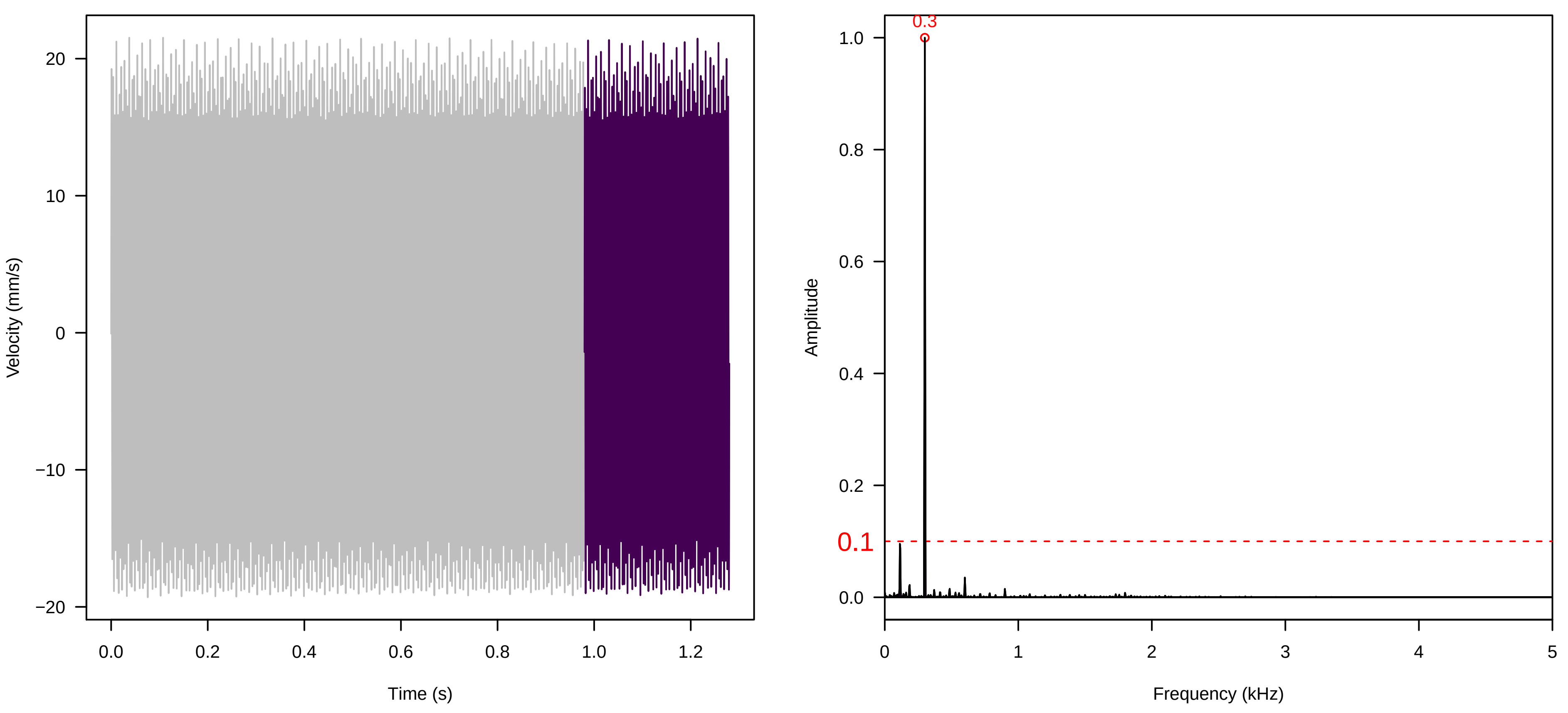


**Vel. =  0.014 ; Str. =  PA ; Axis =  y ; Fl. accession = 10−s−81−1AA**

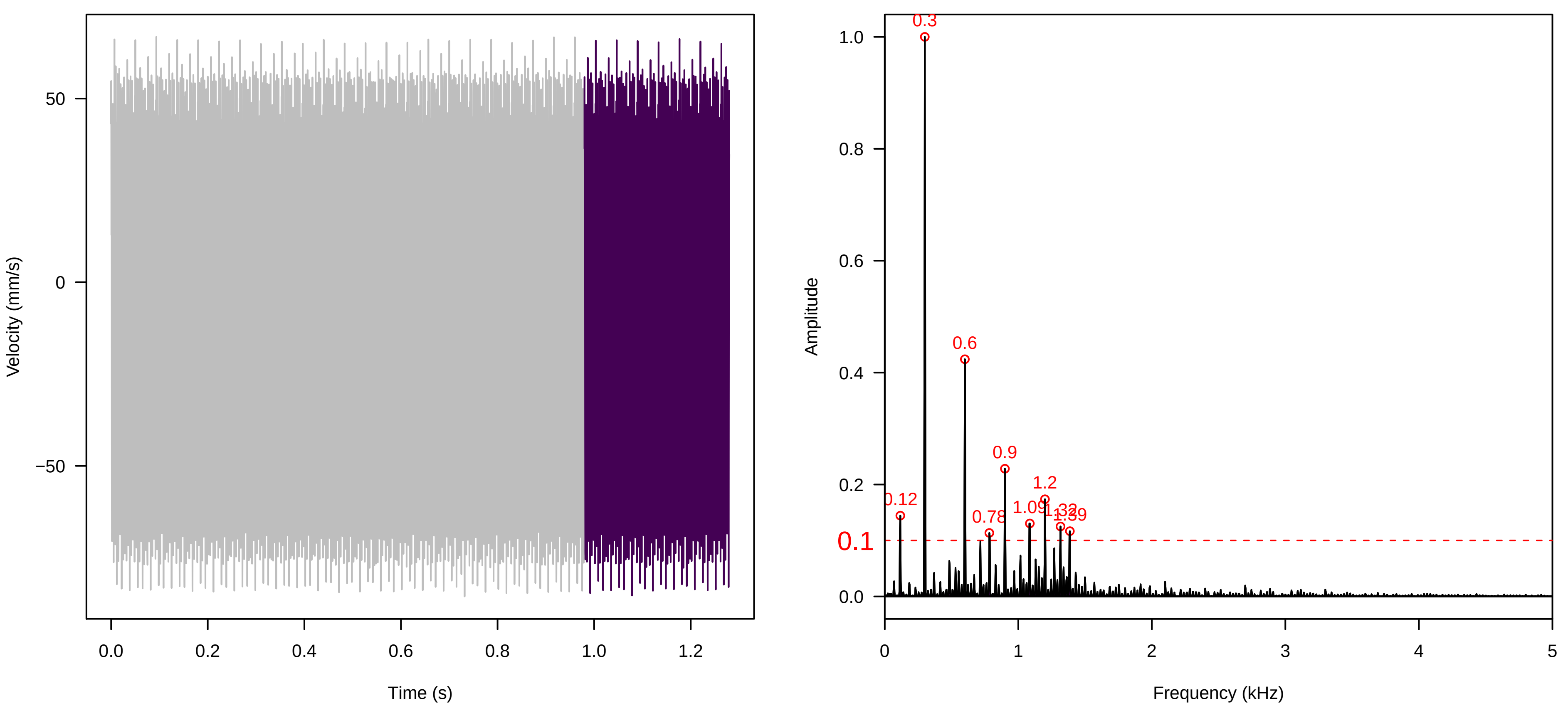


**Vel. =  0.014 ; Str. =  Receptacle ; Axis =  y ; Fl. accession = 10−s−81−1AA**

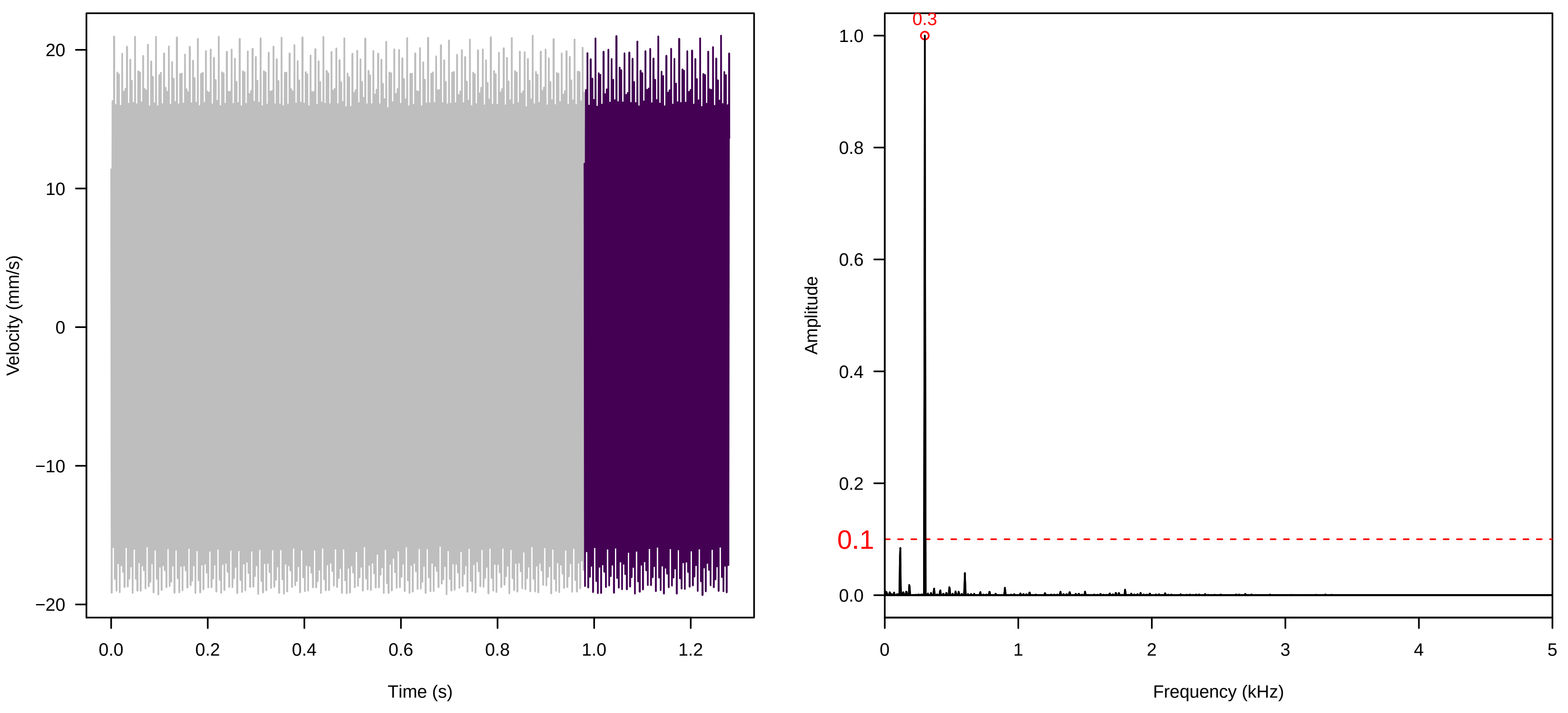

**Vel. = 0.028 ; Str. = PA ; Axis = y ; Fl. accession = 10−s−81−1AA**

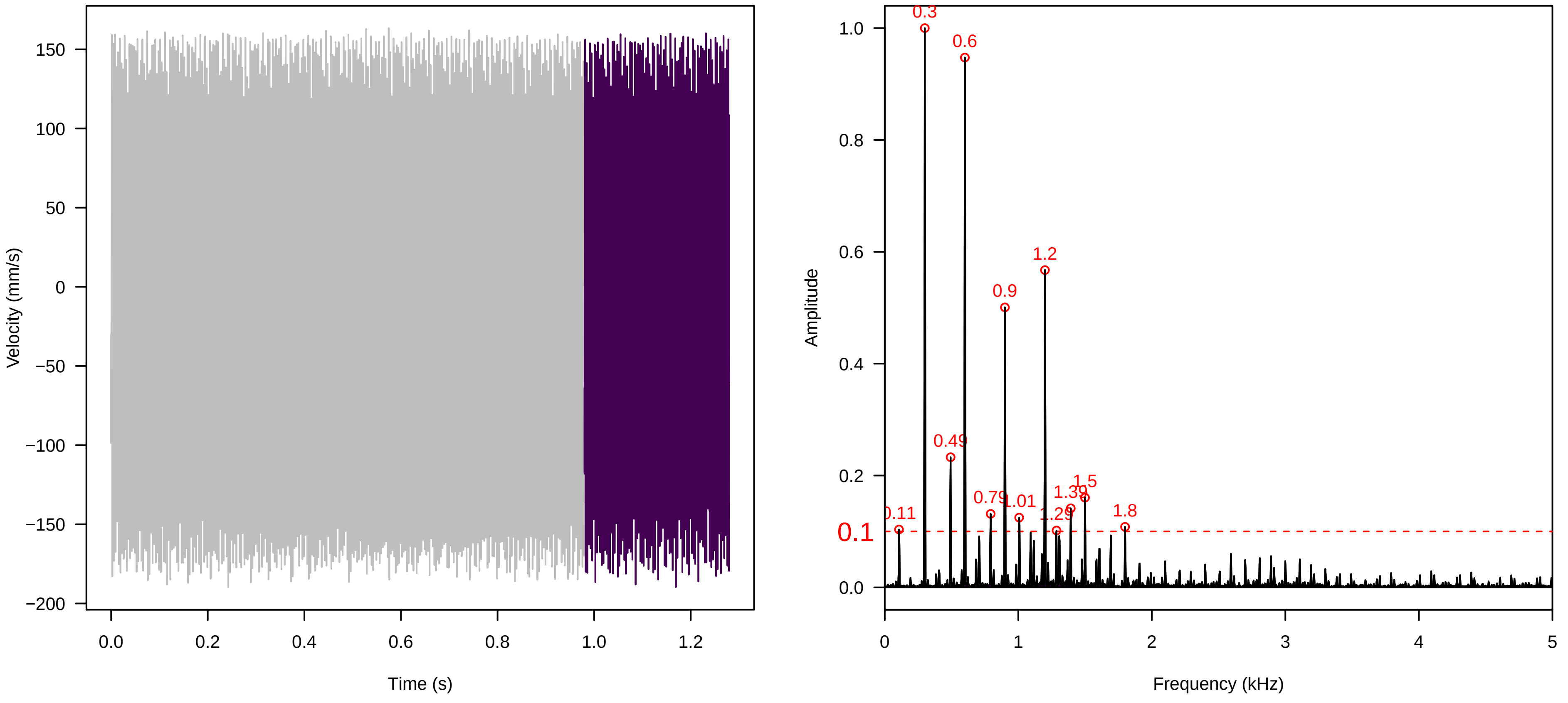


**Vel. = 0.028 ; Str. = Receptacle ; Axis = y ; Fl. accession = 10−s−81−1AA**

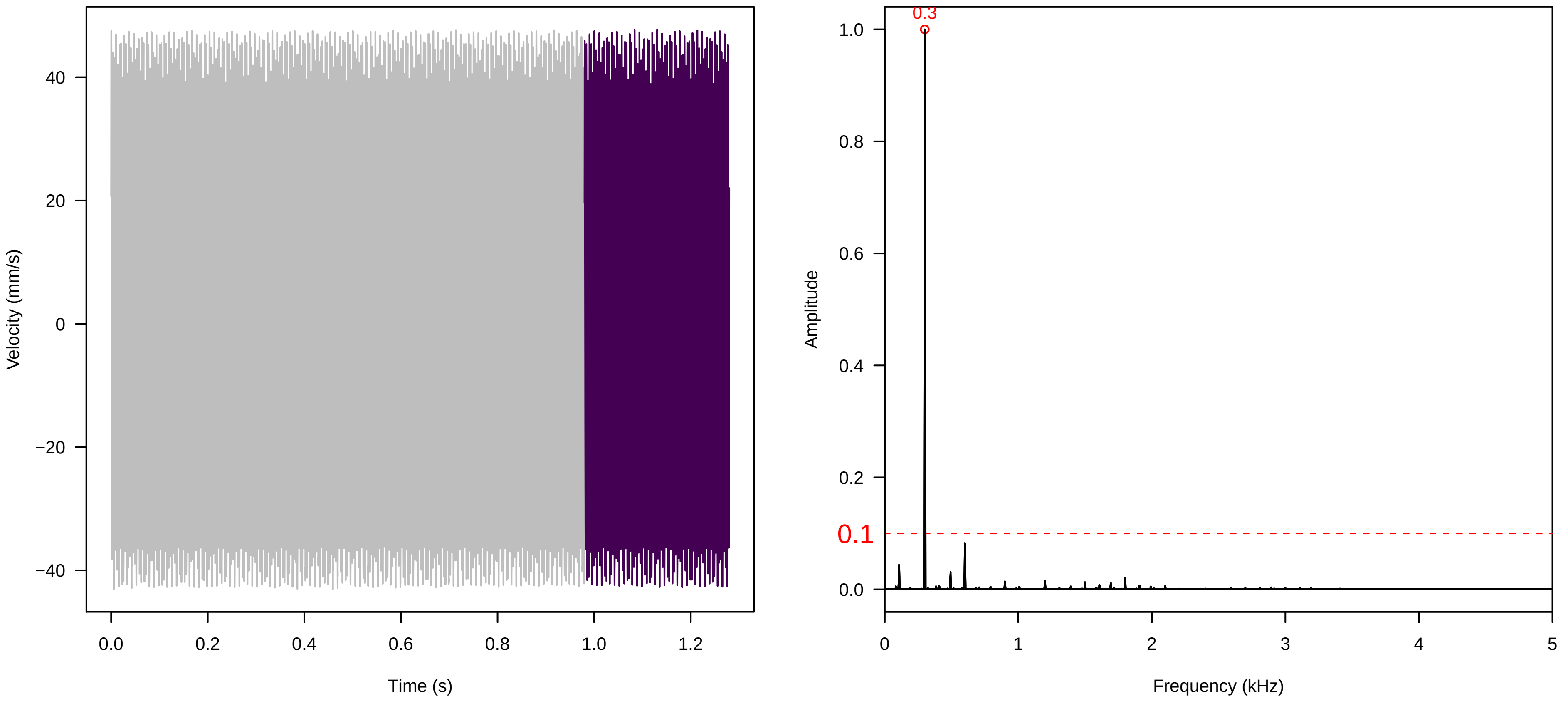


**Vel. = 0.028 ; Str. = FA ; Axis = y ; Fl. accession = 10−s−81−1AA**

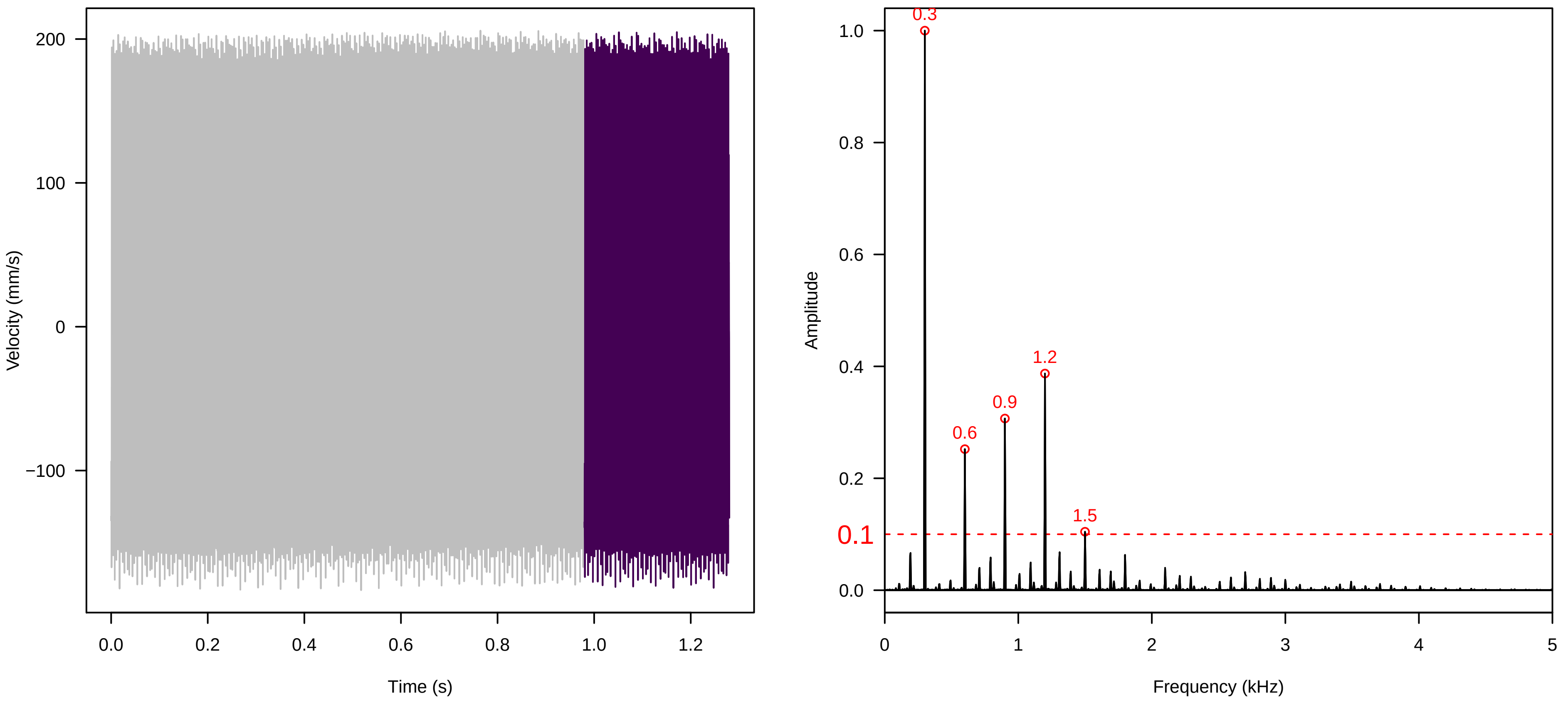

**Vel. = 0.028 ; Str. = Receptacle ; Axis = y ; Fl. accession = 10−s−81−1AA**

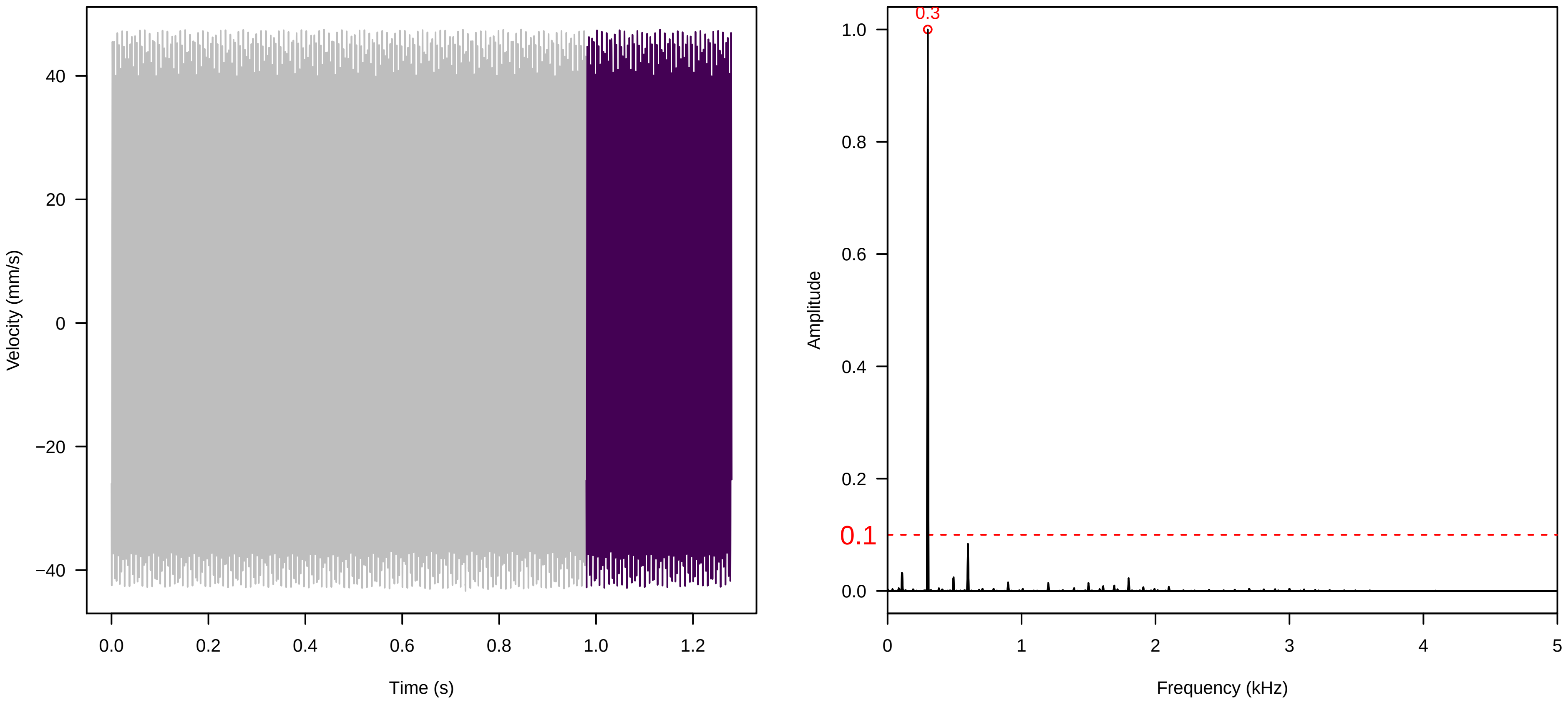


**Vel. = 0.028 ; Str. = Corolla ; Axis = y ; Fl. accession = 10−s−81−1AA**

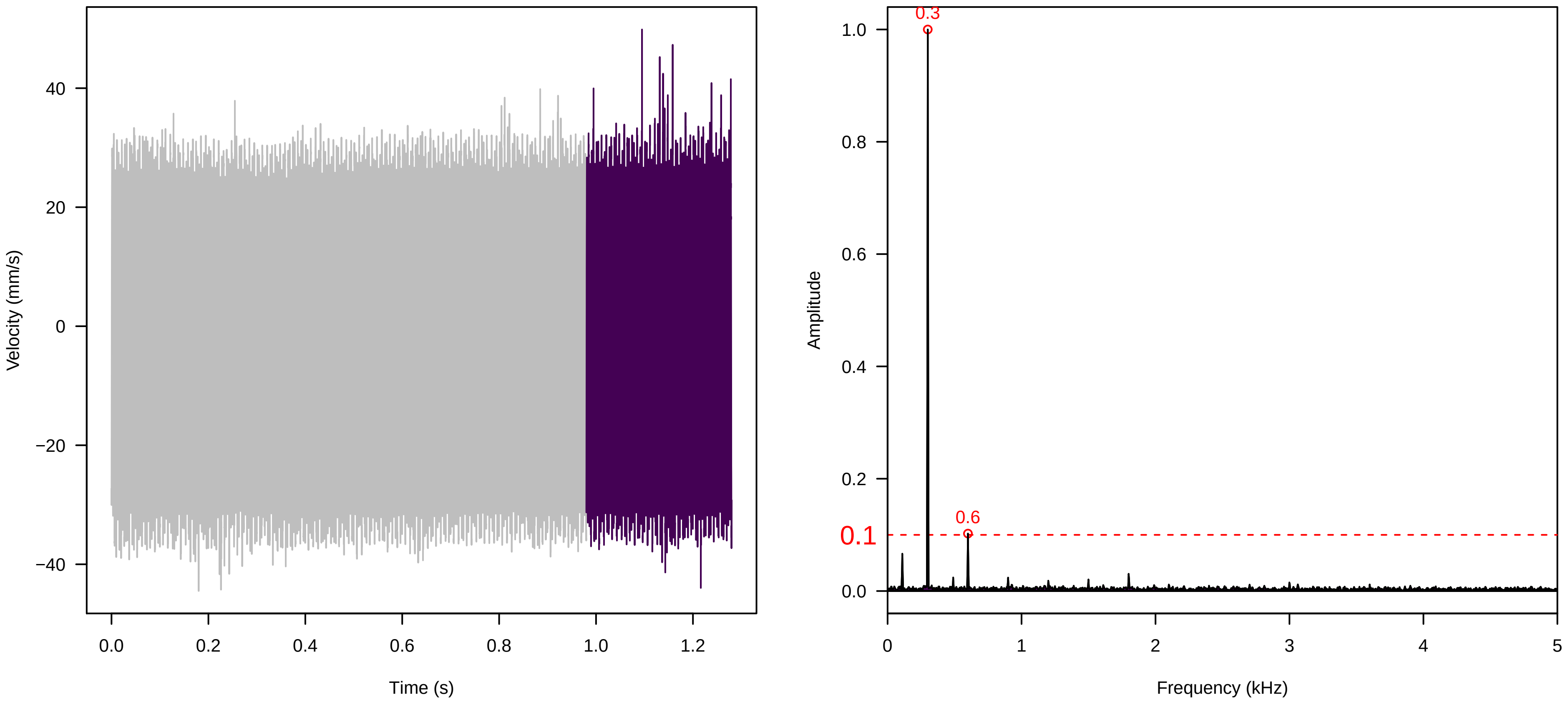


**Vel. = 0.028 ; Str. = Receptacle ; Axis = y ; Fl. accession = 10−s−81−1AA**

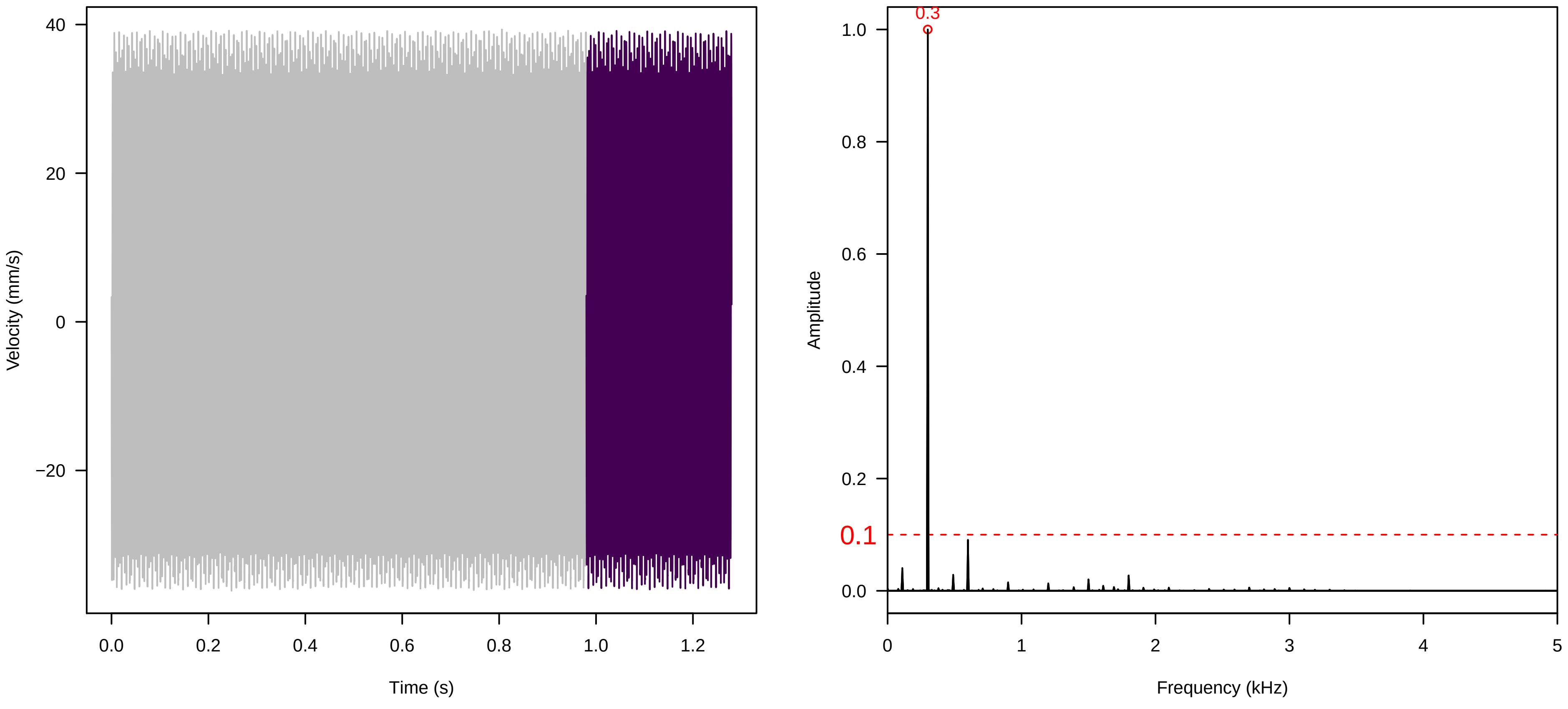

**Vel. = 0.057 ; Str. = Corolla ; Axis = y ; Fl. accession = 10−s−81−1AA**

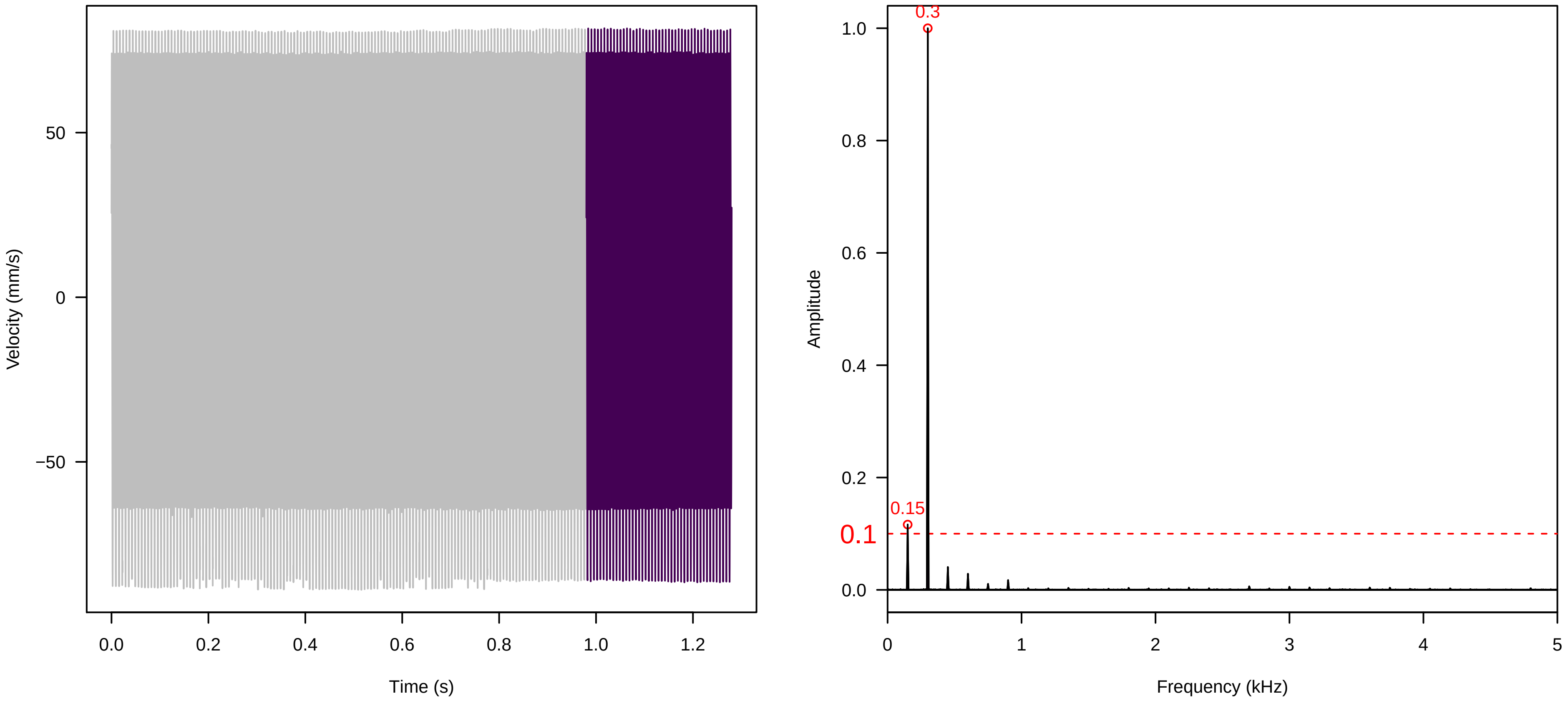


**Vel. = 0.057 ; Str. = Receptacle ; Axis = y ; Fl. accession = 10−s−81−1AA**

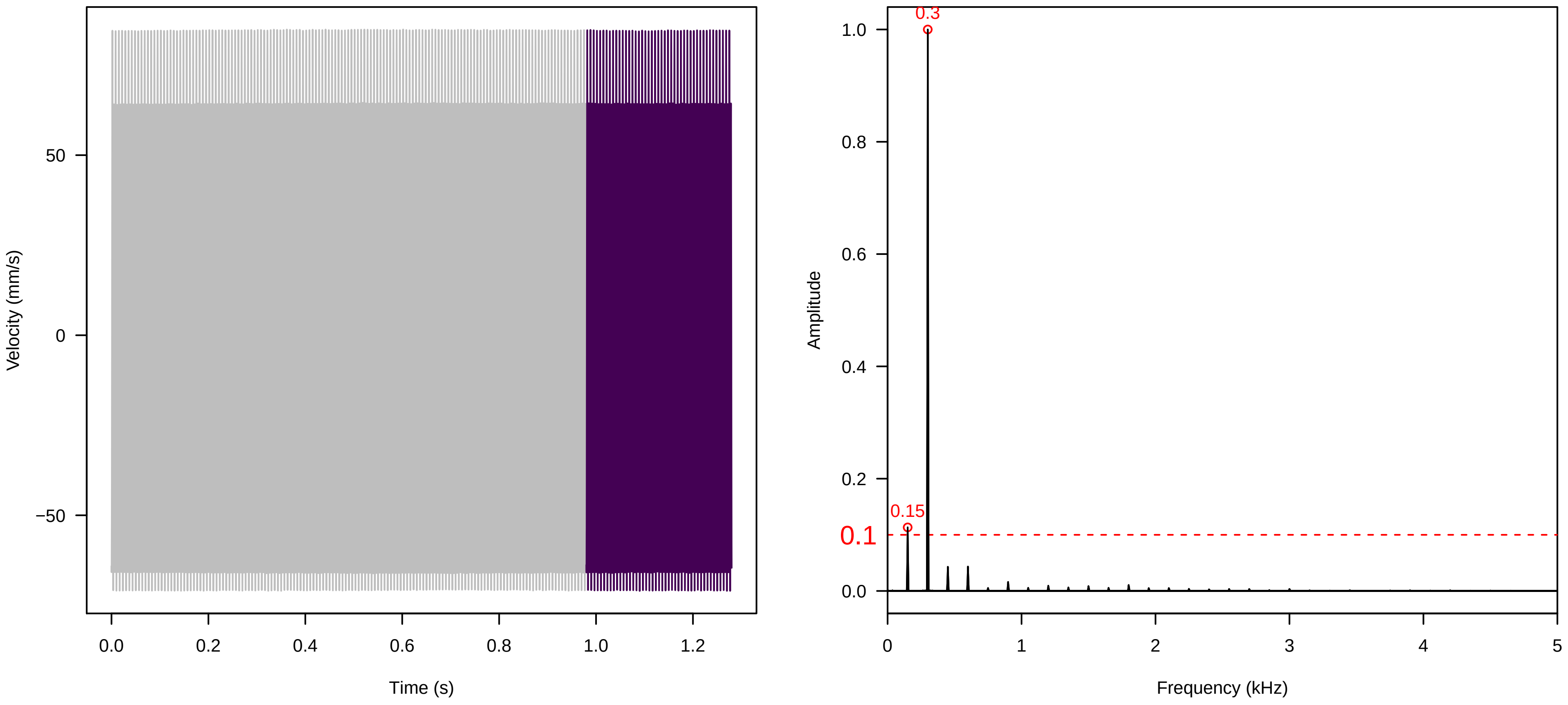


**Vel. = 0.057 ; Str. = FA ; Axis = y ; Fl. accession = 10−s−81−1AA**

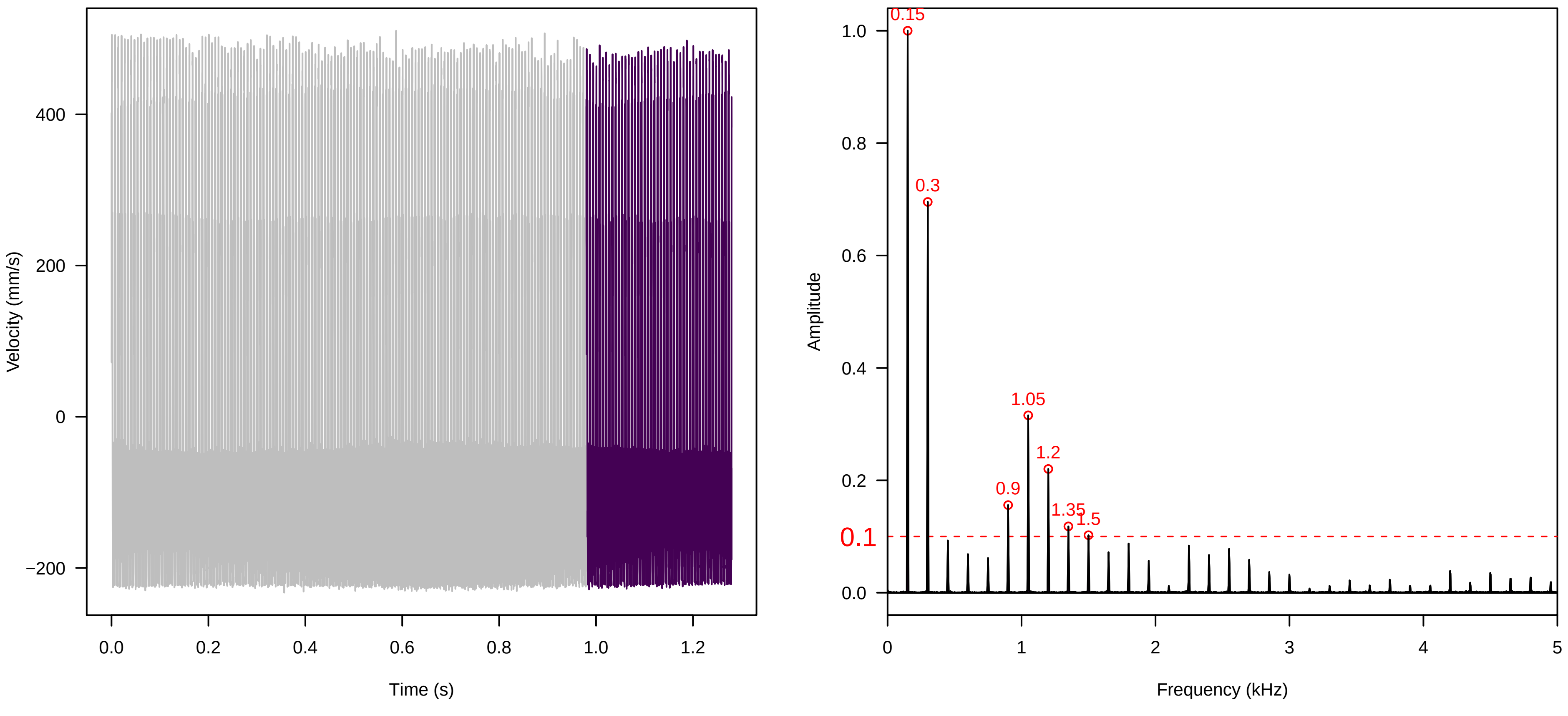

Vel. =  0.057 ; Str. =  Receptacle ; Axis =  y ; Fl. accession = 10−s−81−1AA

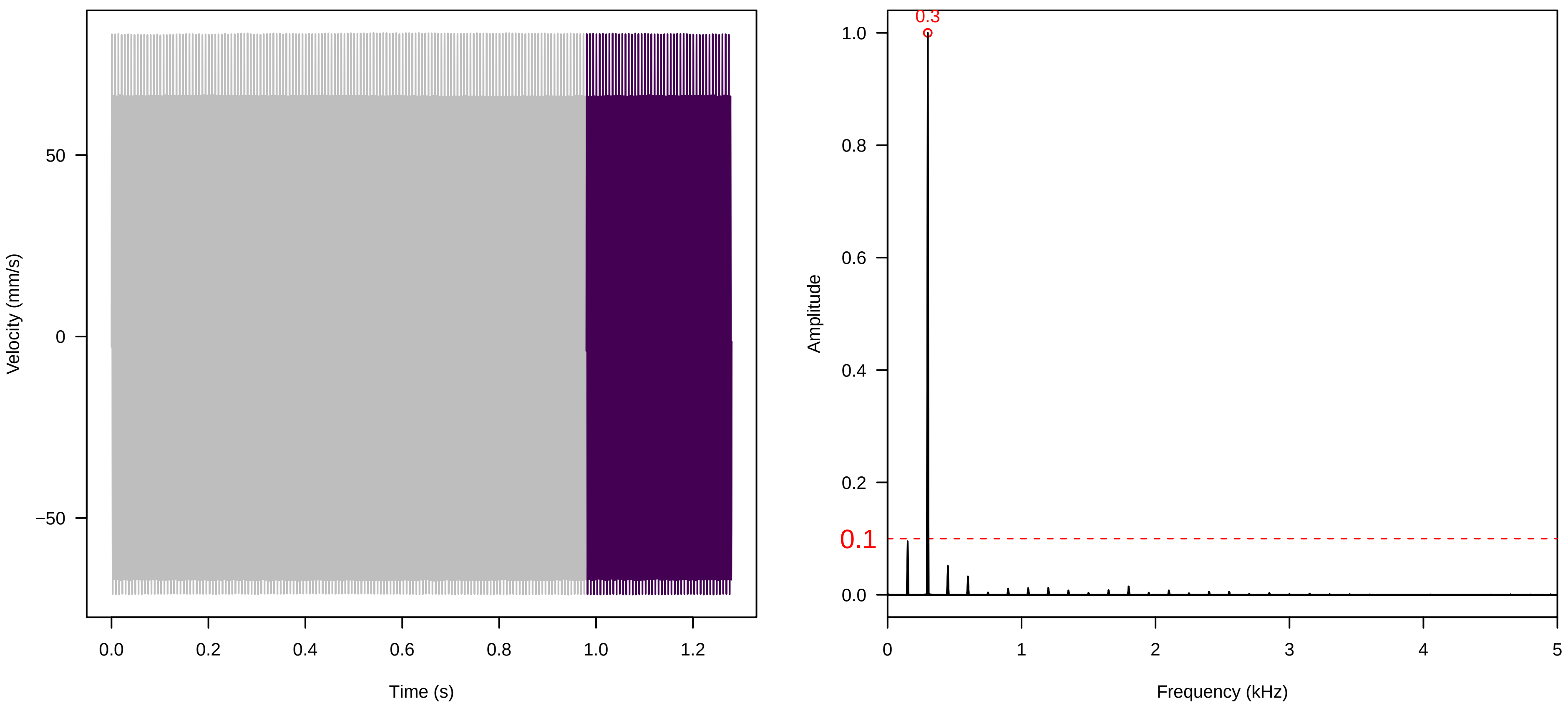


Vel. =  0.057 ; Str. =  PA ; Axis =  y ; Fl. accession = 10−s−81−1AA

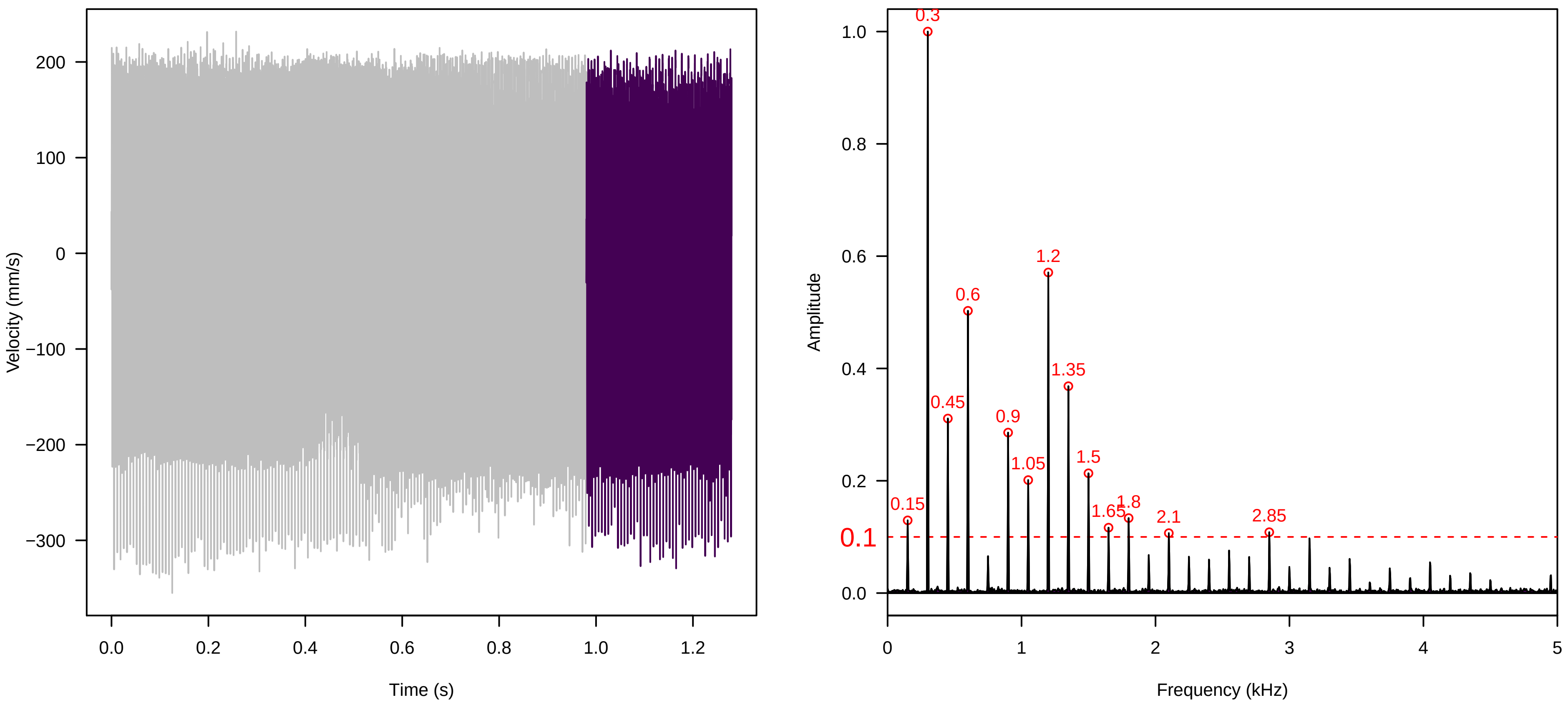


Vel. =  0.057 ; Str. =  Receptacle ; Axis =  y ; Fl. accession = 10−s−81−1AA

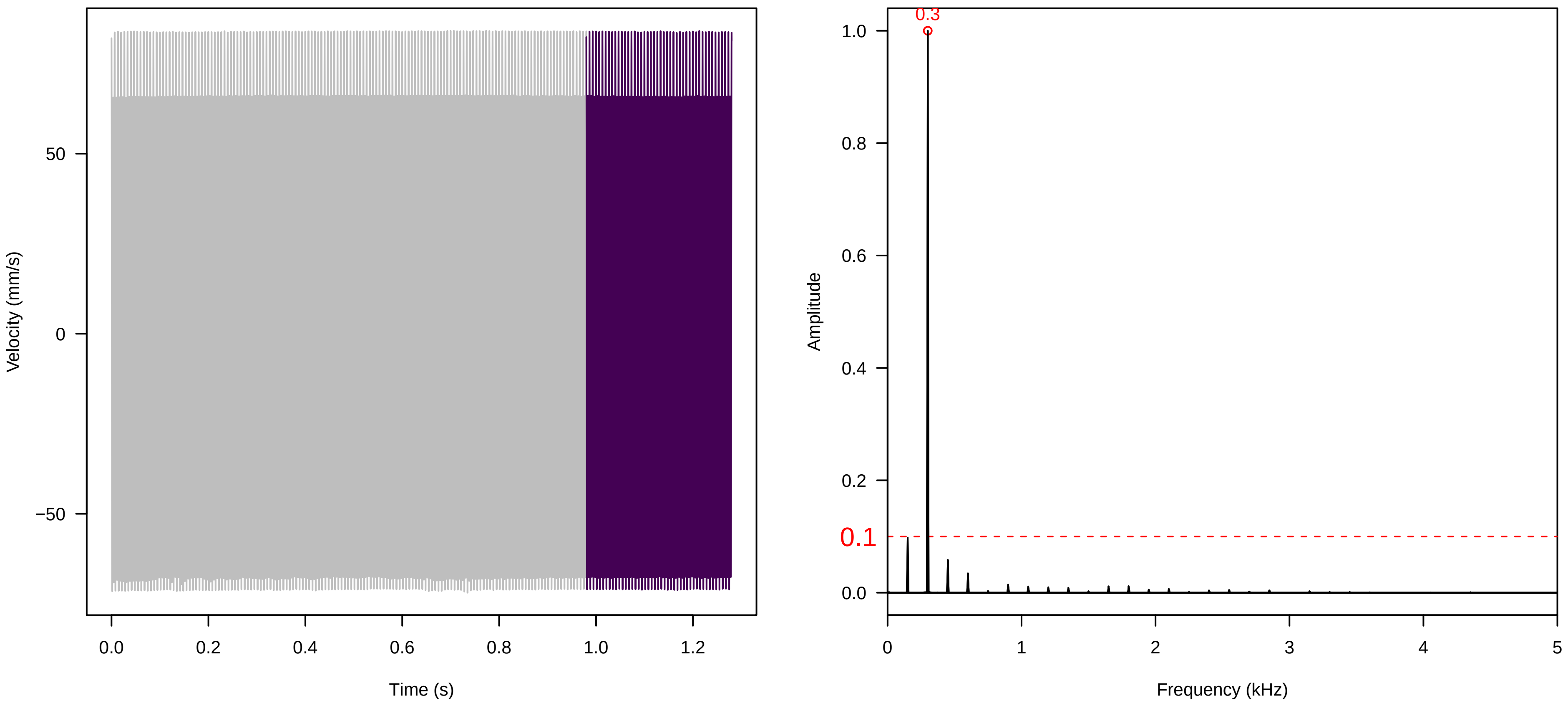

**Vel. =  0.057 ; Str. =  Corolla ; Axis =  y ; Fl. accession = 10−s−81−1AB**

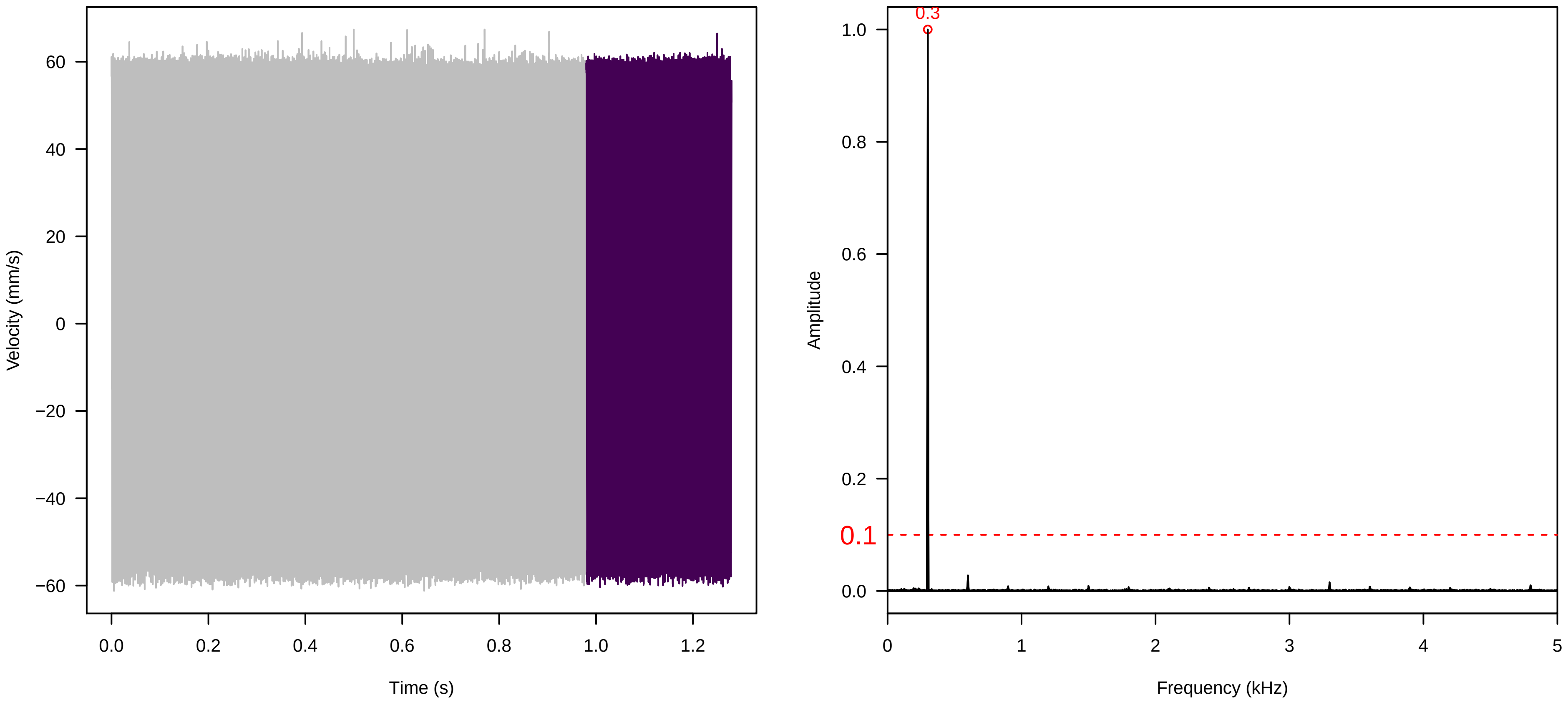


**Vel. =  0.057 ; Str. =  Receptacle ; Axis =  y ; Fl. accession = 10−s−81−1AB**

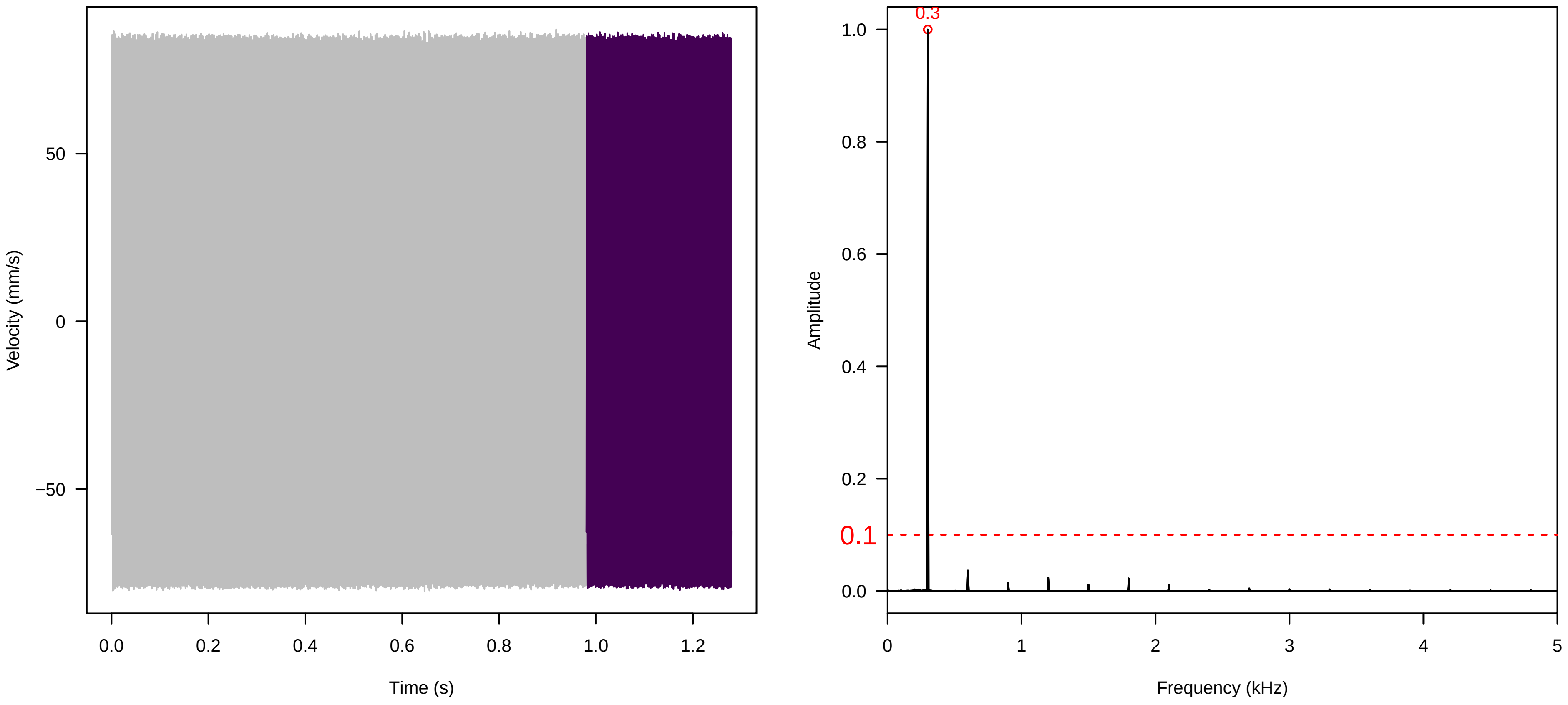


**Vel. =  0.057 ; Str. =  FA ; Axis =  y ; Fl. accession = 10−s−81−1AB**

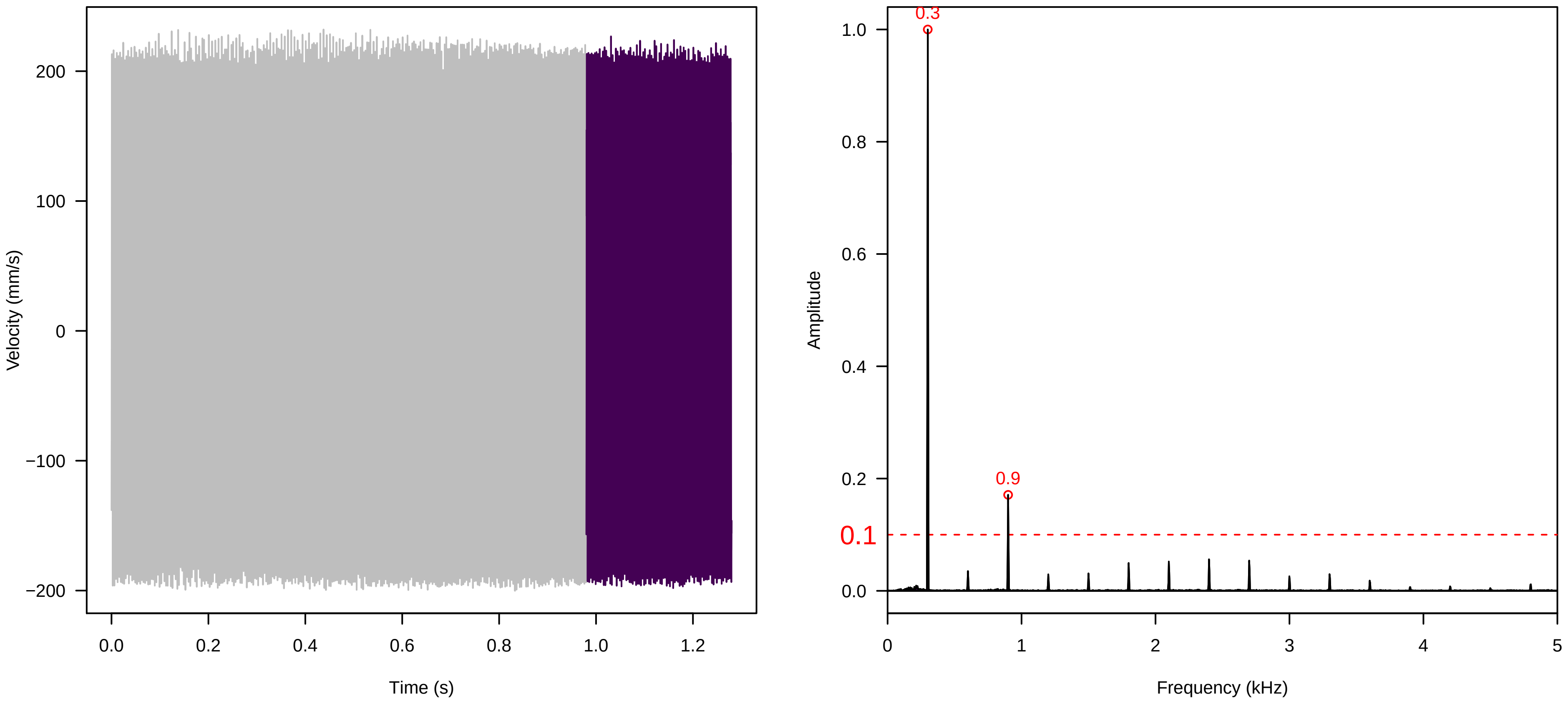

**Vel. =  0.057 ; Str. =  Receptacle ; Axis =  y ; Fl. accession = 10−s−81−1AB**

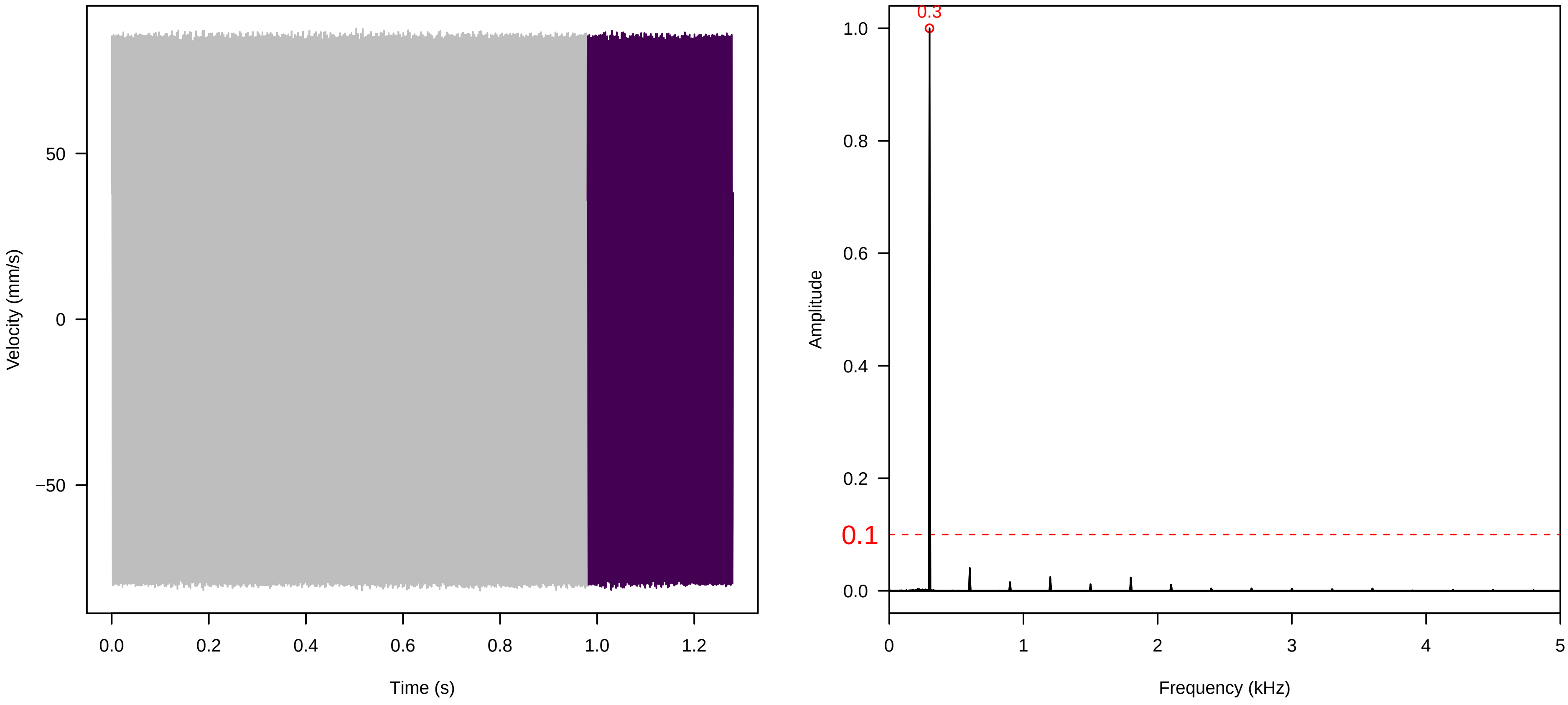


**Vel. =  0.057 ; Str. =  PA ; Axis =  y ; Fl. accession = 10−s−81−1AB**

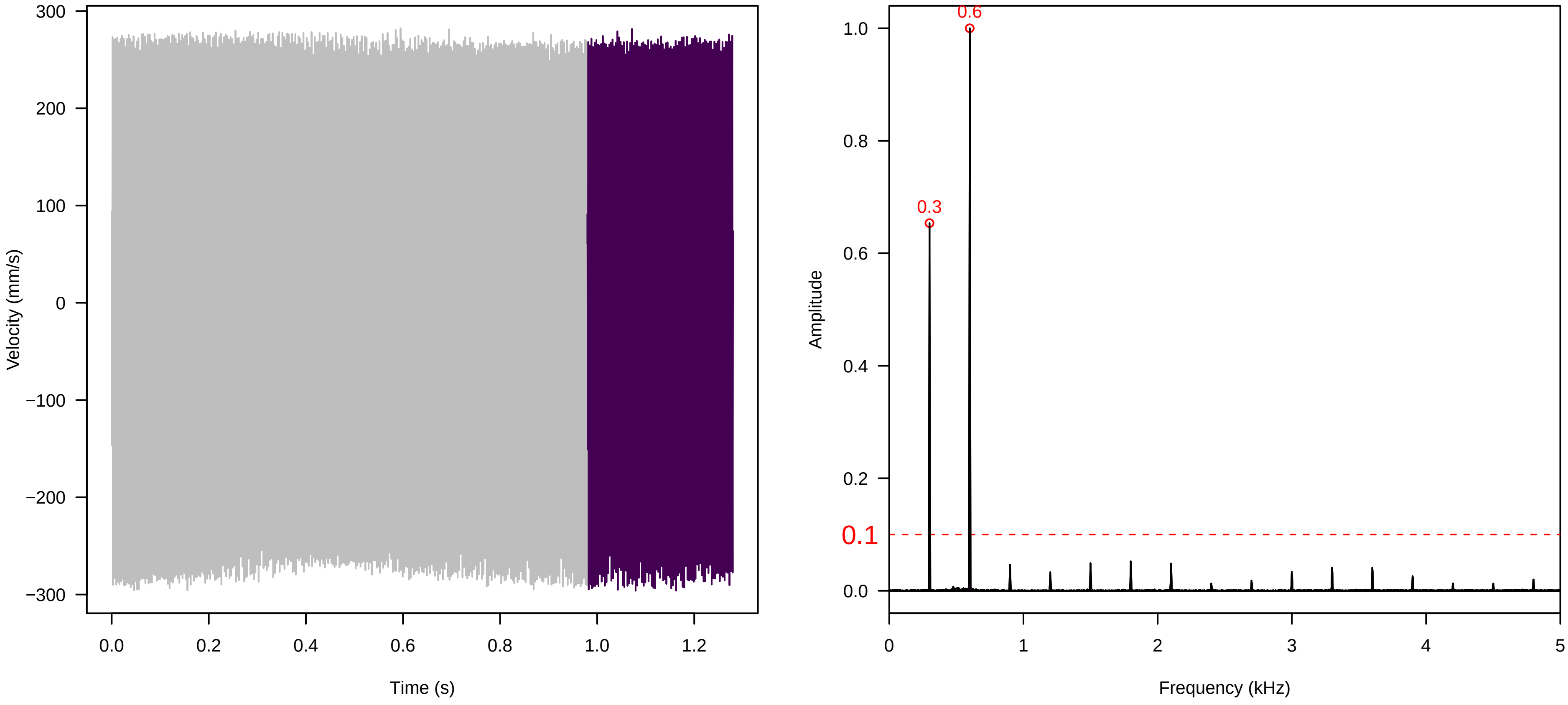


**Vel. =  0.057 ; Str. =  Receptacle ; Axis =  y ; Fl. accession = 10−s−81−1AB**

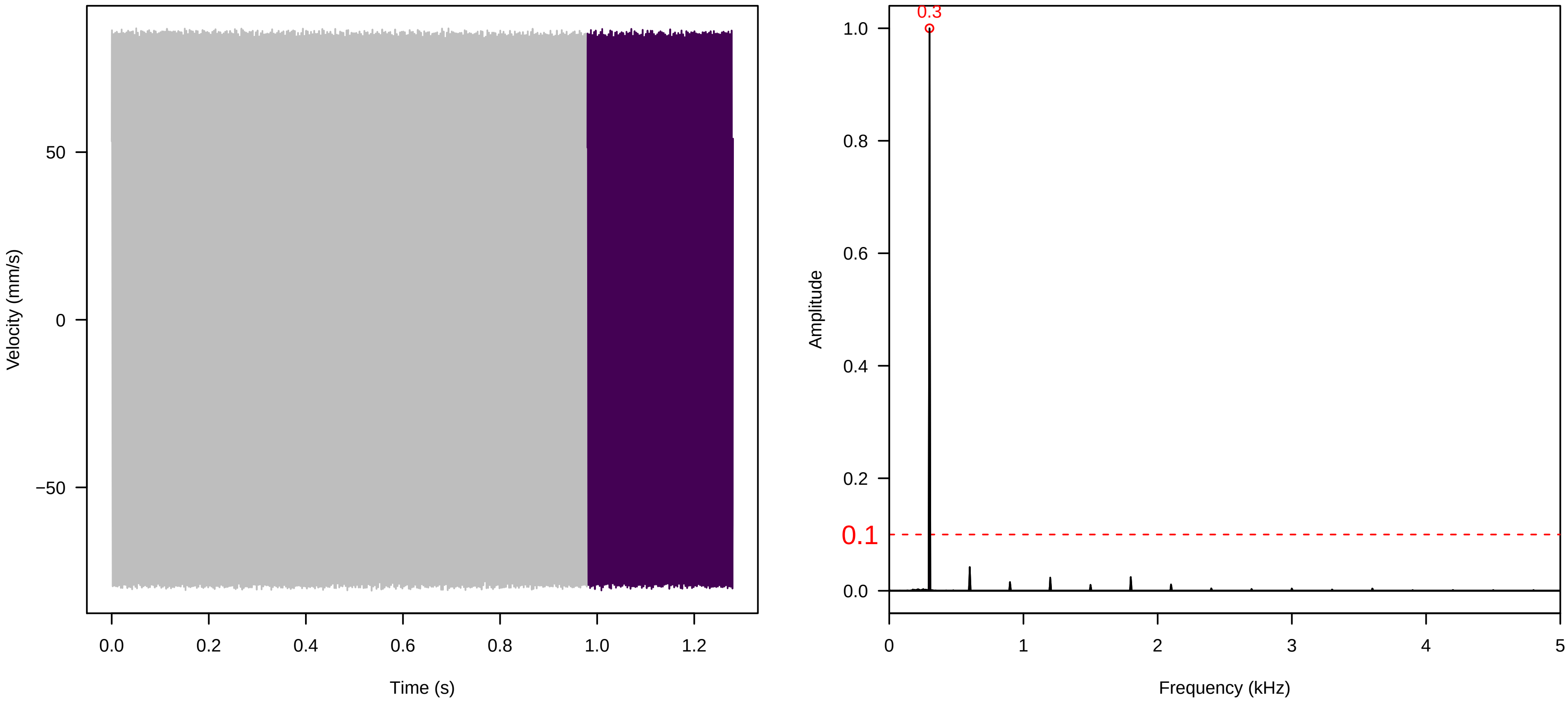

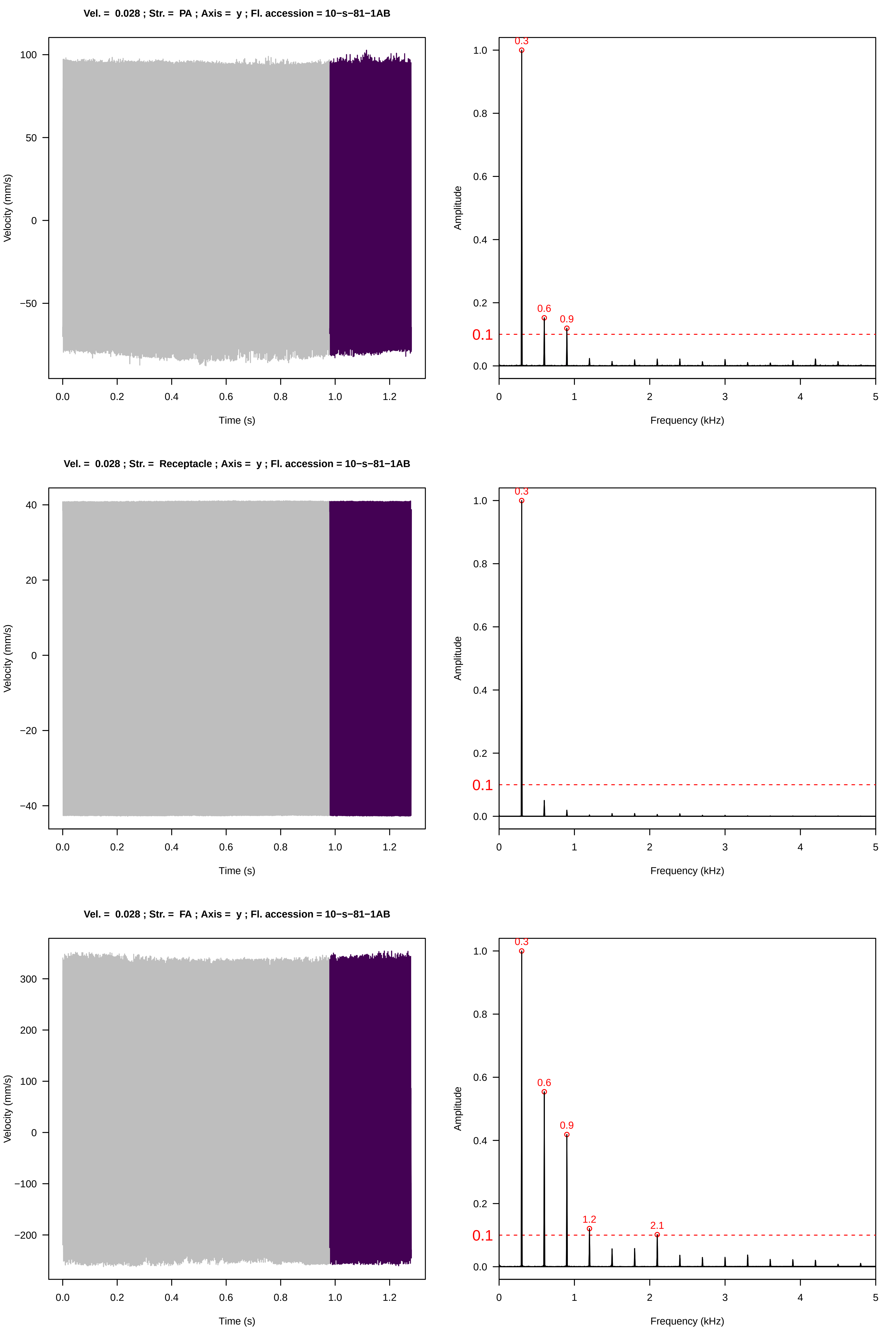

Vel. = 0.028 ; Str. = PA ; Axis = y ; Fl. accession = 10−s−81−1AB
Velocity (mm/s)
Time (s)
Amplitude
Frequency (kHz)
0.3
0.6
0.9
0.1
Vel. = 0.028 ; Str. = Receptacle ; Axis = y ; Fl. accession = 10−s−81−1AB
Velocity (mm/s)
Time (s)
Amplitude
Frequency (kHz)
0.3
0.1
Vel. = 0.028 ; Str. = FA ; Axis = y ; Fl. accession = 10−s−81−1AB
Velocity (mm/s)
Time (s)
Amplitude
Frequency (kHz)
0.3
0.6
0.9
1.2
2.1
0.1

**Vel. = 0.028 ; Str. = Receptacle ; Axis = y ; Fl. accession = 10−s−81−1AB**

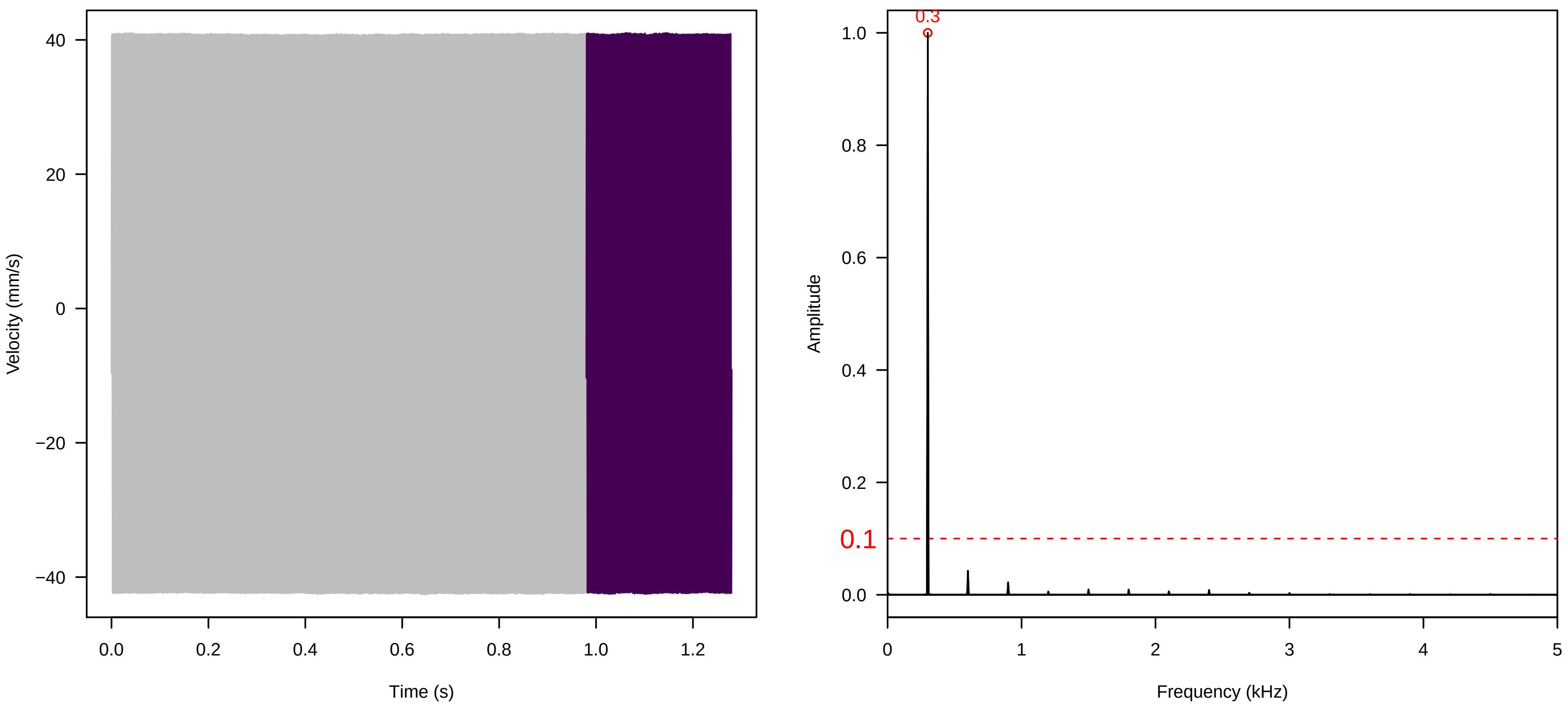


**Vel. = 0.028 ; Str. = Corolla ; Axis = y ; Fl. accession = 10−s−81−1AB**

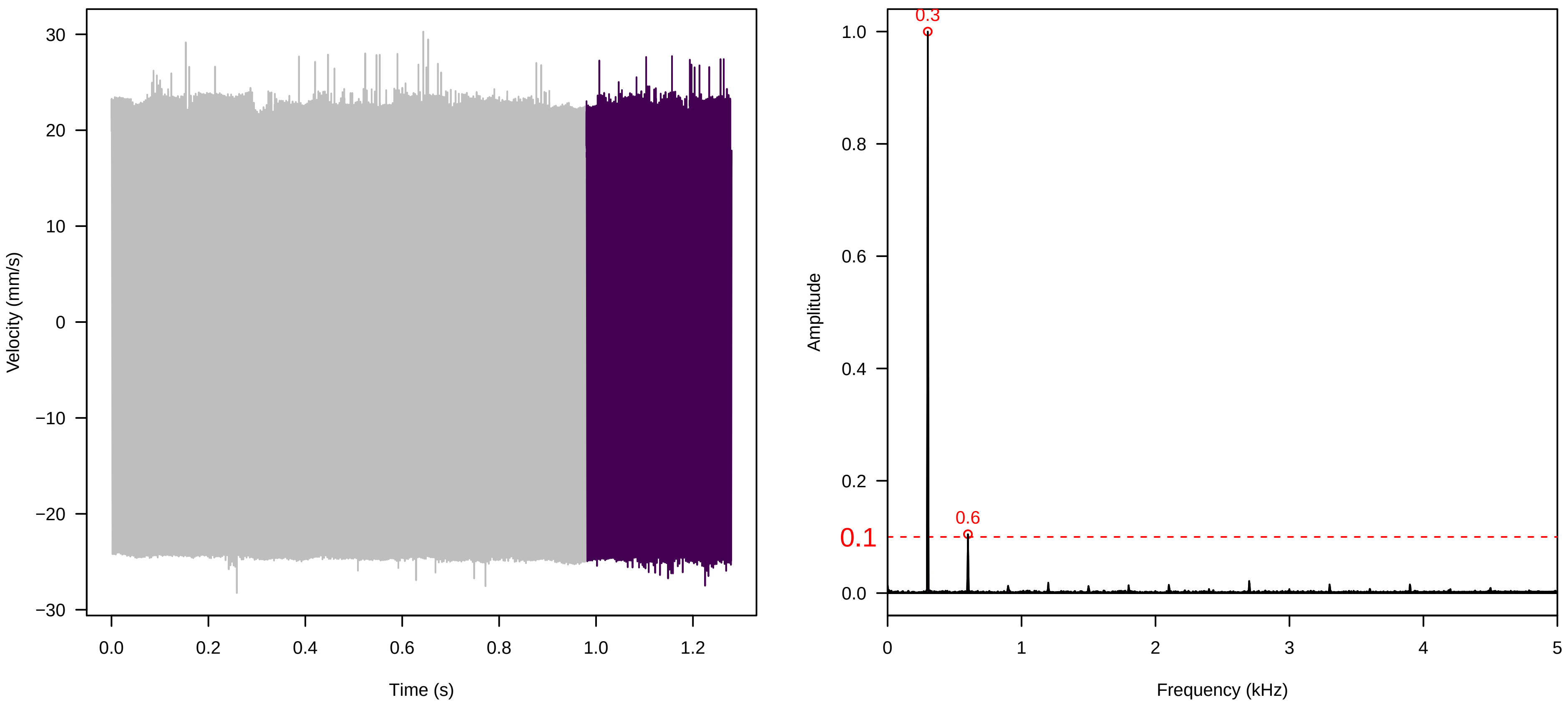


**Vel. = 0.028 ; Str. = Receptacle ; Axis = y ; Fl. accession = 10−s−81−1AB**

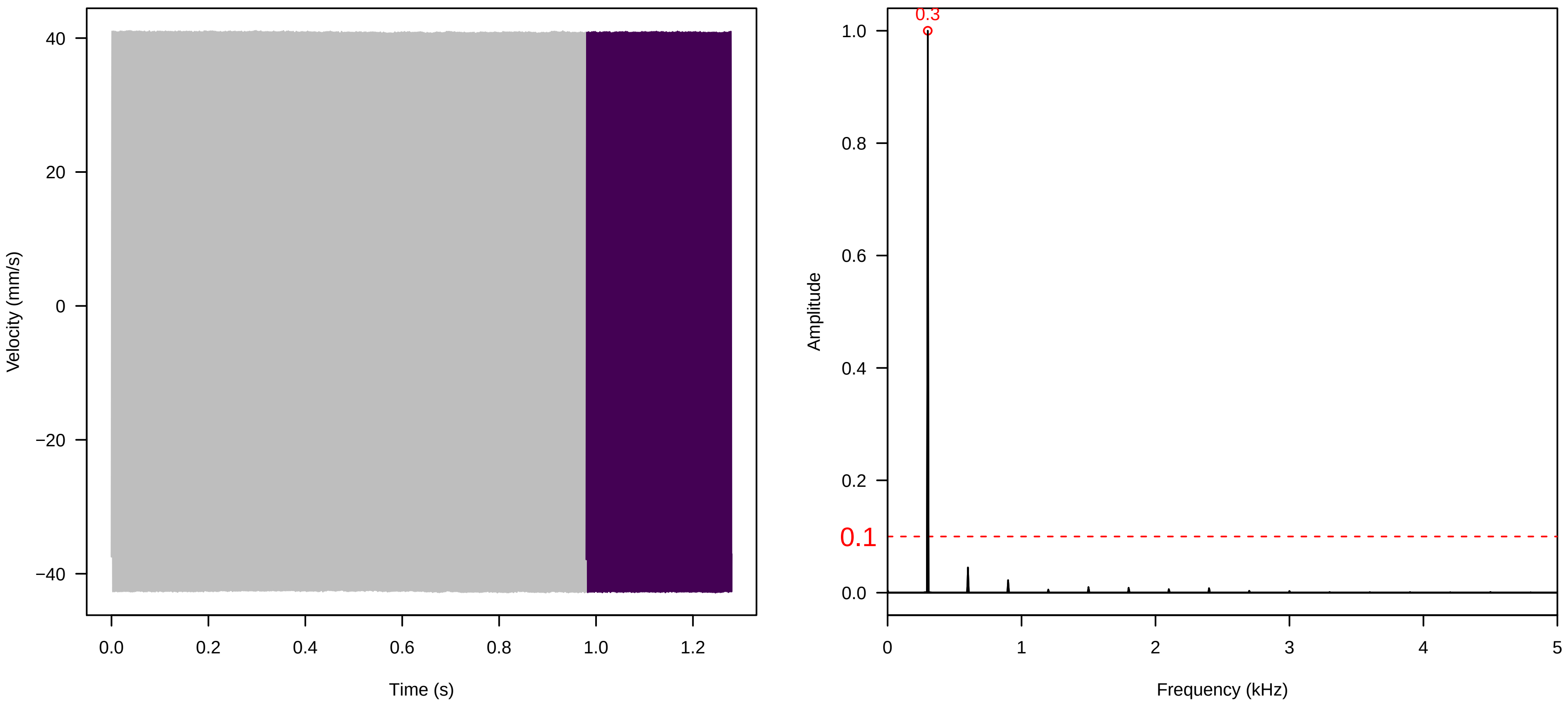

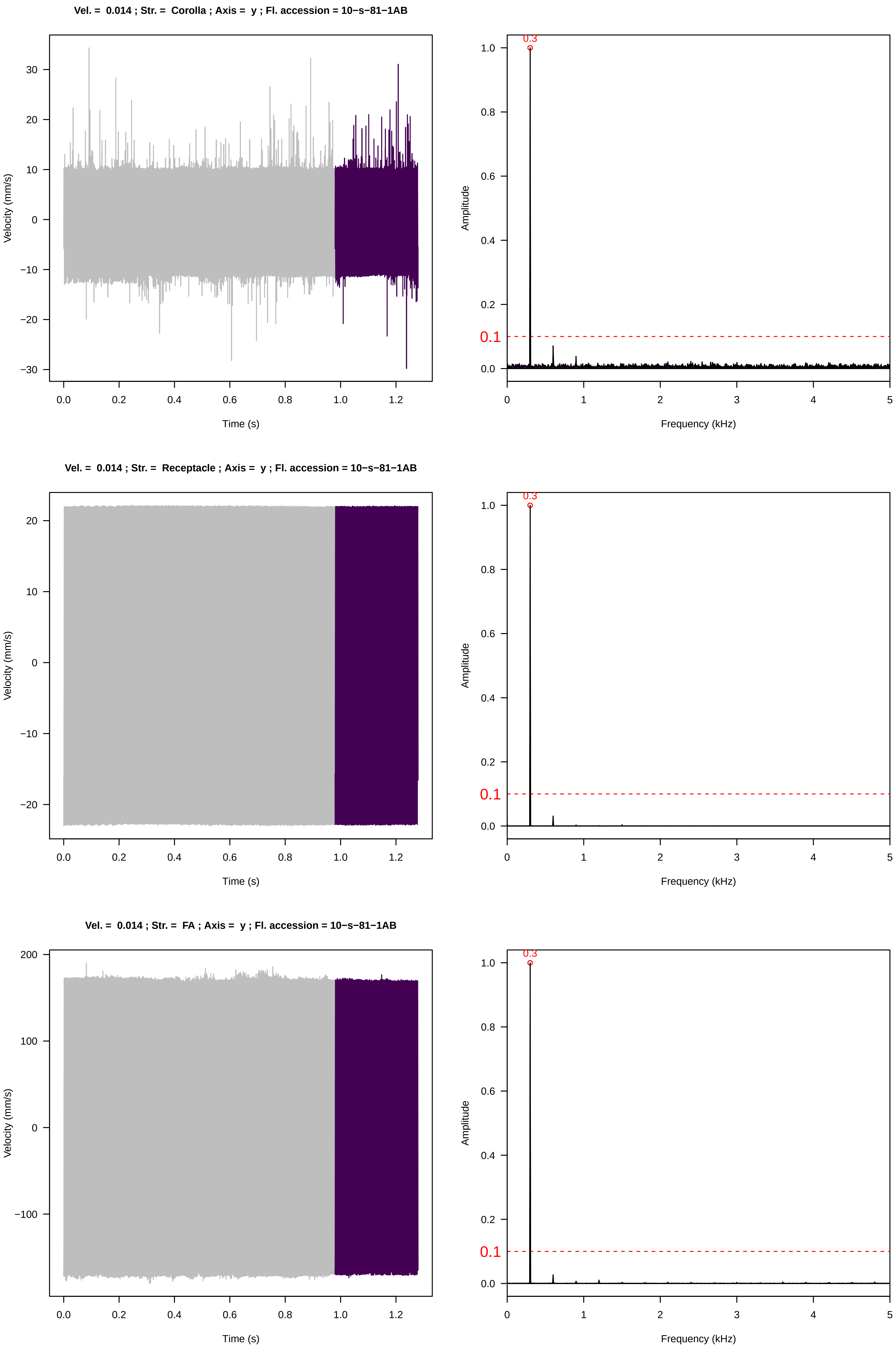
Vel. = 0.014 ; Str. = Corolla ; Axis = y ; Fl. accession = 10−s−81−1AB
Velocity (mm/s)
Time (s)
Amplitude
Frequency (kHz)
0.3
0.1
Vel. = 0.014 ; Str. = Receptacle ; Axis = y ; Fl. accession = 10−s−81−1AB
Velocity (mm/s)
Time (s)
Amplitude
Frequency (kHz)
0.3
0.1
Vel. = 0.014 ; Str. = FA ; Axis = y ; Fl. accession = 10−s−81−1AB
Velocity (mm/s)
Time (s)
Amplitude
Frequency (kHz)
0.3
0.1

**Vel. = 0.014 ; Str. = Receptacle ; Axis = y ; Fl. accession = 10−s−81−1AB**

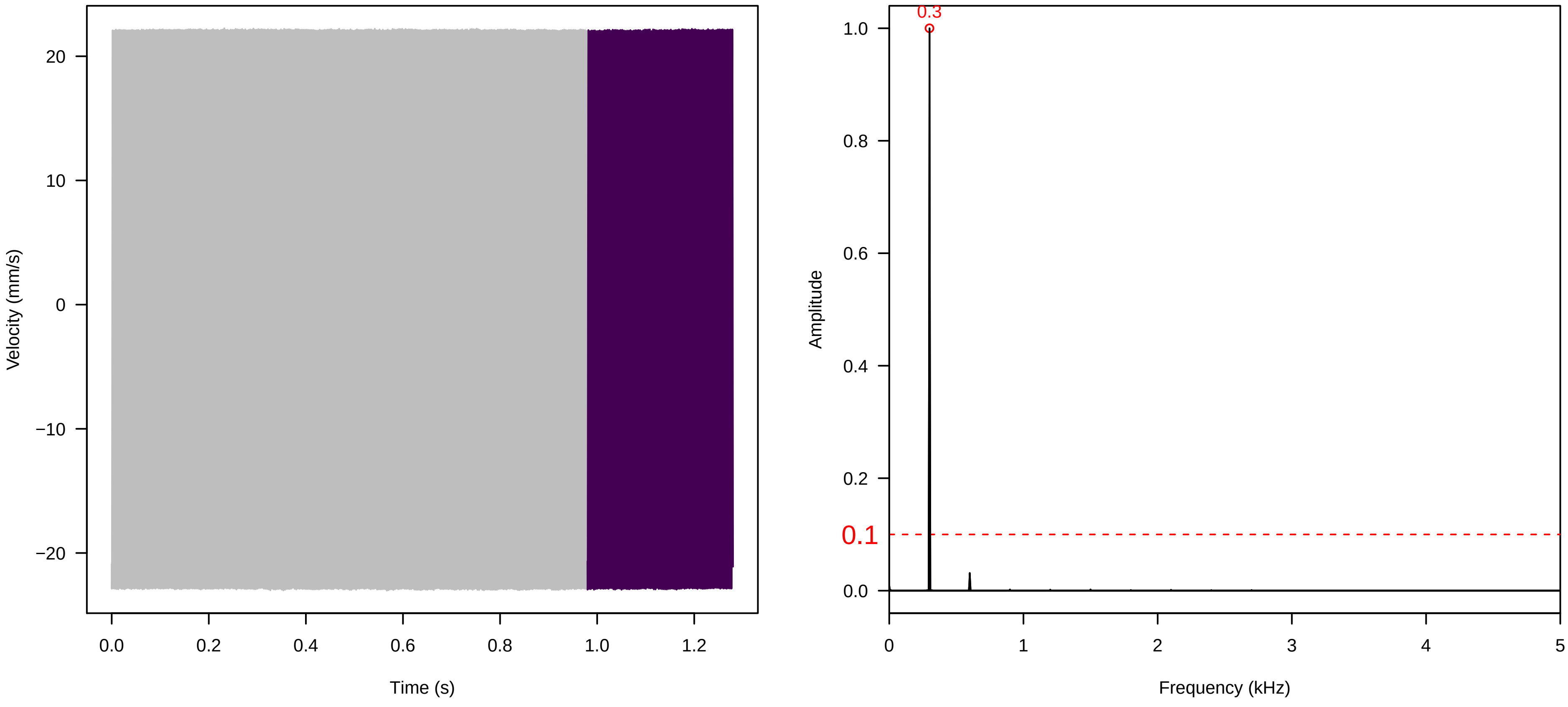


**Vel. = 0.014 ; Str. = PA ; Axis = y ; Fl. accession = 10−s−81−1AB**

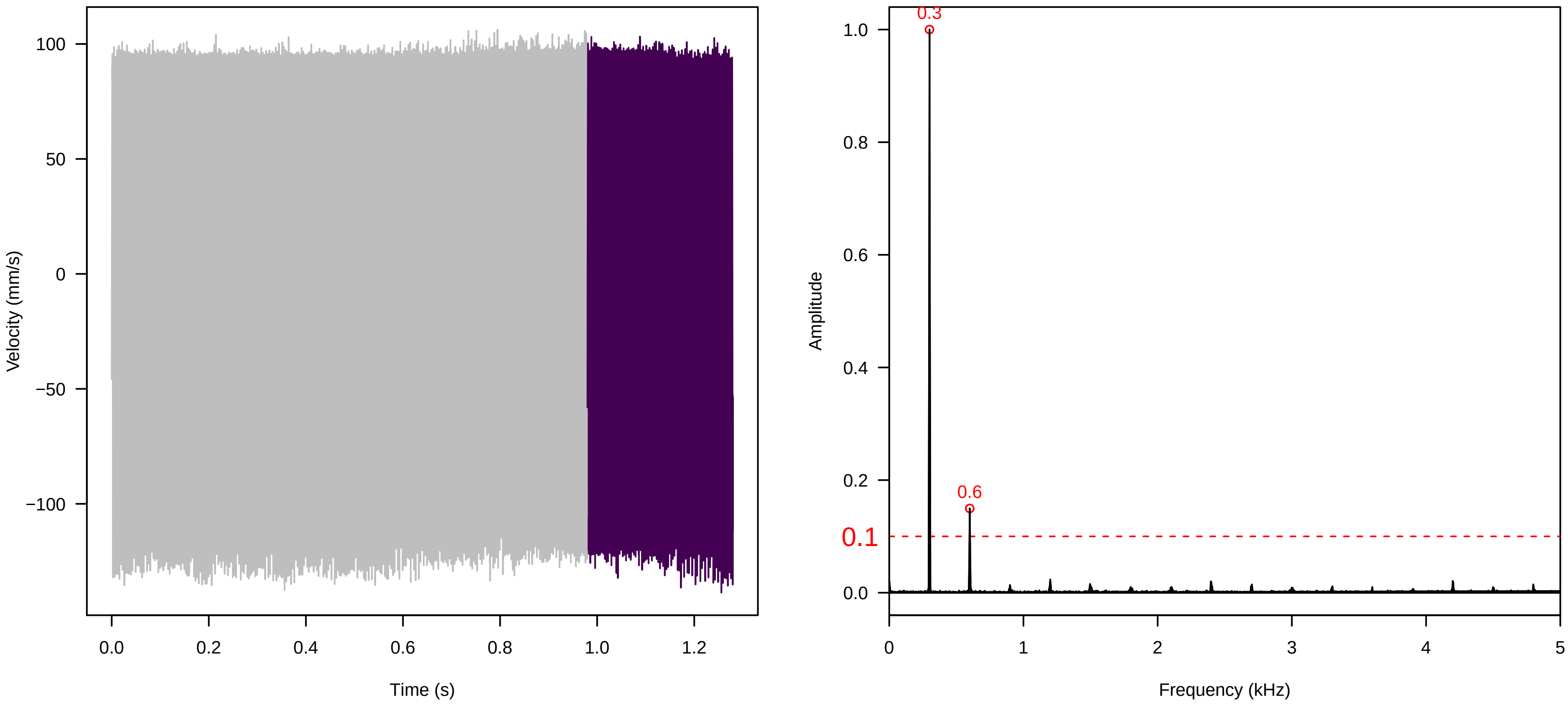


**Vel. = 0.014 ; Str. = Receptacle ; Axis = y ; Fl. accession = 10−s−81−1AB**

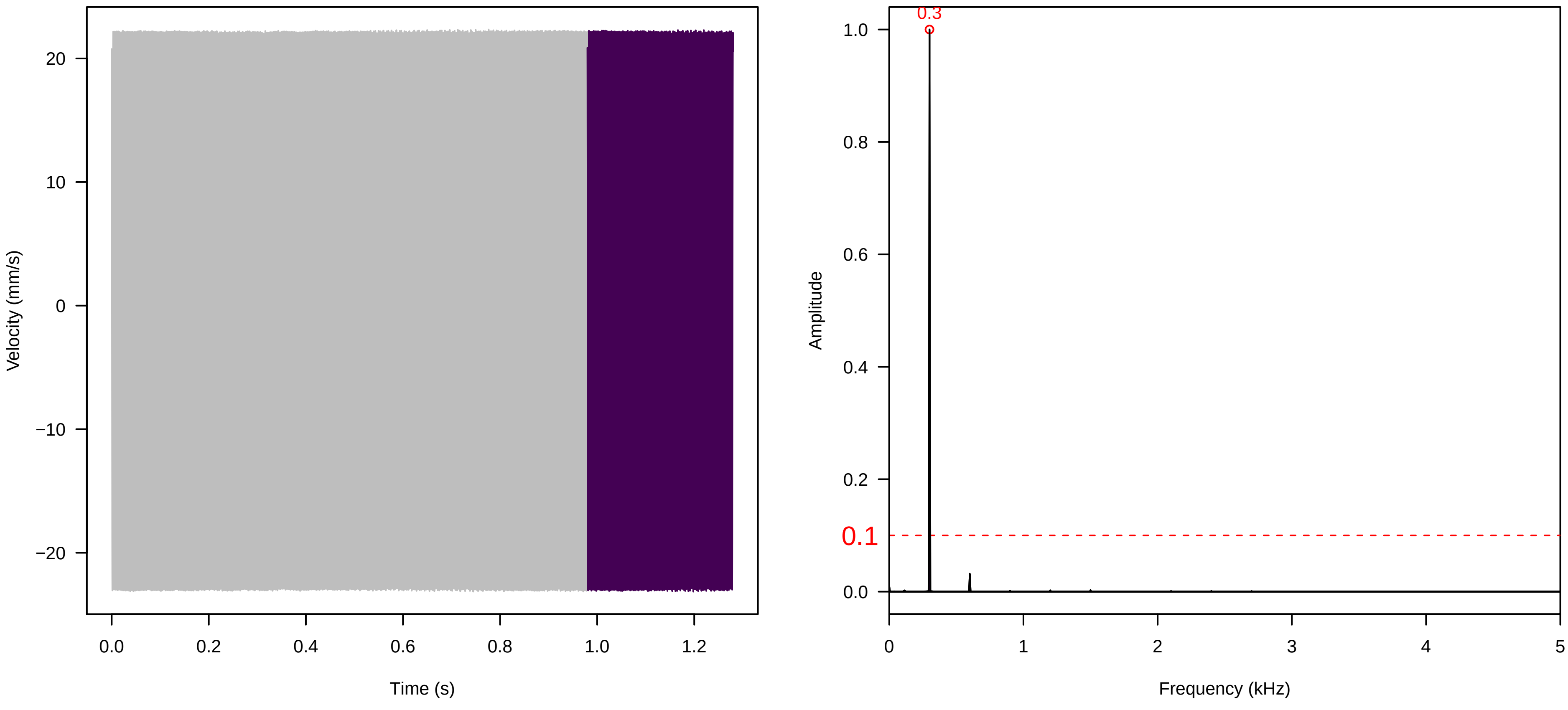

**Vel. = 0.014 ; Str. = Corolla ; Axis = z ; Fl. accession = 10−s−81−1AB**

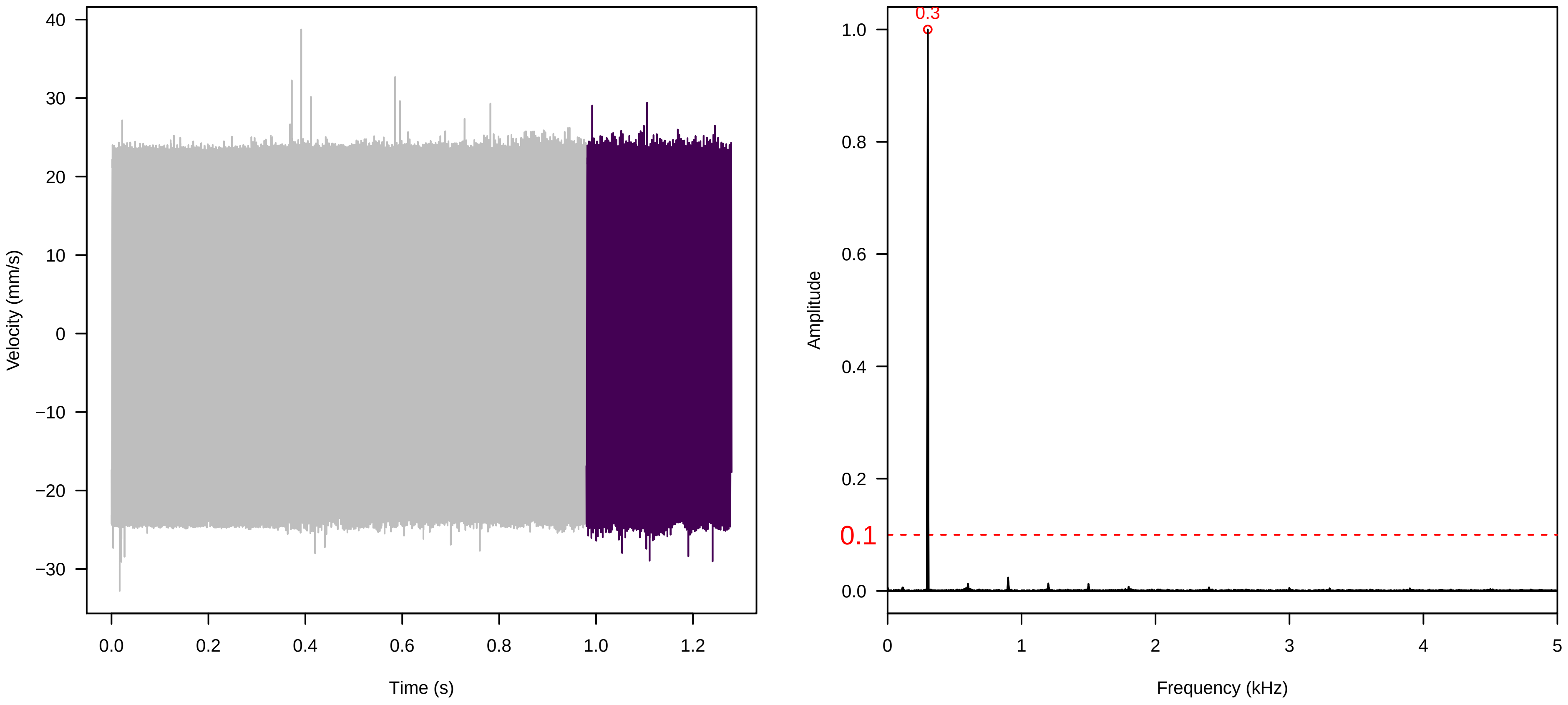


**Vel. = 0.014 ; Str. = Receptacle ; Axis = z ; Fl. accession = 10−s−81−1AB**

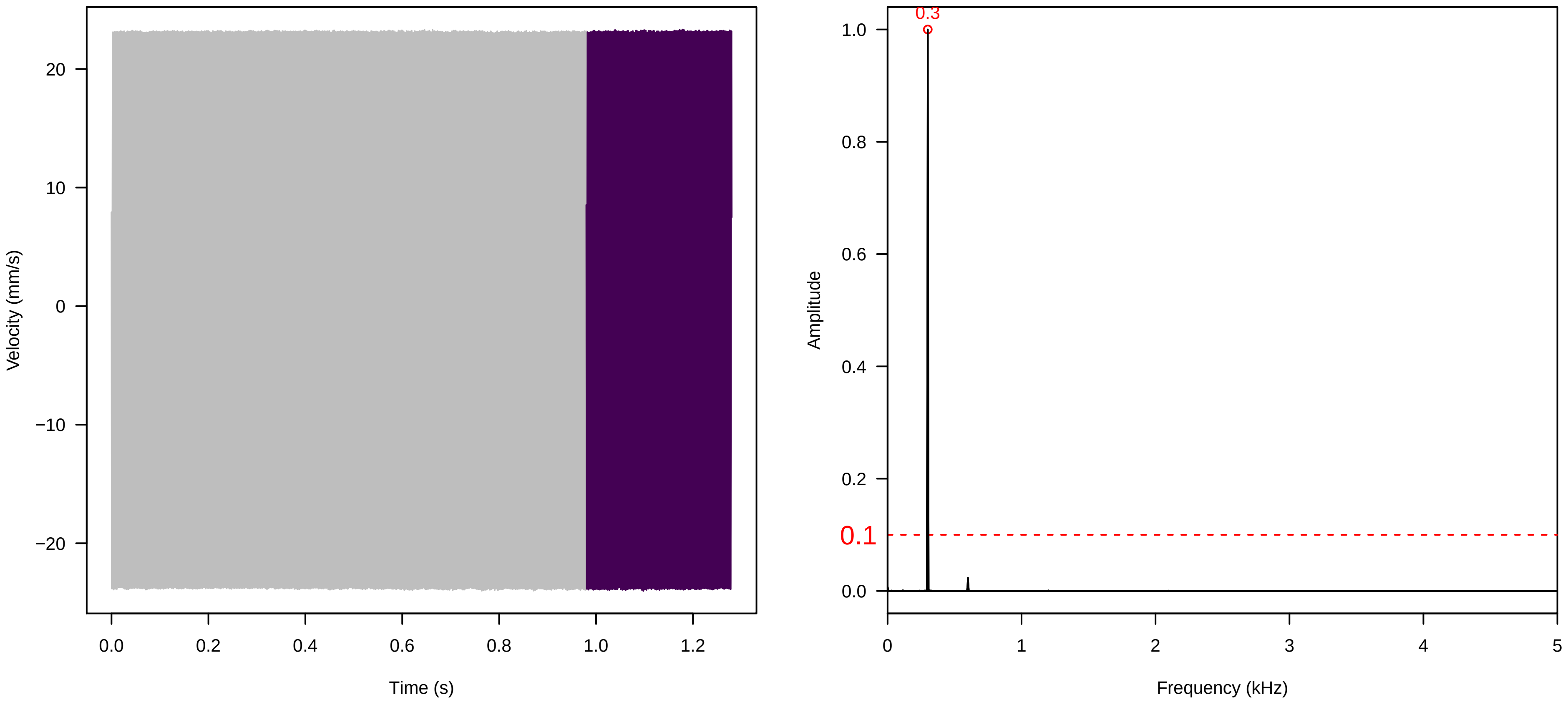


**Vel. = 0.014 ; Str. = FA ; Axis = z ; Fl. accession = 10−s−81−1AB**

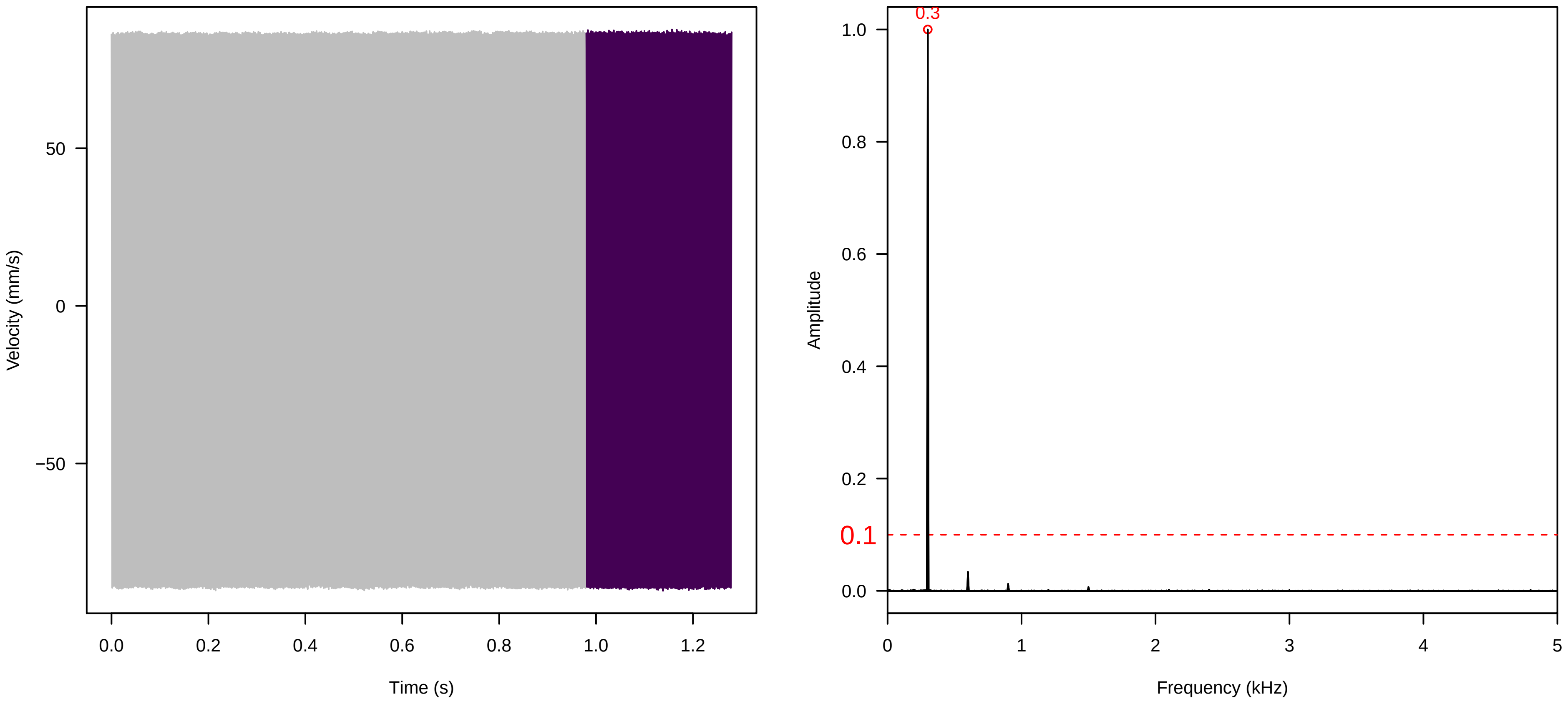

**Vel. =  0.014 ; Str. =  Receptacle ; Axis =  z ; Fl. accession = 10−s−81−1AB**

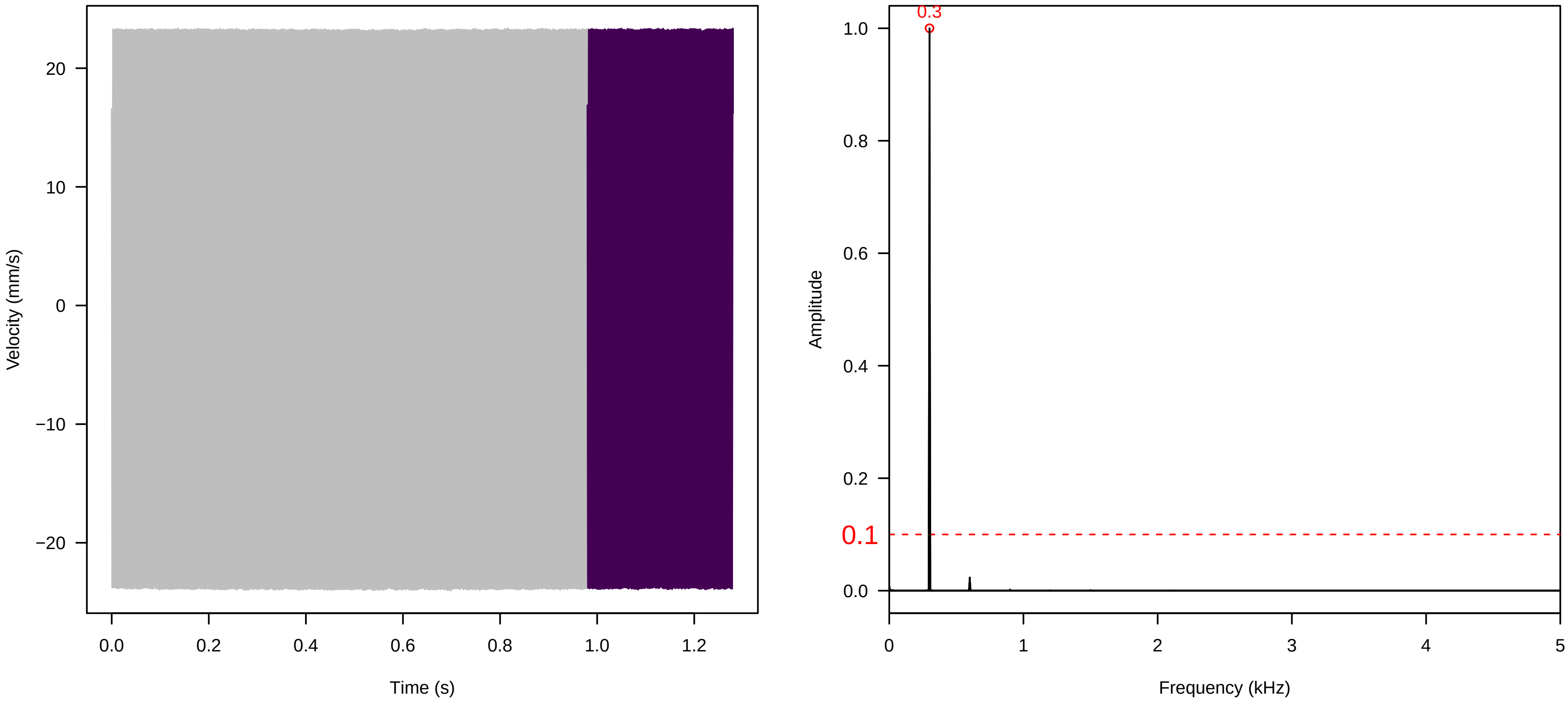


**Vel. =  0.014 ; Str. =  PA ; Axis =  z ; Fl. accession = 10−s−81−1AB**

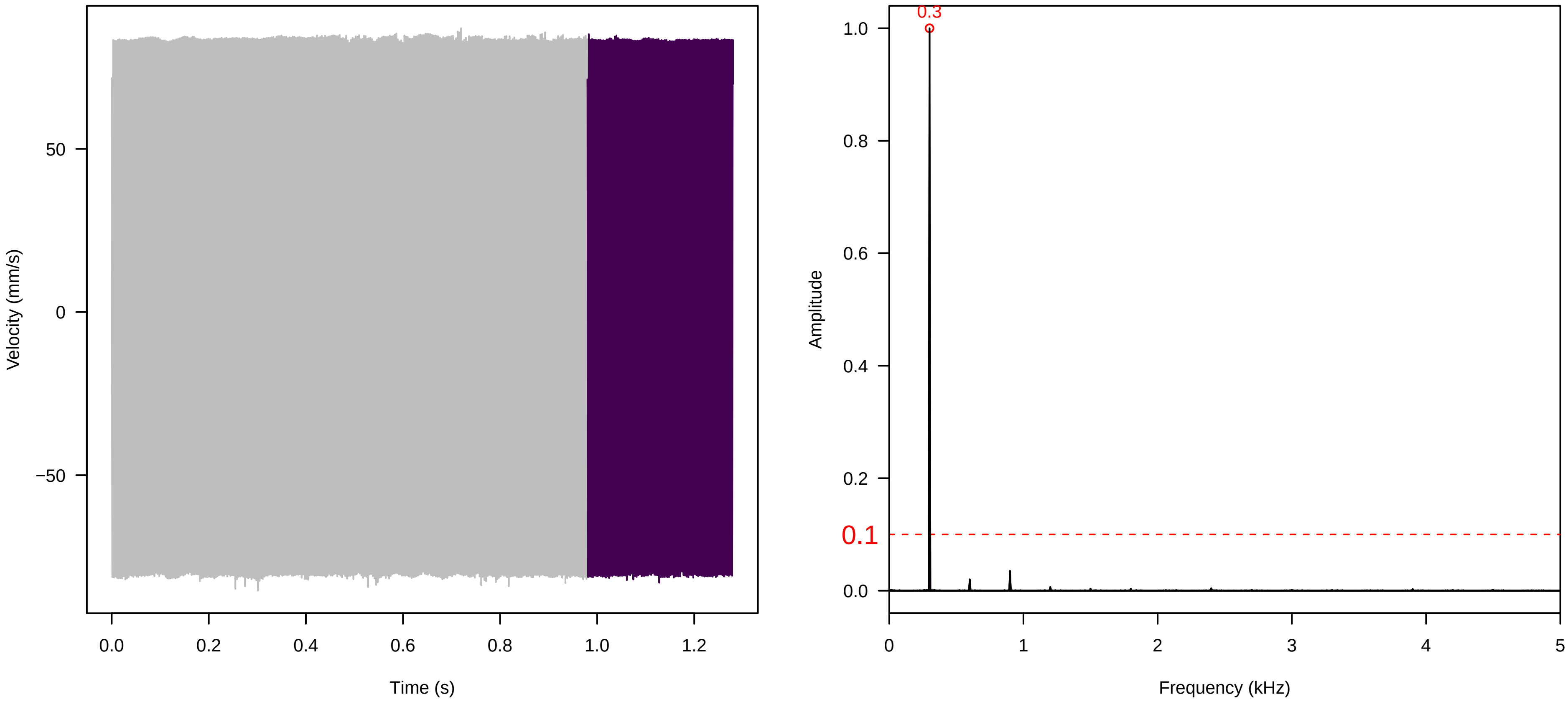


**Vel. =  0.014 ; Str. =  Receptacle ; Axis =  z ; Fl. accession = 10−s−81−1AB**

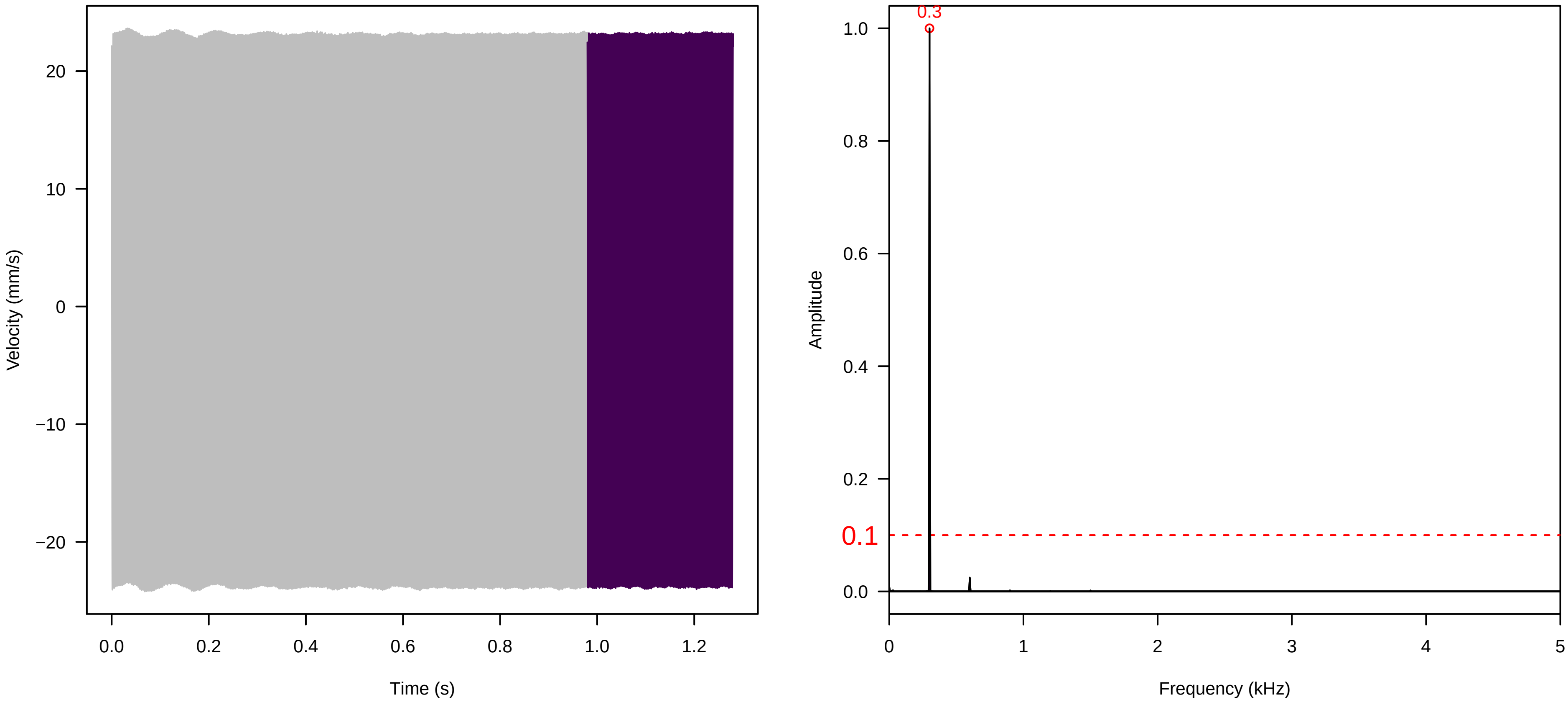

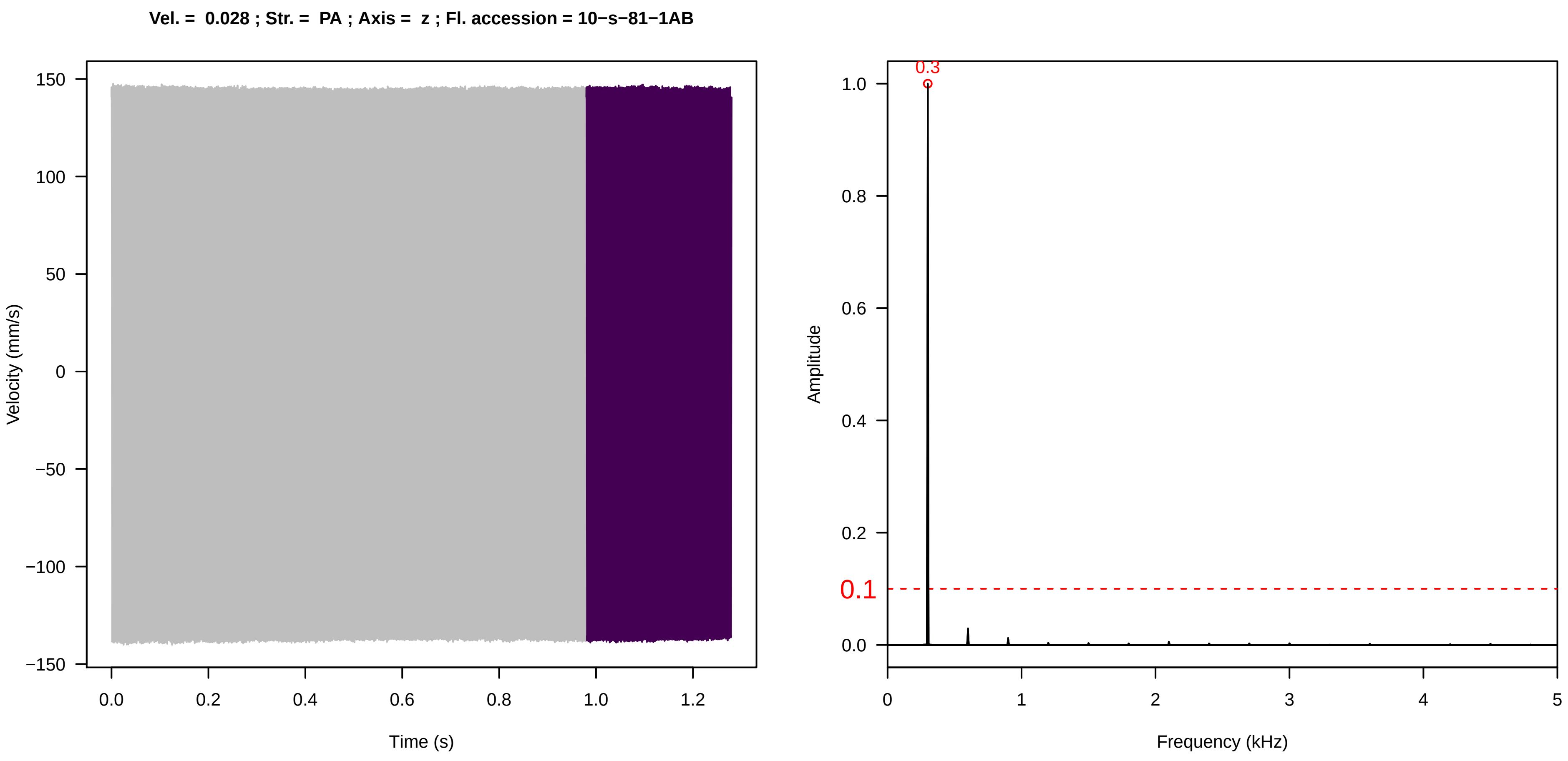
Vel. = 0.028 ; Str. = PA ; Axis = z ; Fl. accession = 10−s−81−1AB
Velocity (mm/s)
Time (s)
Amplitude
Frequency (kHz)
0.3
0.1

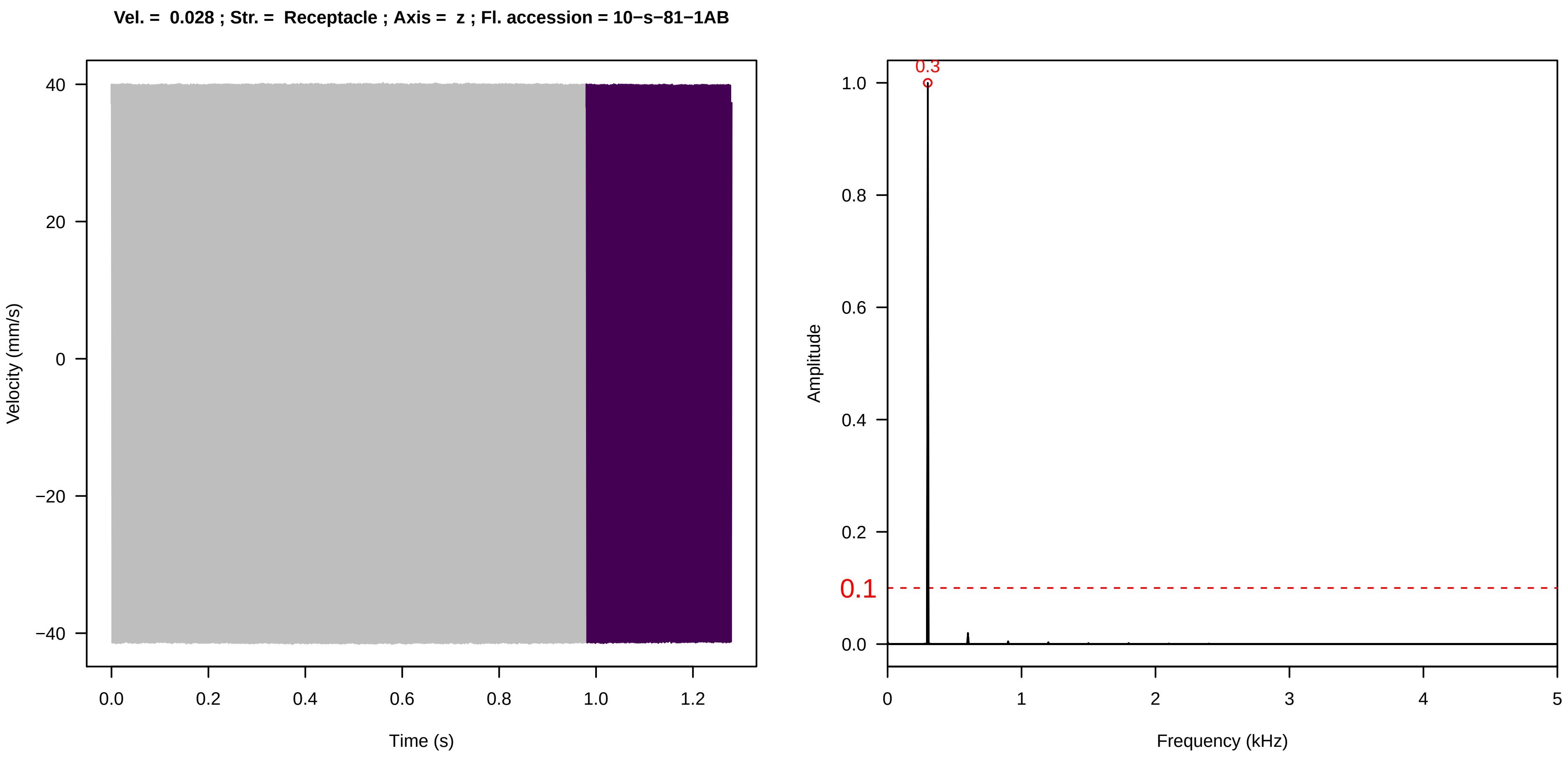
Vel. = 0.028 ; Str. = Receptacle ; Axis = z ; Fl. accession = 10−s−81−1AB
Velocity (mm/s)
Time (s)
Amplitude
Frequency (kHz)
0.3
0.1

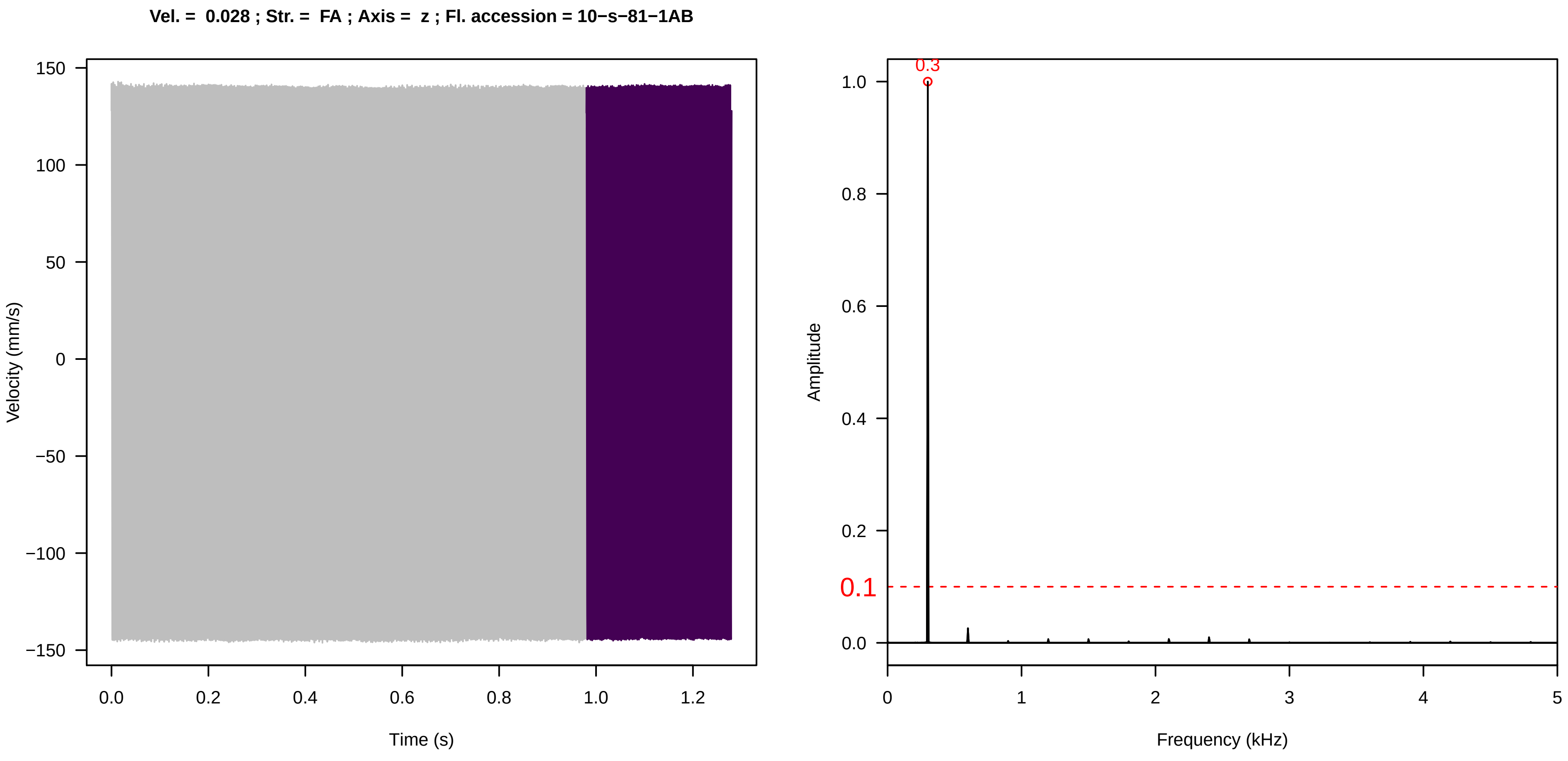
Vel. = 0.028 ; Str. = FA ; Axis = z ; Fl. accession = 10−s−81−1AB
Velocity (mm/s)
Time (s)
Amplitude
Frequency (kHz)
0.3
0.1

**Vel. =  0.028 ; Str. =  Receptacle ; Axis =  z ; Fl. accession = 10−s−81−1AB**

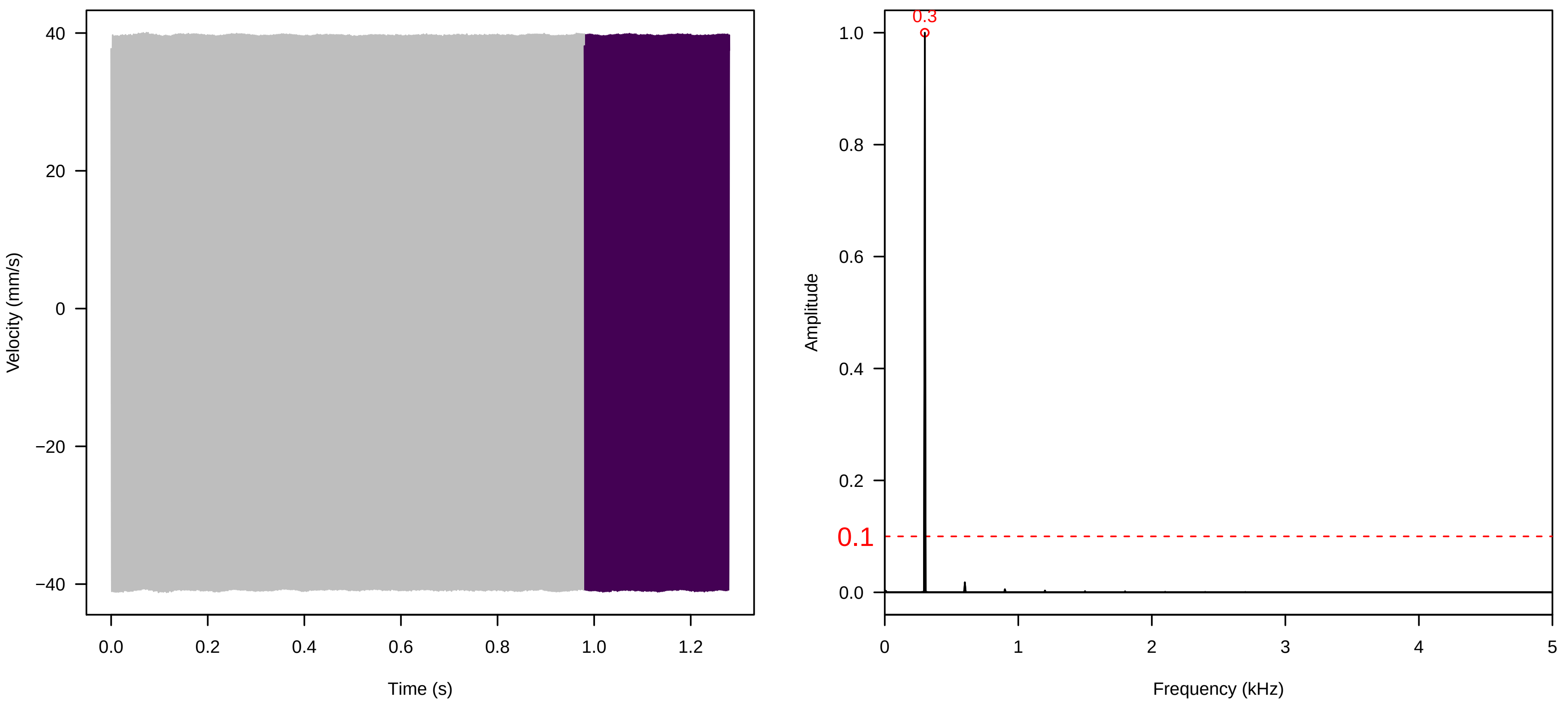


**Vel. =  0.028 ; Str. =  Corolla ; Axis =  z ; Fl. accession = 10−s−81−1AB**

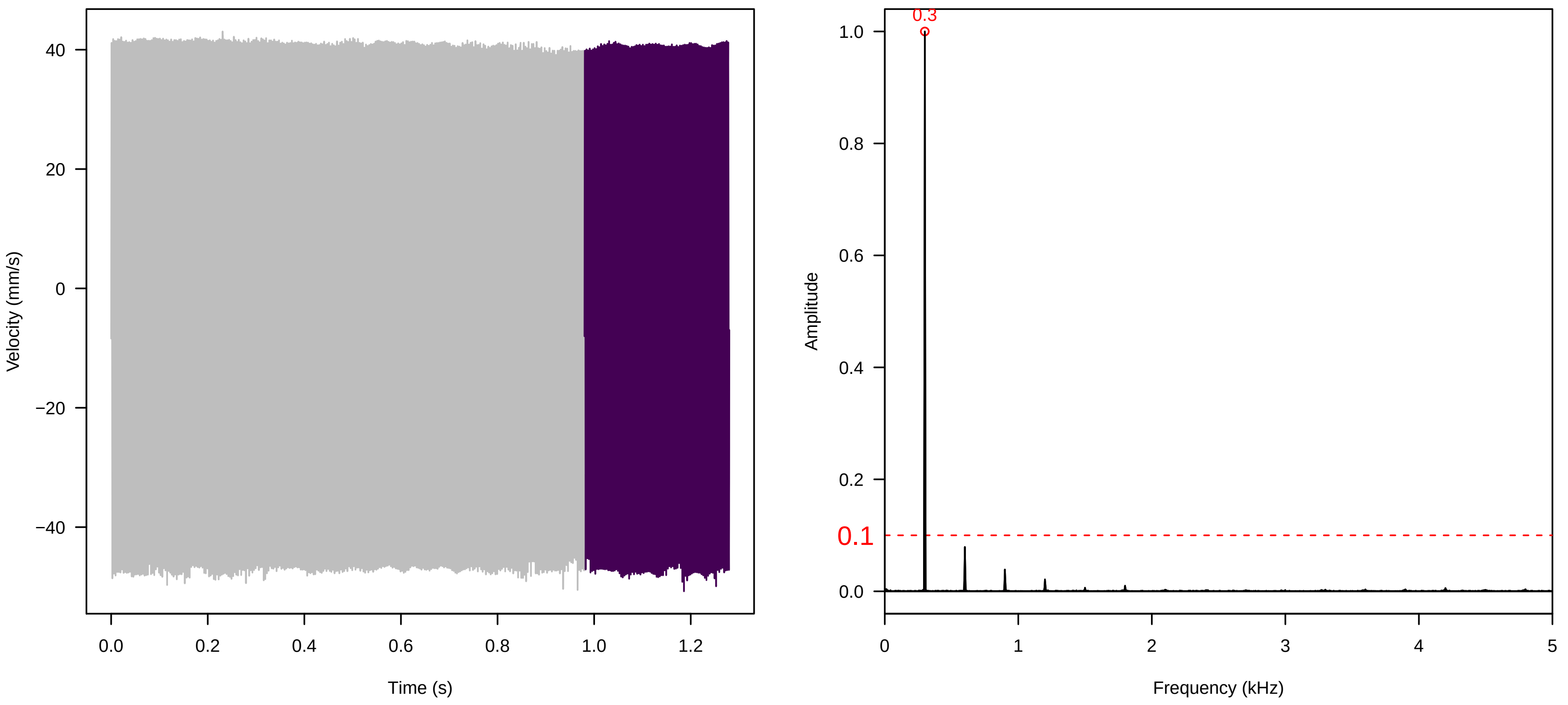


**Vel. =  0.028 ; Str. =  Receptacle ; Axis =  z ; Fl. accession = 10−s−81−1AB**

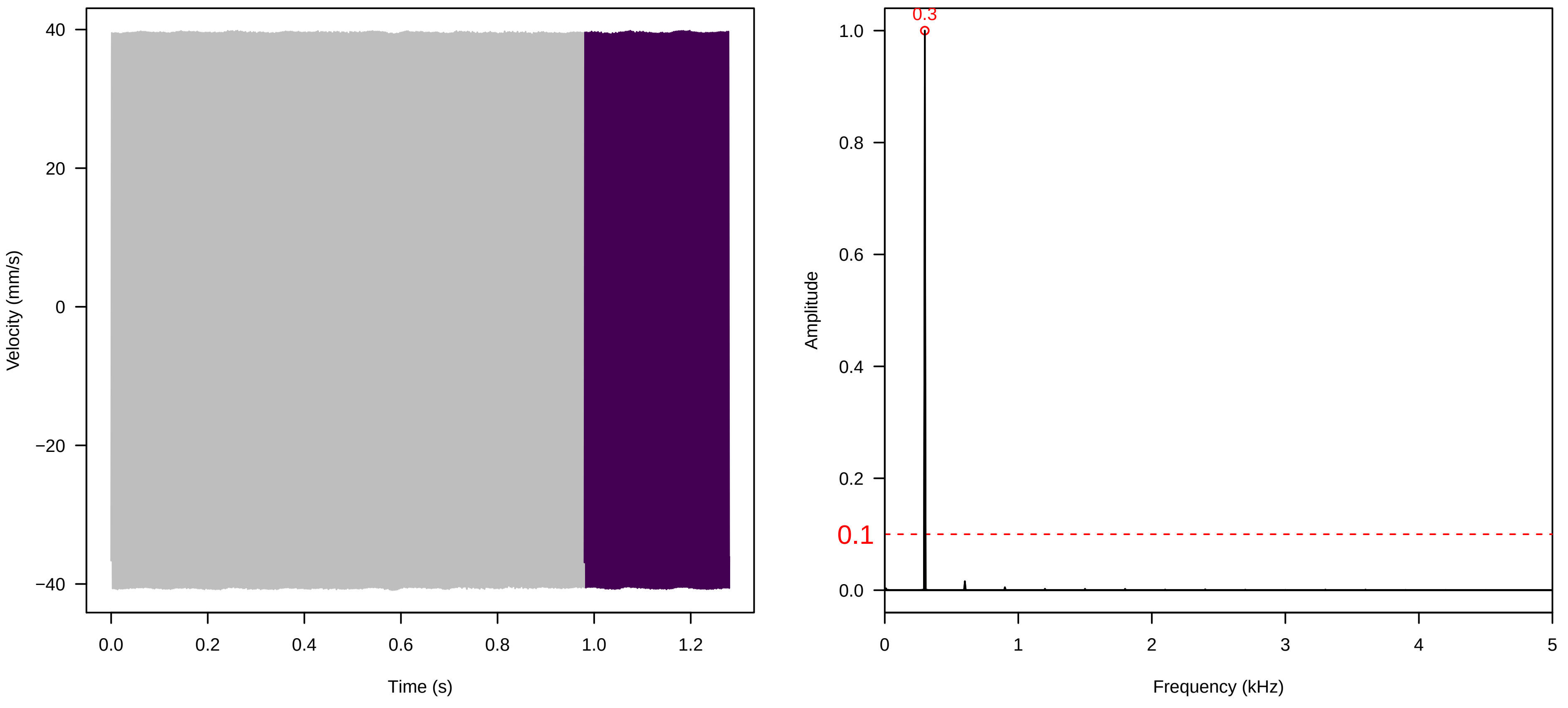

**Vel. = 0.057 ; Str. = Corolla ; Axis = z ; Fl. accession = 10−s−81−1AB**

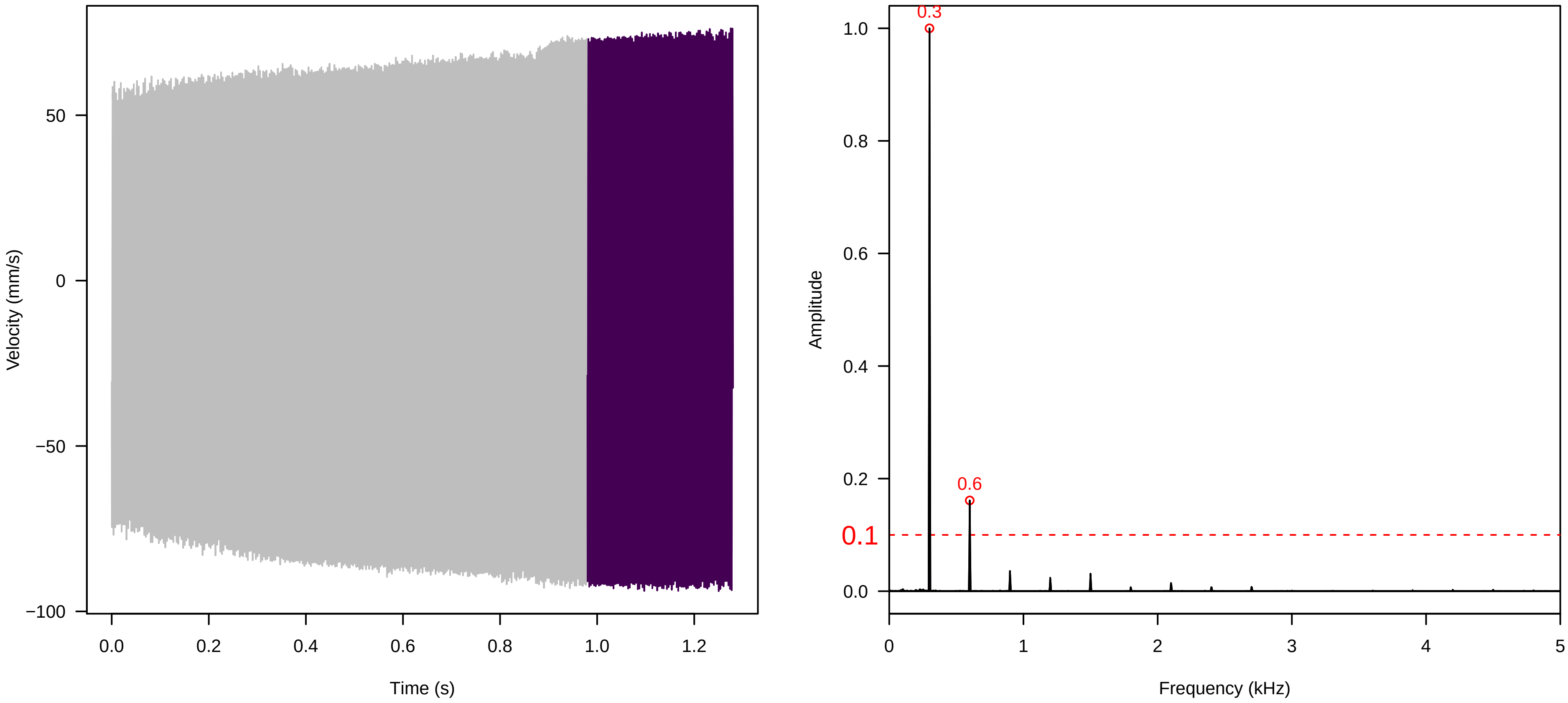


**Vel. = 0.057 ; Str. = Receptacle ; Axis = z ; Fl. accession = 10−s−81−1AB**

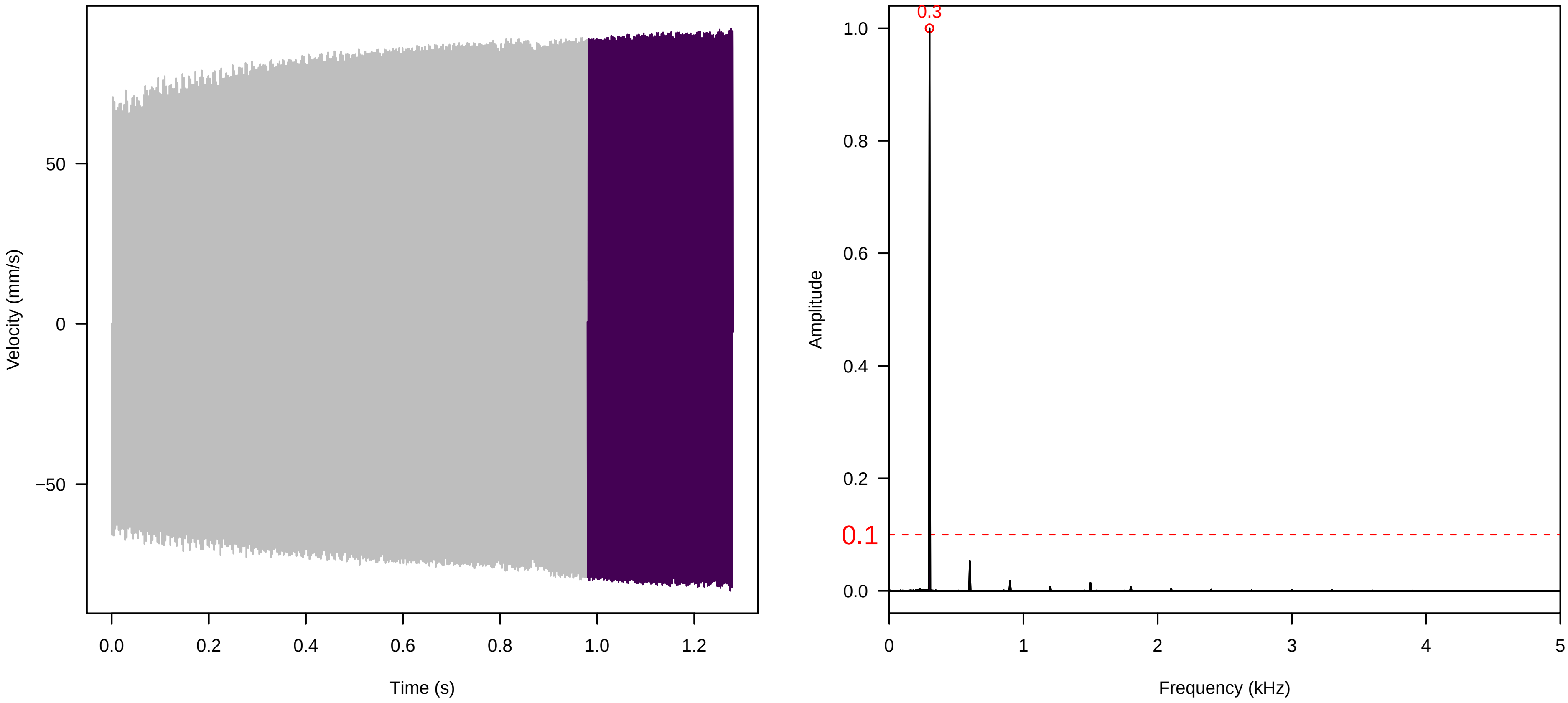


**Vel. = 0.057 ; Str. = FA ; Axis = z ; Fl. accession = 10−s−81−1AB**

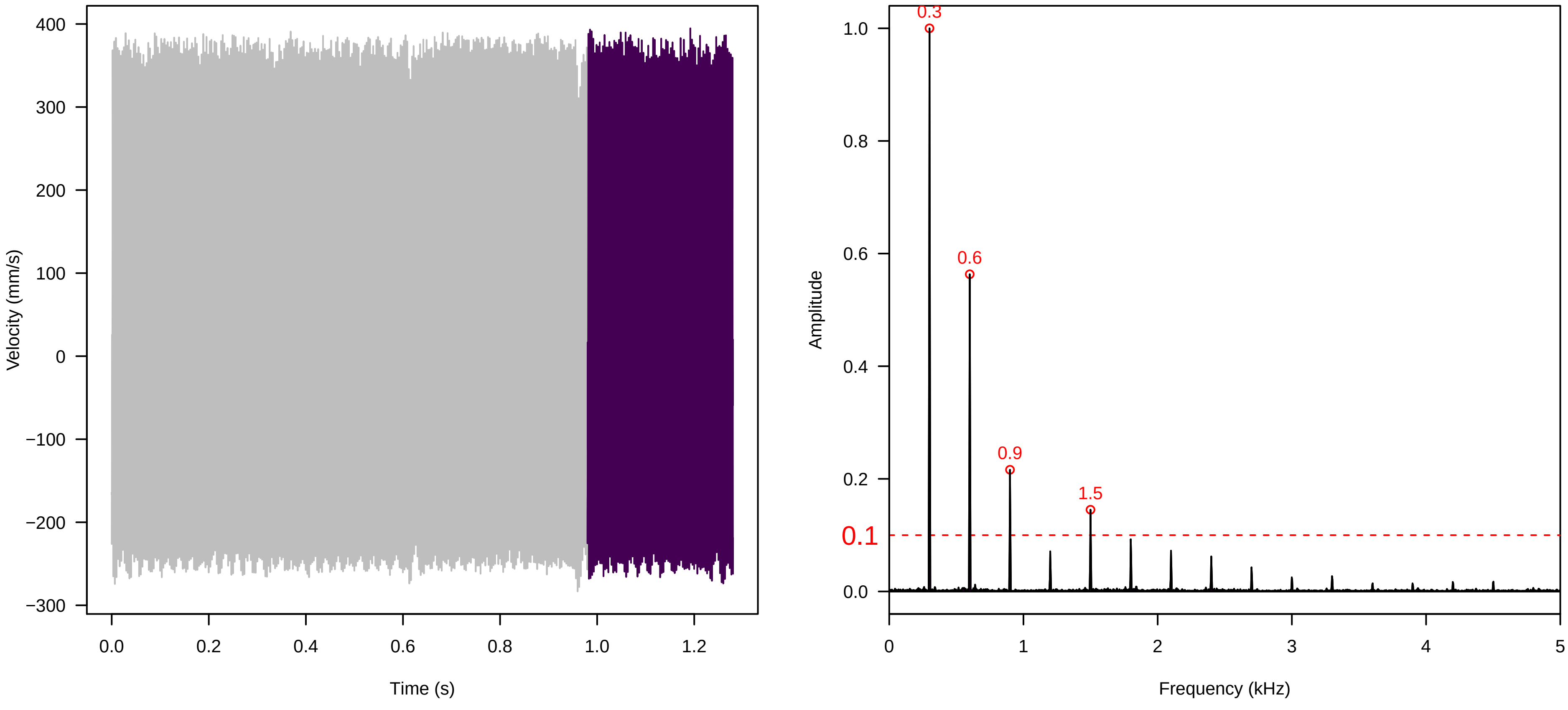

**Vel. = 0.057 ; Str. = Receptacle ; Axis = z ; Fl. accession = 10−s−81−1AB**

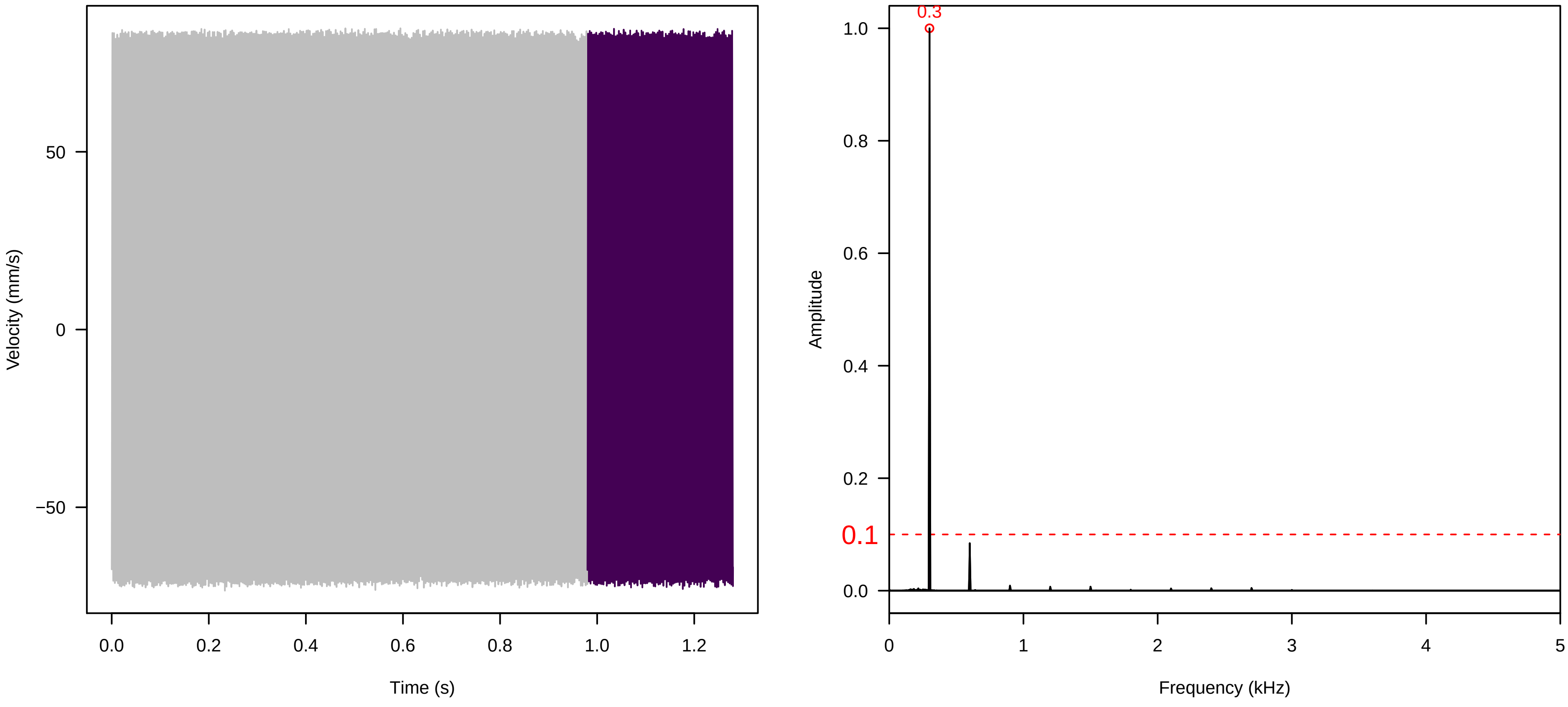


**Vel. = 0.057 ; Str. = PA ; Axis = z ; Fl. accession = 10−s−81−1AB**

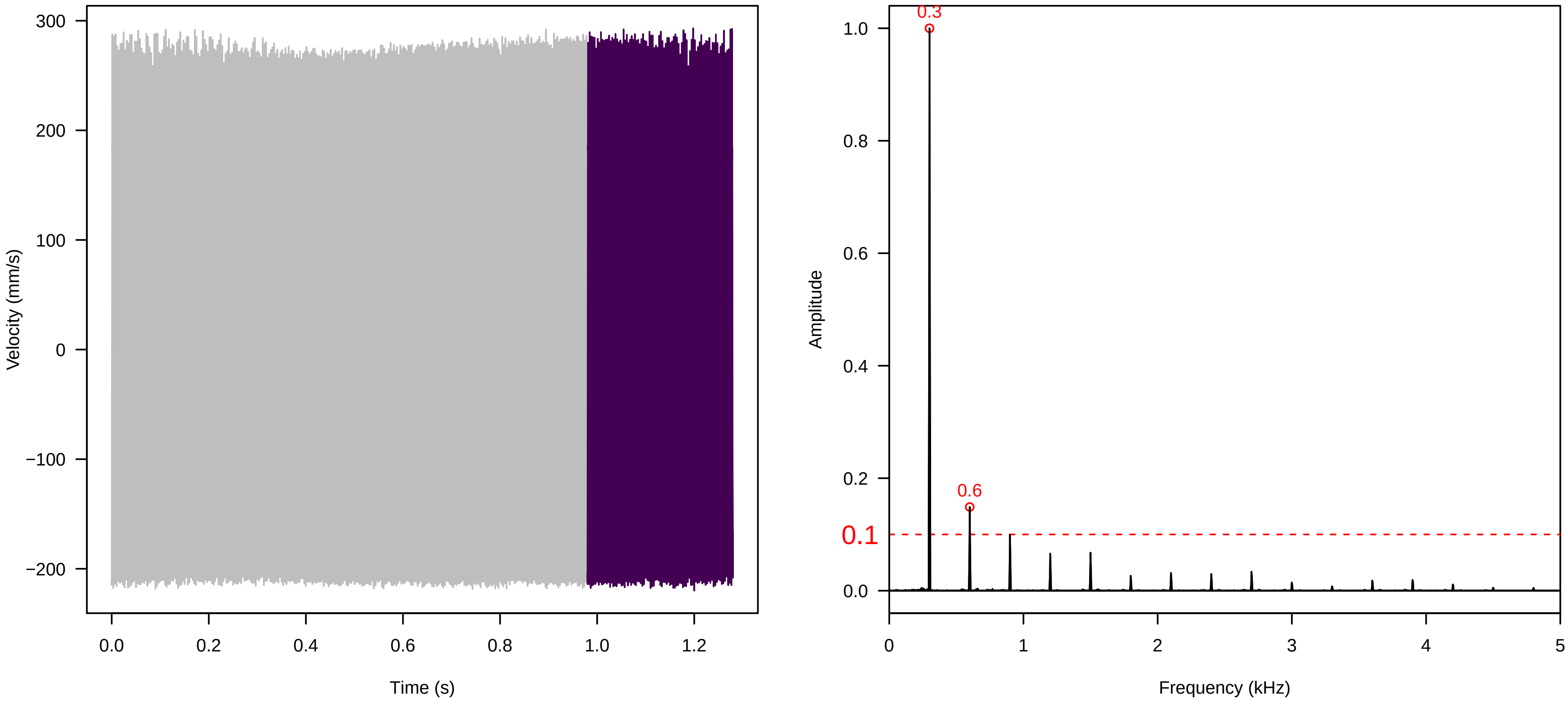


**Vel. = 0.057 ; Str. = Receptacle ; Axis = z ; Fl. accession = 10−s−81−1AB**

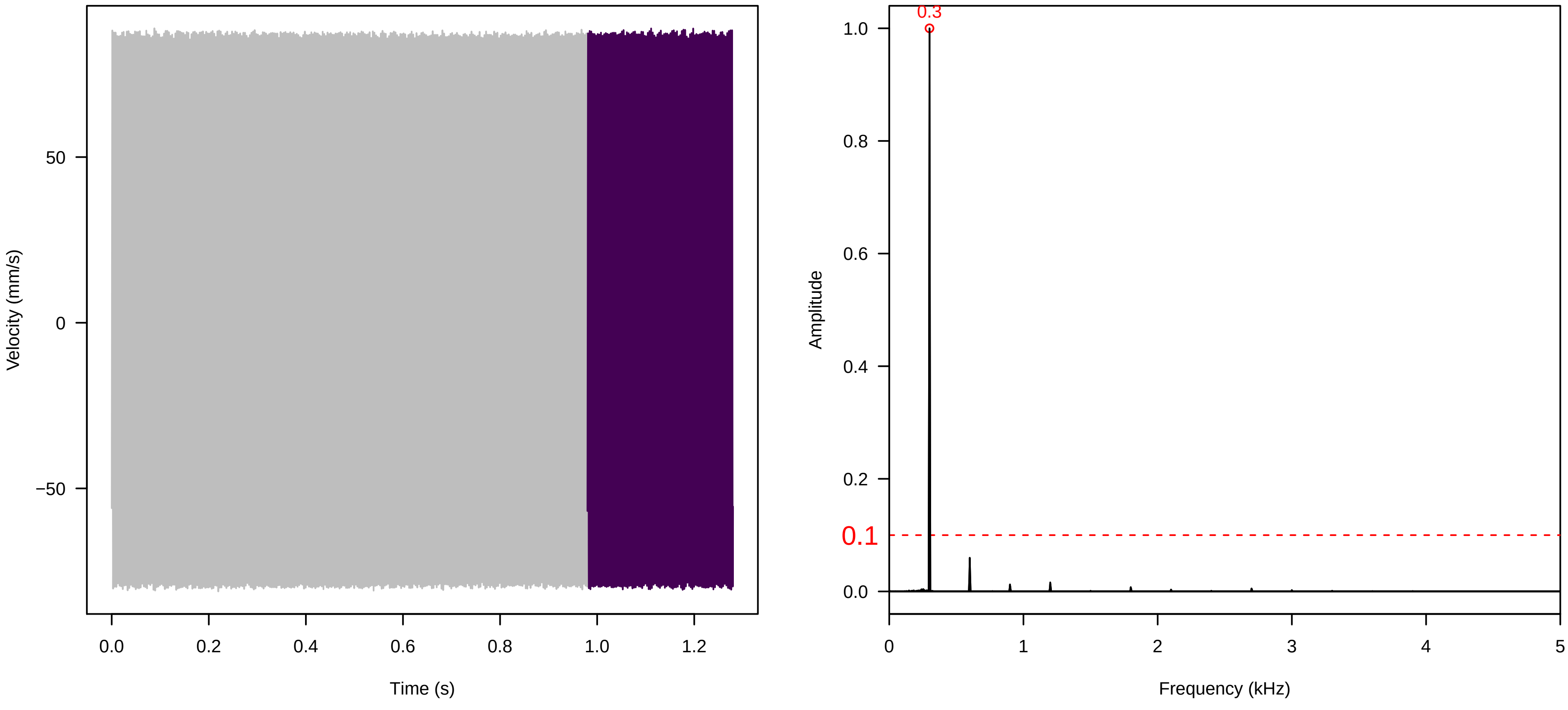

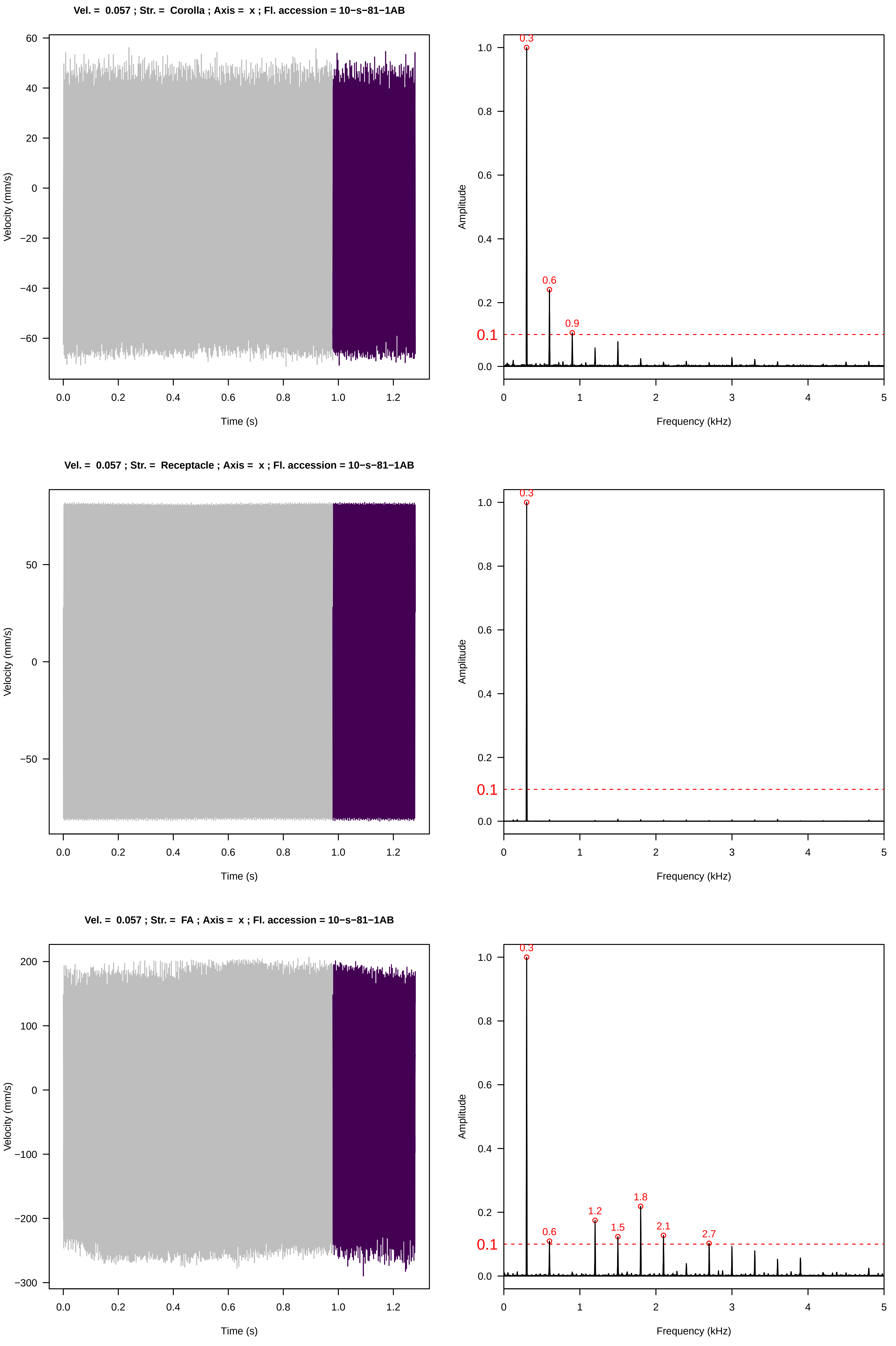
Vel. = 0.057 ; Str. = Corolla ; Axis = x ; Fl. accession = 10−s−81−1AB
Velocity (mm/s)
Time (s)
Amplitude
Frequency (kHz)
0.3
0.6
0.9
0.1
Vel. = 0.057 ; Str. = Receptacle ; Axis = x ; Fl. accession = 10−s−81−1AB
Velocity (mm/s)
Time (s)
Amplitude
Frequency (kHz)
0.3
0.1
Vel. = 0.057 ; Str. = FA ; Axis = x ; Fl. accession = 10−s−81−1AB
Velocity (mm/s)
Time (s)
Amplitude
Frequency (kHz)
0.3
0.6
1.2
1.5
1.8
2.1
2.7
0.1

**Vel. = 0.057 ; Str. = Receptacle ; Axis = x ; Fl. accession = 10−s−81−1AB**

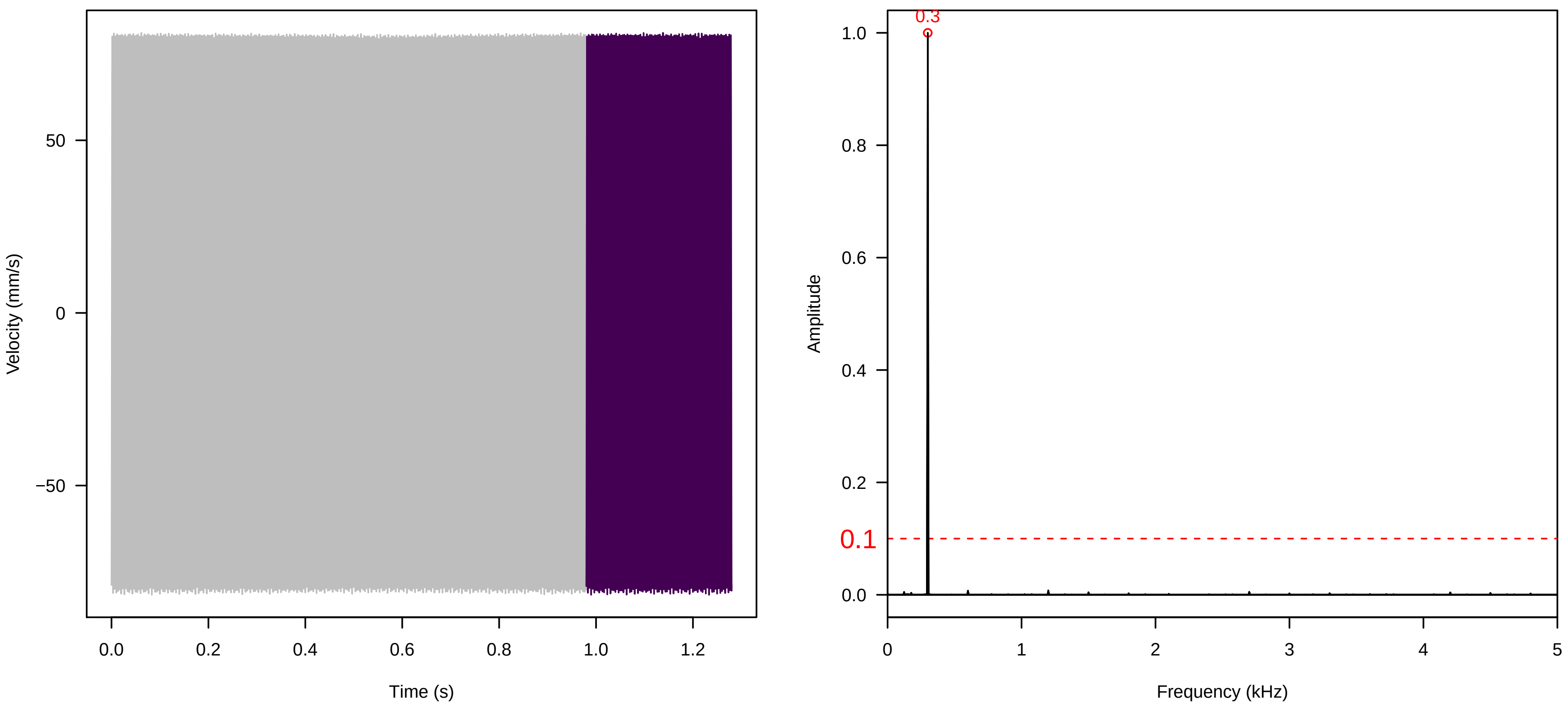


**Vel. = 0.057 ; Str. = PA ; Axis = x ; Fl. accession = 10−s−81−1AB**

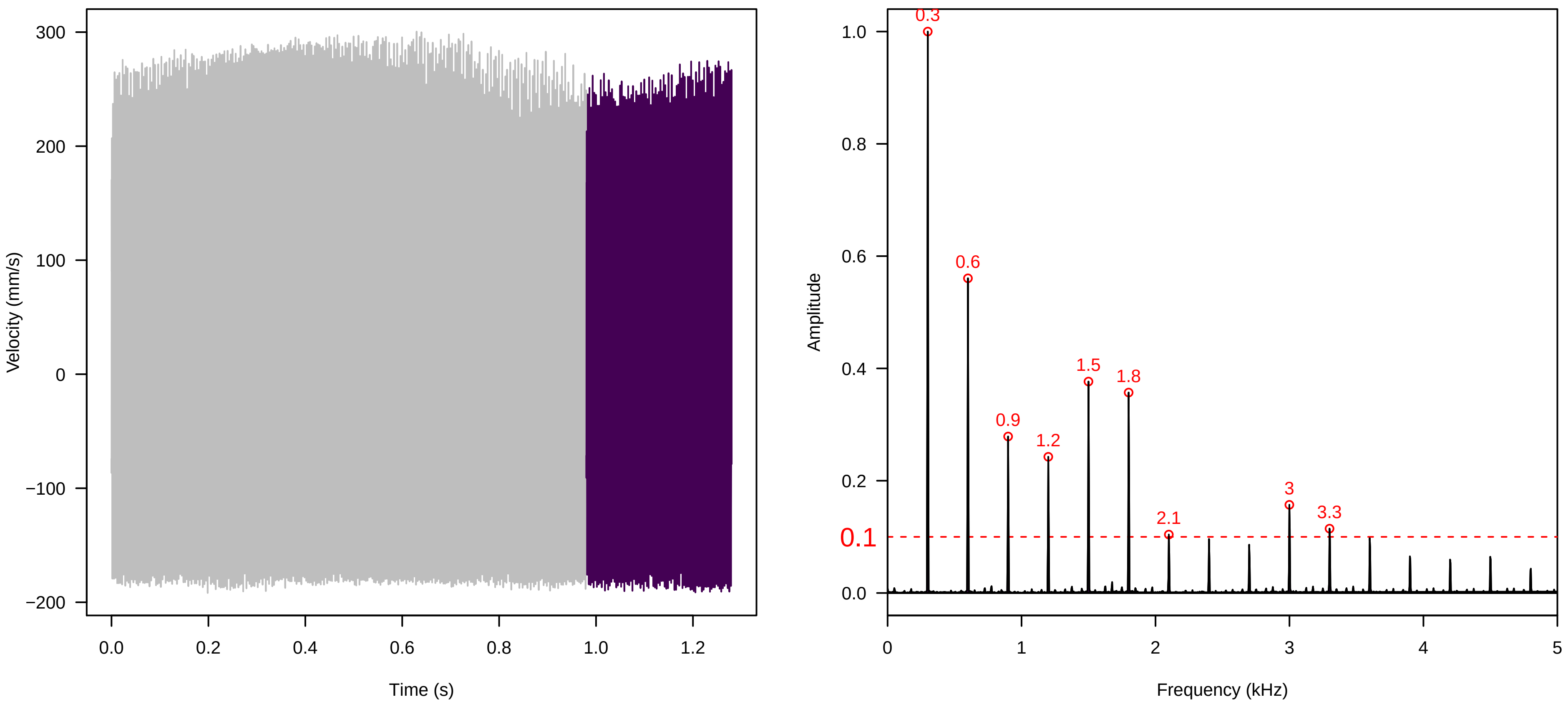


**Vel. = 0.057 ; Str. = Receptacle ; Axis = x ; Fl. accession = 10−s−81−1AB**

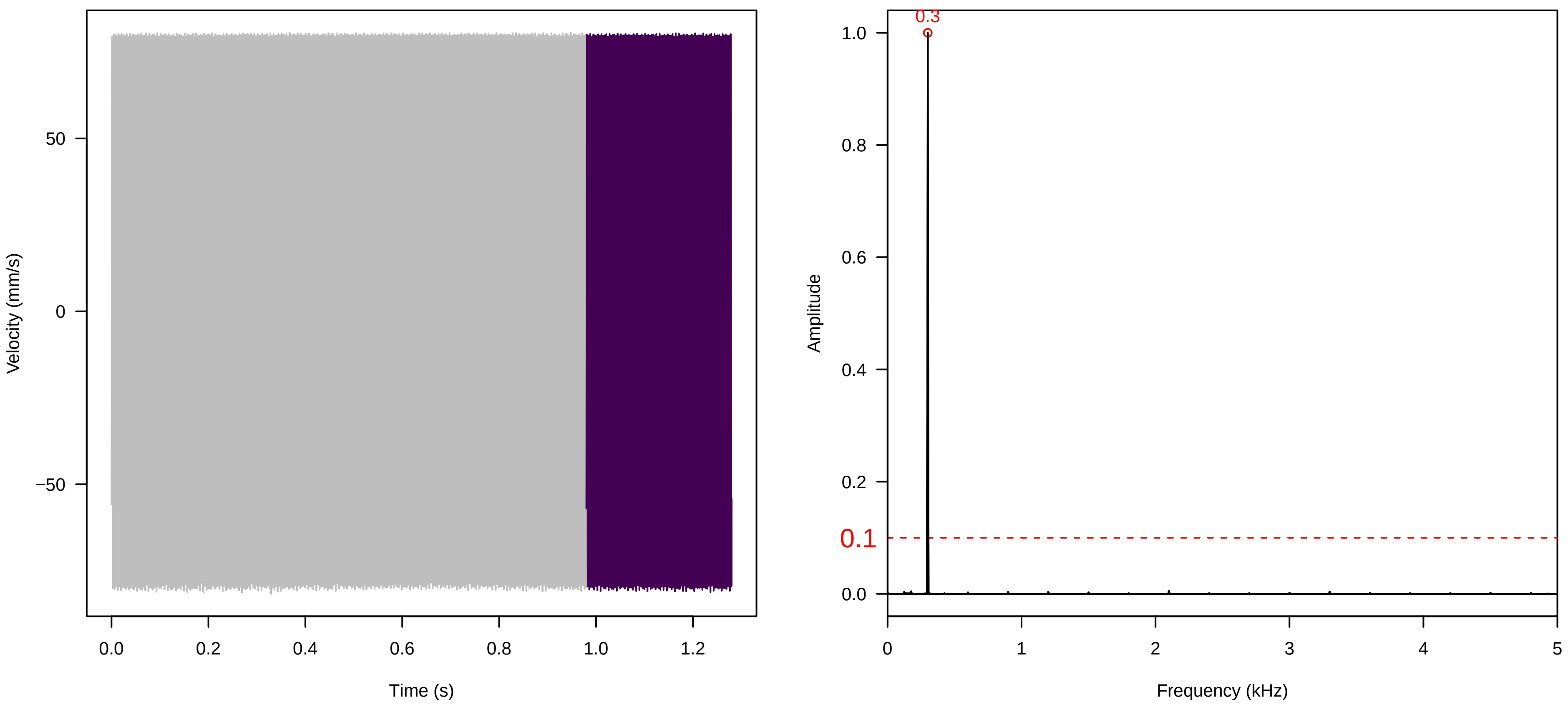

**Vel. = 0.028 ; Str. = PA ; Axis = x ; Fl. accession = 10−s−81−1AB**

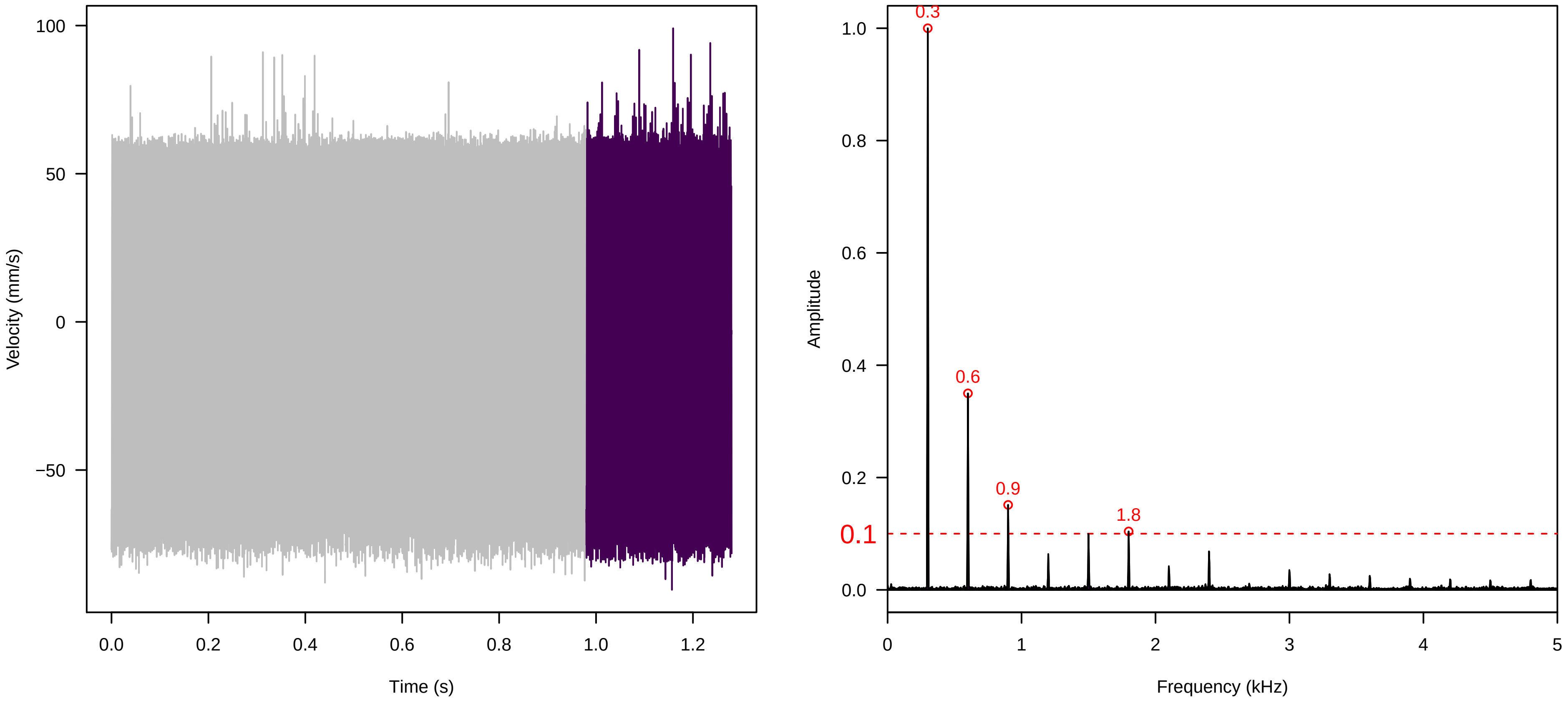


**Vel. = 0.028 ; Str. = Receptacle ; Axis = x ; Fl. accession = 10−s−81−1AB**

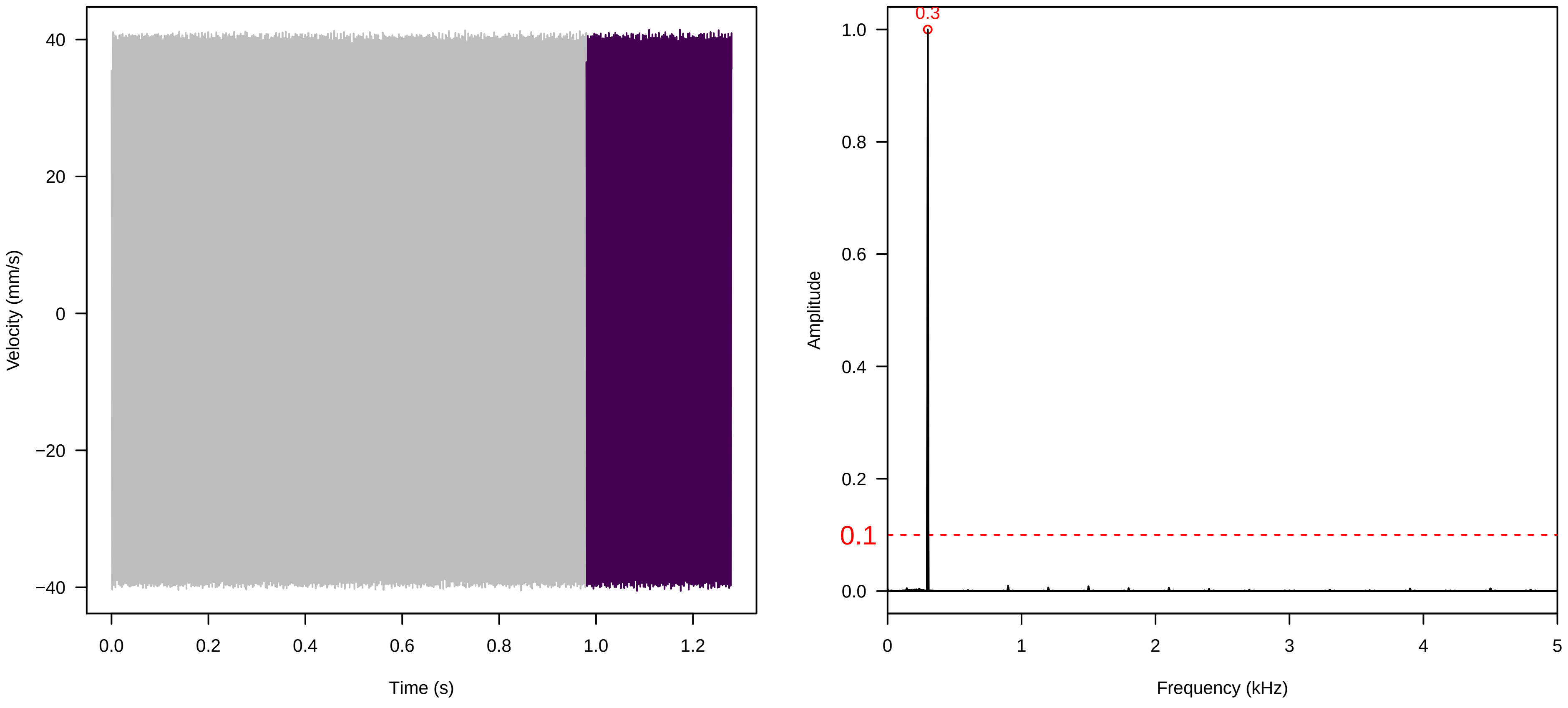


**Vel. = 0.028 ; Str. = FA ; Axis = x ; Fl. accession = 10−s−81−1AB**

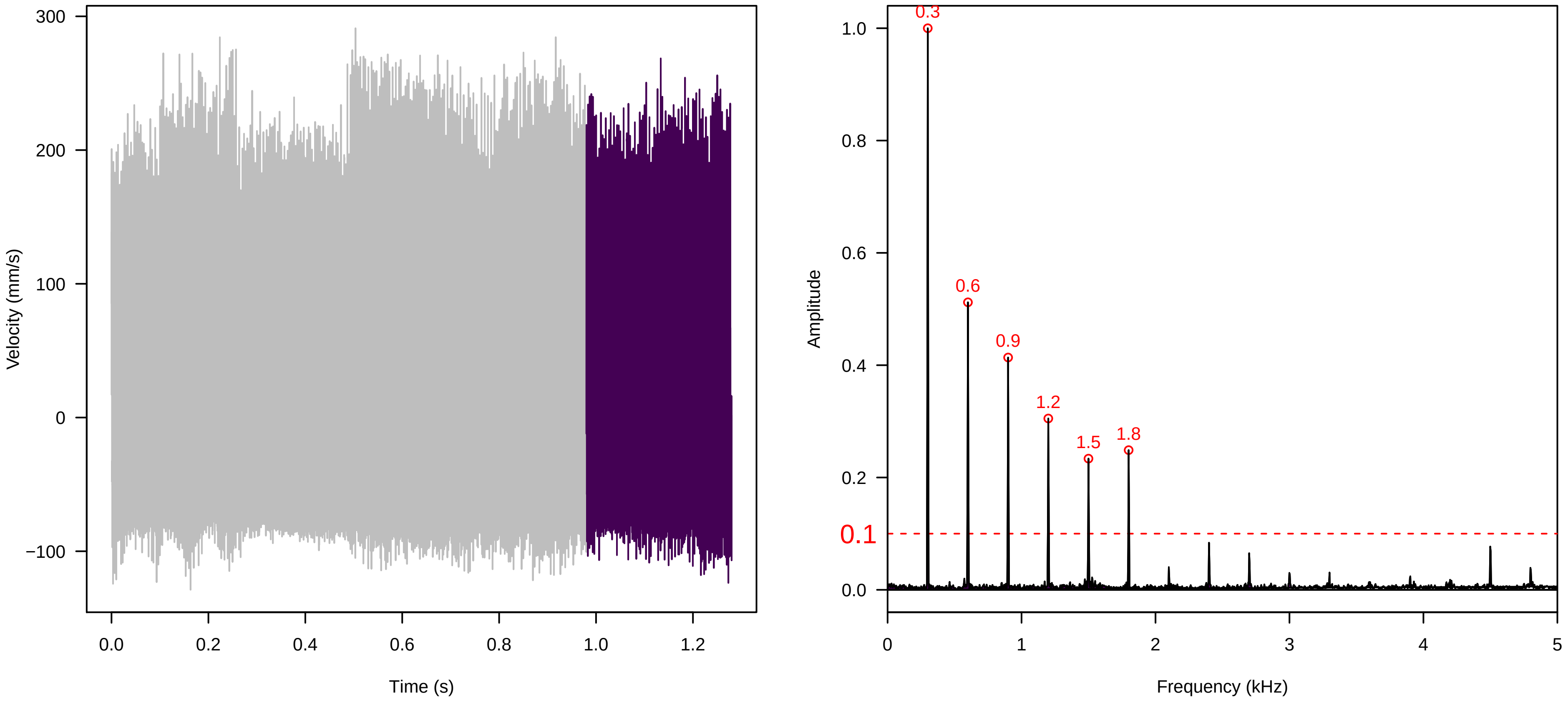

**Vel. =  0.028 ; Str. =  Receptacle ; Axis =  x ; Fl. accession = 10−s−81−1AB**

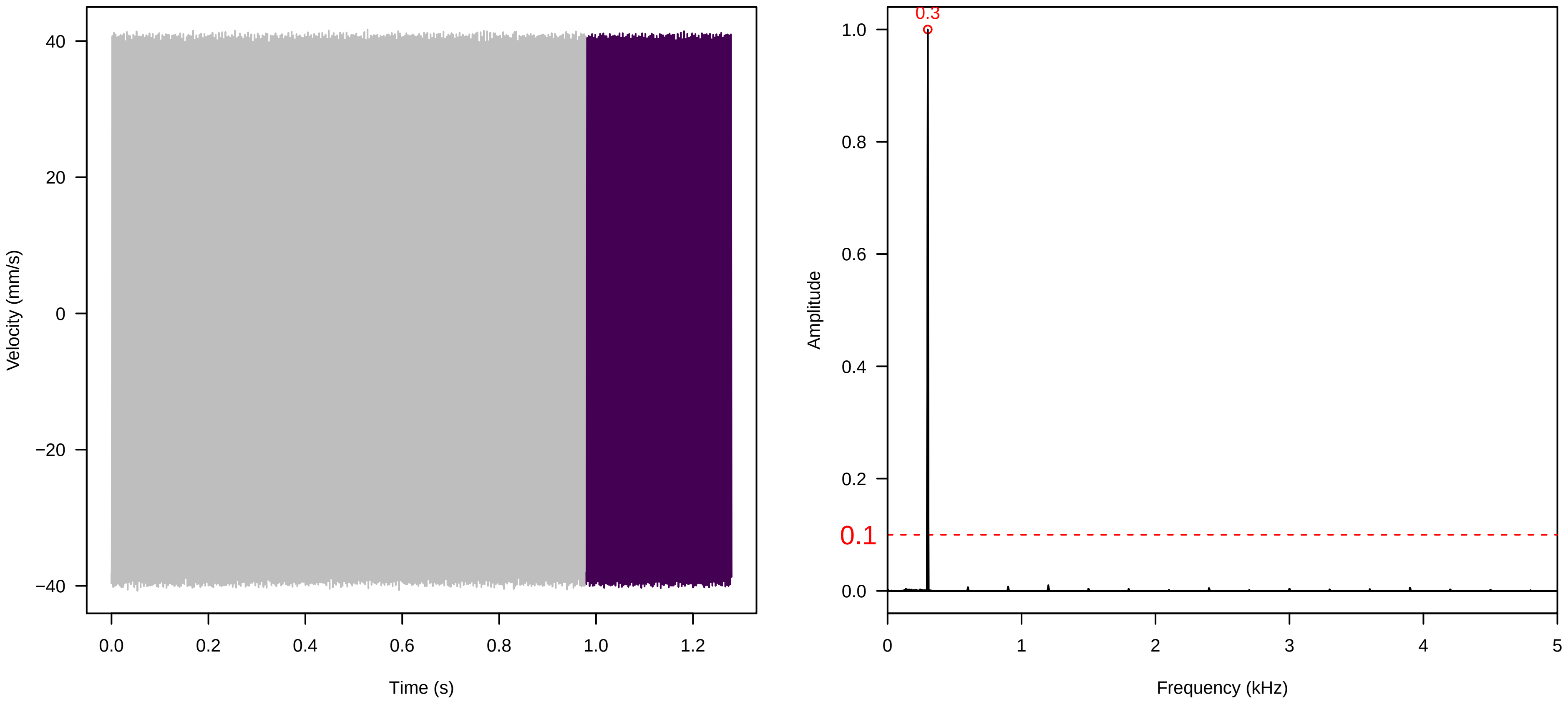


**Vel. =  0.028 ; Str. =  Corolla ; Axis =  x ; Fl. accession = 10−s−81−1AB**

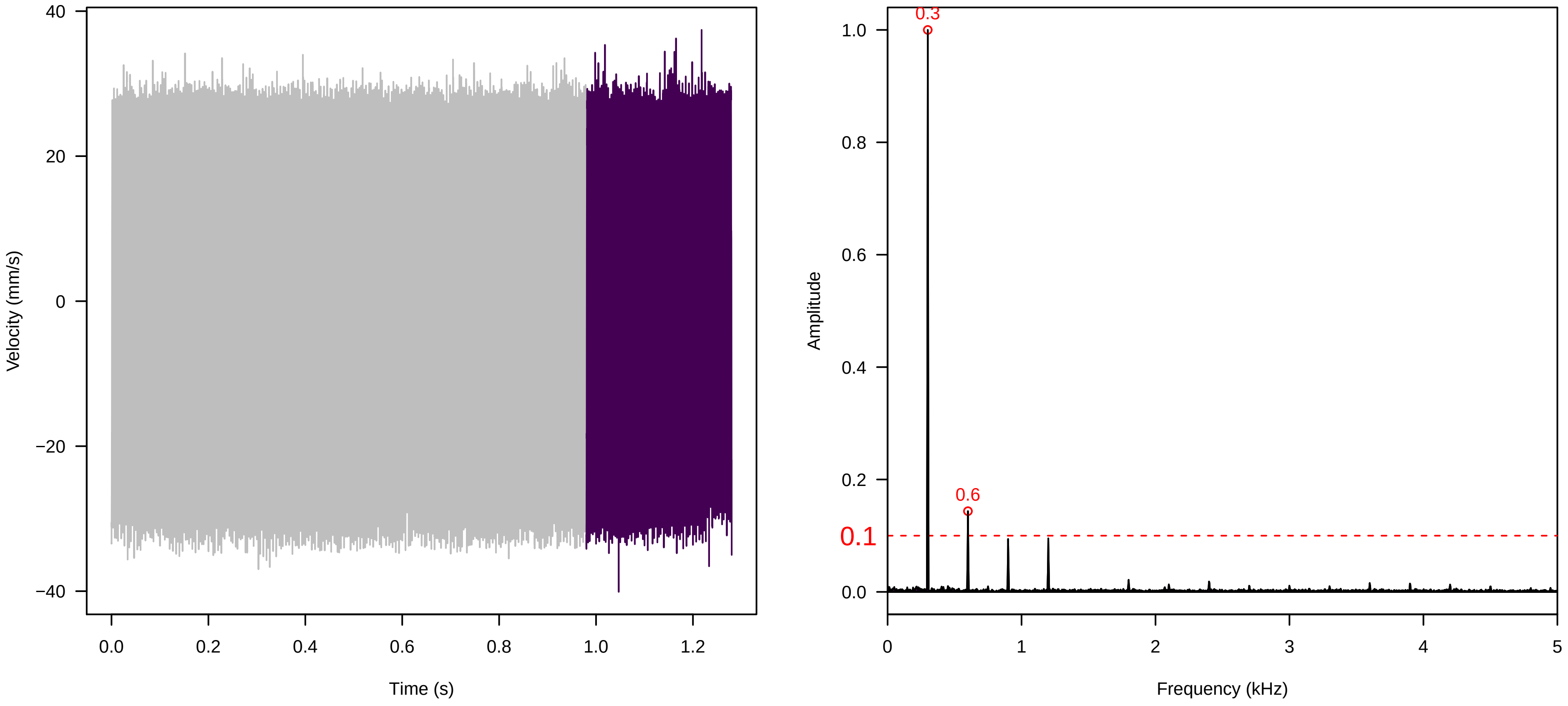


**Vel. =  0.028 ; Str. =  Receptacle ; Axis =  x ; Fl. accession = 10−s−81−1AB**

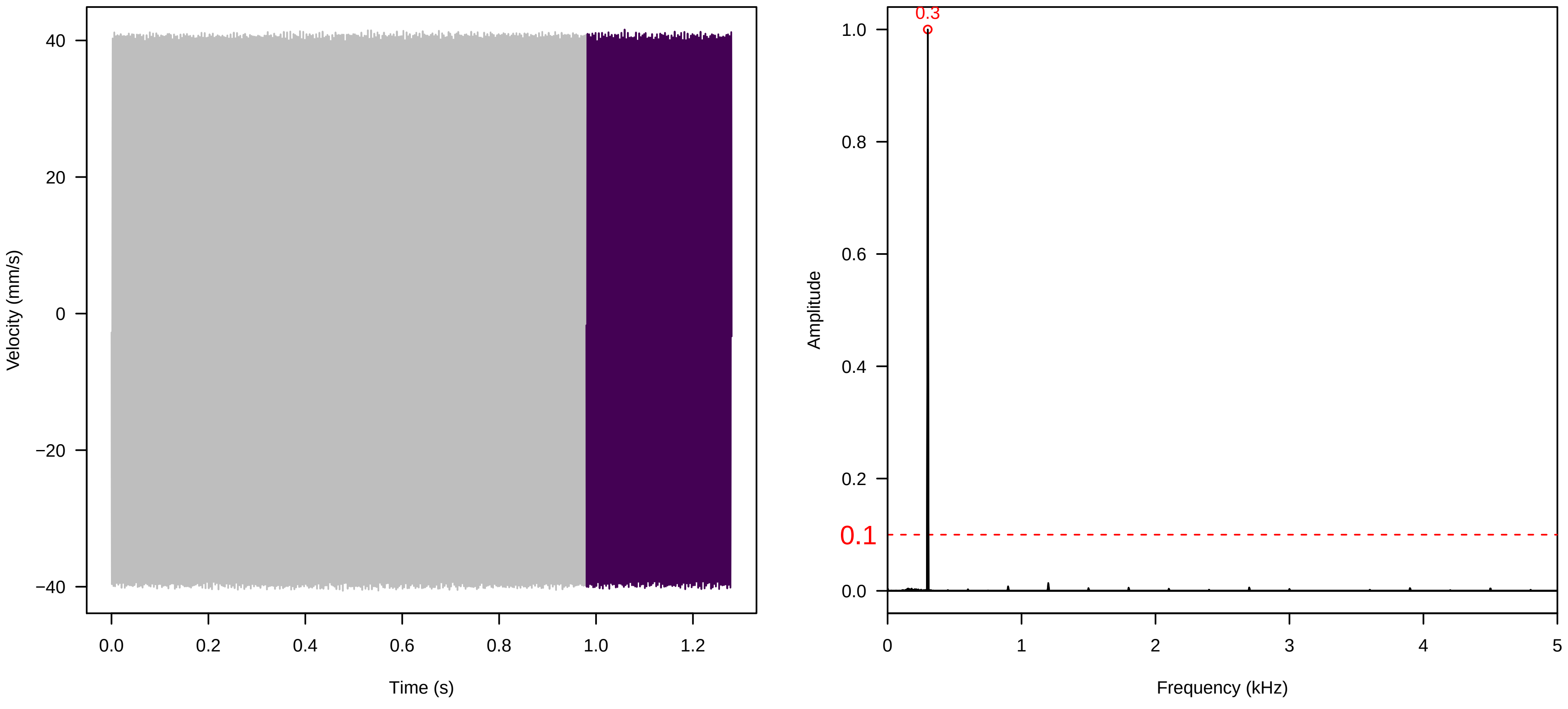

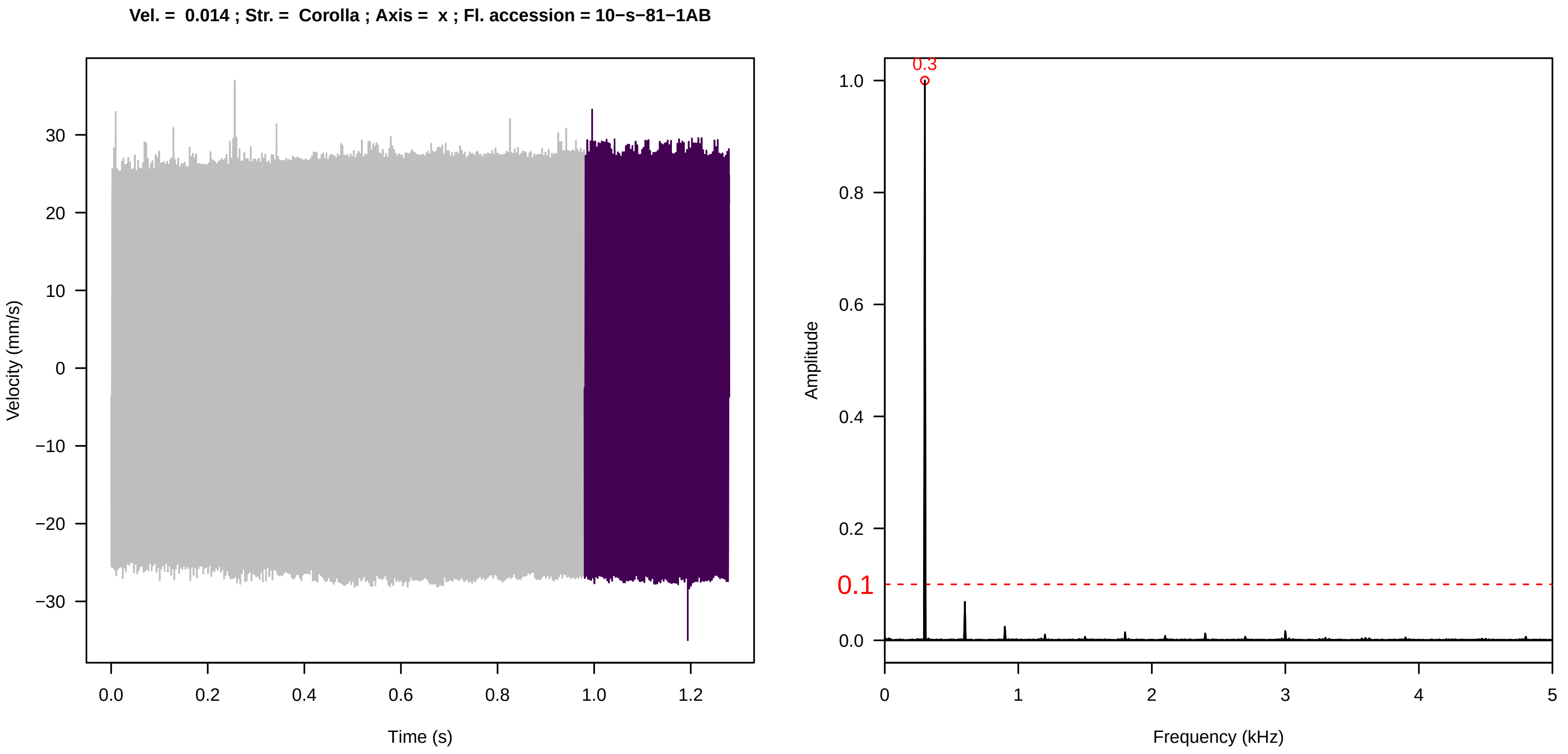
Vel. = 0.014 ; Str. = Corolla ; Axis = x ; Fl. accession = 10−s−81−1AB
Velocity (mm/s)
Time (s)
Amplitude
Frequency (kHz)
0.3
0.1

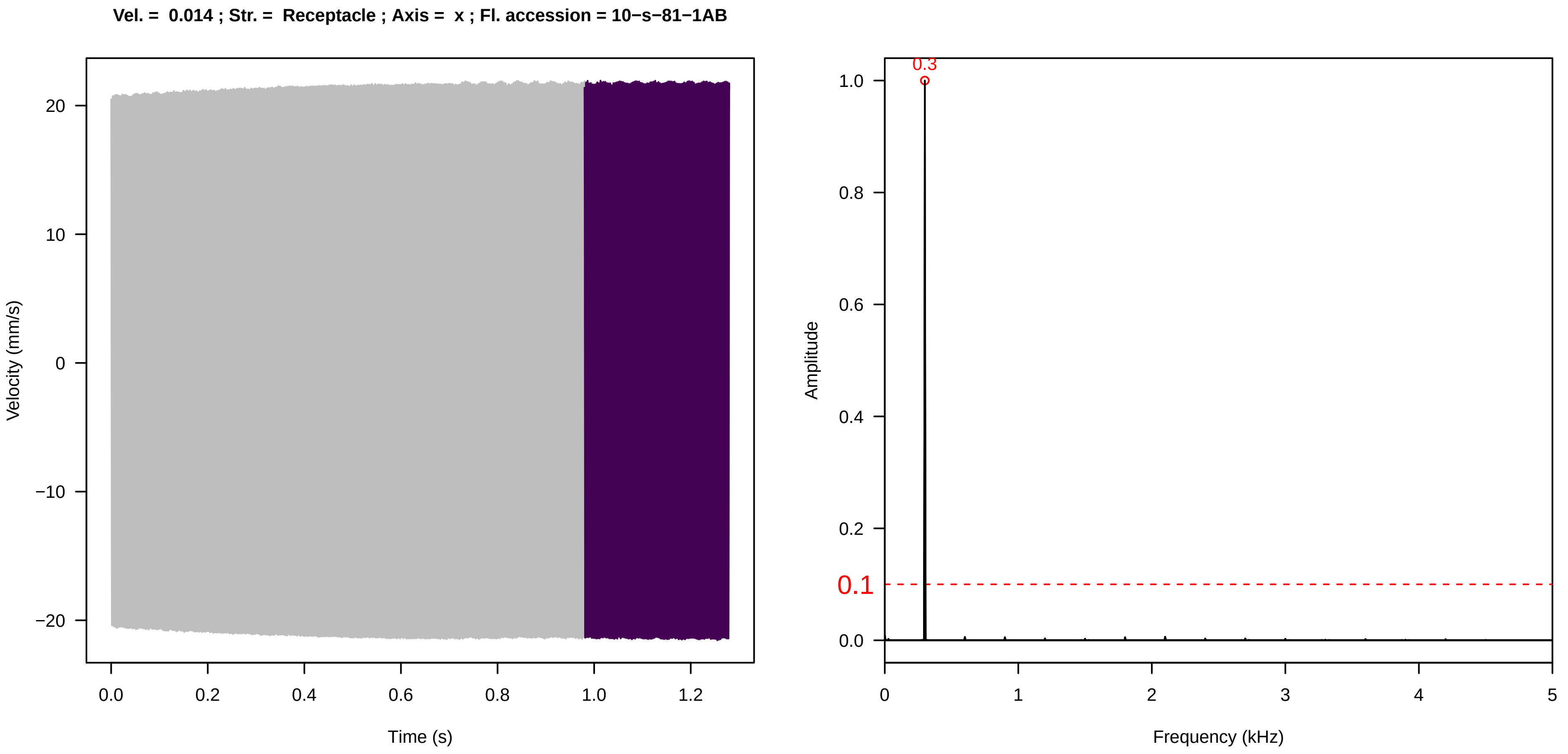
Vel. = 0.014 ; Str. = Receptacle ; Axis = x ; Fl. accession = 10−s−81−1AB
Velocity (mm/s)
Time (s)
Amplitude
Frequency (kHz)
0.3
0.1

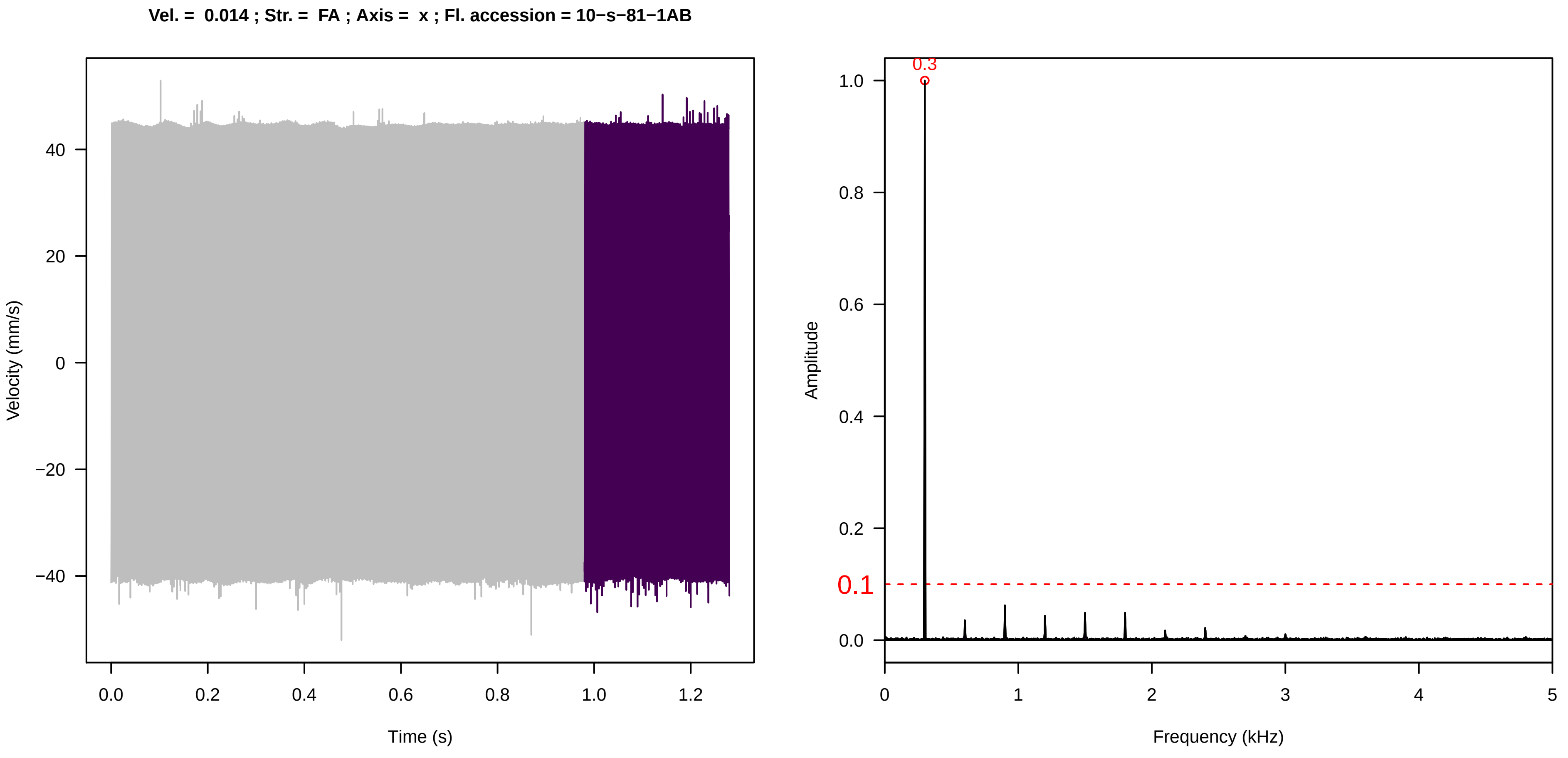
Vel. = 0.014 ; Str. = FA ; Axis = x ; Fl. accession = 10−s−81−1AB
Velocity (mm/s)
Time (s)
Amplitude
Frequency (kHz)
0.3
0.1

**Vel. = 0.014 ; Str. = Receptacle ; Axis = x ; Fl. accession = 10−s−81−1AB**

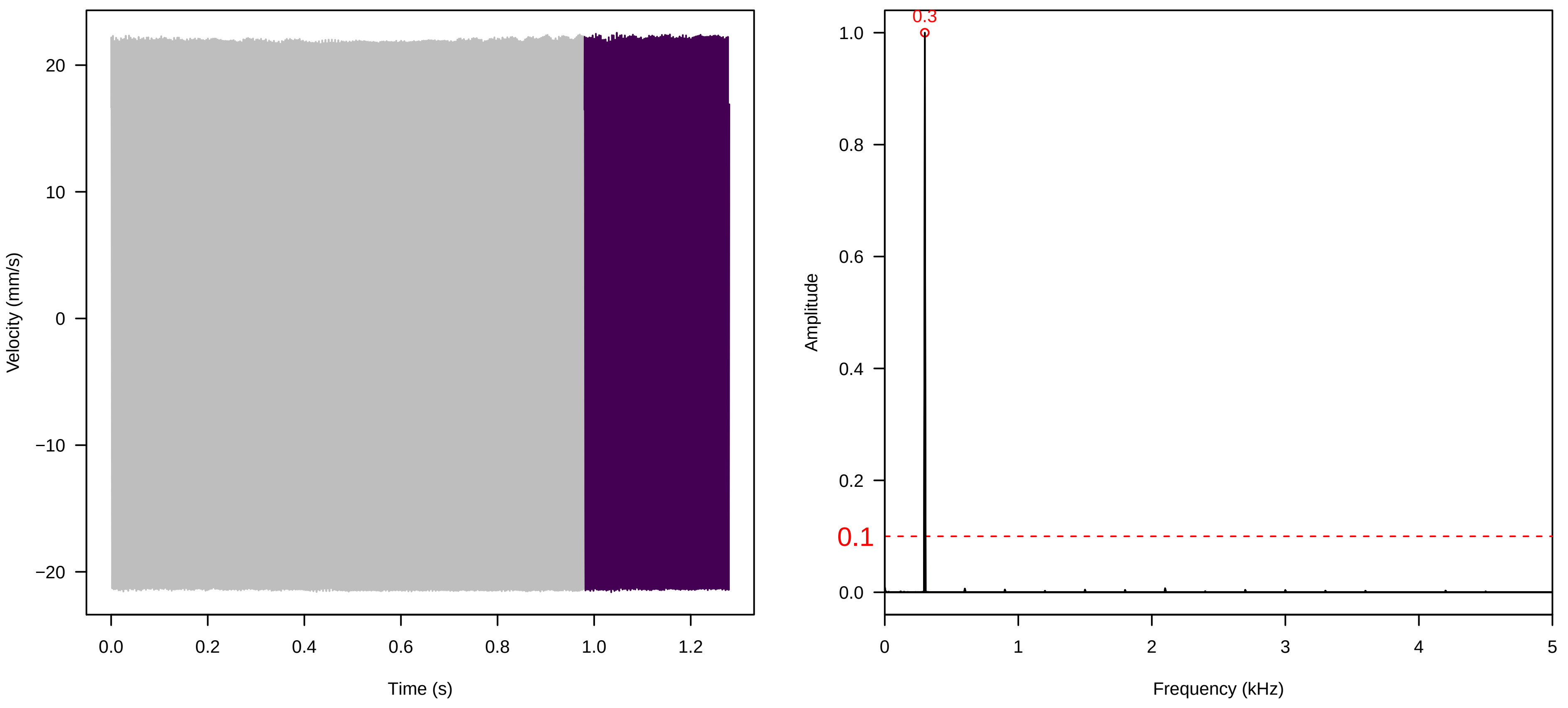


**Vel. = 0.014 ; Str. = PA ; Axis = x ; Fl. accession = 10−s−81−1AB**

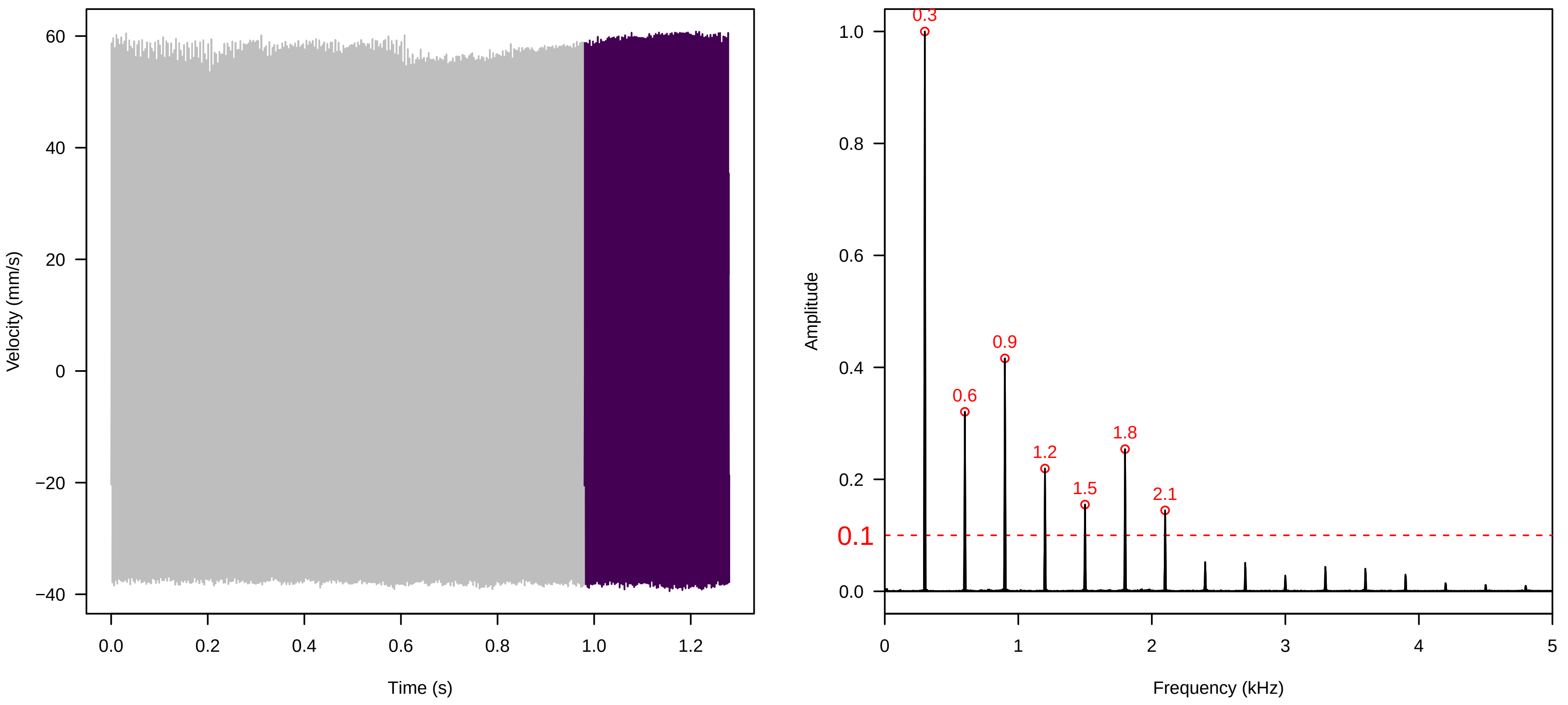


**Vel. = 0.014 ; Str. = Receptacle ; Axis = x ; Fl. accession = 10−s−81−1AB**

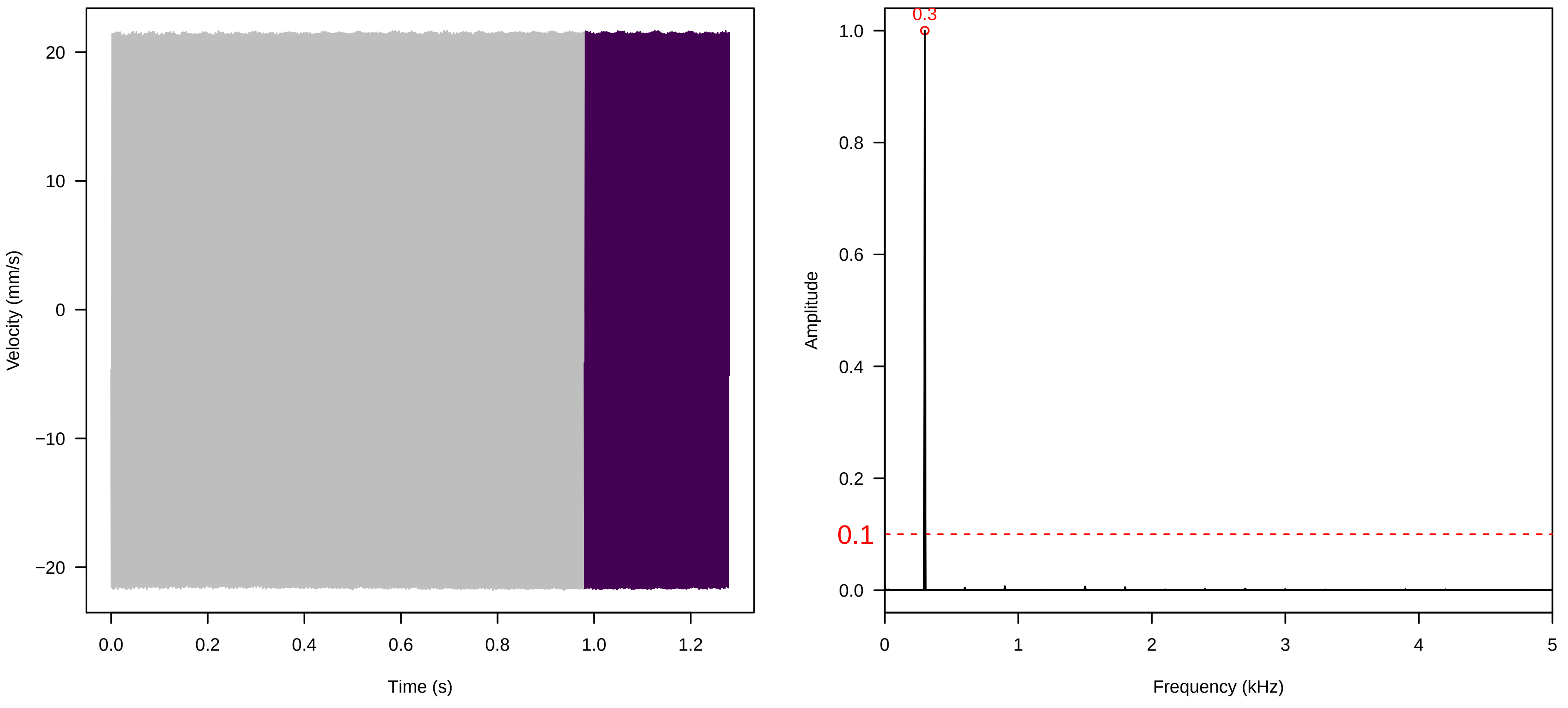

**Vel. = 0.014 ; Str. = Corolla ; Axis = x ; Fl. accession = 10−s−81−8**

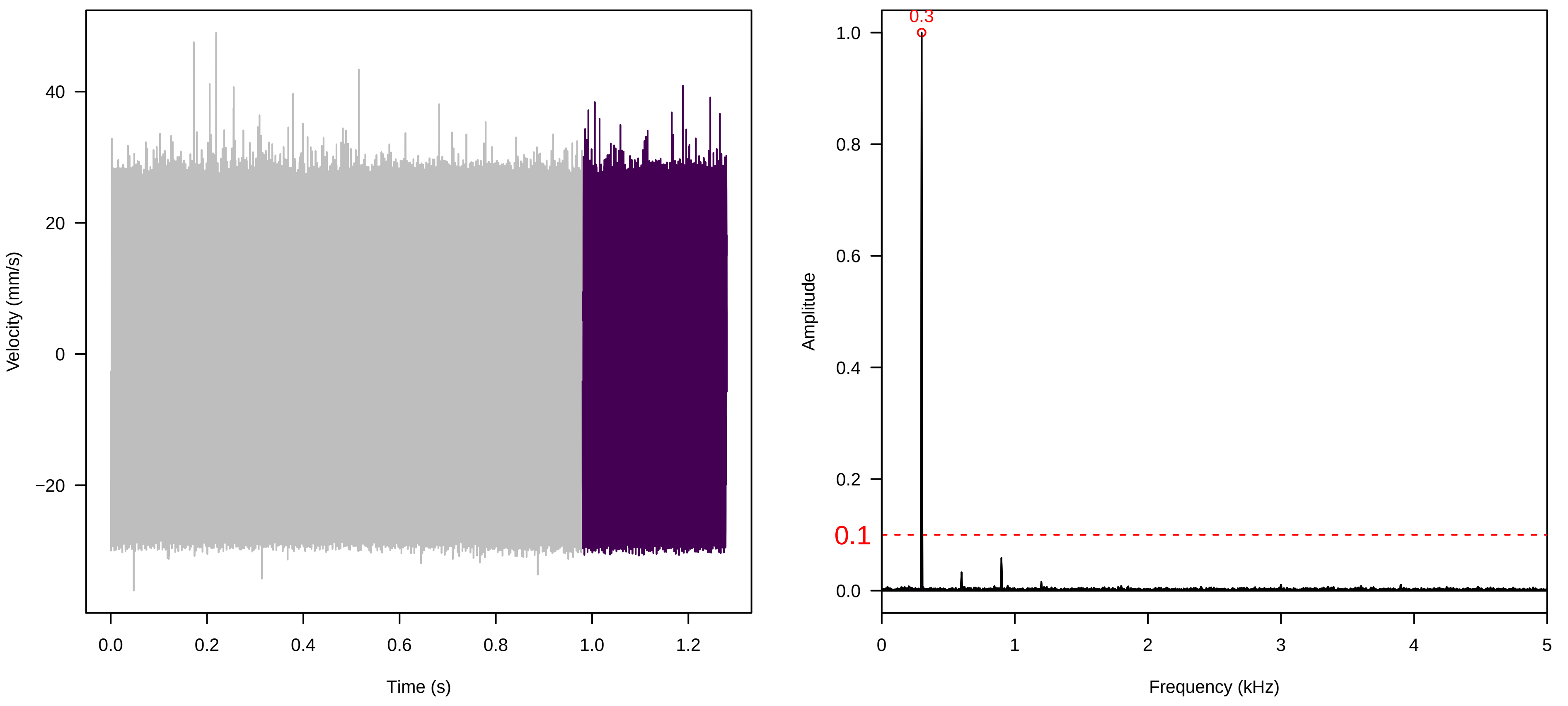


**Vel. = 0.014 ; Str. = Receptacle ; Axis = x ; Fl. accession = 10−s−81−8**

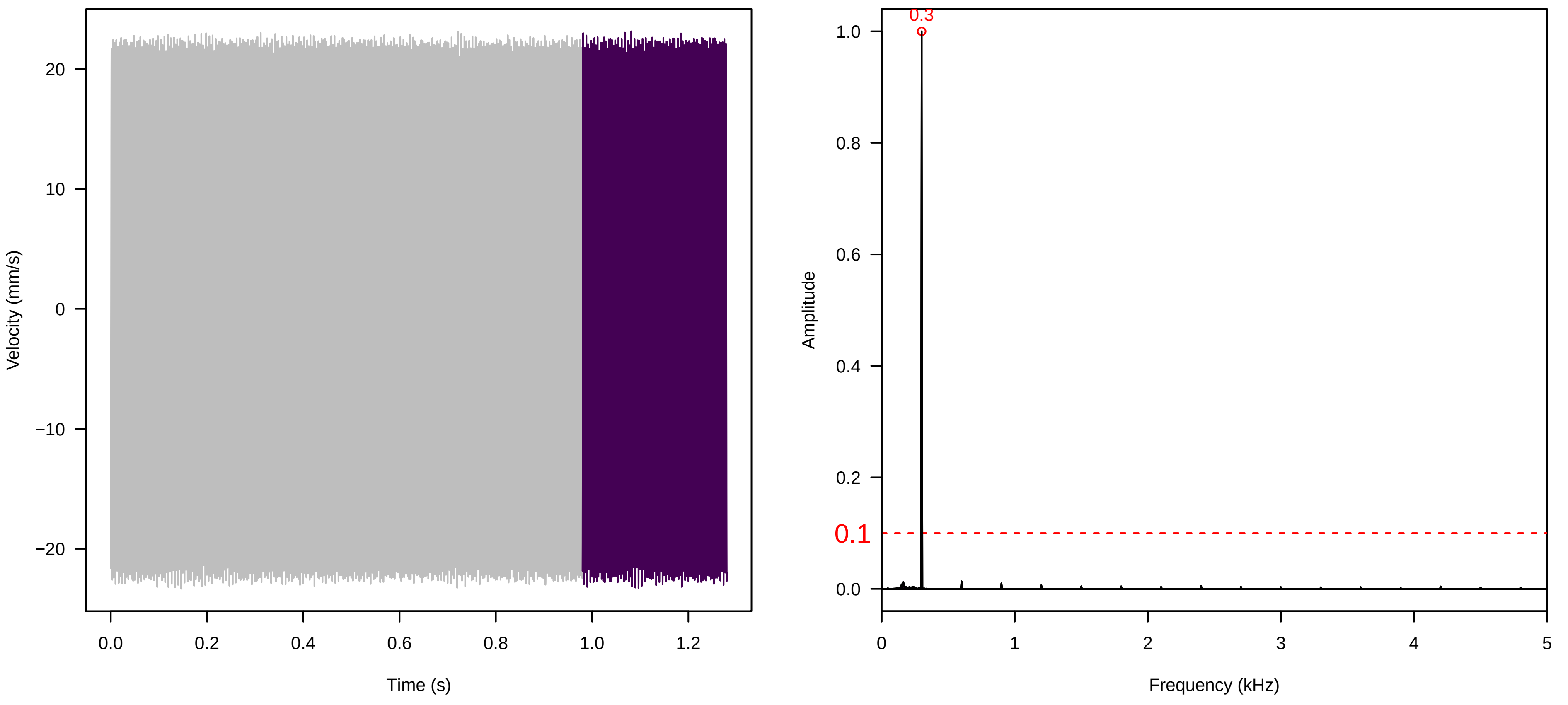


**Vel. = 0.014 ; Str. = FA ; Axis = x ; Fl. accession = 10−s−81−8**

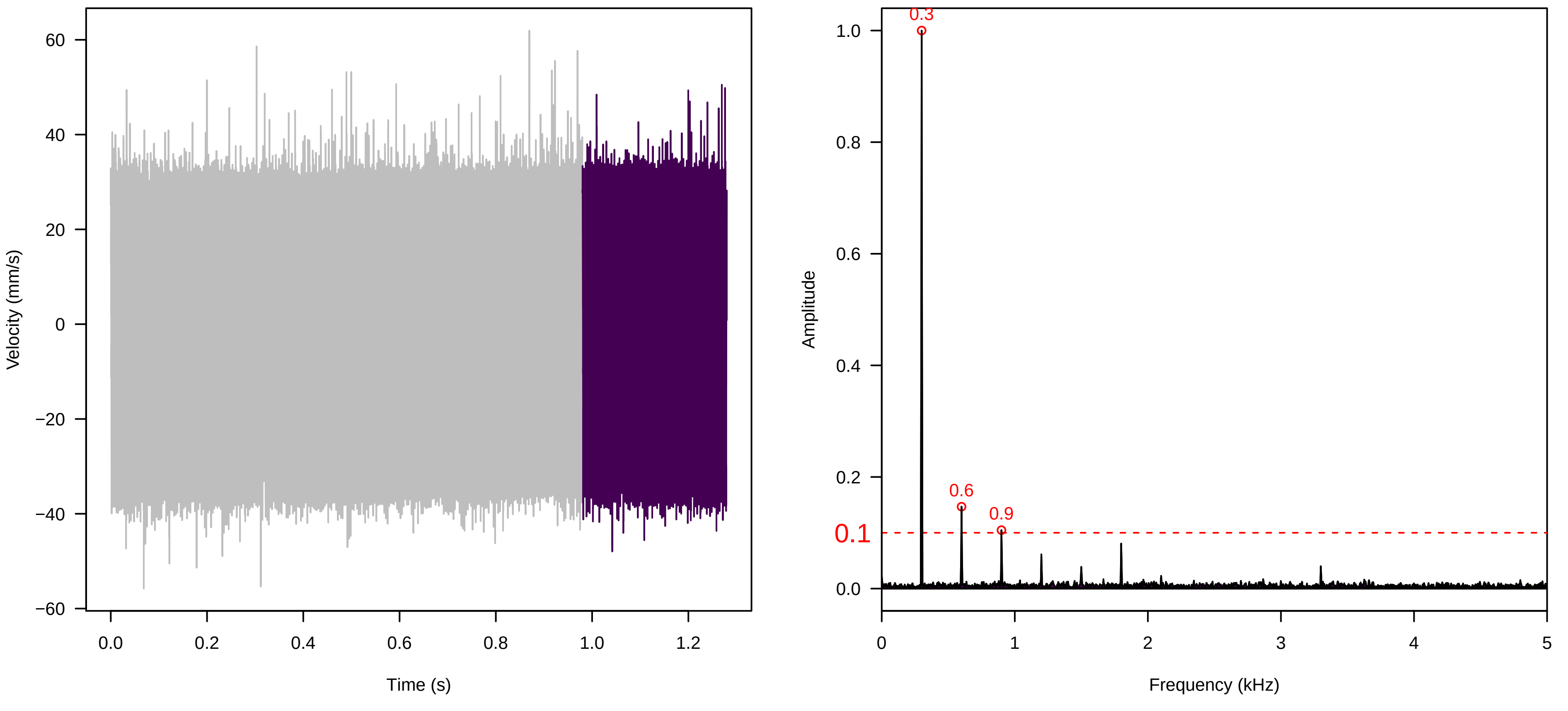

**Vel. = 0.014 ; Str. = Receptacle ; Axis = x ; Fl. accession = 10−s−81−8**

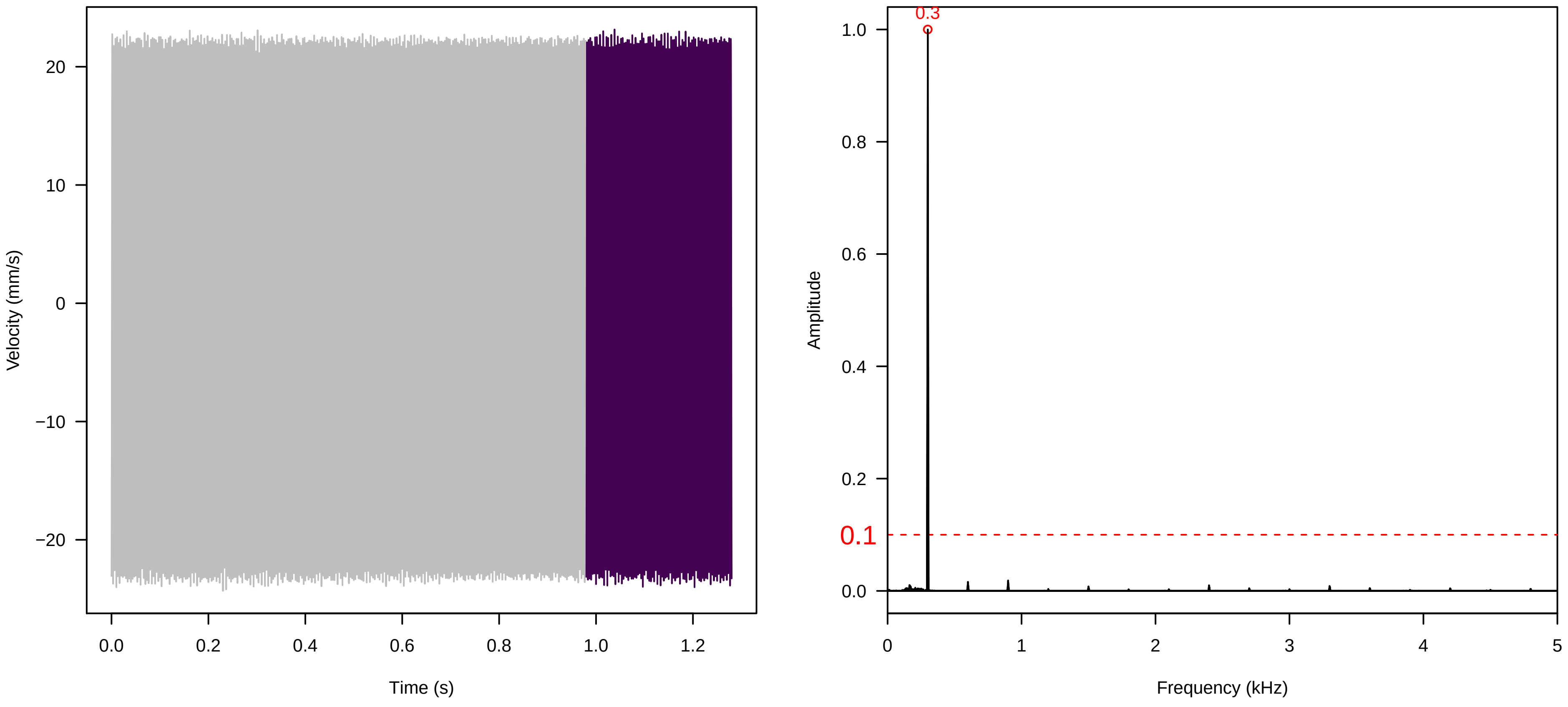


**Vel. = 0.014 ; Str. = PA ; Axis = x ; Fl. accession = 10−s−81−8**

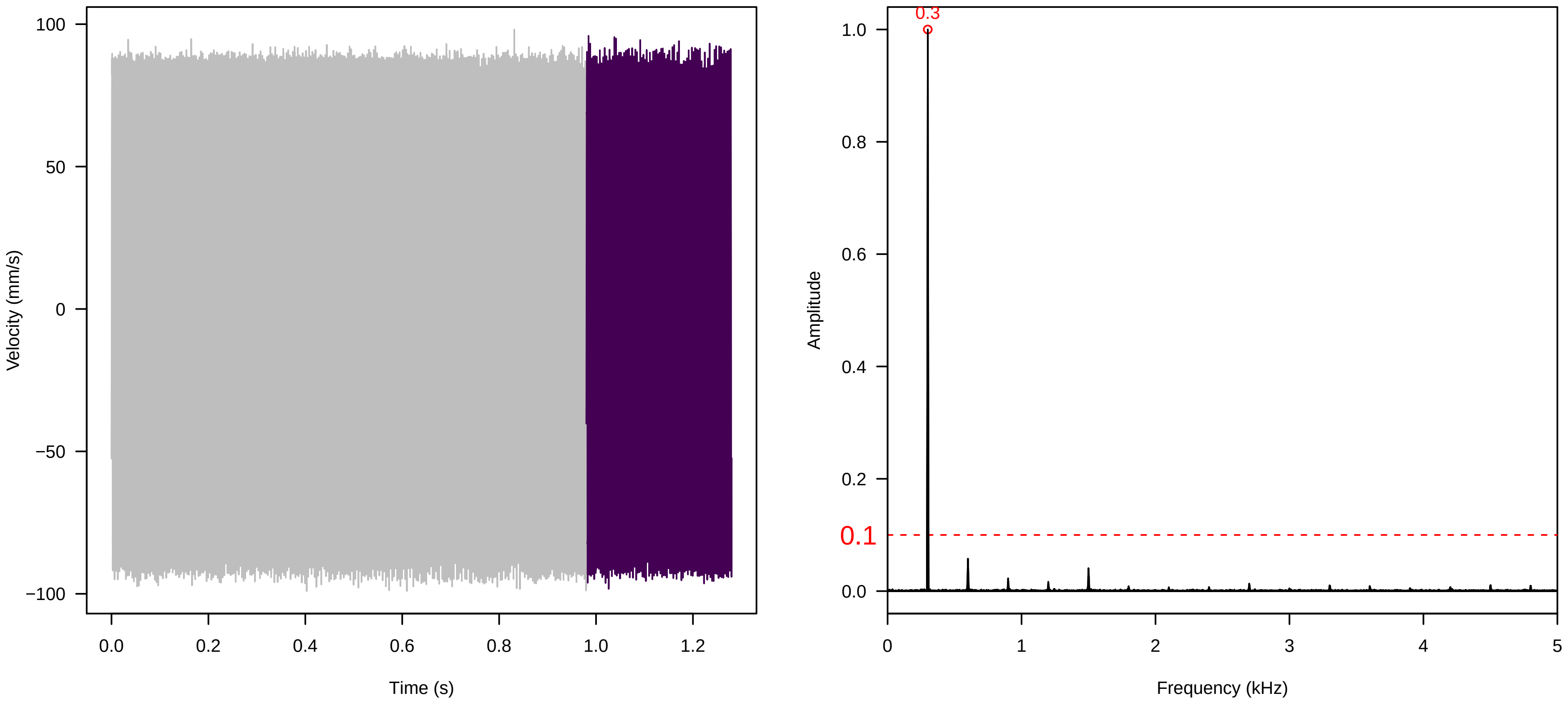


**Vel. = 0.014 ; Str. = Receptacle ; Axis = x ; Fl. accession = 10−s−81−8**

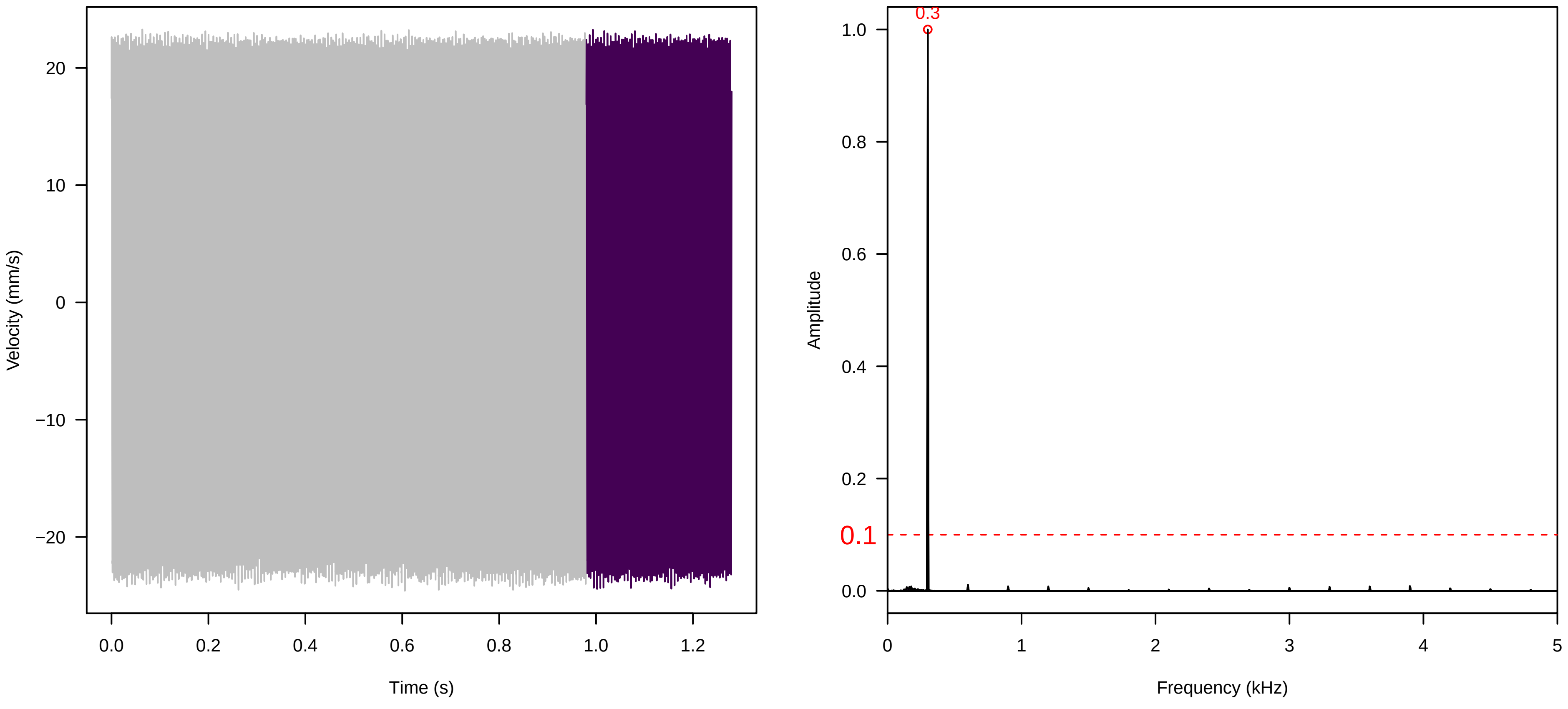

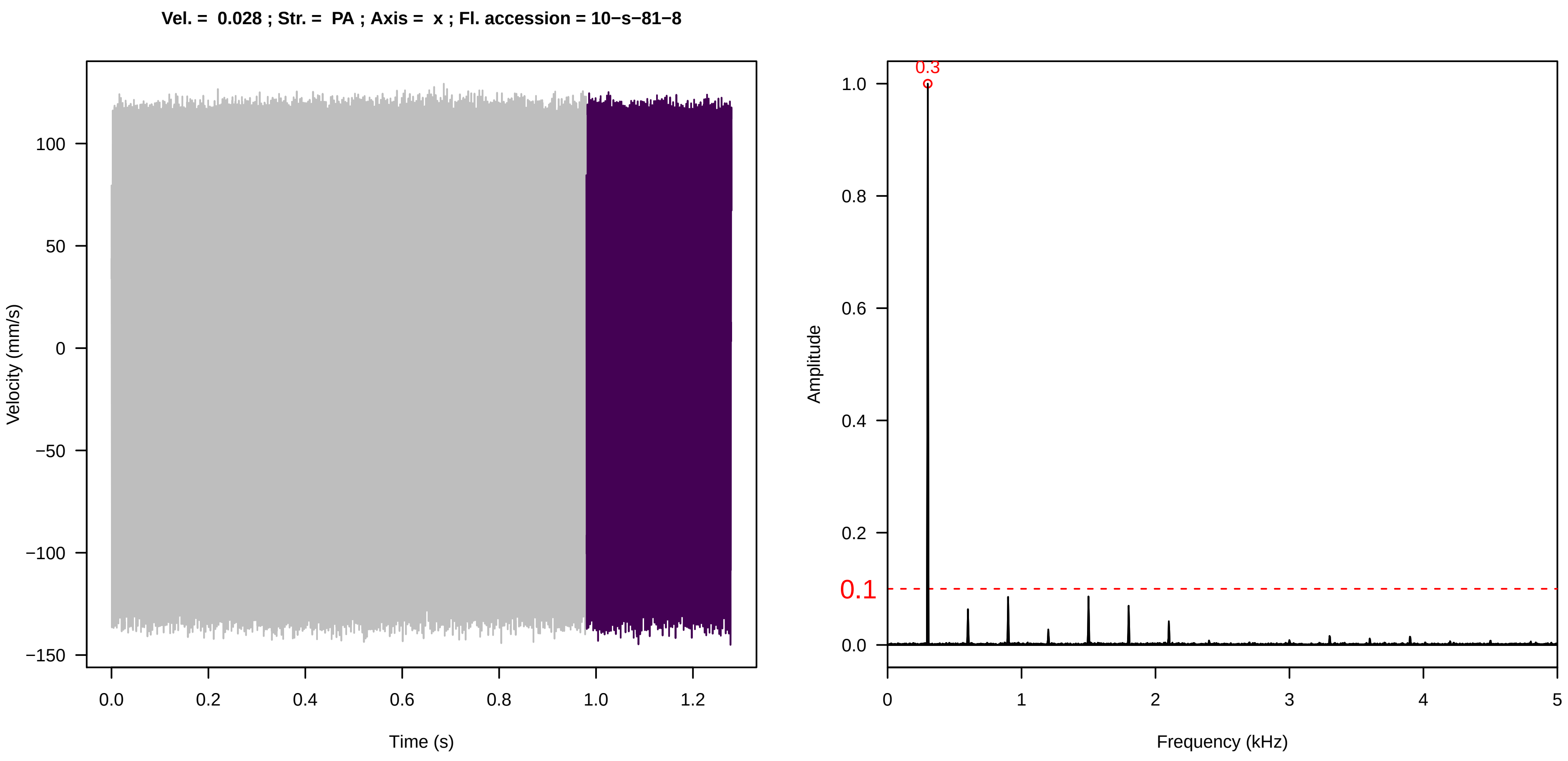
Vel. = 0.028 ; Str. = PA ; Axis = x ; Fl. accession = 10−s−81−8
Velocity (mm/s)
Time (s)
Amplitude
Frequency (kHz)
0.3
0.1

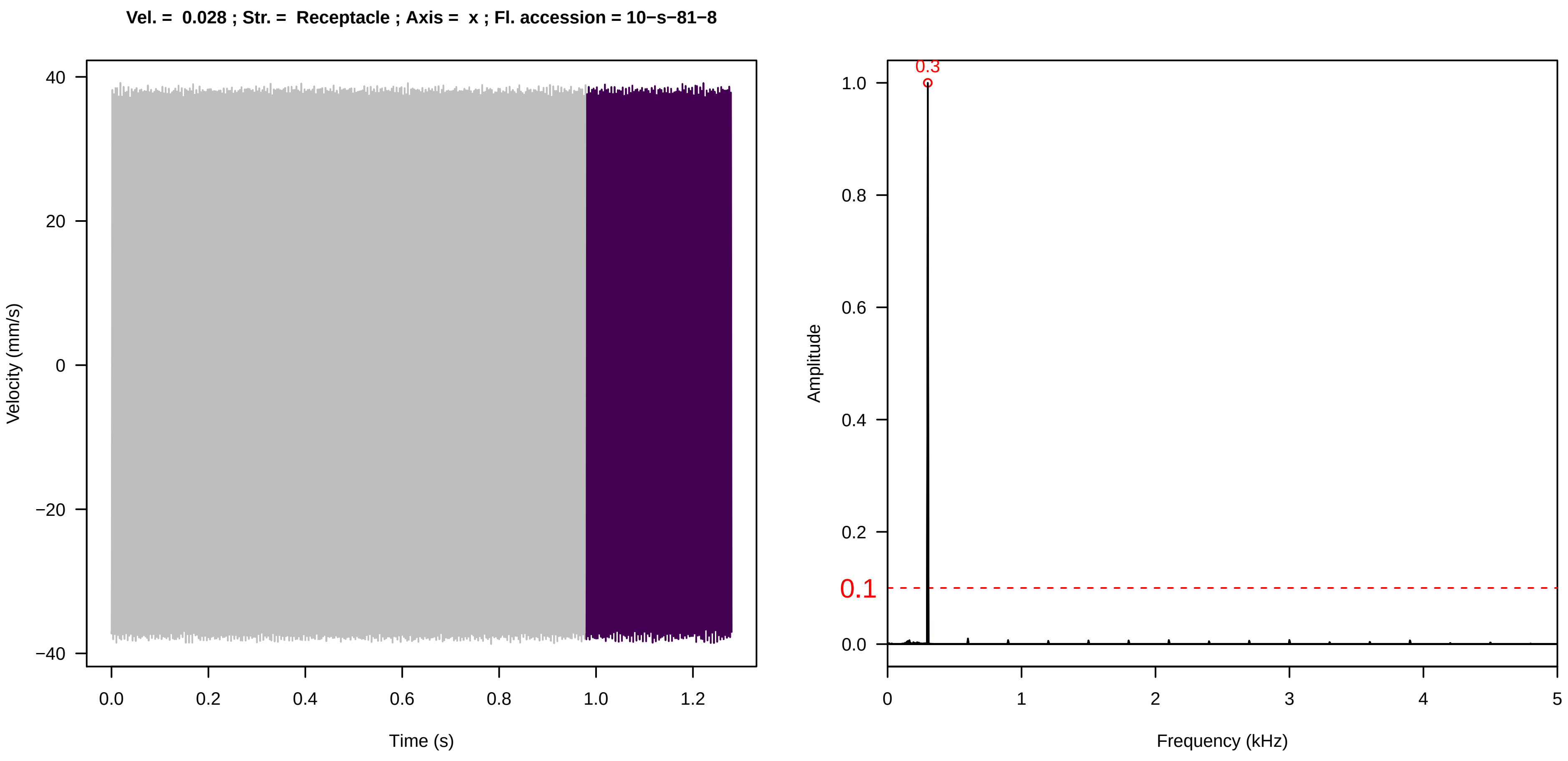
Vel. = 0.028 ; Str. = Receptacle ; Axis = x ; Fl. accession = 10−s−81−8
Velocity (mm/s)
Time (s)
Amplitude
Frequency (kHz)
0.3
0.1

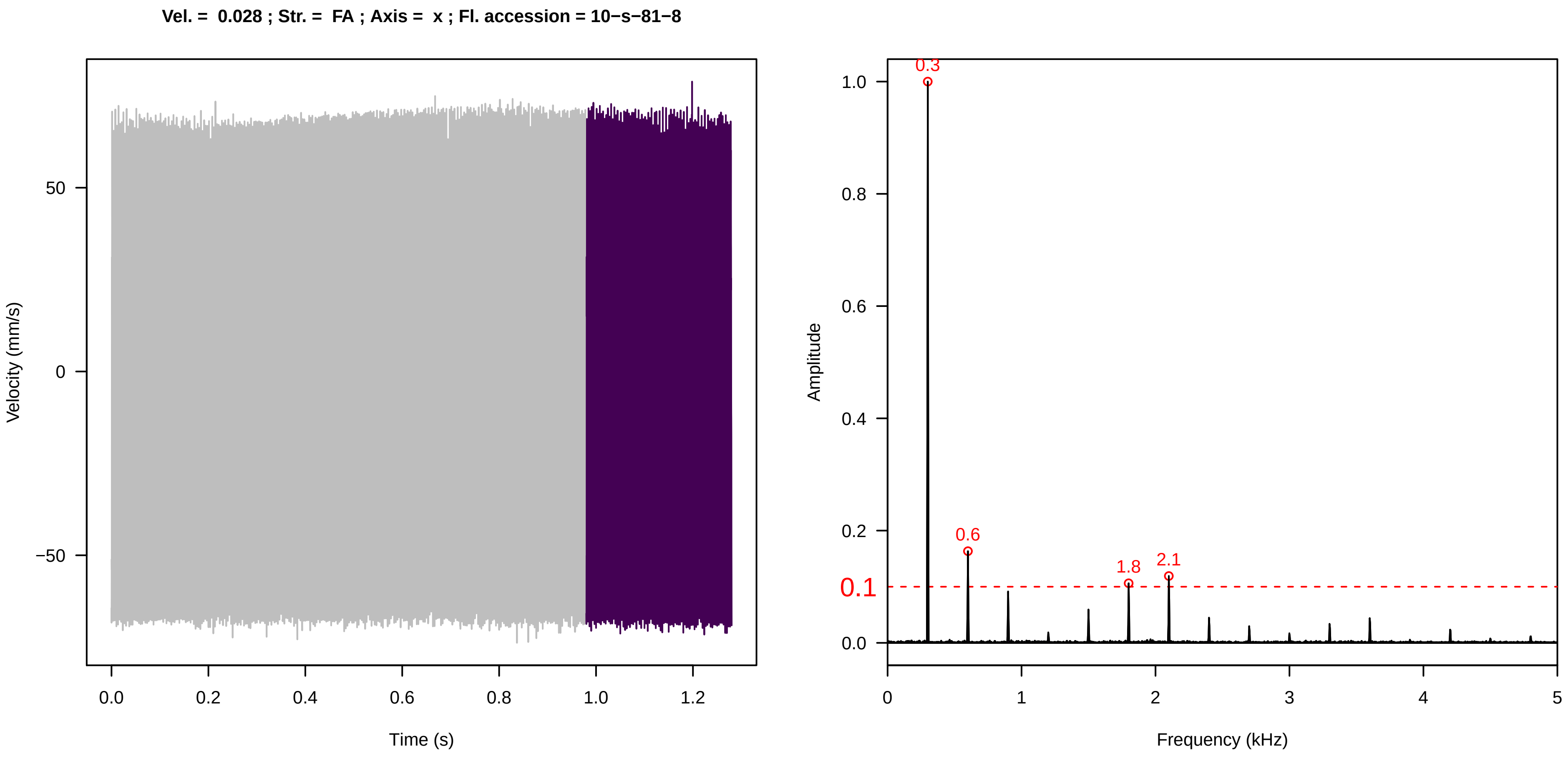
Vel. = 0.028 ; Str. = FA ; Axis = x ; Fl. accession = 10−s−81−8
Velocity (mm/s)
Time (s)
Amplitude
Frequency (kHz)
0.3
0.6
1.8
2.1
0.1

**Vel. = 0.028 ; Str. = Receptacle ; Axis = x ; Fl. accession = 10−s−81−8**

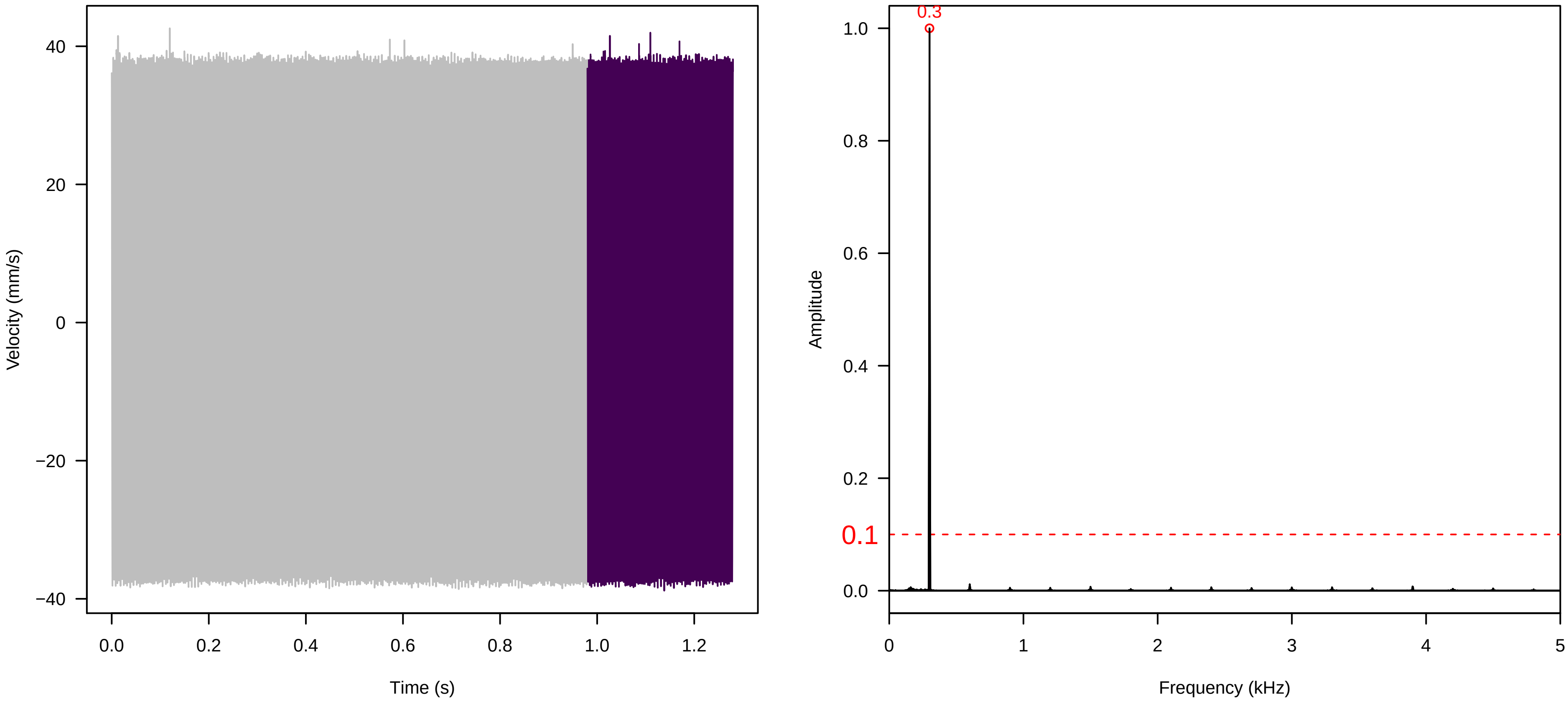


**Vel. = 0.028 ; Str. = Corolla ; Axis = x ; Fl. accession = 10−s−81−8**

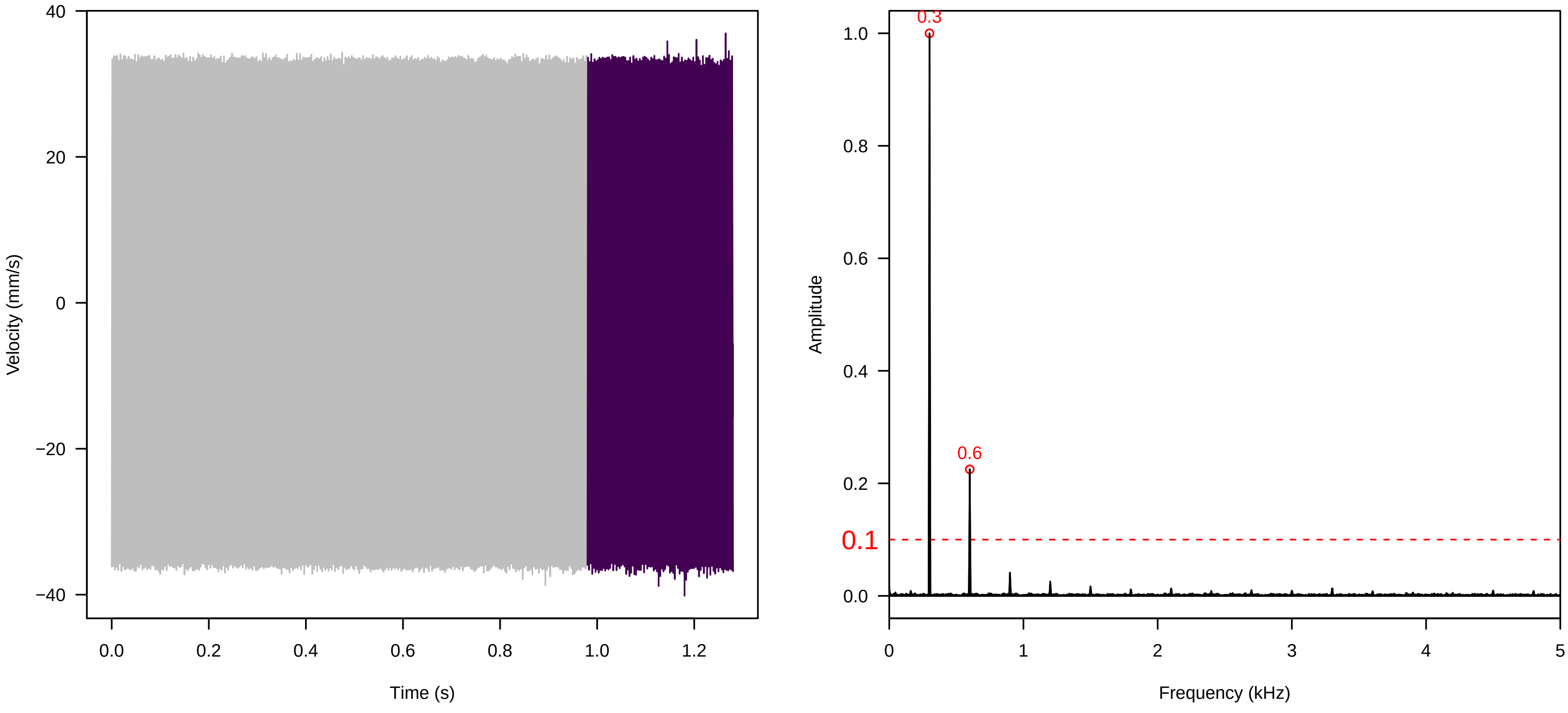


**Vel. = 0.028 ; Str. = Receptacle ; Axis = x ; Fl. accession = 10−s−81−8**

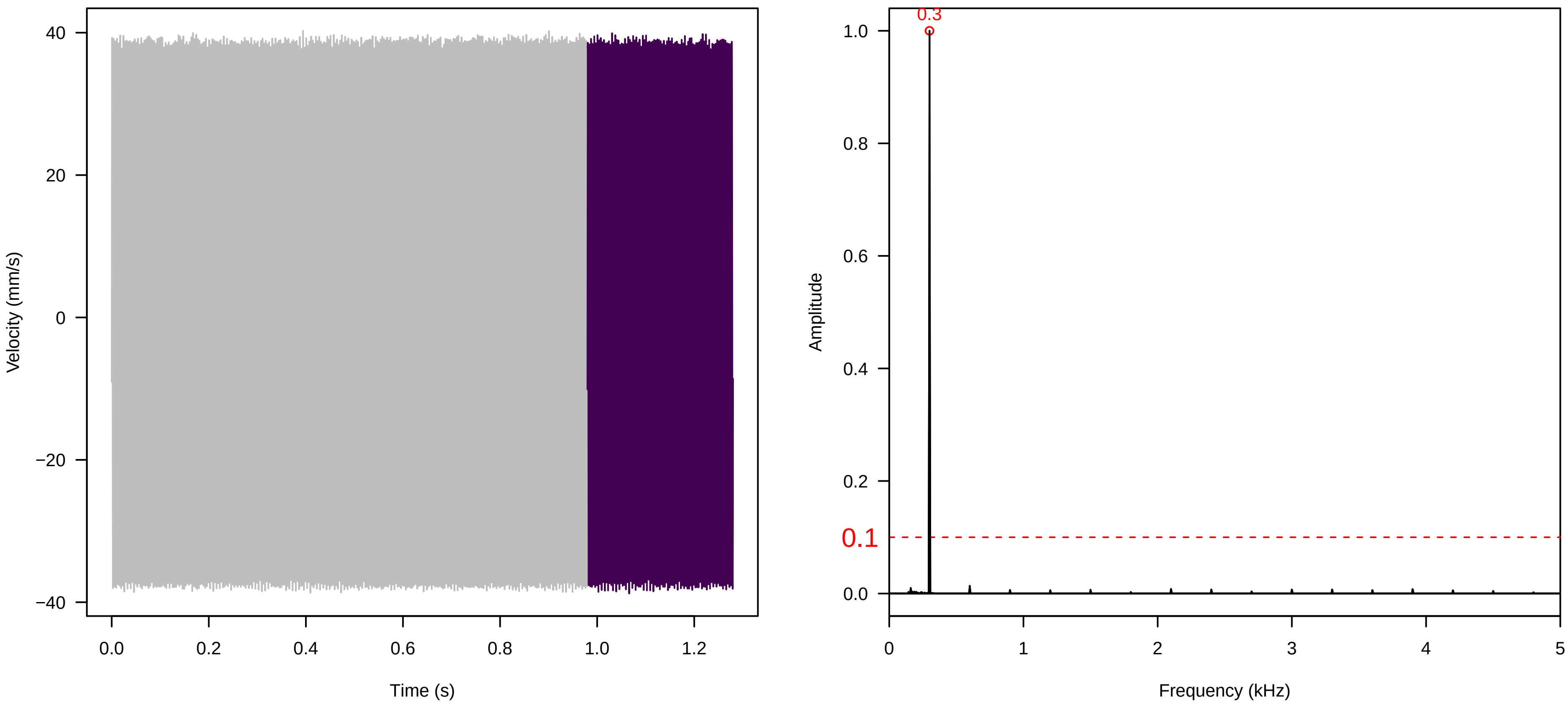

**Vel. =  0.057 ; Str. =  Corolla ; Axis =  x ; Fl. accession = 10−s−81−8**

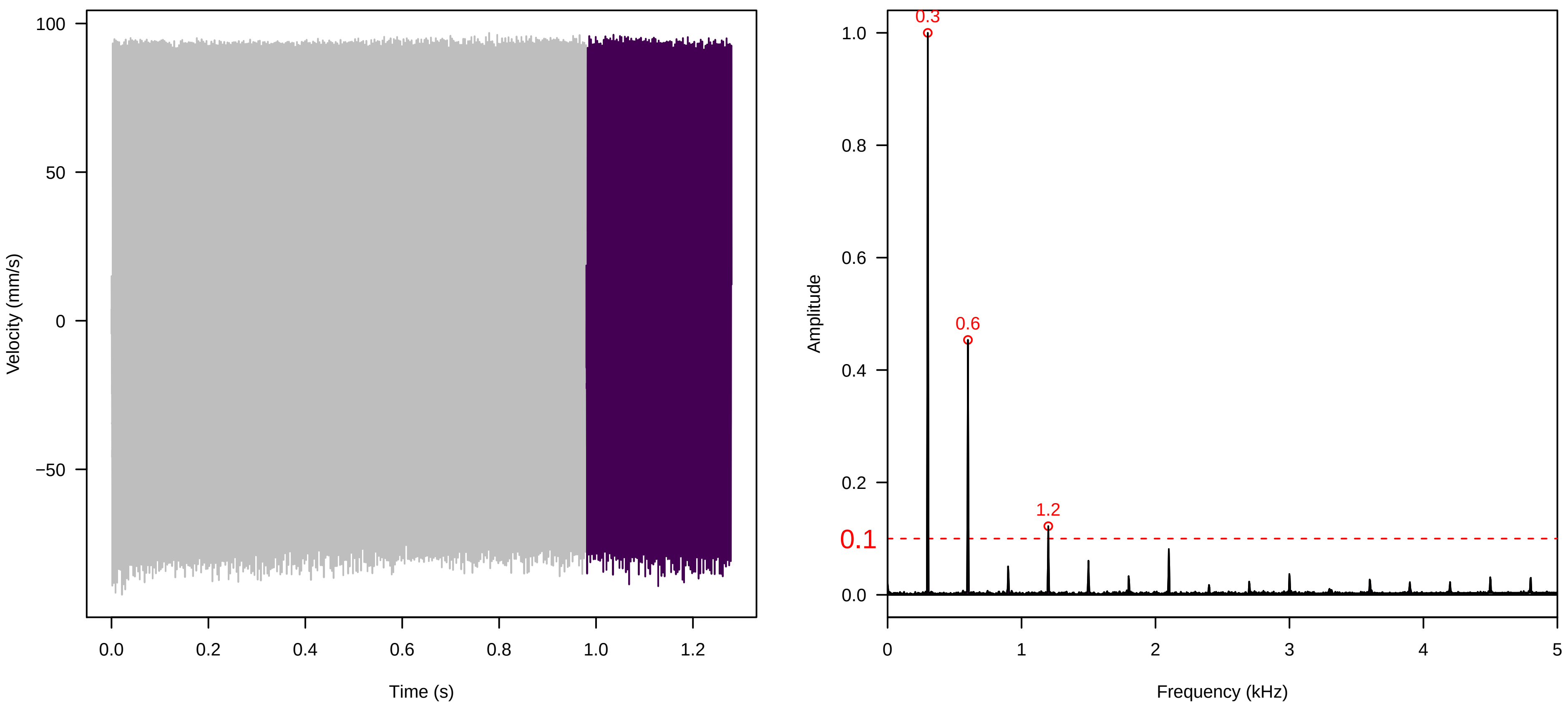


**Vel. =  0.057 ; Str. =  Receptacle ; Axis =  x ; Fl. accession = 10−s−81−8**

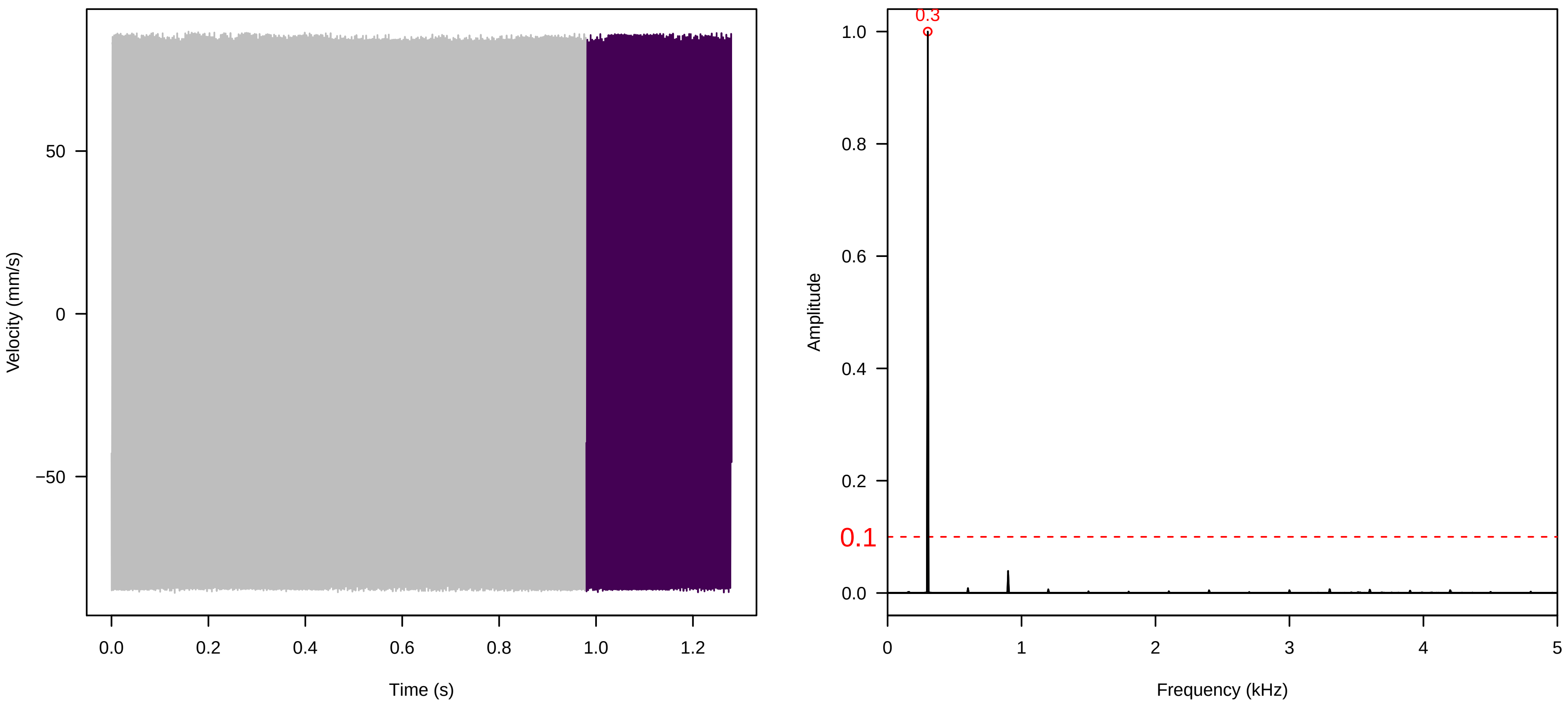


**Vel. =  0.057 ; Str. =  FA ; Axis =  x ; Fl. accession = 10−s−81−8**

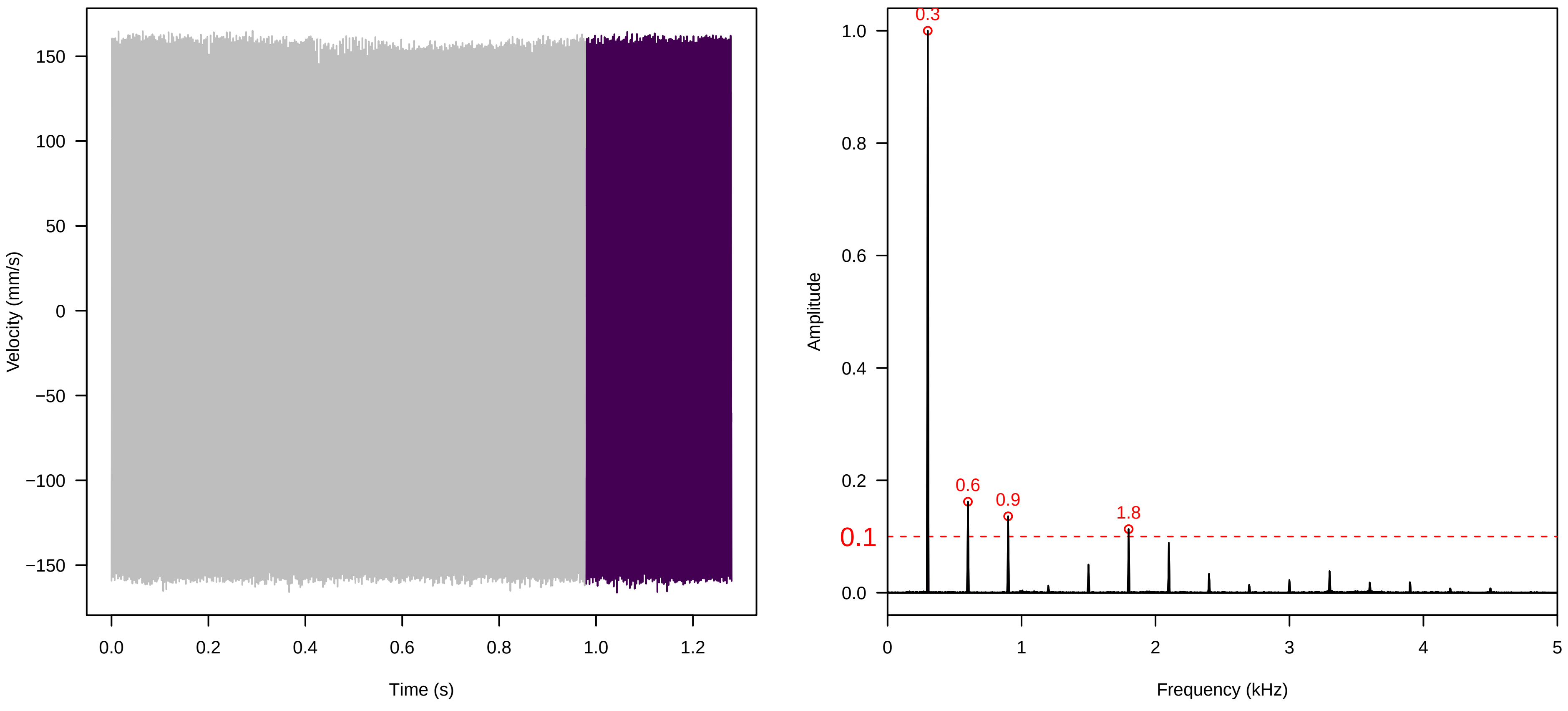

Vel. = 0.057 ; Str. = Receptacle ; Axis = x ; Fl. accession = 10−s−81−8

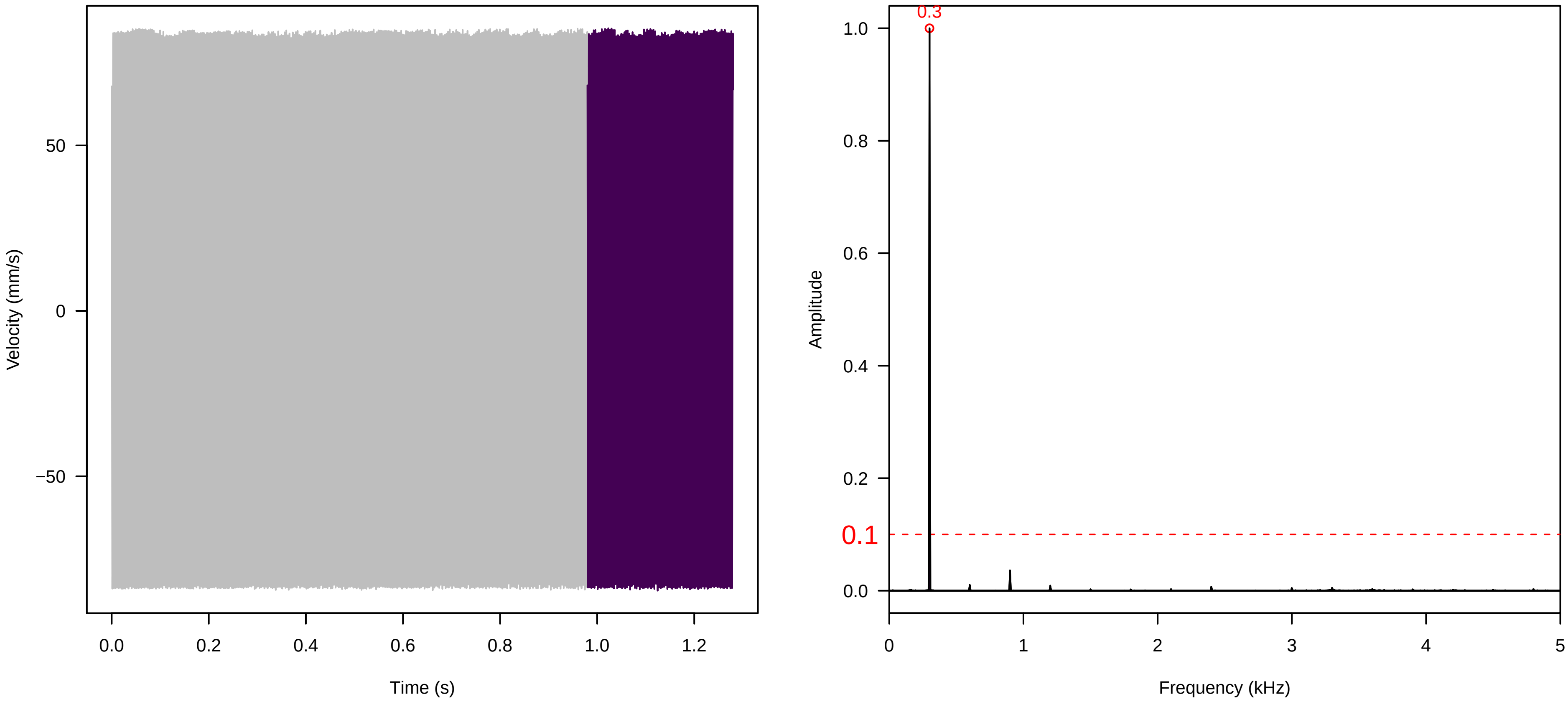


Vel. = 0.057 ; Str. = PA ; Axis = x ; Fl. accession = 10−s−81−8

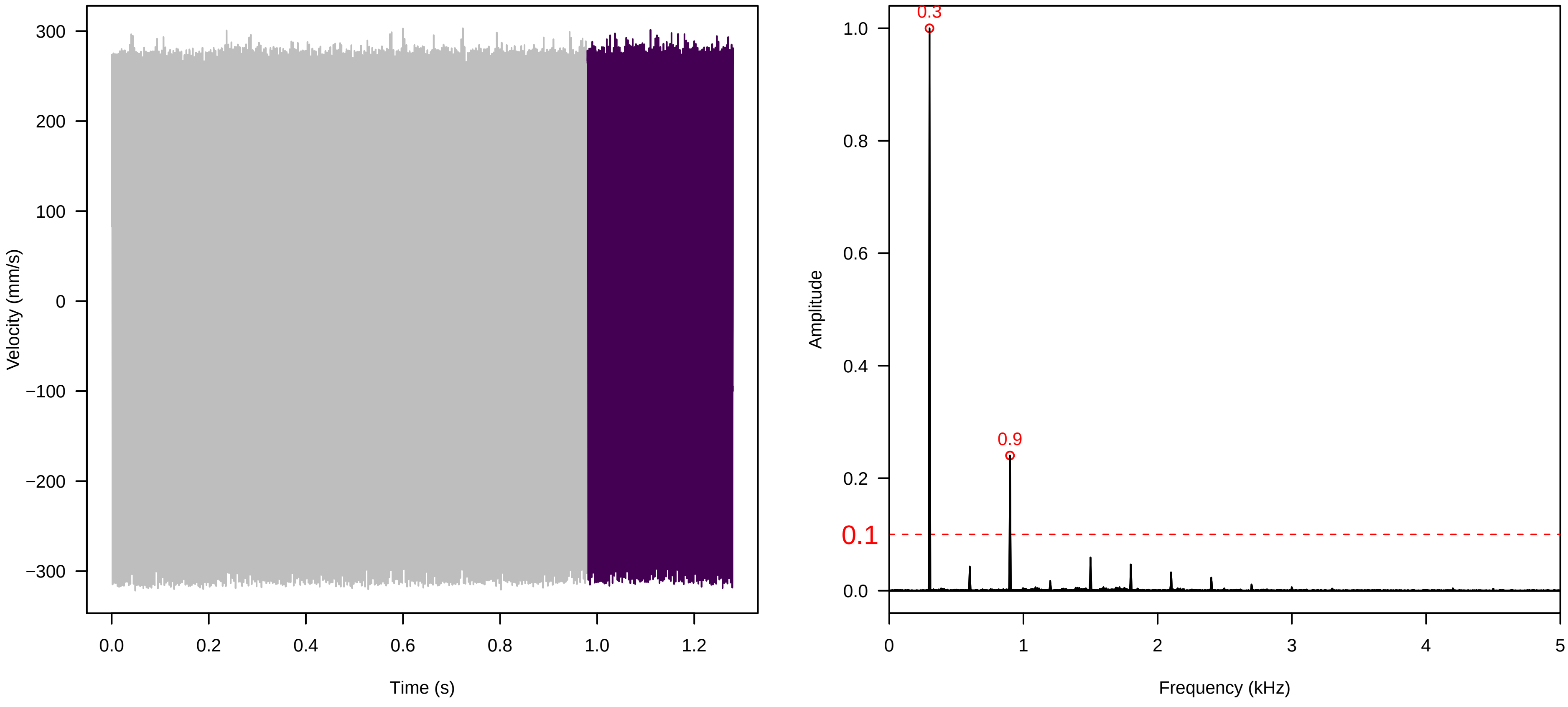


Vel. = 0.057 ; Str. = Receptacle ; Axis = x ; Fl. accession = 10−s−81−8

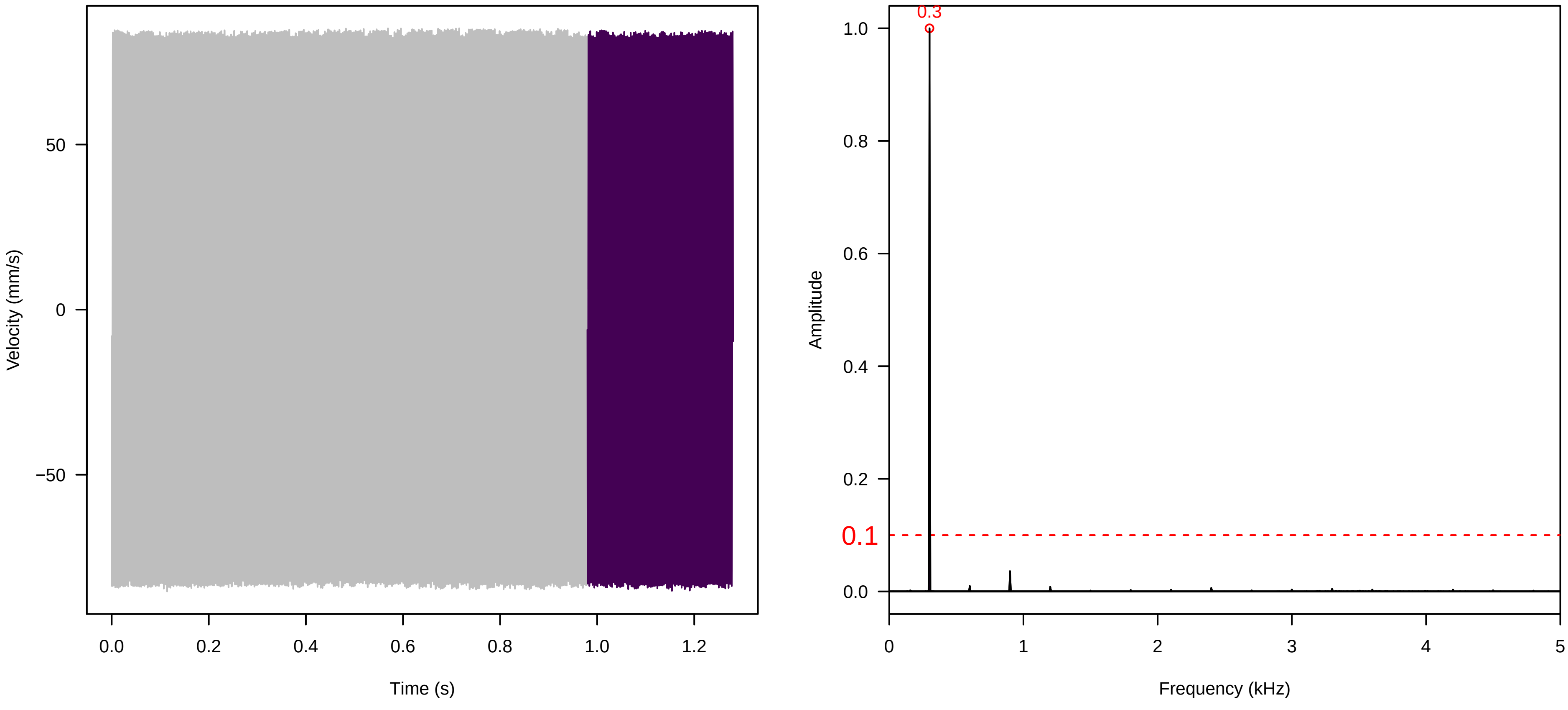

**Vel. =  0.057 ; Str. =  Corolla ; Axis =  z ; Fl. accession = 10−s−81−8**

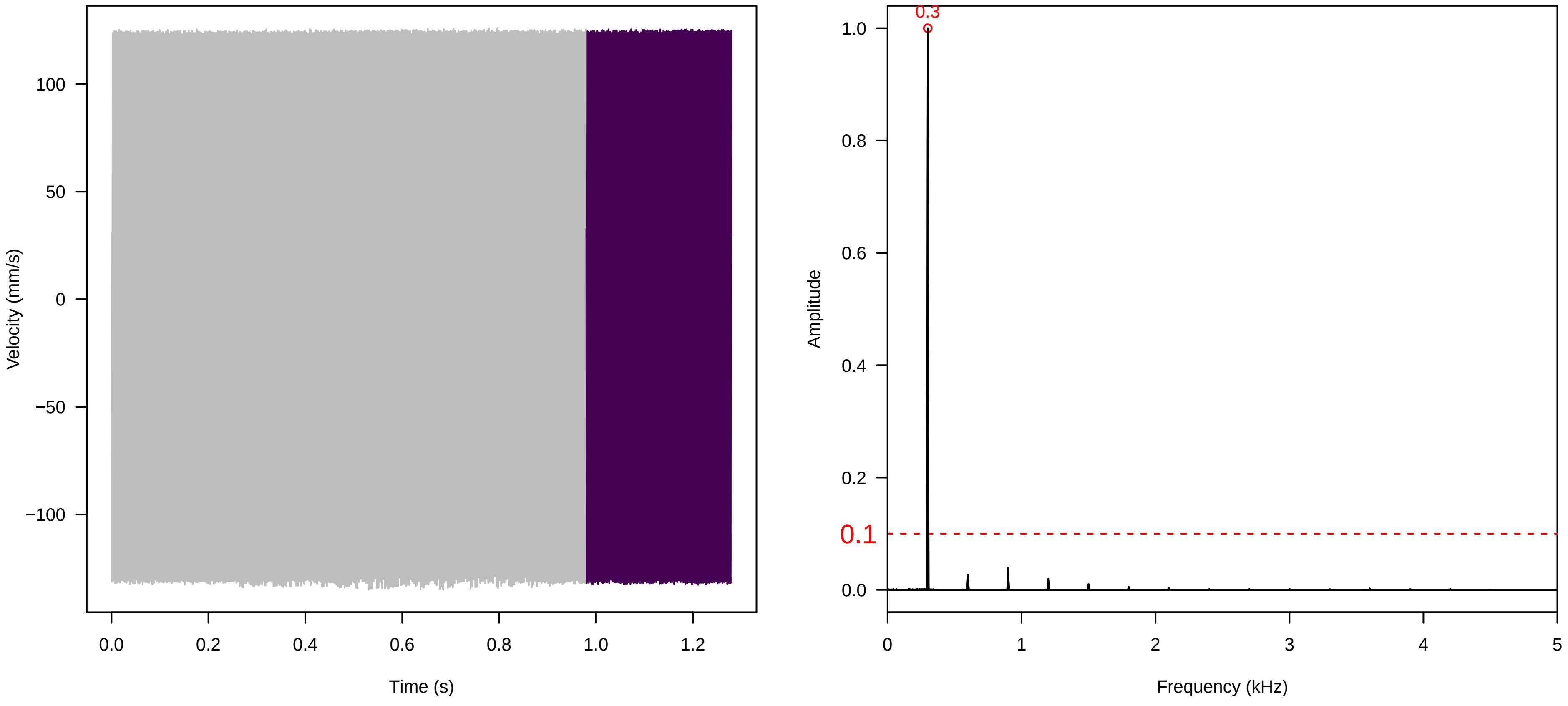


**Vel. =  0.057 ; Str. =  Receptacle ; Axis =  z ; Fl. accession = 10−s−81−8**

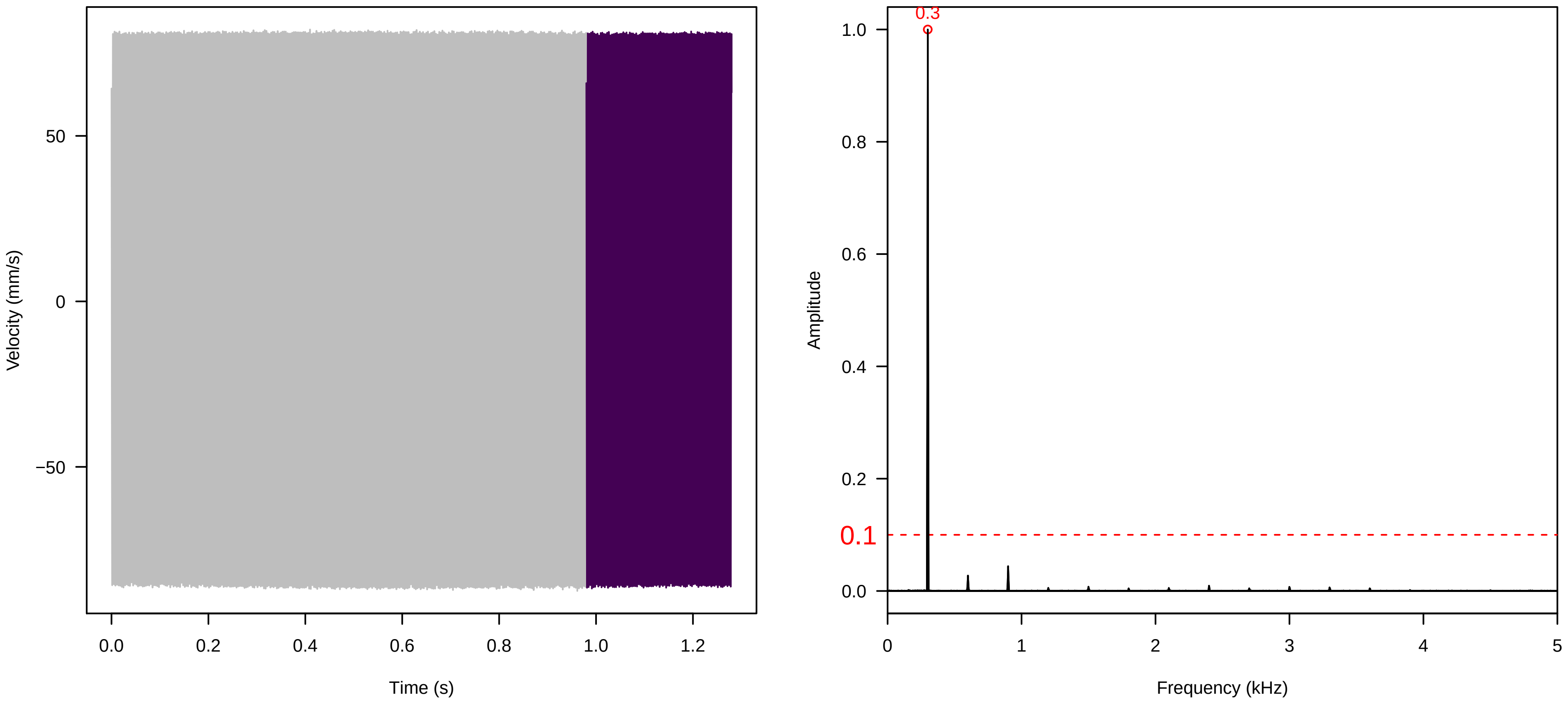


**Vel. =  0.057 ; Str. =  FA ; Axis =  z ; Fl. accession = 10−s−81−8**

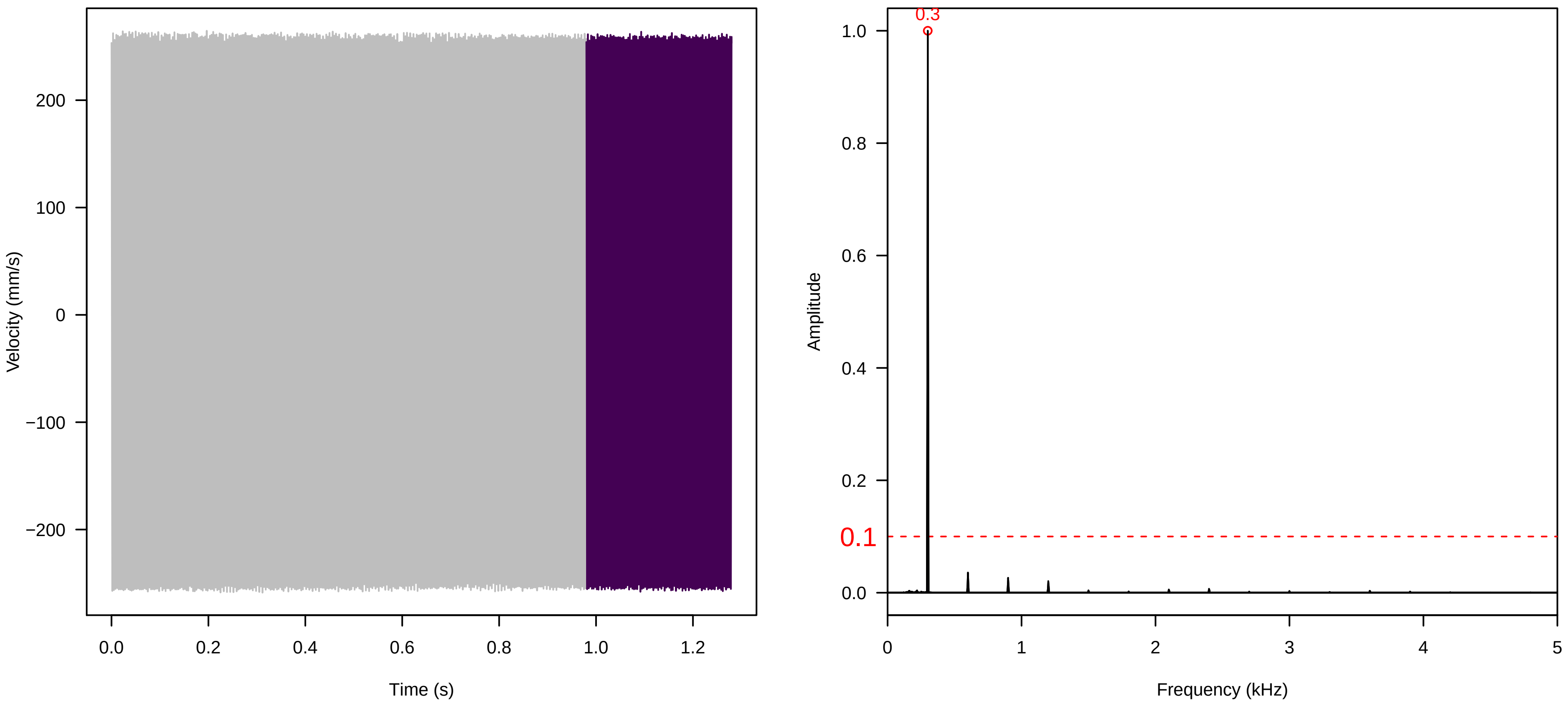

**Vel. = 0.057 ; Str. = Receptacle ; Axis = z ; Fl. accession = 10−s−81−8**

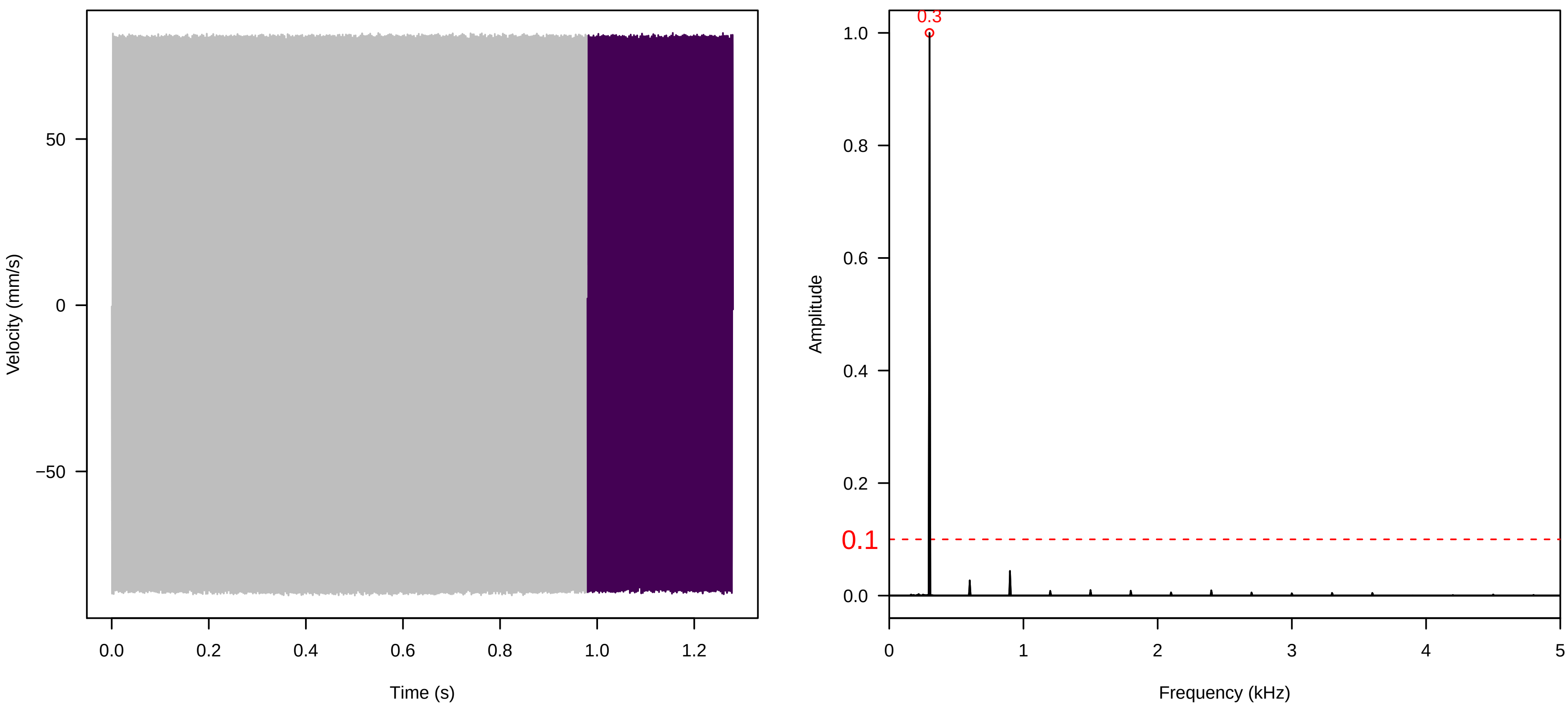


**Vel. = 0.057 ; Str. = PA ; Axis = z ; Fl. accession = 10−s−81−8**

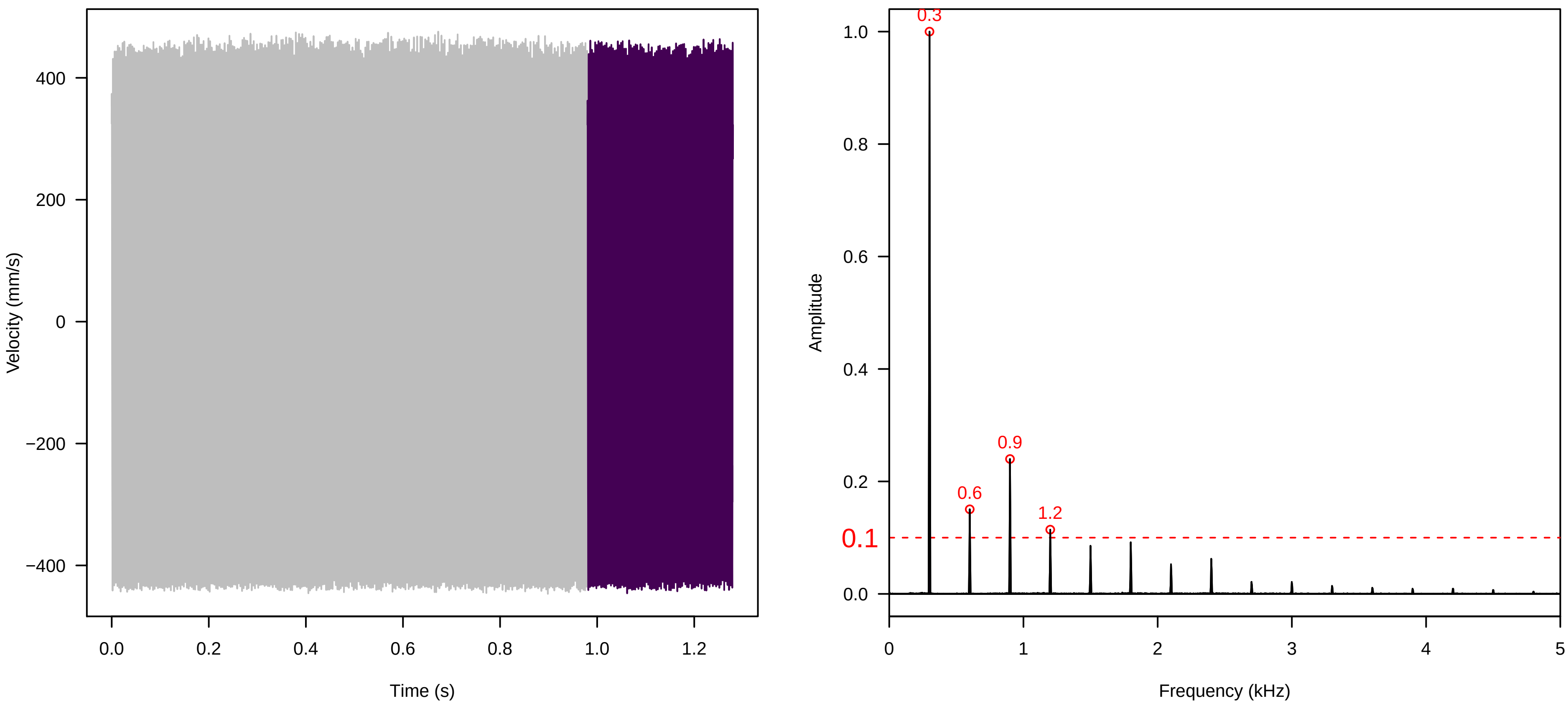


**Vel. = 0.057 ; Str. = Receptacle ; Axis = z ; Fl. accession = 10−s−81−8**

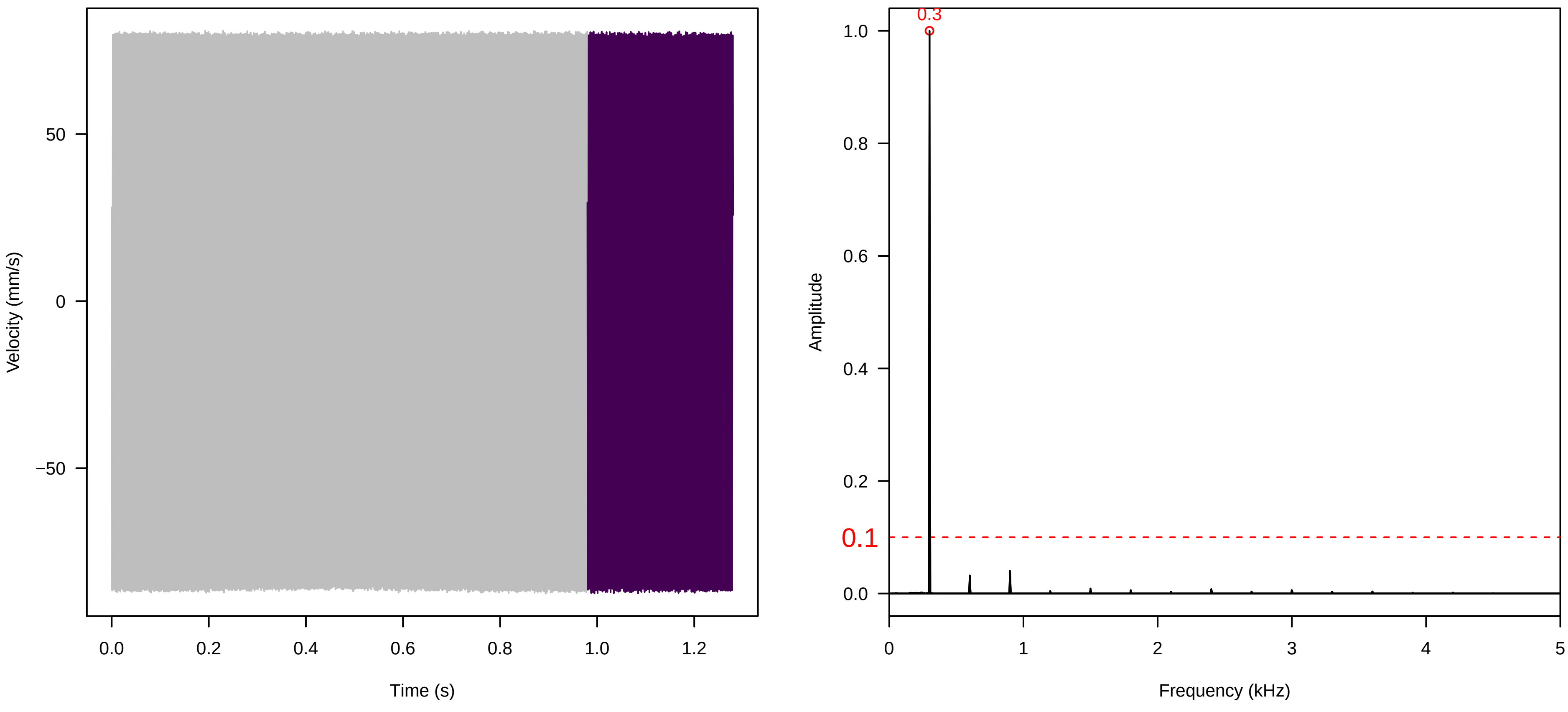

**Vel. = 0.028 ; Str. = PA ; Axis = z ; Fl. accession = 10−s−81−8**

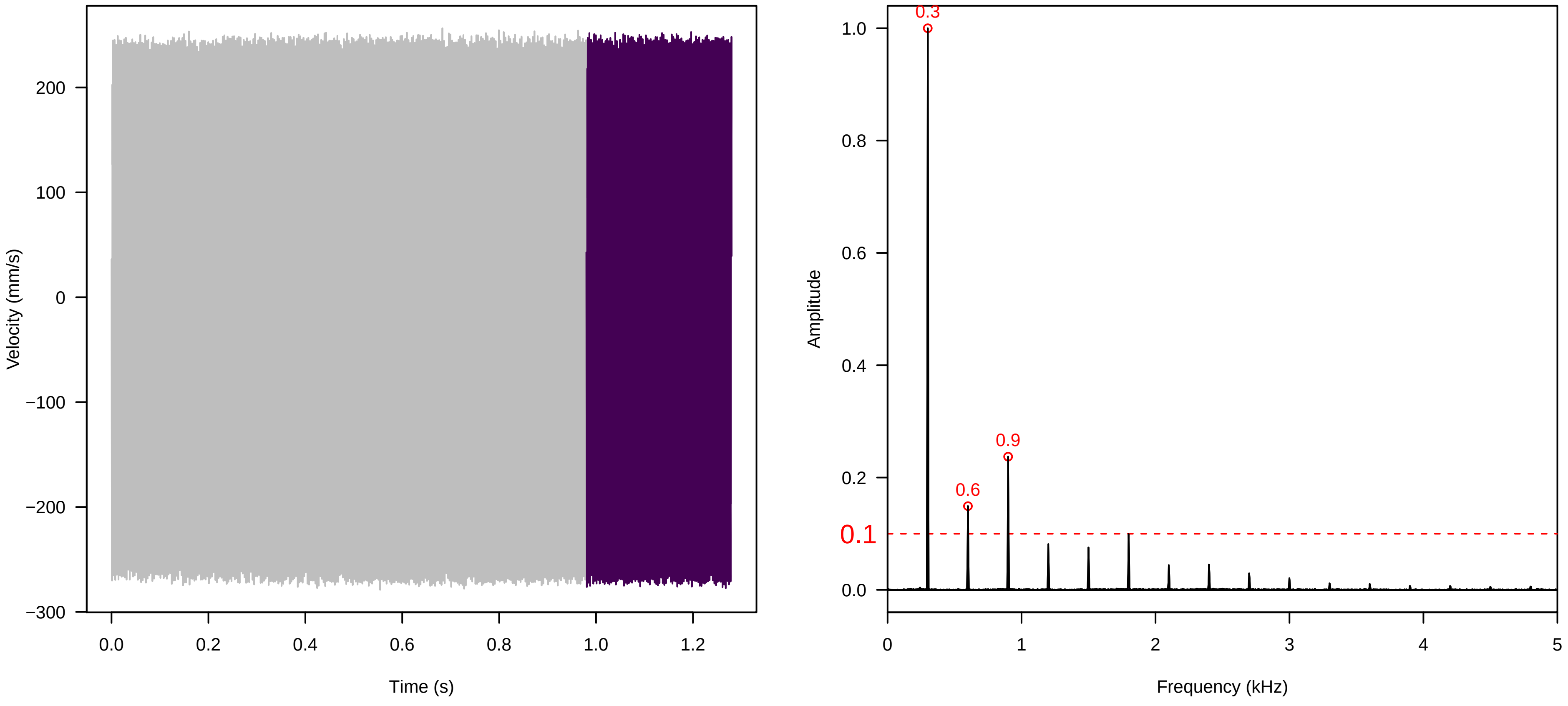


**Vel. = 0.028 ; Str. = Receptacle ; Axis = z ; Fl. accession = 10−s−81−8**

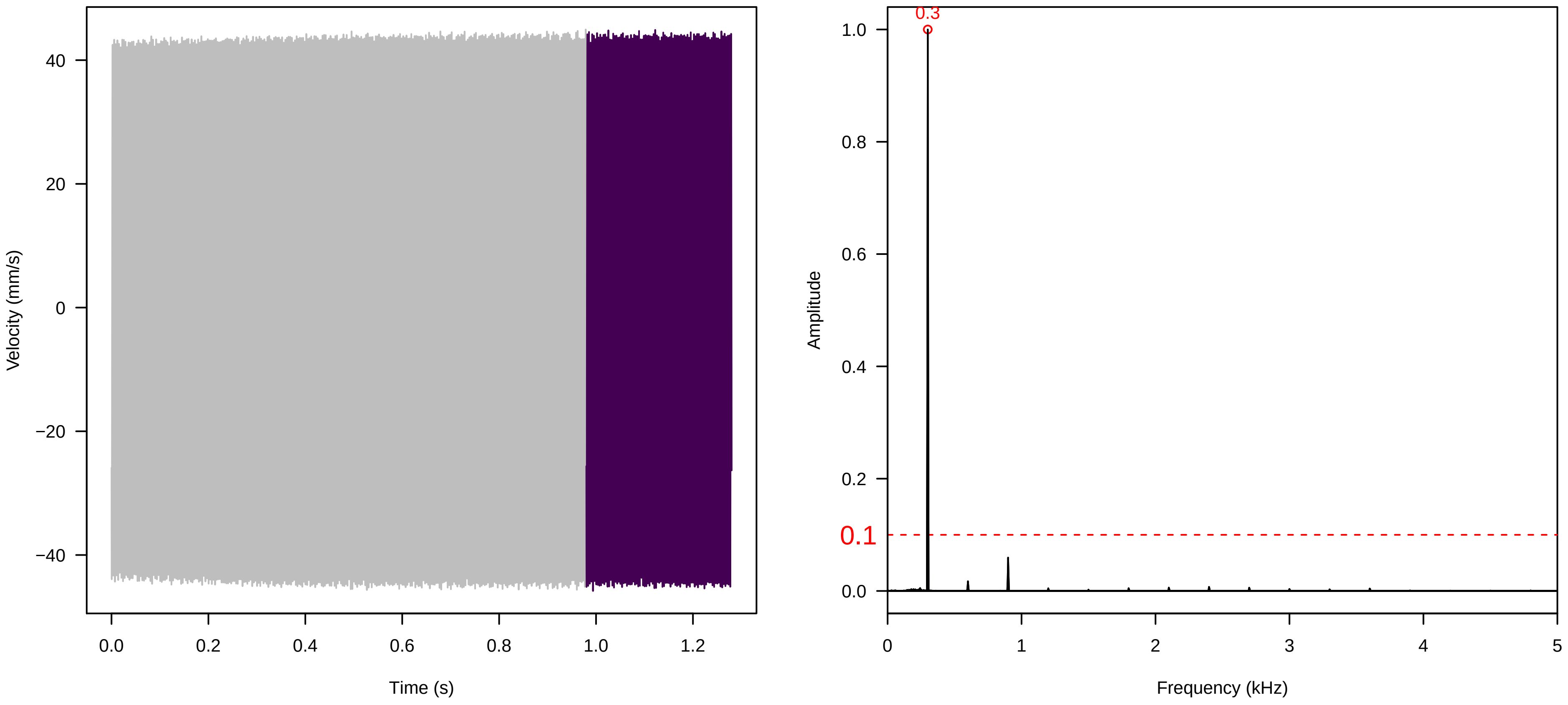


**Vel. = 0.028 ; Str. = FA ; Axis = z ; Fl. accession = 10−s−81−8**

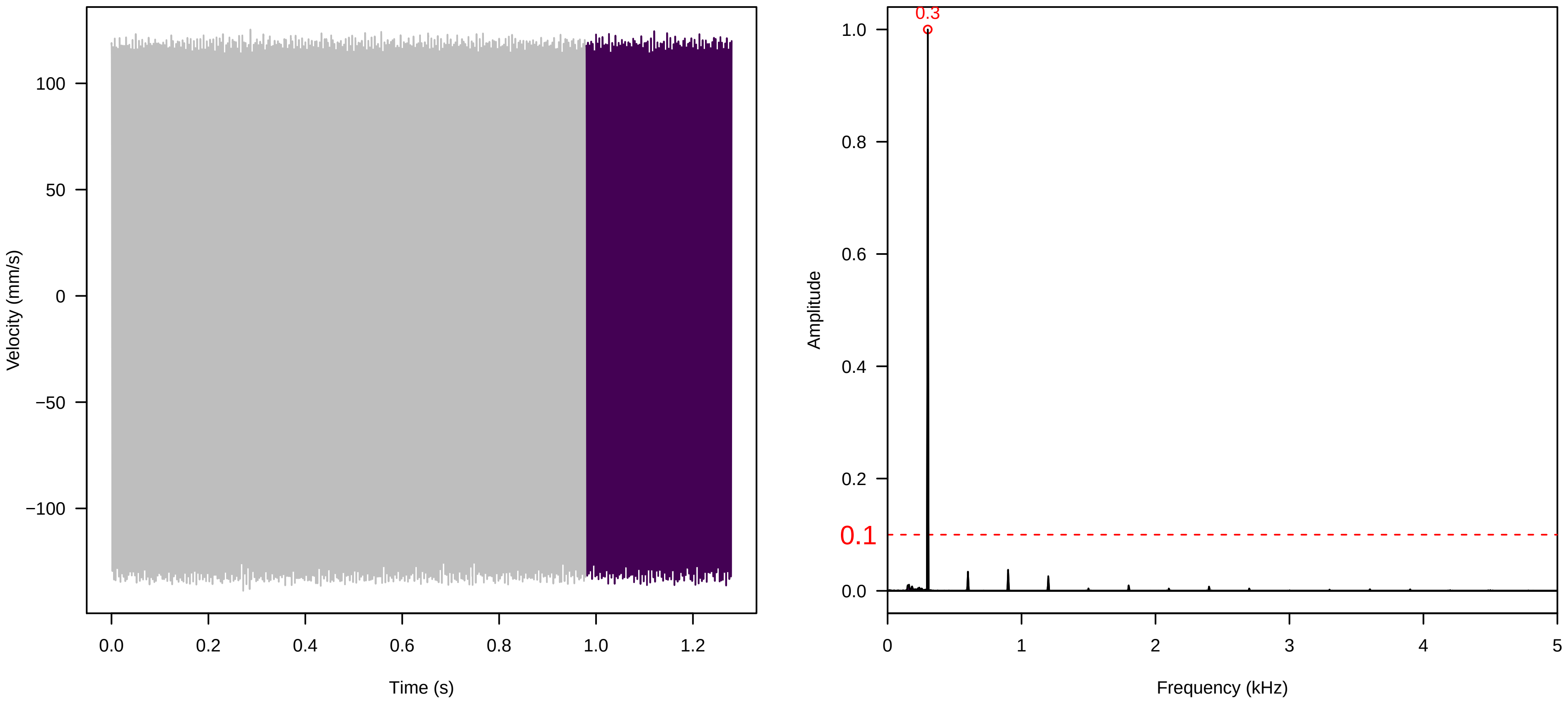

**Vel. = 0.028 ; Str. = Receptacle ; Axis = z ; Fl. accession = 10−s−81−8**

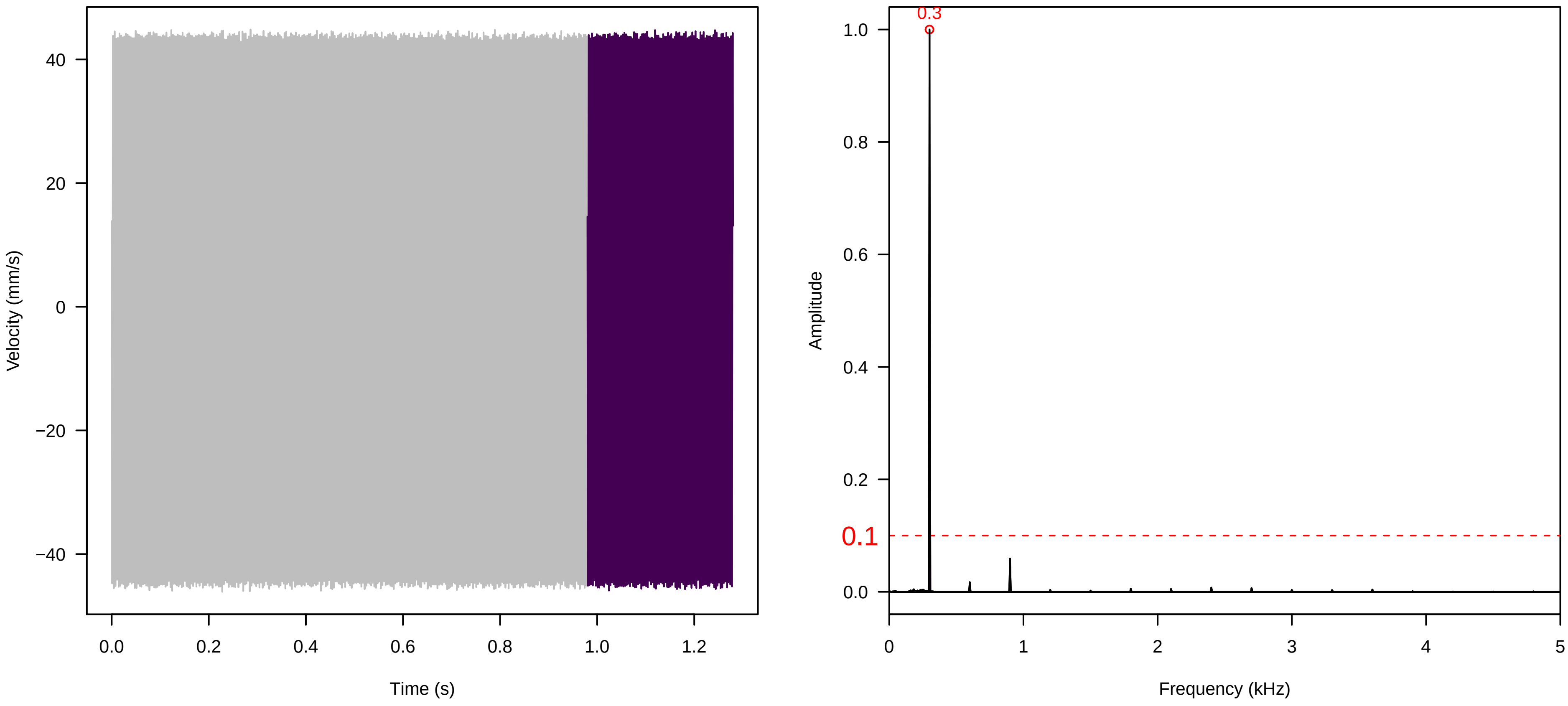


**Vel. = 0.028 ; Str. = Corolla ; Axis = z ; Fl. accession = 10−s−81−8**

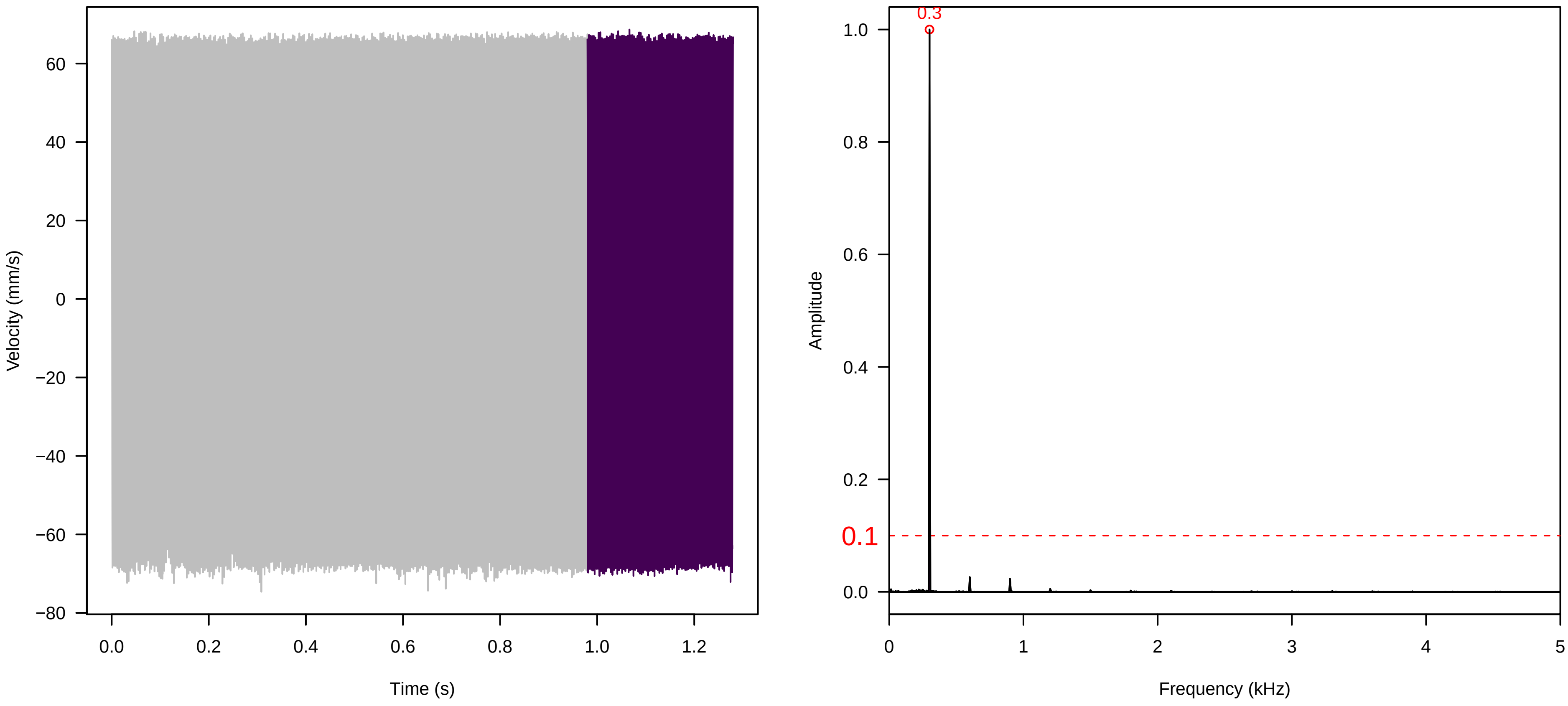


**Vel. = 0.028 ; Str. = Receptacle ; Axis = z ; Fl. accession = 10−s−81−8**

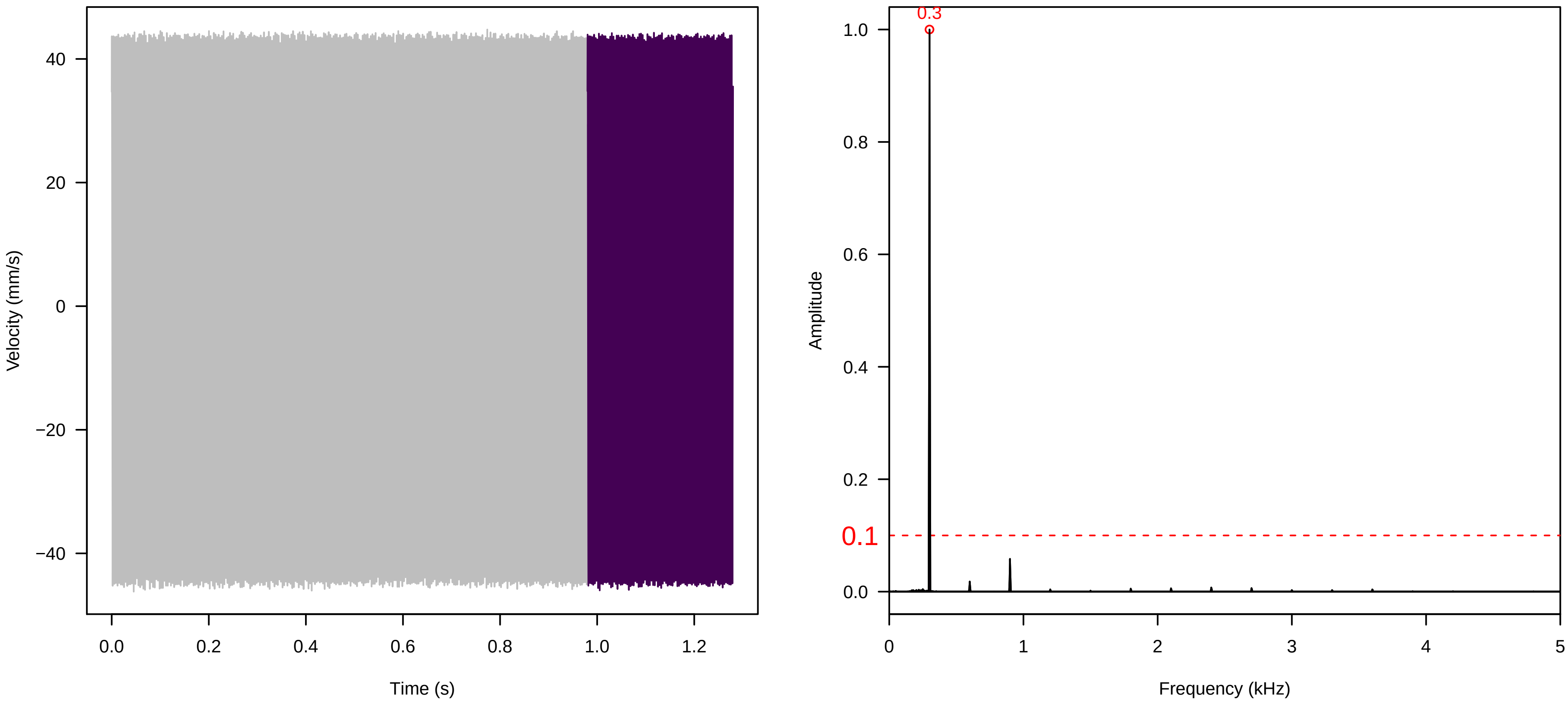

**Vel. = 0.014 ; Str. = Corolla ; Axis = z ; Fl. accession = 10−s−81−8**

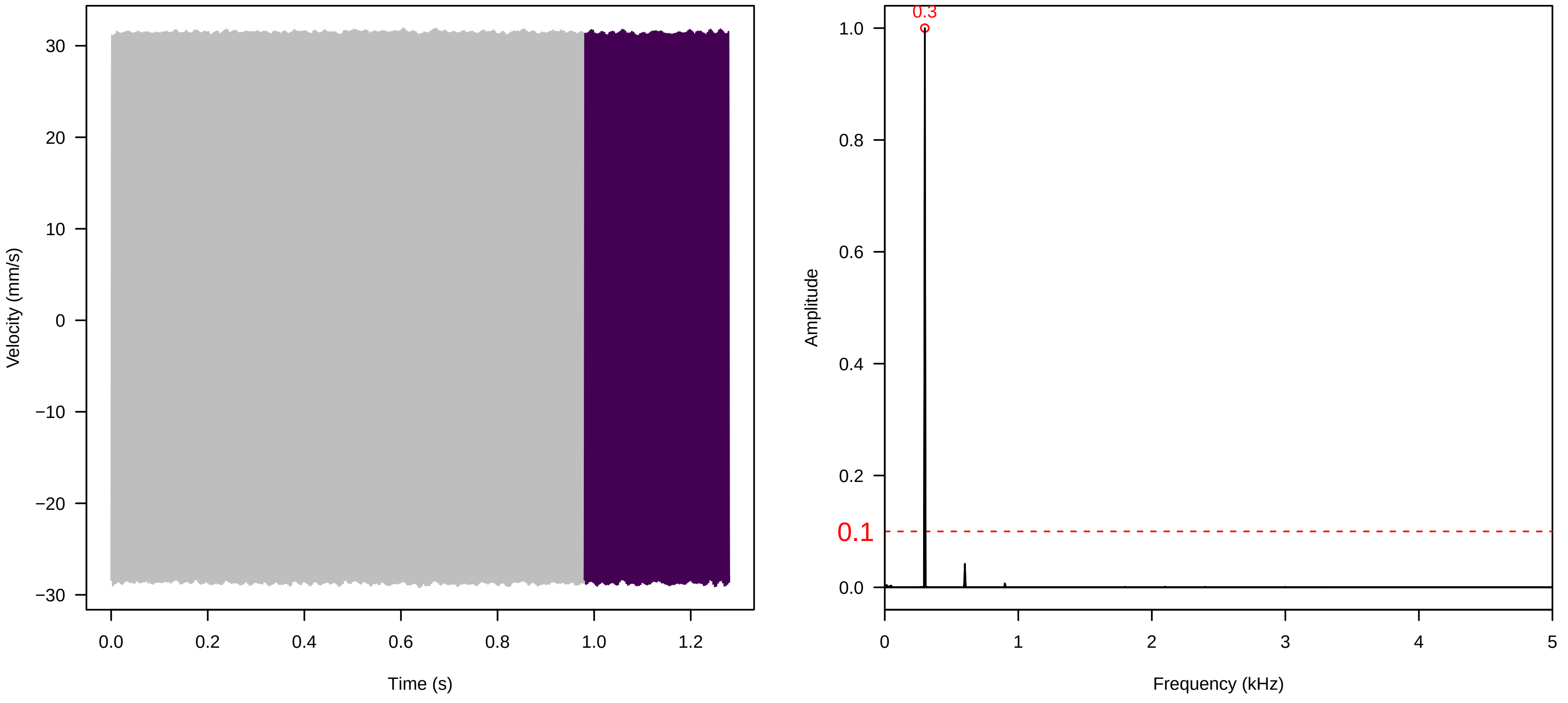


**Vel. = 0.014 ; Str. = Receptacle ; Axis = z ; Fl. accession = 10−s−81−8**

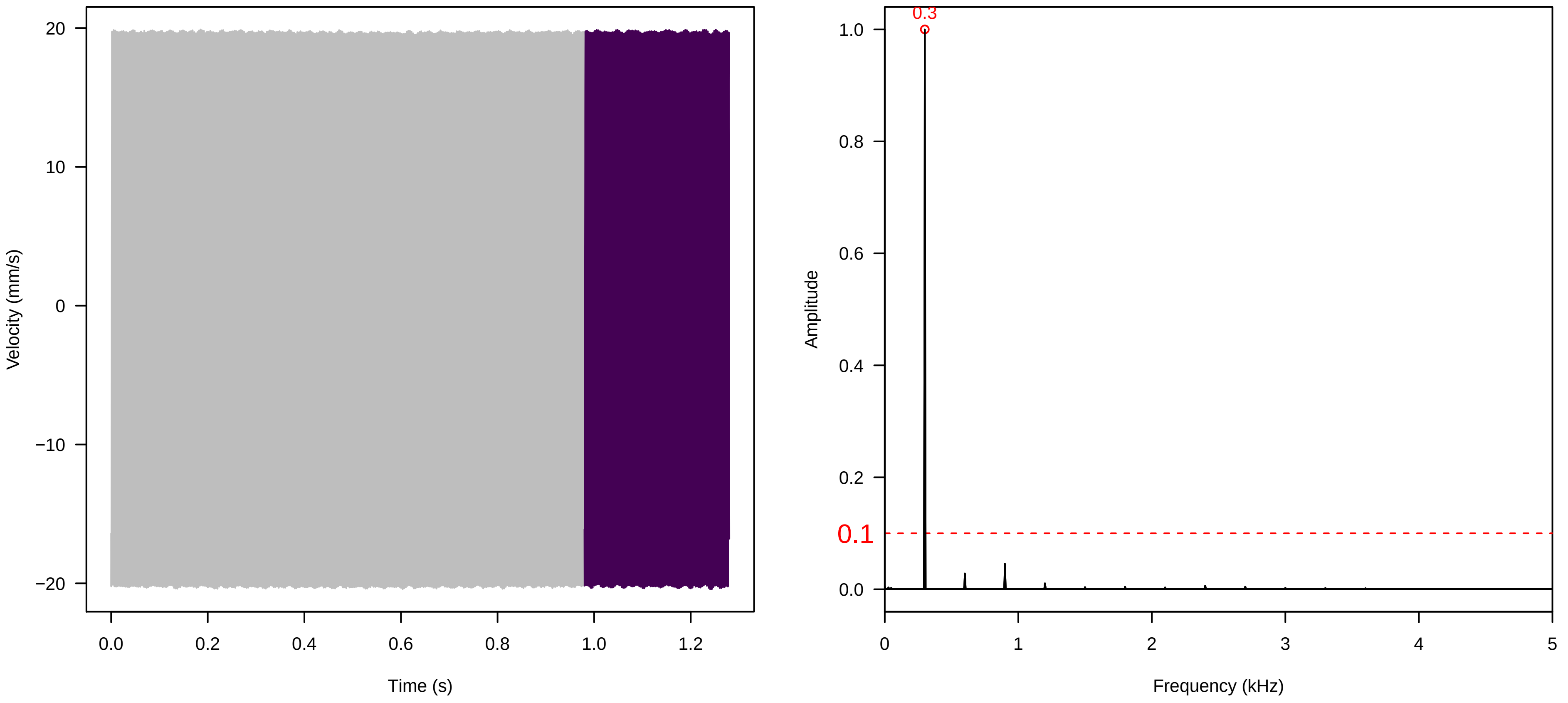


**Vel. = 0.014 ; Str. = FA ; Axis = z ; Fl. accession = 10−s−81−8**

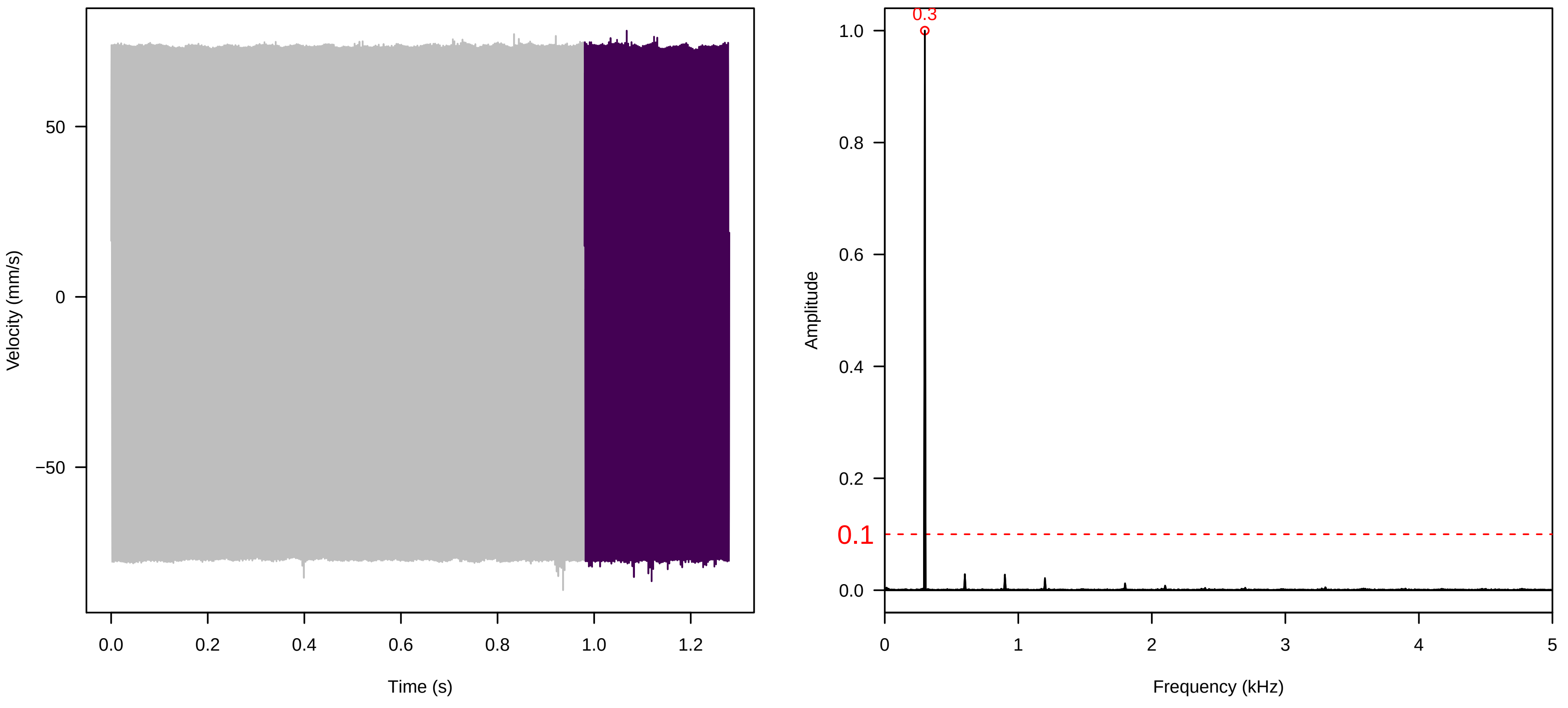

**Vel. = 0.014 ; Str. = Receptacle ; Axis = z ; Fl. accession = 10−s−81−8**

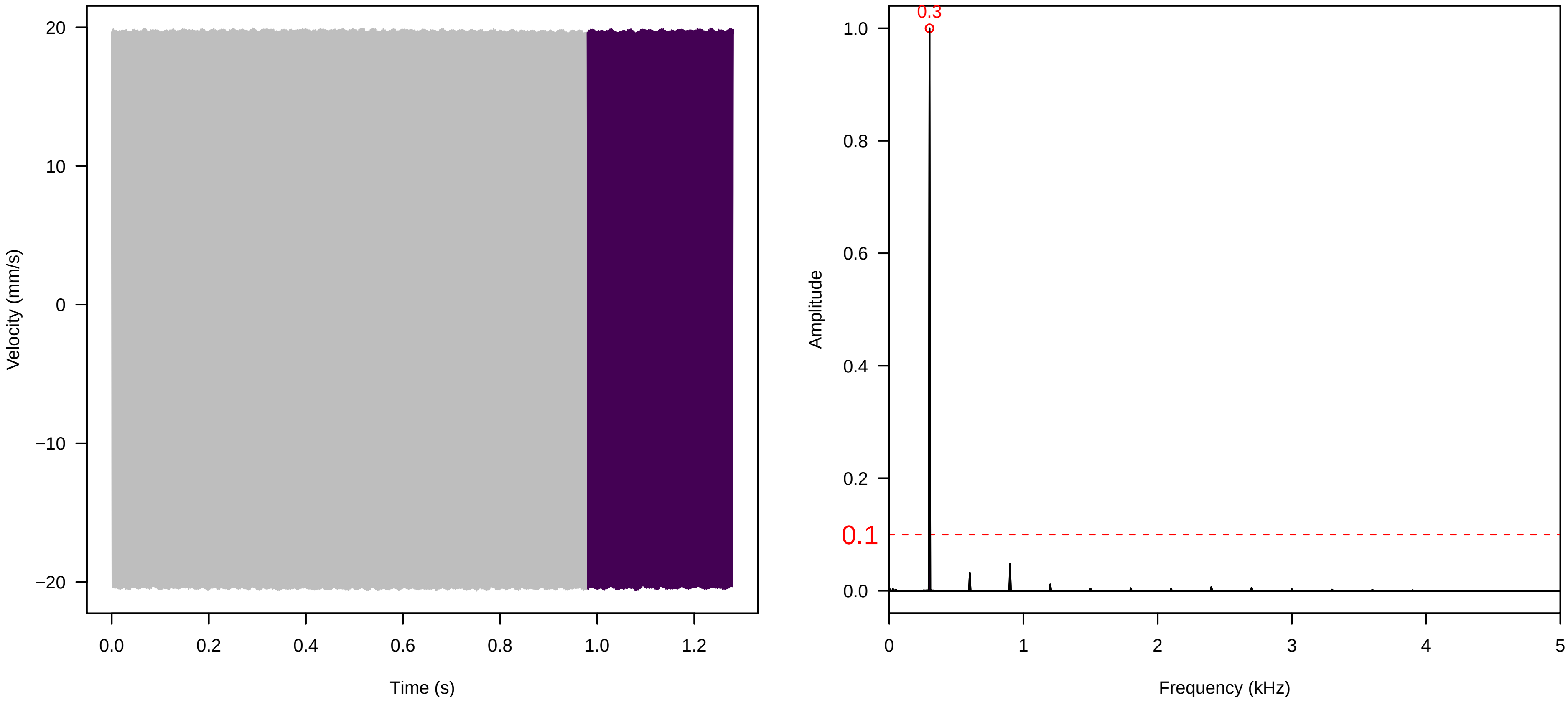


**Vel. = 0.014 ; Str. = PA ; Axis = z ; Fl. accession = 10−s−81−8**

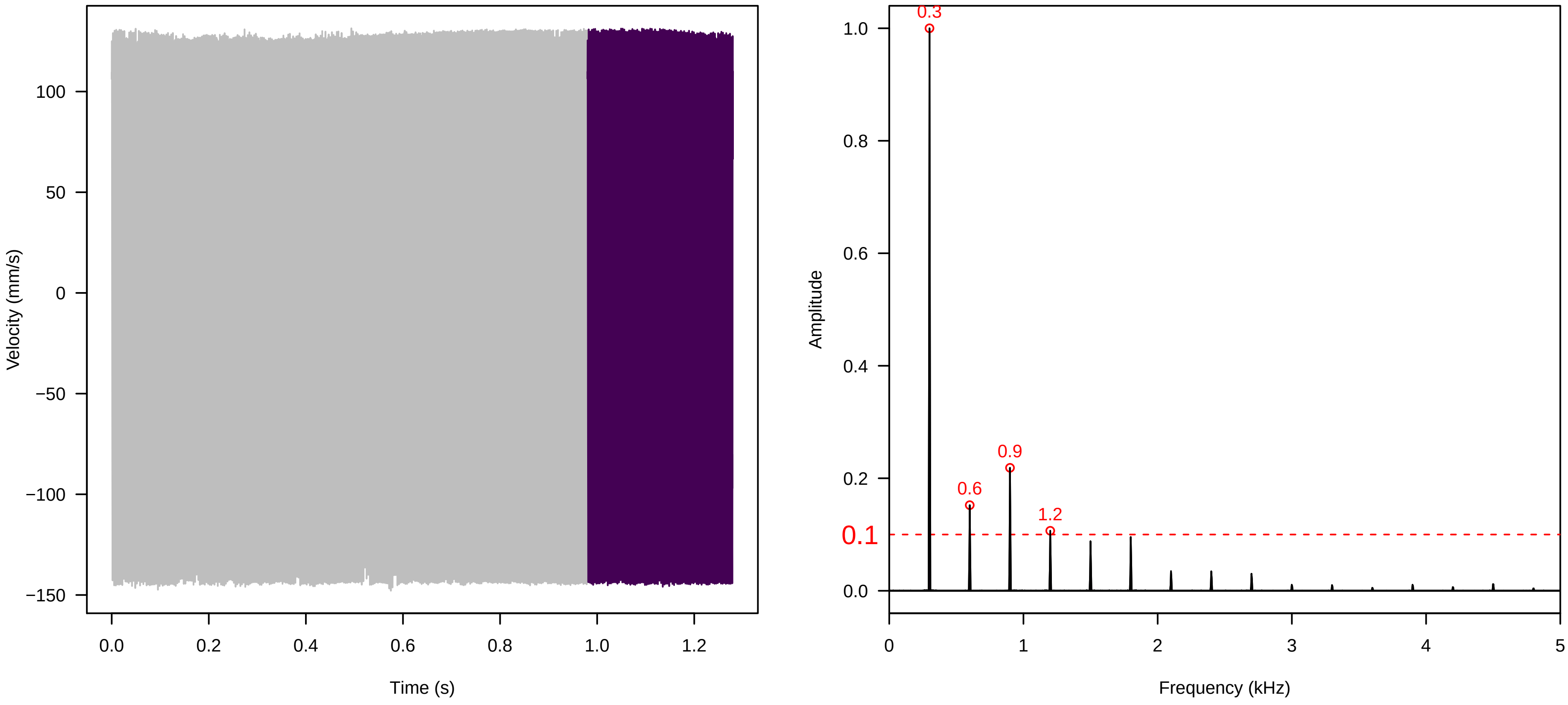


**Vel. = 0.014 ; Str. = Receptacle ; Axis = z ; Fl. accession = 10−s−81−8**

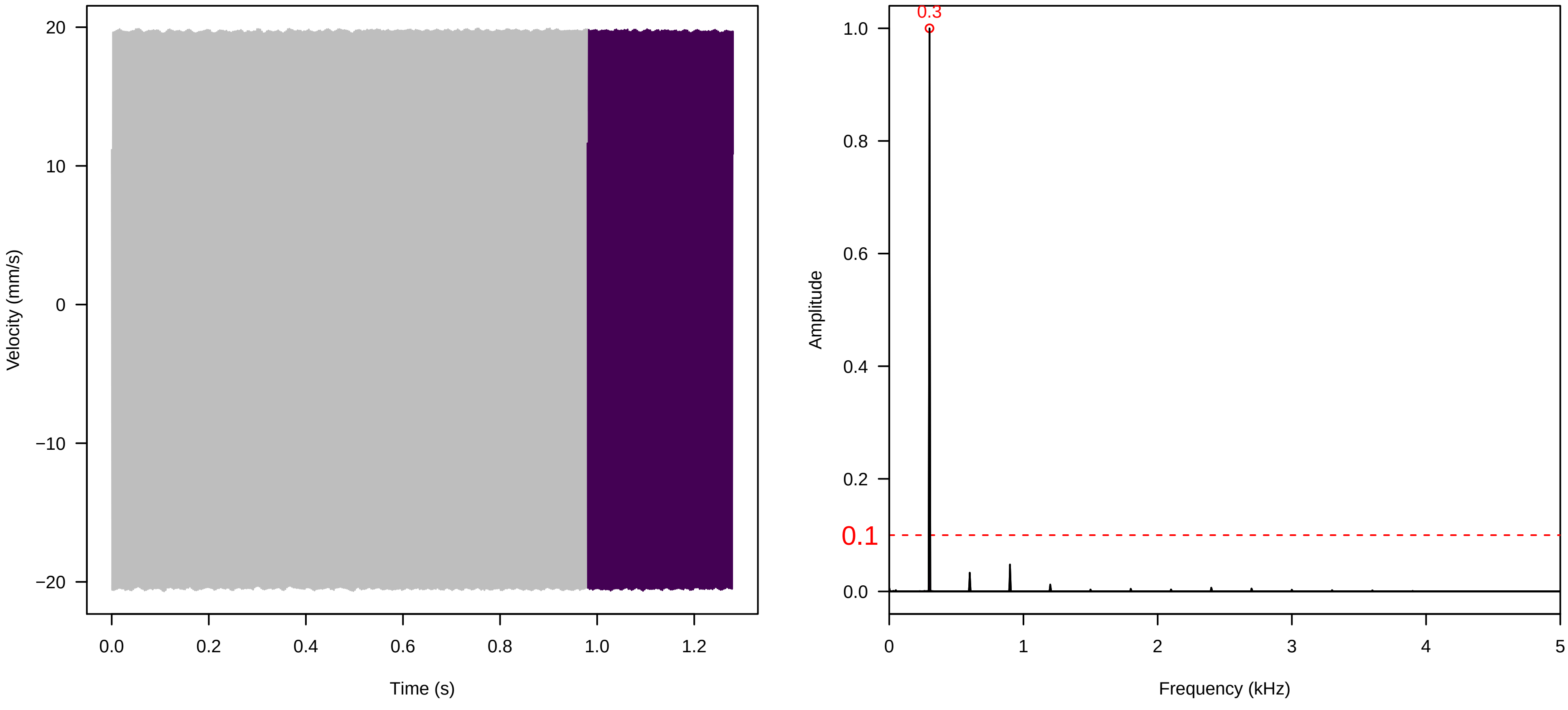

**Vel. = 0.014 ; Str. = Corolla ; Axis = y ; Fl. accession = 10−s−81−8**

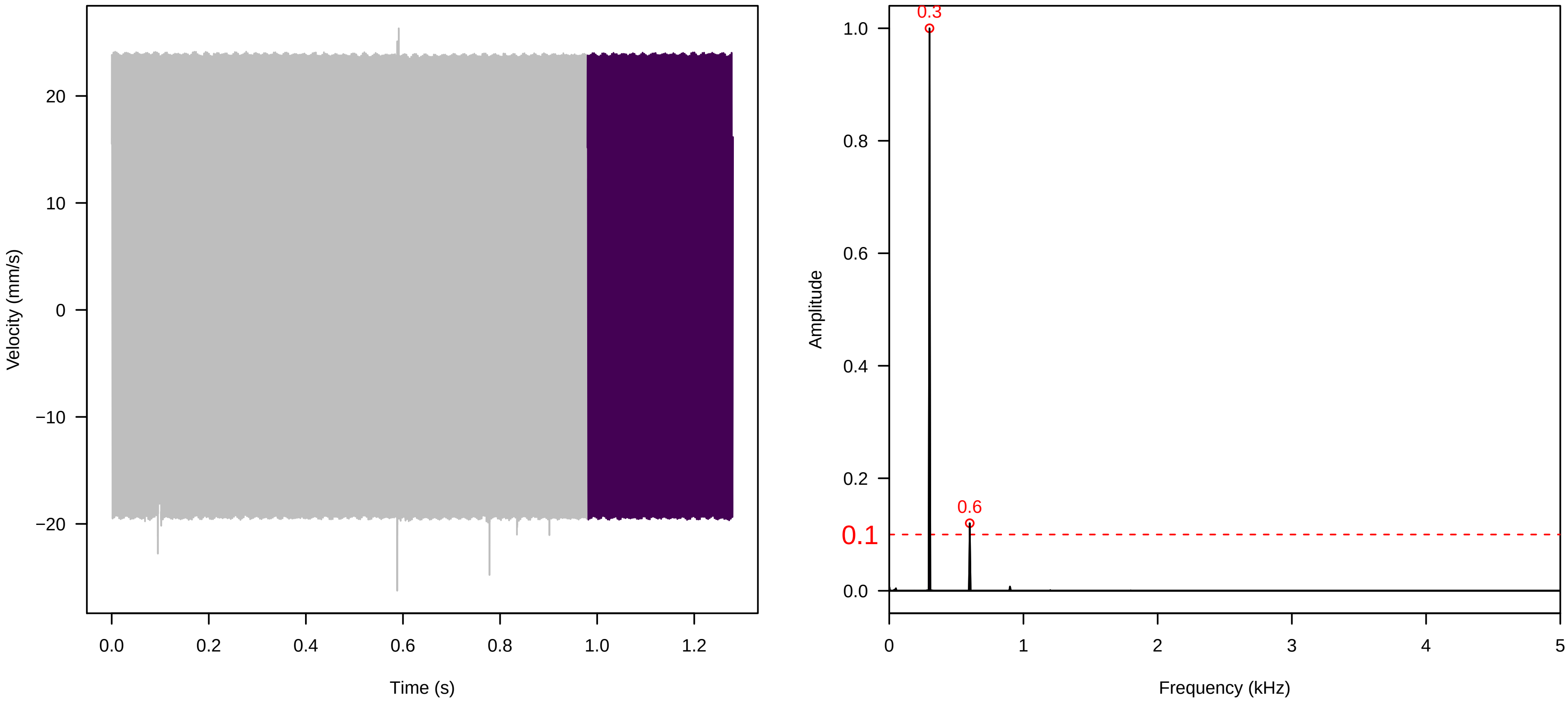


**Vel. = 0.014 ; Str. = Receptacle ; Axis = y ; Fl. accession = 10−s−81−8**

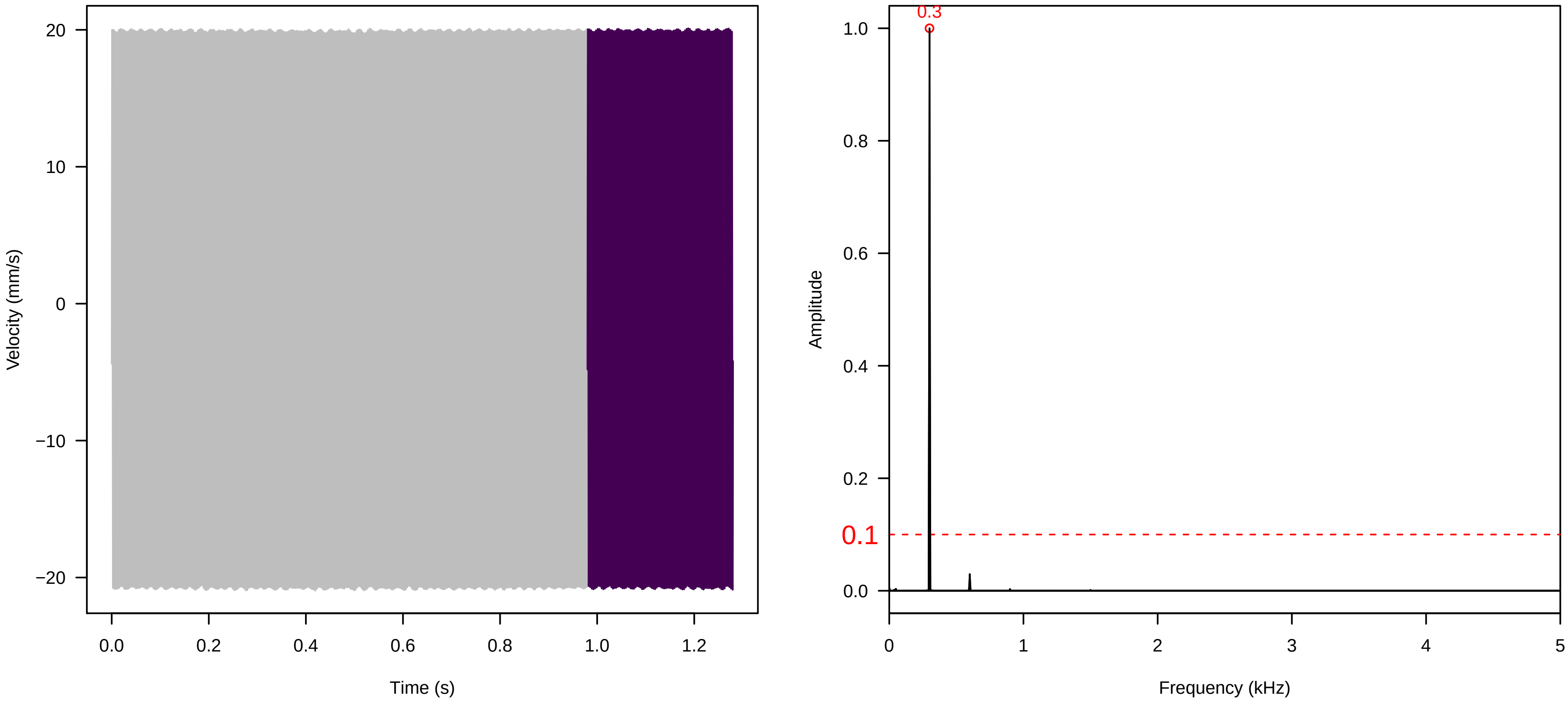


**Vel. = 0.014 ; Str. = FA ; Axis = y ; Fl. accession = 10−s−81−8**

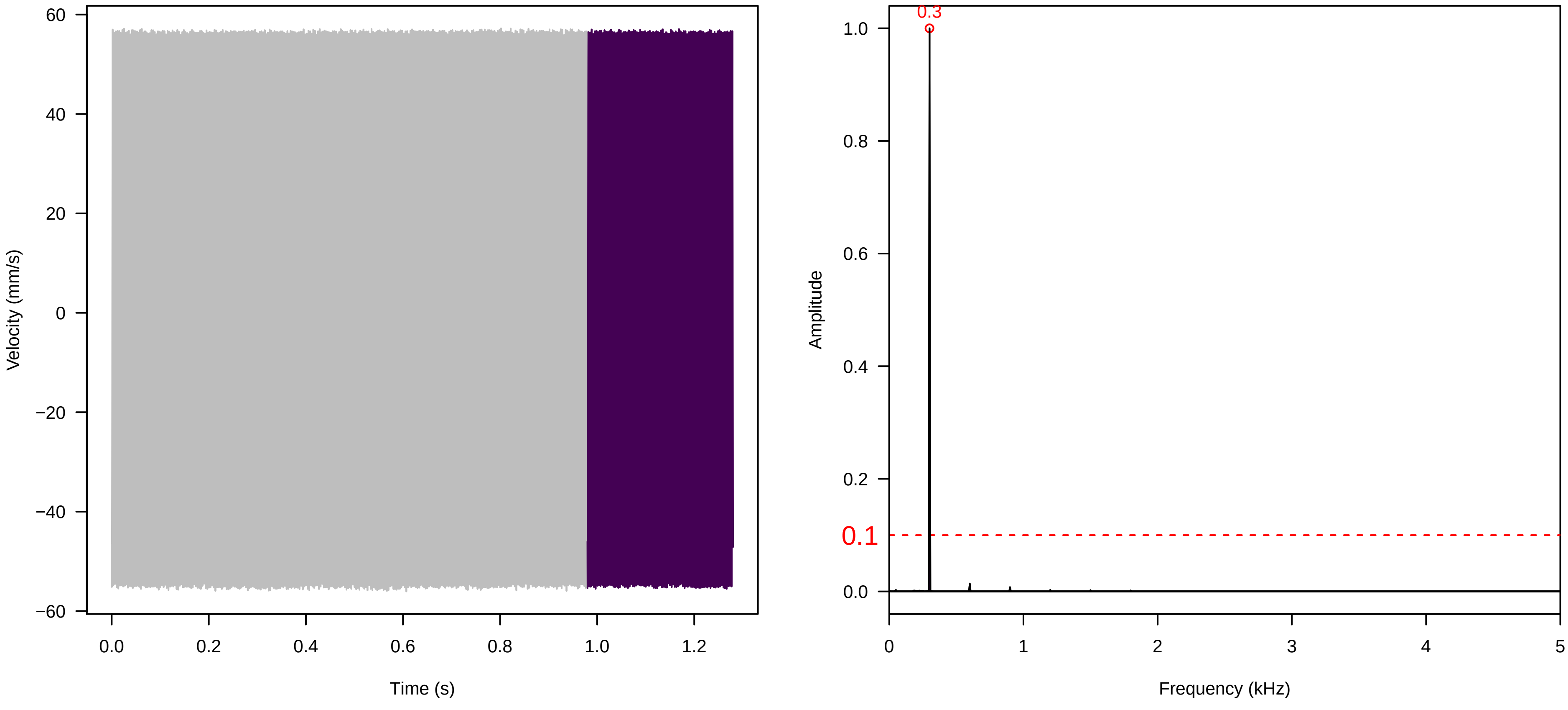

**Vel. = 0.014 ; Str. = Receptacle ; Axis = y ; Fl. accession = 10−s−81−8**

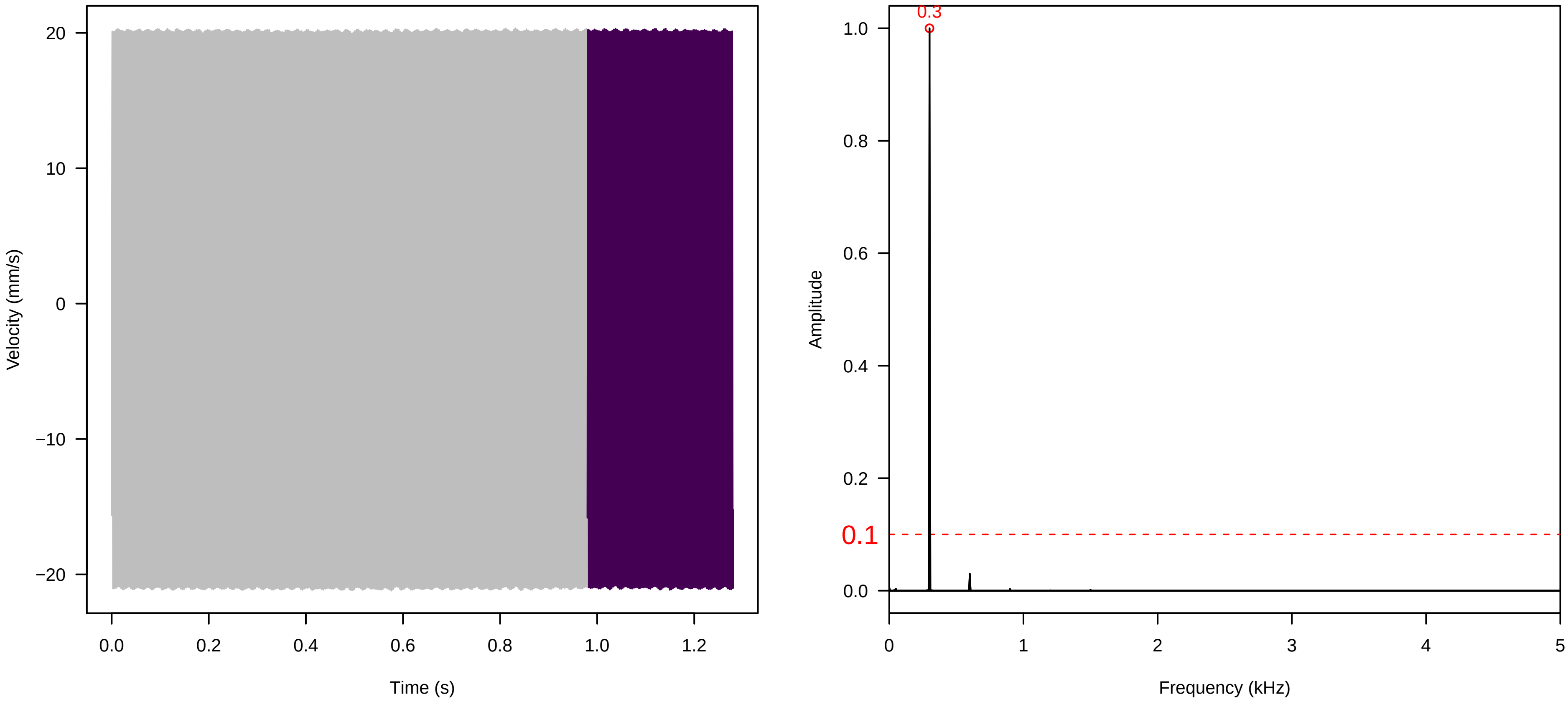


**Vel. = 0.014 ; Str. = PA ; Axis = y ; Fl. accession = 10−s−81−8**

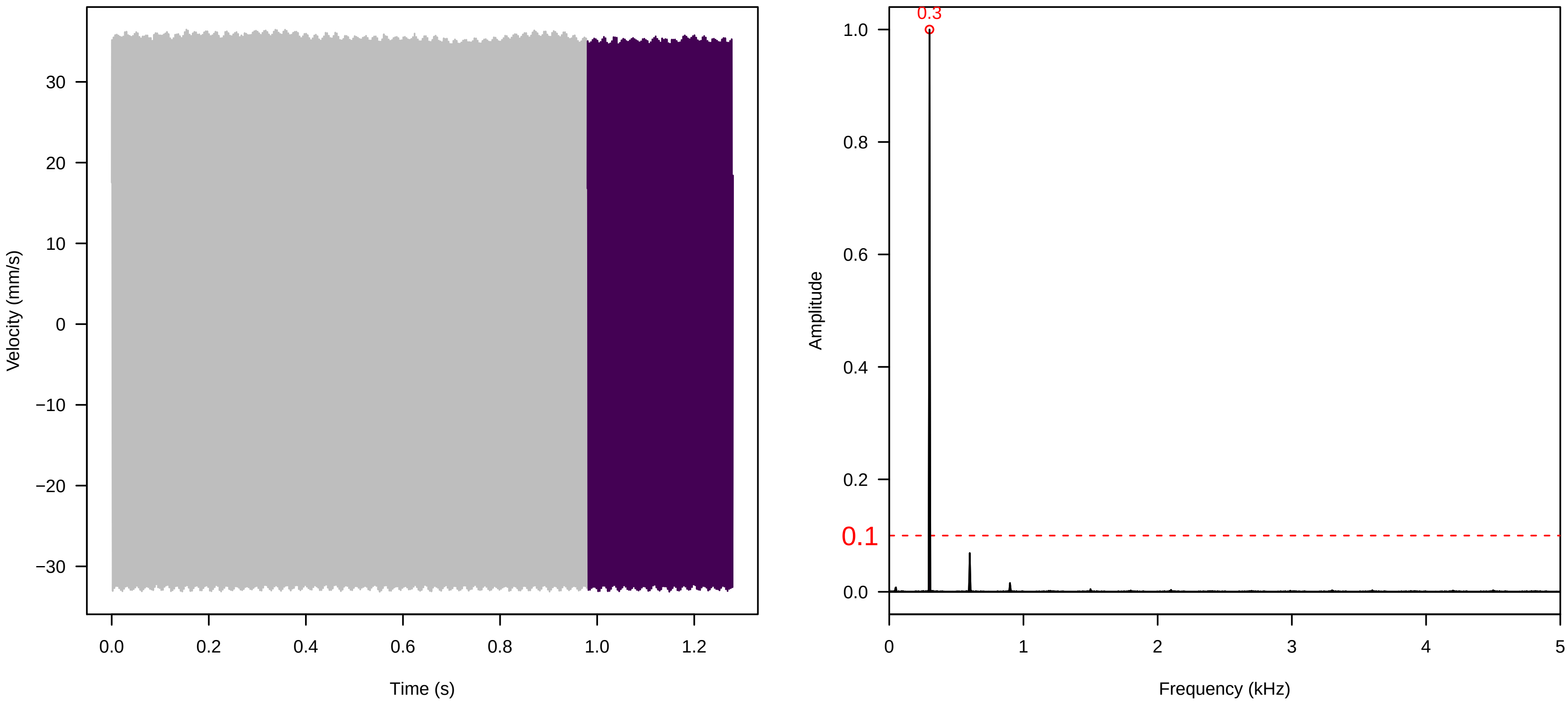


**Vel. = 0.014 ; Str. = Receptacle ; Axis = y ; Fl. accession = 10−s−81−8**

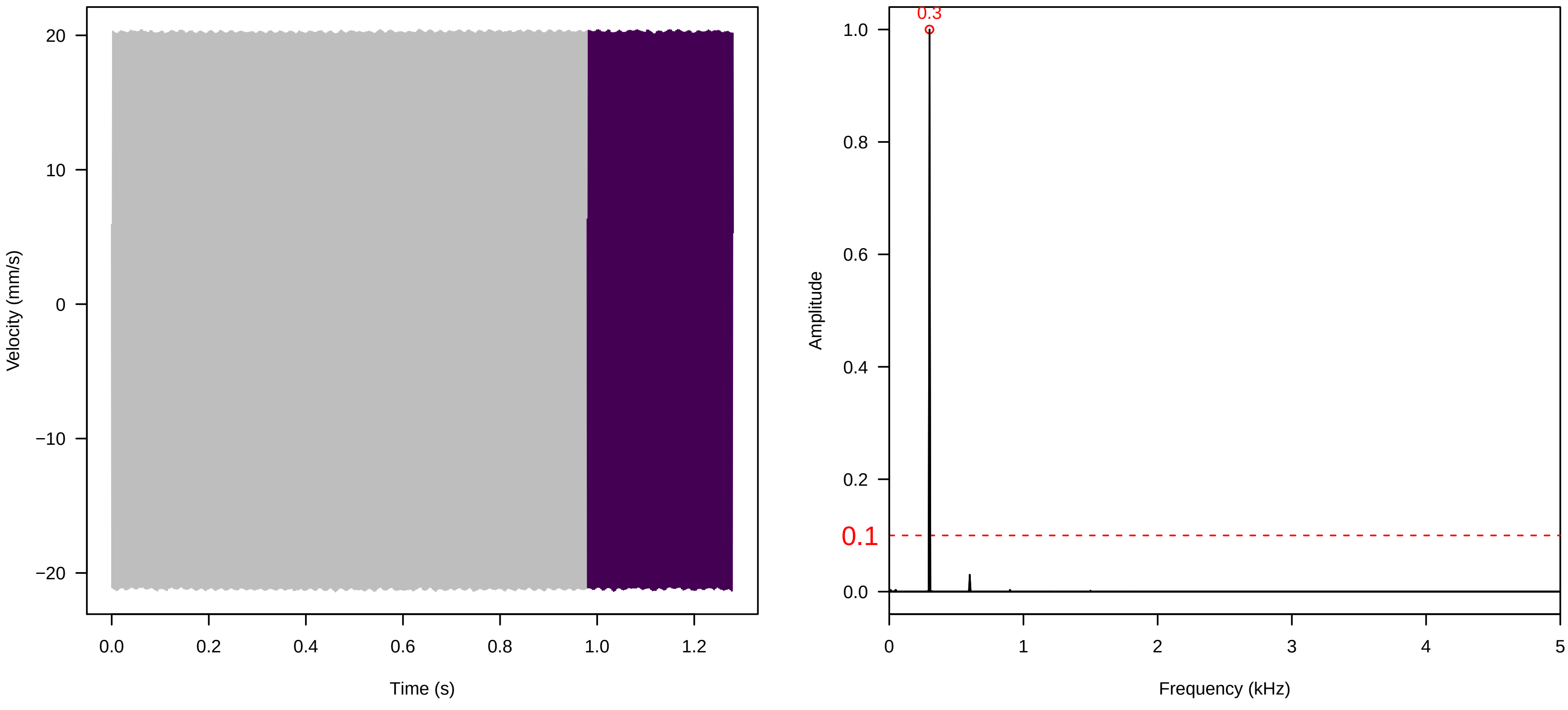

**Vel. = 0.028 ; Str. = PA ; Axis = y ; Fl. accession = 10−s−81−8**

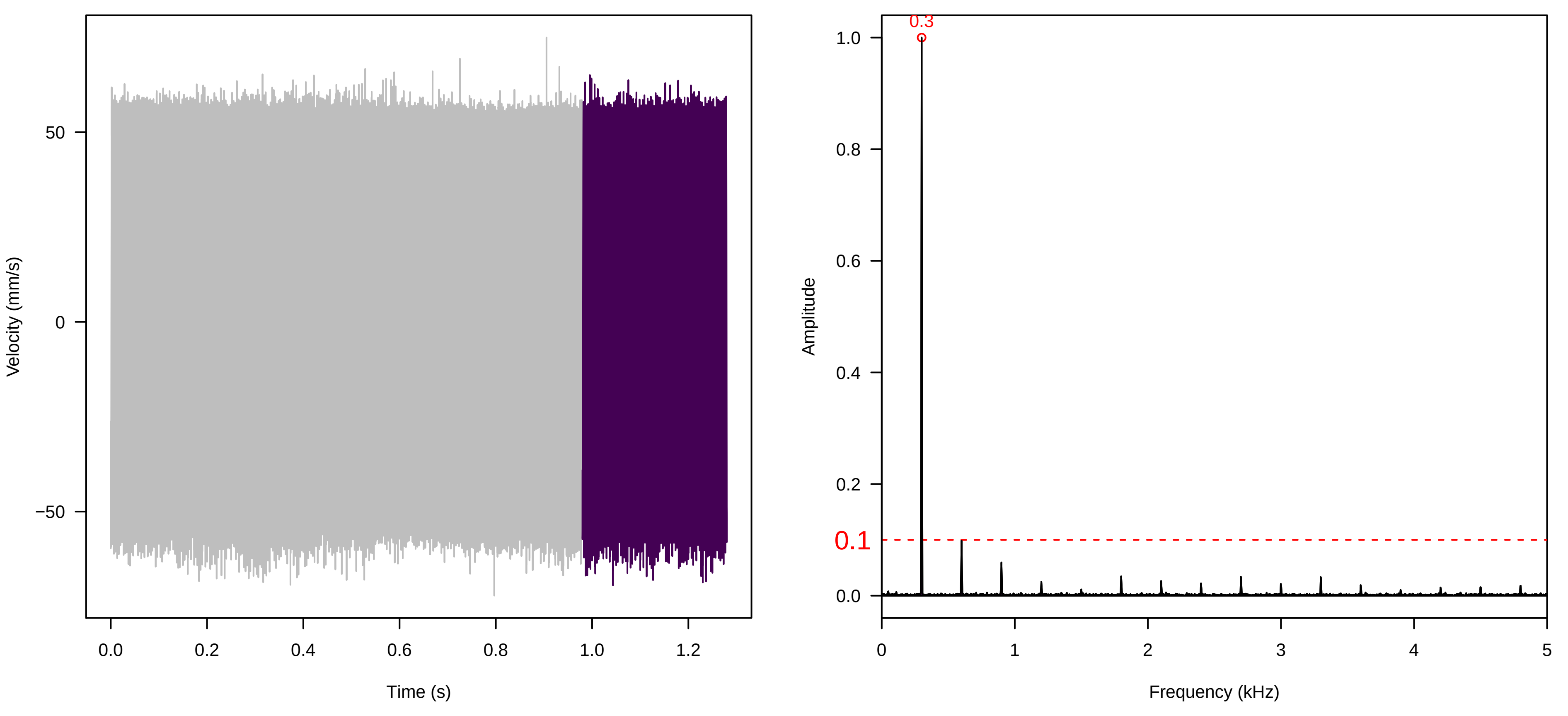


**Vel. = 0.028 ; Str. = Receptacle ; Axis = y ; Fl. accession = 10−s−81−8**

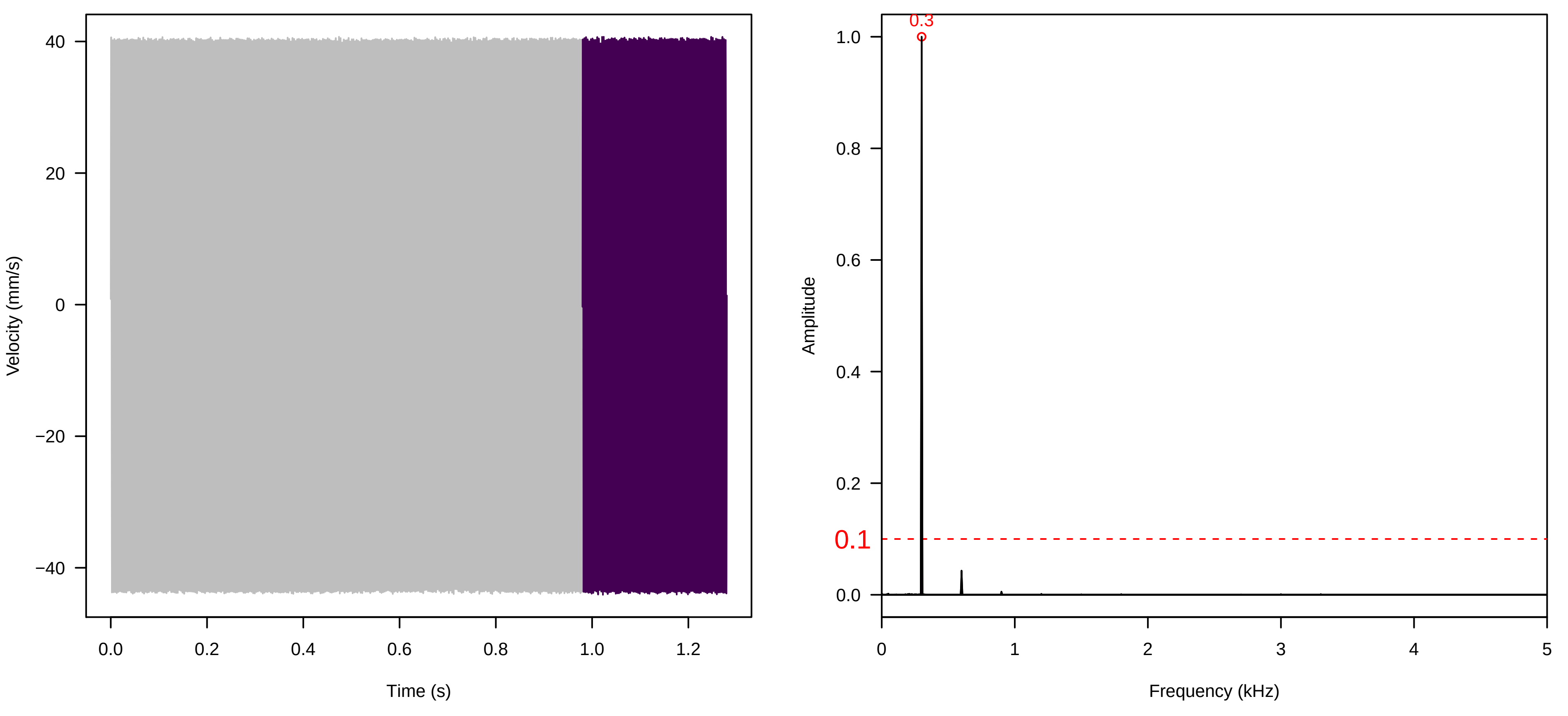


**Vel. = 0.028 ; Str. = FA ; Axis = y ; Fl. accession = 10−s−81−8**

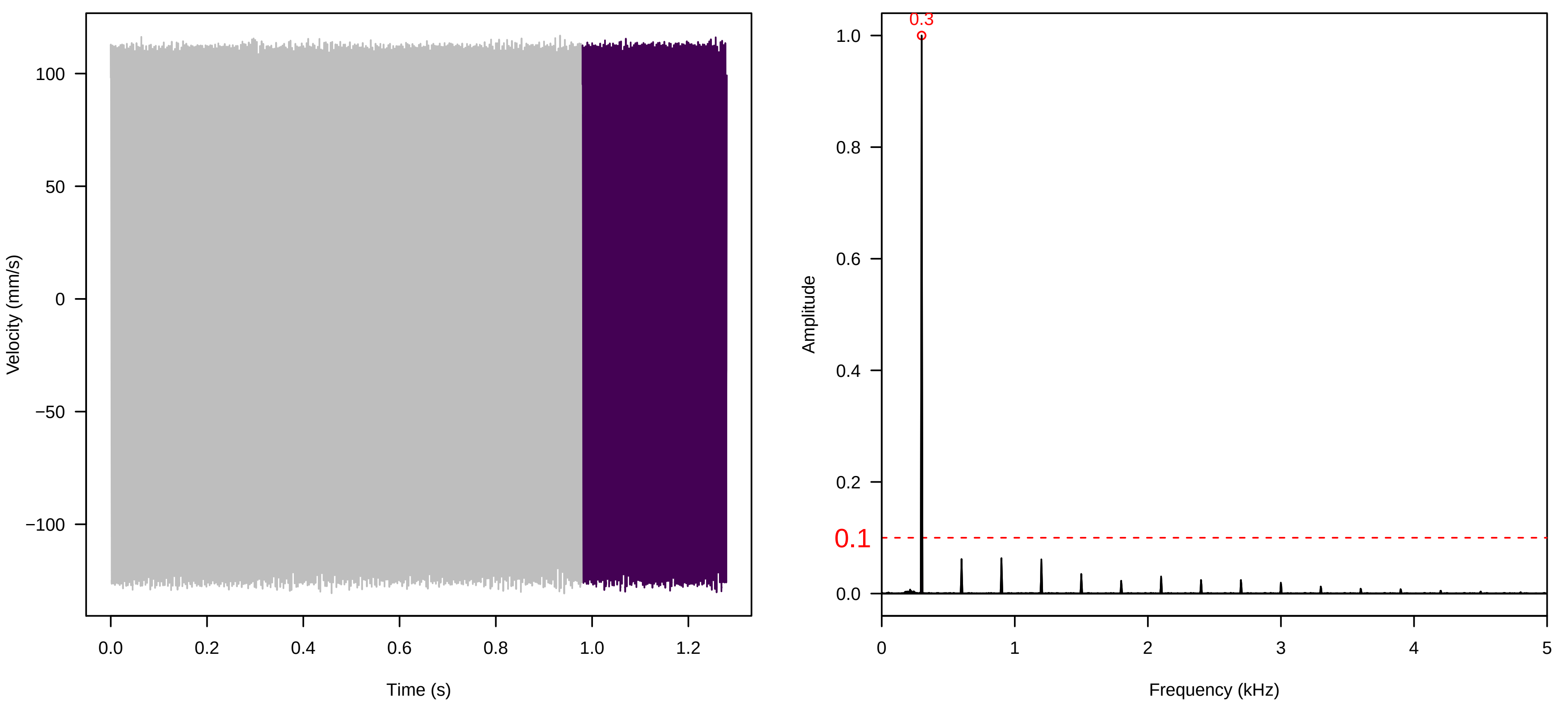

**Vel. = 0.028 ; Str. = Receptacle ; Axis = y ; Fl. accession = 10−s−81−8**

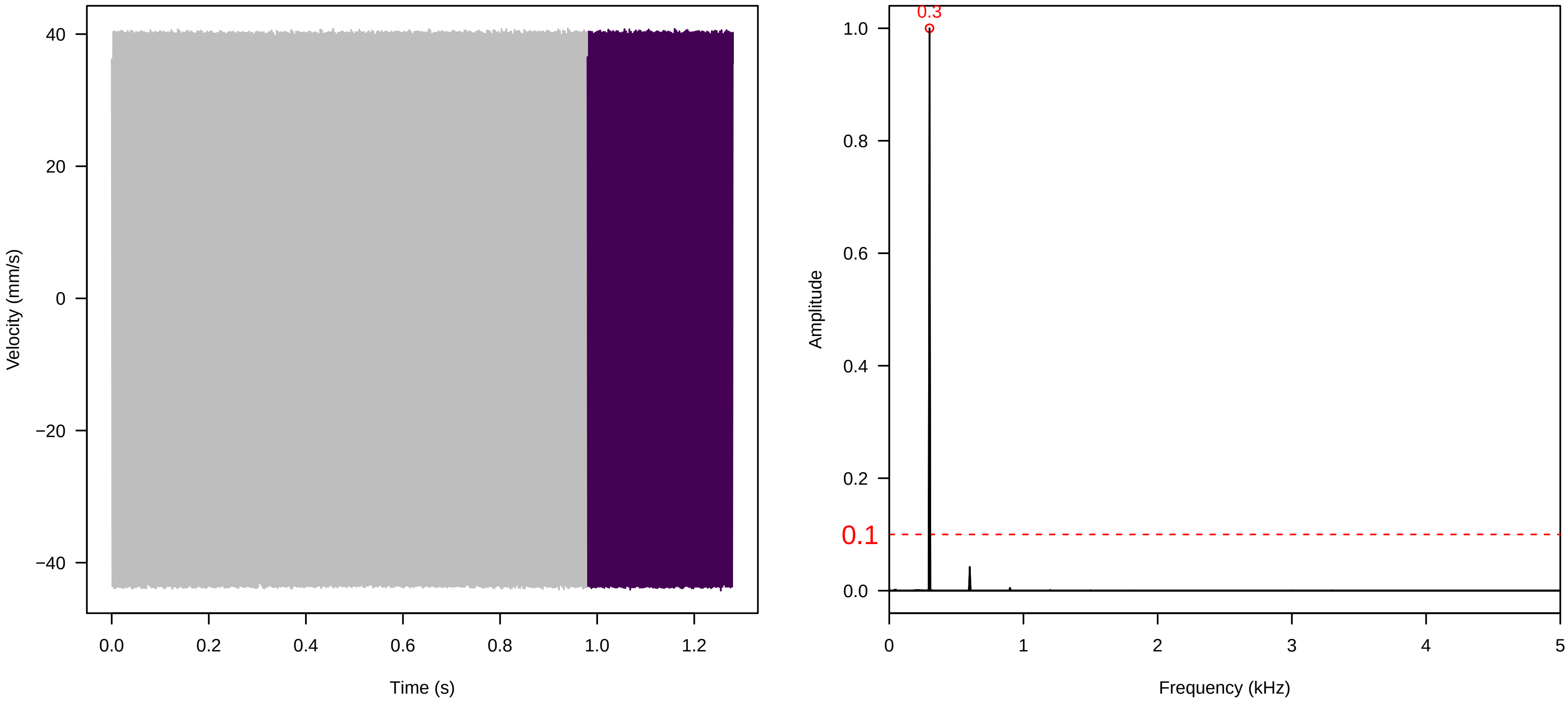


**Vel. = 0.028 ; Str. = Corolla ; Axis = y ; Fl. accession = 10−s−81−8**

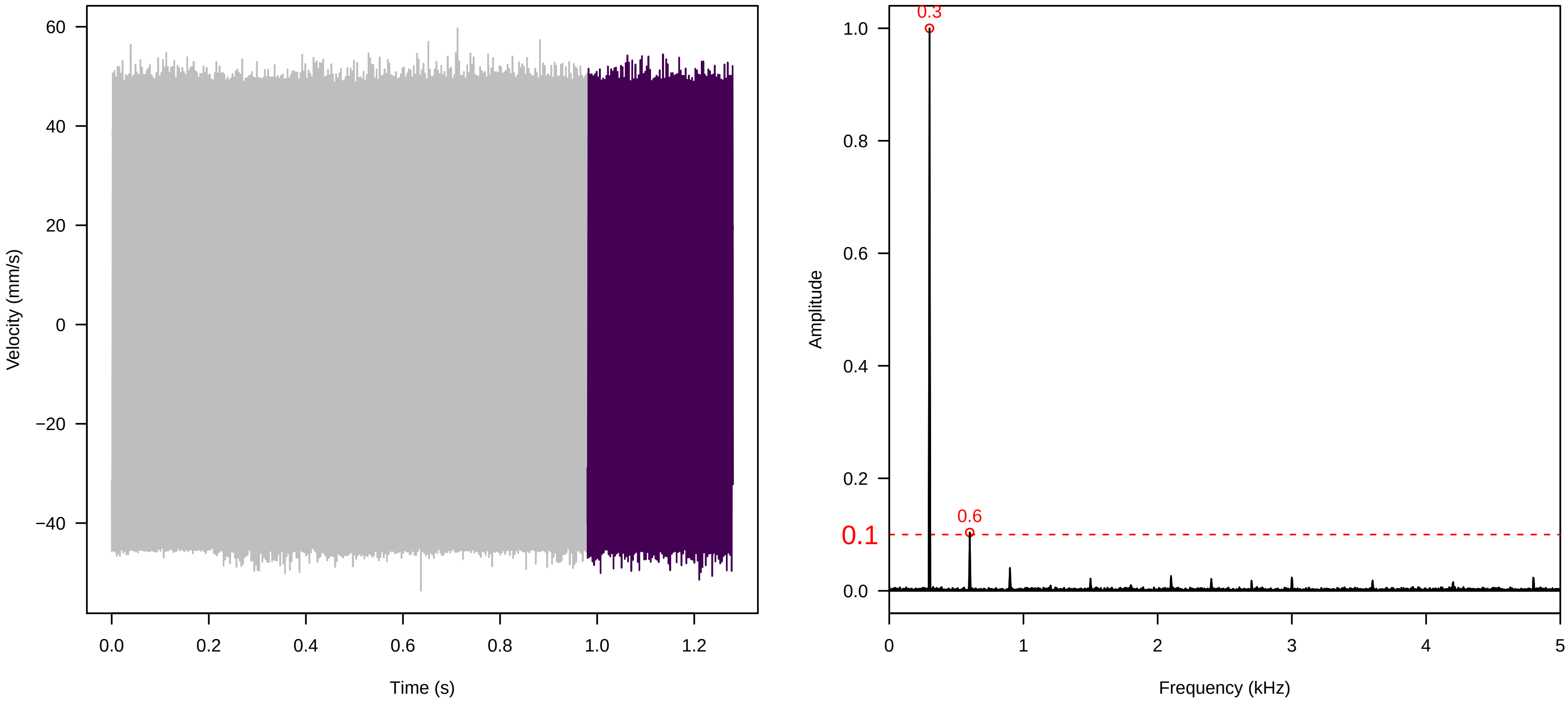


**Vel. = 0.028 ; Str. = Receptacle ; Axis = y ; Fl. accession = 10−s−81−8**

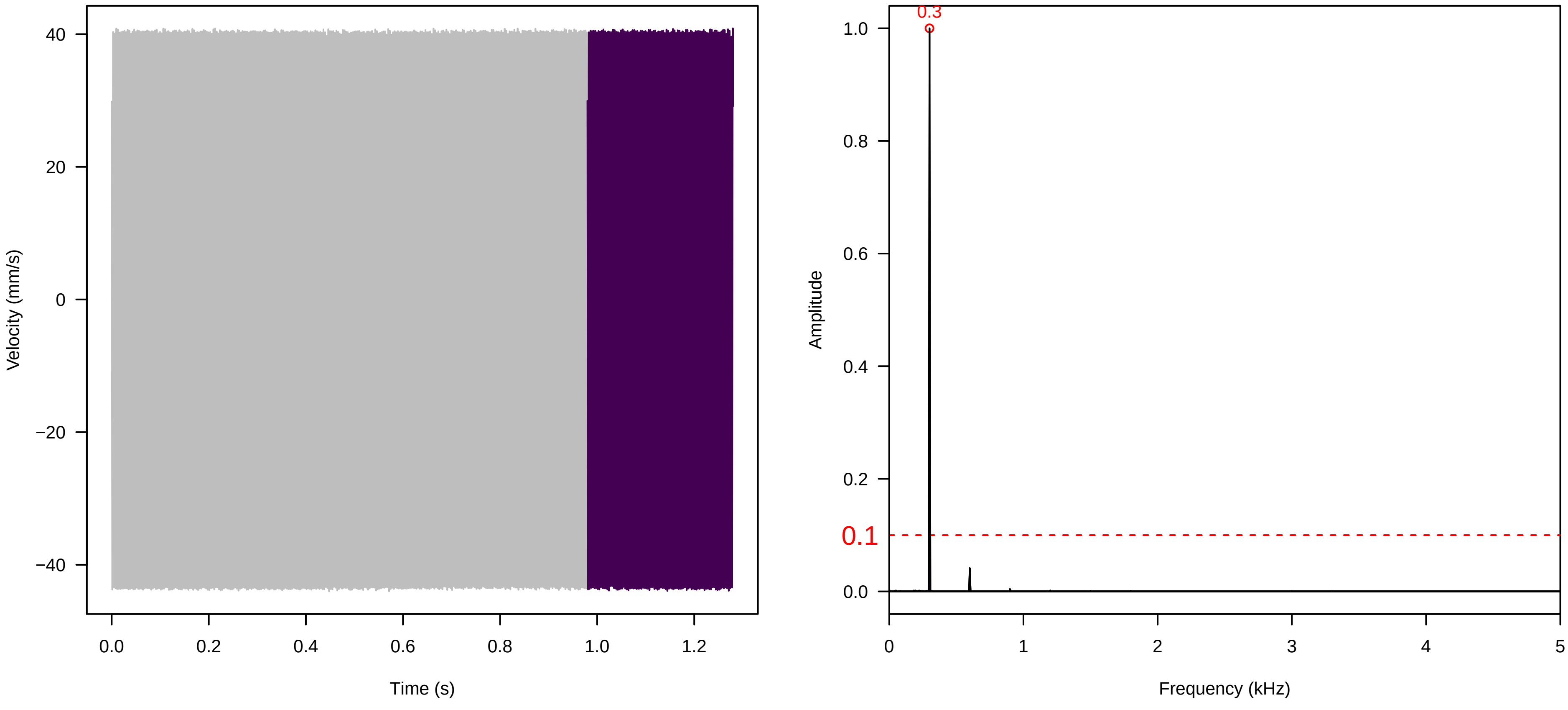

**Vel. = 0.057 ; Str. = Corolla ; Axis = y ; Fl. accession = 10−s−81−8**

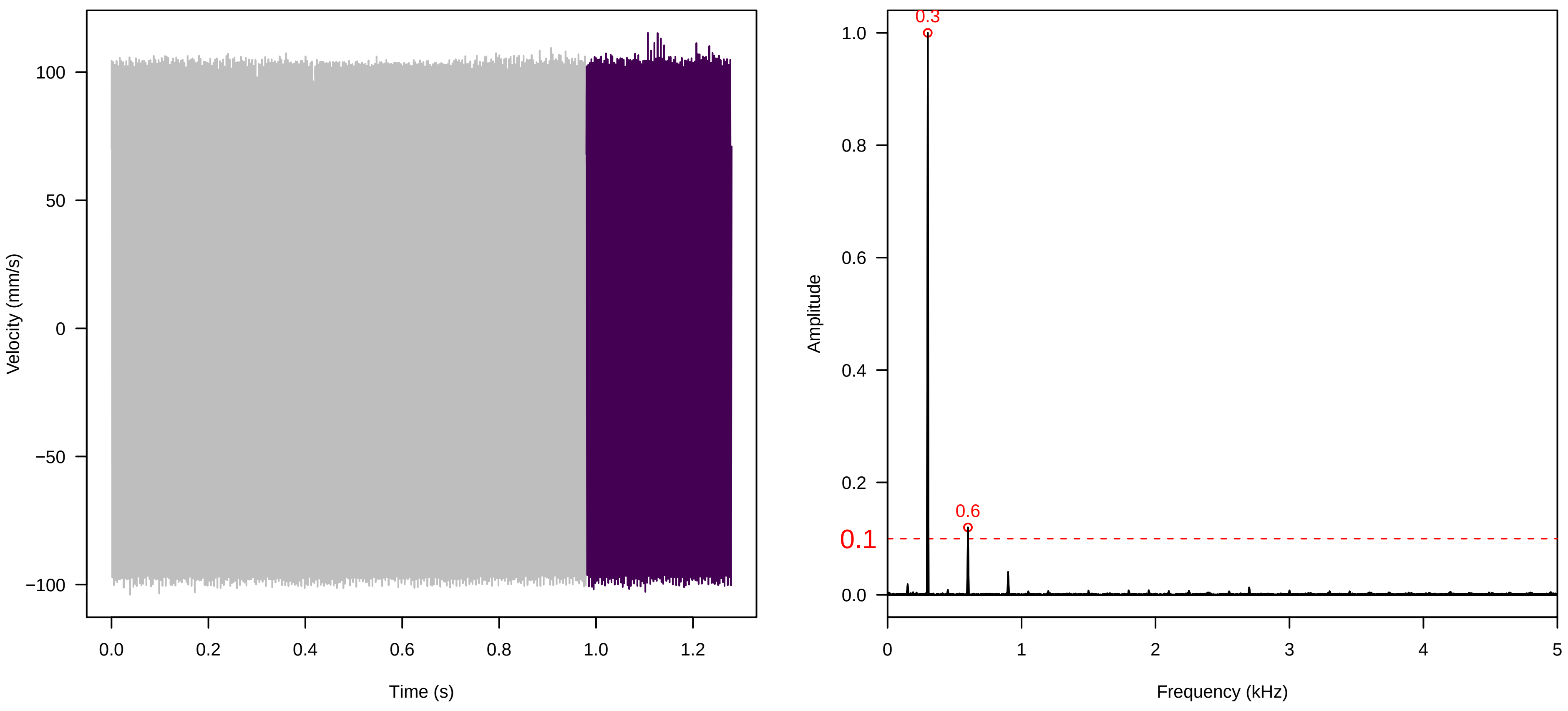


**Vel. = 0.057 ; Str. = Receptacle ; Axis = y ; Fl. accession = 10−s−81−8**

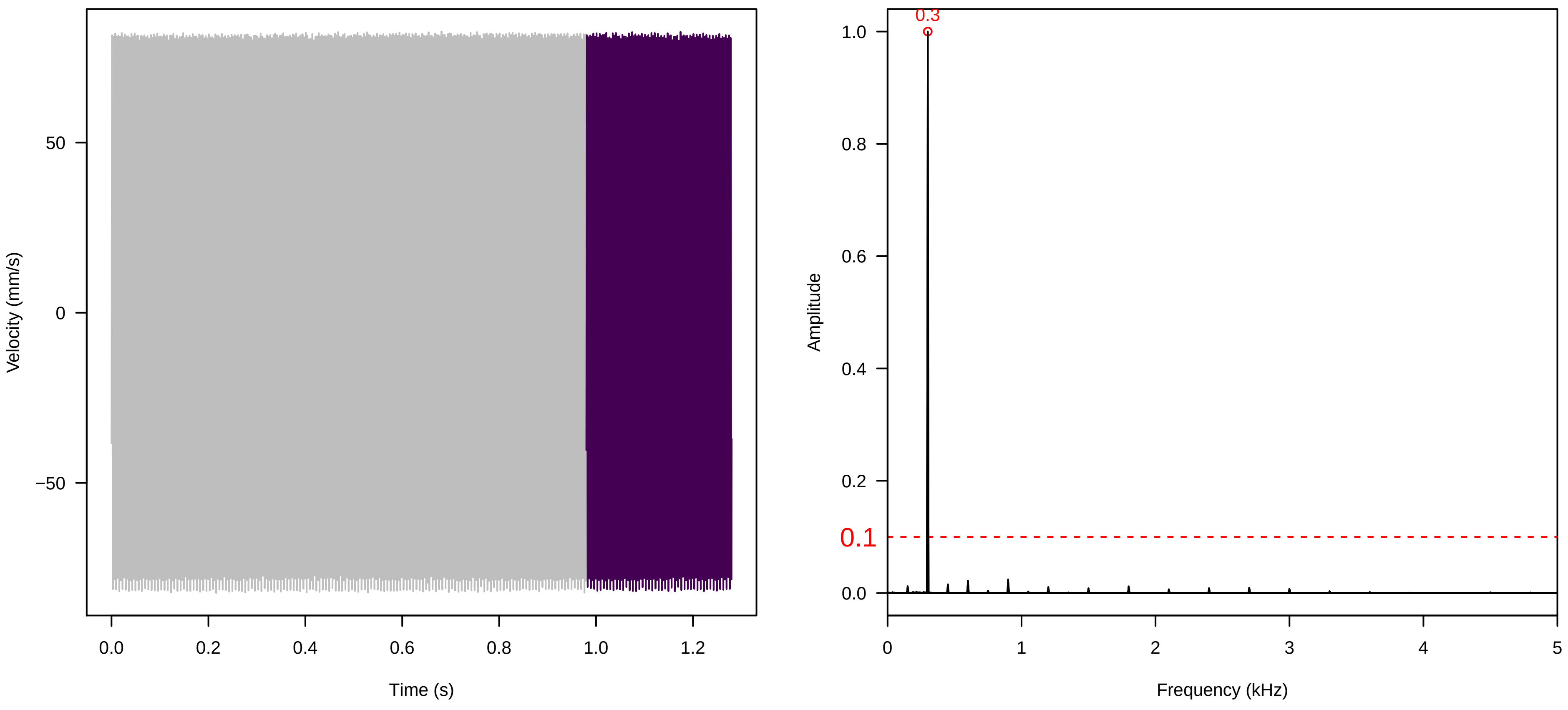


**Vel. = 0.057 ; Str. = FA ; Axis = y ; Fl. accession = 10−s−81−8**

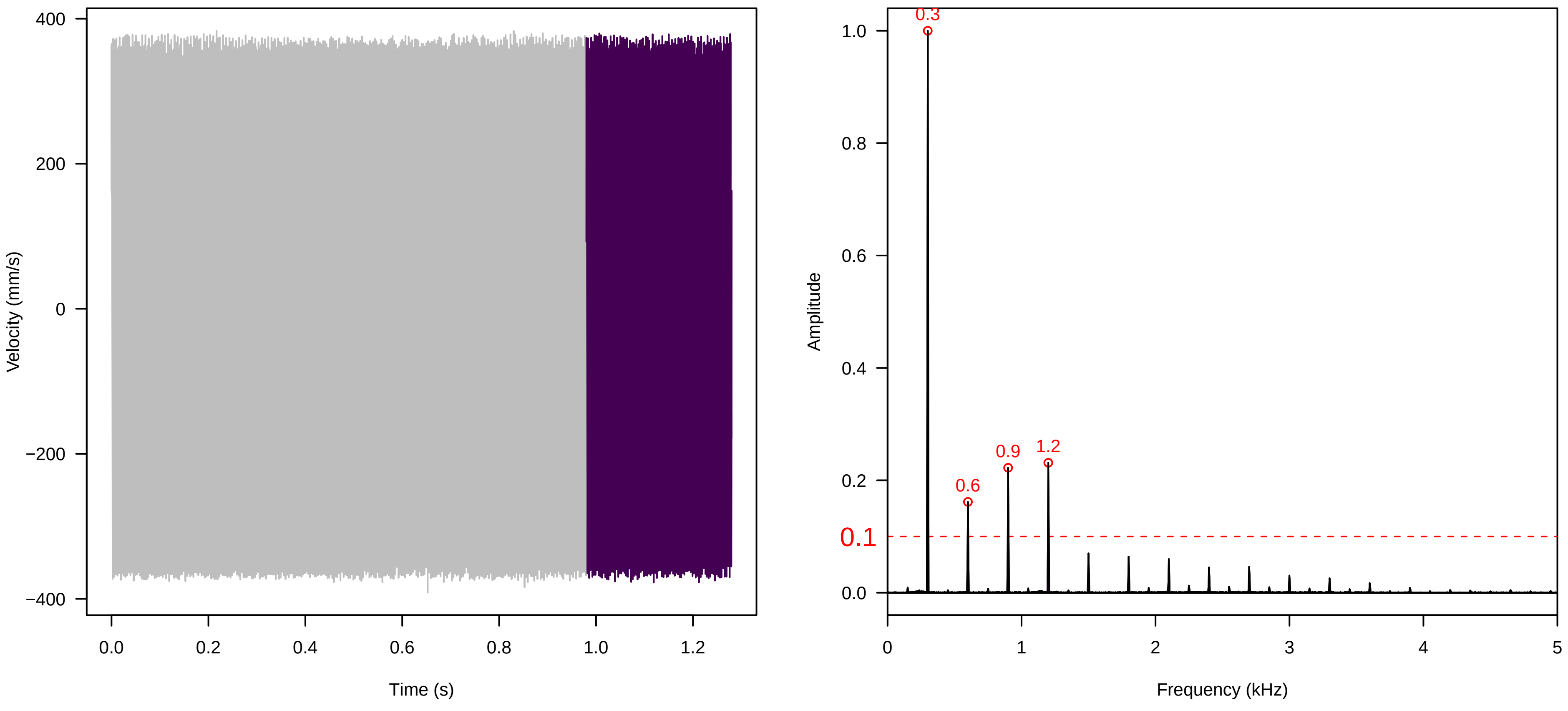

**Vel. = 0.057 ; Str. = Receptacle ; Axis = y ; Fl. accession = 10−s−81−8**

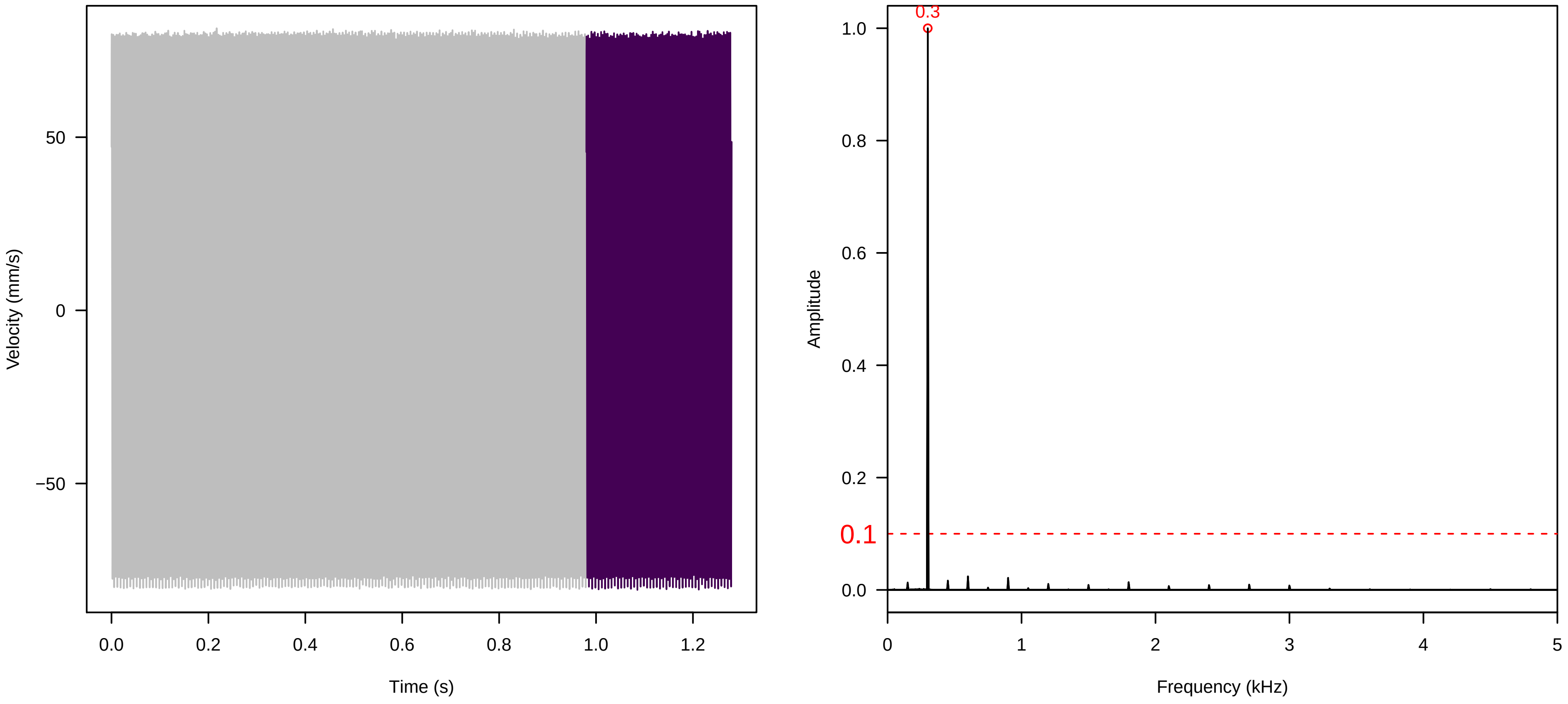


**Vel. = 0.057 ; Str. = PA ; Axis = y ; Fl. accession = 10−s−81−8**

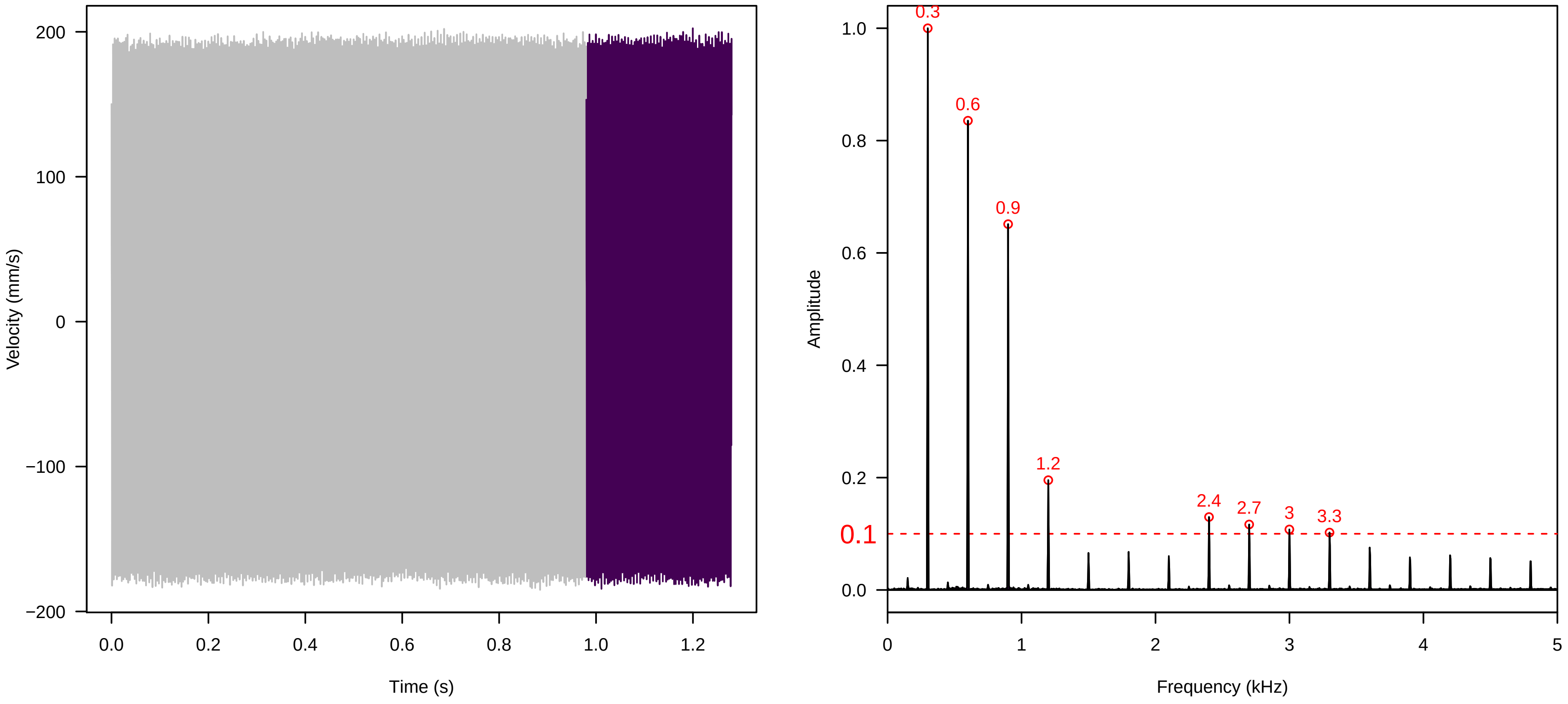


**Vel. = 0.057 ; Str. = Receptacle ; Axis = y ; Fl. accession = 10−s−81−8**

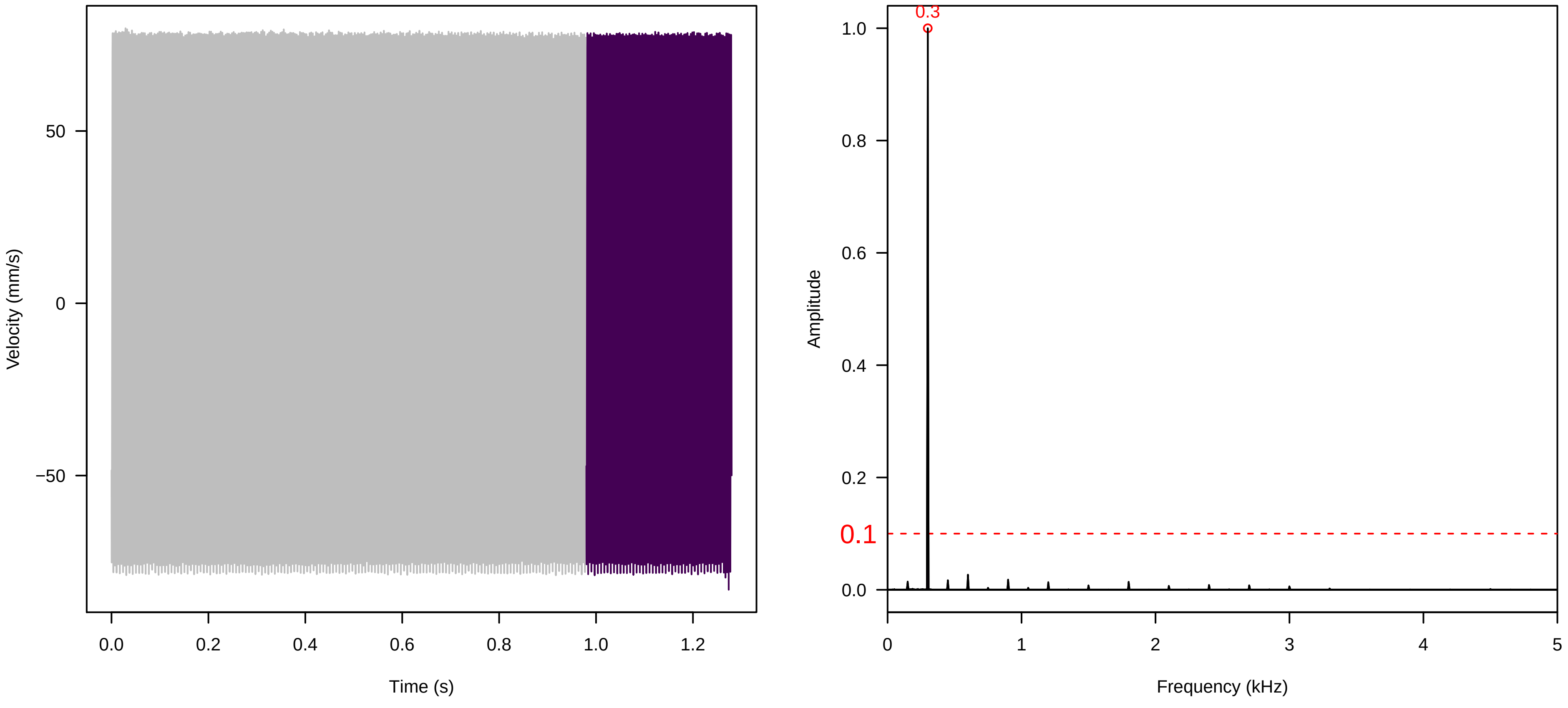

**Vel. = 0.057 ; Str. = Corolla ; Axis = y ; Fl. accession = 10−s−79−2**

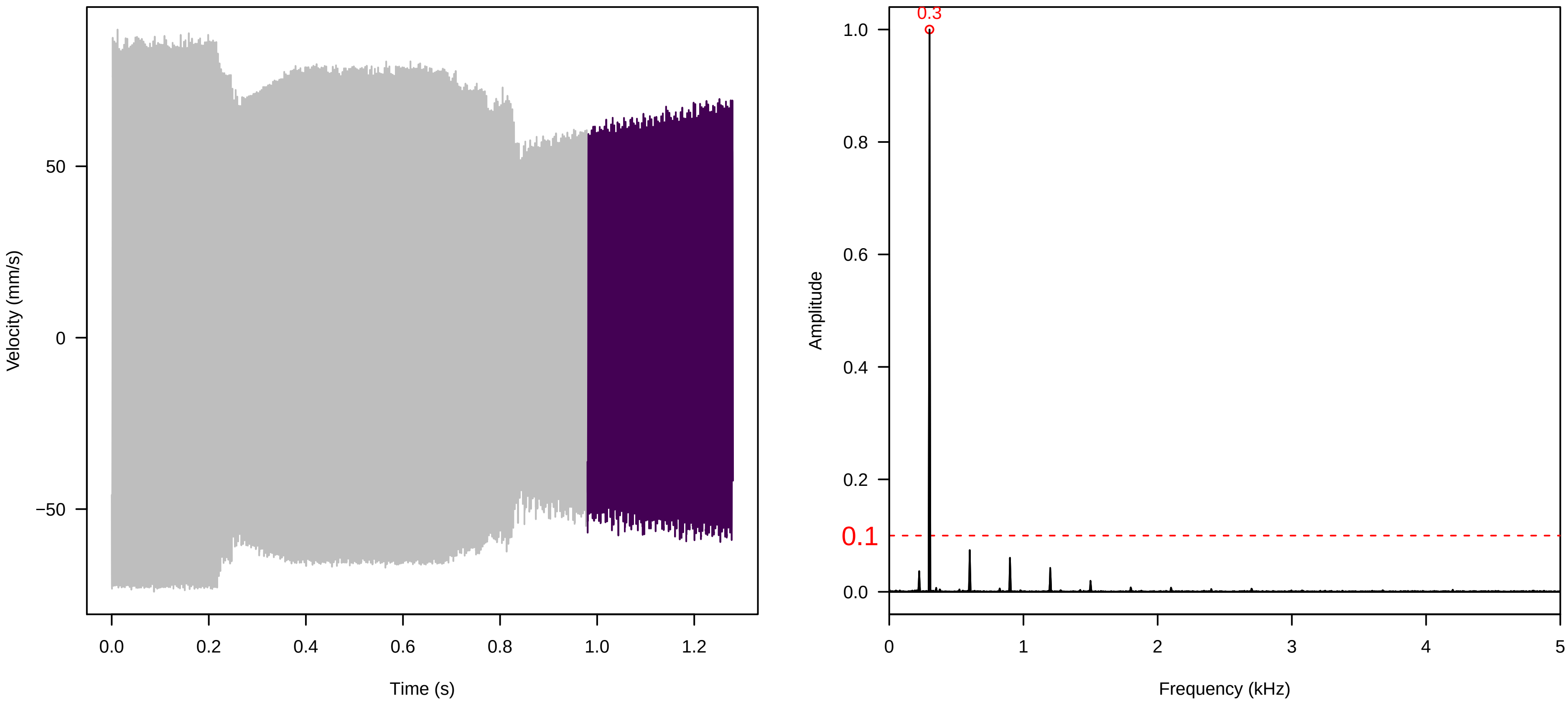


**Vel. = 0.057 ; Str. = Receptacle ; Axis = y ; Fl. accession = 10−s−79−2**

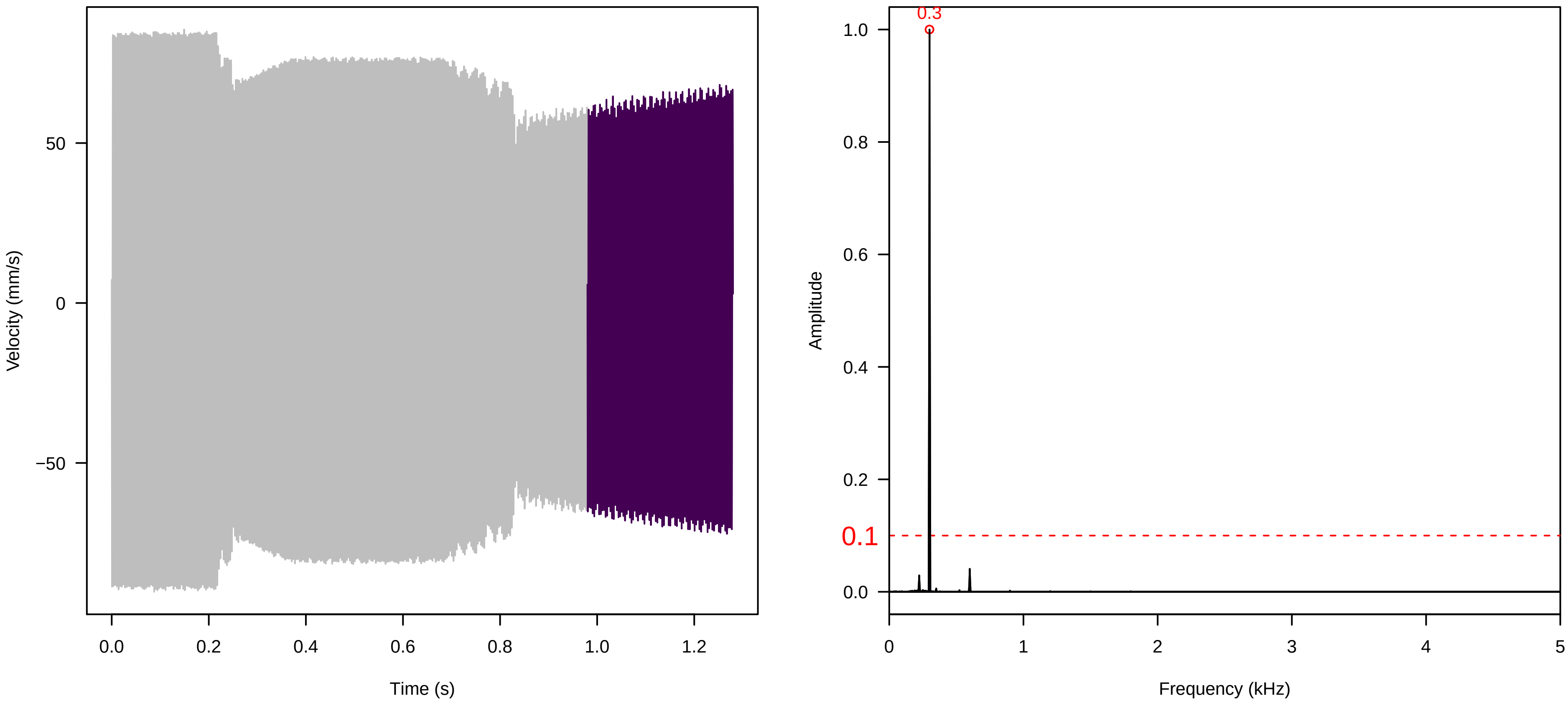


**Vel. = 0.057 ; Str. = FA ; Axis = y ; Fl. accession = 10−s−79−2**

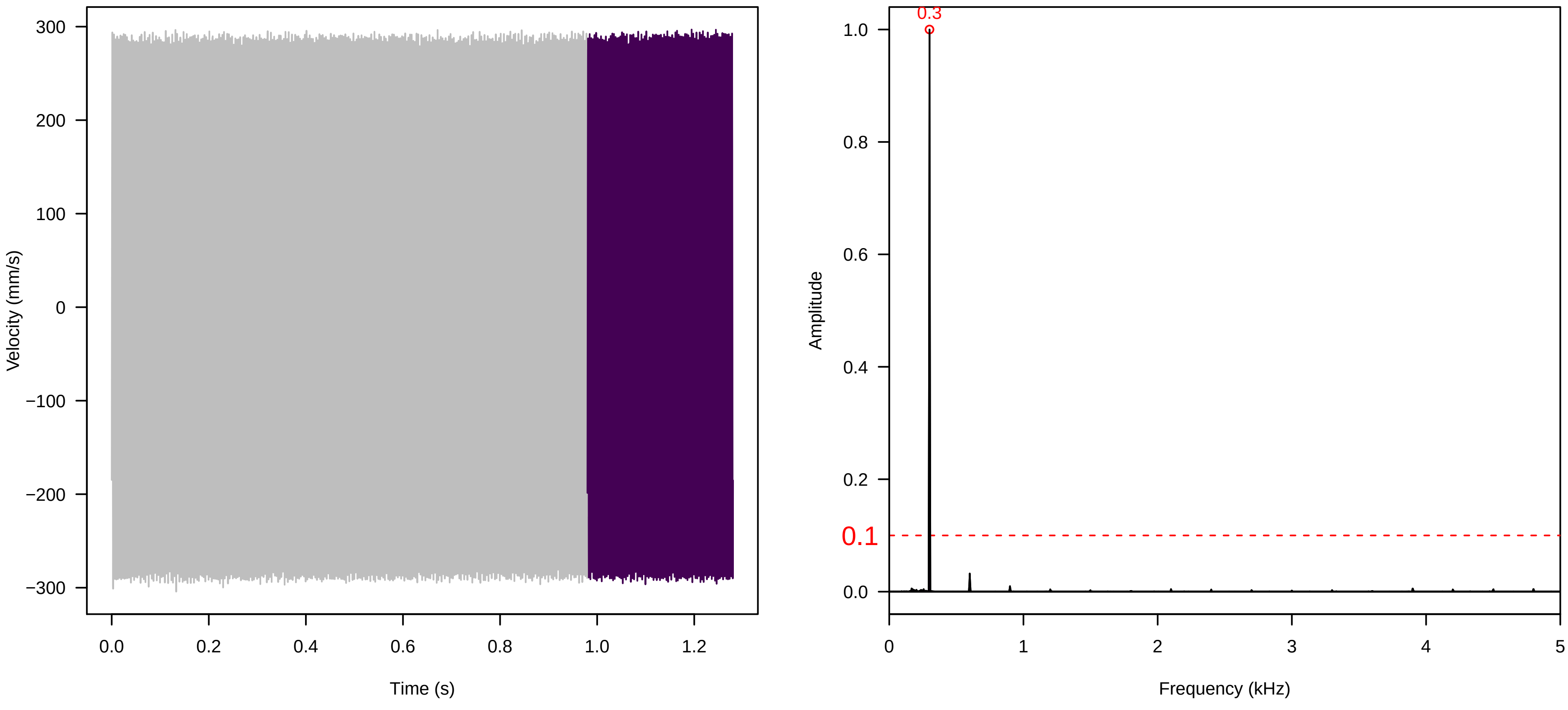

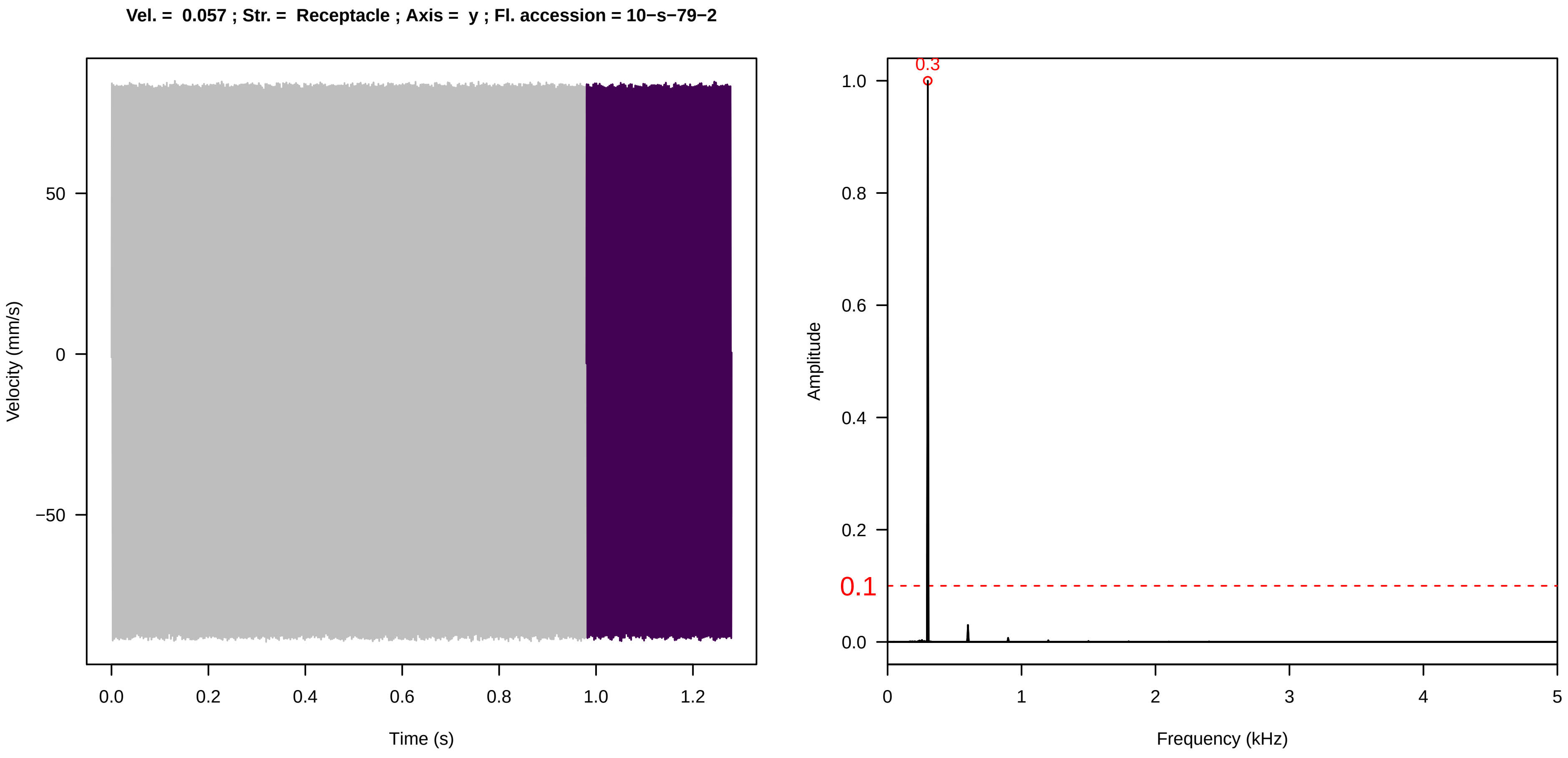
Vel. = 0.057 ; Str. = Receptacle ; Axis = y ; Fl. accession = 10−s−79−2
Velocity (mm/s)
Time (s)
Amplitude
Frequency (kHz)
0.3
0.1

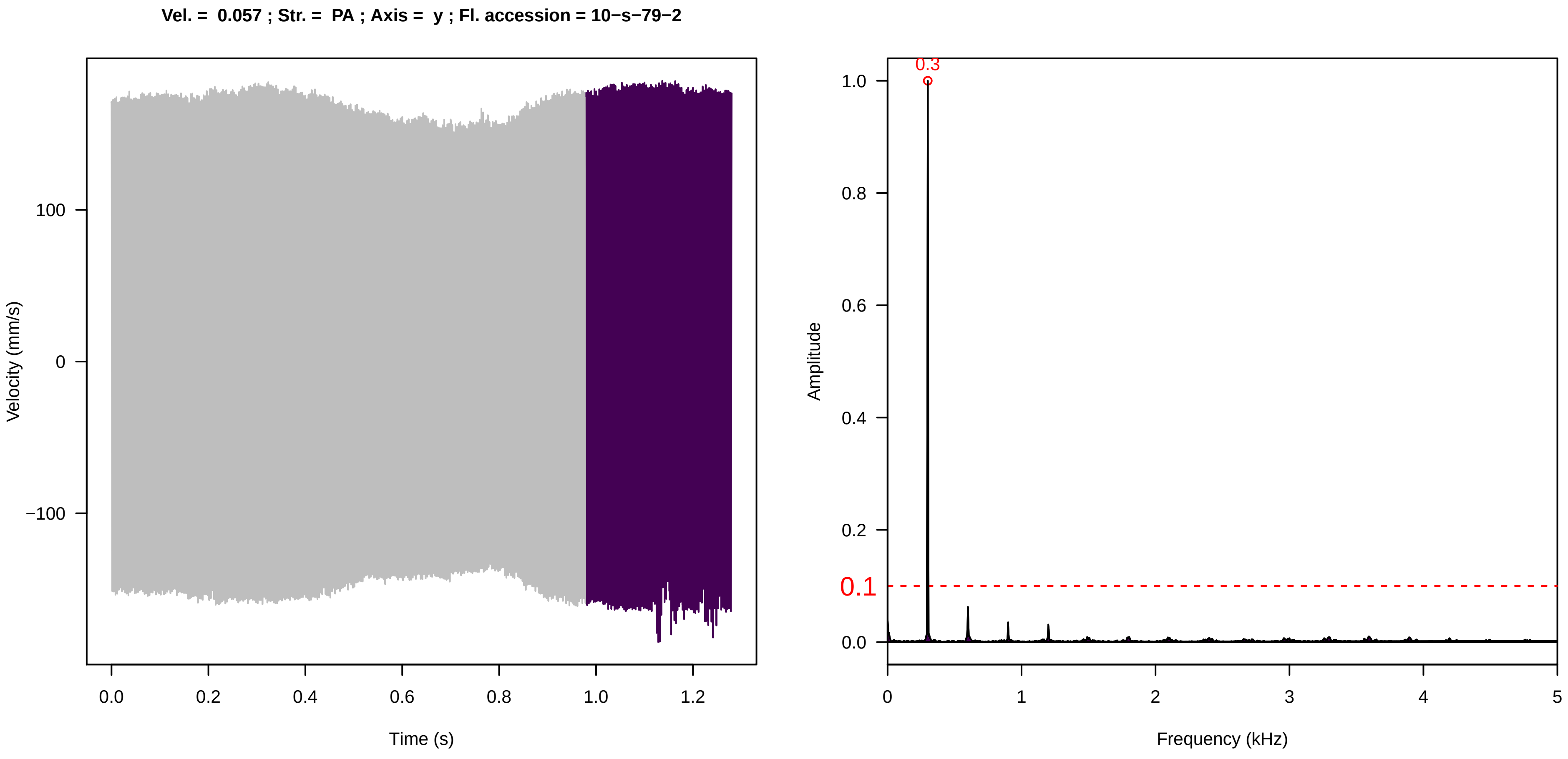
Vel. = 0.057 ; Str. = PA ; Axis = y ; Fl. accession = 10−s−79−2
Velocity (mm/s)
Time (s)
Amplitude
Frequency (kHz)
0.3
0.1

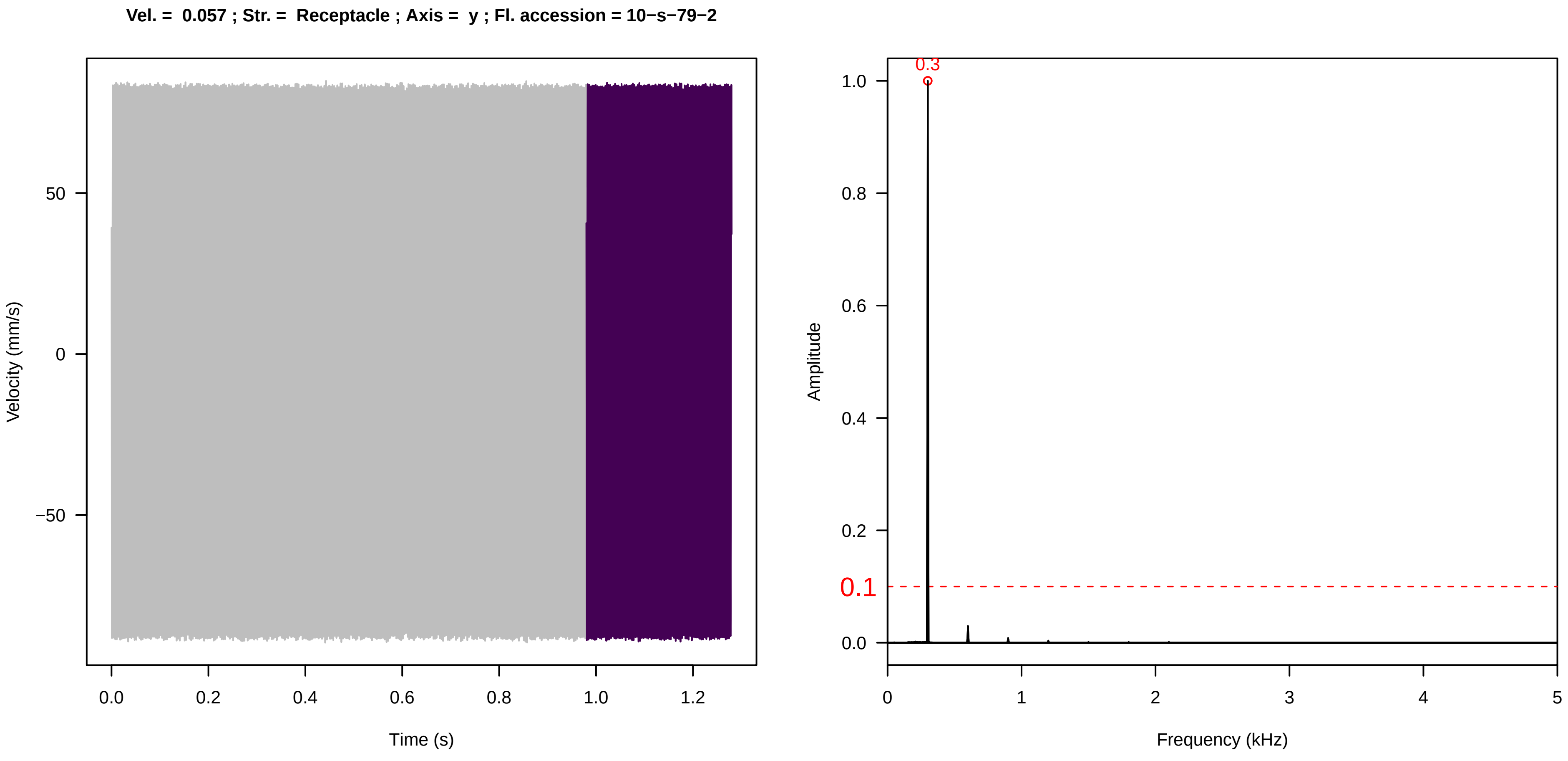
Vel. = 0.057 ; Str. = Receptacle ; Axis = y ; Fl. accession = 10−s−79−2
Velocity (mm/s)
Time (s)
Amplitude
Frequency (kHz)
0.3
0.1

**Vel. = 0.028 ; Str. = PA ; Axis = y ; Fl. accession = 10−s−79−2**

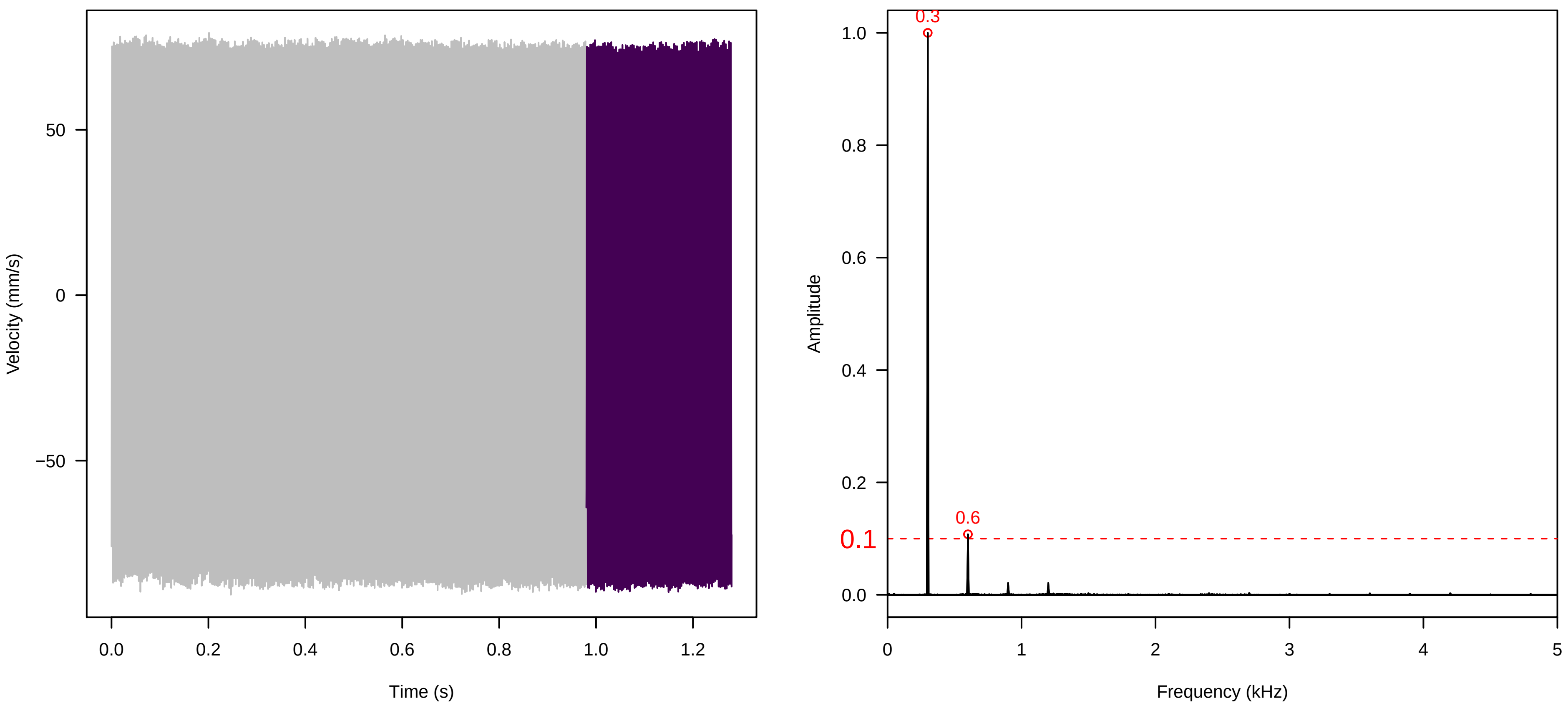


**Vel. = 0.028 ; Str. = Receptacle ; Axis = y ; Fl. accession = 10−s−79−2**

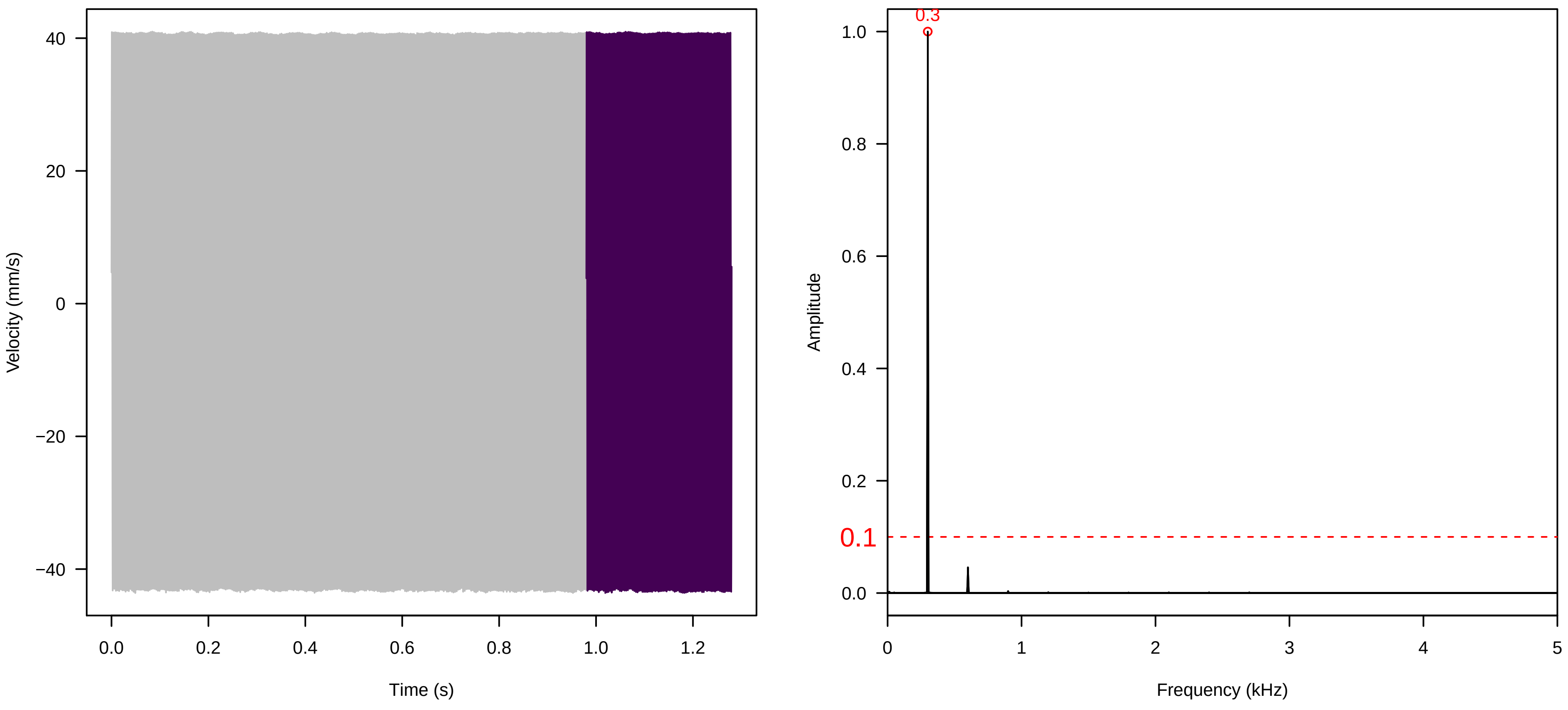


**Vel. = 0.028 ; Str. = FA ; Axis = y ; Fl. accession = 10−s−79−2**

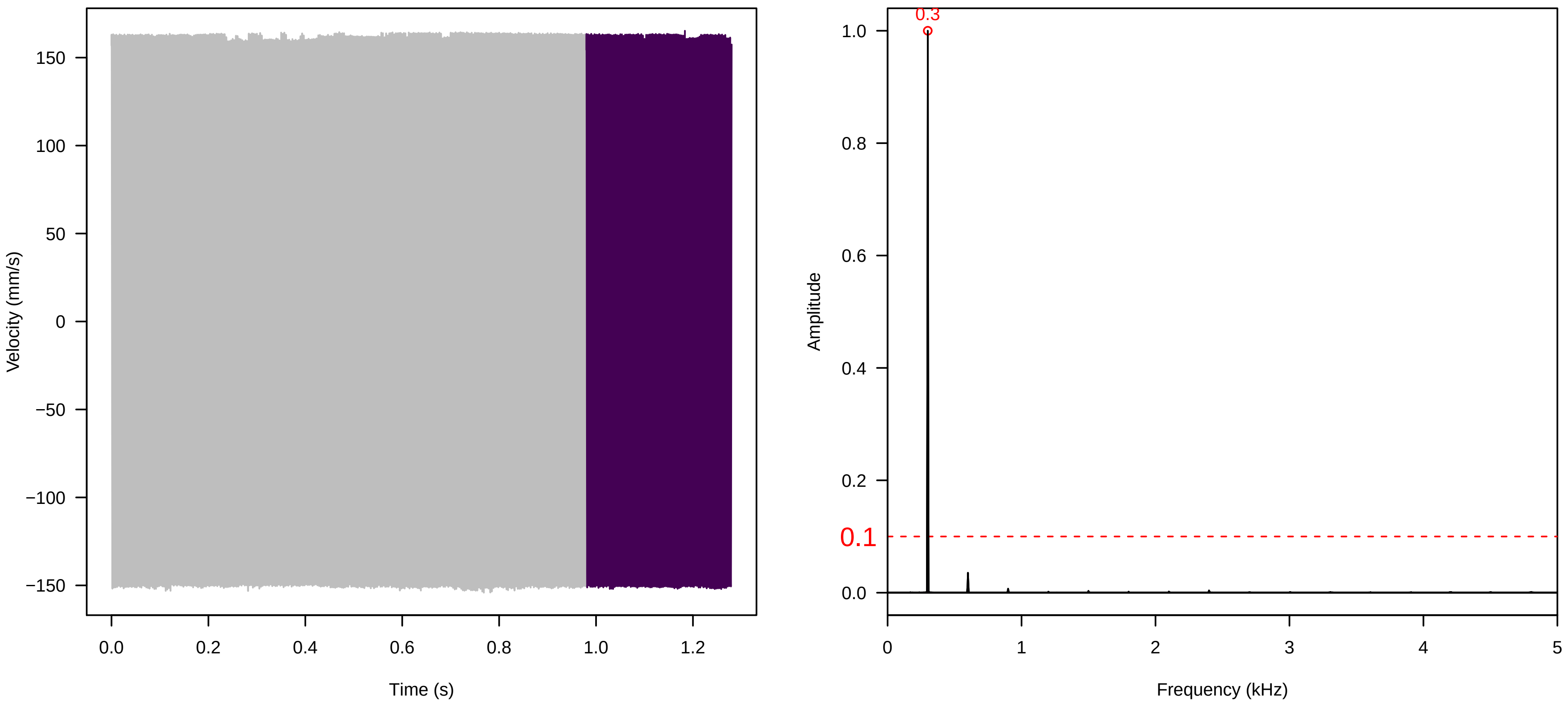

**Vel. = 0.028 ; Str. = Receptacle ; Axis = y ; Fl. accession = 10−s−79−2**

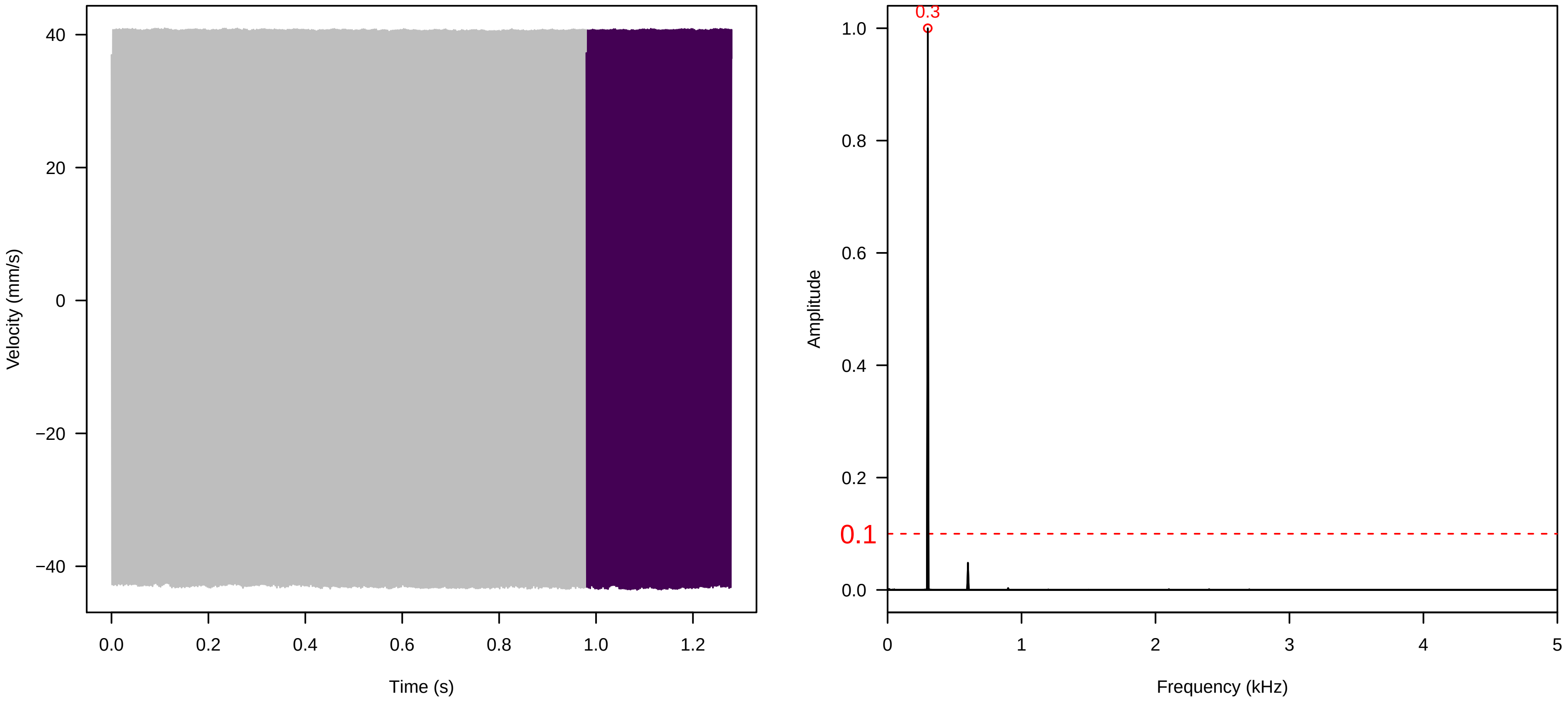


**Vel. = 0.028 ; Str. = Corolla ; Axis = y ; Fl. accession = 10−s−79−2**

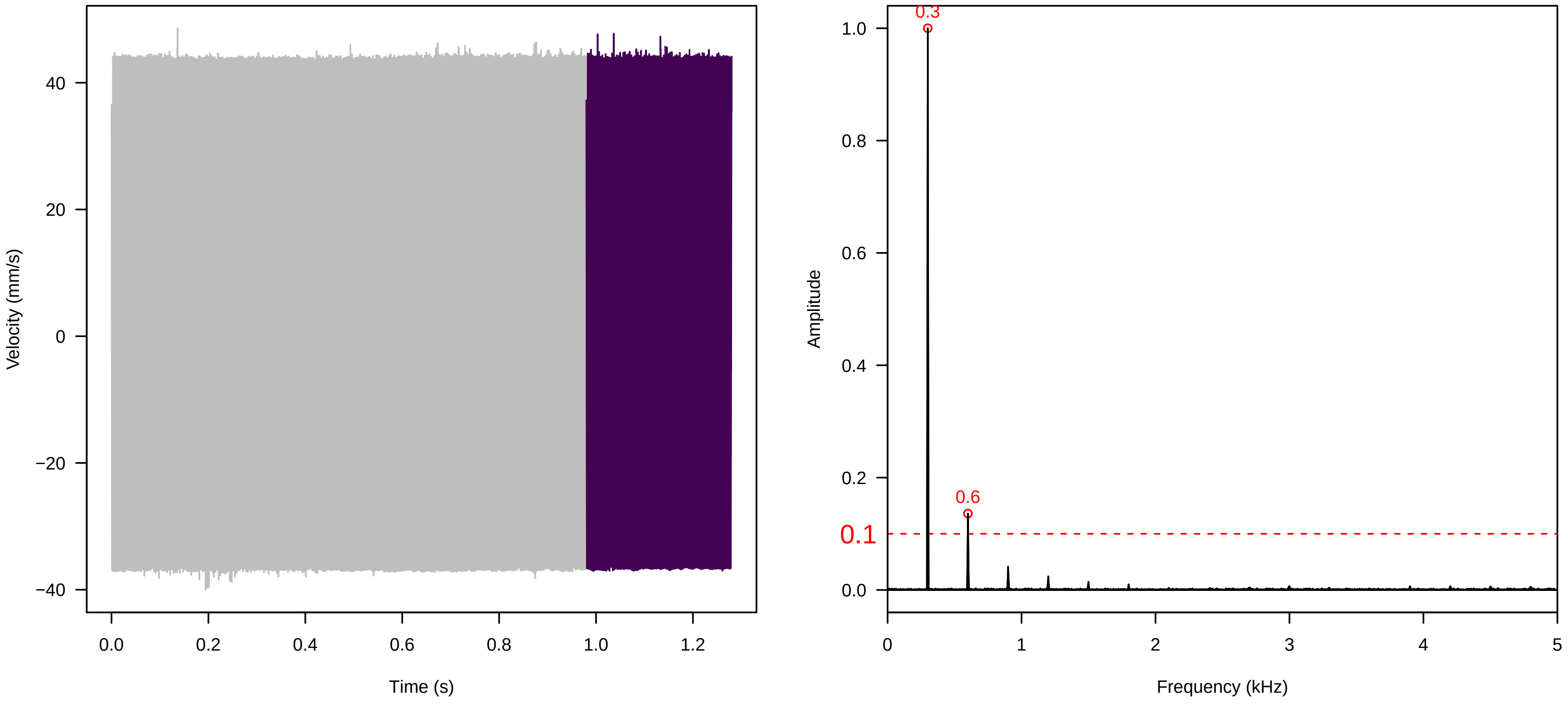


**Vel. = 0.028 ; Str. = Receptacle ; Axis = y ; Fl. accession = 10−s−79−2**

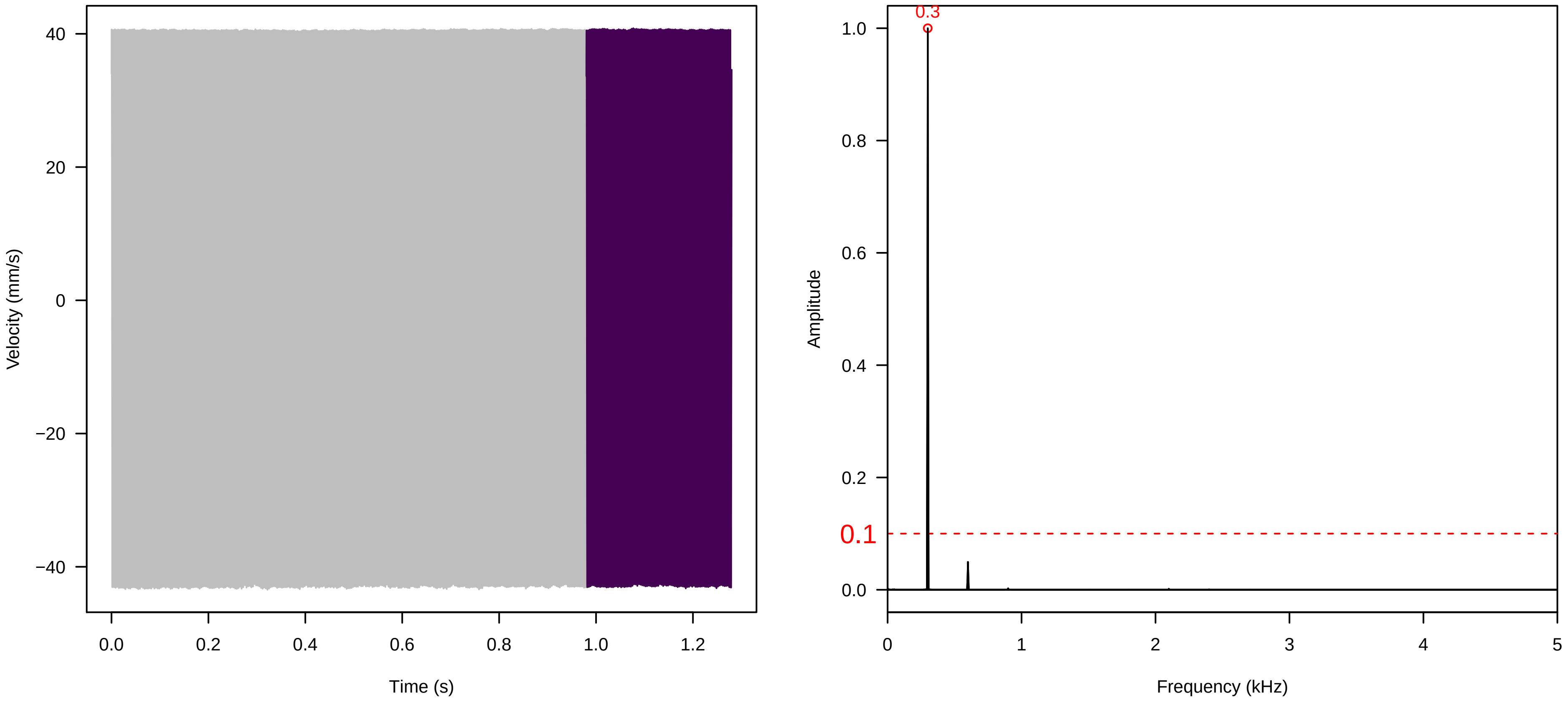

**Vel. = 0.014 ; Str. = Corolla ; Axis = y ; Fl. accession = 10−s−79−2**

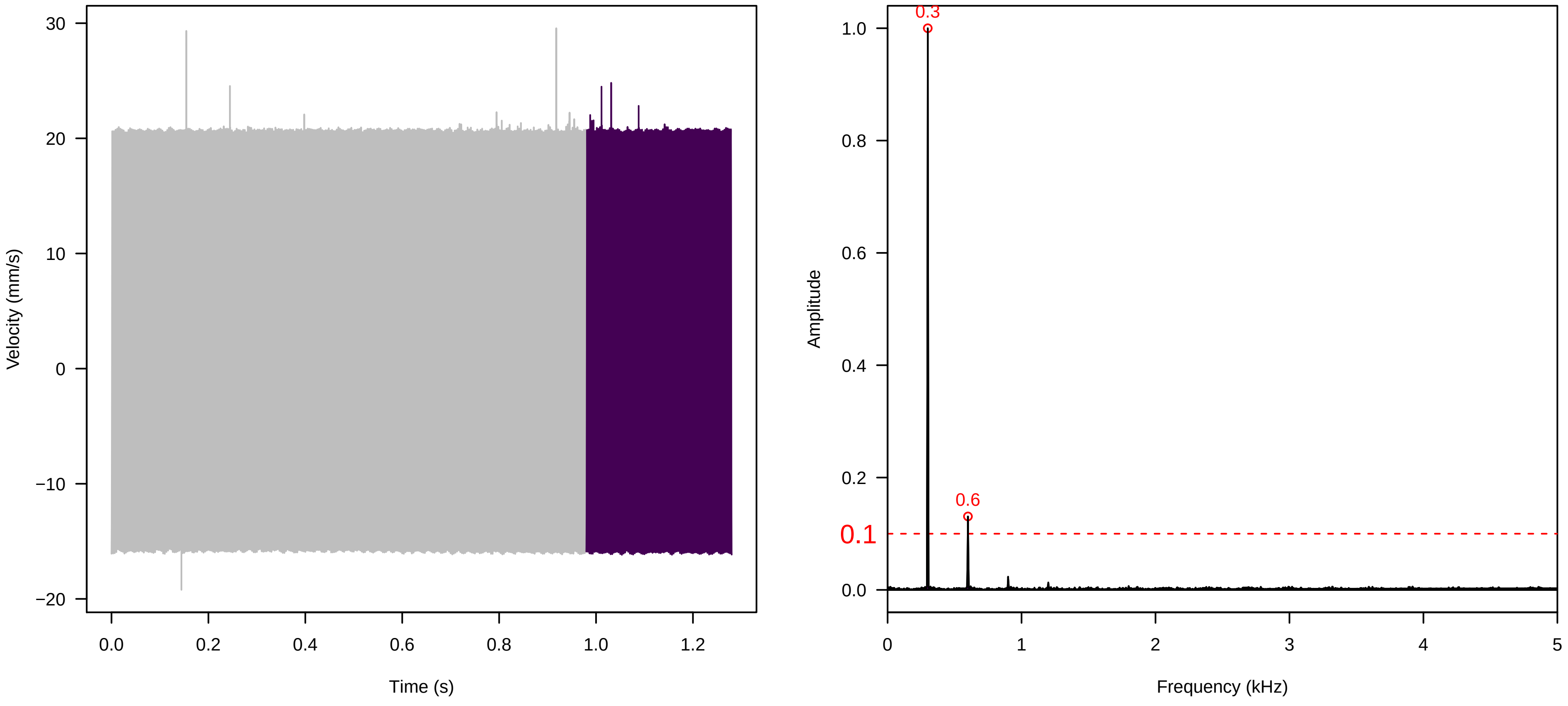


**Vel. = 0.014 ; Str. = Receptacle ; Axis = y ; Fl. accession = 10−s−79−2**

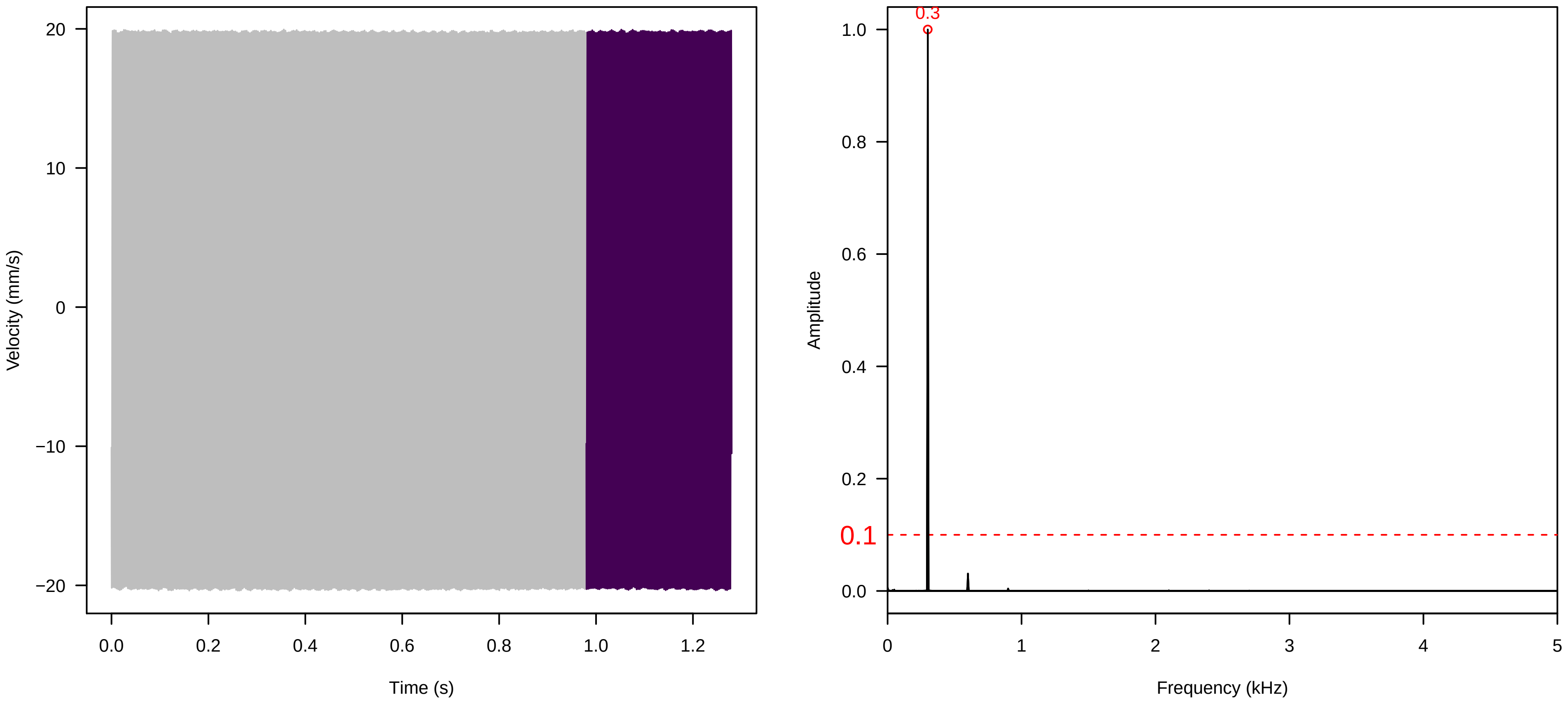


**Vel. = 0.014 ; Str. = FA ; Axis = y ; Fl. accession = 10−s−79−2**

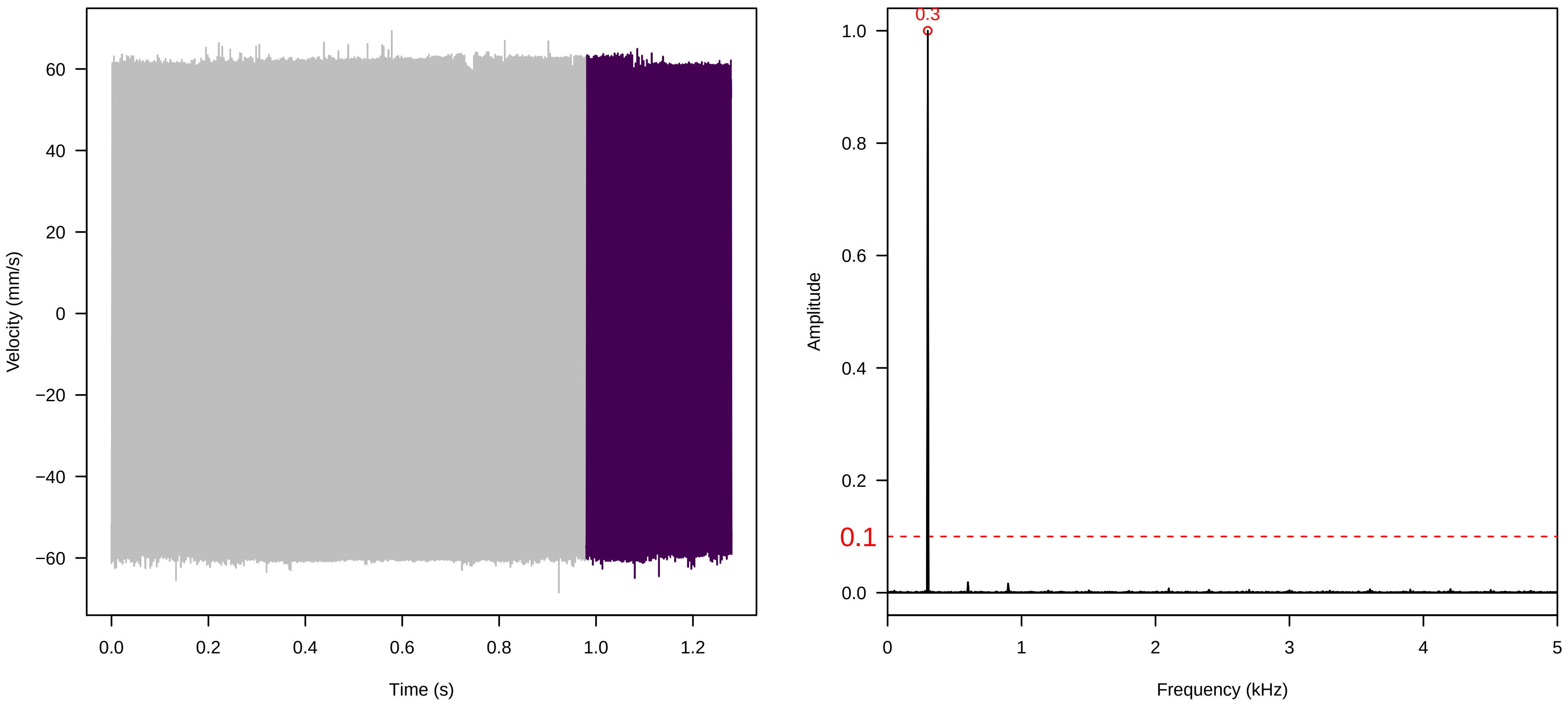

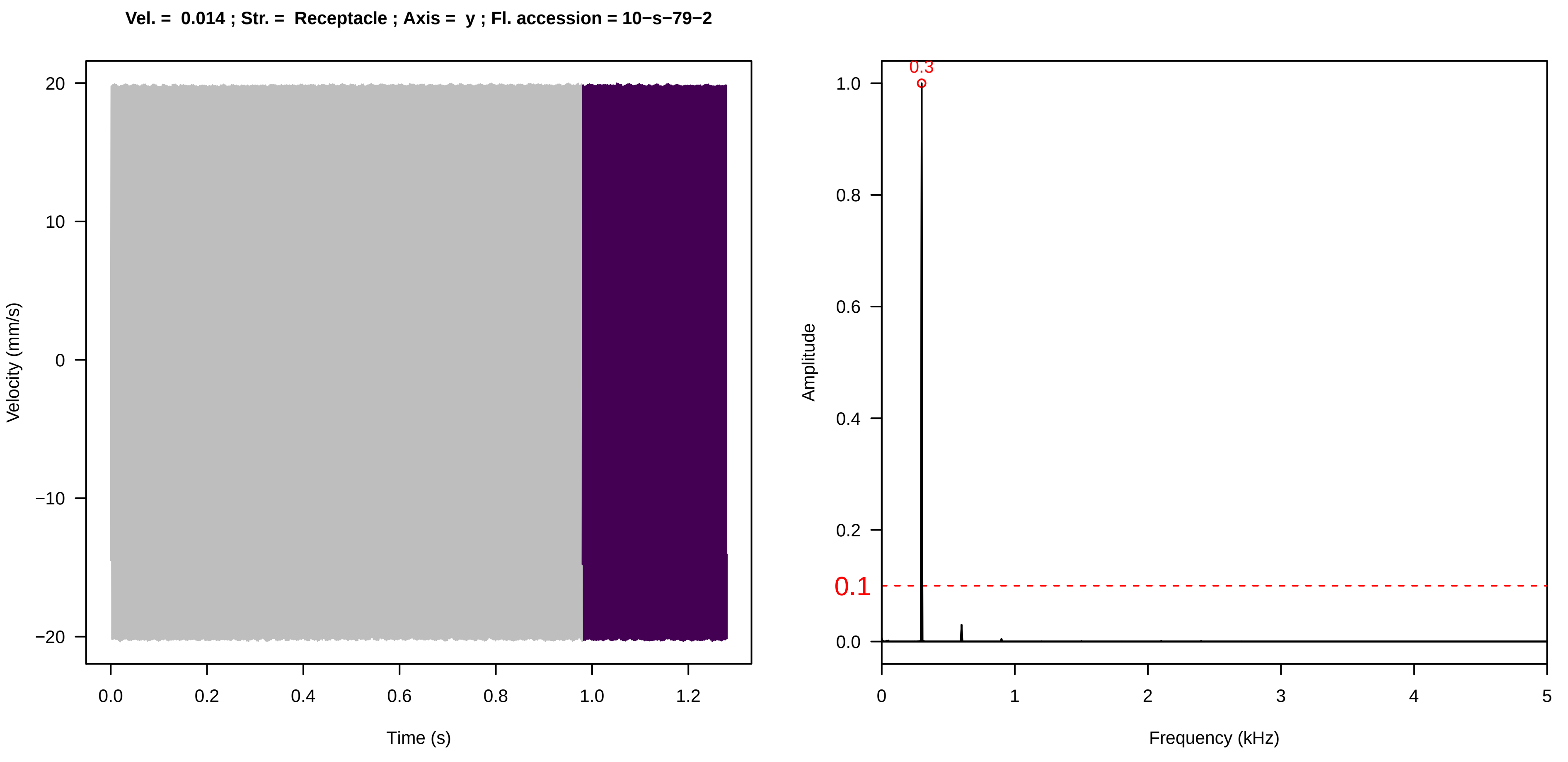
Vel. = 0.014 ; Str. = Receptacle ; Axis = y ; Fl. accession = 10−s−79−2
Velocity (mm/s)
Time (s)
Amplitude
Frequency (kHz)
0.3
0.1

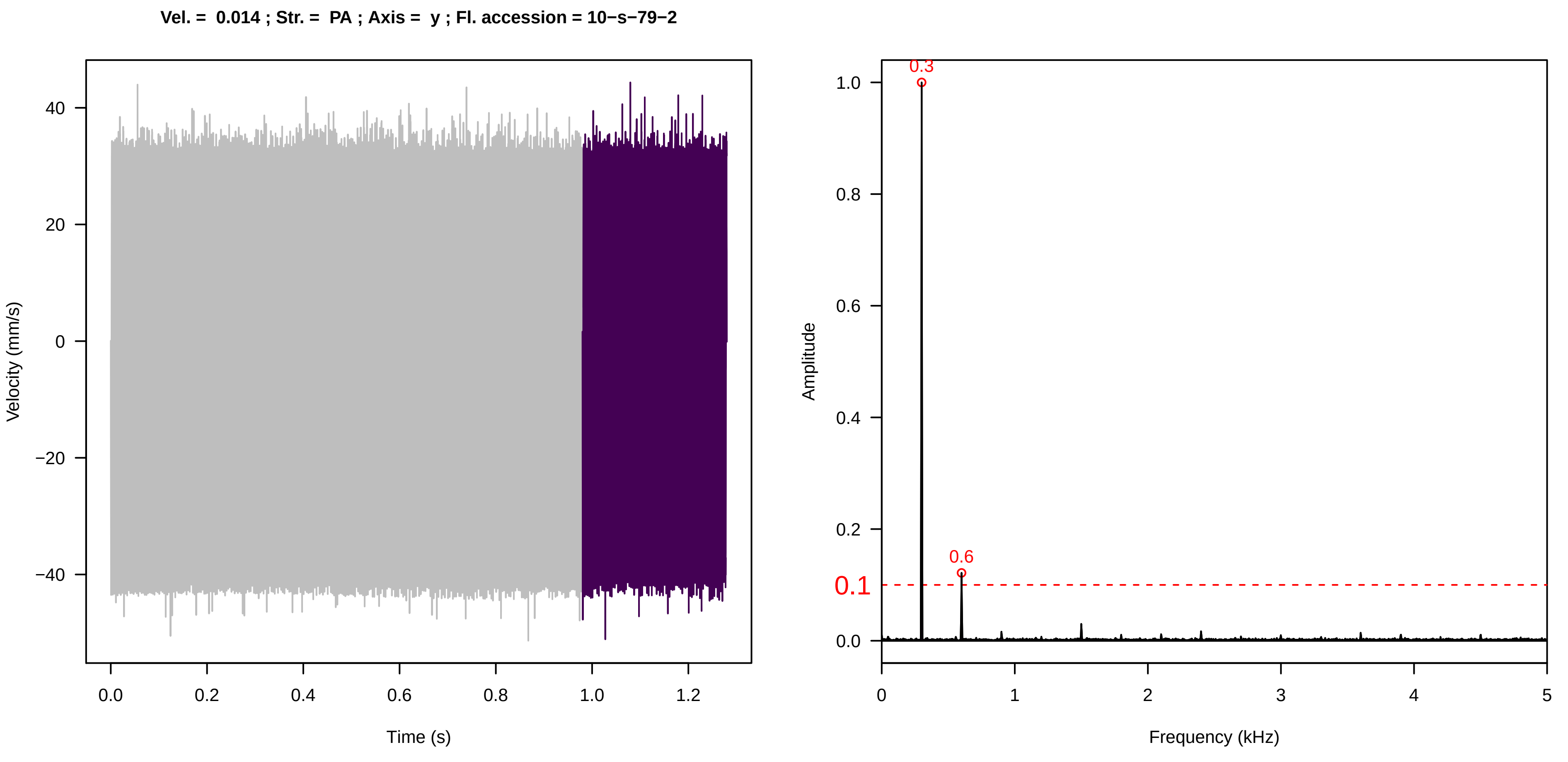
Vel. = 0.014 ; Str. = PA ; Axis = y ; Fl. accession = 10−s−79−2
Velocity (mm/s)
Time (s)
Amplitude
Frequency (kHz)
0.3
0.6
0.1

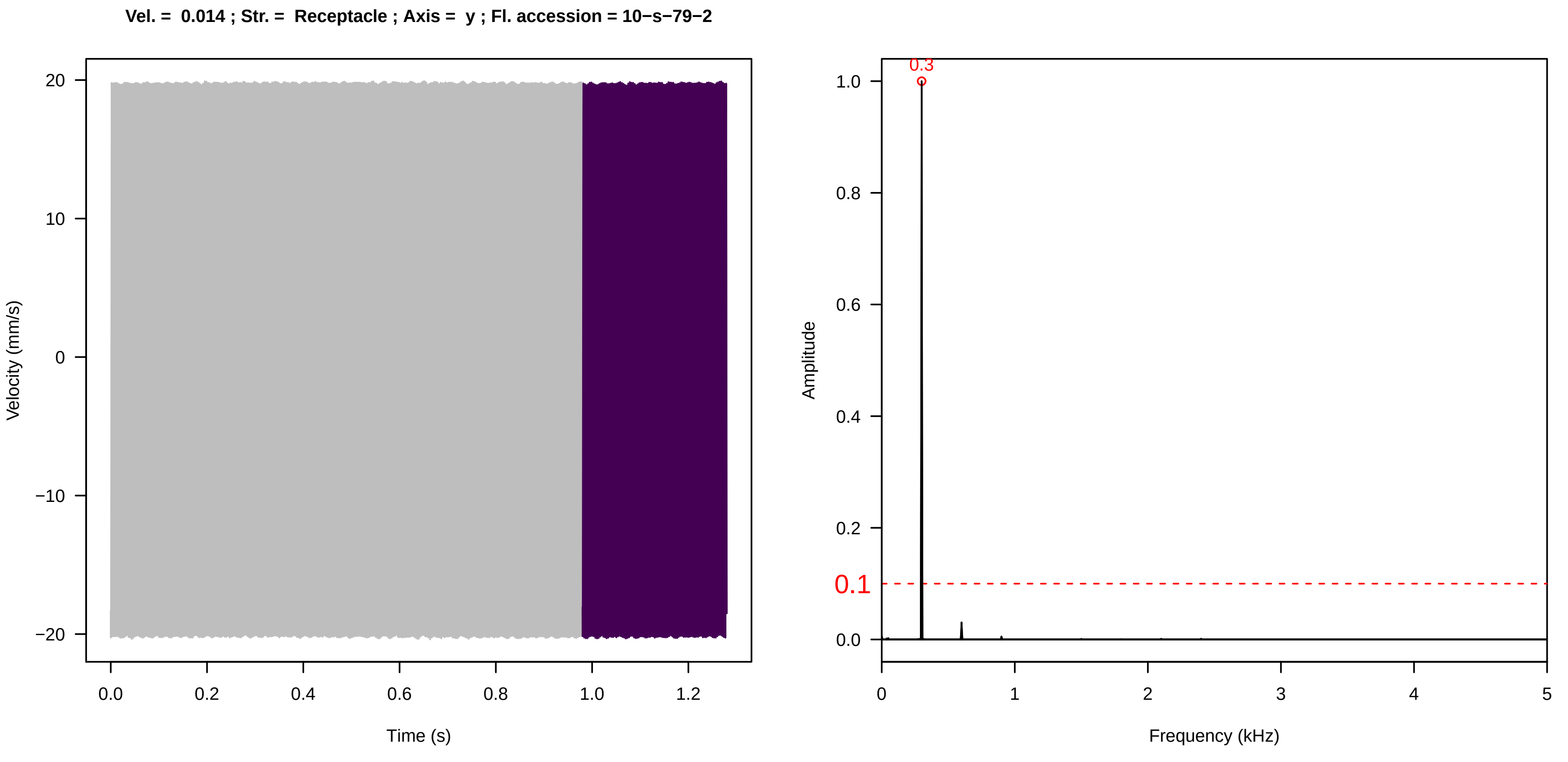
Vel. = 0.014 ; Str. = Receptacle ; Axis = y ; Fl. accession = 10−s−79−2
Velocity (mm/s)
Time (s)
Amplitude
Frequency (kHz)
0.3
0.1

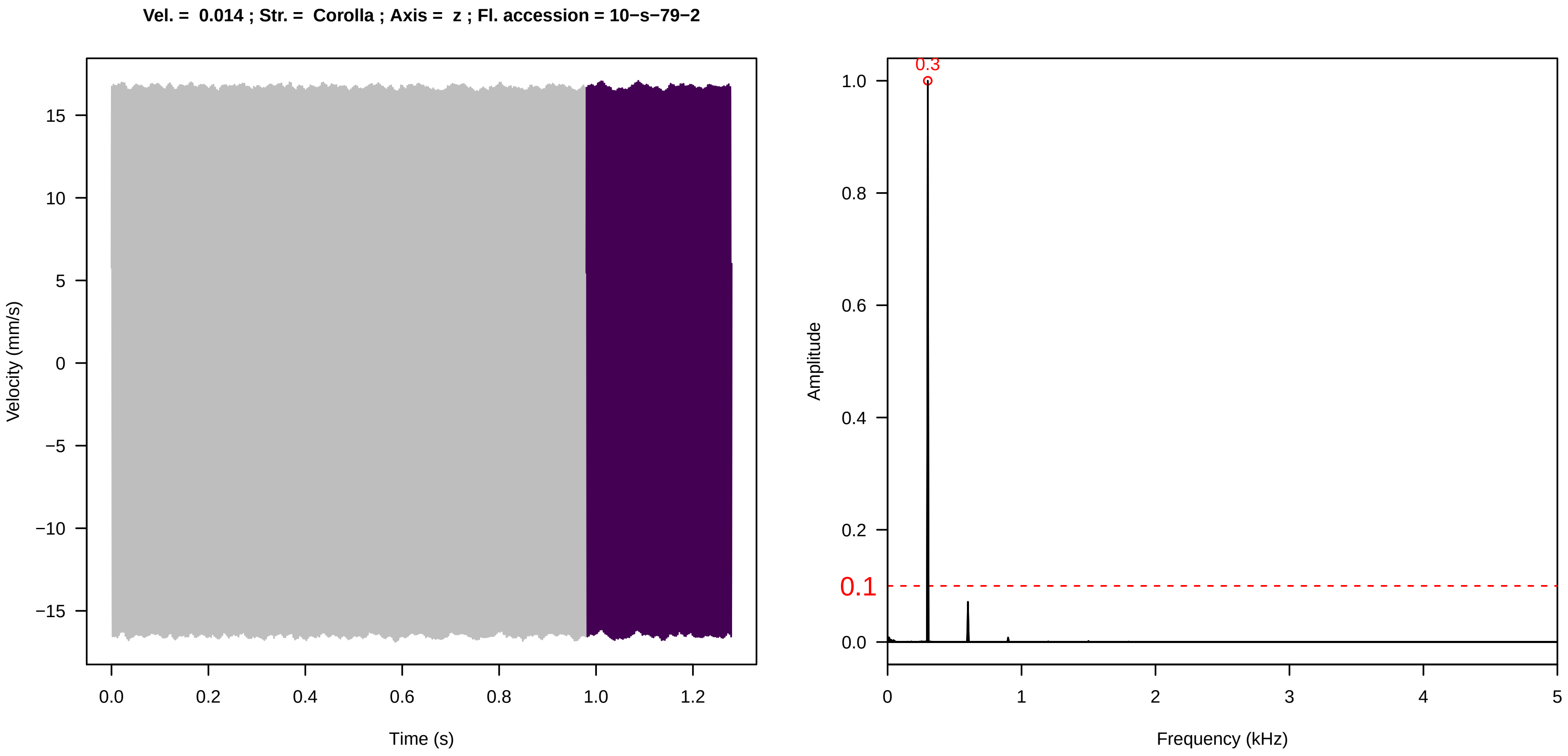
Vel. = 0.014 ; Str. = Corolla ; Axis = z ; Fl. accession = 10−s−79−2
Velocity (mm/s)
Time (s)
0.3
0.1
Amplitude
Frequency (kHz)

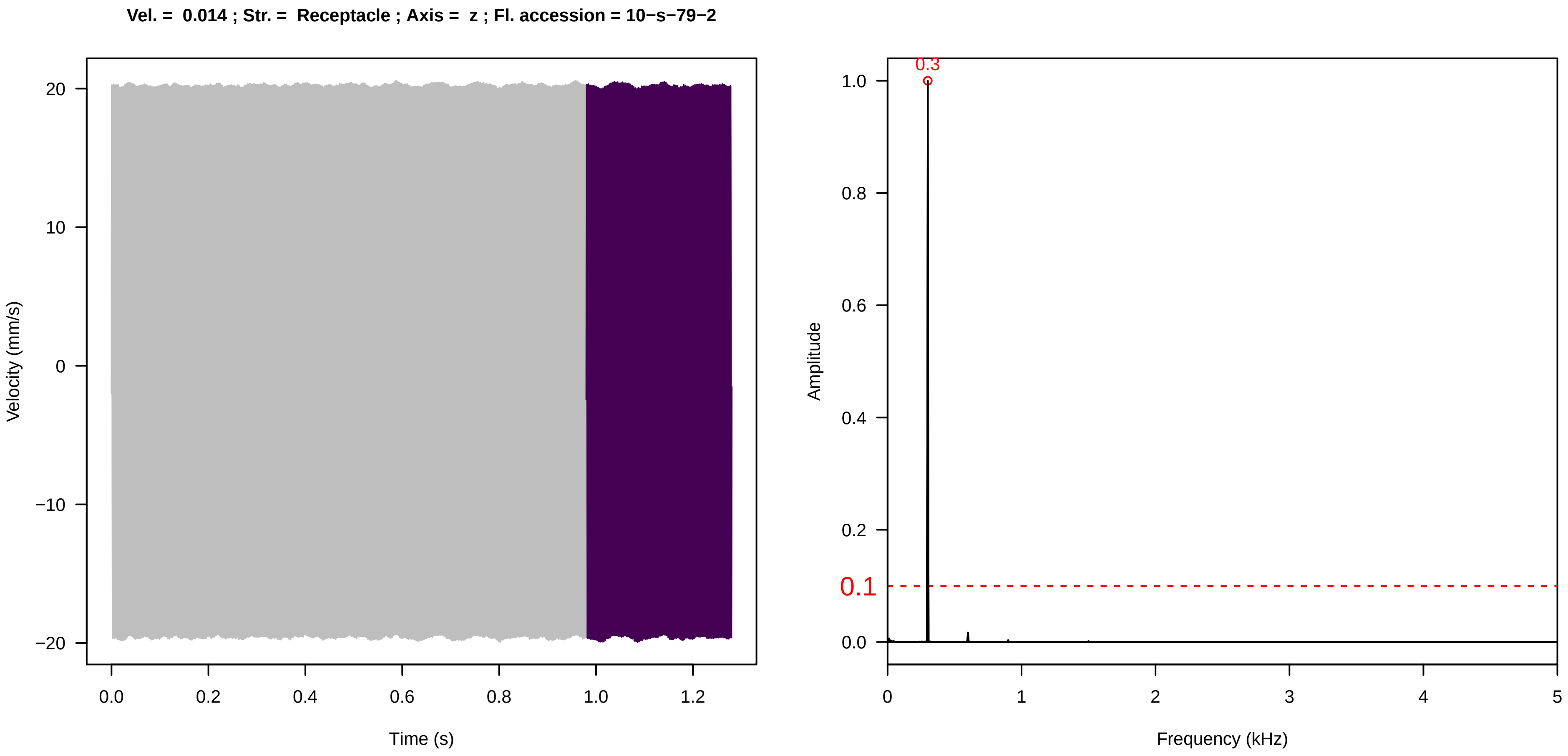
Vel. = 0.014 ; Str. = Receptacle ; Axis = z ; Fl. accession = 10−s−79−2
Velocity (mm/s)
Time (s)
0.3
0.1
Amplitude
Frequency (kHz)

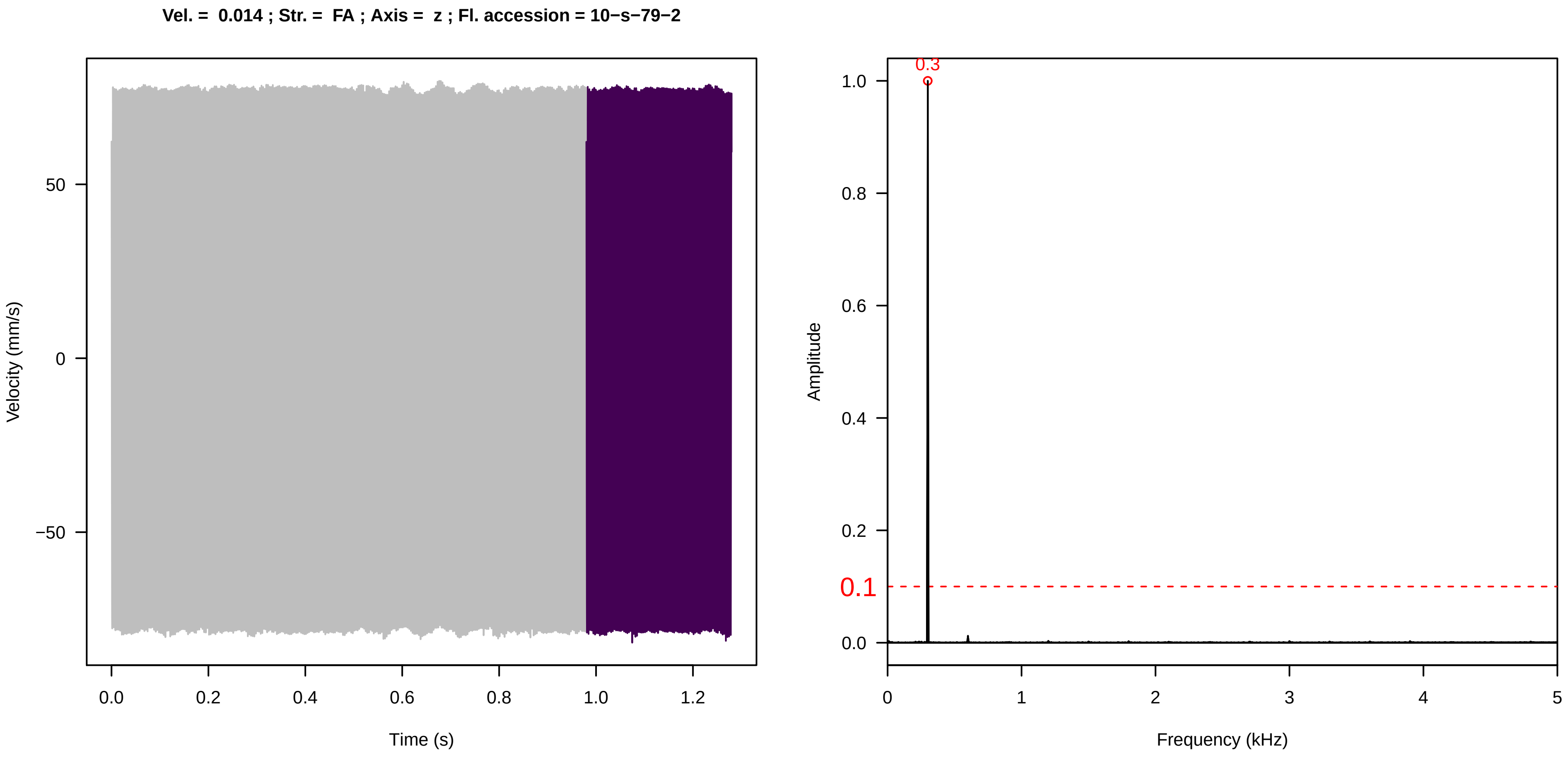
Vel. = 0.014 ; Str. = FA ; Axis = z ; Fl. accession = 10−s−79−2
Velocity (mm/s)
Time (s)
0.3
0.1
Amplitude
Frequency (kHz)

**Vel. = 0.014 ; Str. = Receptacle ; Axis = z ; Fl. accession = 10−s−79−2**

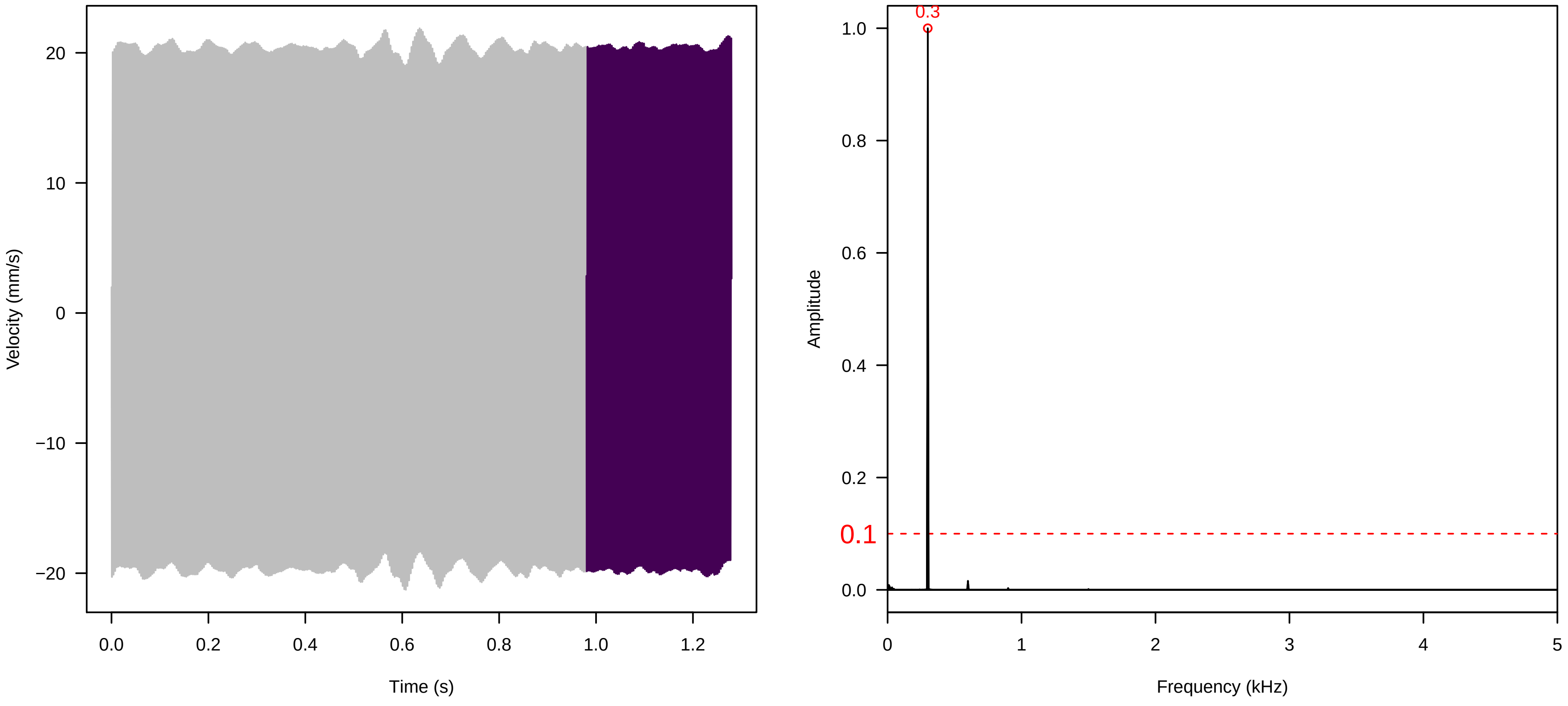


**Vel. = 0.014 ; Str. = PA ; Axis = z ; Fl. accession = 10−s−79−2**

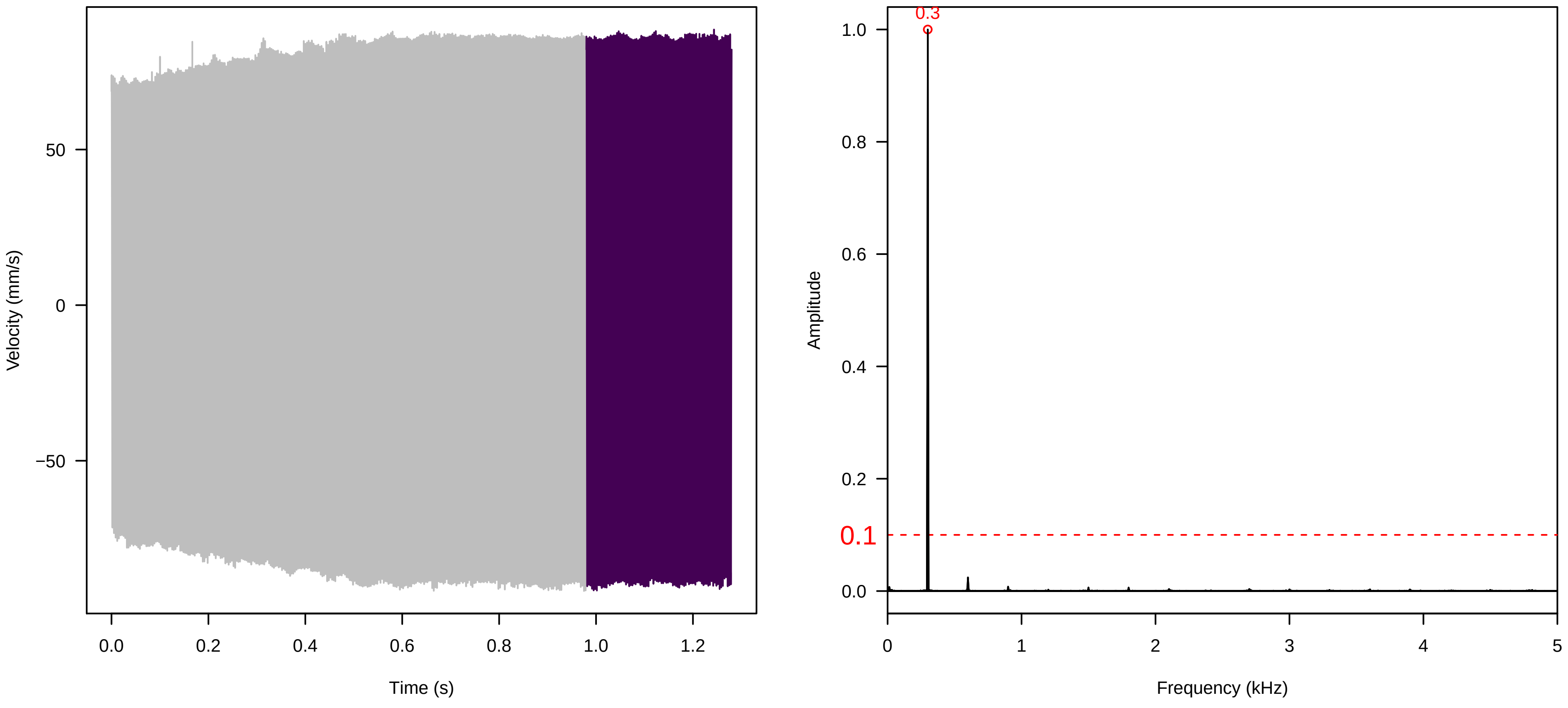


**Vel. = 0.014 ; Str. = Receptacle ; Axis = z ; Fl. accession = 10−s−79−2**

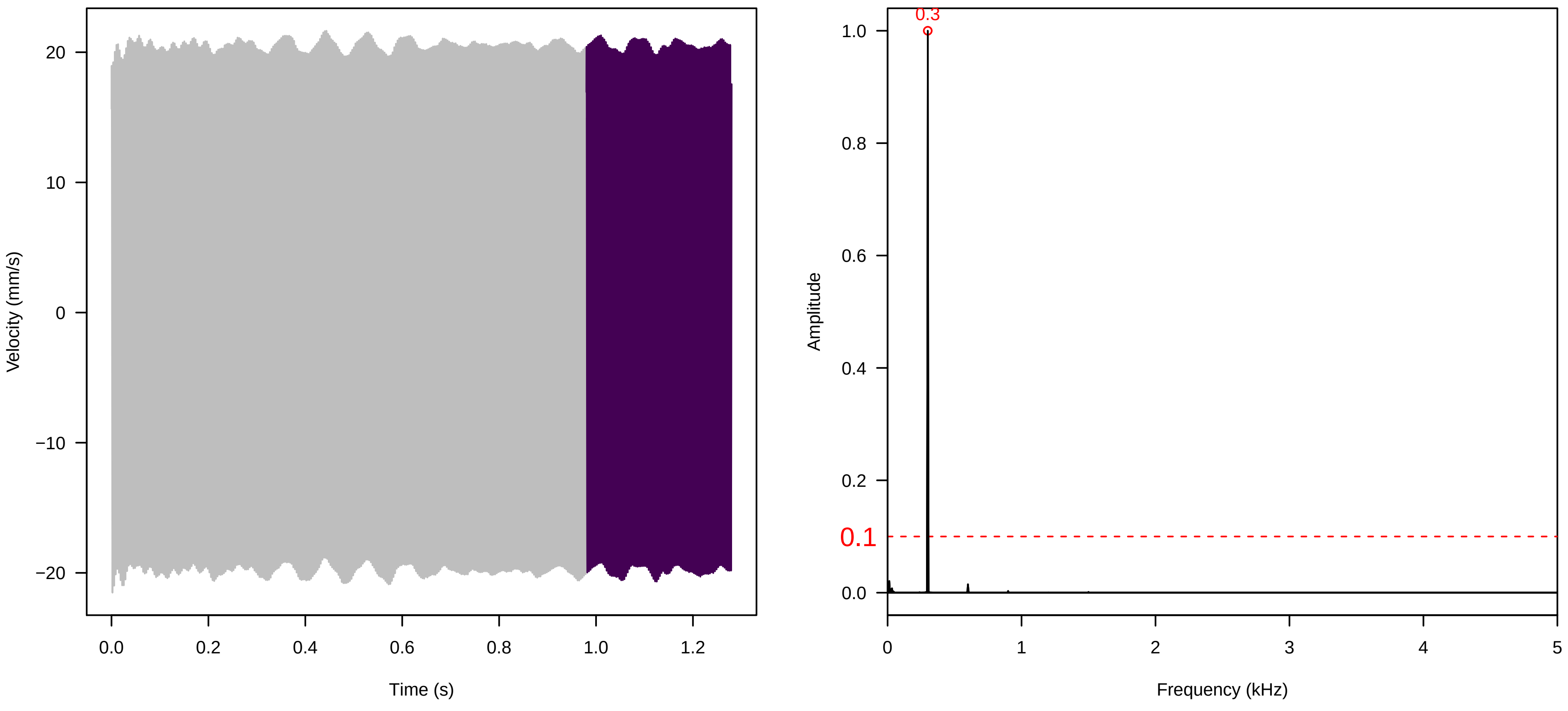

**Vel. = 0.028 ; Str. = PA ; Axis = z ; Fl. accession = 10−s−79−2**

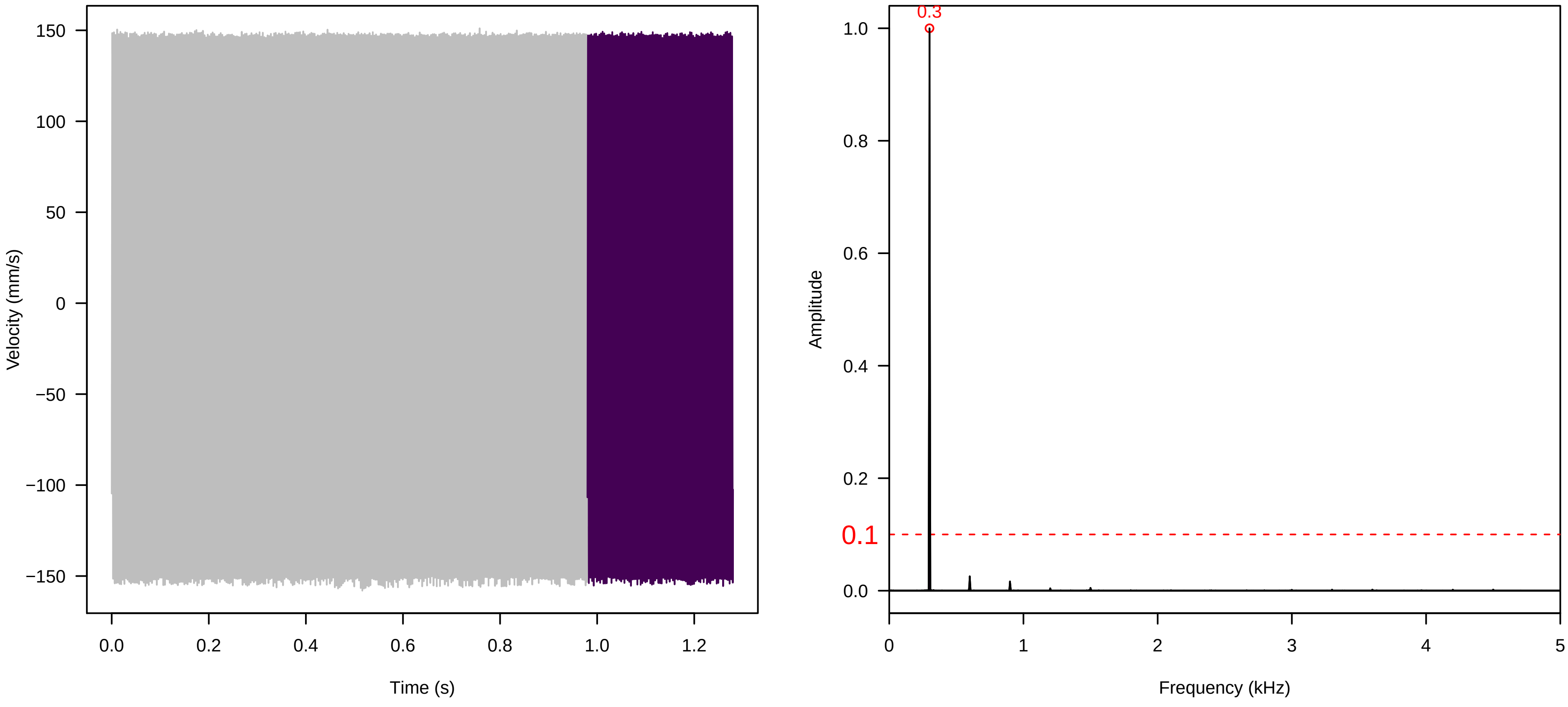


**Vel. = 0.028 ; Str. = Receptacle ; Axis = z ; Fl. accession = 10−s−79−2**

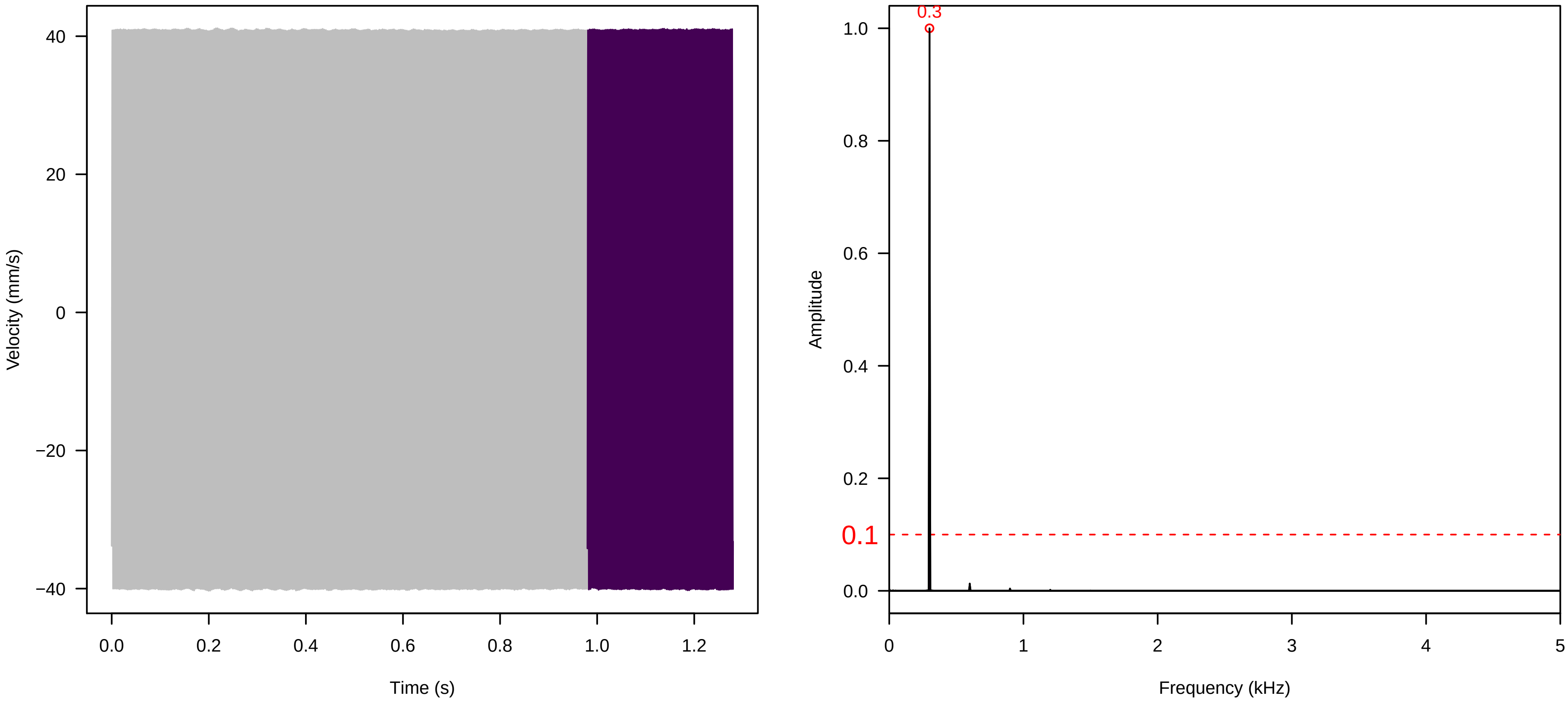


**Vel. = 0.028 ; Str. = FA ; Axis = z ; Fl. accession = 10−s−79−2**

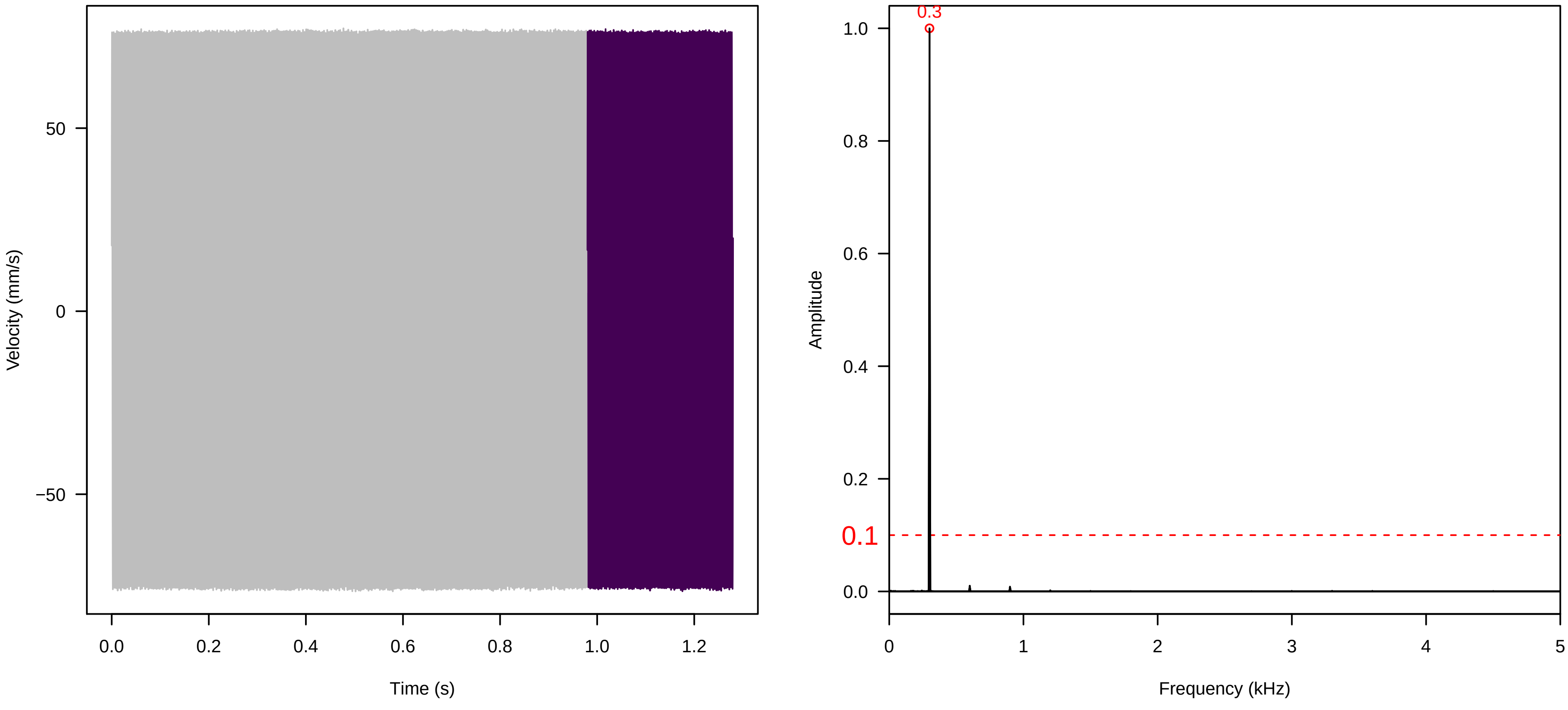

**Vel. = 0.028 ; Str. = Receptacle ; Axis = z ; Fl. accession = 10−s−79−2**

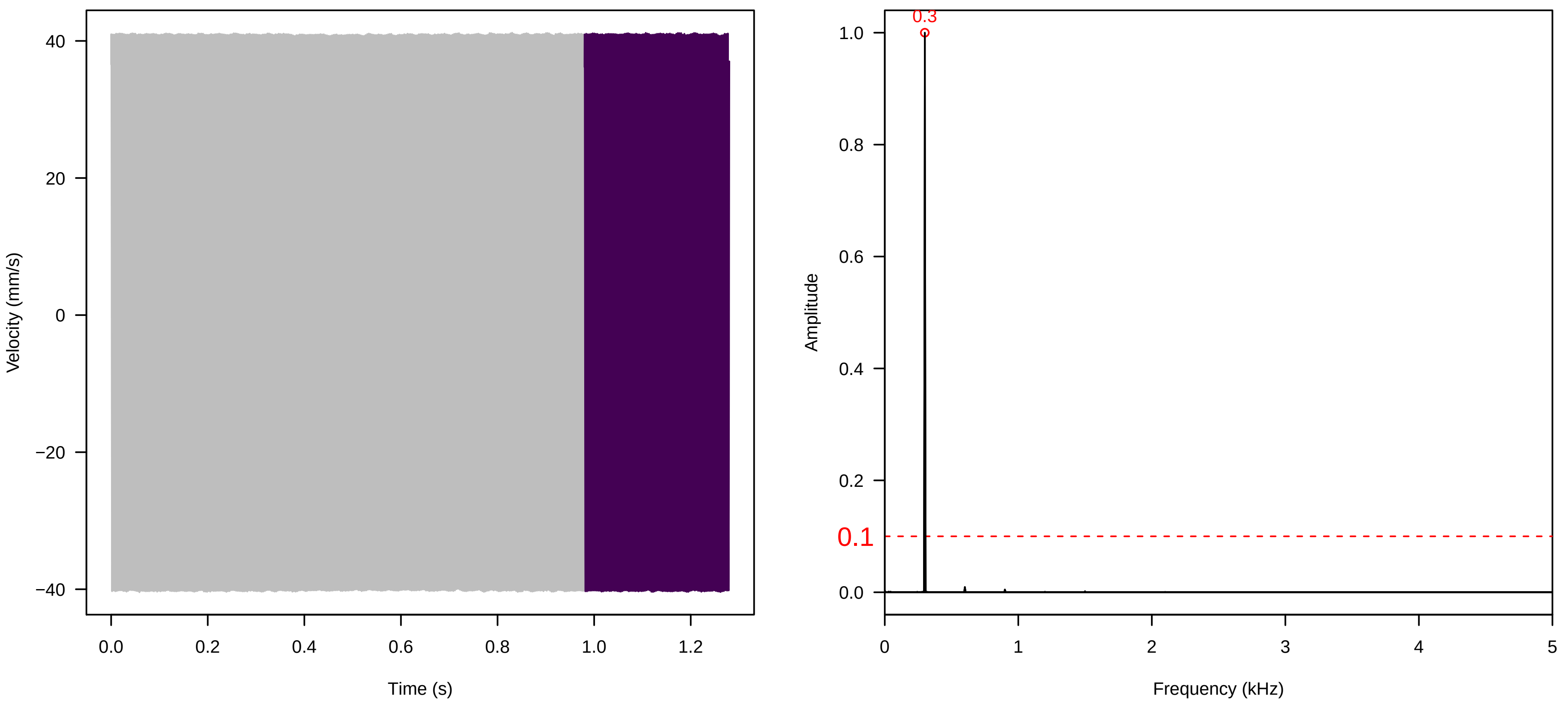


**Vel. = 0.028 ; Str. = Corolla ; Axis = z ; Fl. accession = 10−s−79−2**

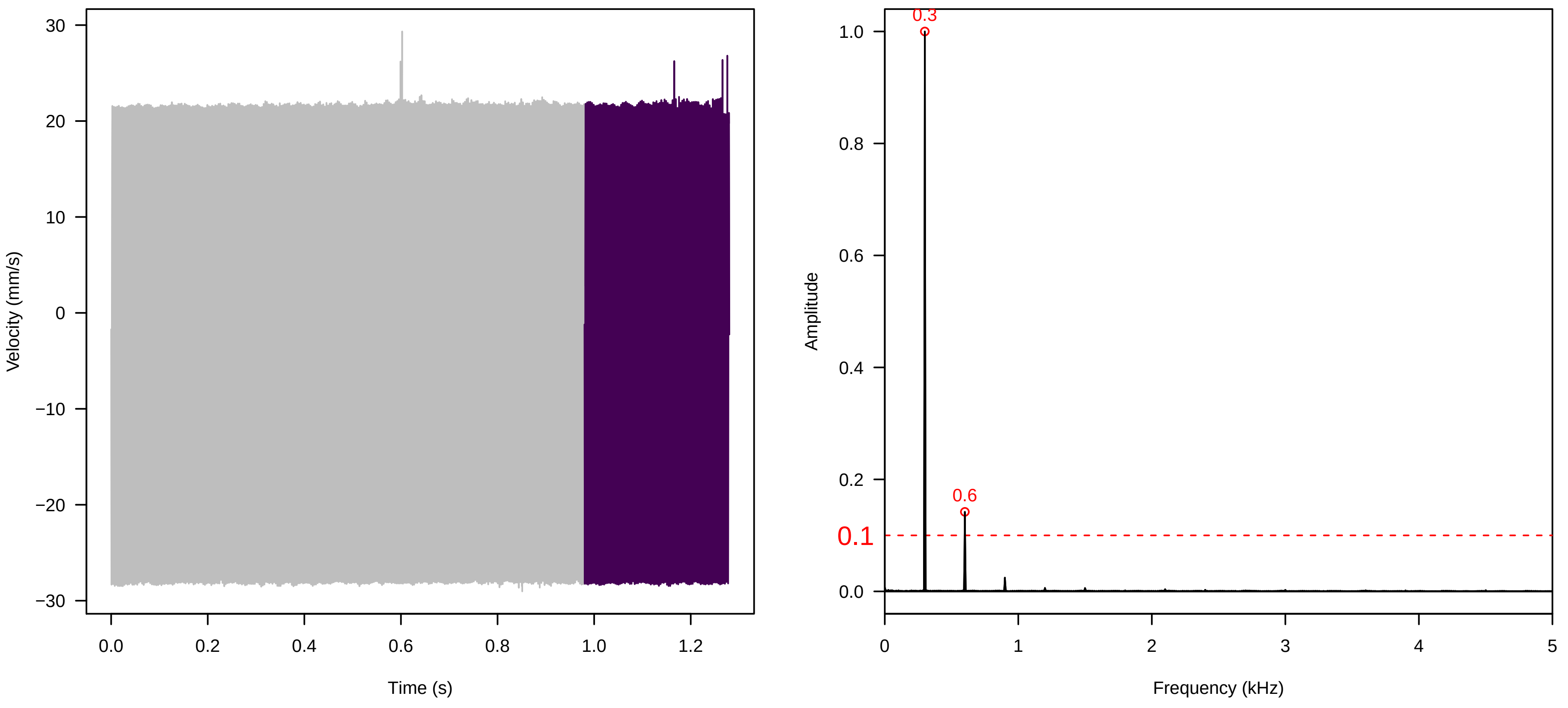


**Vel. = 0.028 ; Str. = Receptacle ; Axis = z ; Fl. accession = 10−s−79−2**

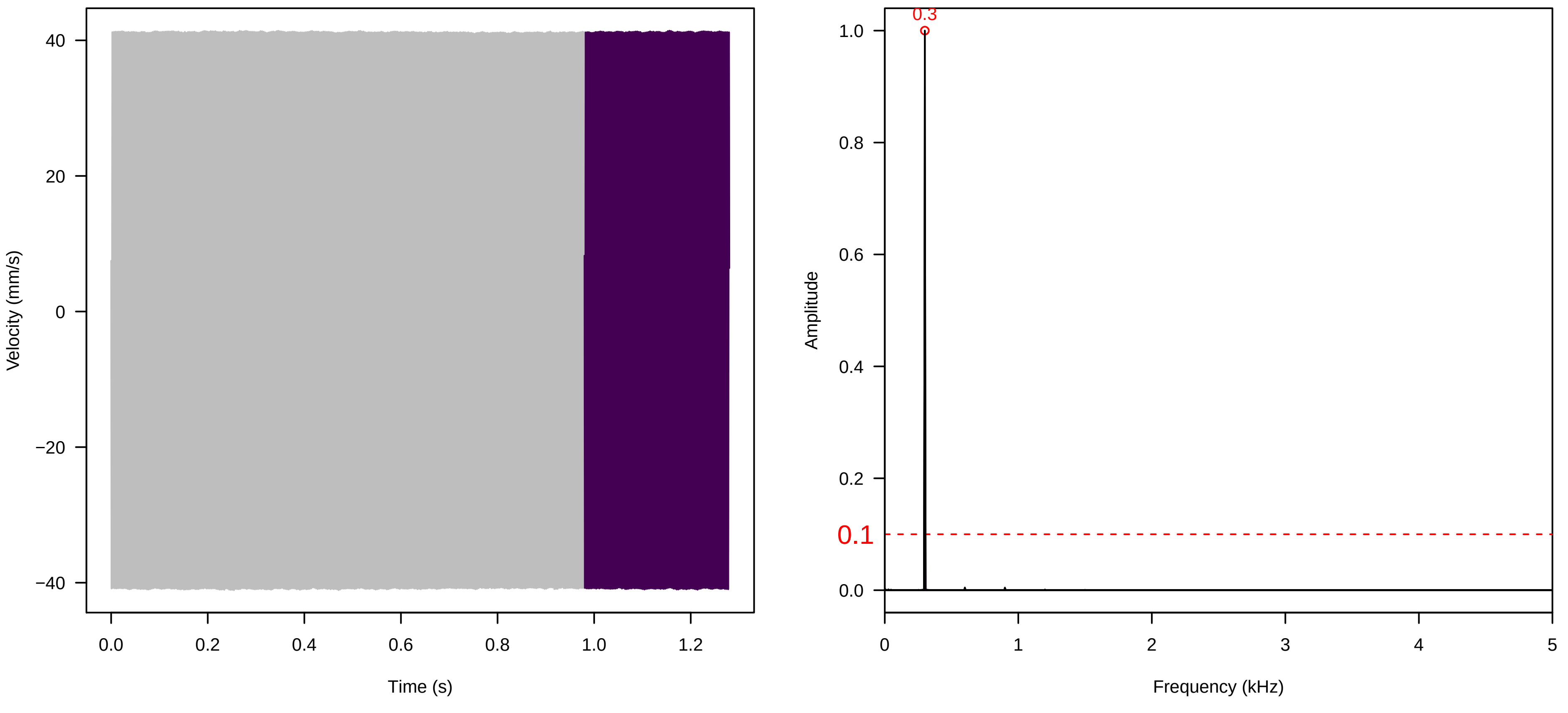

**Vel. = 0.057 ; Str. = Corolla ; Axis = z ; Fl. accession = 10−s−79−2**

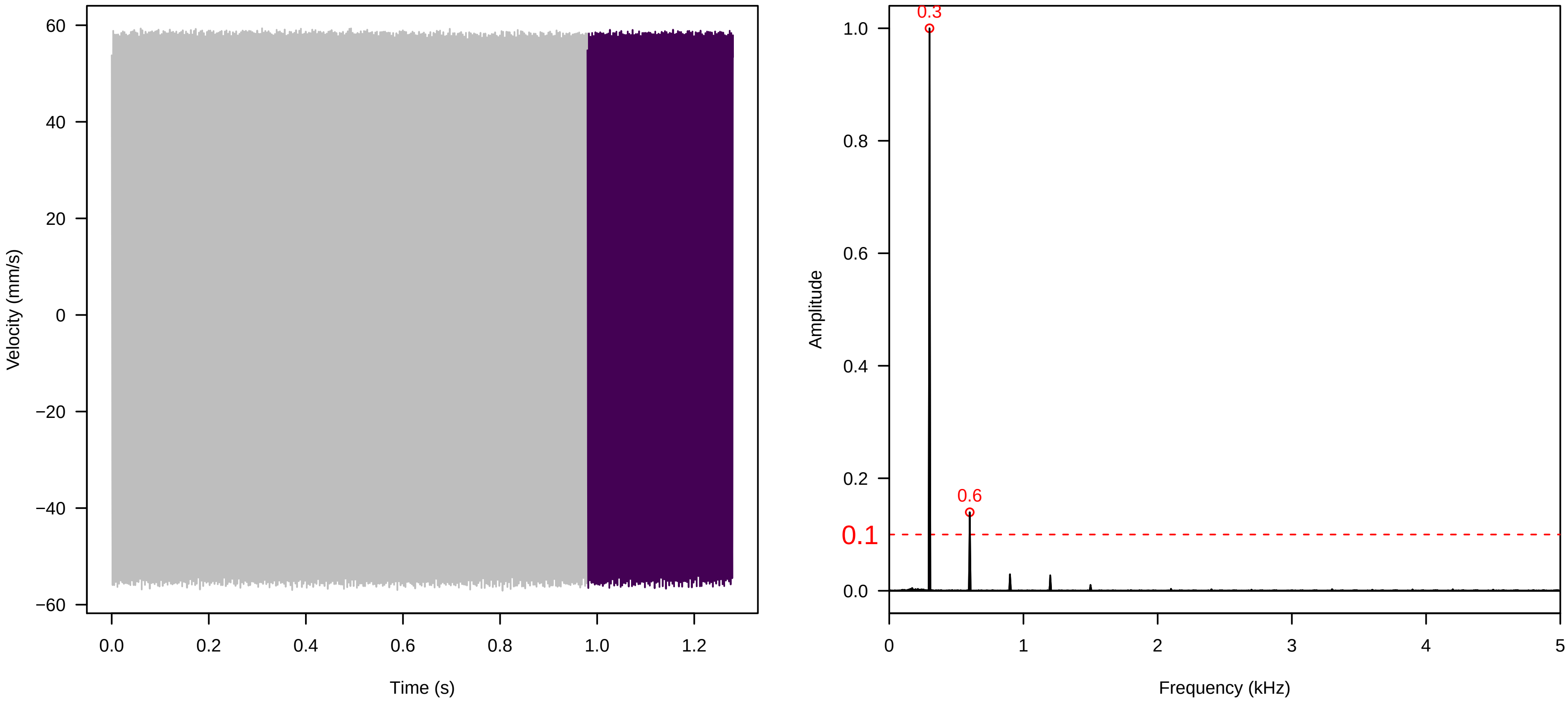


**Vel. = 0.057 ; Str. = Receptacle ; Axis = z ; Fl. accession = 10−s−79−2**

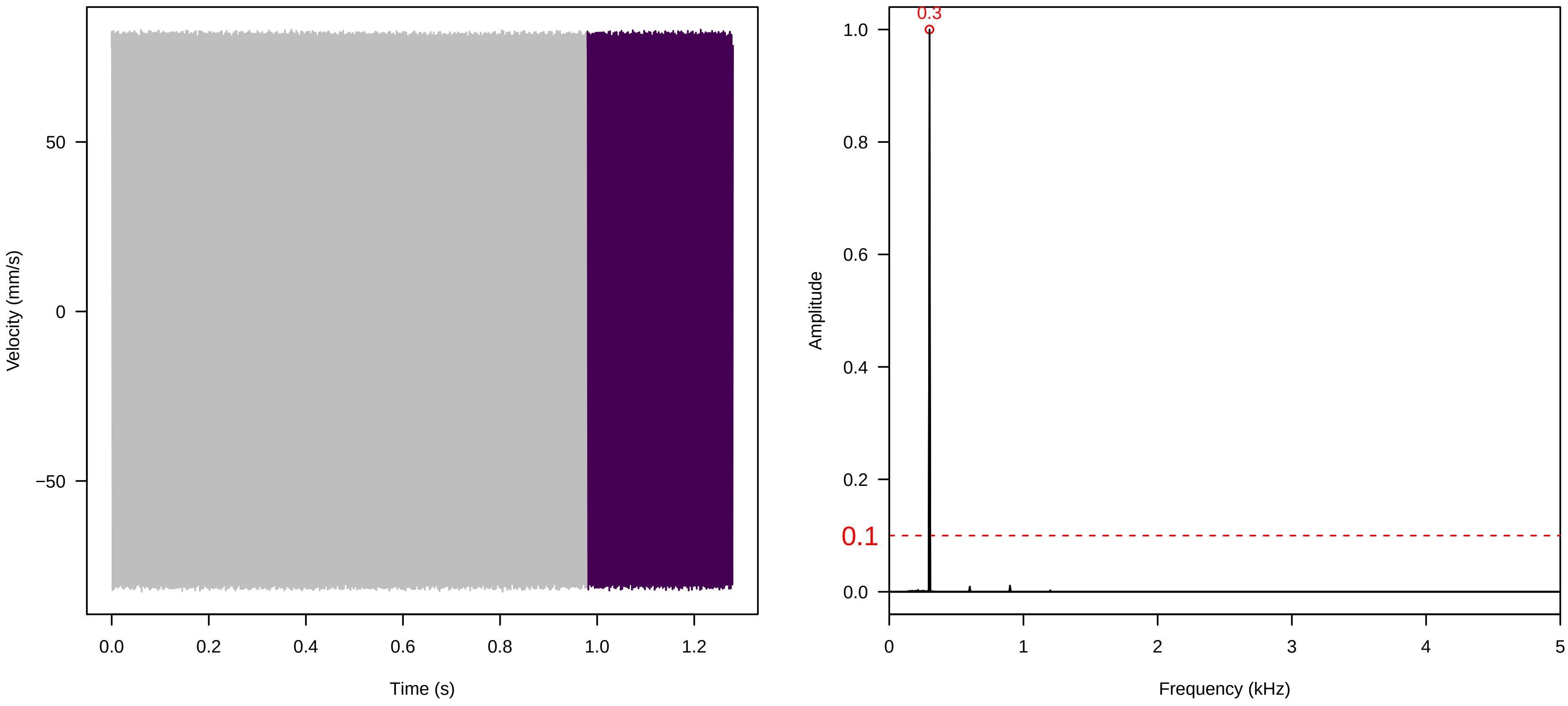


**Vel. = 0.057 ; Str. = FA ; Axis = z ; Fl. accession = 10−s−79−2**

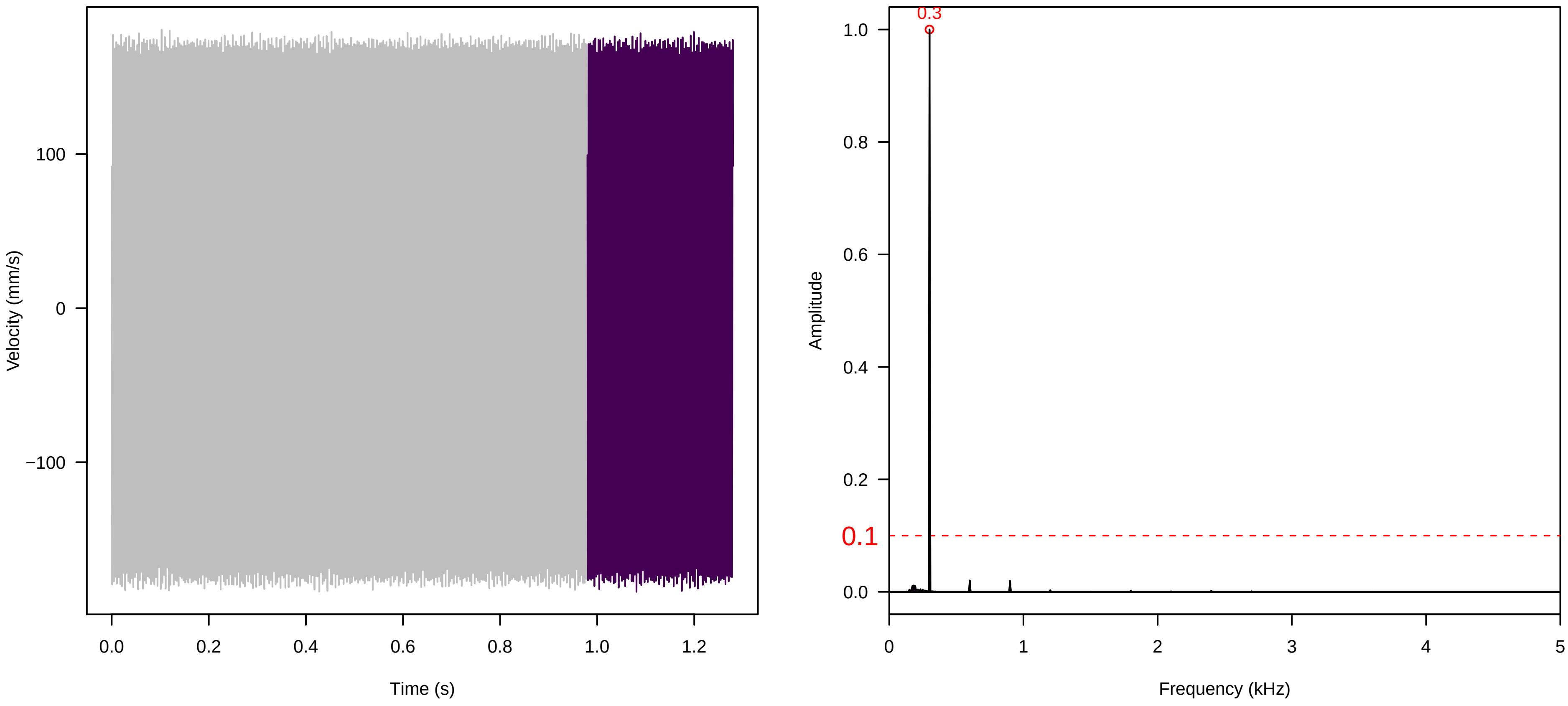

**Vel. = 0.057 ; Str. = Receptacle ; Axis = z ; Fl. accession = 10−s−79−2**

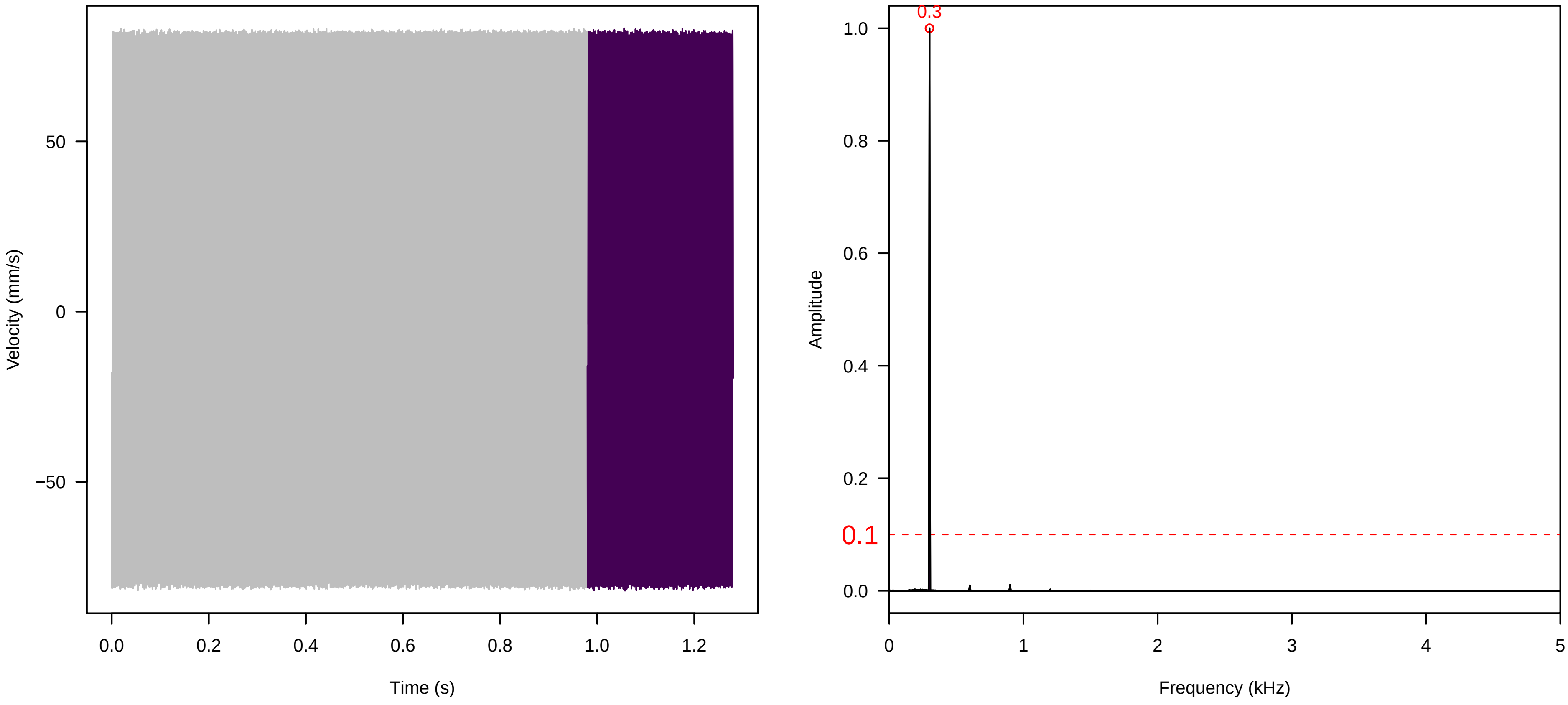


**Vel. = 0.057 ; Str. = PA ; Axis = z ; Fl. accession = 10−s−79−2**

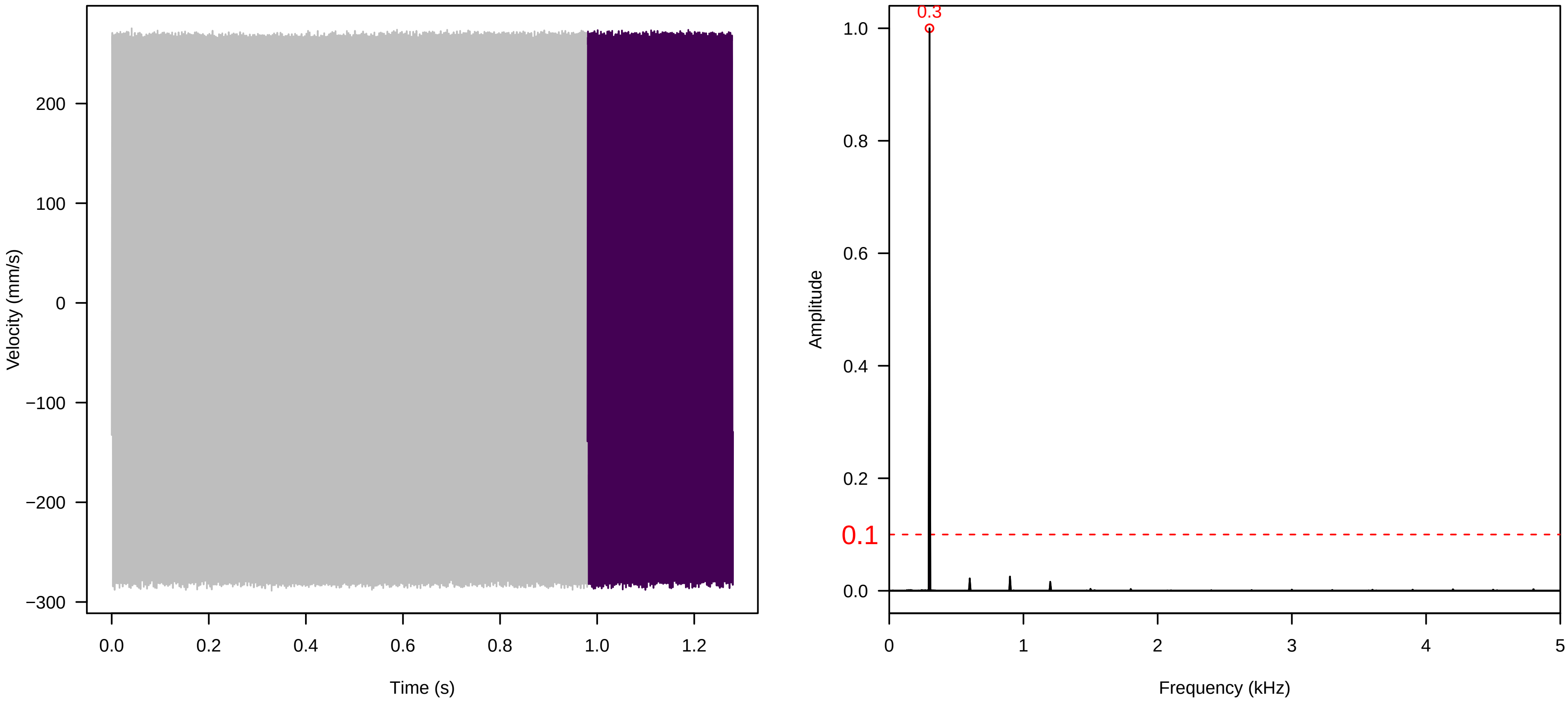


**Vel. = 0.057 ; Str. = Receptacle ; Axis = z ; Fl. accession = 10−s−79−2**

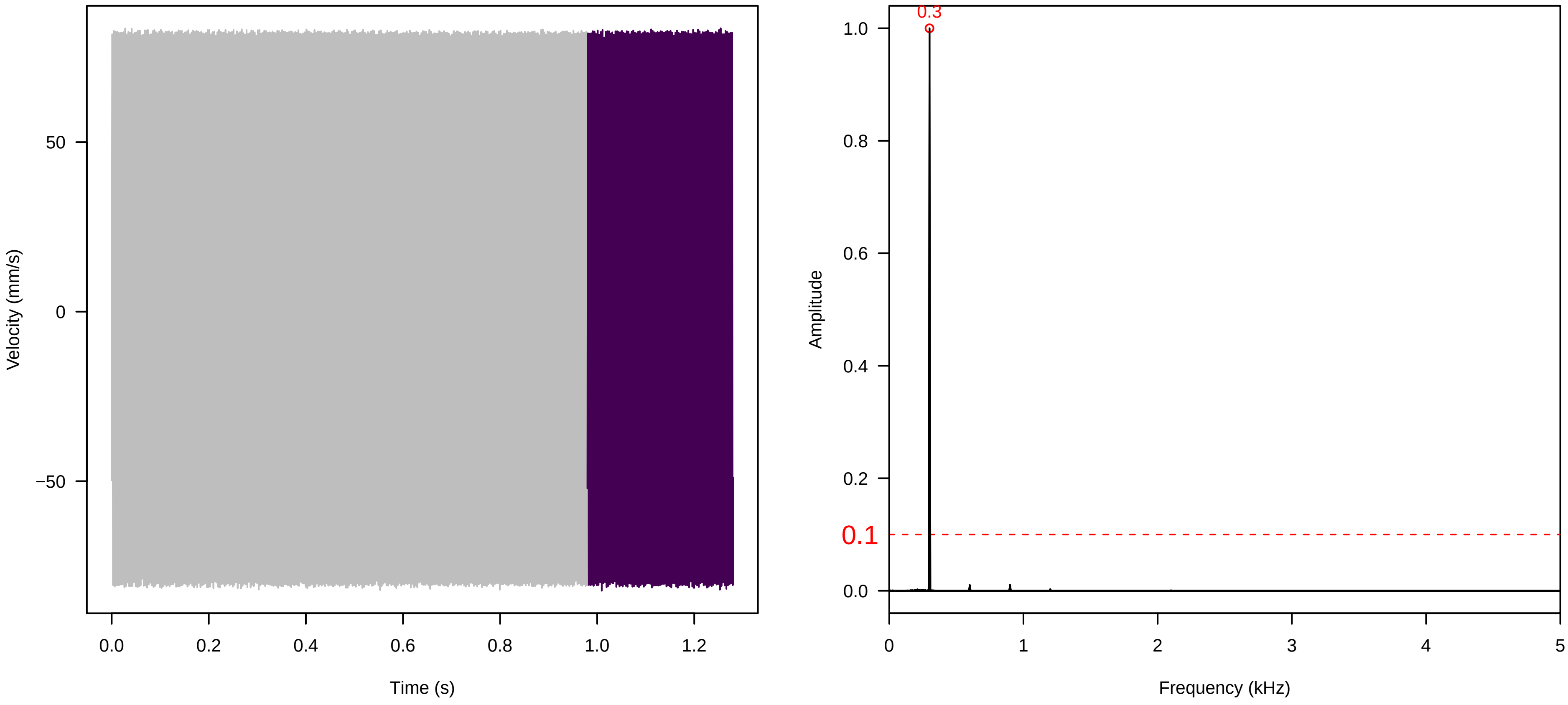

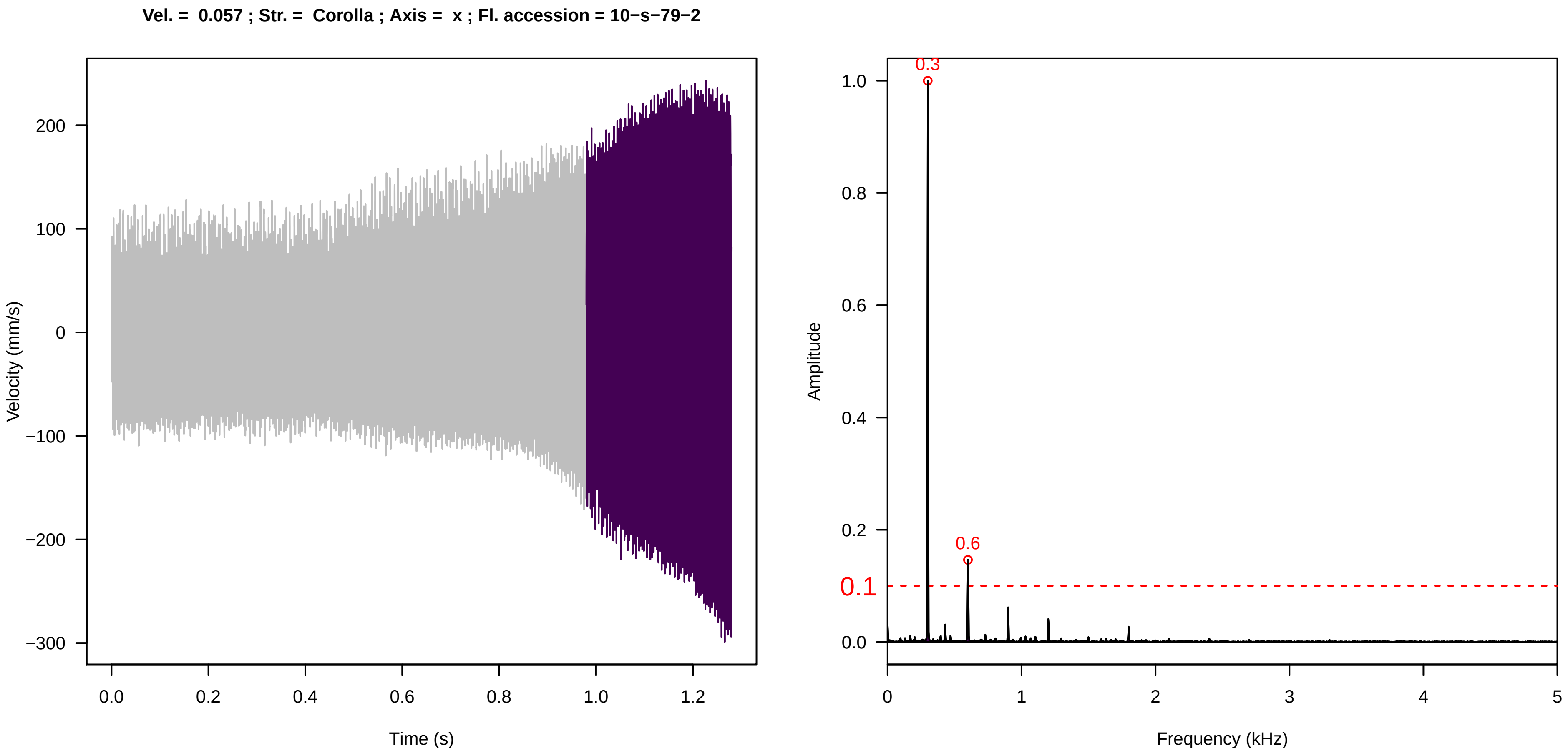
Vel. = 0.057 ; Str. = Corolla ; Axis = x ; Fl. accession = 10−s−79−2
Velocity (mm/s)
Time (s)
Amplitude
Frequency (kHz)
0.3
0.6
0.1

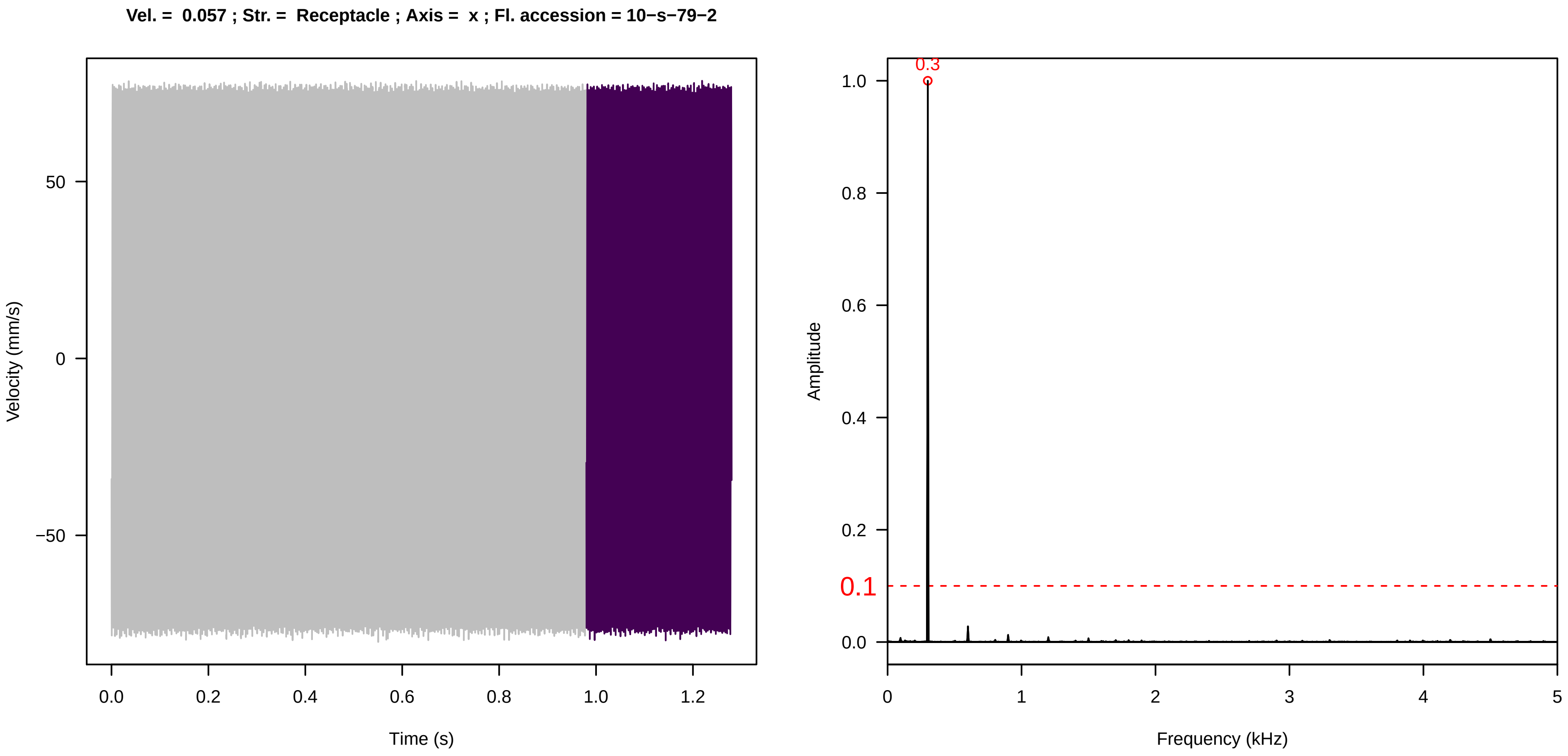
Vel. = 0.057 ; Str. = Receptacle ; Axis = x ; Fl. accession = 10−s−79−2
Velocity (mm/s)
Time (s)
Amplitude
Frequency (kHz)
0.3
0.1

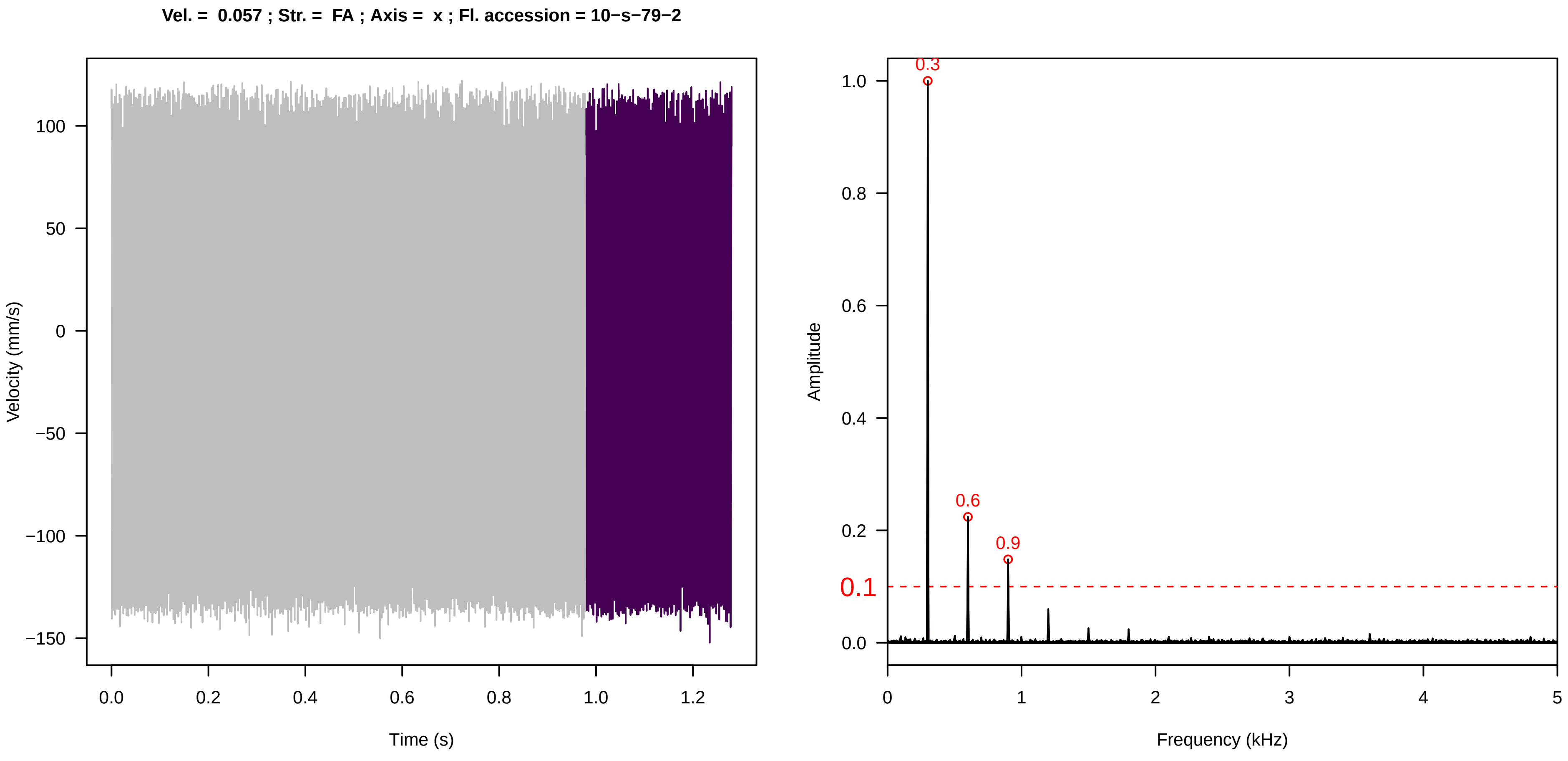
Vel. = 0.057 ; Str. = FA ; Axis = x ; Fl. accession = 10−s−79−2
Velocity (mm/s)
Time (s)
Amplitude
Frequency (kHz)
0.3
0.6
0.9
0.1

**Vel. = 0.057 ; Str. = Receptacle ; Axis = x ; Fl. accession = 10−s−79−2**

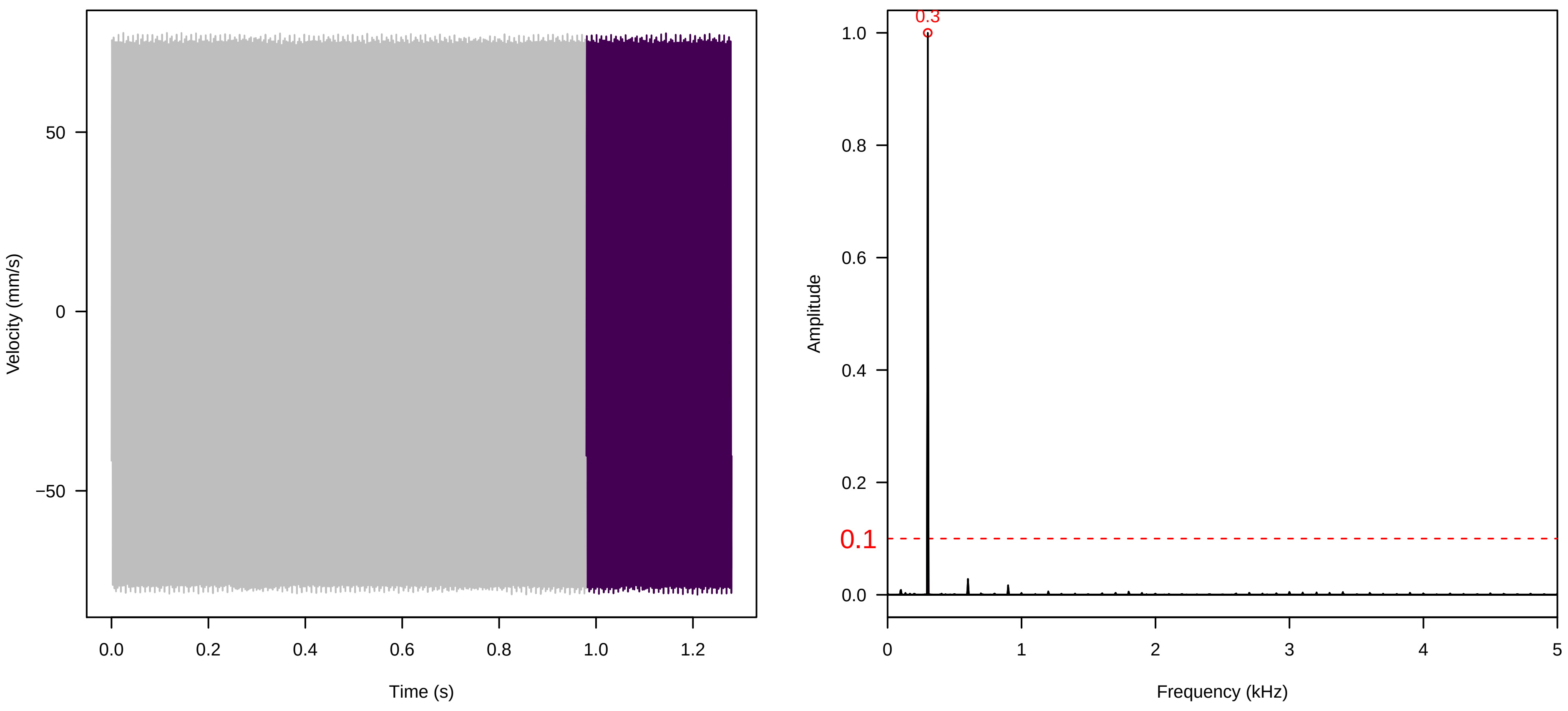


**Vel. = 0.057 ; Str. = PA ; Axis = x ; Fl. accession = 10−s−79−2**

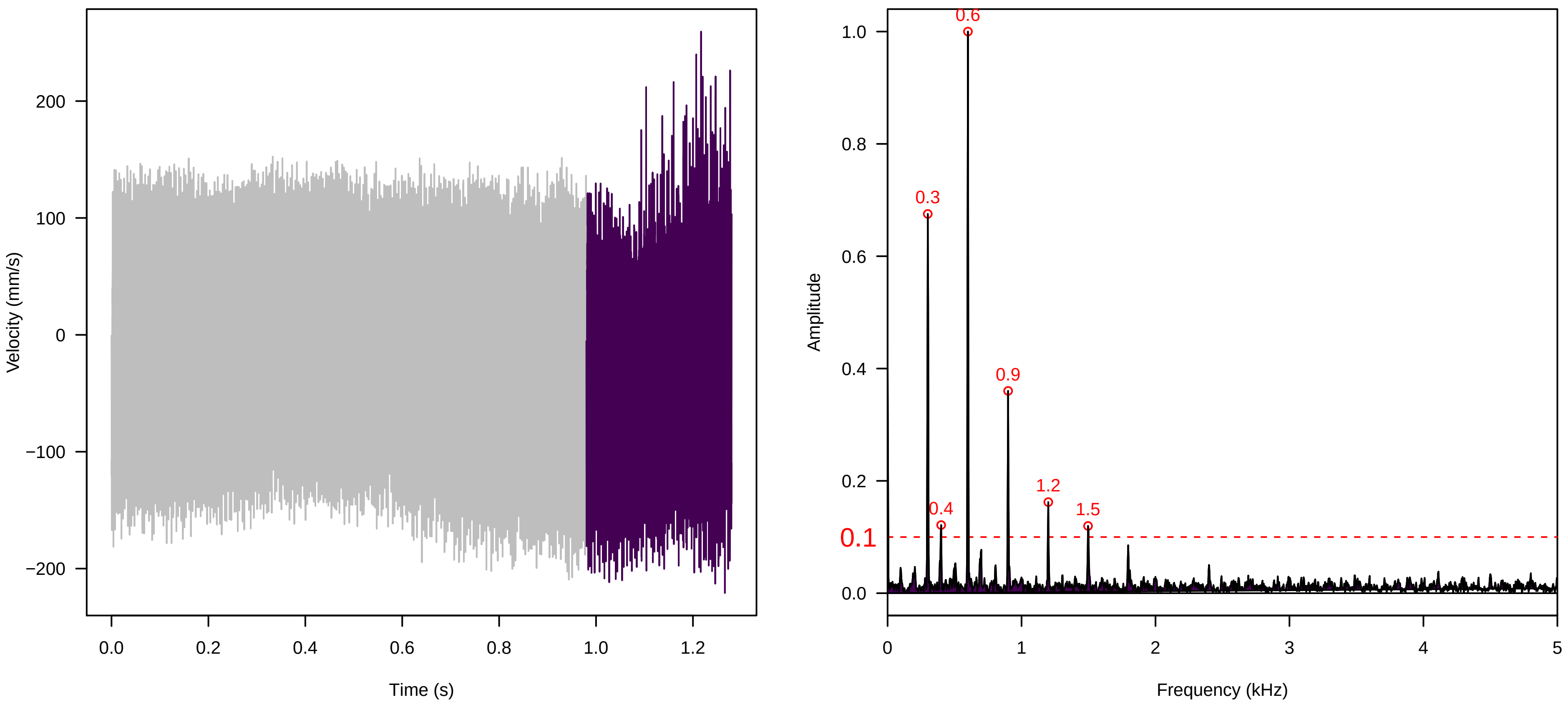


**Vel. = 0.057 ; Str. = Receptacle ; Axis = x ; Fl. accession = 10−s−79−2**

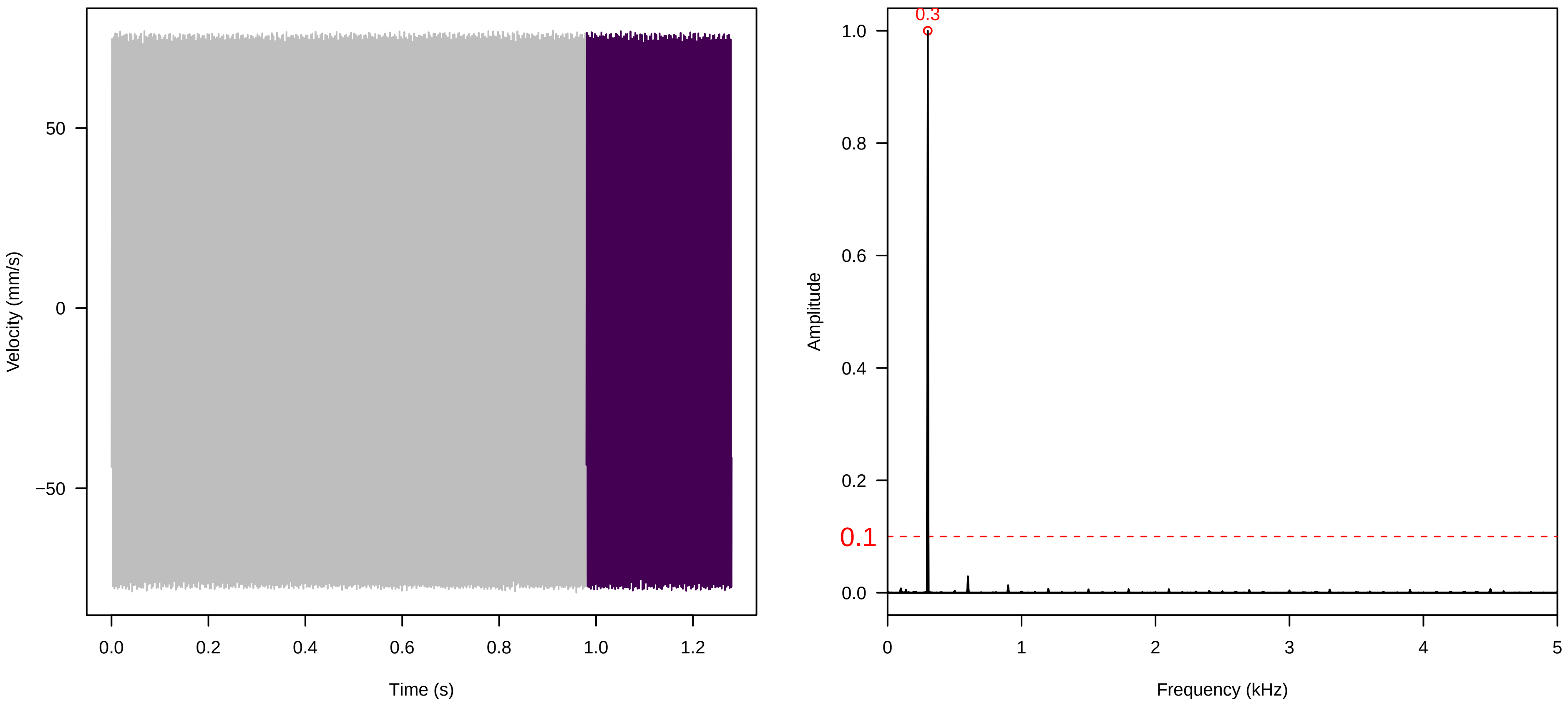

**Vel. = 0.028 ; Str. = PA ; Axis = x ; Fl. accession = 10−s−79−2**

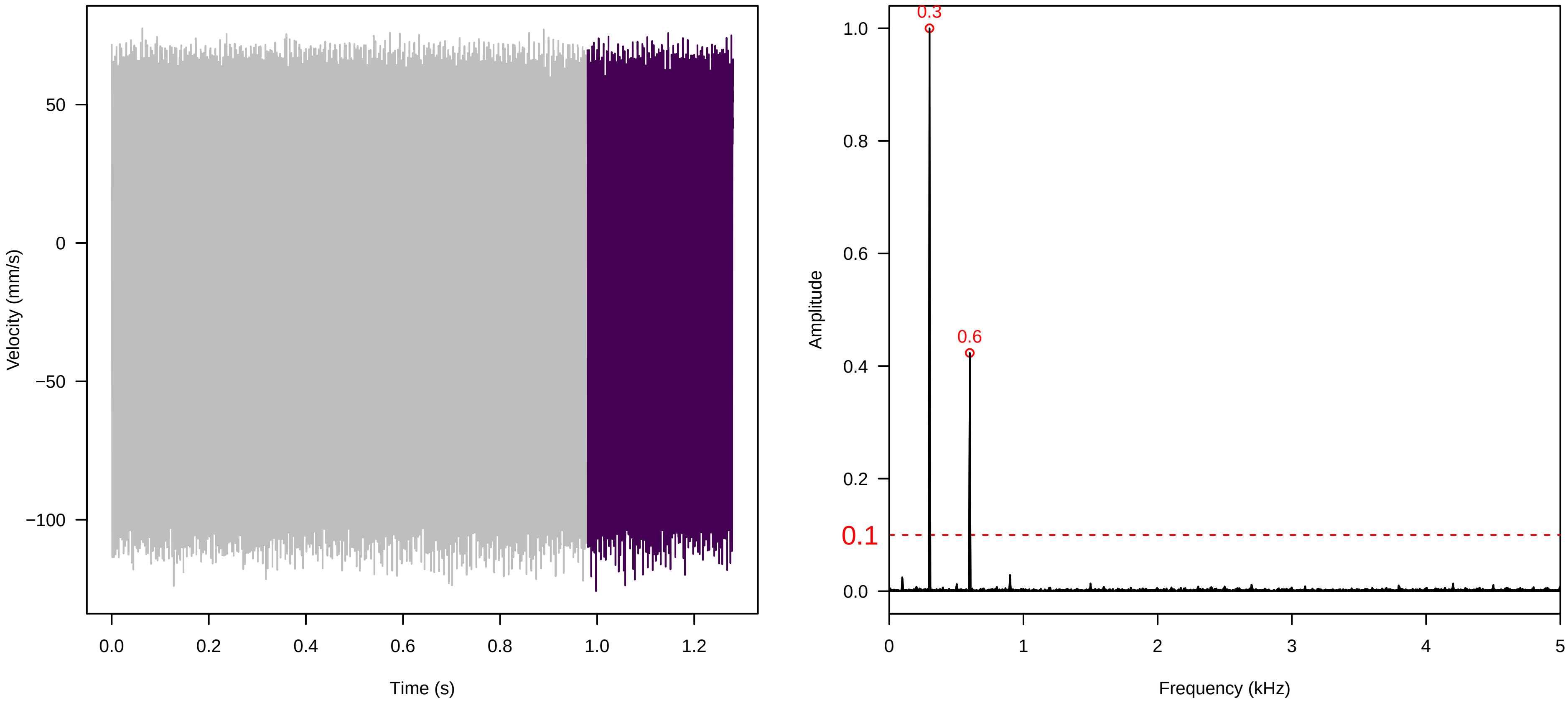


**Vel. = 0.028 ; Str. = Receptacle ; Axis = x ; Fl. accession = 10−s−79−2**

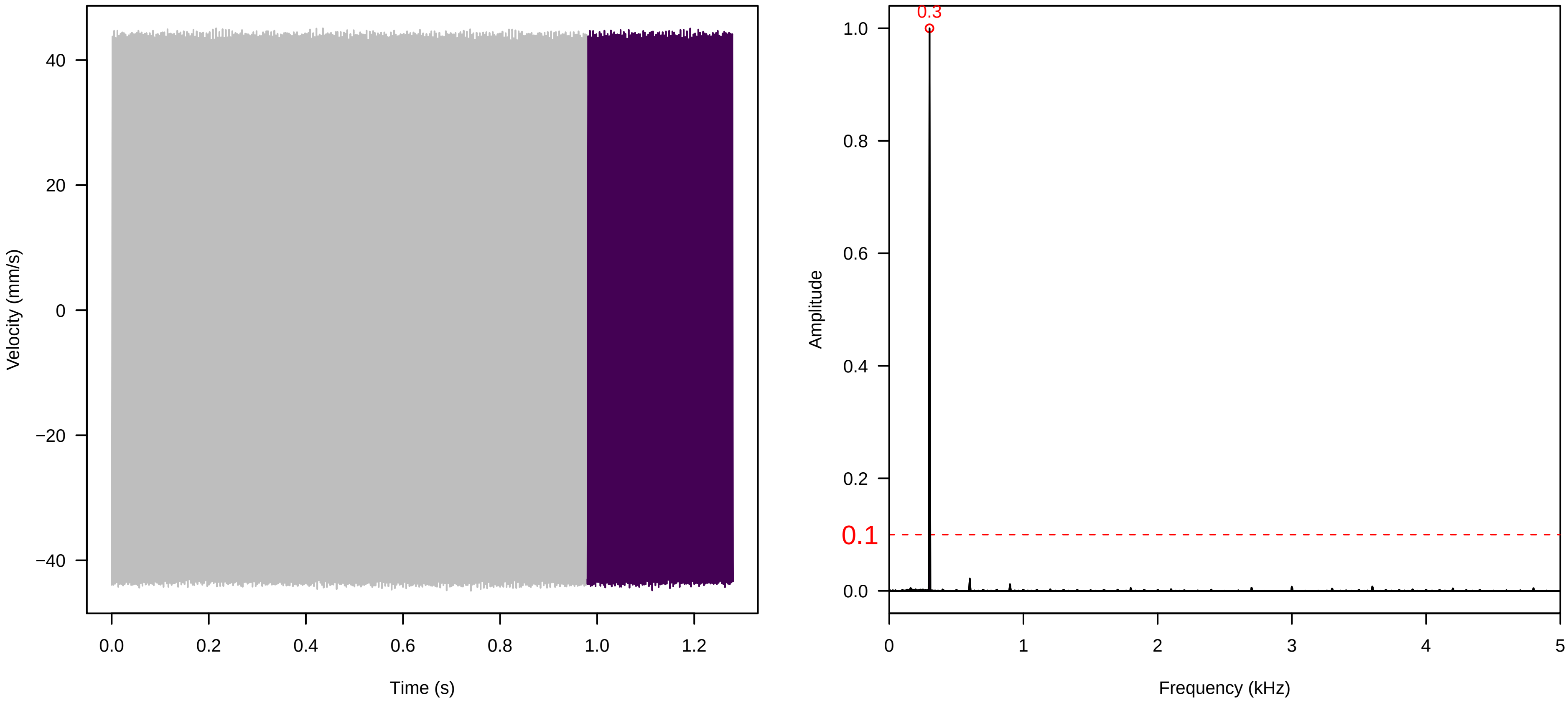


**Vel. = 0.028 ; Str. = FA ; Axis = x ; Fl. accession = 10−s−79−2**

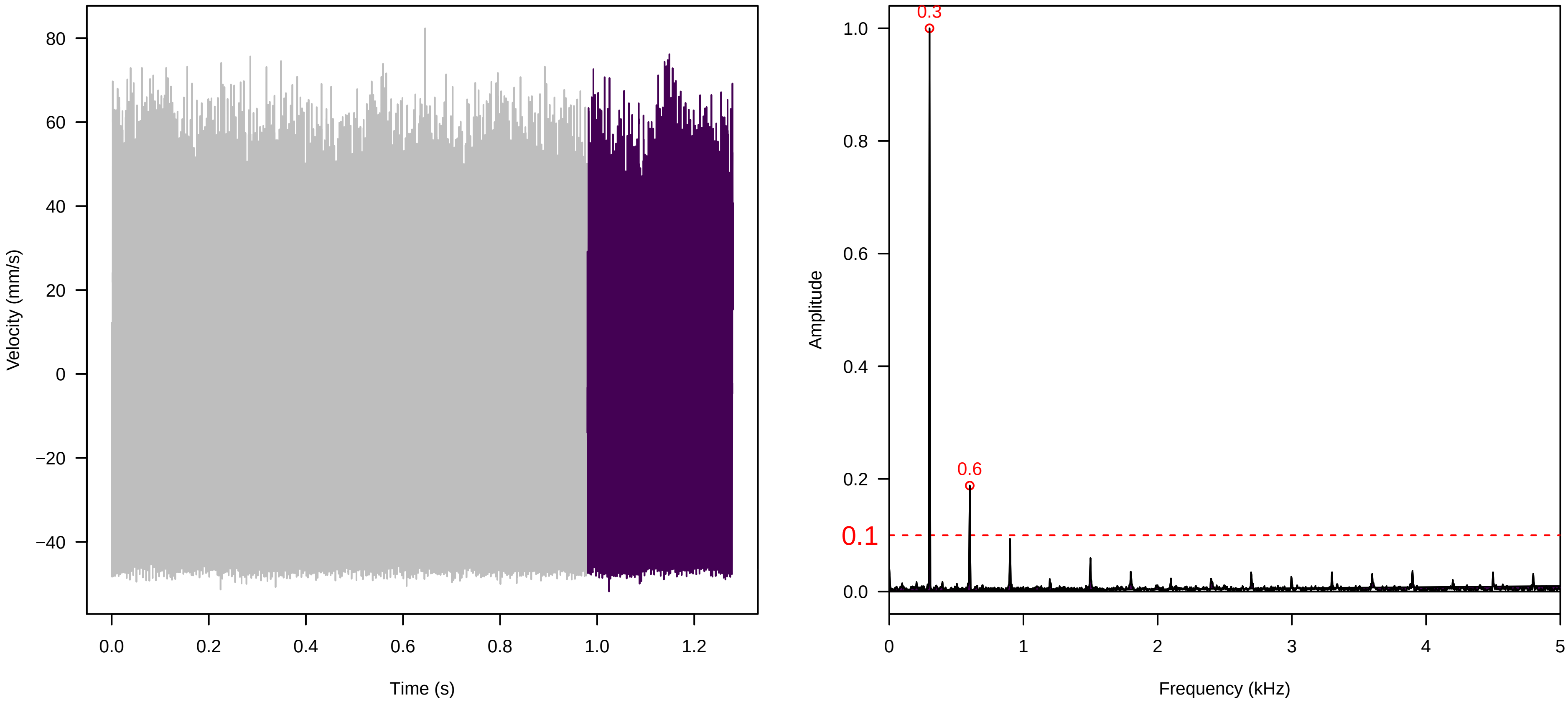

**Vel. = 0.028 ; Str. = Receptacle ; Axis = x ; Fl. accession = 10−s−79−2**

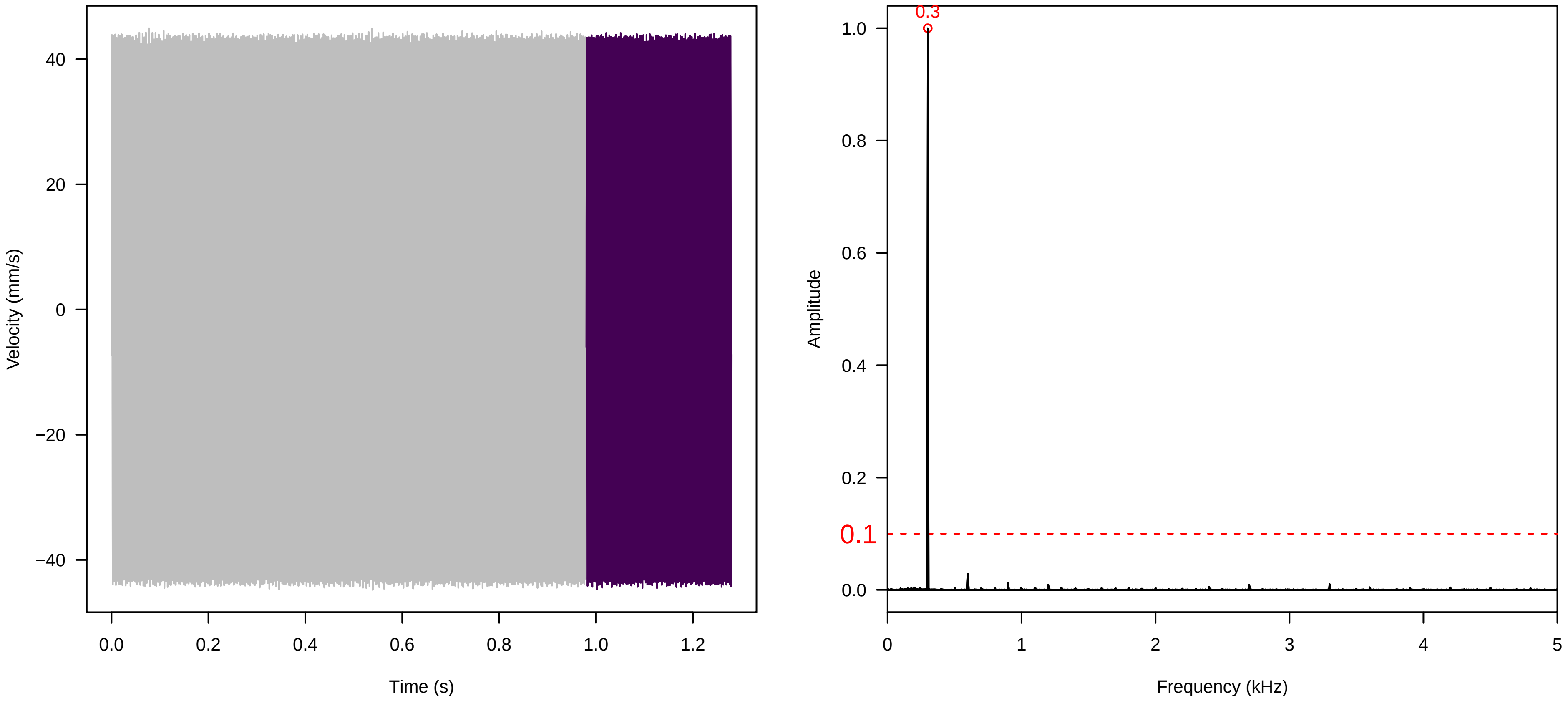


**Vel. = 0.028 ; Str. = Corolla ; Axis = x ; Fl. accession = 10−s−79−2**

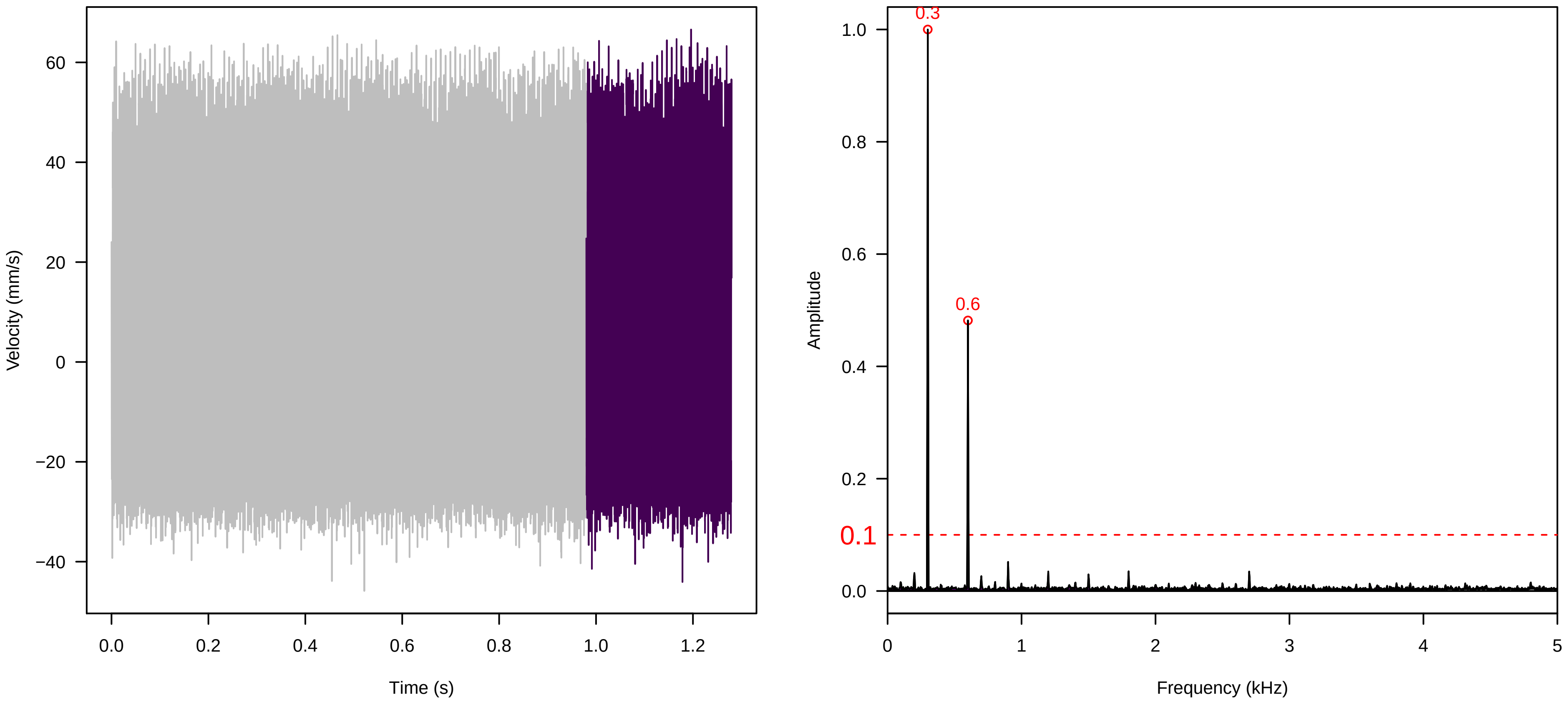


**Vel. = 0.028 ; Str. = Receptacle ; Axis = x ; Fl. accession = 10−s−79−2**

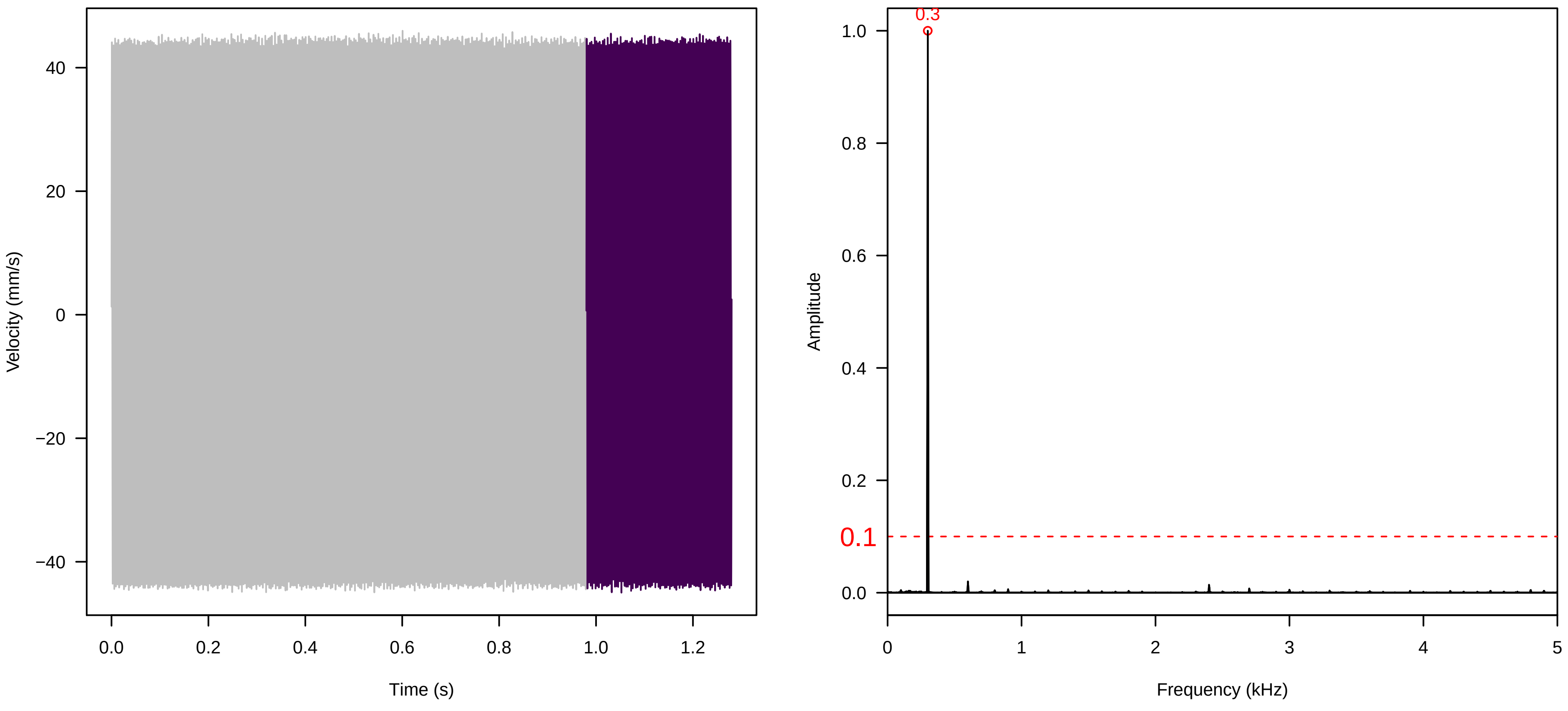

**Vel. = 0.014 ; Str. = Corolla ; Axis = x ; Fl. accession = 10−s−79−2**

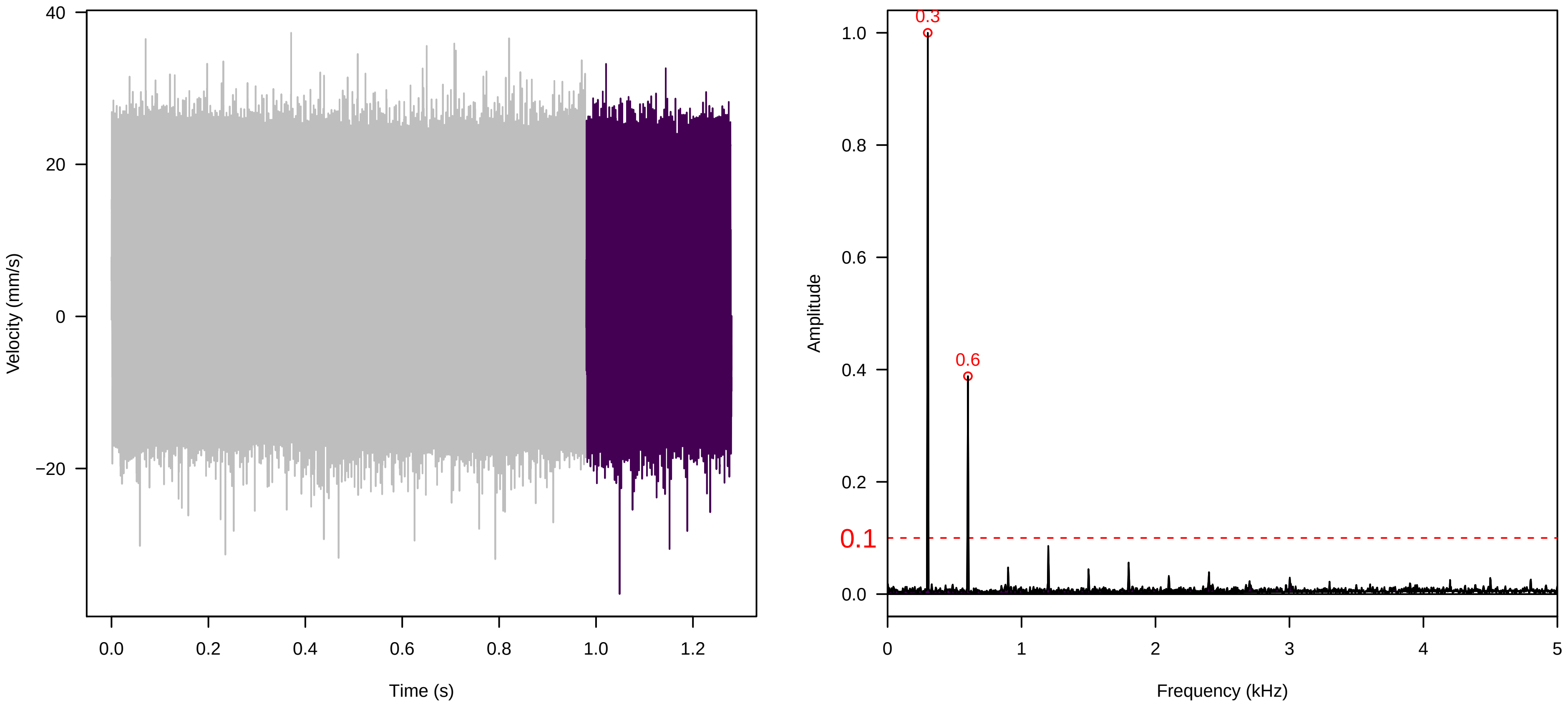


**Vel. = 0.014 ; Str. = Receptacle ; Axis = x ; Fl. accession = 10−s−79−2**

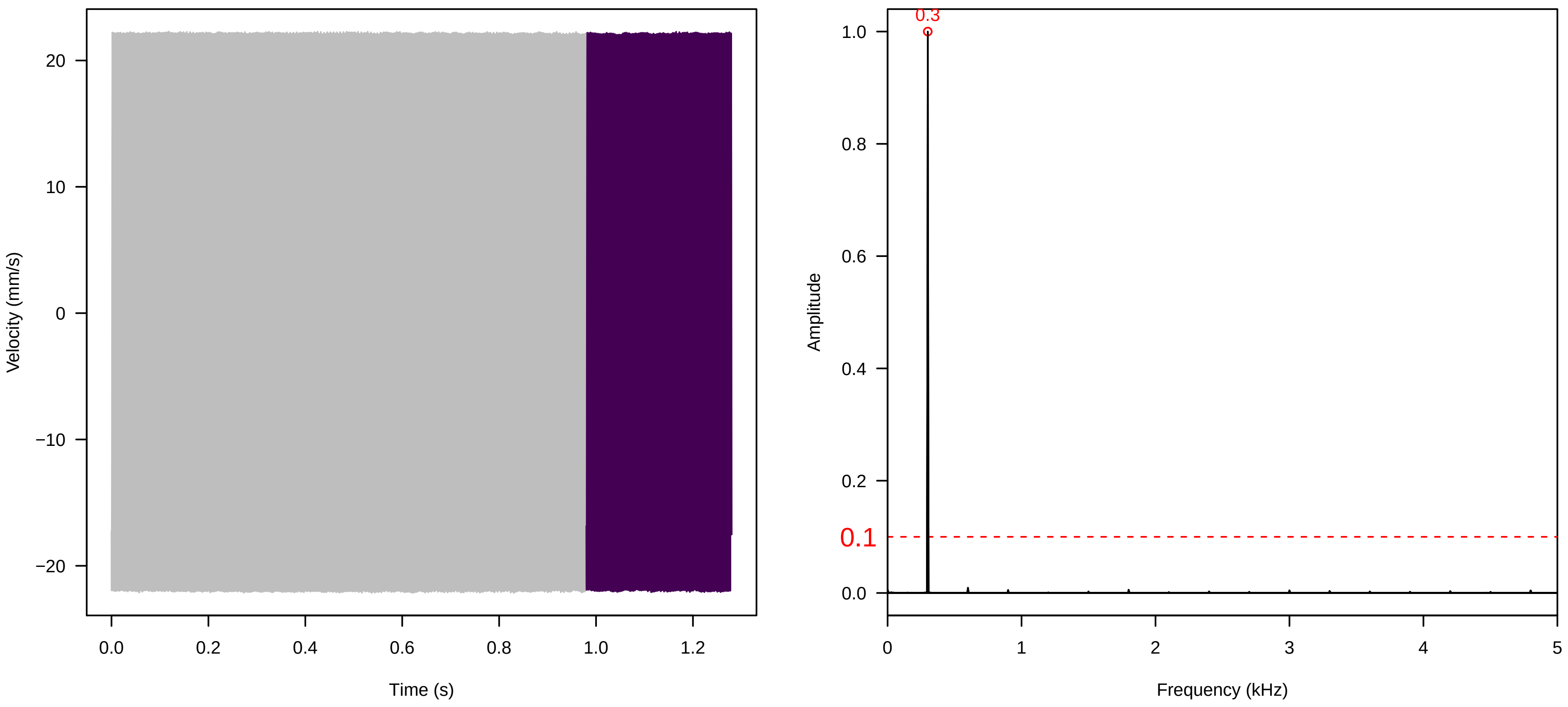


**Vel. = 0.014 ; Str. = FA ; Axis = x ; Fl. accession = 10−s−79−2**

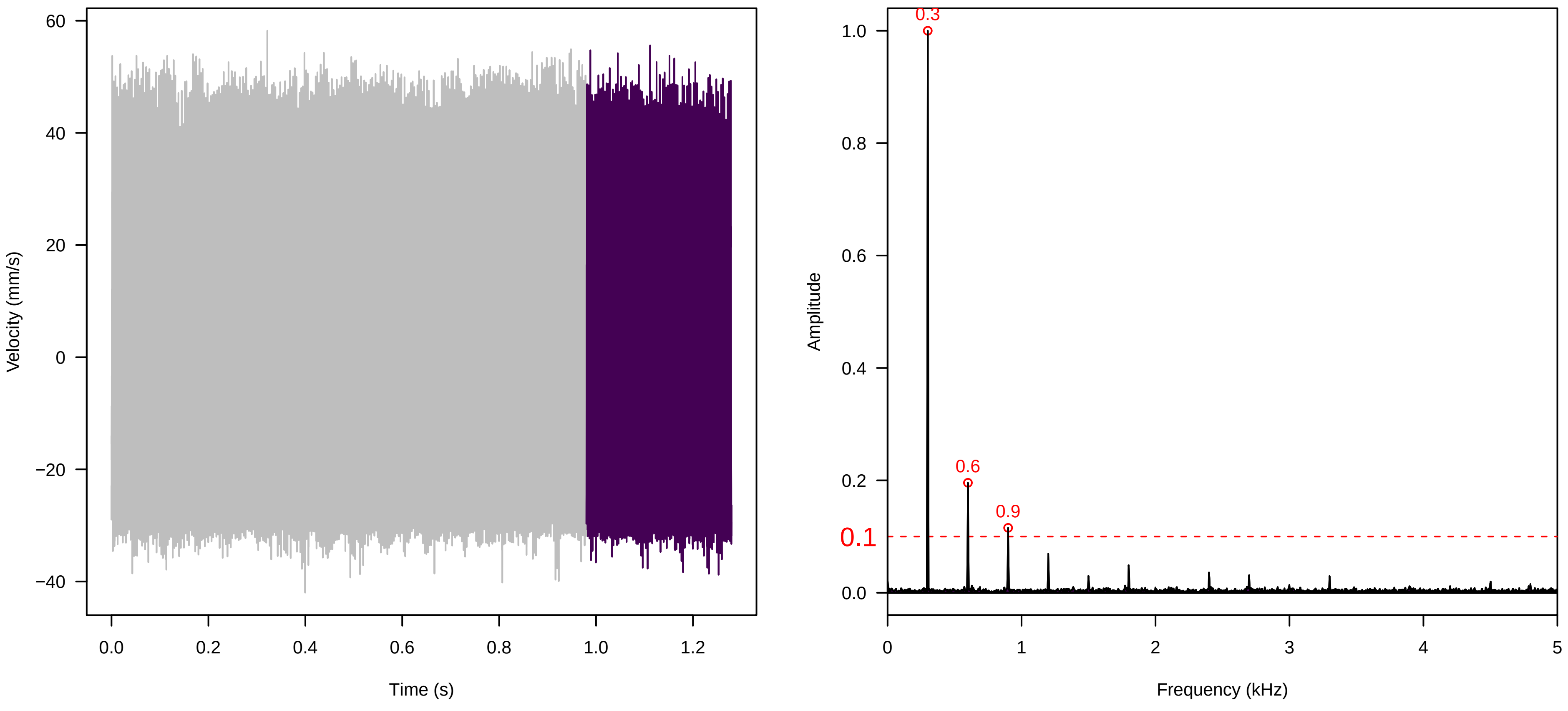

**Vel. =  0.014 ; Str. =  Receptacle ; Axis =  x ; Fl. accession = 10−s−79−2**

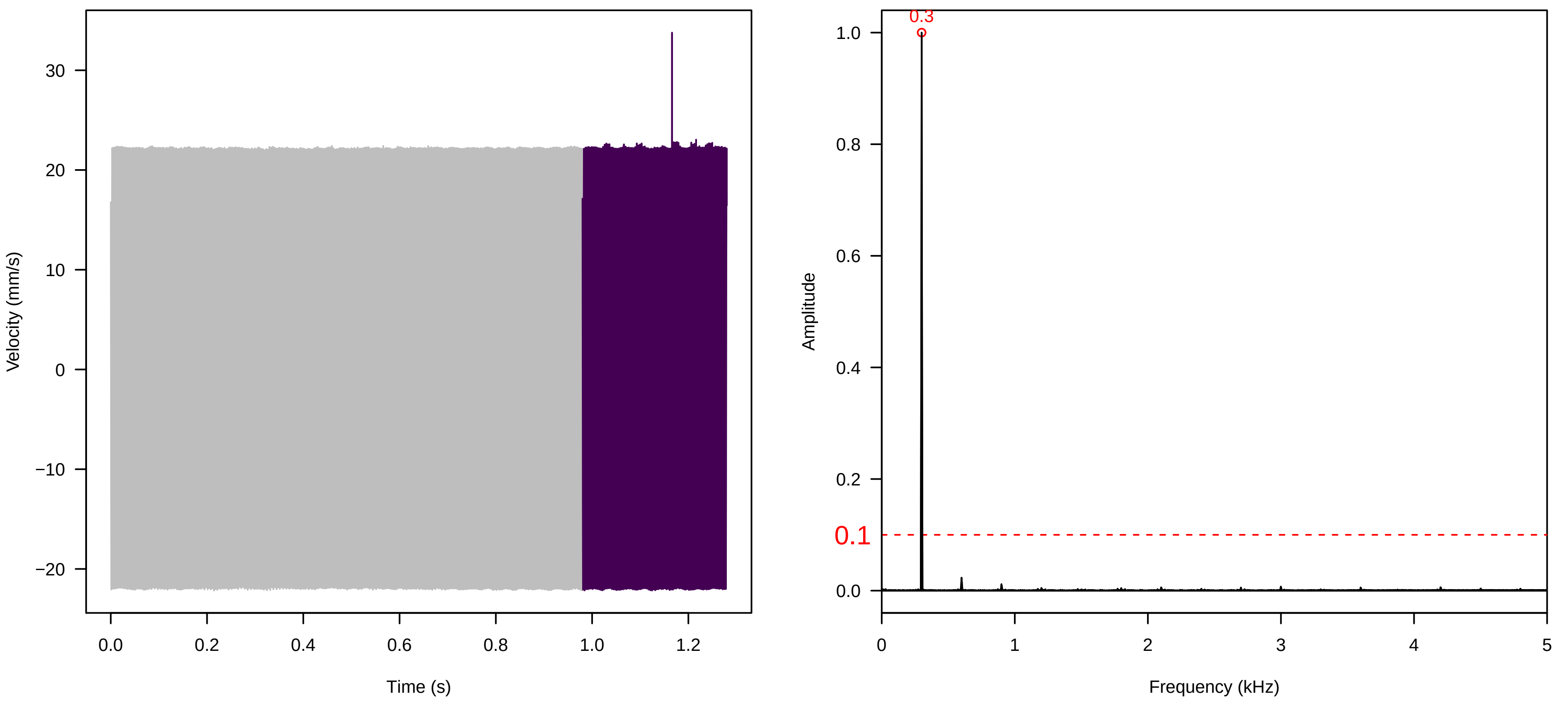


**Vel. =  0.014 ; Str. =  PA ; Axis =  x ; Fl. accession = 10−s−79−2**

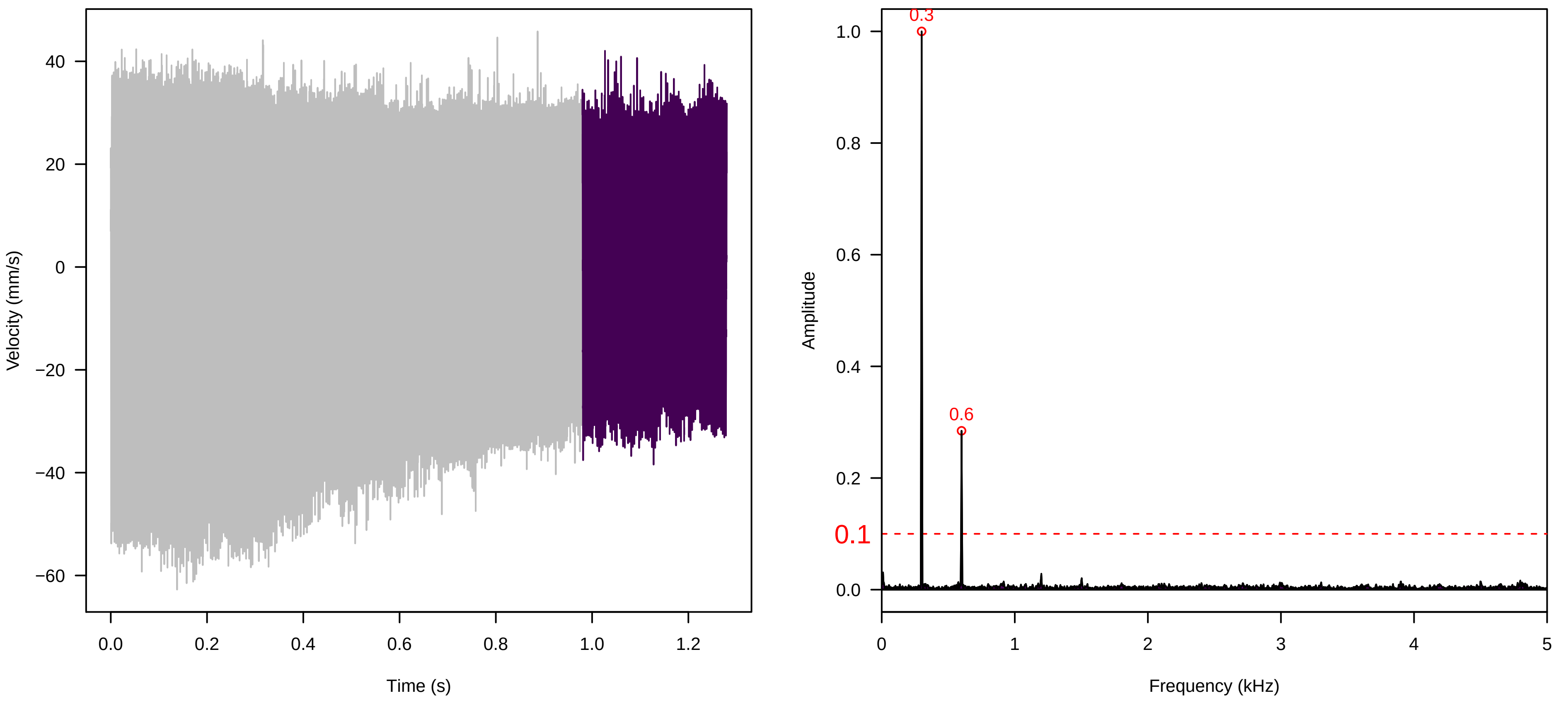


**Vel. =  0.014 ; Str. =  Receptacle ; Axis =  x ; Fl. accession = 10−s−79−2**

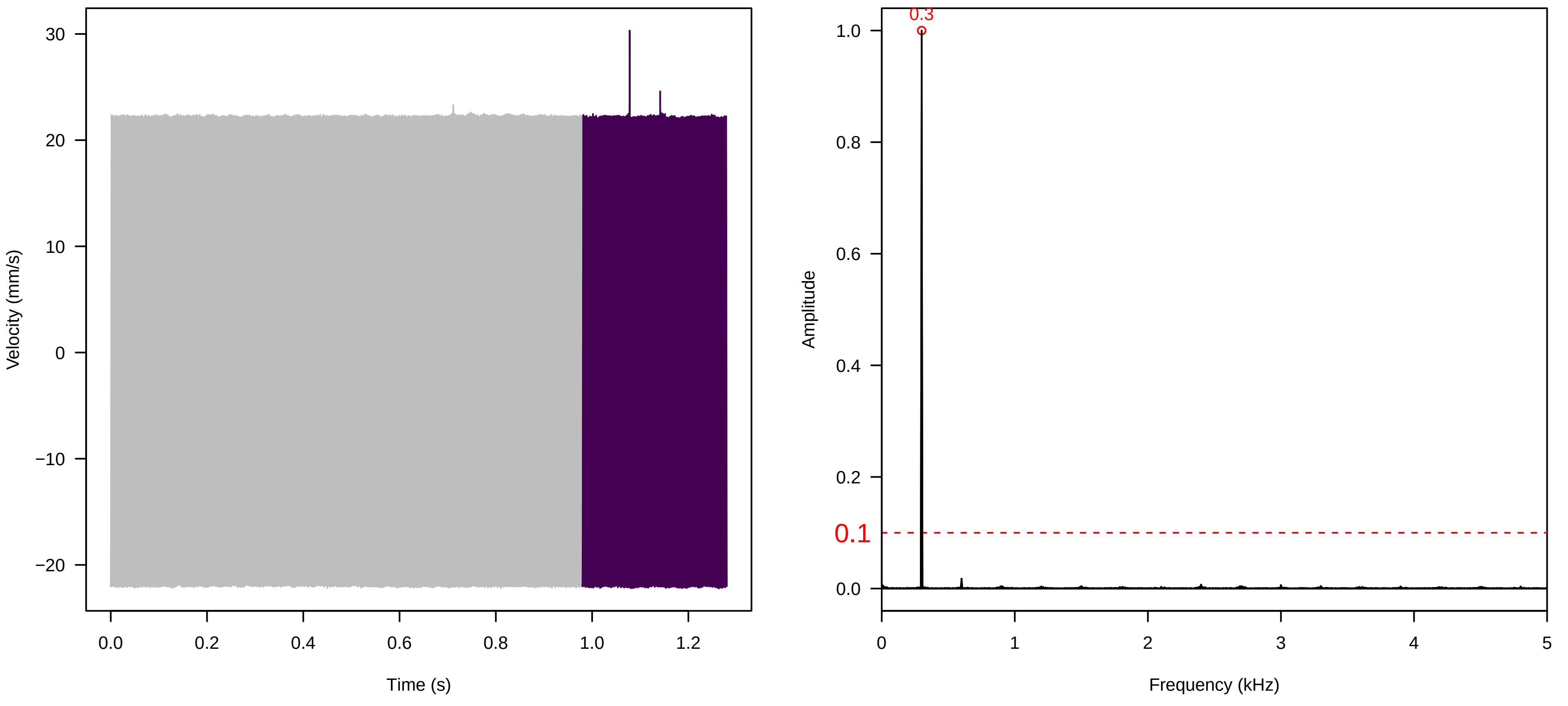

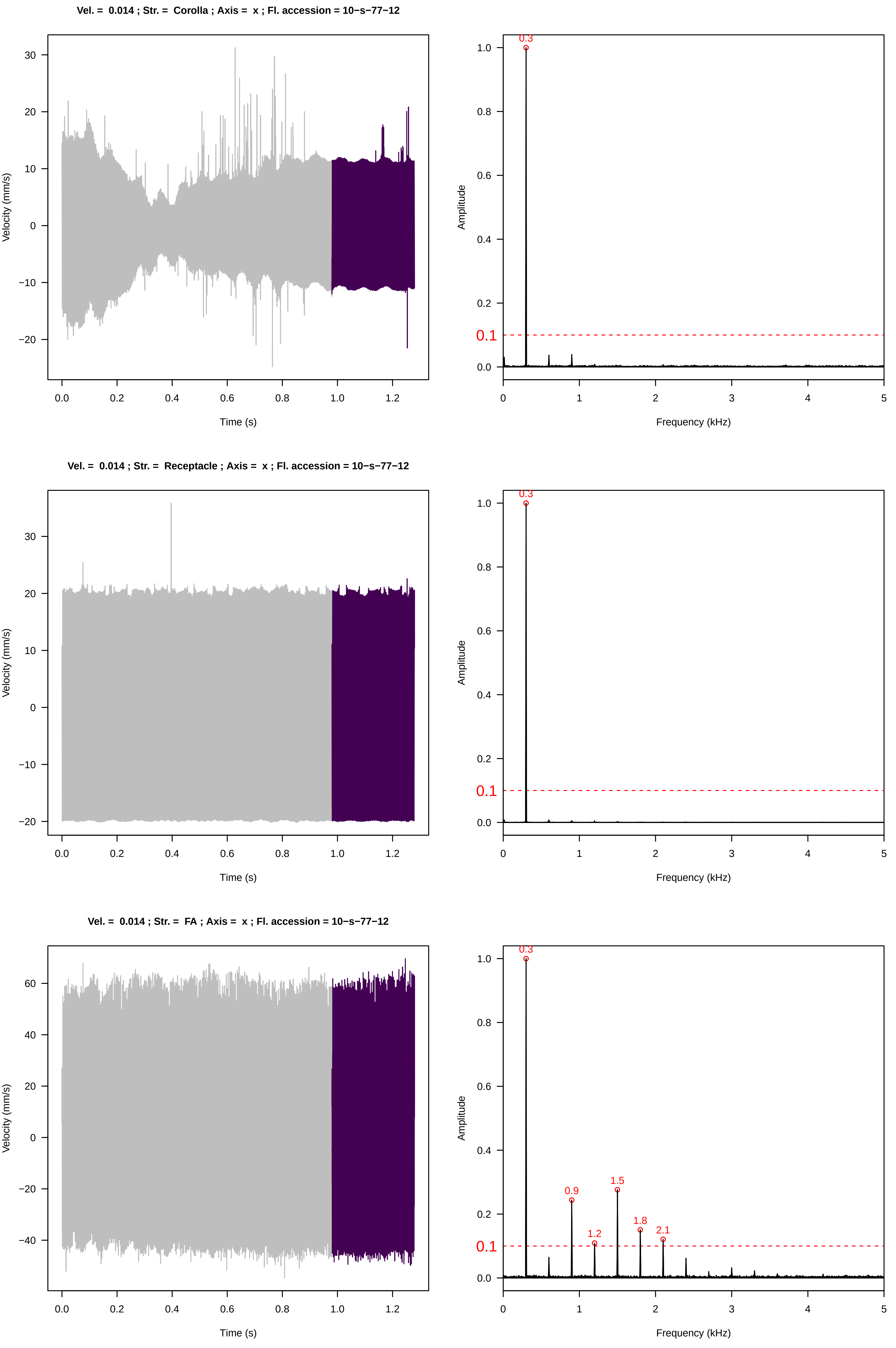
Vel. = 0.014 ; Str. = Corolla ; Axis = x ; Fl. accession = 10−s−77−12
Velocity (mm/s)
Time (s)
Amplitude
Frequency (kHz)
0.3
0.1
Vel. = 0.014 ; Str. = Receptacle ; Axis = x ; Fl. accession = 10−s−77−12
Velocity (mm/s)
Time (s)
Amplitude
Frequency (kHz)
0.3
0.1
Vel. = 0.014 ; Str. = FA ; Axis = x ; Fl. accession = 10−s−77−12
Velocity (mm/s)
Time (s)
Amplitude
Frequency (kHz)
0.3
0.9
1.2
1.5
1.8
2.1
0.1

**Vel. = 0.014 ; Str. = Receptacle ; Axis = x ; Fl. accession = 10−s−77−12**

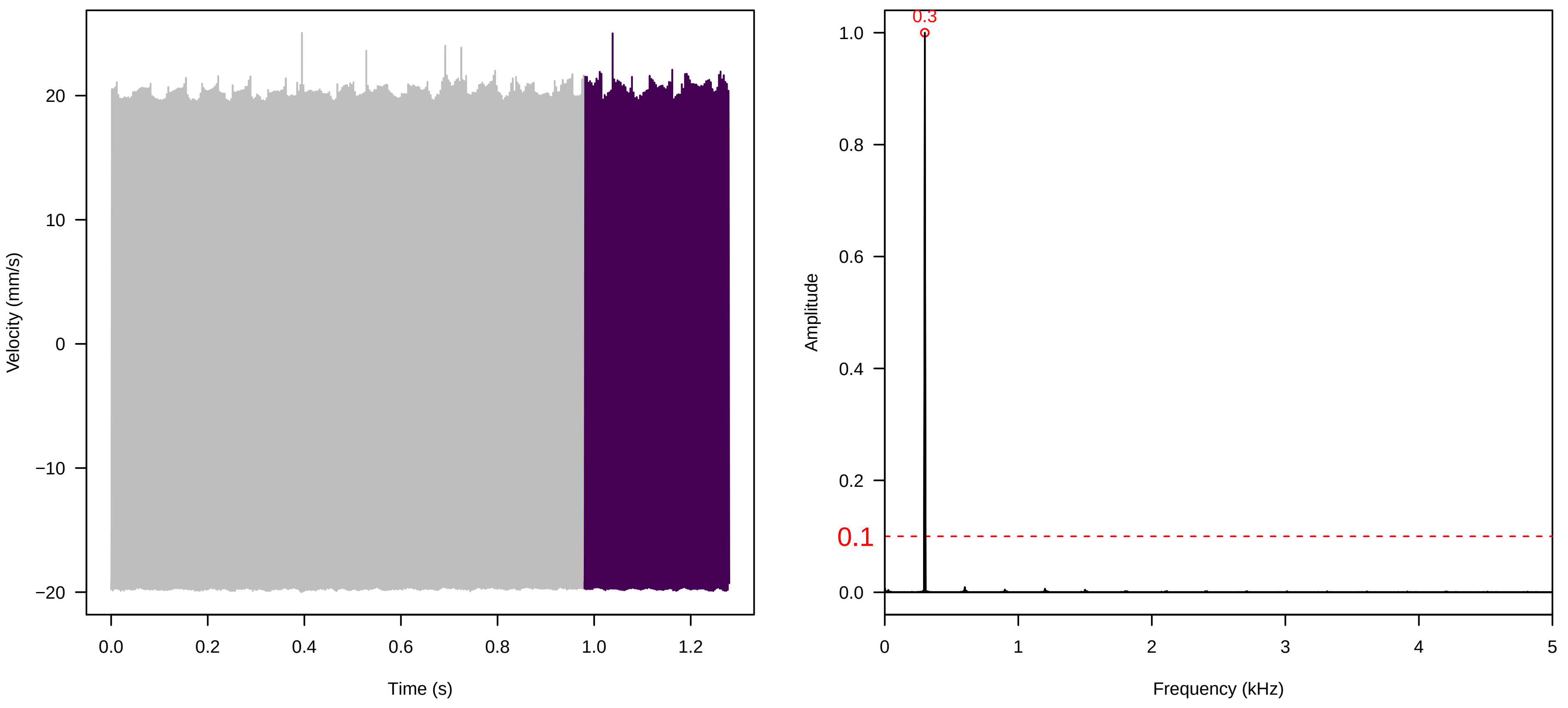


**Vel. = 0.014 ; Str. = PA ; Axis = x ; Fl. accession = 10−s−77−12**

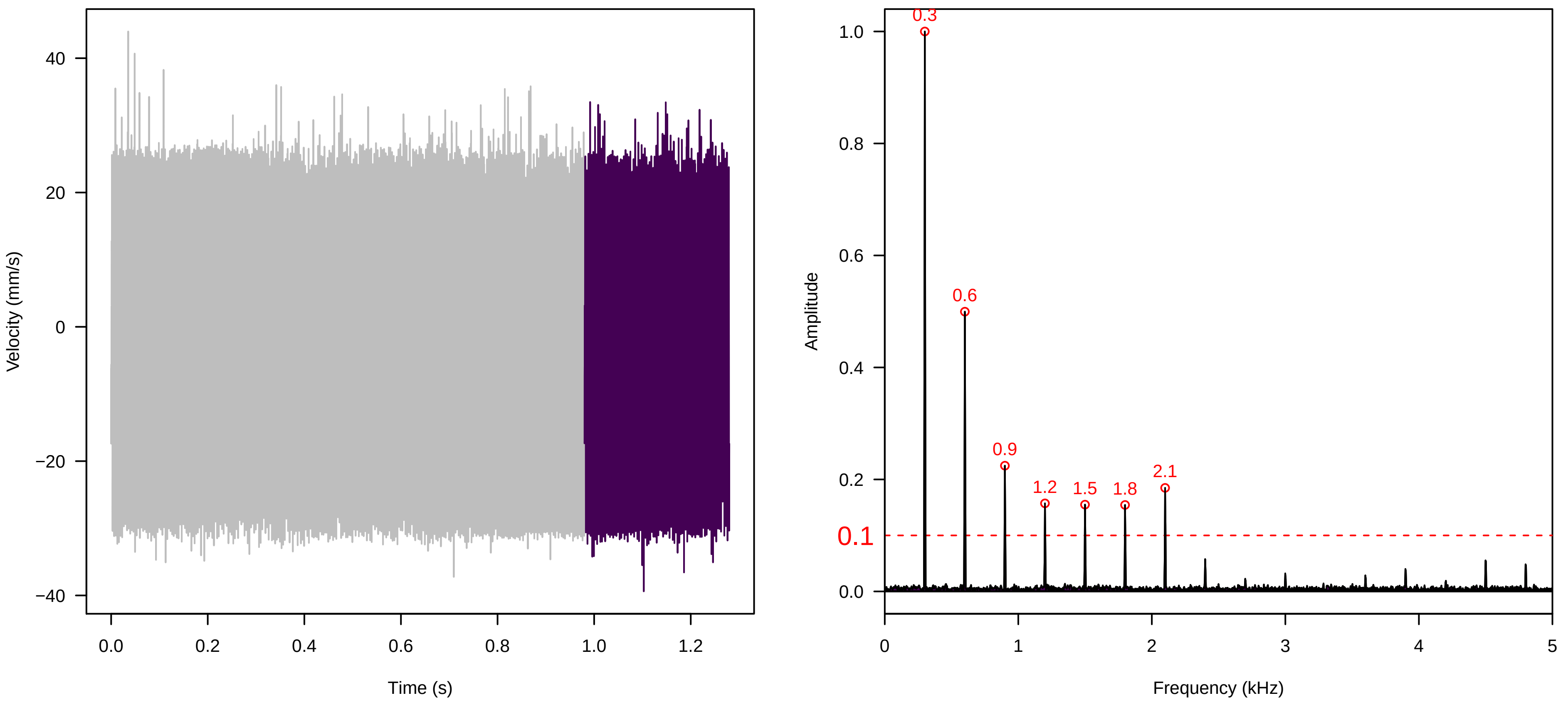


**Vel. = 0.014 ; Str. = Receptacle ; Axis = x ; Fl. accession = 10−s−77−12**

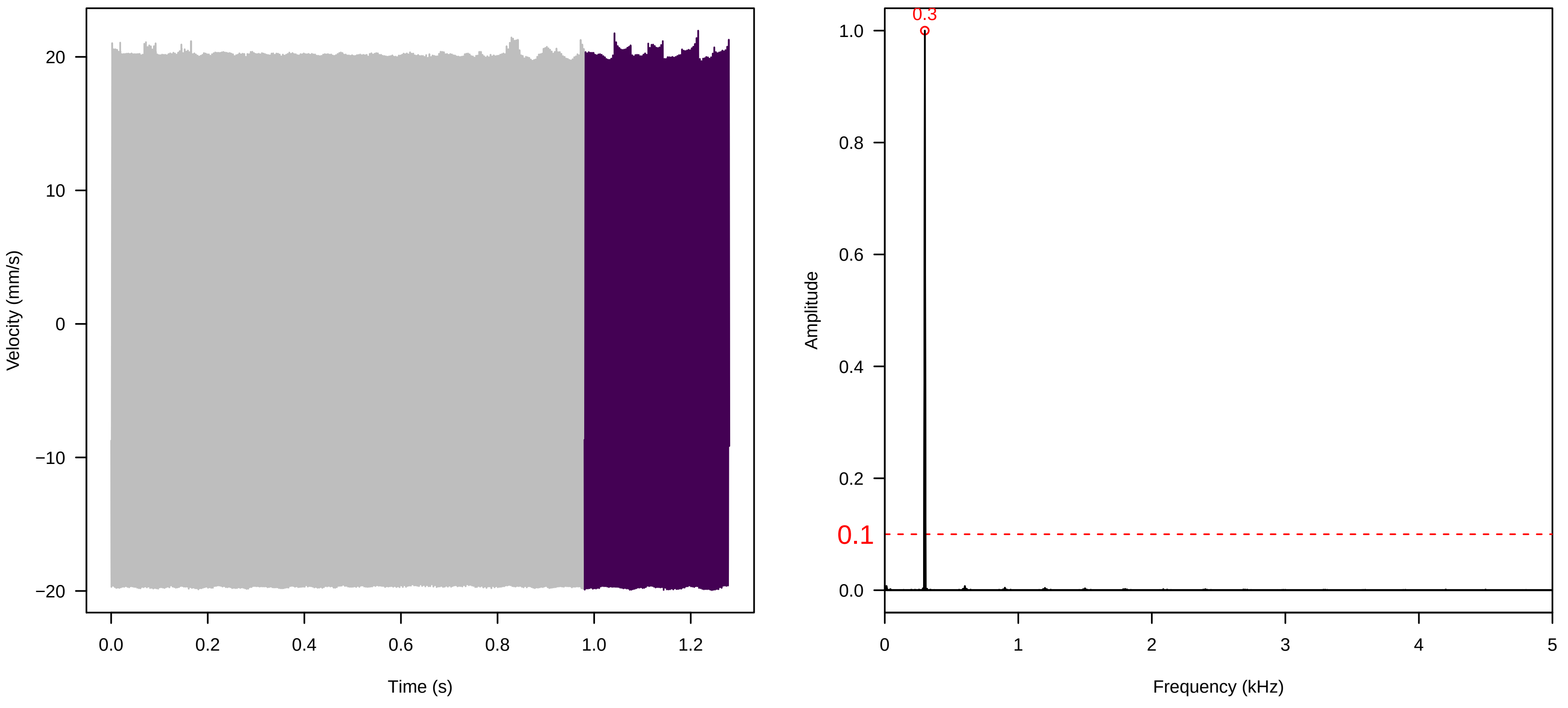

**Vel. =  0.028 ; Str. =  PA ; Axis =  x ; Fl. accession = 10−s−77−12**

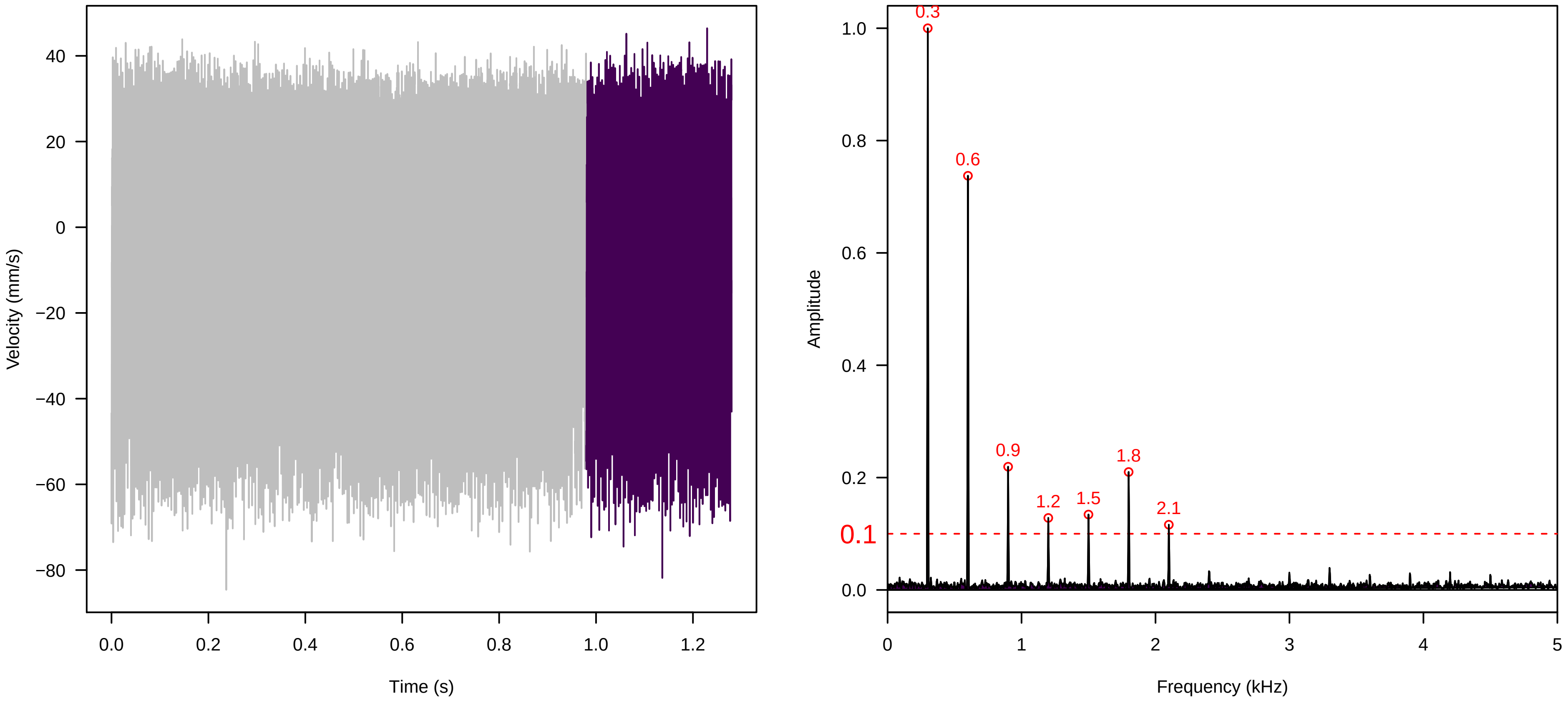


**Vel. =  0.028 ; Str. =  Receptacle ; Axis =  x ; Fl. accession = 10−s−77−12**

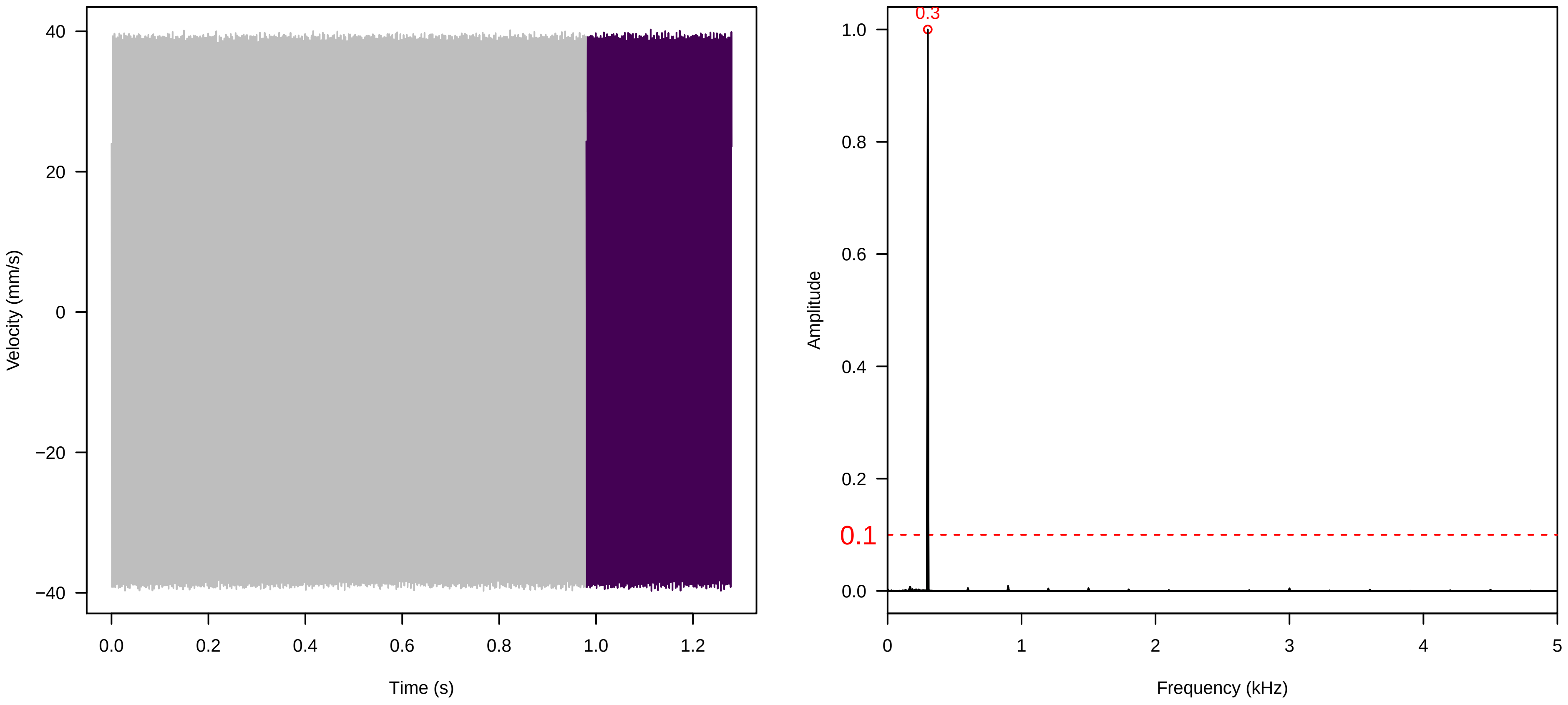


**Vel. =  0.028 ; Str. =  FA ; Axis =  x ; Fl. accession = 10−s−77−12**

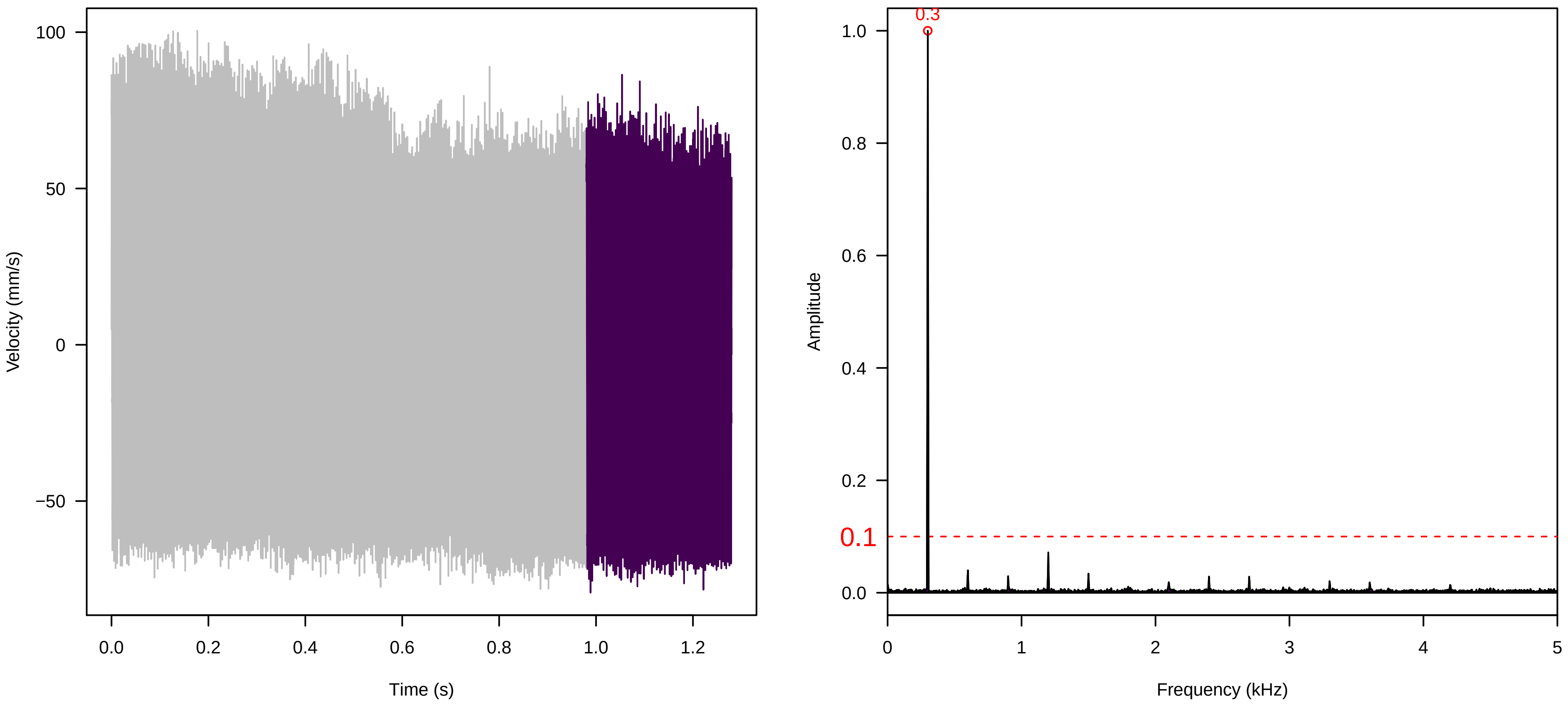

Vel. = 0.028 ; Str. = Receptacle ; Axis = x ; Fl. accession = 10−s−77−12

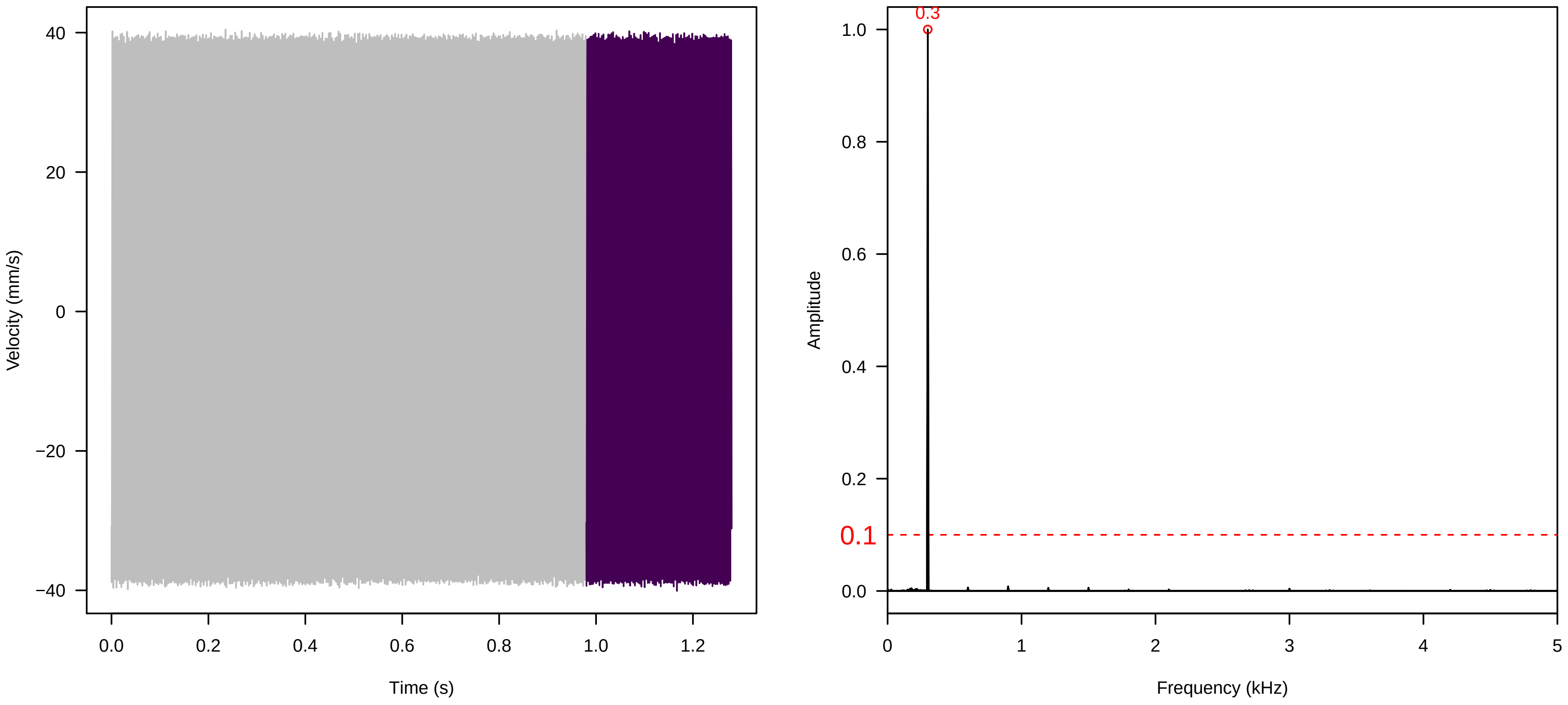


Vel. = 0.028 ; Str. = Corolla ; Axis = x ; Fl. accession = 10−s−77−12

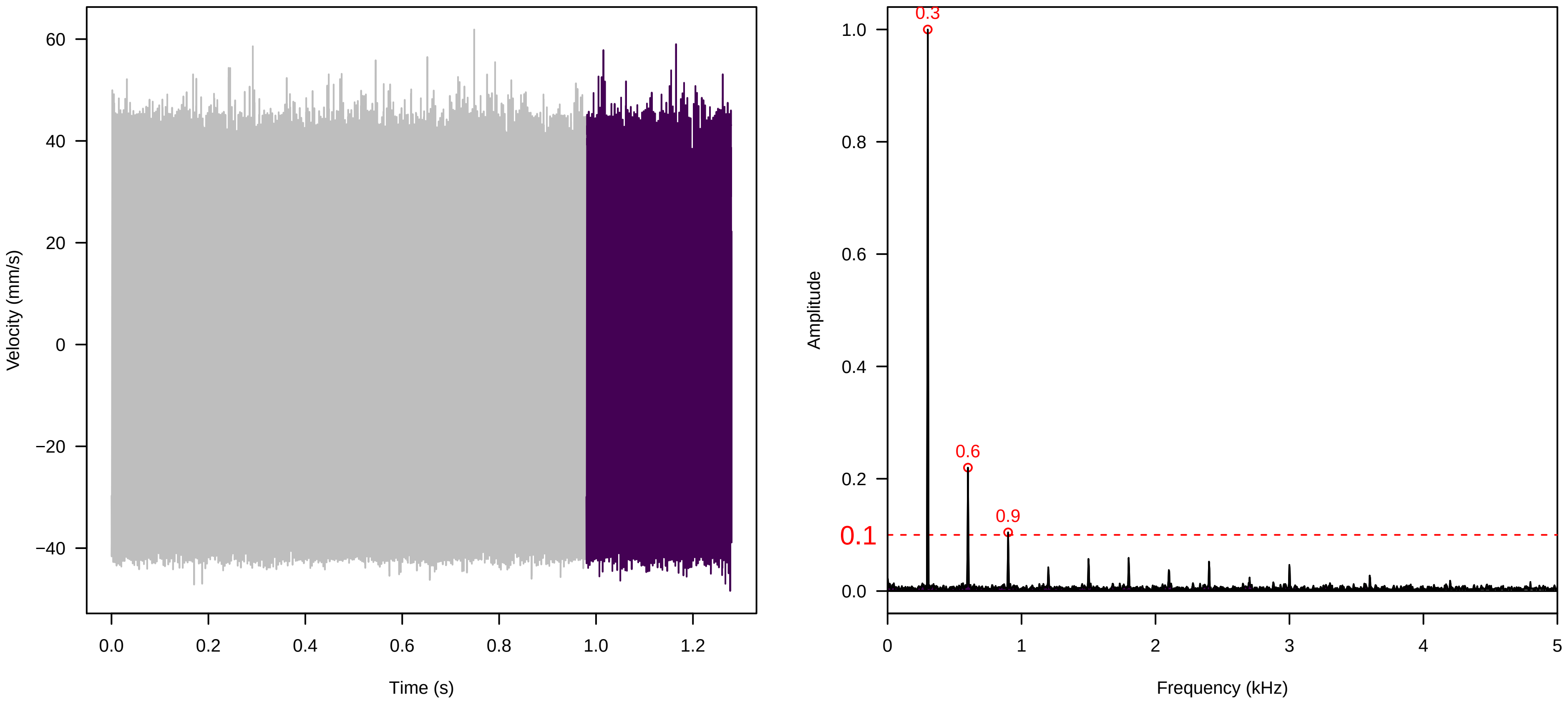


Vel. = 0.028 ; Str. = Receptacle ; Axis = x ; Fl. accession = 10−s−77−12

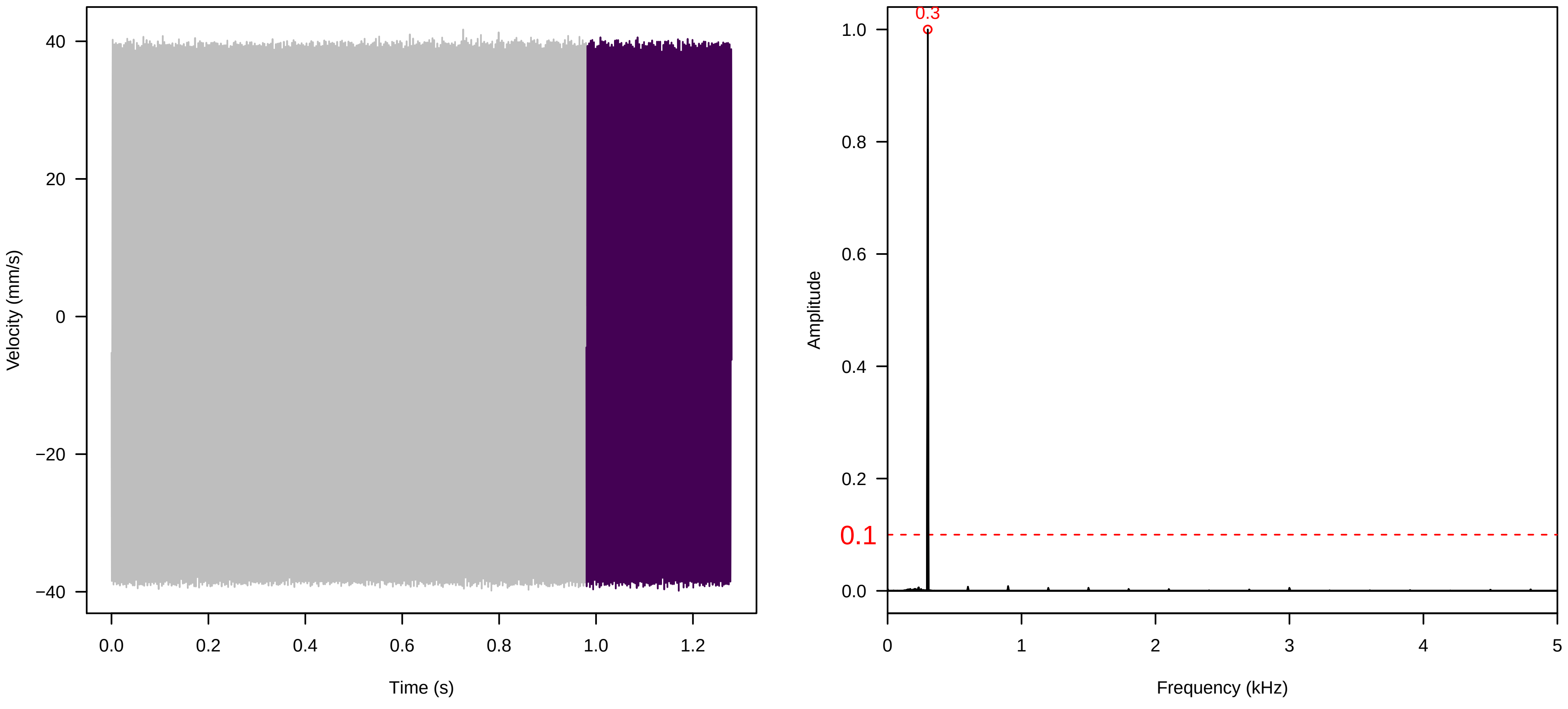

**Vel. = 0.057 ; Str. = Corolla ; Axis = x ; Fl. accession = 10−s−77−12**

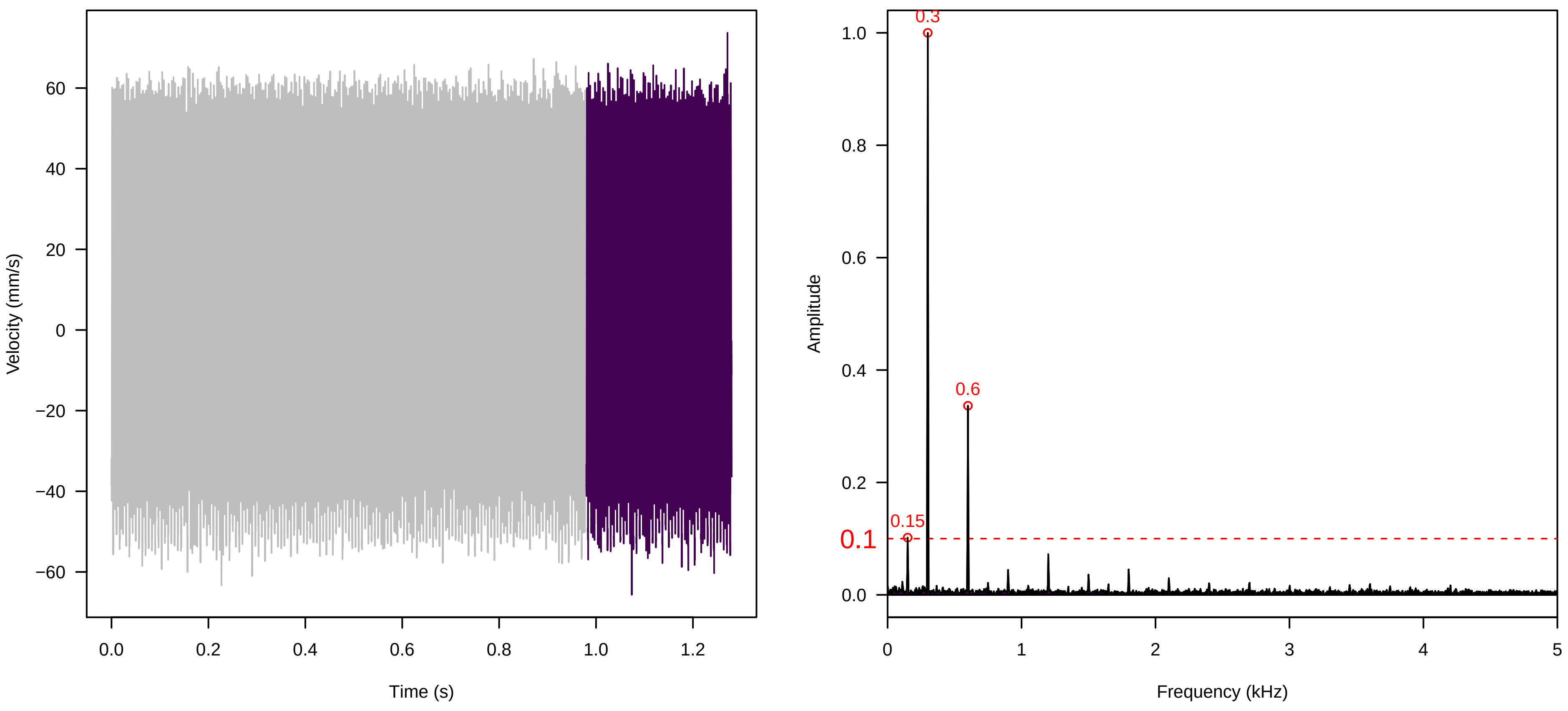


**Vel. = 0.057 ; Str. = Receptacle ; Axis = x ; Fl. accession = 10−s−77−12**

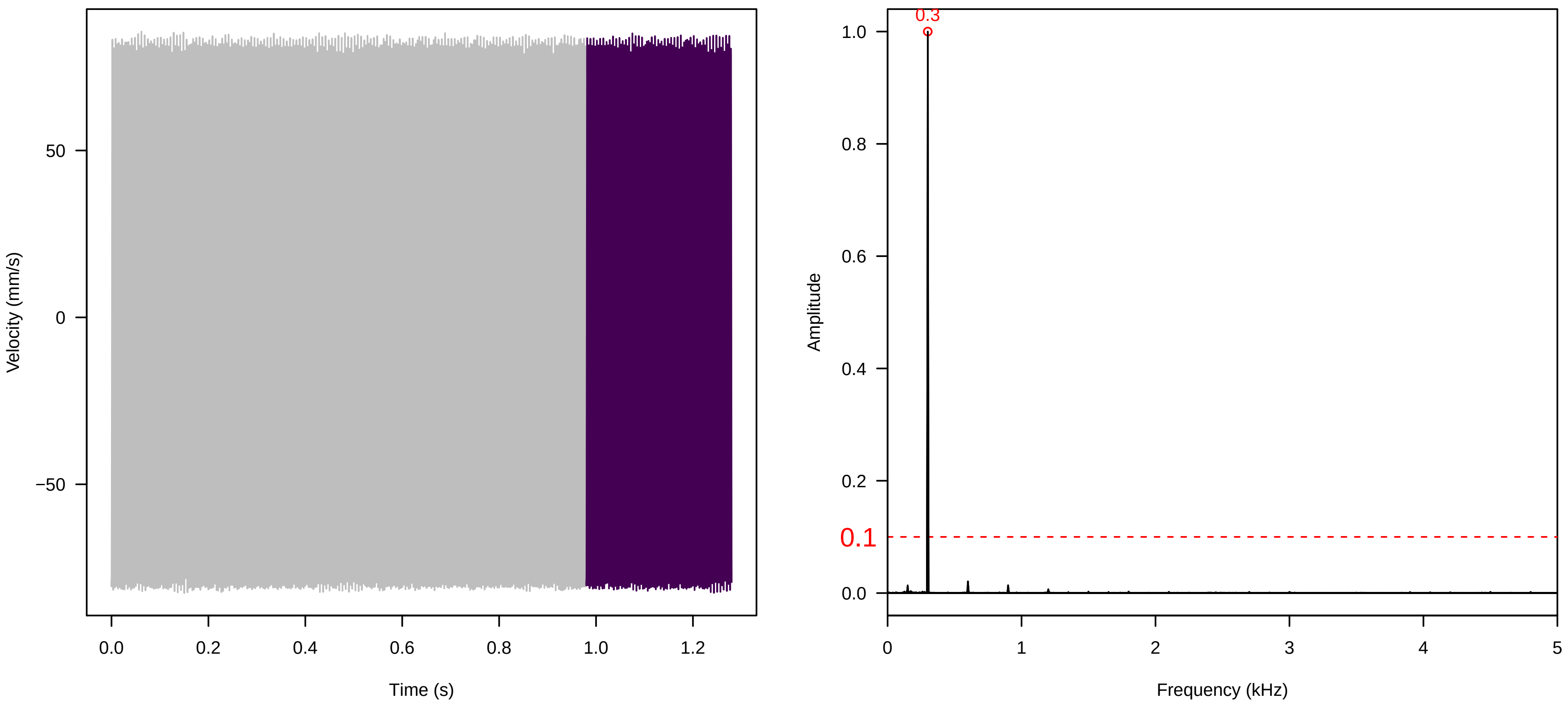


**Vel. = 0.057 ; Str. = FA ; Axis = x ; Fl. accession = 10−s−77−12**

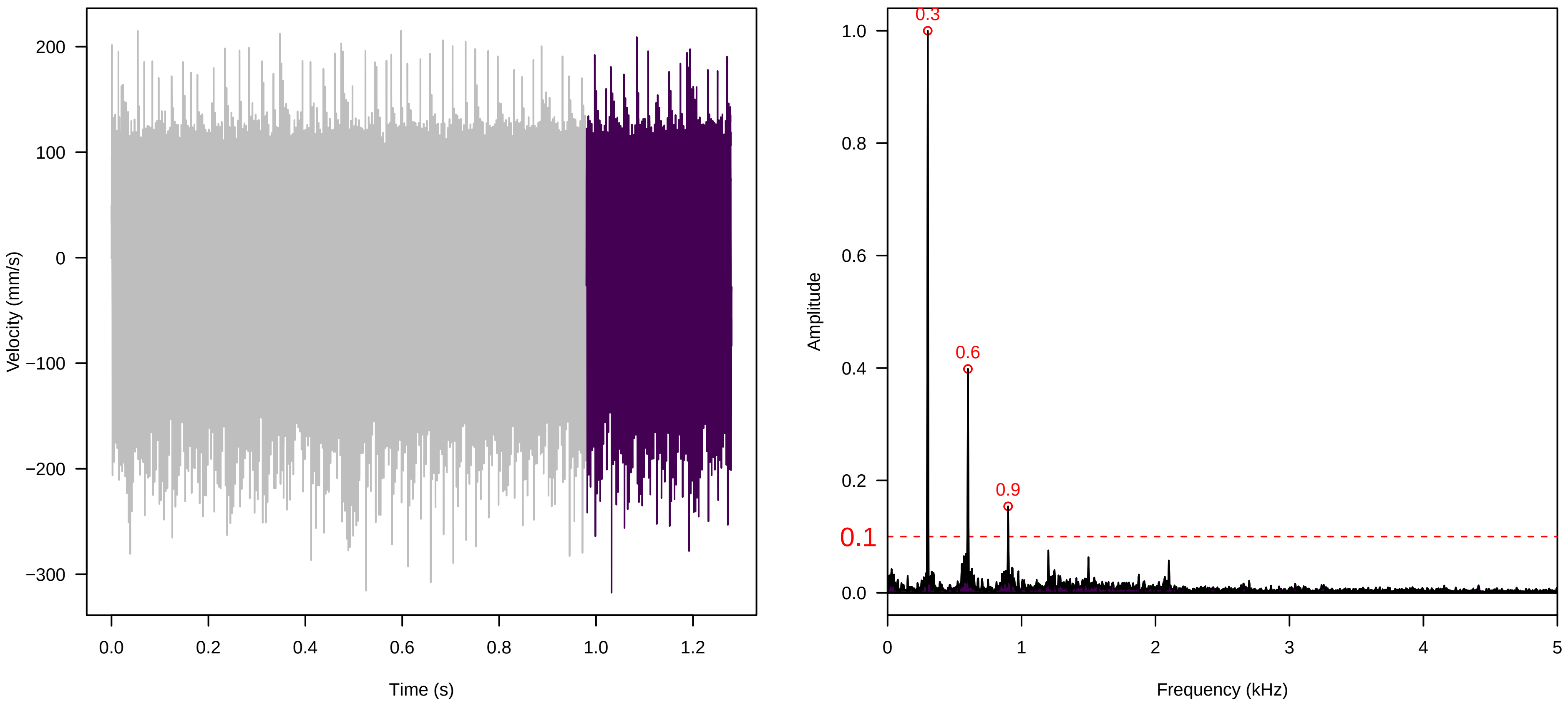

**Vel. =  0.057 ; Str. =  Receptacle ; Axis =  x ; Fl. accession = 10−s−77−12**

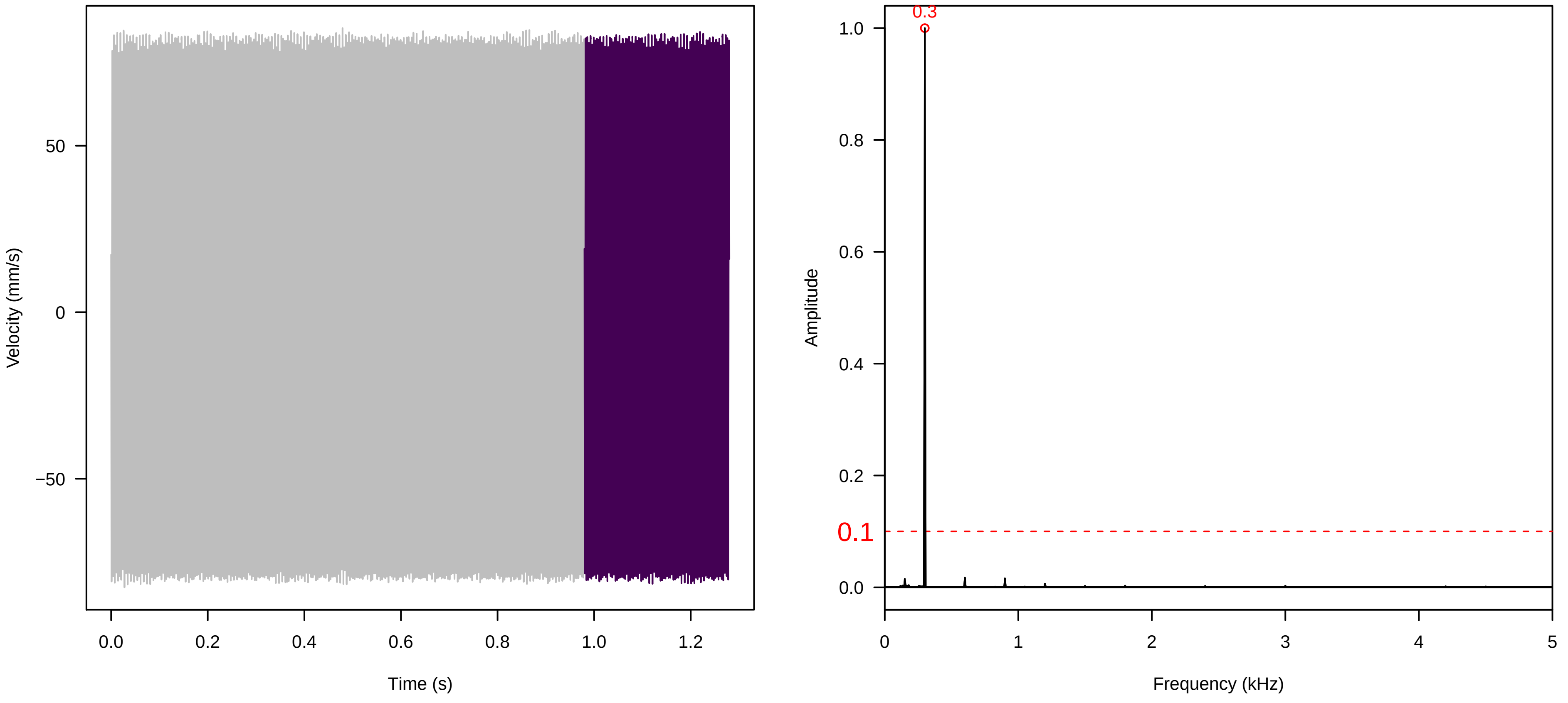


**Vel. =  0.057 ; Str. =  PA ; Axis =  x ; Fl. accession = 10−s−77−12**

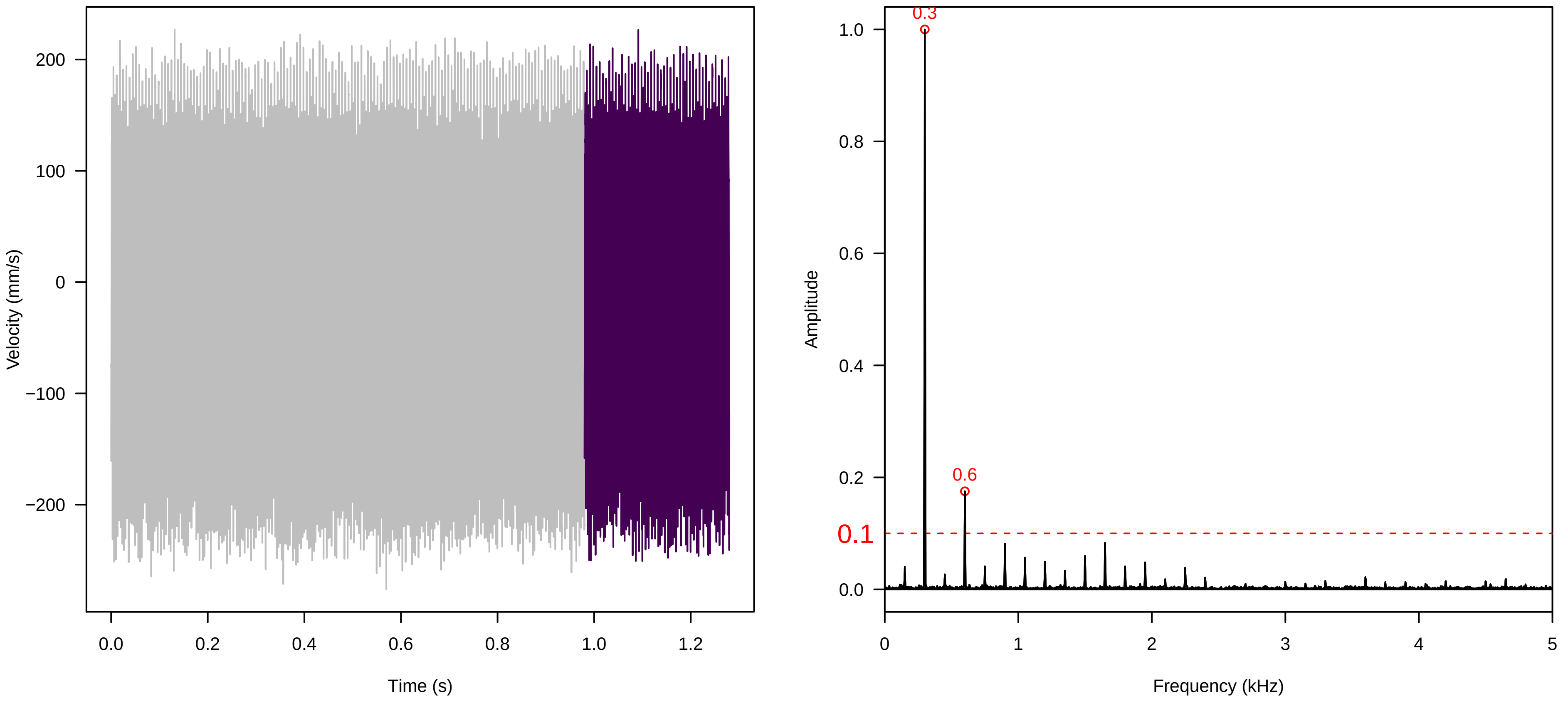


**Vel. =  0.057 ; Str. =  Receptacle ; Axis =  x ; Fl. accession = 10−s−77−12**

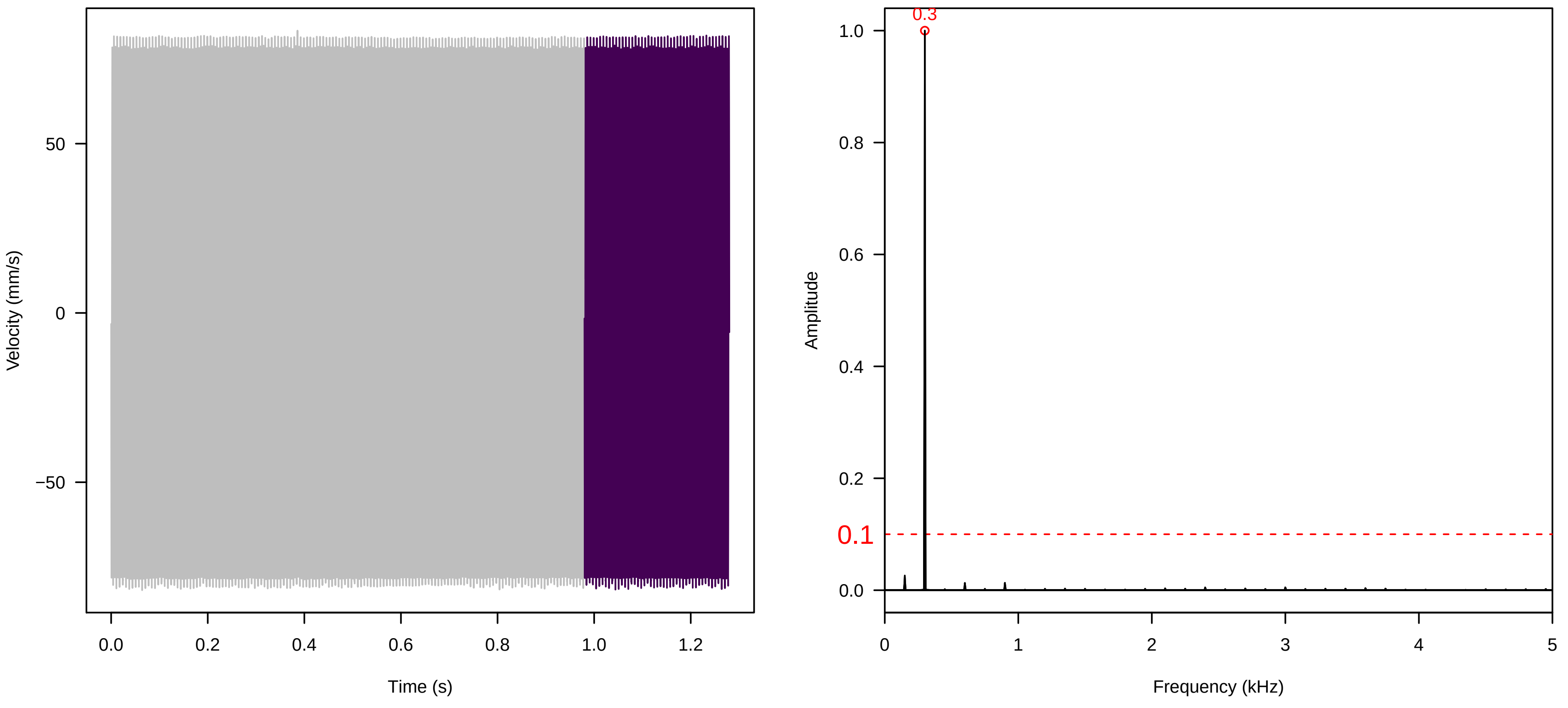

**Vel. = 0.057 ; Str. = Corolla ; Axis = z ; Fl. accession = 10−s−77−12**

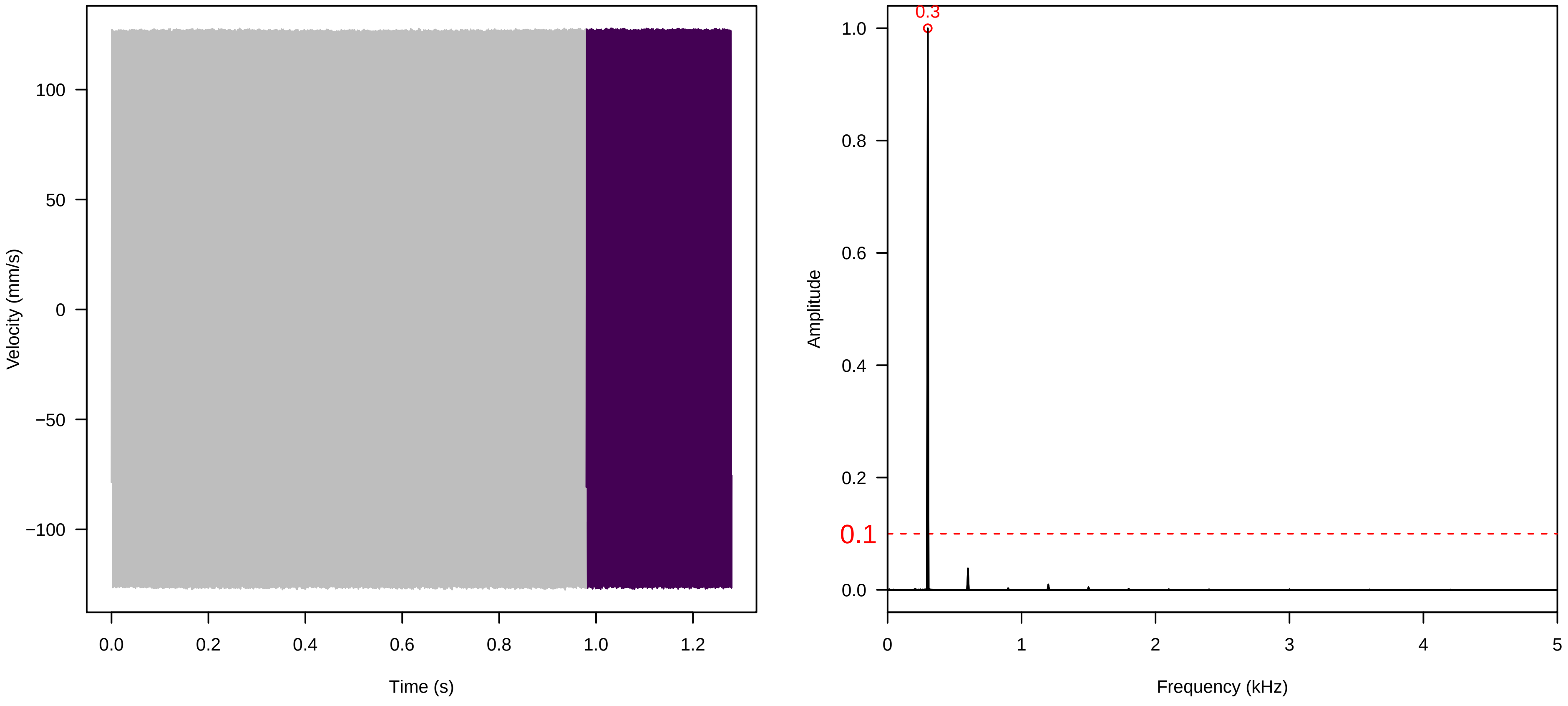


**Vel. = 0.057 ; Str. = Receptacle ; Axis = z ; Fl. accession = 10−s−77−12**

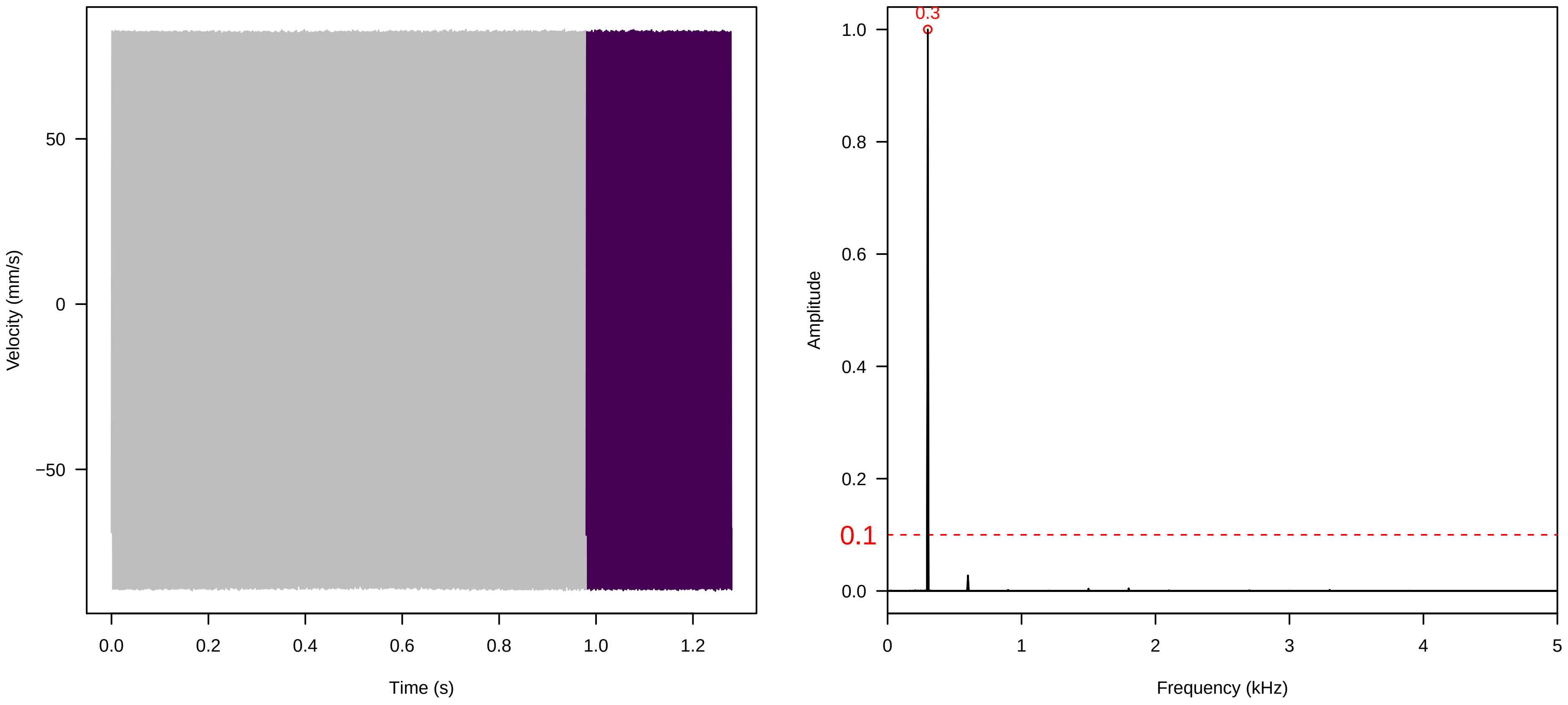


**Vel. = 0.057 ; Str. = FA ; Axis = z ; Fl. accession = 10−s−77−12**

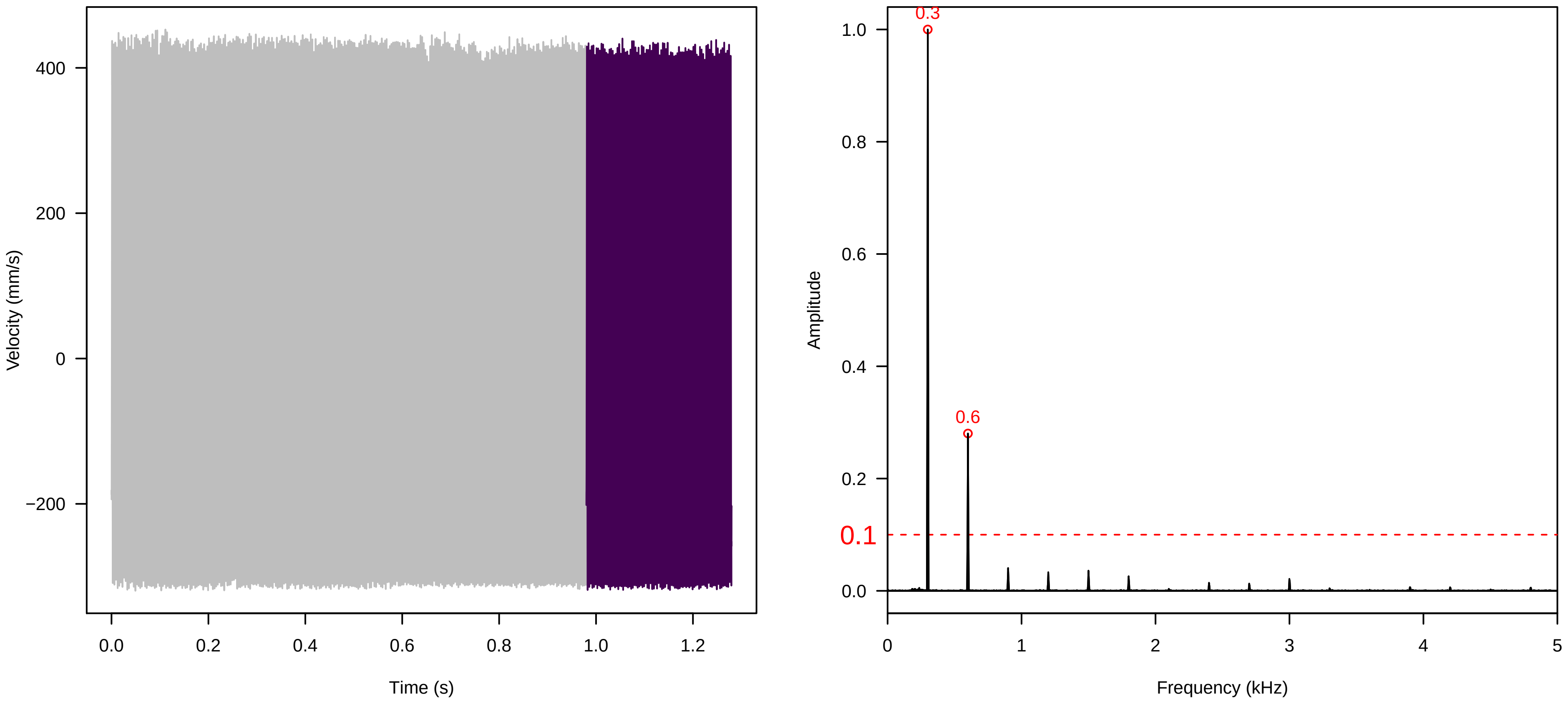

**Vel. = 0.057 ; Str. = Receptacle ; Axis = z ; Fl. accession = 10−s−77−12**

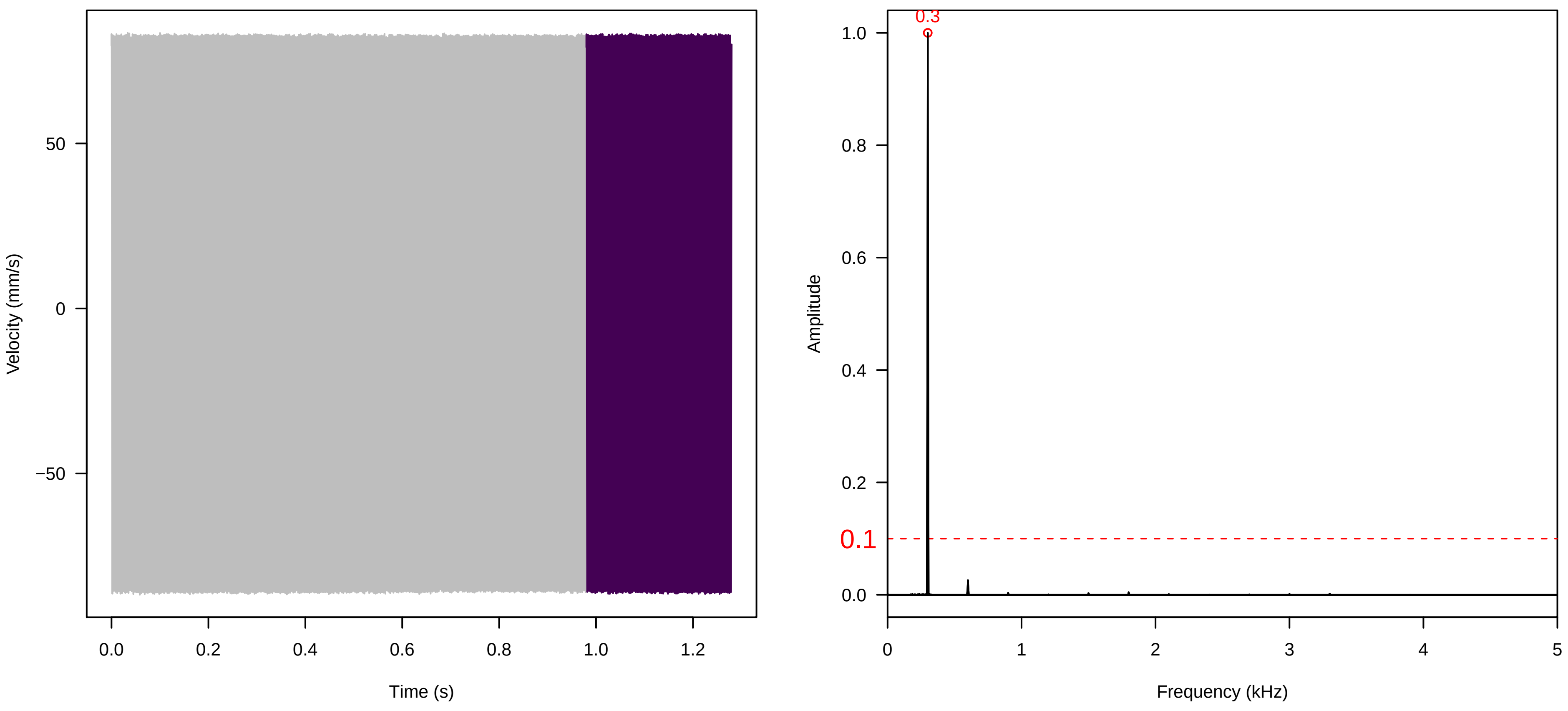


**Vel. = 0.057 ; Str. = PA ; Axis = z ; Fl. accession = 10−s−77−12**

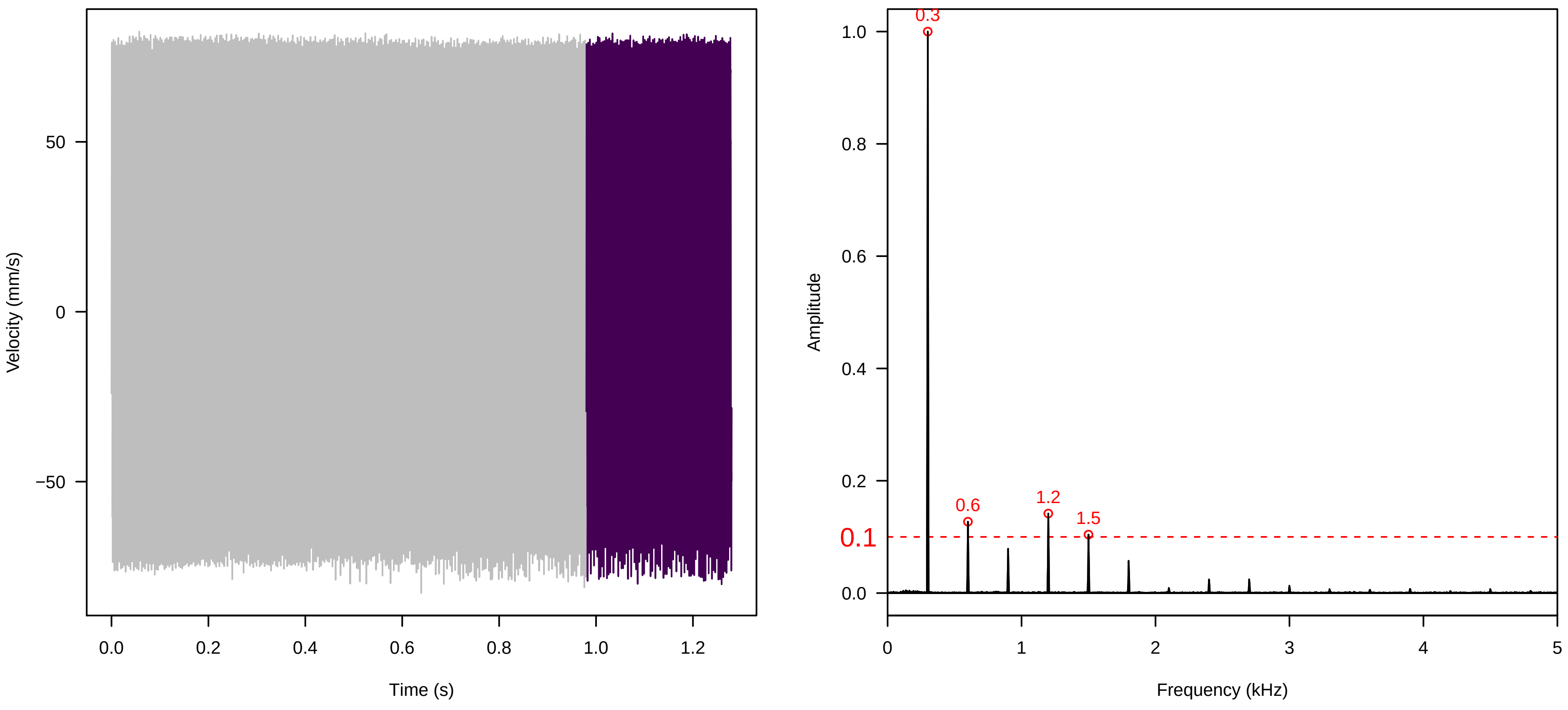


**Vel. = 0.057 ; Str. = Receptacle ; Axis = z ; Fl. accession = 10−s−77−12**

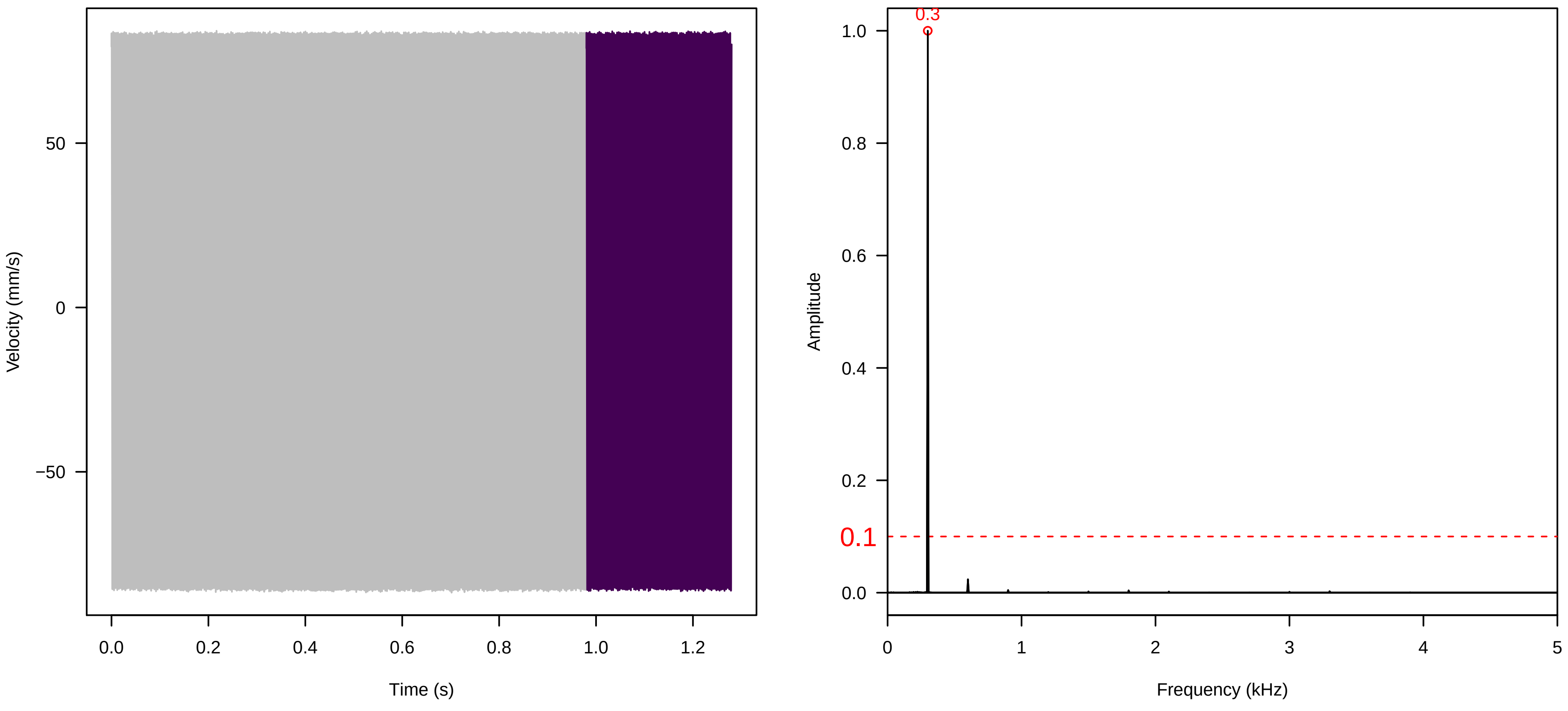

**Vel. = 0.028 ; Str. = PA ; Axis = z ; Fl. accession = 10−s−77−12**

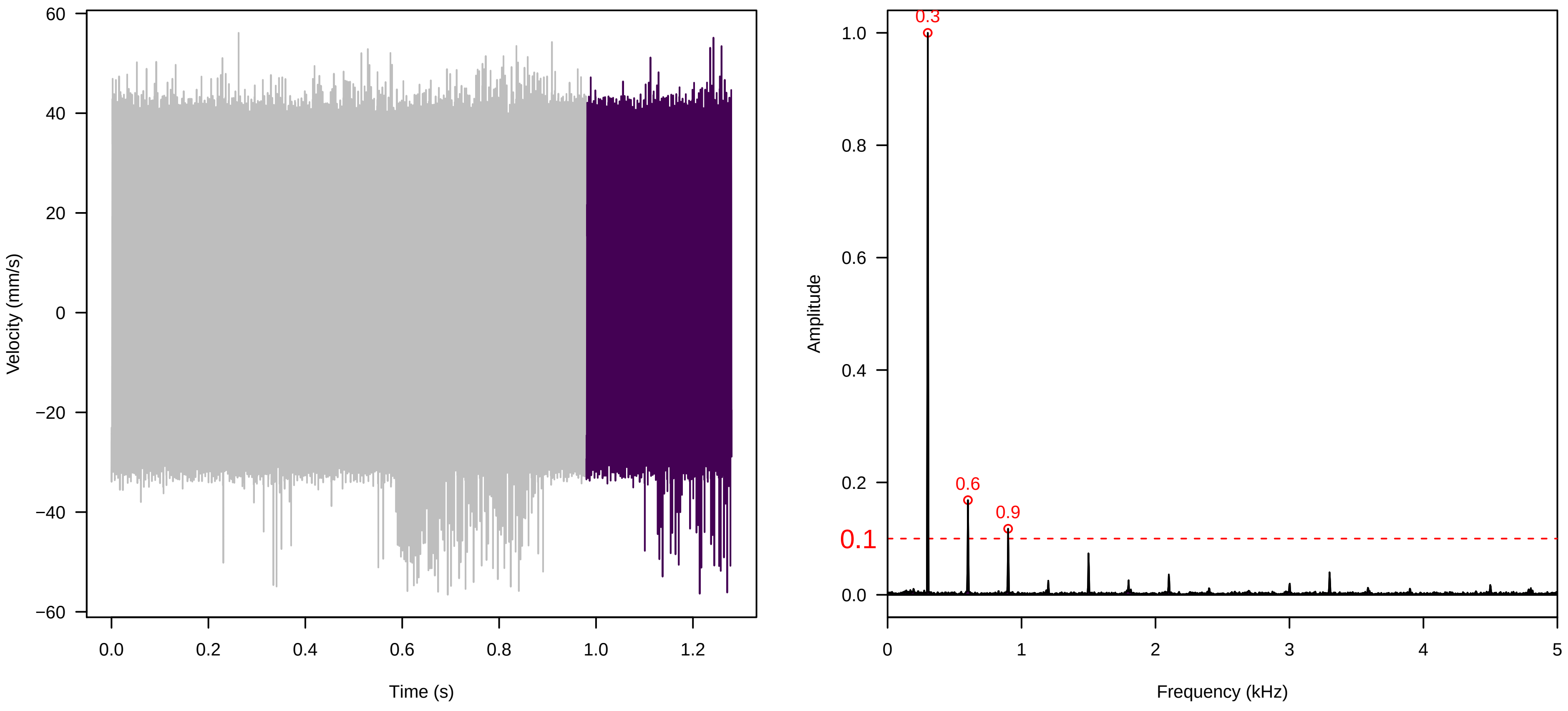


**Vel. = 0.028 ; Str. = Receptacle ; Axis = z ; Fl. accession = 10−s−77−12**

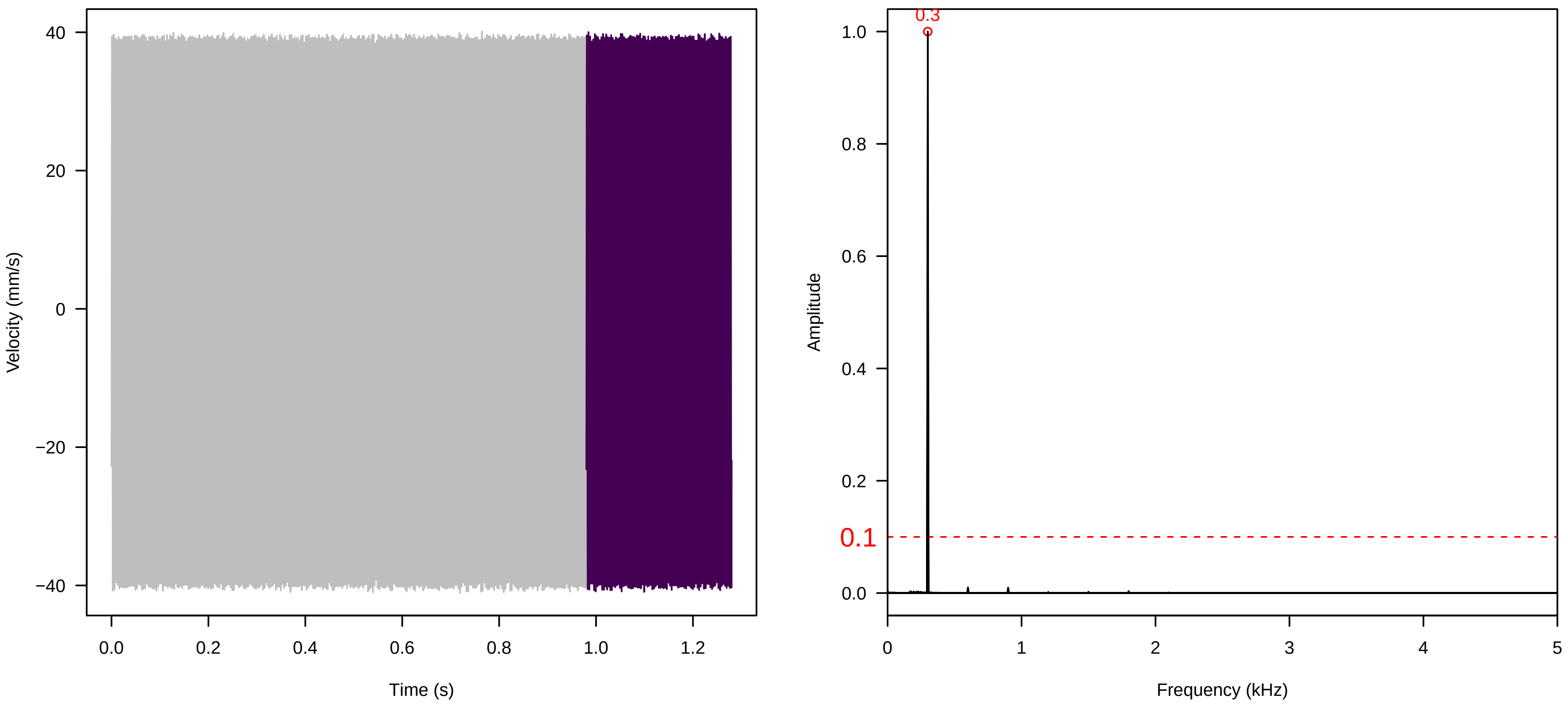


**Vel. = 0.028 ; Str. = FA ; Axis = z ; Fl. accession = 10−s−77−12**

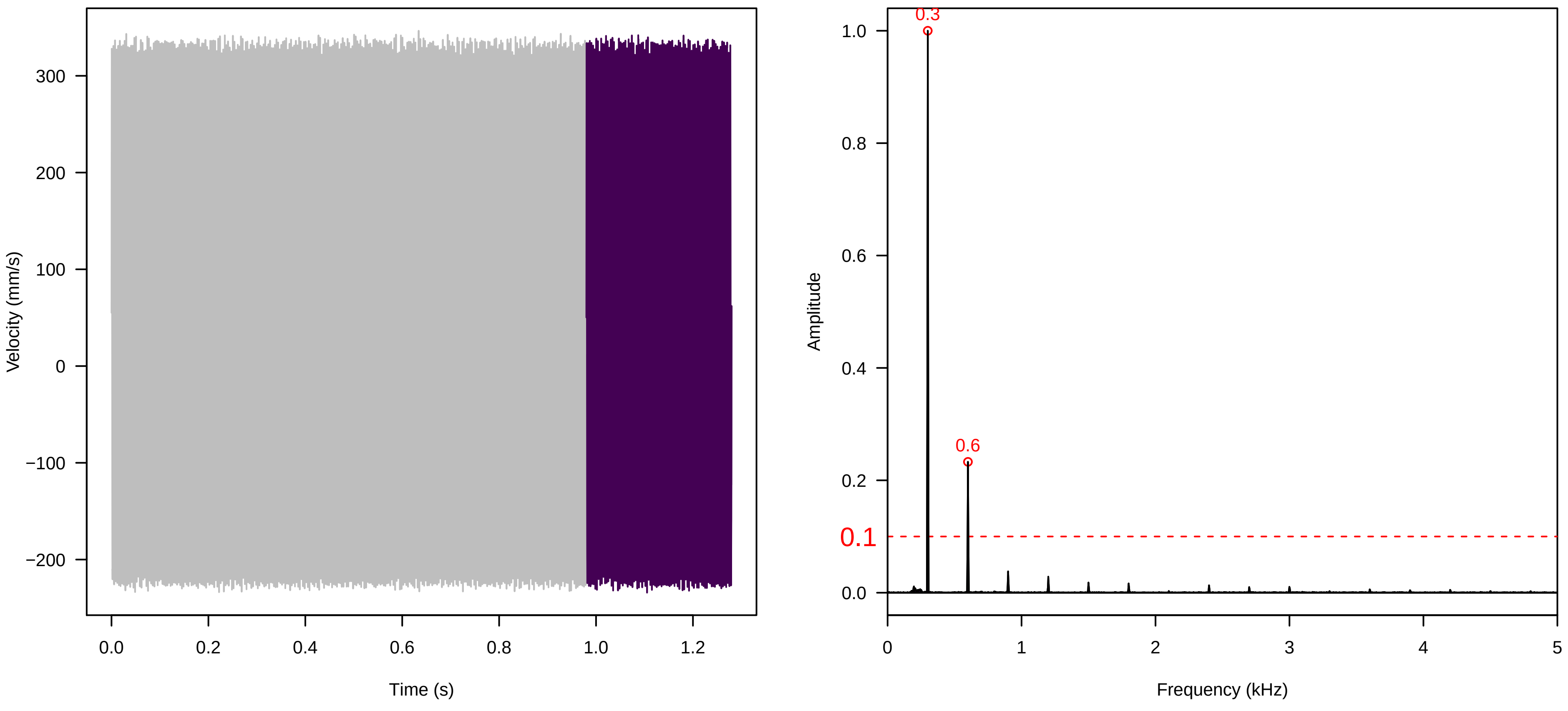

**Vel. = 0.028 ; Str. = Receptacle ; Axis = z ; Fl. accession = 10−s−77−12**

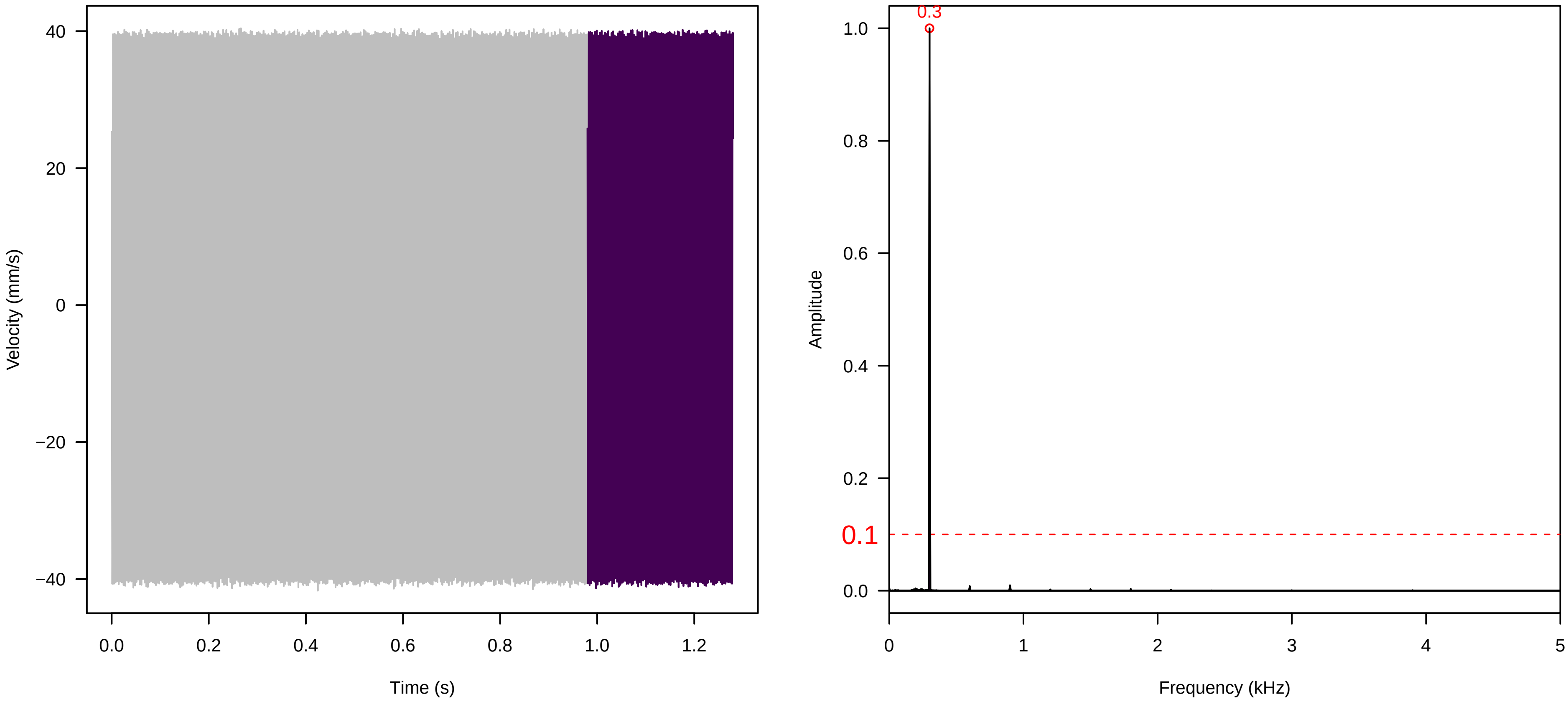


**Vel. = 0.028 ; Str. = Corolla ; Axis = z ; Fl. accession = 10−s−77−12**

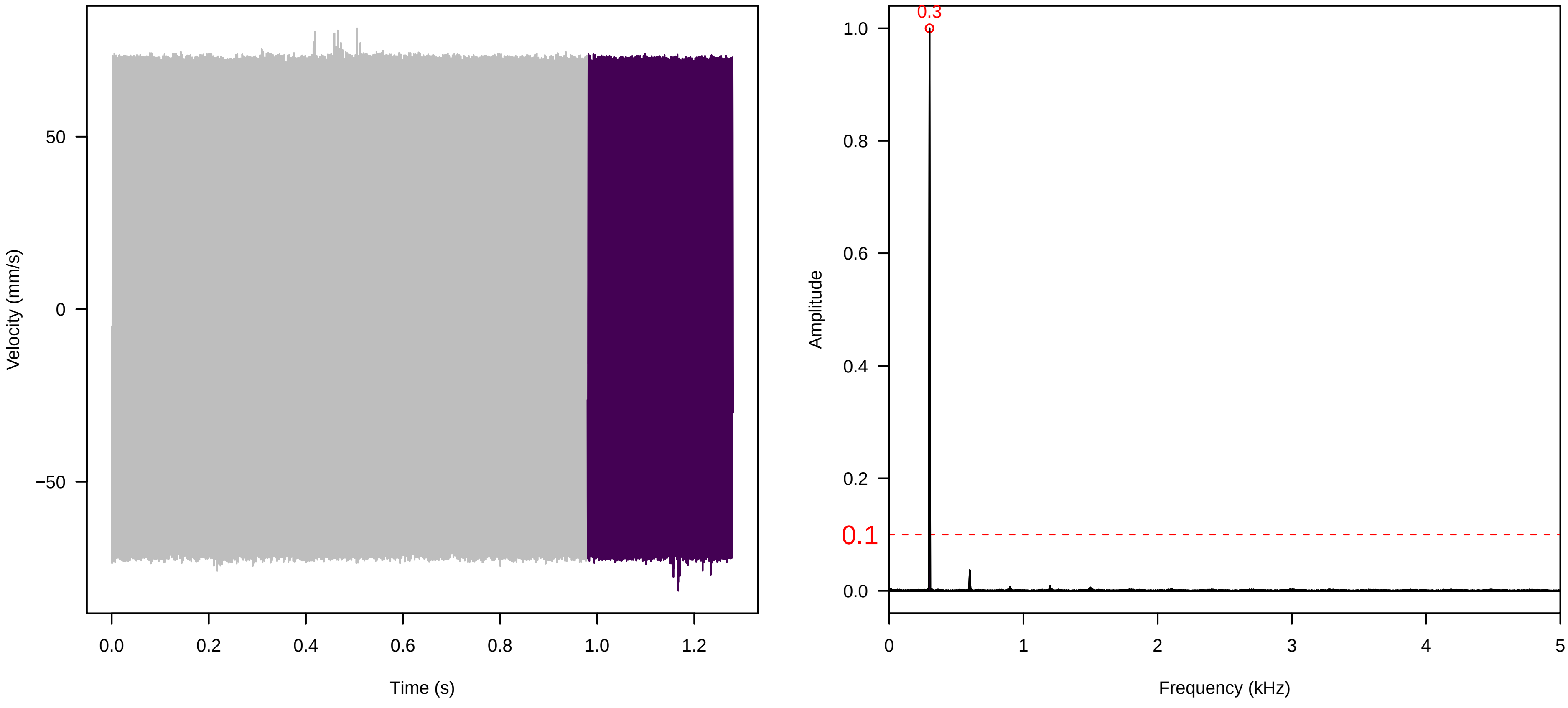


**Vel. = 0.028 ; Str. = Receptacle ; Axis = z ; Fl. accession = 10−s−77−12**

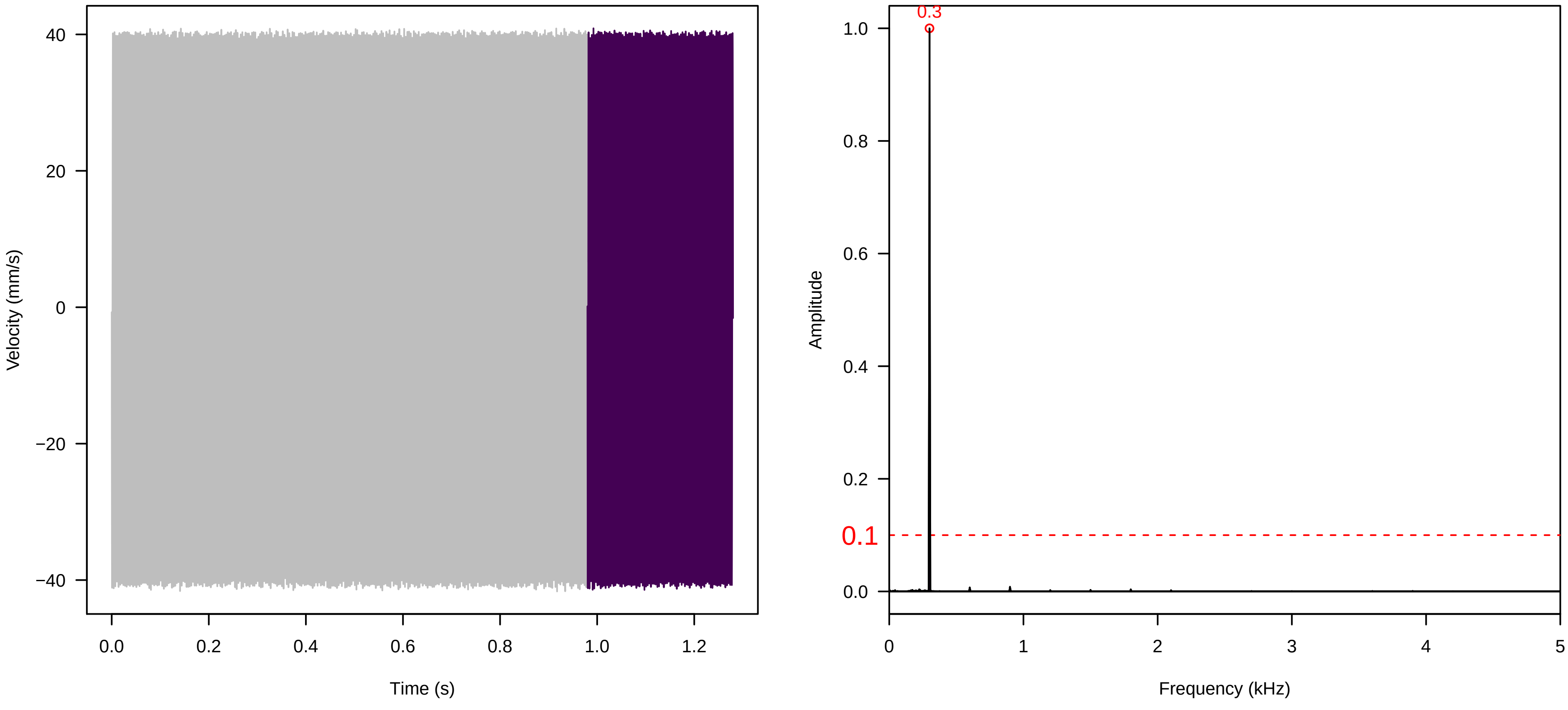

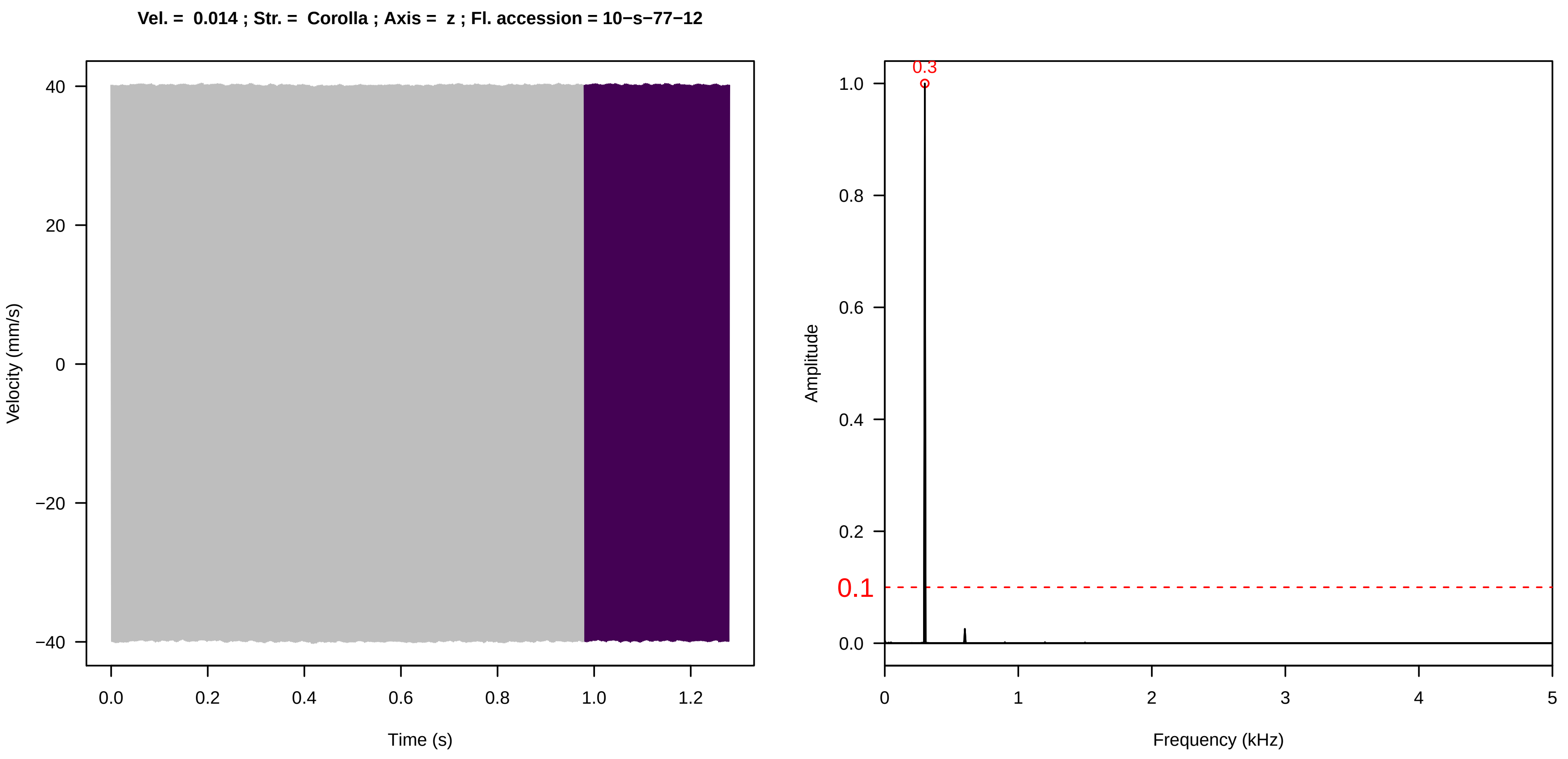
Vel. = 0.014 ; Str. = Corolla ; Axis = z ; Fl. accession = 10−s−77−12
Velocity (mm/s)
Time (s)
Amplitude
Frequency (kHz)
0.3
0.1

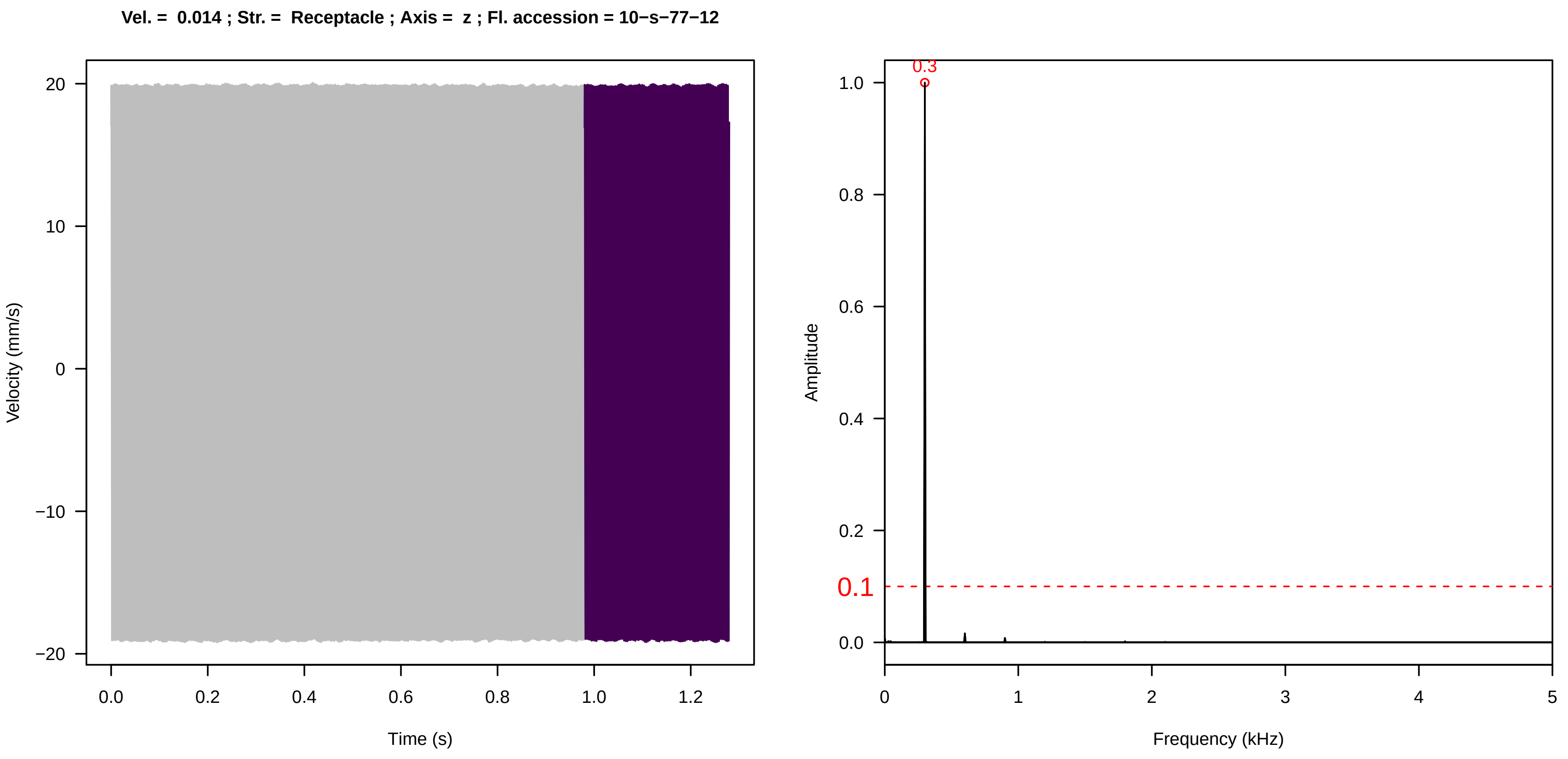
Vel. = 0.014 ; Str. = Receptacle ; Axis = z ; Fl. accession = 10−s−77−12
Velocity (mm/s)
Time (s)
Amplitude
Frequency (kHz)
0.3
0.1

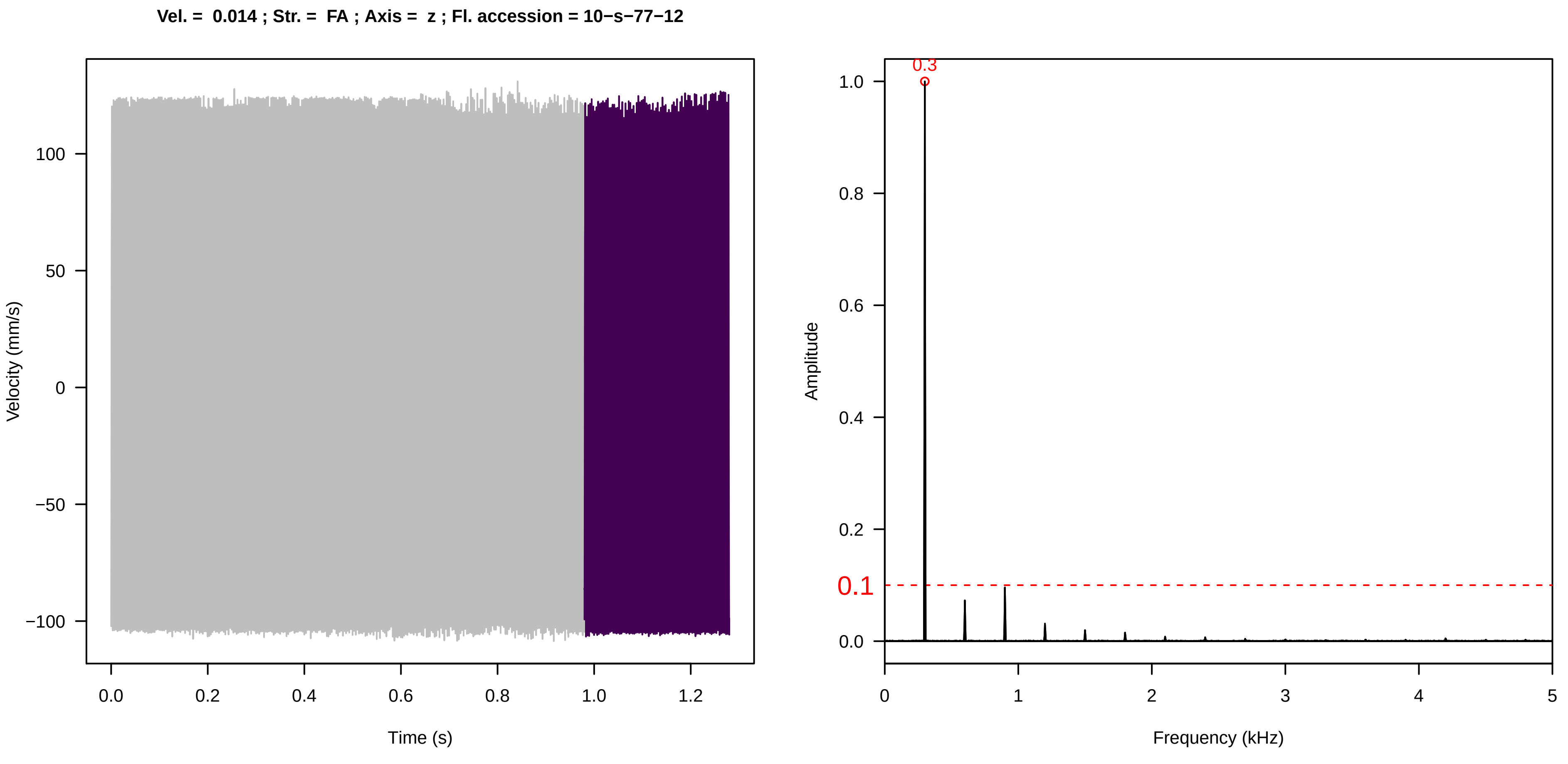
Vel. = 0.014 ; Str. = FA ; Axis = z ; Fl. accession = 10−s−77−12
Velocity (mm/s)
Time (s)
Amplitude
Frequency (kHz)
0.3
0.1

**Vel. = 0.014 ; Str. = Receptacle ; Axis = z ; Fl. accession = 10−s−77−12**

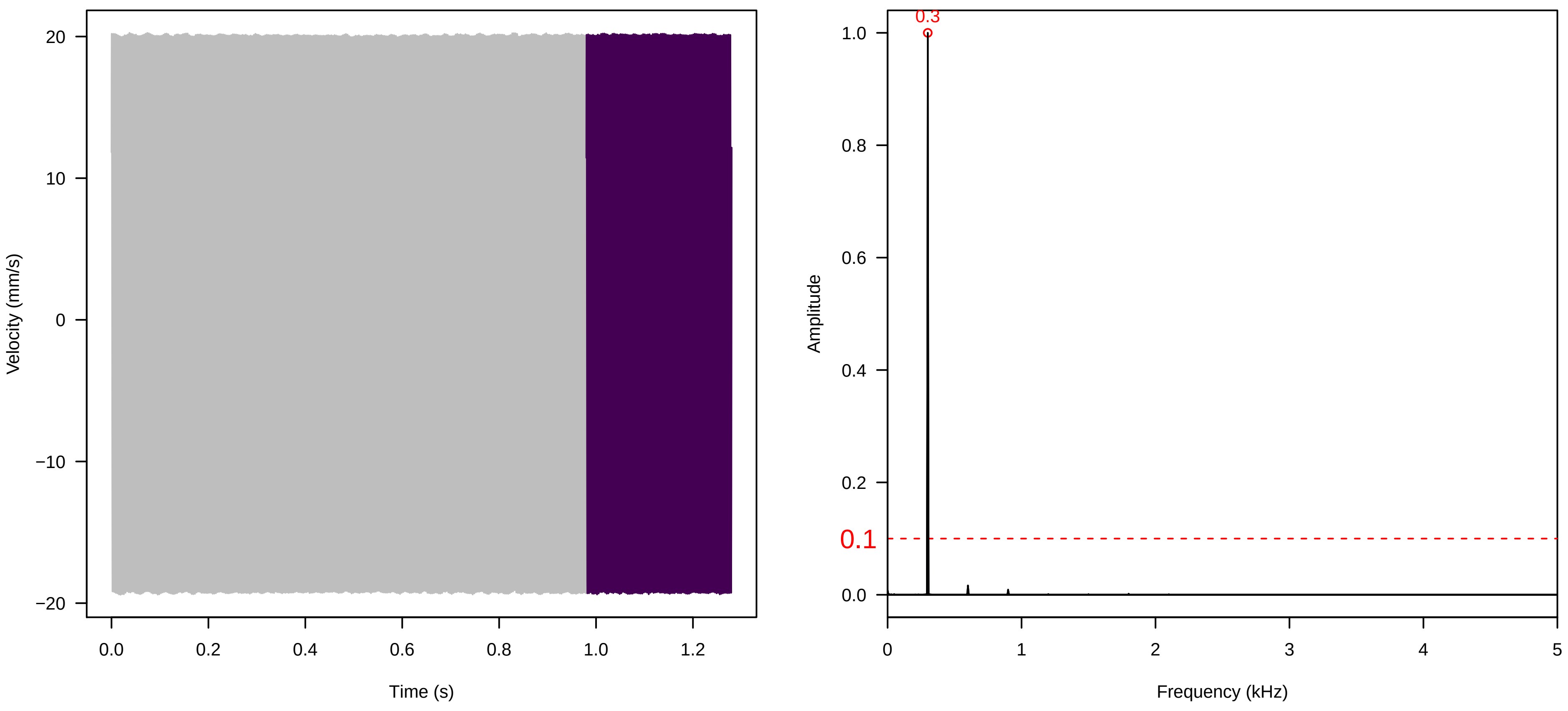


**Vel. = 0.014 ; Str. = PA ; Axis = z ; Fl. accession = 10−s−77−12**

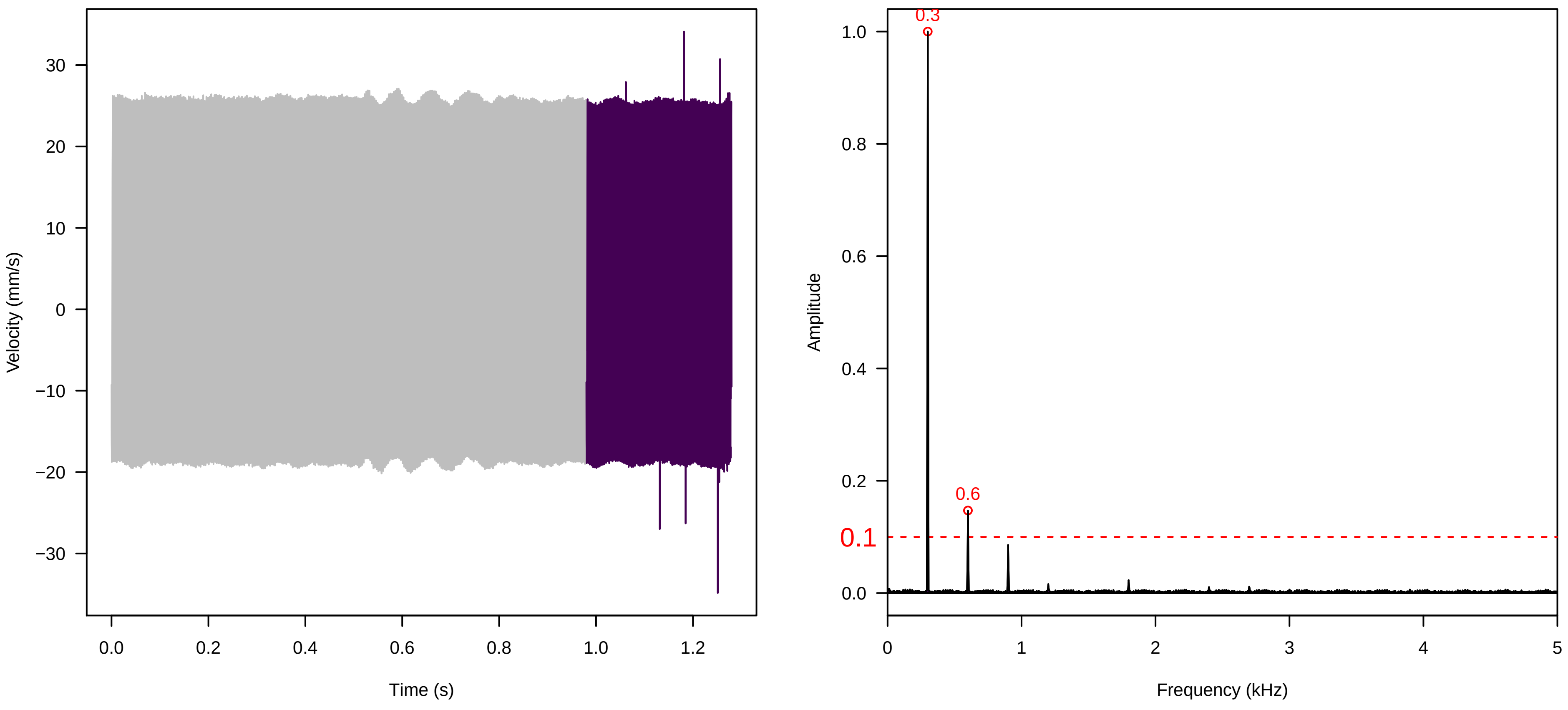


**Vel. = 0.014 ; Str. = Receptacle ; Axis = z ; Fl. accession = 10−s−77−12**

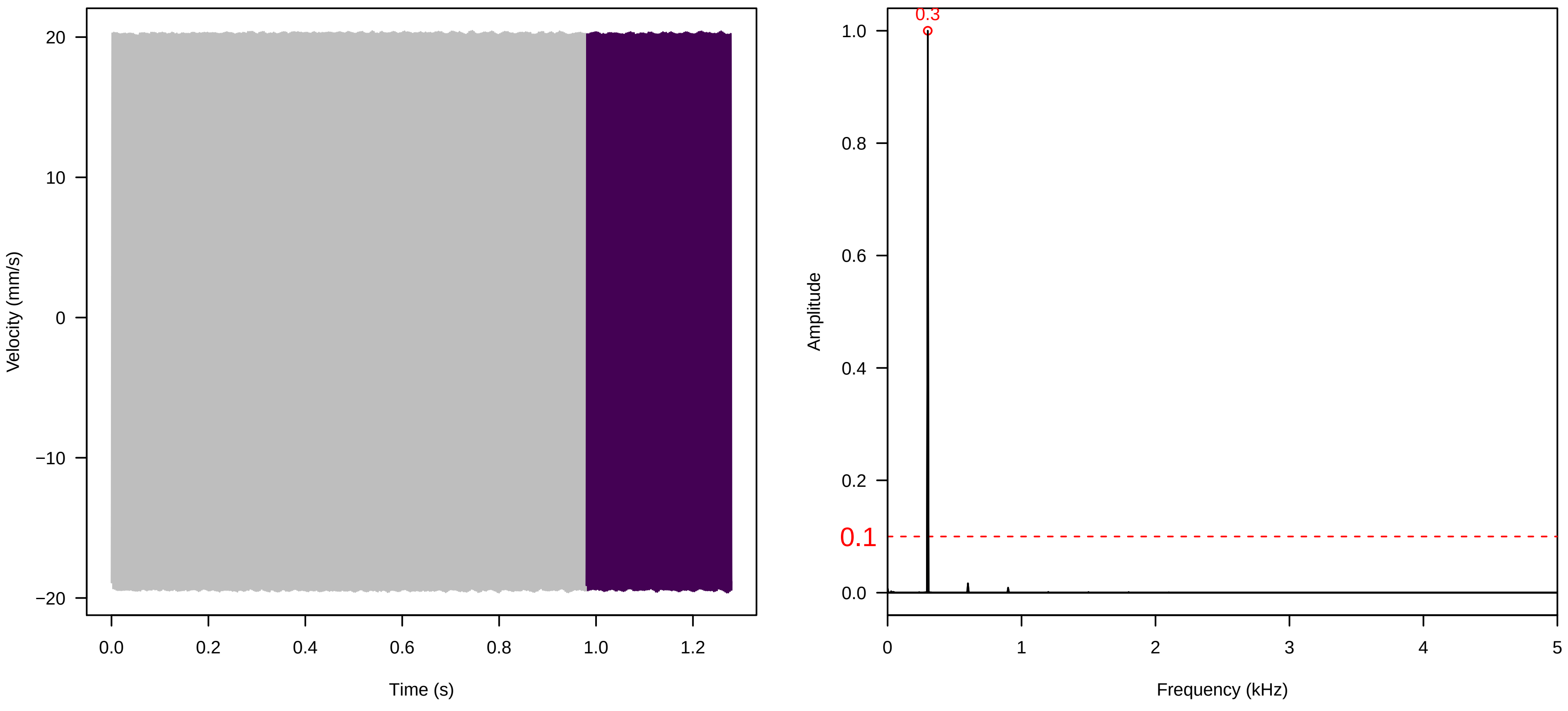

**Vel. =  0.014 ; Str. =  Corolla ; Axis =  y ; Fl. accession = 10−s−77−12**

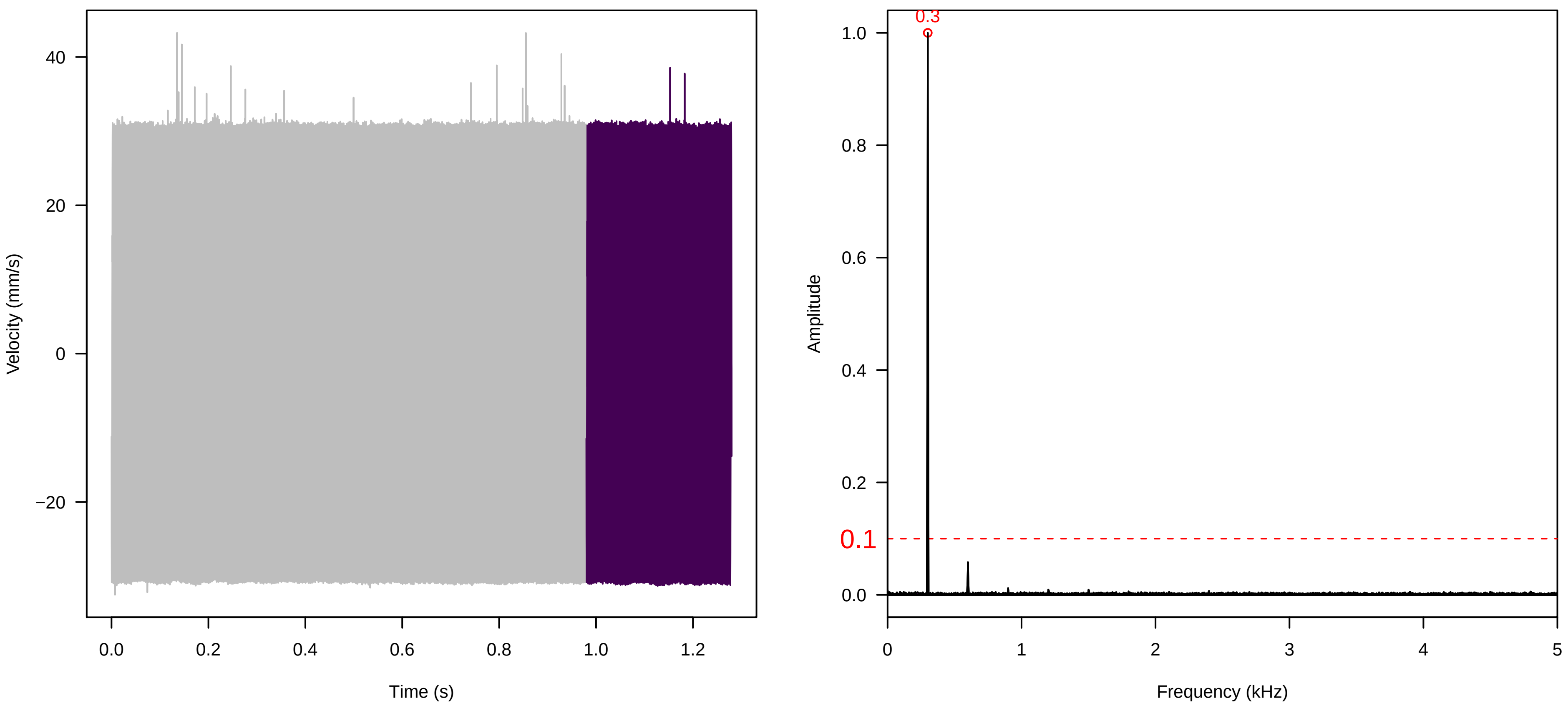


**Vel. =  0.014 ; Str. =  Receptacle ; Axis =  y ; Fl. accession = 10−s−77−12**

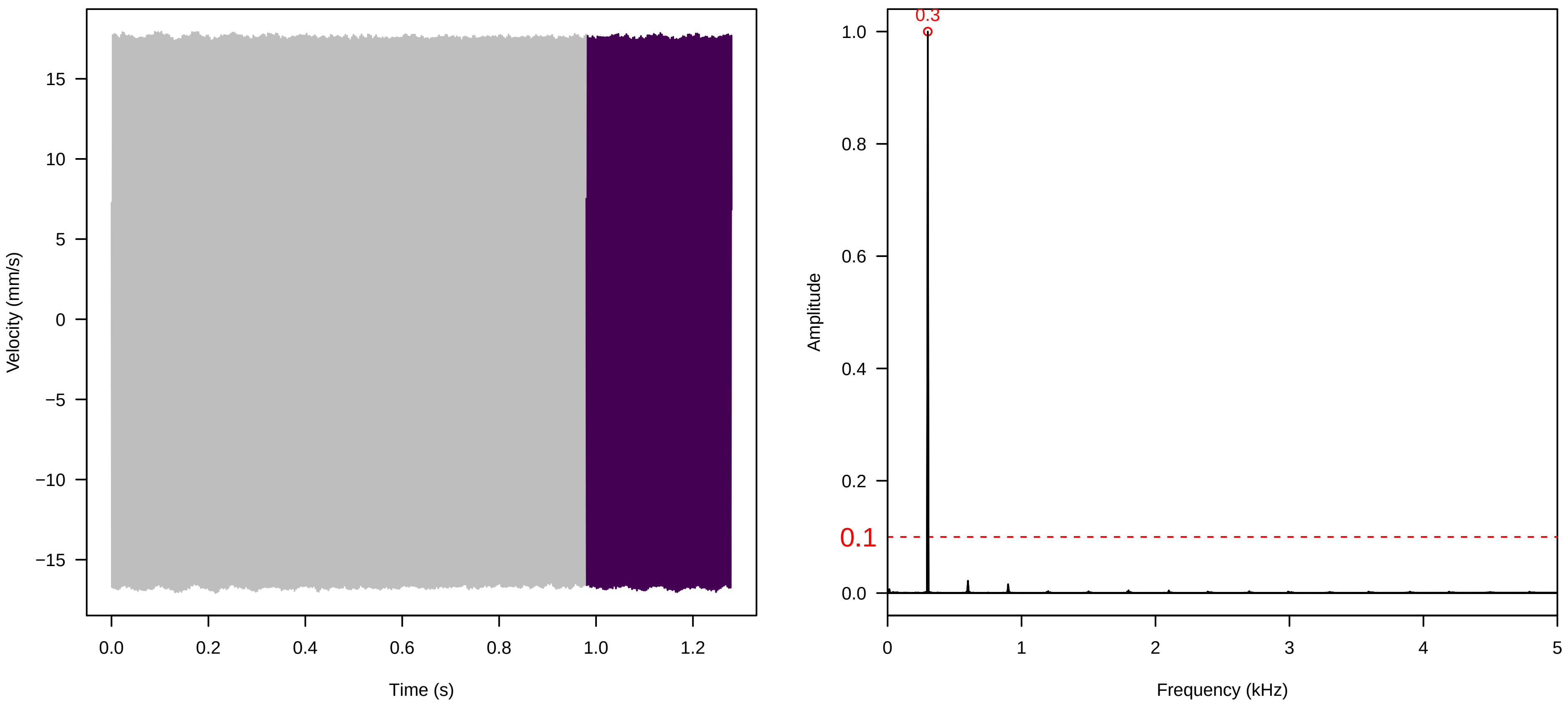


**Vel. =  0.014 ; Str. =  FA ; Axis =  y ; Fl. accession = 10−s−77−12**

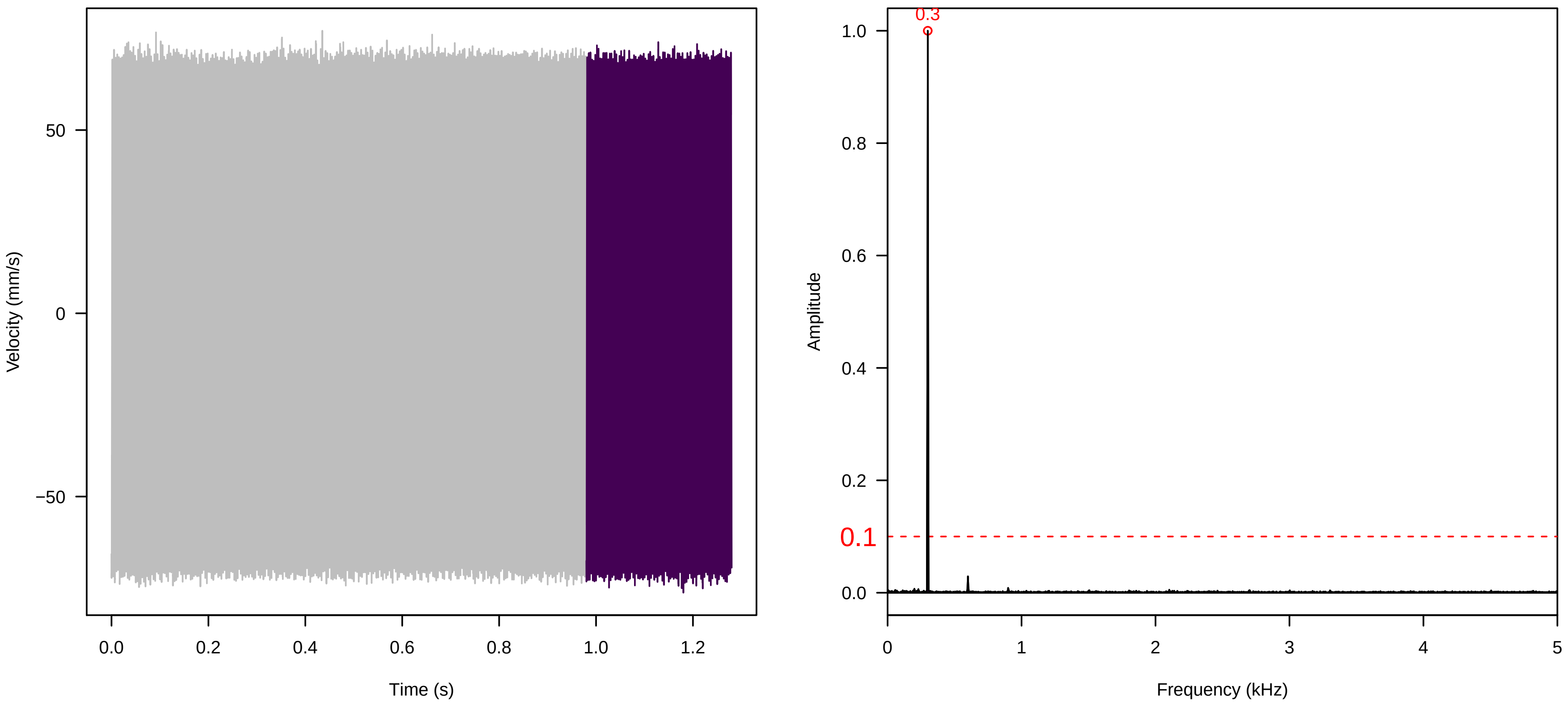

**Vel. = 0.014 ; Str. = Receptacle ; Axis = y ; Fl. accession = 10−s−77−12**

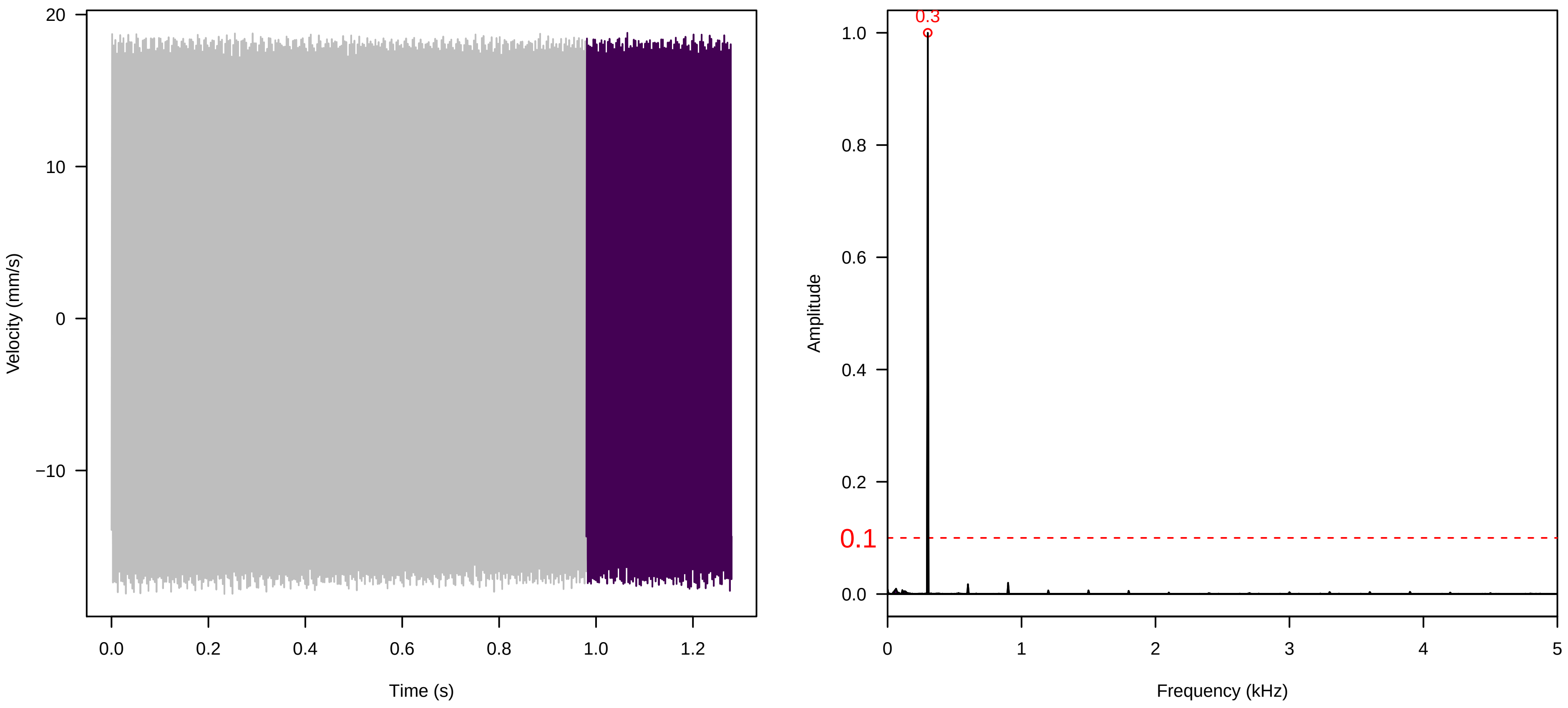


**Vel. = 0.014 ; Str. = PA ; Axis = y ; Fl. accession = 10−s−77−12**

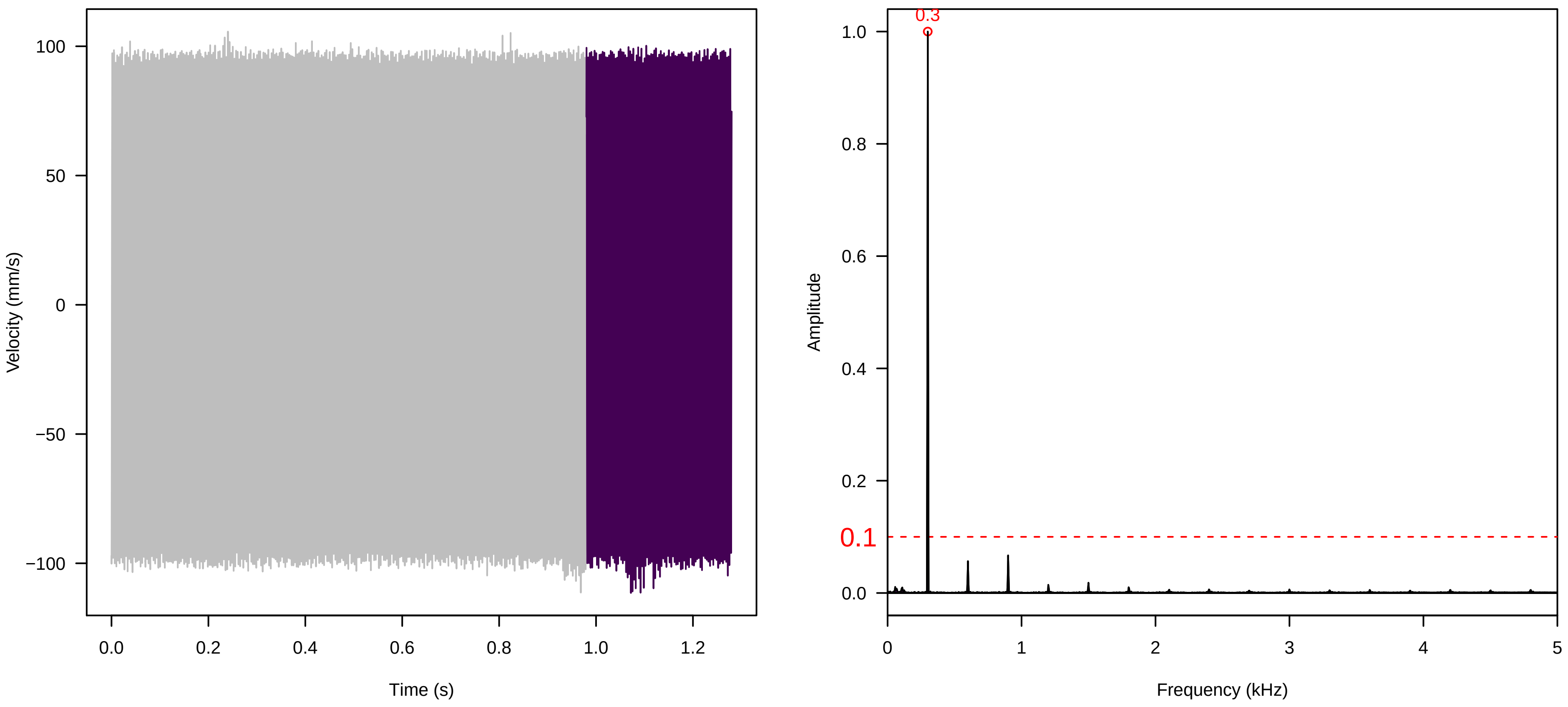


**Vel. = 0.014 ; Str. = Receptacle ; Axis = y ; Fl. accession = 10−s−77−12**

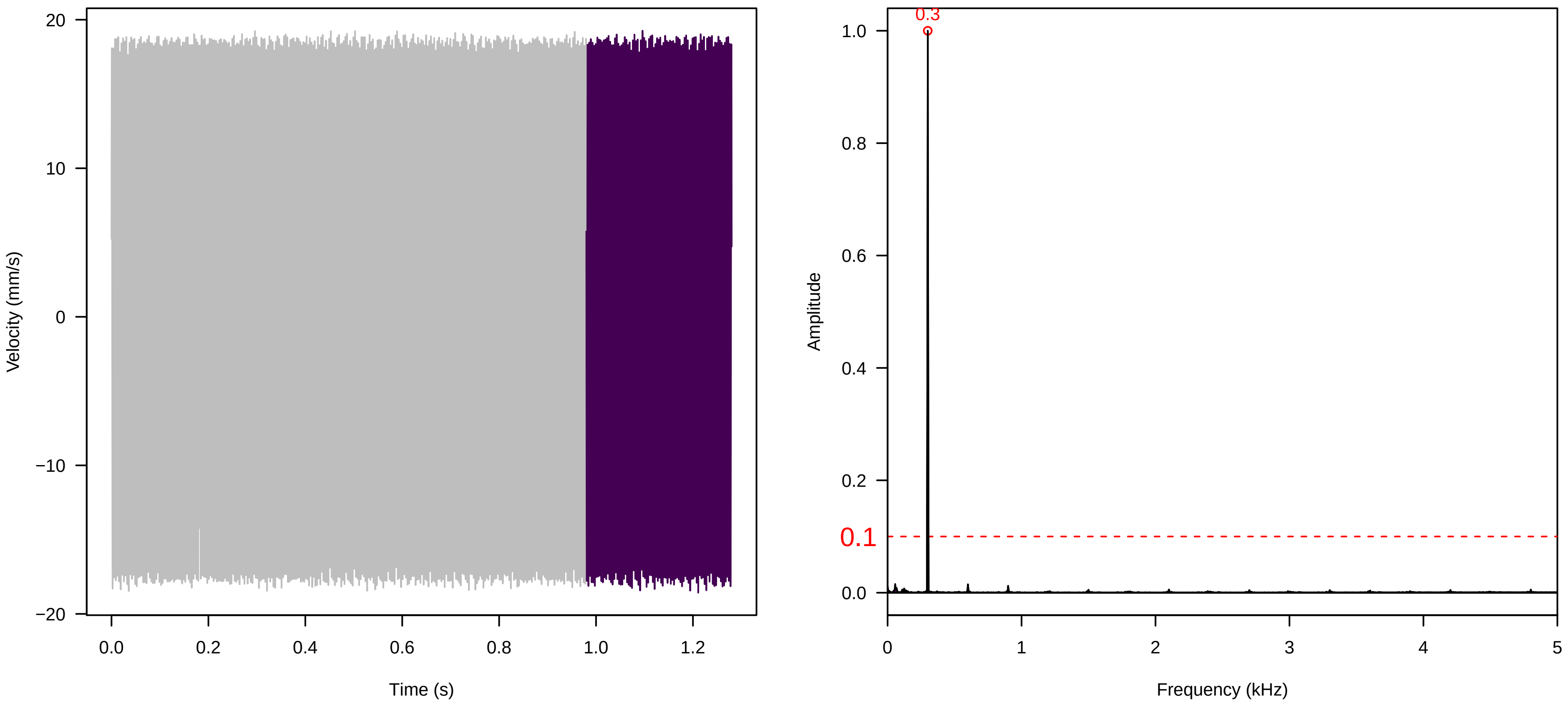

**Vel. = 0.028 ; Str. = PA ; Axis = y ; Fl. accession = 10−s−77−12**

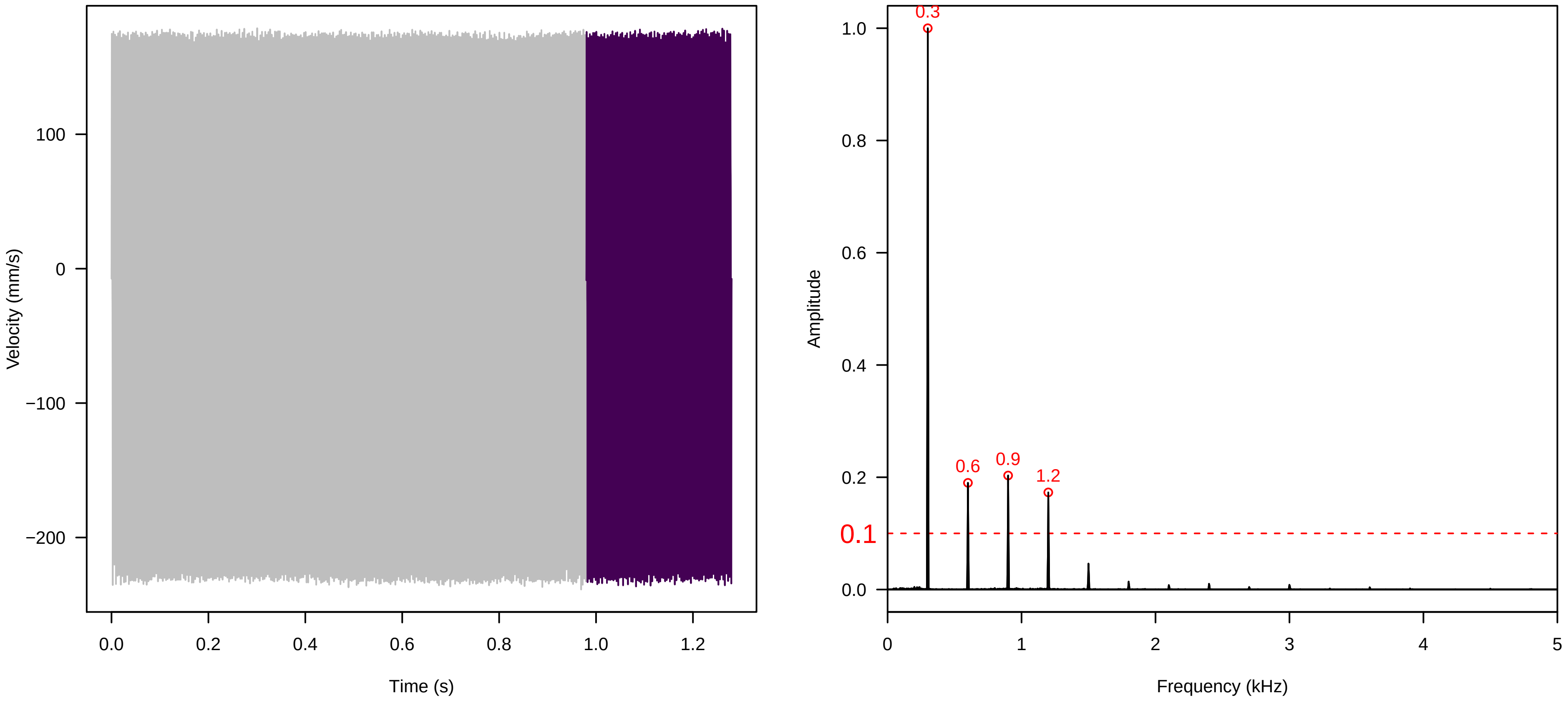


**Vel. = 0.028 ; Str. = Receptacle ; Axis = y ; Fl. accession = 10−s−77−12**

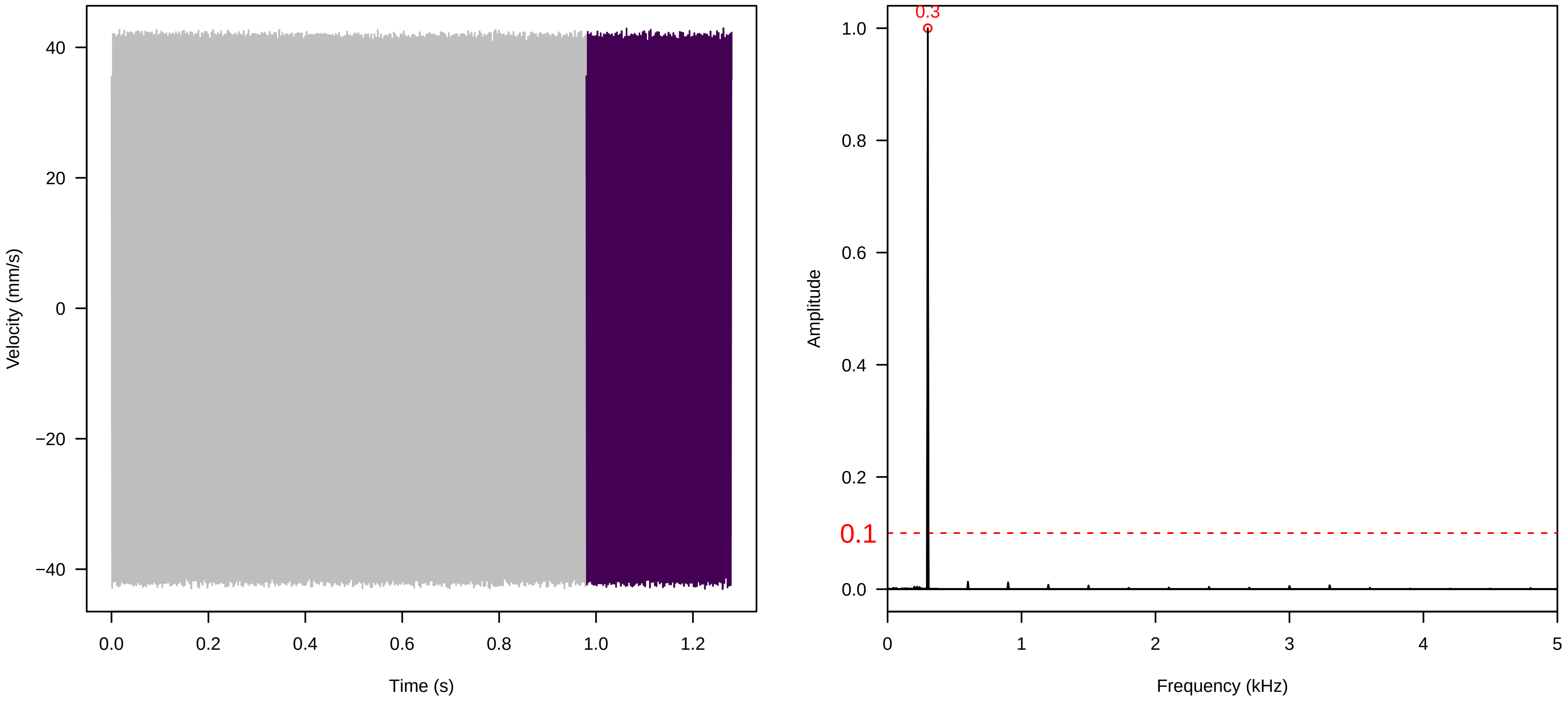


**Vel. = 0.028 ; Str. = FA ; Axis = y ; Fl. accession = 10−s−77−12**

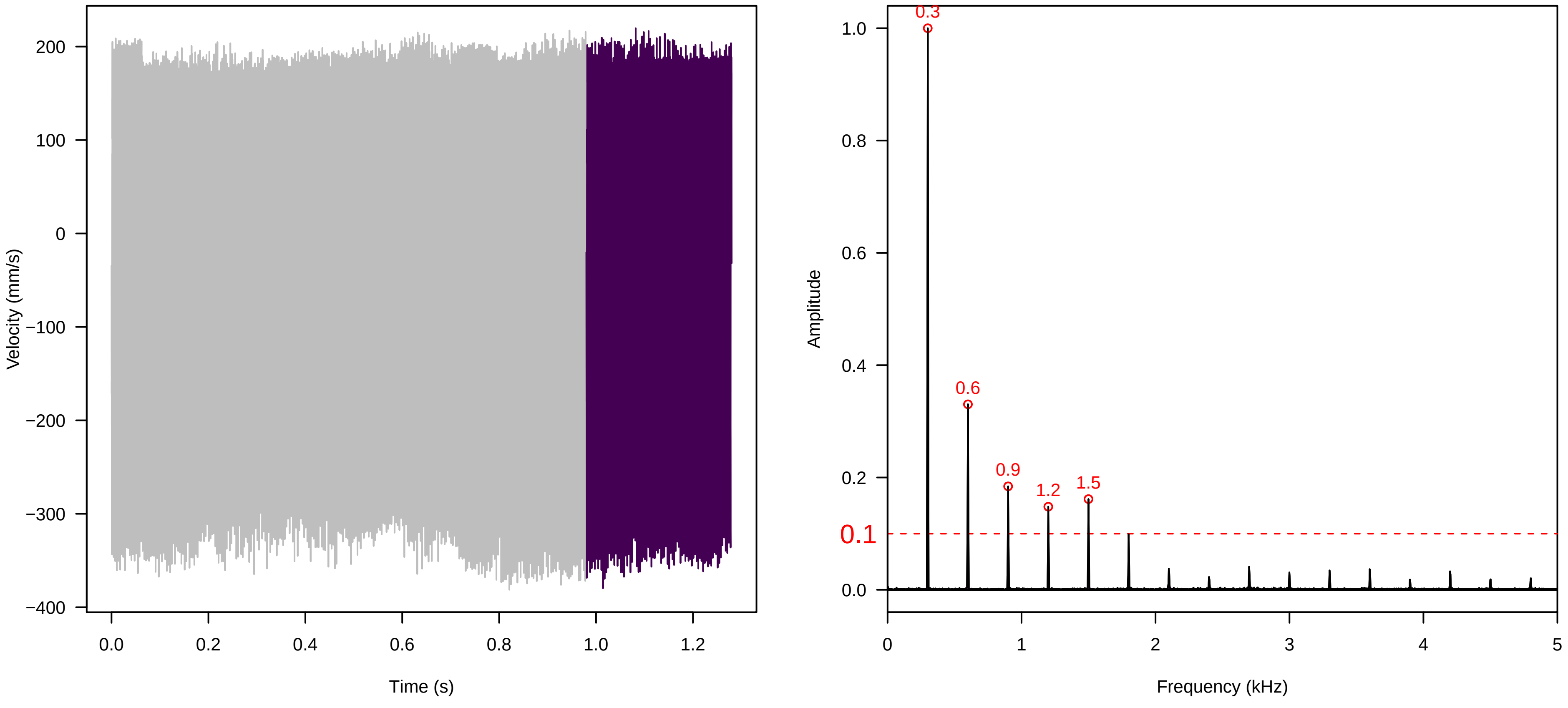

**Vel. = 0.028 ; Str. = Receptacle ; Axis = y ; Fl. accession = 10−s−77−12**

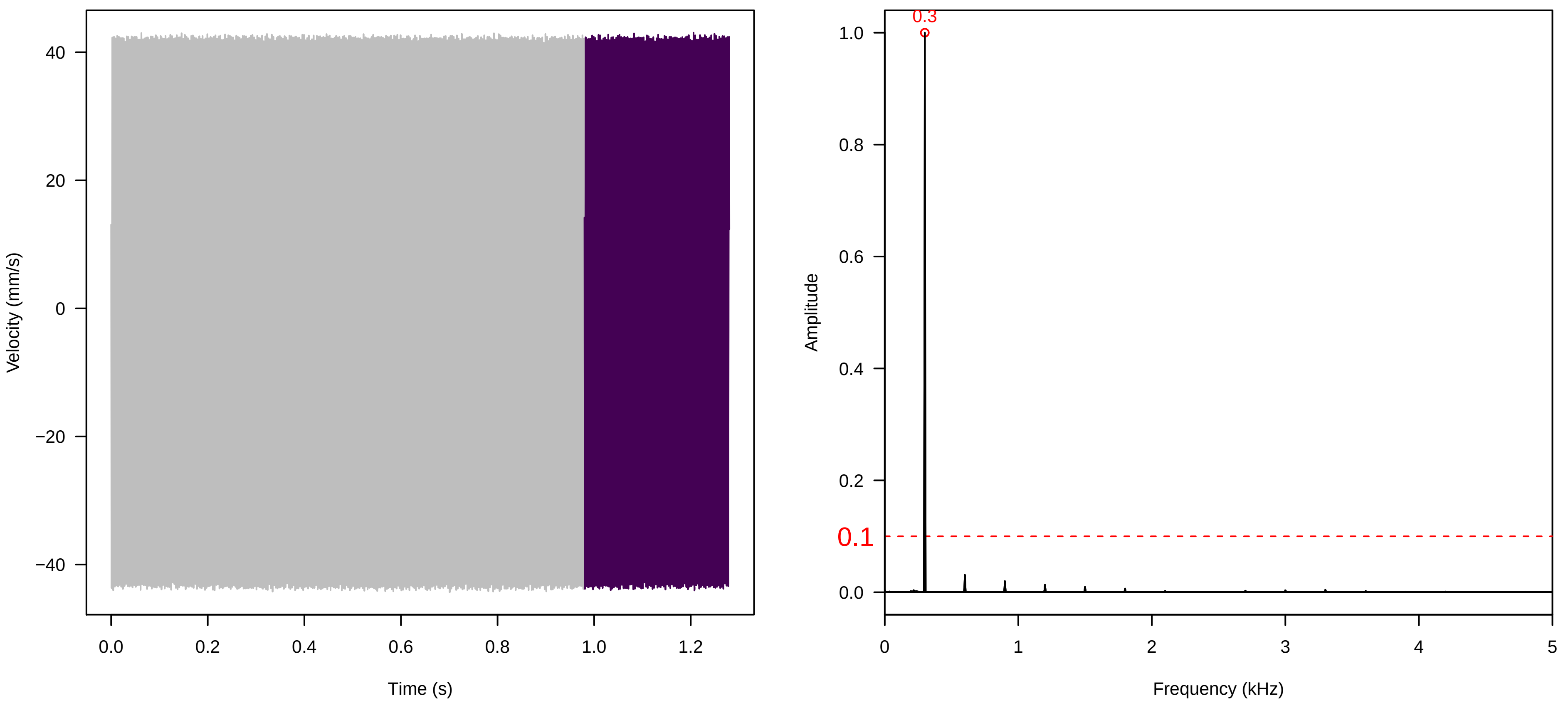


**Vel. = 0.028 ; Str. = Corolla ; Axis = y ; Fl. accession = 10−s−77−12**

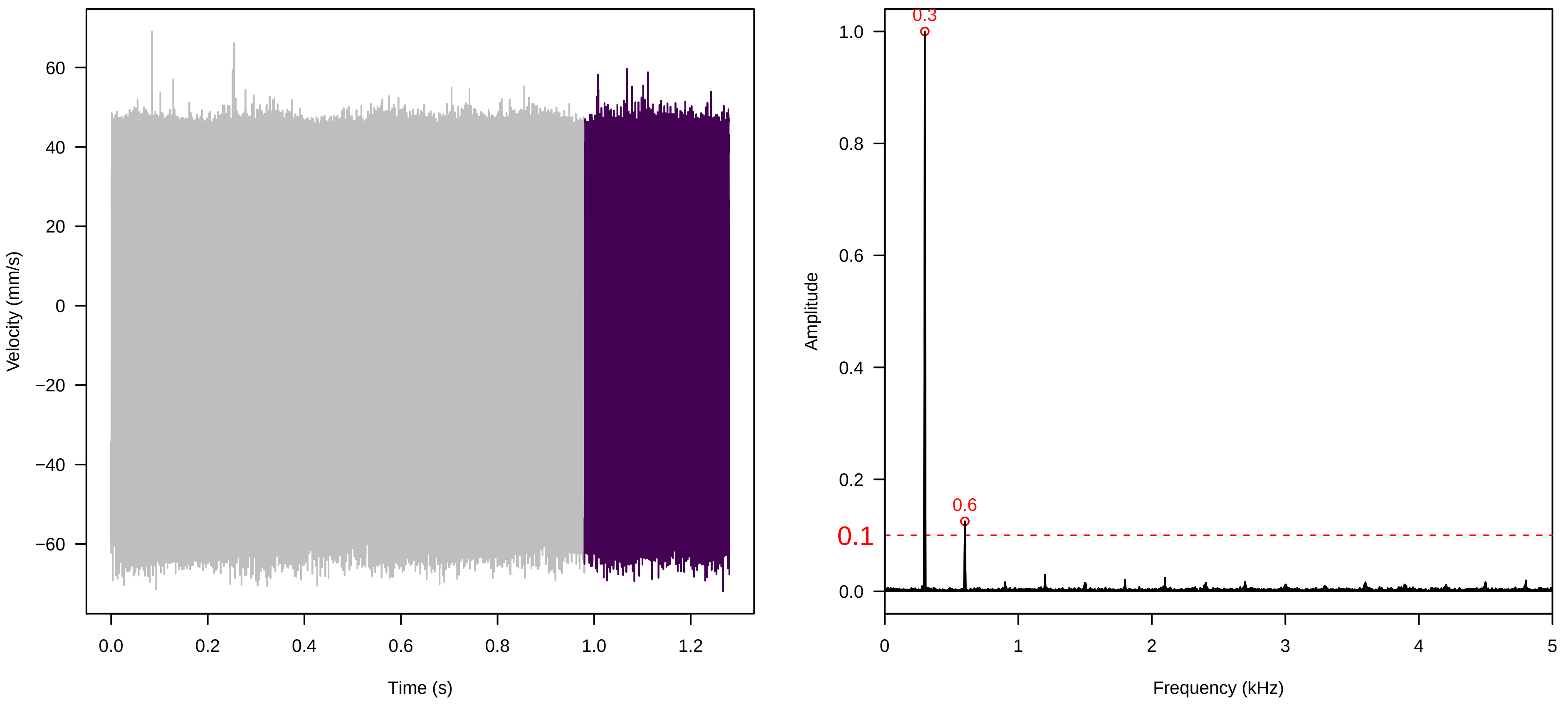


**Vel. = 0.028 ; Str. = Receptacle ; Axis = y ; Fl. accession = 10−s−77−12**

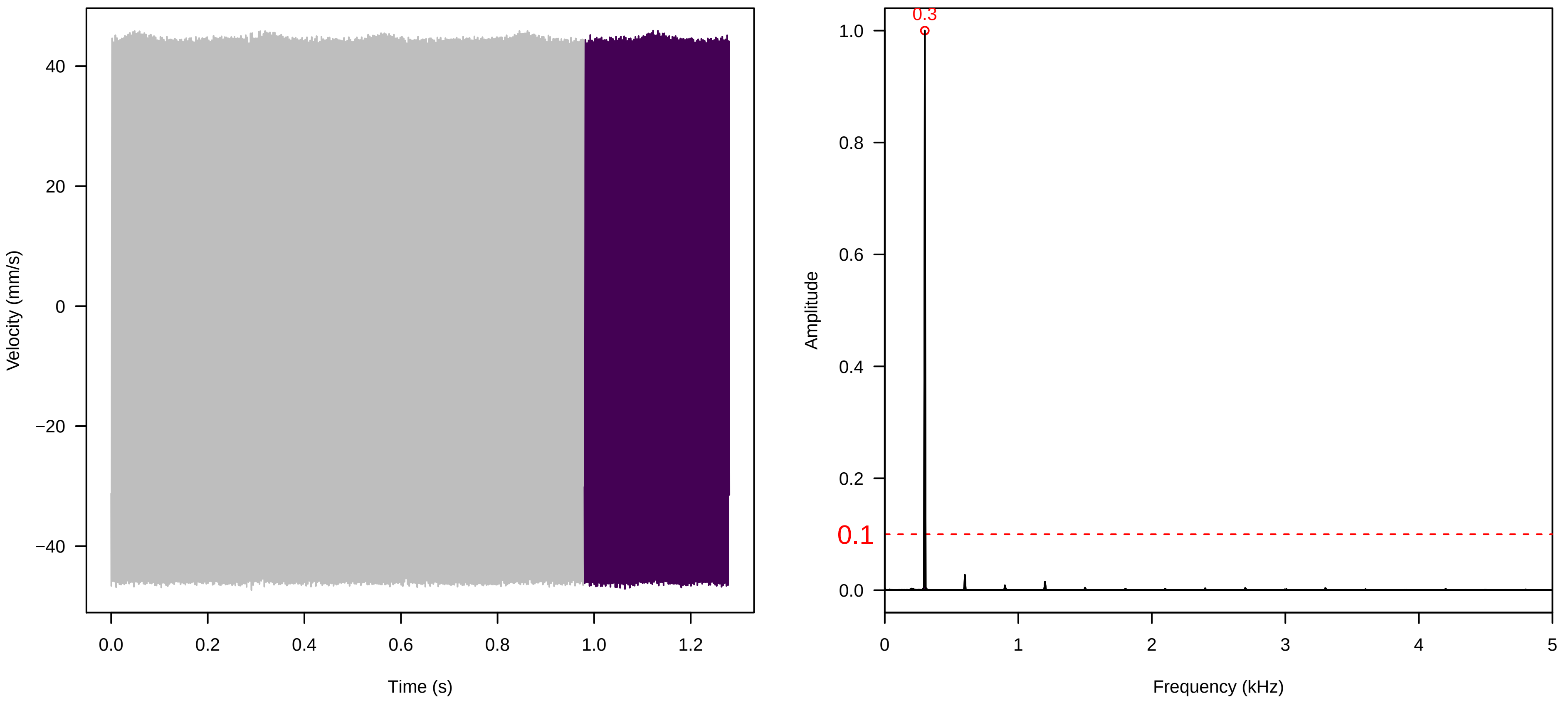

**Vel. = 0.057 ; Str. = Corolla ; Axis = y ; Fl. accession = 10−s−77−12**

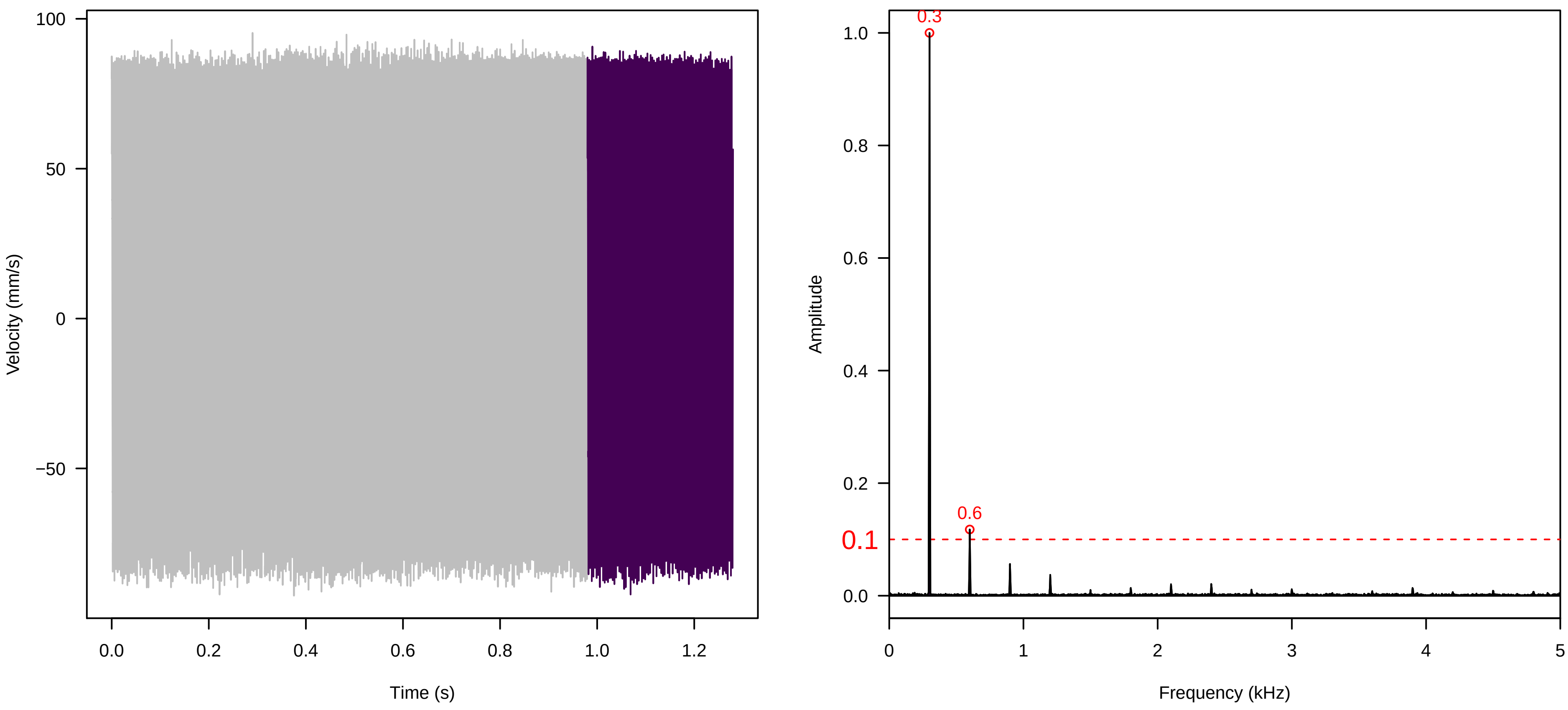


**Vel. = 0.057 ; Str. = Receptacle ; Axis = y ; Fl. accession = 10−s−77−12**

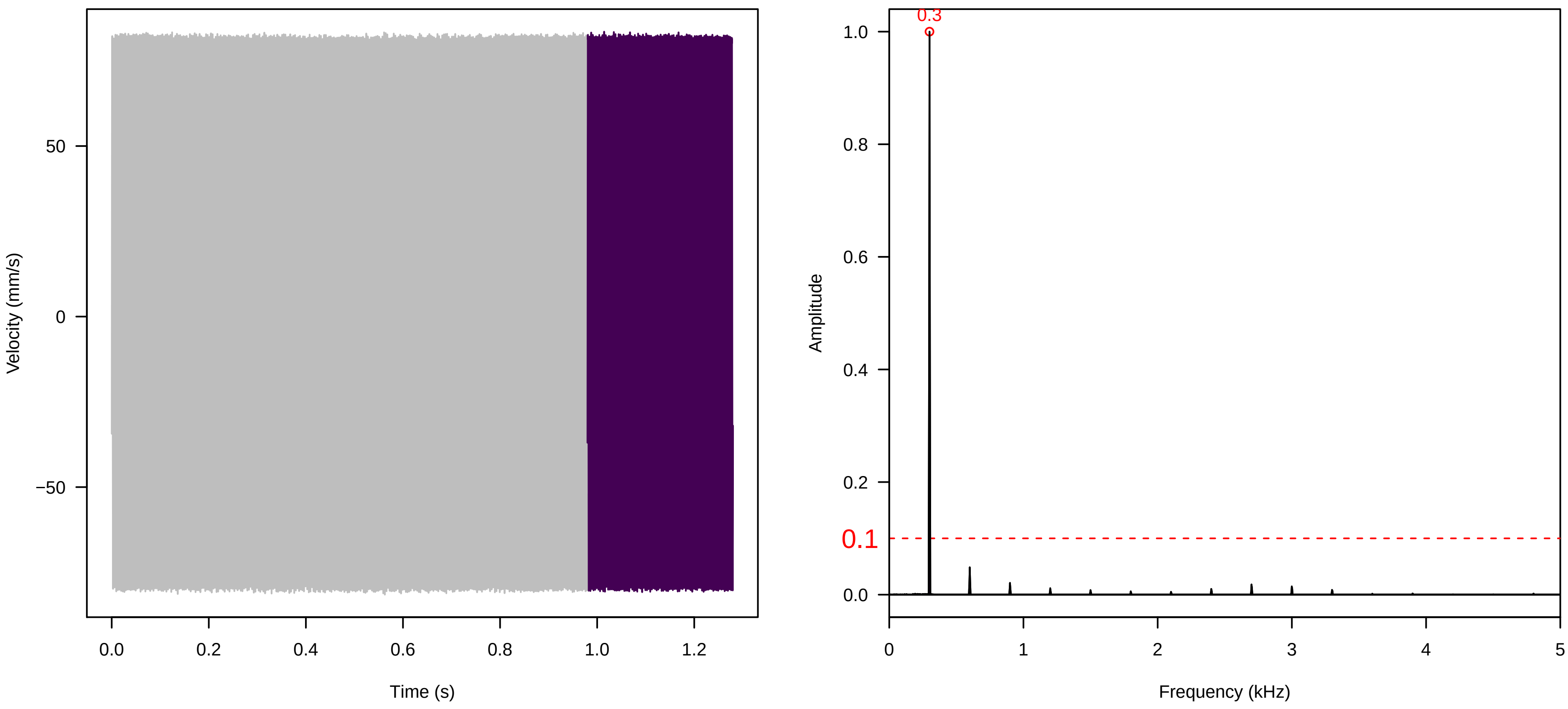


**Vel. = 0.057 ; Str. = FA ; Axis = y ; Fl. accession = 10−s−77−12**

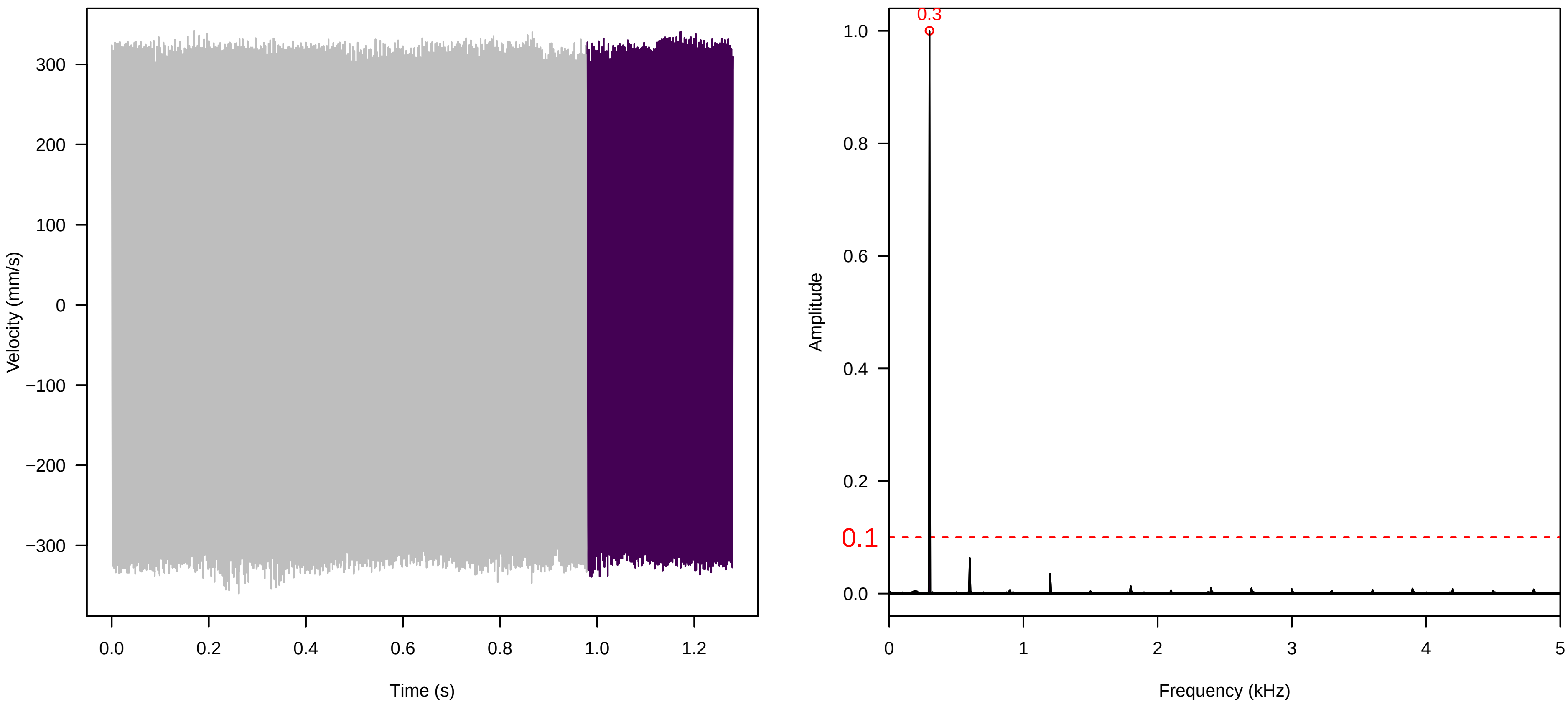

Vel. = 0.057 ; Str. = Receptacle ; Axis = y ; Fl. accession = 10−s−77−12

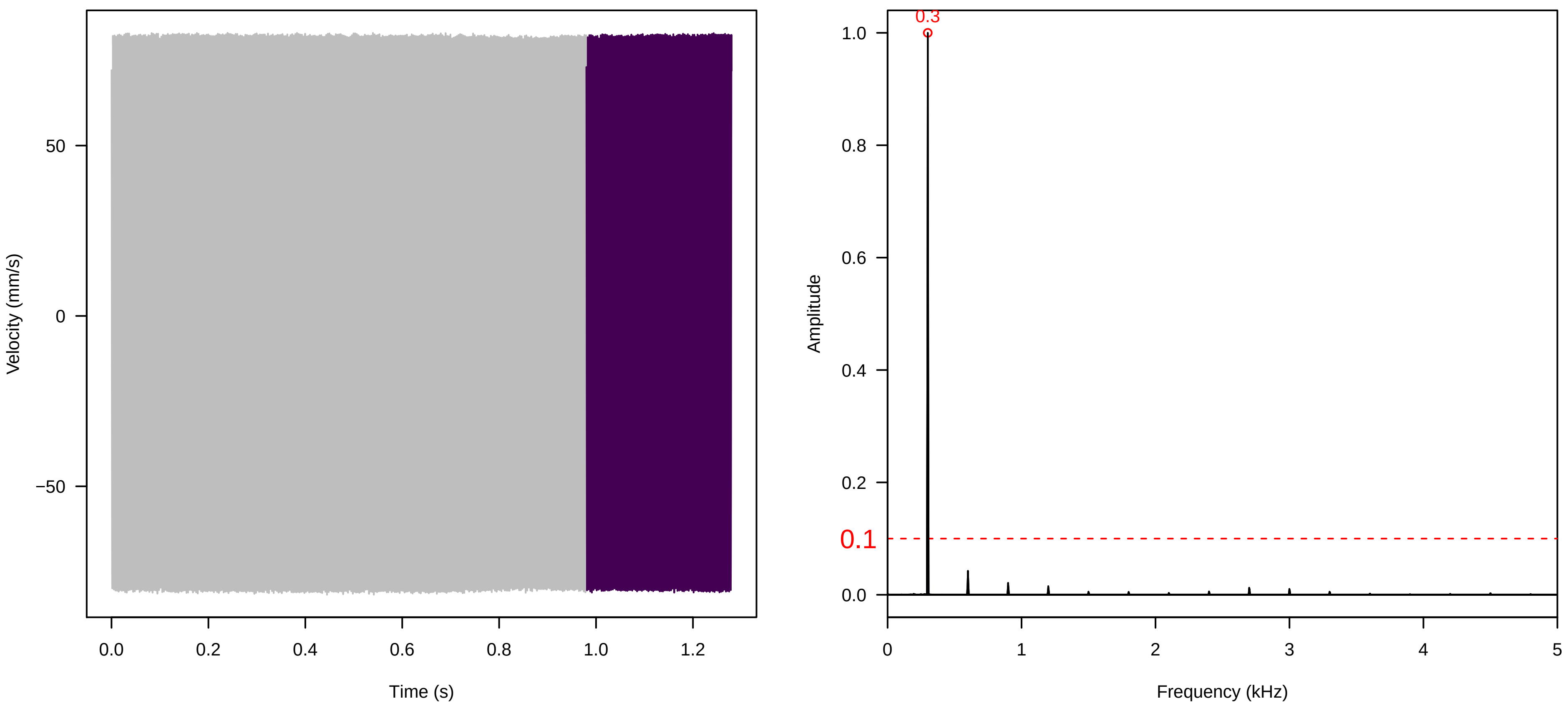


Vel. = 0.057 ; Str. = PA ; Axis = y ; Fl. accession = 10−s−77−12

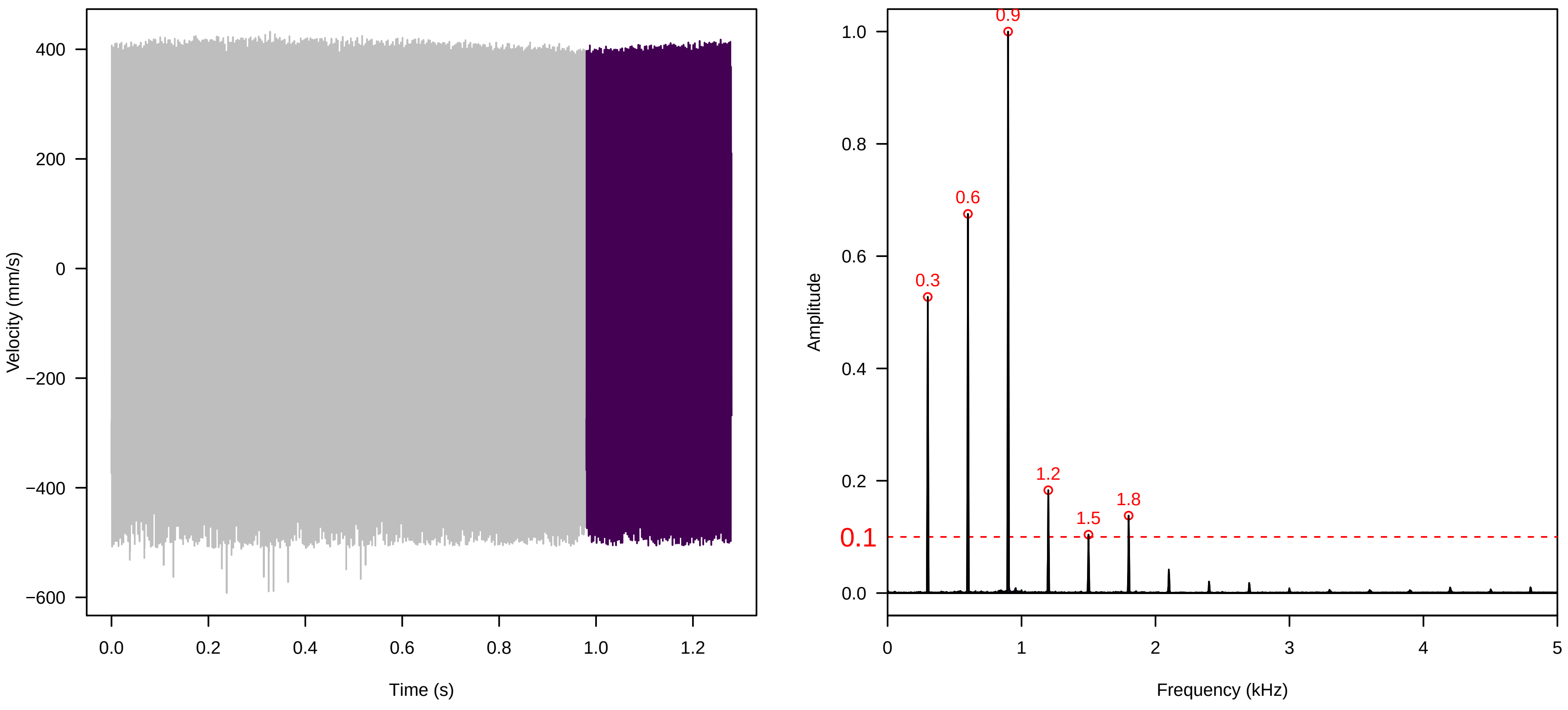


Vel. = 0.057 ; Str. = Receptacle ; Axis = y ; Fl. accession = 10−s−77−12

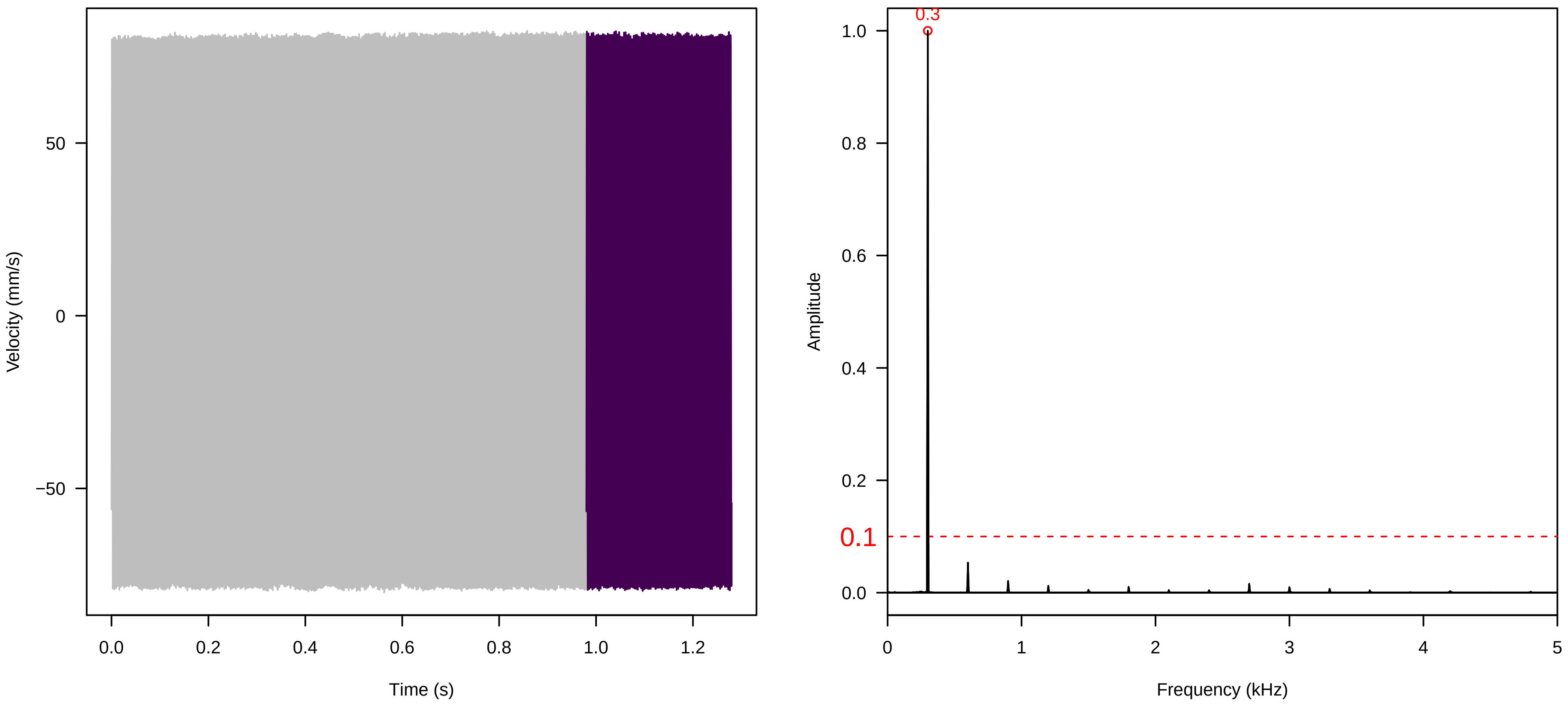

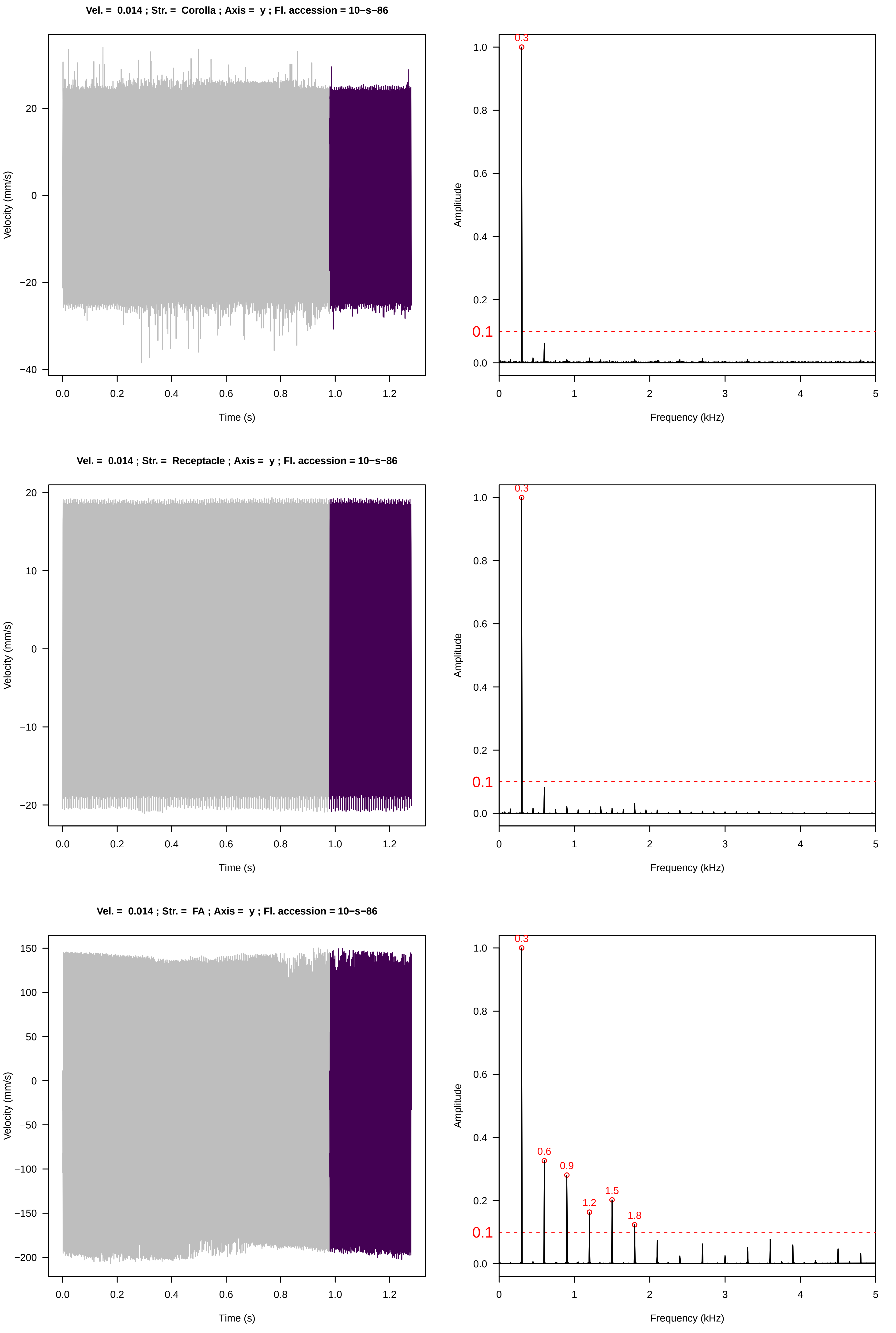
Vel. = 0.014 ; Str. = Corolla ; Axis = y ; Fl. accession = 10−s−86
Velocity (mm/s)
Time (s)
Amplitude
Frequency (kHz)
0.3
0.1
Vel. = 0.014 ; Str. = Receptacle ; Axis = y ; Fl. accession = 10−s−86
Velocity (mm/s)
Time (s)
Amplitude
Frequency (kHz)
0.3
0.1
Vel. = 0.014 ; Str. = FA ; Axis = y ; Fl. accession = 10−s−86
Velocity (mm/s)
Time (s)
Amplitude
Frequency (kHz)
0.3
0.6
0.9
1.2
1.5
1.8
0.1

**Vel. = 0.014 ; Str. = Receptacle ; Axis = y ; Fl. accession = 10−s−86**

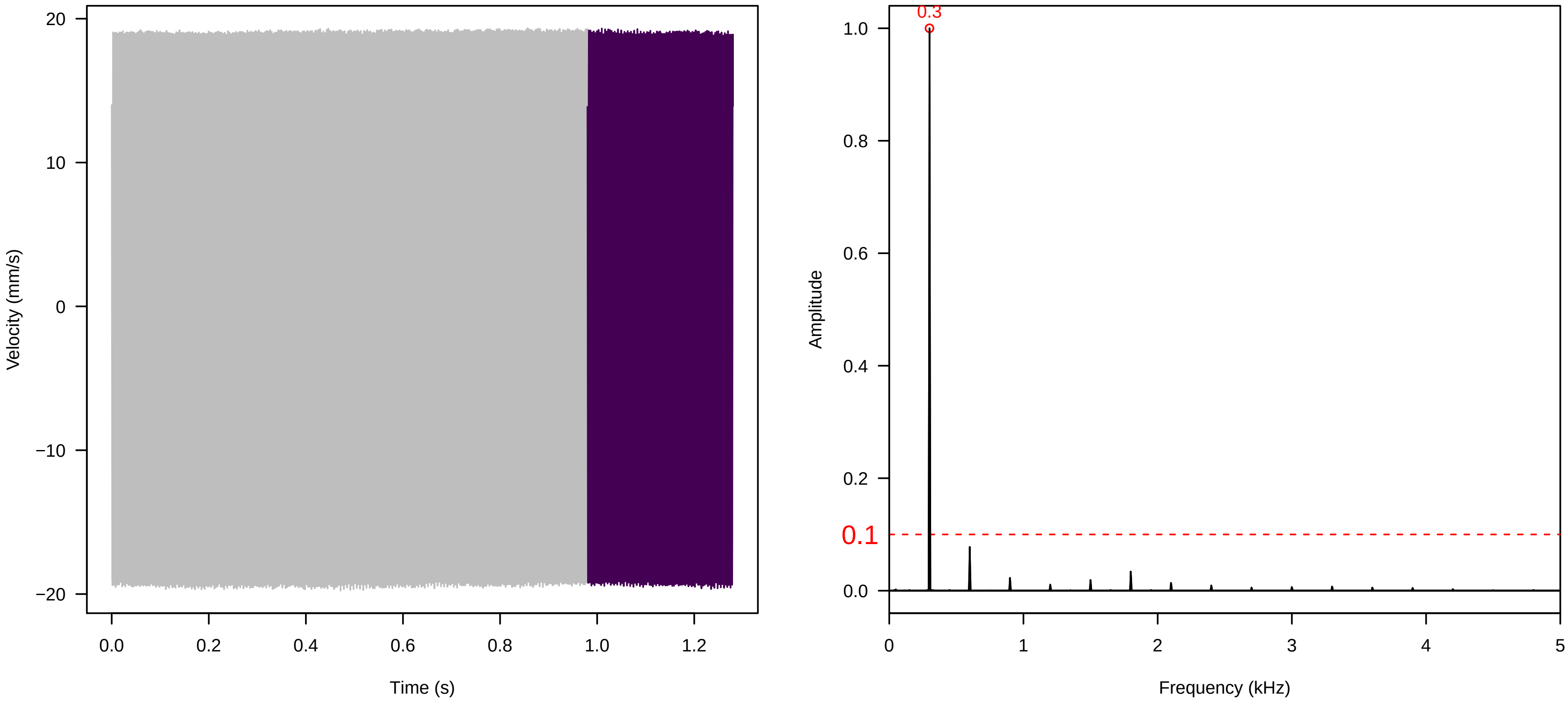


**Vel. = 0.014 ; Str. = PA ; Axis = y ; Fl. accession = 10−s−86**

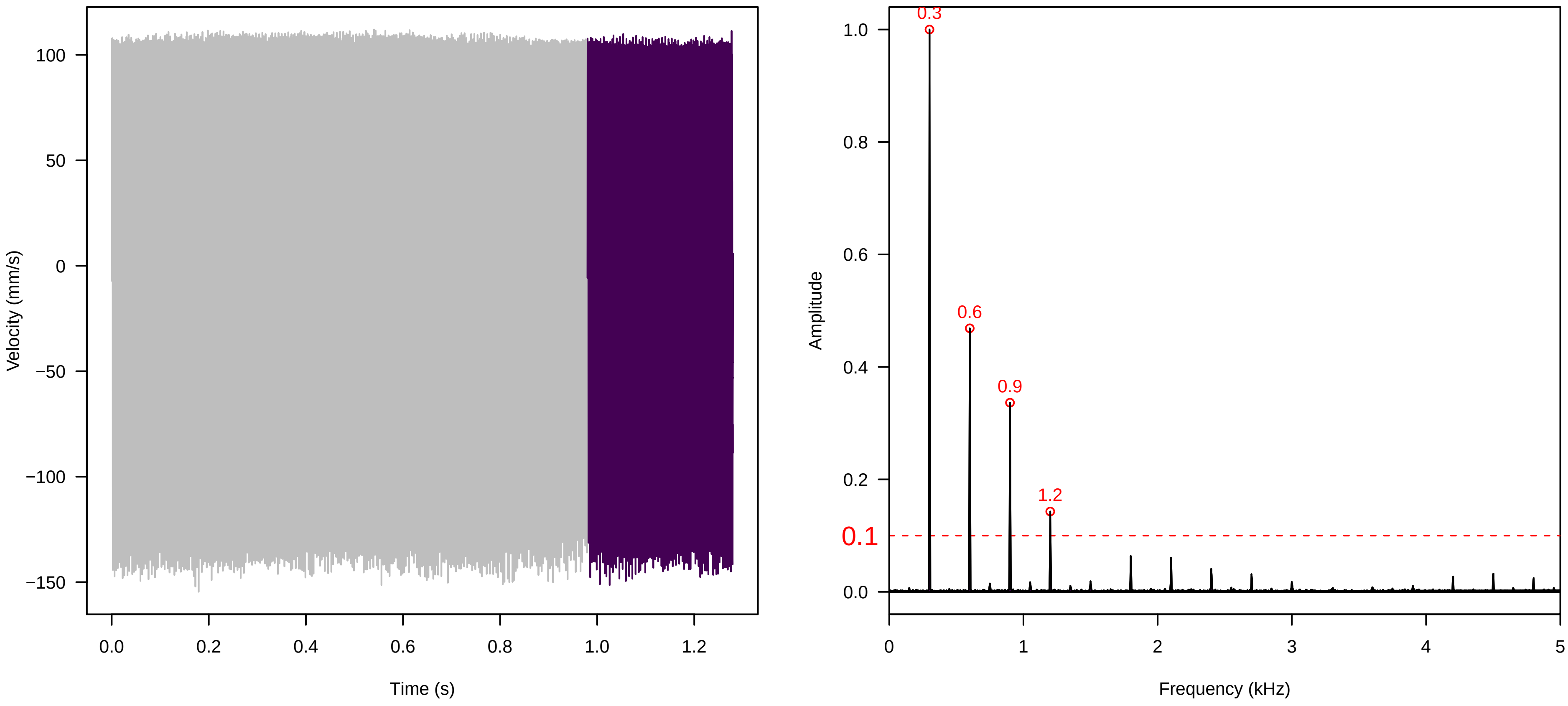


**Vel. = 0.014 ; Str. = Receptacle ; Axis = y ; Fl. accession = 10−s−86**

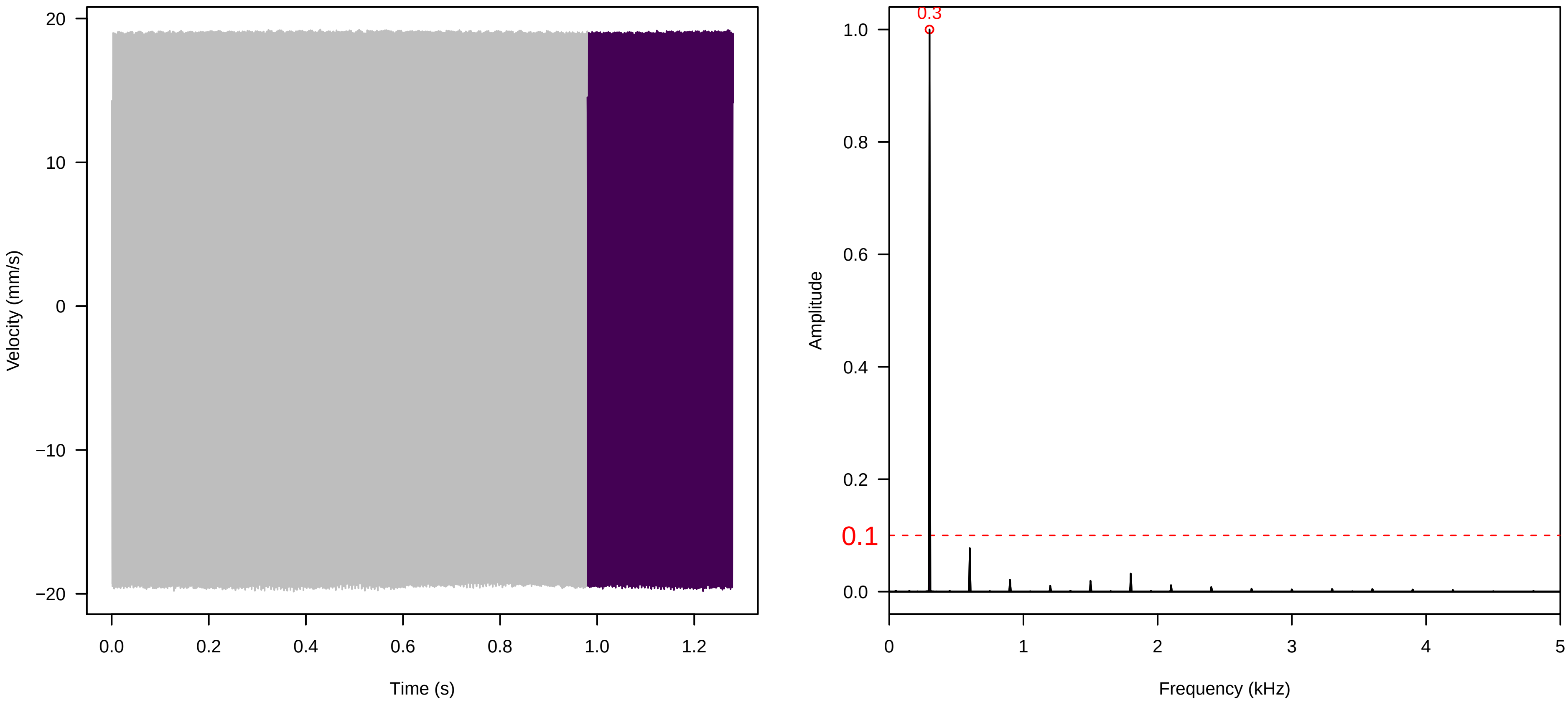

**Vel. = 0.028 ; Str. = PA ; Axis = y ; Fl. accession = 10−s−86**

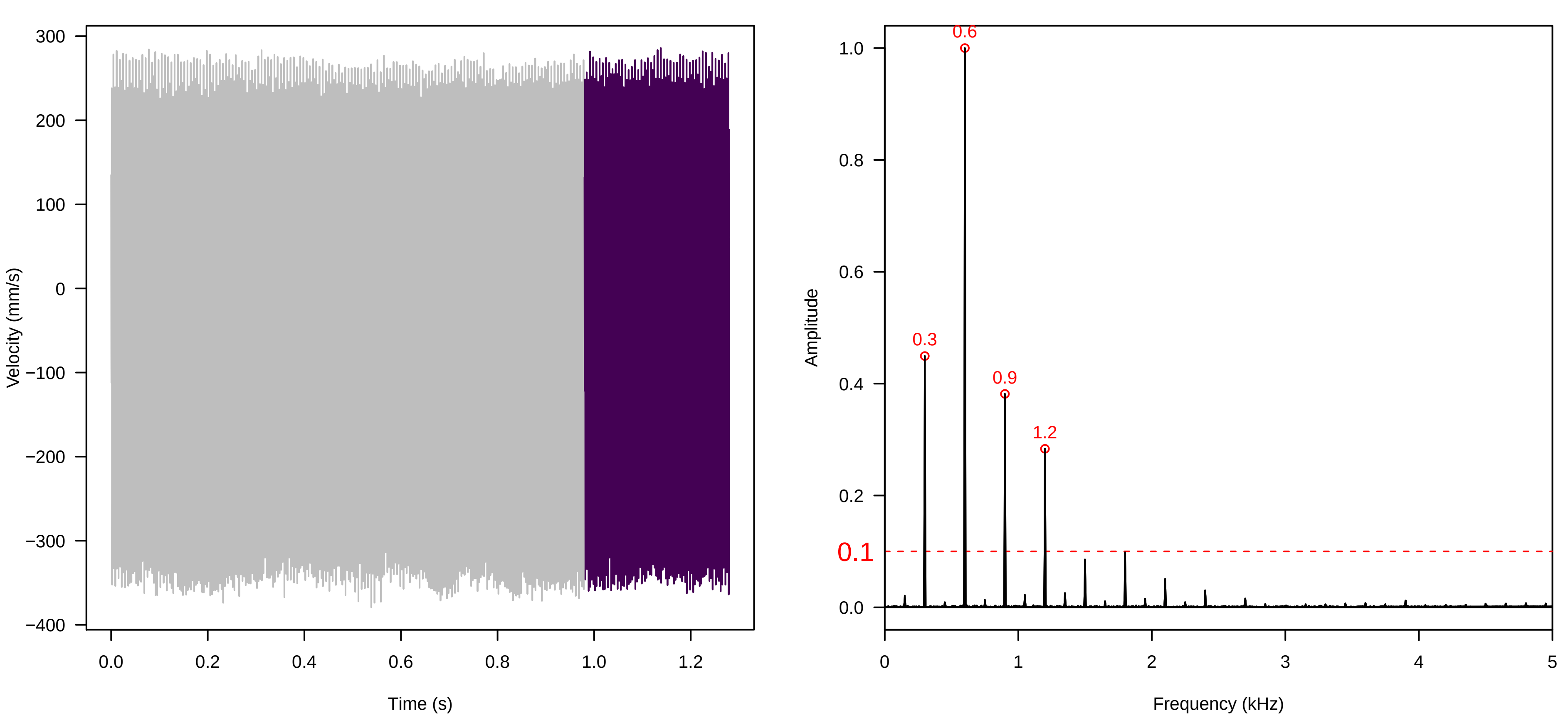


**Vel. = 0.028 ; Str. = Receptacle ; Axis = y ; Fl. accession = 10−s−86**

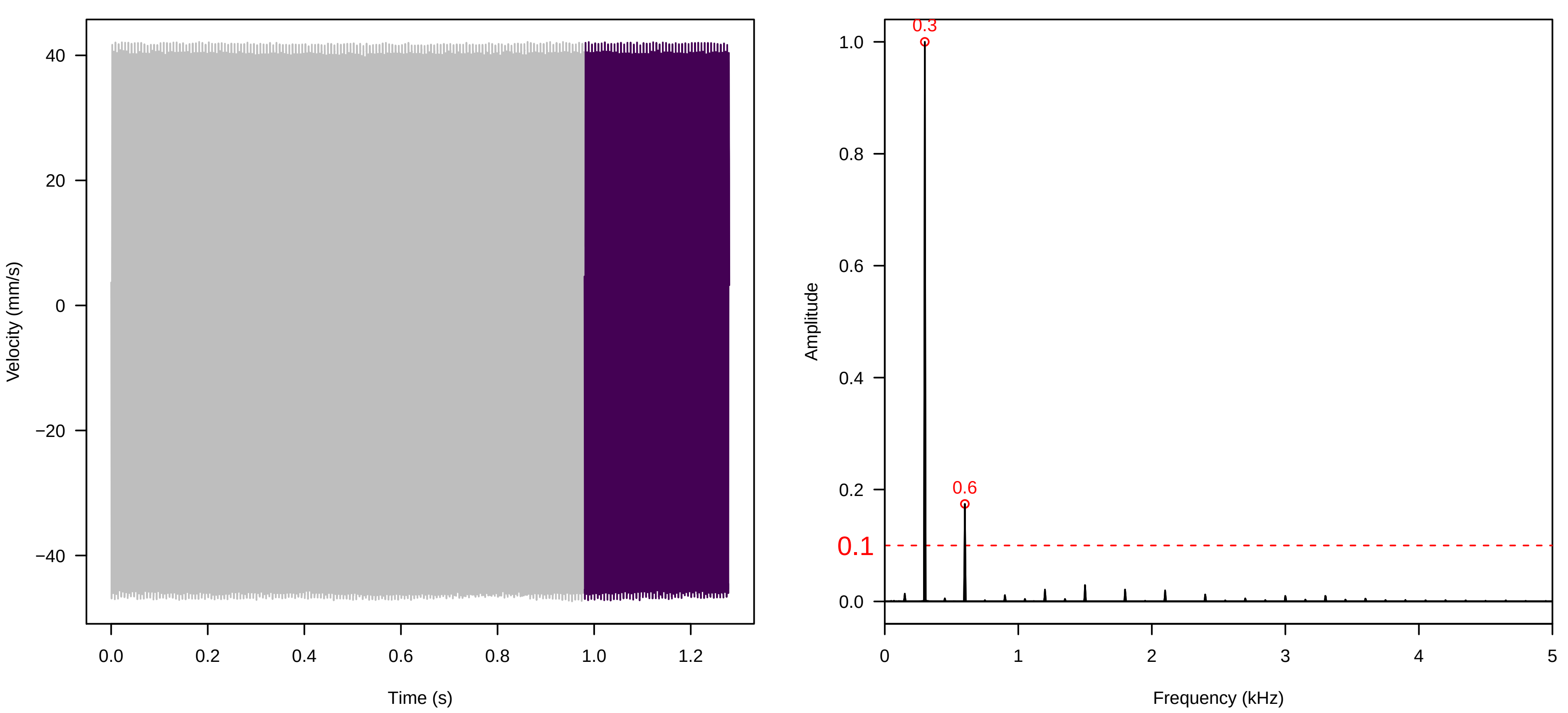


**Vel. = 0.028 ; Str. = FA ; Axis = y ; Fl. accession = 10−s−86**

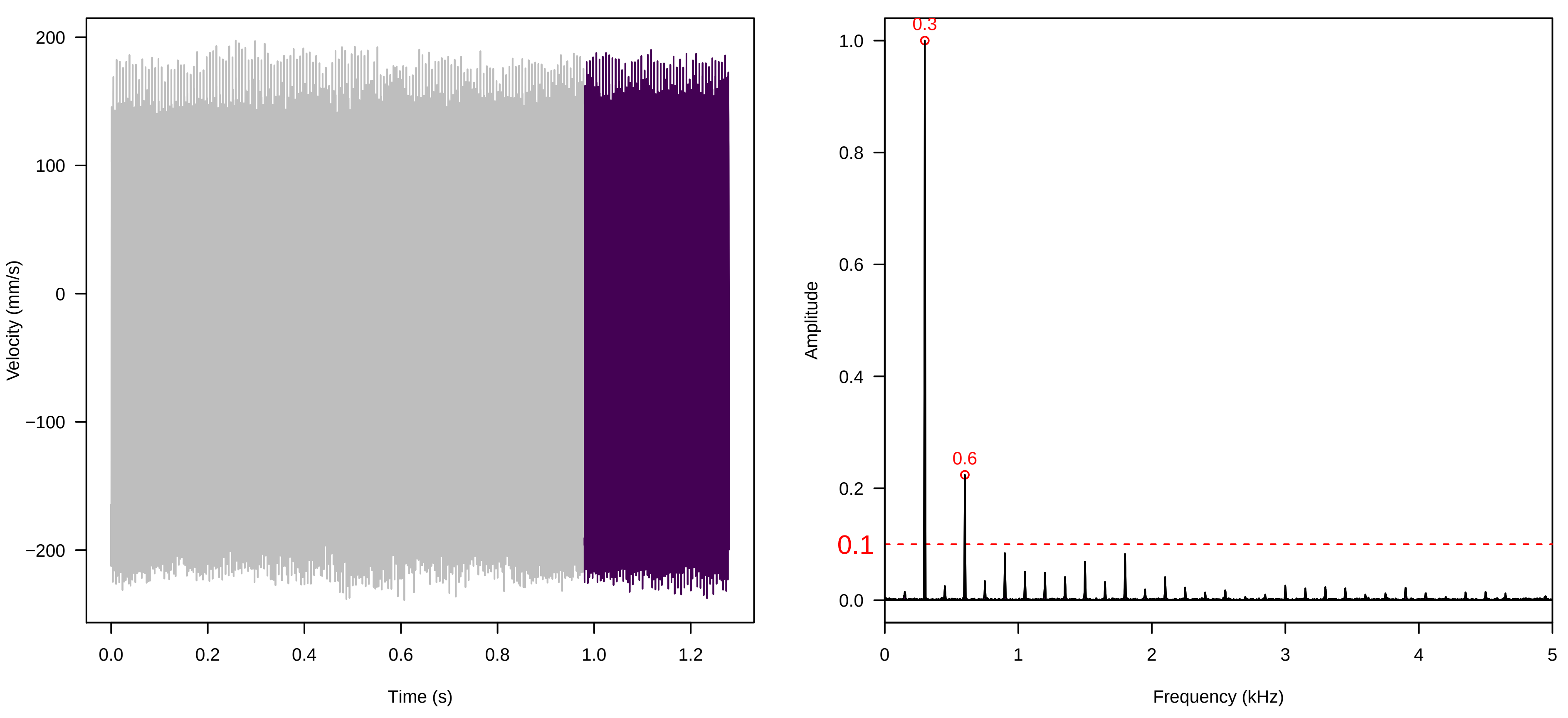

**Vel. =  0.028 ; Str. =  Receptacle ; Axis =  y ; Fl. accession = 10−s−86**

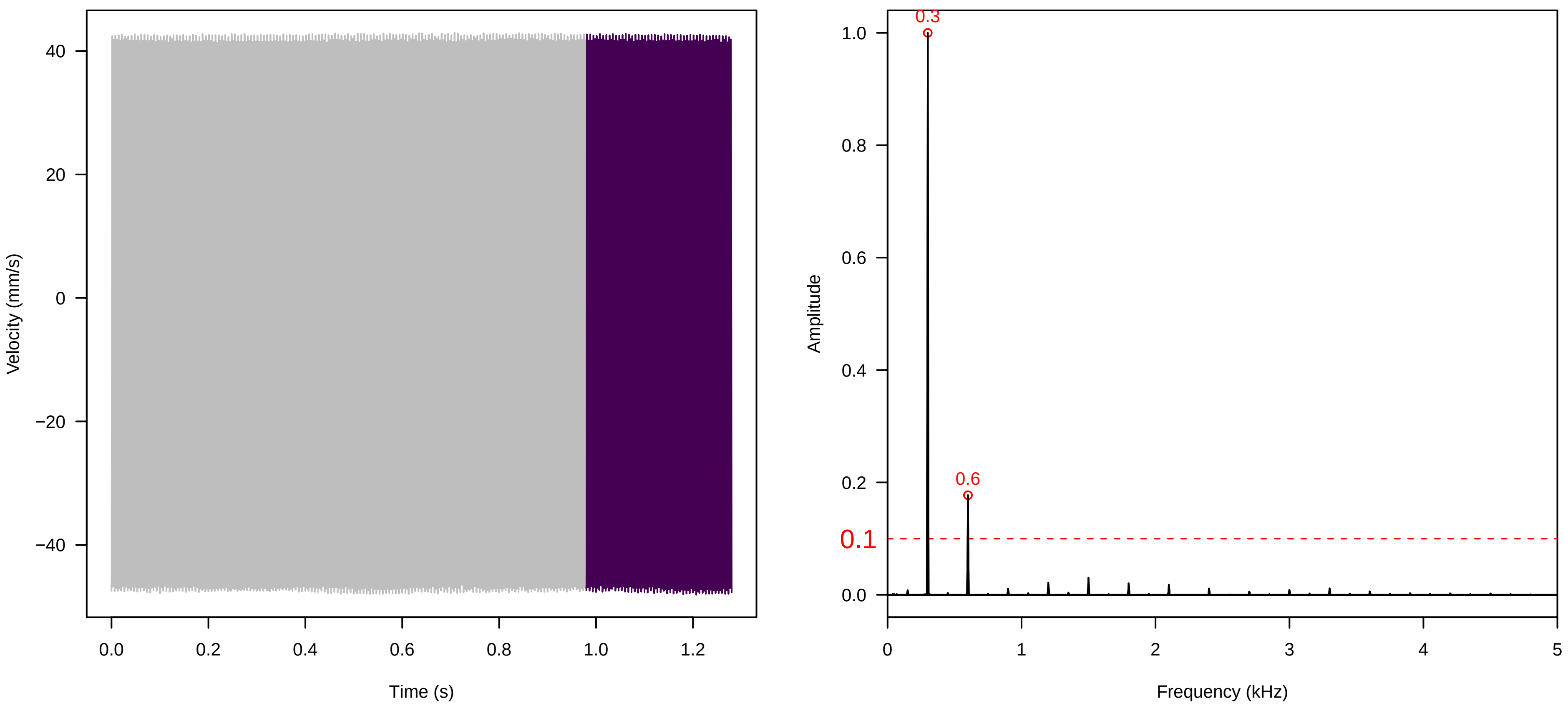


**Vel. =  0.028 ; Str. =  Corolla ; Axis =  y ; Fl. accession = 10−s−86**

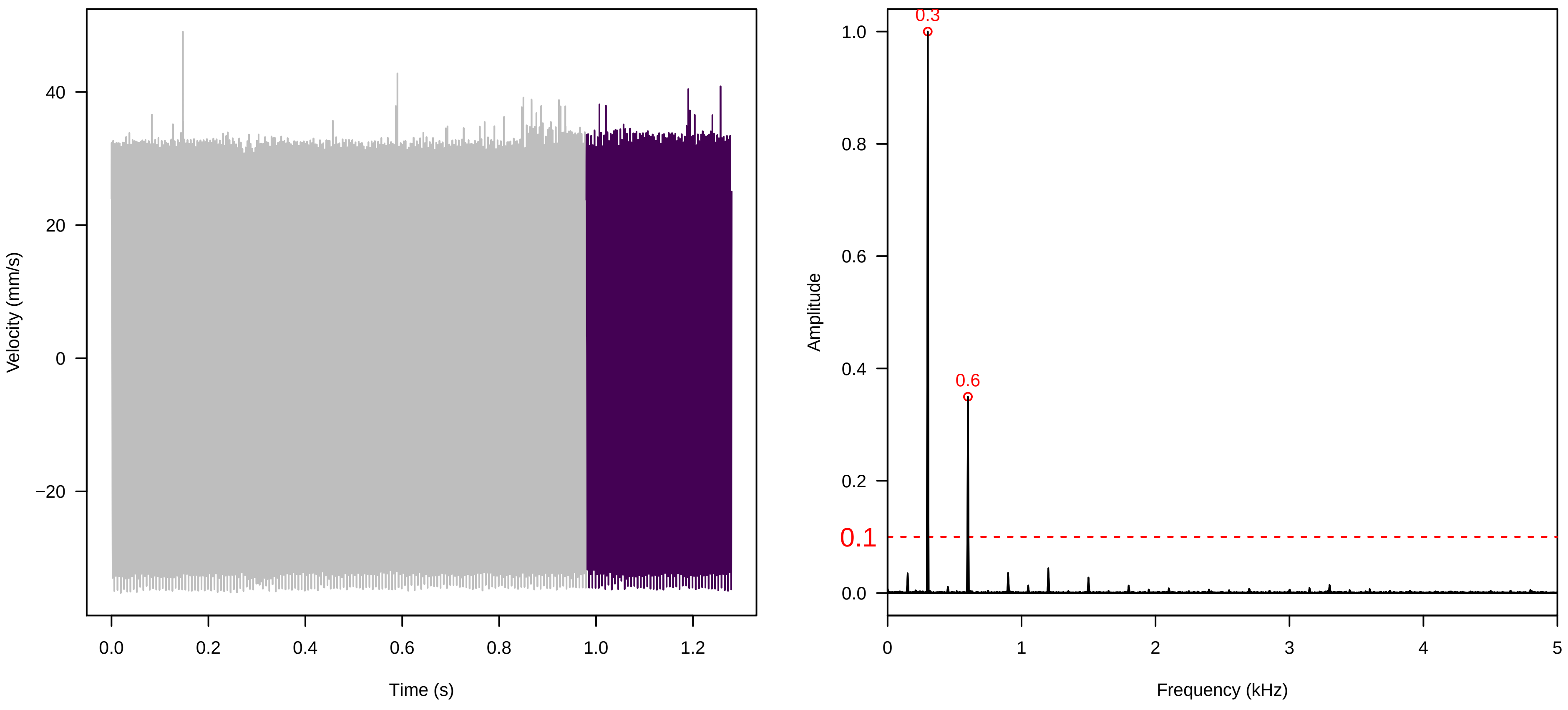


**Vel. =  0.028 ; Str. =  Receptacle ; Axis =  y ; Fl. accession = 10−s−86**

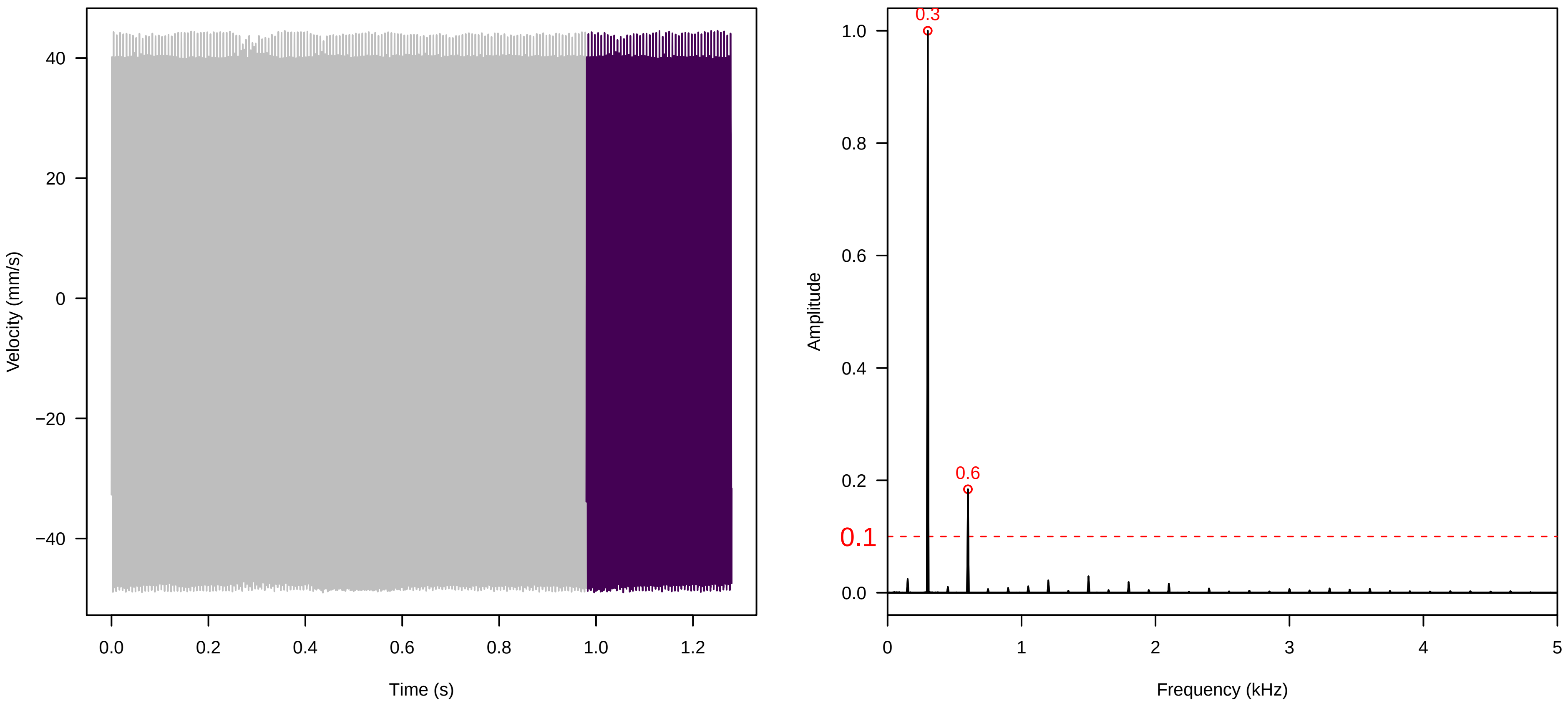

**Vel. =  0.057 ; Str. =  Corolla ; Axis =  y ; Fl. accession = 10−s−86**

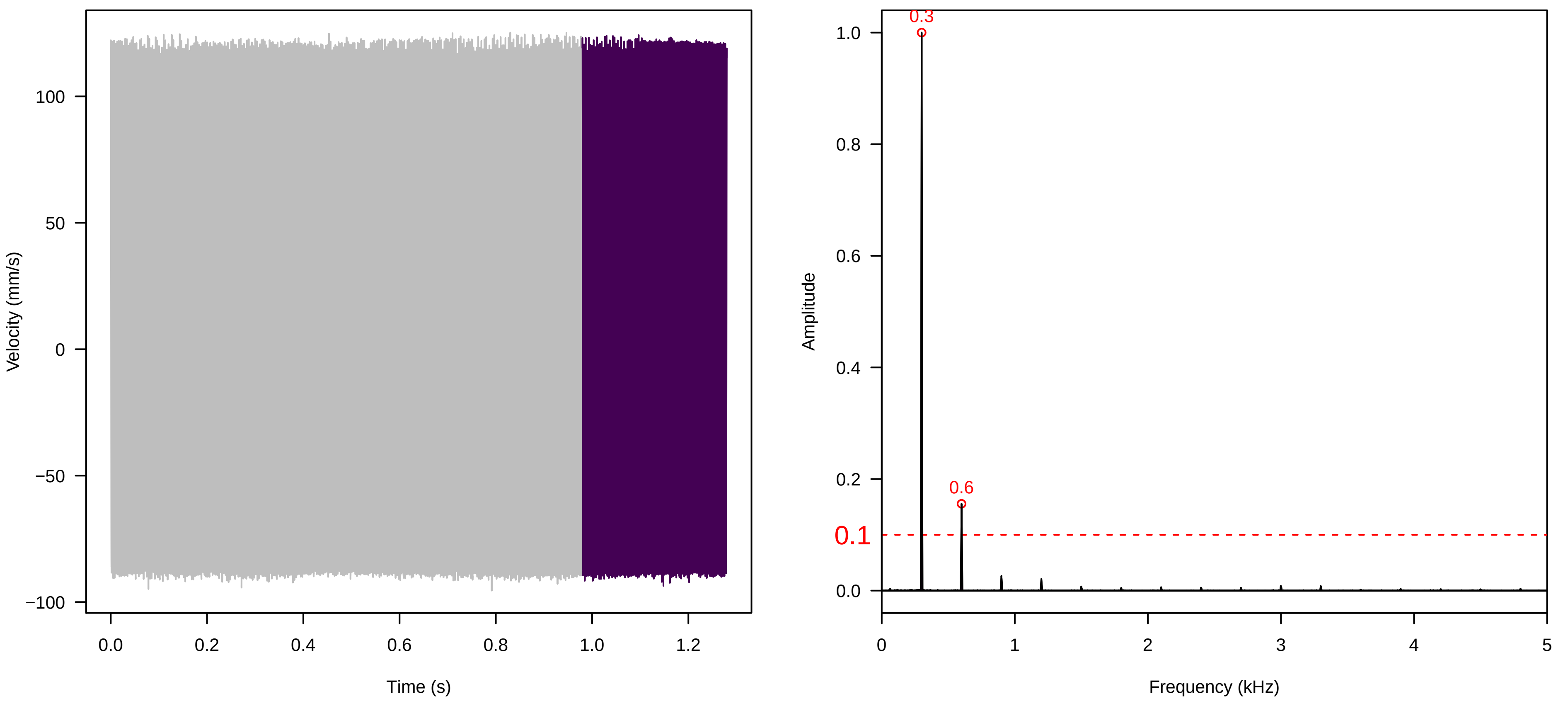


**Vel. =  0.057 ; Str. =  Receptacle ; Axis =  y ; Fl. accession = 10−s−86**

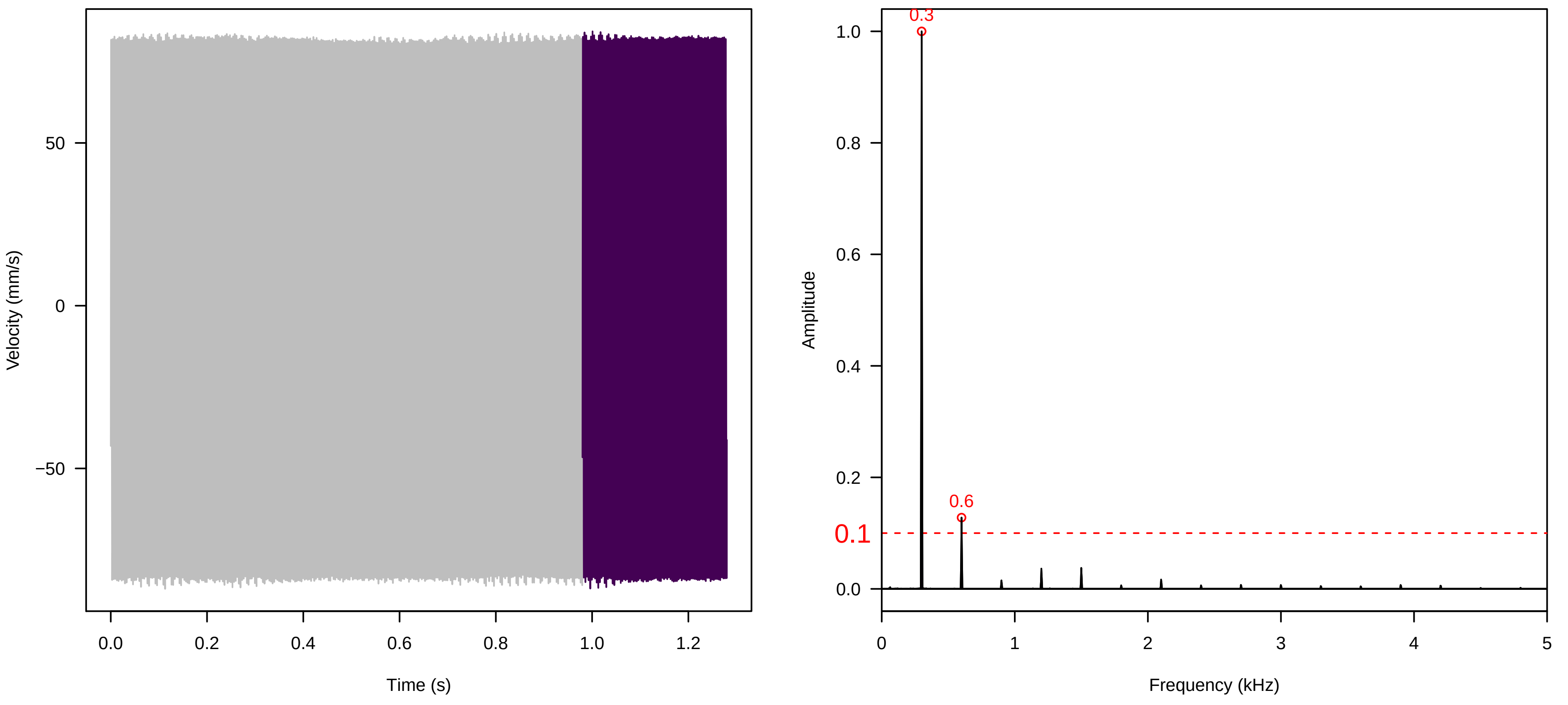


**Vel. =  0.057 ; Str. =  FA ; Axis =  y ; Fl. accession = 10−s−86**

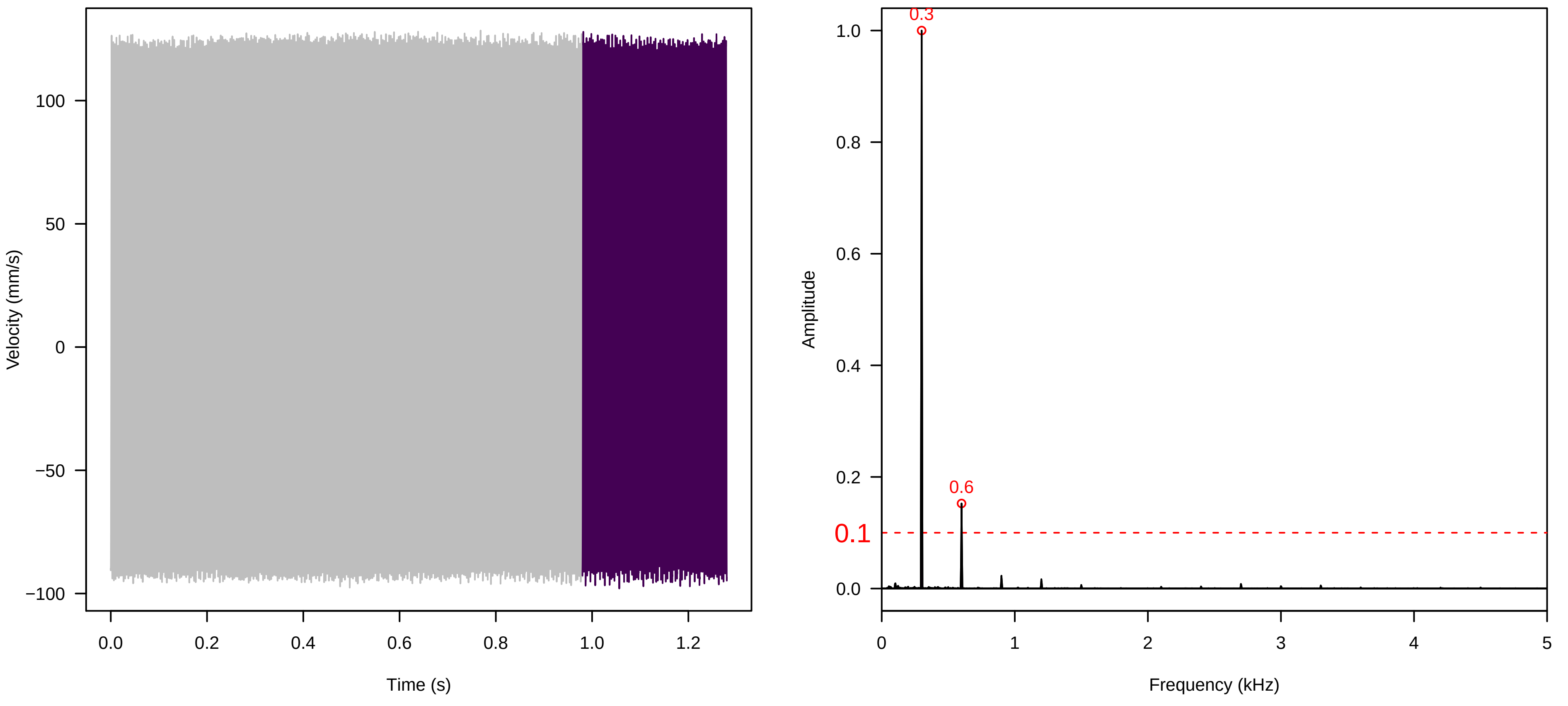

**Vel. = 0.057 ; Str. = Receptacle ; Axis = y ; Fl. accession = 10−s−86**

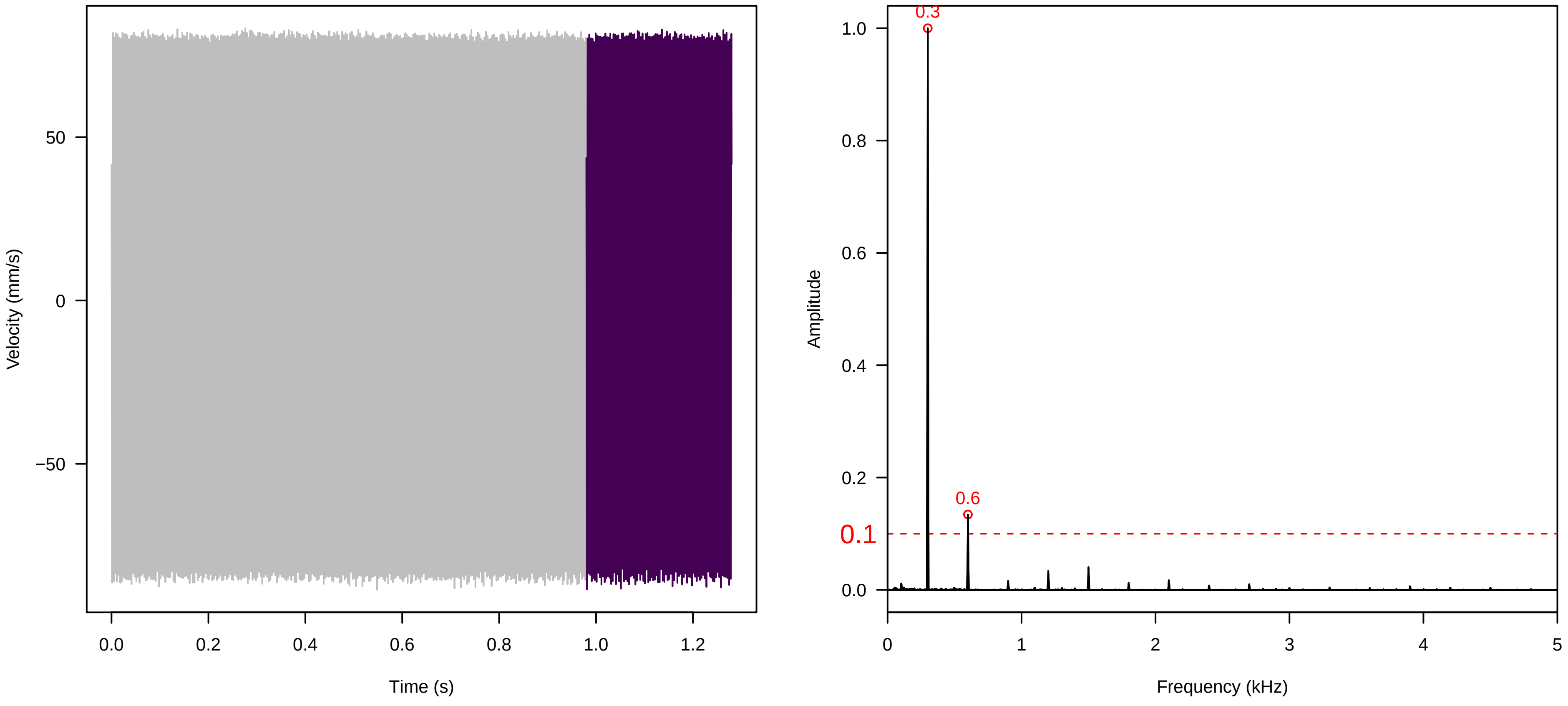


**Vel. = 0.057 ; Str. = PA ; Axis = y ; Fl. accession = 10−s−86**

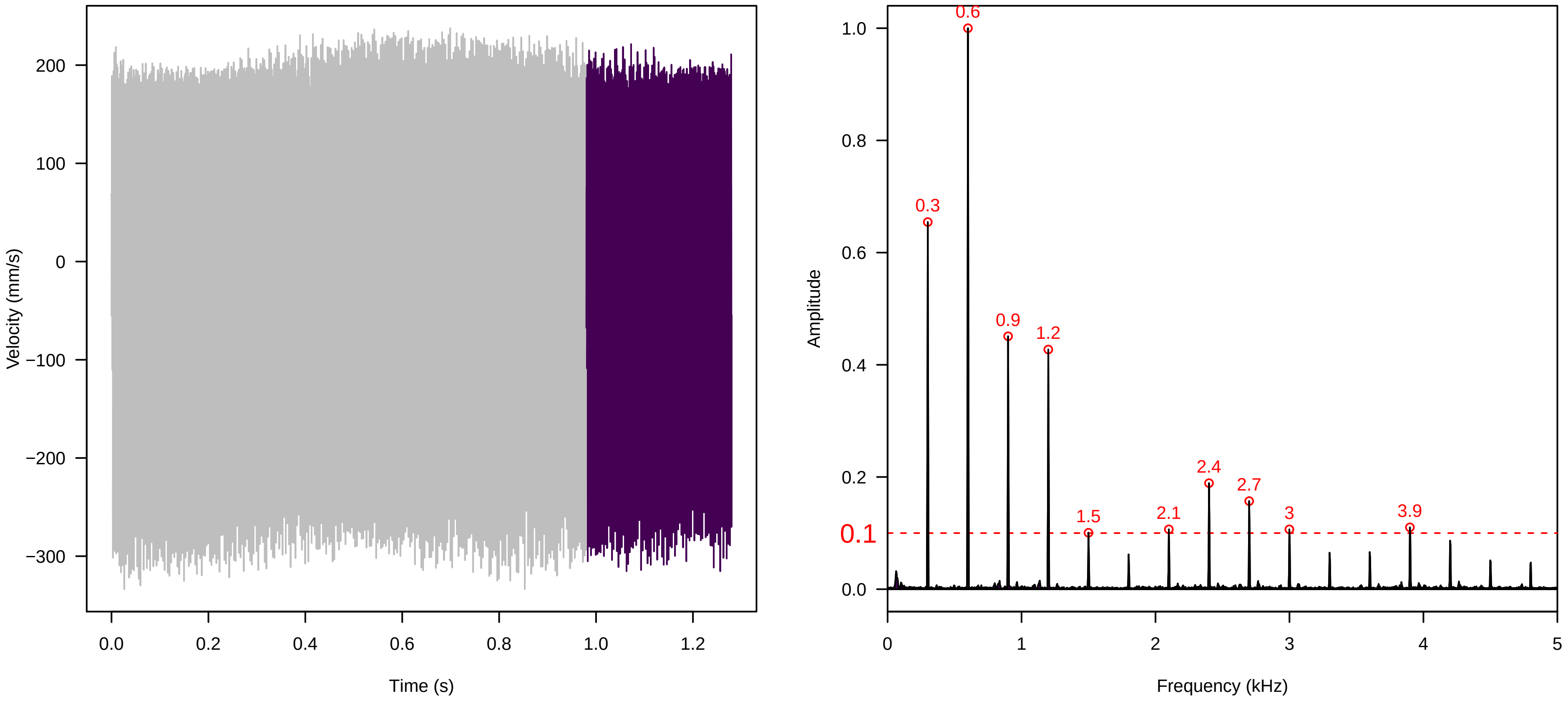


**Vel. = 0.057 ; Str. = Receptacle ; Axis = y ; Fl. accession = 10−s−86**

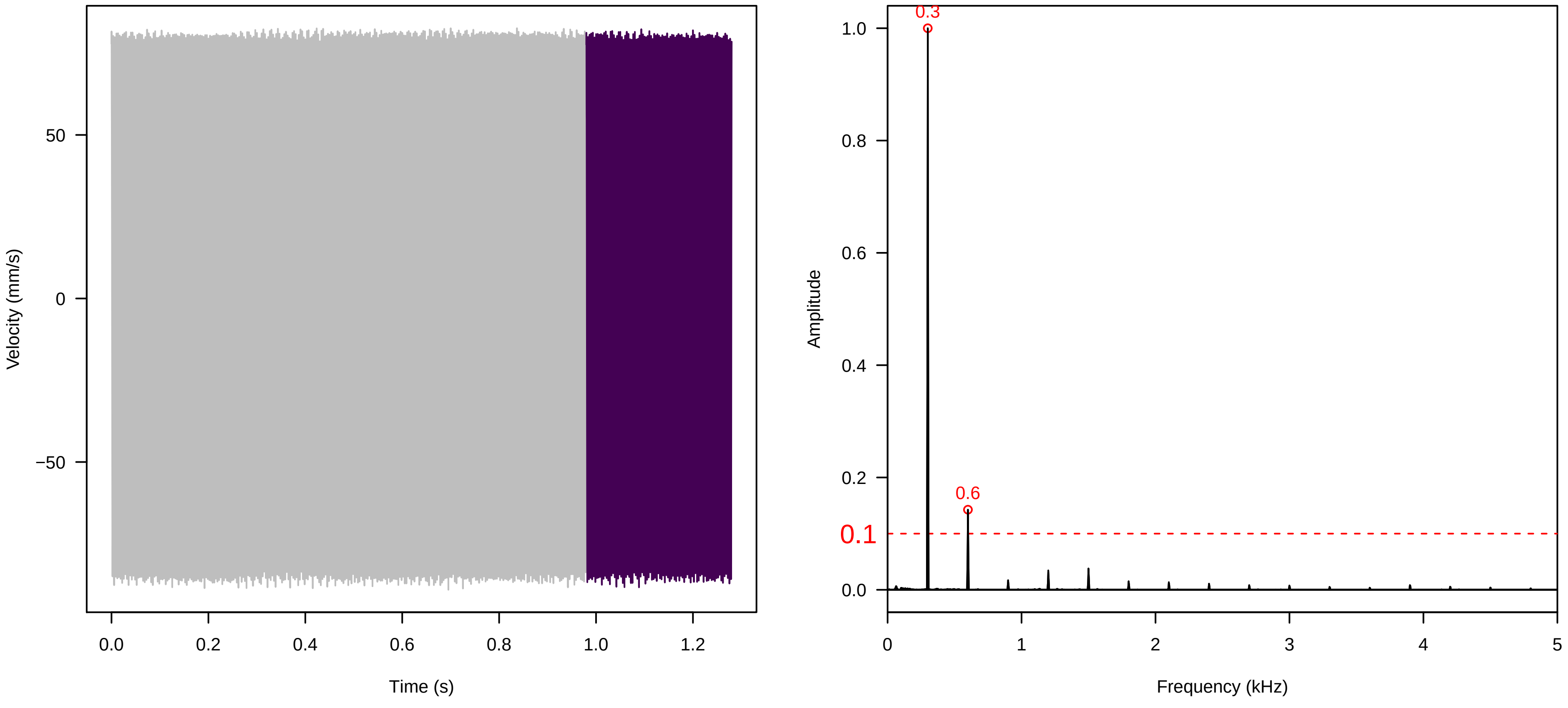

Vel. =  0.014 ; Str. =  Corolla ; Axis =  z ; Fl. accession = 10−s−86

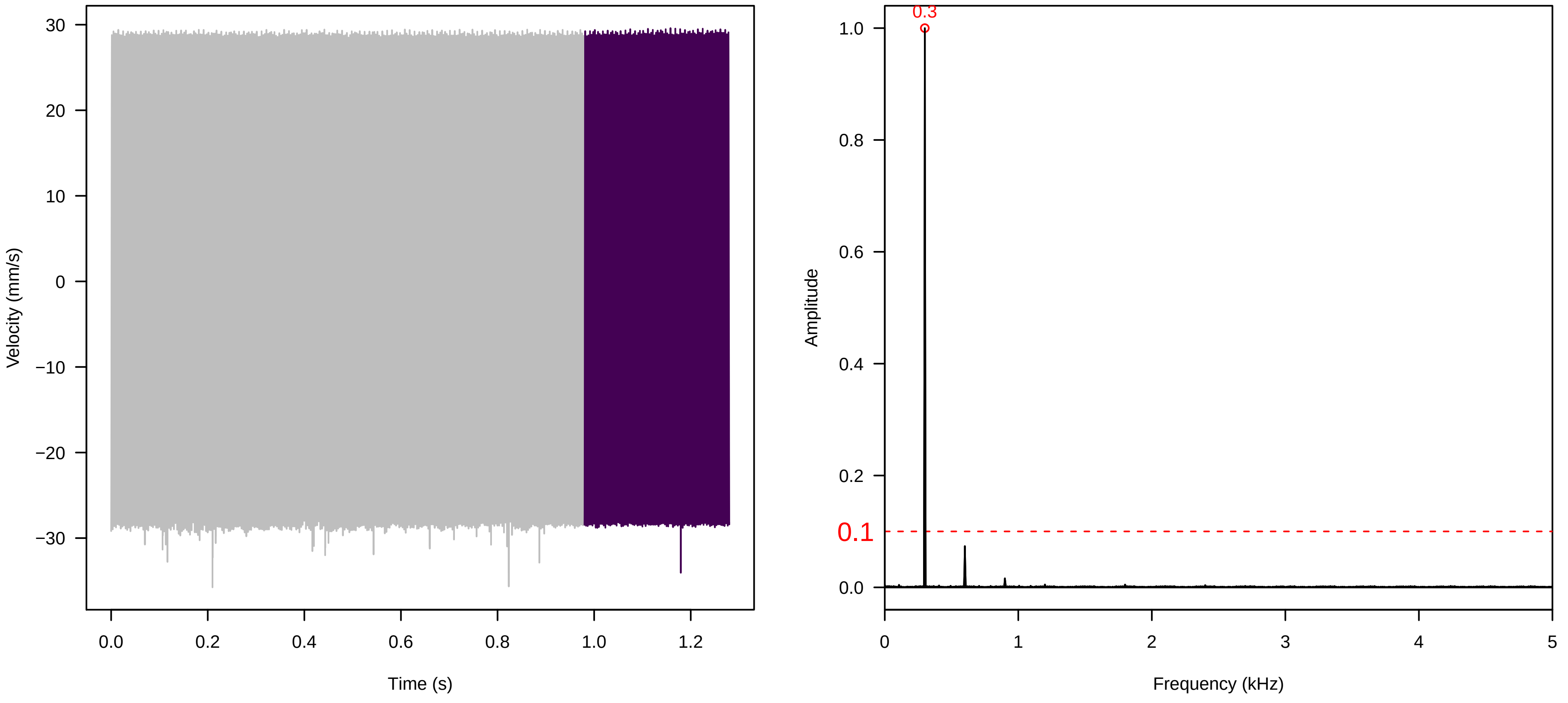


Vel. =  0.014 ; Str. =  Receptacle ; Axis =  z ; Fl. accession = 10−s−86

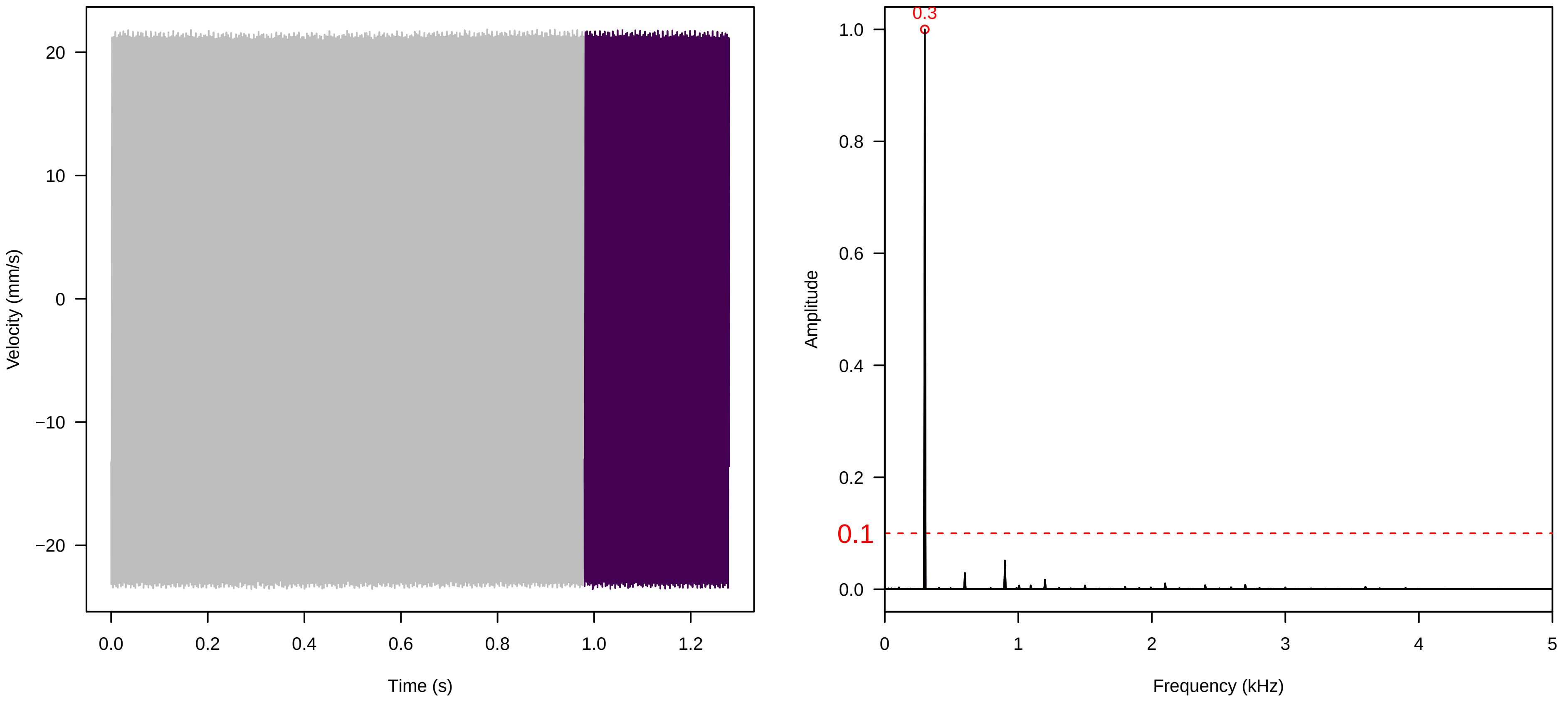


Vel. =  0.014 ; Str. =  FA ; Axis =  z ; Fl. accession = 10−s−86

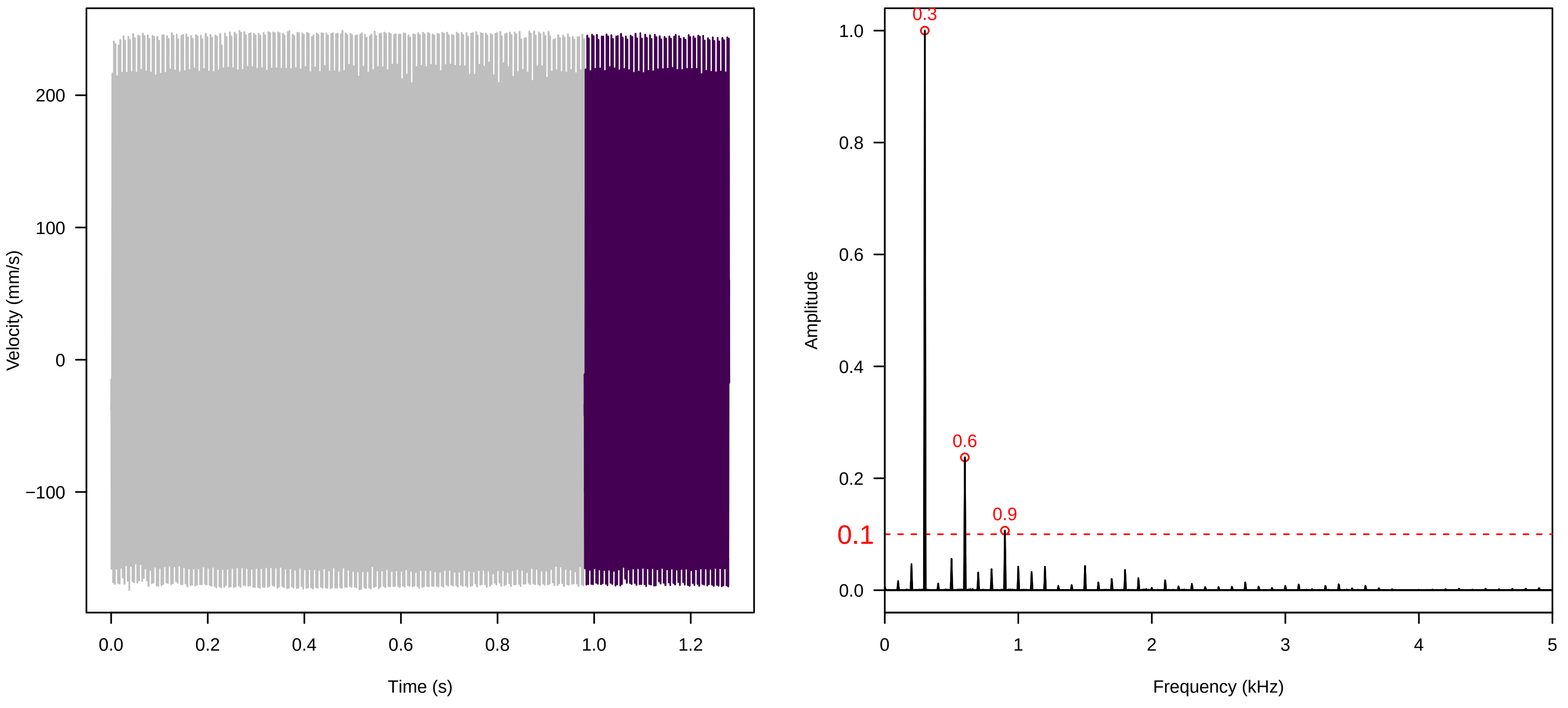

**Vel. =  0.014 ; Str. =  Receptacle ; Axis =  z ; Fl. accession = 10−s−86**

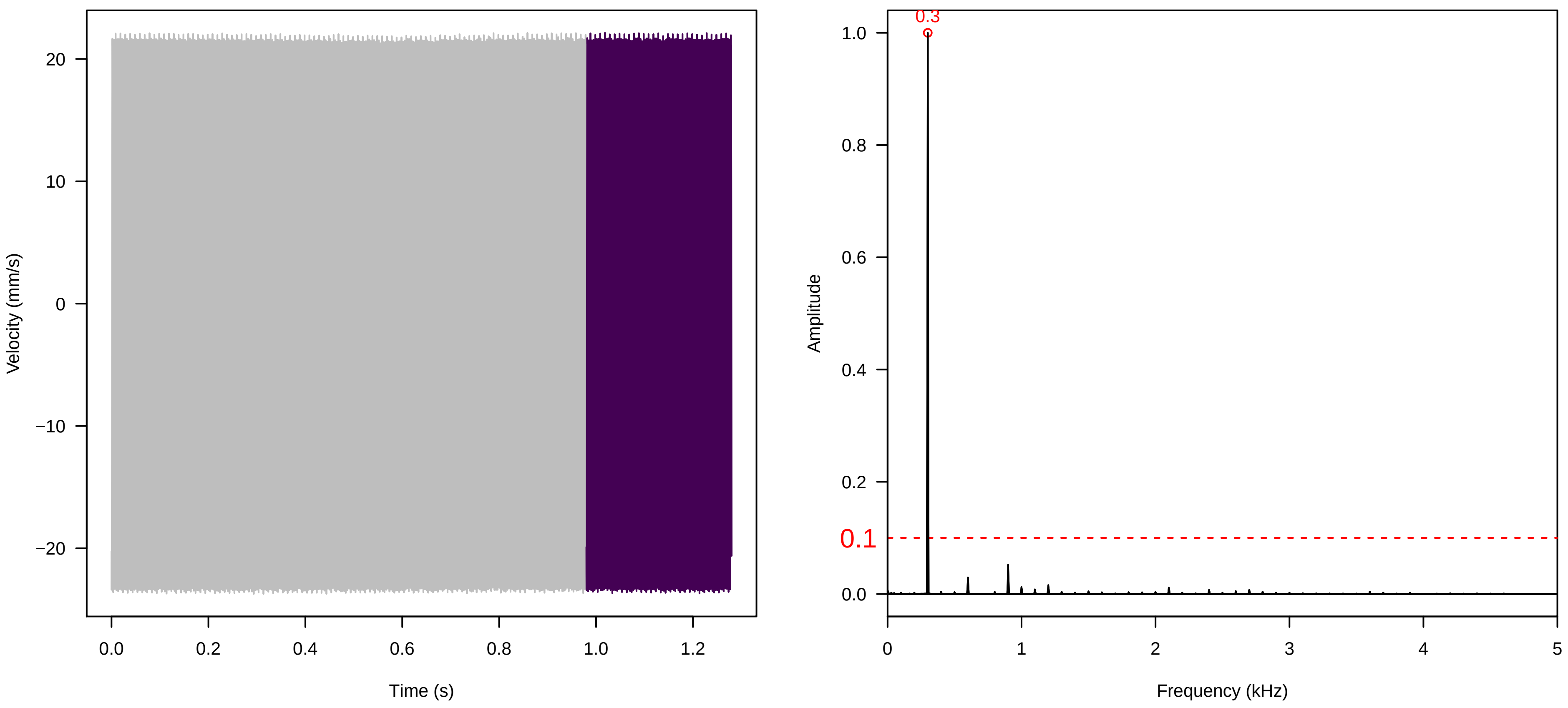


**Vel. =  0.014 ; Str. =  PA ; Axis =  z ; Fl. accession = 10−s−86**

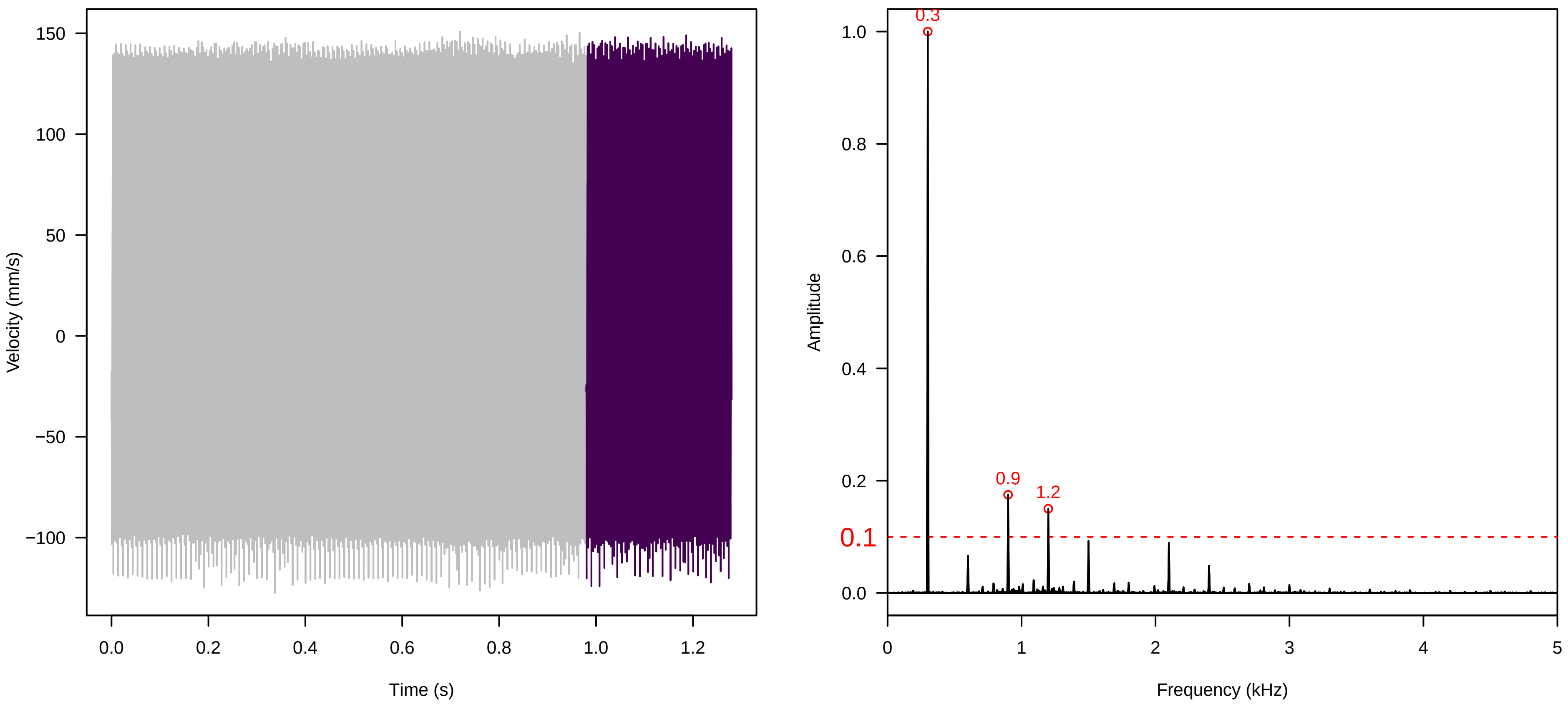


**Vel. =  0.014 ; Str. =  Receptacle ; Axis =  z ; Fl. accession = 10−s−86**

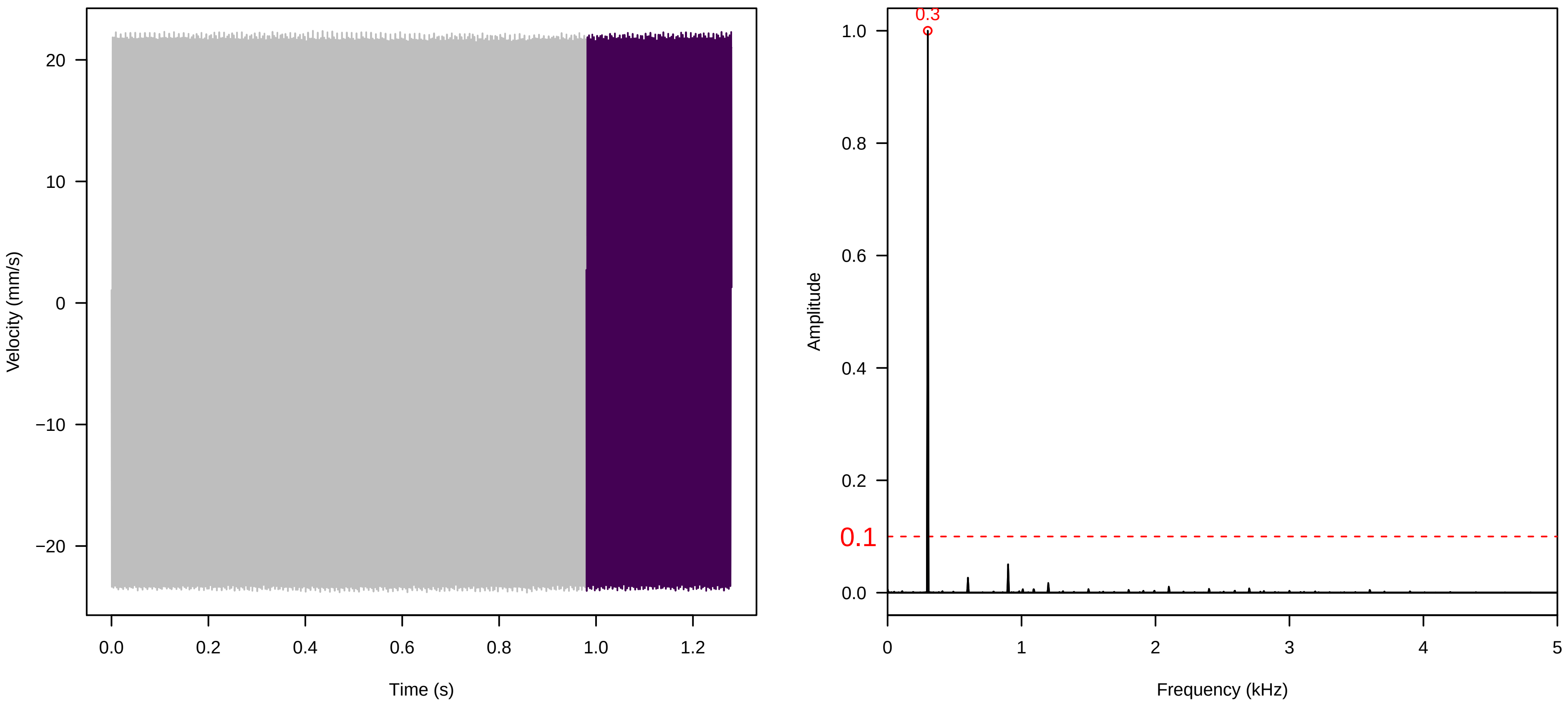

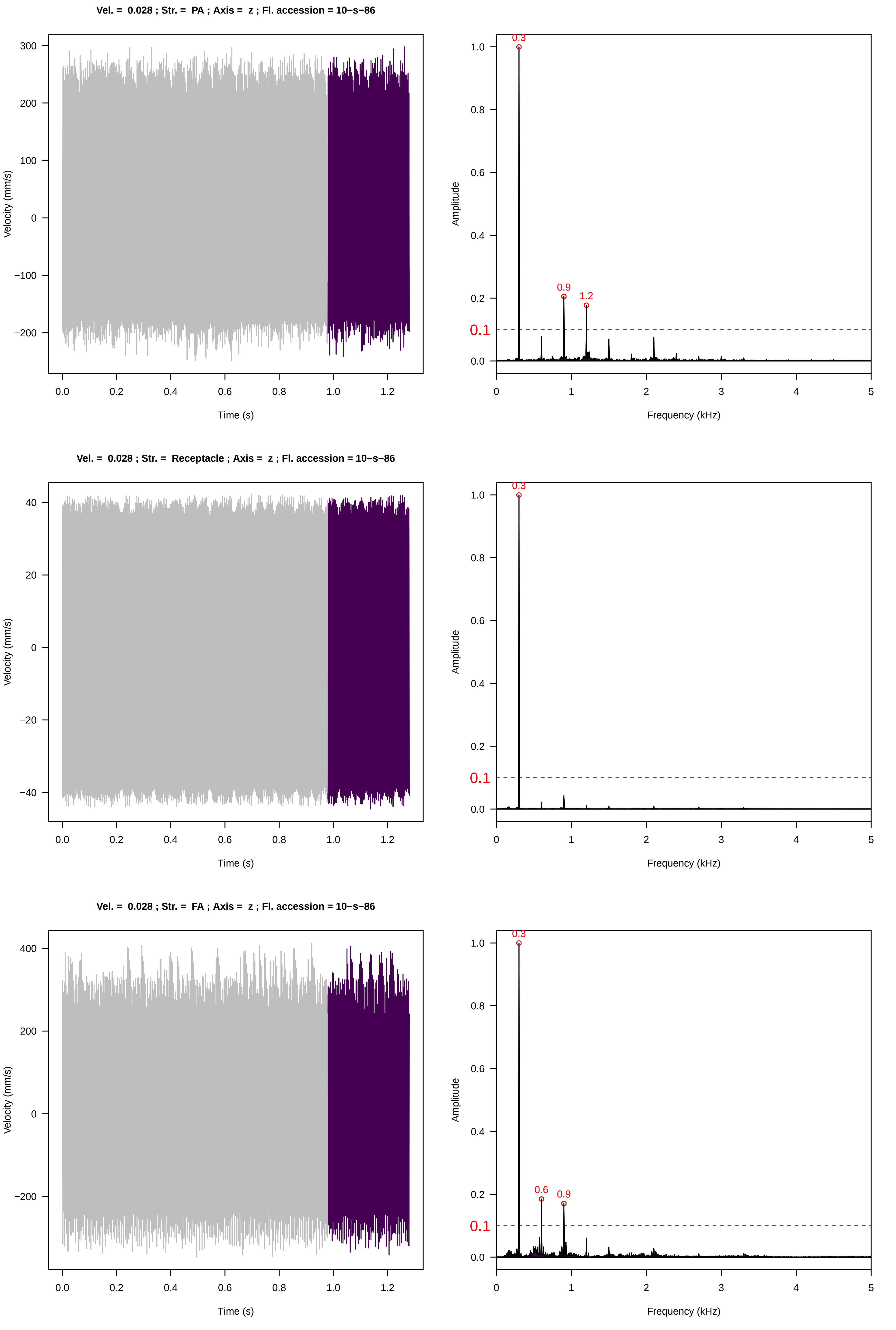

Vel. = 0.028 ; Str. = PA ; Axis = z ; Fl. accession = 10−s−86
Velocity (mm/s)
Time (s)
Amplitude
Frequency (kHz)
0.3
0.9
1.2
0.1
Vel. = 0.028 ; Str. = Receptacle ; Axis = z ; Fl. accession = 10−s−86
Velocity (mm/s)
Time (s)
Amplitude
Frequency (kHz)
0.3
0.1
Vel. = 0.028 ; Str. = FA ; Axis = z ; Fl. accession = 10−s−86
Velocity (mm/s)
Time (s)
Amplitude
Frequency (kHz)
0.3
0.6
0.9
0.1

**Vel. = 0.028 ; Str. = Receptacle ; Axis = z ; Fl. accession = 10−s−86**

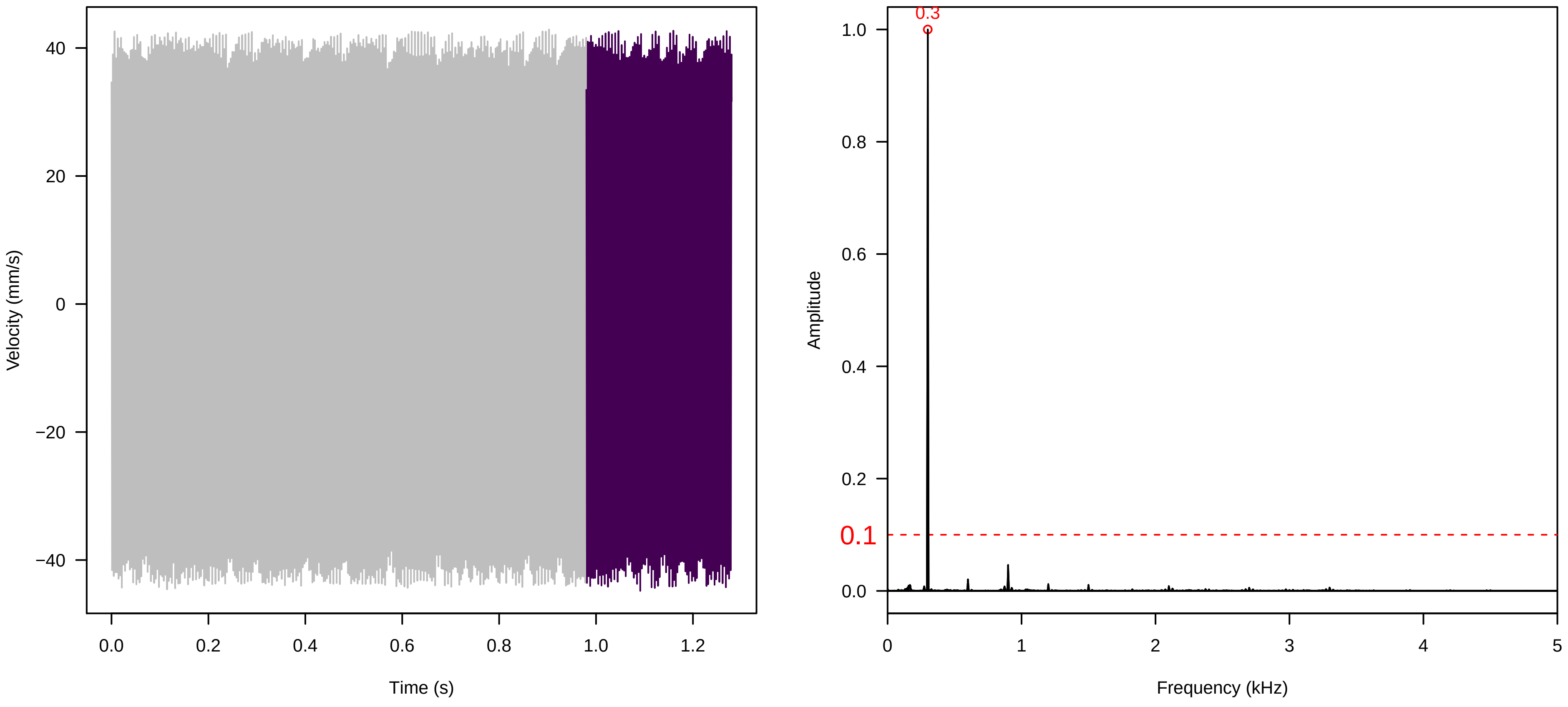


**Vel. = 0.028 ; Str. = Corolla ; Axis = z ; Fl. accession = 10−s−86**

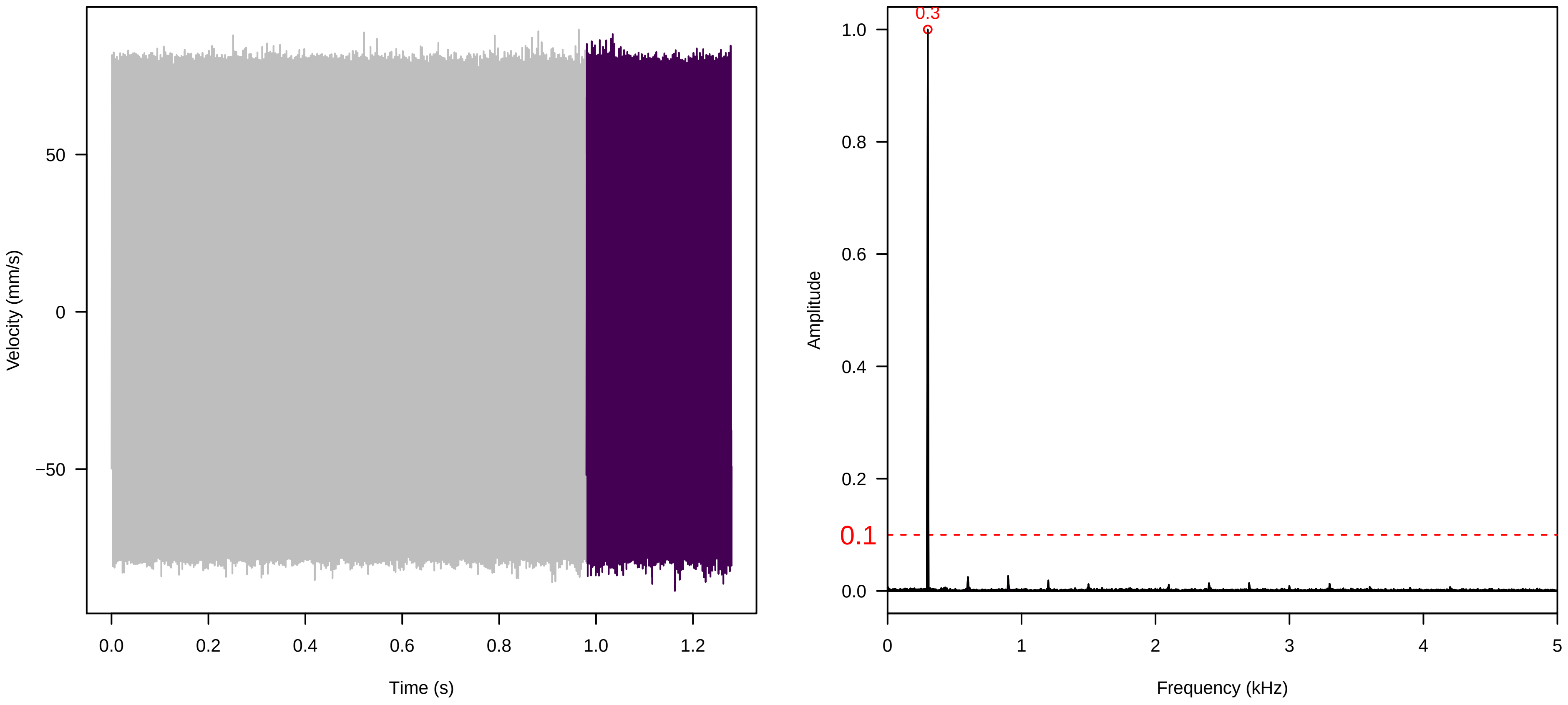


**Vel. = 0.028 ; Str. = Receptacle ; Axis = z ; Fl. accession = 10−s−86**

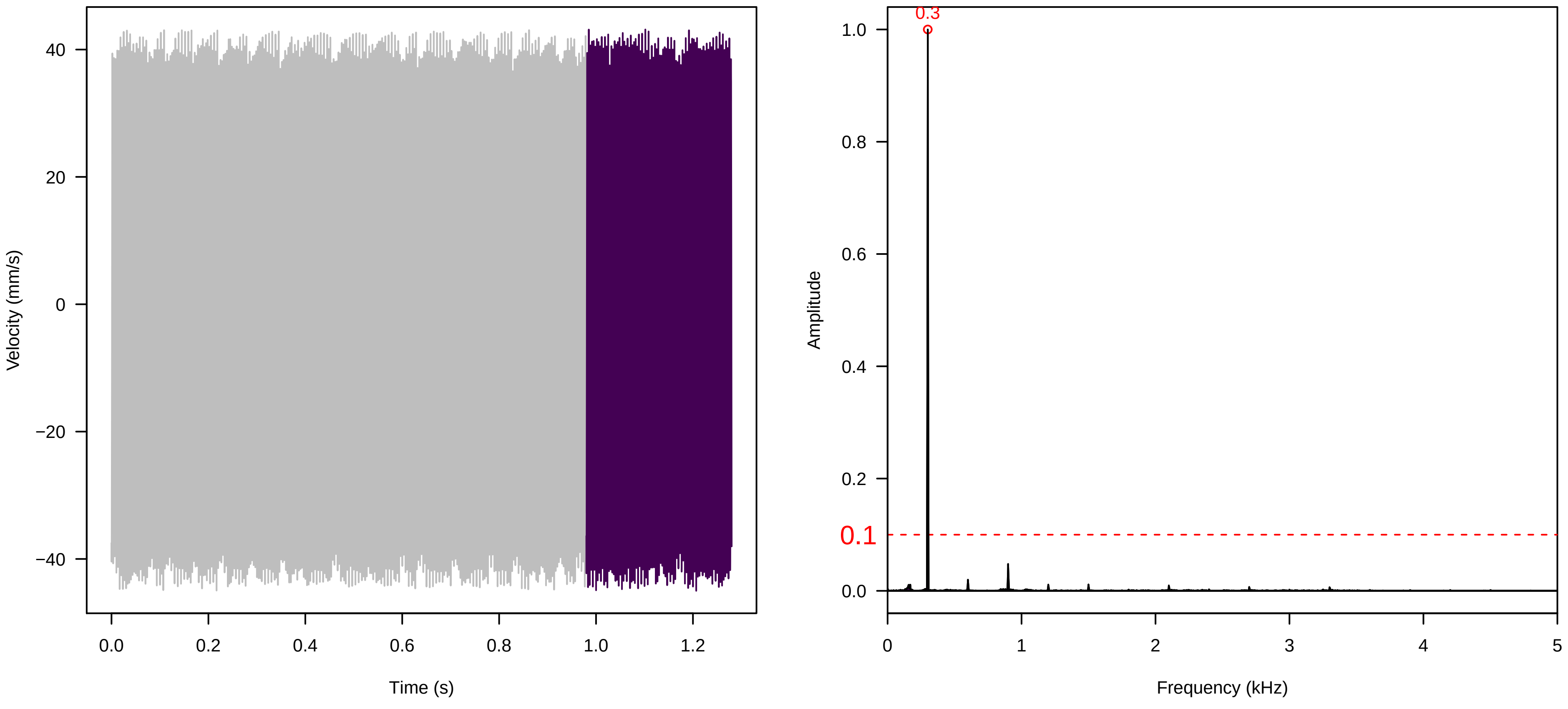

**Vel. = 0.057 ; Str. = Corolla ; Axis = z ; Fl. accession = 10−s−86**

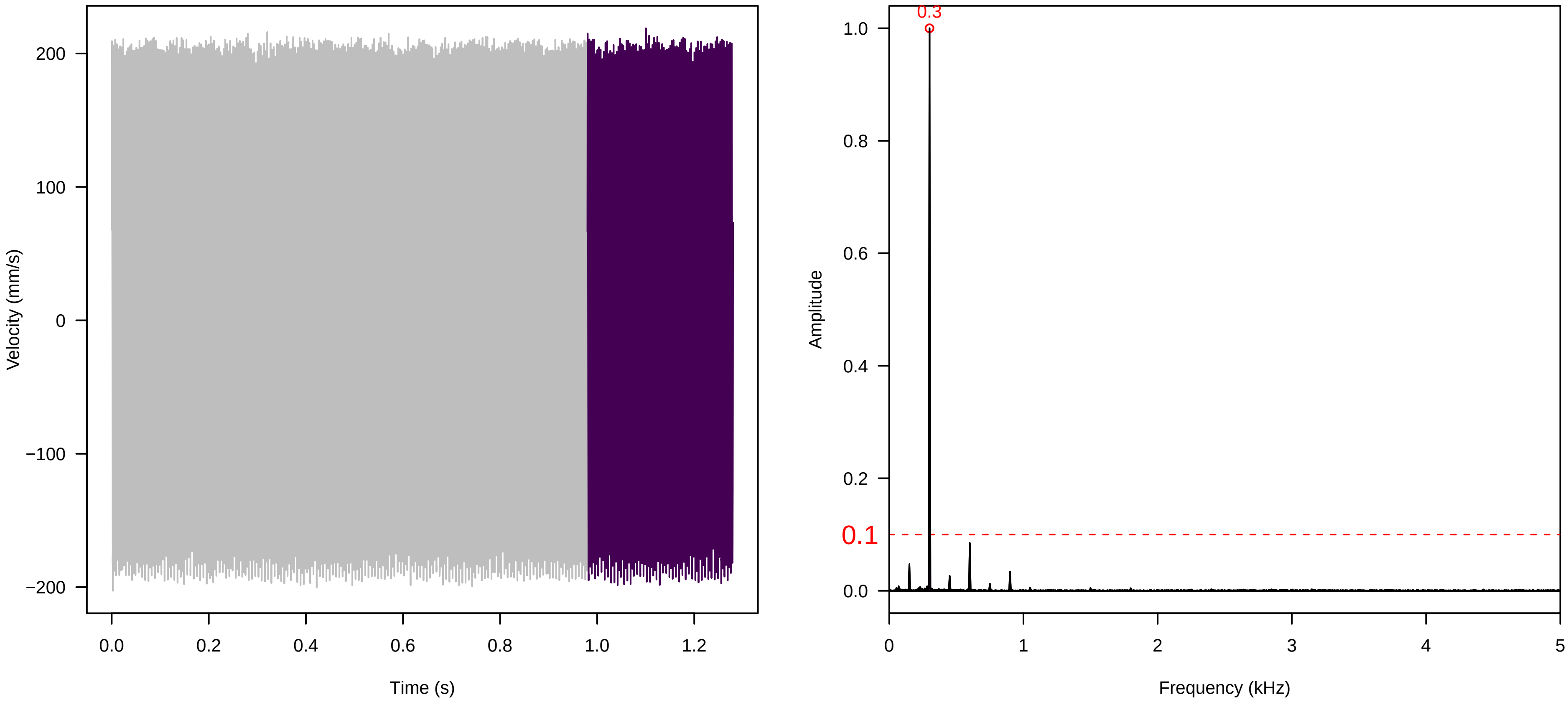


**Vel. = 0.057 ; Str. = Receptacle ; Axis = z ; Fl. accession = 10−s−86**

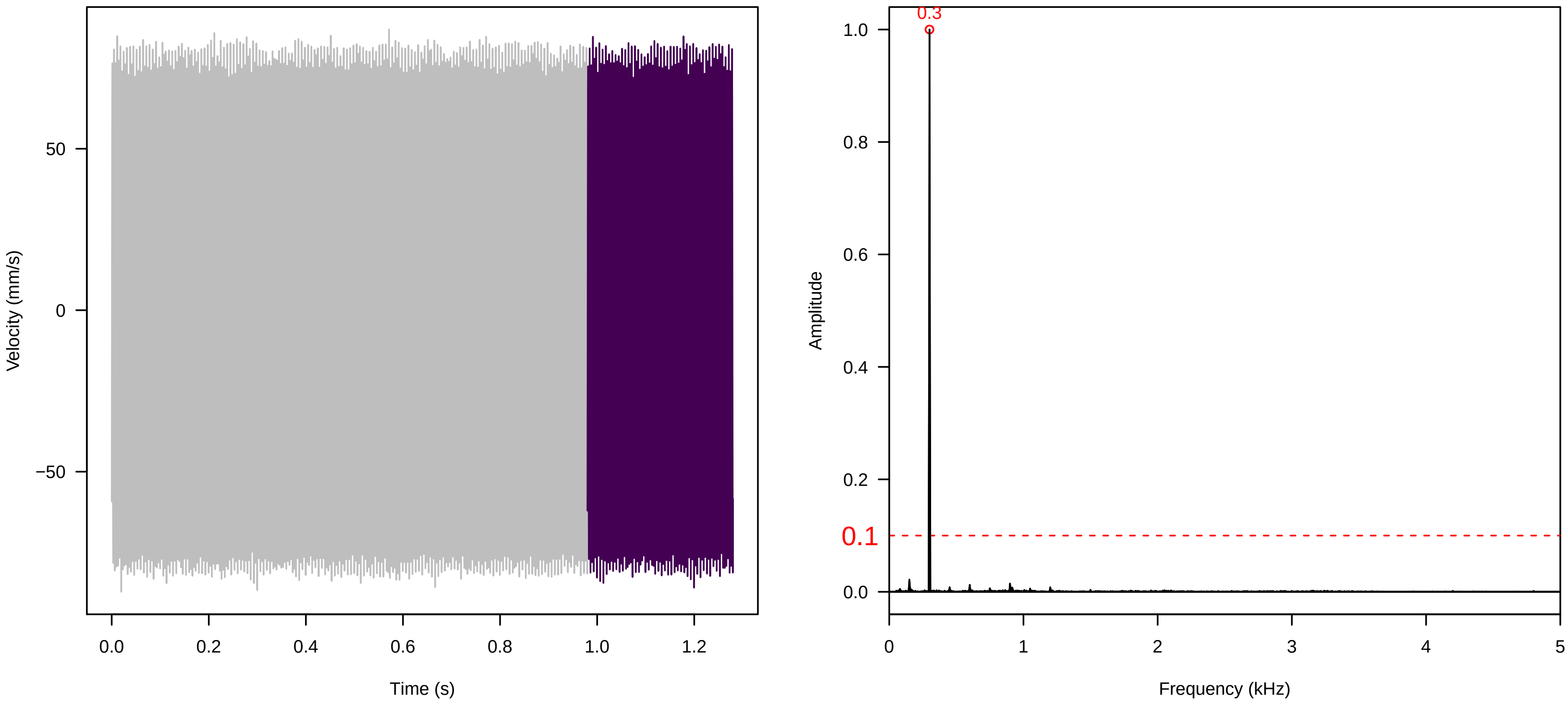


**Vel. = 0.057 ; Str. = FA ; Axis = z ; Fl. accession = 10−s−86**

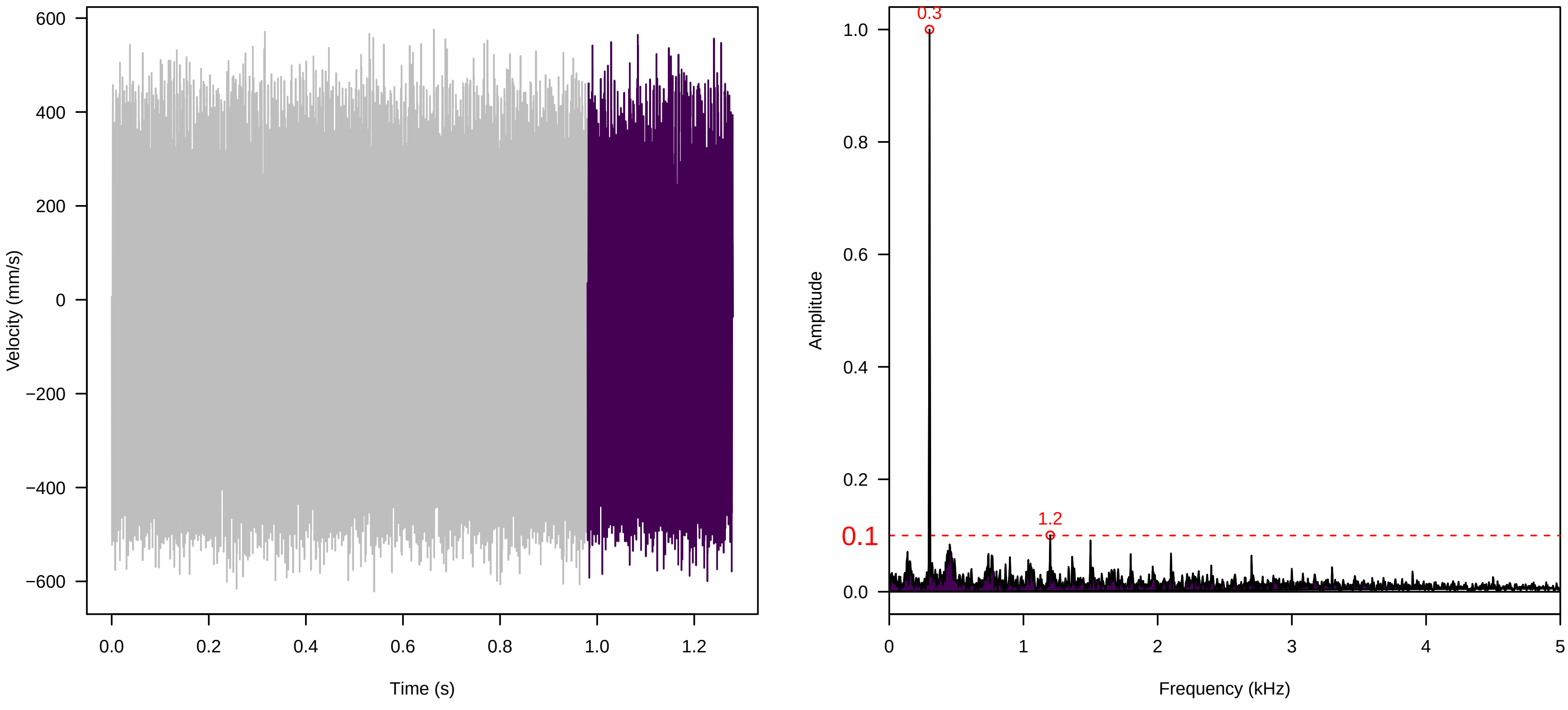

**Vel. = 0.057 ; Str. = Receptacle ; Axis = z ; Fl. accession = 10−s−86**

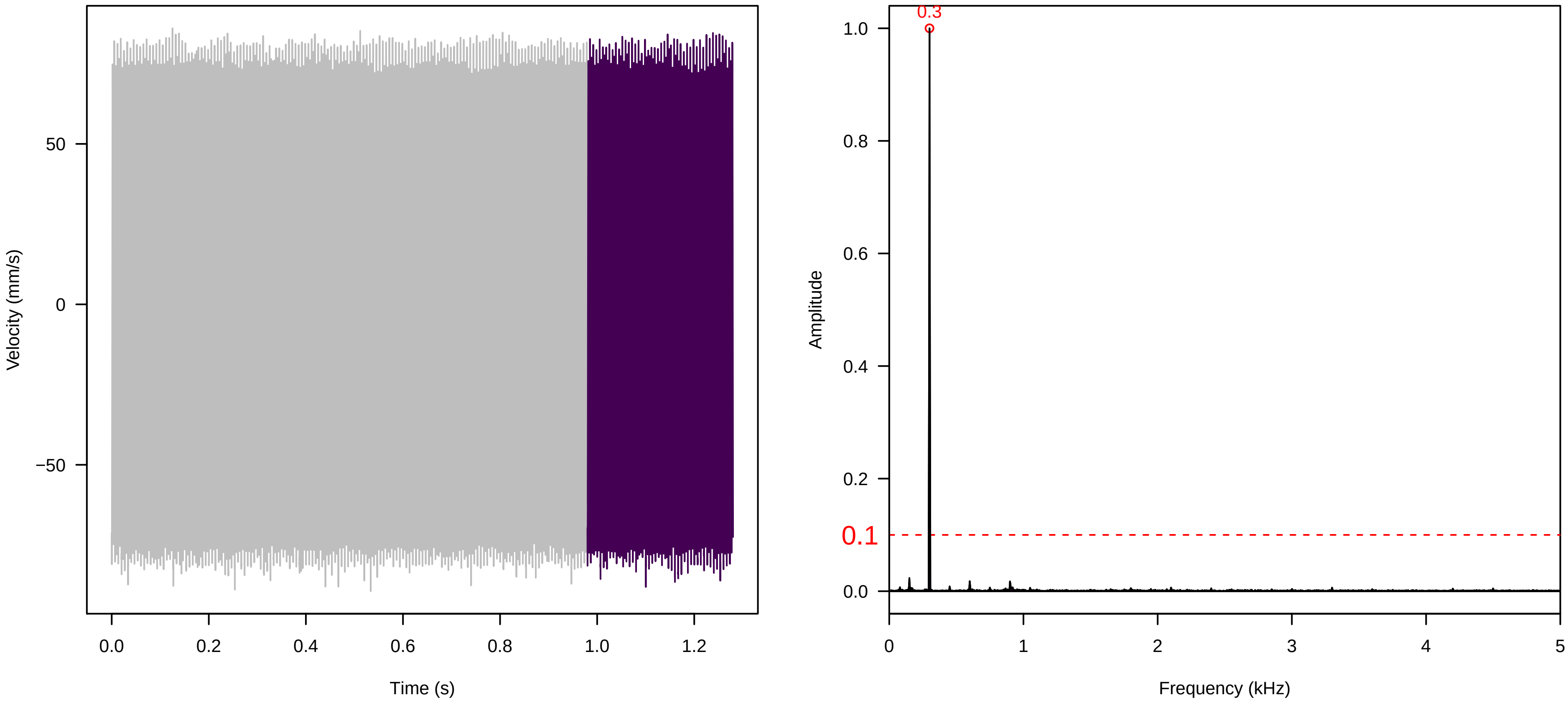


**Vel. = 0.057 ; Str. = PA ; Axis = z ; Fl. accession = 10−s−86**

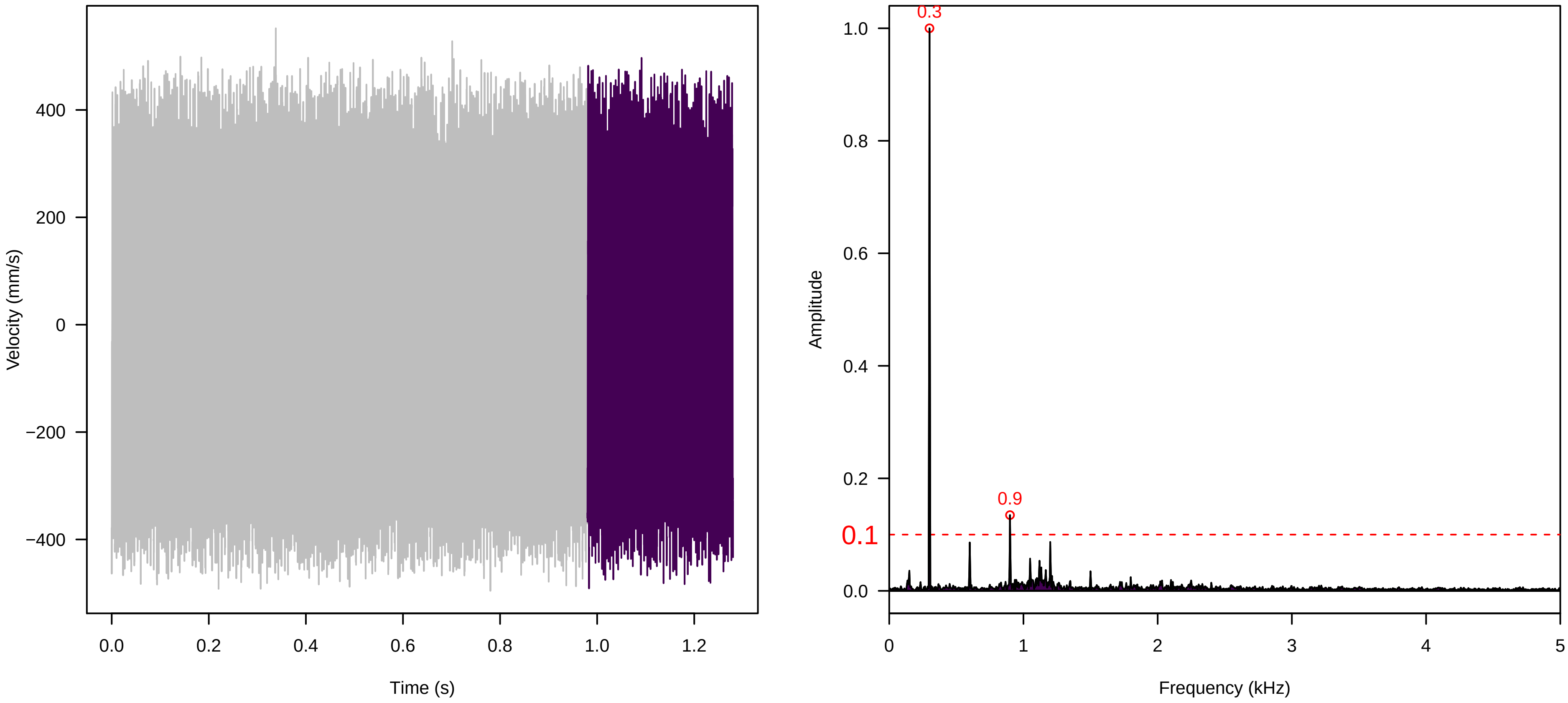


**Vel. = 0.057 ; Str. = Receptacle ; Axis = z ; Fl. accession = 10−s−86**

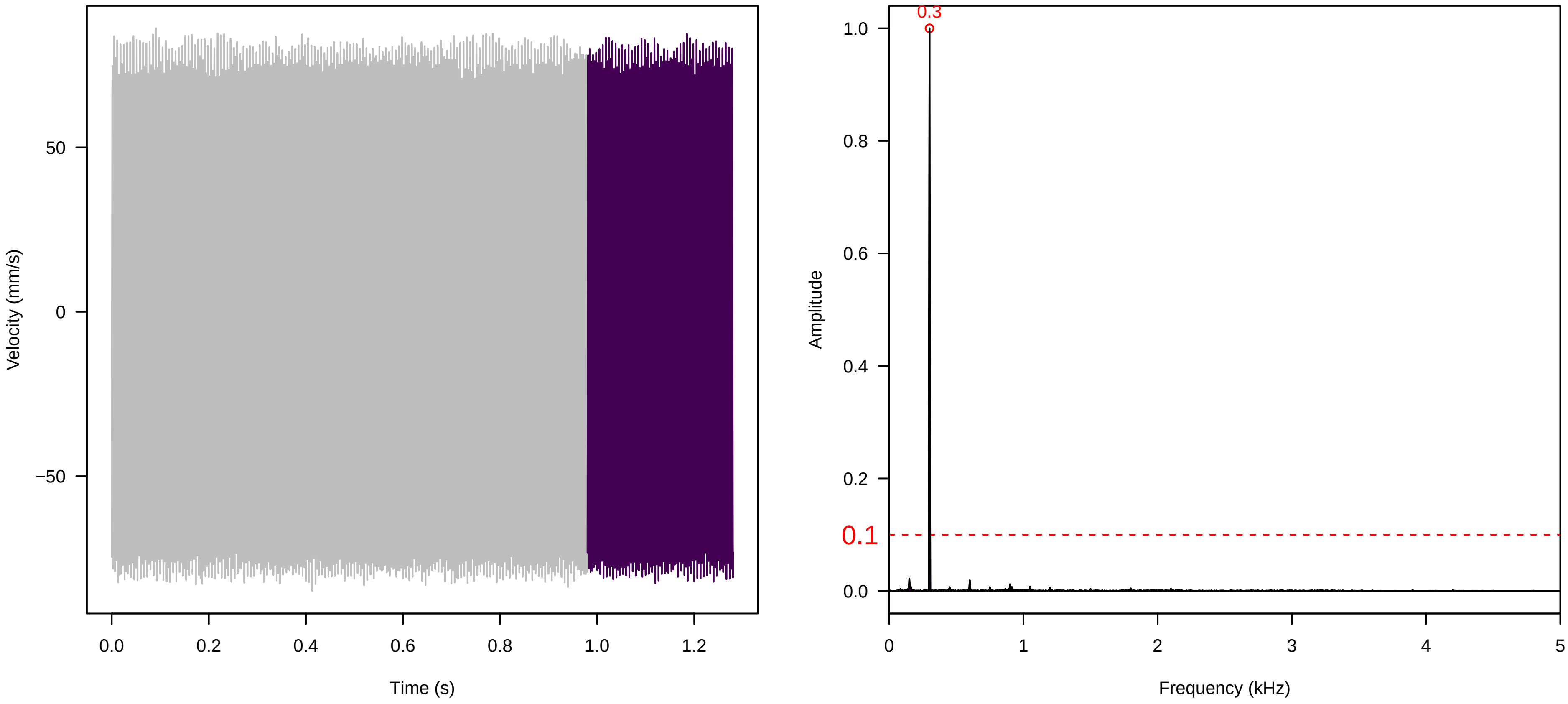

**Vel. =  0.014 ; Str. =  Corolla ; Axis =  x ; Fl. accession = 10−s−86**

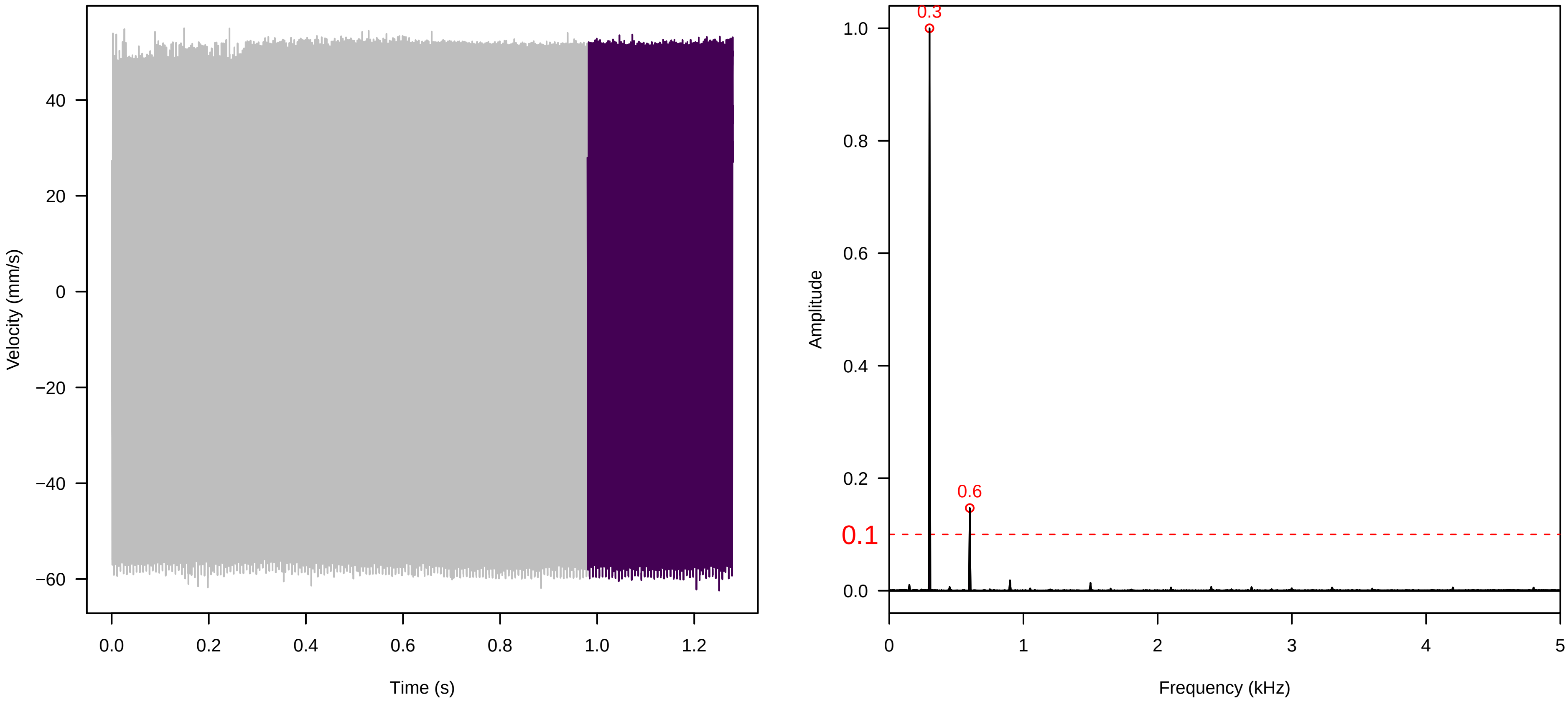


**Vel. =  0.014 ; Str. =  Receptacle ; Axis =  x ; Fl. accession = 10−s−86**

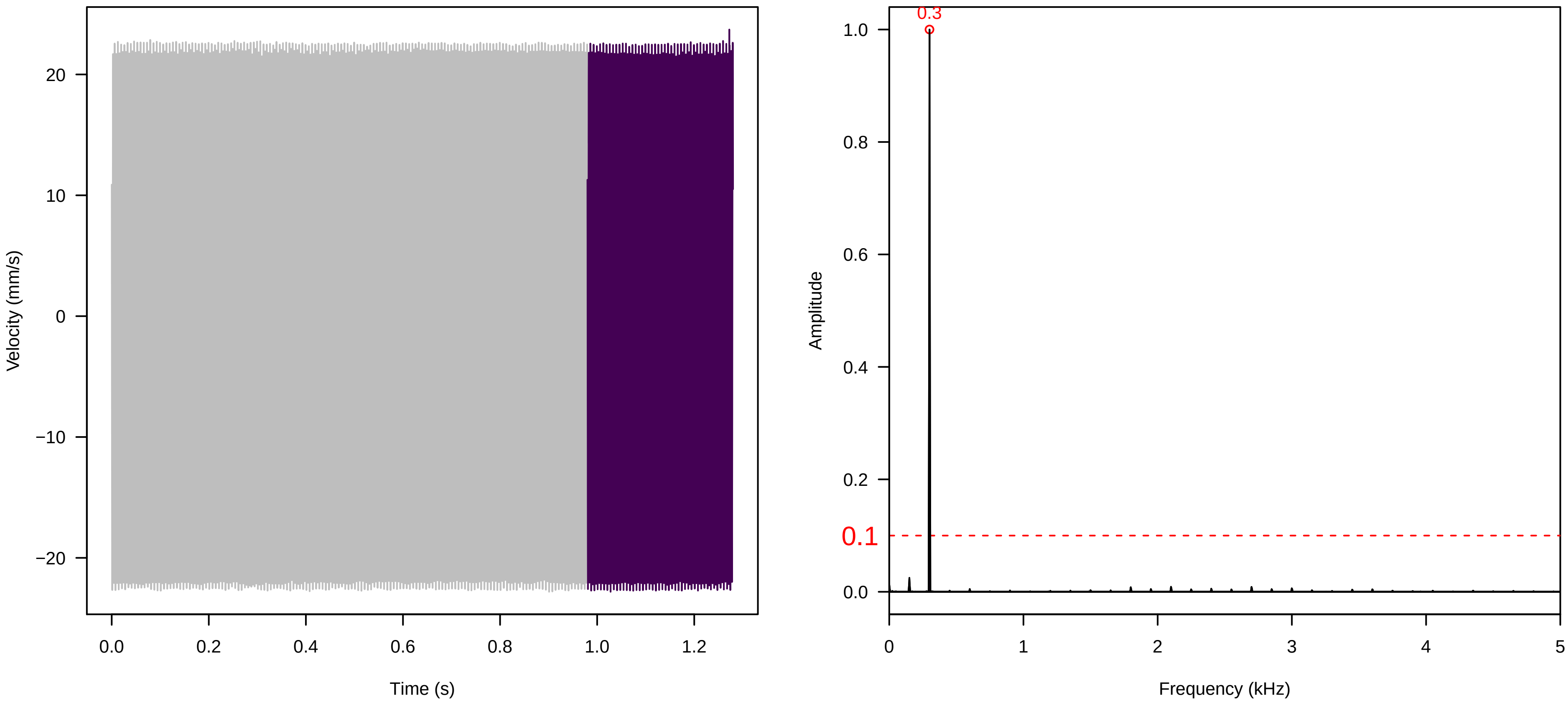


**Vel. =  0.014 ; Str. =  FA ; Axis =  x ; Fl. accession = 10−s−86**

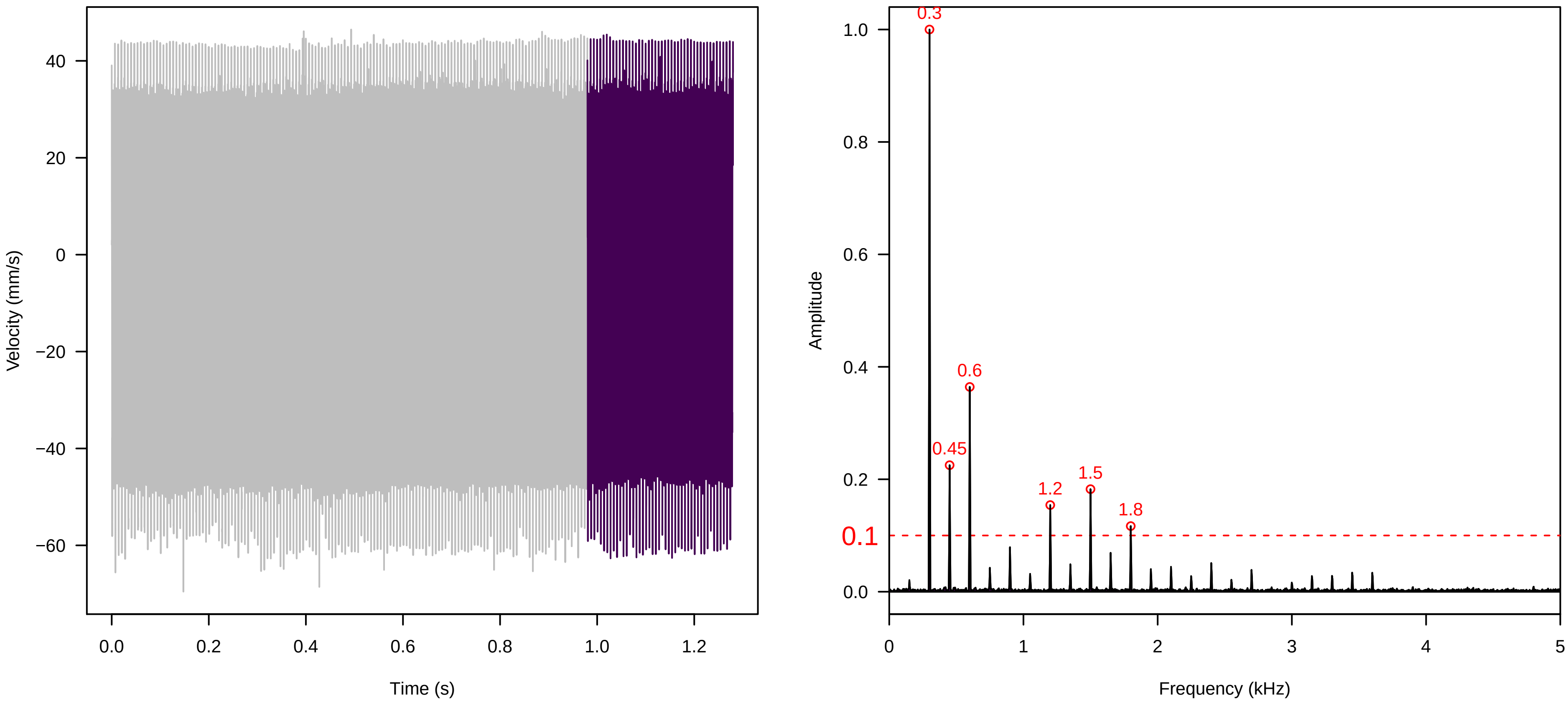

**Vel. = 0.014 ; Str. = Receptacle ; Axis = x ; Fl. accession = 10−s−86**

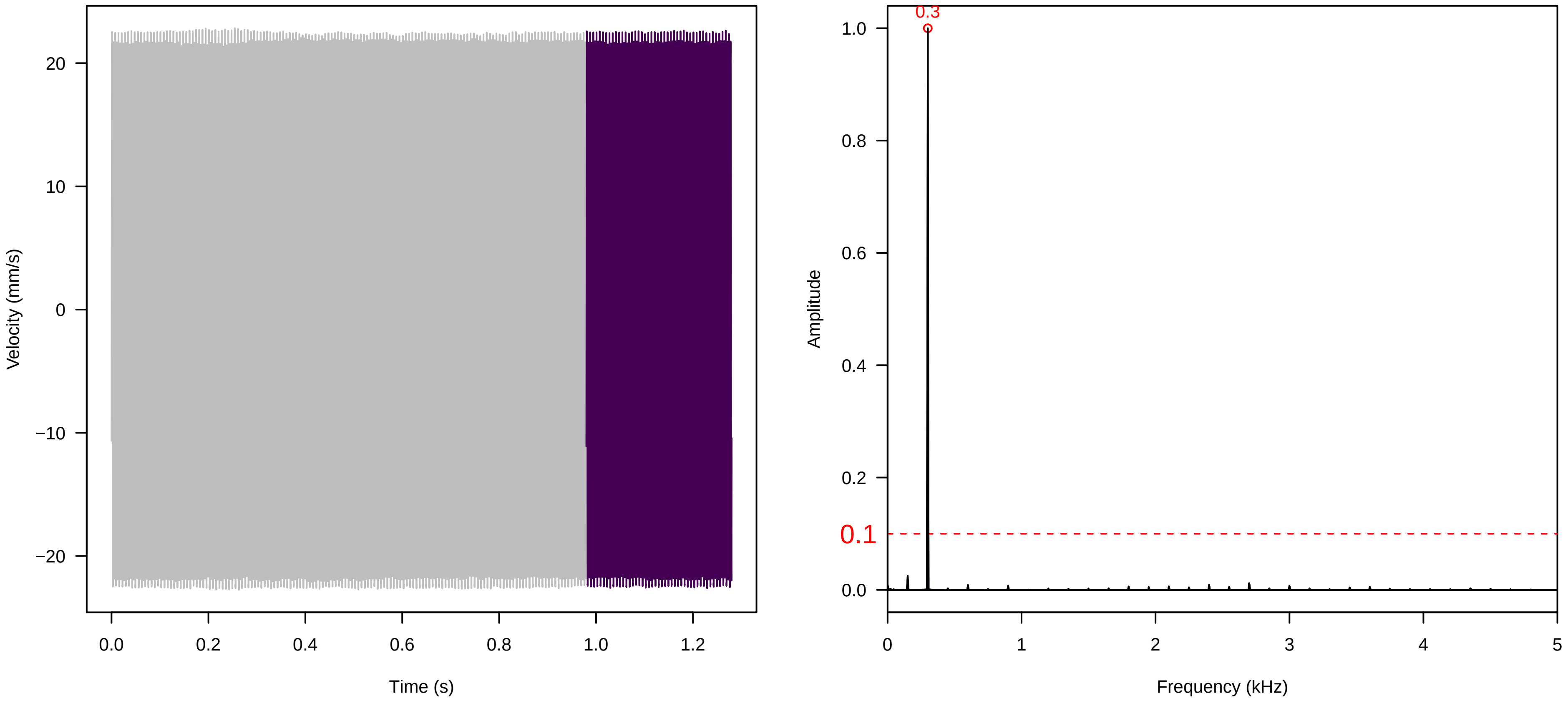


**Vel. = 0.014 ; Str. = PA ; Axis = x ; Fl. accession = 10−s−86**

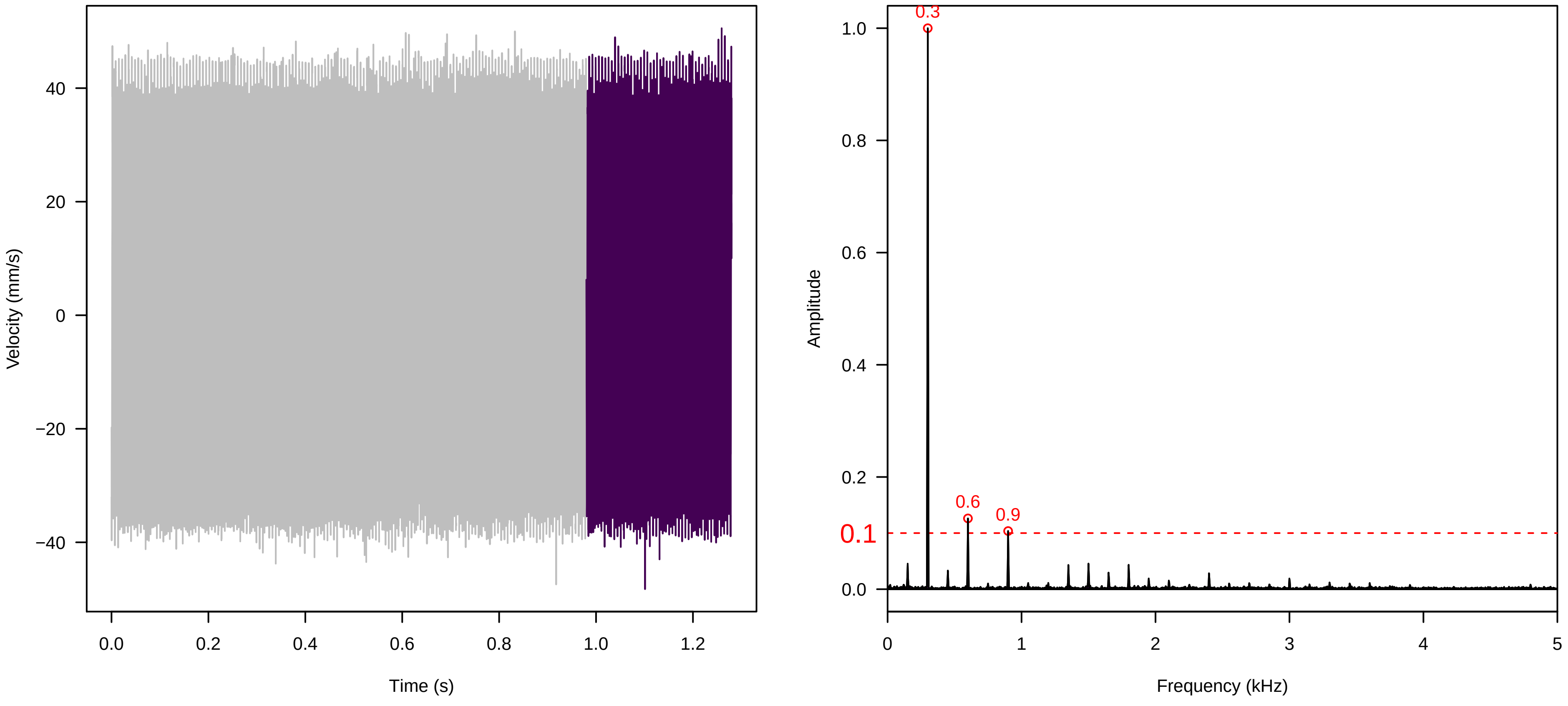


**Vel. = 0.014 ; Str. = Receptacle ; Axis = x ; Fl. accession = 10−s−86**

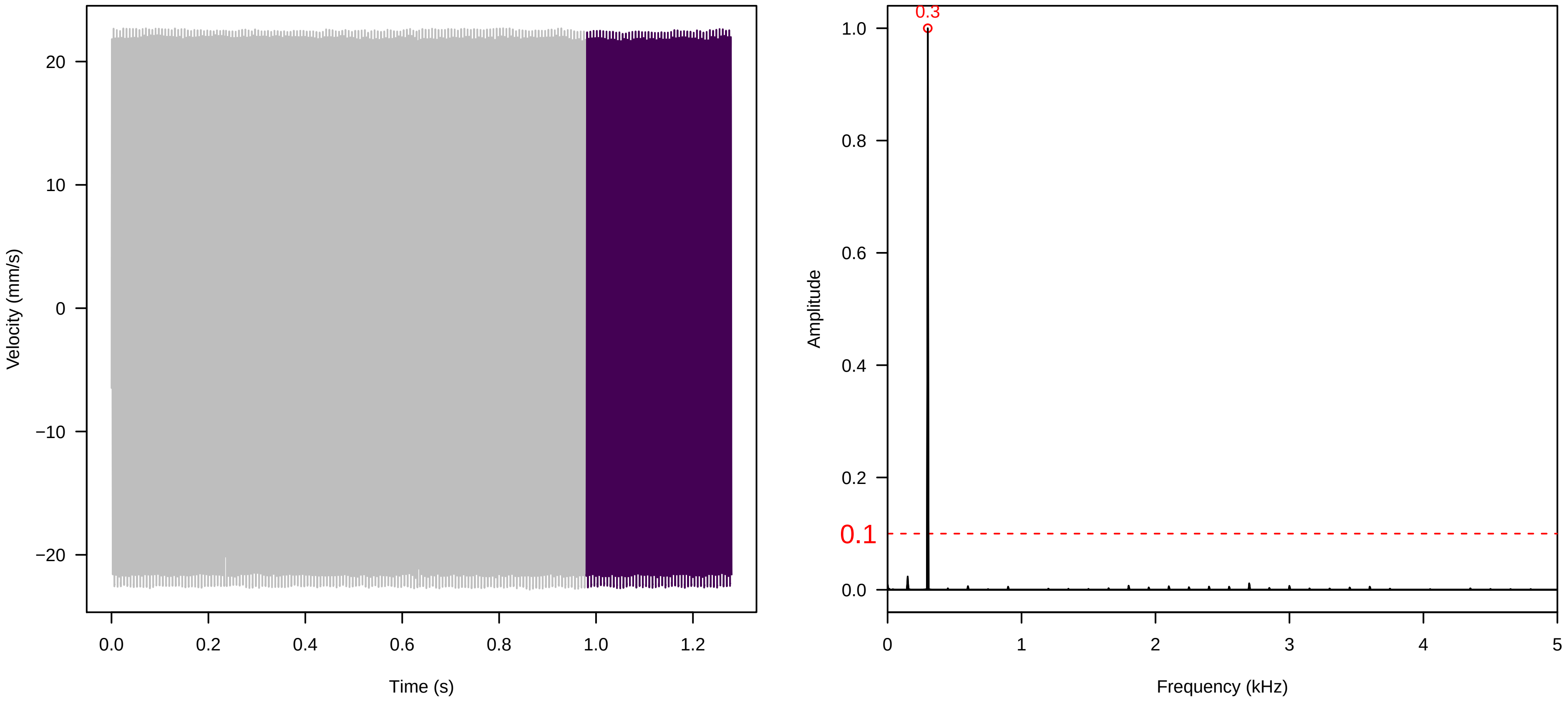

**Vel. = 0.028 ; Str. = PA ; Axis = x ; Fl. accession = 10−s−86**

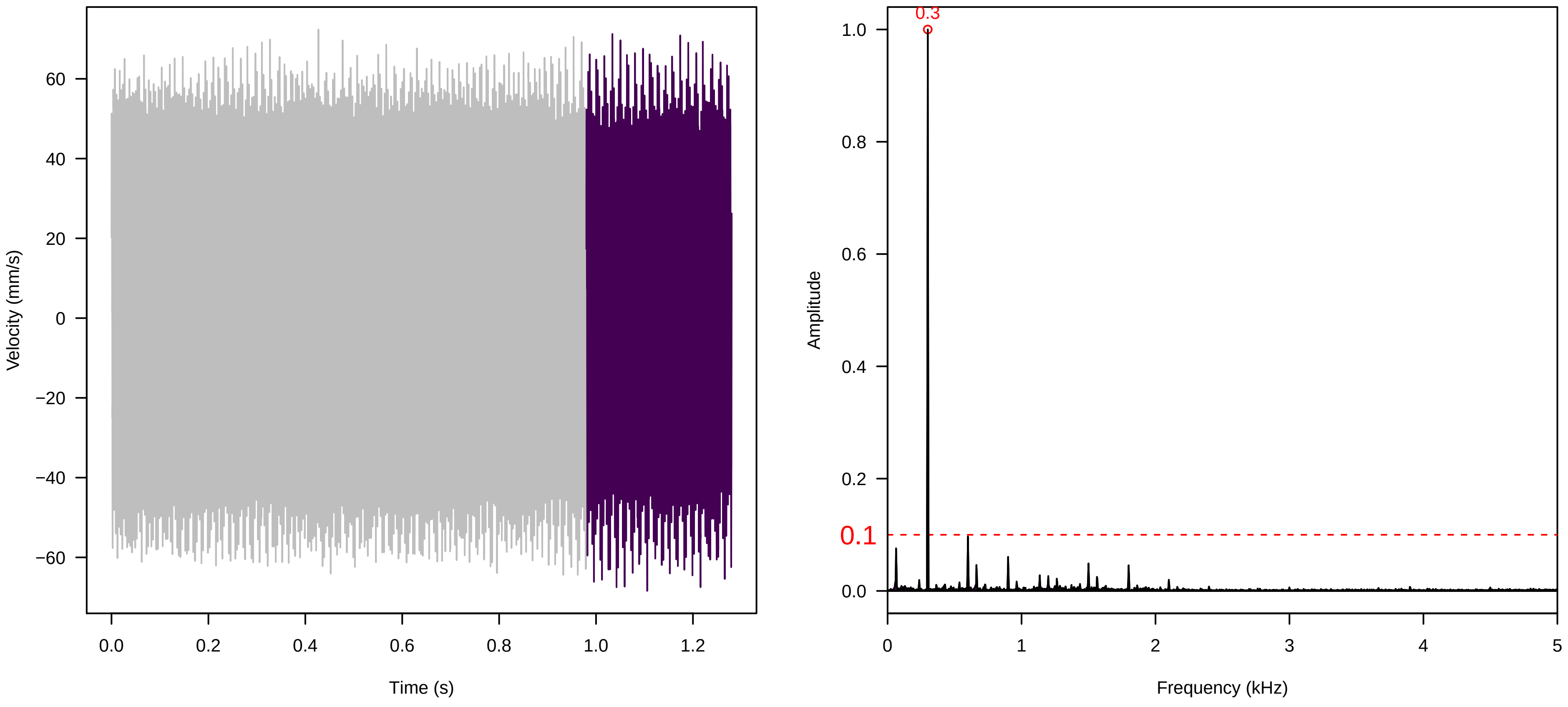


**Vel. = 0.028 ; Str. = Receptacle ; Axis = x ; Fl. accession = 10−s−86**

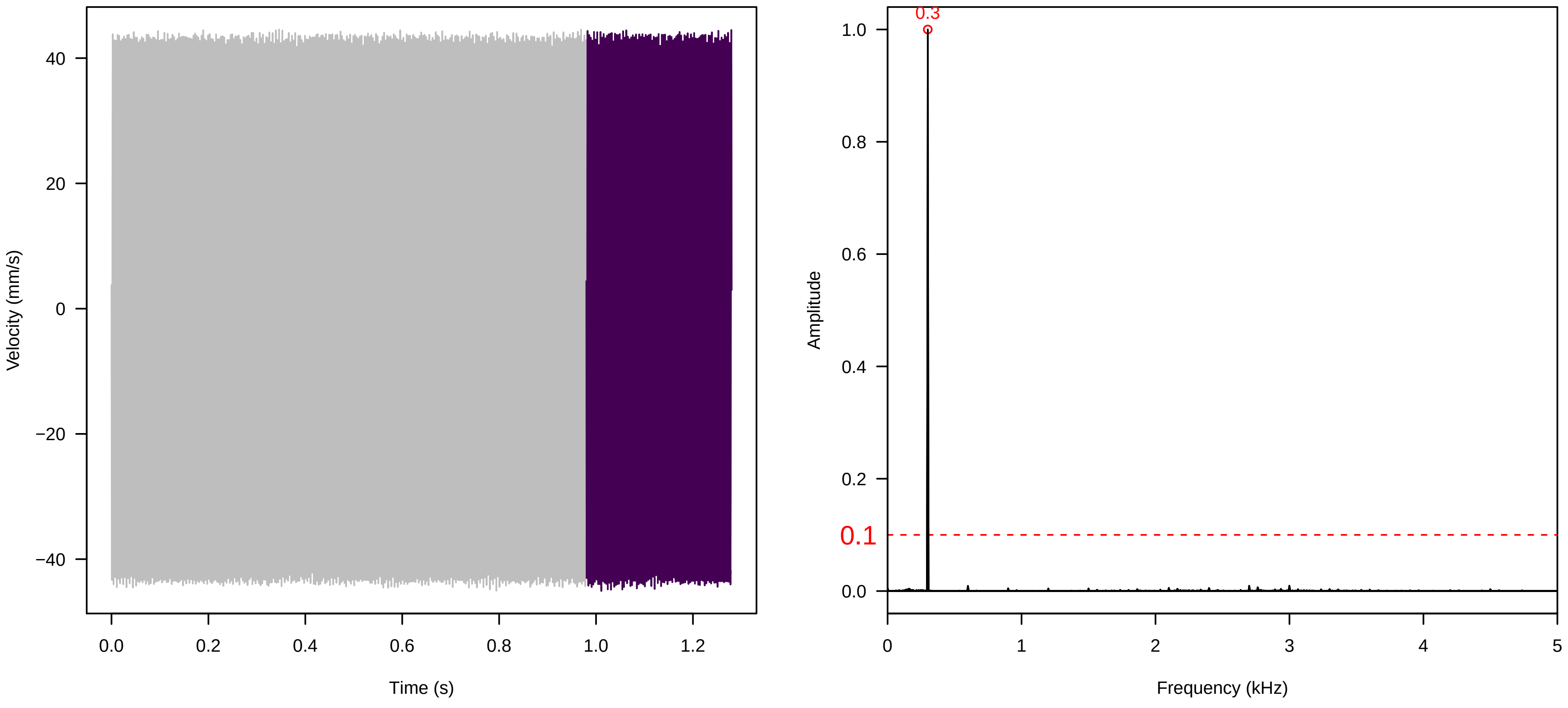


**Vel. = 0.028 ; Str. = FA ; Axis = x ; Fl. accession = 10−s−86**

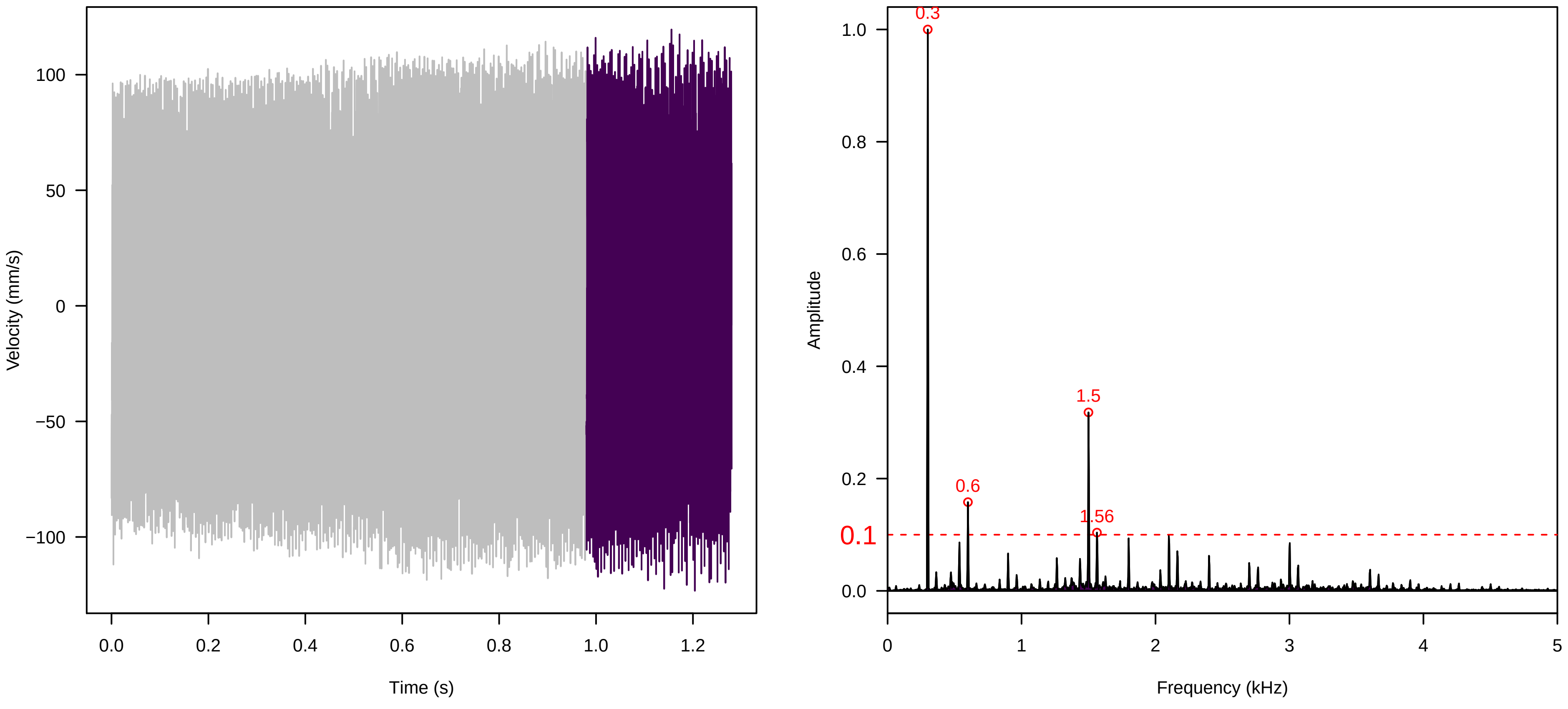

**Vel. =  0.028 ; Str. =  Receptacle ; Axis =  x ; Fl. accession = 10−s−86**

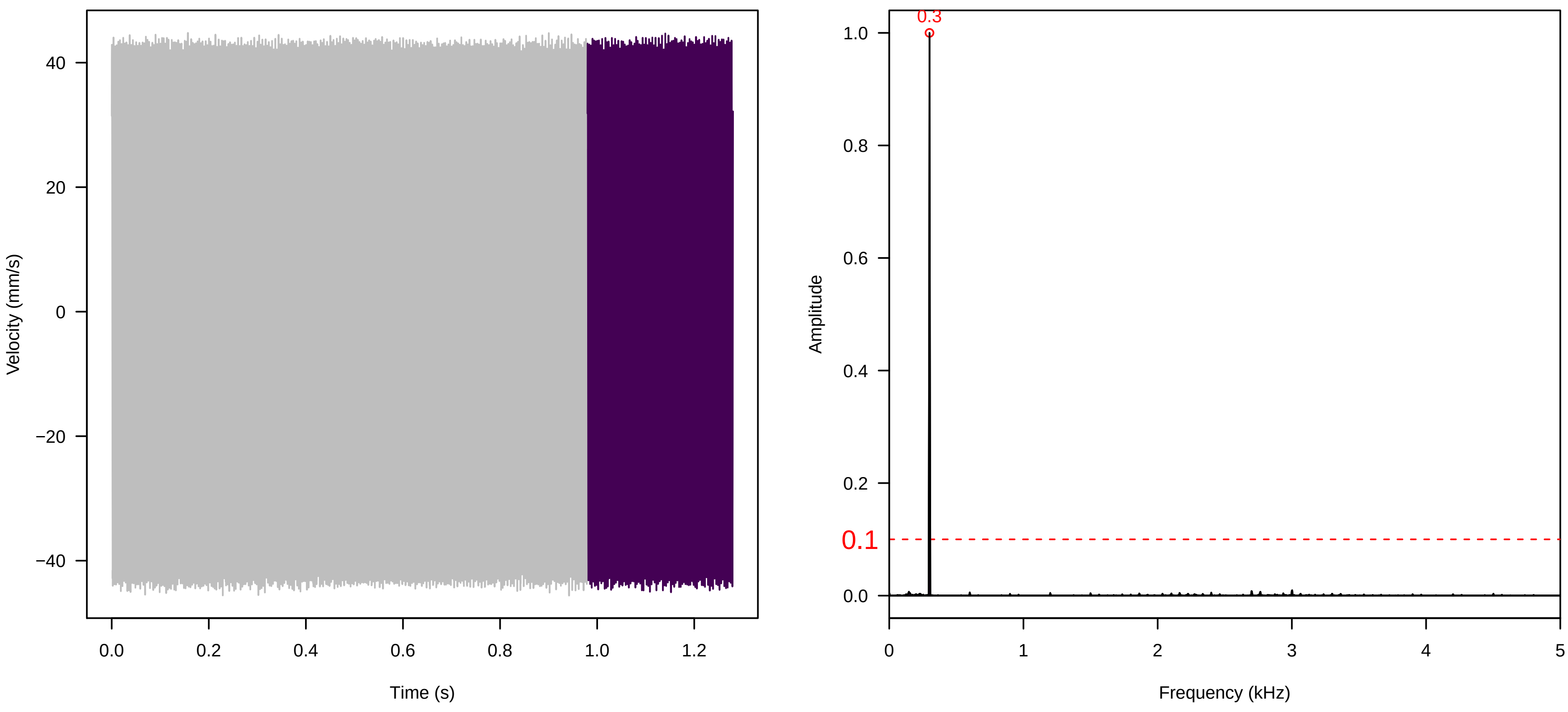


**Vel. =  0.028 ; Str. =  Corolla ; Axis =  x ; Fl. accession = 10−s−86**

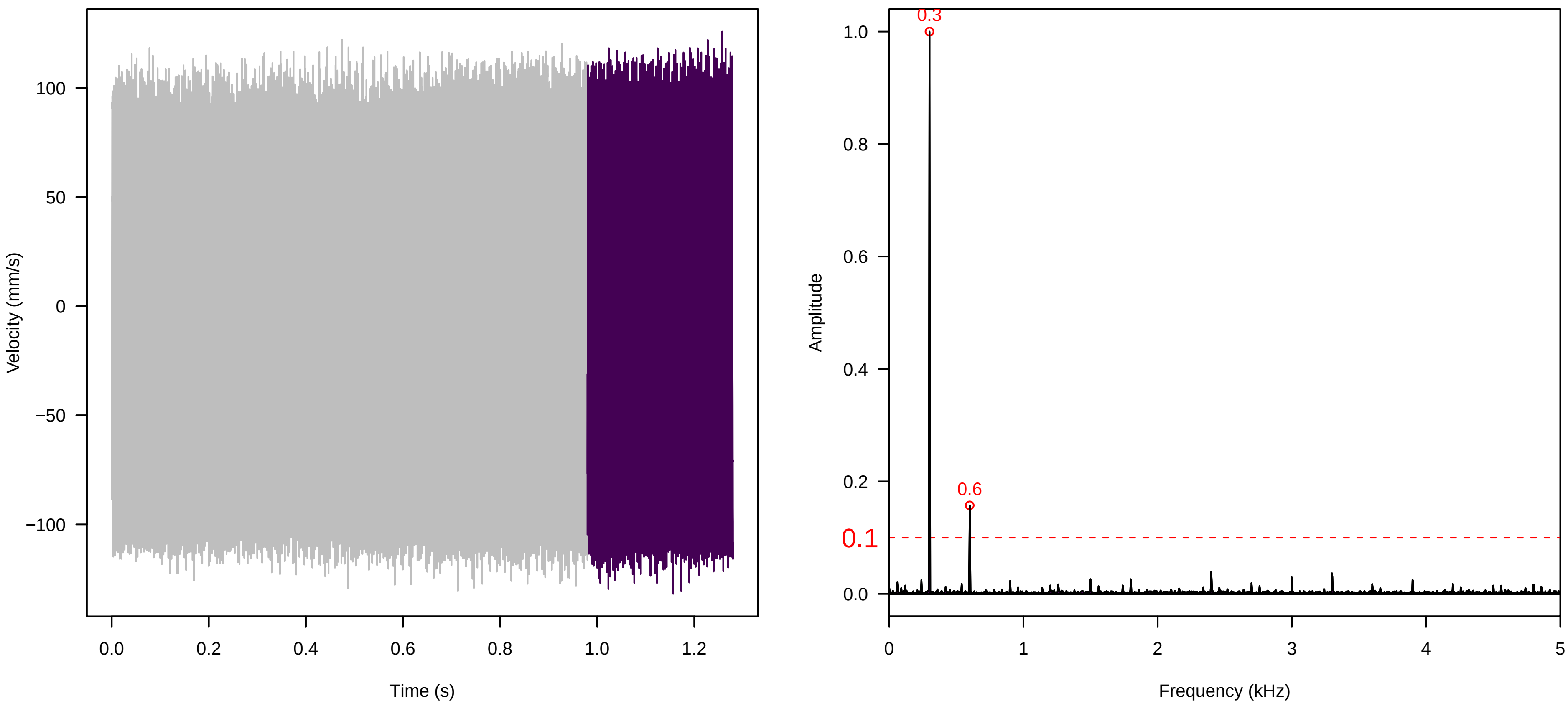


**Vel. =  0.028 ; Str. =  Receptacle ; Axis =  x ; Fl. accession = 10−s−86**

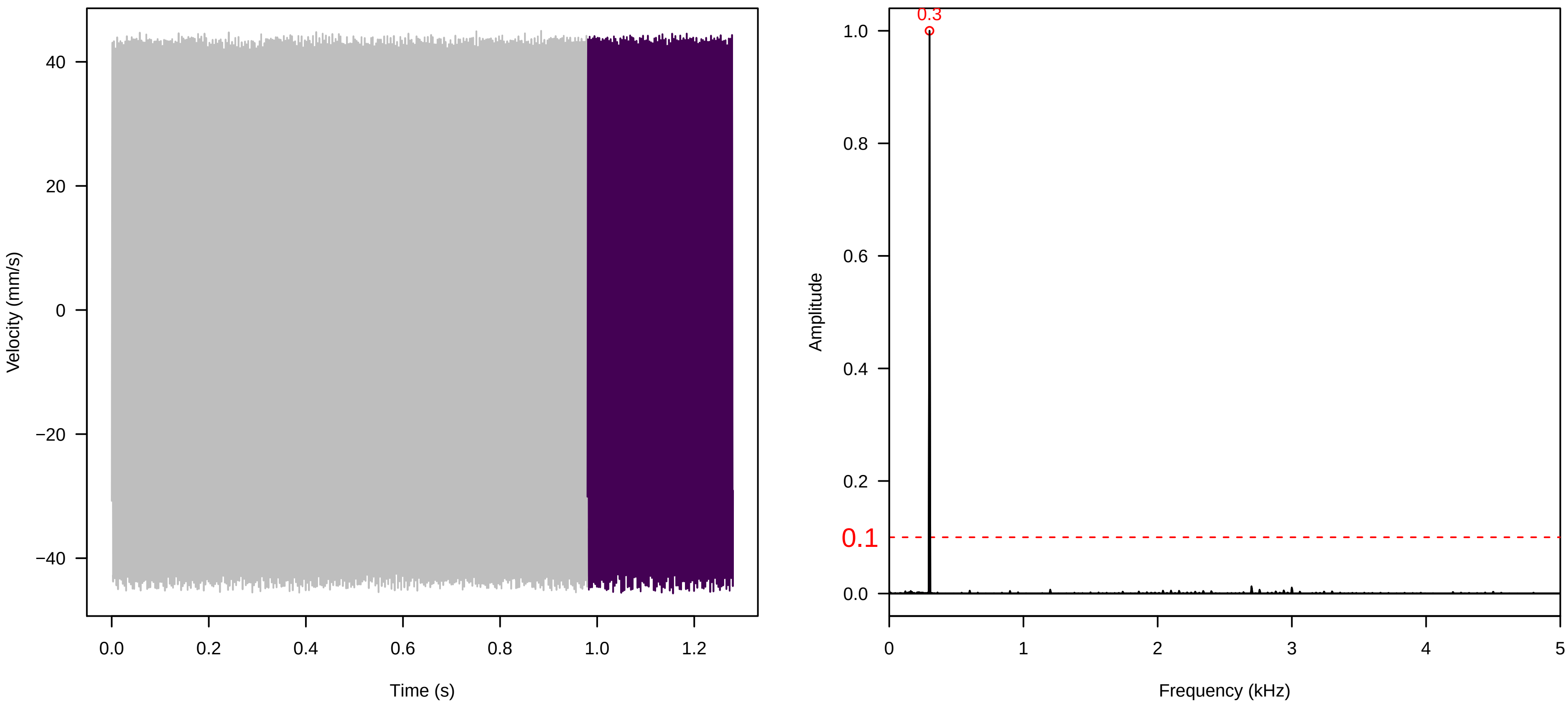

**Vel. =  0.057 ; Str. =  Corolla ; Axis =  x ; Fl. accession = 10−s−86**

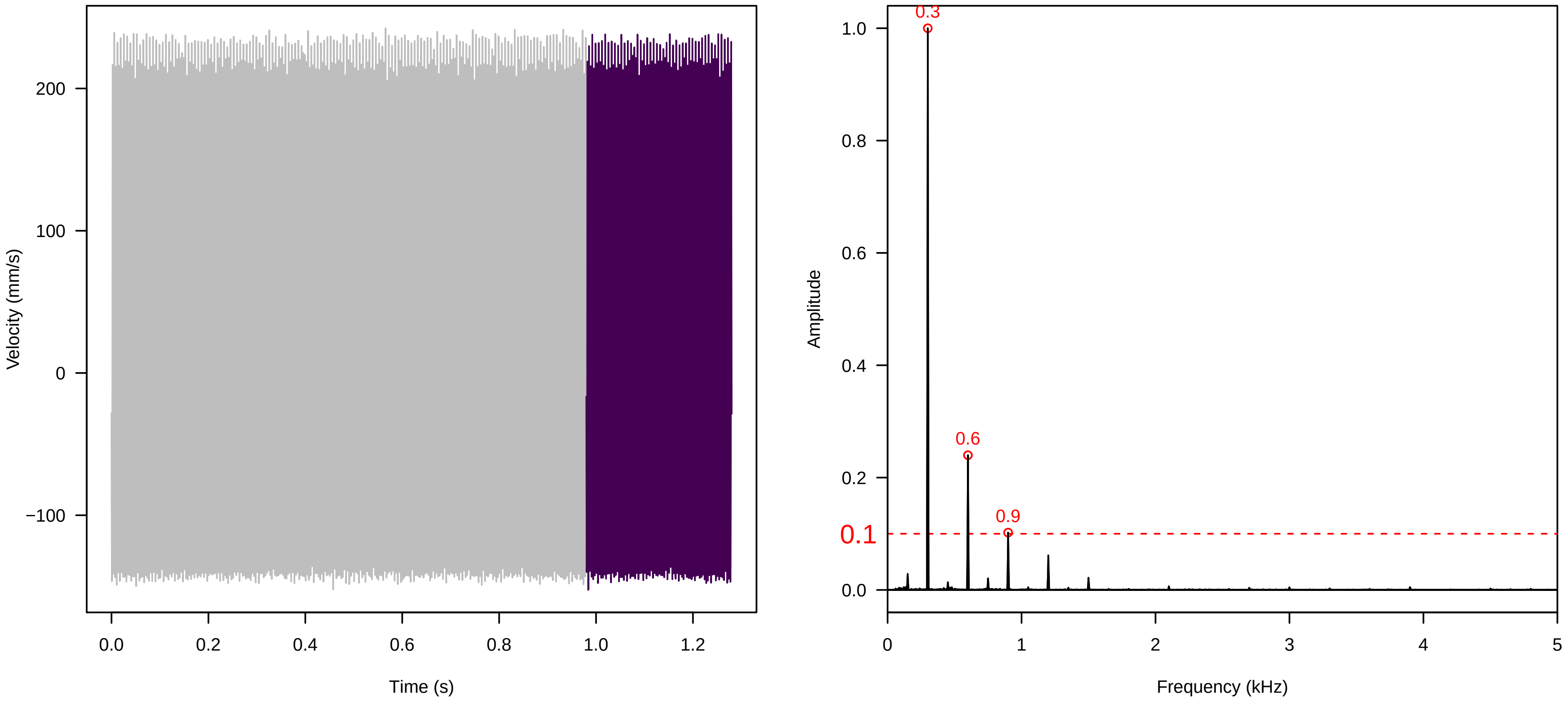


**Vel. =  0.057 ; Str. =  Receptacle ; Axis =  x ; Fl. accession = 10−s−86**

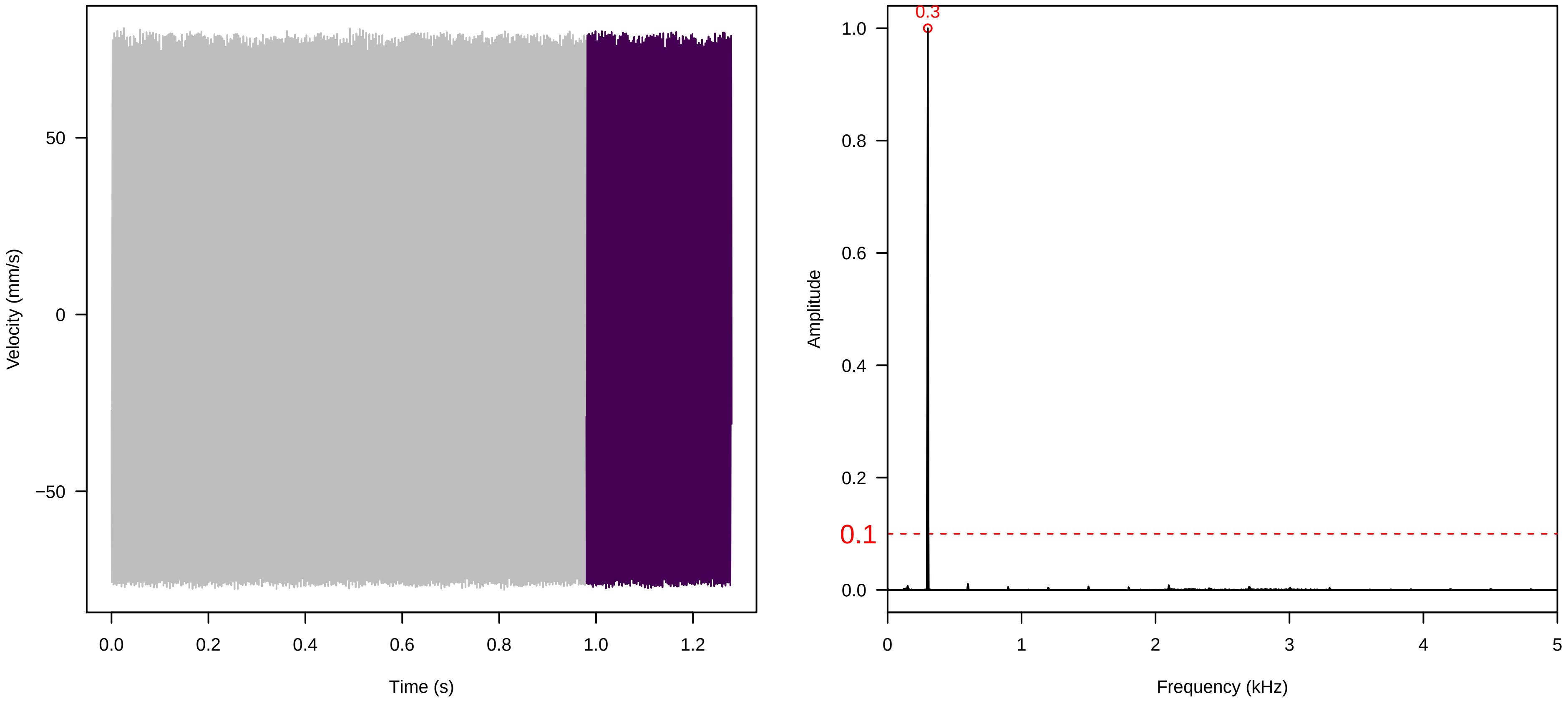


**Vel. =  0.057 ; Str. =  FA ; Axis =  x ; Fl. accession = 10−s−86**

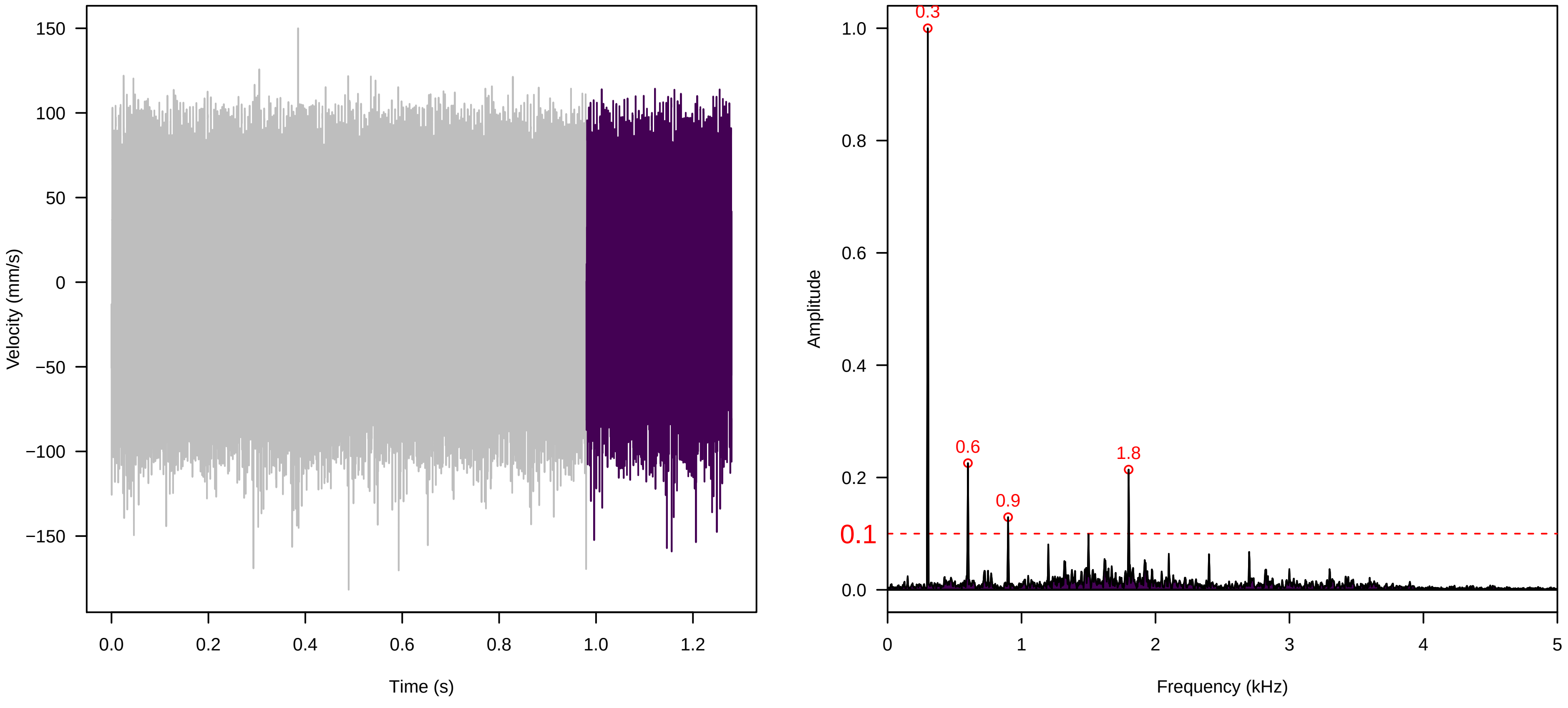

**Vel. = 0.057 ; Str. = Receptacle ; Axis = x ; Fl. accession = 10−s−86**

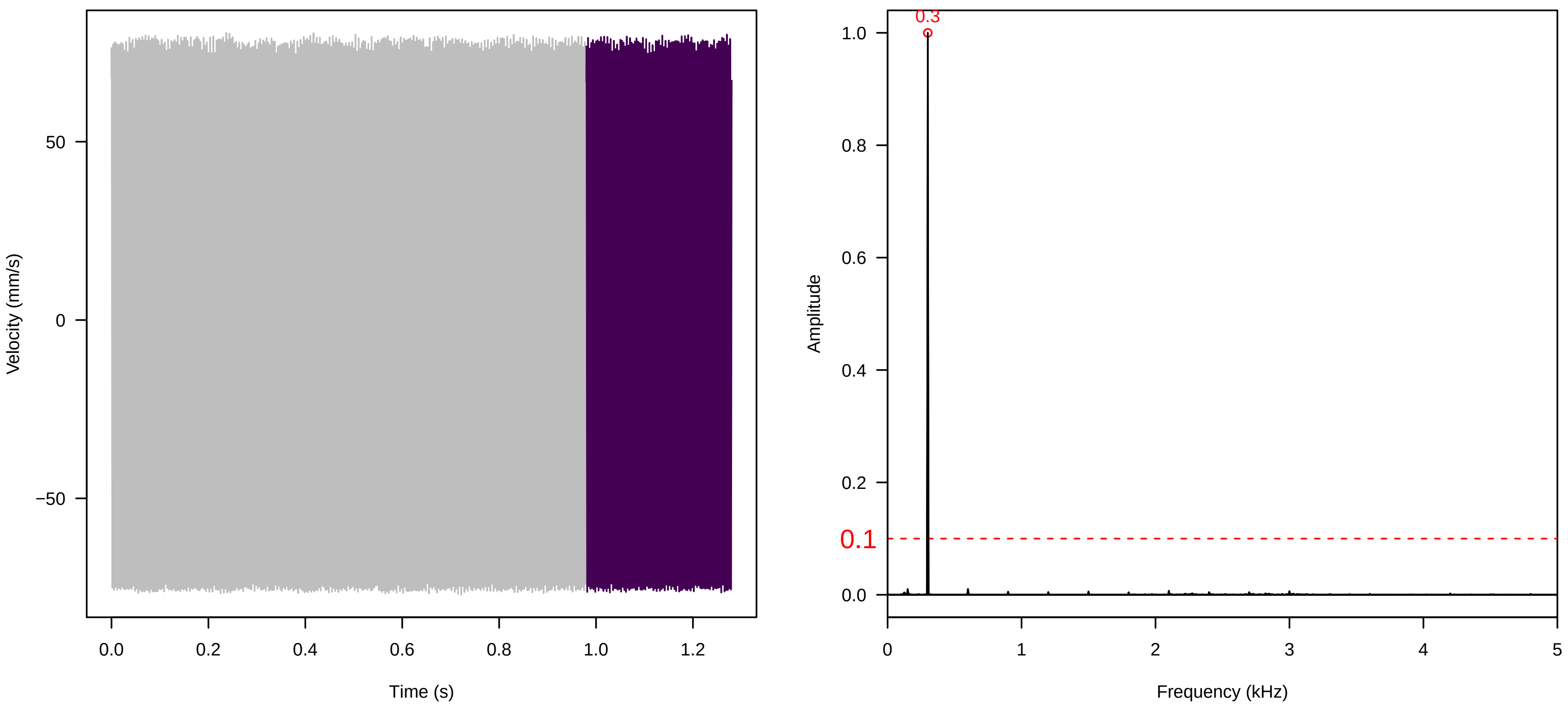


**Vel. = 0.057 ; Str. = PA ; Axis = x ; Fl. accession = 10−s−86**

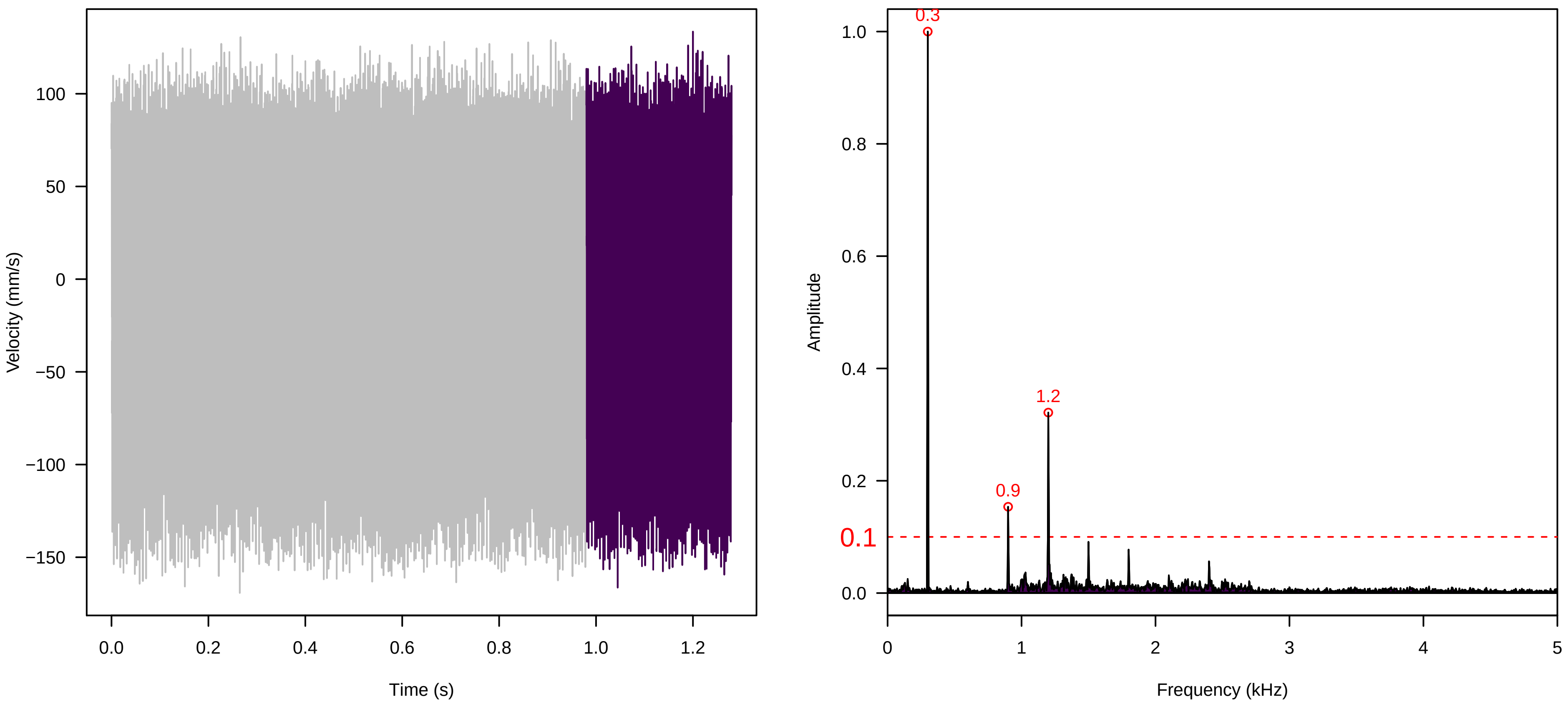


**Vel. = 0.057 ; Str. = Receptacle ; Axis = x ; Fl. accession = 10−s−86**

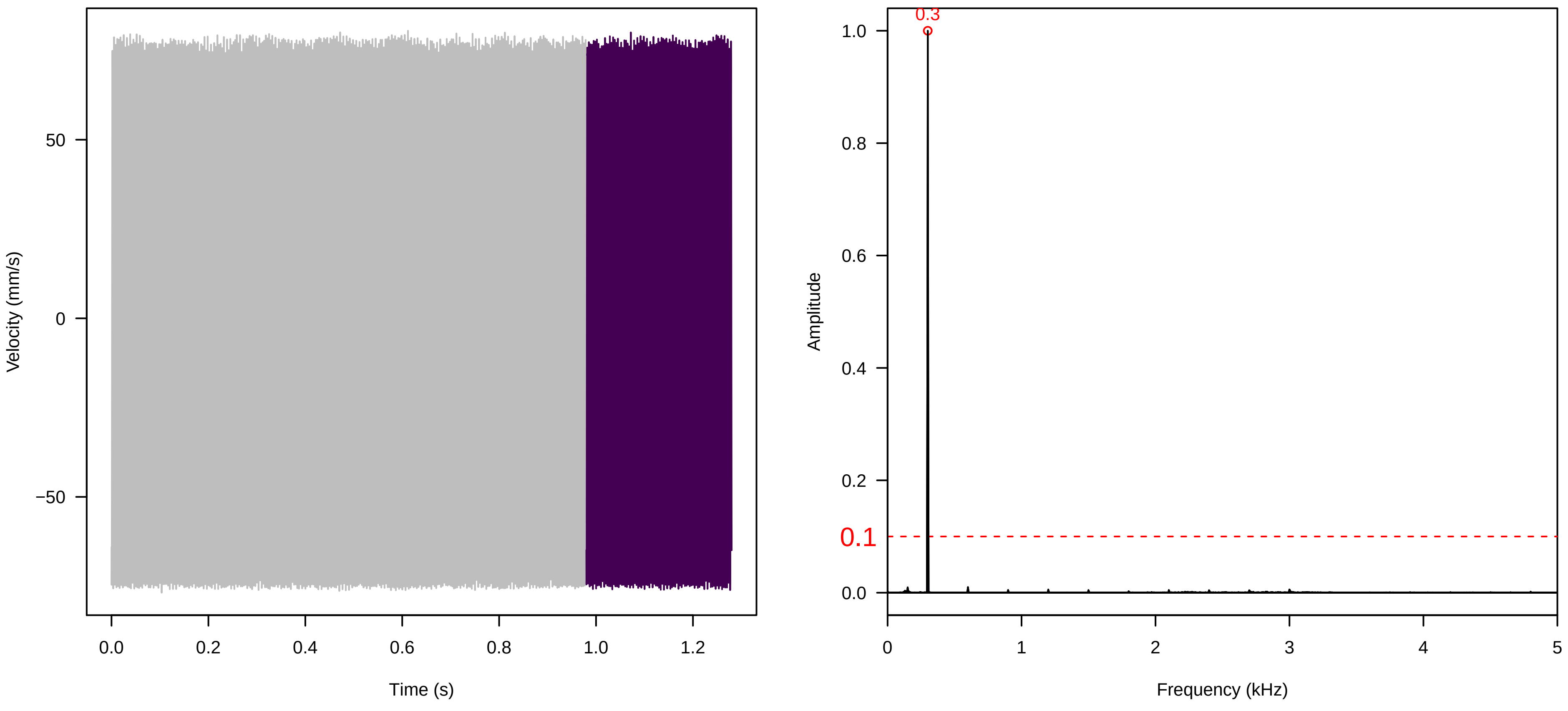

**Vel. = 0.014 ; Str. = Corolla ; Axis = x ; Fl. accession = 10−s−77−19**

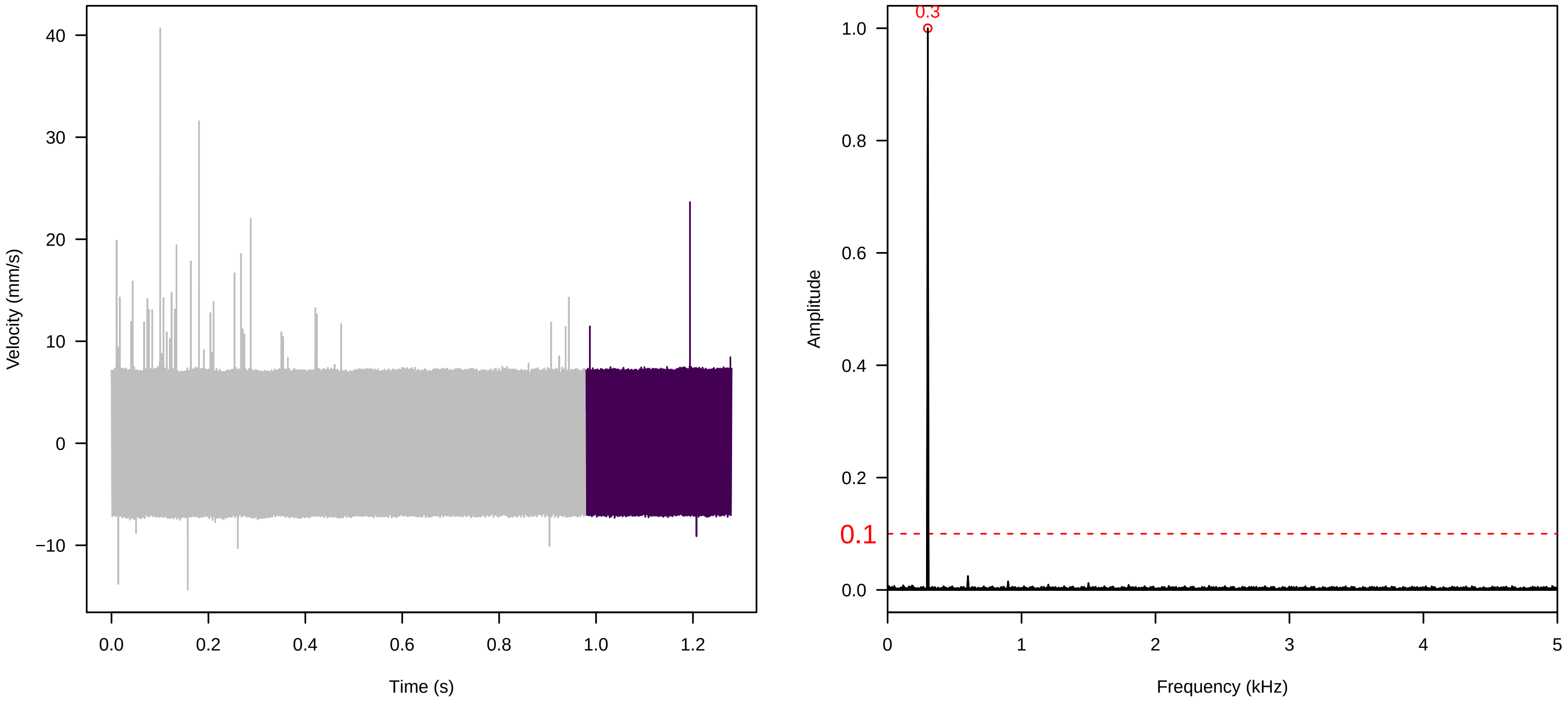


**Vel. = 0.014 ; Str. = Receptacle ; Axis = x ; Fl. accession = 10−s−77−19**

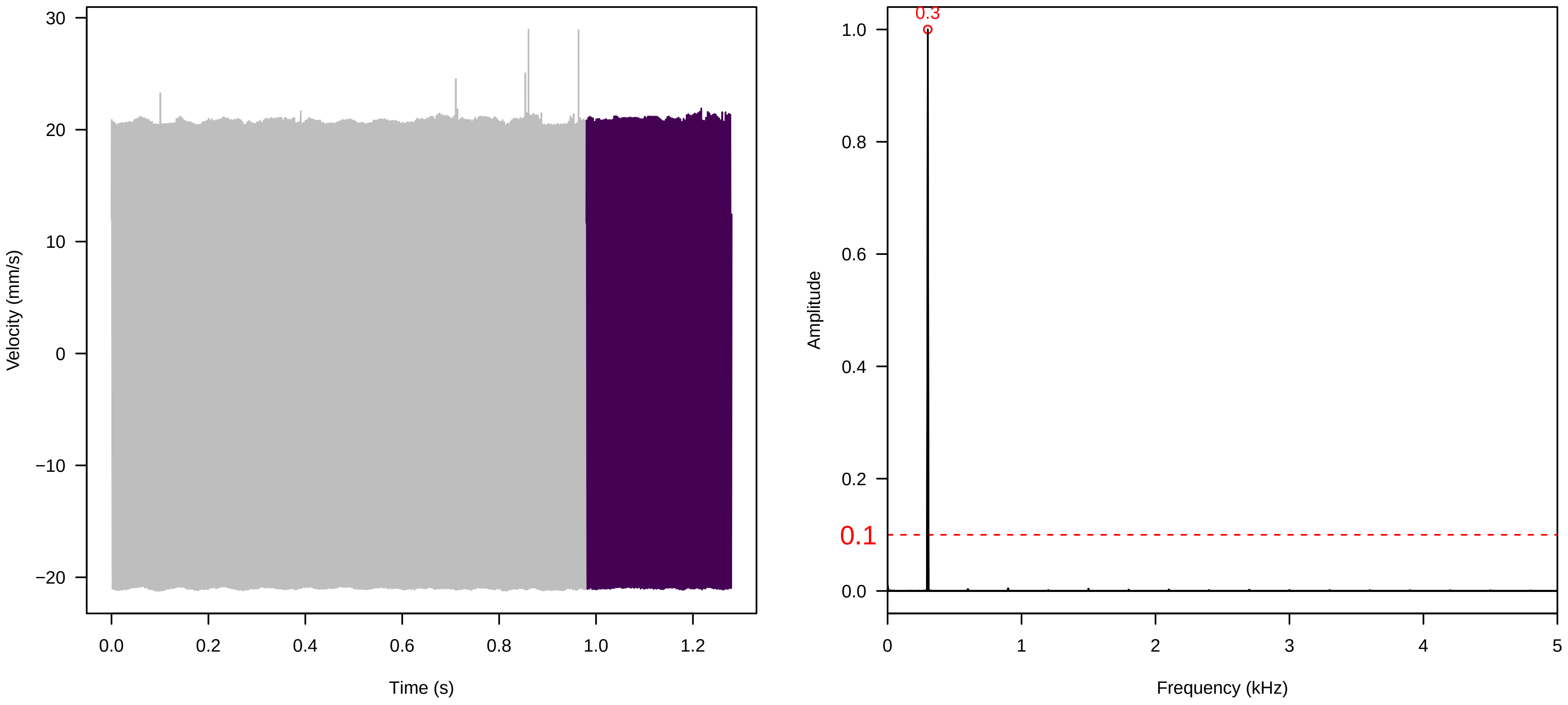


**Vel. = 0.014 ; Str. = FA ; Axis = x ; Fl. accession = 10−s−77−19**

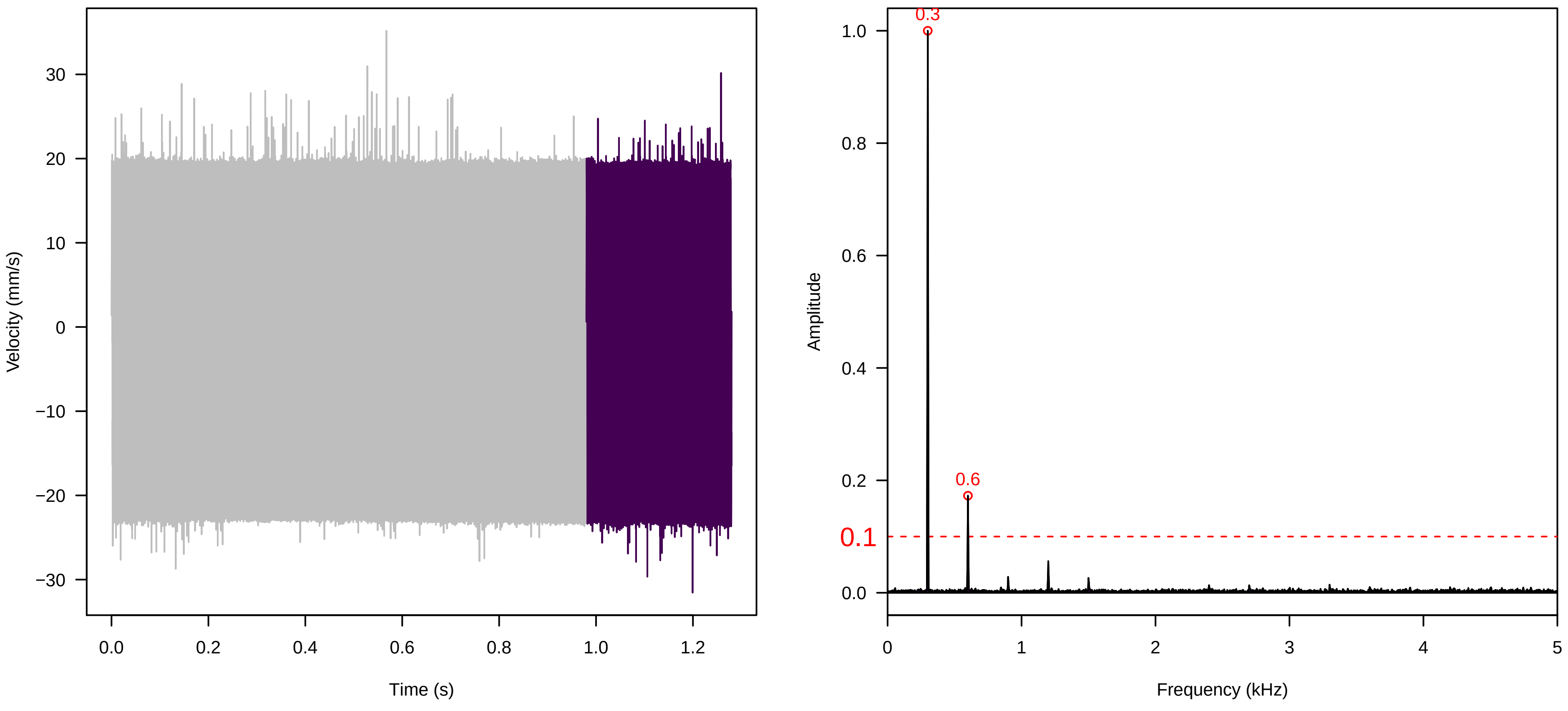

**Vel. = 0.014 ; Str. = Receptacle ; Axis = x ; Fl. accession = 10−s−77−19**

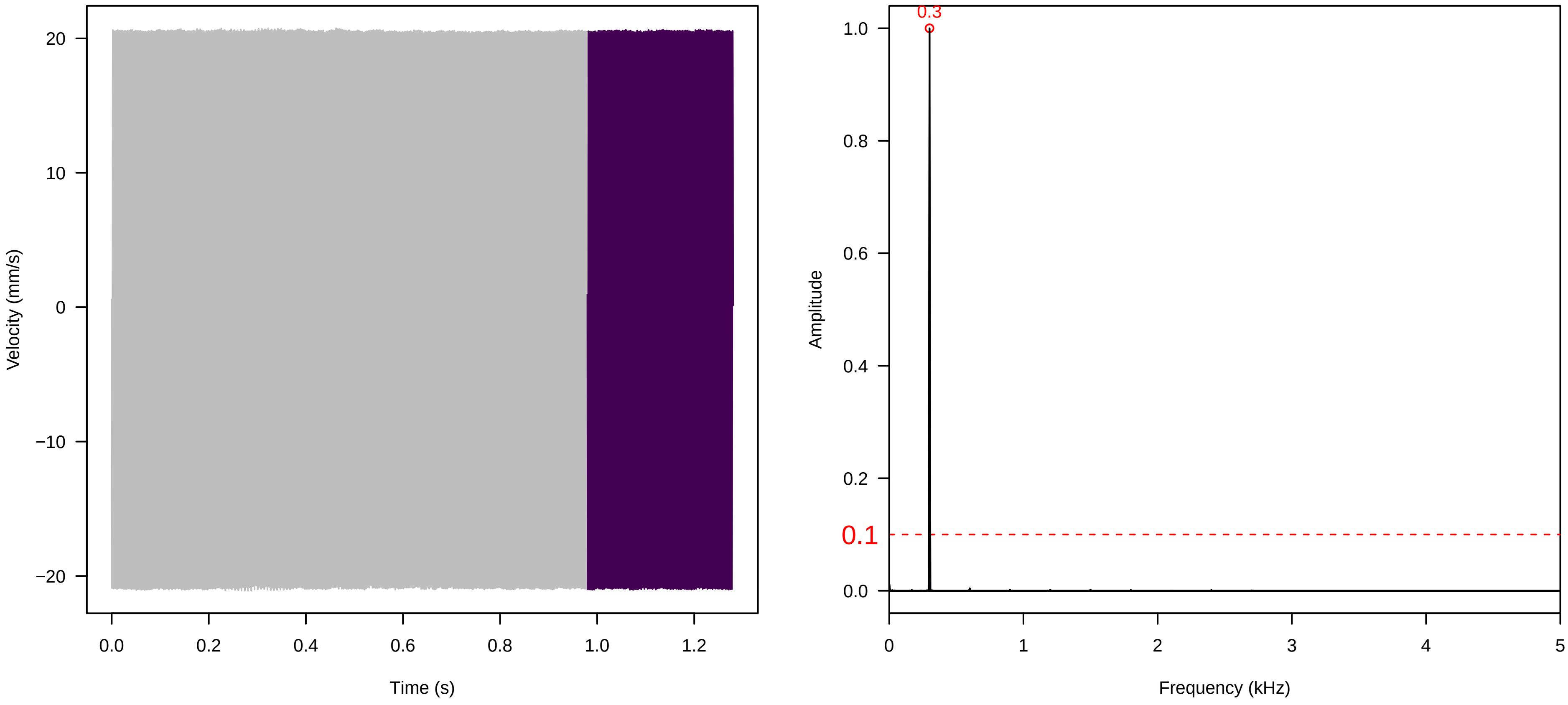


**Vel. = 0.014 ; Str. = PA ; Axis = x ; Fl. accession = 10−s−77−19**

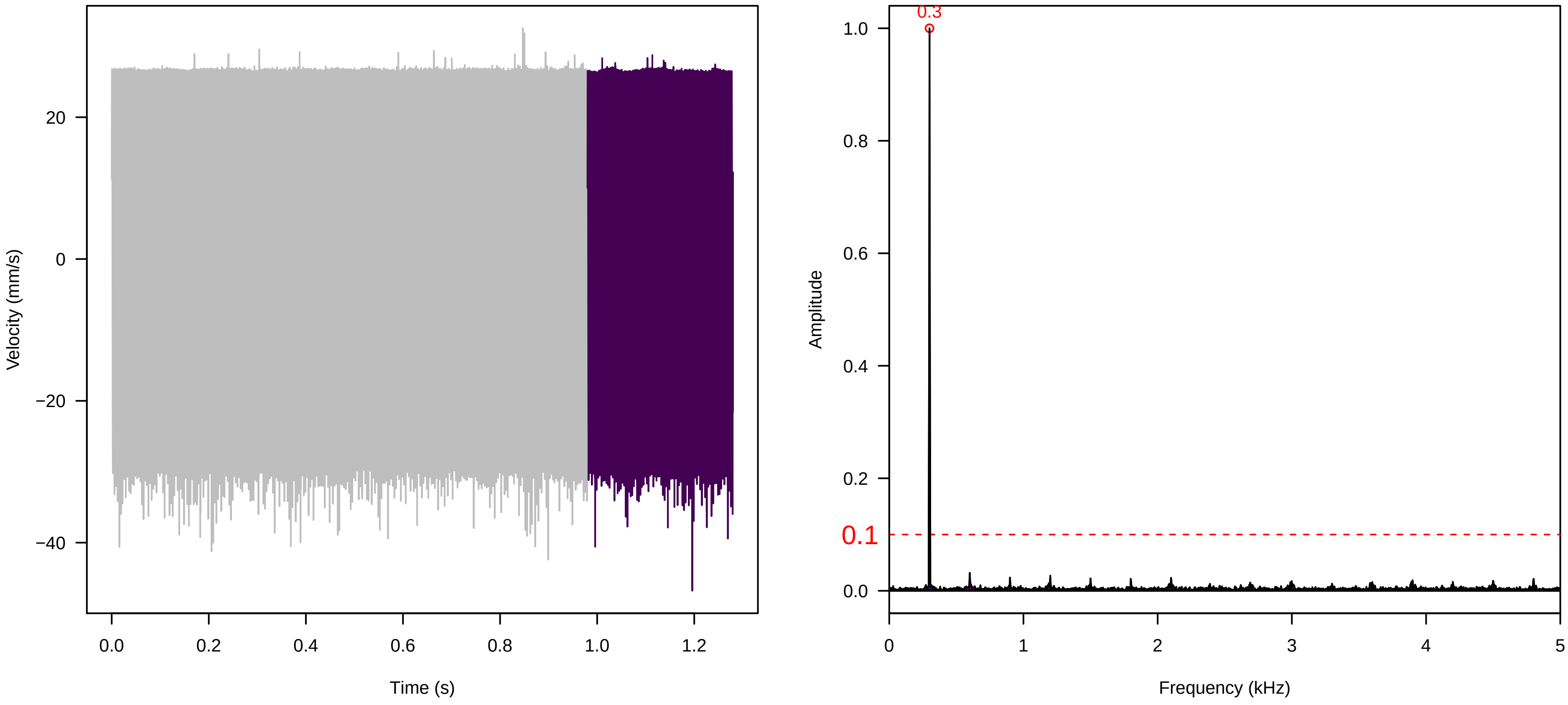


**Vel. = 0.014 ; Str. = Receptacle ; Axis = x ; Fl. accession = 10−s−77−19**

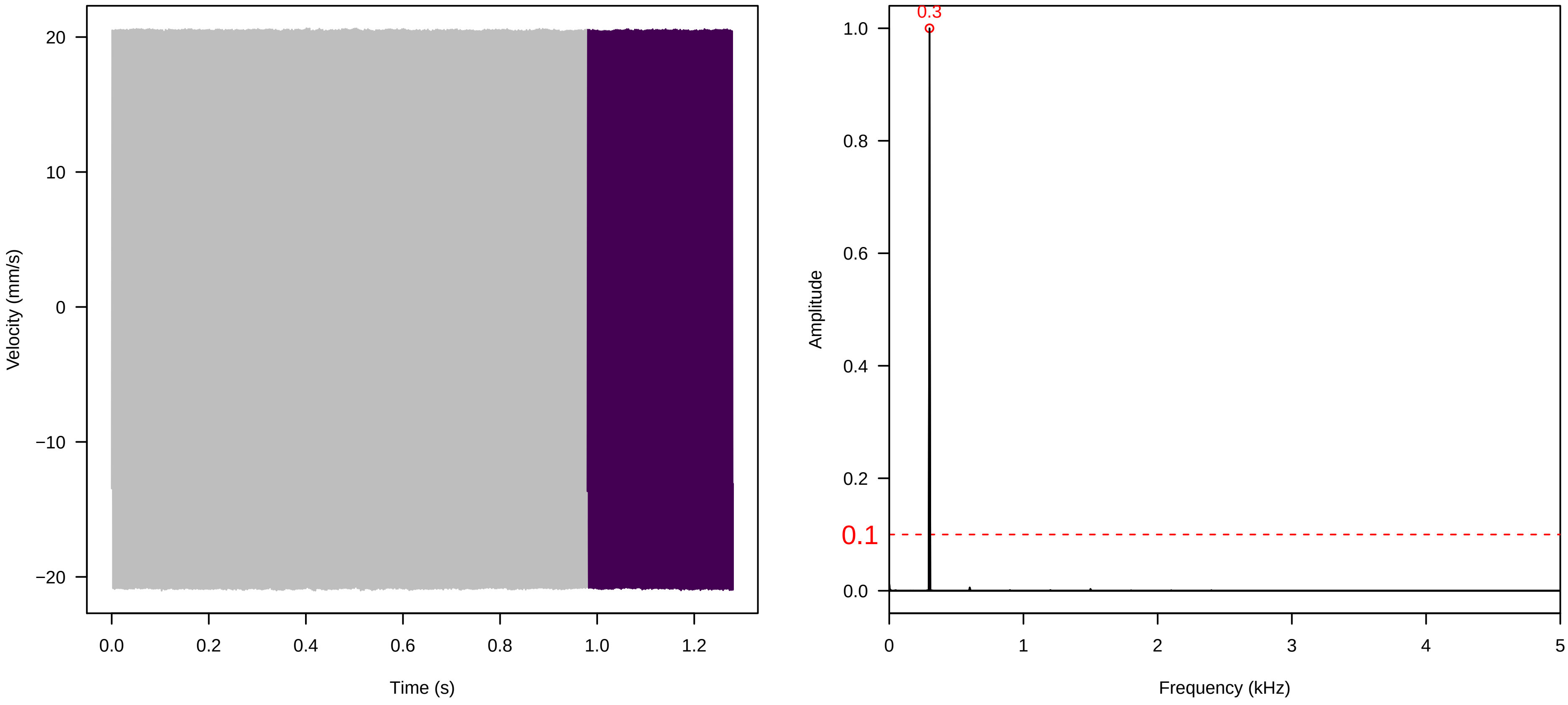

**Vel. = 0.028 ; Str. = PA ; Axis = x ; Fl. accession = 10−s−77−19**

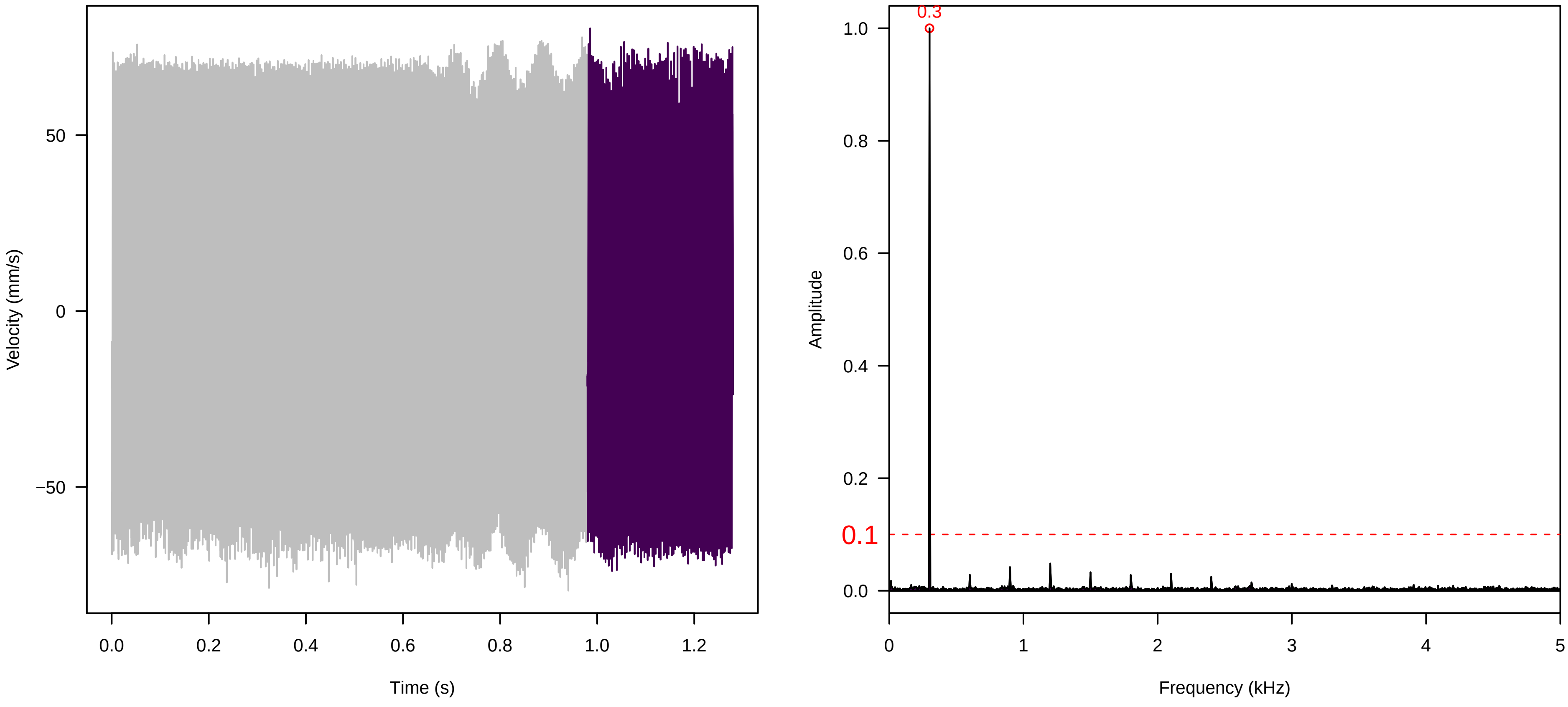


**Vel. = 0.028 ; Str. = Receptacle ; Axis = x ; Fl. accession = 10−s−77−19**

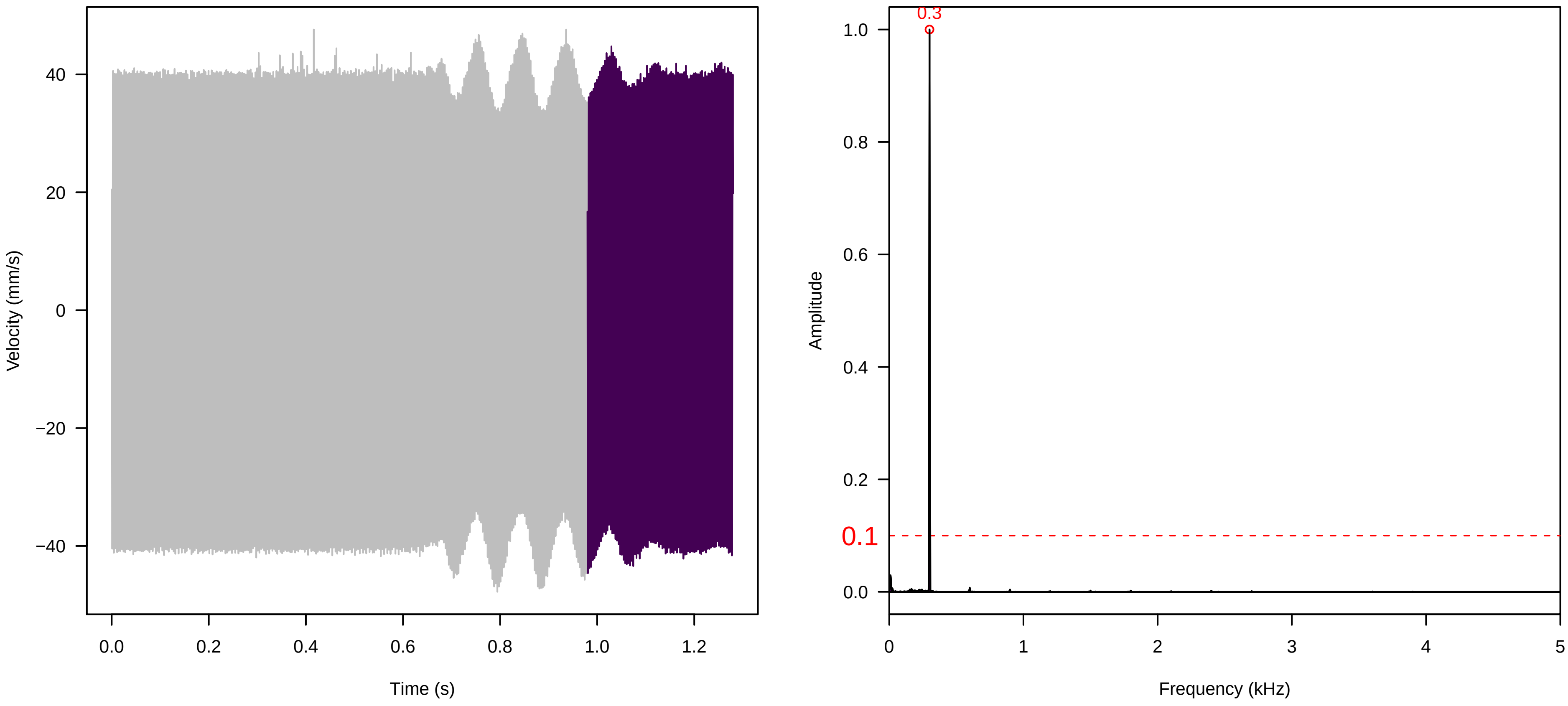


**Vel. = 0.028 ; Str. = FA ; Axis = x ; Fl. accession = 10−s−77−19**

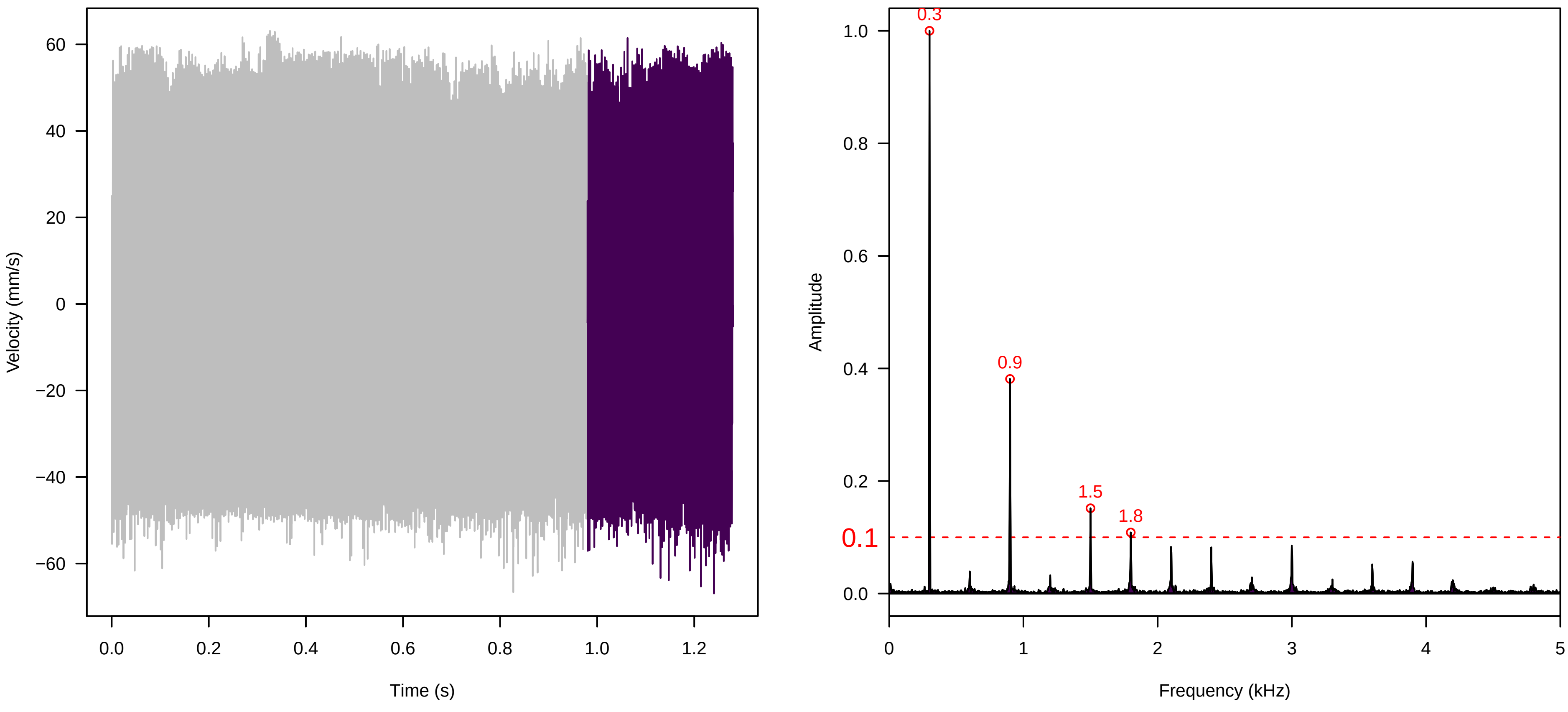

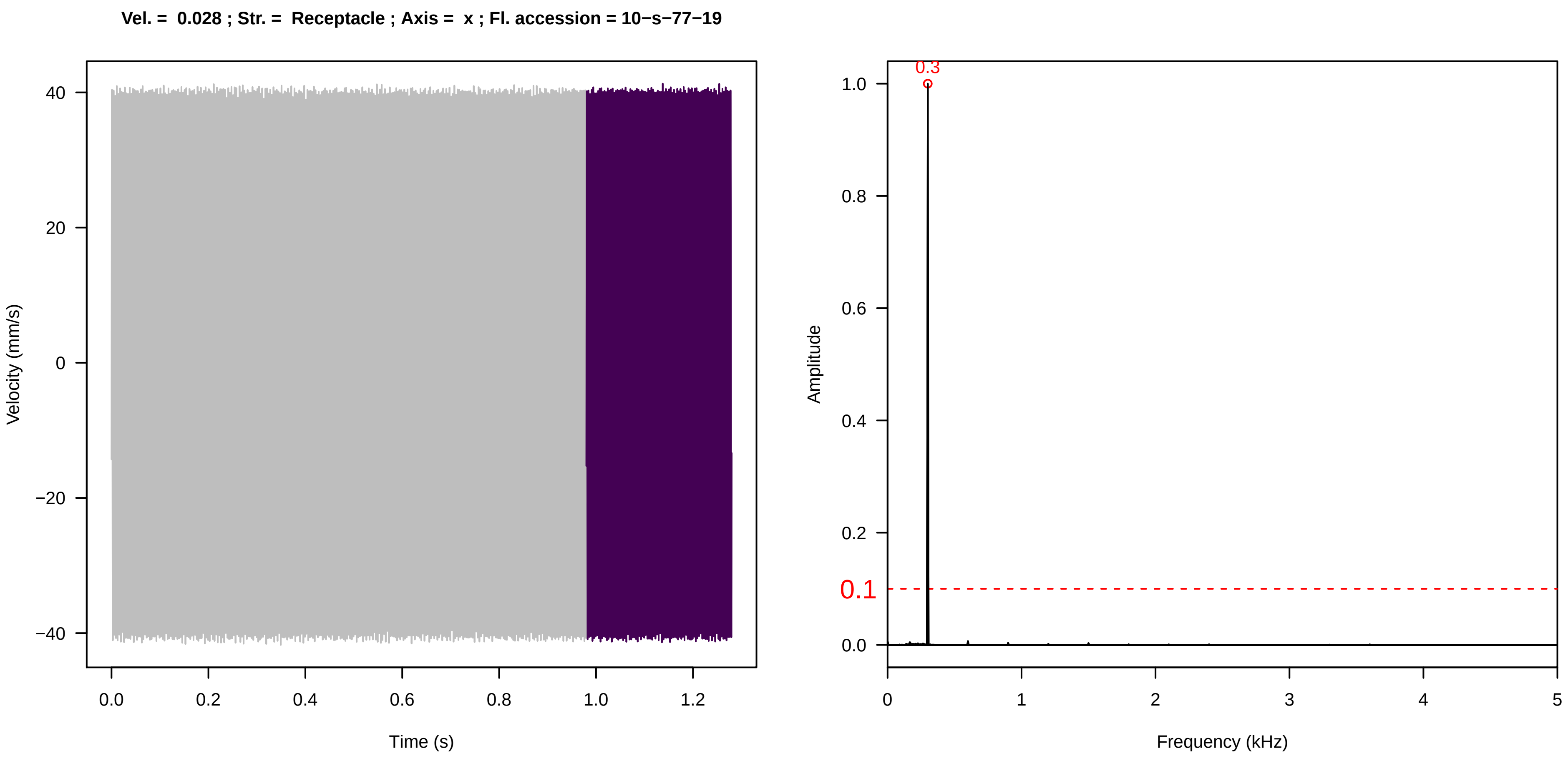
Vel. = 0.028 ; Str. = Receptacle ; Axis = x ; Fl. accession = 10−s−77−19
Velocity (mm/s)
Time (s)
Amplitude
Frequency (kHz)
0.3
0.1

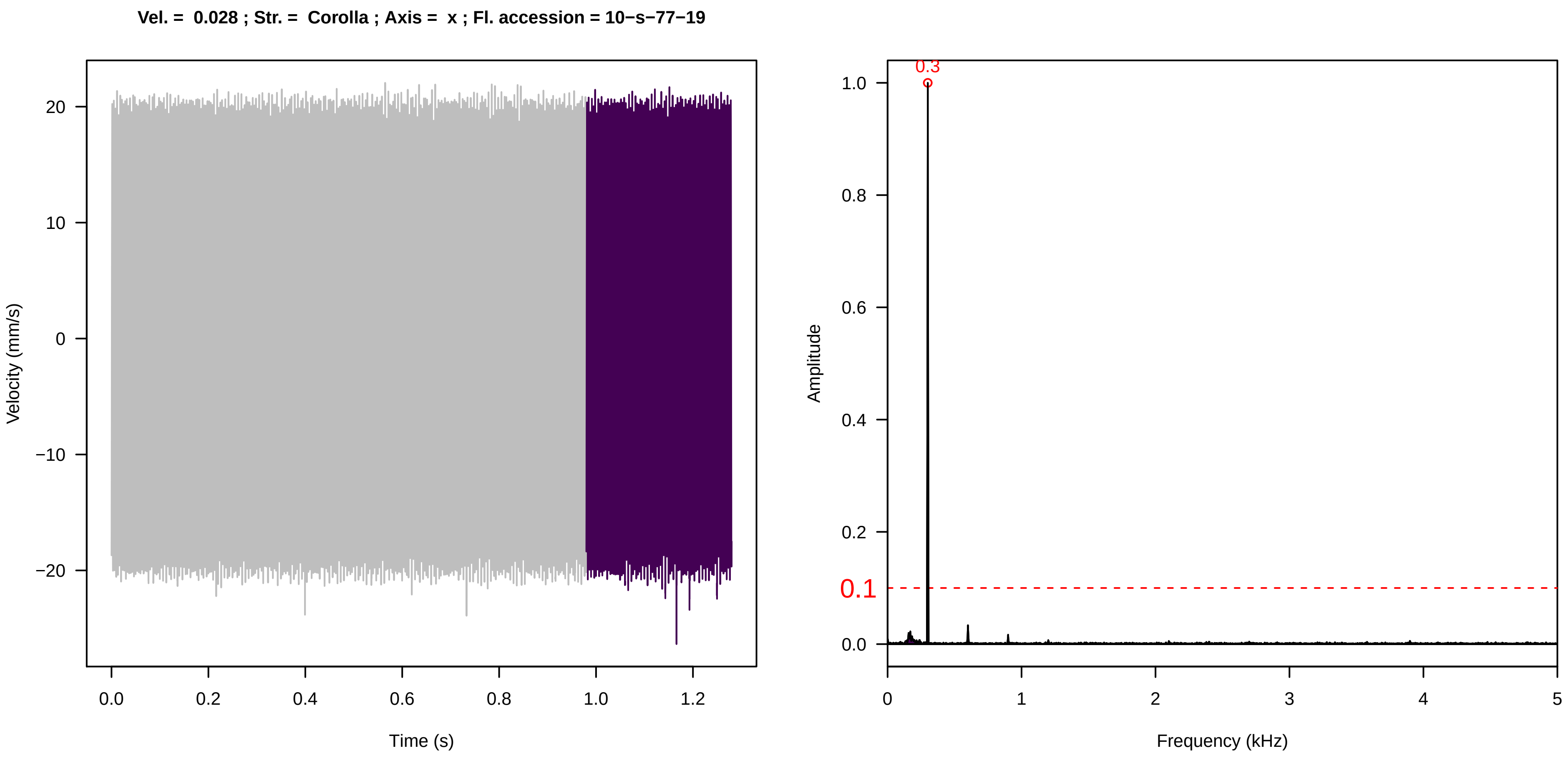
Vel. = 0.028 ; Str. = Corolla ; Axis = x ; Fl. accession = 10−s−77−19
Velocity (mm/s)
Time (s)
Amplitude
Frequency (kHz)
0.3
0.1

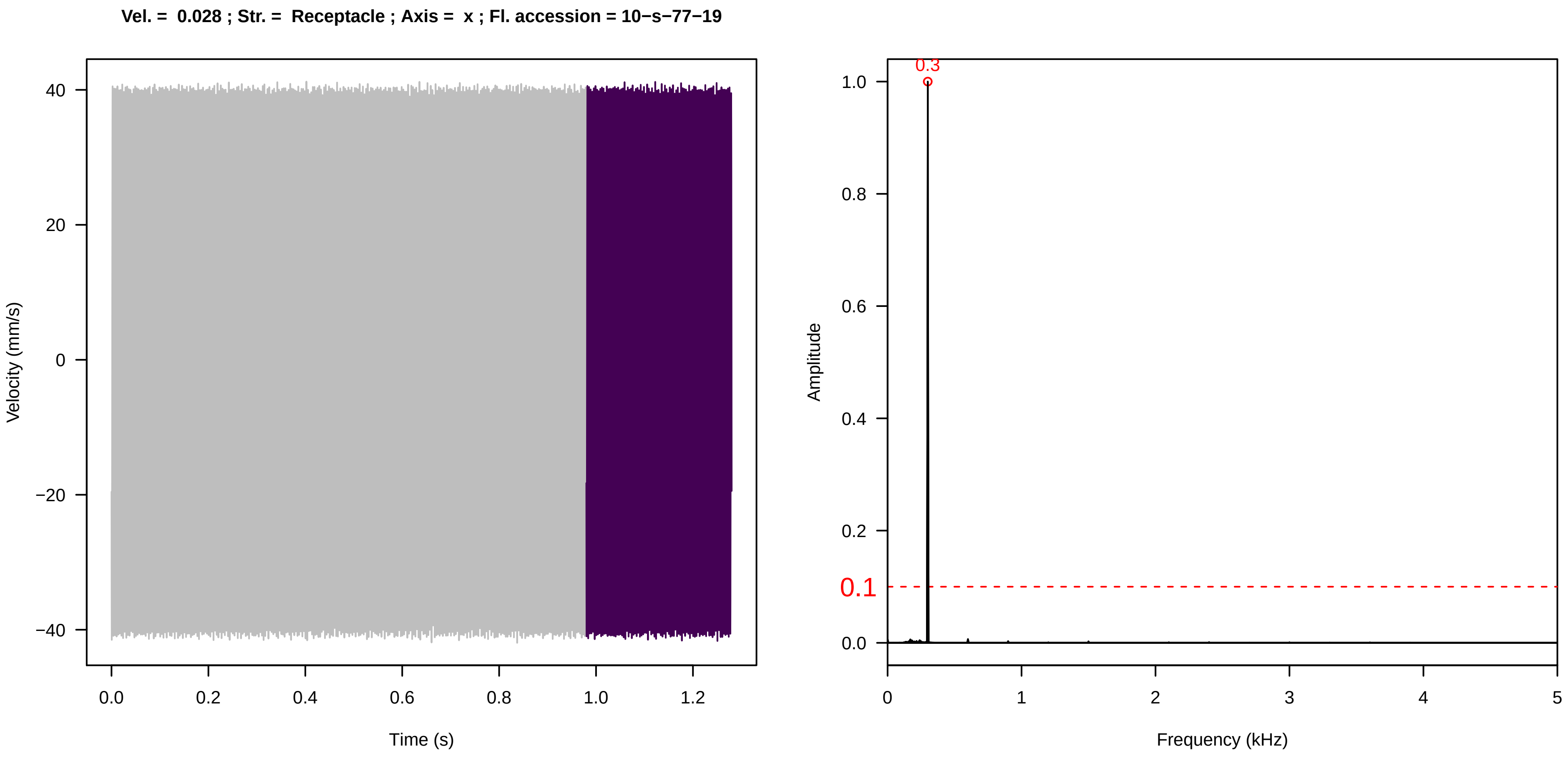
Vel. = 0.028 ; Str. = Receptacle ; Axis = x ; Fl. accession = 10−s−77−19
Velocity (mm/s)
Time (s)
Amplitude
Frequency (kHz)
0.3
0.1

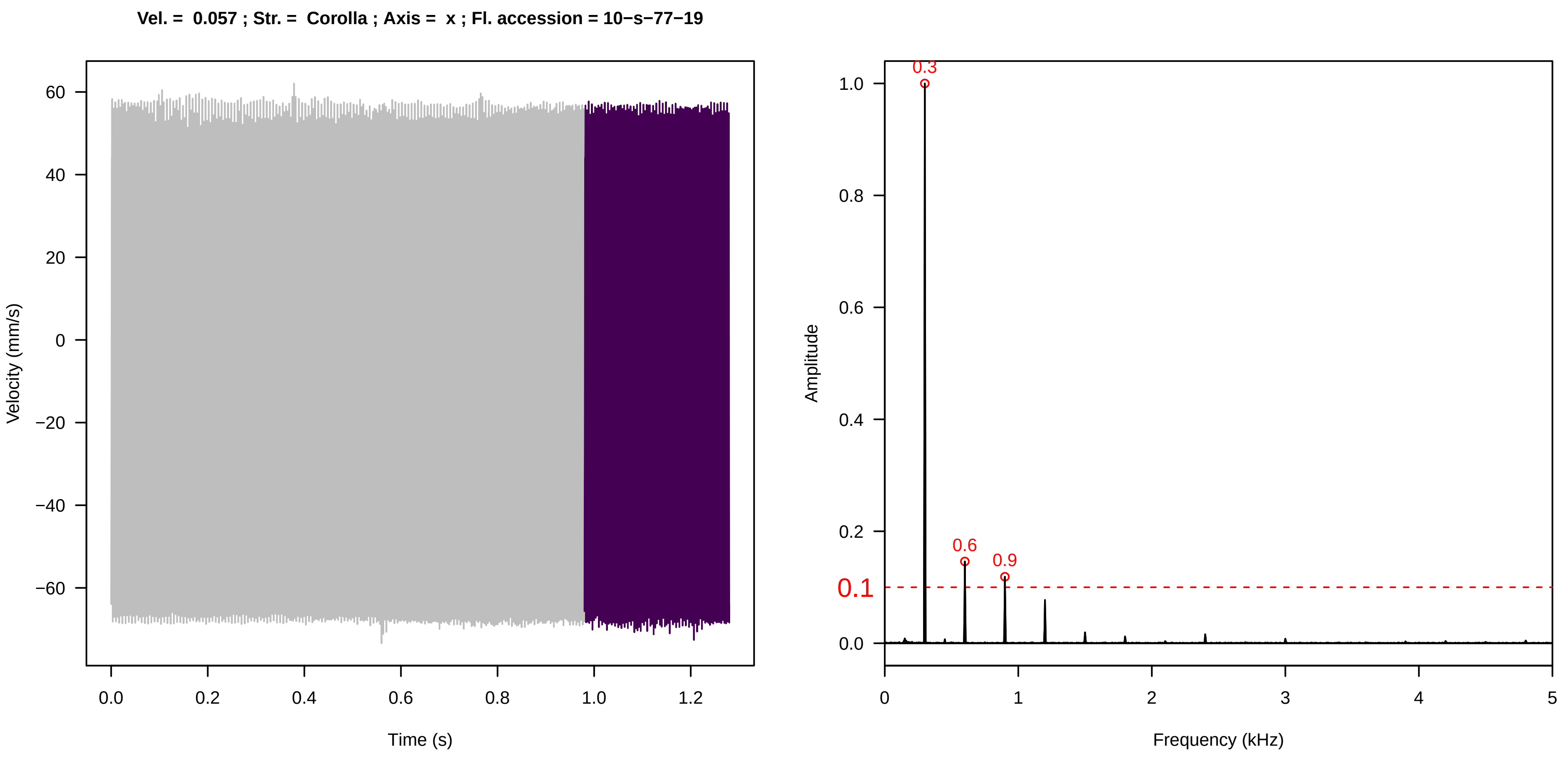
Vel. = 0.057 ; Str. = Corolla ; Axis = x ; Fl. accession = 10−s−77−19
Velocity (mm/s)
Time (s)
Amplitude
Frequency (kHz)
0.3
0.6
0.9
0.1

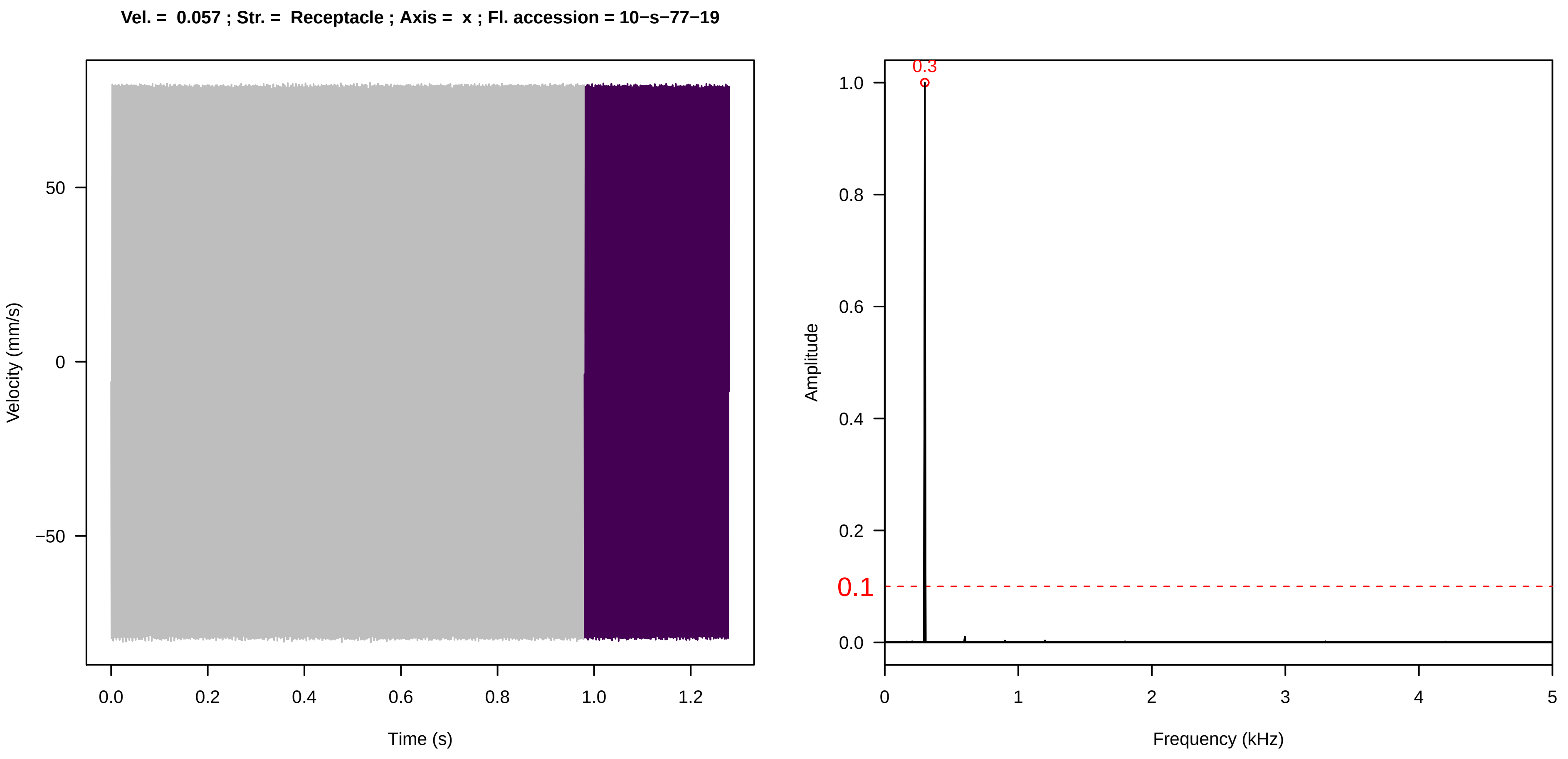
Vel. = 0.057 ; Str. = Receptacle ; Axis = x ; Fl. accession = 10−s−77−19
Velocity (mm/s)
Time (s)
Amplitude
Frequency (kHz)
0.3
0.1

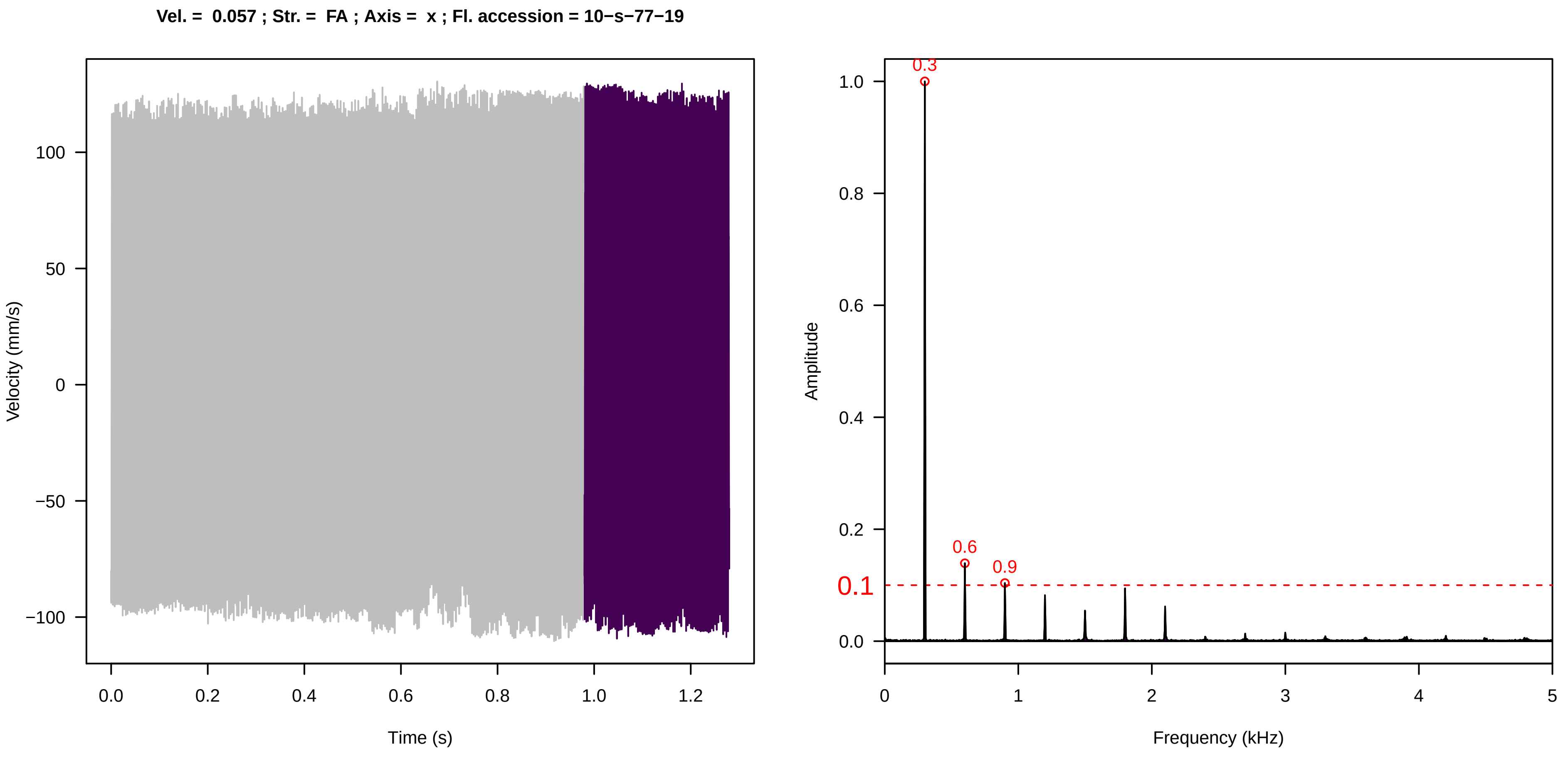
Vel. = 0.057 ; Str. = FA ; Axis = x ; Fl. accession = 10−s−77−19
Velocity (mm/s)
Time (s)
Amplitude
Frequency (kHz)
0.3
0.6
0.9
0.1

**Vel. =  0.057 ; Str. =  Receptacle ; Axis =  x ; Fl. accession = 10−s−77−19**

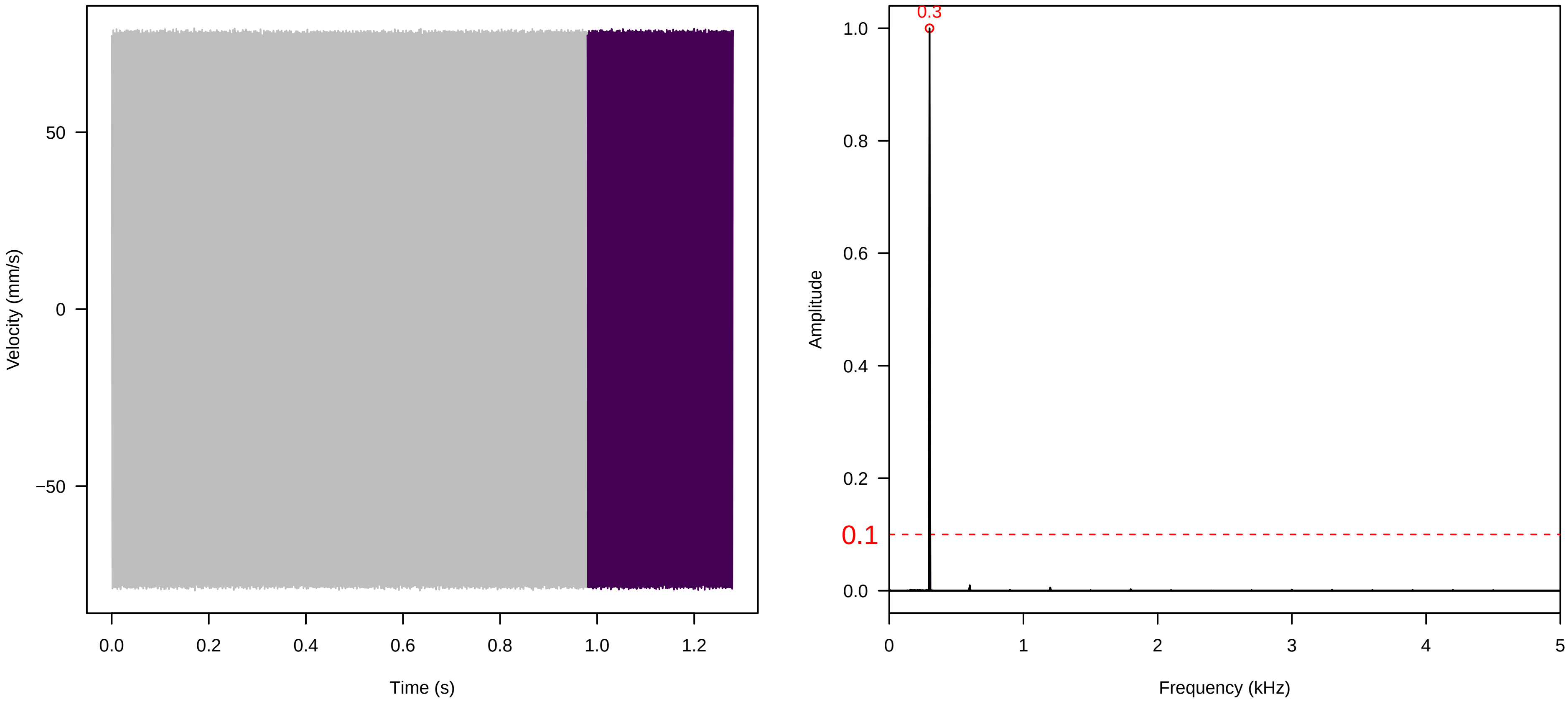


**Vel. =  0.057 ; Str. =  PA ; Axis =  x ; Fl. accession = 10−s−77−19**

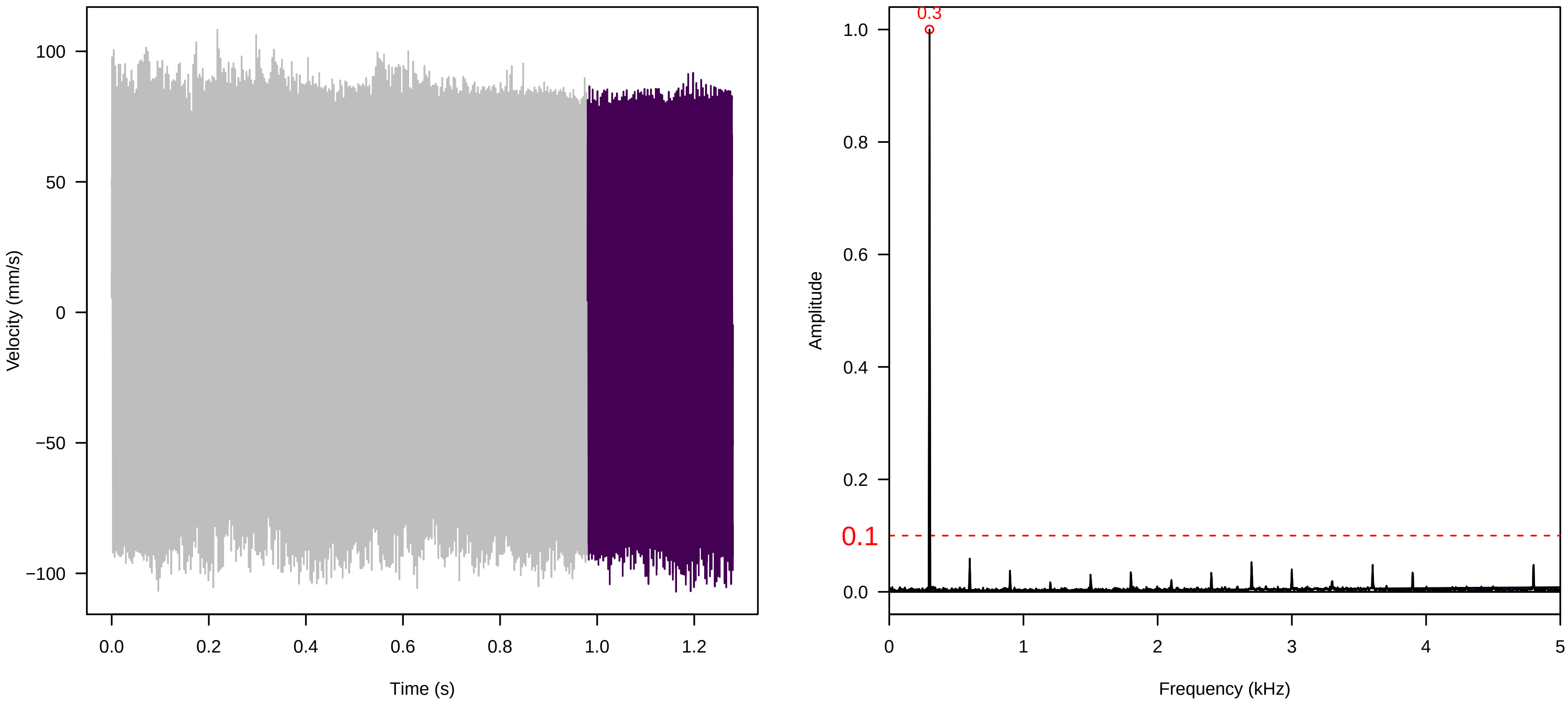


**Vel. =  0.057 ; Str. =  Receptacle ; Axis =  x ; Fl. accession = 10−s−77−19**

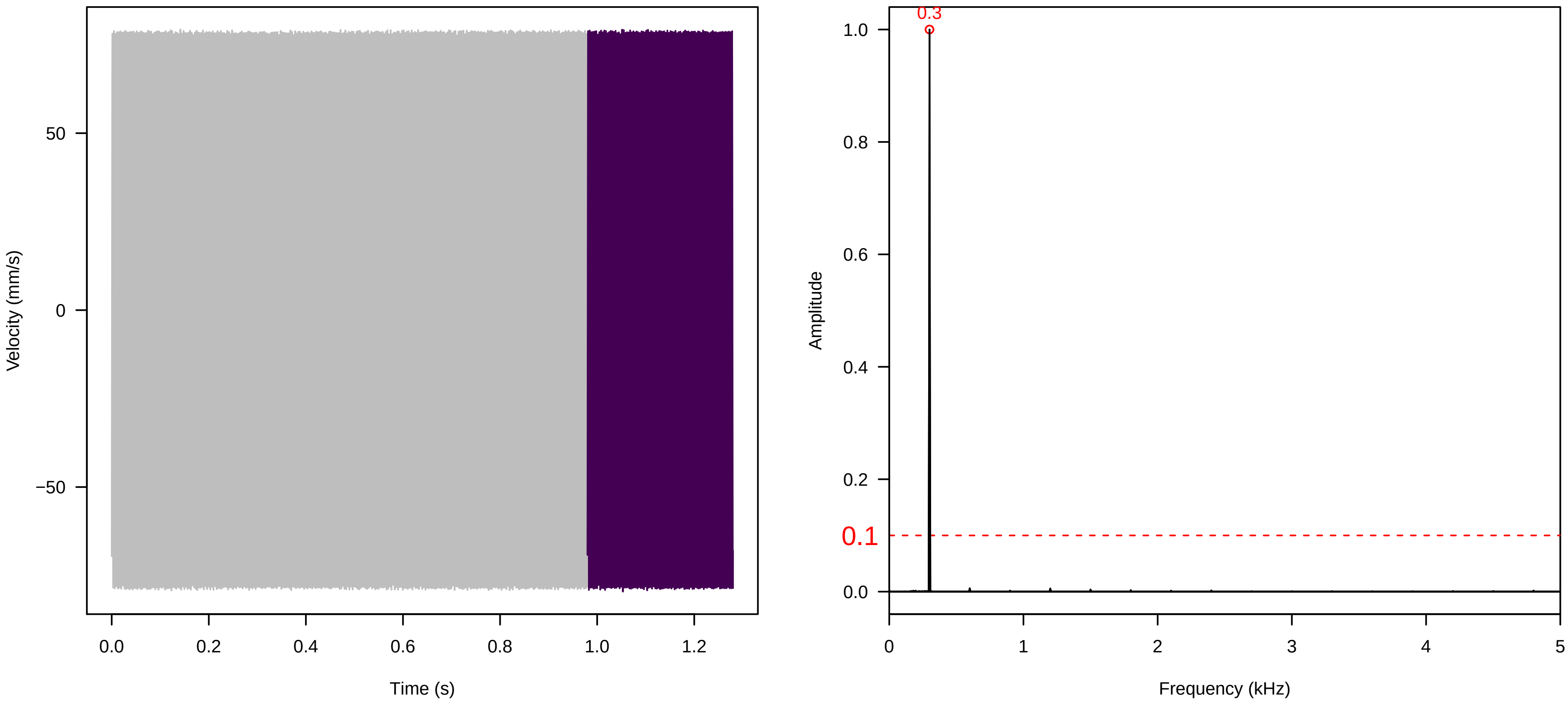

**Vel. = 0.057 ; Str. = Corolla ; Axis = z ; Fl. accession = 10−s−77−19**

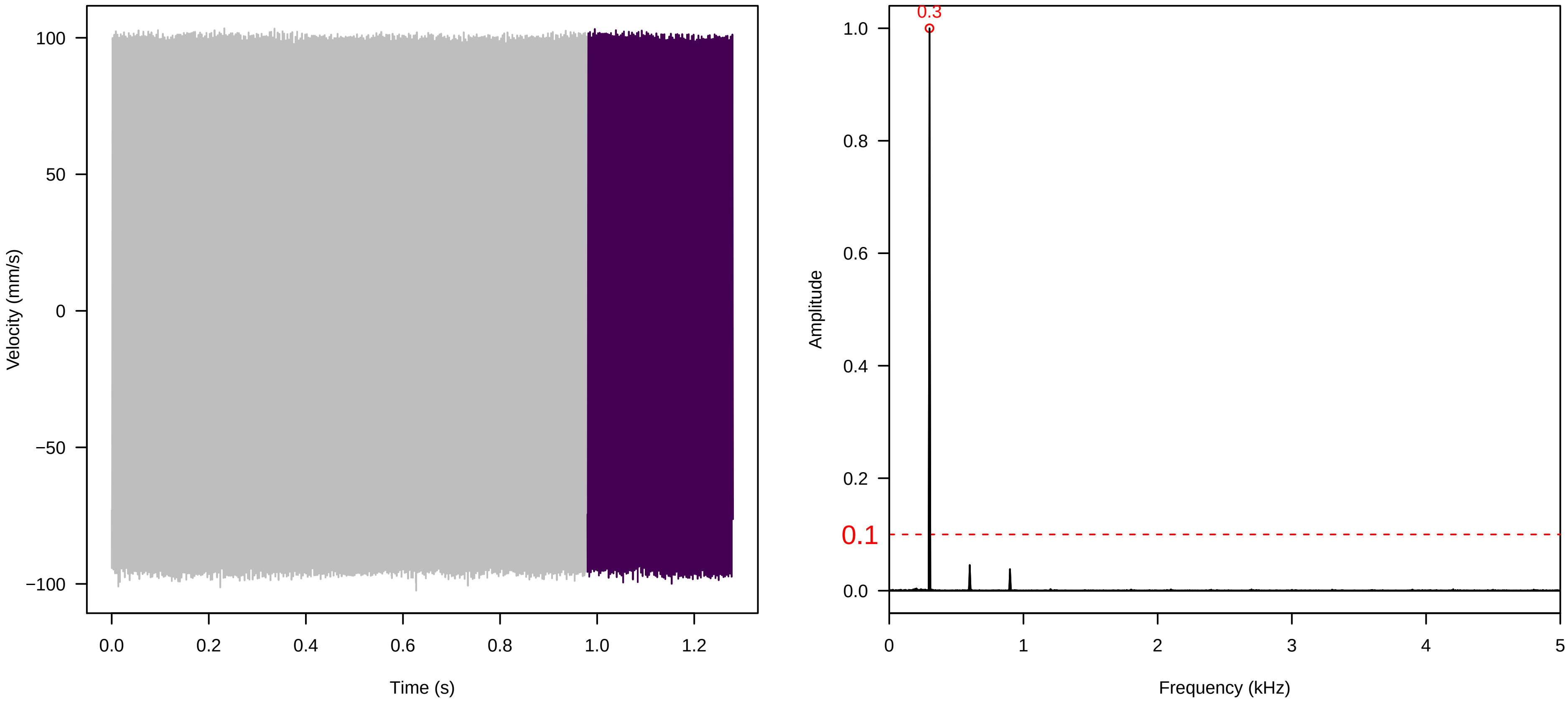


**Vel. = 0.057 ; Str. = Receptacle ; Axis = z ; Fl. accession = 10−s−77−19**

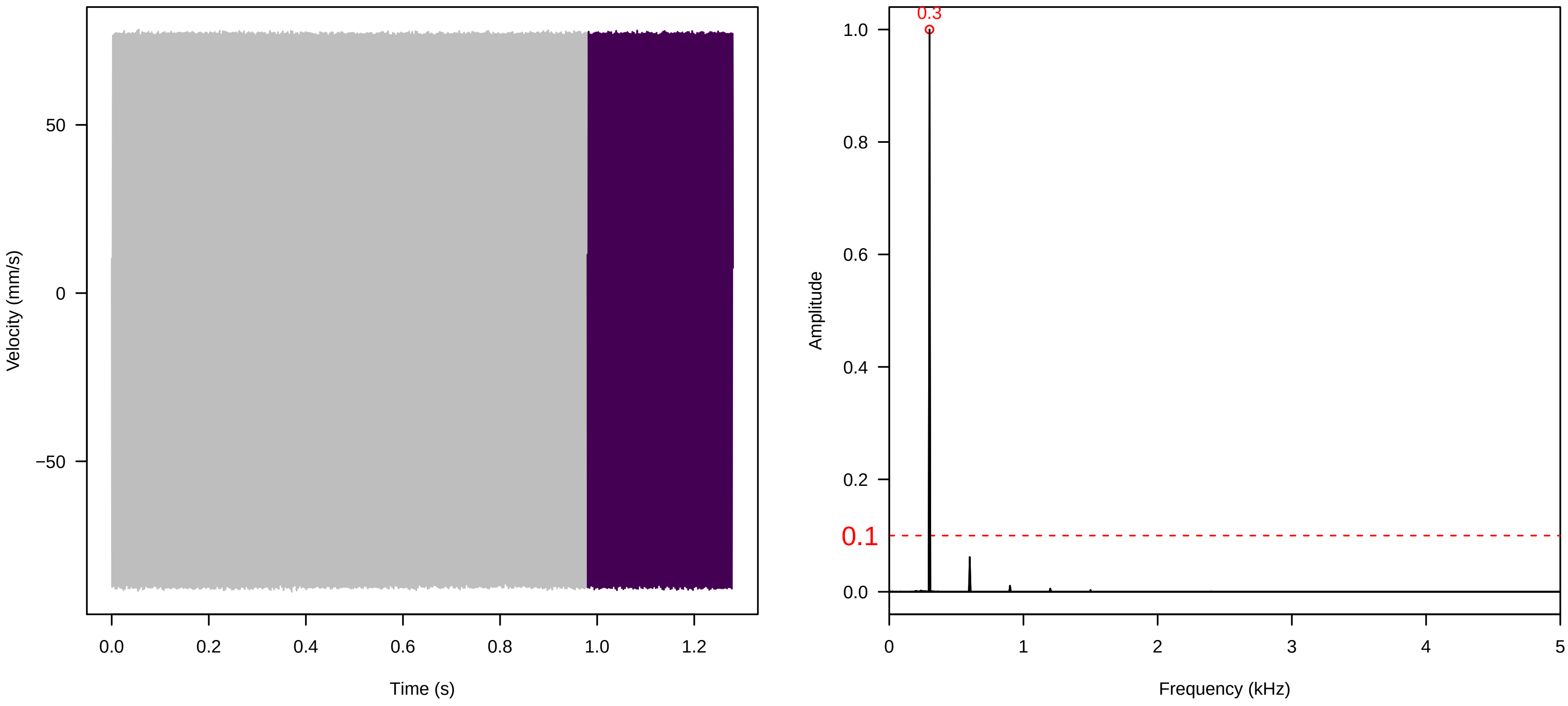


**Vel. = 0.057 ; Str. = FA ; Axis = z ; Fl. accession = 10−s−77−19**

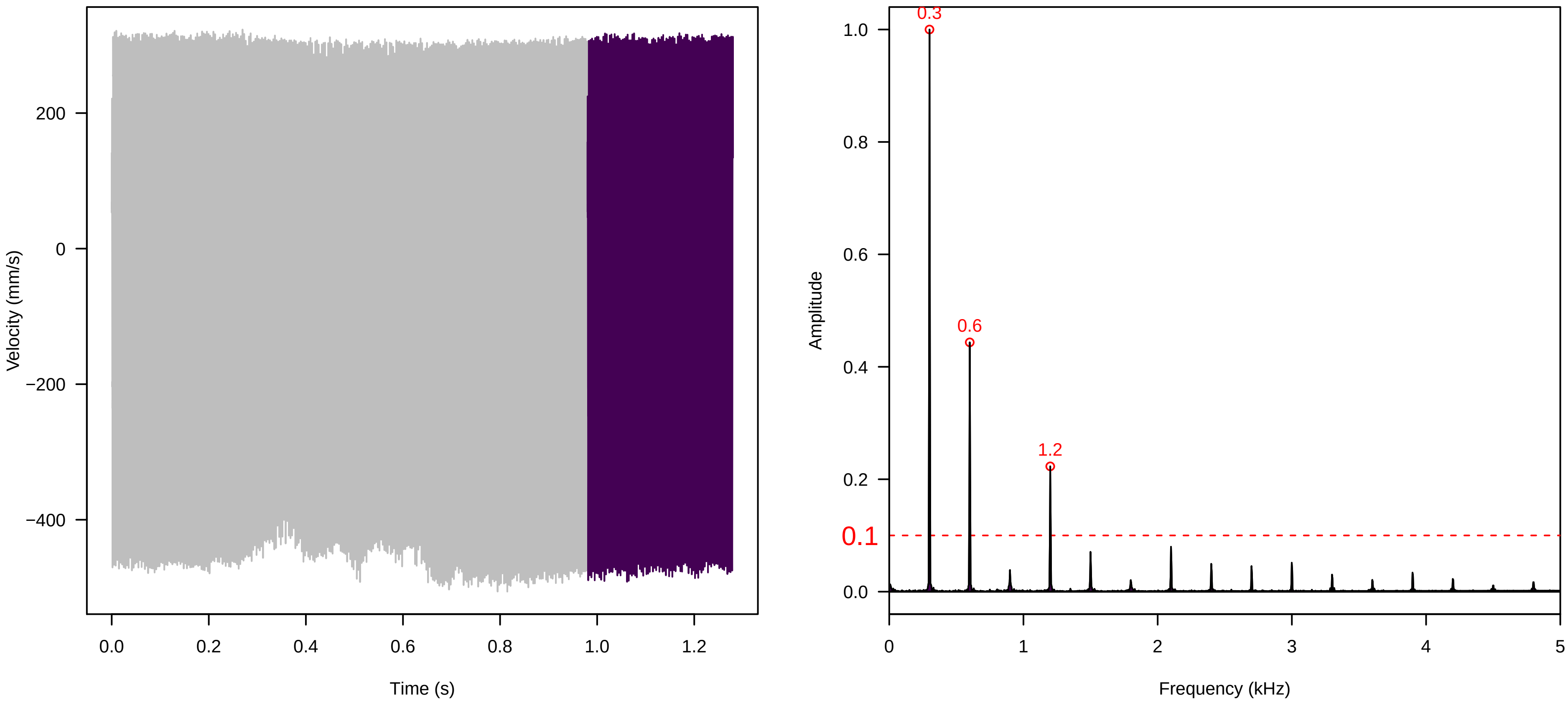

**Vel. = 0.057 ; Str. = Receptacle ; Axis = z ; Fl. accession = 10−s−77−19**

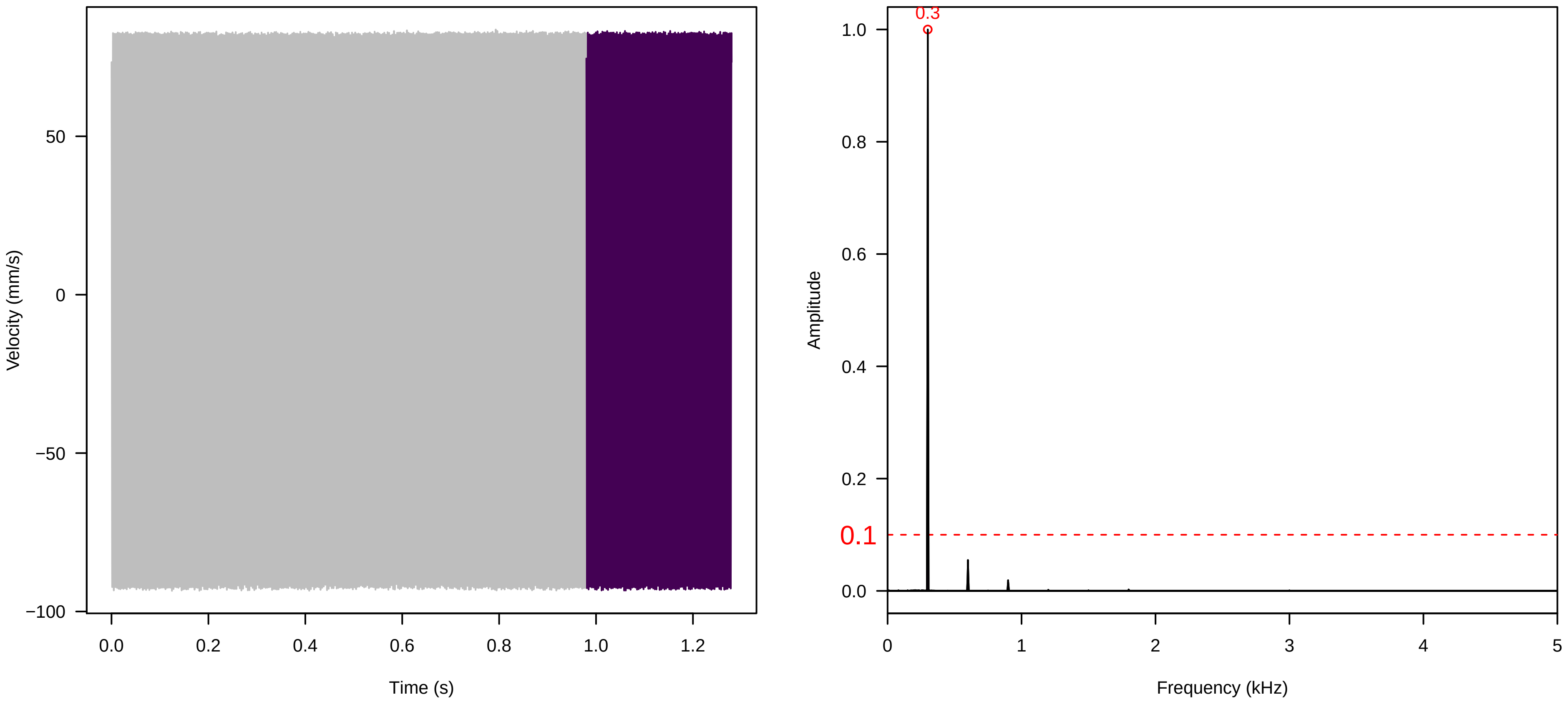


**Vel. = 0.057 ; Str. = PA ; Axis = z ; Fl. accession = 10−s−77−19**

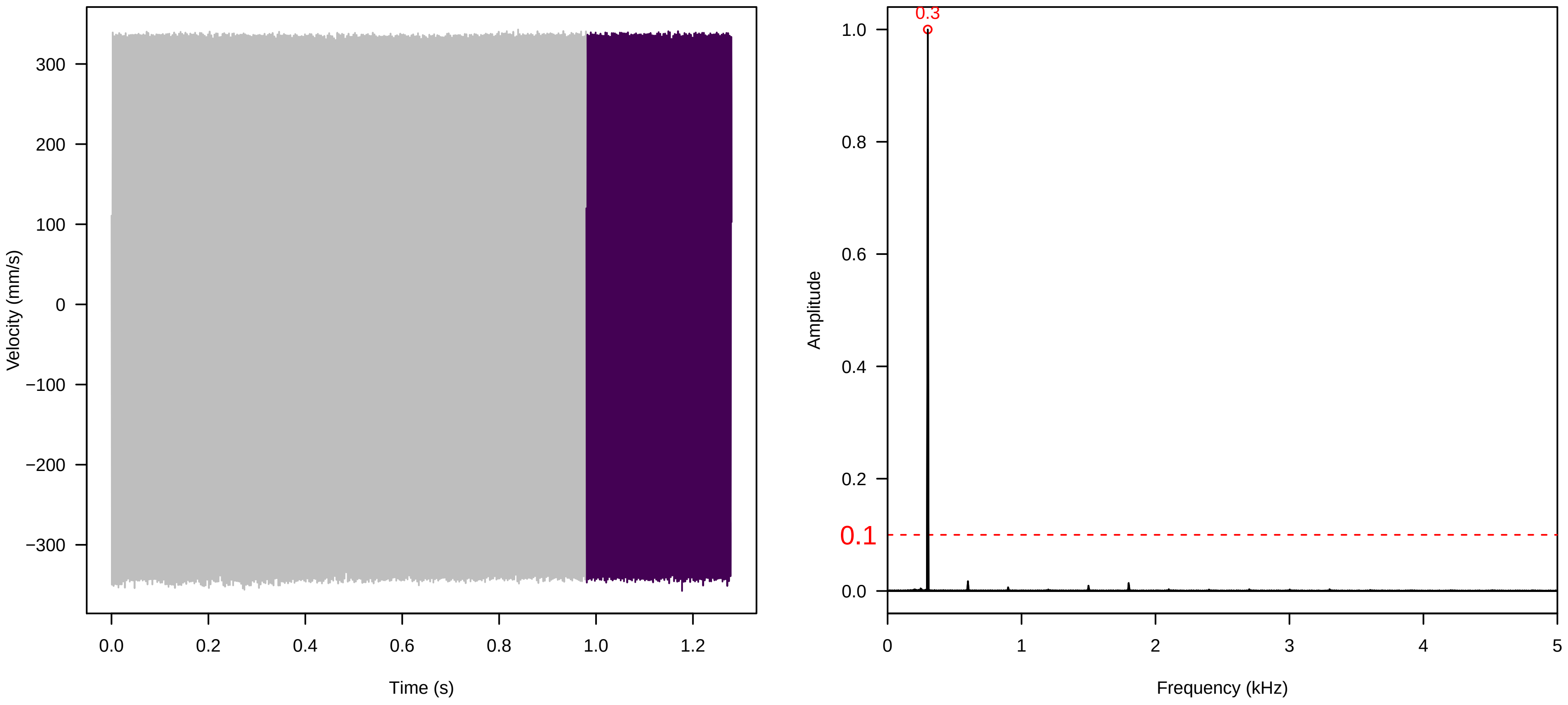


**Vel. = 0.057 ; Str. = Receptacle ; Axis = z ; Fl. accession = 10−s−77−19**

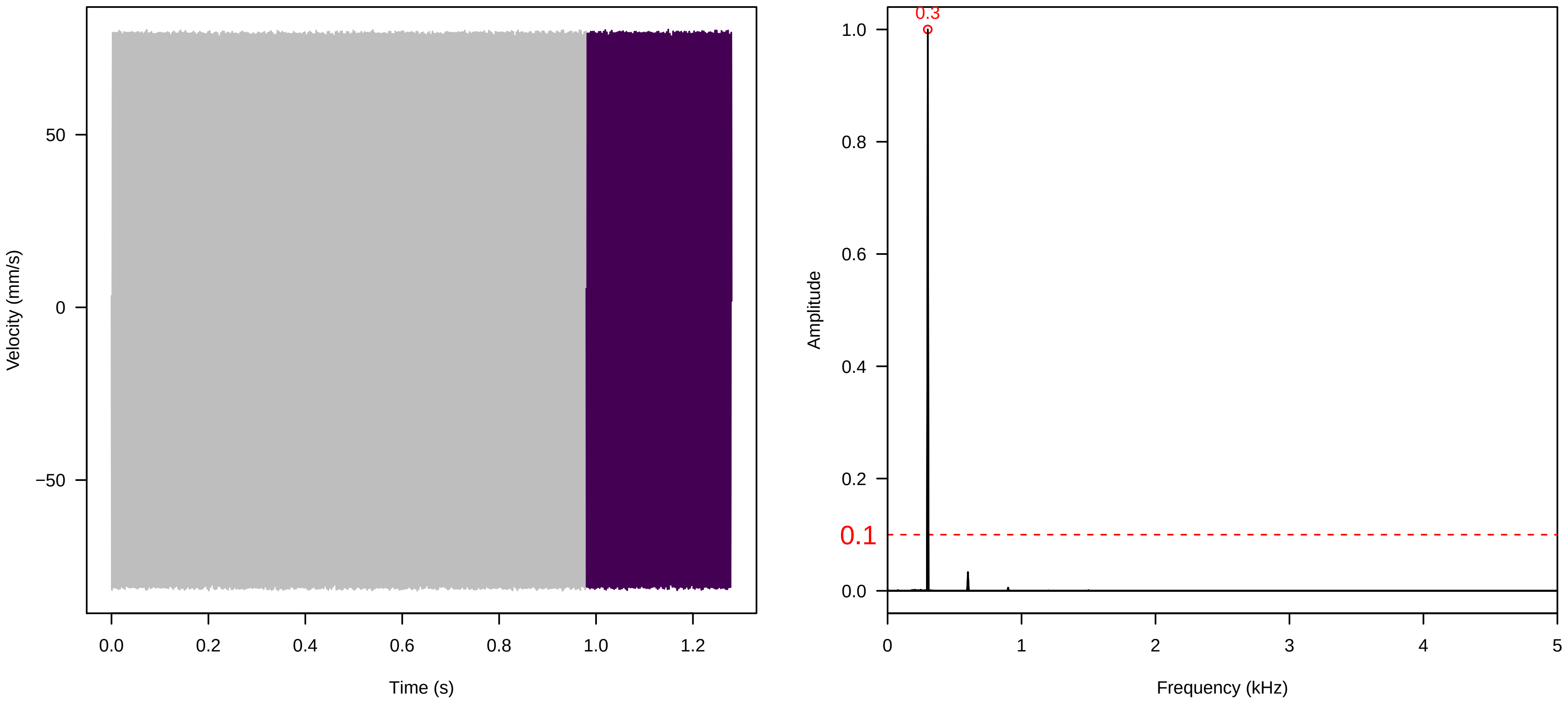

**Vel. = 0.028 ; Str. = PA ; Axis = z ; Fl. accession = 10−s−77−19**

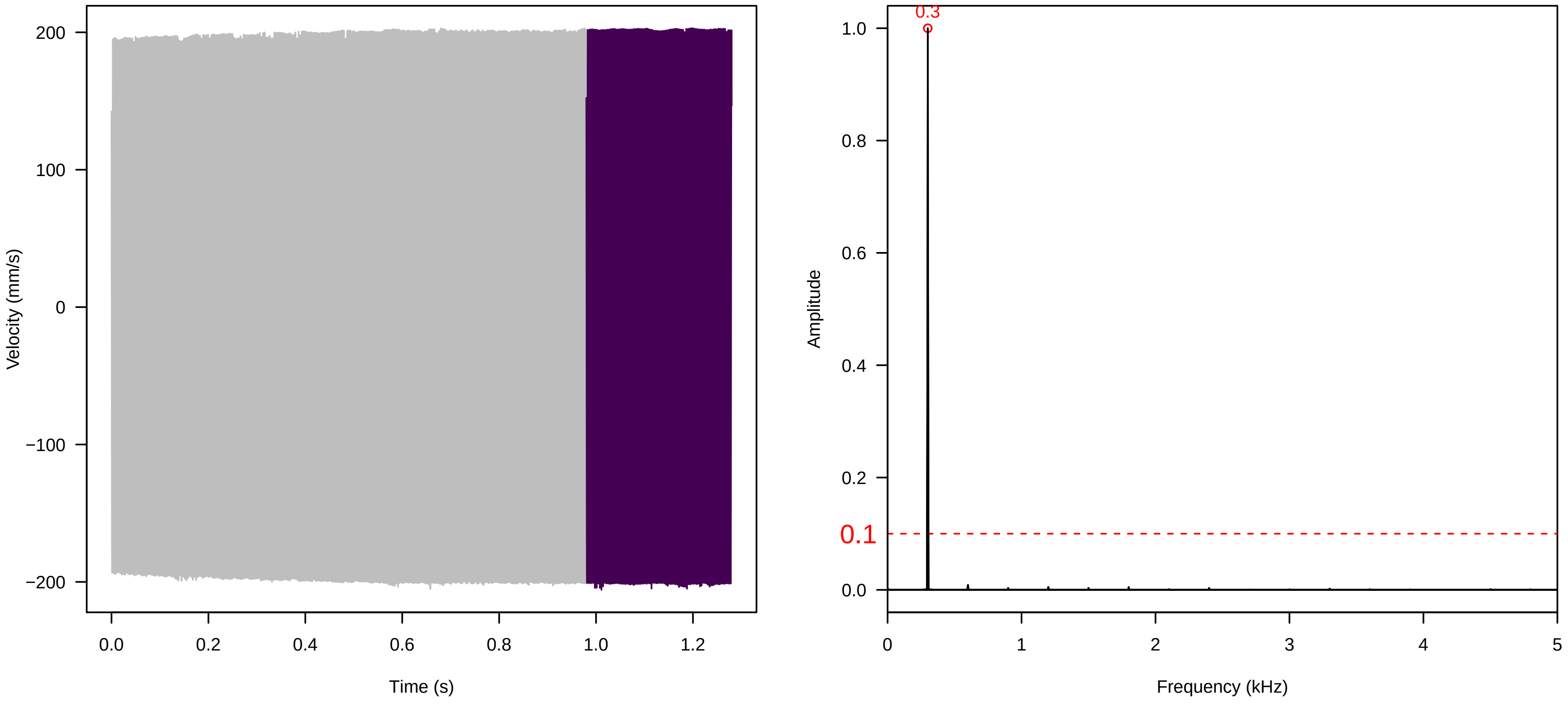


**Vel. = 0.028 ; Str. = Receptacle ; Axis = z ; Fl. accession = 10−s−77−19**

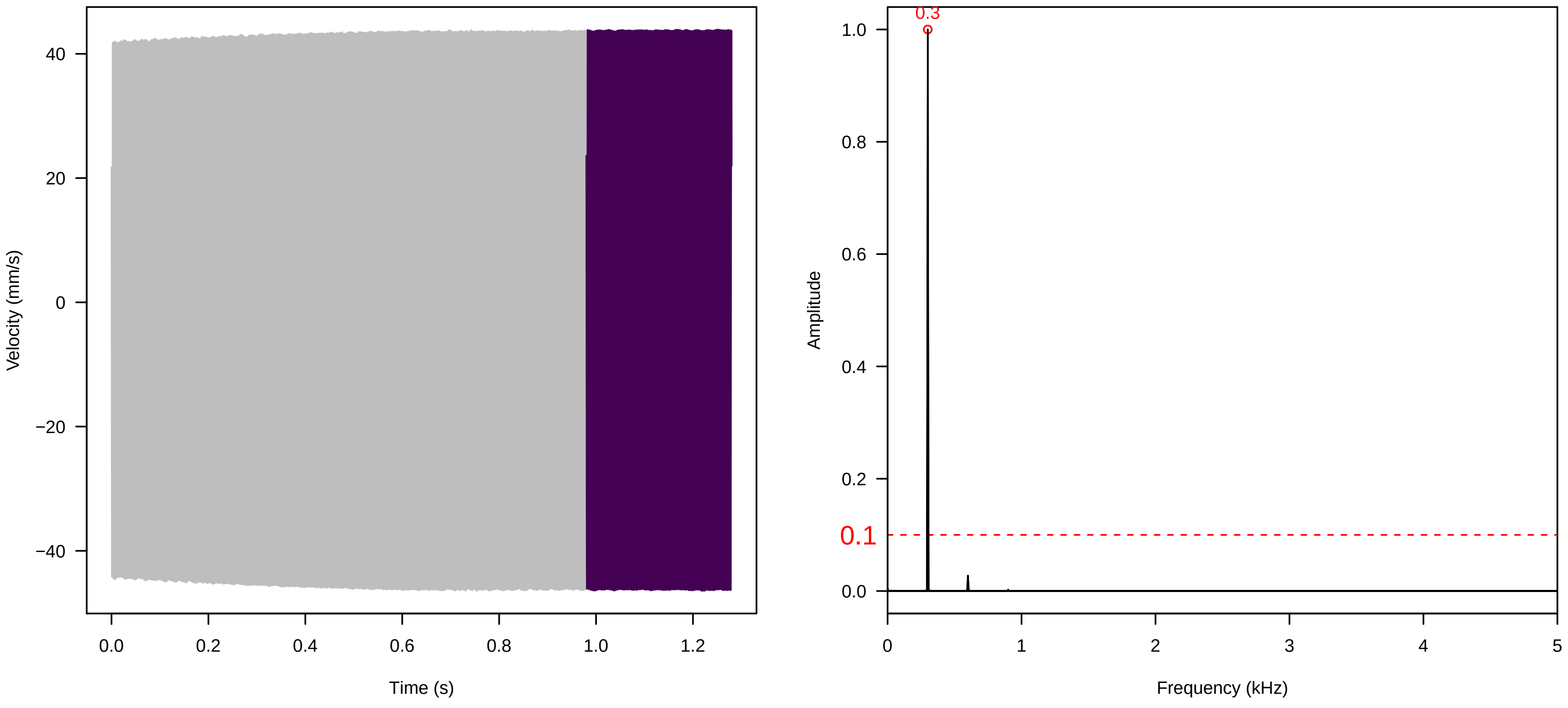


**Vel. = 0.028 ; Str. = FA ; Axis = z ; Fl. accession = 10−s−77−19**

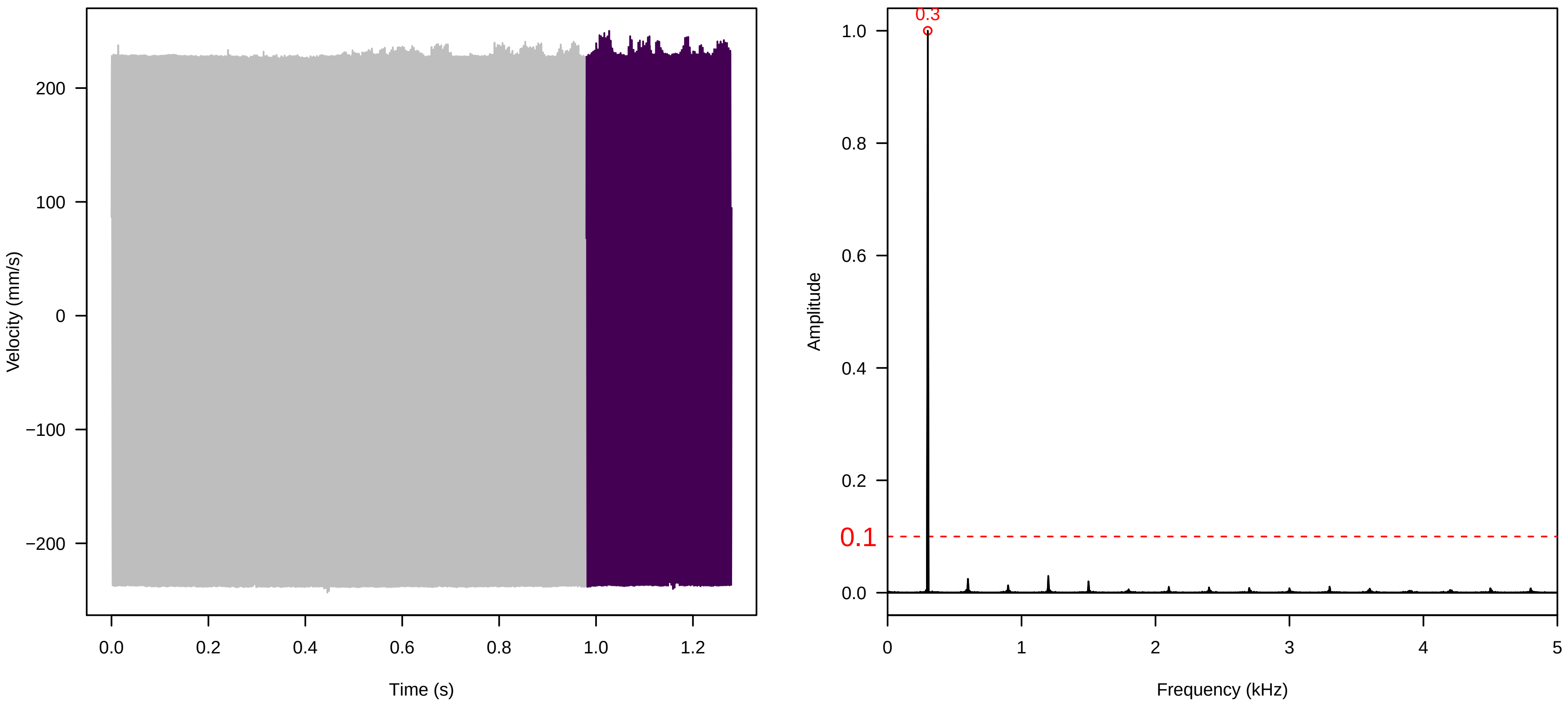

**Vel. =  0.028 ; Str. =  Receptacle ; Axis =  z ; Fl. accession = 10−s−77−19**

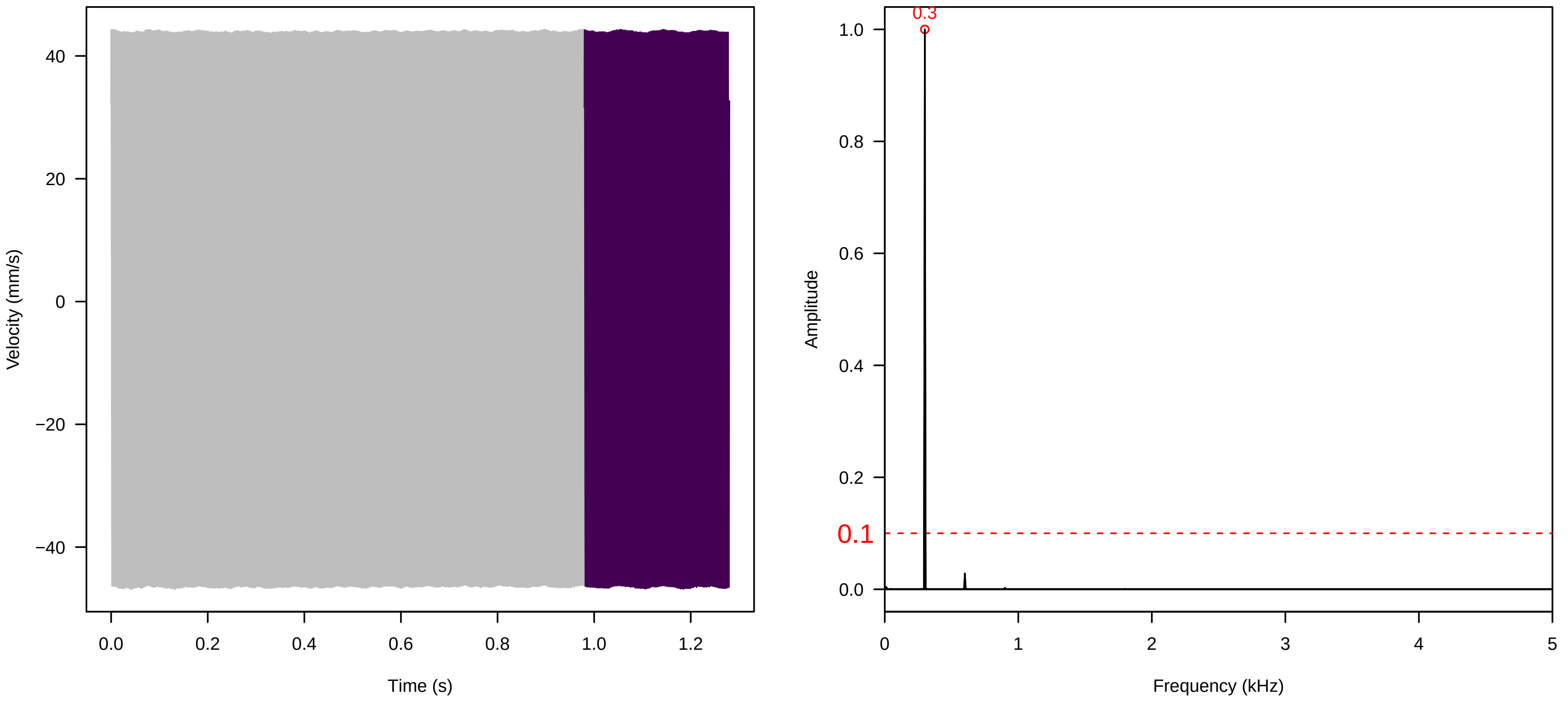


**Vel. =  0.028 ; Str. =  Corolla ; Axis =  z ; Fl. accession = 10−s−77−19**

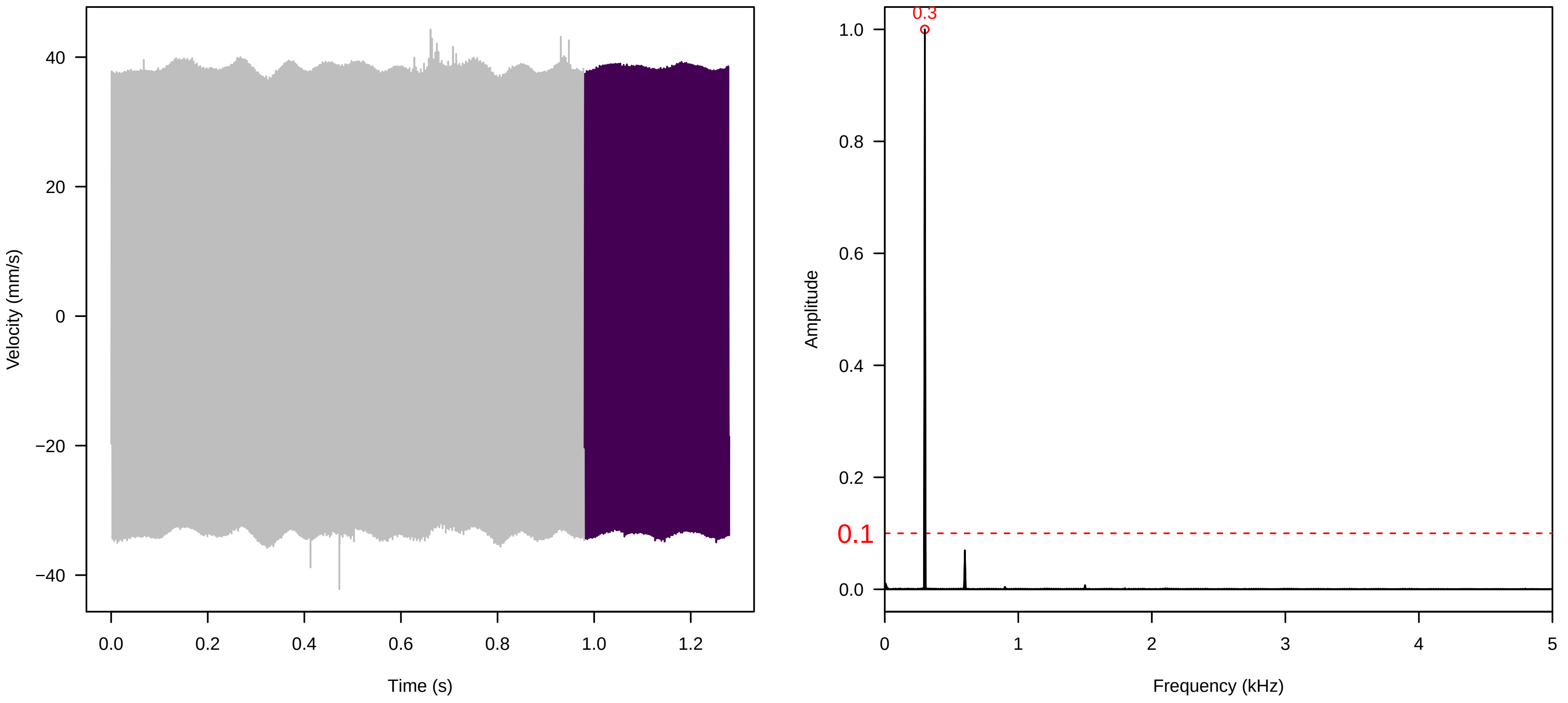


**Vel. =  0.028 ; Str. =  Receptacle ; Axis =  z ; Fl. accession = 10−s−77−19**

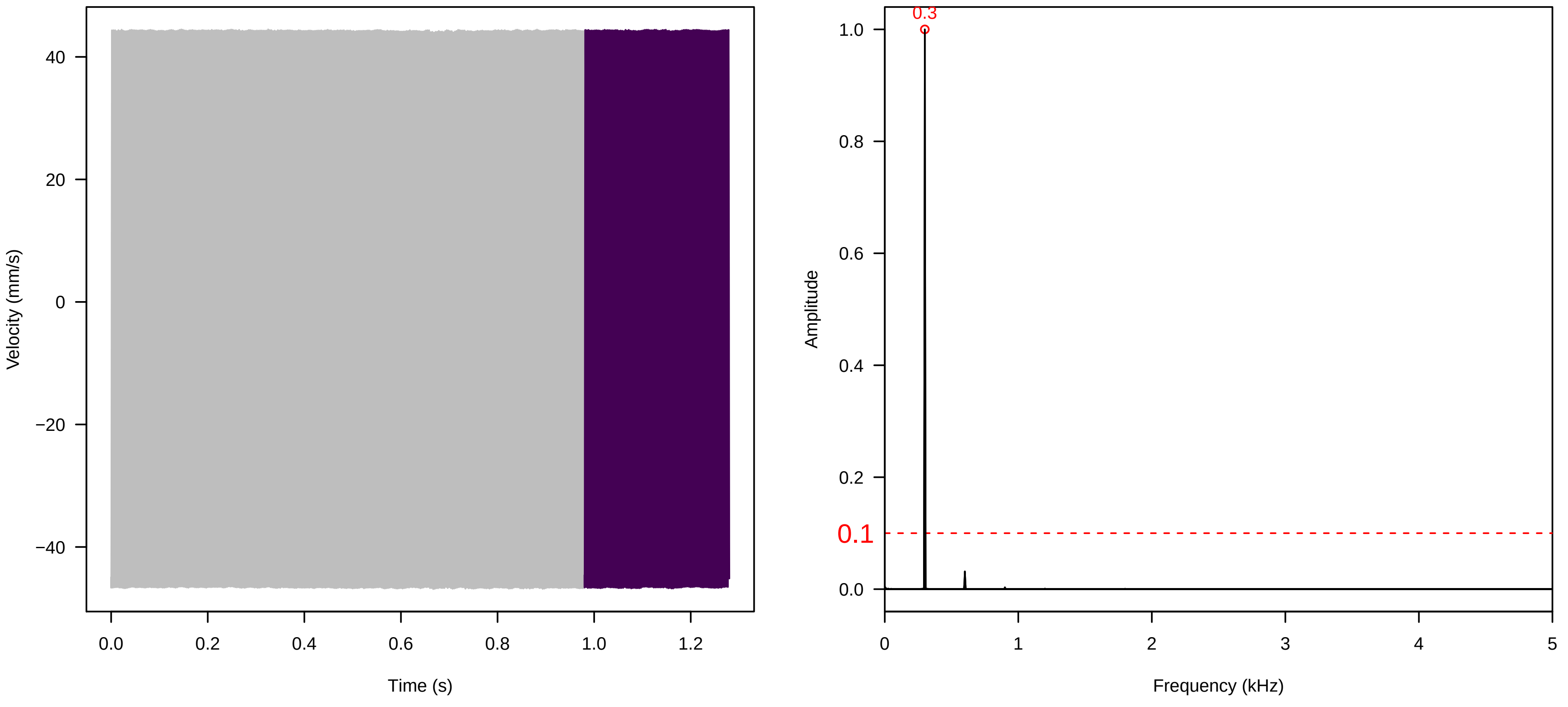

**Vel. = 0.014 ; Str. = Corolla ; Axis = z ; Fl. accession = 10−s−77−19**

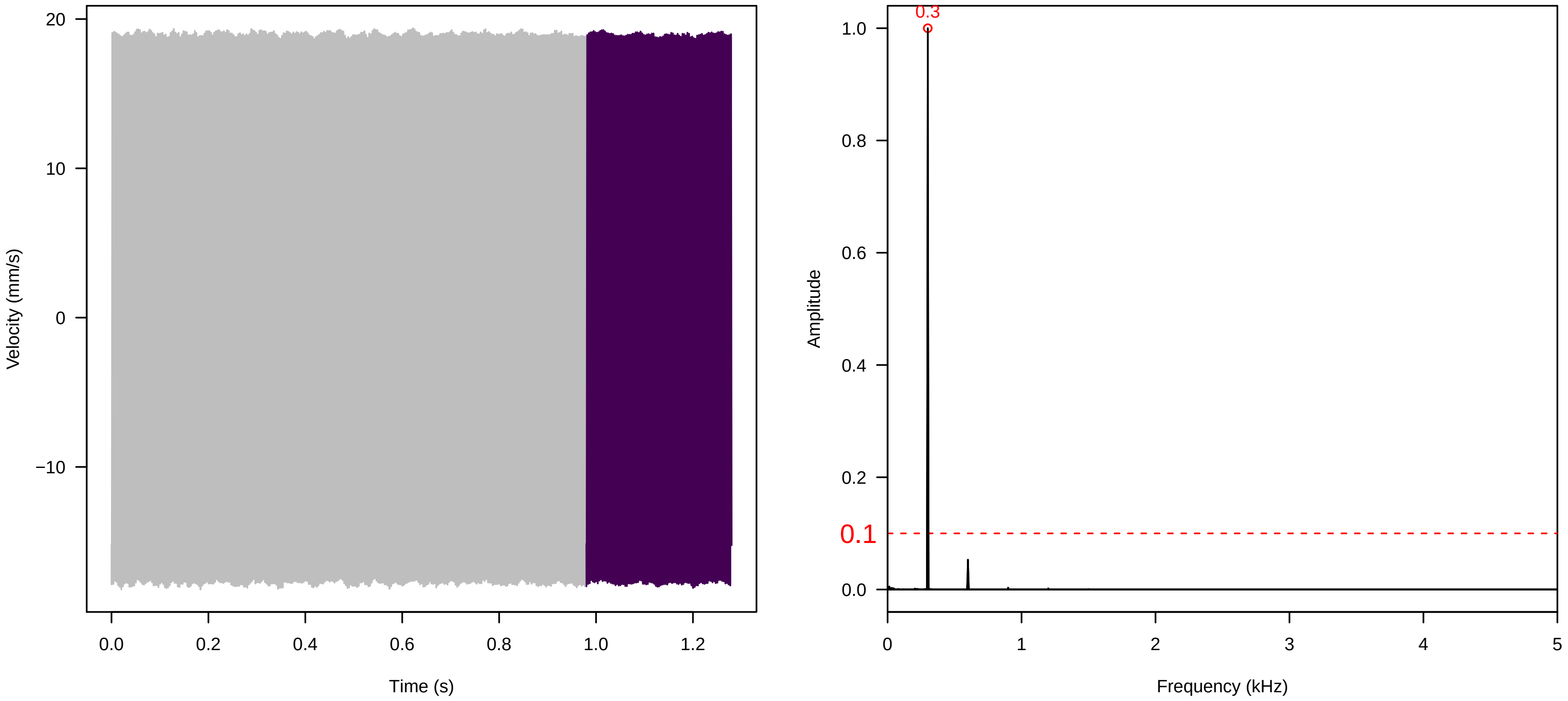


**Vel. = 0.014 ; Str. = Receptacle ; Axis = z ; Fl. accession = 10−s−77−19**

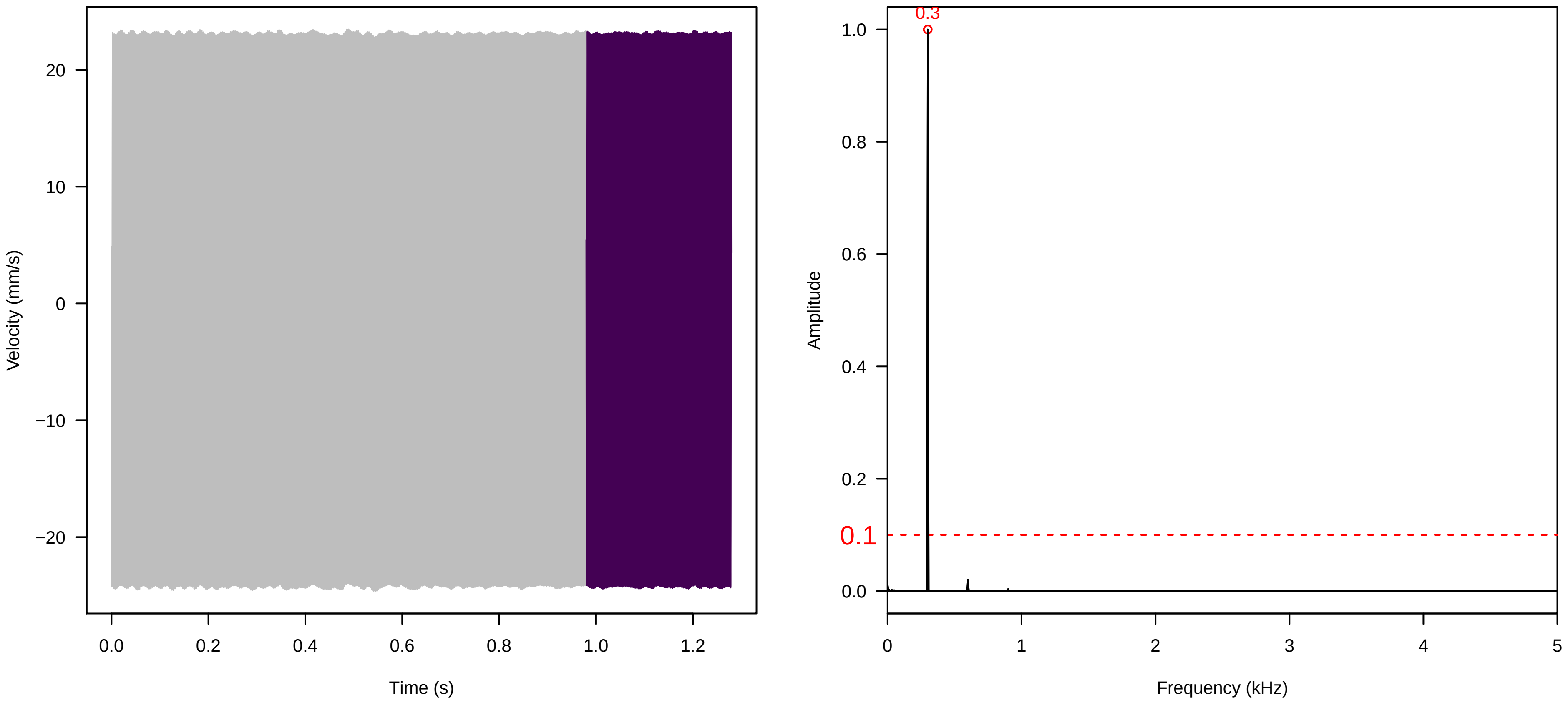


**Vel. = 0.014 ; Str. = FA ; Axis = z ; Fl. accession = 10−s−77−19**

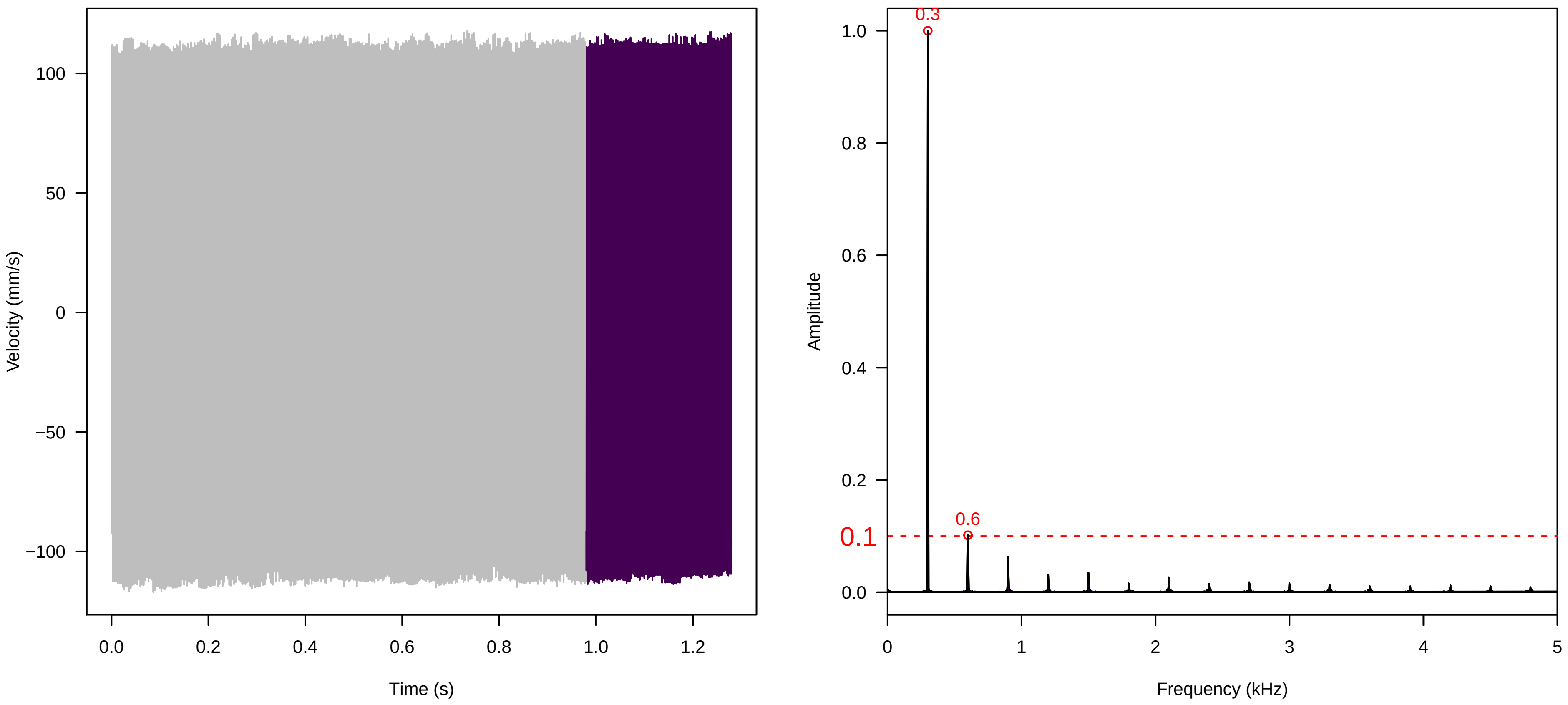

Vel. =  0.014 ; Str. =  Receptacle ; Axis =  z ; Fl. accession = 10−s−77−19

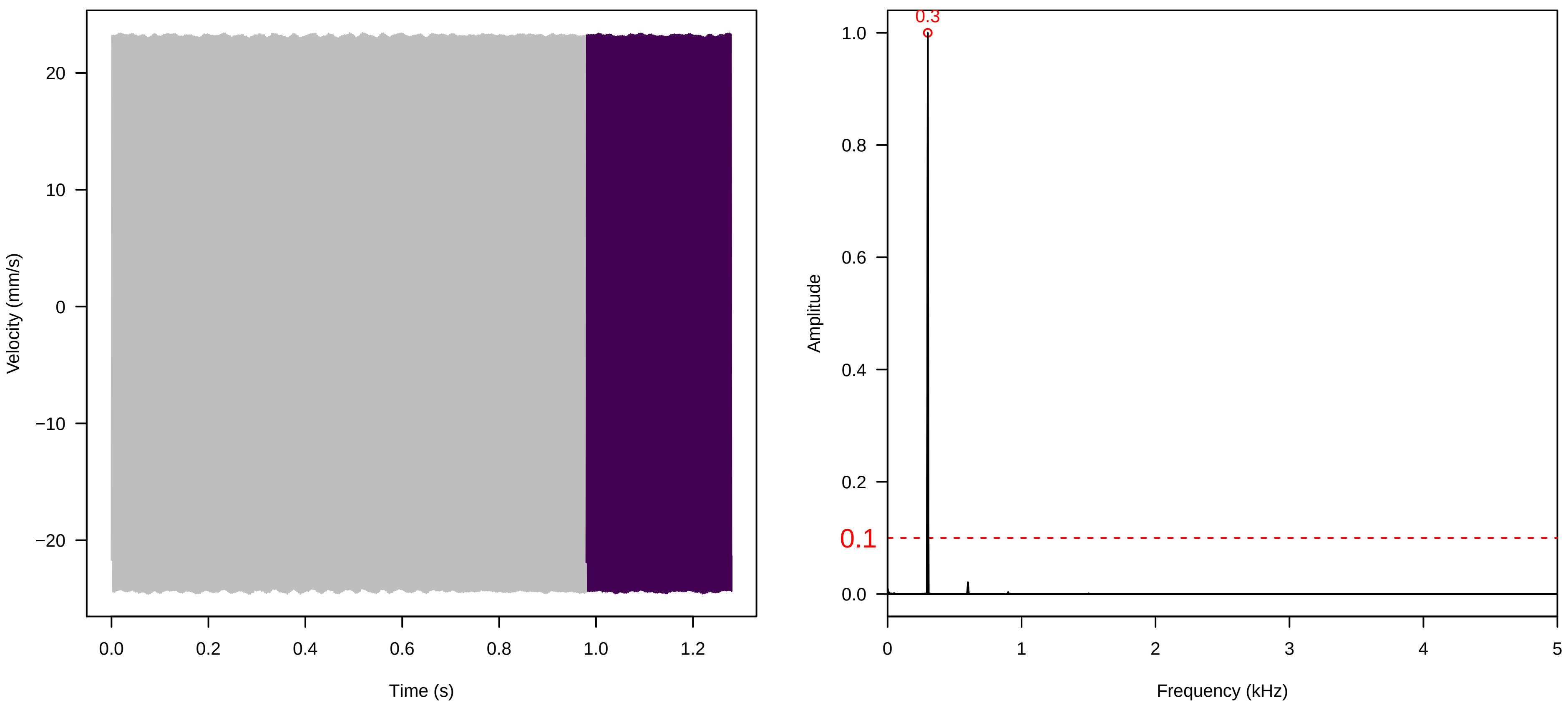


Vel. =  0.014 ; Str. =  PA ; Axis =  z ; Fl. accession = 10−s−77−19

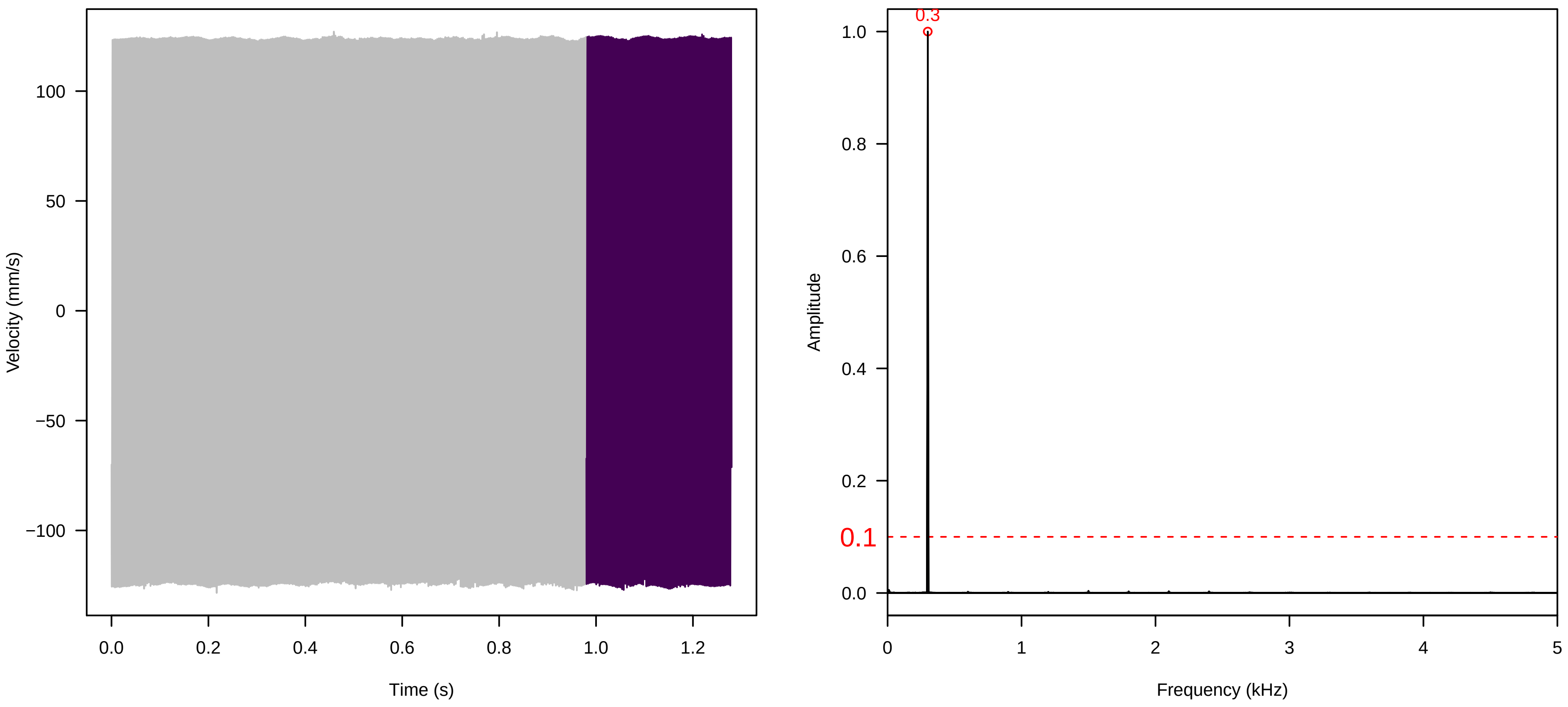


Vel. =  0.014 ; Str. =  Receptacle ; Axis =  z ; Fl. accession = 10−s−77−19

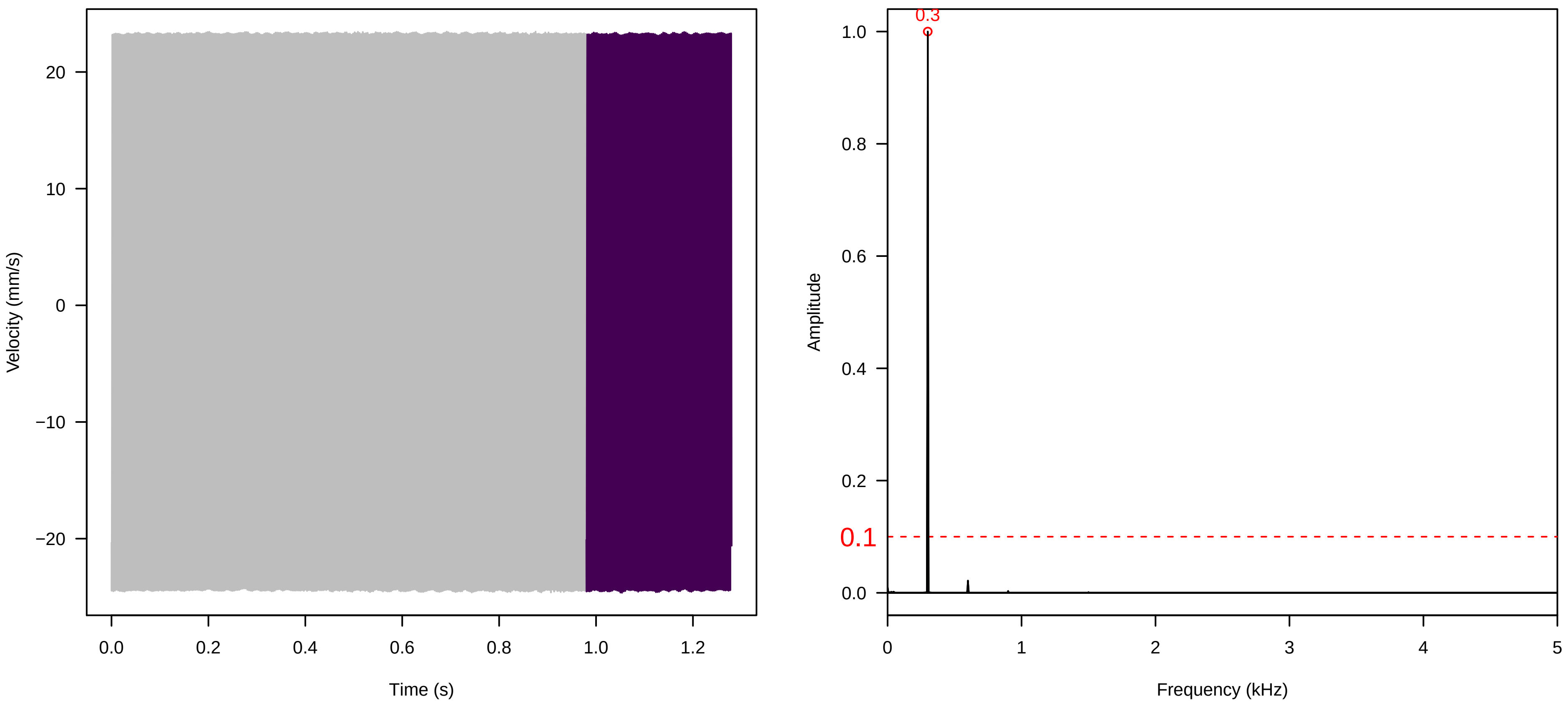

**Vel. = 0.014 ; Str. = Corolla ; Axis = y ; Fl. accession = 10−s−77−19**

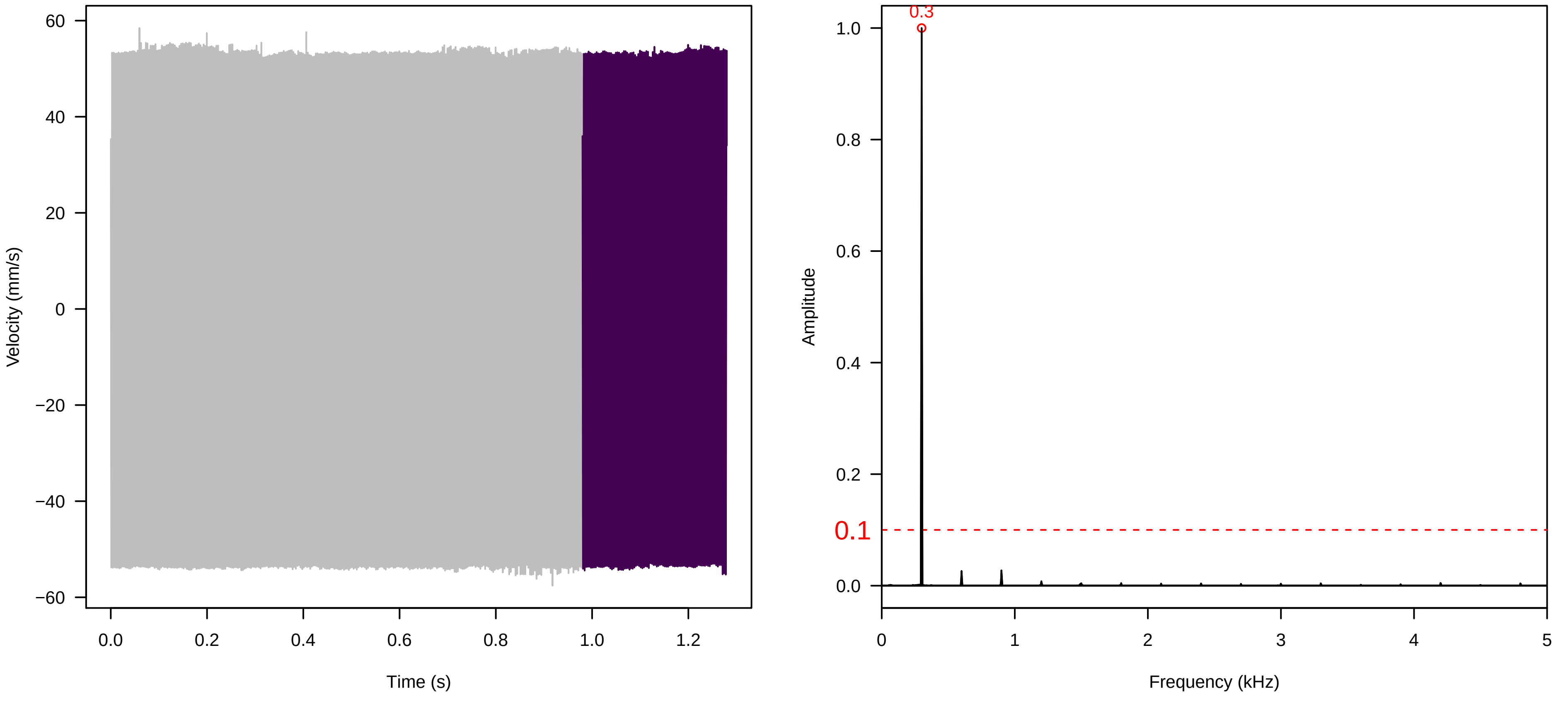


**Vel. = 0.014 ; Str. = Receptacle ; Axis = y ; Fl. accession = 10−s−77−19**

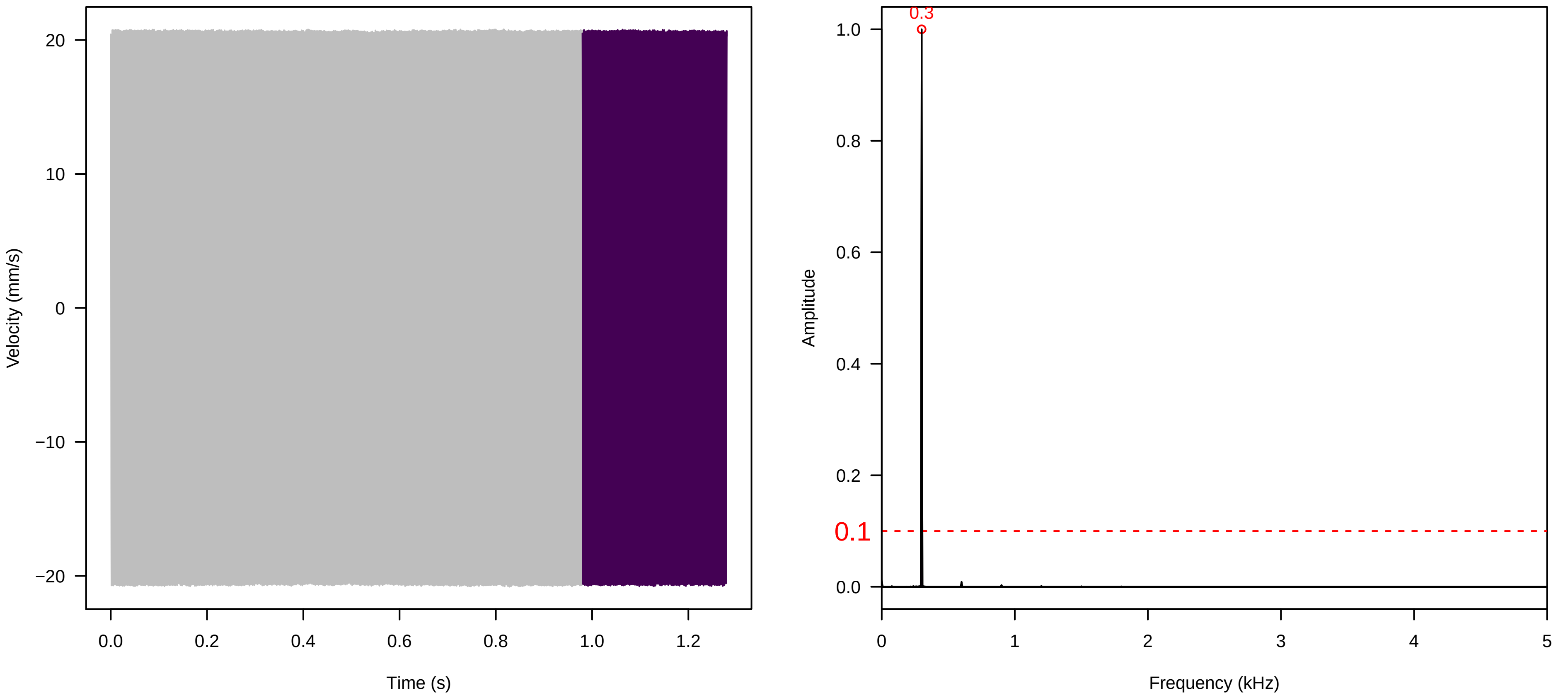


**Vel. = 0.014 ; Str. = FA ; Axis = y ; Fl. accession = 10−s−77−19**

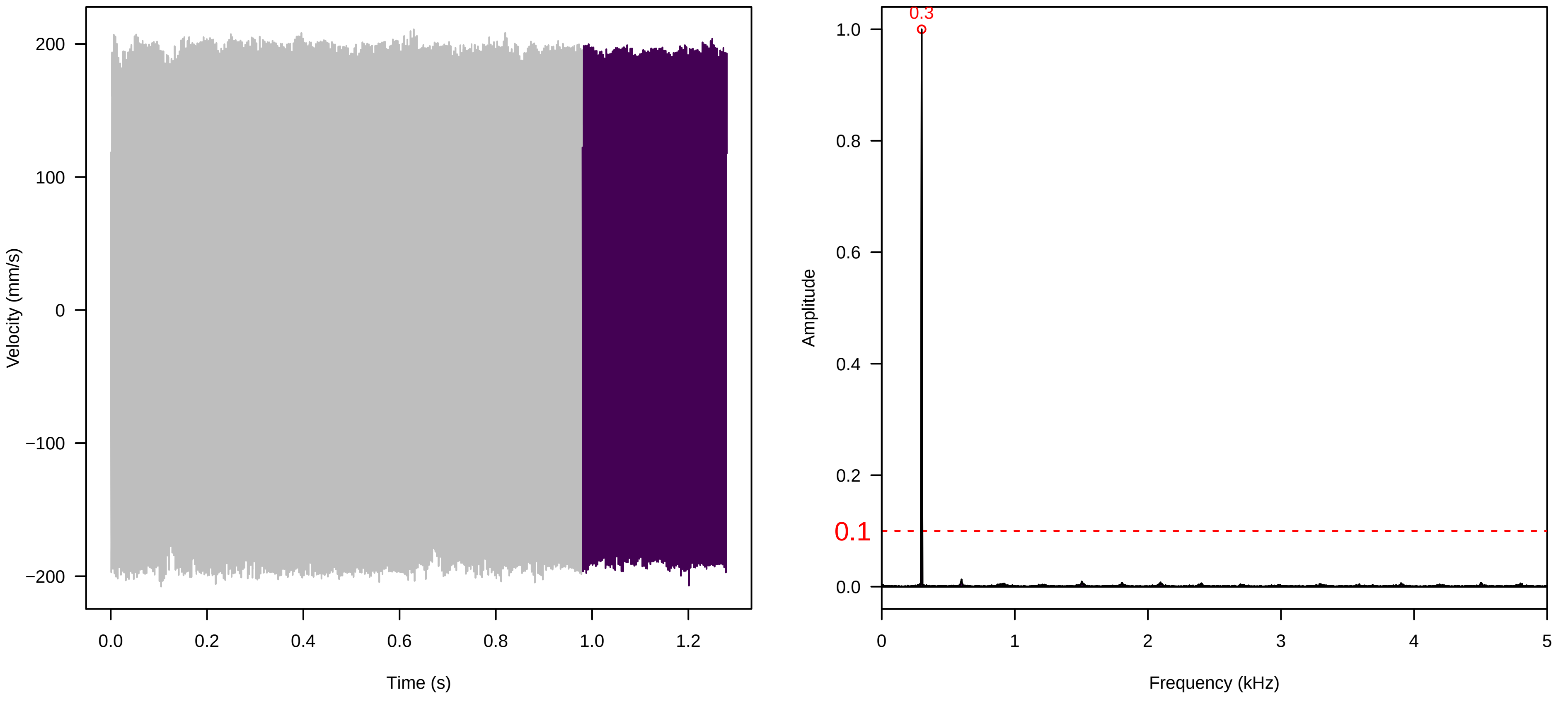

**Vel. = 0.014 ; Str. = Receptacle ; Axis = y ; Fl. accession = 10−s−77−19**

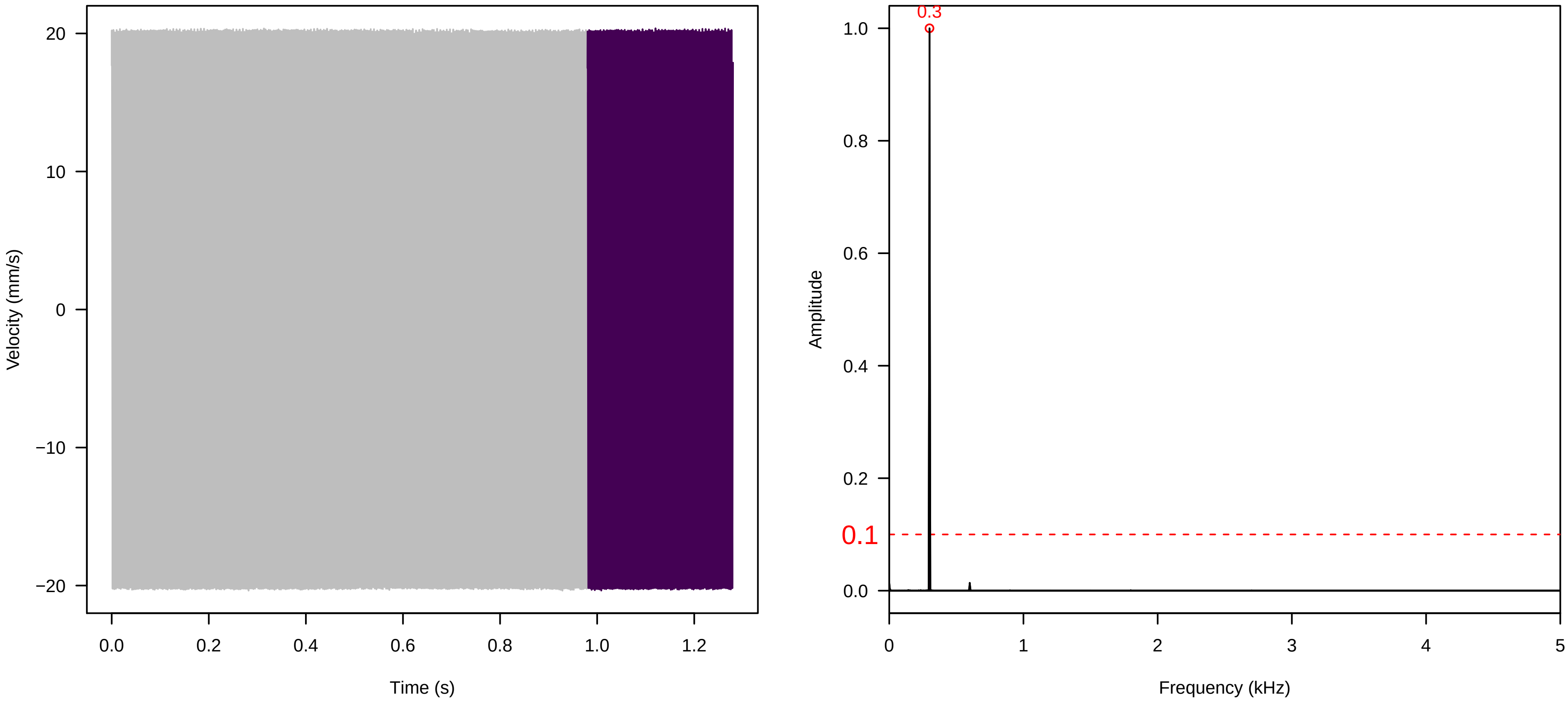


**Vel. = 0.014 ; Str. = PA ; Axis = y ; Fl. accession = 10−s−77−19**

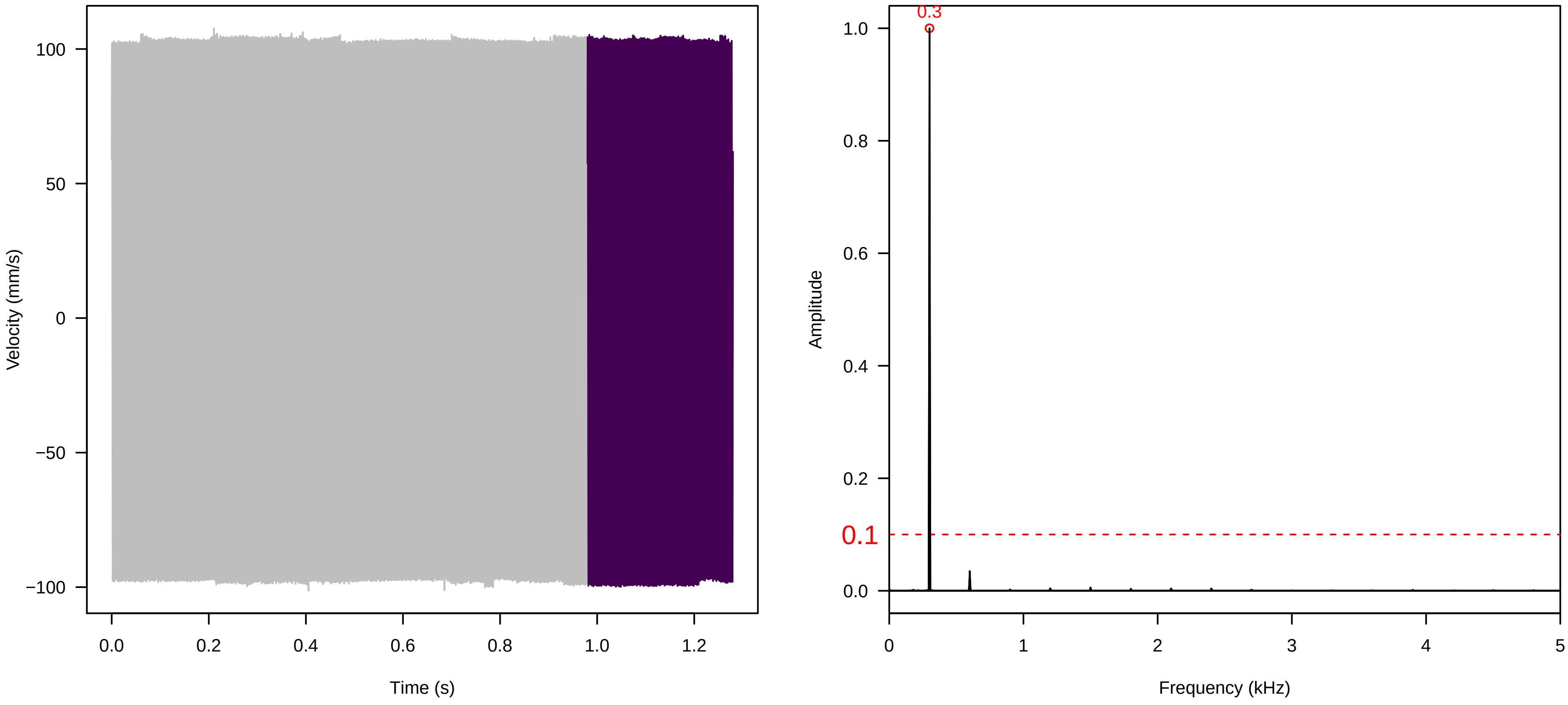


**Vel. = 0.014 ; Str. = Receptacle ; Axis = y ; Fl. accession = 10−s−77−19**

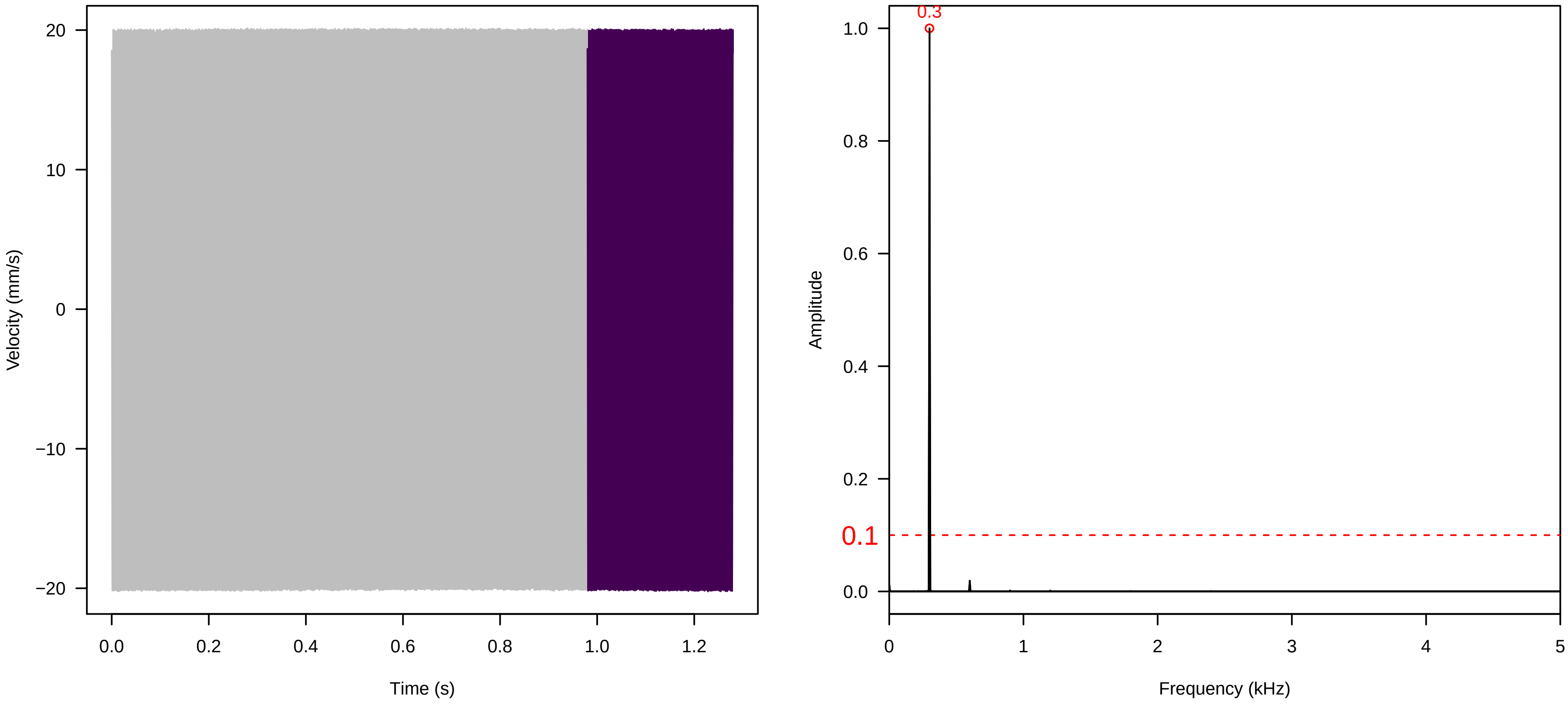

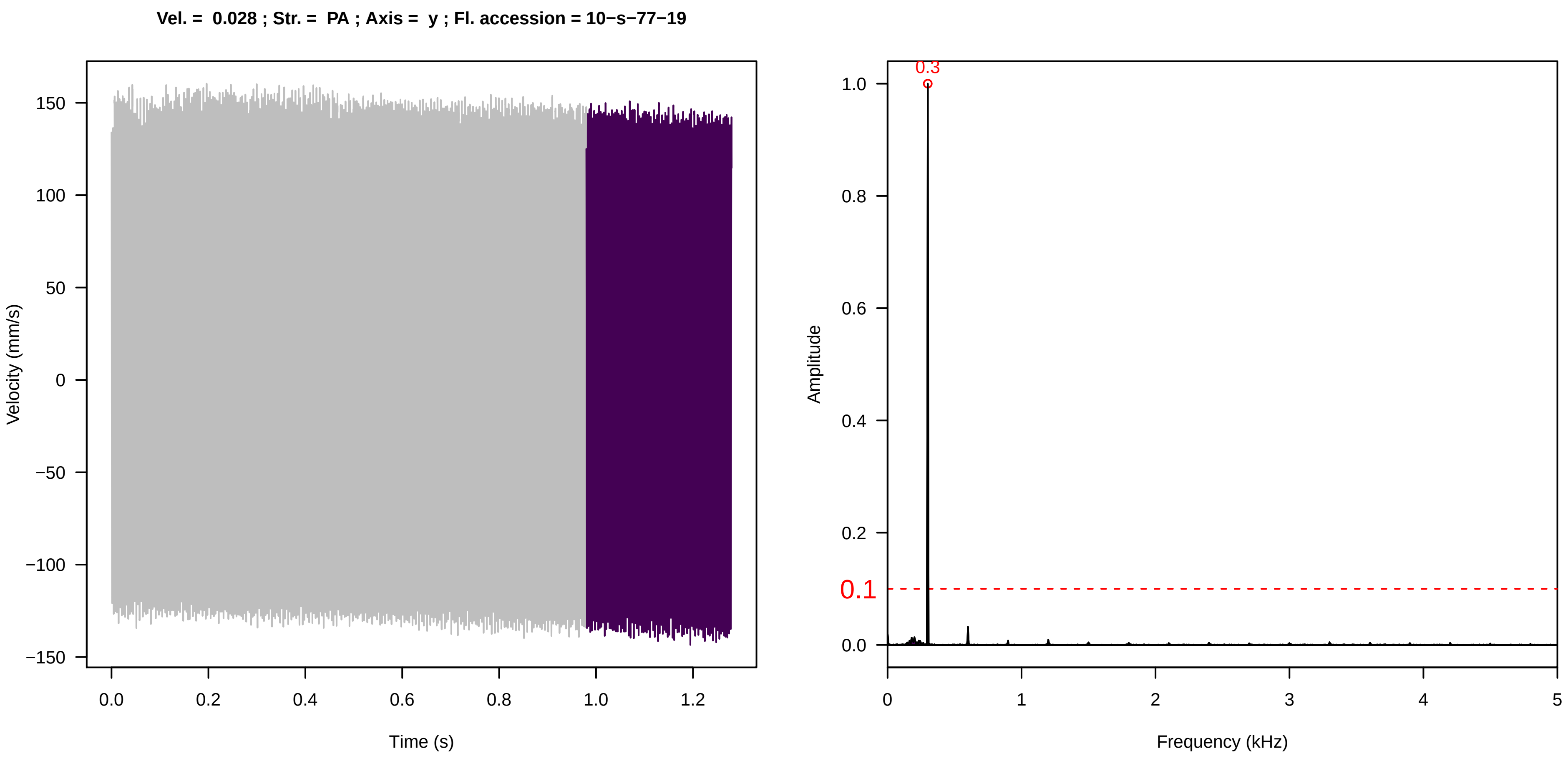
Vel. = 0.028 ; Str. = PA ; Axis = y ; Fl. accession = 10−s−77−19
Velocity (mm/s)
Time (s)
Amplitude
Frequency (kHz)
0.3
0.1

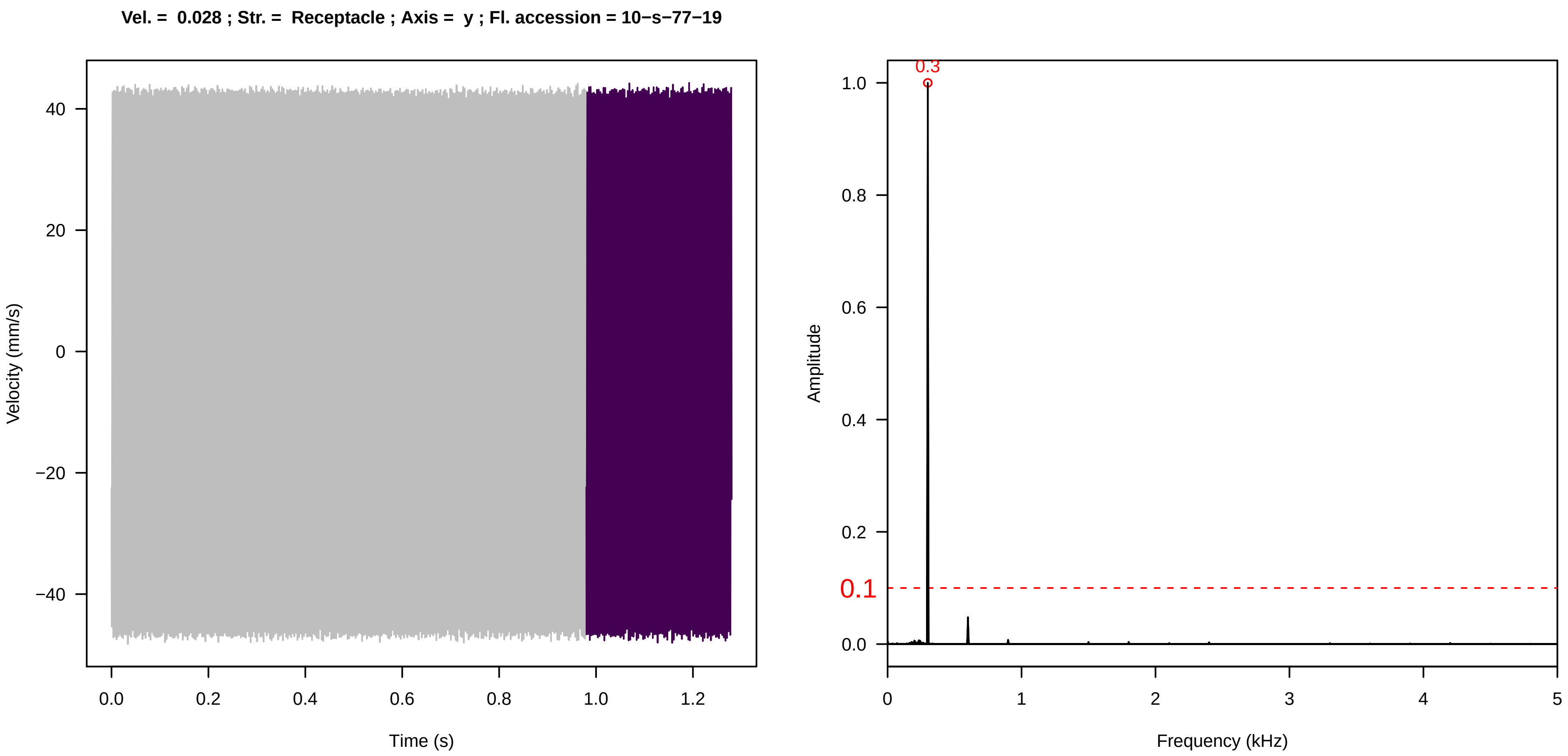
Vel. = 0.028 ; Str. = Receptacle ; Axis = y ; Fl. accession = 10−s−77−19
Velocity (mm/s)
Time (s)
Amplitude
Frequency (kHz)
0.3
0.1

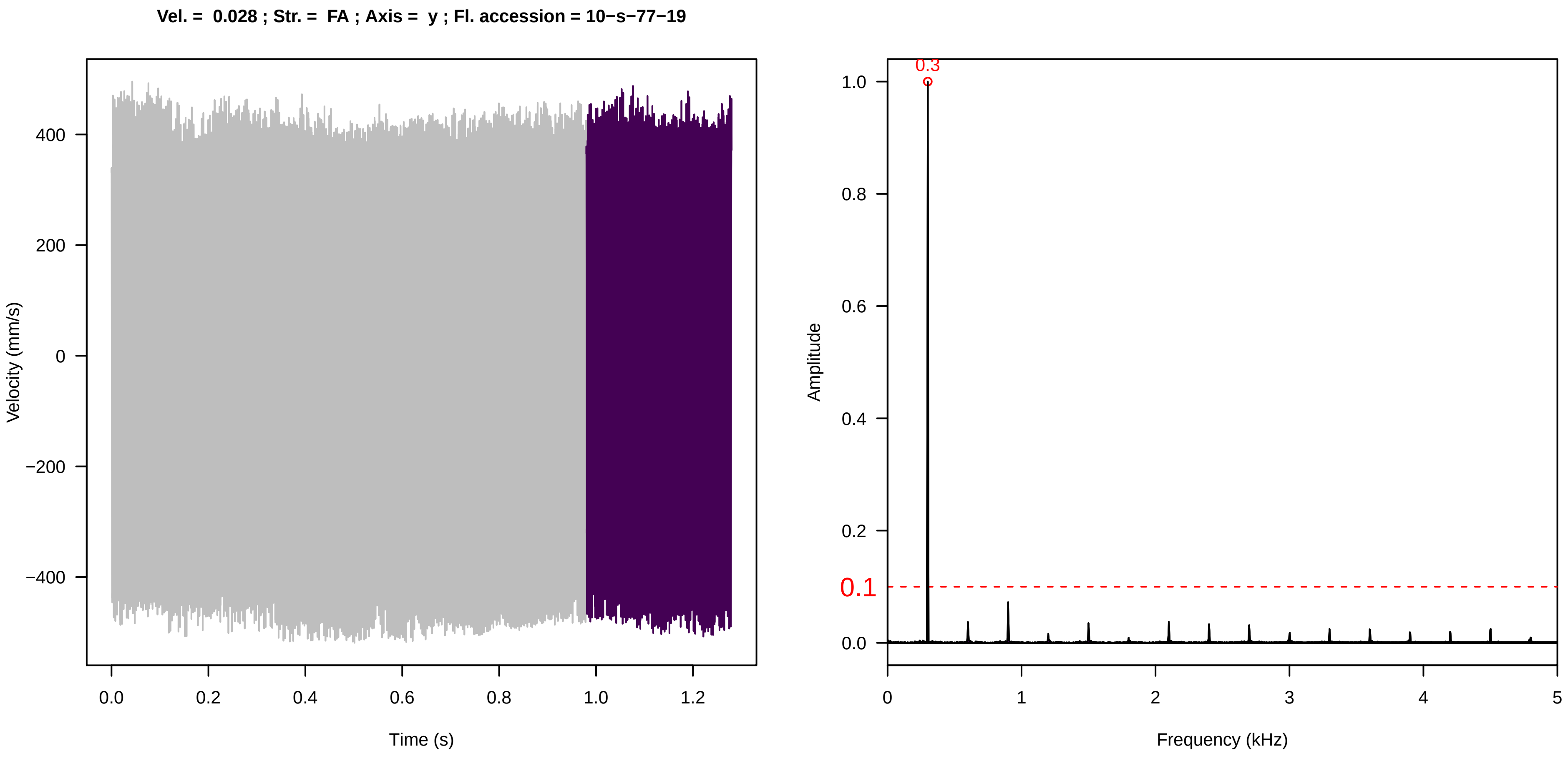
Vel. = 0.028 ; Str. = FA ; Axis = y ; Fl. accession = 10−s−77−19
Velocity (mm/s)
Time (s)
Amplitude
Frequency (kHz)
0.3
0.1

**Vel. =  0.028 ; Str. =  Receptacle ; Axis =  y ; Fl. accession = 10−s−77−19**

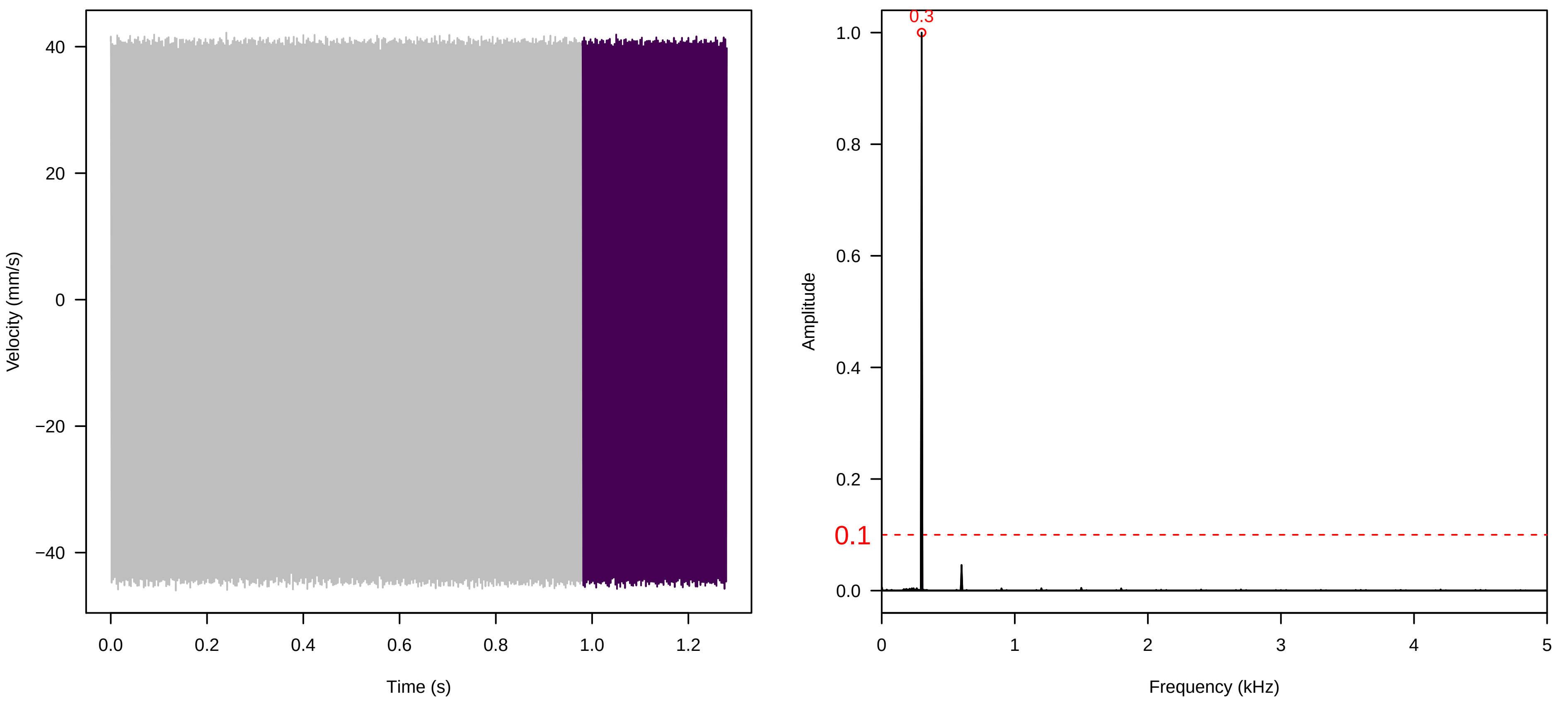


**Vel. =  0.028 ; Str. =  Corolla ; Axis =  y ; Fl. accession = 10−s−77−19**

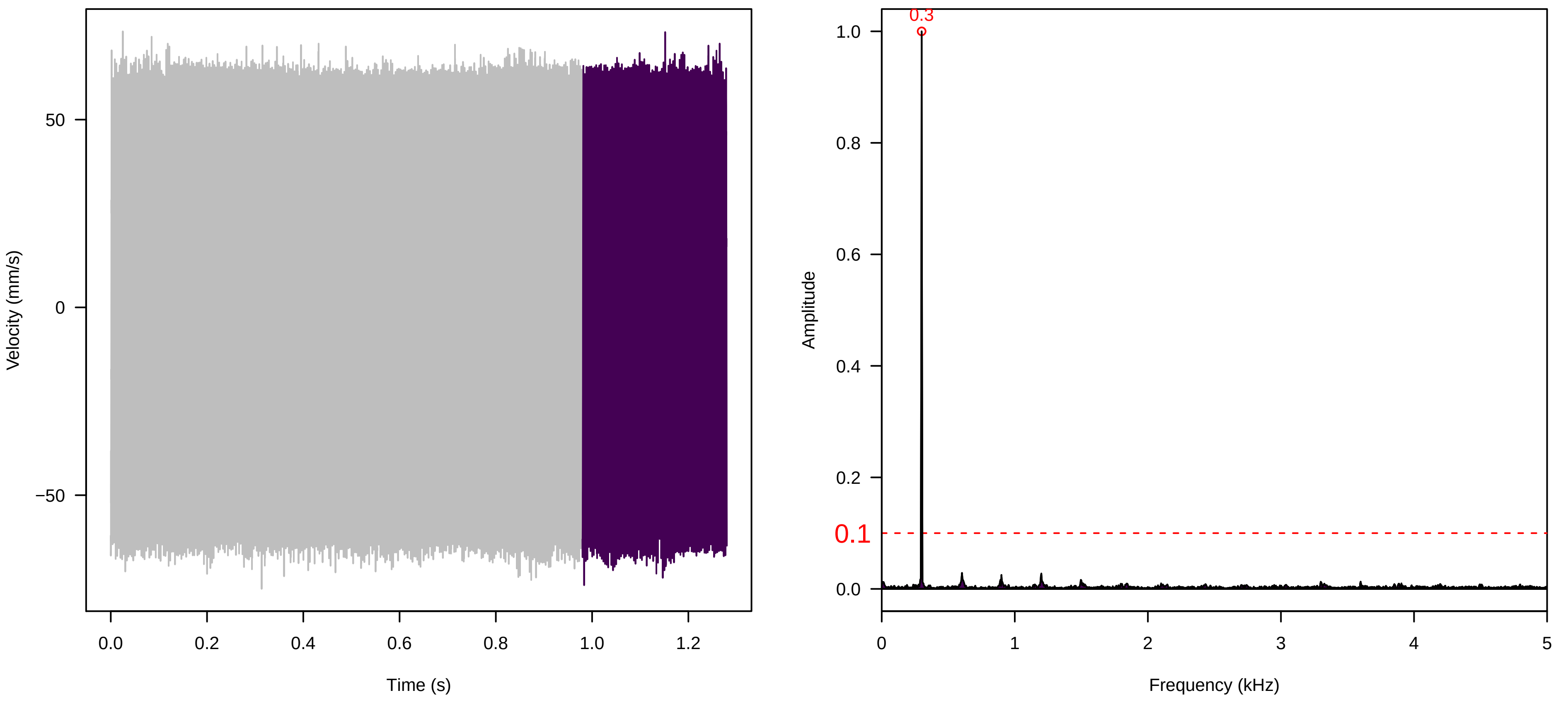


**Vel. =  0.028 ; Str. =  Receptacle ; Axis =  y ; Fl. accession = 10−s−77−19**

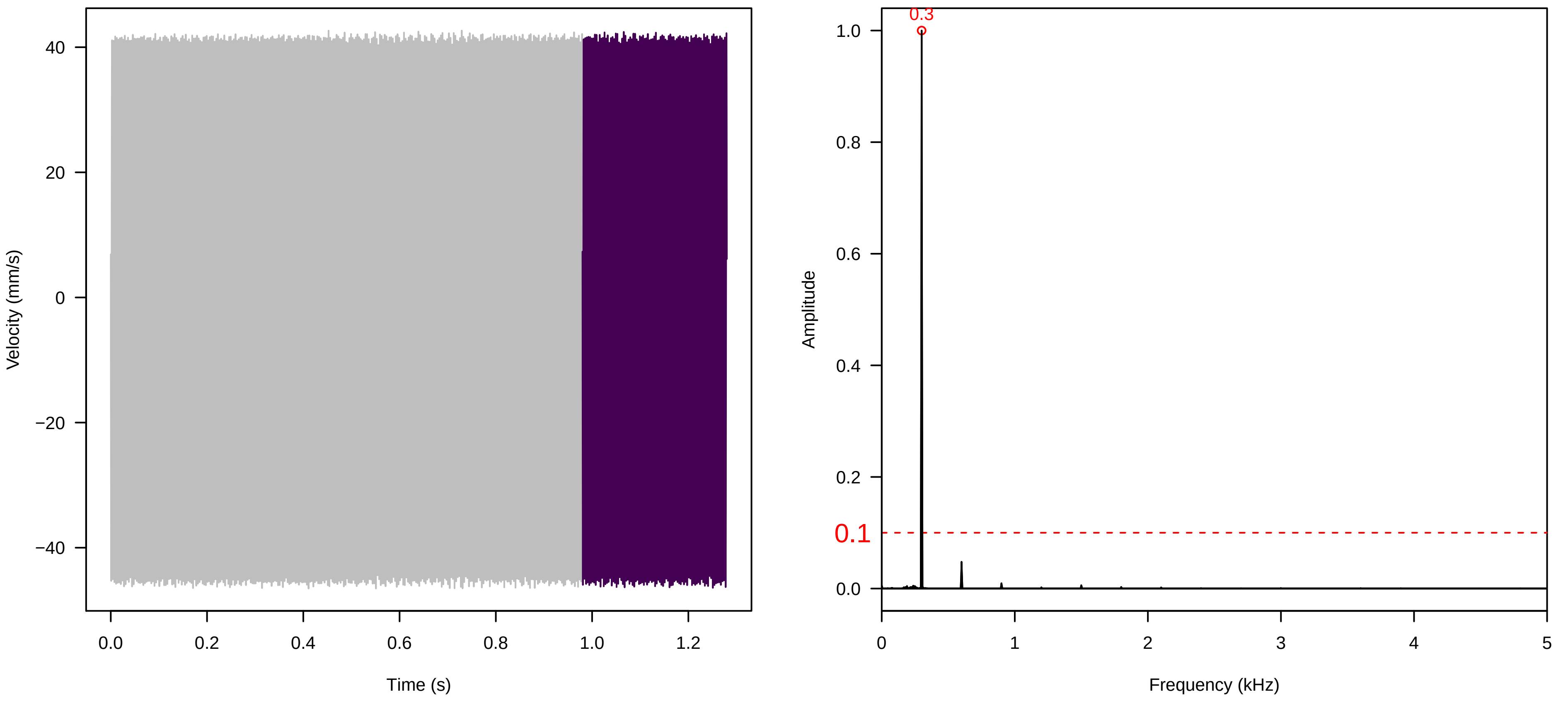

**Vel. = 0.057 ; Str. = Corolla ; Axis = y ; Fl. accession = 10−s−77−19**

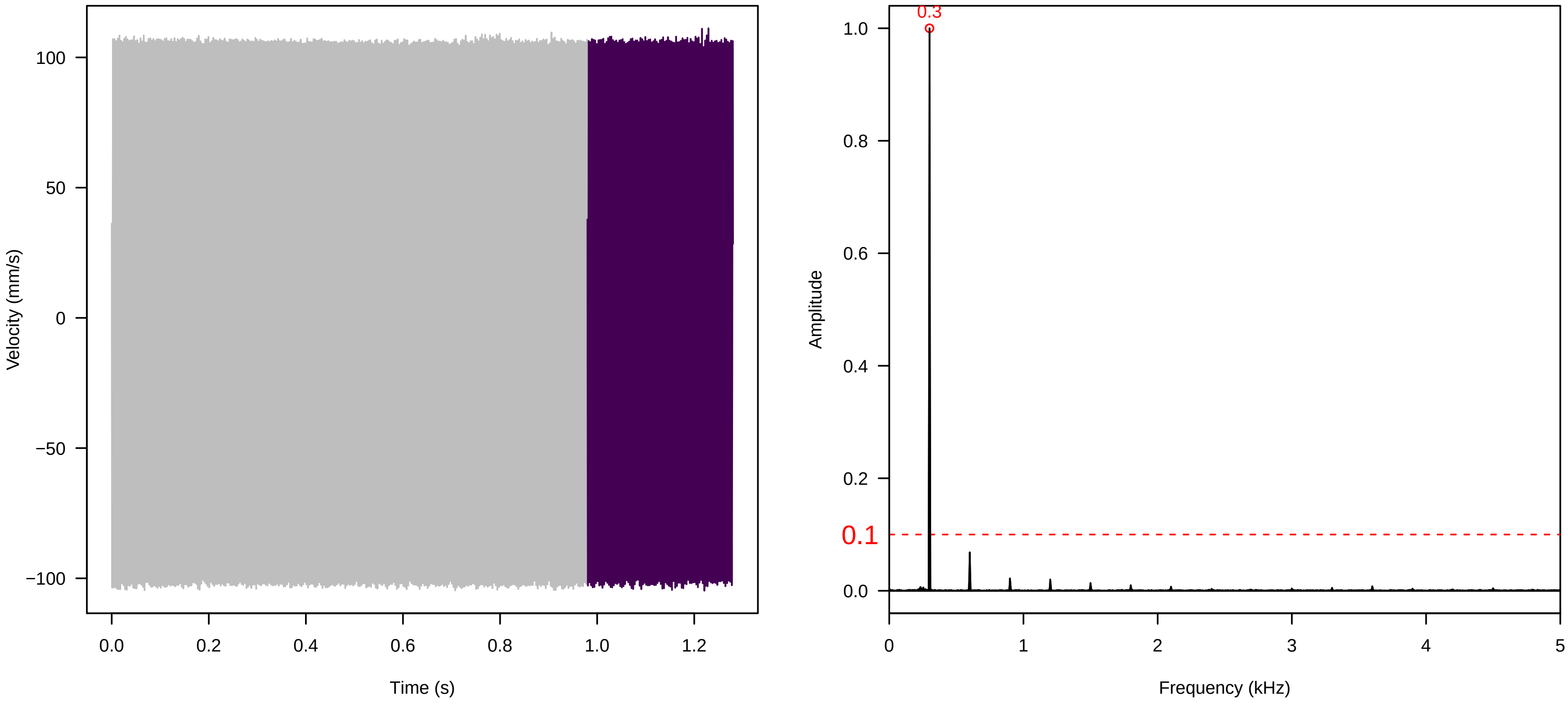


**Vel. = 0.057 ; Str. = Receptacle ; Axis = y ; Fl. accession = 10−s−77−19**

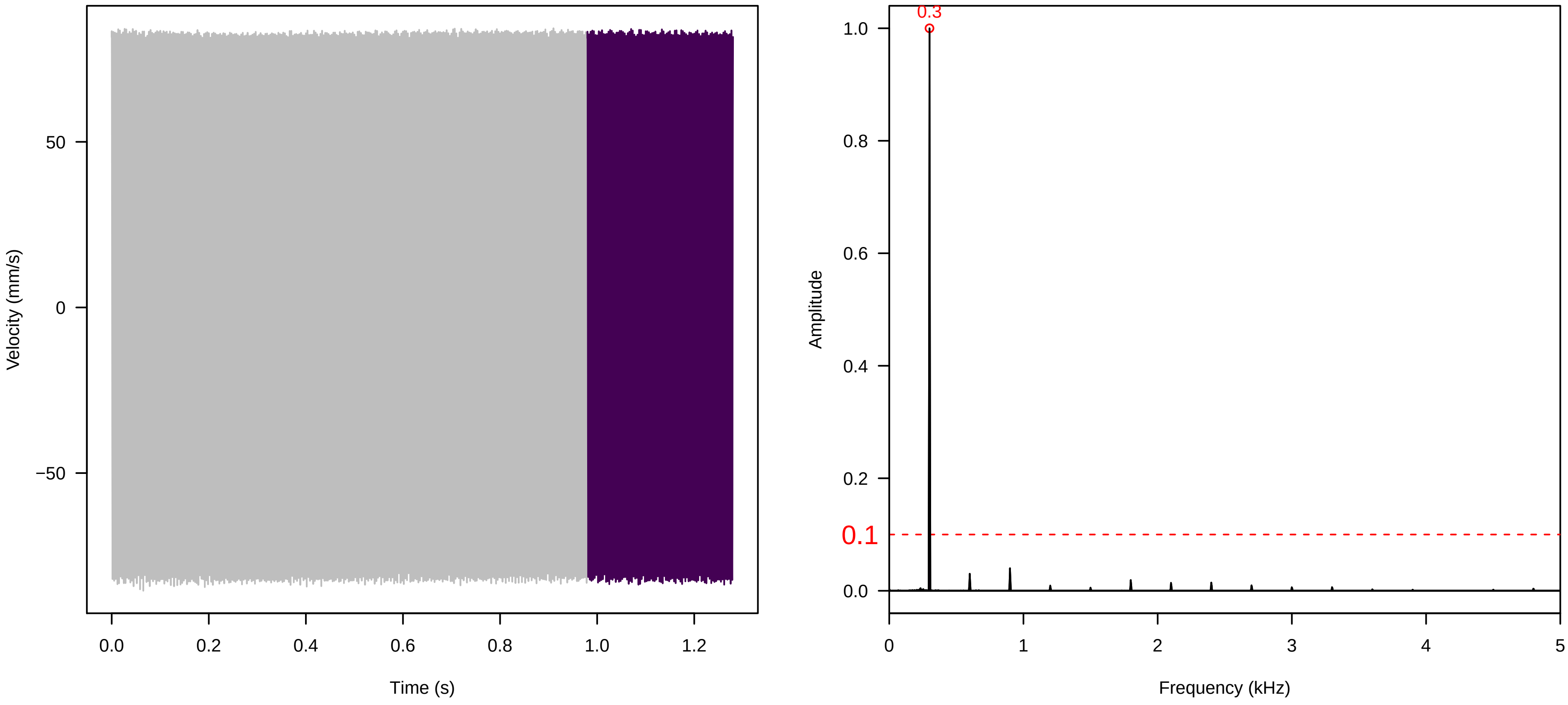


**Vel. = 0.057 ; Str. = Receptacle ; Axis = y ; Fl. accession = 10−s−77−19**

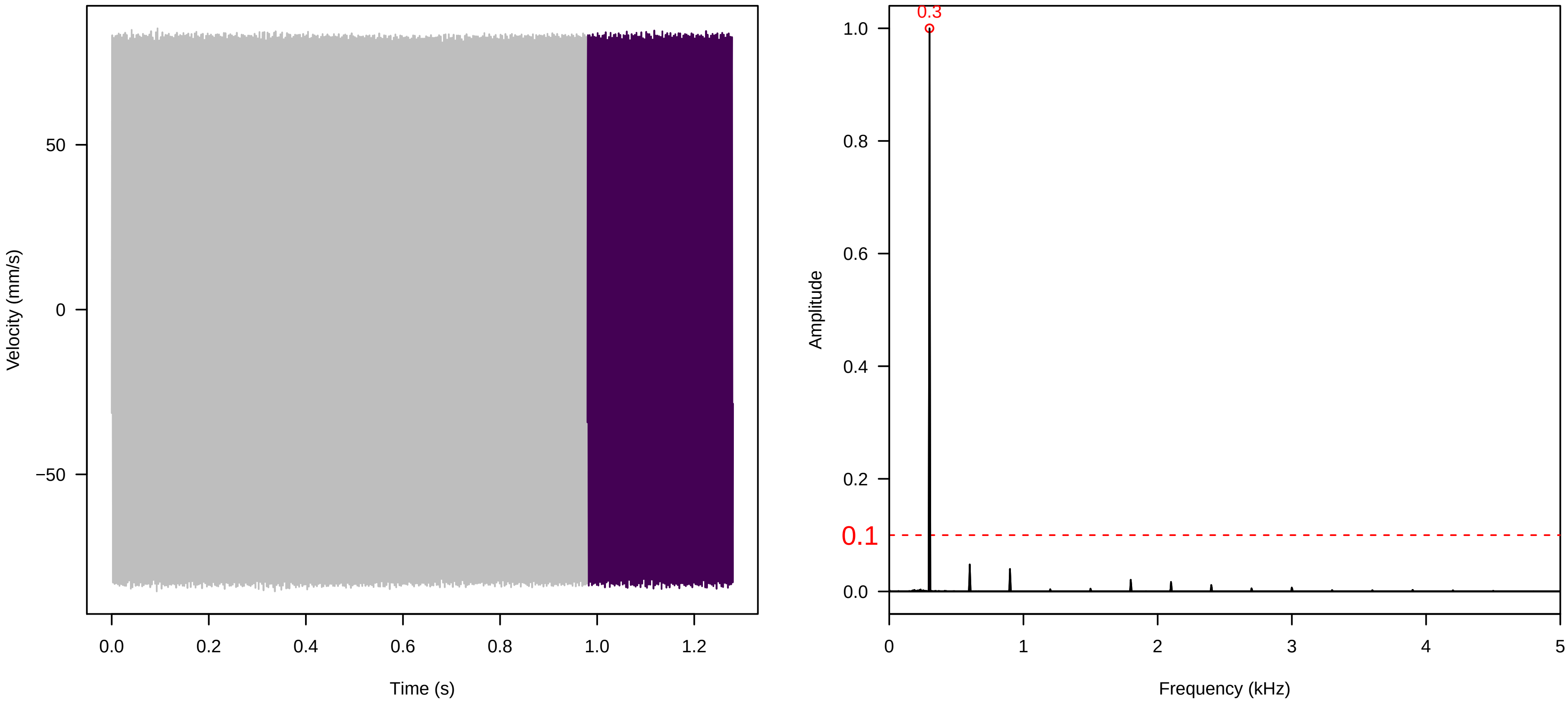

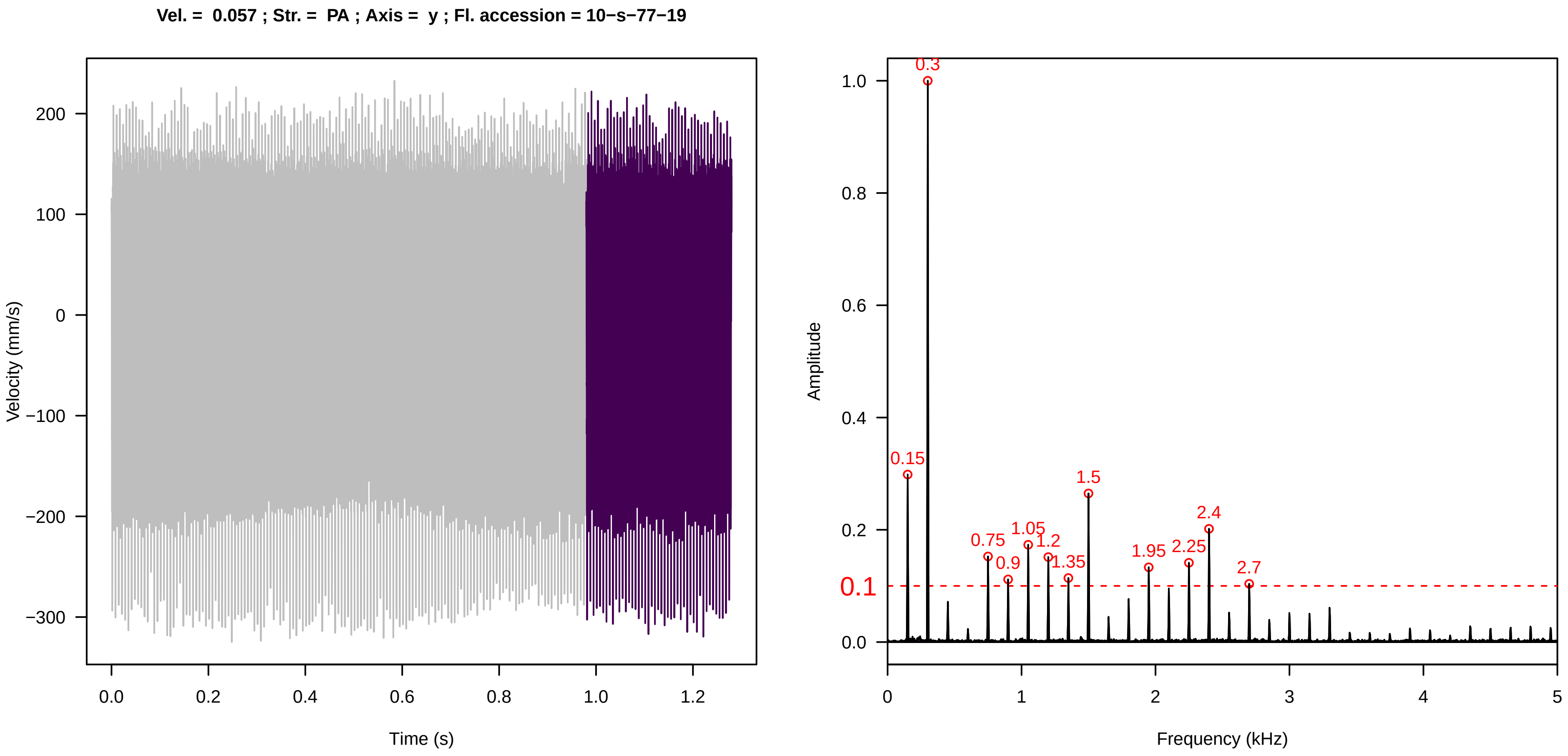
Vel. = 0.057 ; Str. = PA ; Axis = y ; Fl. accession = 10−s−77−19
Velocity (mm/s)
Time (s)
Amplitude
Frequency (kHz)
0.3
0.15
1.5
2.4
0.75
1.05
1.2
0.9
1.35
1.95
2.25
2.7
0.1

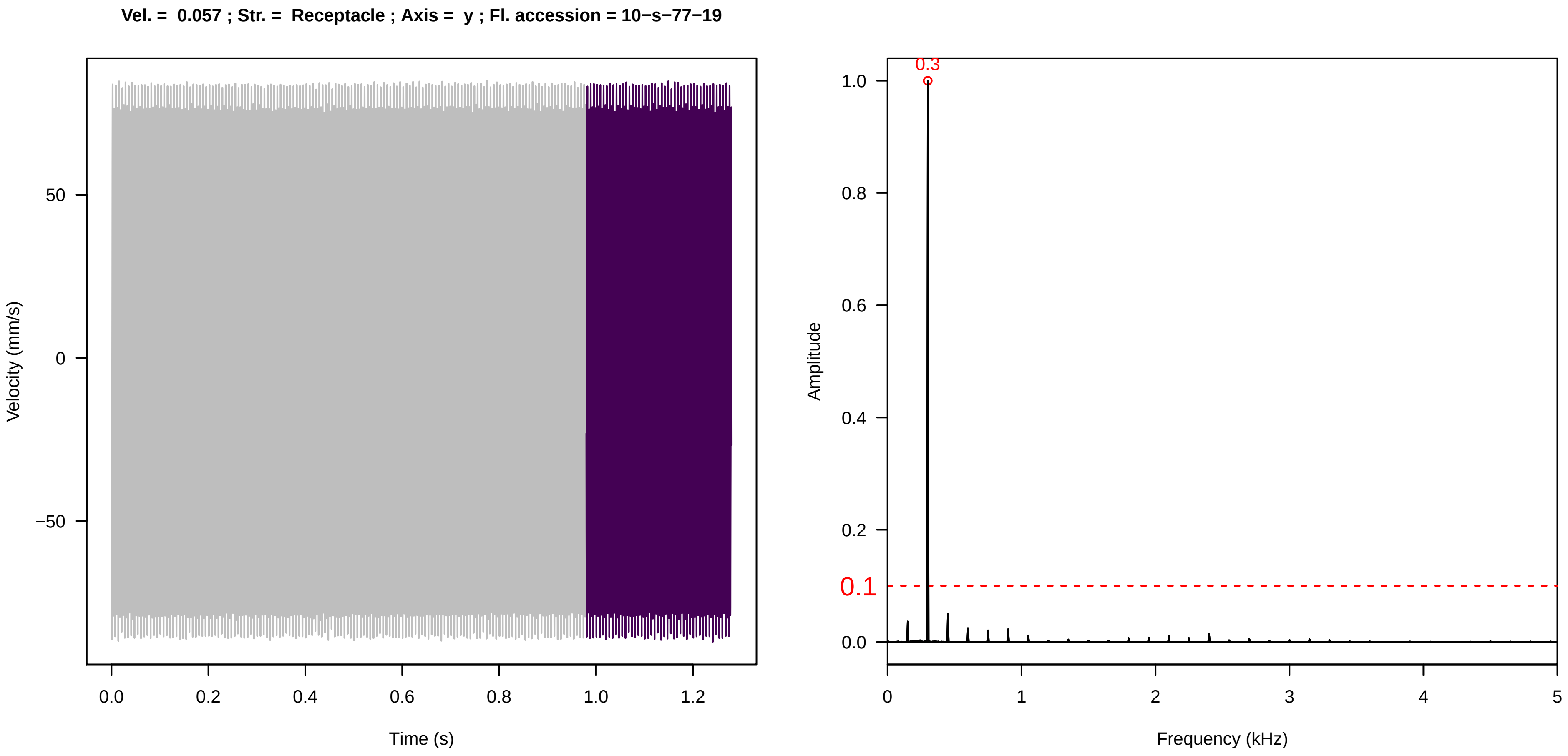
Vel. = 0.057 ; Str. = Receptacle ; Axis = y ; Fl. accession = 10−s−77−19
Velocity (mm/s)
Time (s)
Amplitude
Frequency (kHz)
0.3
0.1

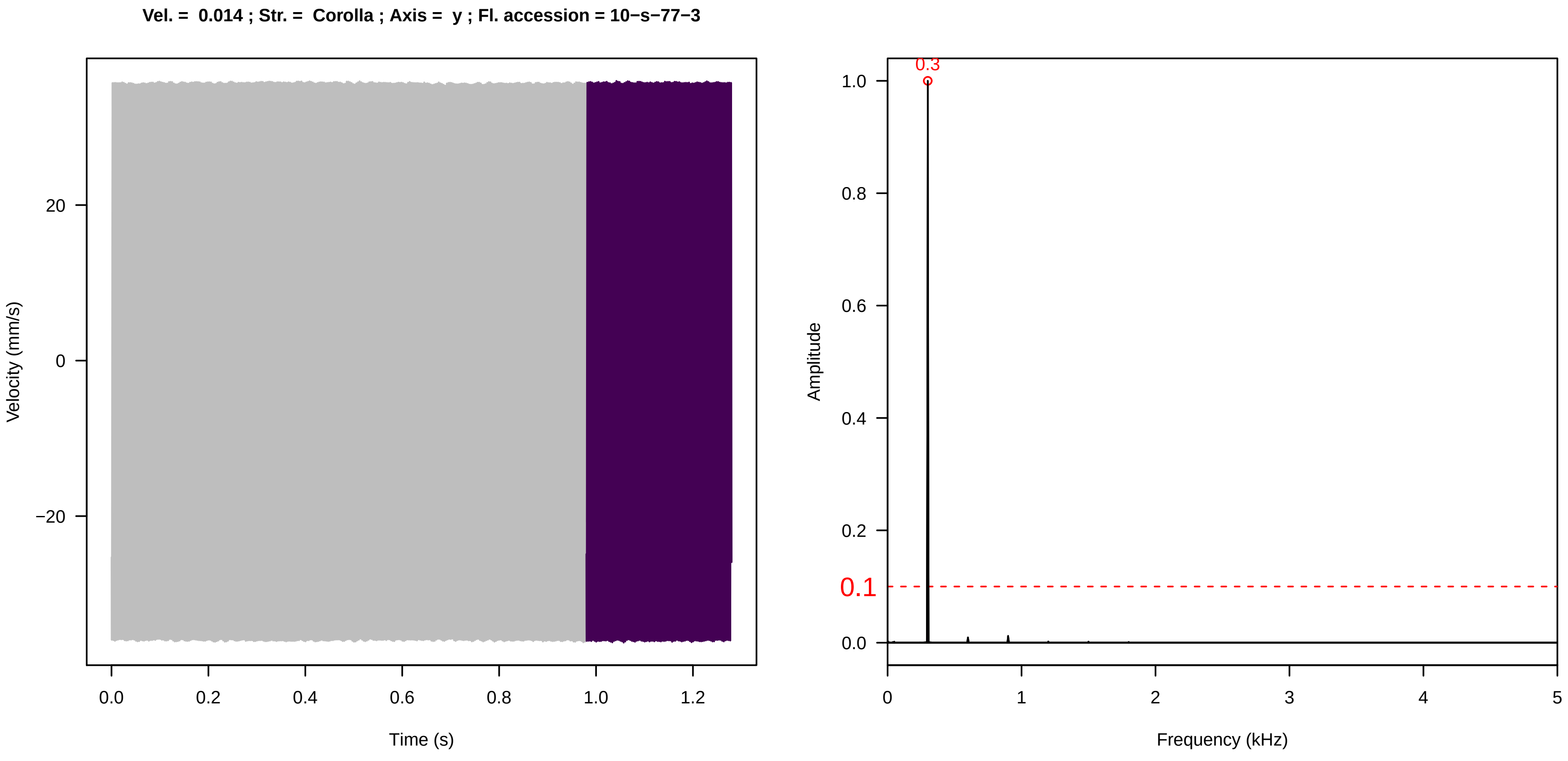
Vel. = 0.014 ; Str. = Corolla ; Axis = y ; Fl. accession = 10−s−77−3
Velocity (mm/s)
Time (s)
Amplitude
Frequency (kHz)
0.3
0.1

**Vel. =  0.014 ; Str. =  Receptacle ; Axis =  y ; Fl. accession = 10−s−77−3**

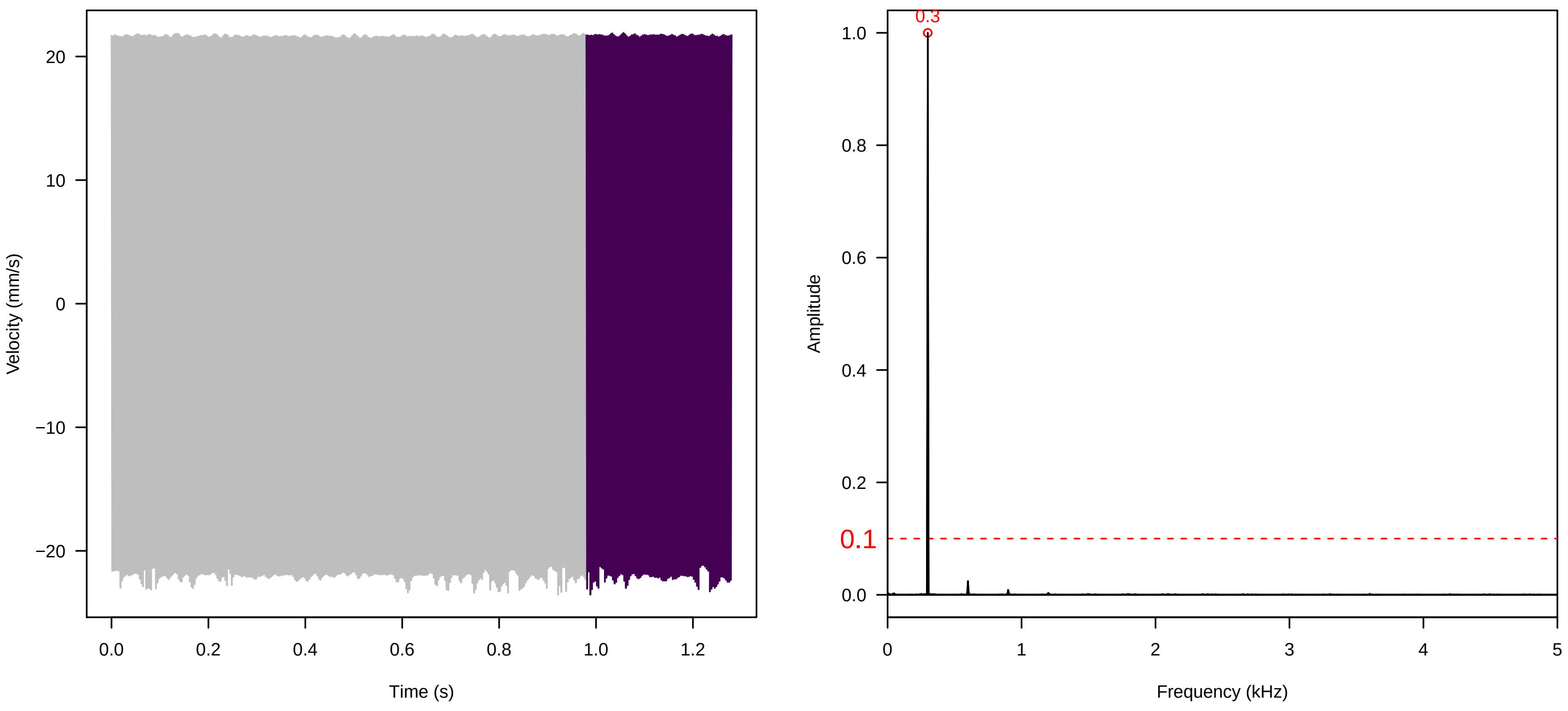


**Vel. =  0.014 ; Str. =  FA ; Axis =  y ; Fl. accession = 10−s−77−3**

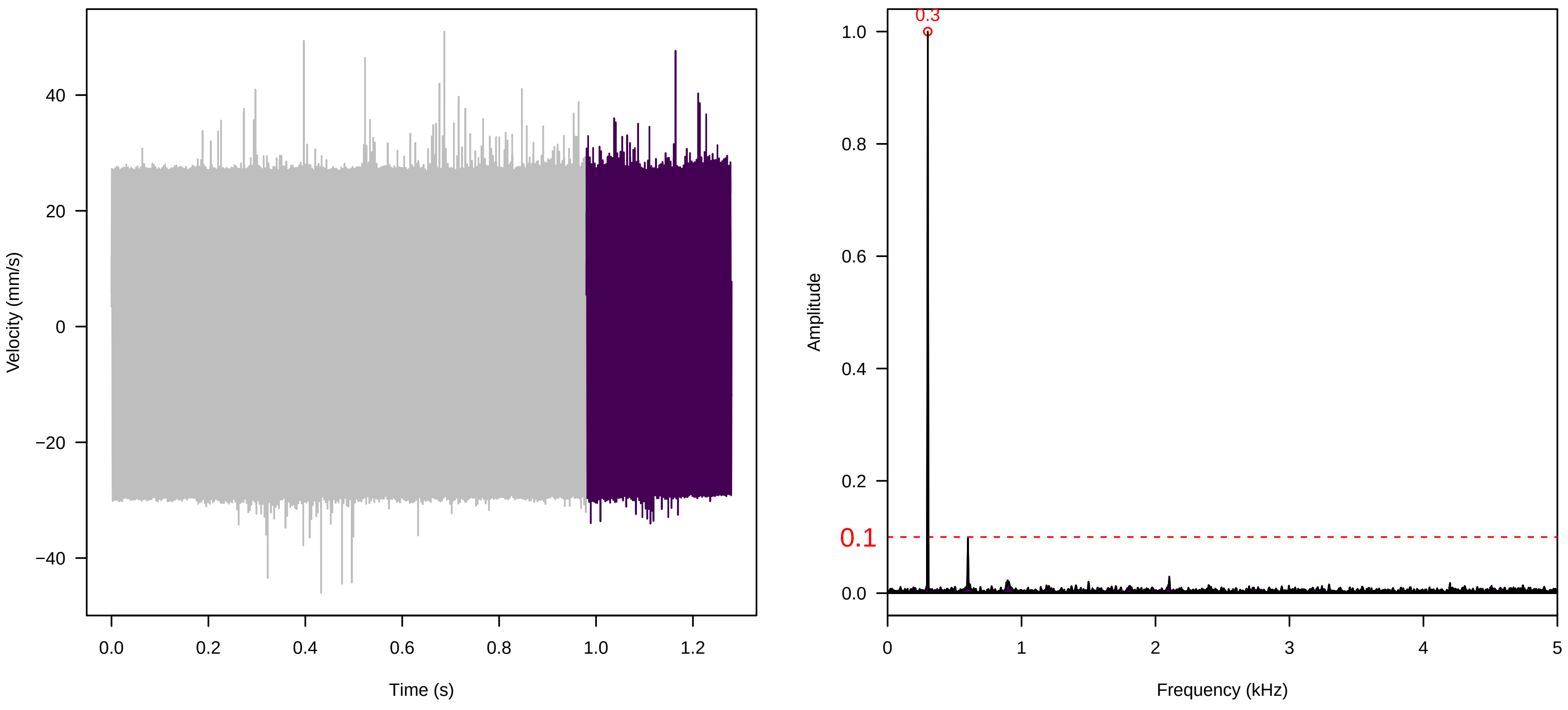


**Vel. =  0.014 ; Str. =  Receptacle ; Axis =  y ; Fl. accession = 10−s−77−3**

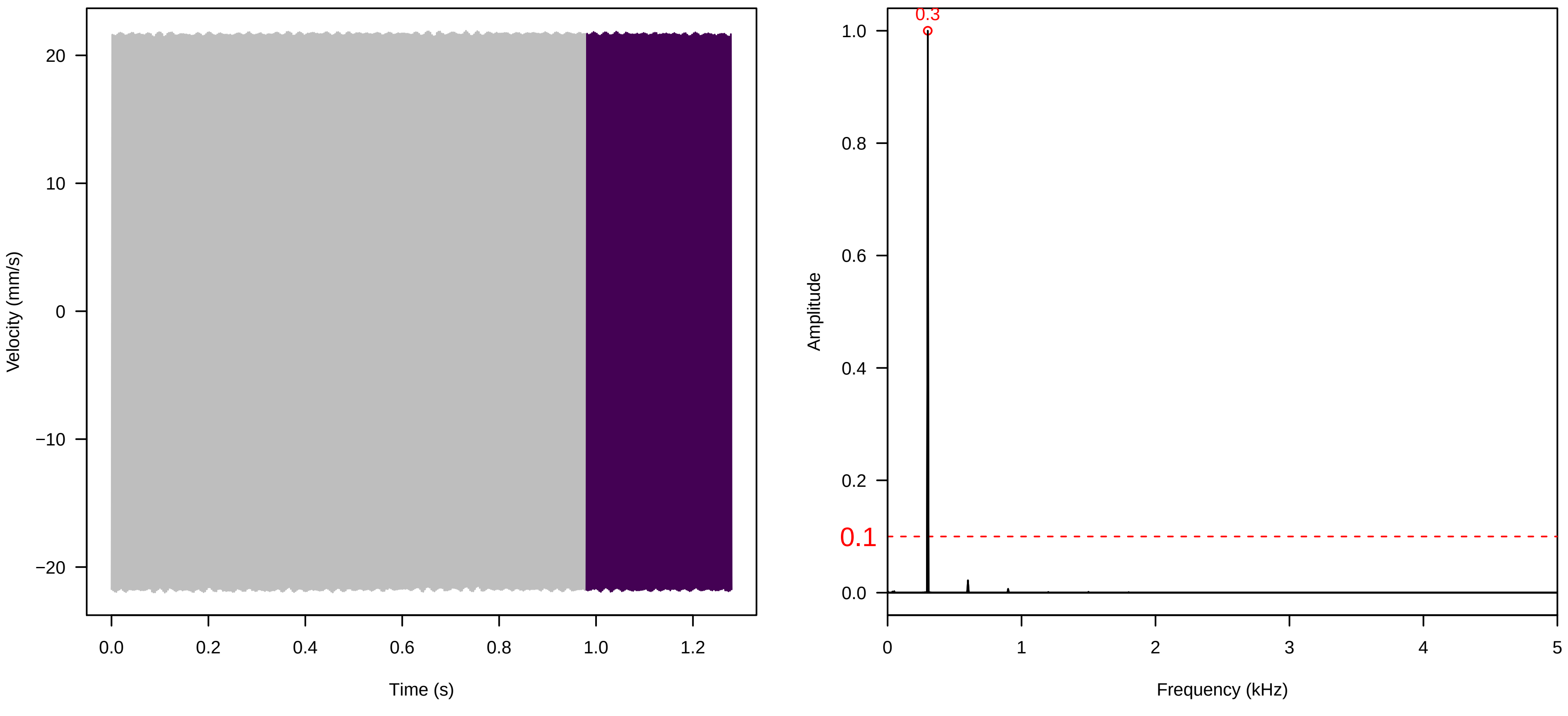

**Vel. = 0.014 ; Str. = PA ; Axis = y ; Fl. accession = 10−s−77−3**

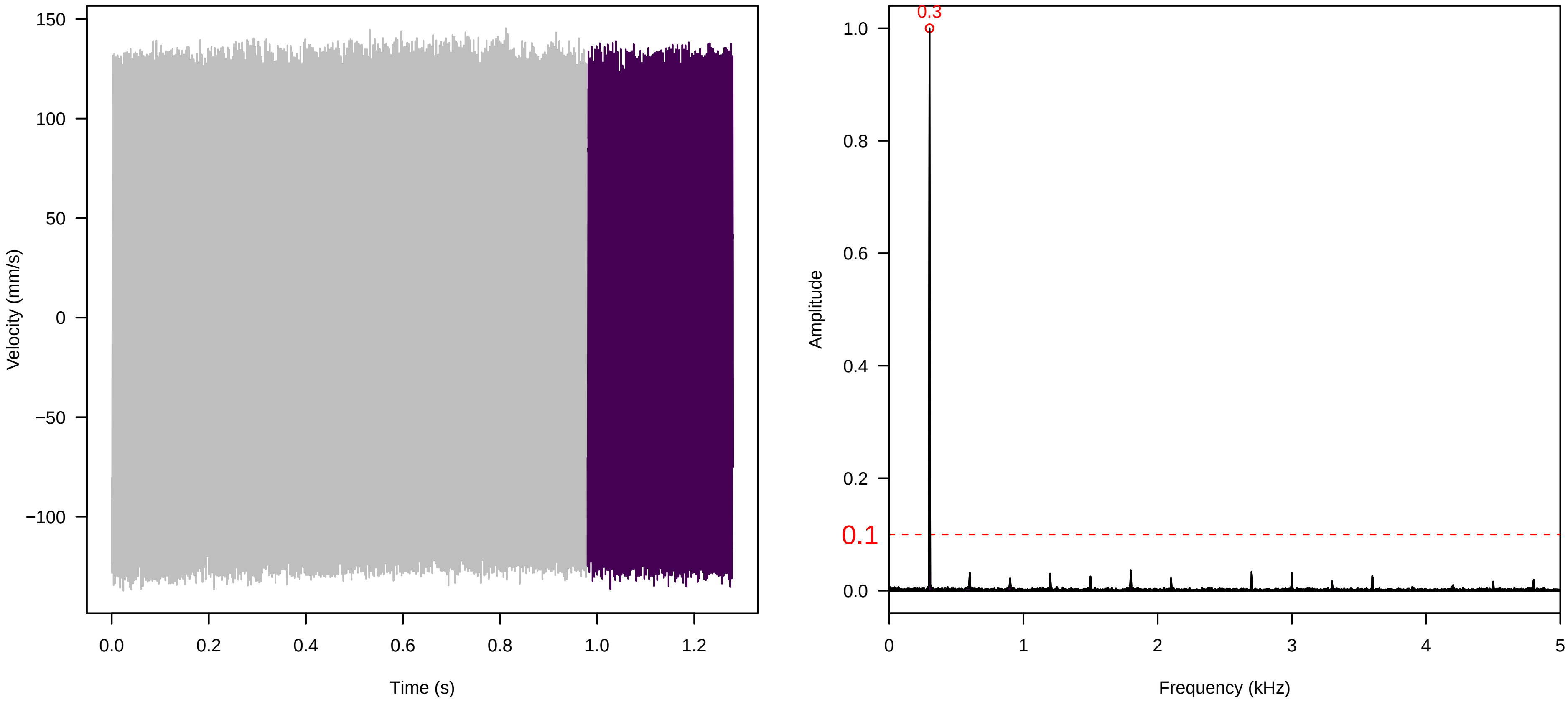


**Vel. = 0.014 ; Str. = Receptacle ; Axis = y ; Fl. accession = 10−s−77−3**

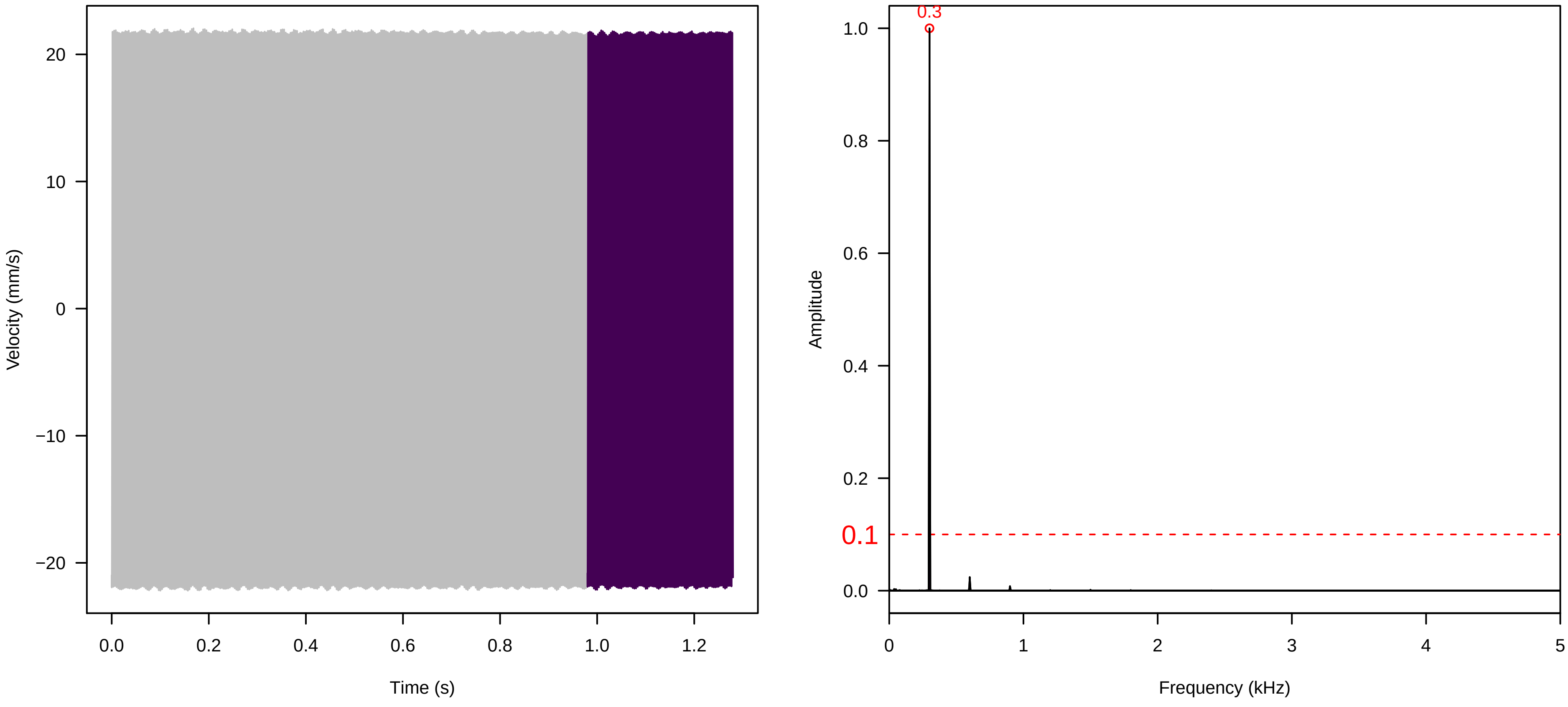


**Vel. = 0.028 ; Str. = PA ; Axis = y ; Fl. accession = 10−s−77−3**

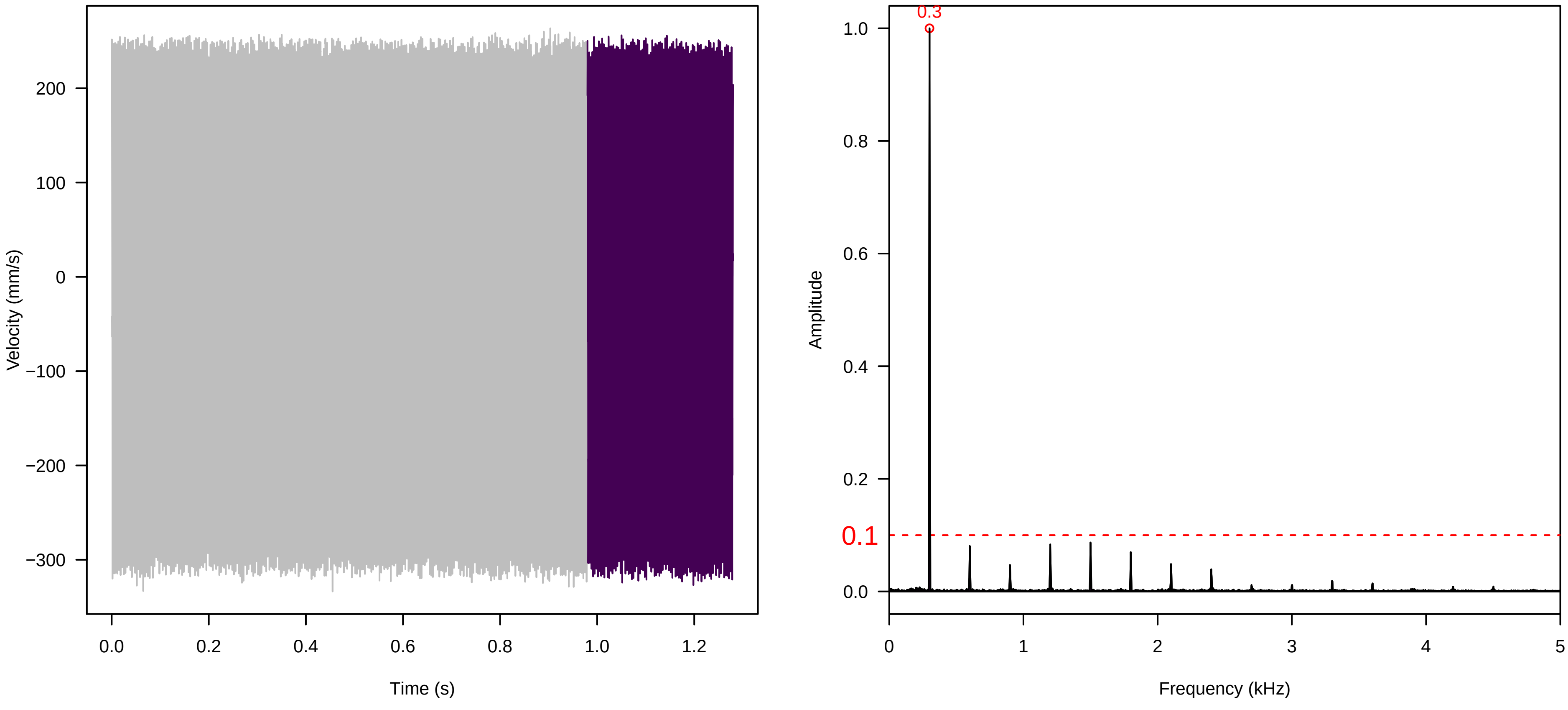

**Vel. = 0.028 ; Str. = Receptacle ; Axis = y ; Fl. accession = 10−s−77−3**

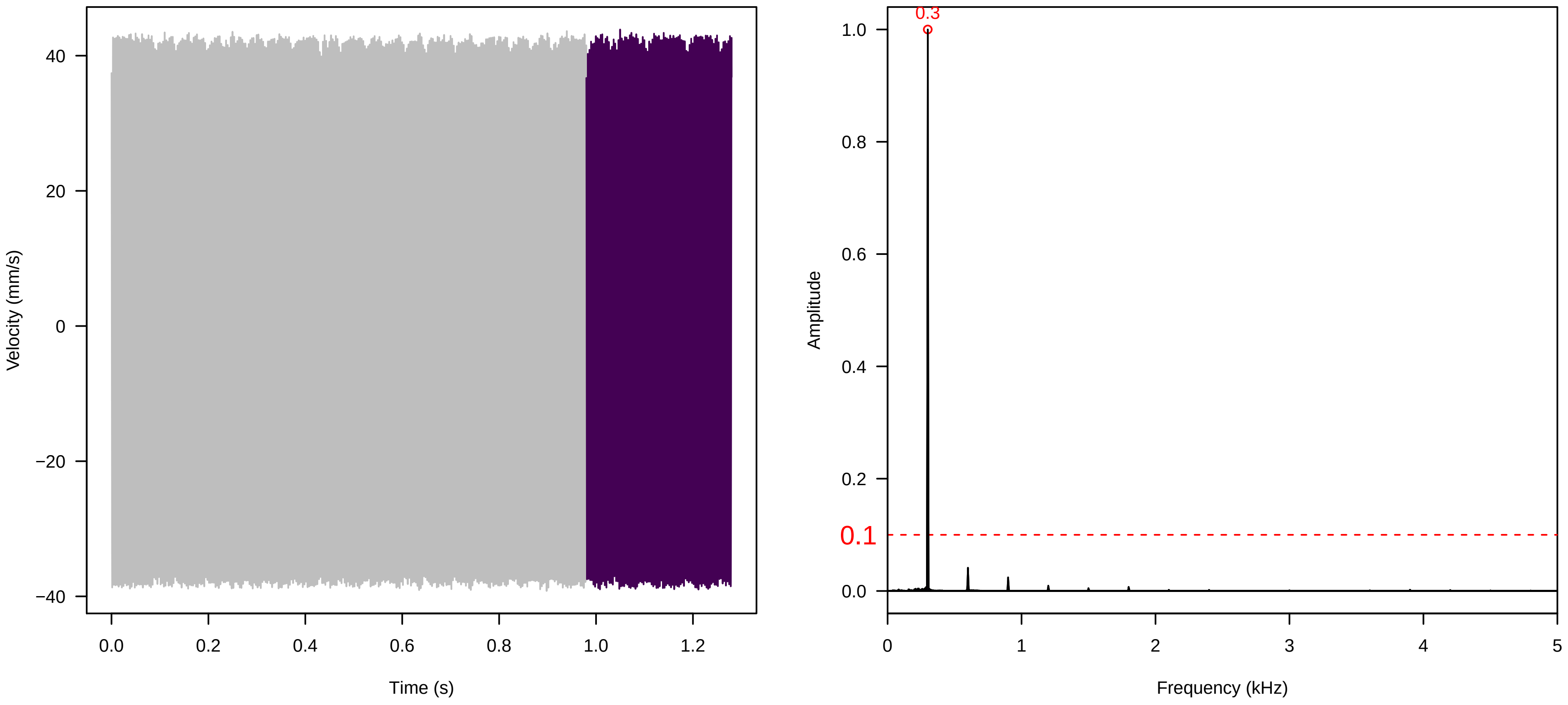


**Vel. = 0.028 ; Str. = FA ; Axis = y ; Fl. accession = 10−s−77−3**

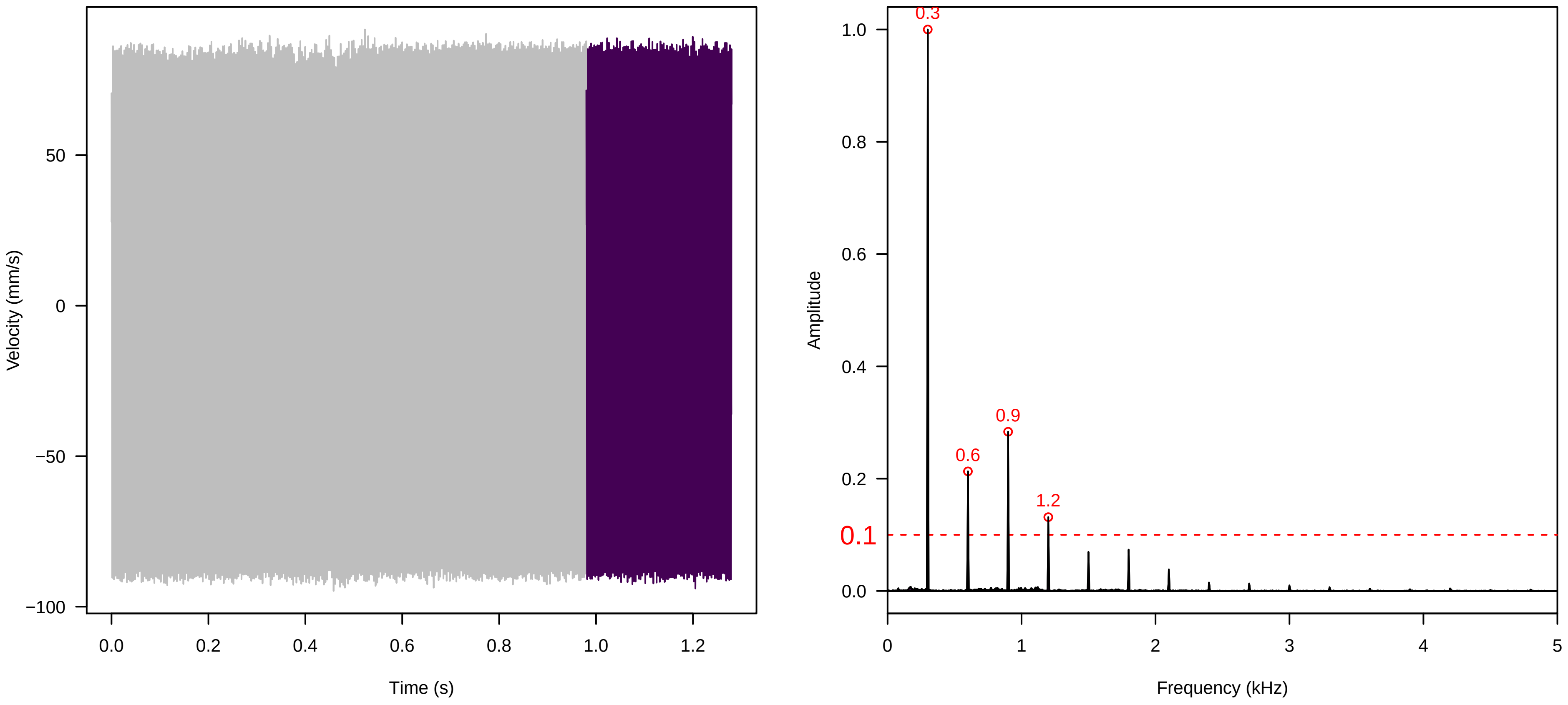


**Vel. = 0.028 ; Str. = Receptacle ; Axis = y ; Fl. accession = 10−s−77−3**

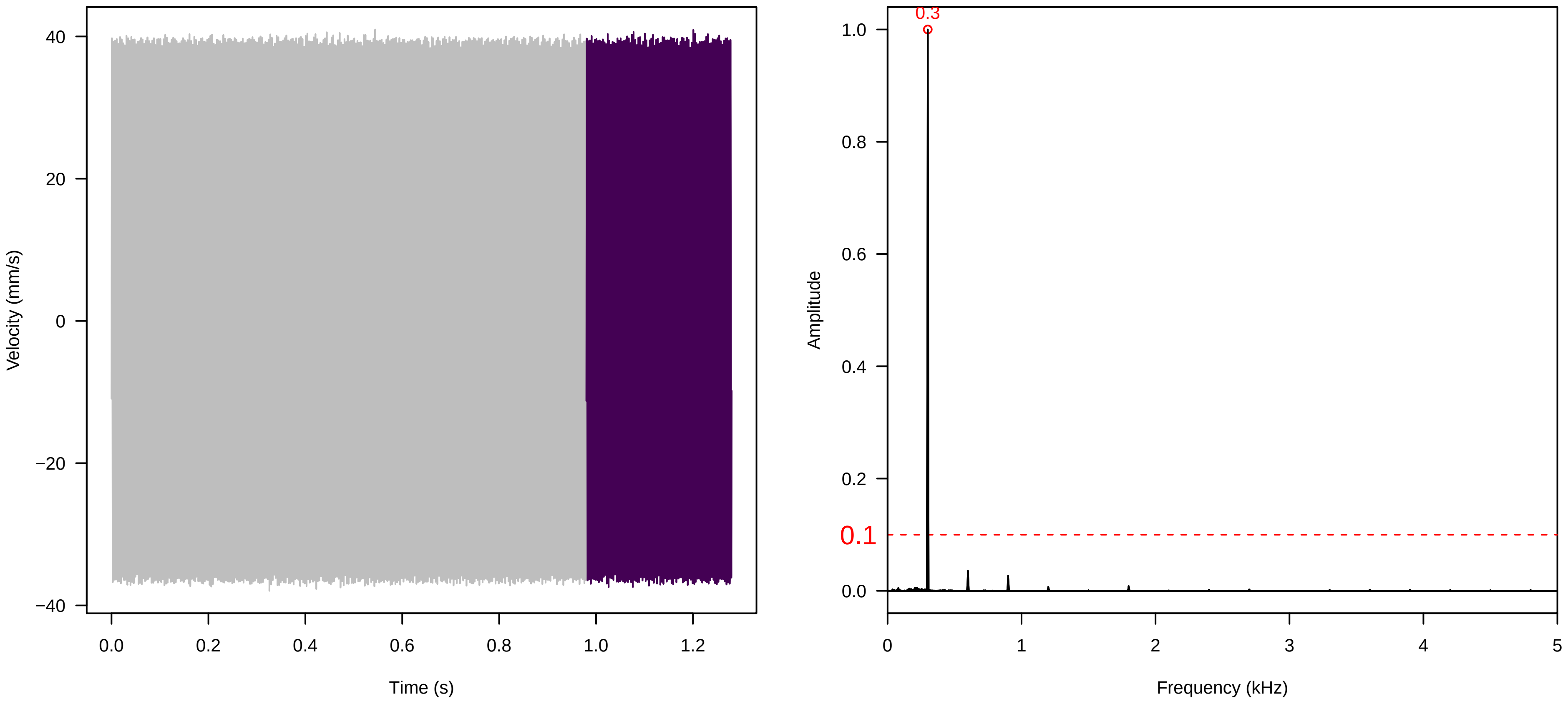

**Vel. = 0.028 ; Str. = Corolla ; Axis = y ; Fl. accession = 10−s−77−3**

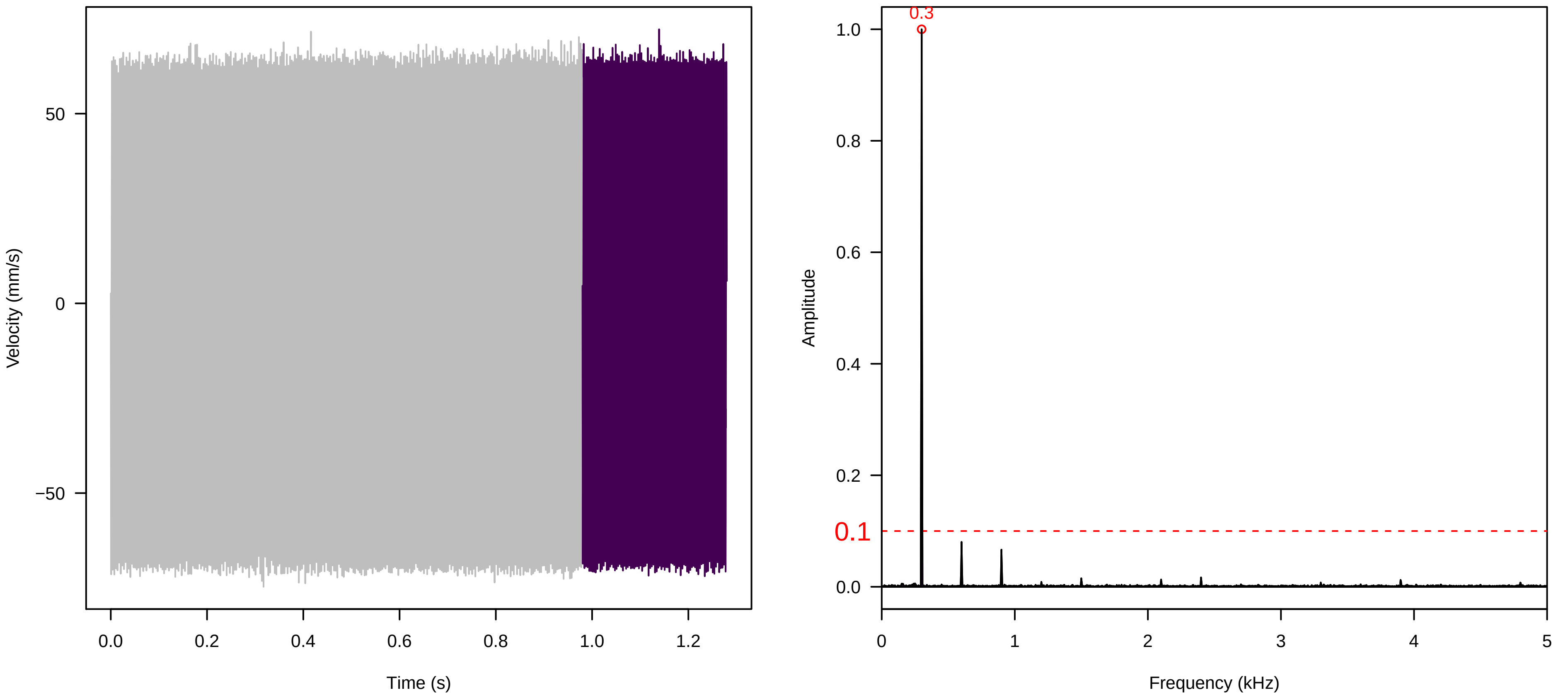


**Vel. = 0.028 ; Str. = Receptacle ; Axis = y ; Fl. accession = 10−s−77−3**

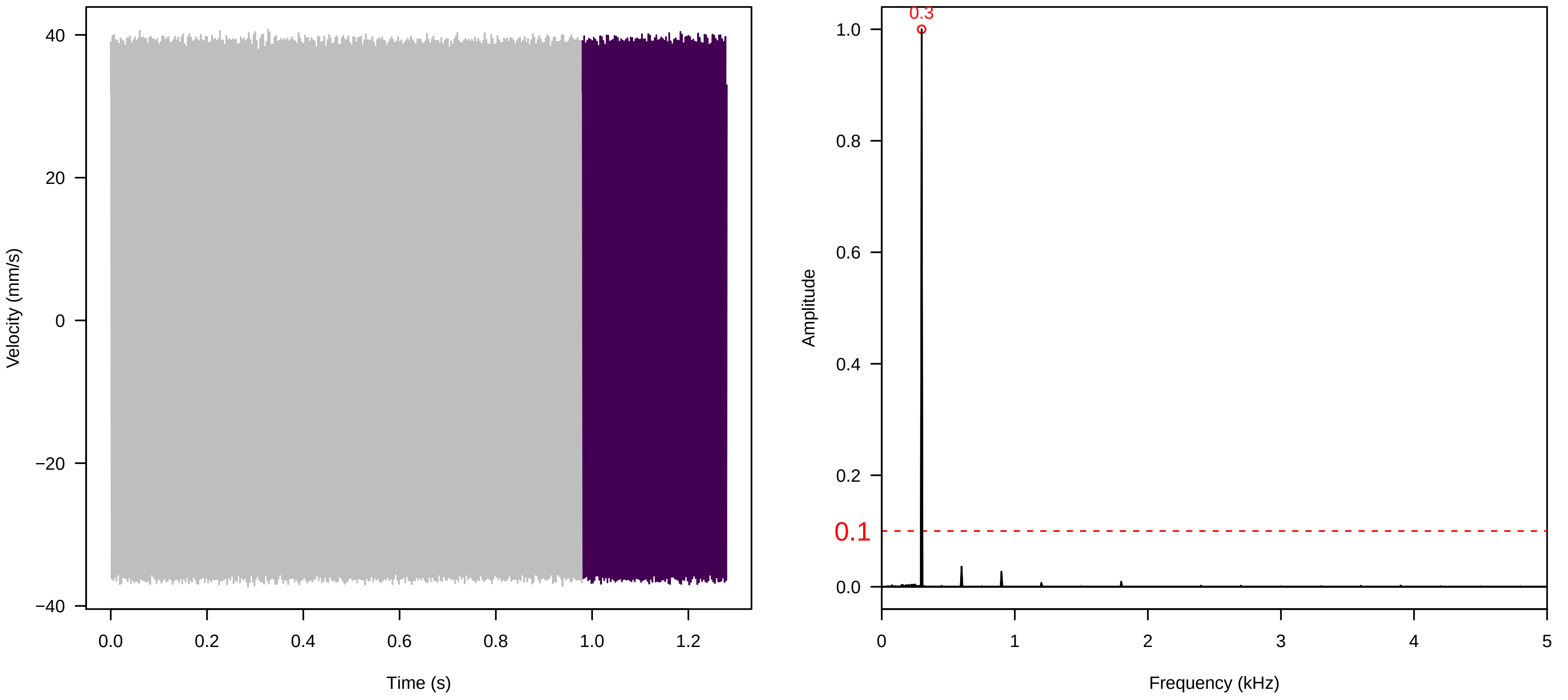


**Vel. = 0.057 ; Str. = Corolla ; Axis = y ; Fl. accession = 10−s−77−3**

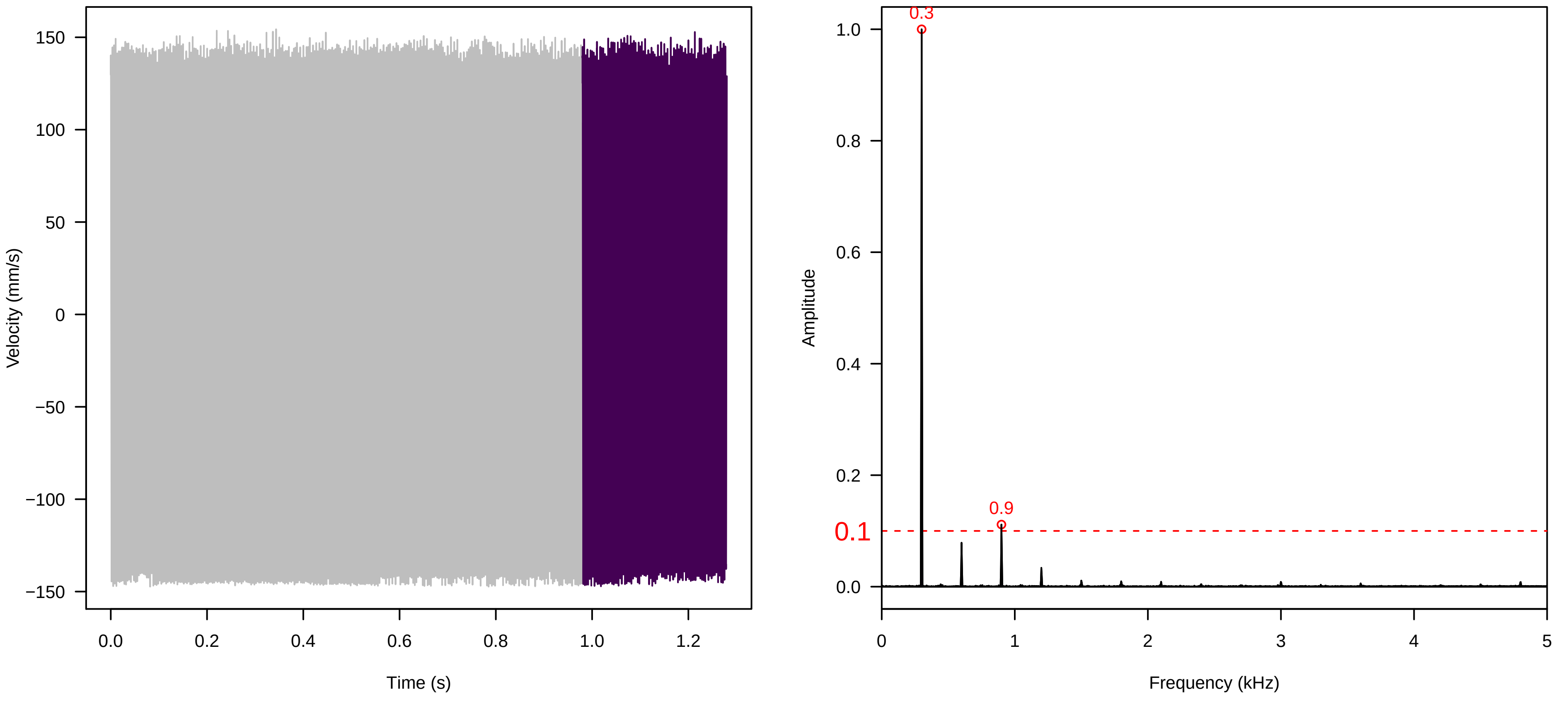

Vel. =  0.057 ; Str. =  Receptacle ; Axis =  y ; Fl. accession = 10−s−77−3

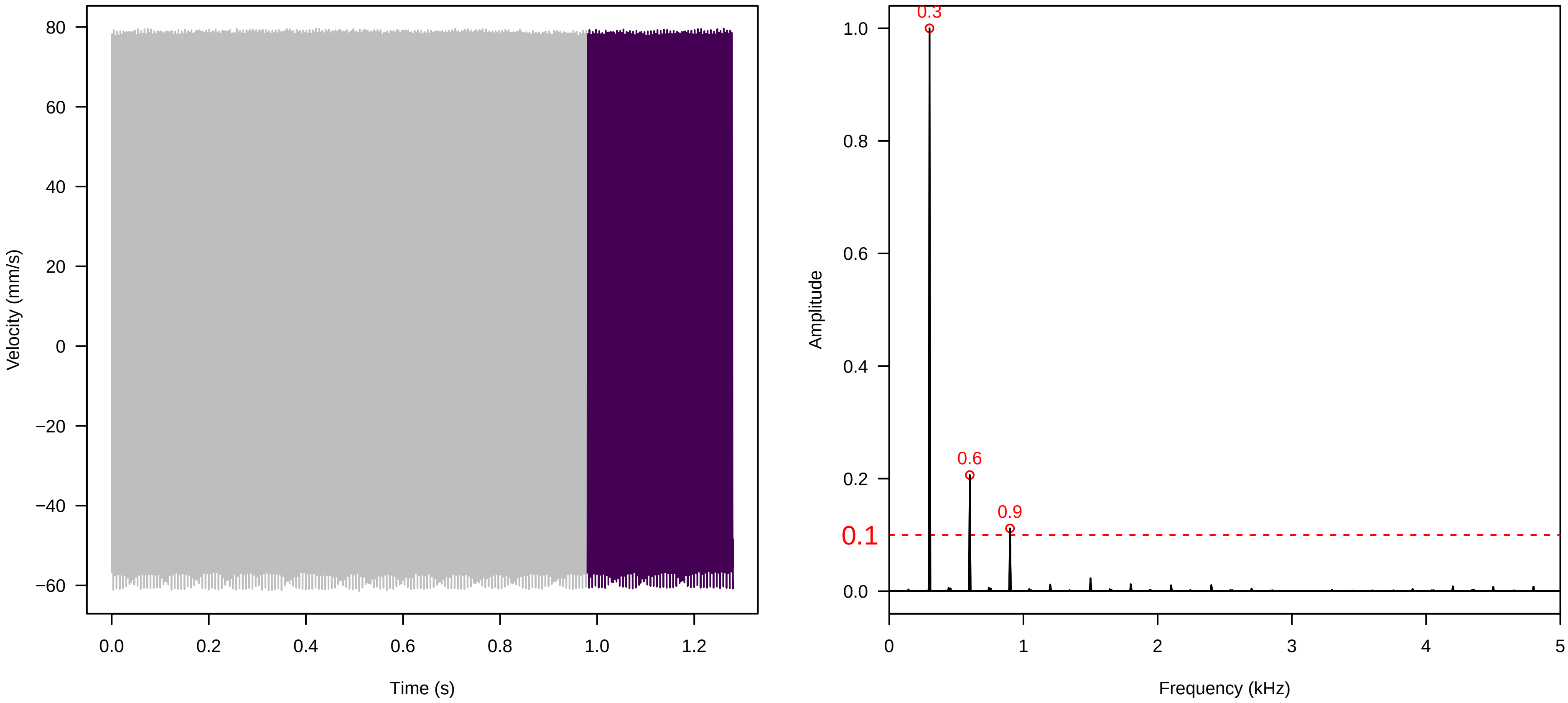


Vel. =  0.057 ; Str. =  FA ; Axis =  y ; Fl. accession = 10−s−77−3

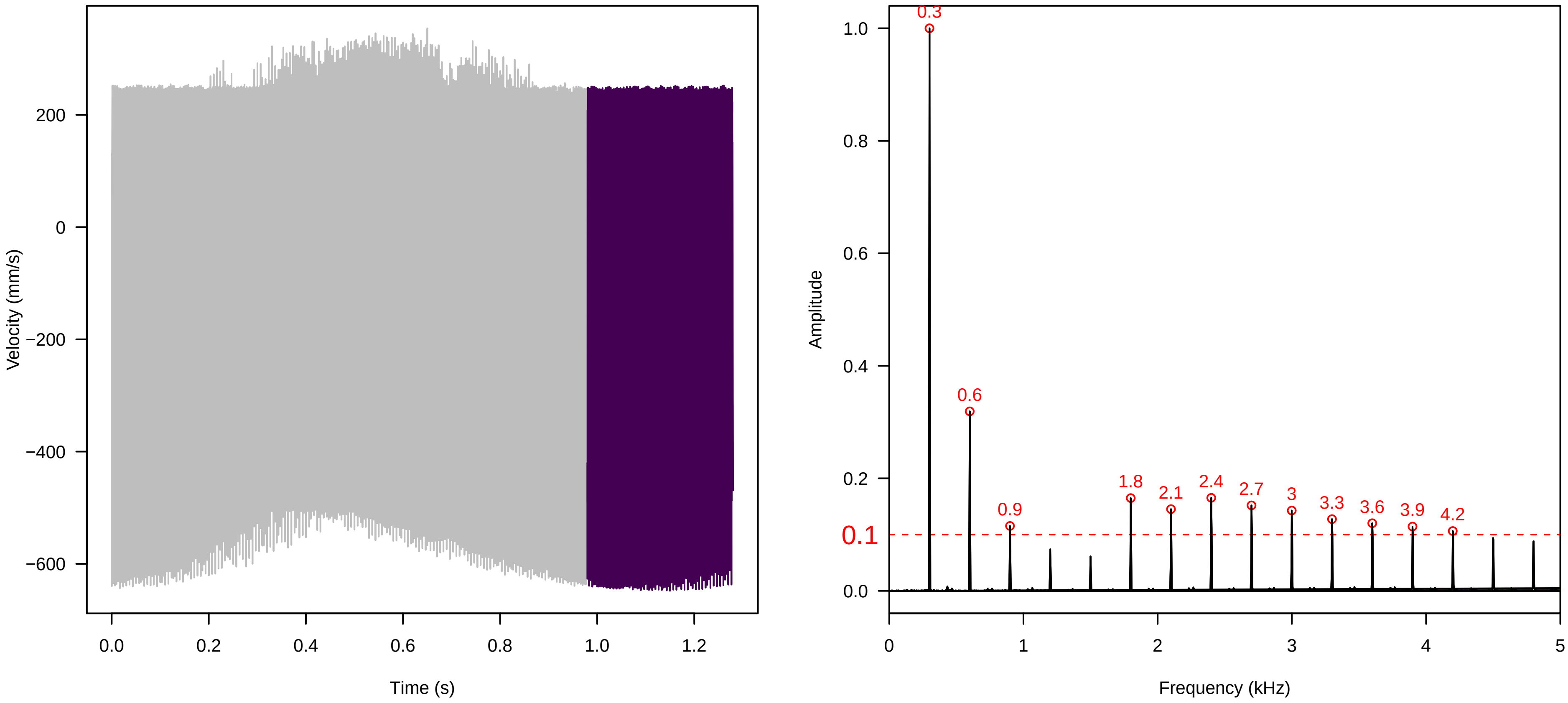


Vel. =  0.057 ; Str. =  Receptacle ; Axis =  y ; Fl. accession = 10−s−77−3

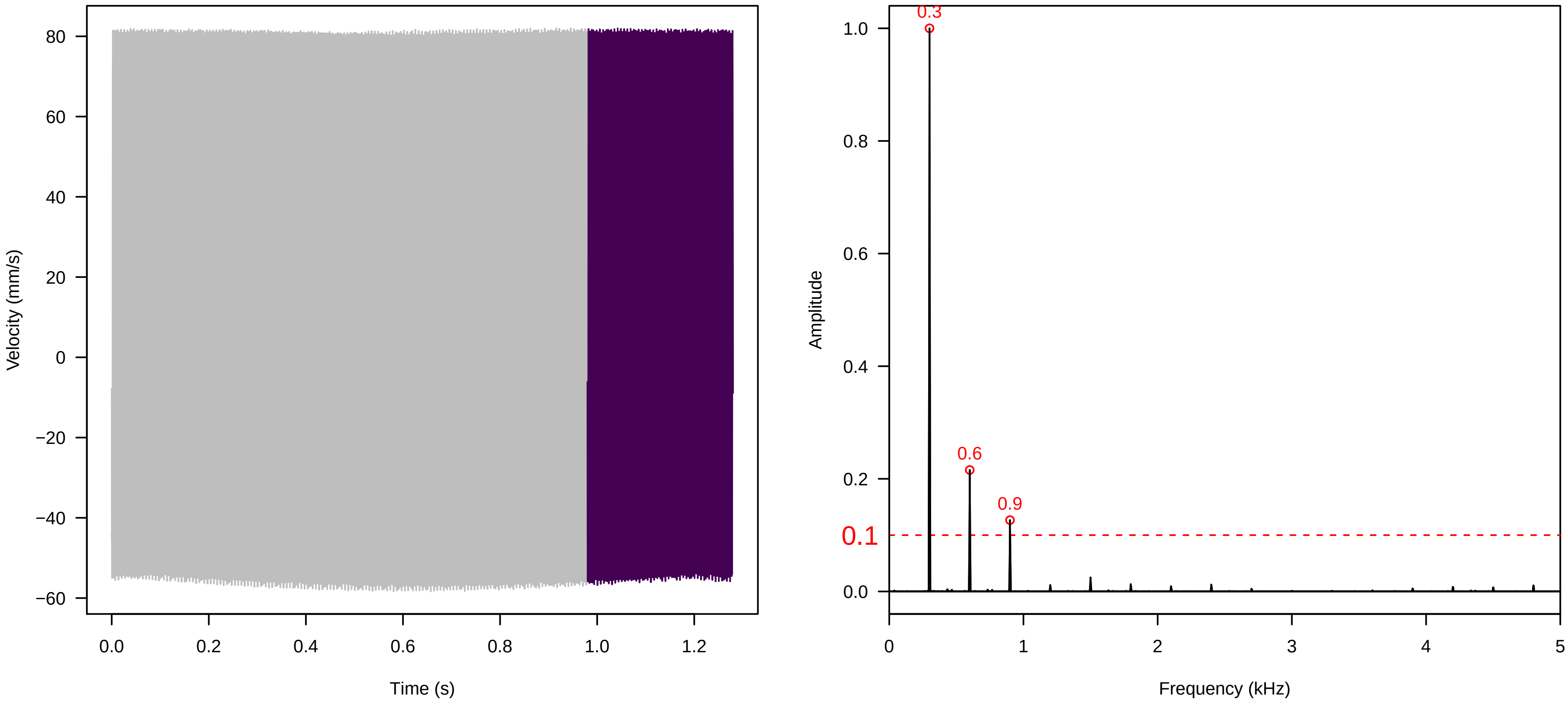

**Vel. = 0.057 ; Str. = PA ; Axis = y ; Fl. accession = 10−s−77−3**

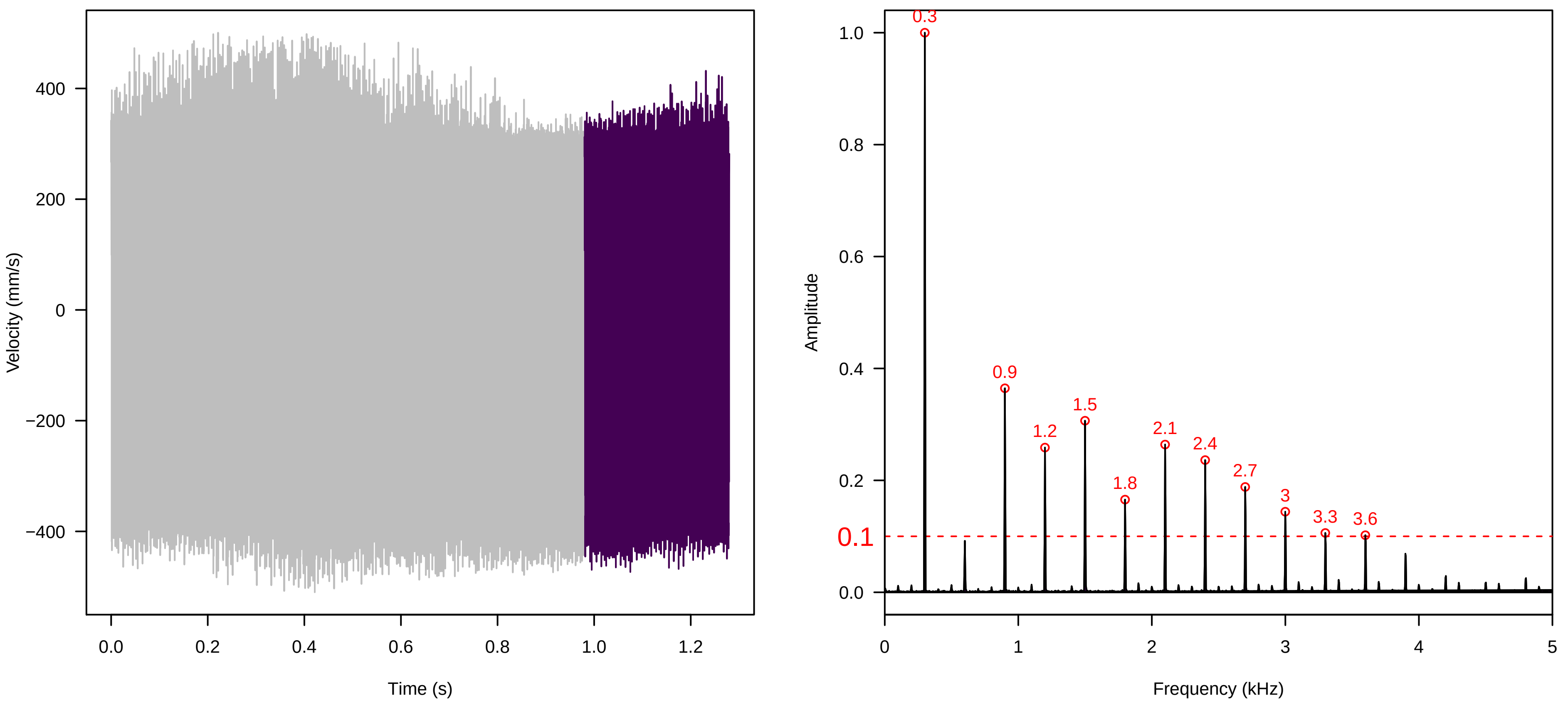


**Vel. = 0.057 ; Str. = Receptacle ; Axis = y ; Fl. accession = 10−s−77−3**

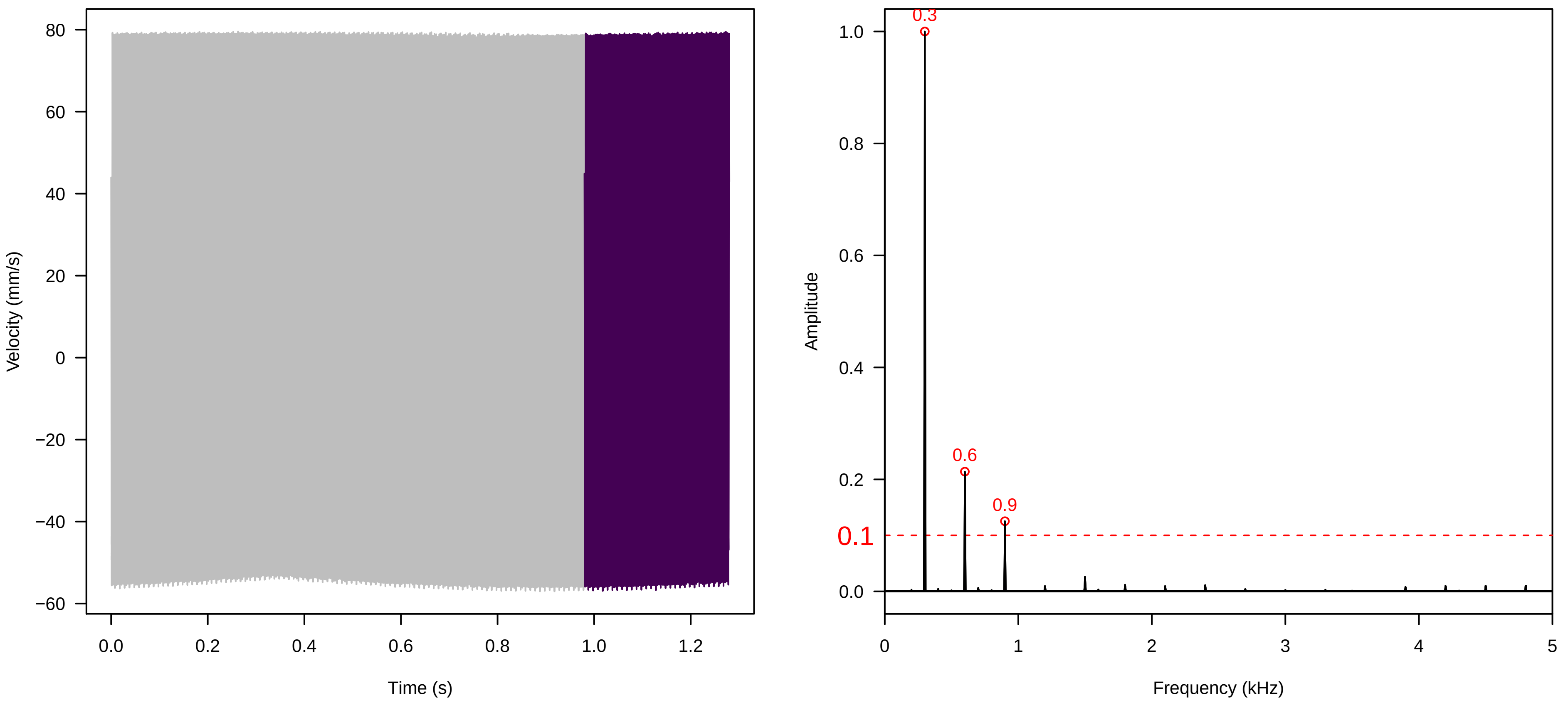


**Vel. = 0.057 ; Str. = Corolla ; Axis = z ; Fl. accession = 10−s−77−3**

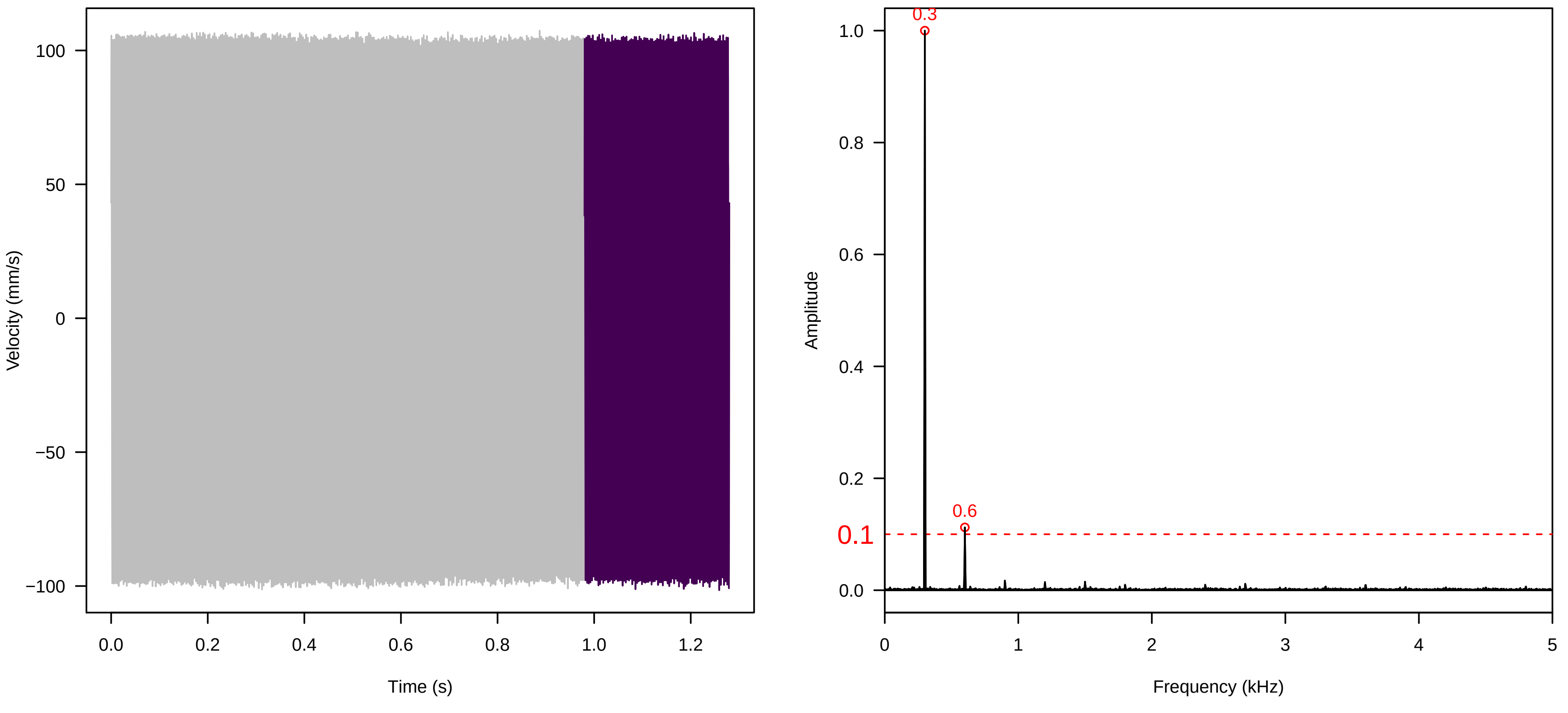

**Vel. = 0.057 ; Str. = Receptacle ; Axis = z ; Fl. accession = 10−s−77−3**

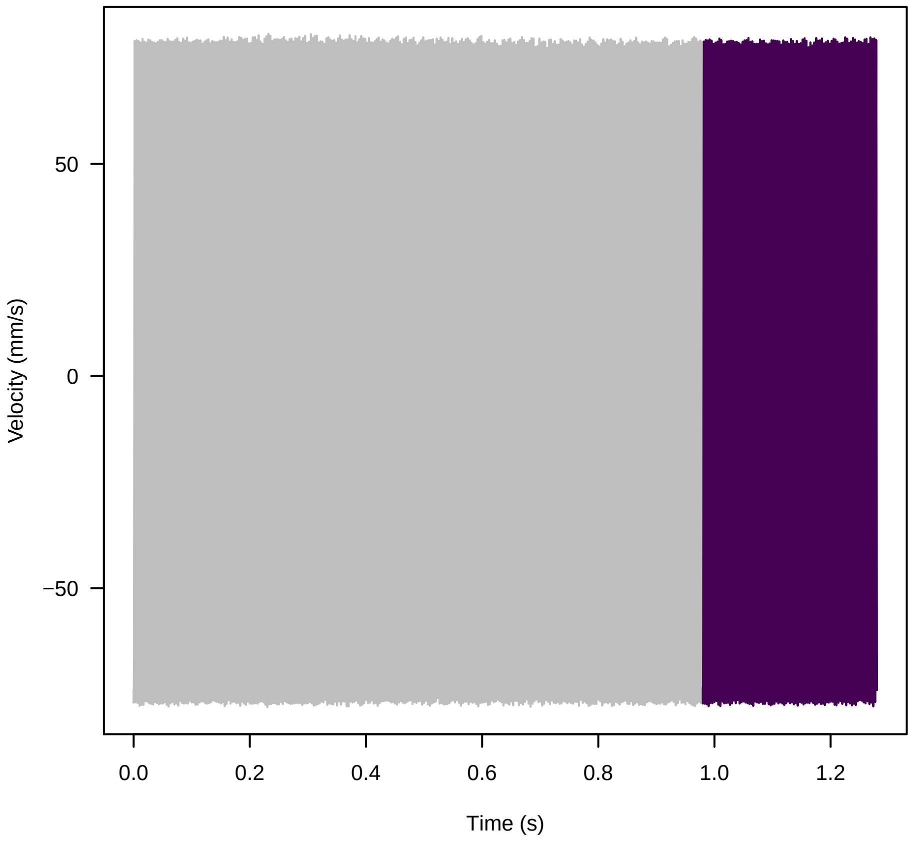


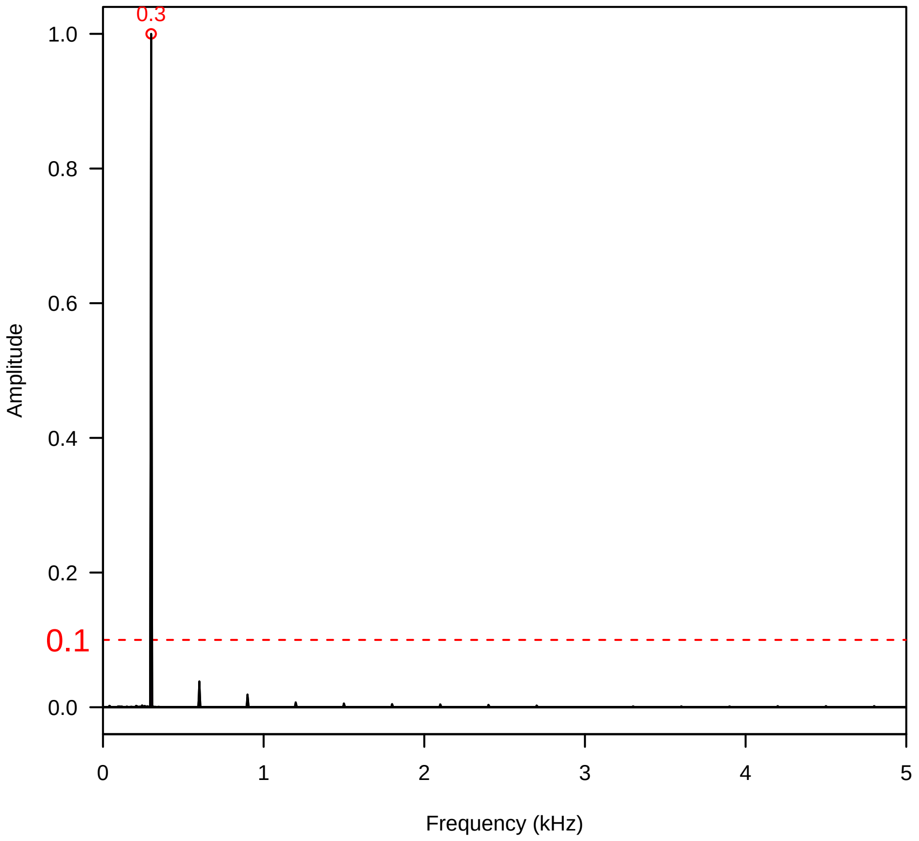


**Vel. = 0.057 ; Str. = FA ; Axis = z ; Fl. accession = 10−s−77−3**

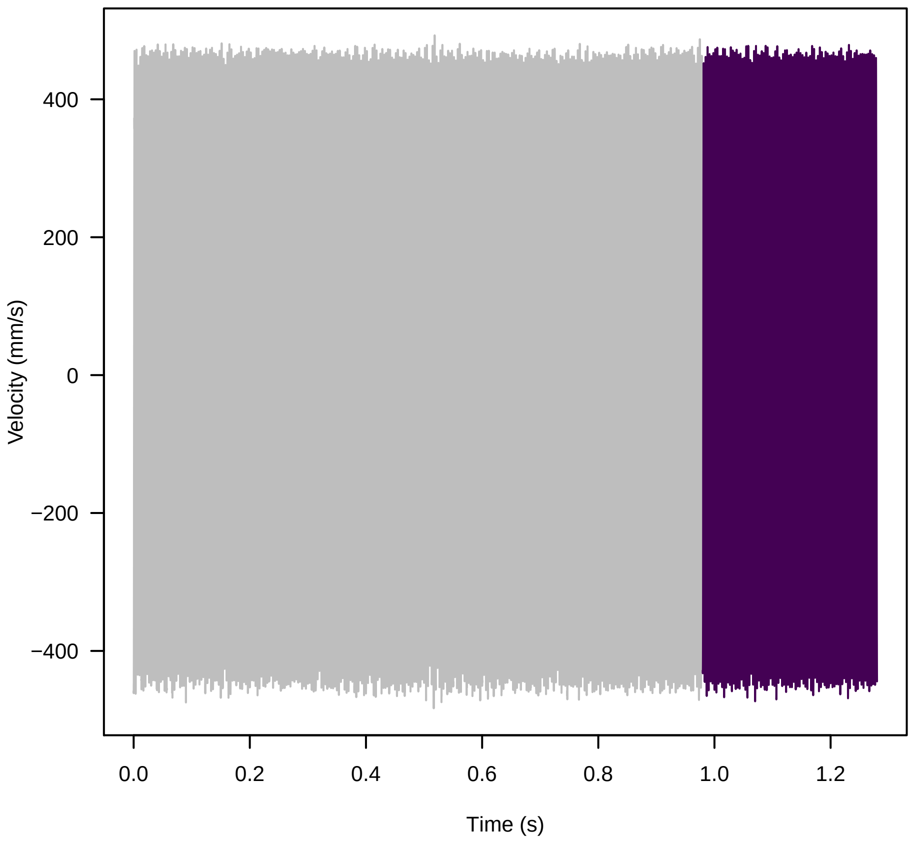


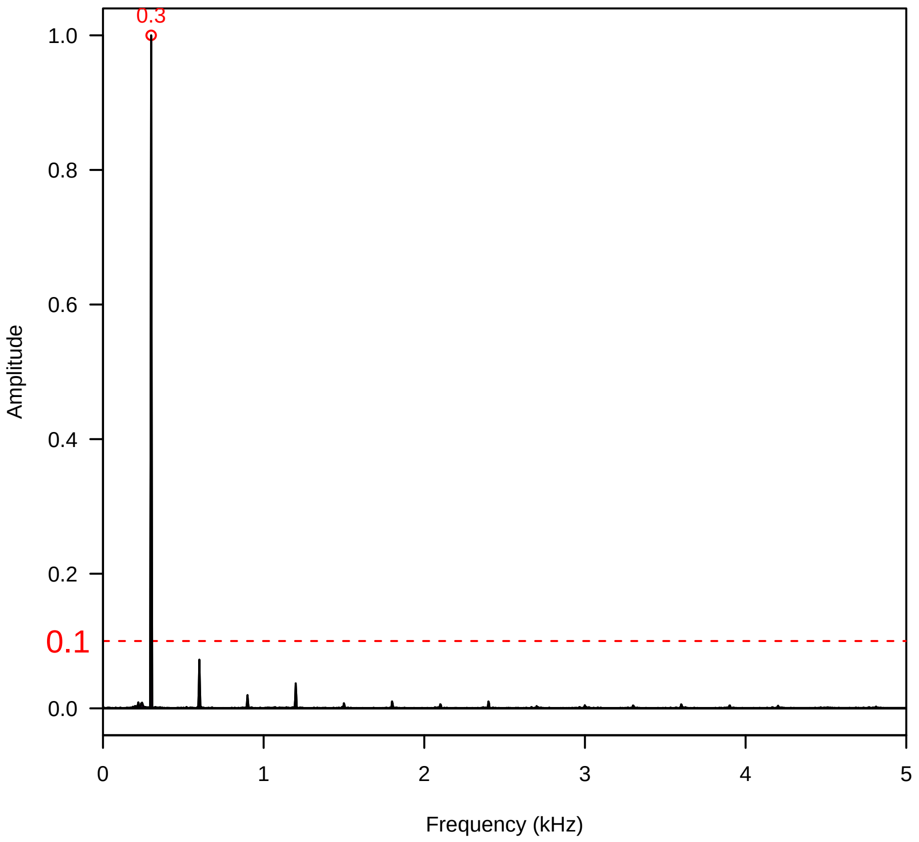


**Vel. = 0.057 ; Str. = Receptacle ; Axis = z ; Fl. accession = 10−s−77−3**

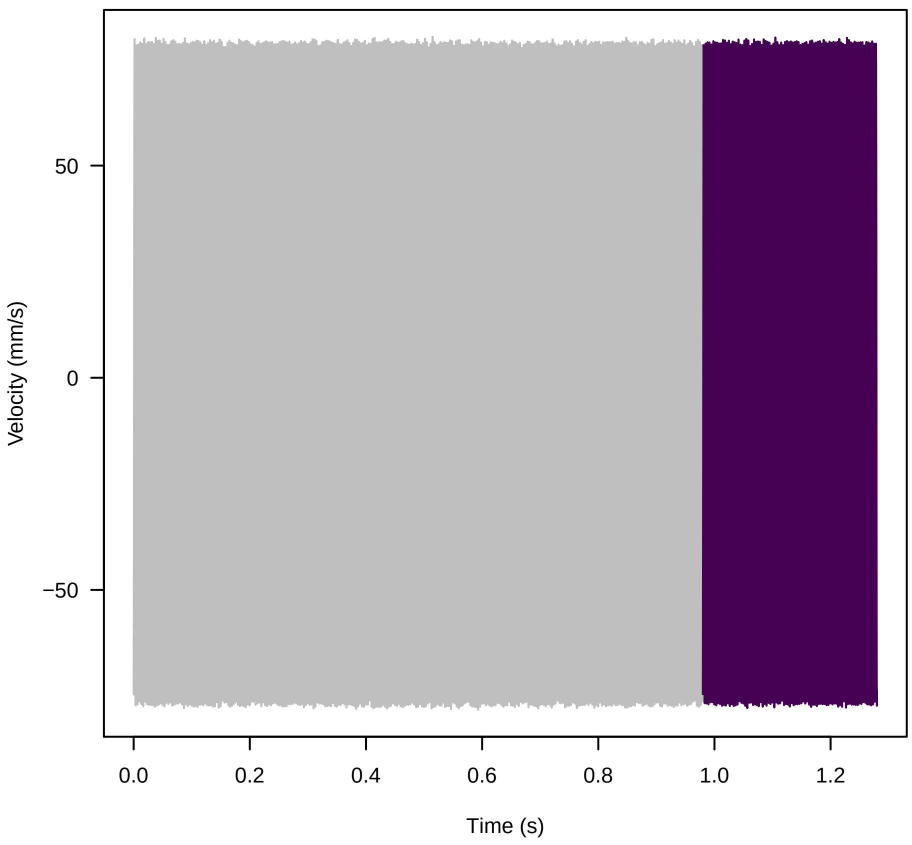


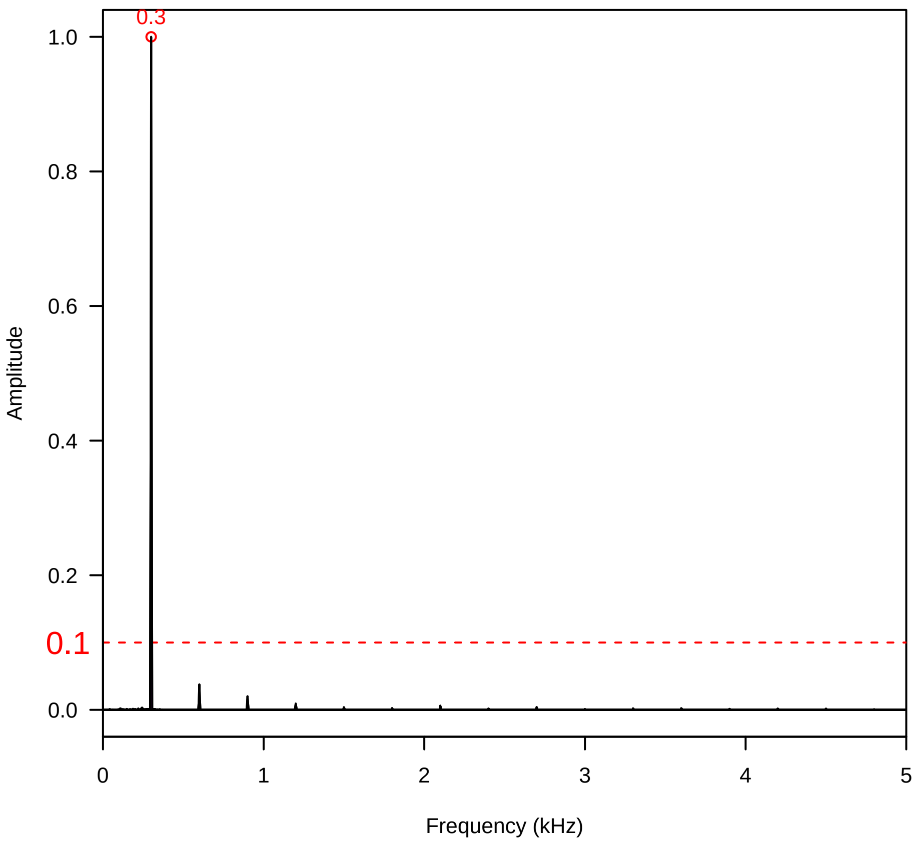

**Vel. = 0.057 ; Str. = PA ; Axis = z ; Fl. accession = 10−s−77−3**

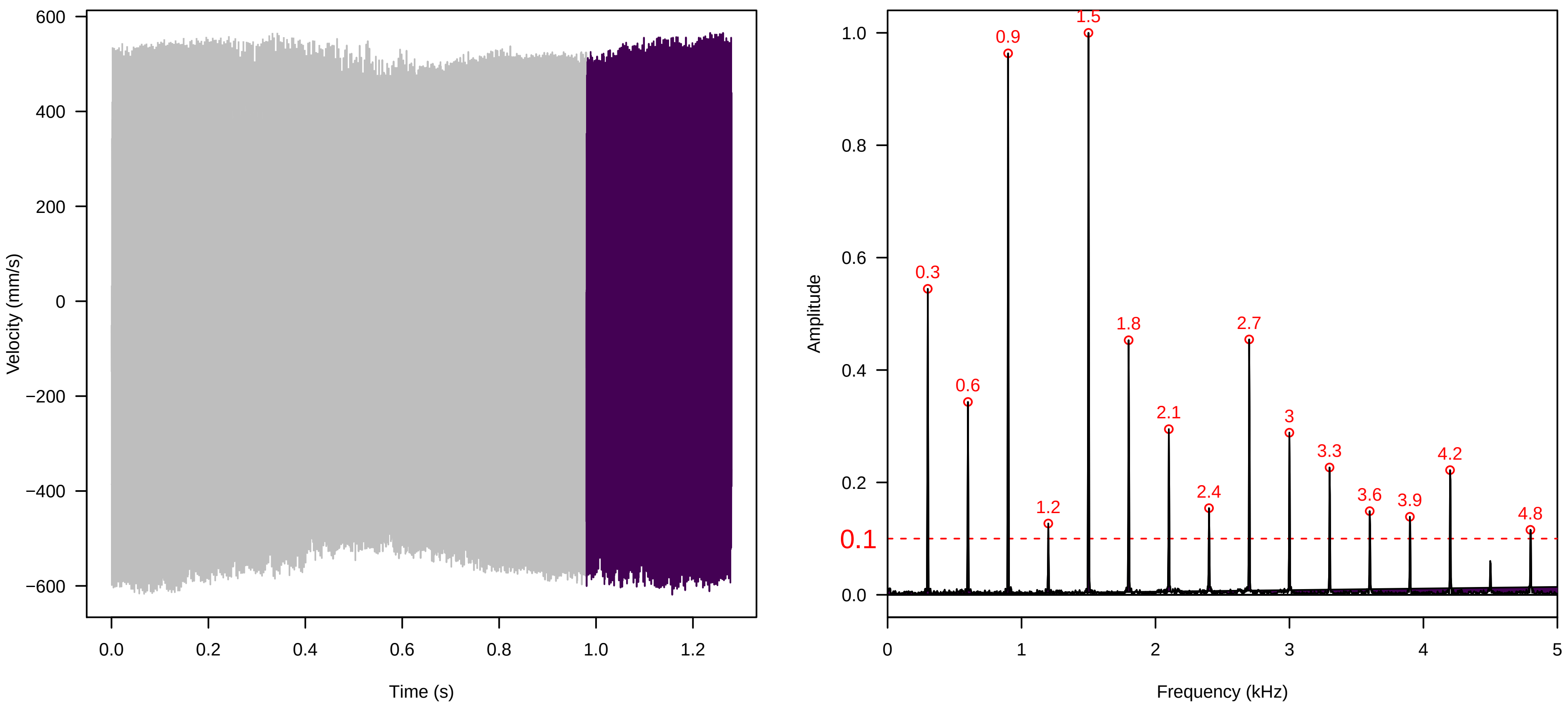


**Vel. = 0.057 ; Str. = Receptacle ; Axis = z ; Fl. accession = 10−s−77−3**

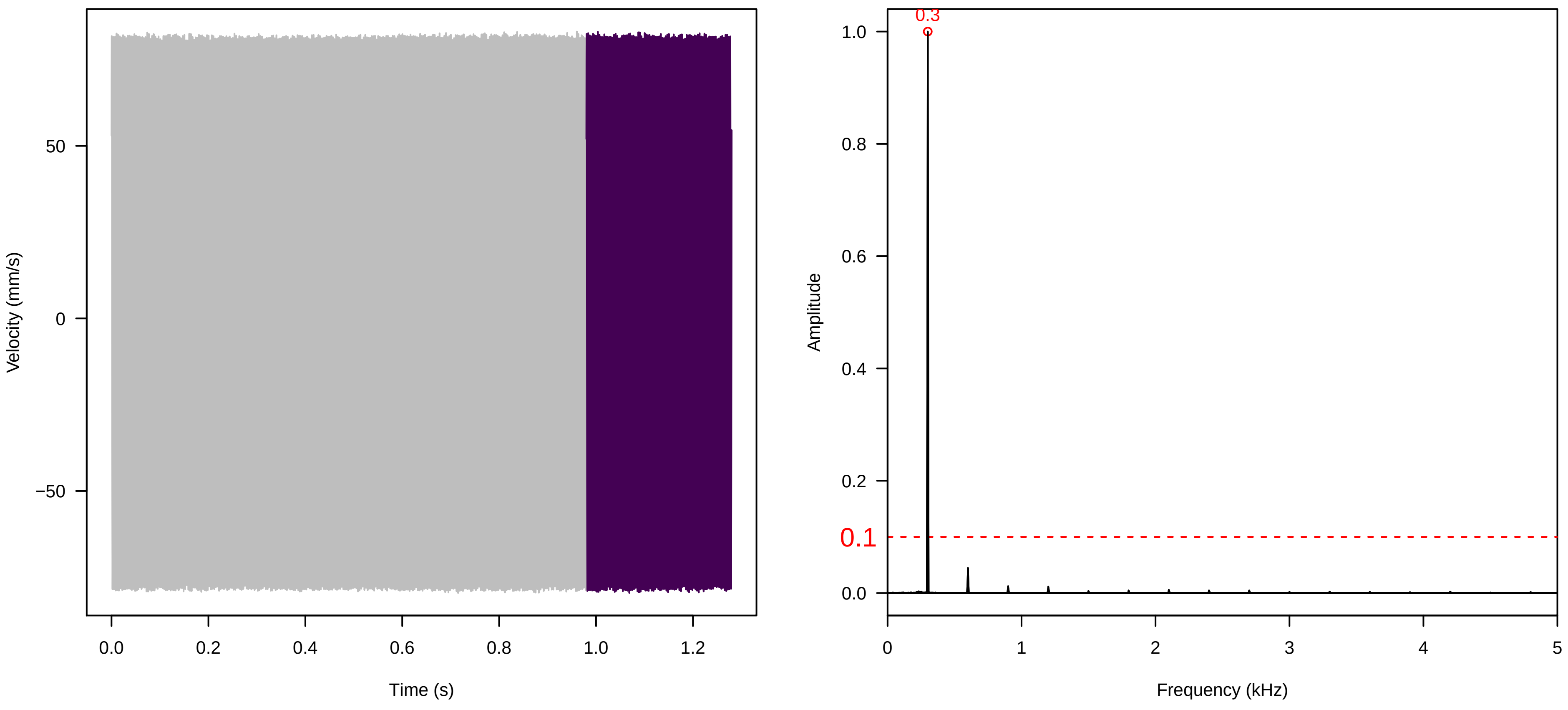


**Vel. = 0.028 ; Str. = PA ; Axis = z ; Fl. accession = 10−s−77−3**

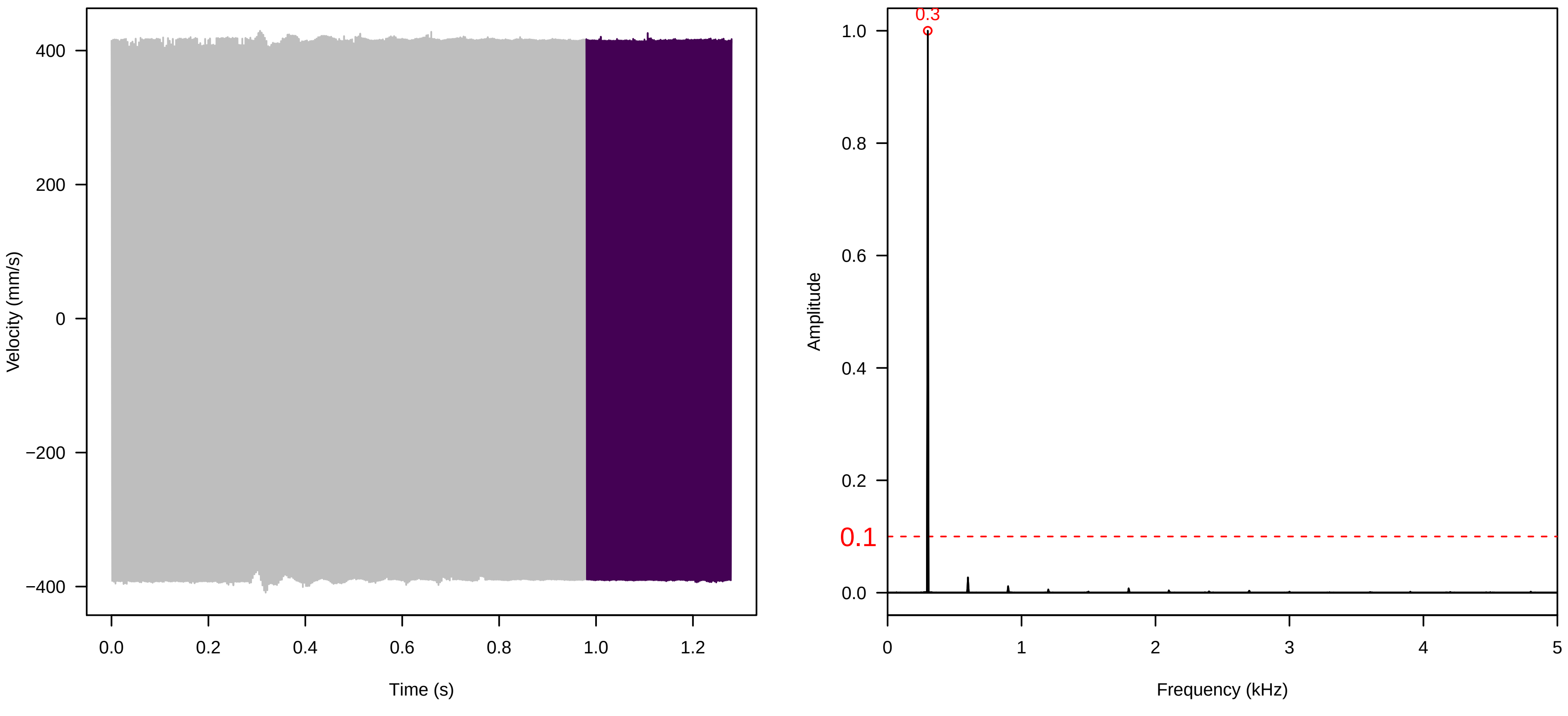

**Vel. =  0.028 ; Str. =  Receptacle ; Axis =  z ; Fl. accession = 10−s−77−3**

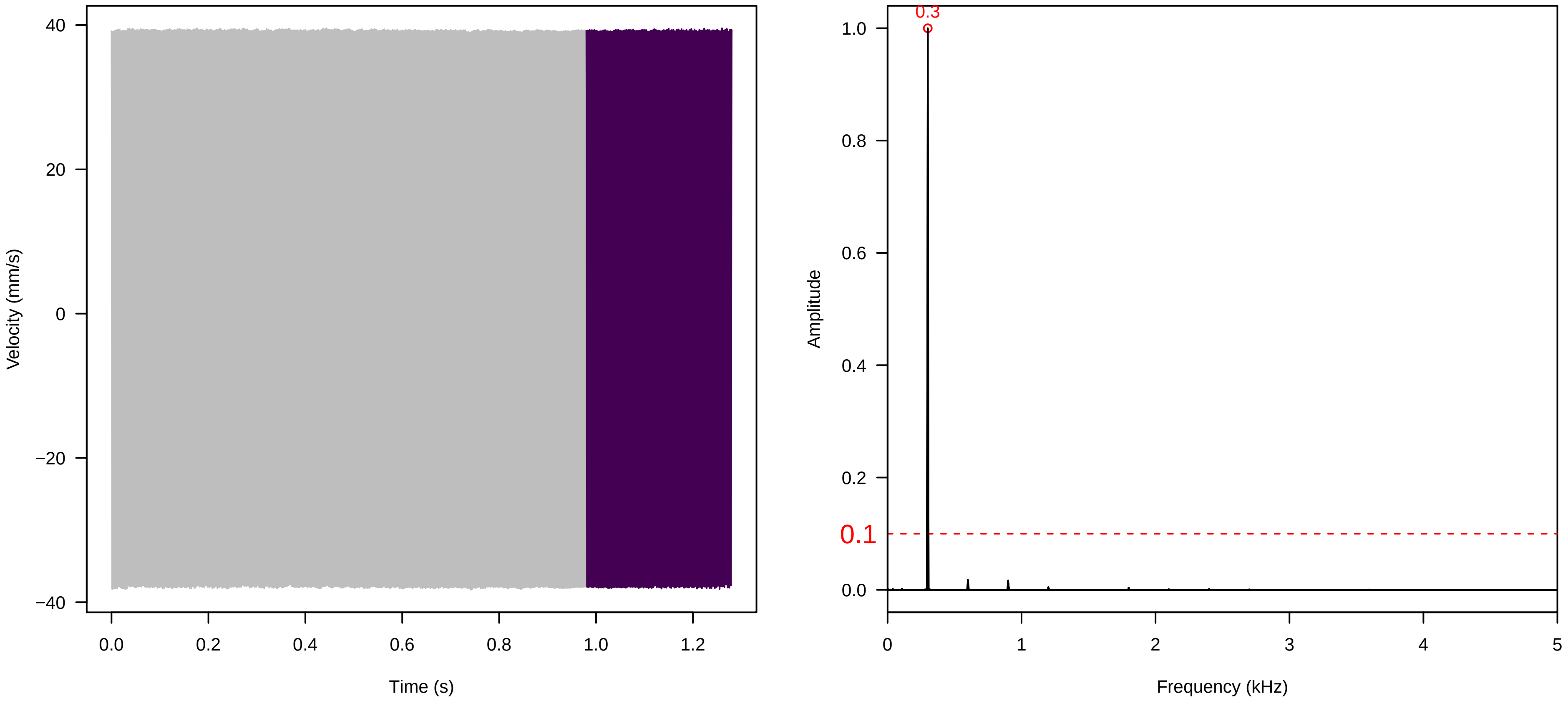


**Vel. =  0.028 ; Str. =  FA ; Axis =  z ; Fl. accession = 10−s−77−3**

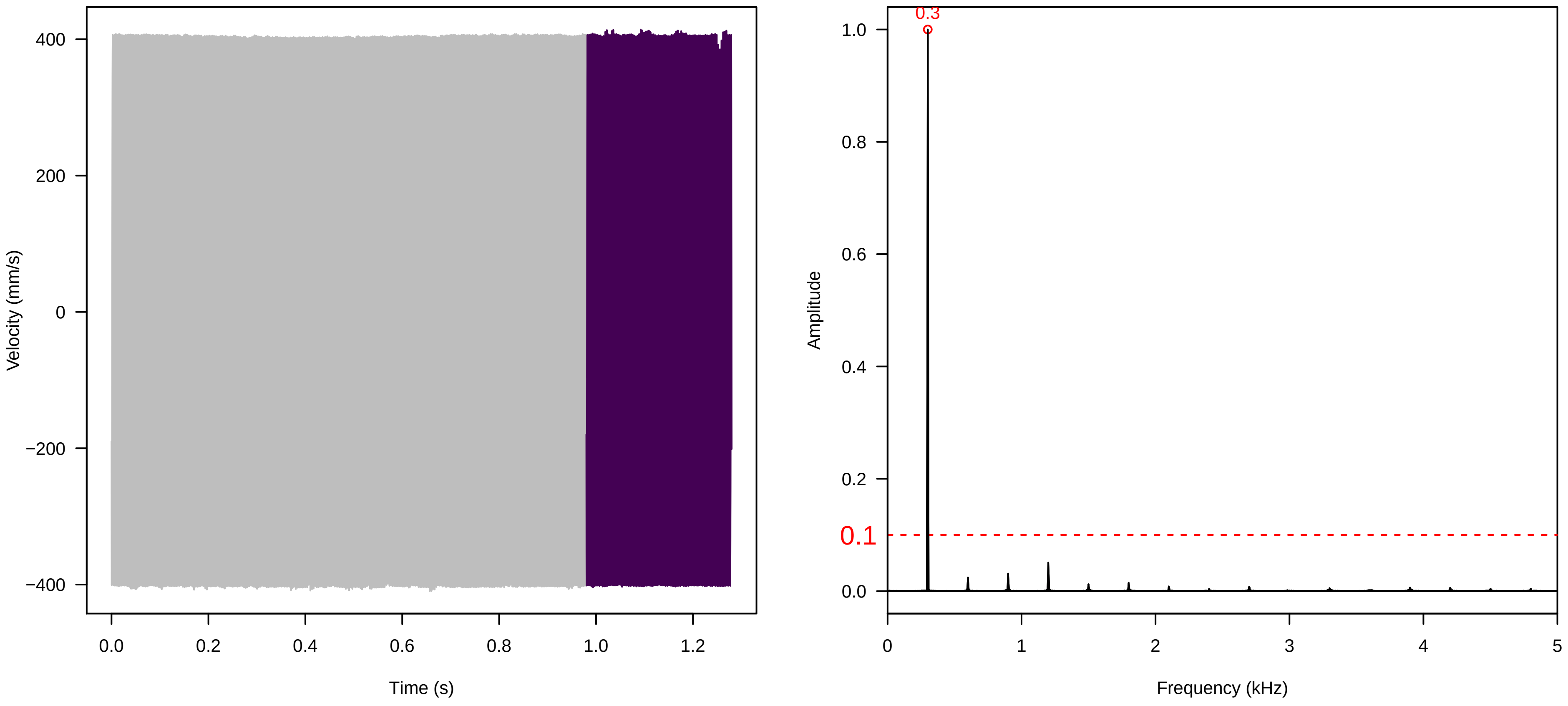


**Vel. =  0.028 ; Str. =  Receptacle ; Axis =  z ; Fl. accession = 10−s−77−3**

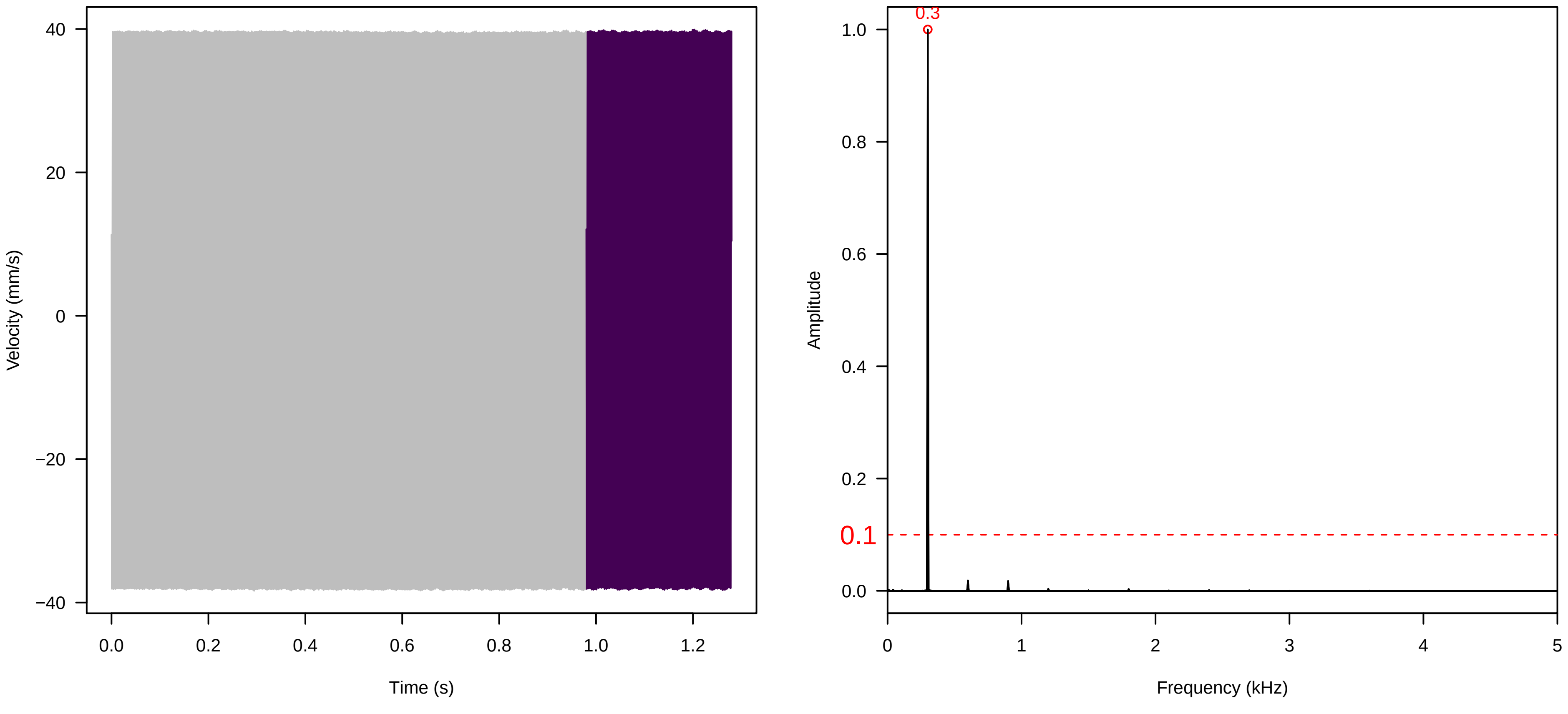

**Vel. = 0.028 ; Str. = Corolla ; Axis = z ; Fl. accession = 10−s−77−3**

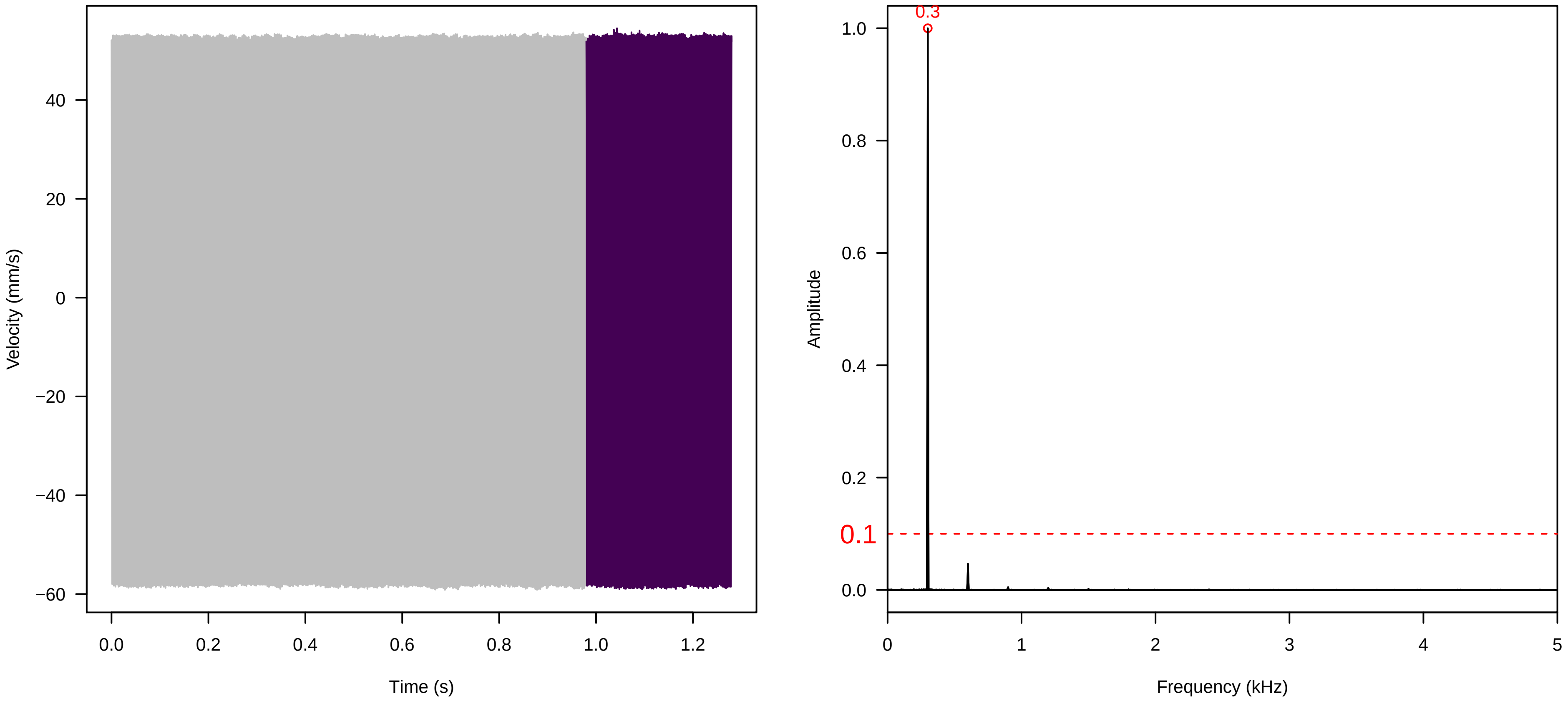


**Vel. = 0.028 ; Str. = Receptacle ; Axis = z ; Fl. accession = 10−s−77−3**

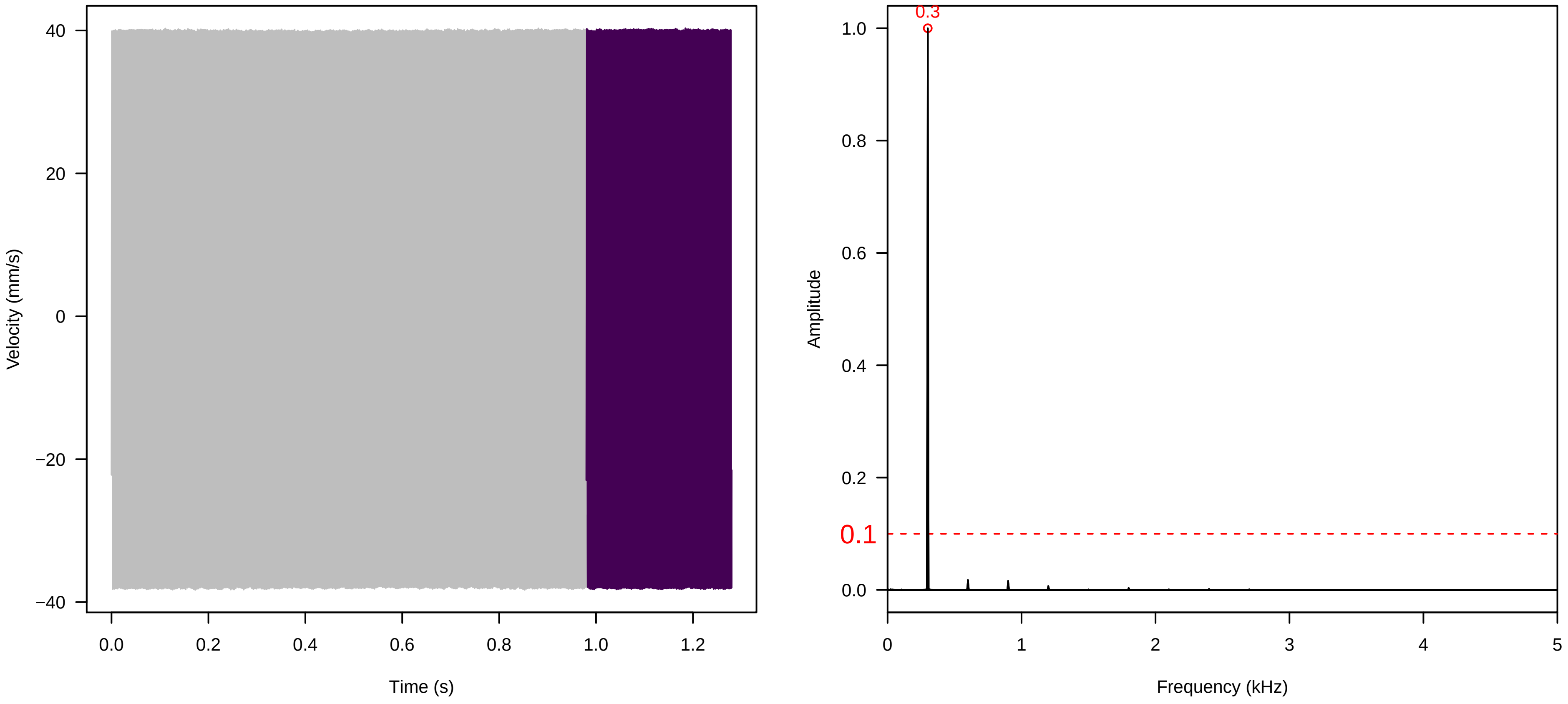


**Vel. = 0.014 ; Str. = Corolla ; Axis = z ; Fl. accession = 10−s−77−3**

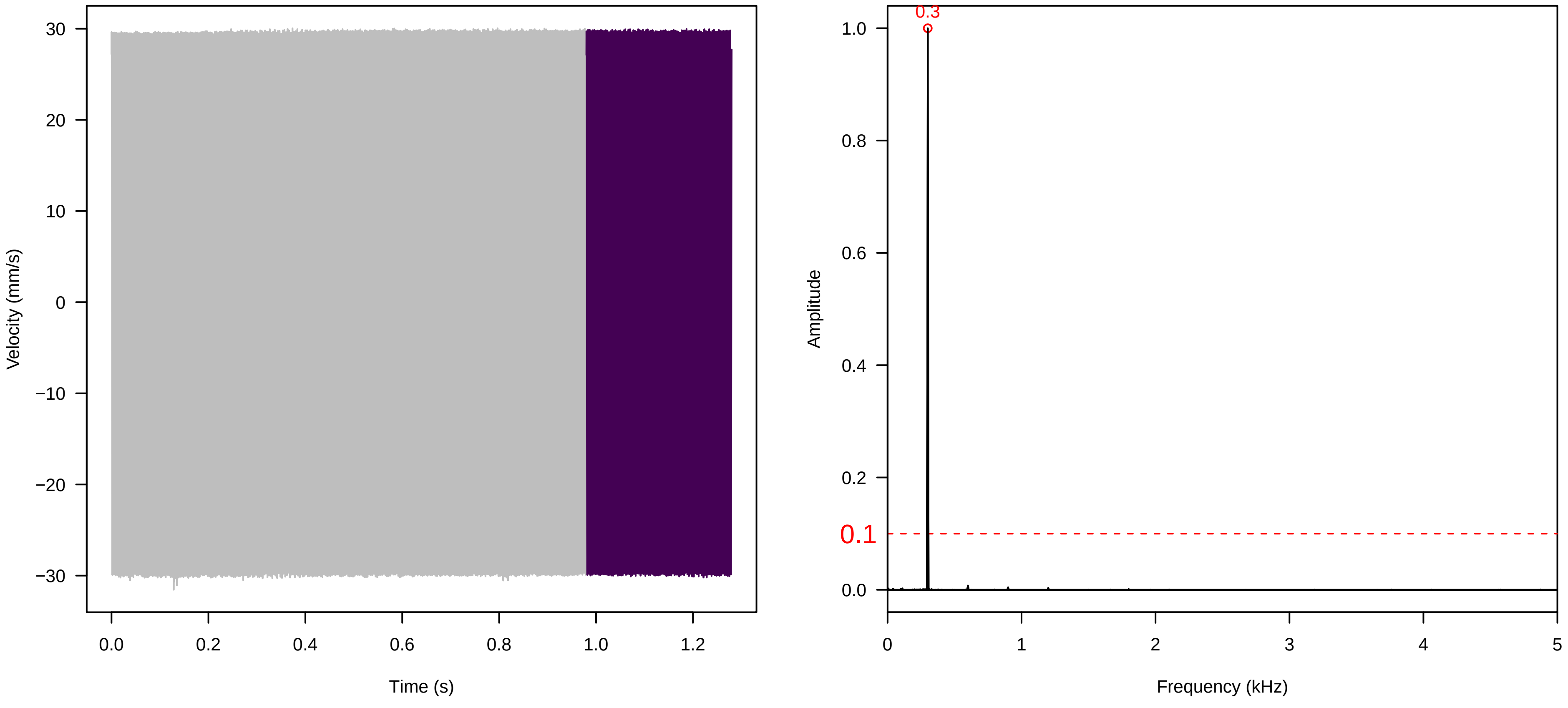

Vel. =  0.014 ; Str. =  Receptacle ; Axis =  z ; Fl. accession = 10−s−77−3

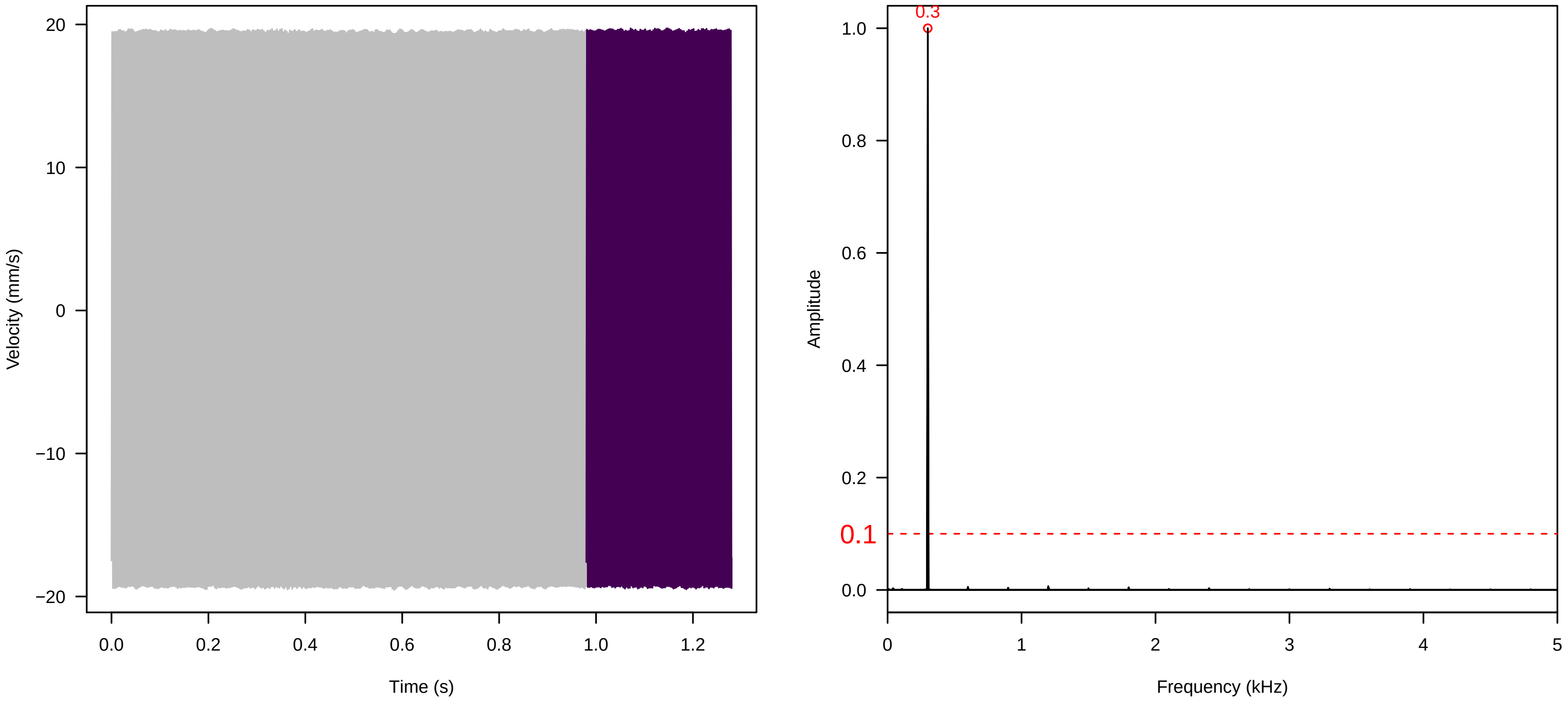


Vel. =  0.014 ; Str. =  FA ; Axis =  z ; Fl. accession = 10−s−77−3

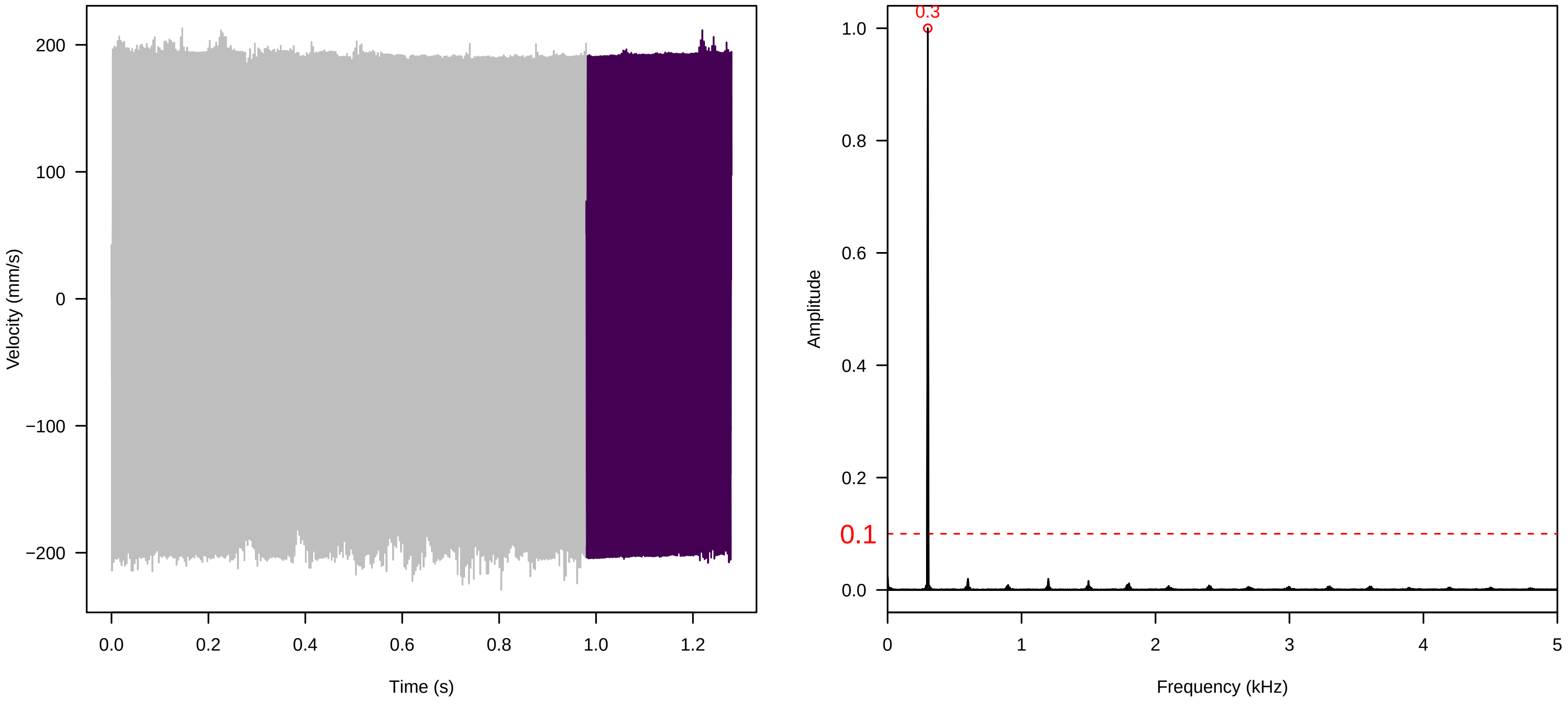


Vel. =  0.014 ; Str. =  Receptacle ; Axis =  z ; Fl. accession = 10−s−77−3

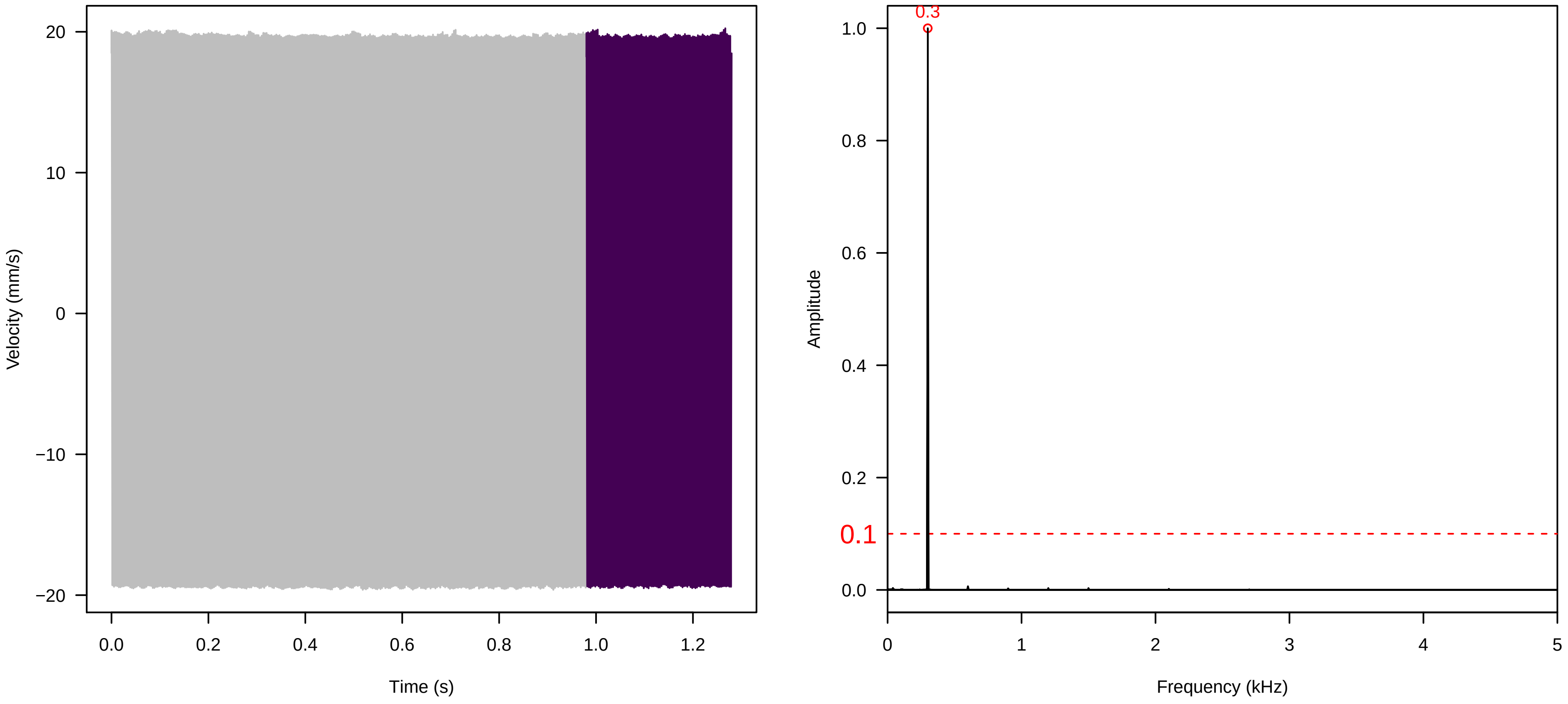

**Vel. = 0.014 ; Str. = PA ; Axis = z ; Fl. accession = 10−s−77−3**

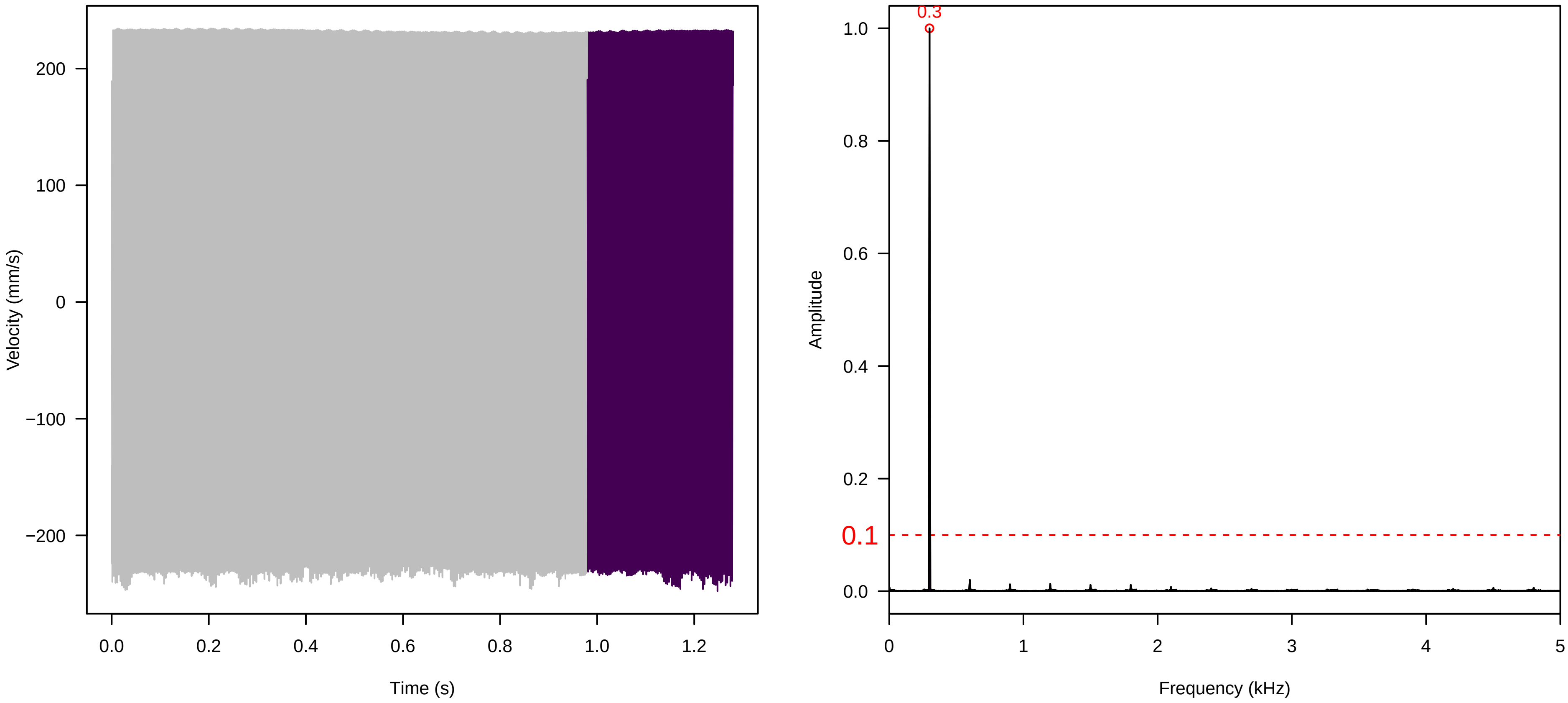


**Vel. = 0.014 ; Str. = Receptacle ; Axis = z ; Fl. accession = 10−s−77−3**

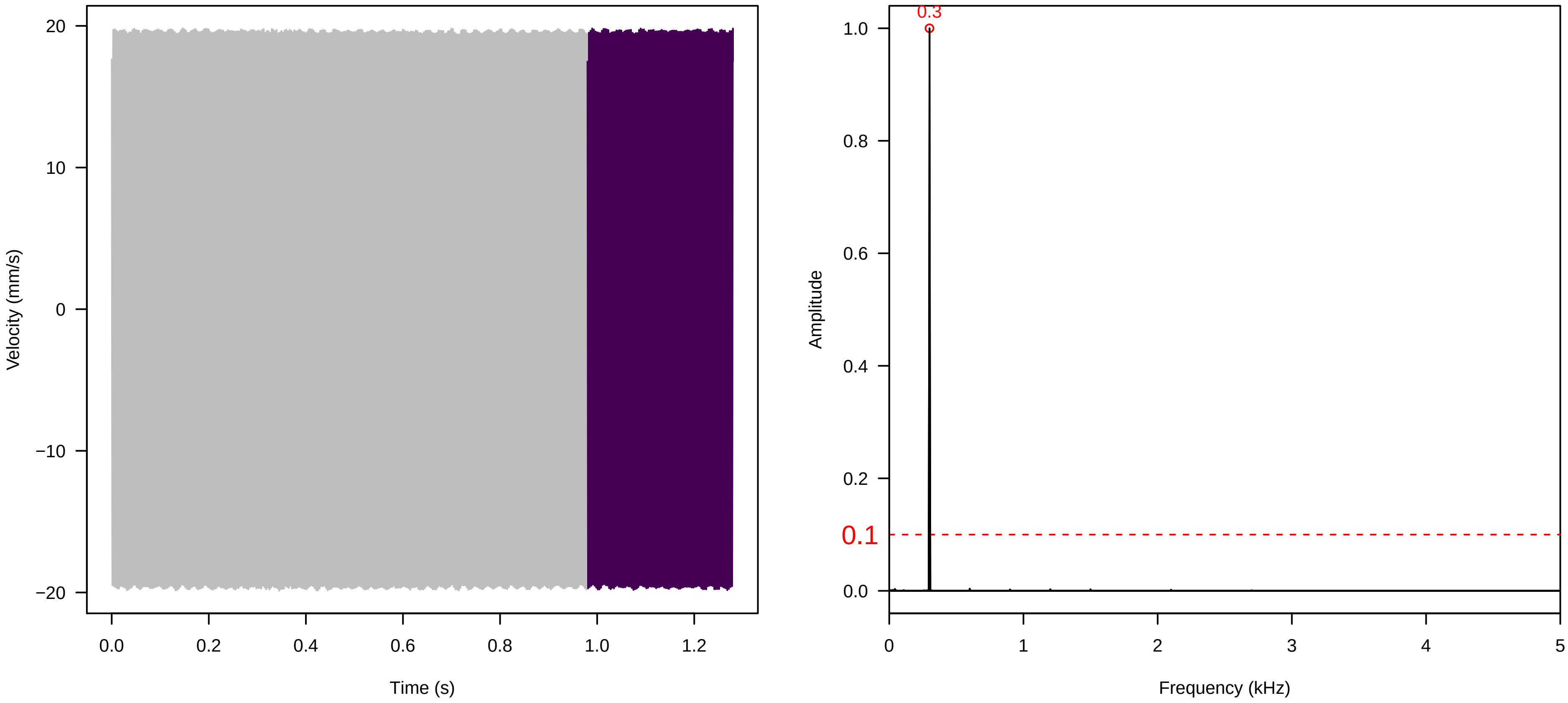


**Vel. = 0.014 ; Str. = Corolla ; Axis = x ; Fl. accession = 10−s−77−3**

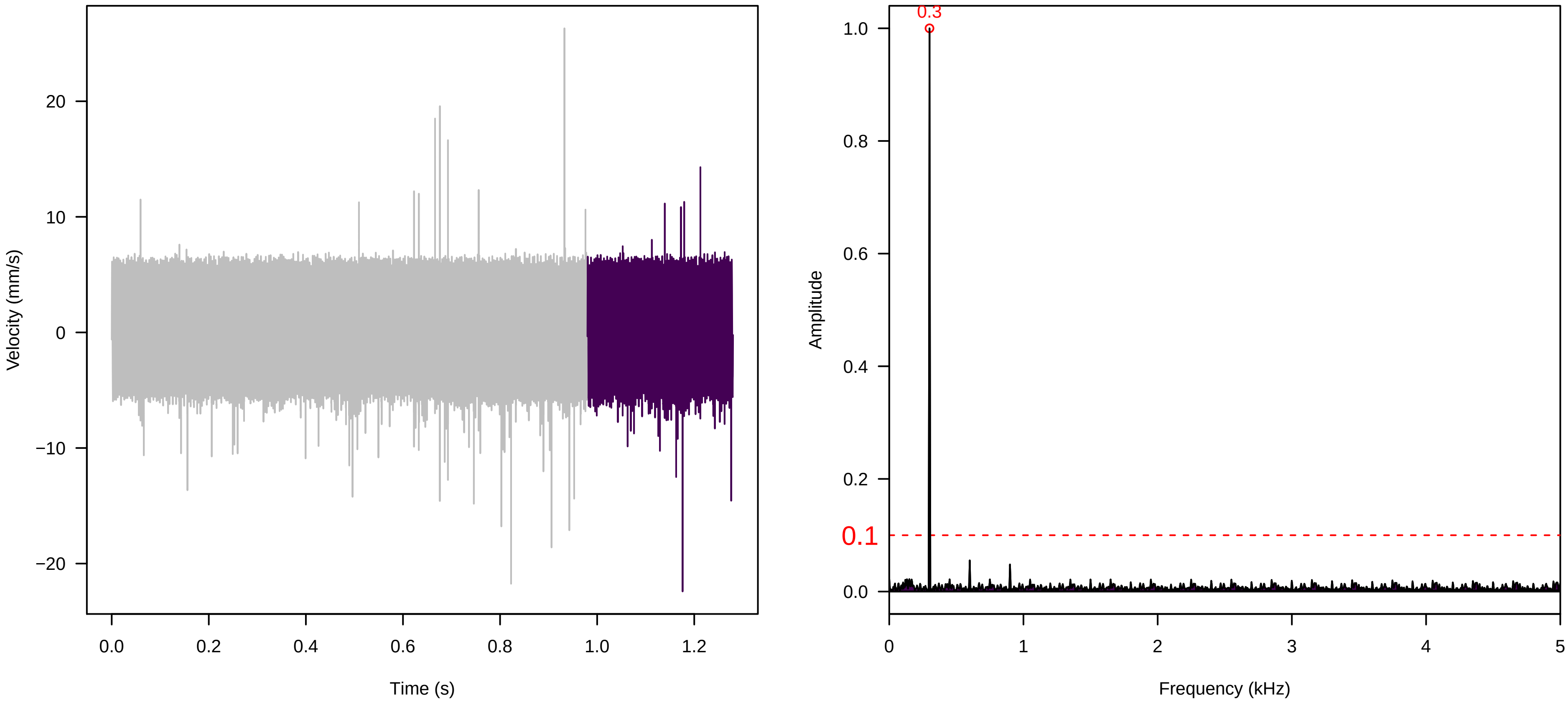

**Vel. =  0.014 ; Str. =  Receptacle ; Axis =  x ; Fl. accession = 10−s−77−3**

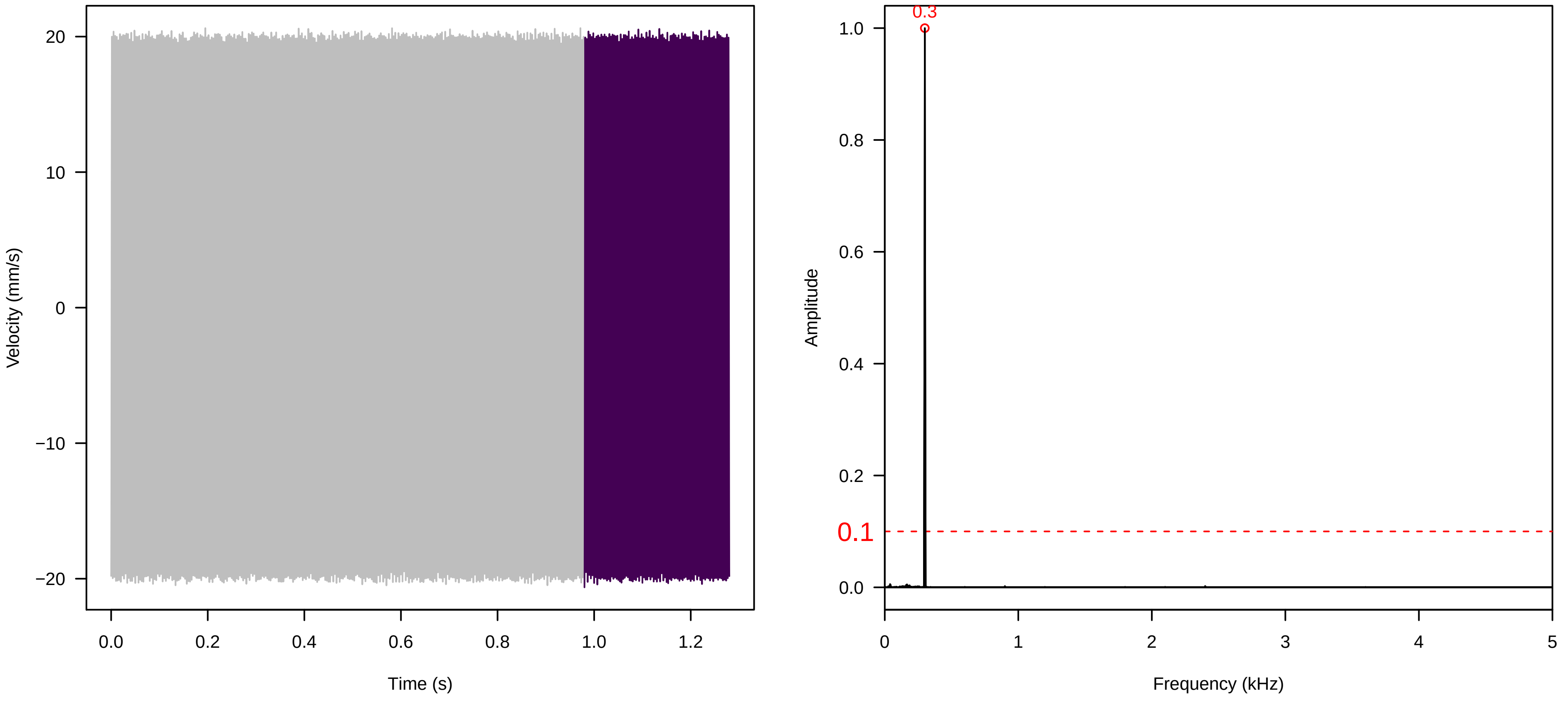


**Vel. =  0.014 ; Str. =  FA ; Axis =  x ; Fl. accession = 10−s−77−3**

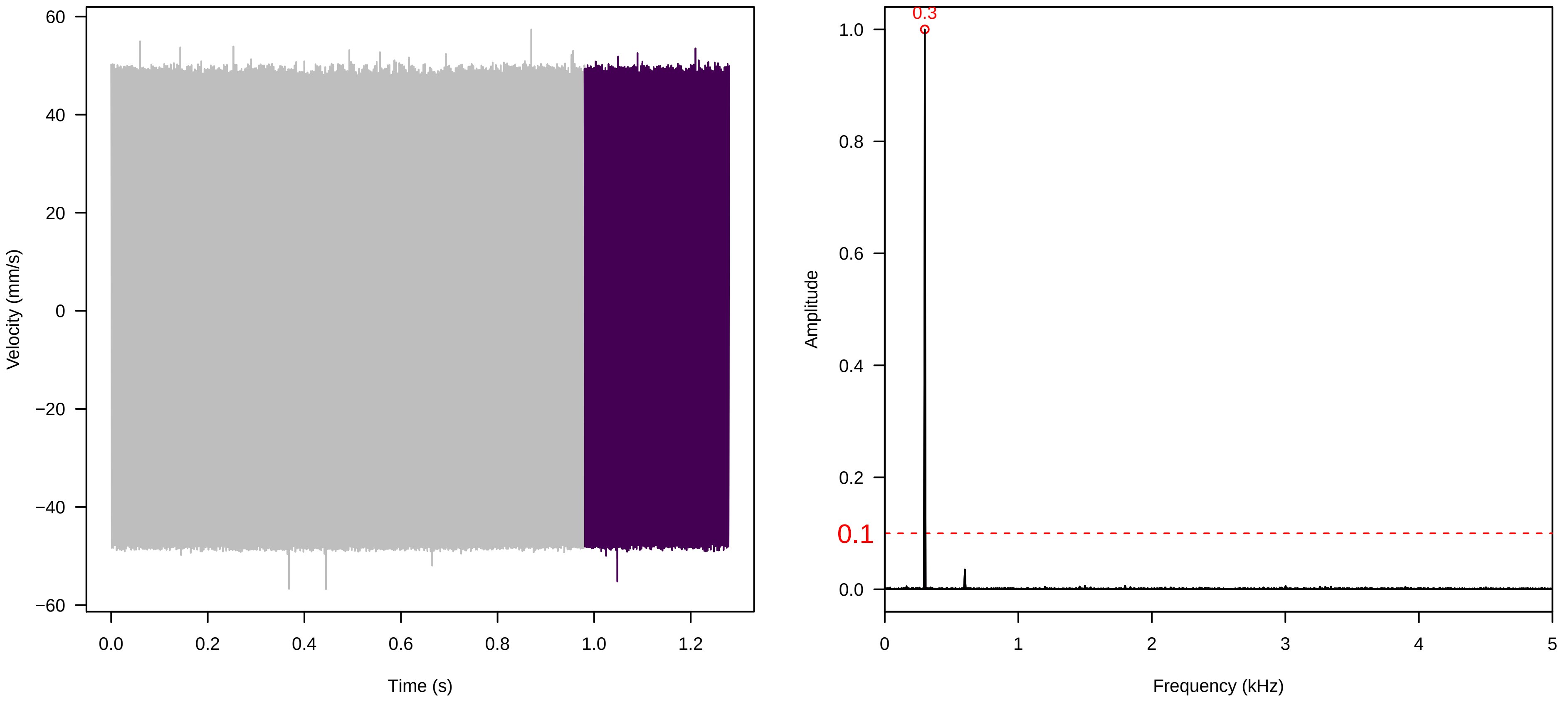


**Vel. =  0.014 ; Str. =  Receptacle ; Axis =  x ; Fl. accession = 10−s−77−3**

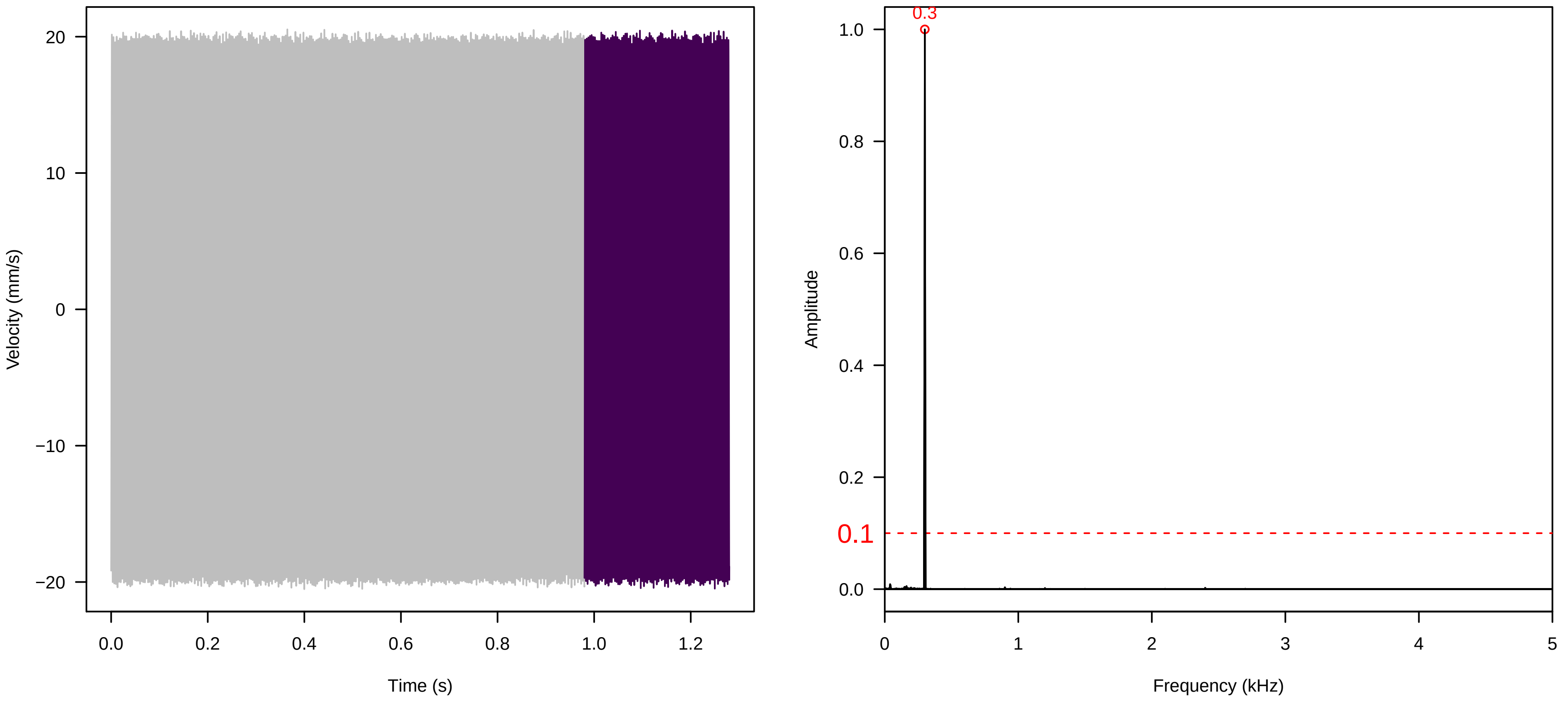

**Vel. = 0.014 ; Str. = PA ; Axis = x ; Fl. accession = 10−s−77−3**

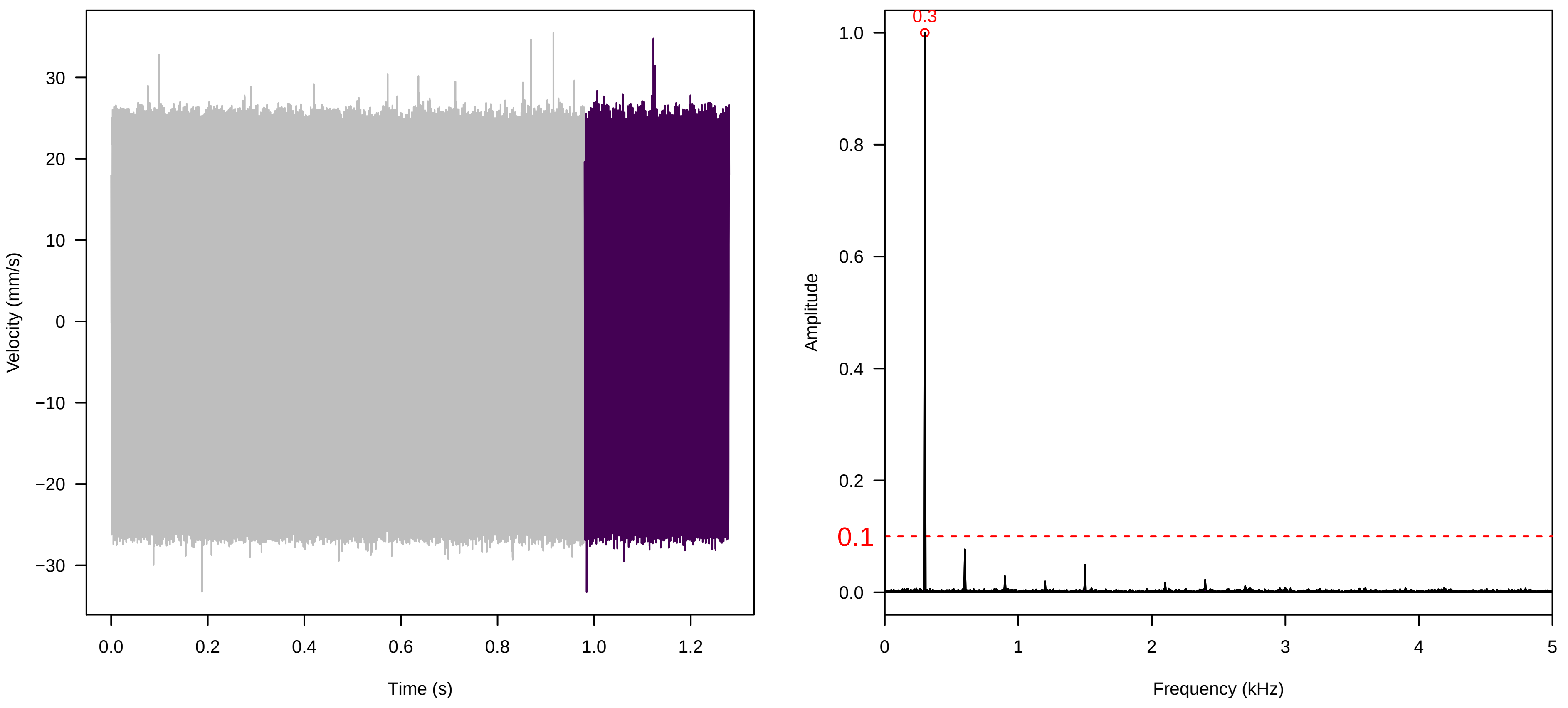


**Vel. = 0.014 ; Str. = Receptacle ; Axis = x ; Fl. accession = 10−s−77−3**

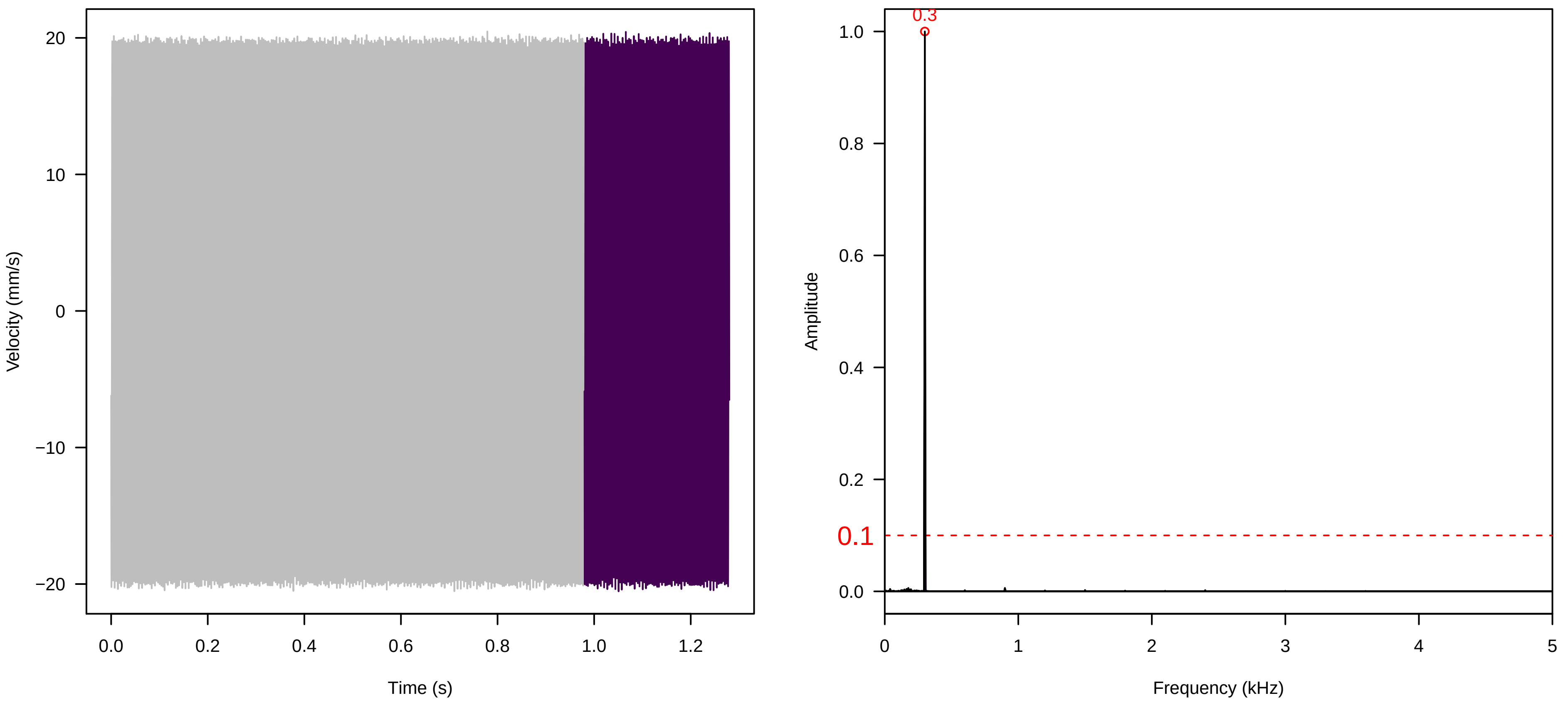


**Vel. = 0.028 ; Str. = PA ; Axis = x ; Fl. accession = 10−s−77−3**

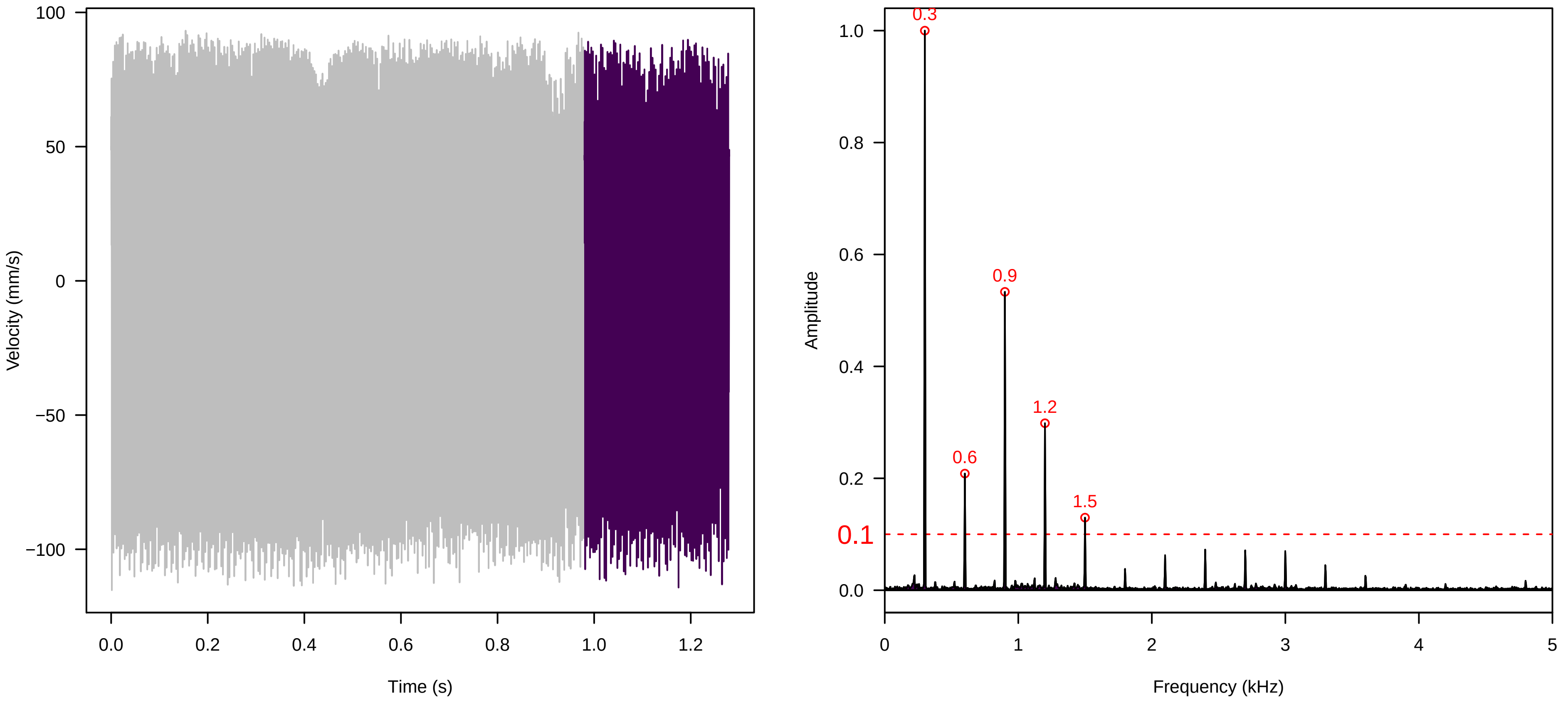

**Vel. =  0.028 ; Str. =  Receptacle ; Axis =  x ; Fl. accession = 10−s−77−3**

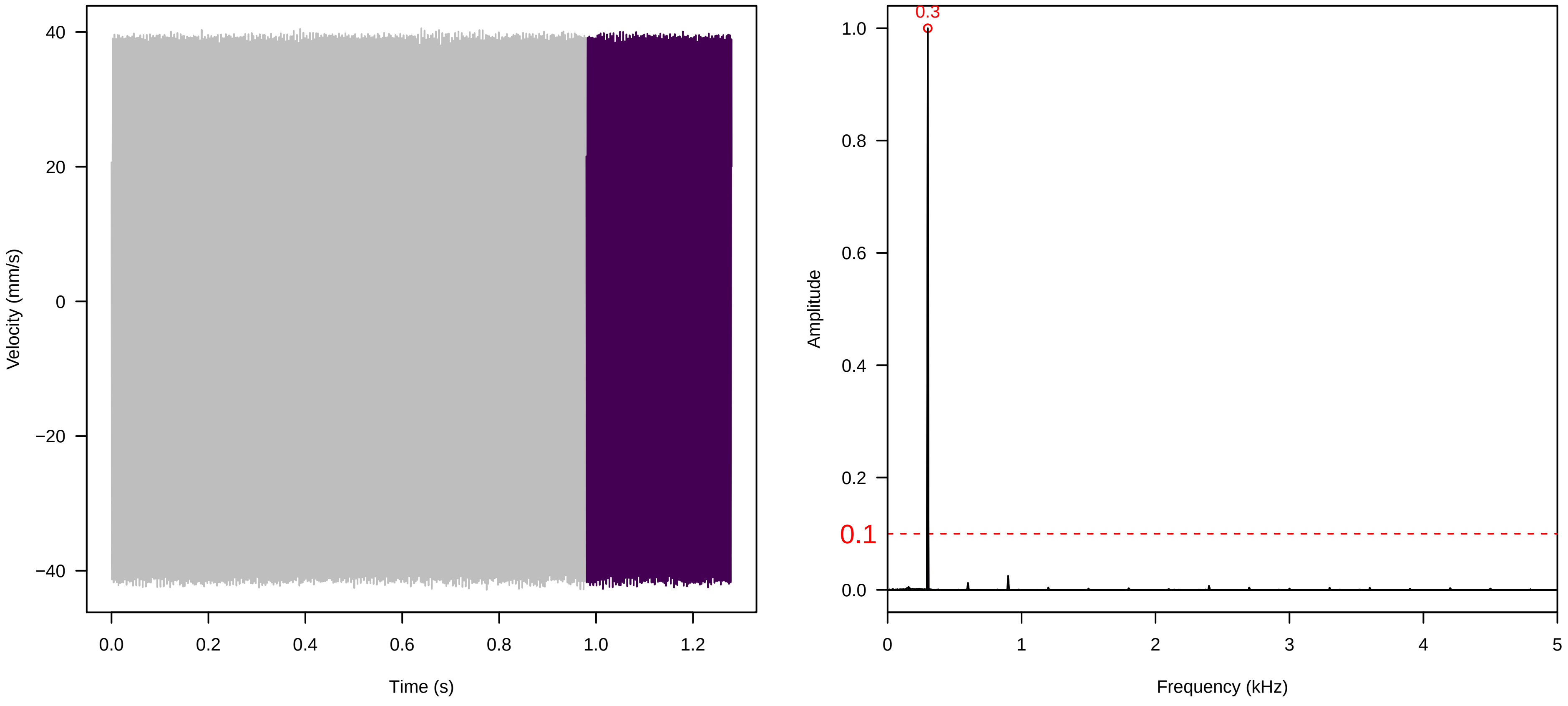


**Vel. =  0.028 ; Str. =  FA ; Axis =  x ; Fl. accession = 10−s−77−3**

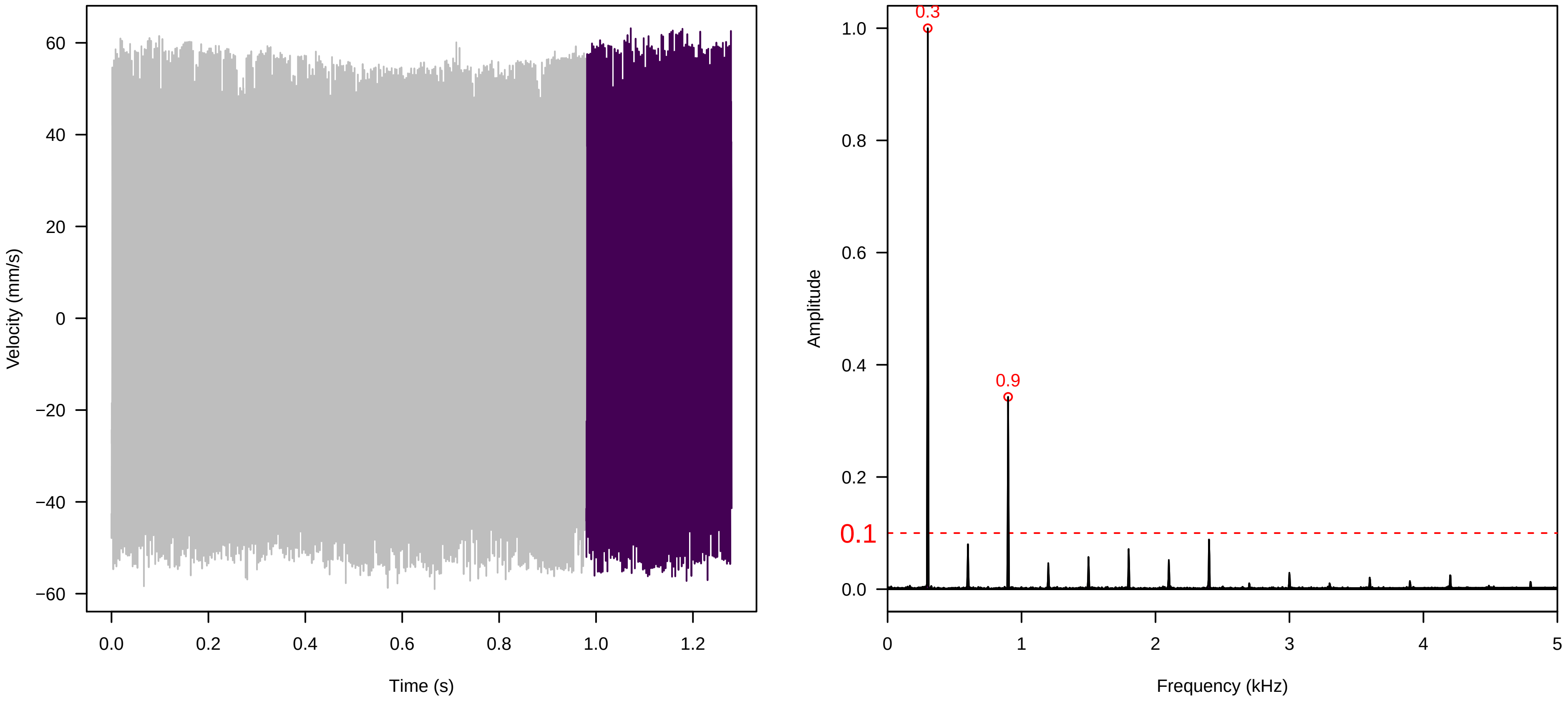


**Vel. =  0.028 ; Str. =  Receptacle ; Axis =  x ; Fl. accession = 10−s−77−3**

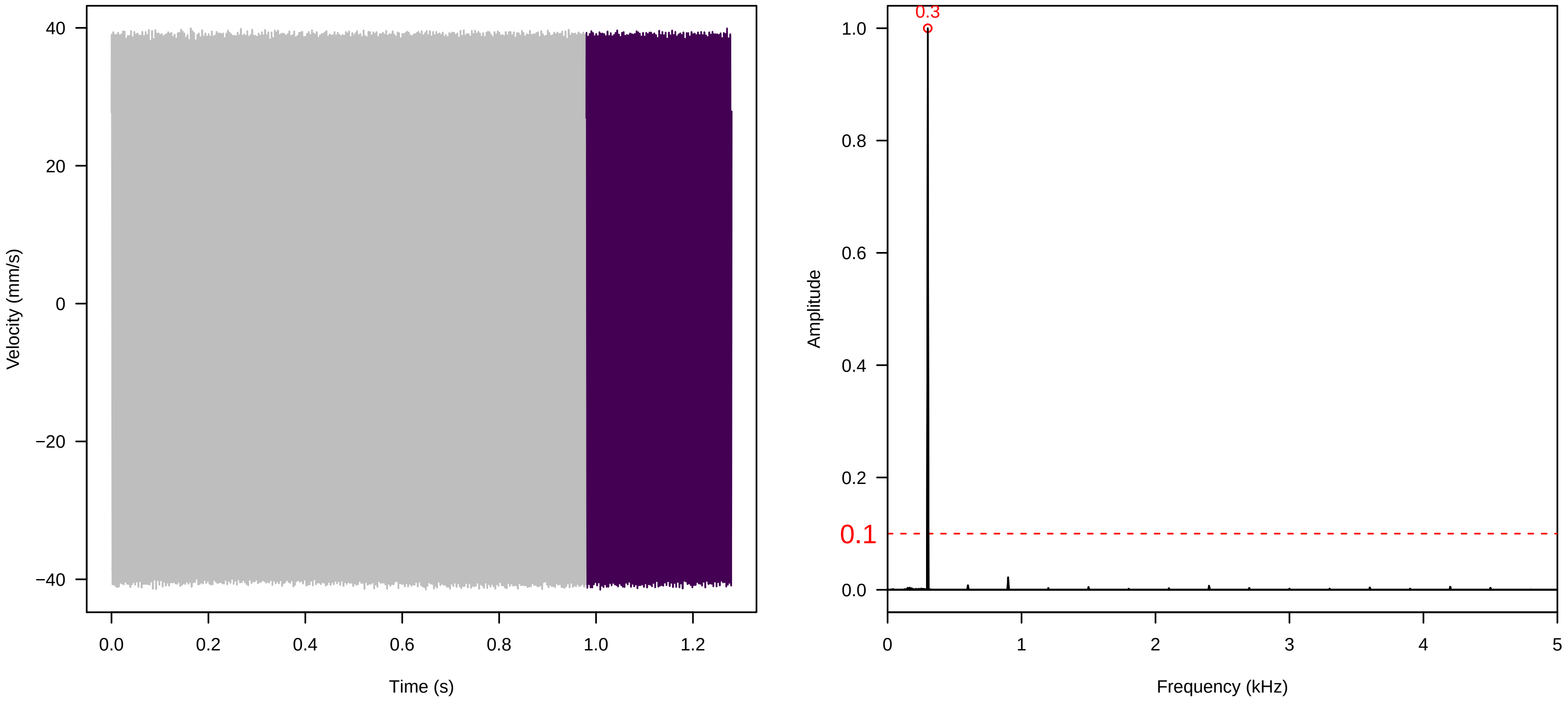

**Vel. =  0.028 ; Str. =  Corolla ; Axis =  x ; Fl. accession = 10−s−77−3**

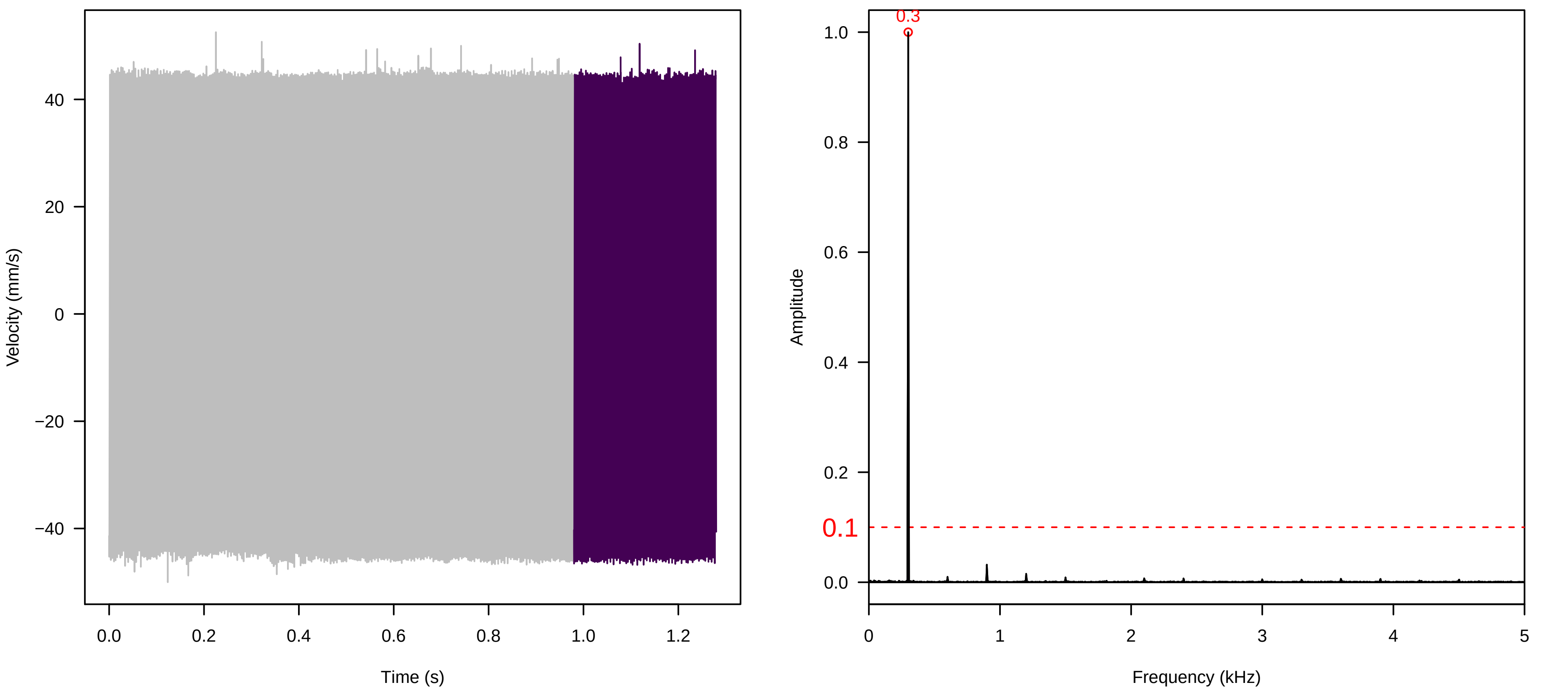


**Vel. =  0.028 ; Str. =  Receptacle ; Axis =  x ; Fl. accession = 10−s−77−3**

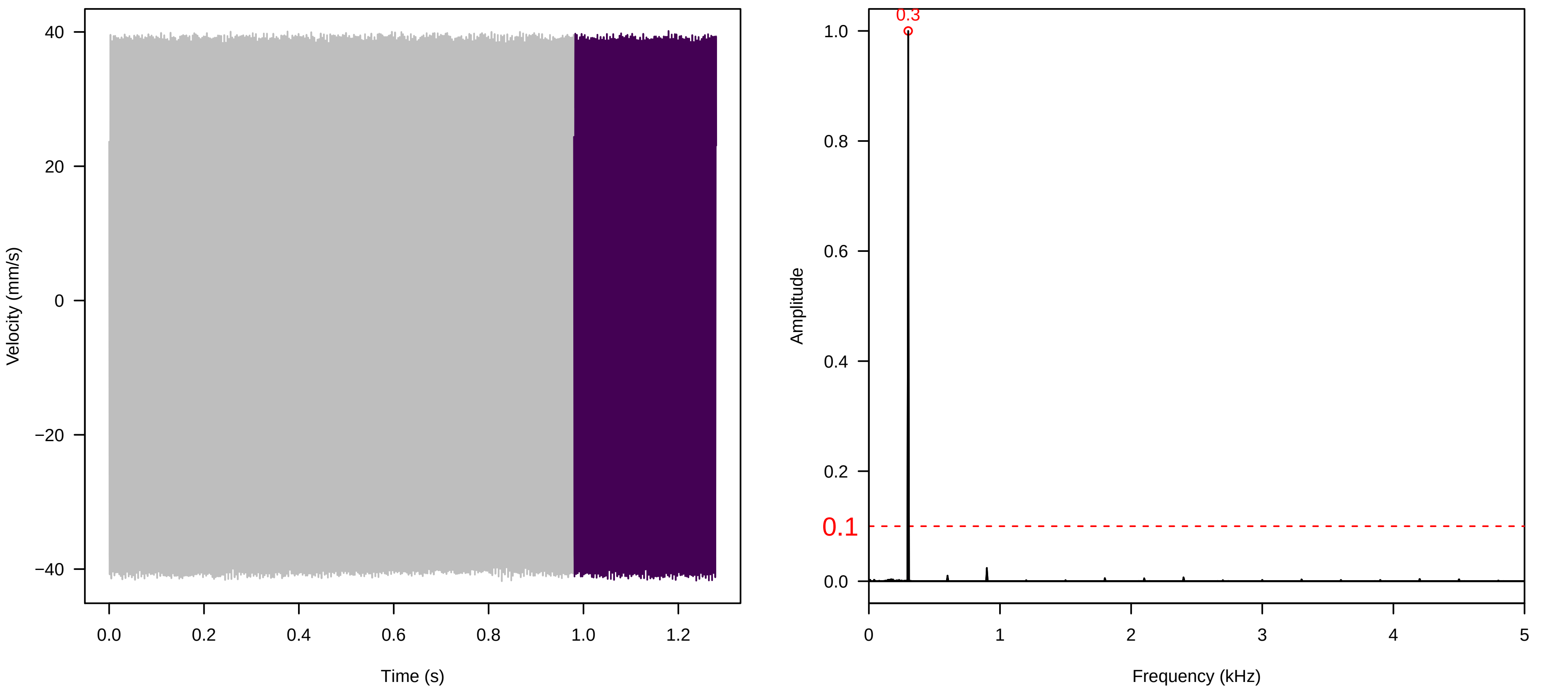


**Vel. =  0.057 ; Str. =  Corolla ; Axis =  x ; Fl. accession = 10−s−77−3**

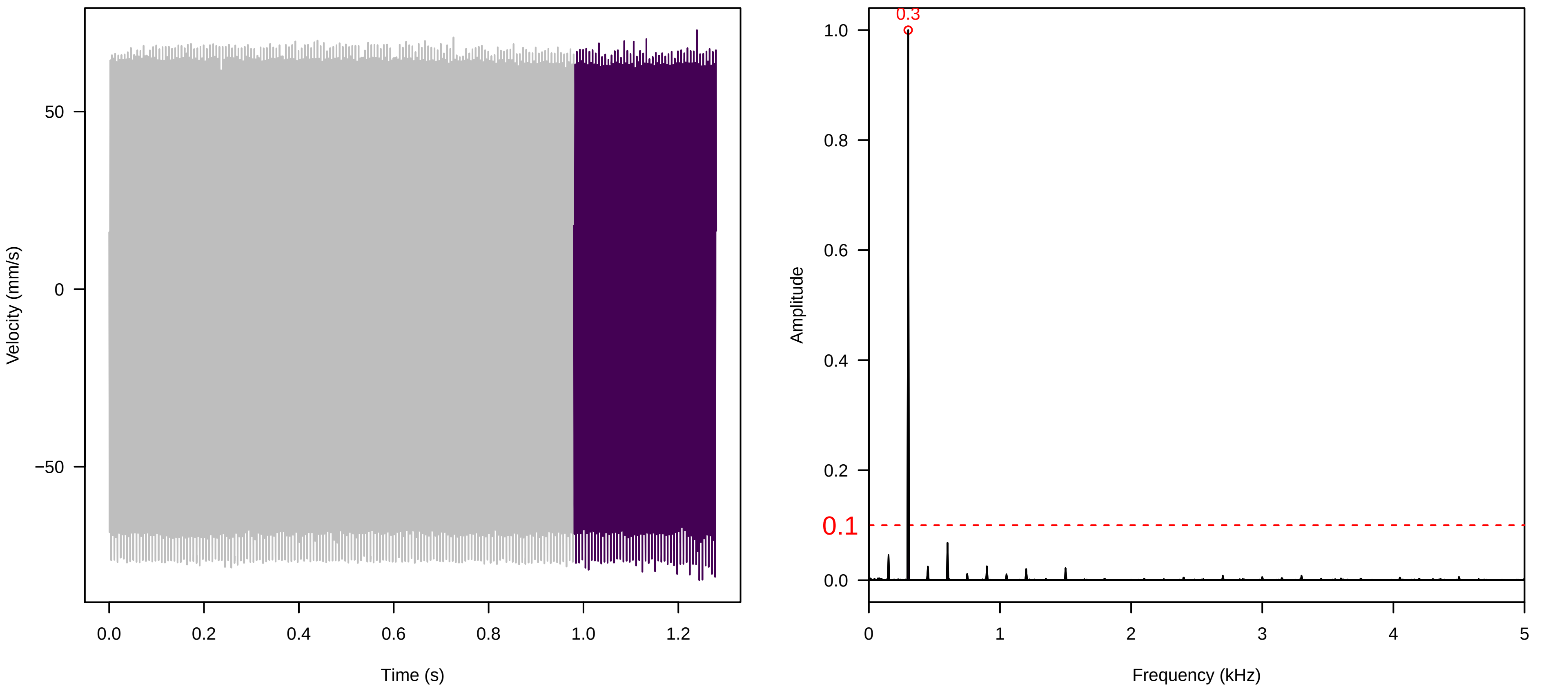

**Vel. = 0.057 ; Str. = Receptacle ; Axis = x ; Fl. accession = 10−s−77−3**

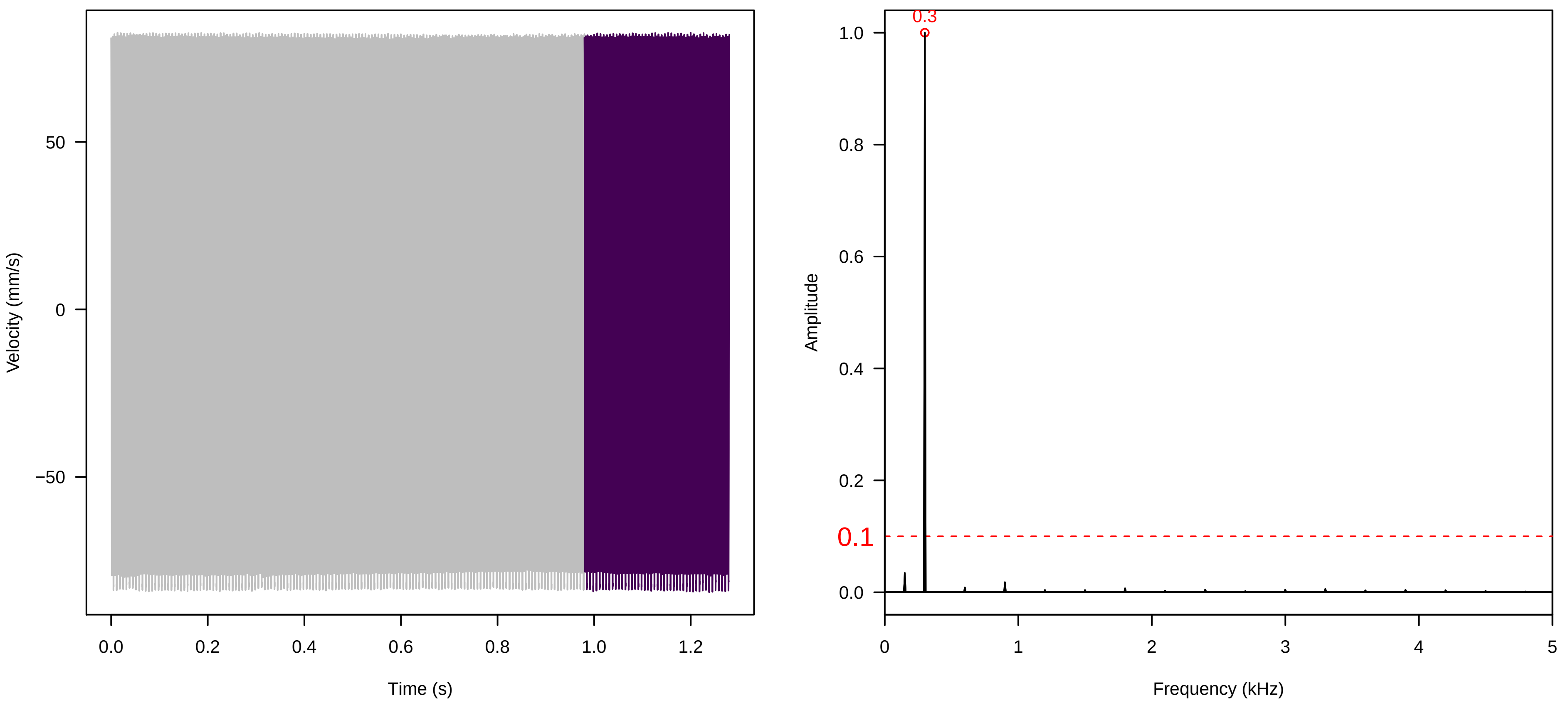


**Vel. = 0.057 ; Str. = FA ; Axis = x ; Fl. accession = 10−s−77−3**

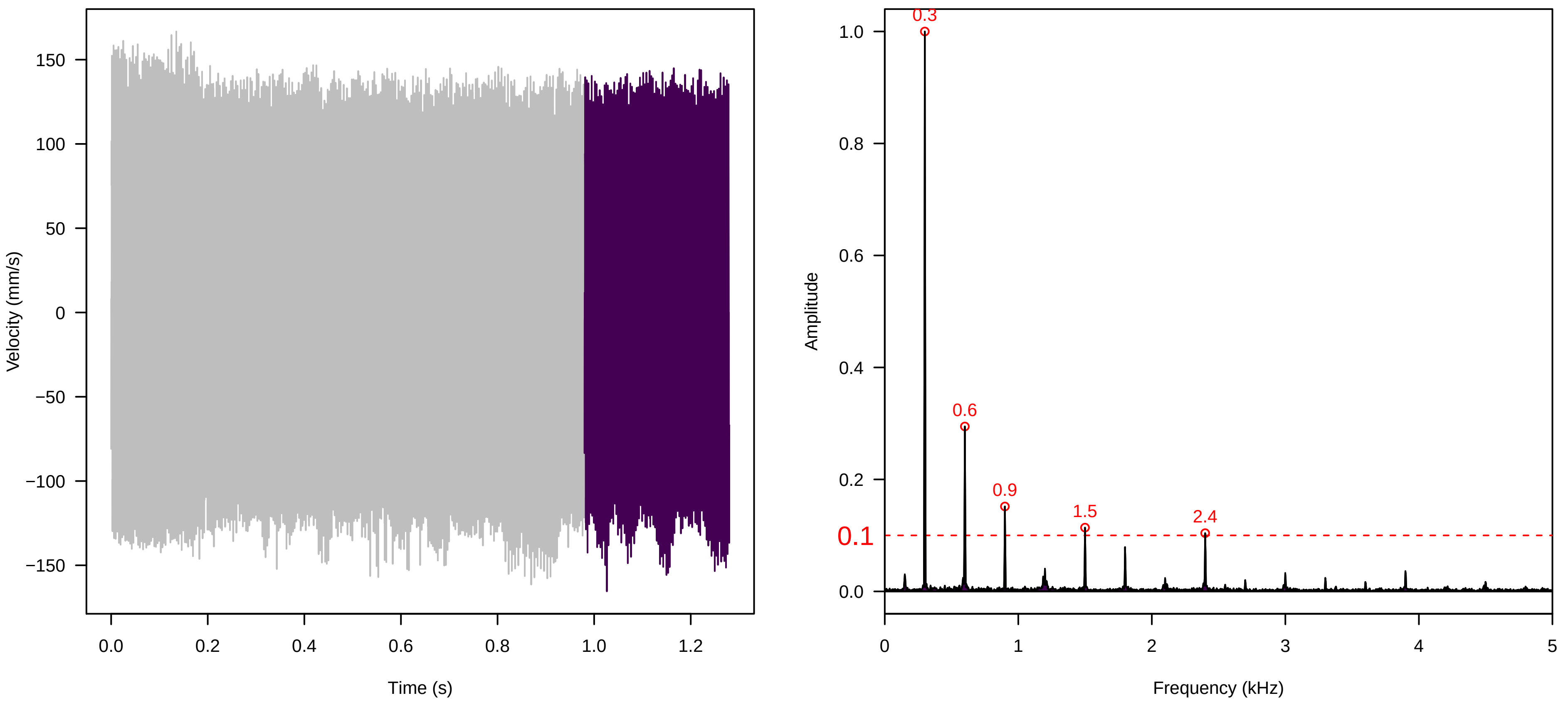


**Vel. = 0.057 ; Str. = Receptacle ; Axis = x ; Fl. accession = 10−s−77−3**

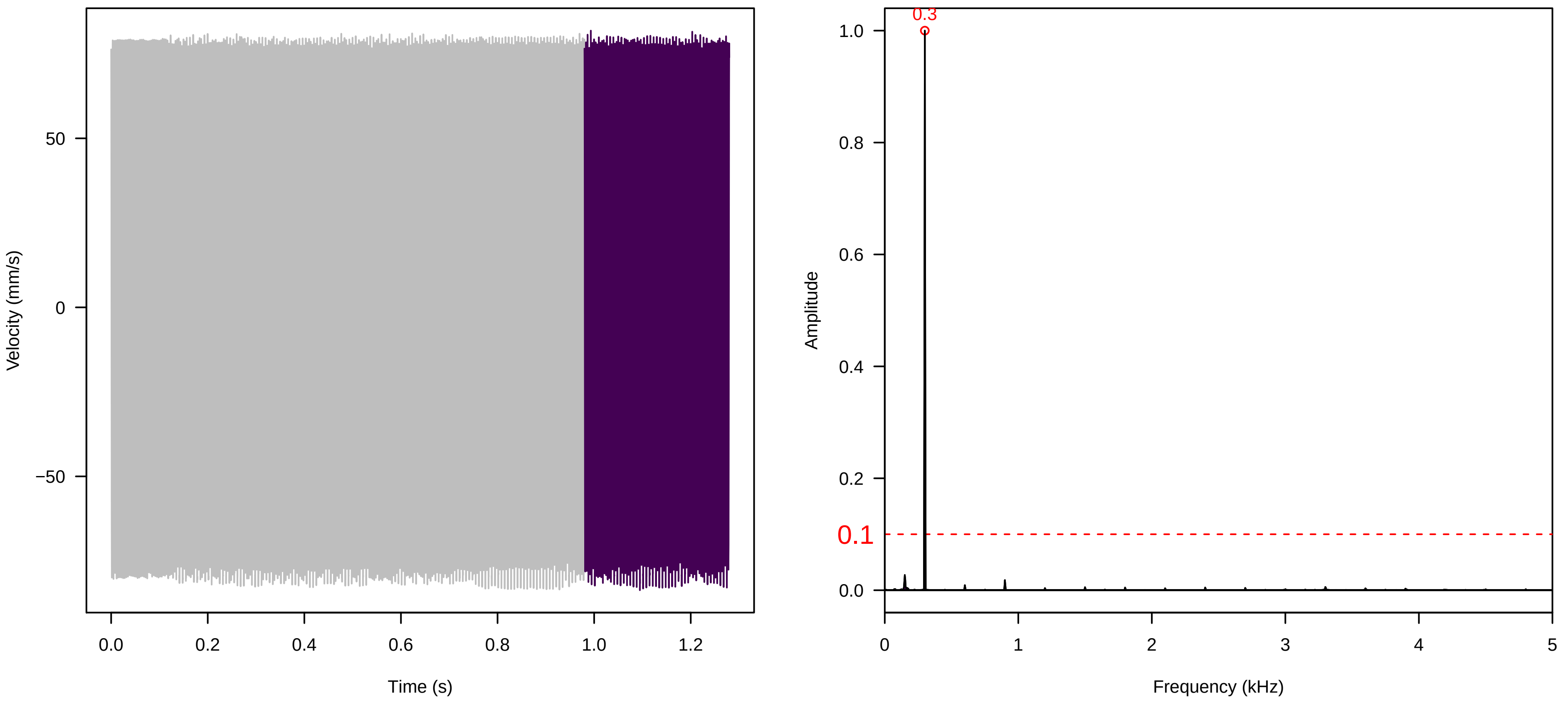

Vel. = 0.057 ; Str. = PA ; Axis = x ; Fl. accession = 10−s−77−3

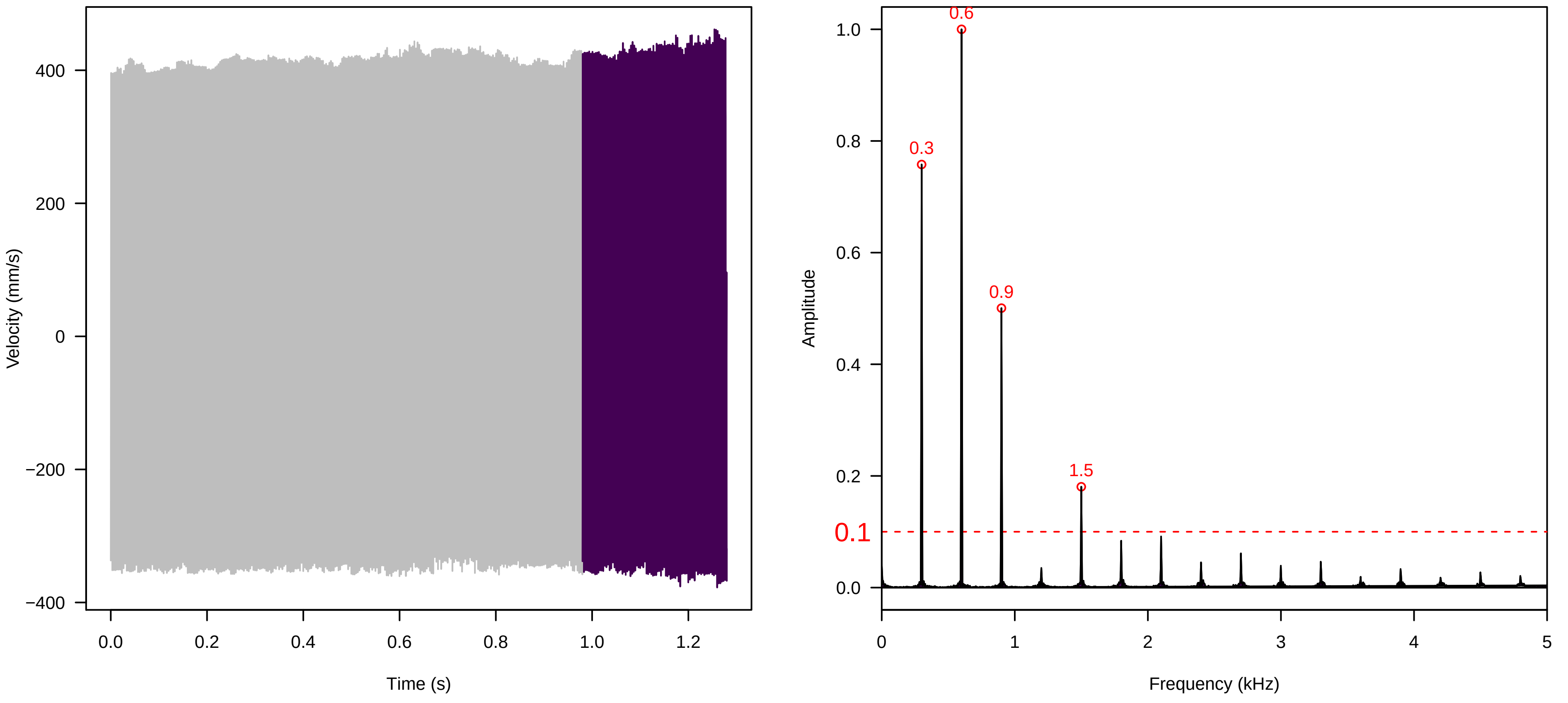


Vel. = 0.057 ; Str. = Receptacle ; Axis = x ; Fl. accession = 10−s−77−3

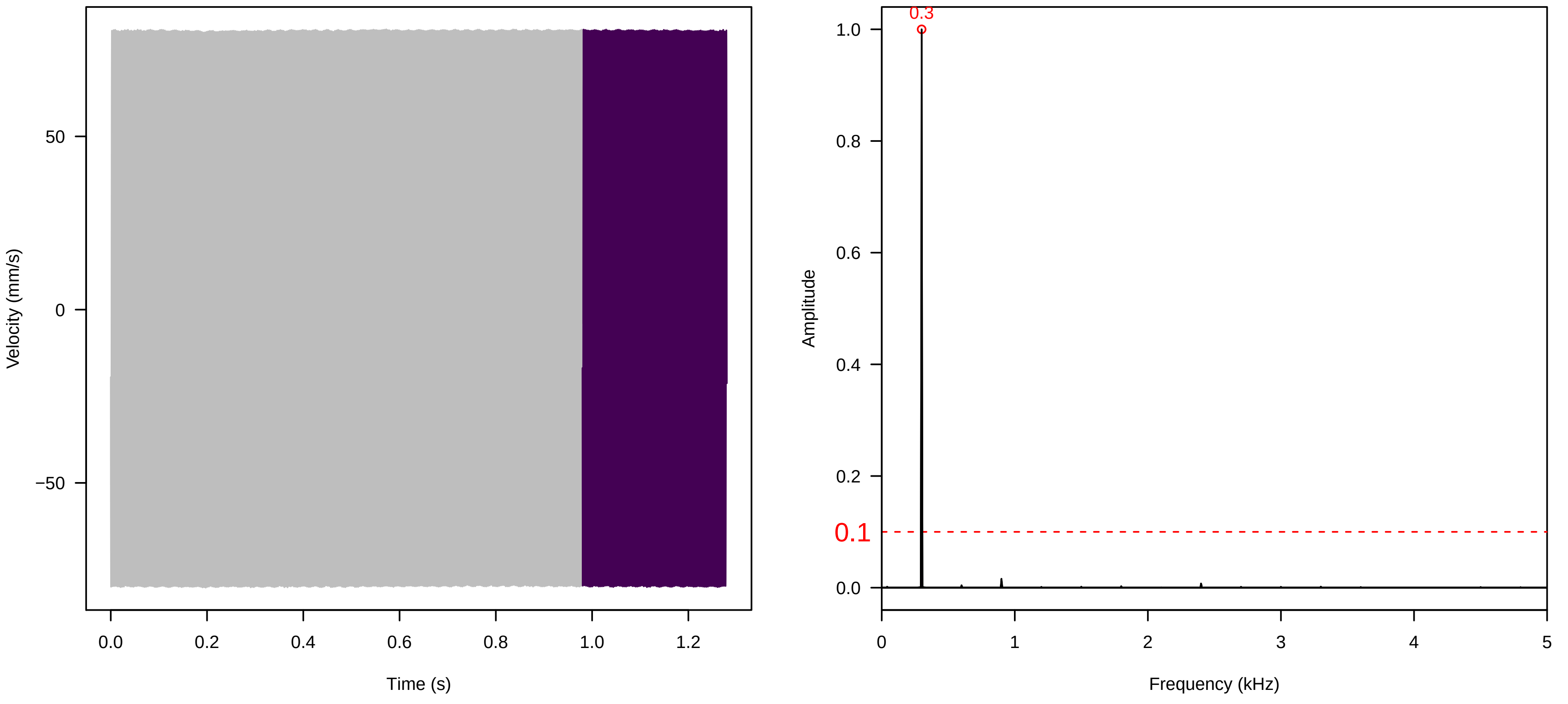


Vel. = 0.014 ; Str. = Corolla ; Axis = x ; Fl. accession = 10−s−79−27

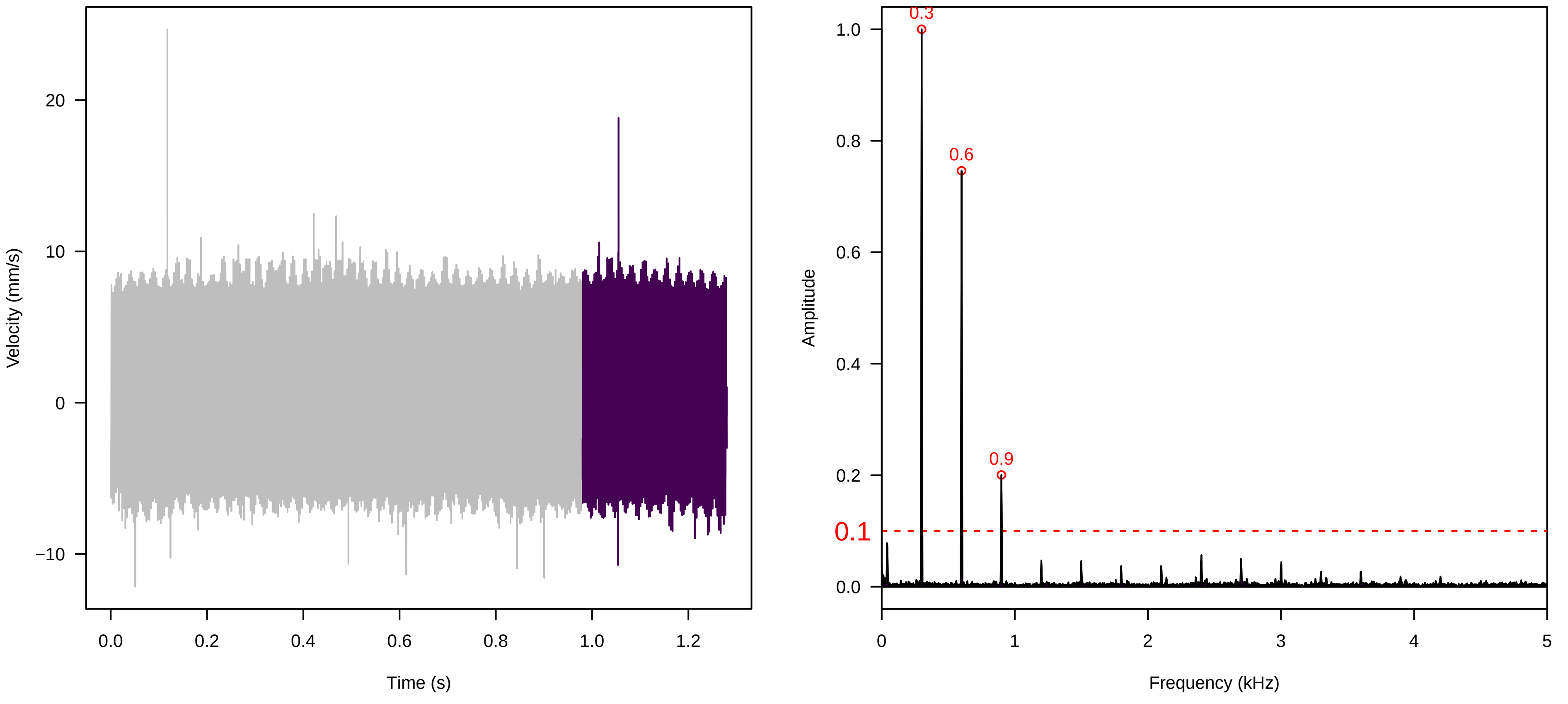

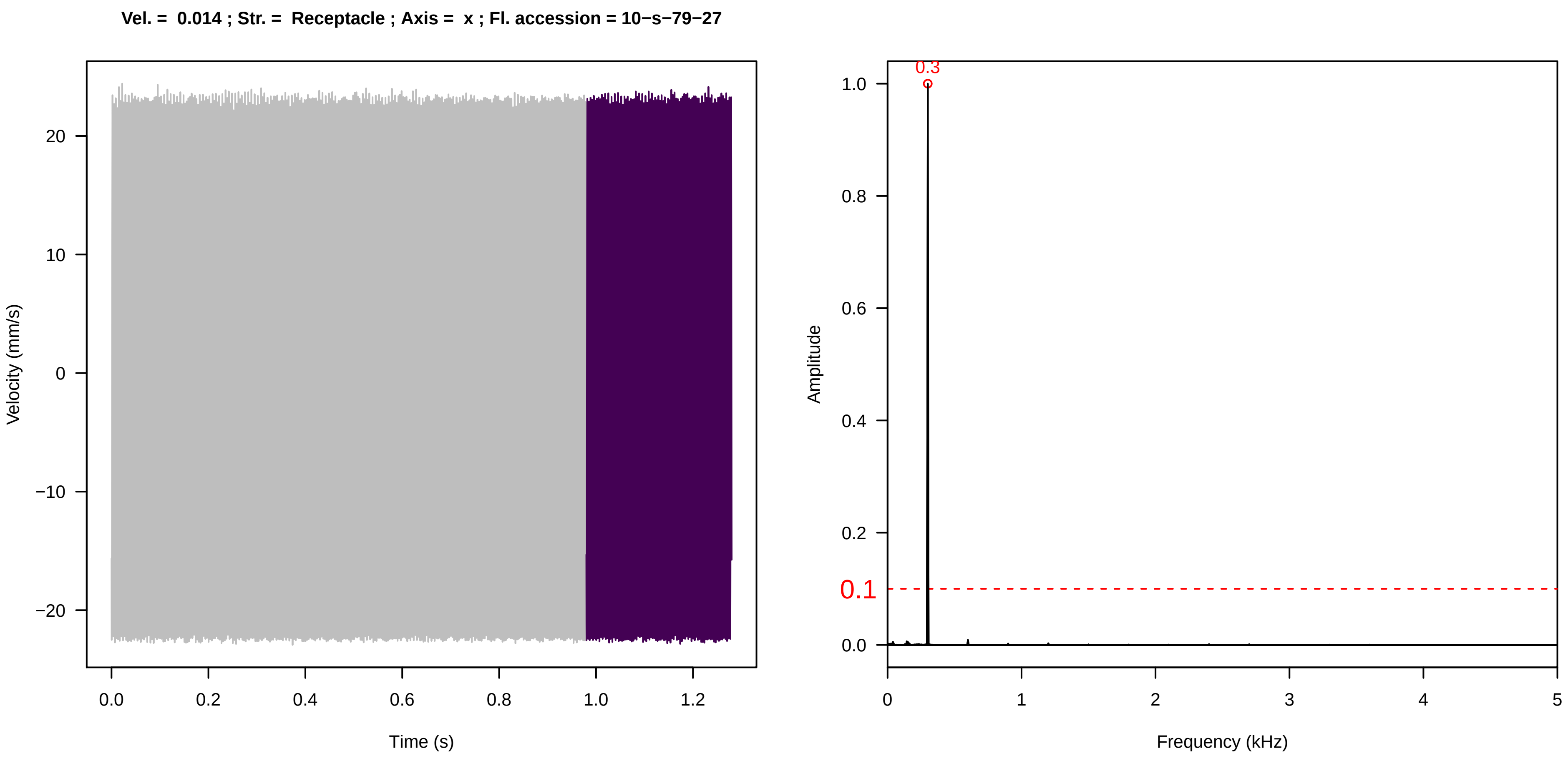
Vel. = 0.014 ; Str. = Receptacle ; Axis = x ; Fl. accession = 10−s−79−27
Velocity (mm/s)
Time (s)
Amplitude
Frequency (kHz)
0.3
0.1

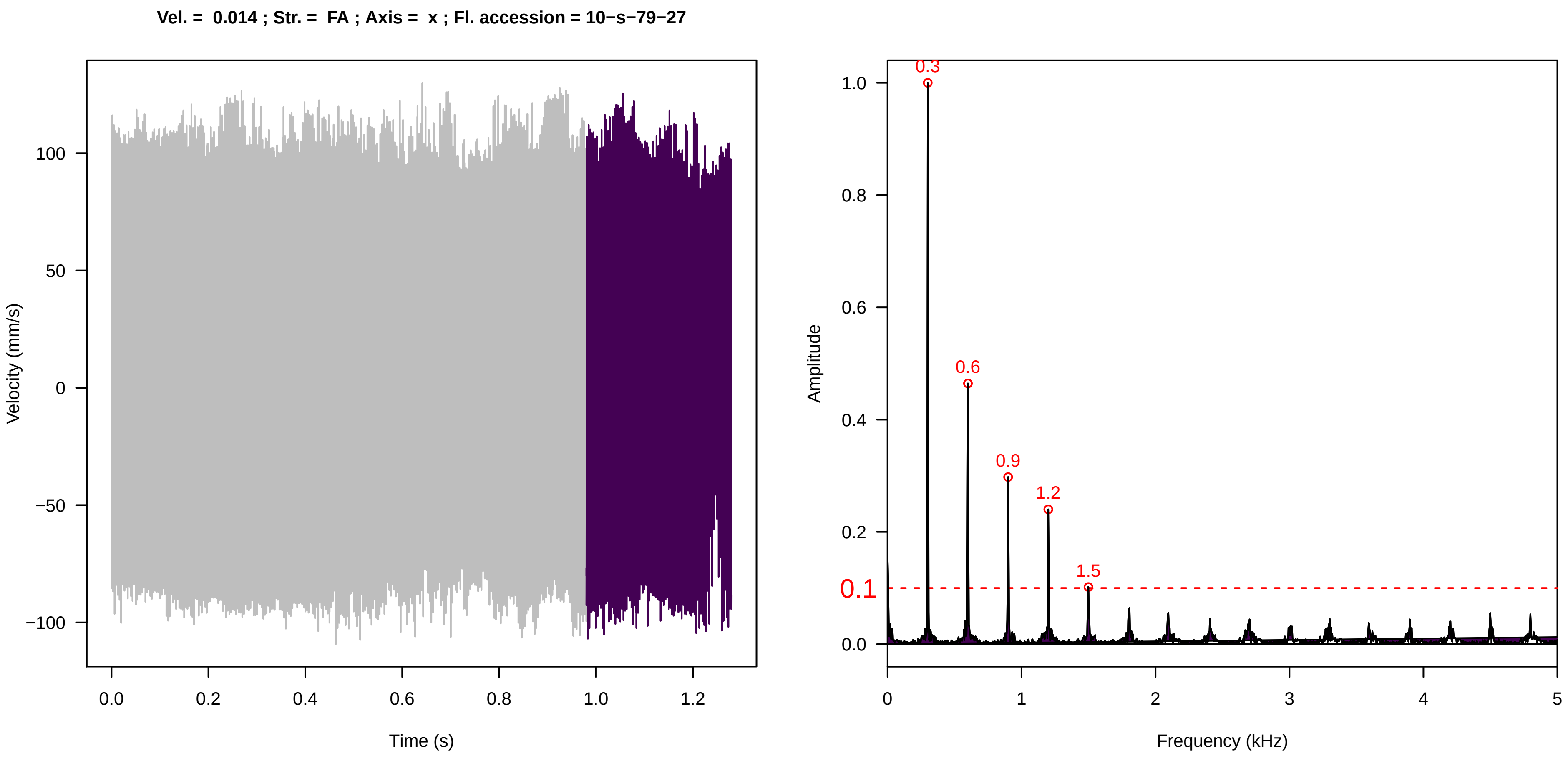
Vel. = 0.014 ; Str. = FA ; Axis = x ; Fl. accession = 10−s−79−27
Velocity (mm/s)
Time (s)
Amplitude
Frequency (kHz)
0.3
0.6
0.9
1.2
1.5
0.1

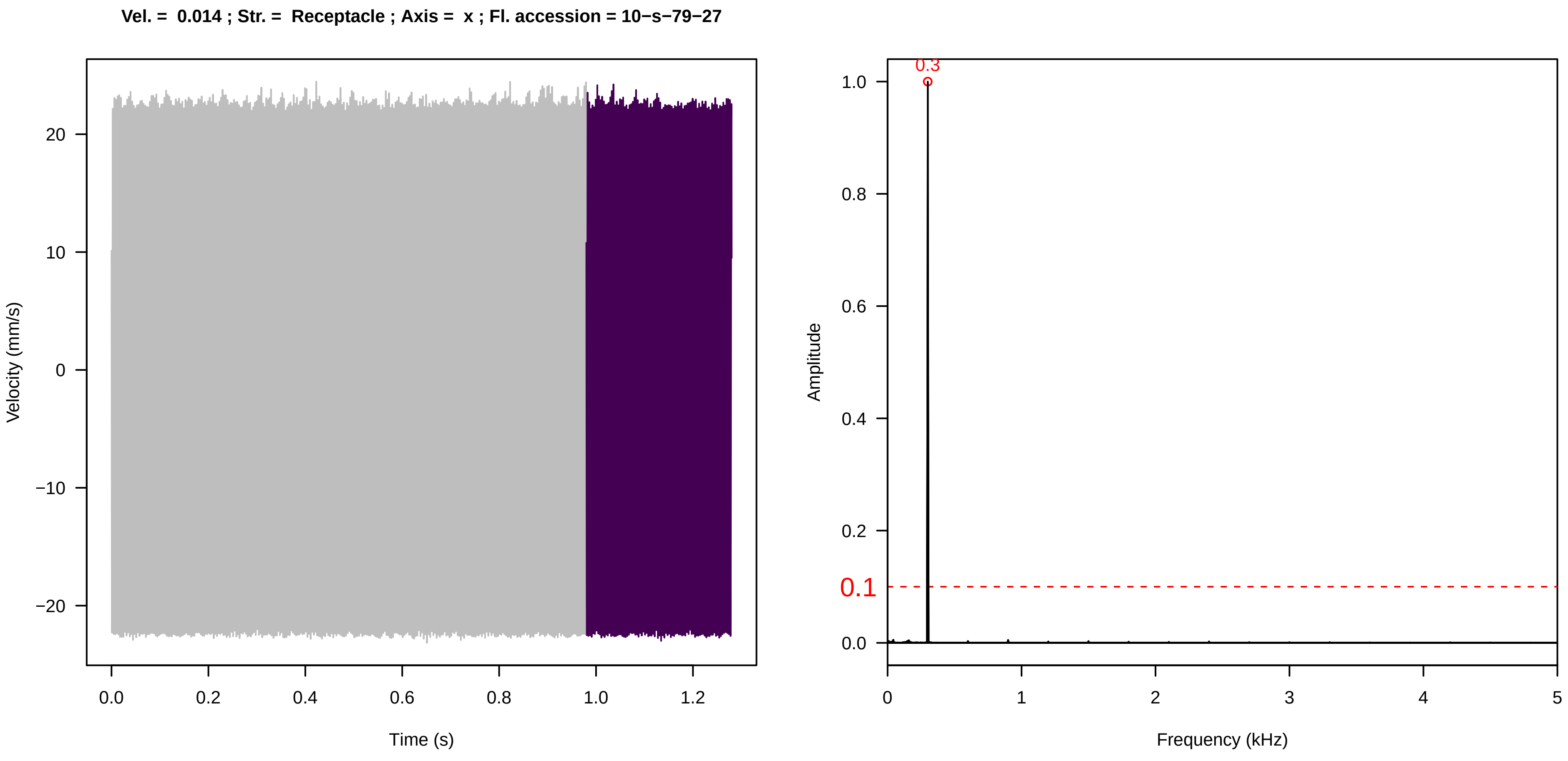
Vel. = 0.014 ; Str. = Receptacle ; Axis = x ; Fl. accession = 10−s−79−27
Velocity (mm/s)
Time (s)
Amplitude
Frequency (kHz)
0.3
0.1

**Vel. =  0.014 ; Str. =  PA ; Axis =  x ; Fl. accession = 10−s−79−27**

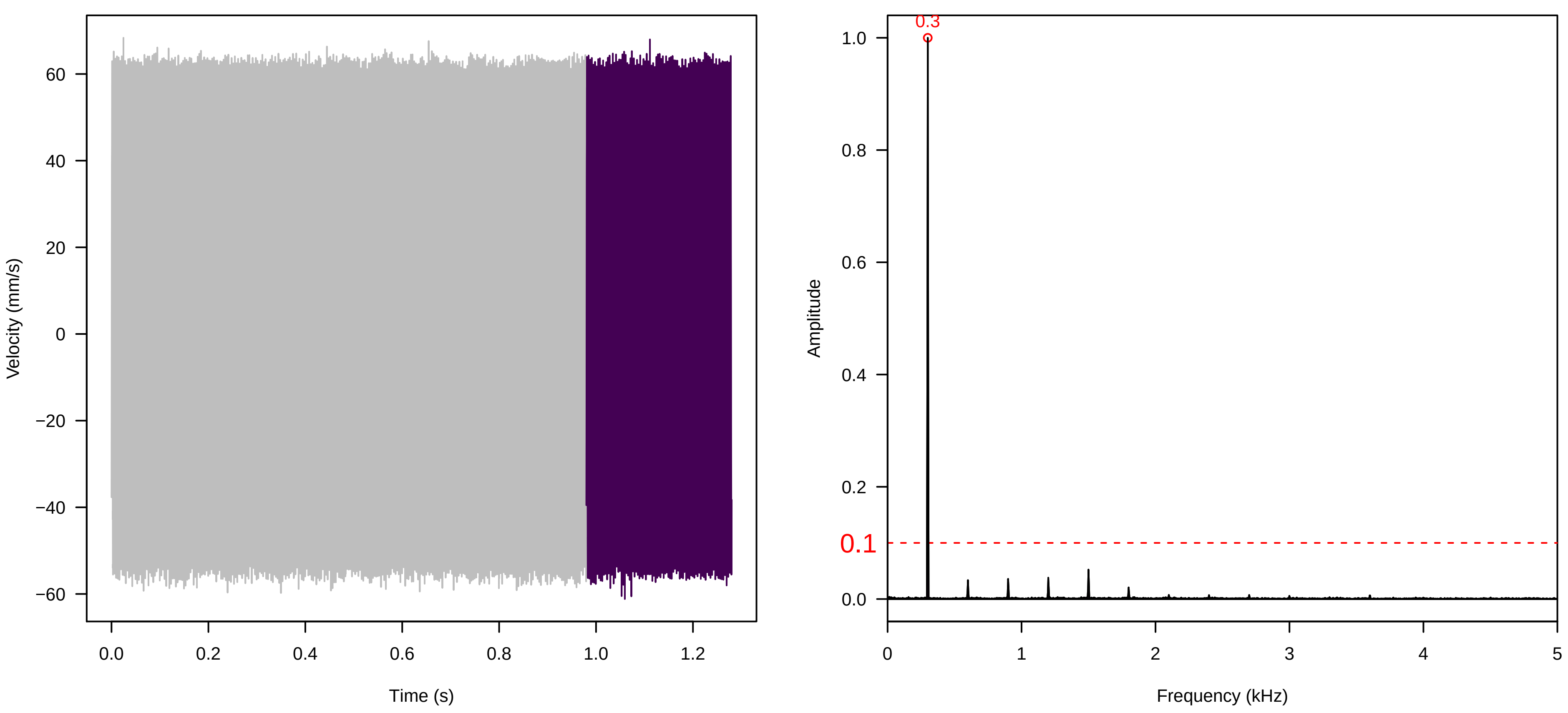


**Vel. =  0.014 ; Str. =  Receptacle ; Axis =  x ; Fl. accession = 10−s−79−27**

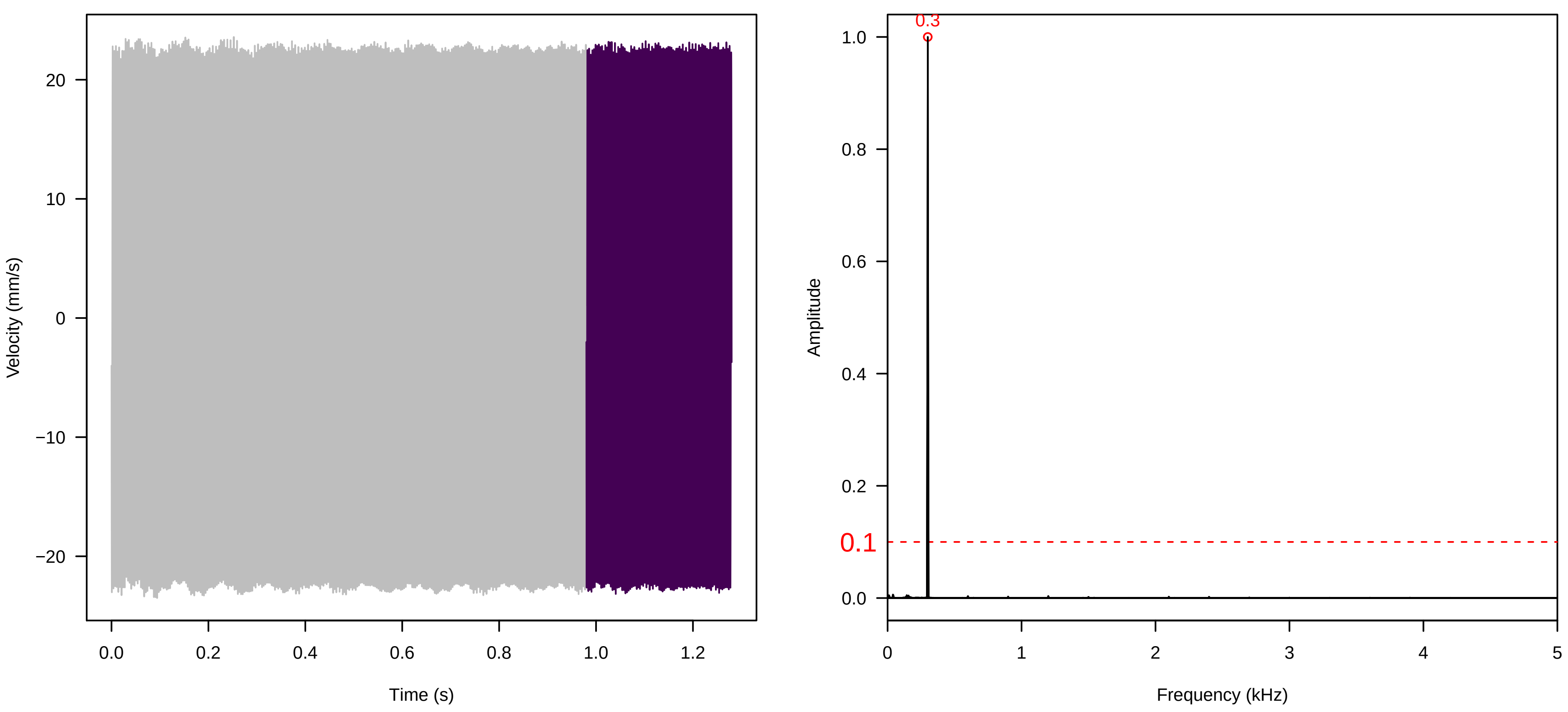


**Vel. =  0.028 ; Str. =  PA ; Axis =  x ; Fl. accession = 10−s−79−27**

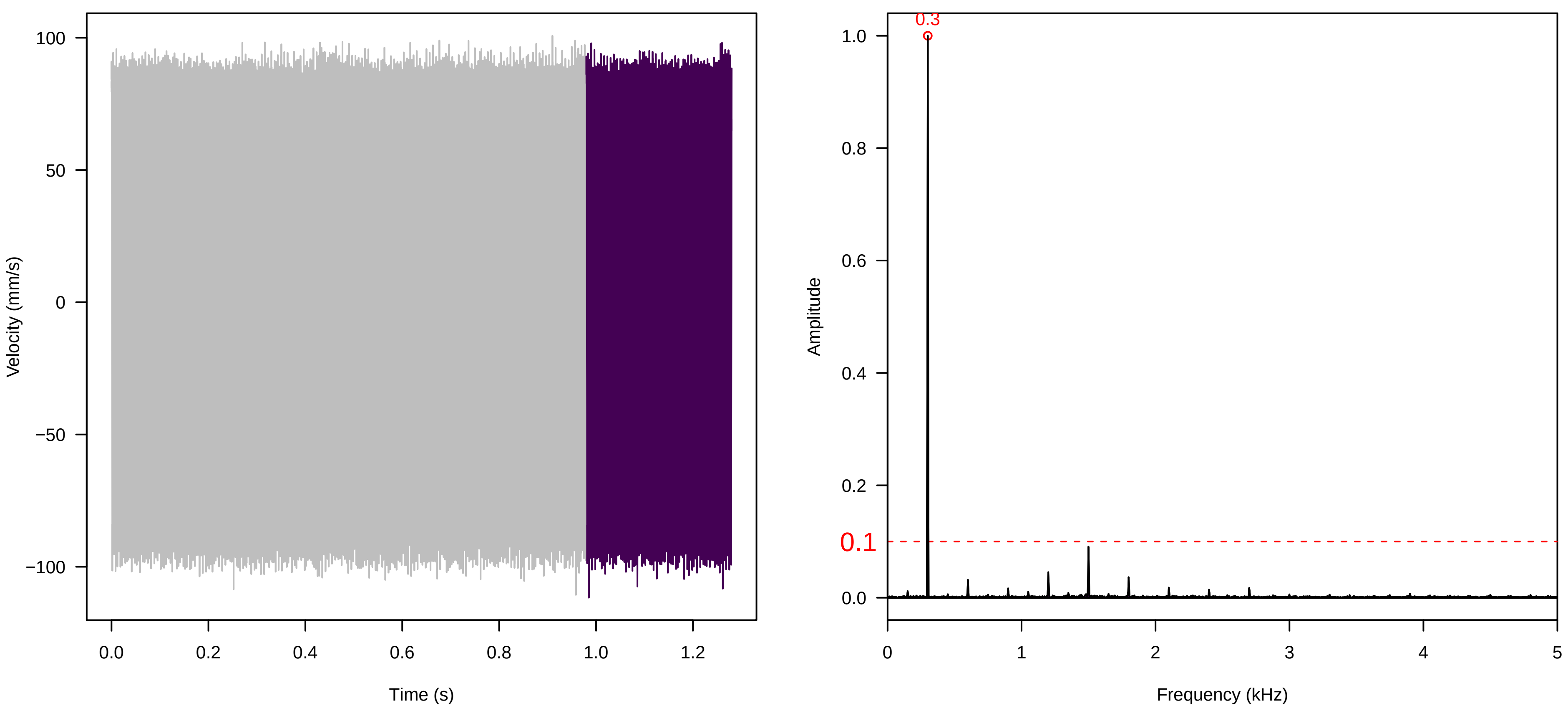

**Vel. = 0.028 ; Str. = Receptacle ; Axis = x ; Fl. accession = 10−s−79−27**

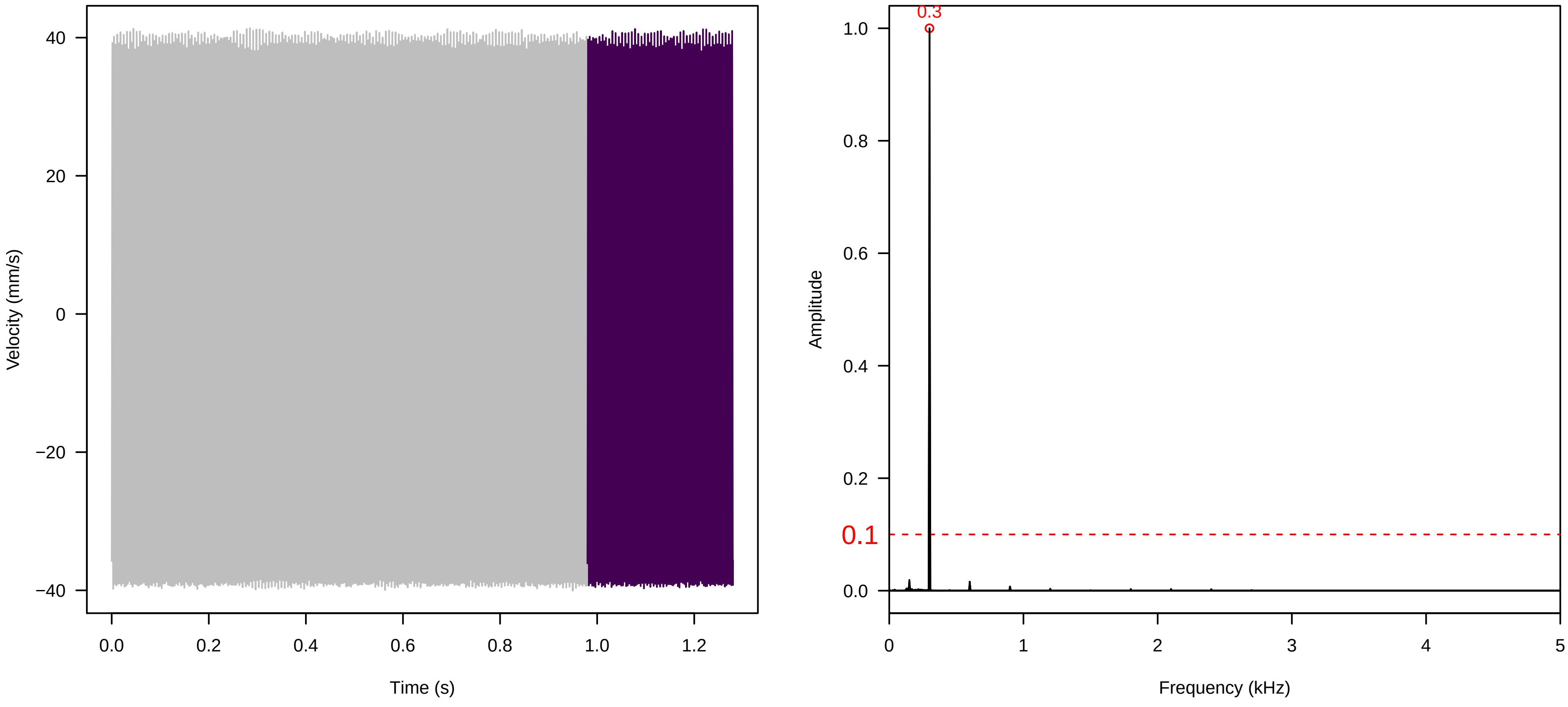


**Vel. = 0.028 ; Str. = FA ; Axis = x ; Fl. accession = 10−s−79−27**

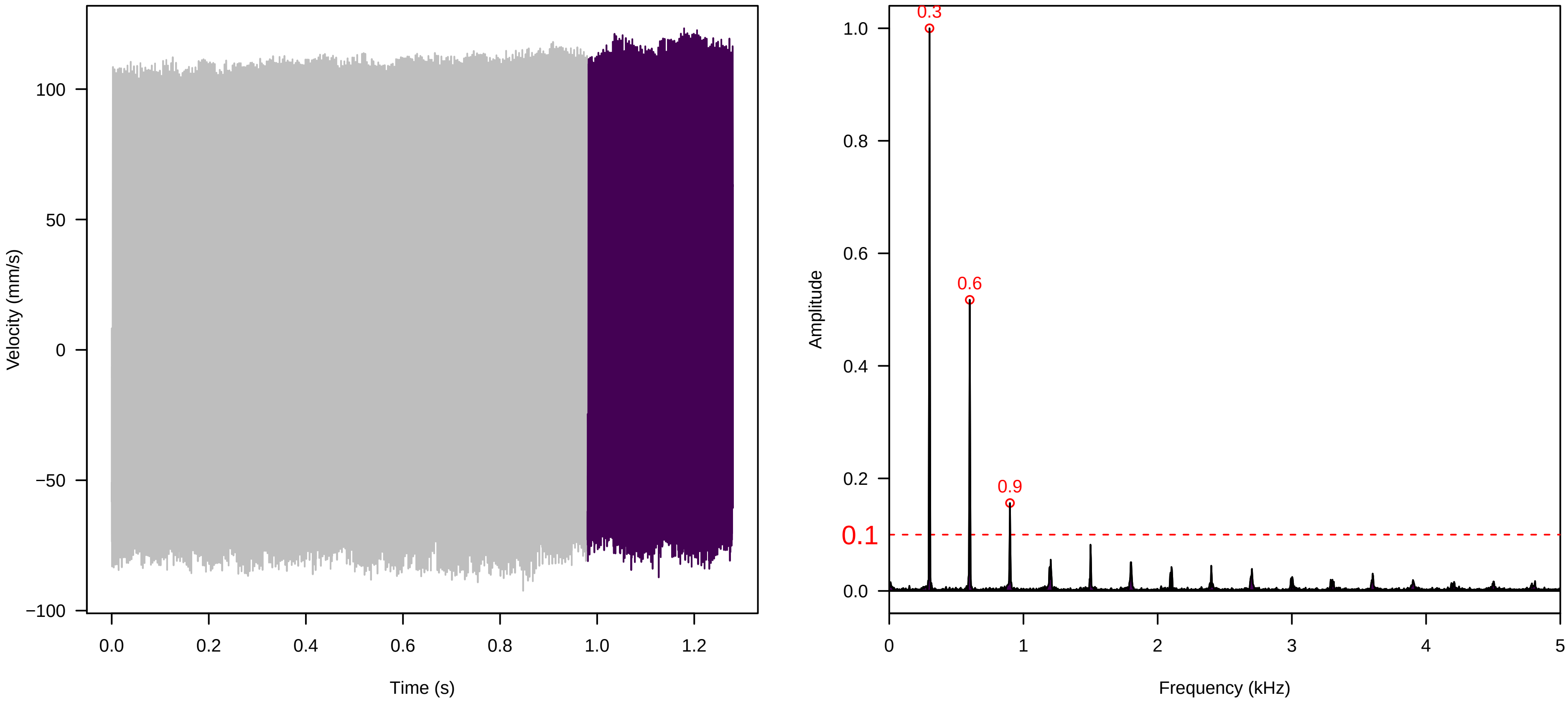


**Vel. = 0.028 ; Str. = Receptacle ; Axis = x ; Fl. accession = 10−s−79−27**

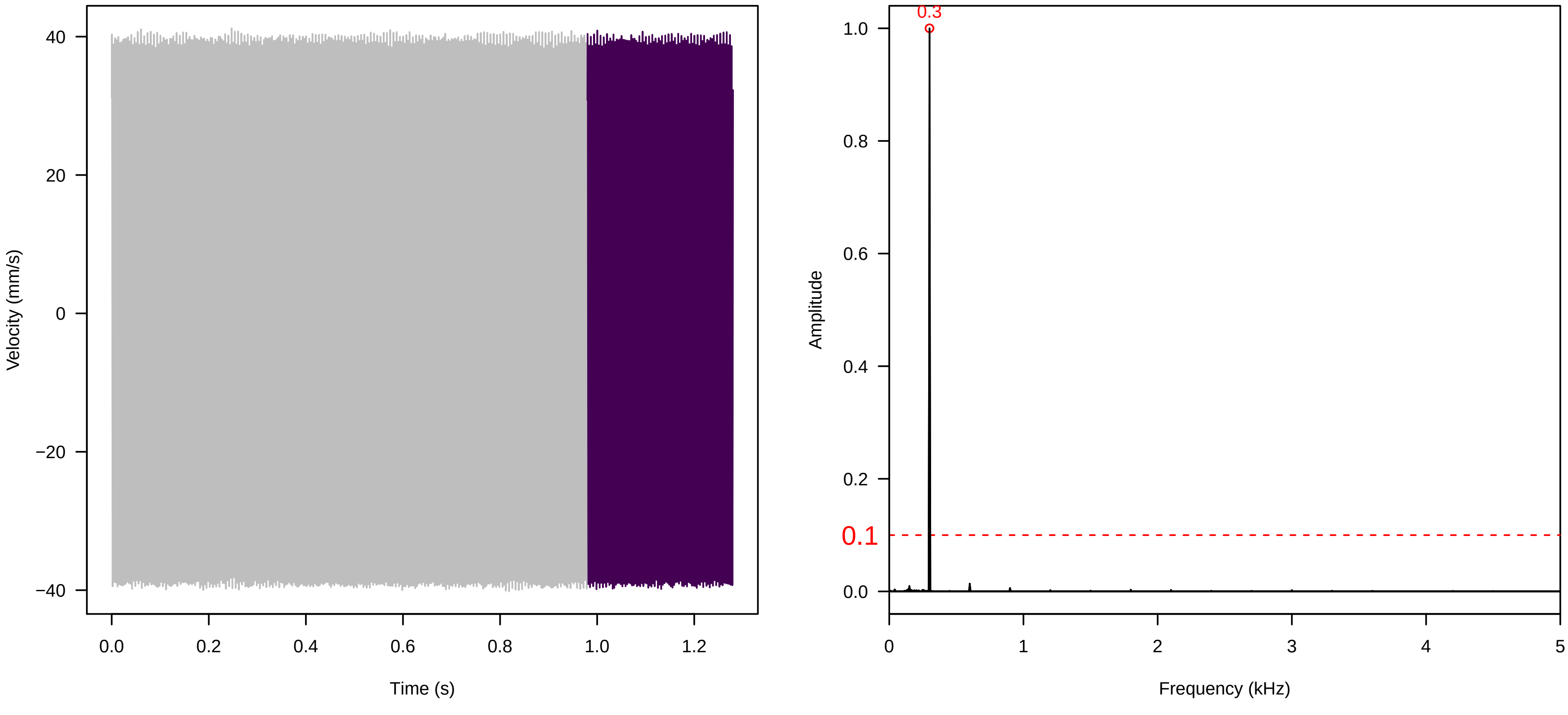

Vel. = 0.028 ; Str. = Corolla ; Axis = x ; Fl. accession = 10−s−79−27

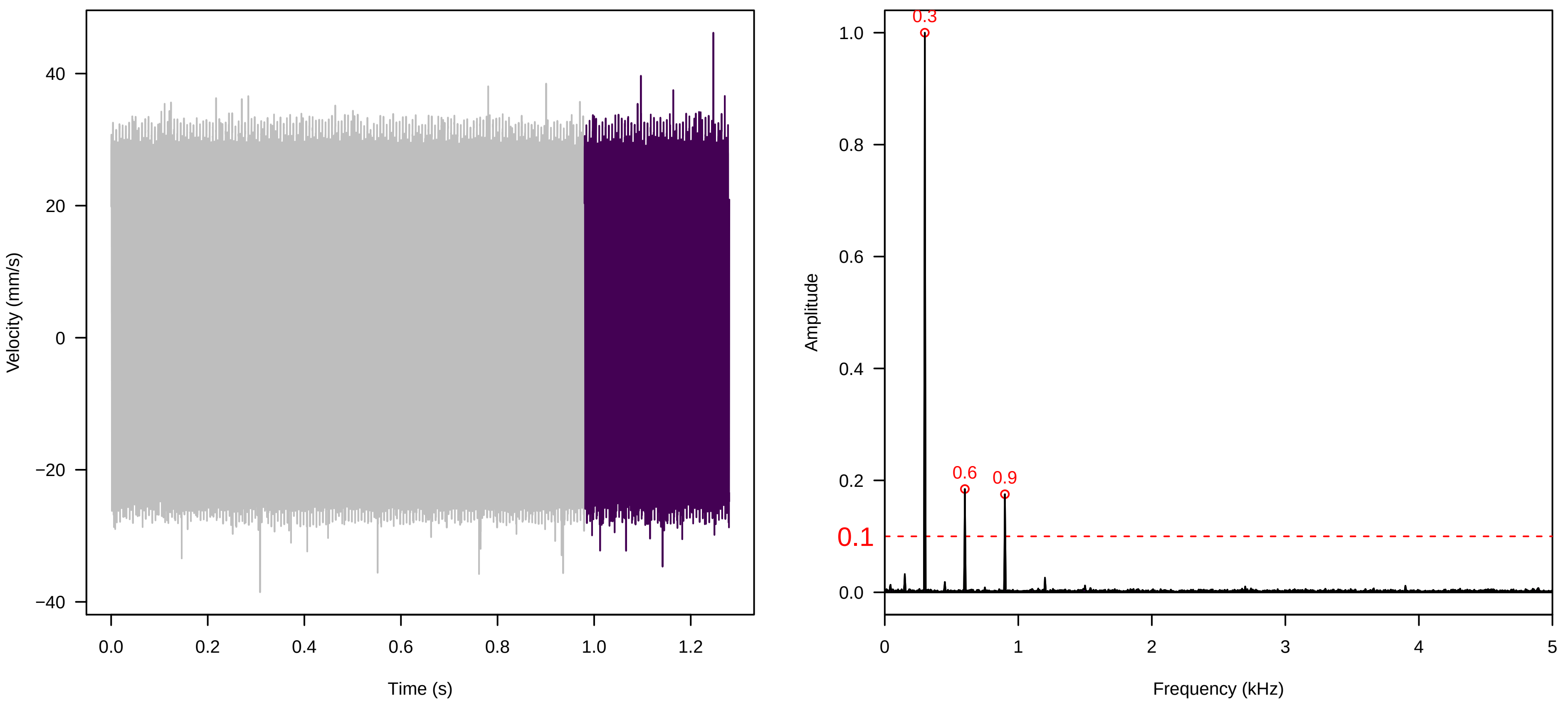


Vel. = 0.028 ; Str. = Receptacle ; Axis = x ; Fl. accession = 10−s−79−27

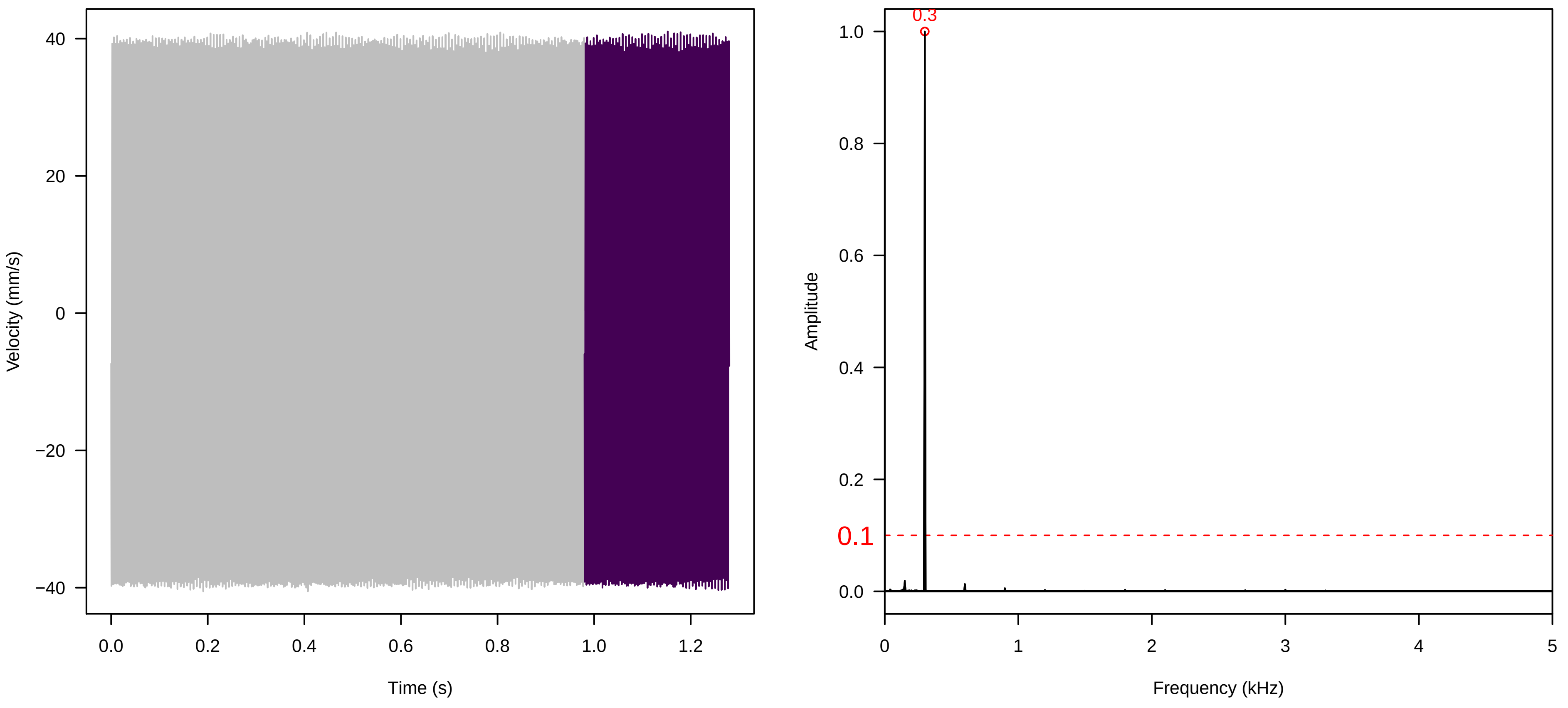


Vel. = 0.057 ; Str. = Corolla ; Axis = x ; Fl. accession = 10−s−79−27

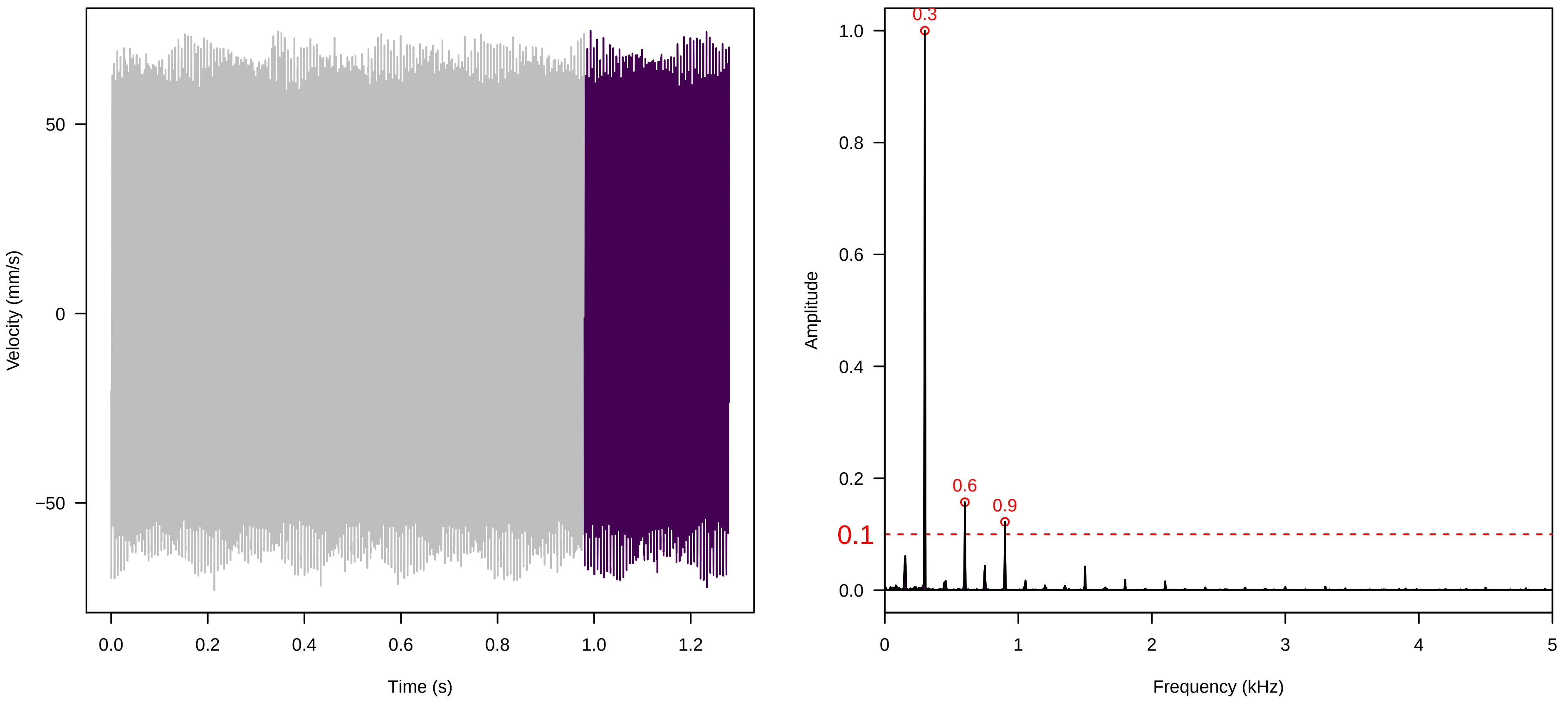

Vel. = 0.057 ; Str. = Receptacle ; Axis = x ; Fl. accession = 10−s−79−27

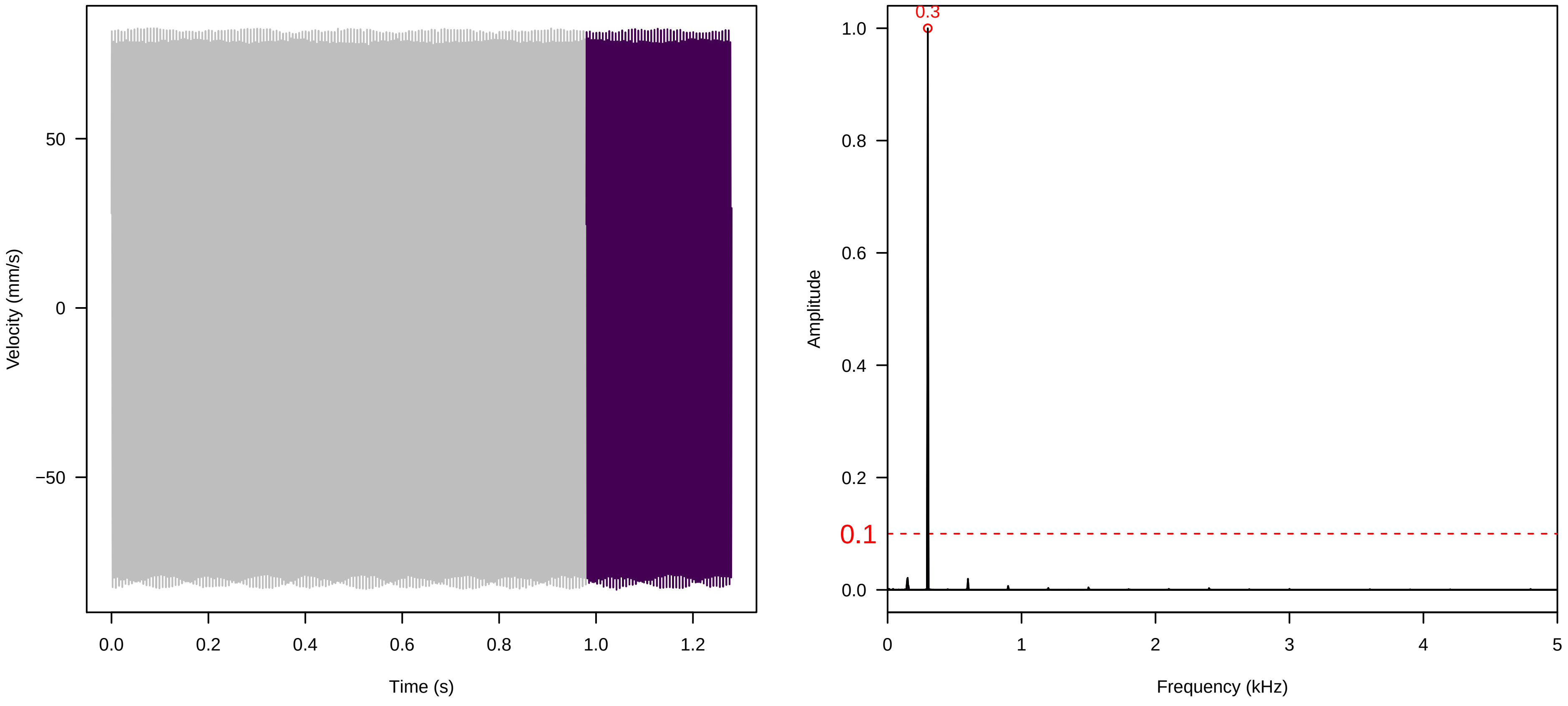


Vel. = 0.057 ; Str. = FA ; Axis = x ; Fl. accession = 10−s−79−27

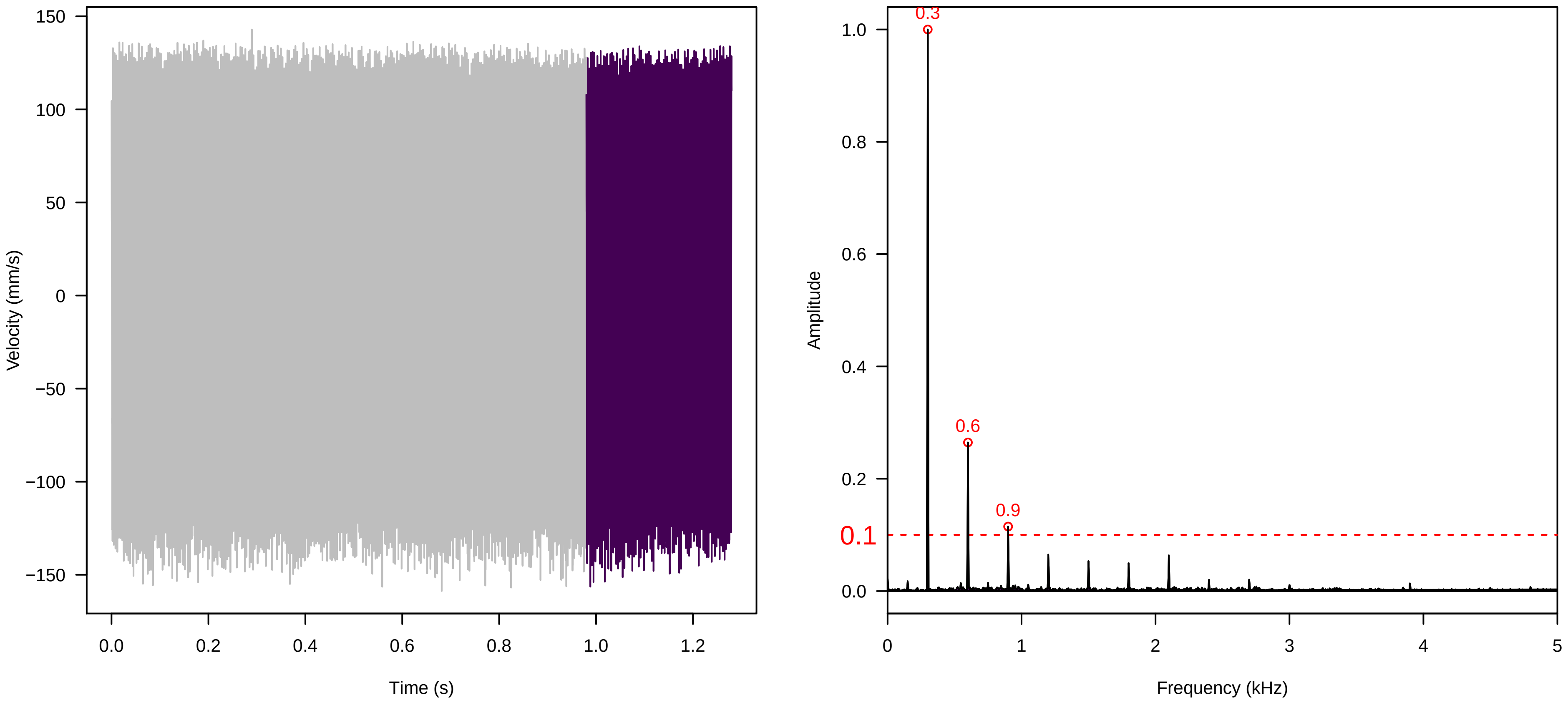


Vel. = 0.057 ; Str. = Receptacle ; Axis = x ; Fl. accession = 10−s−79−27

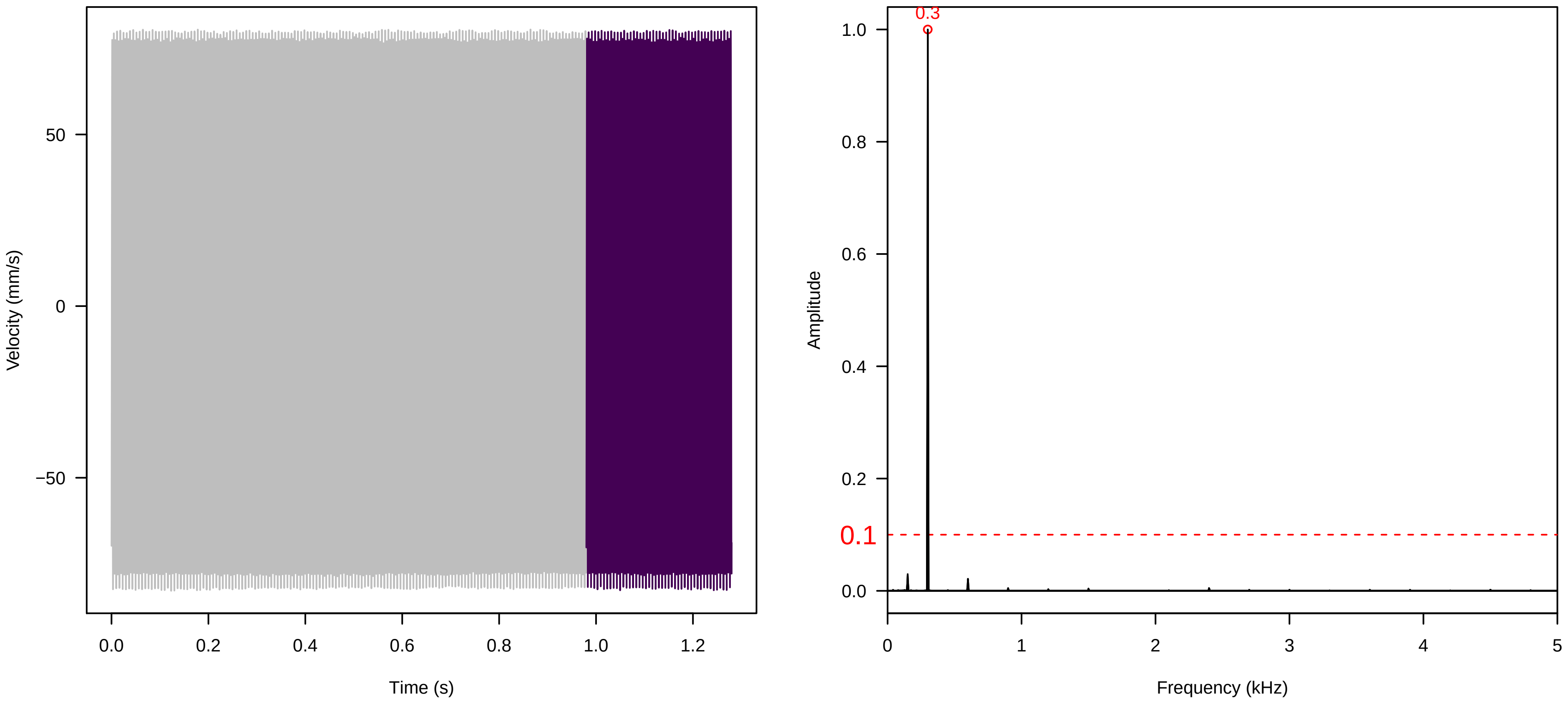

**Vel. = 0.057 ; Str. = PA ; Axis = x ; Fl. accession = 10−s−79−27**

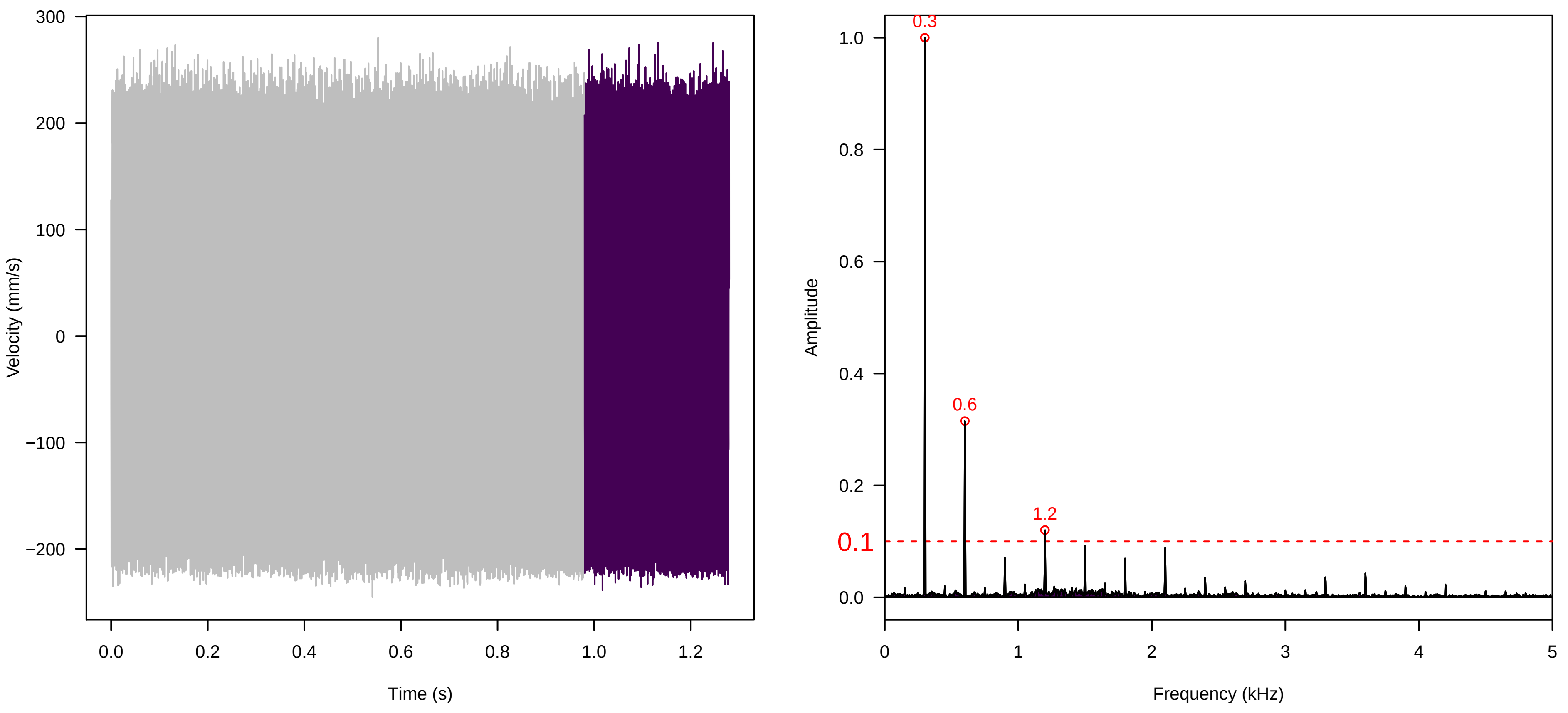


**Vel. = 0.057 ; Str. = Receptacle ; Axis = x ; Fl. accession = 10−s−79−27**

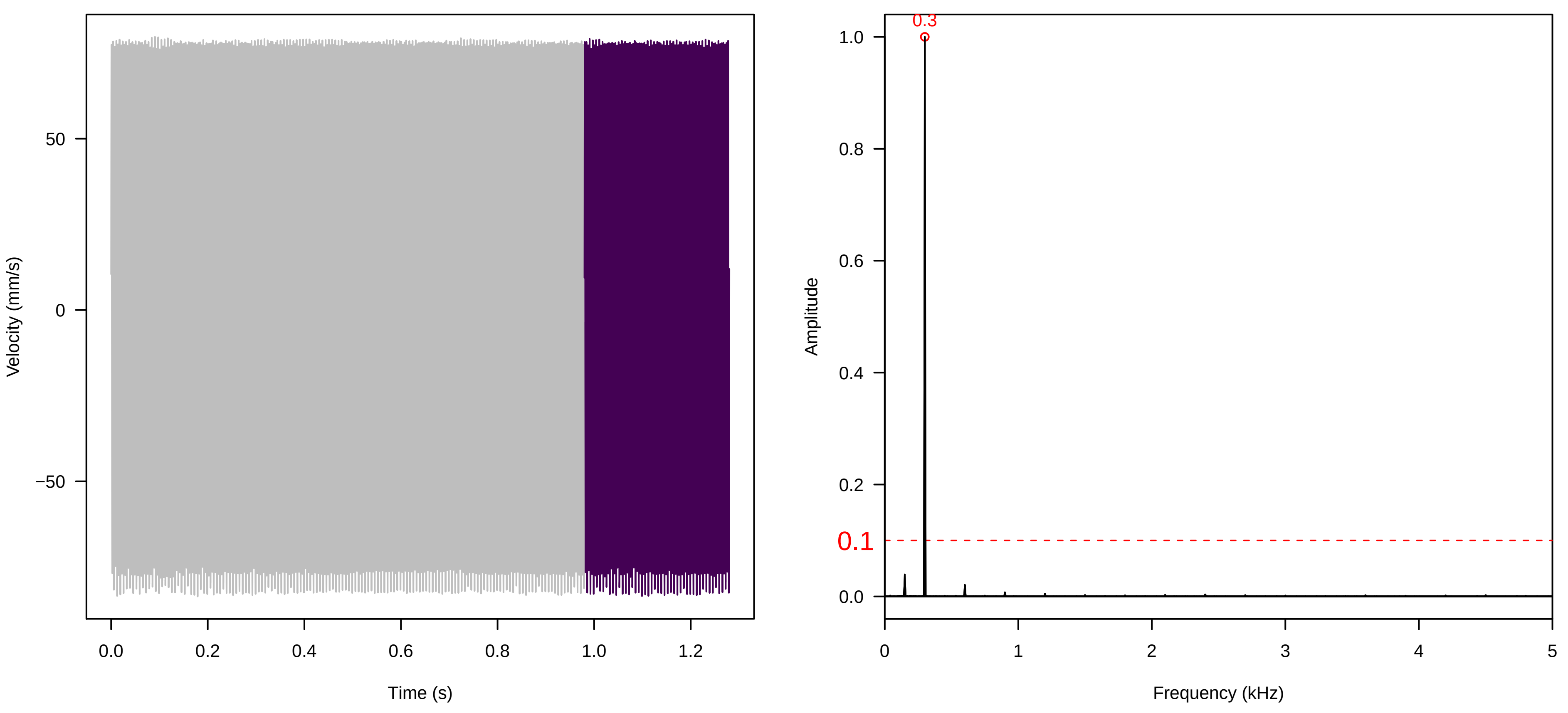


**Vel. = 0.057 ; Str. = Corolla ; Axis = z ; Fl. accession = 10−s−79−27**

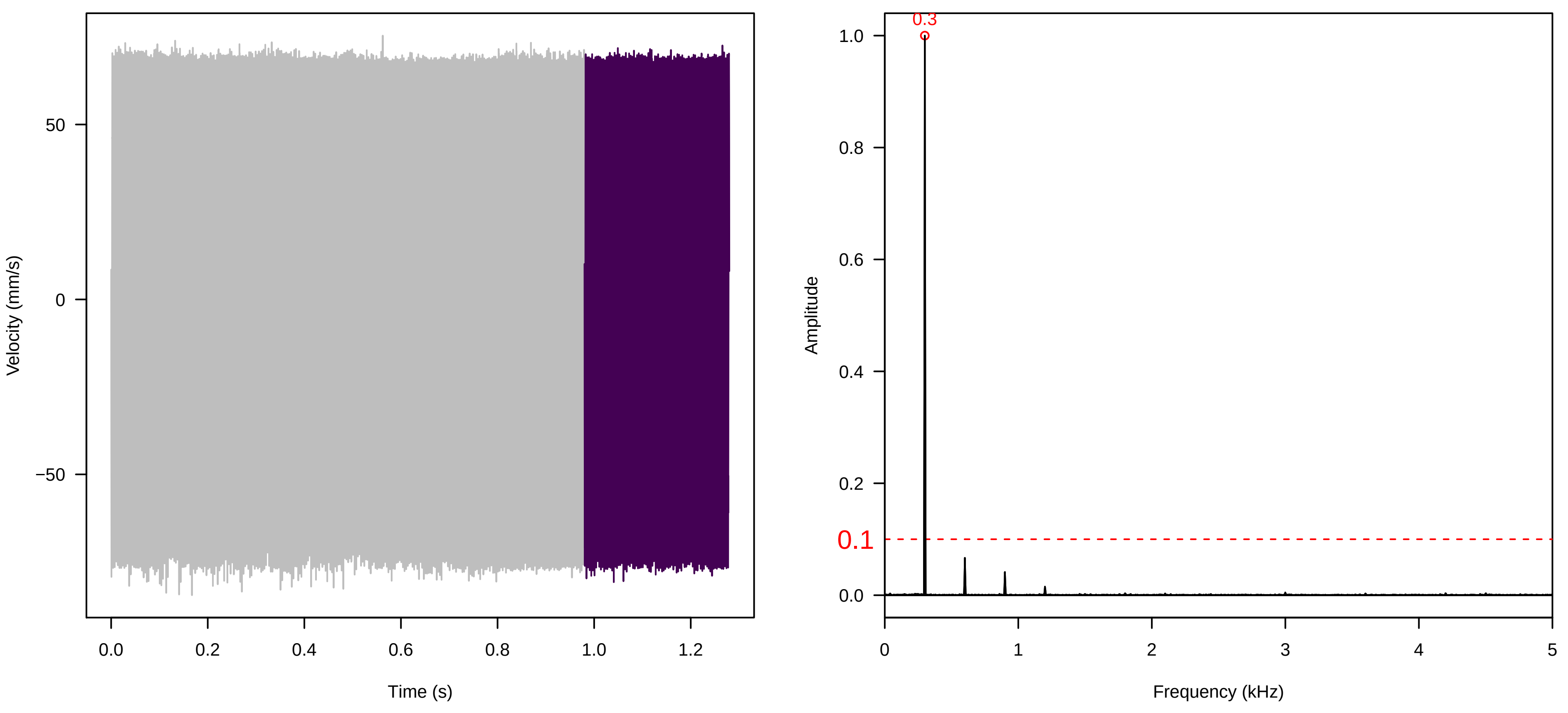

**Vel. = 0.057 ; Str. = Receptacle ; Axis = z ; Fl. accession = 10−s−79−27**

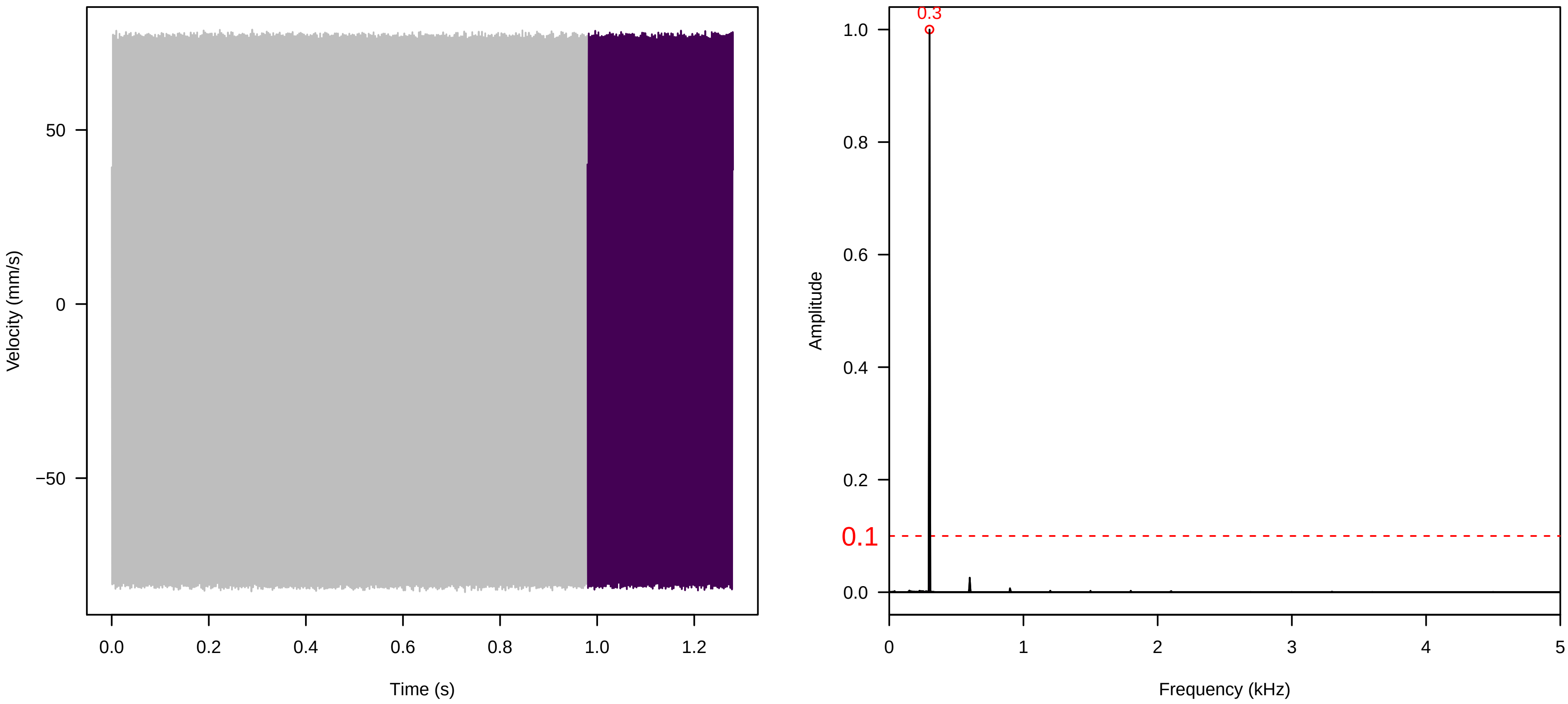


**Vel. = 0.057 ; Str. = FA ; Axis = z ; Fl. accession = 10−s−79−27**

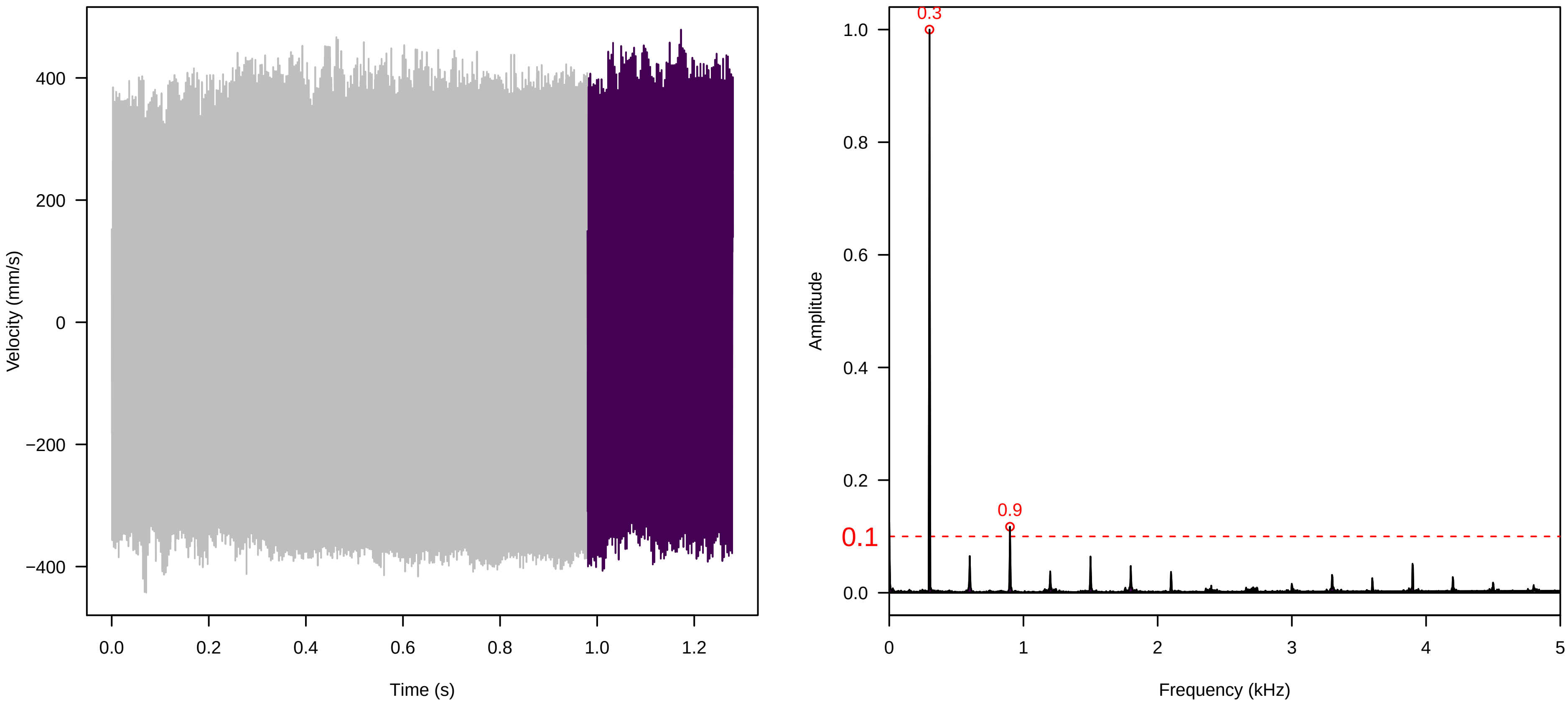


**Vel. = 0.057 ; Str. = Receptacle ; Axis = z ; Fl. accession = 10−s−79−27**

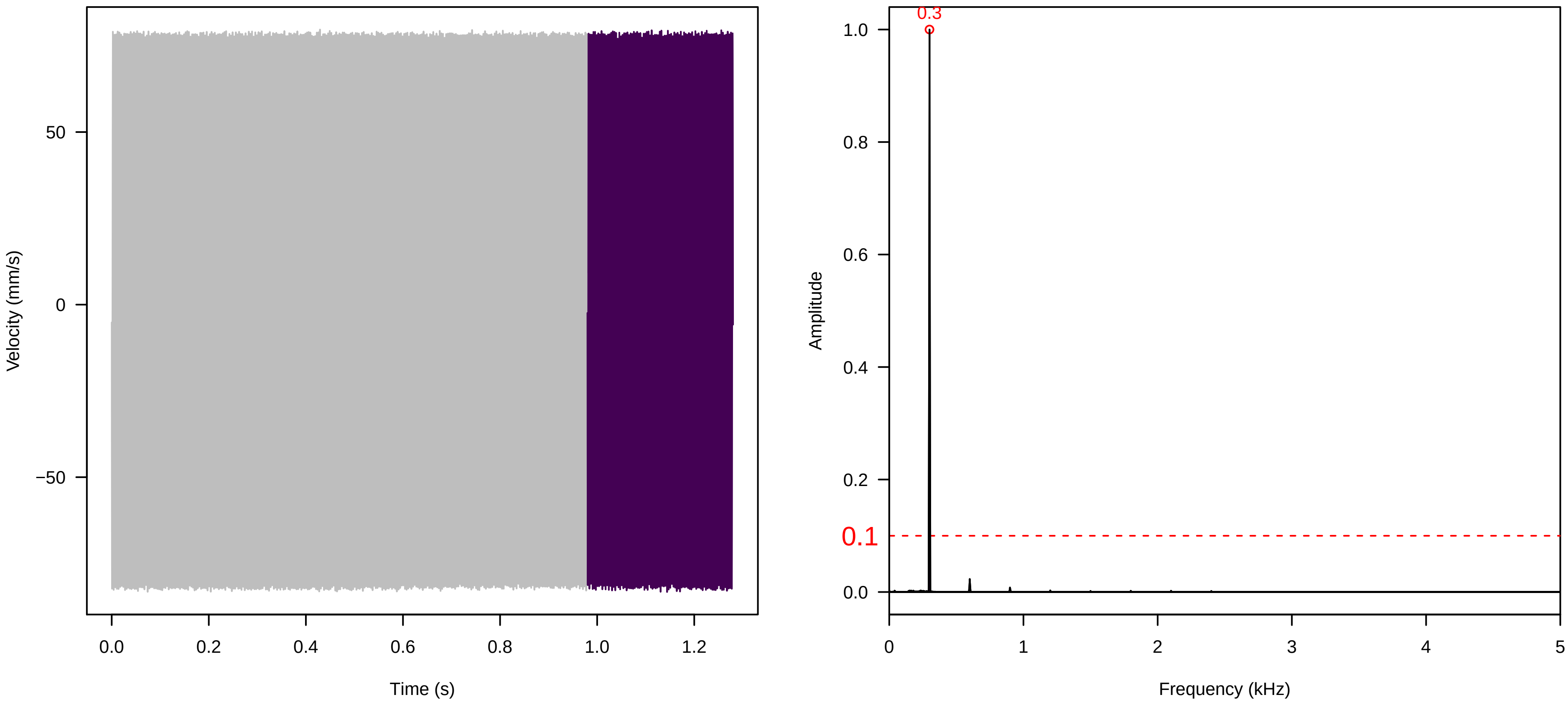

**Vel. = 0.057 ; Str. = PA ; Axis = z ; Fl. accession = 10−s−79−27**

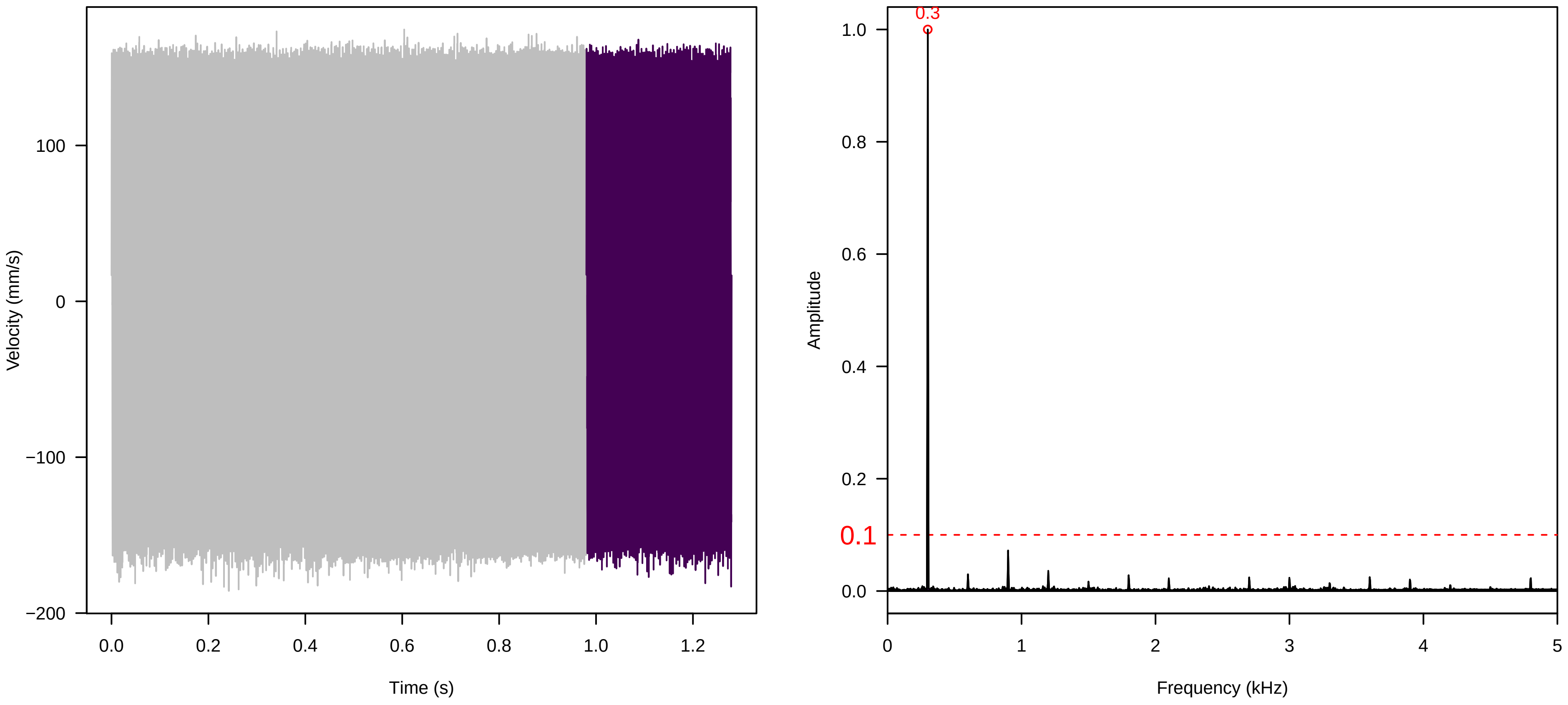


**Vel. = 0.057 ; Str. = Receptacle ; Axis = z ; Fl. accession = 10−s−79−27**

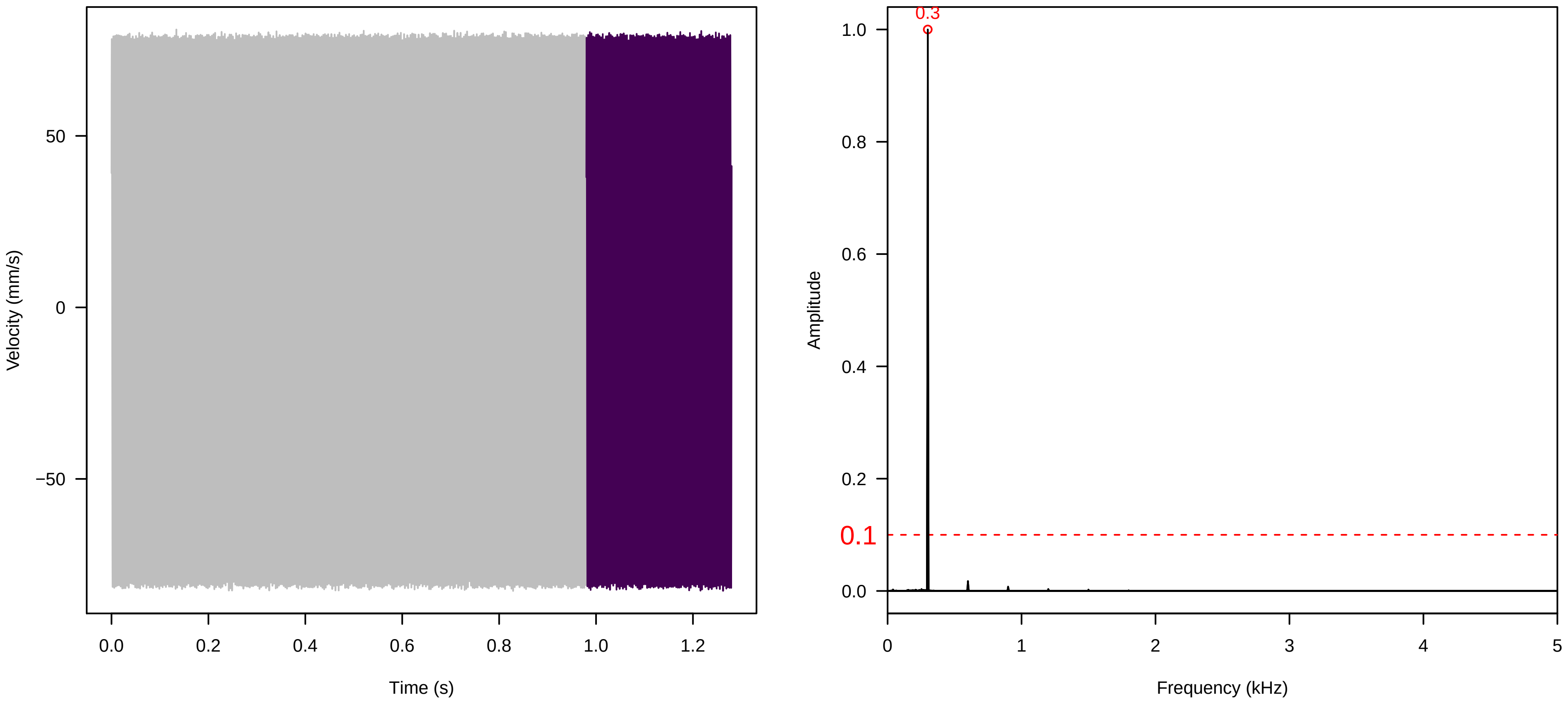


**Vel. = 0.028 ; Str. = PA ; Axis = z ; Fl. accession = 10−s−79−27**

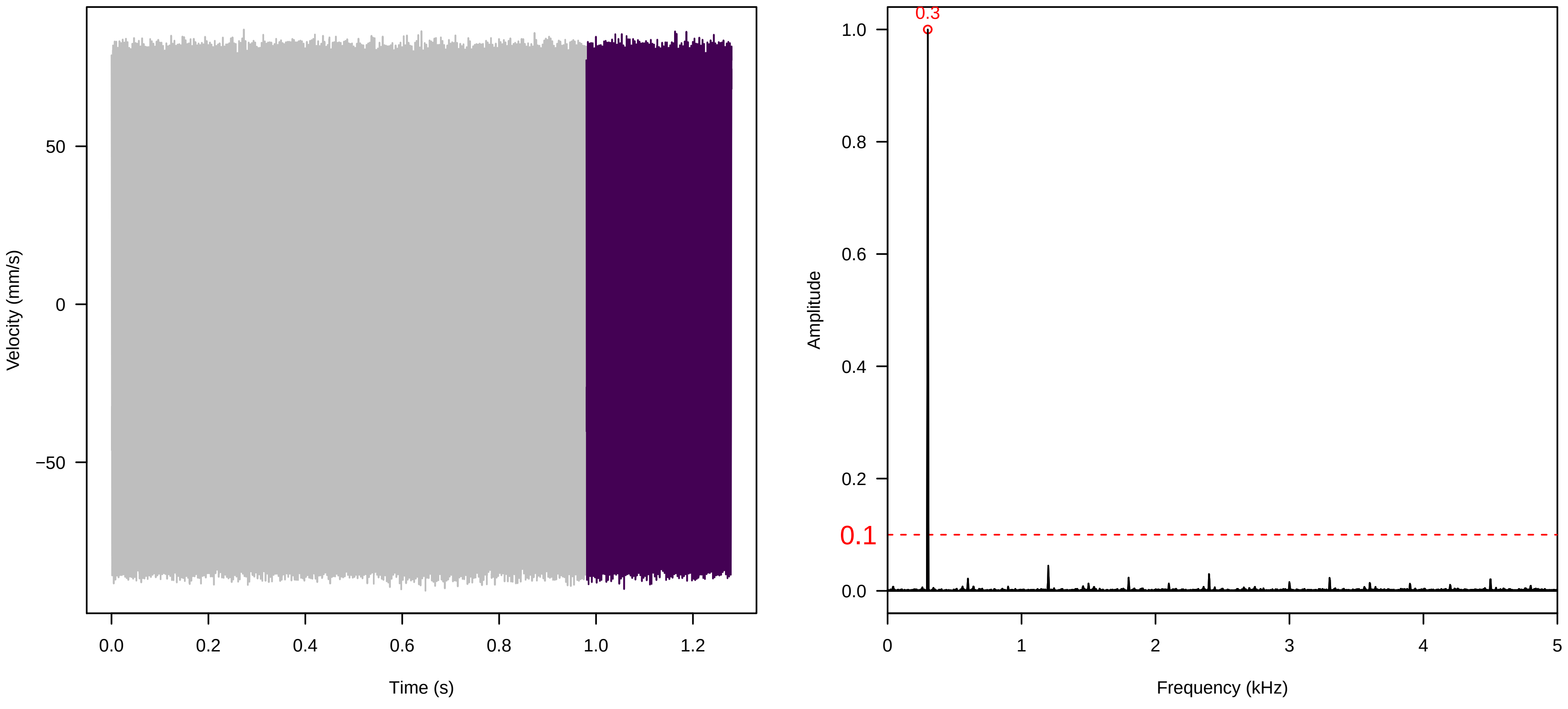

**Vel. = 0.028 ; Str. = Receptacle ; Axis = z ; Fl. accession = 10−s−79−27**

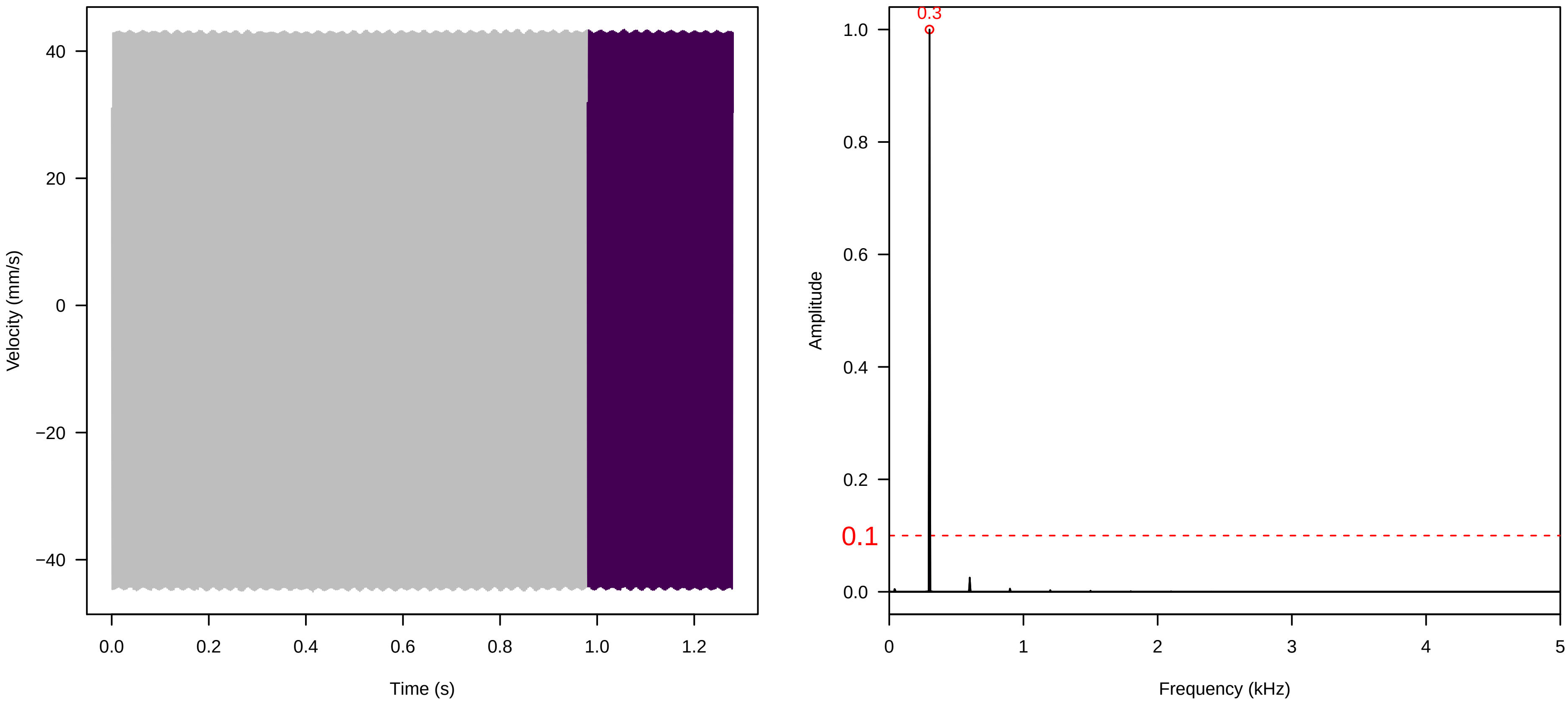


**Vel. = 0.028 ; Str. = FA ; Axis = z ; Fl. accession = 10−s−79−27**

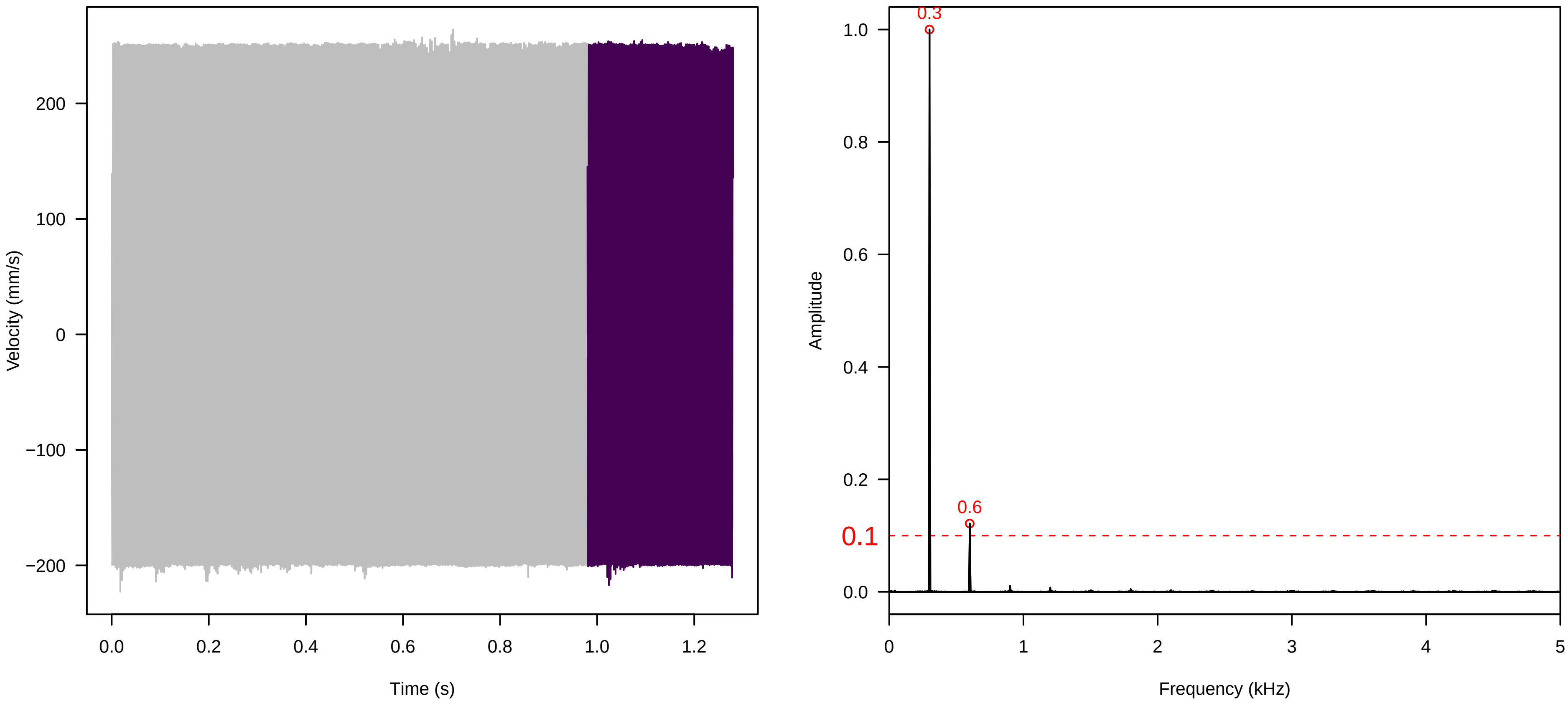


**Vel. = 0.028 ; Str. = Receptacle ; Axis = z ; Fl. accession = 10−s−79−27**

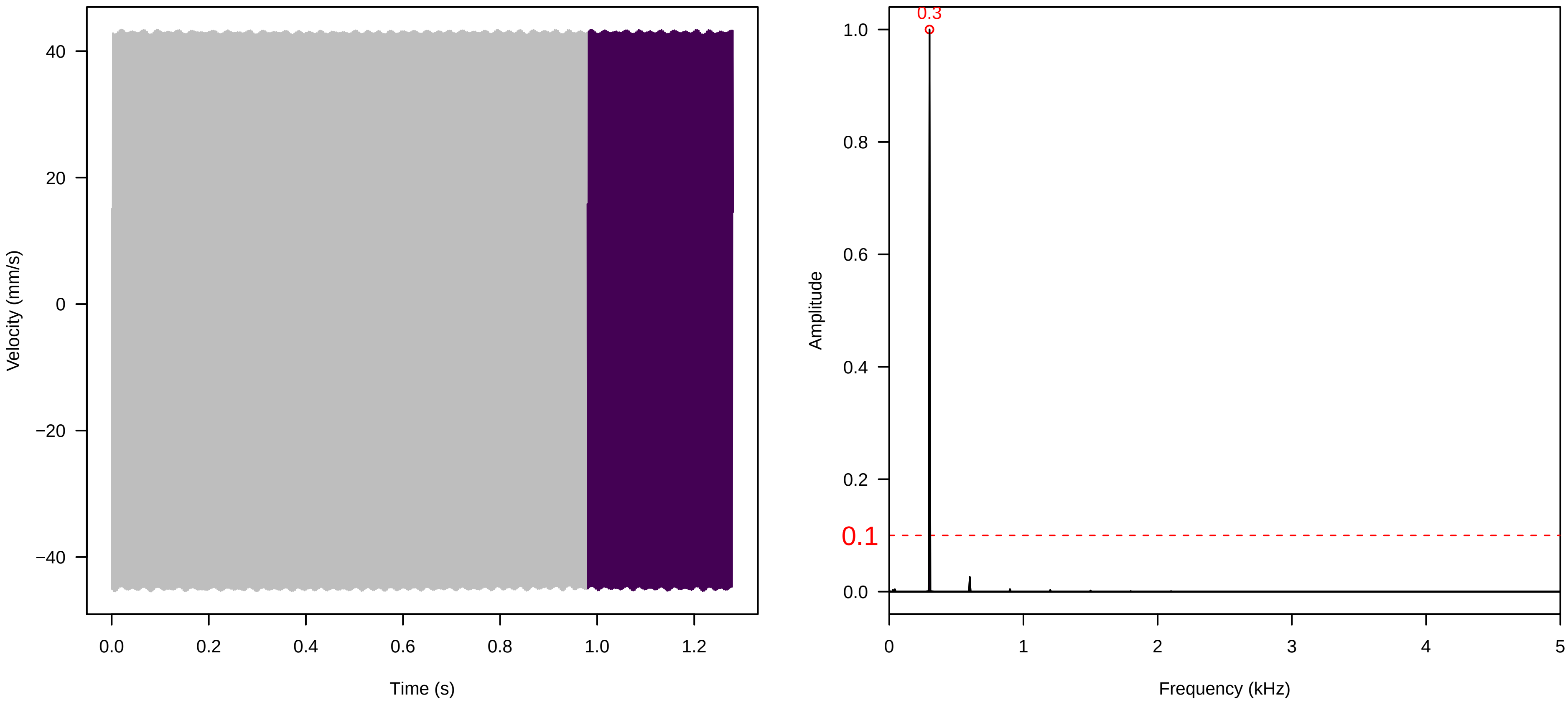

**Vel. = 0.028 ; Str. = Corolla ; Axis = z ; Fl. accession = 10−s−79−27**

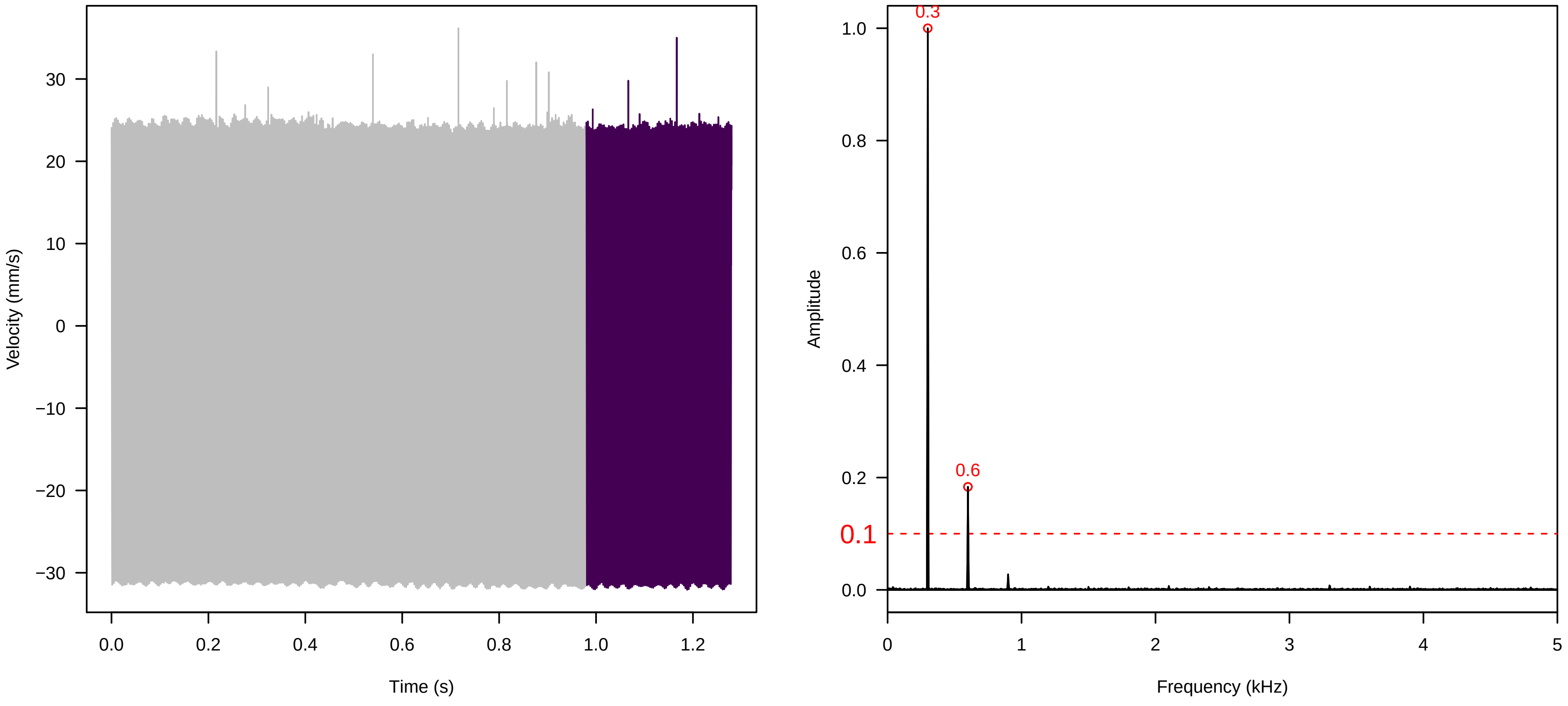


**Vel. = 0.028 ; Str. = Receptacle ; Axis = z ; Fl. accession = 10−s−79−27**

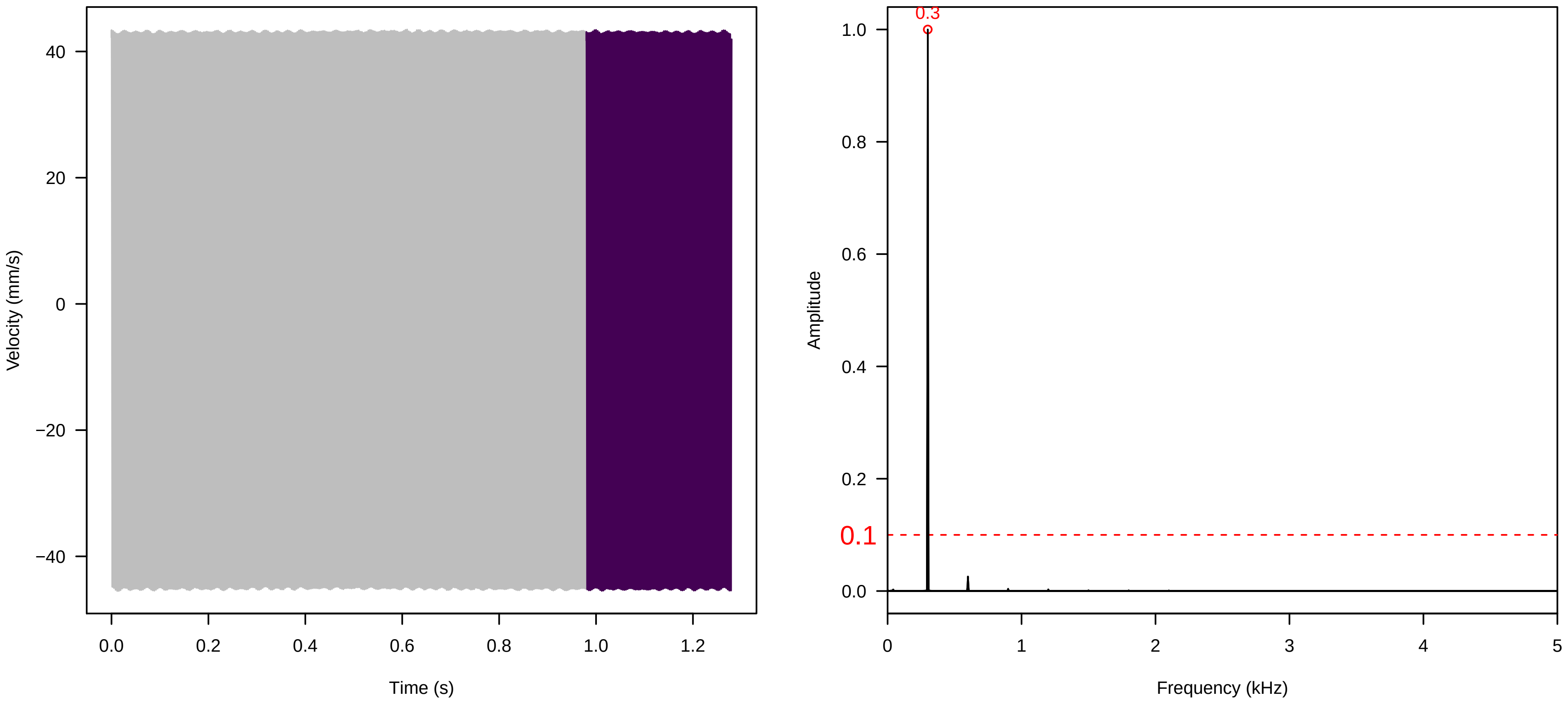


**Vel. = 0.014 ; Str. = Corolla ; Axis = z ; Fl. accession = 10−s−79−27**

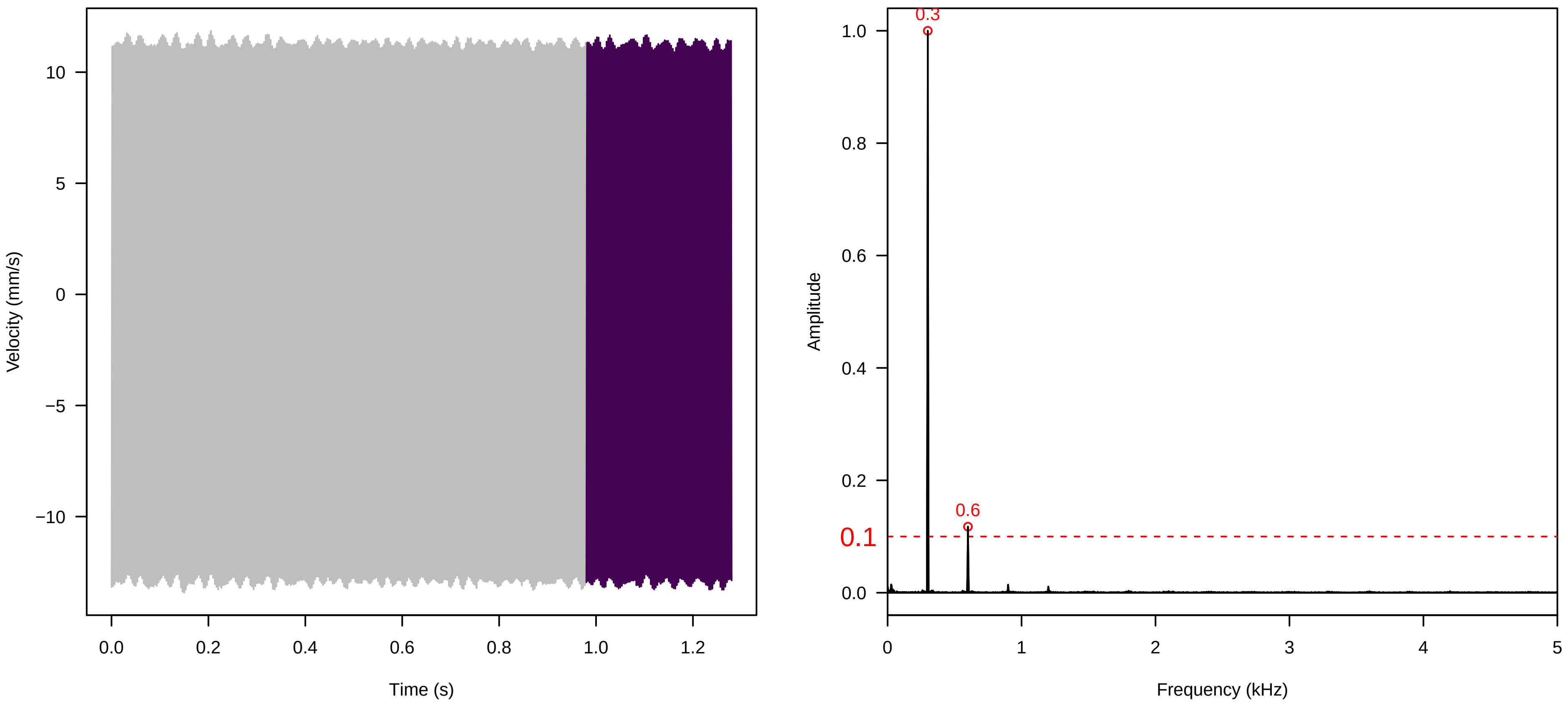

**Vel. = 0.014 ; Str. = Receptacle ; Axis = z ; Fl. accession = 10−s−79−27**

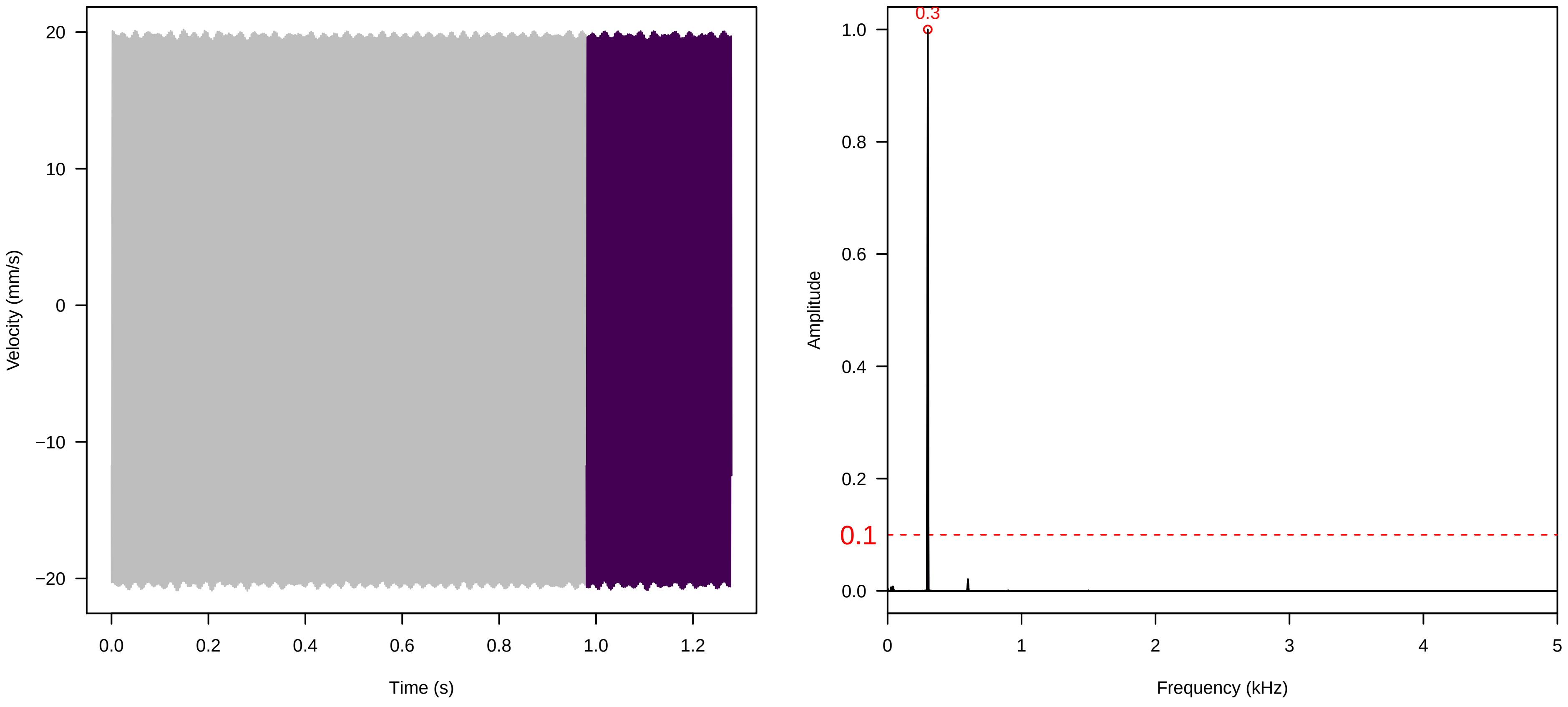


**Vel. = 0.014 ; Str. = FA ; Axis = z ; Fl. accession = 10−s−79−27**

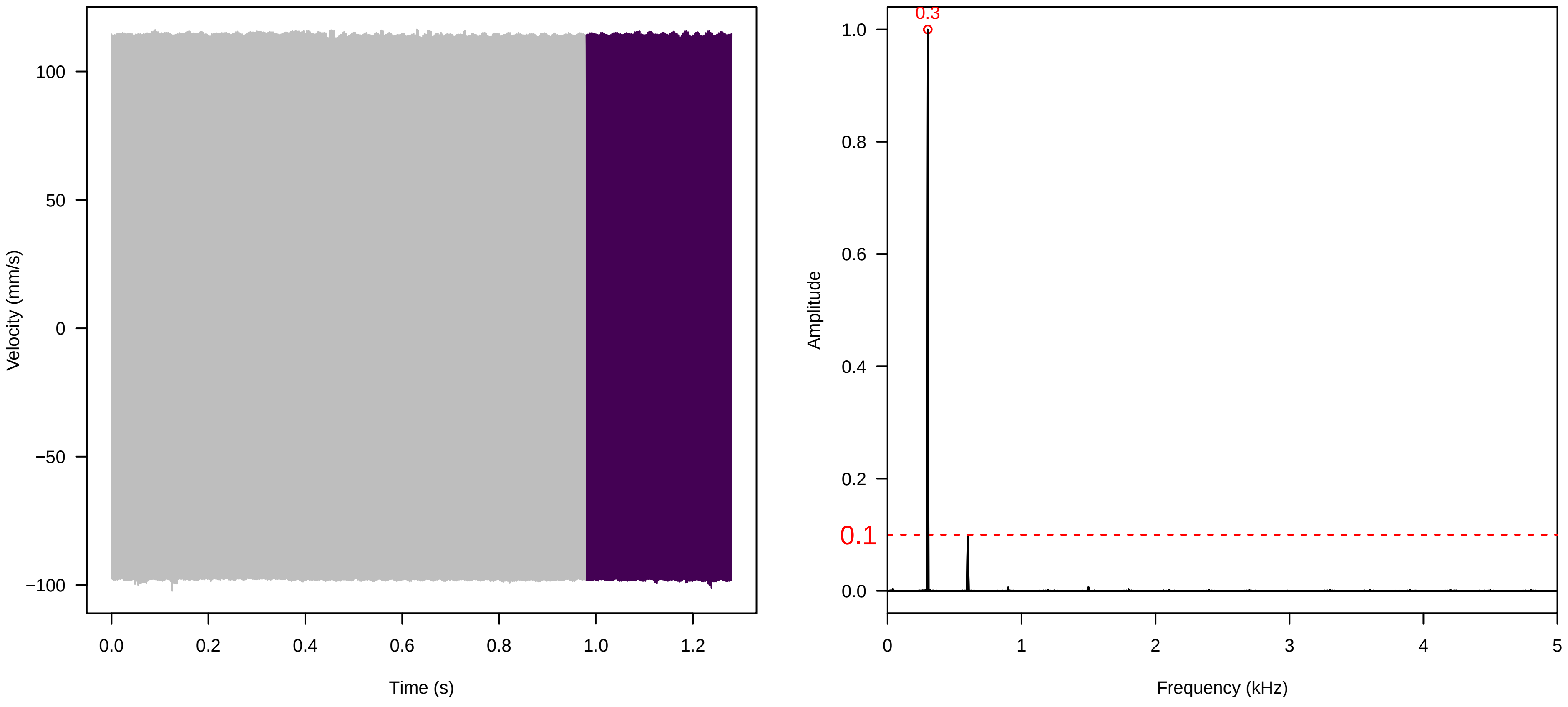


**Vel. = 0.014 ; Str. = Receptacle ; Axis = z ; Fl. accession = 10−s−79−27**

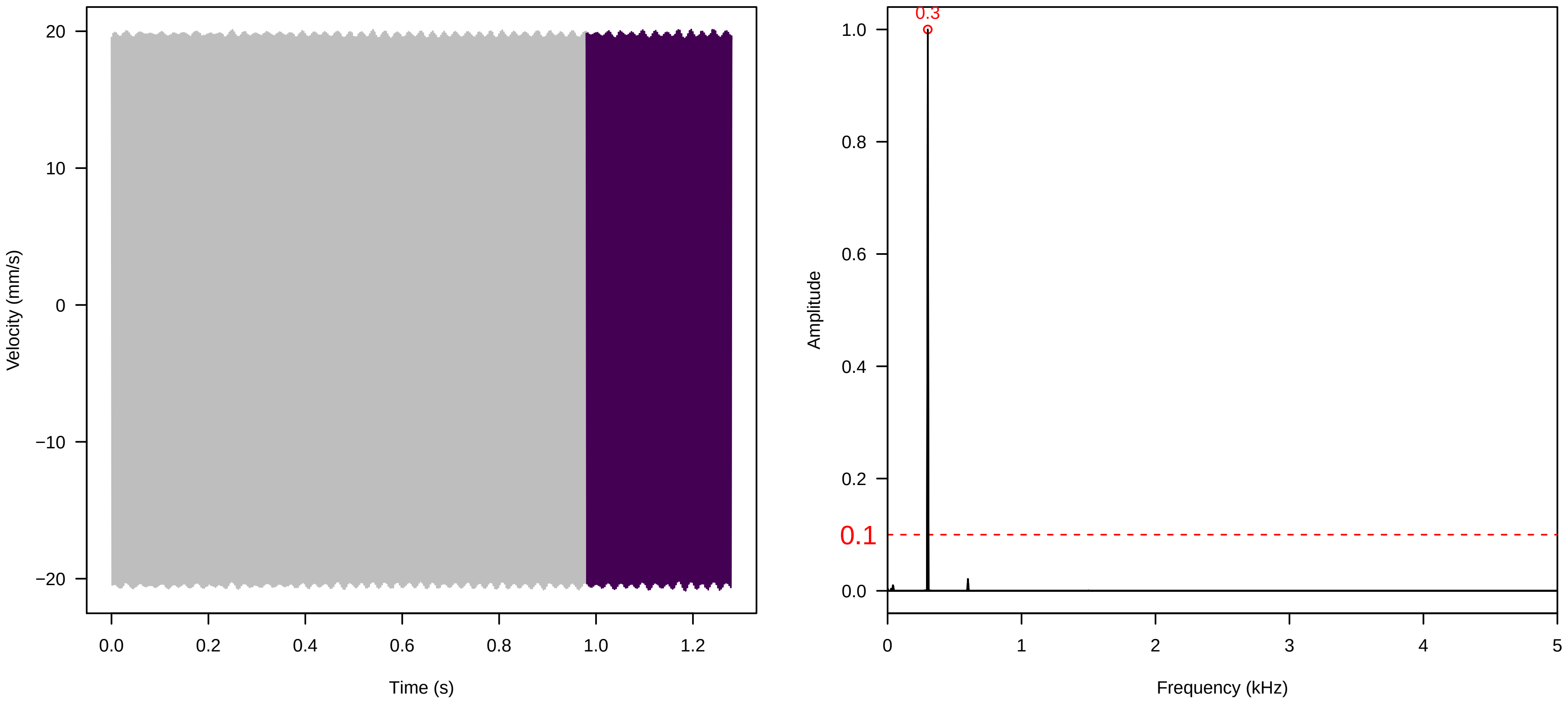

**Vel. = 0.014 ; Str. = PA ; Axis = z ; Fl. accession = 10−s−79−27**

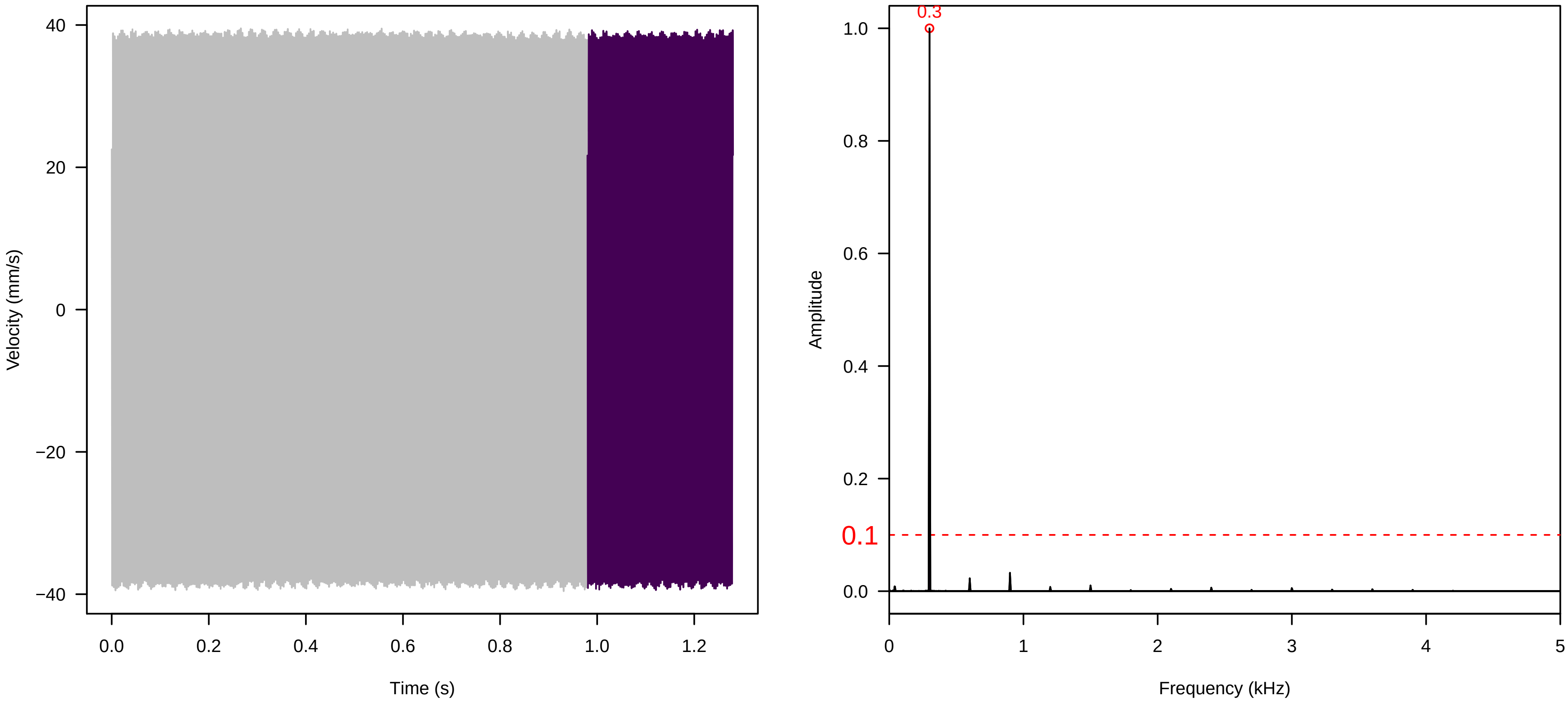


**Vel. = 0.014 ; Str. = Receptacle ; Axis = z ; Fl. accession = 10−s−79−27**

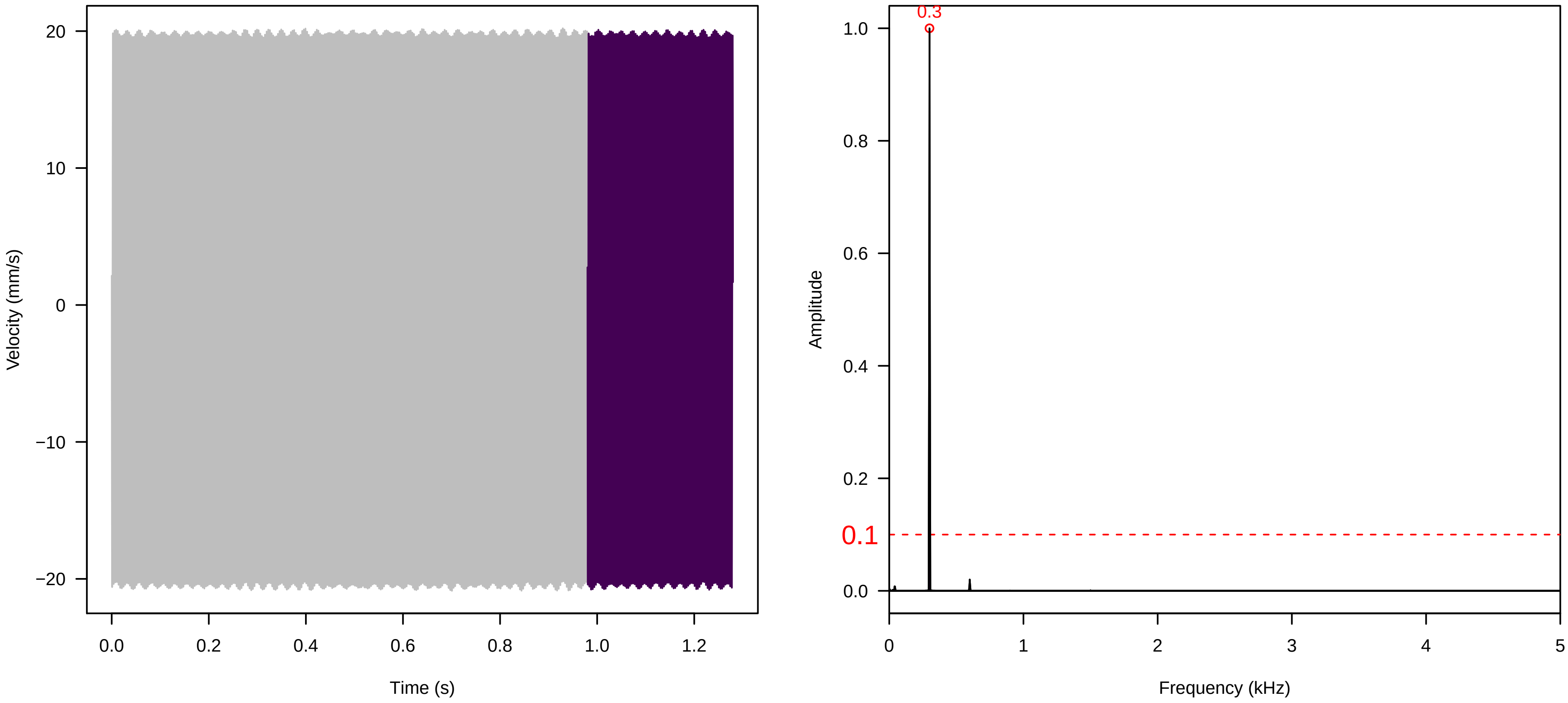


**Vel. = 0.014 ; Str. = Corolla ; Axis = y ; Fl. accession = 10−s−79−27**

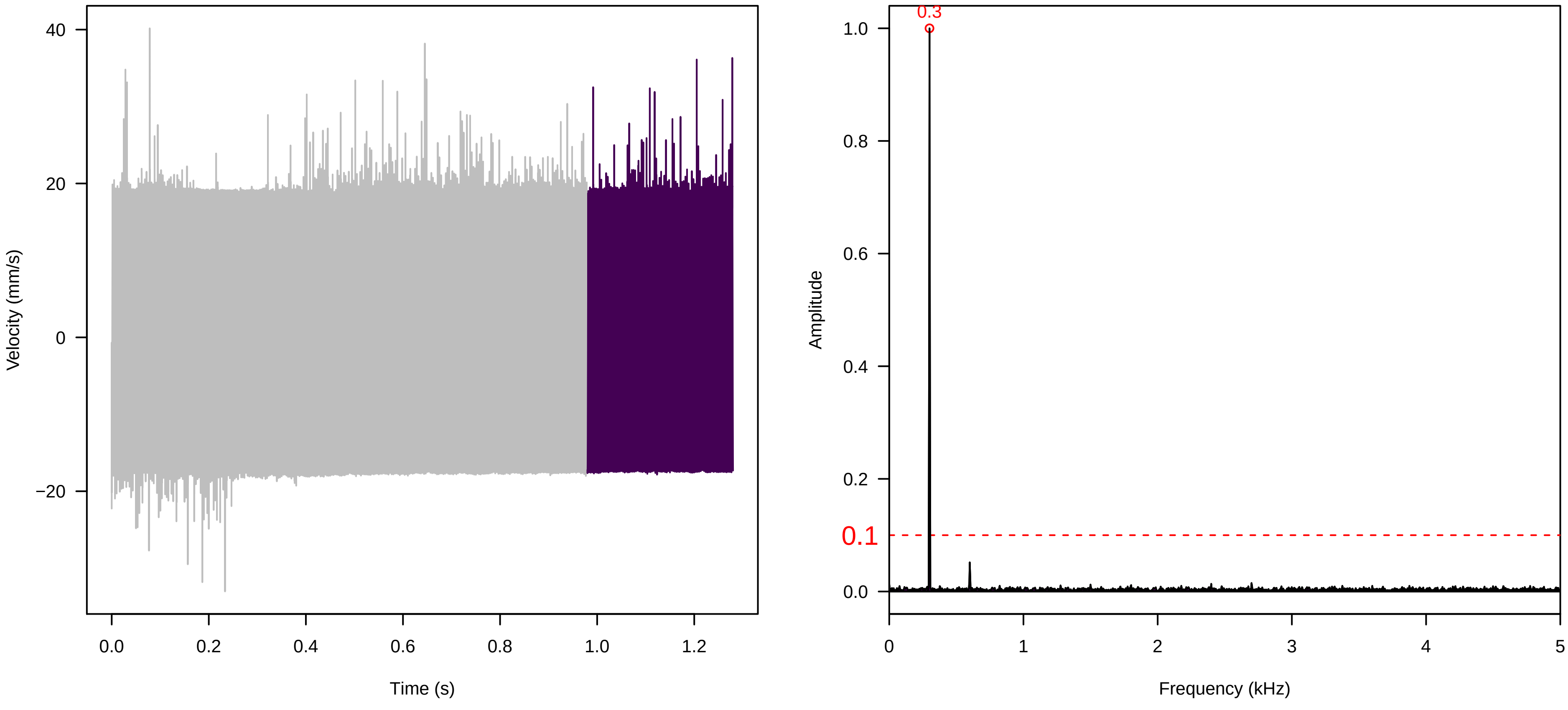

**Vel. = 0.014 ; Str. = Receptacle ; Axis = y ; Fl. accession = 10−s−79−27**

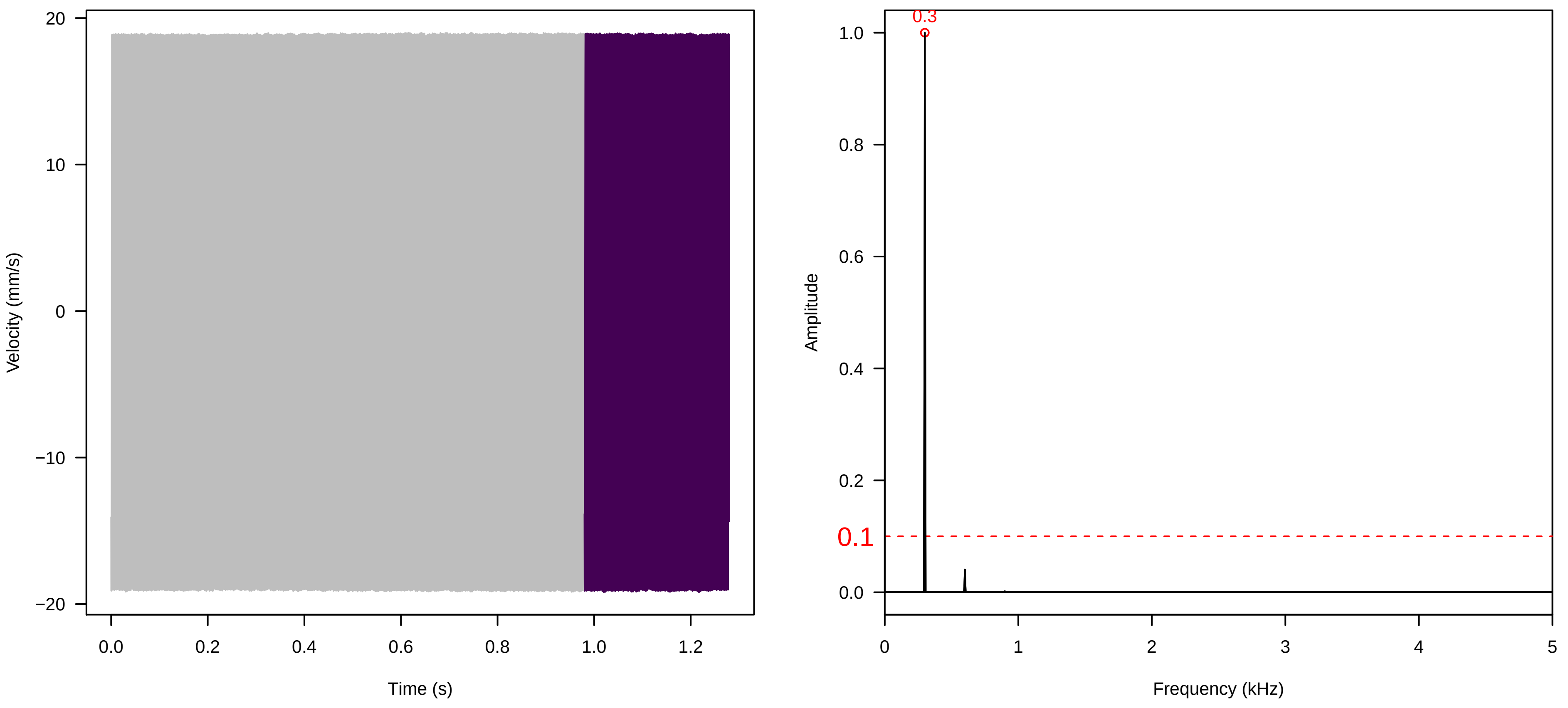


**Vel. = 0.014 ; Str. = FA ; Axis = y ; Fl. accession = 10−s−79−27**

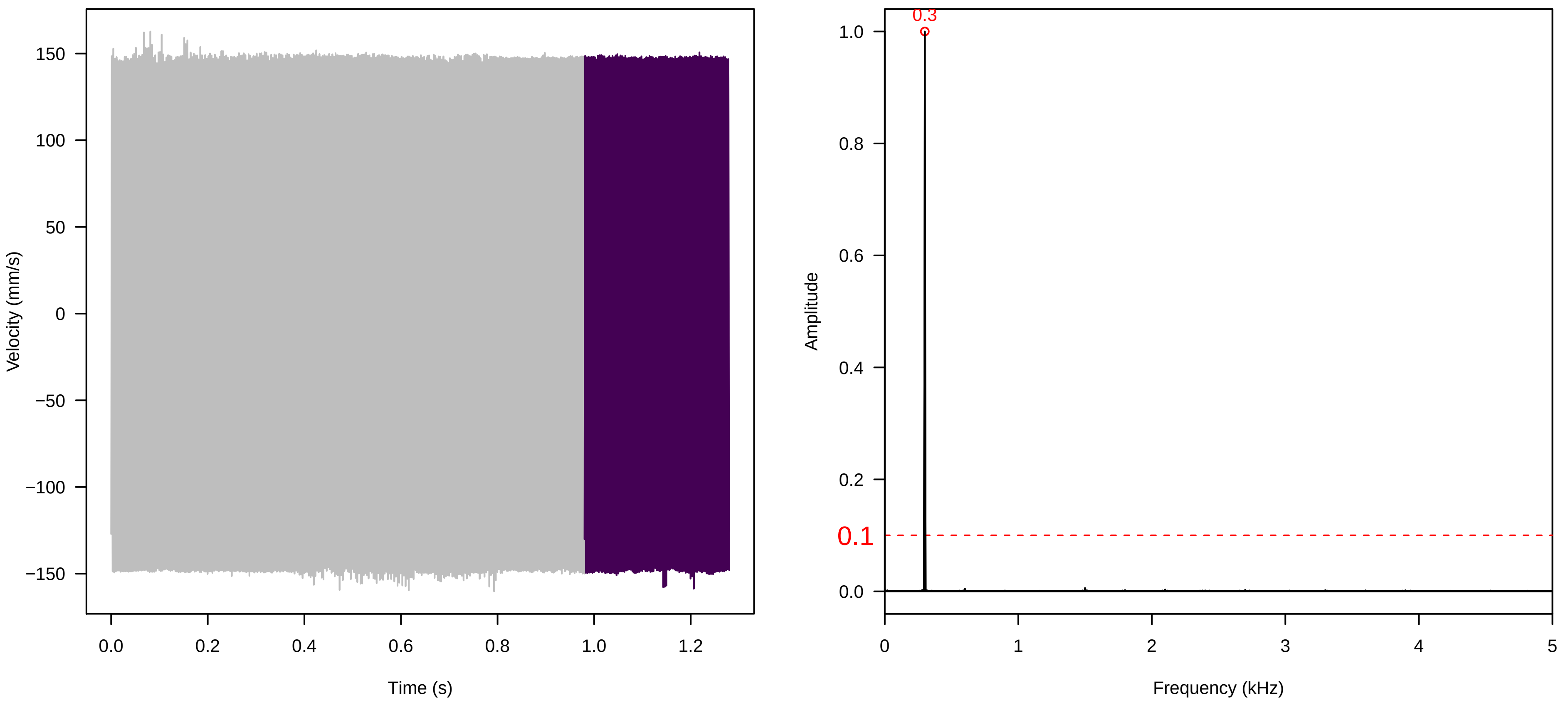


**Vel. = 0.014 ; Str. = Receptacle ; Axis = y ; Fl. accession = 10−s−79−27**

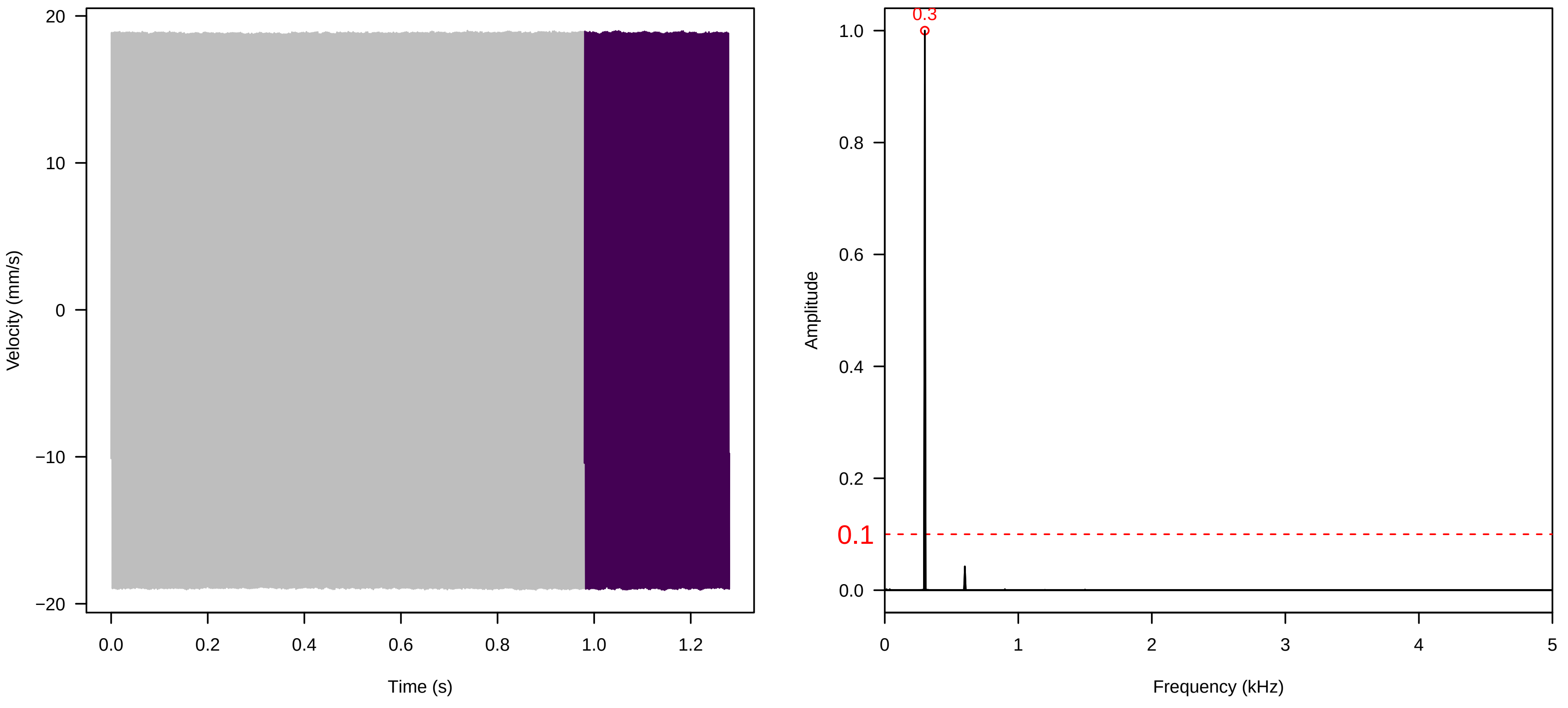

**Vel. = 0.014 ; Str. = PA ; Axis = y ; Fl. accession = 10−s−79−27**

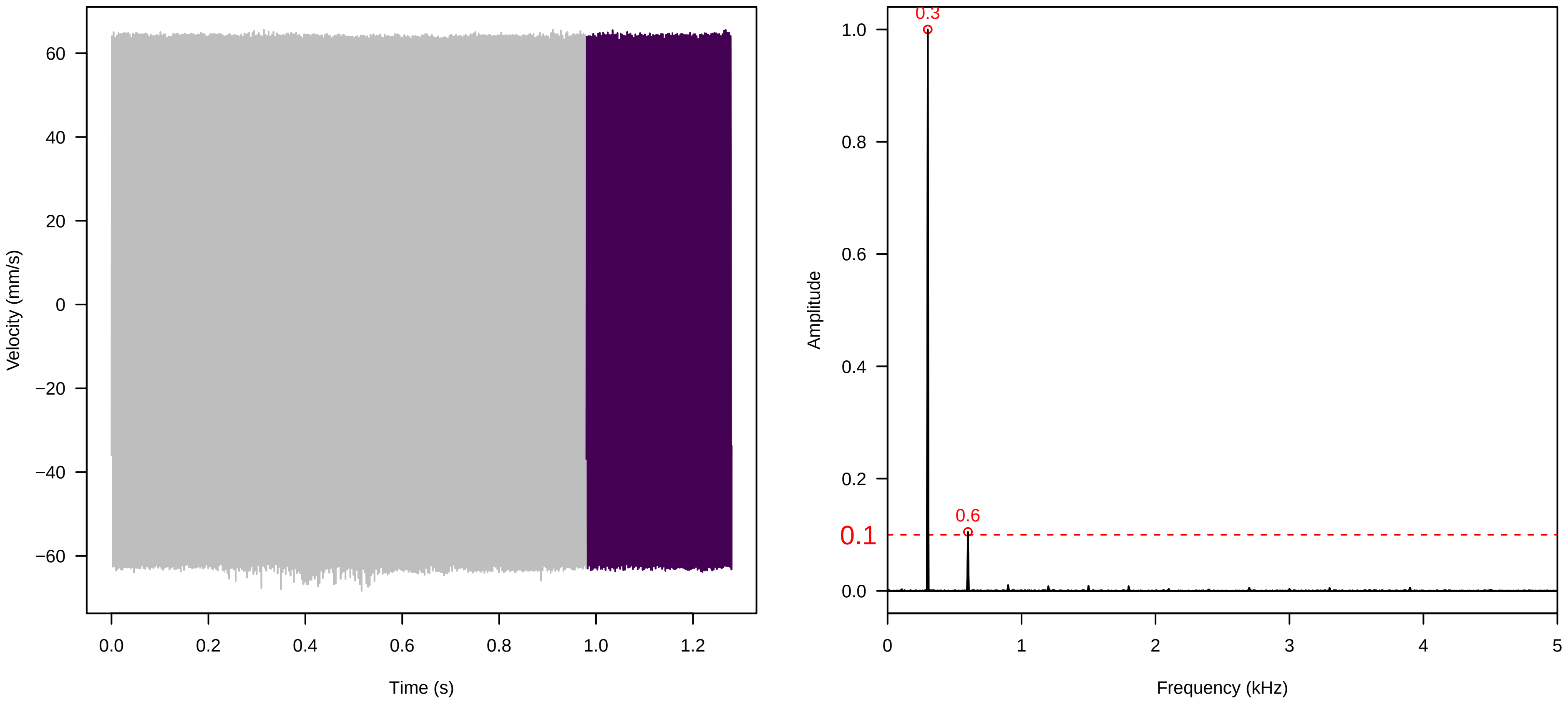


**Vel. = 0.014 ; Str. = Receptacle ; Axis = y ; Fl. accession = 10−s−79−27**

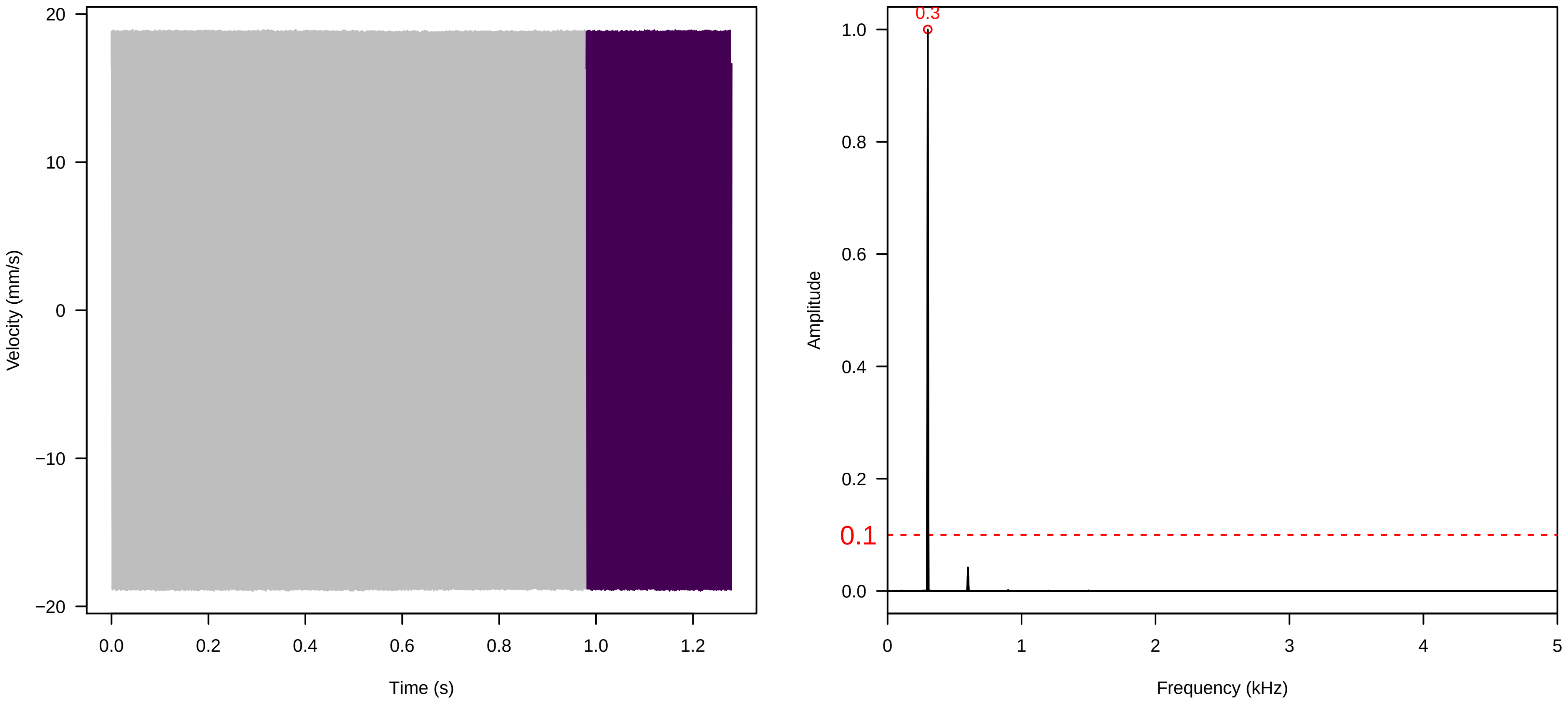


**Vel. = 0.028 ; Str. = PA ; Axis = y ; Fl. accession = 10−s−79−27**

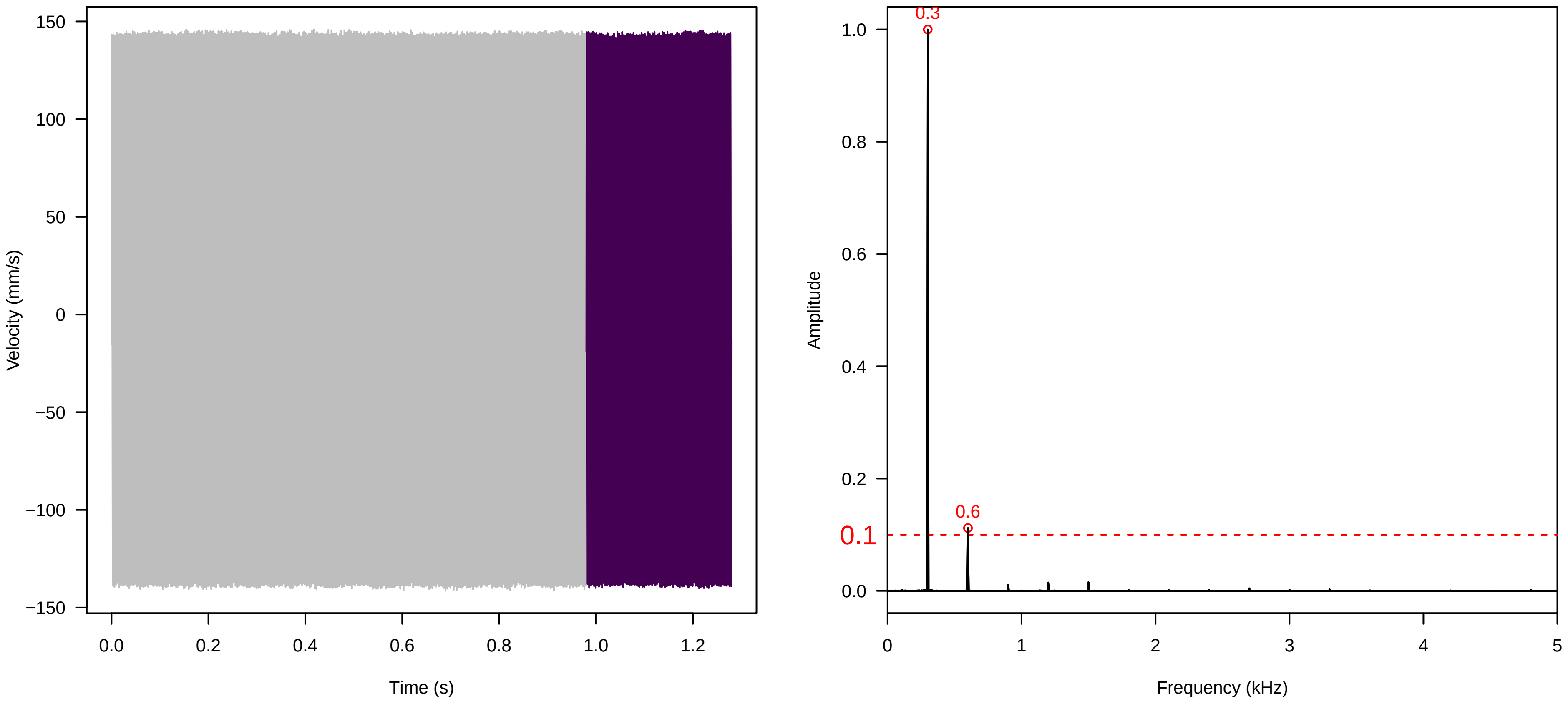

**Vel. =  0.028 ; Str. =  Receptacle ; Axis =  y ; Fl. accession = 10−s−79−27**

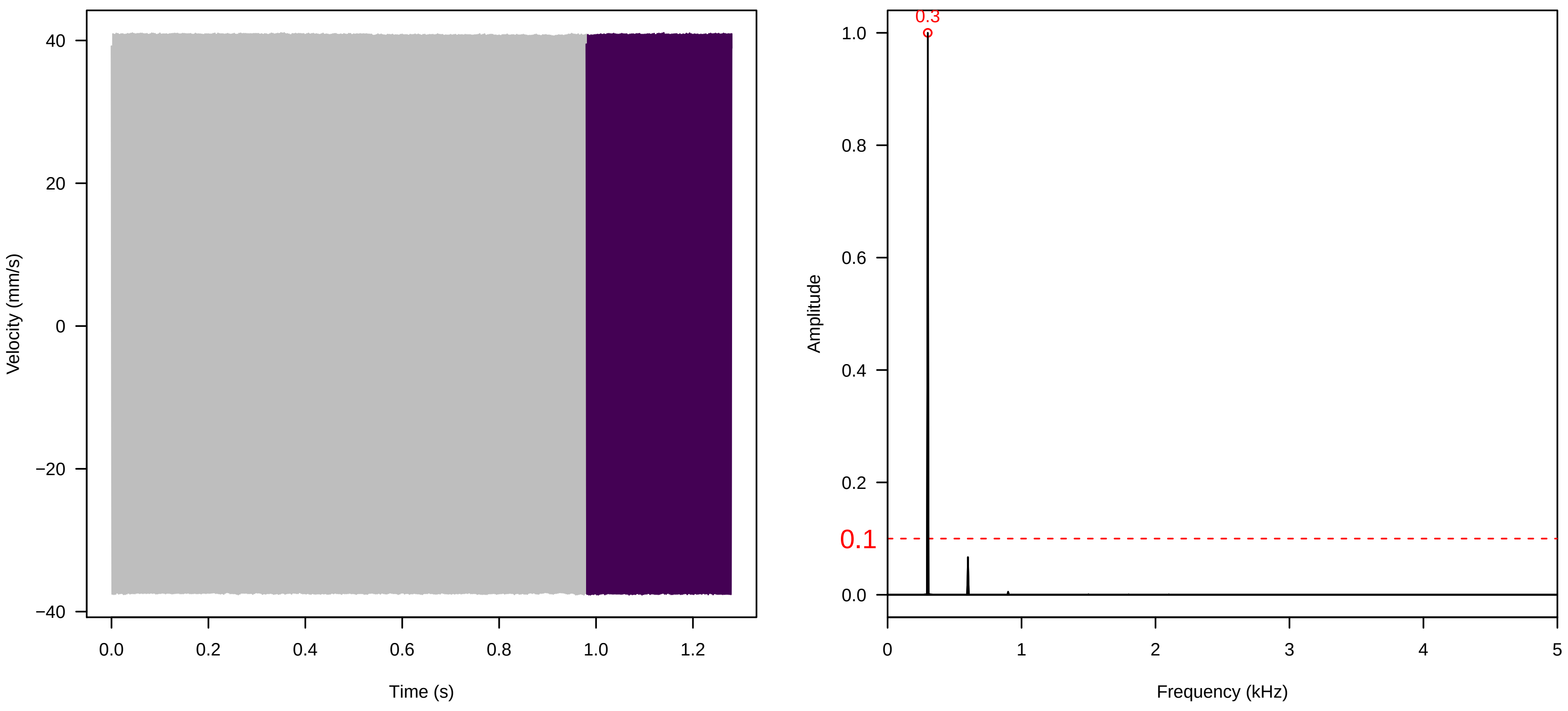


**Vel. =  0.028 ; Str. =  FA ; Axis =  y ; Fl. accession = 10−s−79−27**

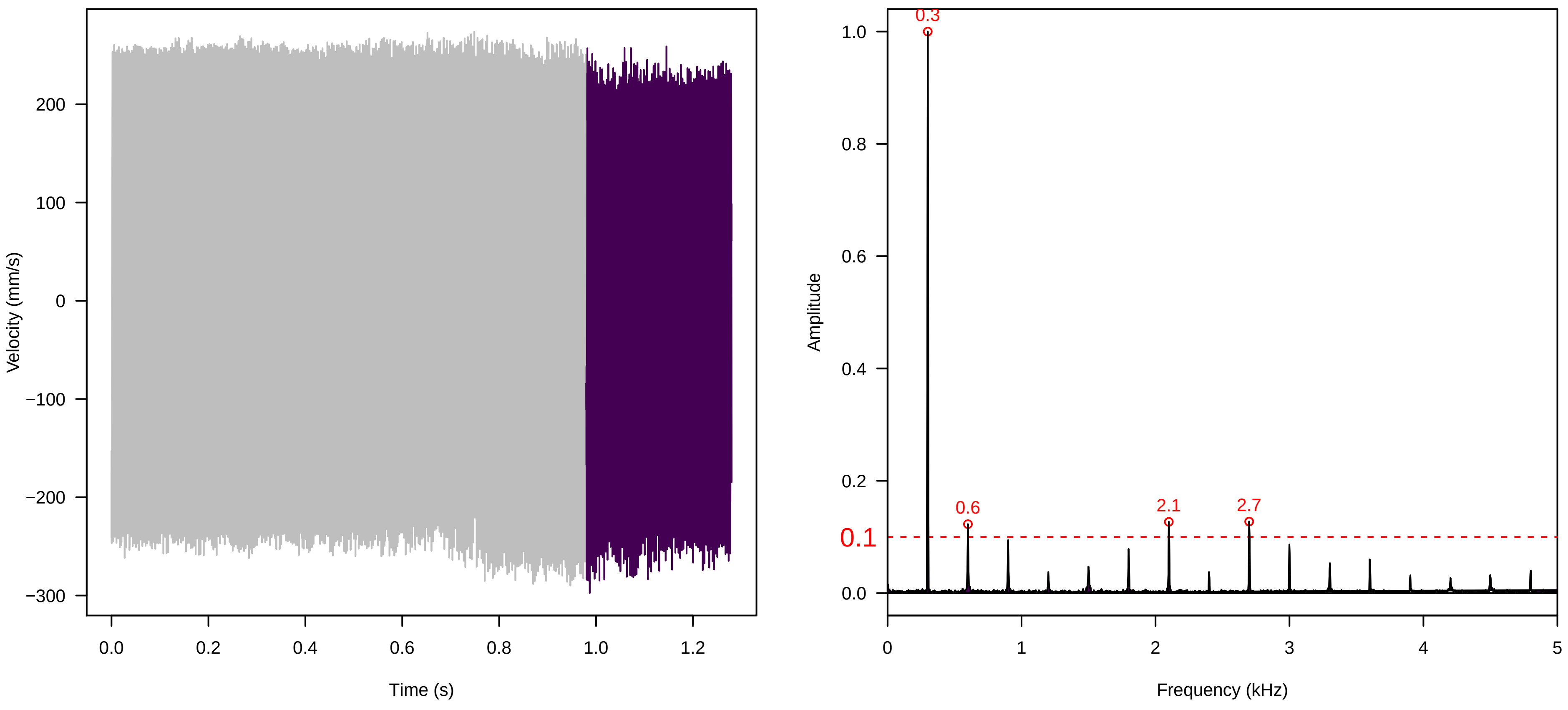


**Vel. =  0.028 ; Str. =  Receptacle ; Axis =  y ; Fl. accession = 10−s−79−27**

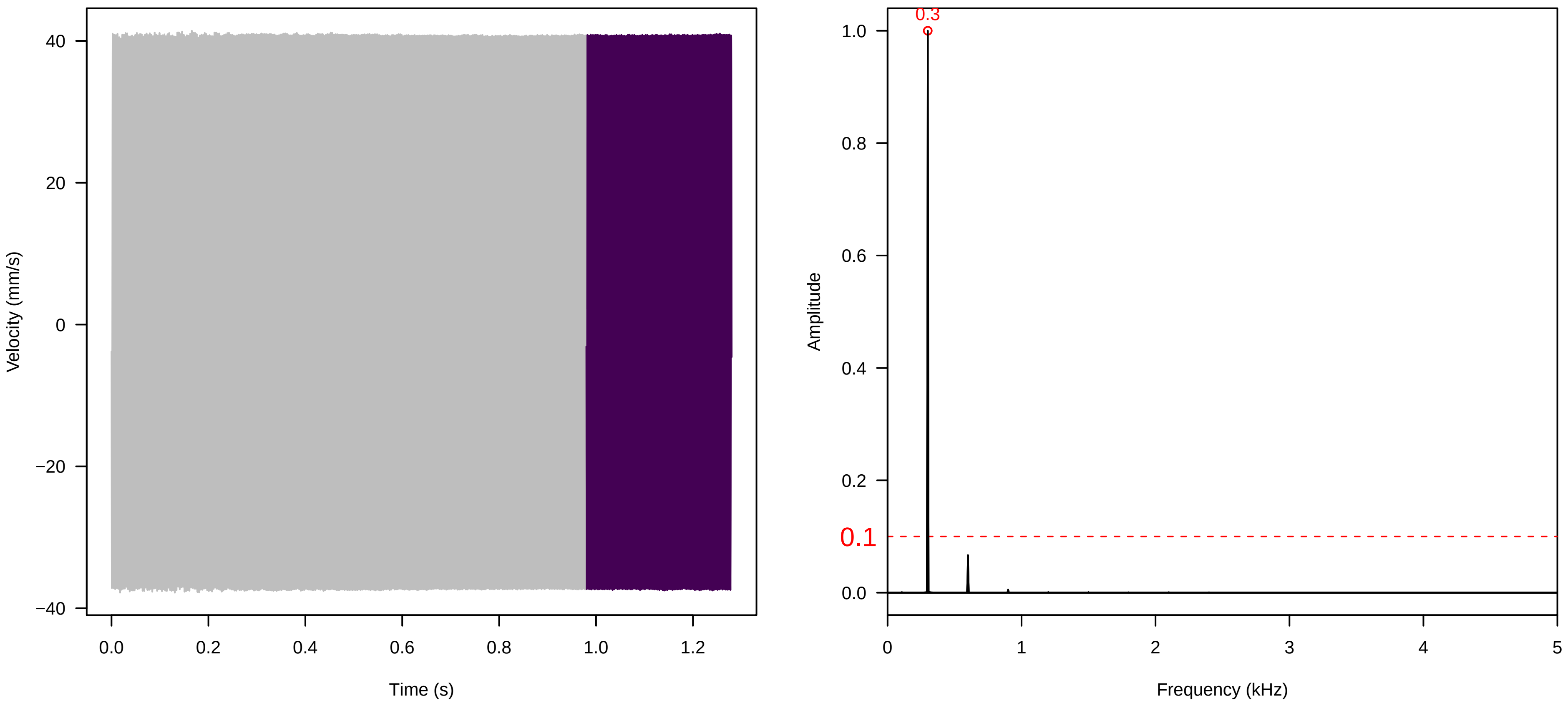

**Vel. =  0.028 ; Str. =  Corolla ; Axis =  y ; Fl. accession = 10−s−79−27**

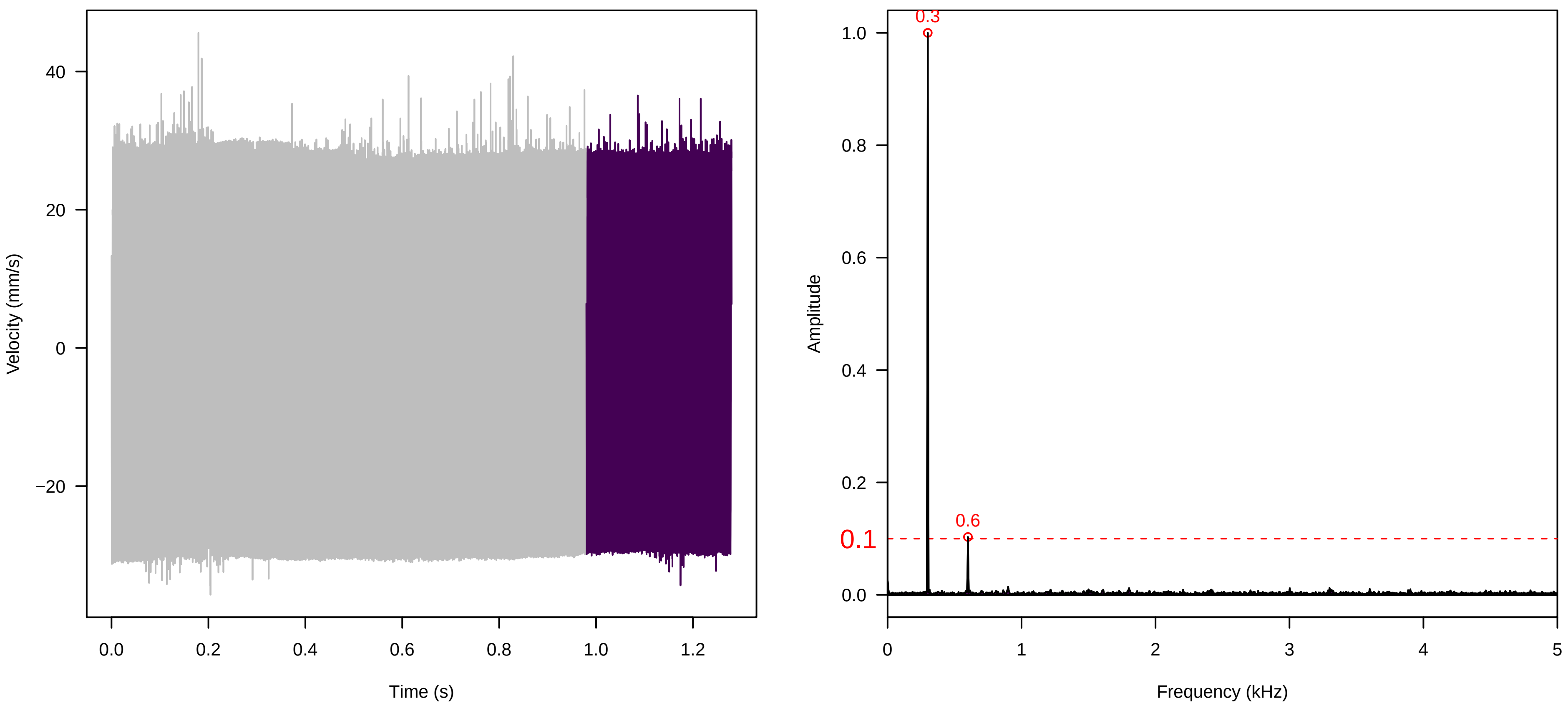


**Vel. =  0.028 ; Str. =  Receptacle ; Axis =  y ; Fl. accession = 10−s−79−27**

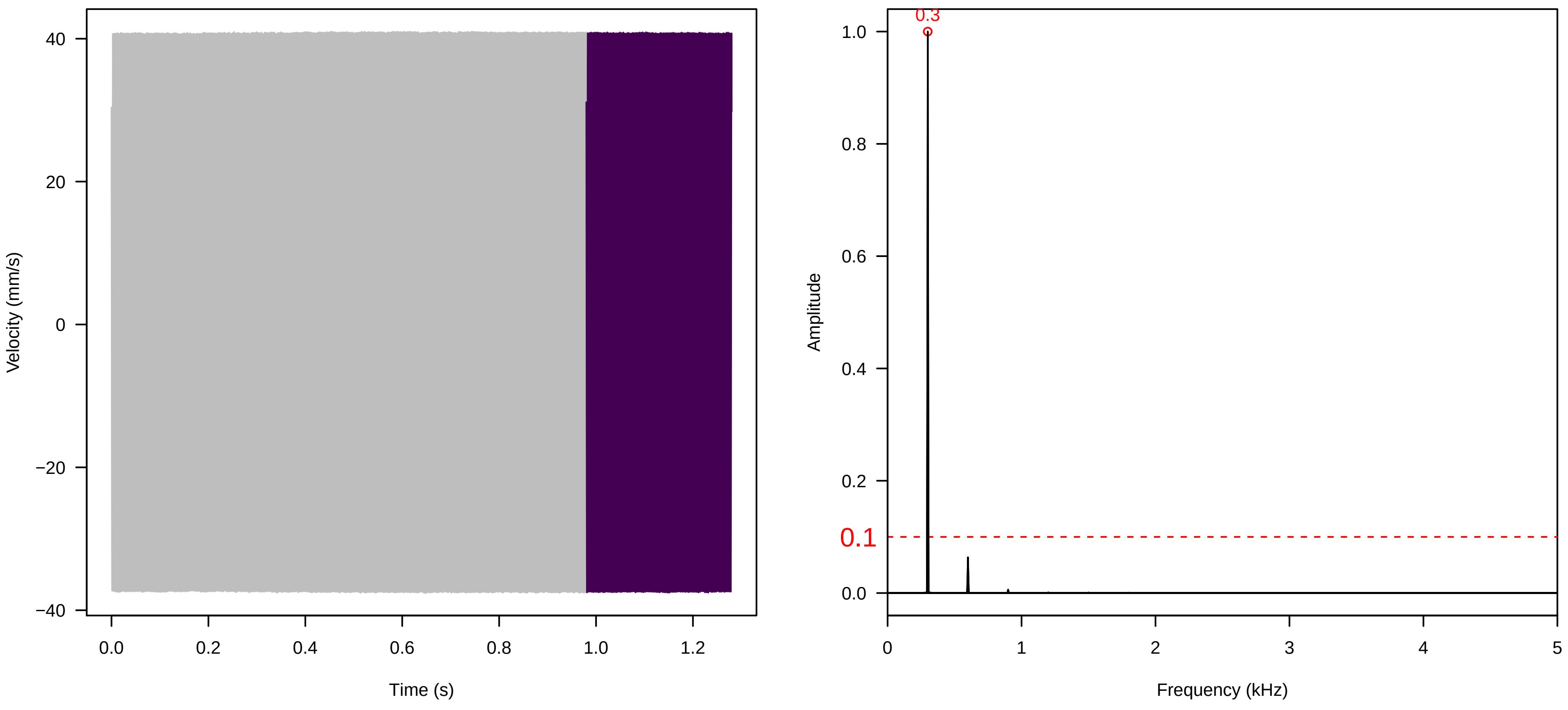


**Vel. =  0.057 ; Str. =  Corolla ; Axis =  y ; Fl. accession = 10−s−79−27**

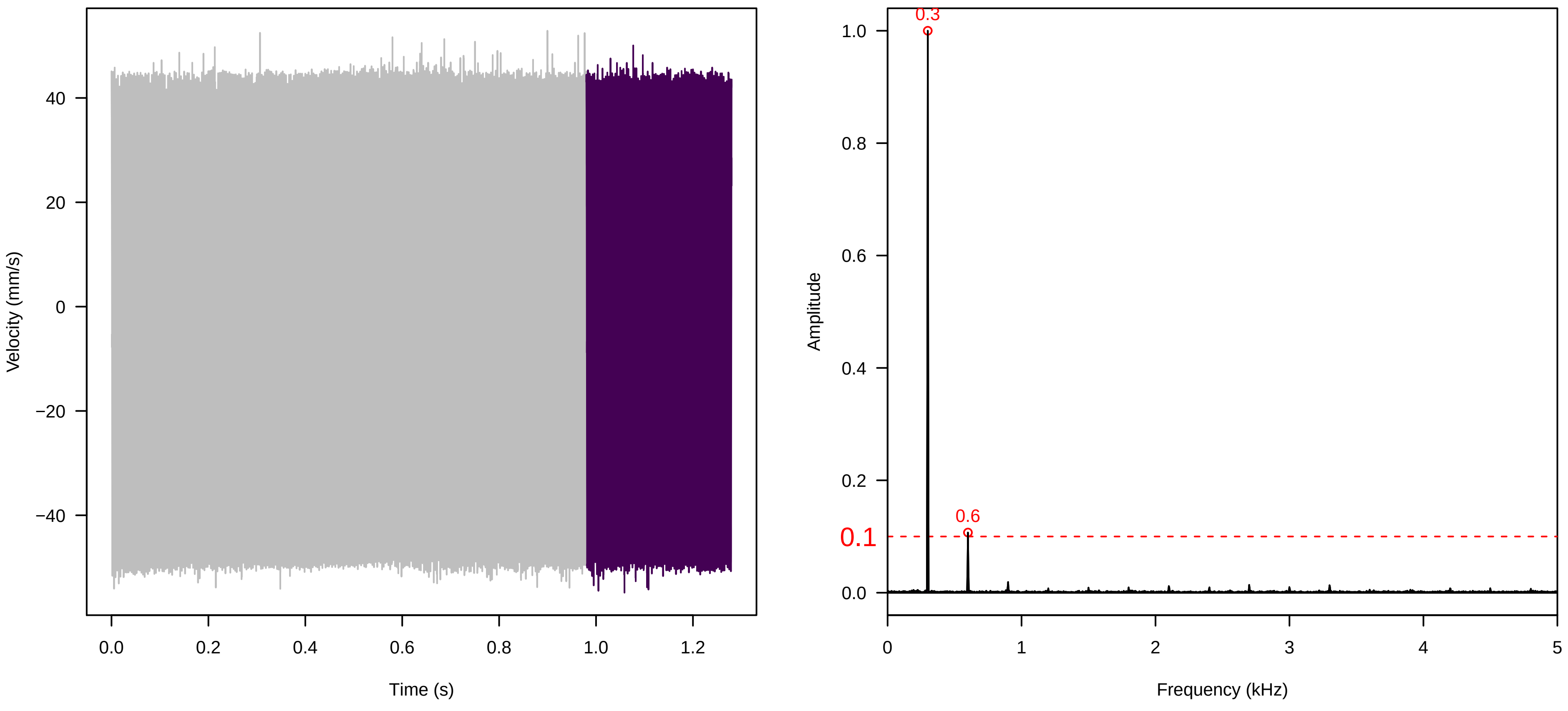

**Vel. =  0.057 ; Str. =  Receptacle ; Axis =  y ; Fl. accession = 10−s−79−27**

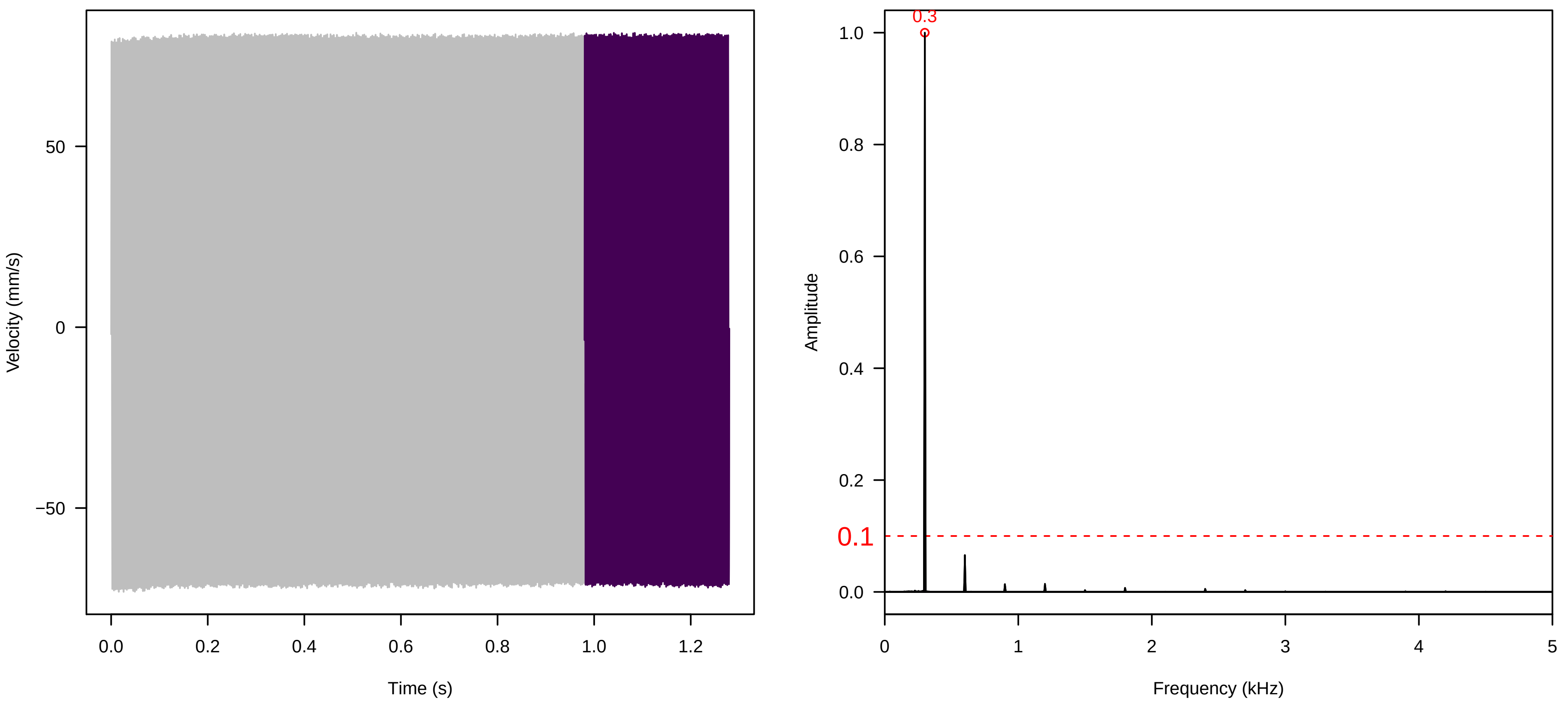


**Vel. =  0.057 ; Str. =  FA ; Axis =  y ; Fl. accession = 10−s−79−27**

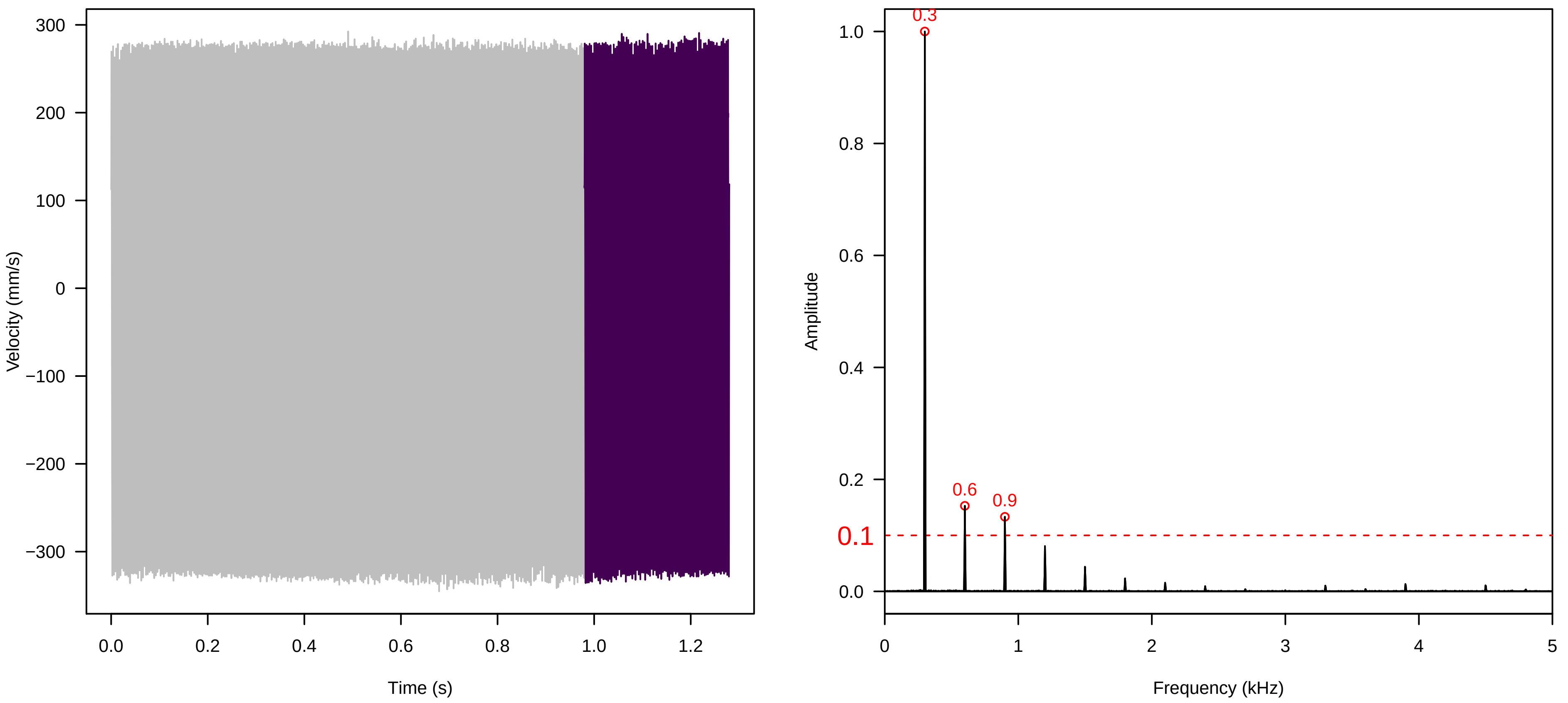


**Vel. =  0.057 ; Str. =  Receptacle ; Axis =  y ; Fl. accession = 10−s−79−27**

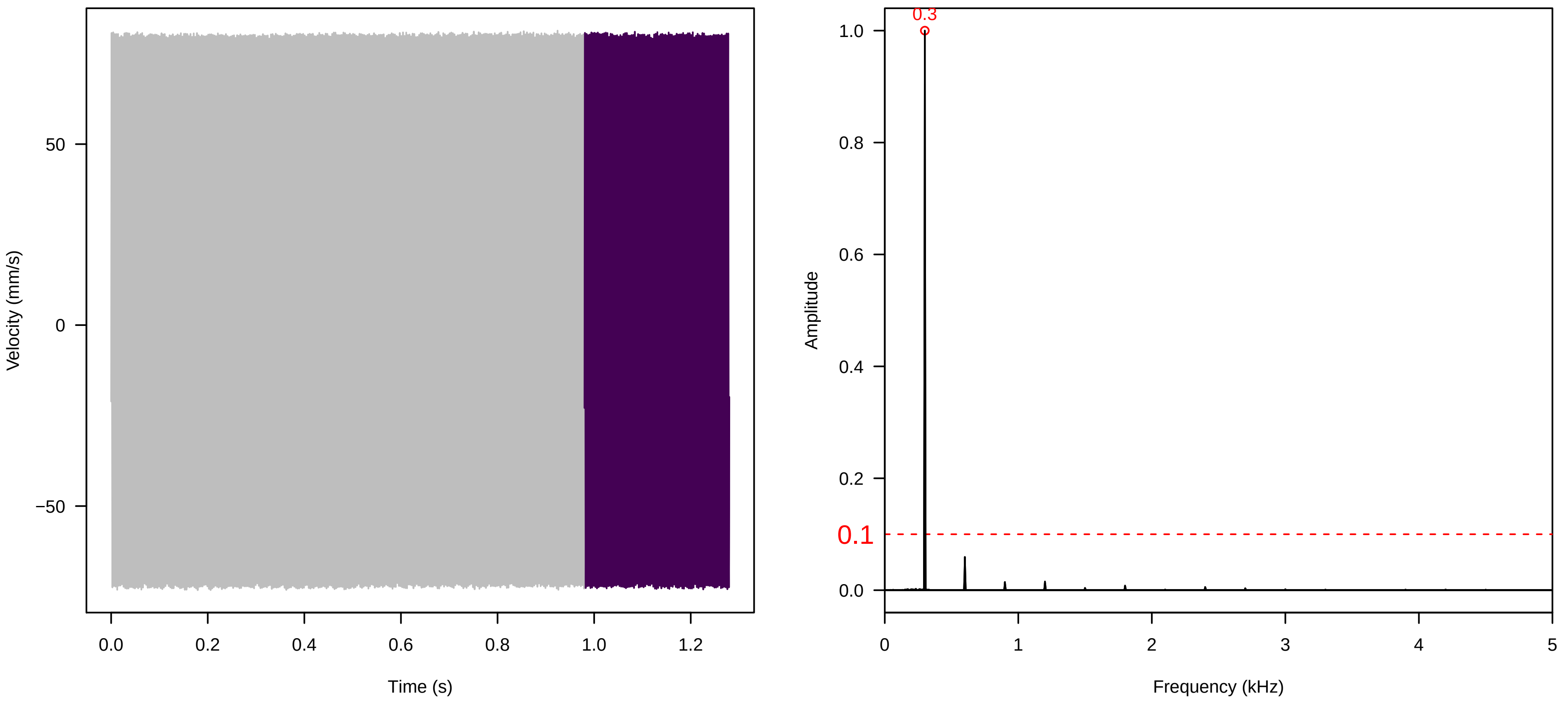

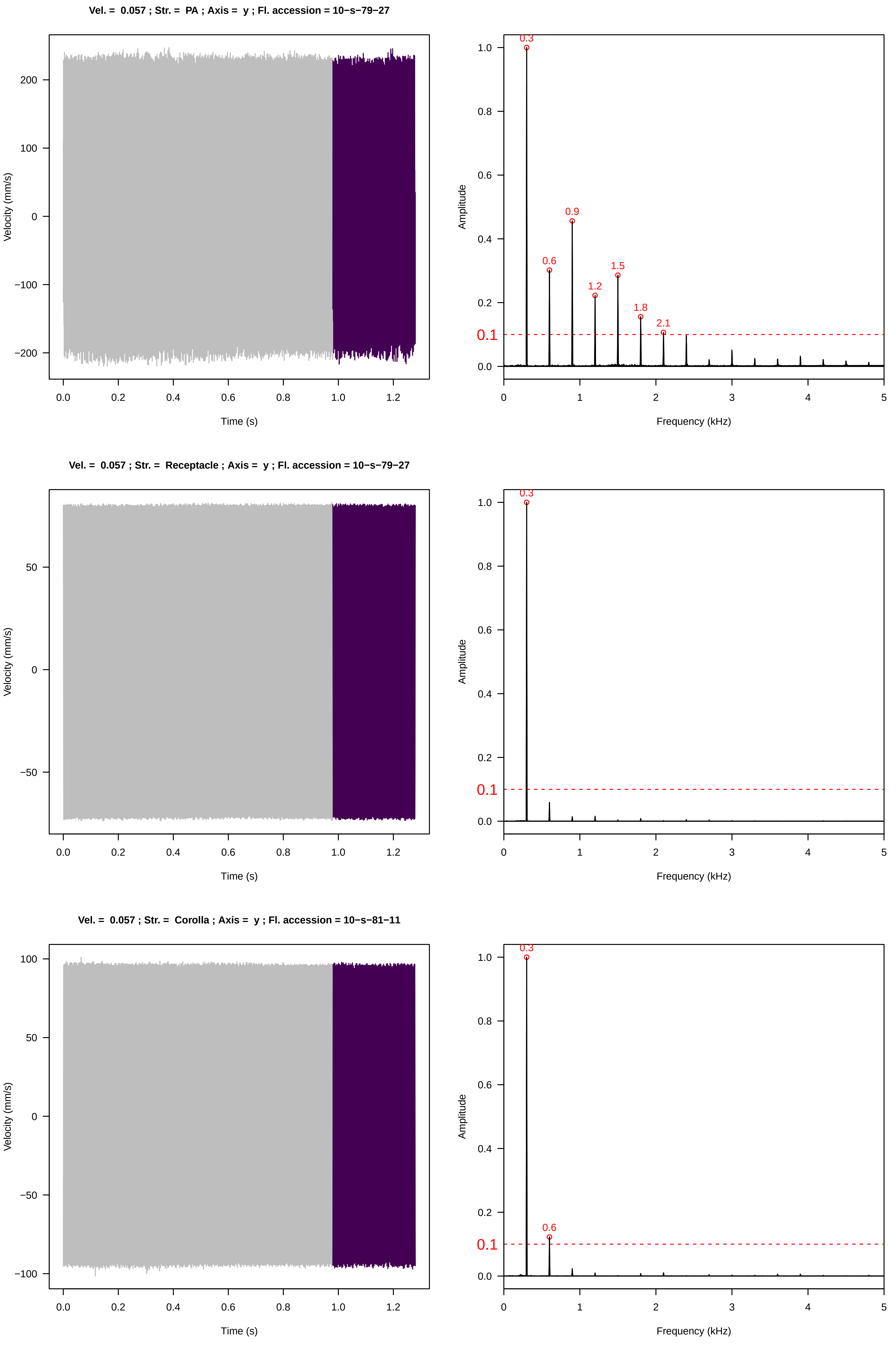
Vel. = 0.057 ; Str. = PA ; Axis = y ; Fl. accession = 10−s−79−27
Velocity (mm/s)
Time (s)
Amplitude
Frequency (kHz)
0.3
0.6
0.9
1.2
1.5
1.8
2.1
0.1
Vel. = 0.057 ; Str. = Receptacle ; Axis = y ; Fl. accession = 10−s−79−27
Velocity (mm/s)
Time (s)
Amplitude
Frequency (kHz)
0.3
0.1
Vel. = 0.057 ; Str. = Corolla ; Axis = y ; Fl. accession = 10−s−81−11
Velocity (mm/s)
Time (s)
Amplitude
Frequency (kHz)
0.3
0.6
0.1

**Vel. = 0.057 ; Str. = Receptacle ; Axis = y ; Fl. accession = 10−s−81−11**

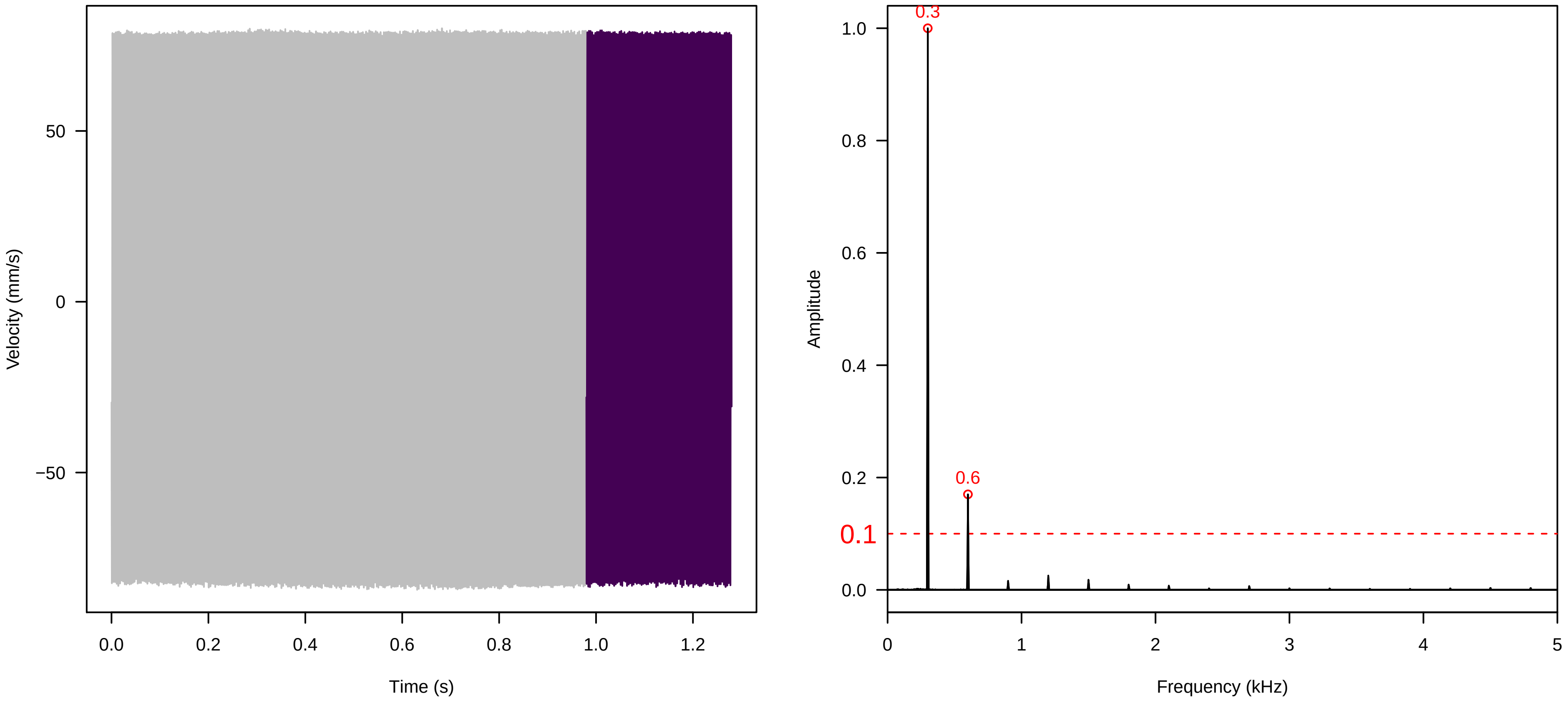


**Vel. = 0.057 ; Str. = FA ; Axis = y ; Fl. accession = 10−s−81−11**

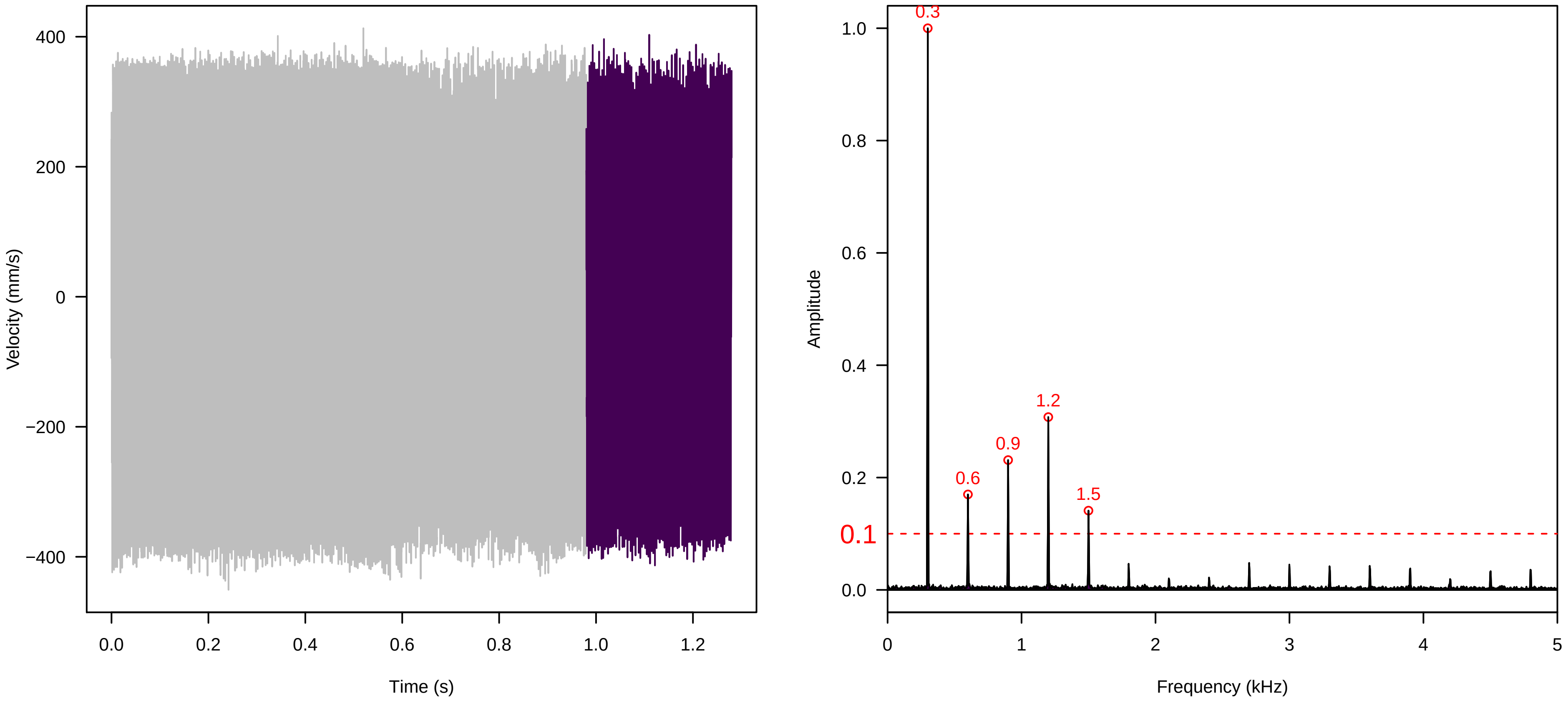


**Vel. = 0.057 ; Str. = Receptacle ; Axis = y ; Fl. accession = 10−s−81−11**

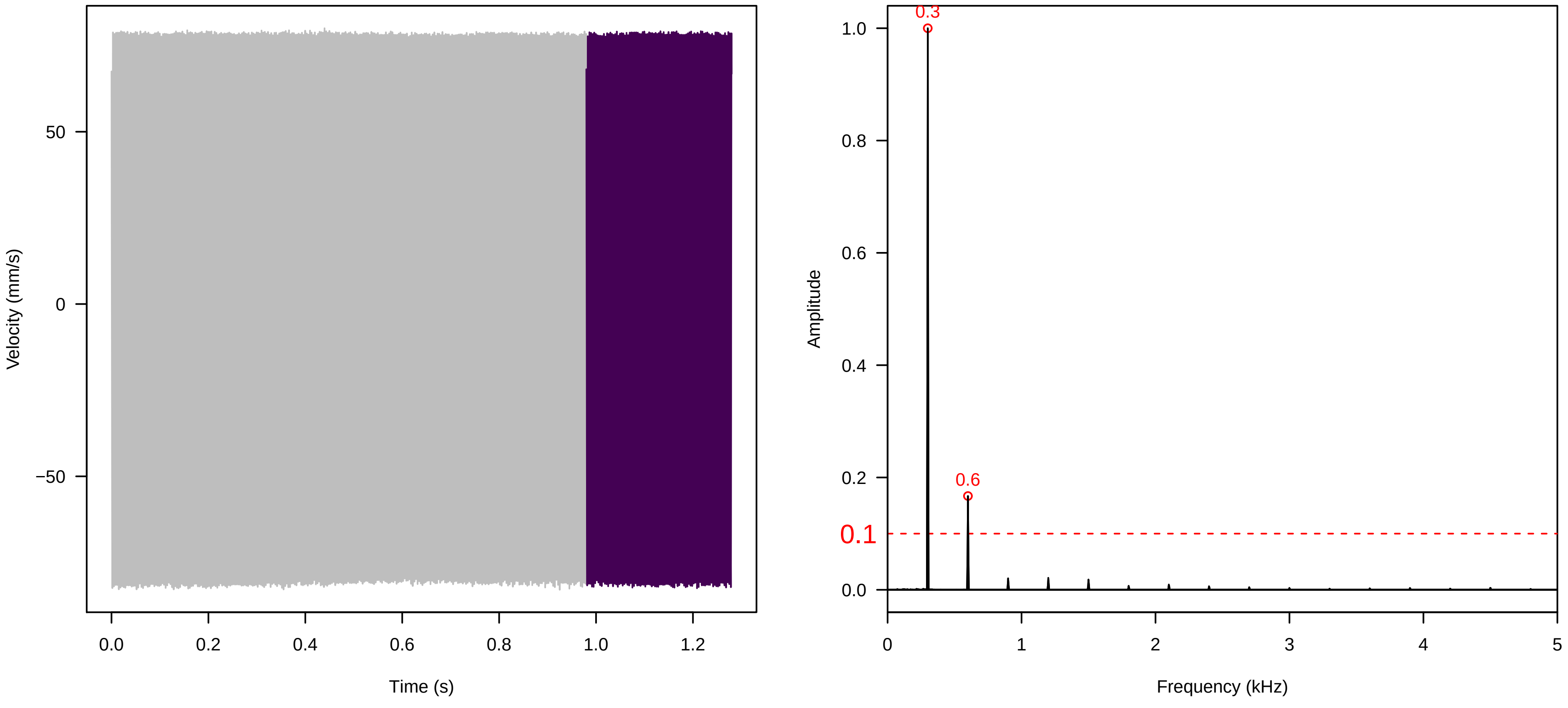

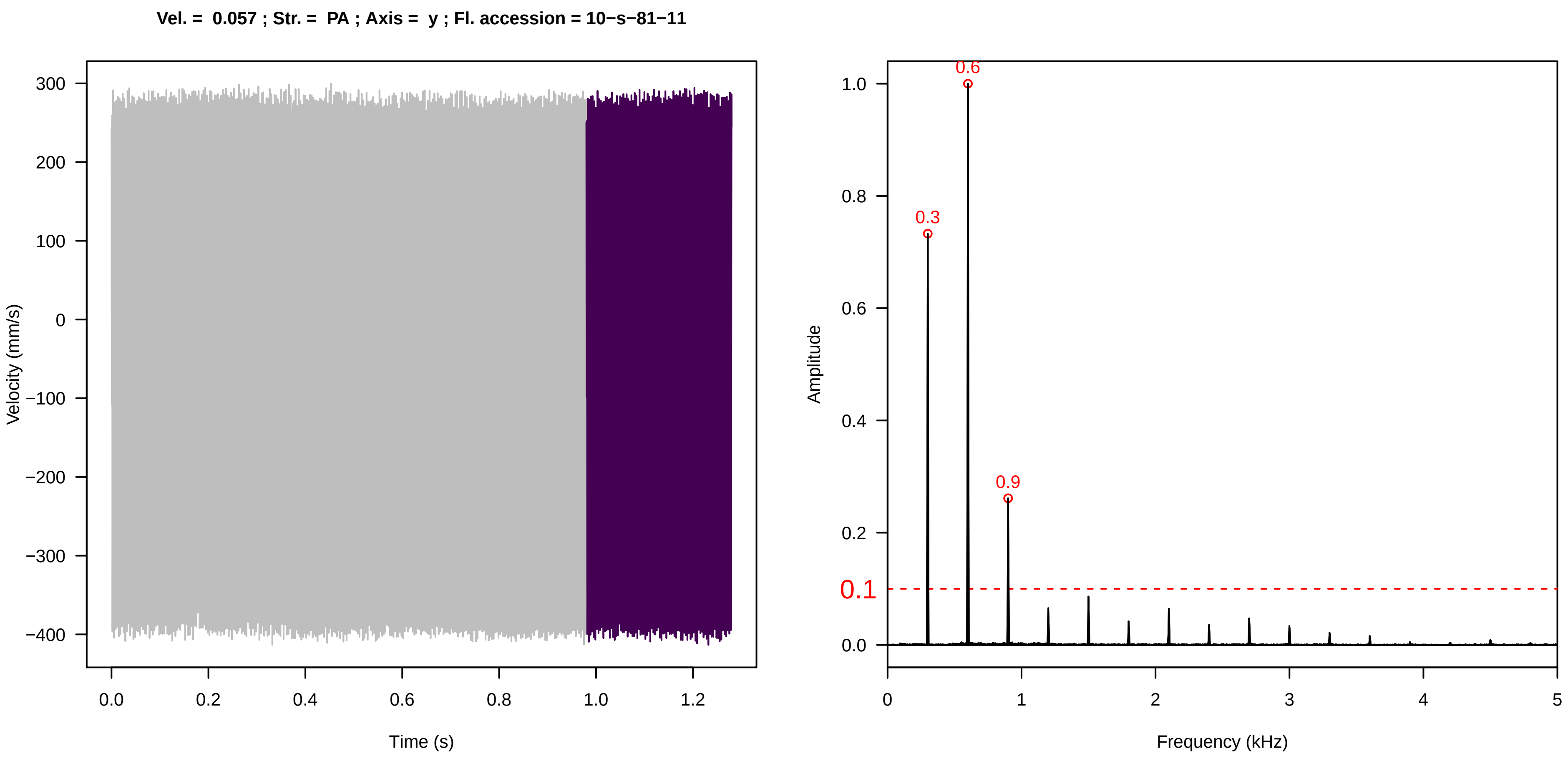
Vel. = 0.057 ; Str. = PA ; Axis = y ; Fl. accession = 10−s−81−11
Velocity (mm/s)
Time (s)
Amplitude
Frequency (kHz)
0.3
0.6
0.9
0.1

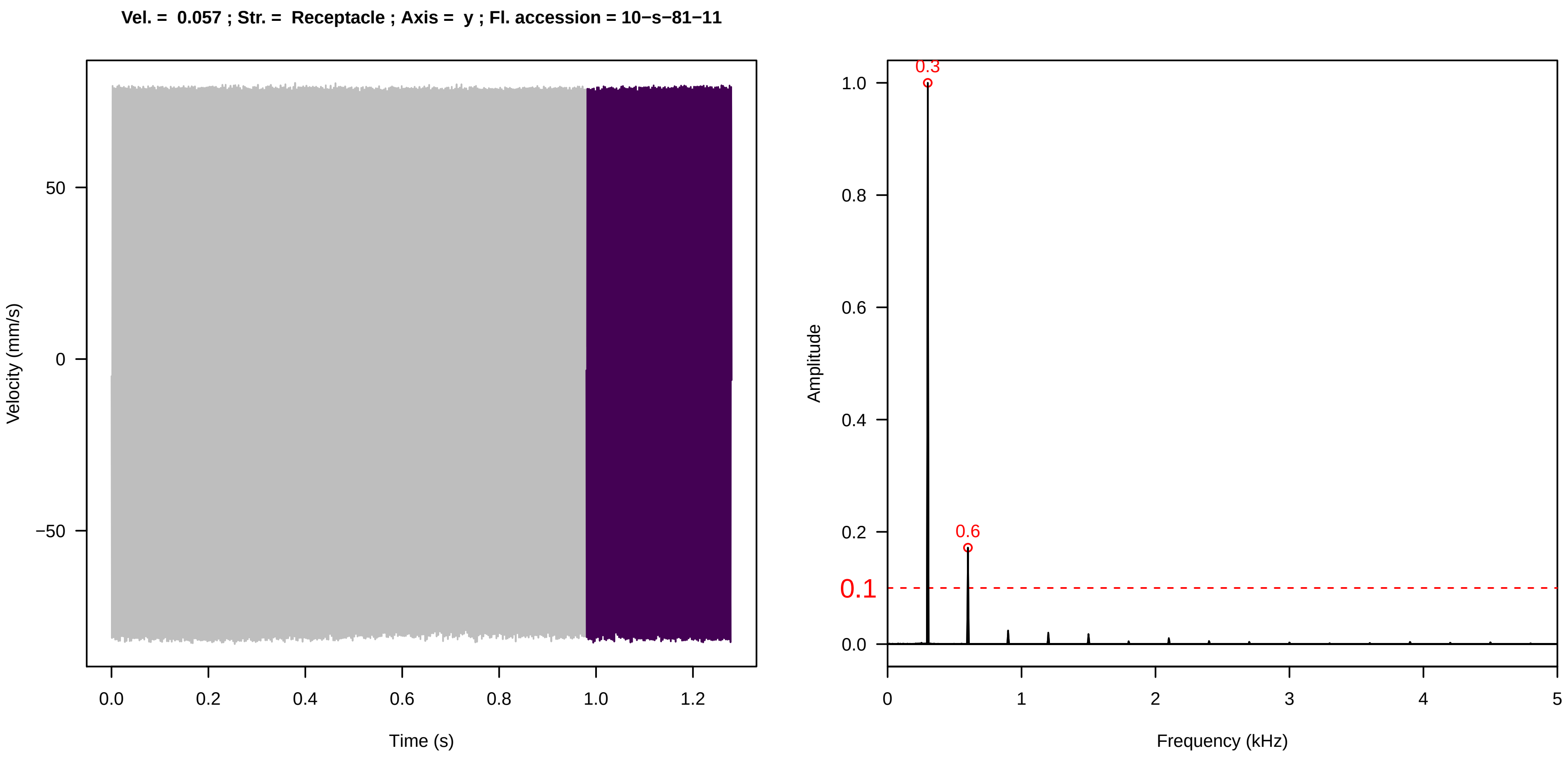
Vel. = 0.057 ; Str. = Receptacle ; Axis = y ; Fl. accession = 10−s−81−11
Velocity (mm/s)
Time (s)
Amplitude
Frequency (kHz)
0.3
0.6
0.1

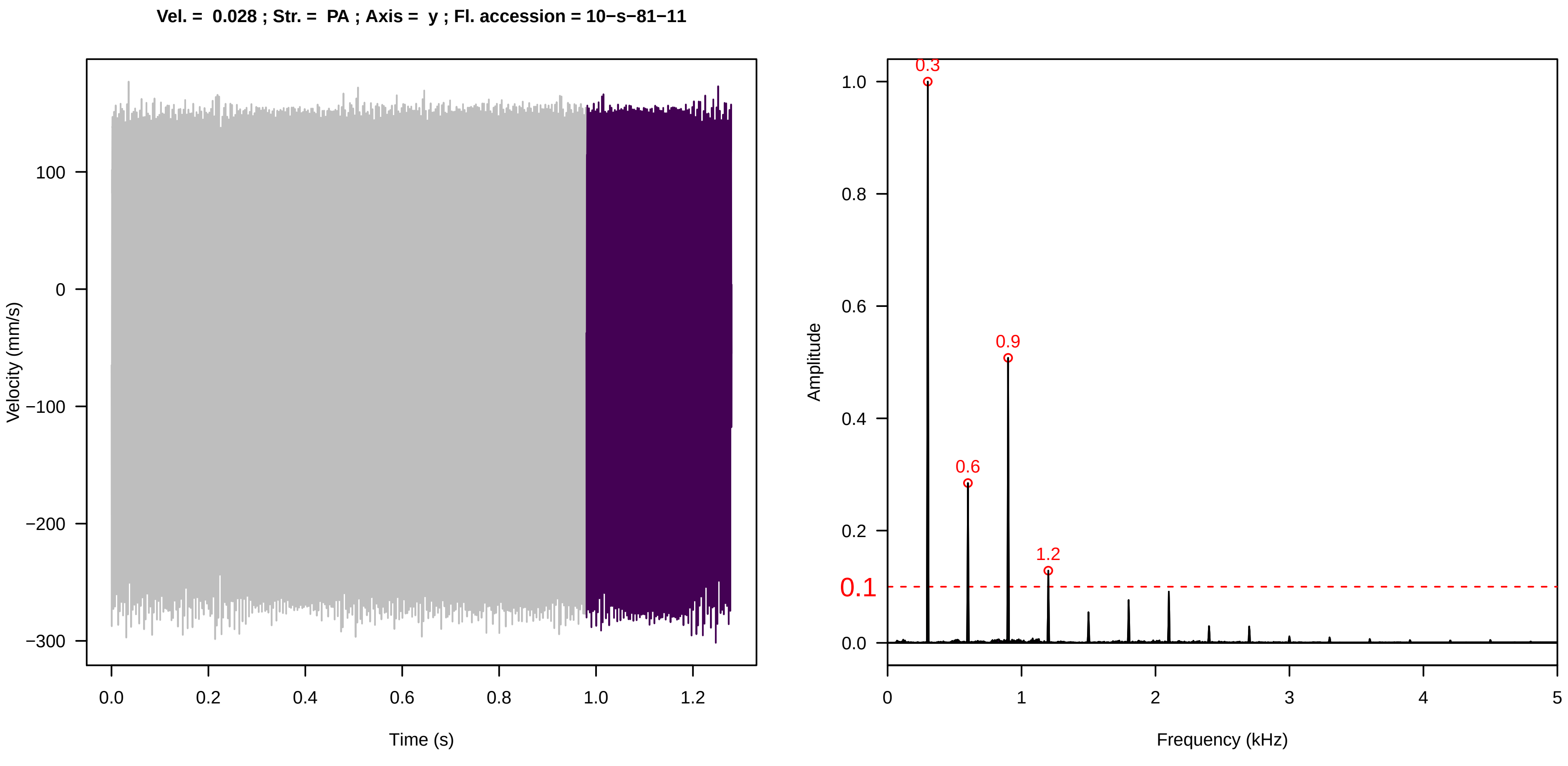
Vel. = 0.028 ; Str. = PA ; Axis = y ; Fl. accession = 10−s−81−11
Velocity (mm/s)
Time (s)
Amplitude
Frequency (kHz)
0.3
0.6
0.9
1.2
0.1

**Vel. = 0.028 ; Str. = Receptacle ; Axis = y ; Fl. accession = 10−s−81−11**

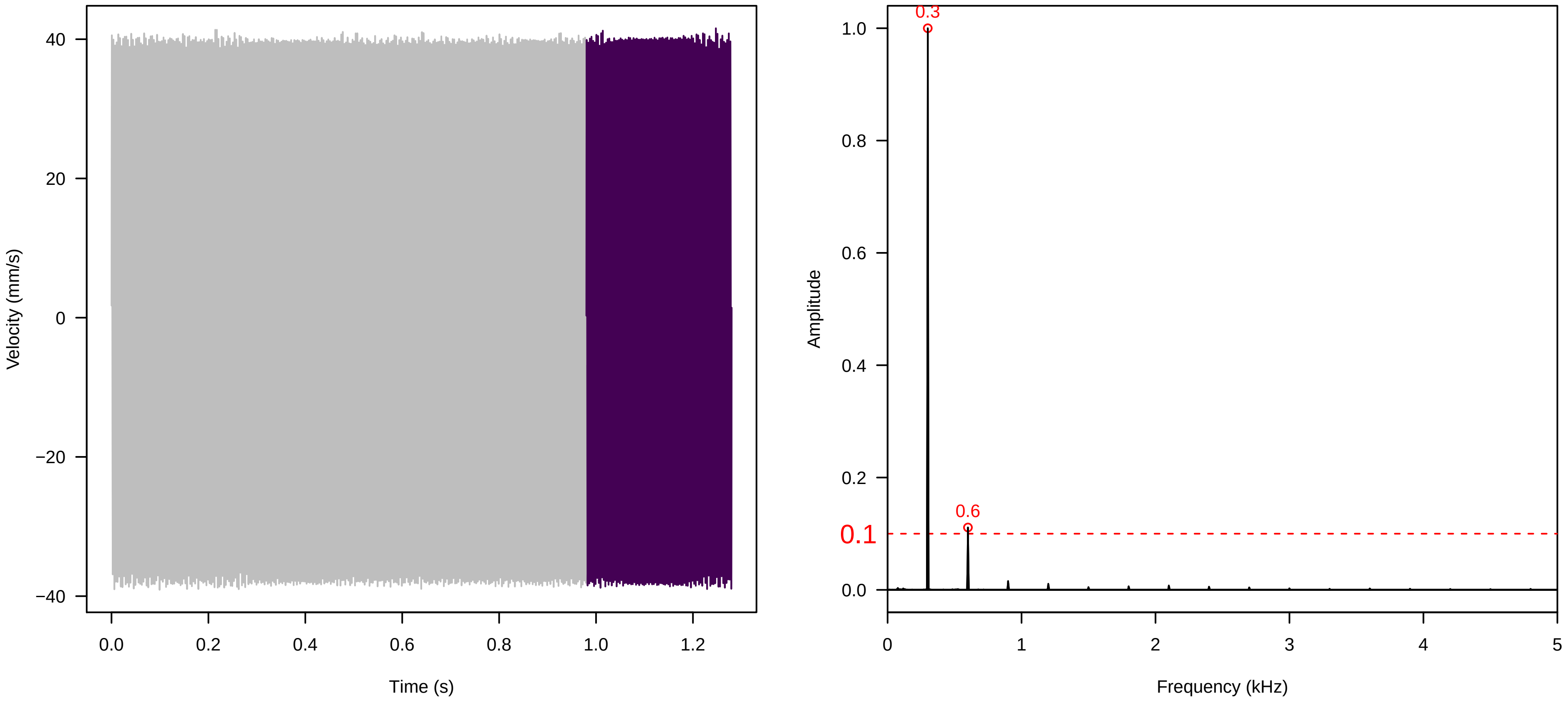


**Vel. = 0.028 ; Str. = FA ; Axis = y ; Fl. accession = 10−s−81−11**

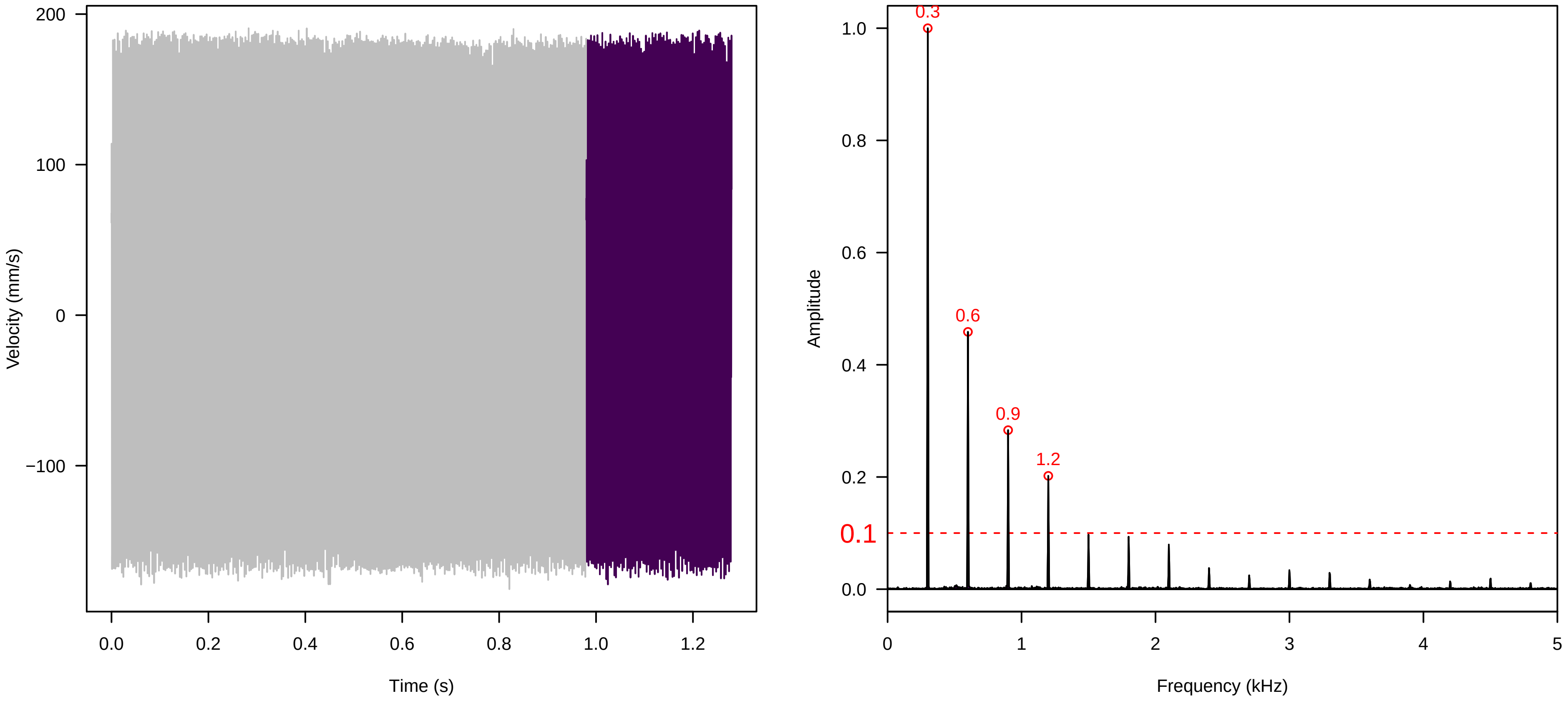


**Vel. = 0.028 ; Str. = Receptacle ; Axis = y ; Fl. accession = 10−s−81−11**

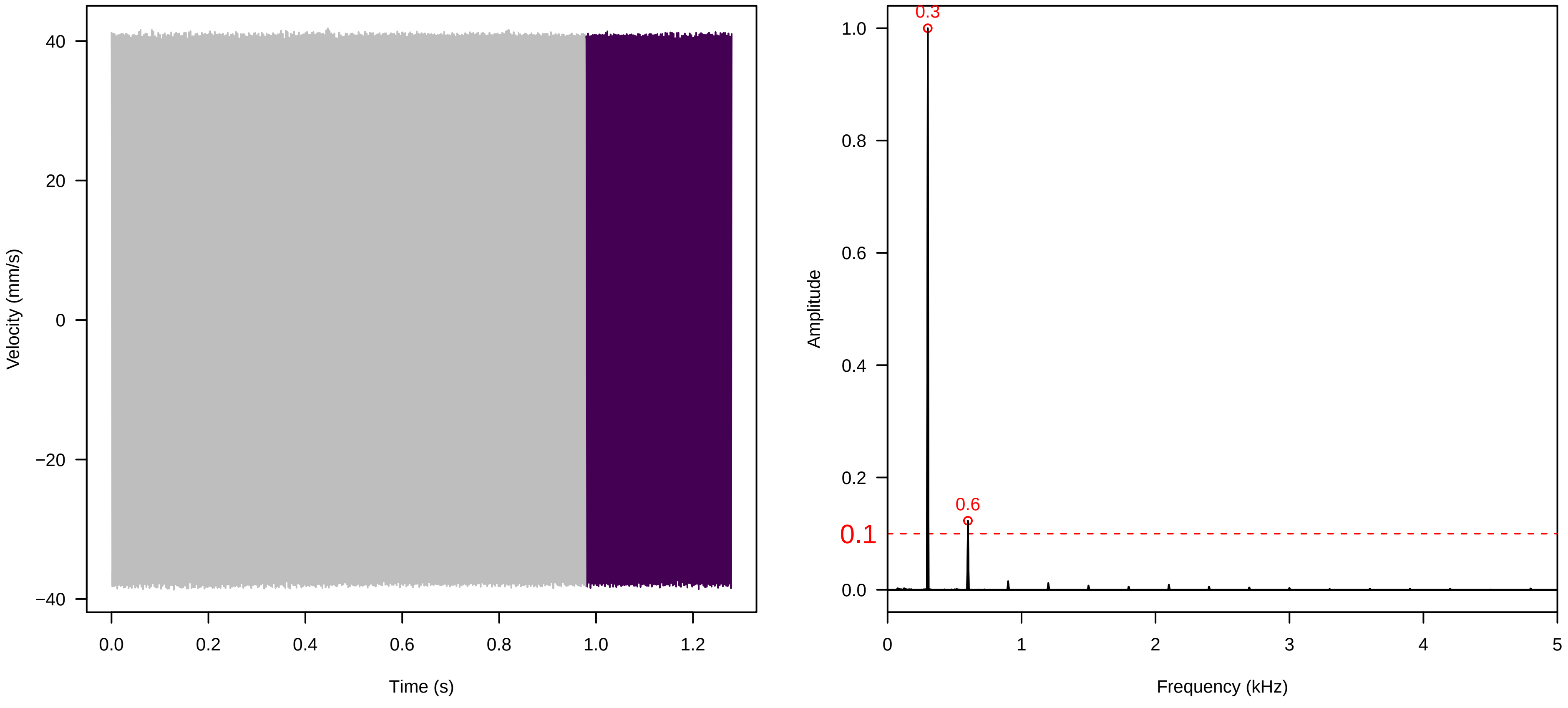

**Vel. = 0.028 ; Str. = Corolla ; Axis = y ; Fl. accession = 10−s−81−11**

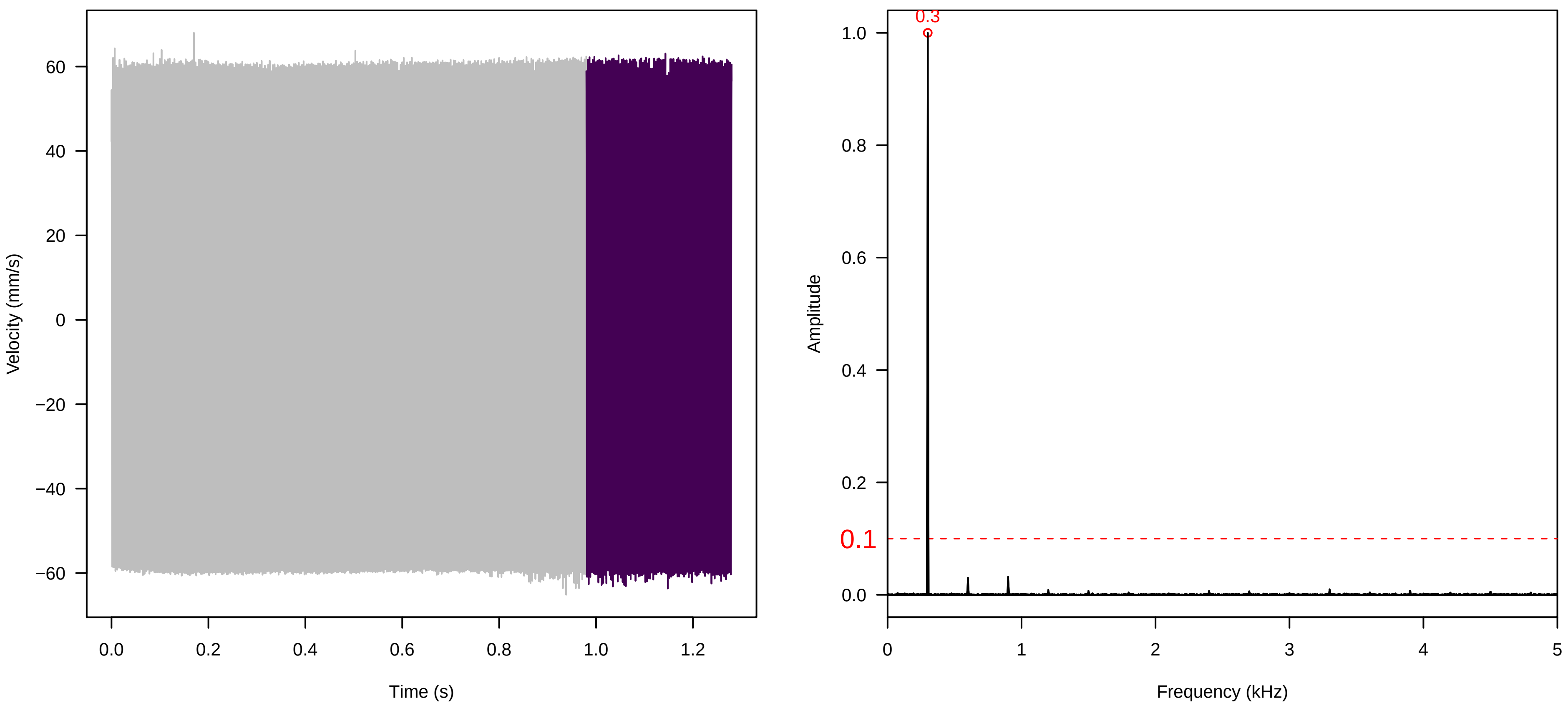


**Vel. = 0.028 ; Str. = Receptacle ; Axis = y ; Fl. accession = 10−s−81−11**

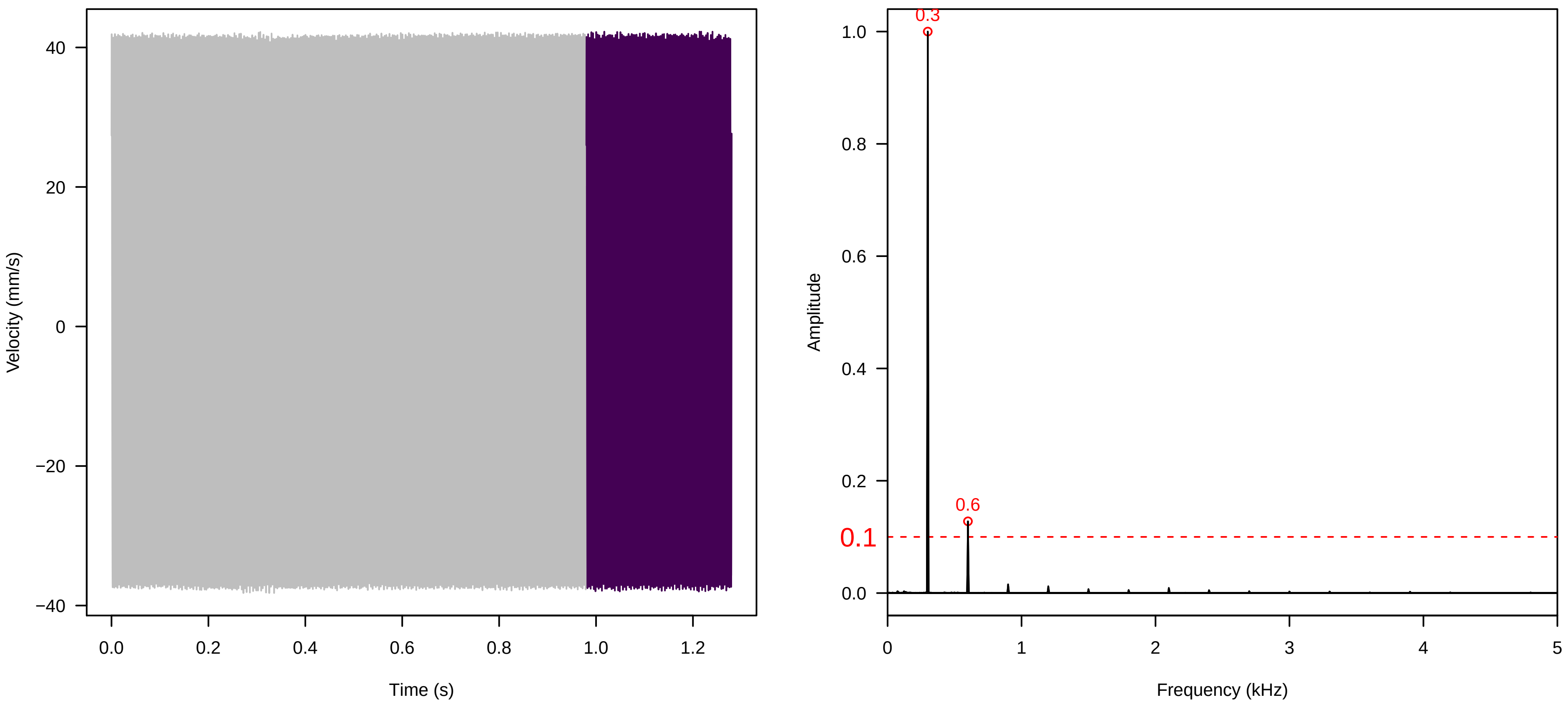


**Vel. = 0.014 ; Str. = Corolla ; Axis = y ; Fl. accession = 10−s−81−11**

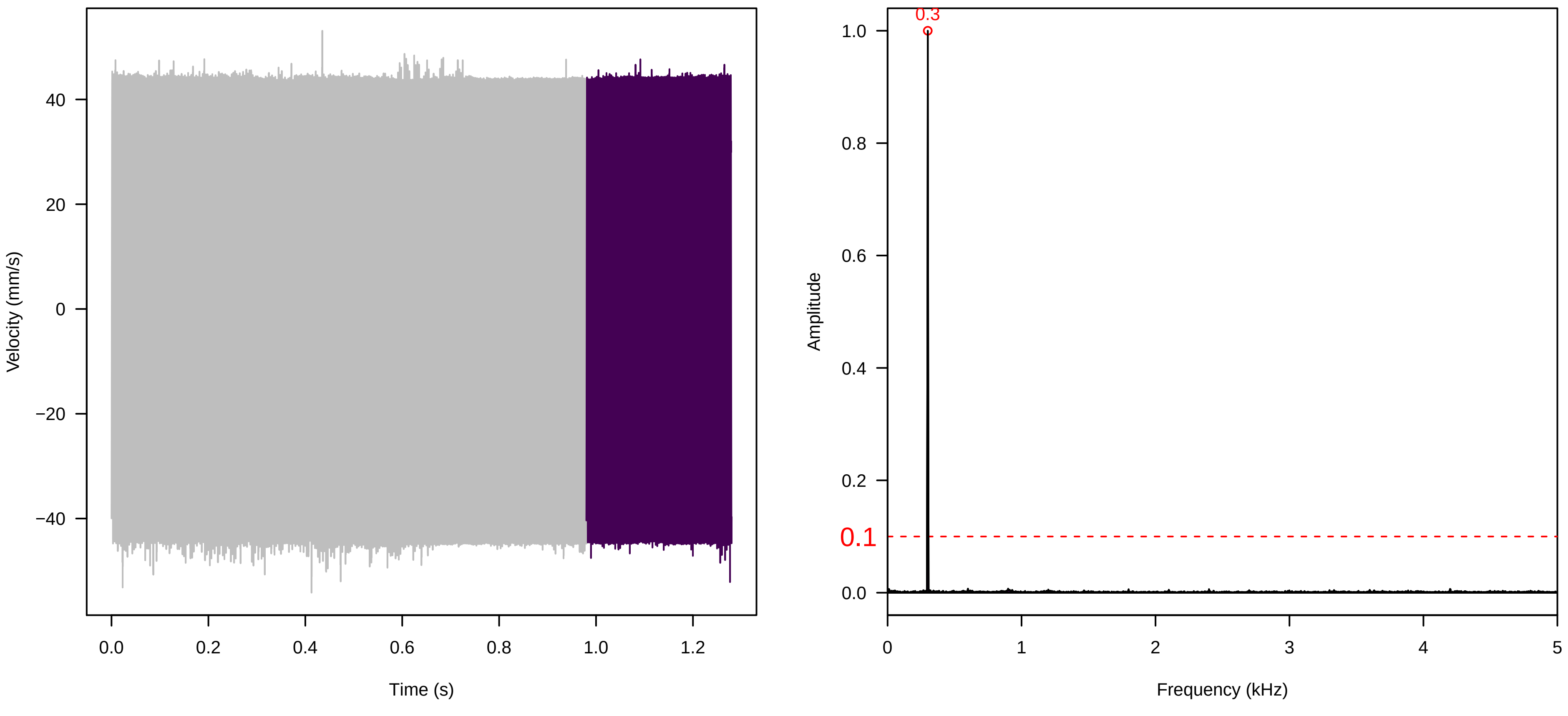

**Vel. = 0.014 ; Str. = Receptacle ; Axis = y ; Fl. accession = 10−s−81−11**

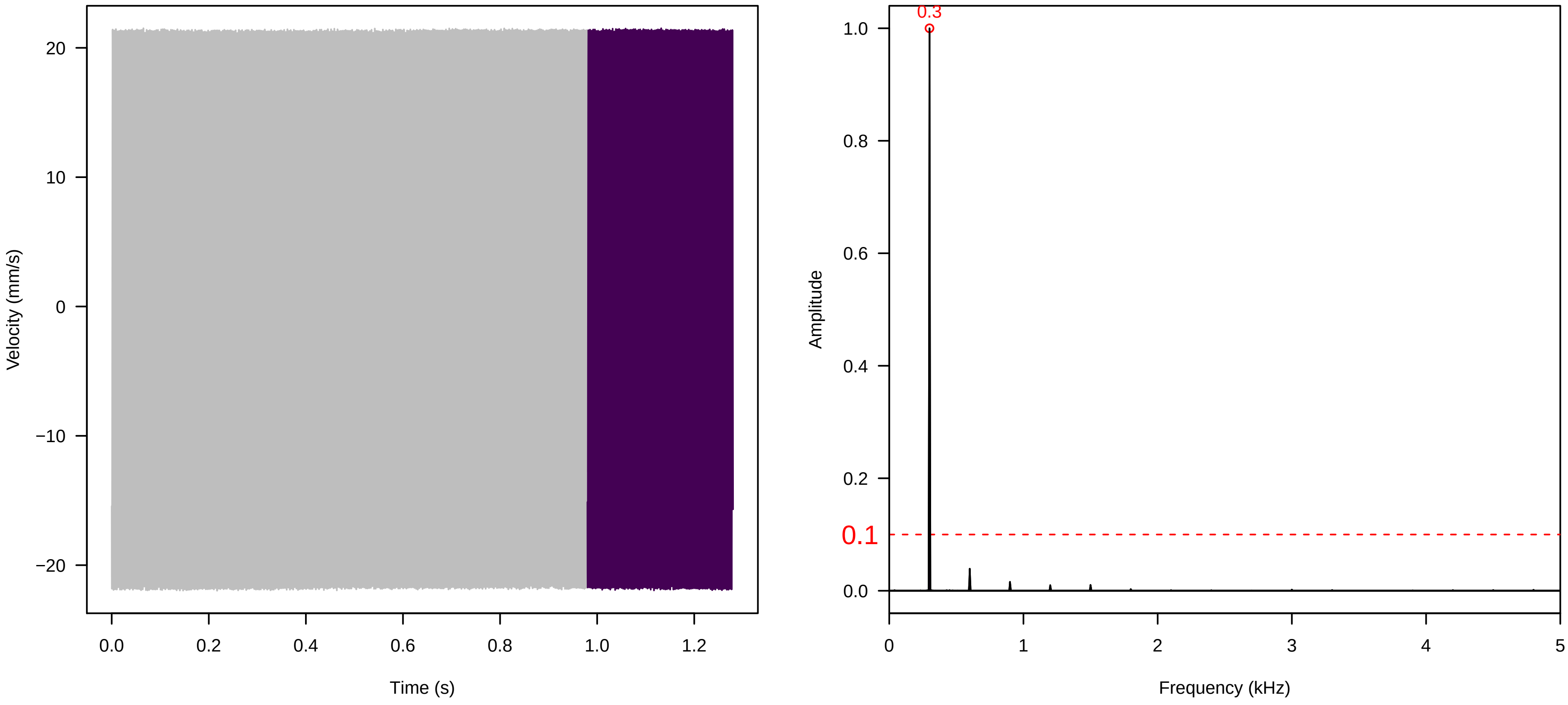


**Vel. = 0.014 ; Str. = FA ; Axis = y ; Fl. accession = 10−s−81−11**

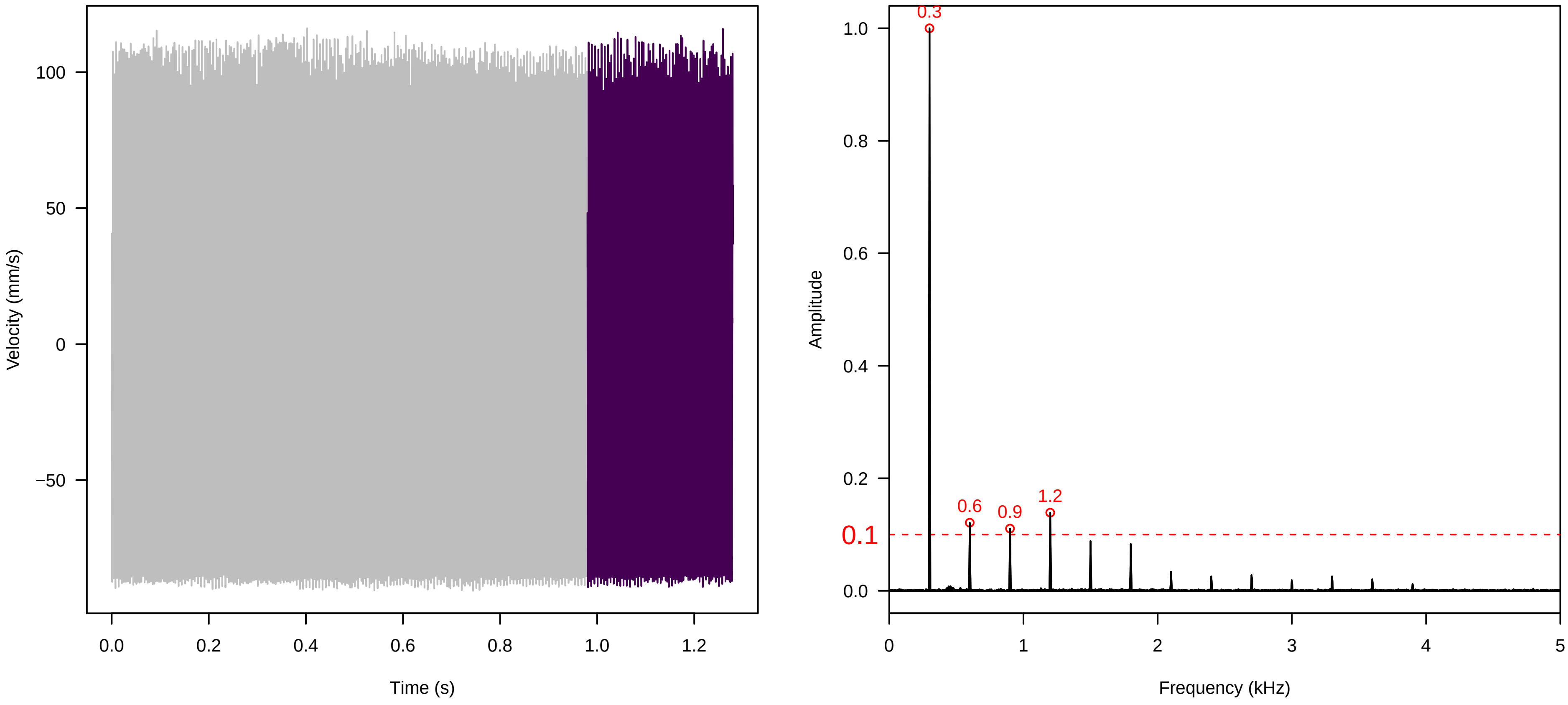


**Vel. = 0.014 ; Str. = Receptacle ; Axis = y ; Fl. accession = 10−s−81−11**

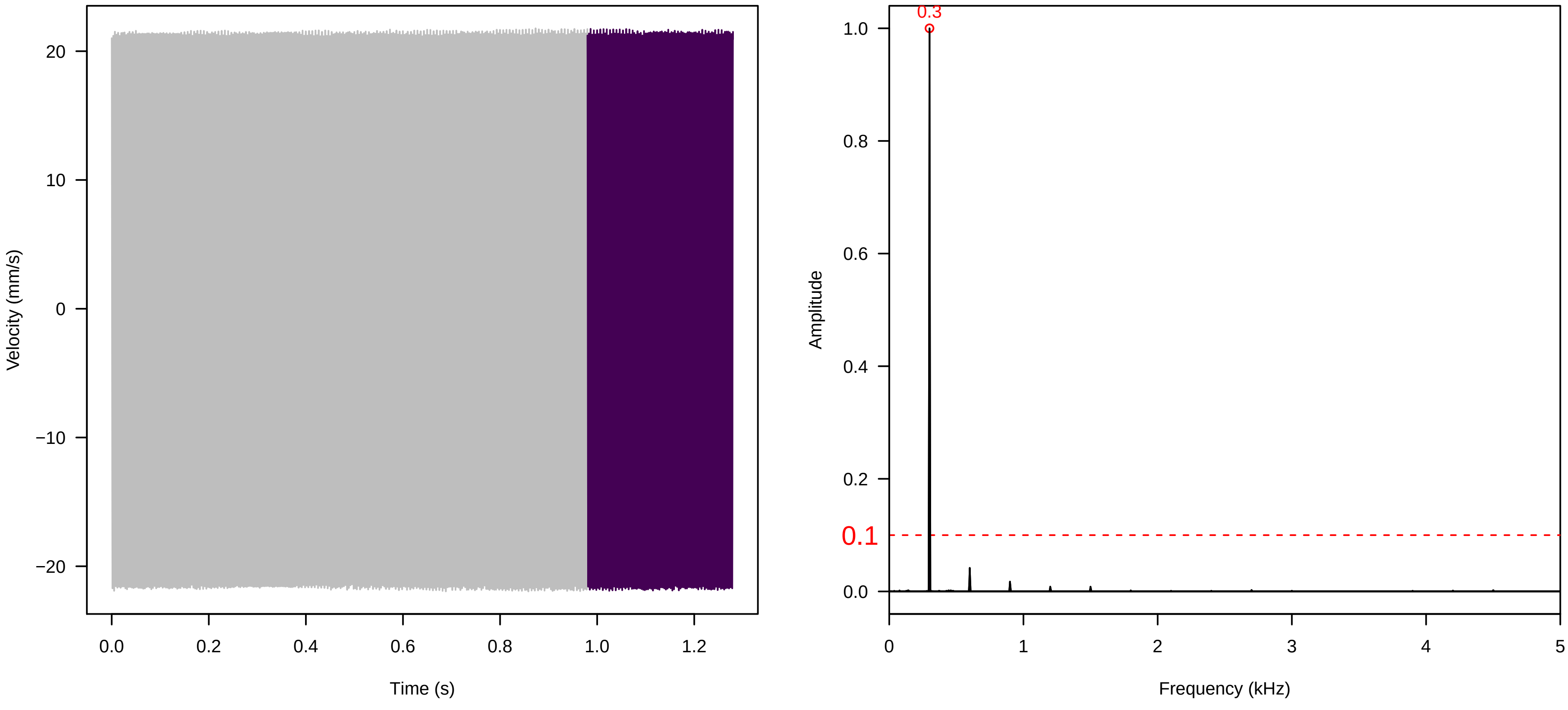

**Vel. = 0.014 ; Str. = PA ; Axis = y ; Fl. accession = 10−s−81−11**

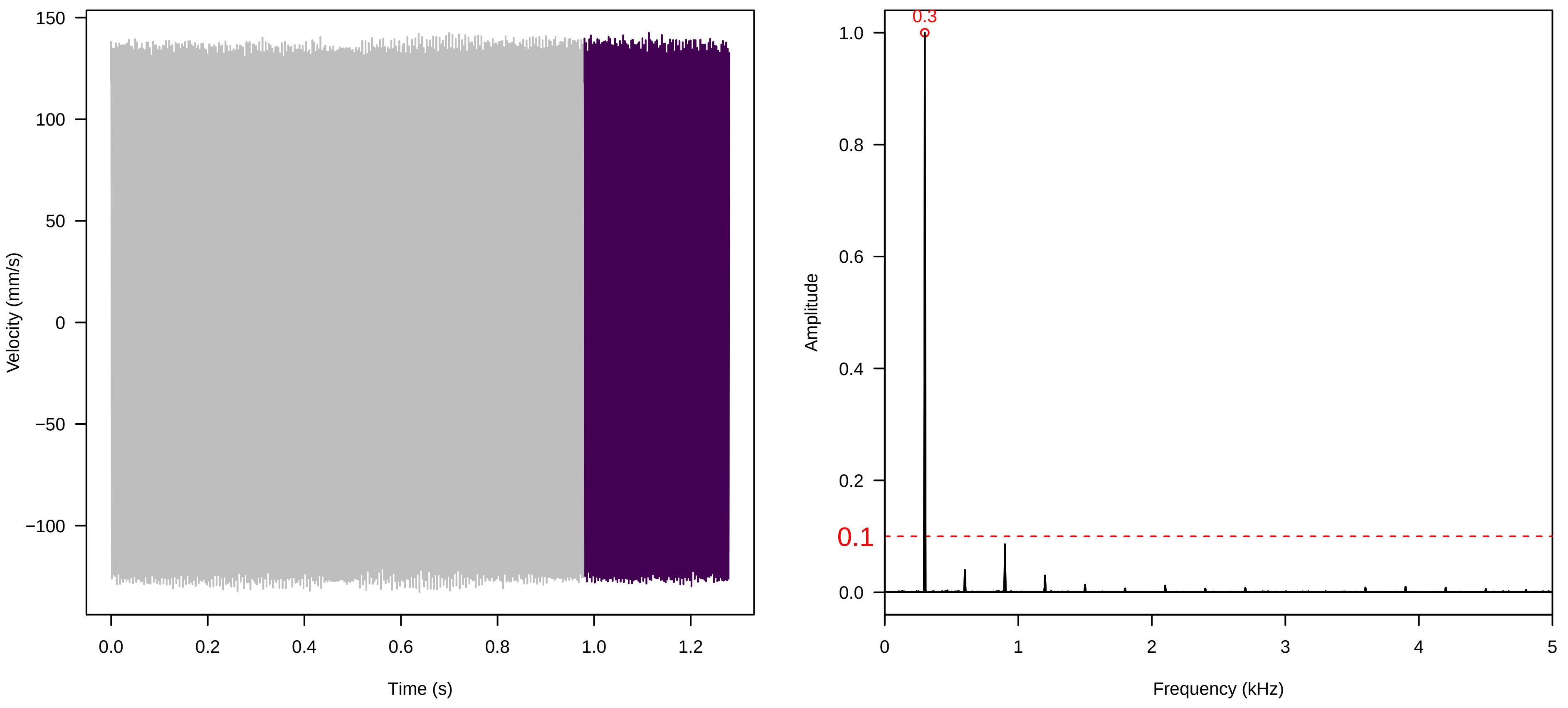


**Vel. = 0.014 ; Str. = Receptacle ; Axis = y ; Fl. accession = 10−s−81−11**

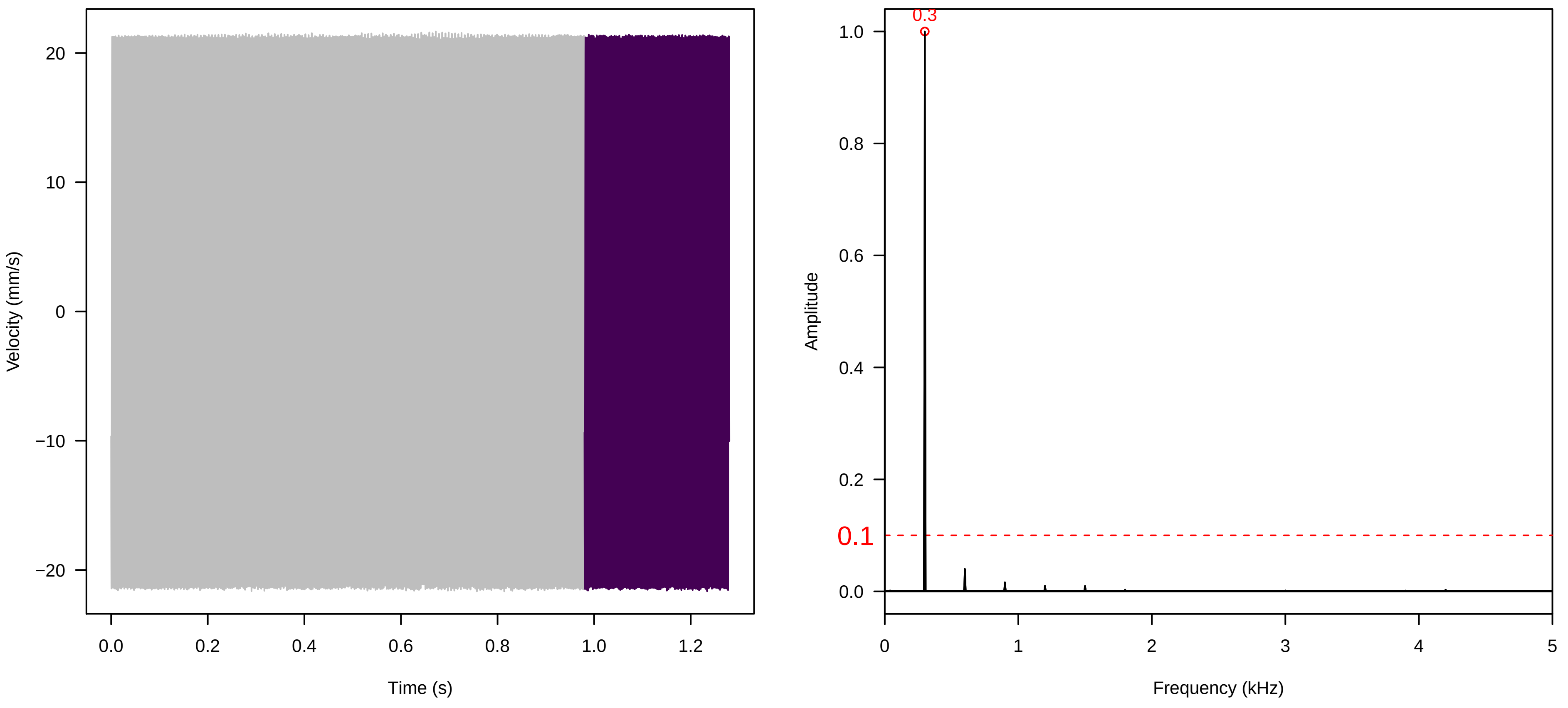


**Vel. = 0.014 ; Str. = Corolla ; Axis = z ; Fl. accession = 10−s−81−11**

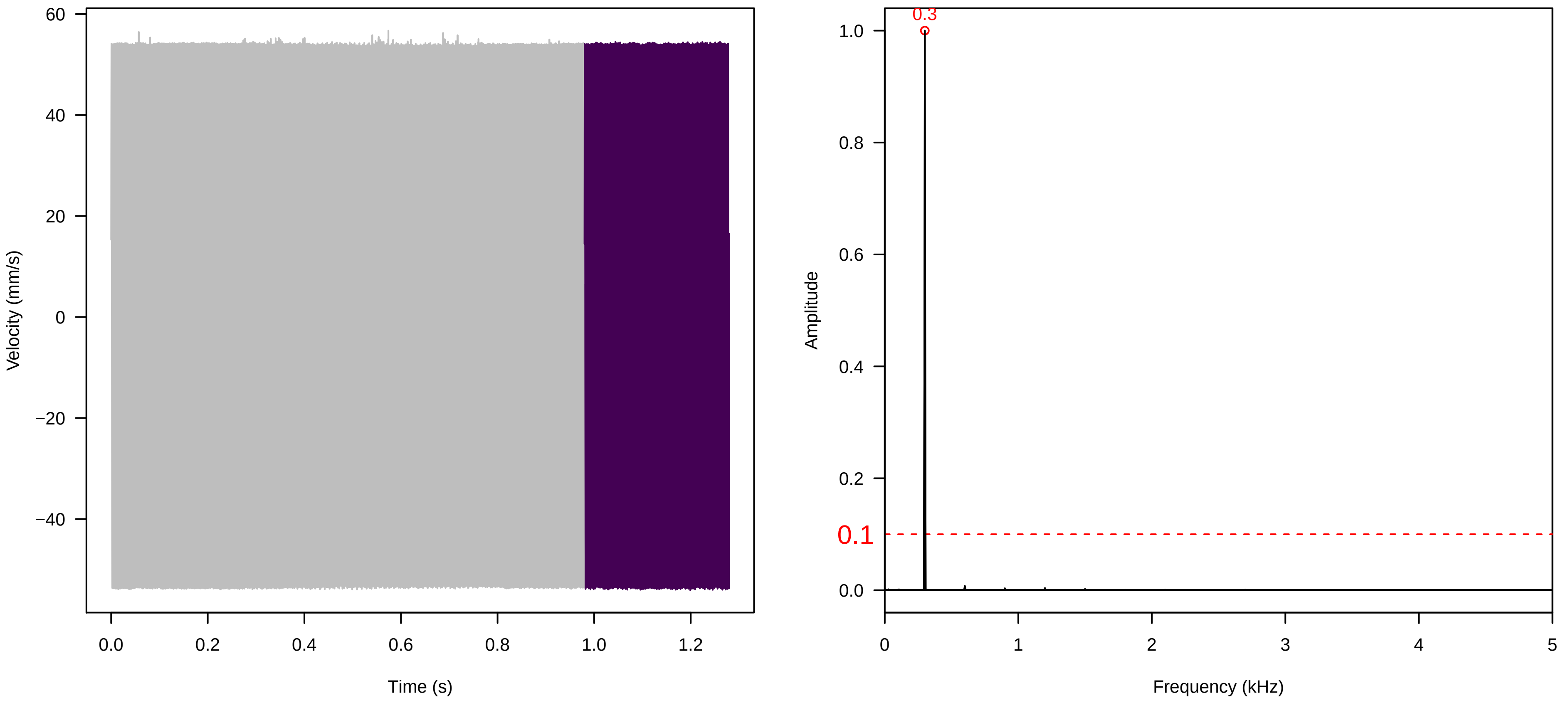

**Vel. = 0.014 ; Str. = Receptacle ; Axis = z ; Fl. accession = 10−s−81−11**

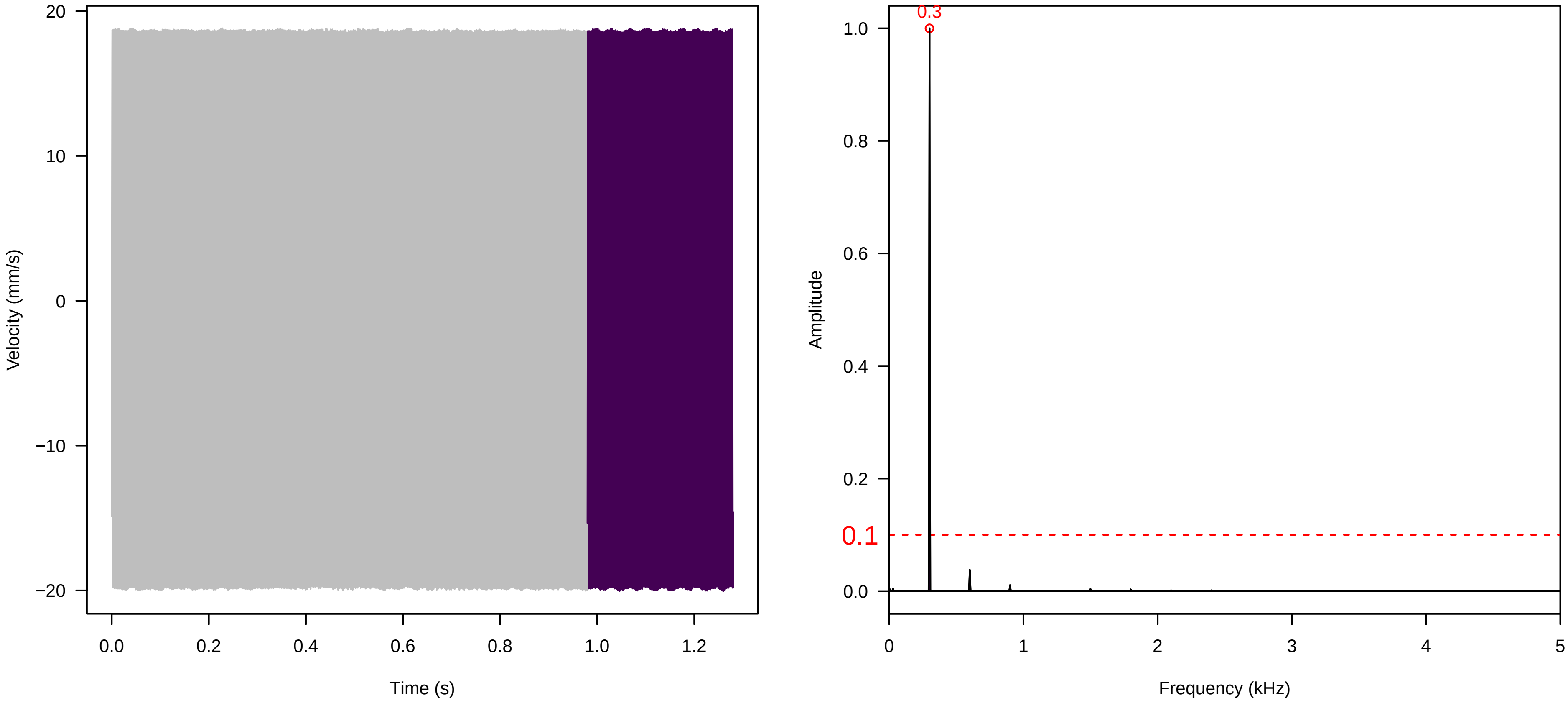


**Vel. = 0.014 ; Str. = FA ; Axis = z ; Fl. accession = 10−s−81−11**

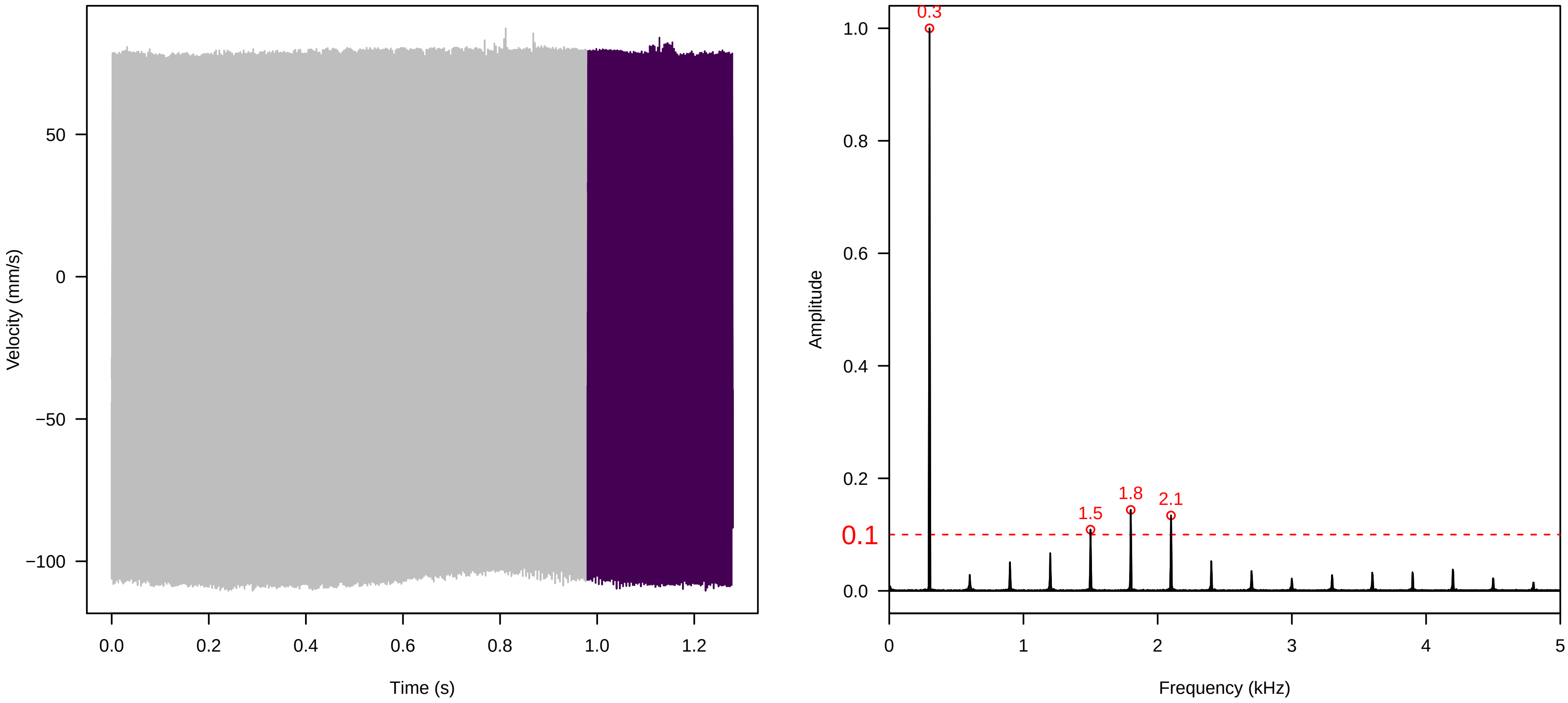


**Vel. = 0.014 ; Str. = Receptacle ; Axis = z ; Fl. accession = 10−s−81−11**

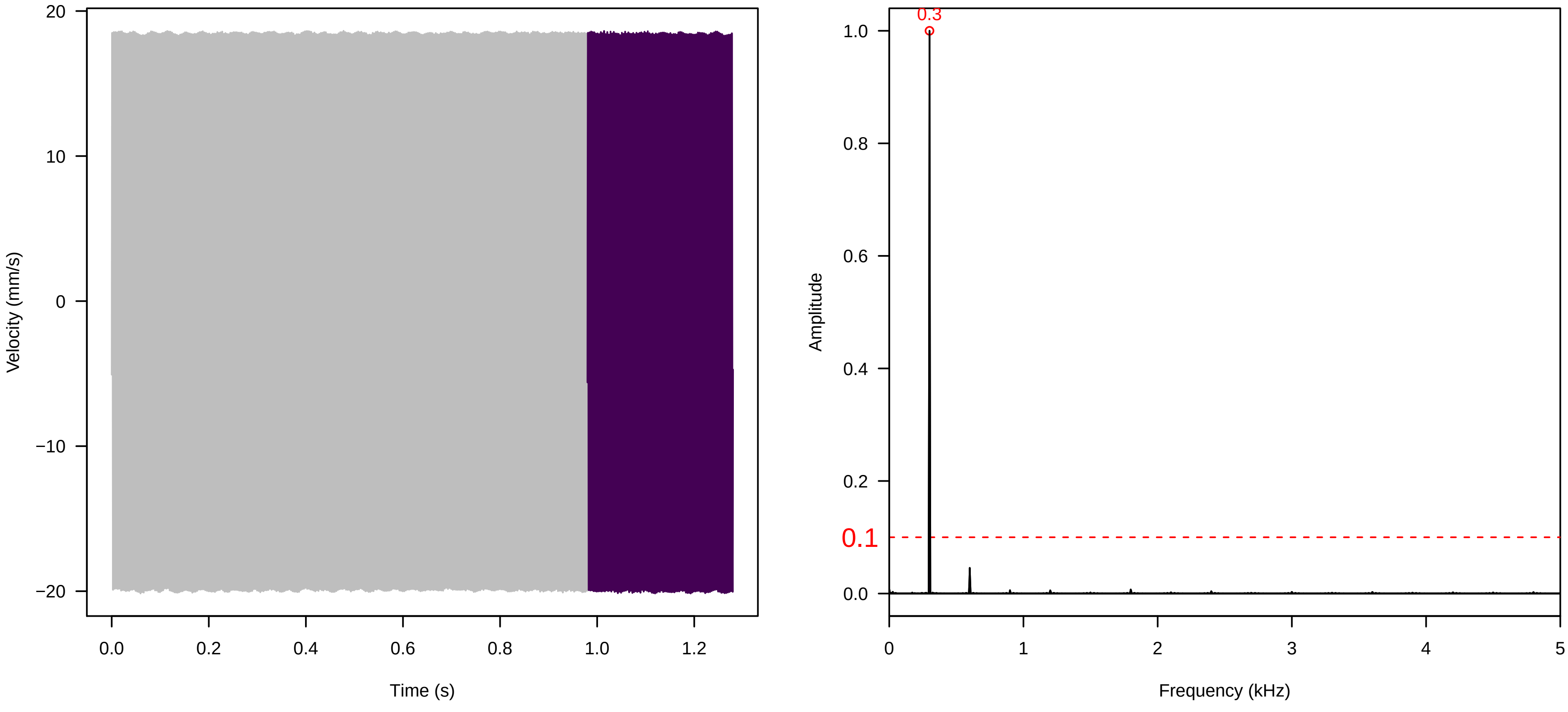

**Vel. = 0.014 ; Str. = PA ; Axis = z ; Fl. accession = 10−s−81−11**

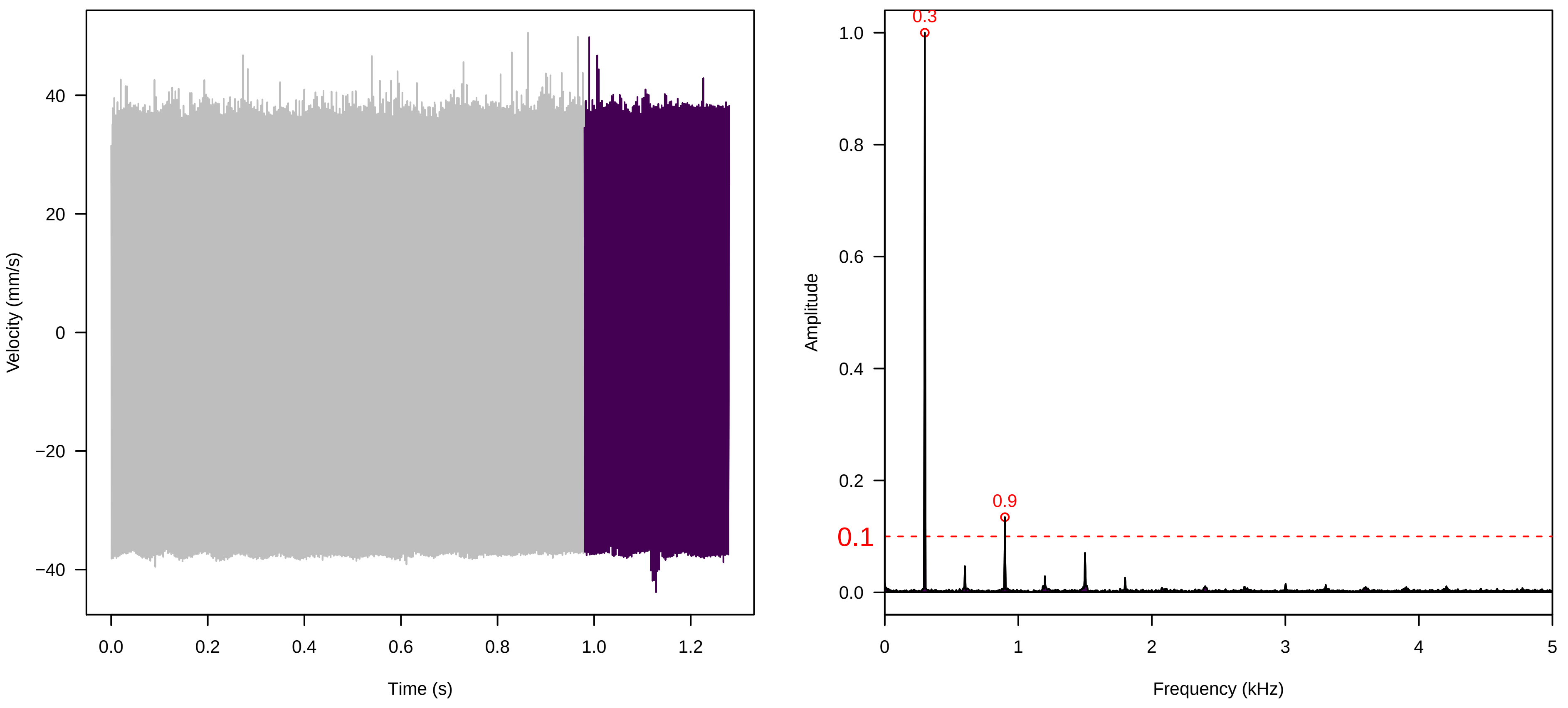


**Vel. = 0.014 ; Str. = Receptacle ; Axis = z ; Fl. accession = 10−s−81−11**

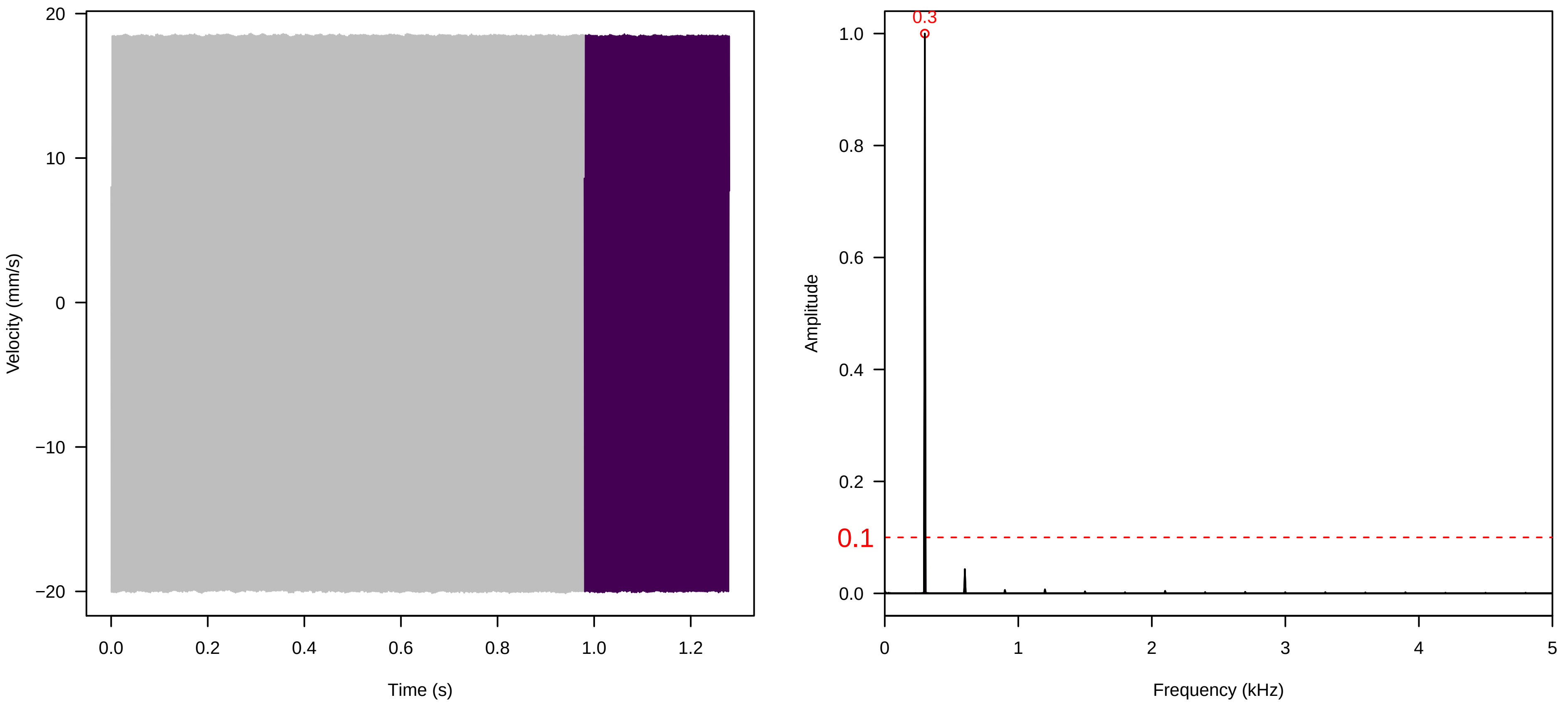


**Vel. = 0.028 ; Str. = PA ; Axis = z ; Fl. accession = 10−s−81−11**

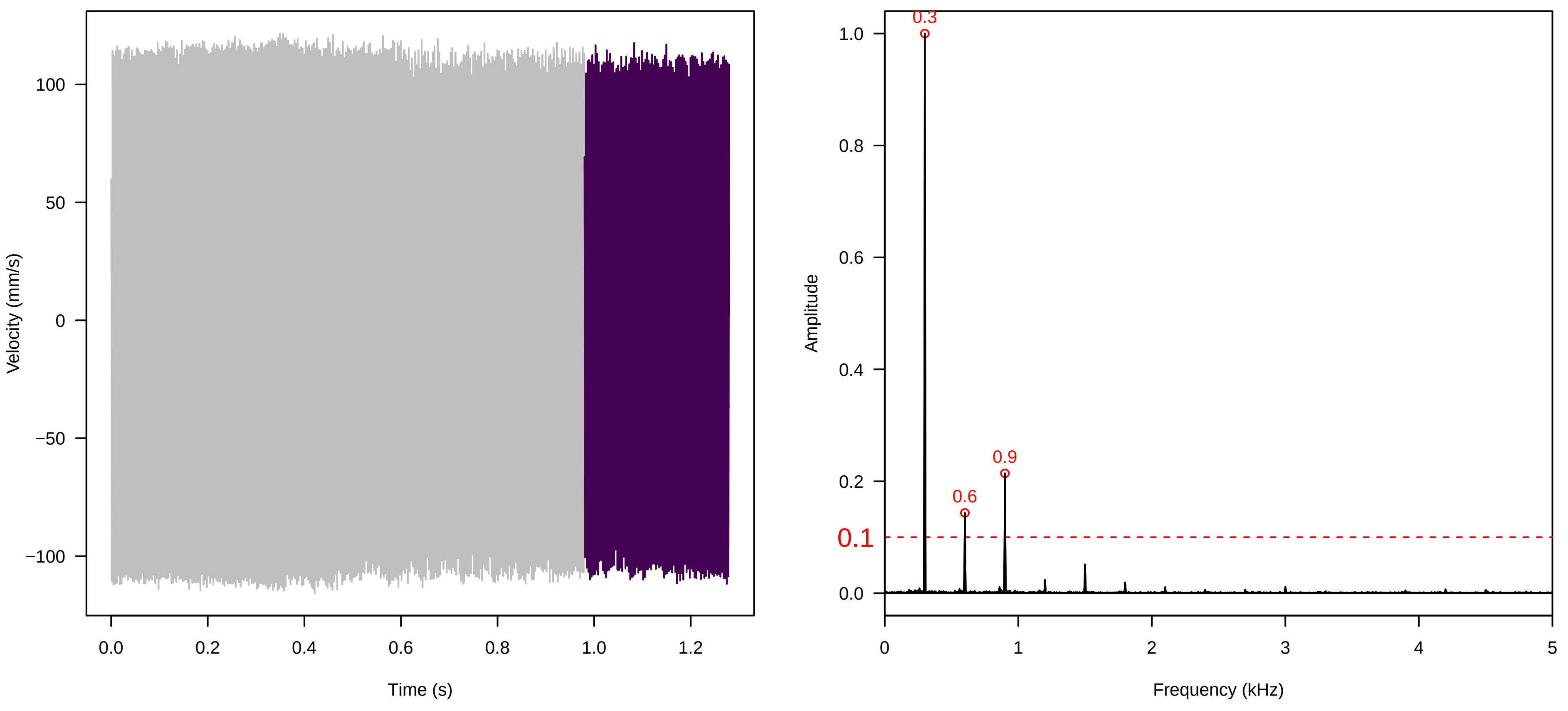

**Vel. = 0.028 ; Str. = Receptacle ; Axis = z ; Fl. accession = 10−s−81−11**

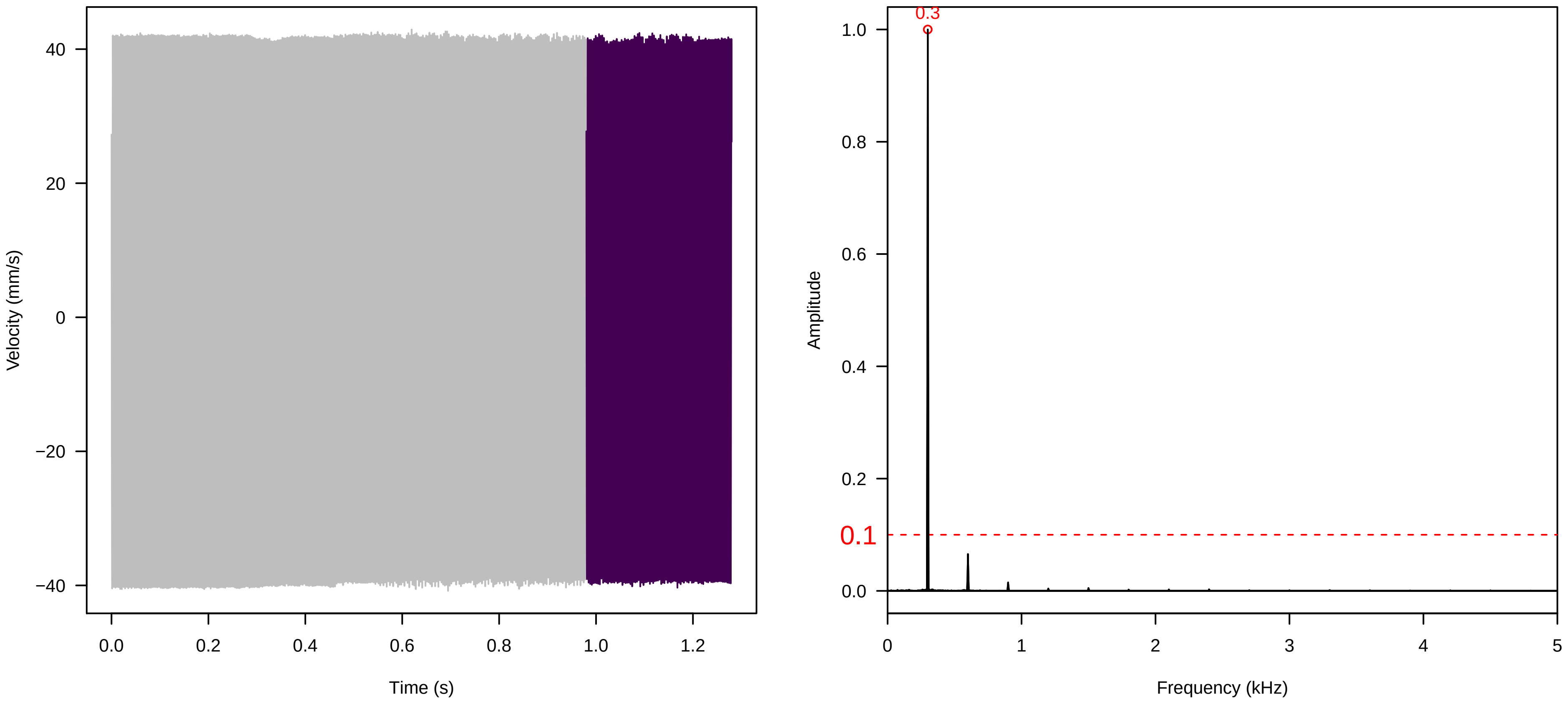


**Vel. = 0.028 ; Str. = FA ; Axis = z ; Fl. accession = 10−s−81−11**

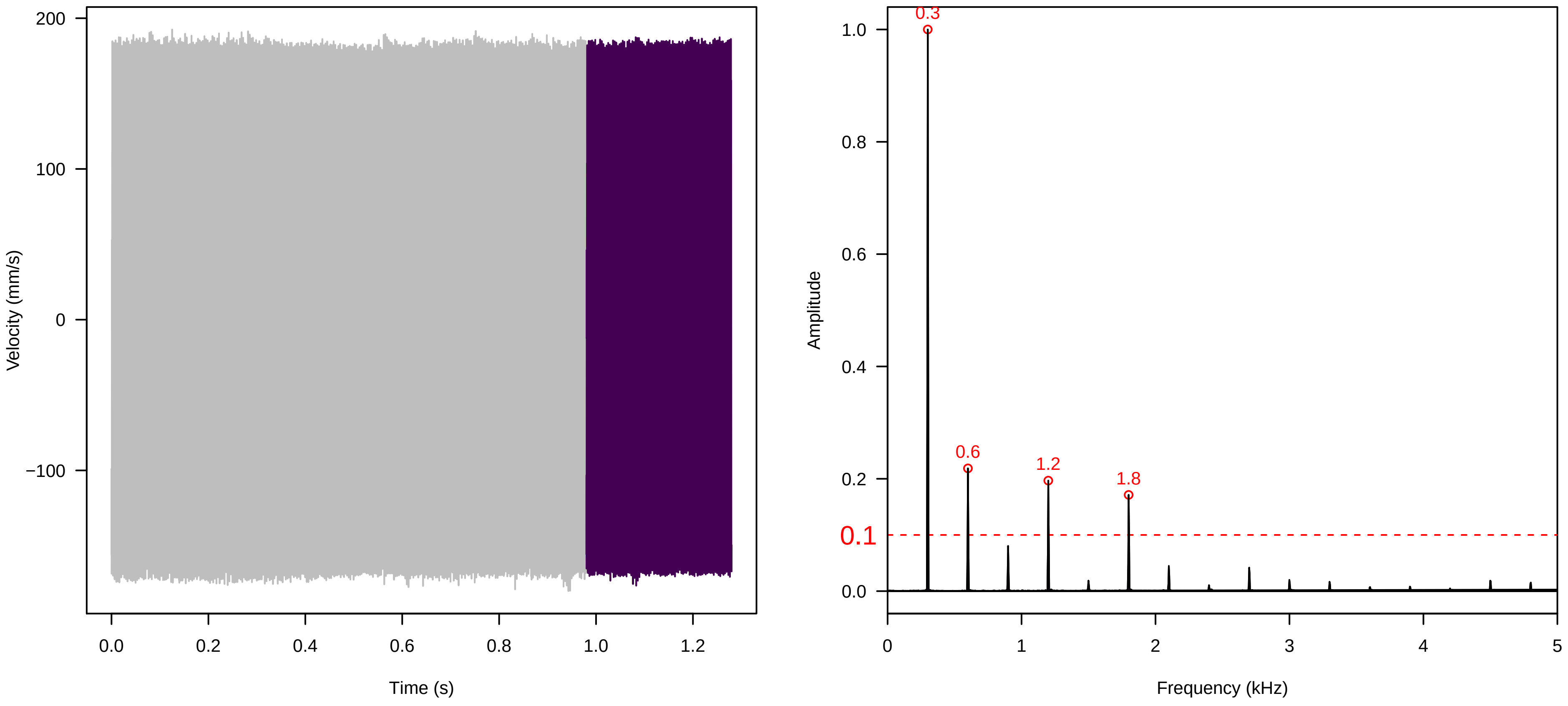


**Vel. = 0.028 ; Str. = Receptacle ; Axis = z ; Fl. accession = 10−s−81−11**

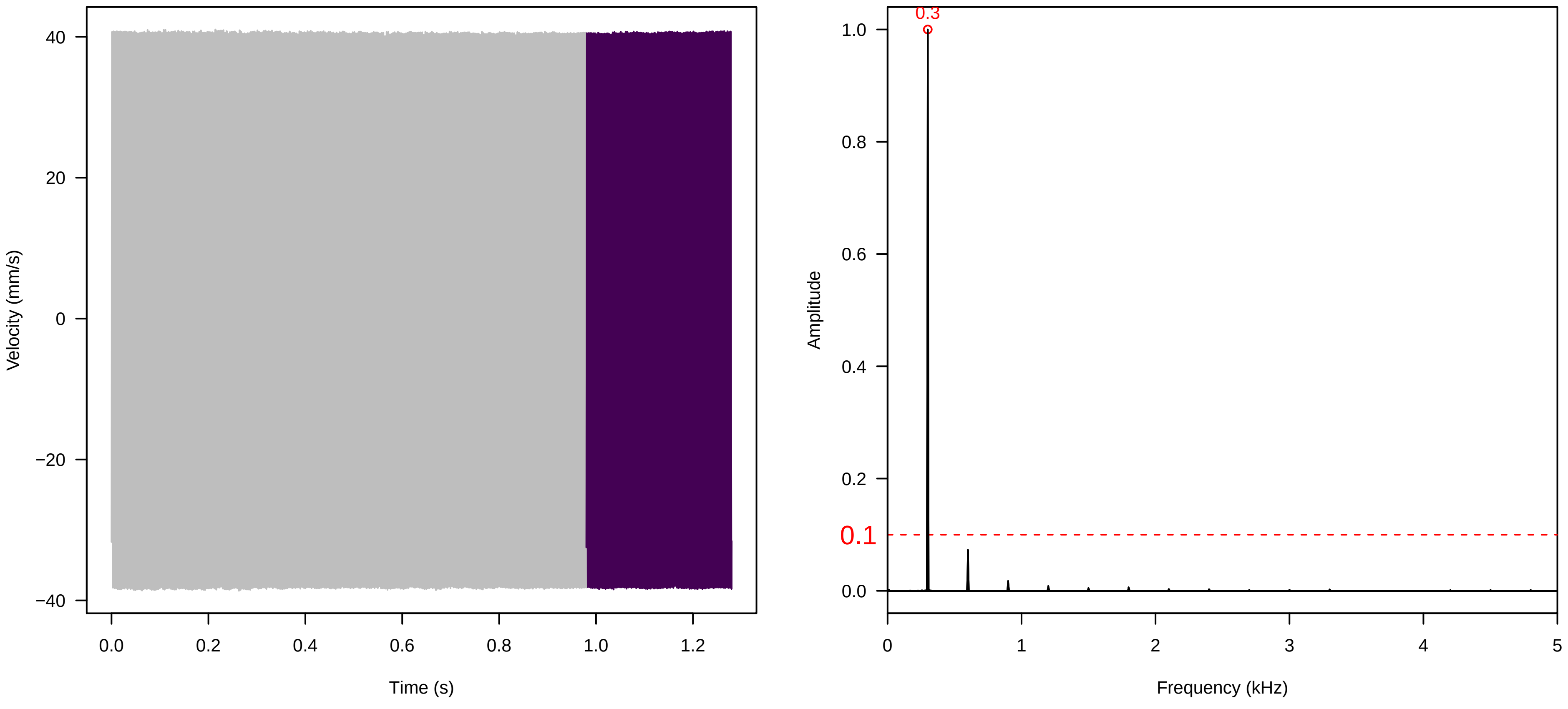

**Vel. = 0.028 ; Str. = Corolla ; Axis = z ; Fl. accession = 10−s−81−11**

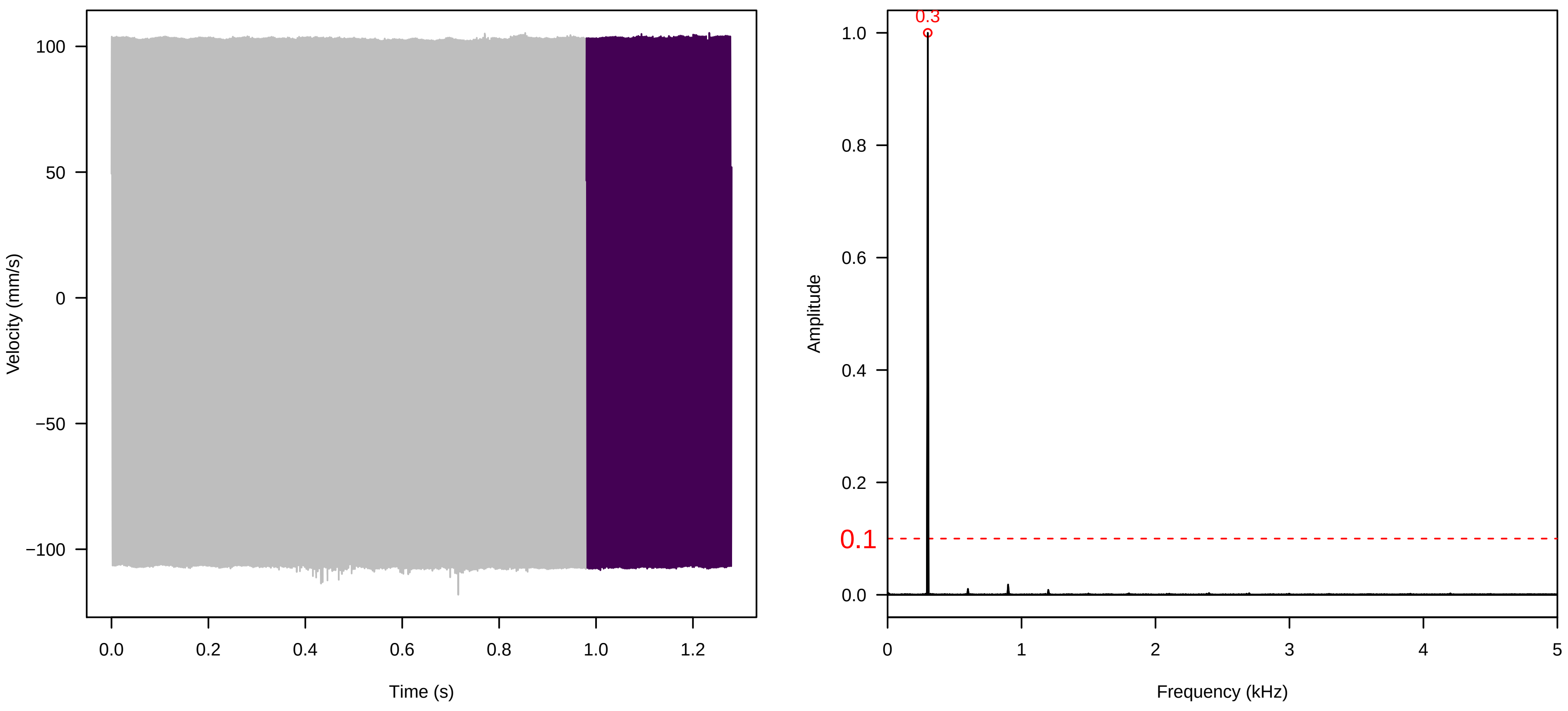


**Vel. = 0.028 ; Str. = Receptacle ; Axis = z ; Fl. accession = 10−s−81−11**

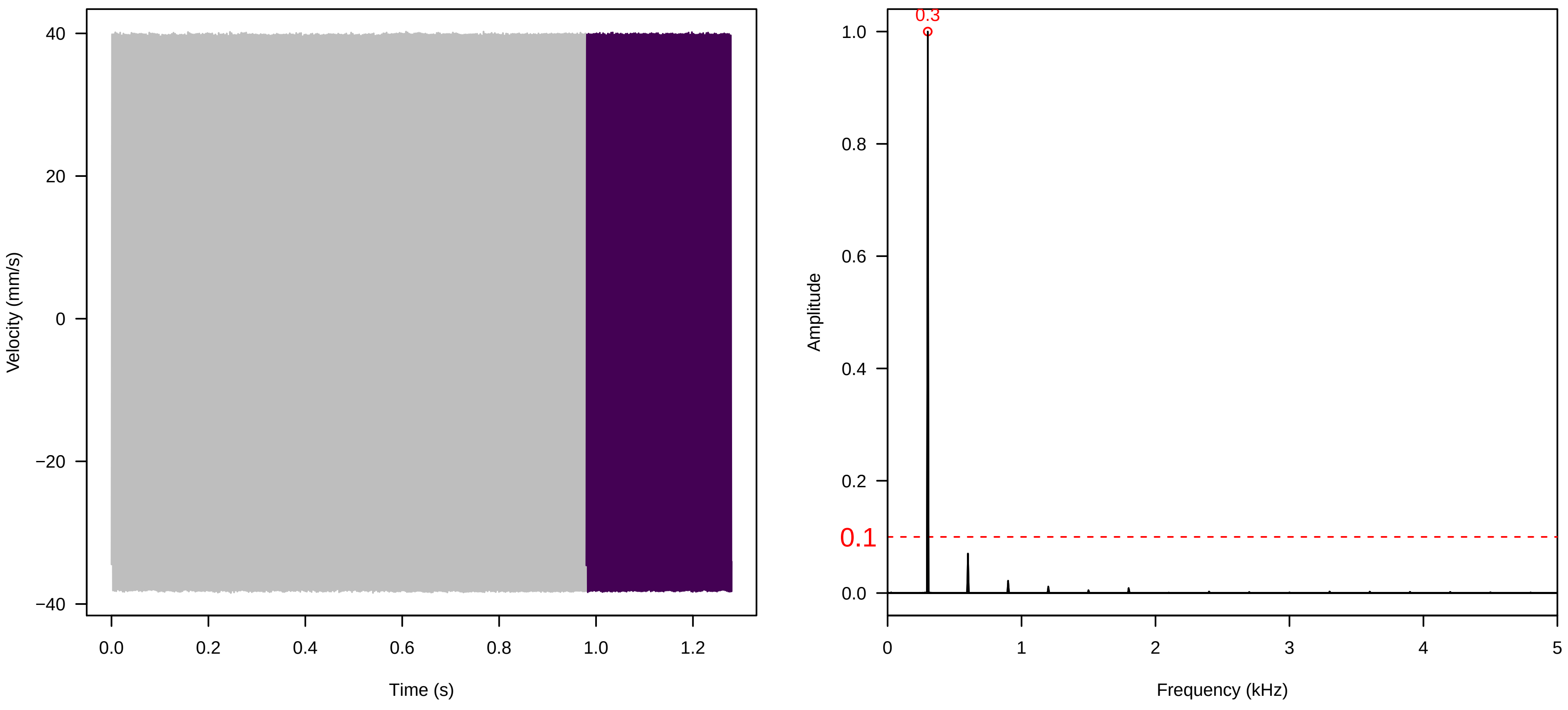


**Vel. = 0.057 ; Str. = Corolla ; Axis = z ; Fl. accession = 10−s−81−11**

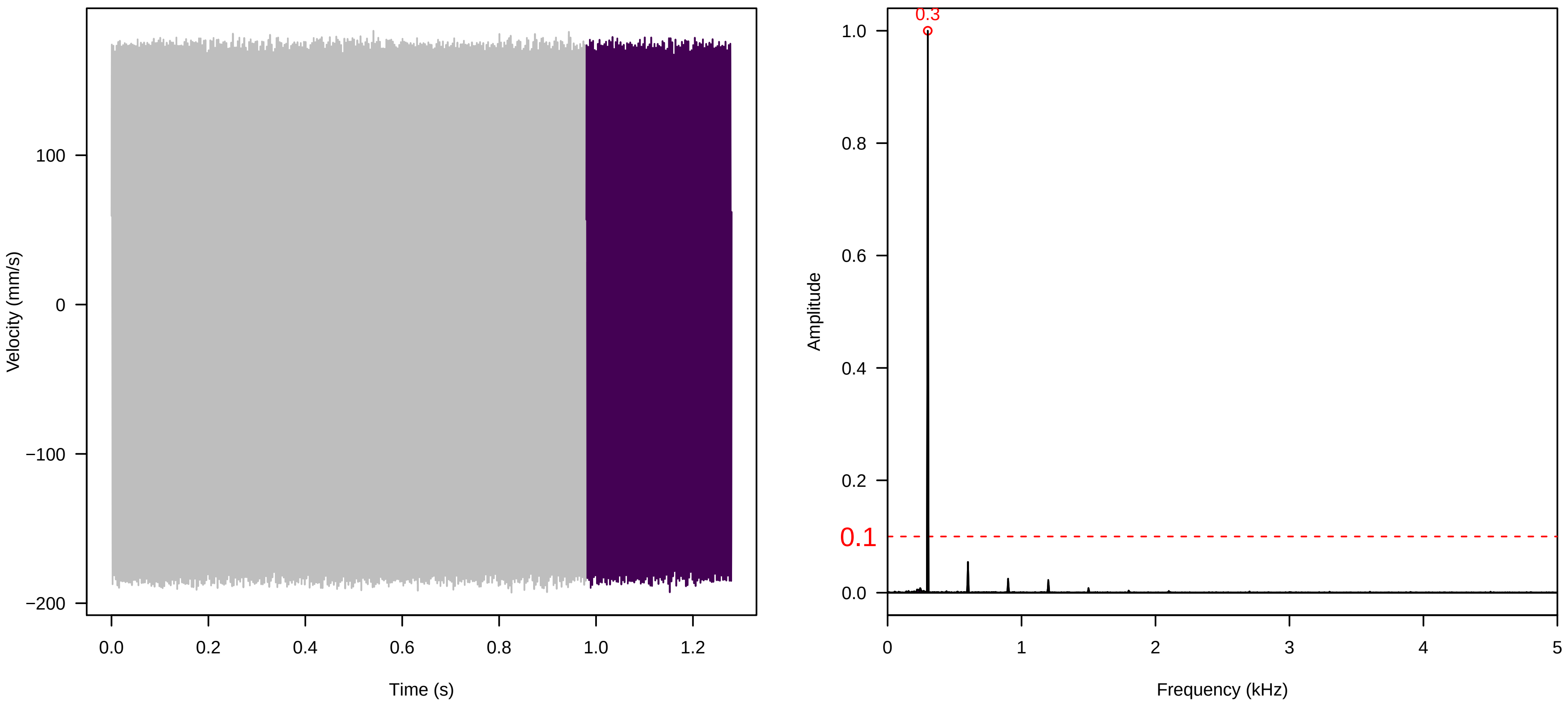

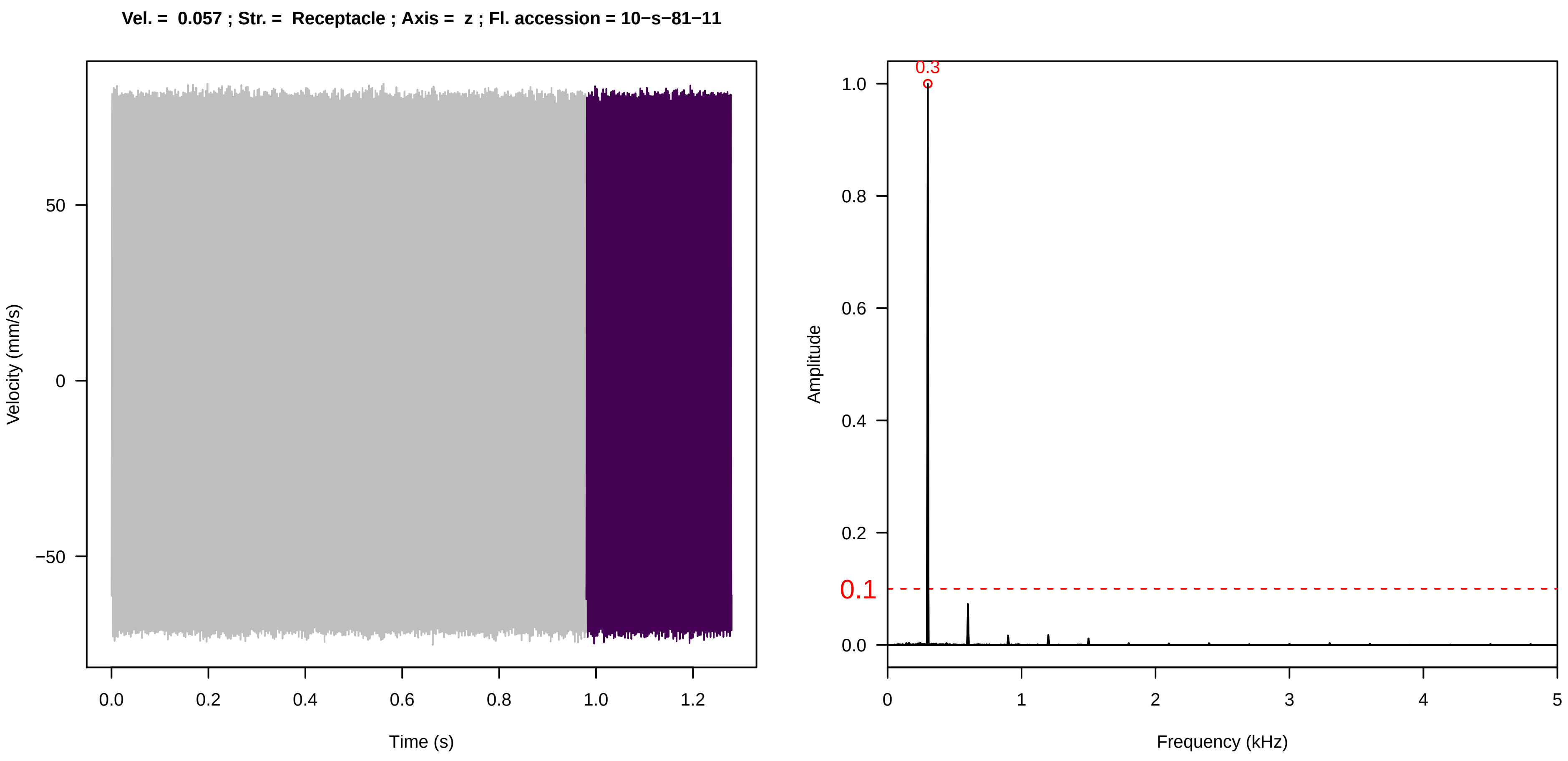

Vel. = 0.057 ; Str. = Receptacle ; Axis = z ; Fl. accession = 10−s−81−11
Velocity (mm/s)
Time (s)
Amplitude
Frequency (kHz)
0.3
0.1


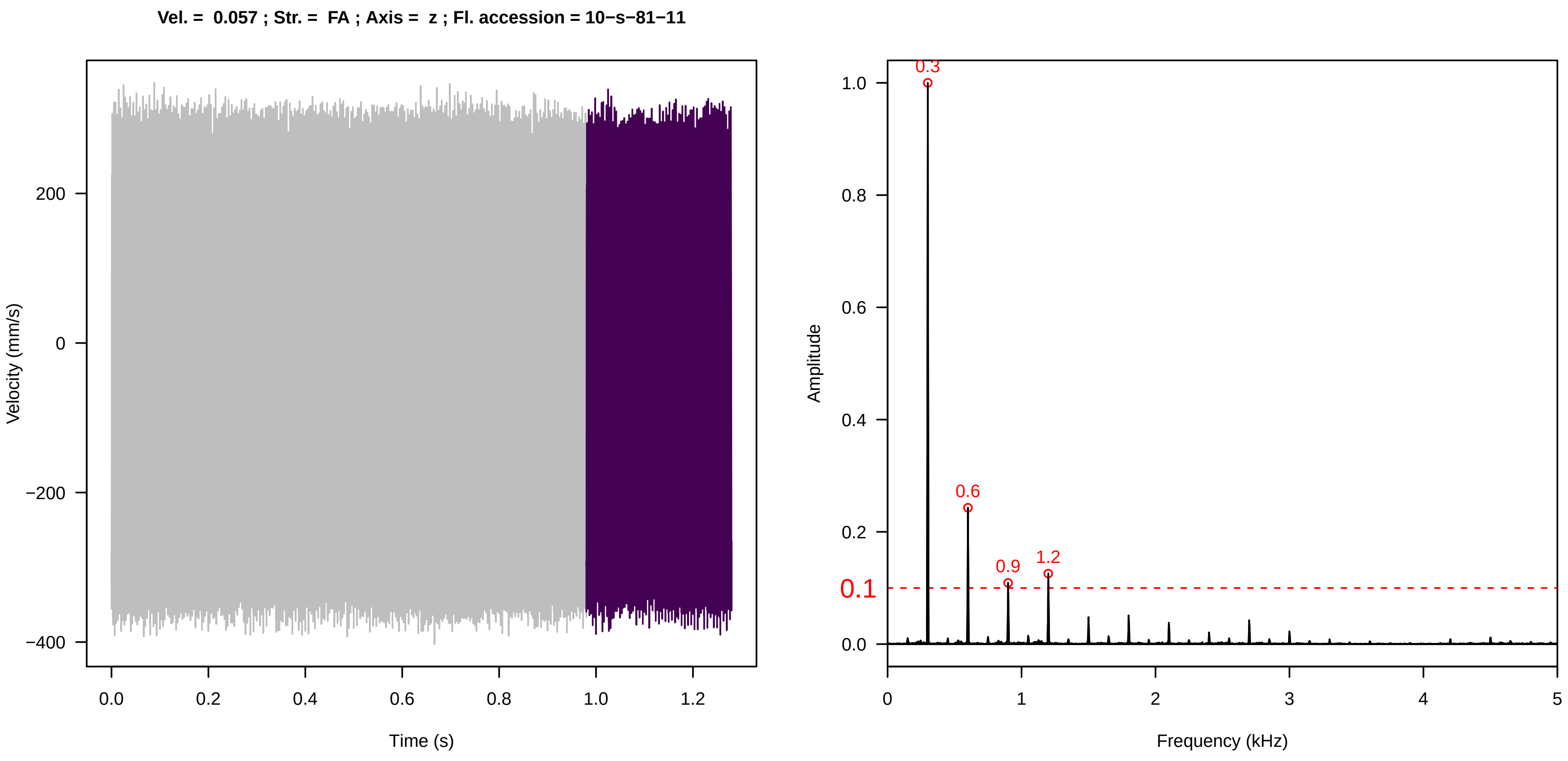

Vel. = 0.057 ; Str. = FA ; Axis = z ; Fl. accession = 10−s−81−11
Velocity (mm/s)
Time (s)
Amplitude
Frequency (kHz)
0.3
0.6
0.9
1.2
0.1


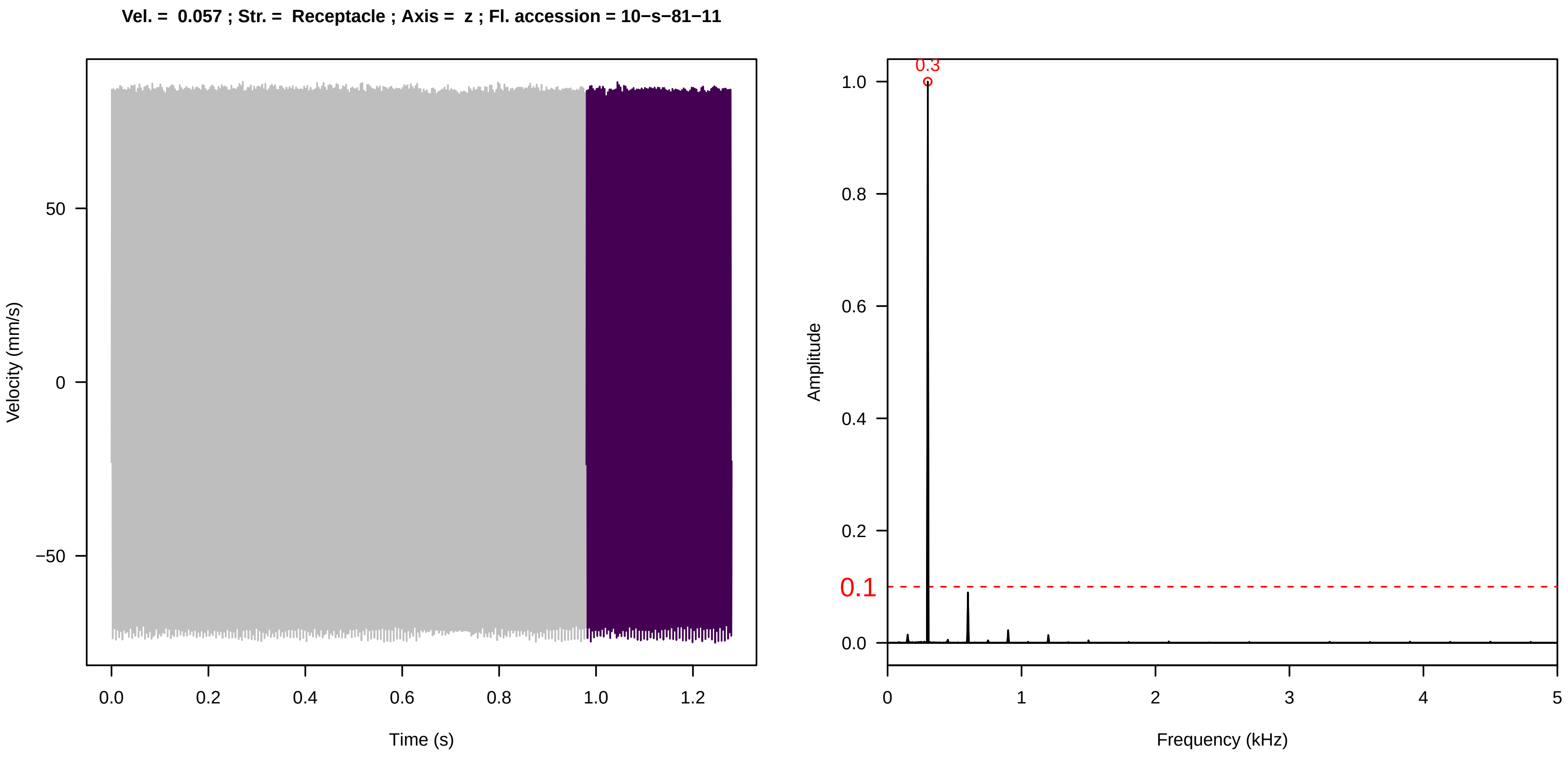

Vel. = 0.057 ; Str. = Receptacle ; Axis = z ; Fl. accession = 10−s−81−11
Velocity (mm/s)
Time (s)
Amplitude
Frequency (kHz)
0.3
0.1

Vel. = 0.057 ; Str. = PA ; Axis = z ; Fl. accession = 10−s−81−11

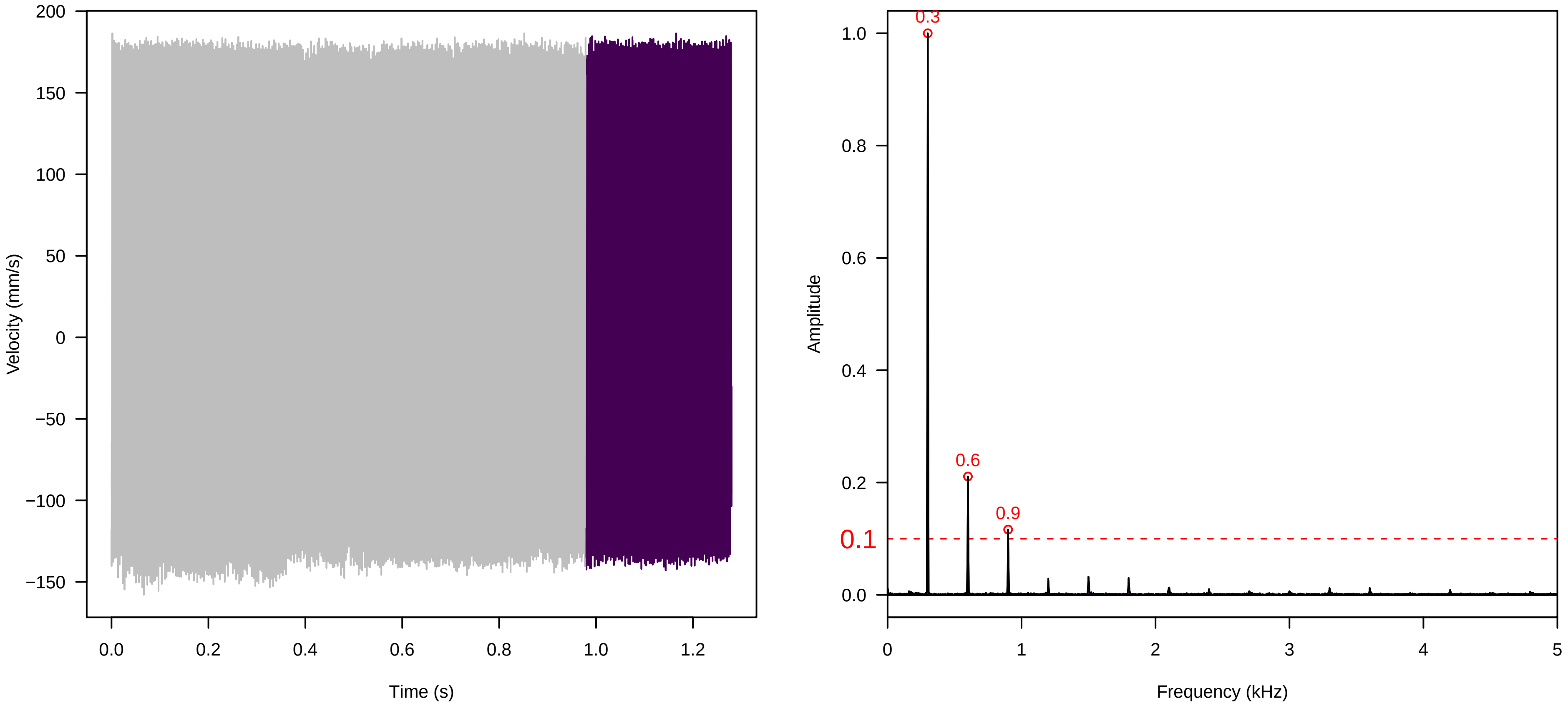


Vel. = 0.057 ; Str. = Receptacle ; Axis = z ; Fl. accession = 10−s−81−11

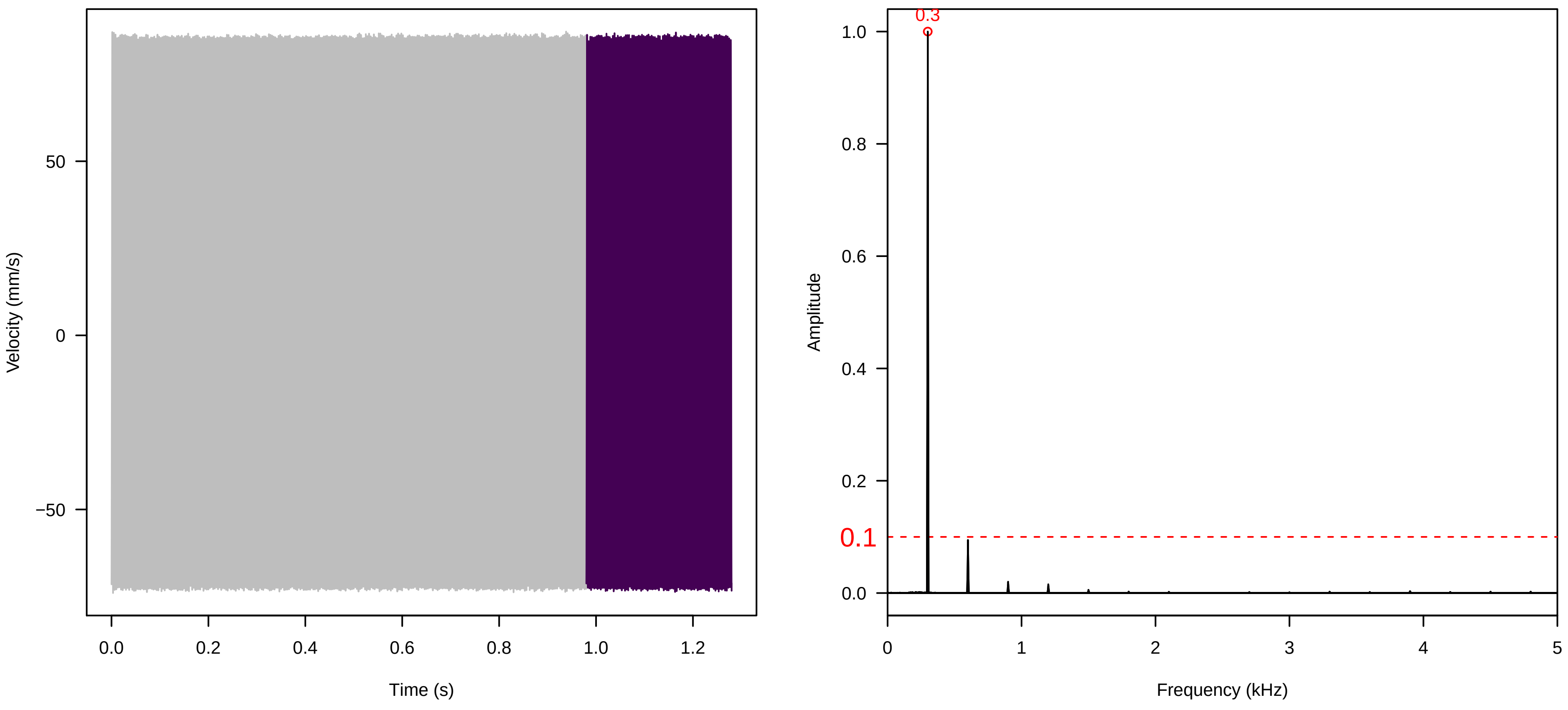


Vel. = 0.014 ; Str. = Corolla ; Axis = x ; Fl. accession = 10−s−81−11

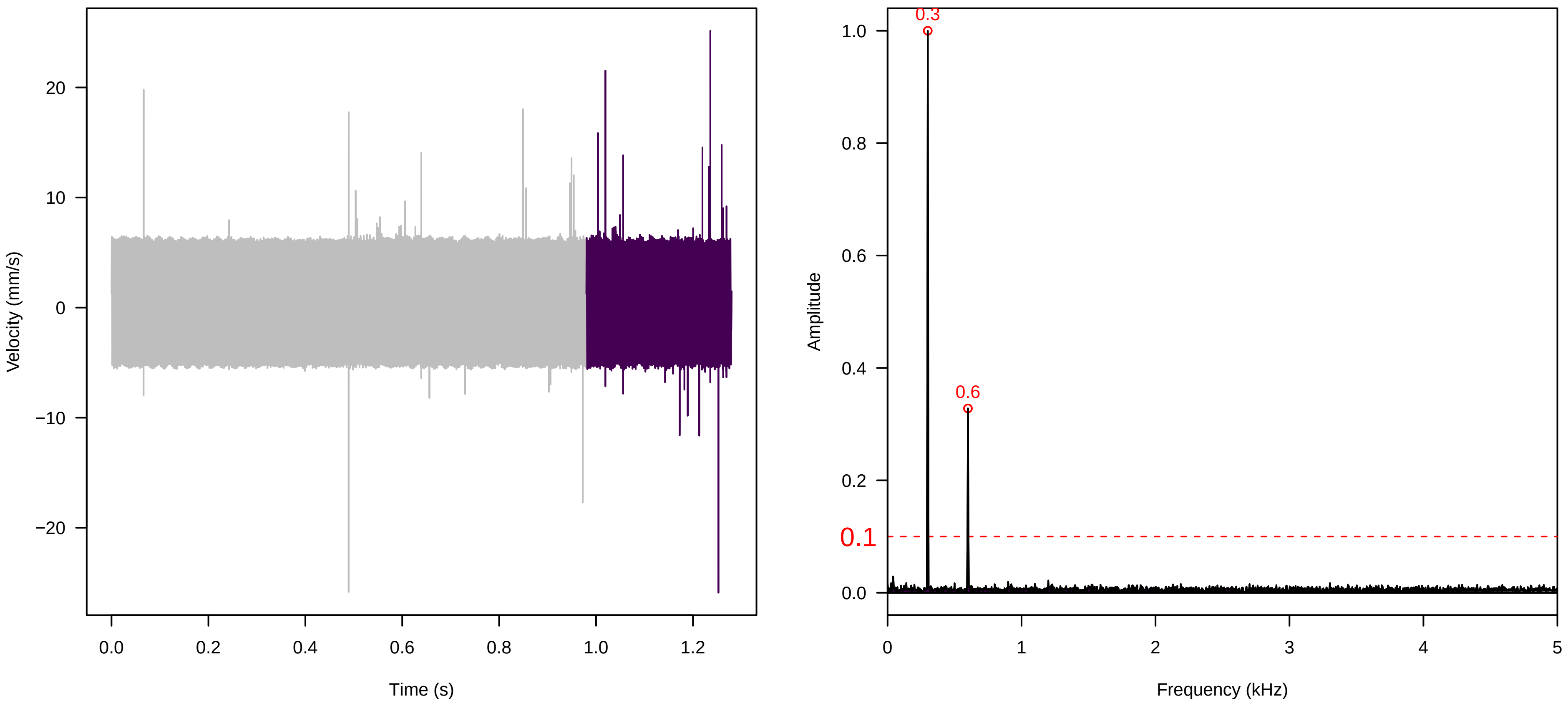

**Vel. = 0.014 ; Str. = Receptacle ; Axis = x ; Fl. accession = 10−s−81−11**

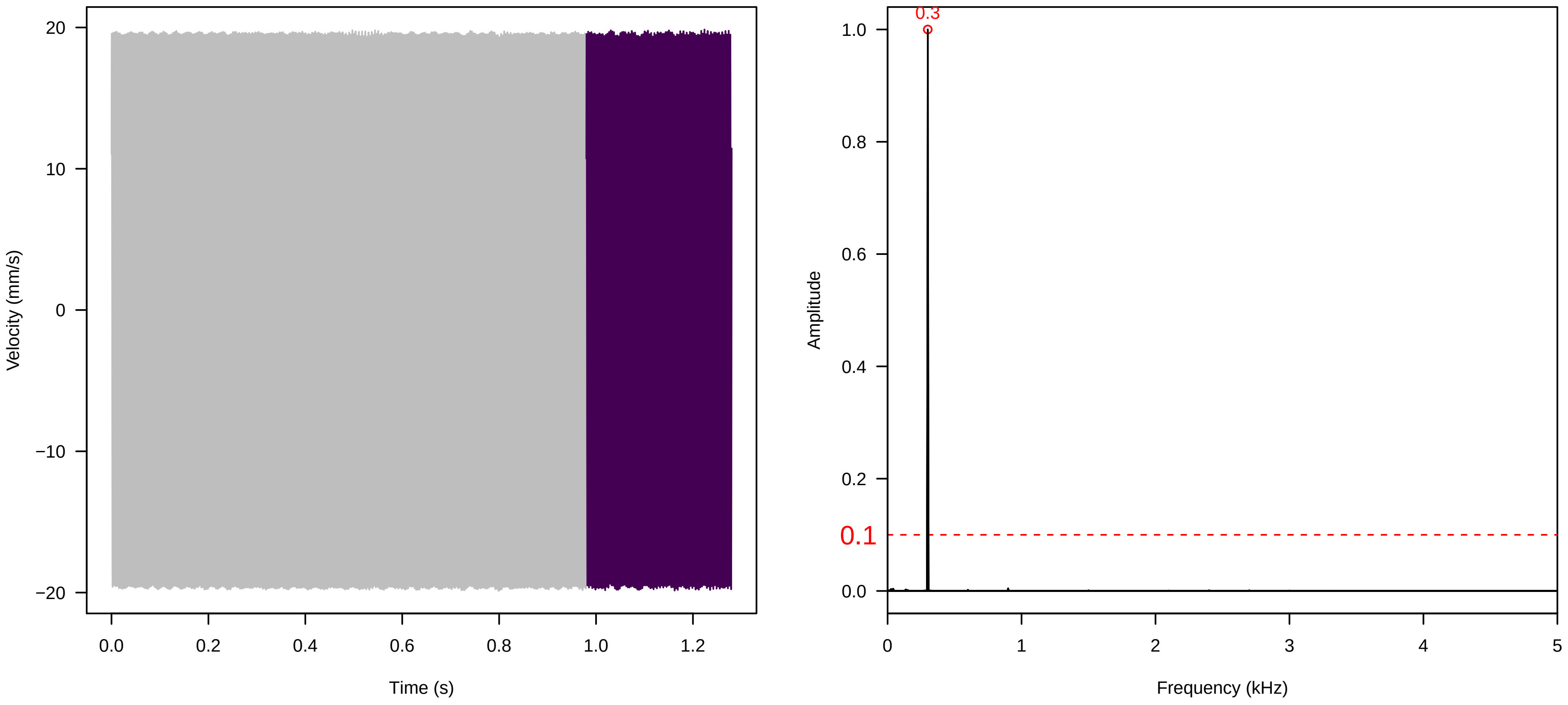


**Vel. = 0.014 ; Str. = FA ; Axis = x ; Fl. accession = 10−s−81−11**

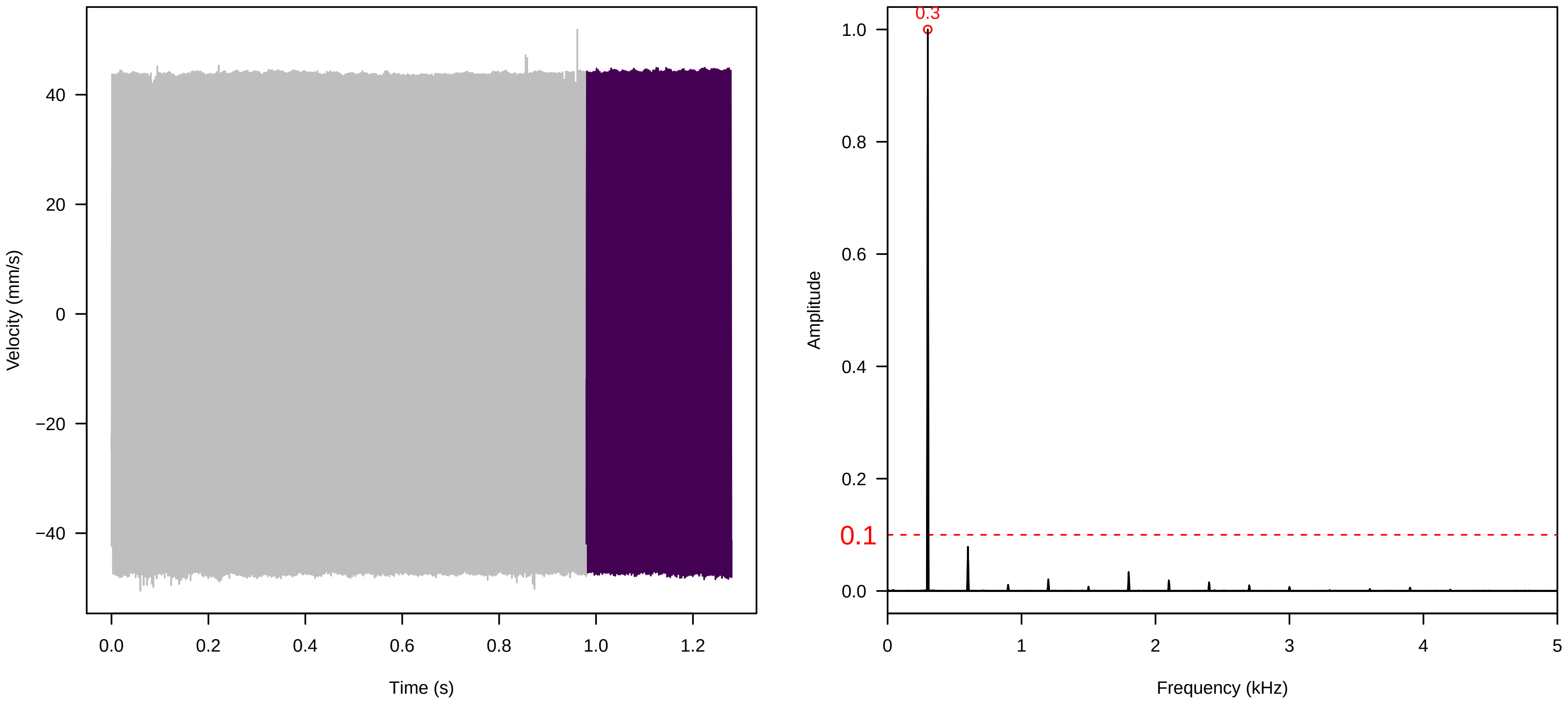


**Vel. = 0.014 ; Str. = Receptacle ; Axis = x ; Fl. accession = 10−s−81−11**

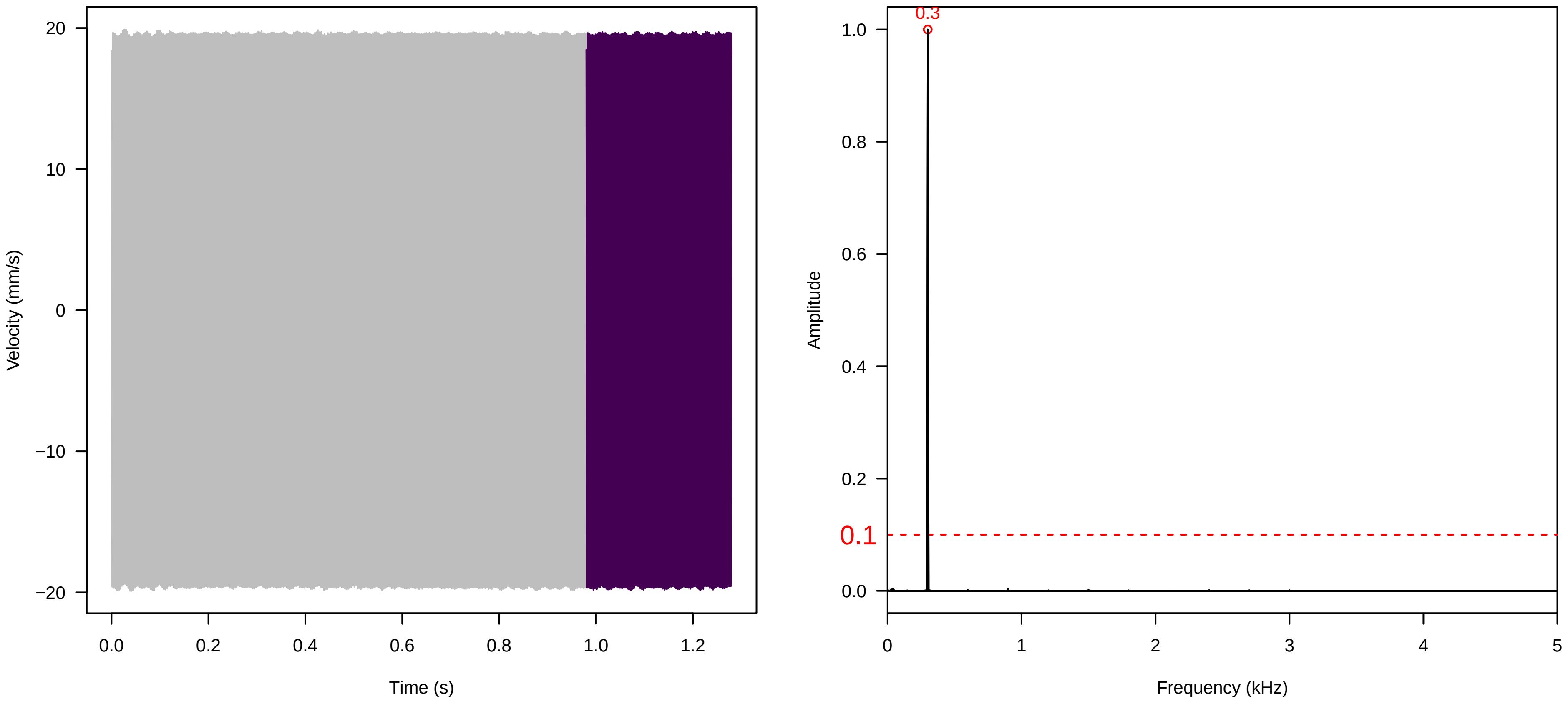

**Vel. = 0.014 ; Str. = PA ; Axis = x ; Fl. accession = 10−s−81−11**

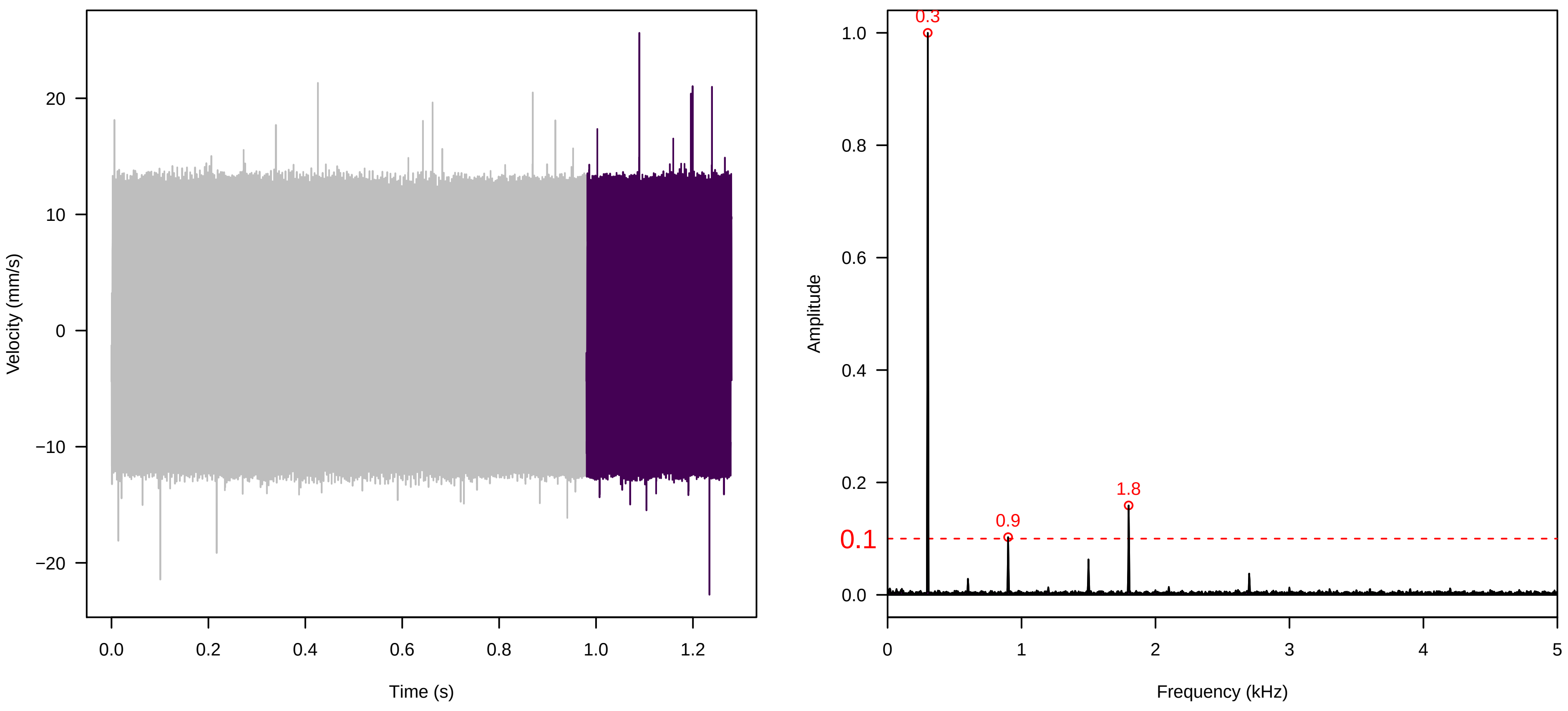


**Vel. = 0.014 ; Str. = Receptacle ; Axis = x ; Fl. accession = 10−s−81−11**

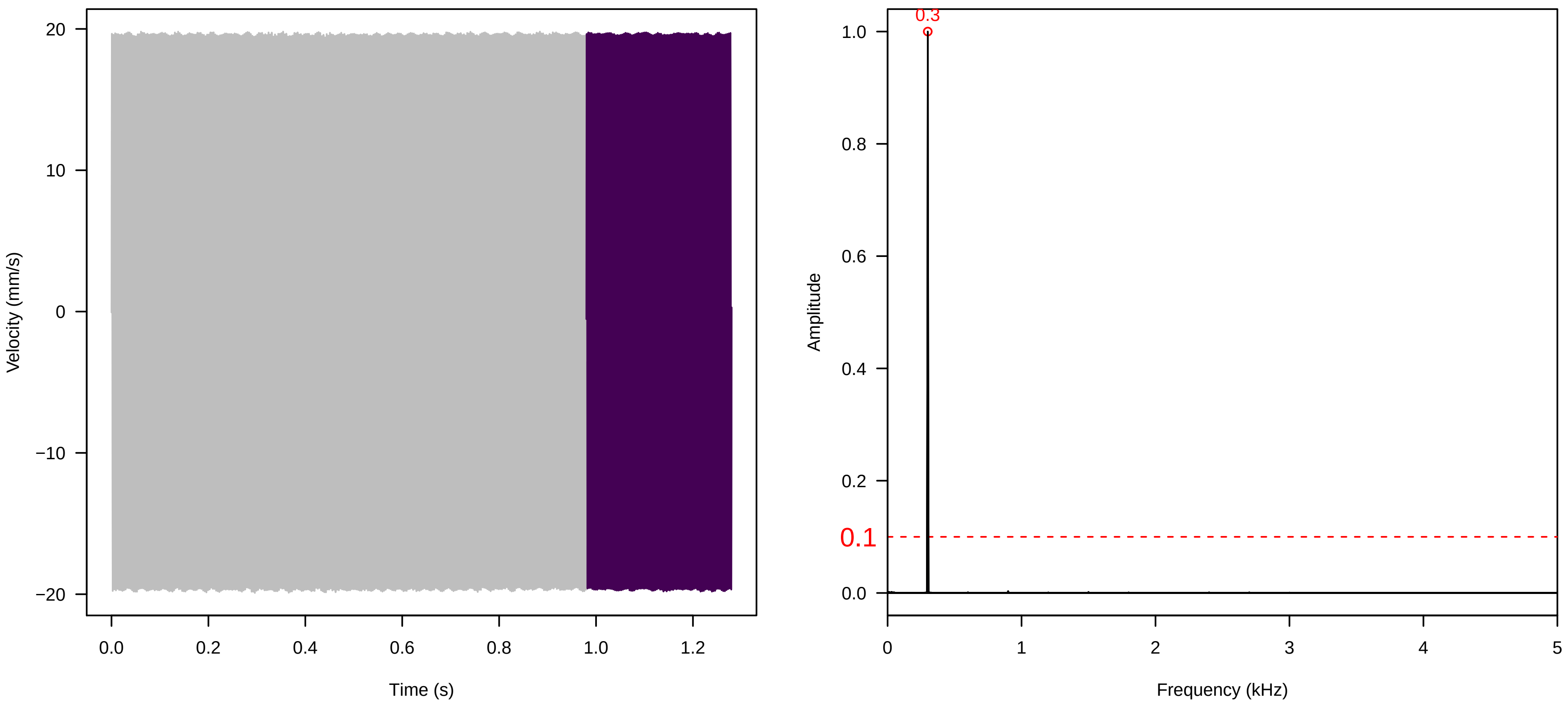


**Vel. = 0.028 ; Str. = PA ; Axis = x ; Fl. accession = 10−s−81−11**

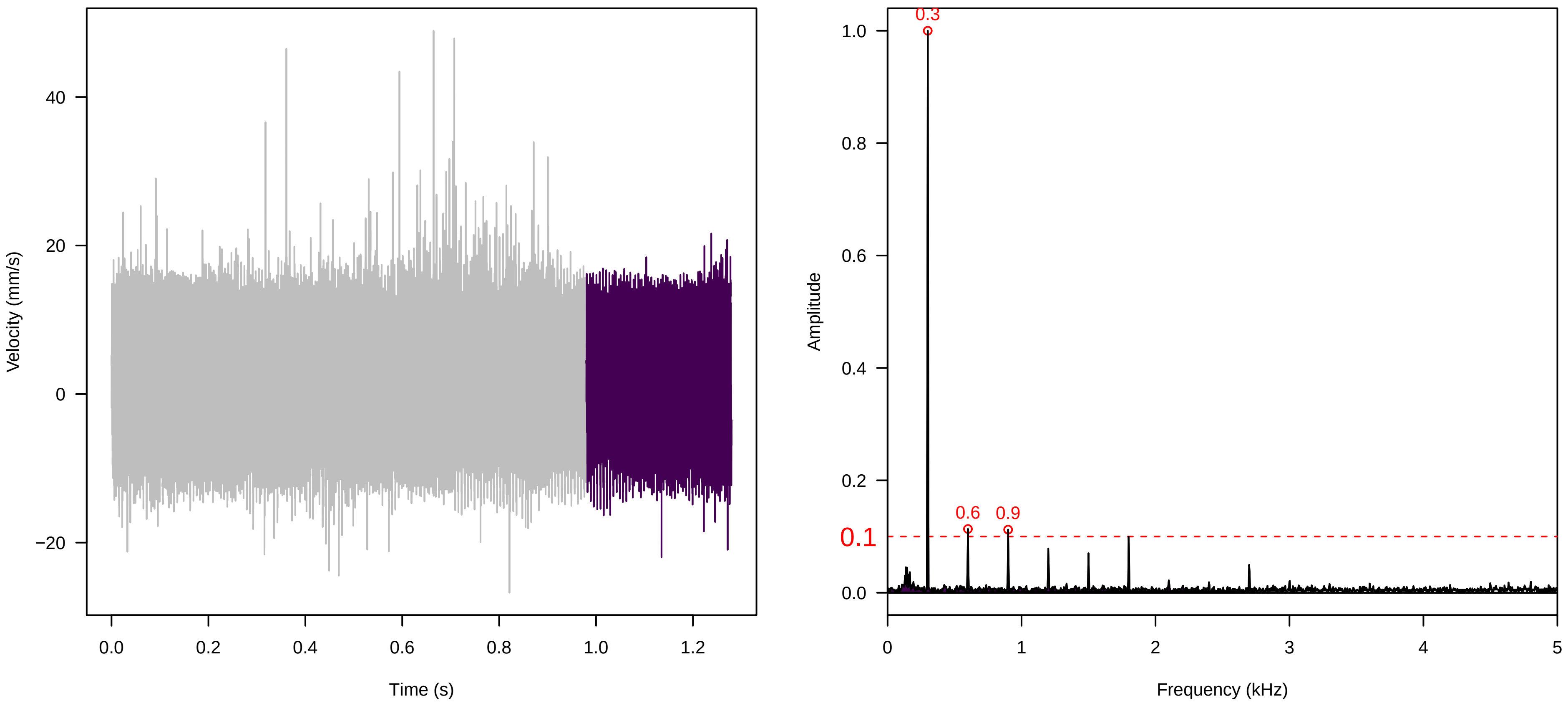

**Vel. = 0.028 ; Str. = Receptacle ; Axis = x ; Fl. accession = 10−s−81−11**

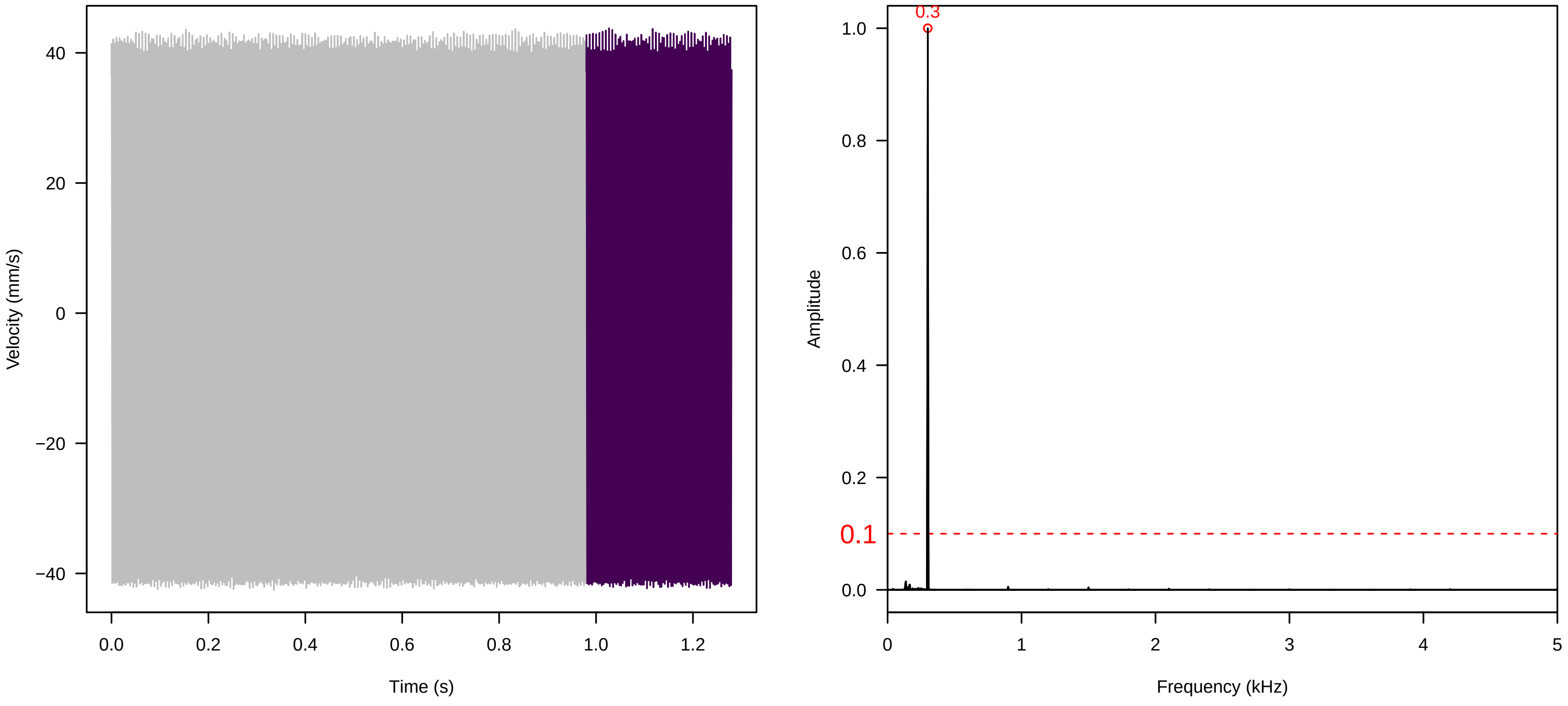


**Vel. = 0.028 ; Str. = FA ; Axis = x ; Fl. accession = 10−s−81−11**

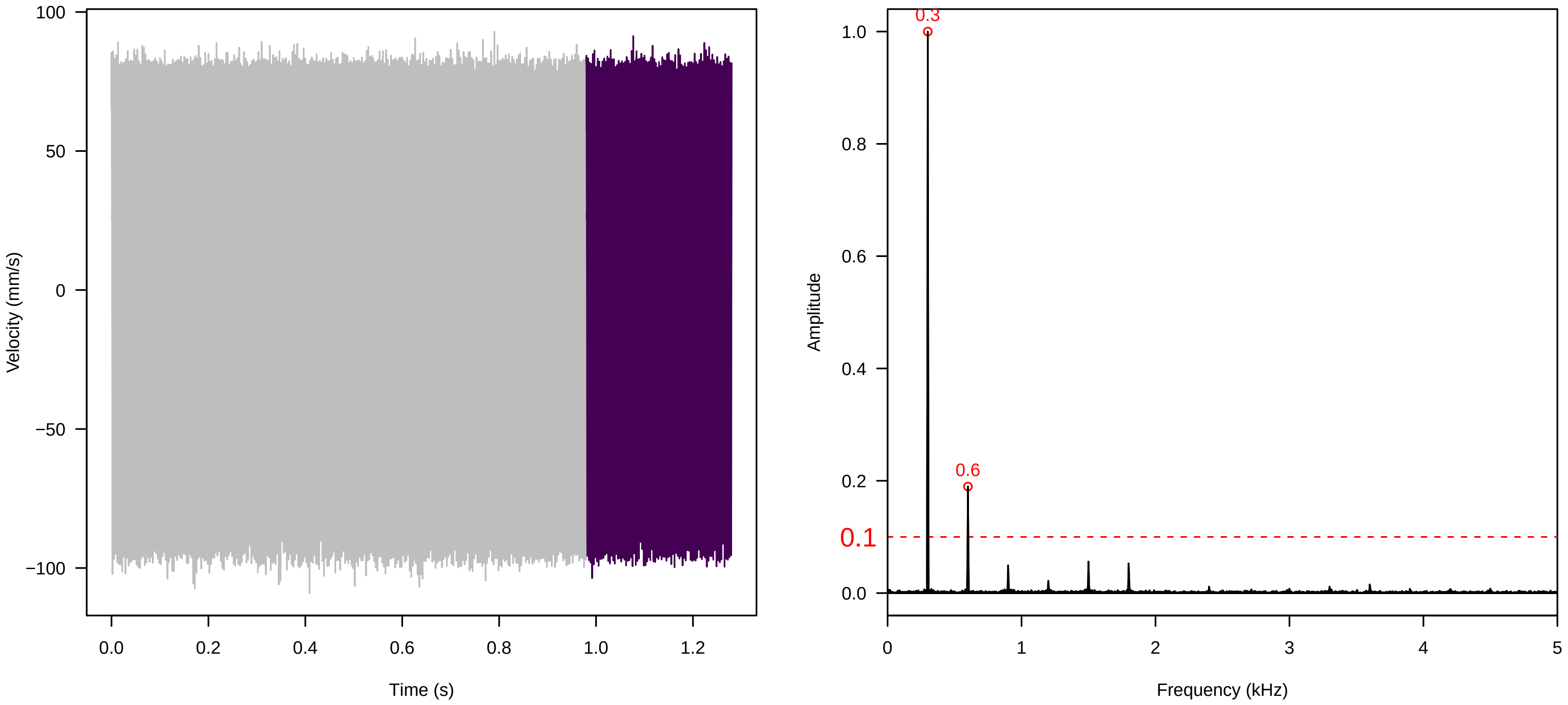


**Vel. = 0.028 ; Str. = Receptacle ; Axis = x ; Fl. accession = 10−s−81−11**

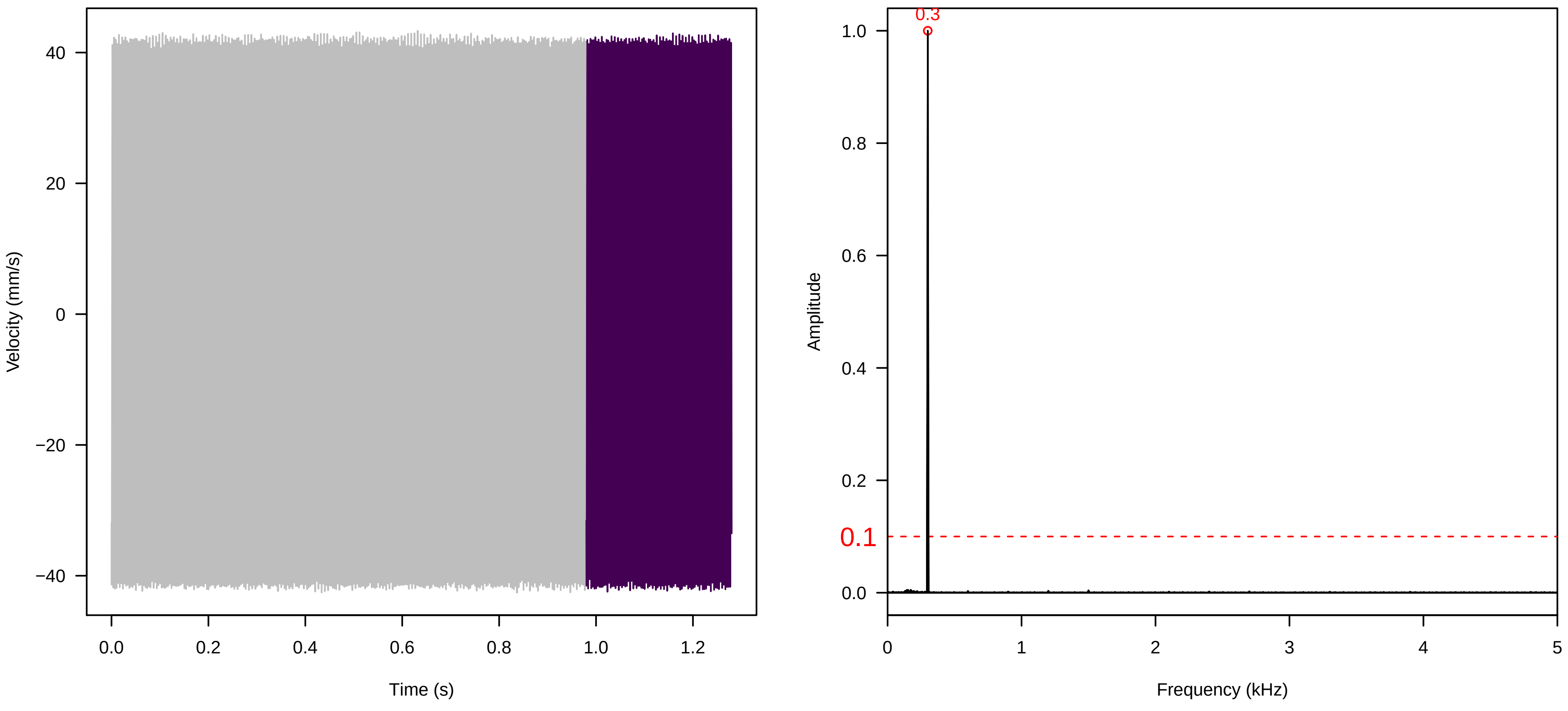

**Vel. = 0.028 ; Str. = Corolla ; Axis = x ; Fl. accession = 10−s−81−11**

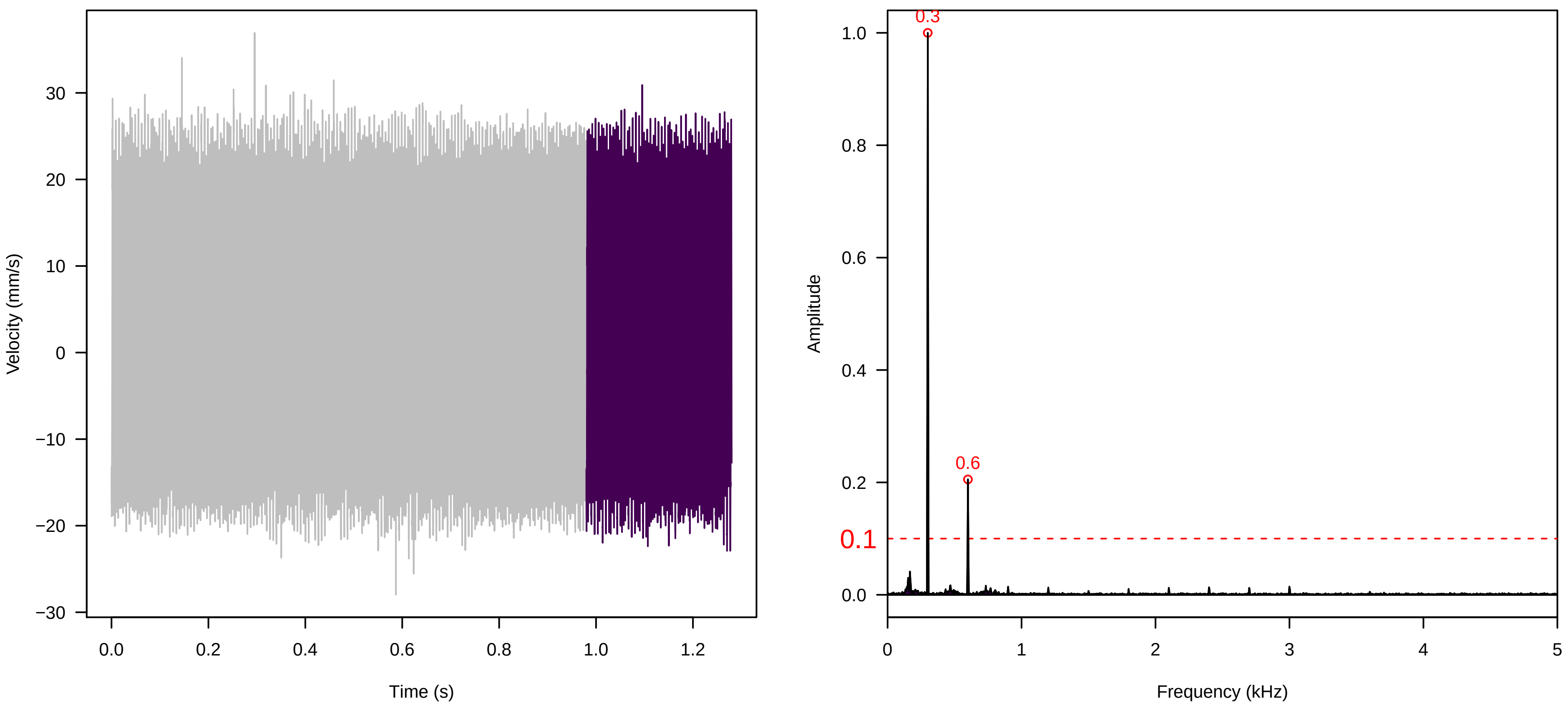


**Vel. = 0.028 ; Str. = Receptacle ; Axis = x ; Fl. accession = 10−s−81−11**

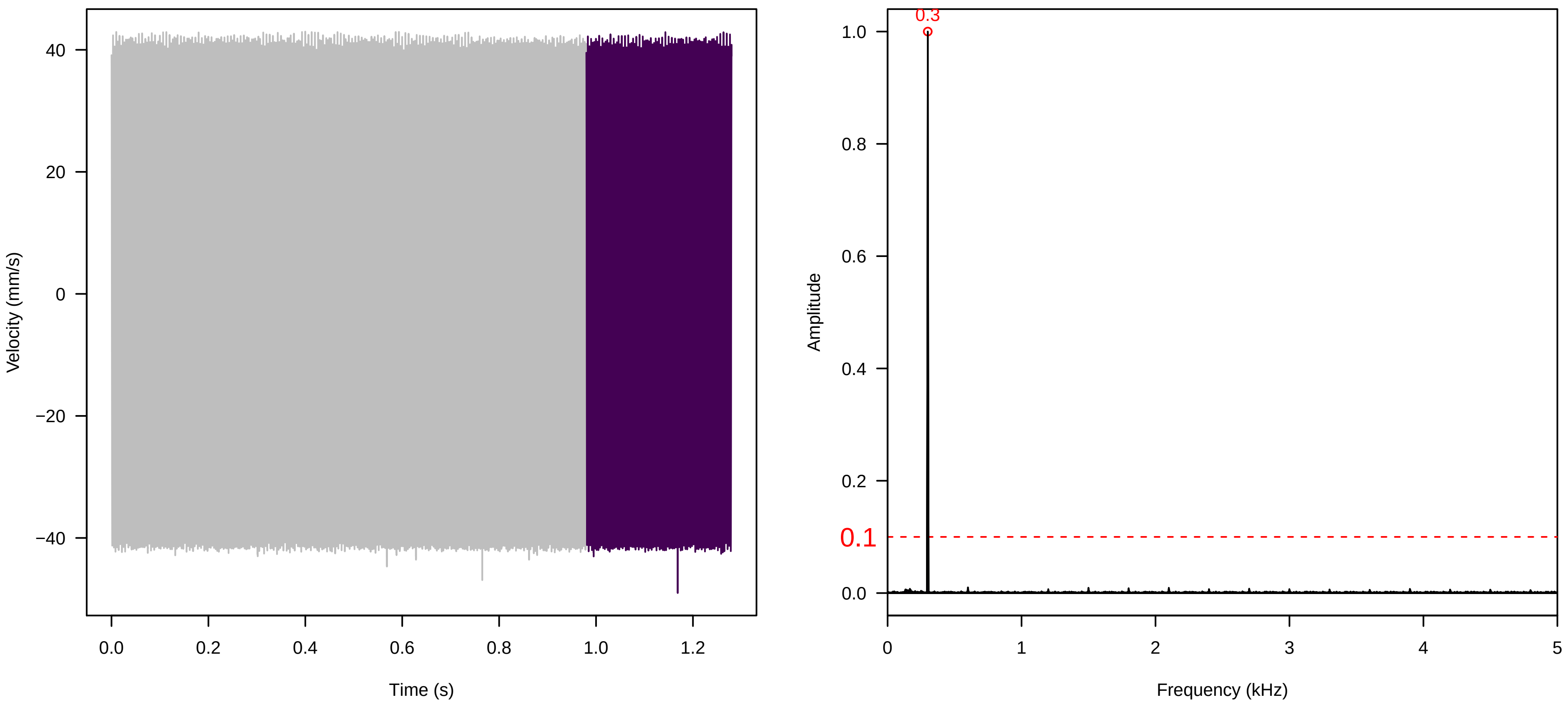


**Vel. = 0.057 ; Str. = Corolla ; Axis = x ; Fl. accession = 10−s−81−11**

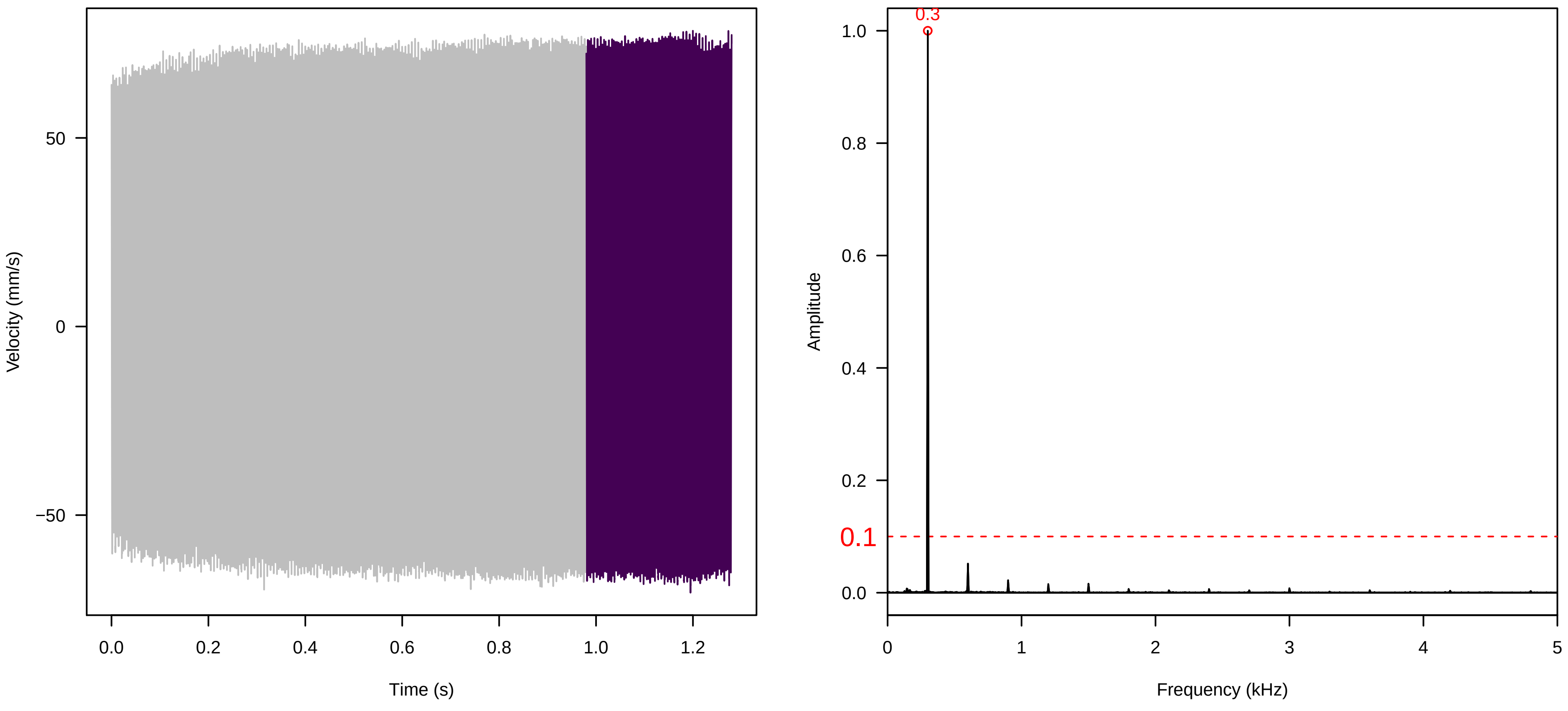

**Vel. = 0.057 ; Str. = Receptacle ; Axis = x ; Fl. accession = 10−s−81−11**

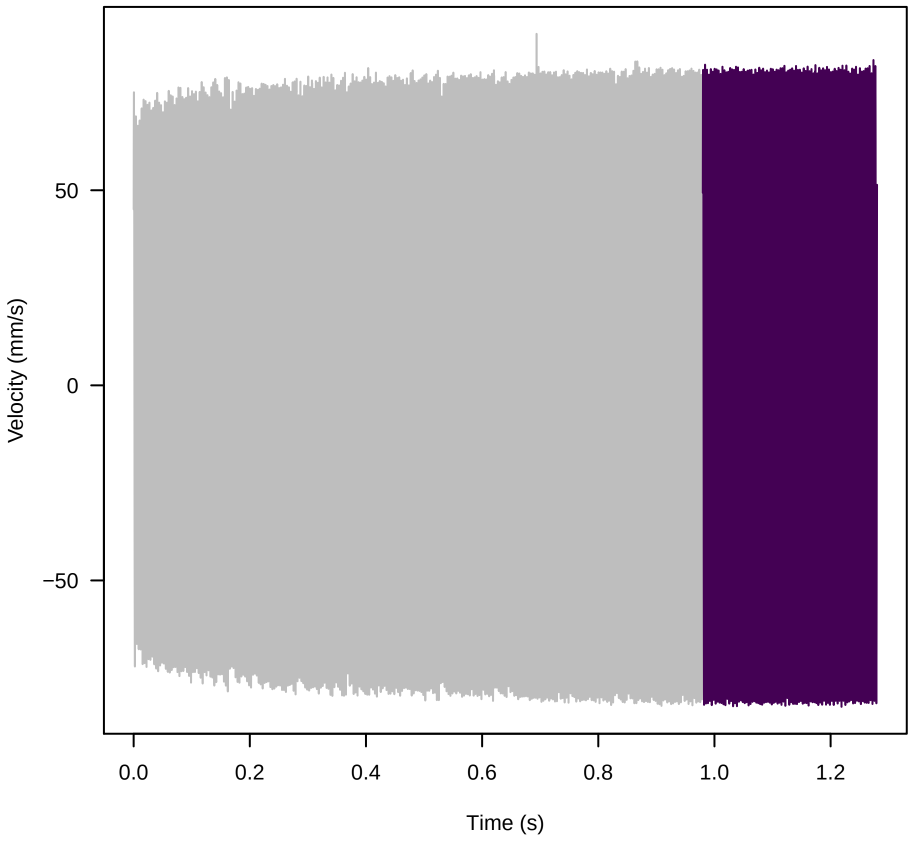


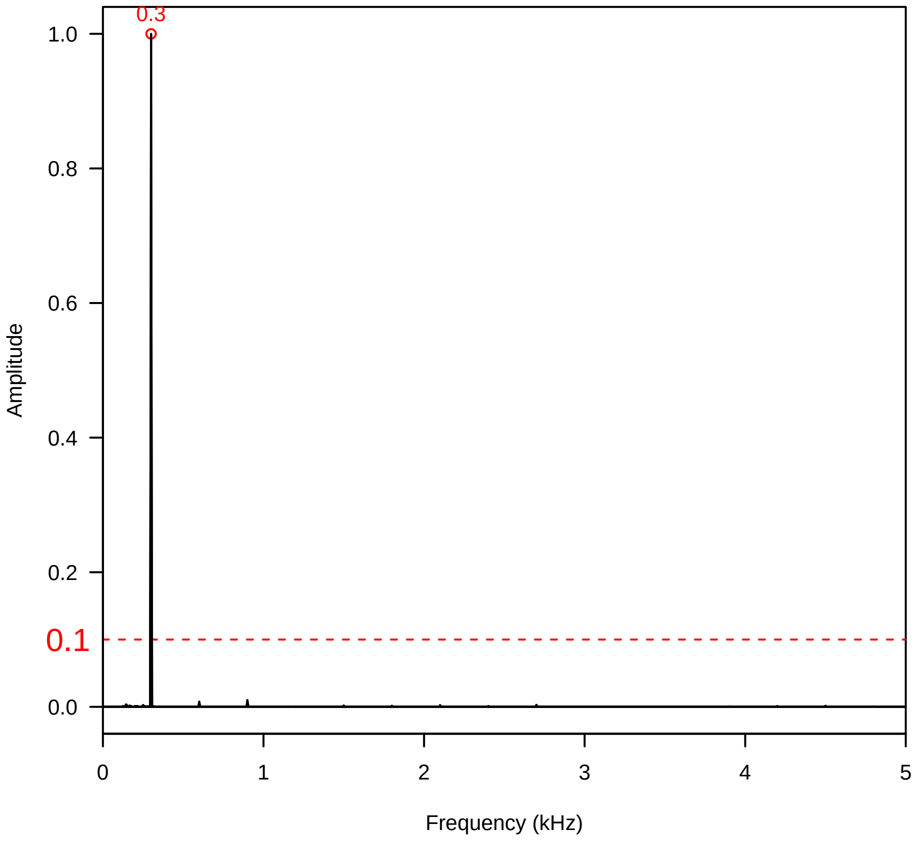


**Vel. = 0.057 ; Str. = FA ; Axis = x ; Fl. accession = 10−s−81−11**

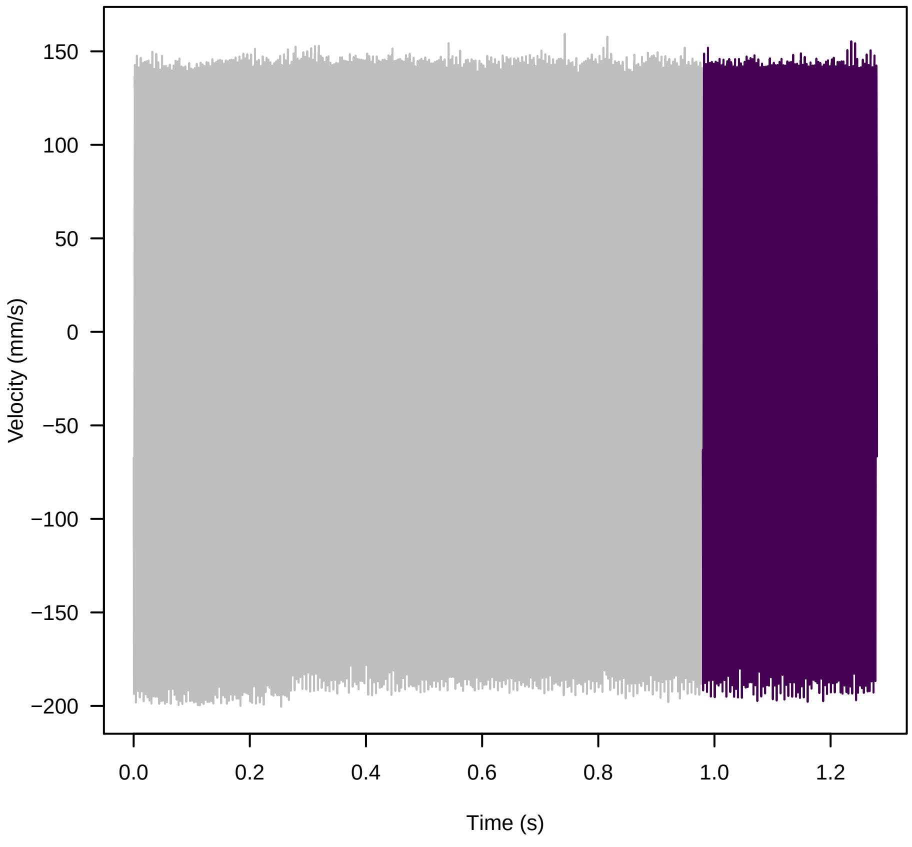


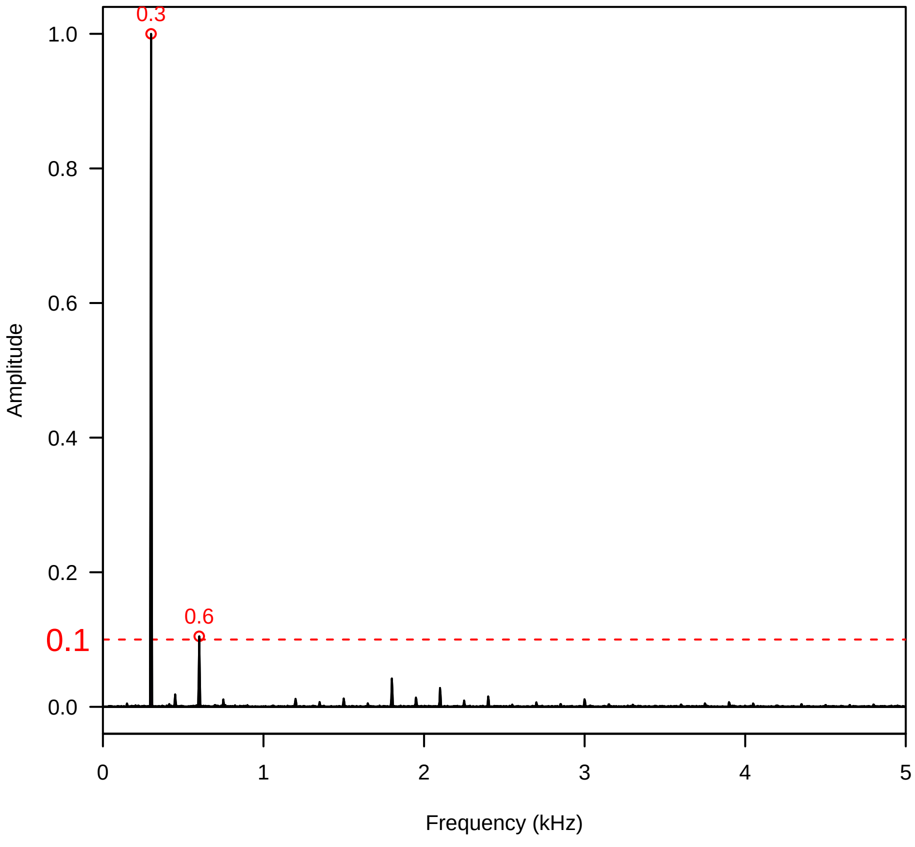


**Vel. = 0.057 ; Str. = Receptacle ; Axis = x ; Fl. accession = 10−s−81−11**

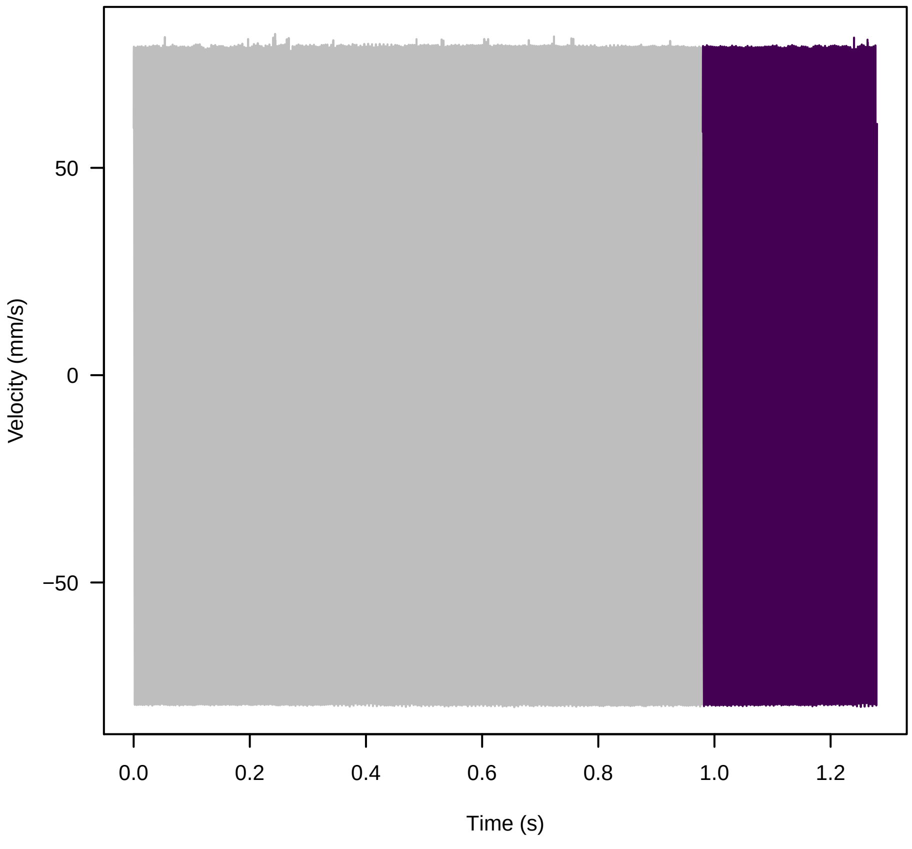


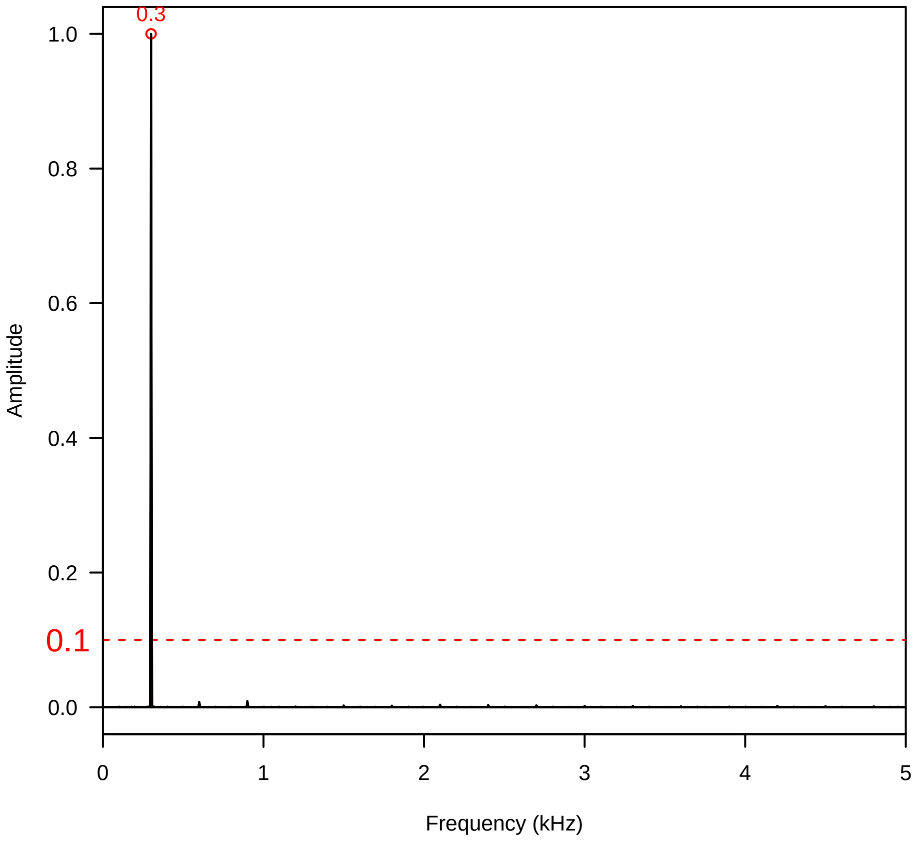

**Vel. =  0.057 ; Str. =  PA ; Axis =  x ; Fl. accession = 10−s−81−11**

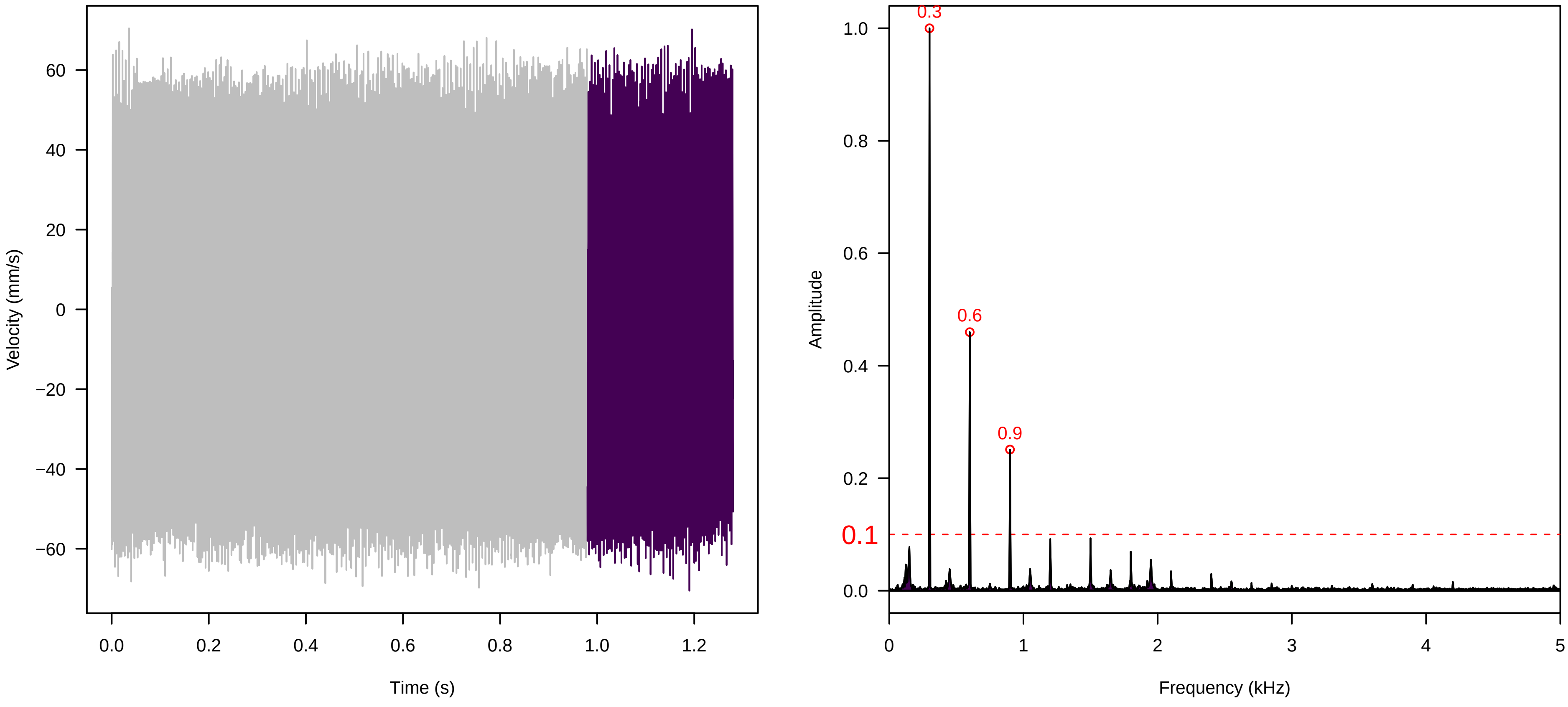


**Vel. =  0.057 ; Str. =  Receptacle ; Axis =  x ; Fl. accession = 10−s−81−11**

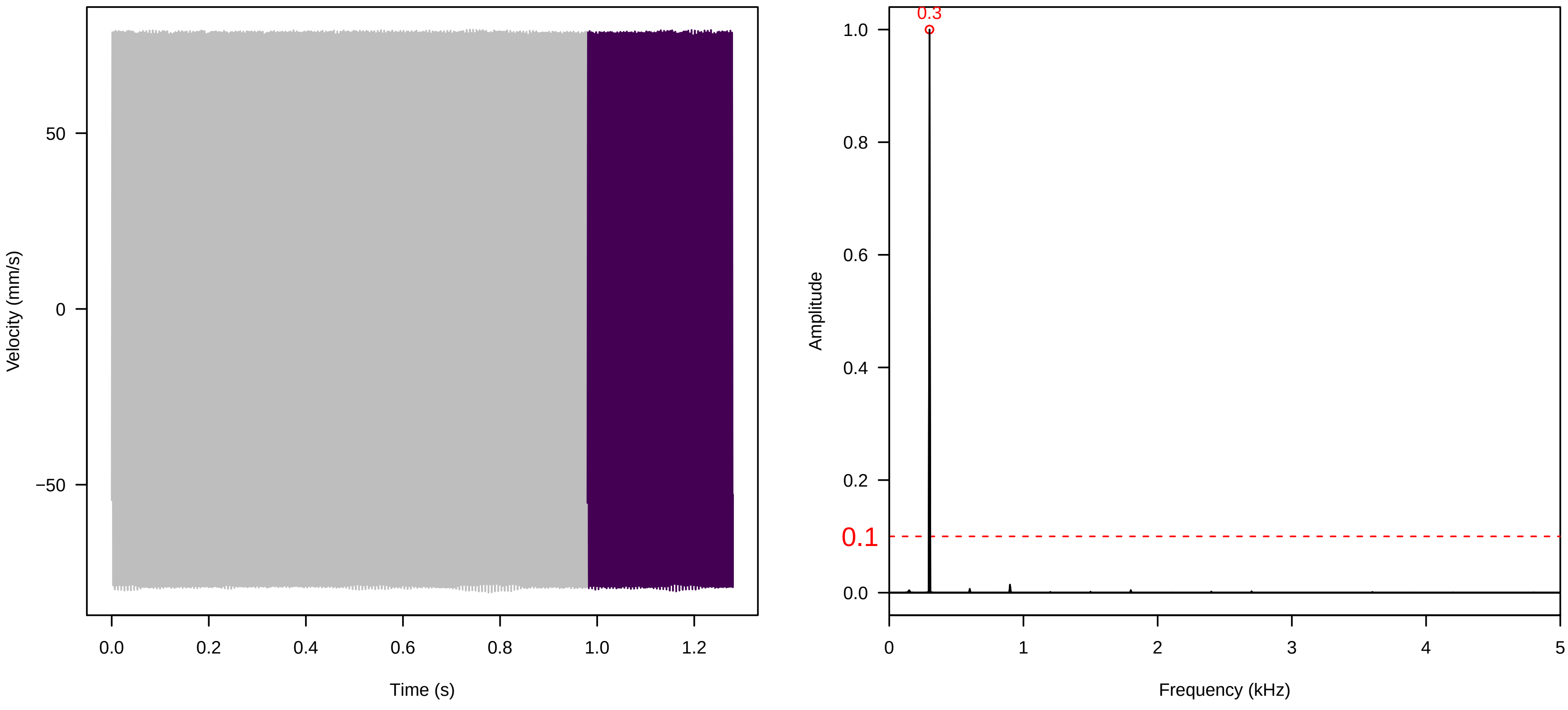
